# A single cysteine residue in vimentin regulates long non-coding RNA *XIST* to suppress epithelial-mesenchymal transition and stemness in breast cancer

**DOI:** 10.1101/2024.11.13.623301

**Authors:** Saima Usman, W. Andrew Yeudall, Muy-Teck Teh, Fatemah Ghloum, Hemanth Tummala, Ahmad Waseem

## Abstract

Vimentin is a type III intermediate filament (IF) protein, that is induced in a large number of solid tumours. A single cysteine at position 328 in vimentin plays a crucial role in assembly, organisation and stability of IFs. However, its exact function during epithelial mesenchymal transition (EMT) and cancer progression has not been investigated. To investigate this, we have transduced wildtype (WT) and C328S vimentin separately in MCF-7 cells that lack endogenous vimentin. The expression of C328-VIM impacted vimentin-actin interactions and induced EMT-like features that include enhanced cell proliferation, migration, invasion accompanied by reduced cell adhesion when compared to the wildtype cells. Functional transcriptomic studies confirmed the upregulation of EMT and mesenchymal markers, downregulation of epithelial markers as well as acquisition of signatures associated with cancer stemness (*CD56, Oct4, PROCR and CD49f*) thus transforming MCF-7 cells from oestrogen positive to triple reduced (*ESR1, PGR, and HER2*) status. We also observed a stark increase in the expression of long non-coding RNA, *XIST* in MCF-7 cells expressing C328-VIM. Targeting the mutant vimentin or *XIST* by RNA interference partially reversed the phenotypes in C328-VIM expressing MCF-7 cells. Furthermore, introduction of C328-VIM cells into nude mice promoted tumour growth by increasing cancer stemness in an oestrogen independent manner. Altogether, our studies provide insight into how cysteine 328 in vimentin dictates mechano-transduction signals to remodel actin cytoskeleton and protect against EMT and cancer growth via modulating lncRNA *XIST*. Therefore, targeting vimentin and/or *XIST* via RNA interference should be a promising therapeutic strategy for breast cancer treatment.

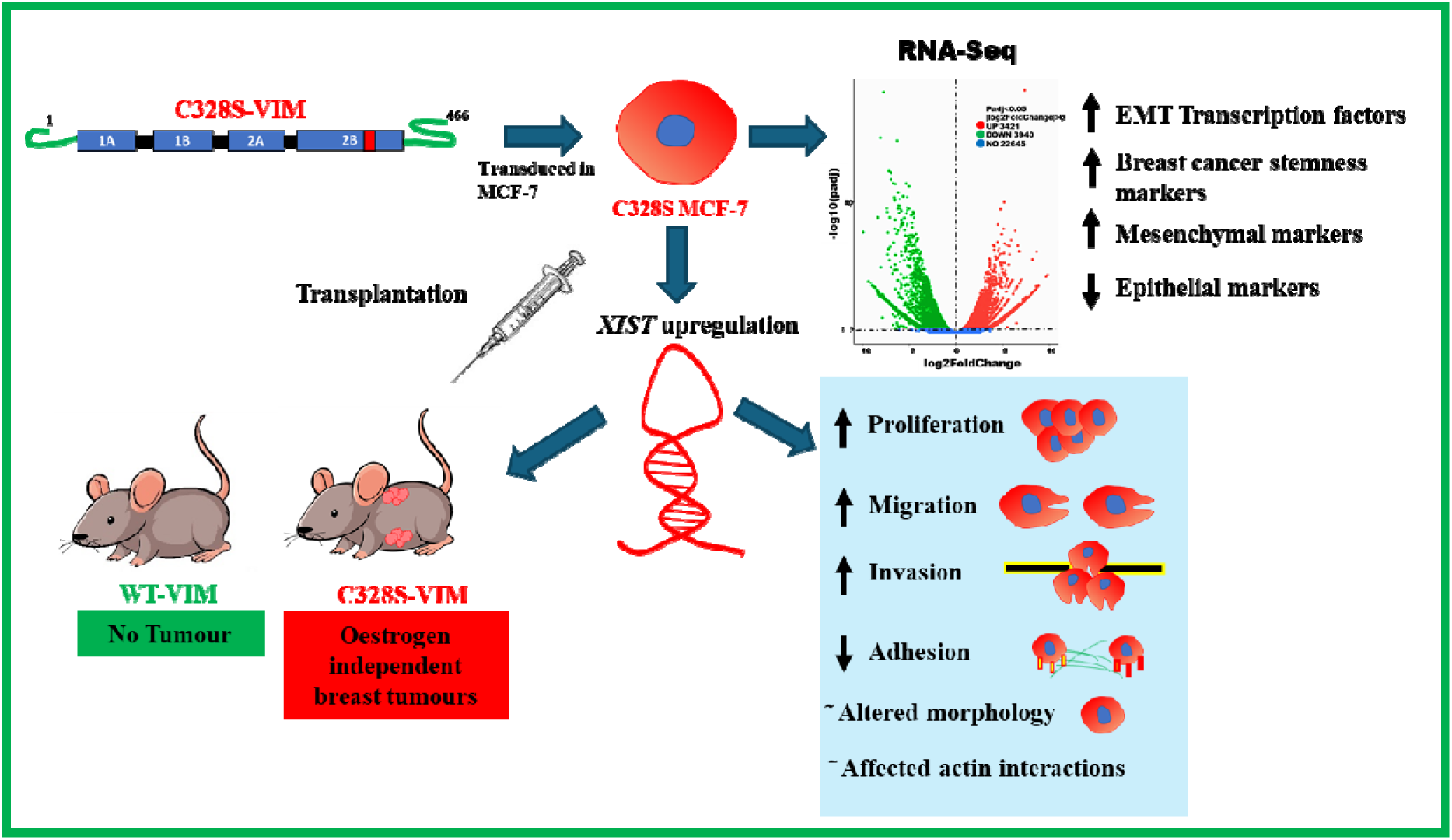

## Introduction

Vimentin is a type III intermediate filament (IF) protein normally expressed in mesenchymal cells. Due to its diverse pathophysiological role(s), it is one of the most extensively studied IFs. It has been proposed to play a role in epithelial-to-mesenchymal transition (EMT) and cancer metastasis [1]. It is a structurally dynamic protein active in cell mechanics [2], normal positioning of cell organelles, nuclear and DNA integrity [3], stress response [4], autophagy [5], cell proliferation, migration [6], adhesion, invasion [7], signalling, angiogenesis [8] and immune responses [9]. The complete crystal structure of vimentin is still not available; however, crystal structures of small segments have been investigated [10]. Structurally, vimentin monomer has a central α-helical rod domain flanked by non-helical head and tail domains on each side [11]. Studies have described its polymerisation in which monomers assemble in parallel to form dimers, with two dimers assembling in antiparallel to form tetramers [12]. Eight tetramers combine to form unit length filaments (ULFs) that further assemble head to tail into 10 nm compact mature filaments [13]. IFs lack polarity and in mature filaments subunits continue to assemble at both ends and they can be exchanged anywhere along the whole filament length, a process that requires ATP [14]. Vimentin directly interacts with actin through its tail domain to maintain mechanical integrity of cell and cytoskeletal crosstalk [15].

Vimentin has a single cysteine residue at position 328 in the rod domain, which is reported to be a stress sensor induced by oxidants and electrophiles, resulting in its zinc mediated modifications and consequently disassembly or rearrangement of filaments as a stress response [4, 16]. Different modifications of C328 that have been reported during cellular senescence, rheumatoid arthritis, cataract and atherosclerosis [17]. Any mutation or modification in C328 can hamper the intracellular stress response of vimentin. As a result, multiple cellular functions can be disrupted due to inability of the cell to sense and respond to stress. Therefore, this residue has been reported to have a widespread pathophysiological implications and is considered a flash point for stress ignition [17]. Although the role of vimenetin C328 in response to cellular stress has been studied in some detail, its role in regulating EMT, tumour growth and progression has not been studied.

To investigate the functional implications of vimentin residue C328 in regulating EMT, cancer progression and stemness, we expressed C328S vimentin in MCF-7 cells, a vimentin deficient cell line [18] commonly employed to study EMT [19] [20]. Our findings reveal that C328S-VIM altered cell morphology with disorganized F-actin and induced EMT and cancer stem cell characeristics. A highly significant observation was the upregulation of long non-coding RNA (lncRNA), *XIST,* in C328S-VIM expressing MCF-7 cells, which are normally oestrogen-dependent for tumorogenesis [21], became oestrogen-independent by C328S-VIM in nude mice. We show that C328S-VIM can trigger an EMT program and enhance stemness in MCF-7 cells, in part via *XIST* upregulation. These findings have far reaching implications, particularly considering the numerous vimentin variants reported in the vicinity of C328 residue in several solid tumours [1].

## Materials and methods

### Cell culture and cell lines

MCF-7, A431, HFF cells were obtained from the cell bank of Cancer Research UK and characterised by the expression of key biomarkers using immunocytochemistry and reverse transcription (RT)/quantitative polymerase chain reaction (qPCR). They were cultured in Dulbecco’s Modified Eagle Medium (DMEM), containing 10% (v/v) foetal calf serum (FCS), 50 units/mL penicillin and 50μg/mL streptomycin (complete medium) and maintained in the incubator in an atmosphere of 5% CO_2_ + 95% air at 37°C.

### In silico analysis

Using PyMOL, an in-silico analysis was conducted on wild type and mutant vimentin to assess their interactions with wildtype actin. The Protein Data Bank (PDB) file for the wildtype vimentin (PdB id: P08670) was acquired from www.rcsb.org, and the mutation (C328S) was introduced by subsequent energy minimisation steps.

### cDNA synthesis and qPCR

Cells were cultured in six well plate format in triplicates and lysed in 500µl Dynabeads mRNA lysis buffer when about 70% confluent. A total of 50 ng mRNA from each cell line was used to prepare the cDNA as described previously [18]. qPCR was performed with a LightCycler 480 qPCR System (Roche, Burgress Hill, UK) as described previously [18]. The forward and reverse primers for the genes studied are listed in **Table S1**.

### Protein extraction and western blotting

Cells were cultured in 6 well plates in triplicates and lysed using 250μl/well Laemmli lysis buffer, SDS gel electrophoresis and western blotting were performed as described previously [18]. All antibodies used are listed in **Table S2**. Quantification of the band intensities were performed using *Image J* [22].

### Plasmid constructs, cDNA cloning, retrovirus production and spinfection

The full-length human vimentin cDNA was sub-cloned in pLPChygro as described previously [23] and named as wildtype (WT) VIM. HPLC-purified primers (listed in **Table S3**) were used for site-directed mutagenesis (SDM) at C328 and Y117 as described earlier [24]. For preparing vimentin cDNA with double mutant (DMT), Y117L and C328S mutations were introduced sequencially and confirmed by sequencing.

For vimentin knockdown, cloning of VIM-shRNA along with non-target control (NTC) in *pSiren-Retro-Q* (Clontech, USA) has been described previously [25]. For silencing *XIST* expression, shRNA oligos were designed using the following software freely available online (http://web.stanford.edu/group/markkaylab/cgi-bin/). The forward and reverse primers (listed in **Table S3**) were annealed, phosphorylated and ligated into *pSuper.retro.puro* previously digested with *Bgl*II and *Xho*I as described earlier [26]. All clones were sequenced before use.

For packaging puromycin constructs, a one-step amphotropic retrovirus production was employed using Phoenix-A cells [27]. For hygromycin constructs, a two-step method making ecotropic retrovirus using Phoenix-E in the first step followed by an amphotropic virus production using PT67 [28] in the second step was employed [29].

Fifty thousand MCF-7 cells were seeded in T25 culture flask in complete medium and retroviral transduction by spinfection was performed as described previously [18]. All transduced cell lines are listed in **Table 1**.

**Table 1:** Summary of cell lines used in this study.

| Name of cell line | Insert | Drug selection | Retrovirus vector | Parent cell line |
| --- | --- | --- | --- | --- |
| **MCF-7CV** | None; vector | Hygromycin | pLPChygro | MCF-7 |
| **MCF-7WT** | WT-VIM | Hygromycin | pLPChygro-VIM | MCF-7 |
| **MCF-7C328S** | C328S-VIM | Hygromycin | pLPChygro-C328S-VIM | MCF-7 |
| **MCF-7Y117L** | Y117L-VIM | Hygromycin | pLPChygro-Y117L-VIM | MCF-7 |
| **MCF-7DMT (double mutant)** | Y117L,C328S-VIM | Hygromycin | pLPChygro-DMT-VIM | MCF-7 |
| **MCF-7C328S_shNTC** | Non-target control oligos | Puromycin | pSiren-Retro-Q vector | MCF-7C328S |
| **MCF-7C328S_shVIM** | Vimentin specific short hairpin oligos | Puromycin | pSuper.retro.puro | MCF-7 C328S |
| **MCF-7C328S_shNTC** | Non-targeting control<br>oligos | Puromycin | pSuper.retro.puro | MCF-7 C328S |
| **MCF-7C328S_shXIST-1** | *XIST* specific short<br>hairpin 1 | Puromycin | pSuper.retro.puro | MCF-7 C328S |
| **MCF-7C328S_shXIST-2** | *XIST* specific short<br>hairpin 2 | Puromycin | pSuper.retro.puro | MCF-7 C328S |
| **MCF-7C328S_shXIST-3** | *XIST* specific short<br>hairpin 3 | Puromycin | pSuper.retro.puro | MCF-7 C328S |
| **MCF-7C328S_shXIST-4** | *XIST* specific short<br>hairpin 4 | Puromycin | pSuper.retro.puro | MCF-7 C328S |
| **MCF-7WT+AcGFP-**<br>**C328S-VIM** | C328S-VIM | Puromycin | pLPCpuro-AcGFP-GS10 | MCF-7WT |
| **MCF-7WT+ C328S-VIM** | C328S-VIM | Puromycin | pLPCpuro | MCF-7WT |
| **HFF-AcGFP-C328S-**<br>**VIM** | C328S-VIM | Puromycin | pLPCpuro-AcGFP-GS10 | HFF-1 |
| **HFF-C328S-VIM** | C328S-VIM | Puromycin | pLPCpuro | HFF-1 |
| **A431-C328S-VIM** | C328S-VIM | Puromycin | pLPCpuro | A431 |

### Analysis of cell parameters

For analysing morphological features, 10,000 cells were seeded in 96 well plates in triplicates, and stained with CellMask^TM^ Deep Red (Invitrogen, cat # H32721) (1µl in 200 ml) and DAPI (1μg/ml) for 2 h and washed with PBS 3x times as described earlier [18]. Cell morphology was analysed using an INCA 2200 IN Cell Analyzer GE Widefield System. More than 2000 cells were counted from each cell line using GE IN Carta software (INCarta Cytiva, USA).

### Functional assays

Colony formation assay, CyQUANT^TM^ cell proliferation, CyQUANT^TM^ cell adhesion, chemotactic migration and MTT assays were performed as described earlier [18, 30]. For invasion assay, culture inserts (8 µm pore size) were coated with Matrigel (1:20) in a 24-well plate in serum-free medium in triplicate and the same procedure was followed as described for migration assay [18].

### RNA-Seq analysis and bioinformatics

Cells were plated in 10cm dishes in duplicates and, when they reached 80-90% confluence, they were washed twice with PBS and total RNA was extracted using a Qiagen RNeasy Kit (cat # 74104) according to the manufacturer’s protocol. Samples were processed by Novogene Europe (Cambridge, UK) for library preparation and bioinformatics analyses [18].

### Transplantation assay

All transplantation experiments were carried out at Augusta University under Institutional Animal Care and Use Committee (IACUC) protocol number 2015-0736, in animal biosafety level 2 (ABSL2) conditions. Stably transduced MCF-7 cells (MCF-7CV, MCF-7WT, MCF-7C328S, MCF-7Y117L and MCF-7DMT) were cultured under standard growth conditions as outlined earlier, until they reached 80% confluence. Cells were trypsinized, counted, and resuspended in complete growth medium at a density of 1×10^7^ cells per ml. Subsequently, cells were transplanted subcutaneously and bilaterally into the flanks of 6-week-old female athymic nu/nu mice (2.5×10^6^ cells/0.25ml), 2 mice per cell line. Mice were monitored daily for general health and tumour formation. Once tumors became evident, caliper measurements were made weekly until the experimental endpoint. Tumour volume was determined for all groups using the formula: V = (W^2^ × L)/2, where V is the tumour volume, W is the tumour width and L is the tumour length. Mice were euthanized by CO_2_ inhalation followed by bilateral thoracotomy. Tumours were excised, fixed in formalin, paraffin-embedded, sectioned at 5μm, stained with haematoxylin and eosin, and imaged using a Keyence BZ-X700 microscope.

### Flow cytometry analyses

Cells were harvested by trypsinisation, centrifuged at 600 x g for 5 min, washed 3X in PBS (supplemented with 0.5% BSA), and resuspended in appropriate volume of Flow Cytometry Staining Buffer (FCSB) (eBioscience™ cat. #□00-4222-57) to a the final cell concentration of 1 x 10^7^□cells/mL. Cells were stained for surface antigens CD56 and CD201 using antibodies listed in **Table S1** in FCSB so that the final volume was 100 µL (i.e., 50 µL of cell sample + 50 µL of antibody mix) for 30 min at 2–8°C (in the dark). The labelled cells were washed twice in FCSB and centrifuged at 600 x g for 5 min at RT. The cell pellet was resuspended in azide-free and serum/protein-free PBS and 1 μl of Fixable Viability Dye (FVD) (listed in **Table S1**) was added per 1 mL of suspension. Cells were vortexed and incubated for 30 min at 2–8°C in the dark. These cells were washed twice with FCSB and suspended in 0.5 mL FCSB and analyzed using a BD LSRII Analyzer in the Blizard Flow Cytometry Core Facility at QMUL. Data analyses were carried out using *FlowJo v10*.

### Immunohistochemistry

For the detection of stem cell markers in mouse tumors, 5 μm tissue sections were dewaxed in Safe-Clear (Fisher Scientific, Pittsburgh, PA), rehydrated, and antigen retrieval was performed by incubation in Retrievagen A (BD Biosciences, CA) at 95°C for 30 min. After cooling to ambient temperature, endogenous peroxidase was blocked by incubation in 3% (v/v) hydrogen peroxide for 15 min, slides were washed in Tris-Buffered Saline (TBS) pH8.0, and blocked in normal goat serum (Vector Laboratories, Burlingame, CA). Sections were then incubated with anti-CD56 or anti-CD201 antibodies listed in **Table S1** or the equivalent concentration of normal rabbit or normal goat IgG as control, at 4°C for 16h. Slides were washed in TBS, and then incubated sequentially with biotinylated anti-rabbit/mouse (for CD56) or biotinylated anti-goat (for CD201) secondary antibodies and streptavidin-peroxidase reagent (Vectastain Elite ABC-HRP Kit; Vector Laboratories, Burlingame, CA, cat. # PK-6100) according to the manufacturer’s instructions. Colour development was achieved using 3, 3’-diaminobenzidine (DAB) substrate, and slides were lightly counterstained with Harris hematoxylin, dehydrated, mounted in Permount, and imaged by brightfield microscopy (Keyence Corporation, Itasca, IL). Routine hematoxylin and eosin (H & E) staining of tumor sections was performed by the Electron Microscopy and Histology Core Laboratory, Medical College of Georgia, Augusta University, as described previously [31]. The integrated optical density (IOD) of CD56 and CD201 was calculated as the product of optical intensity of positive cells x area of positive cells using *ImageJ* [32].

### Statistical analyses

All experiments were performed in triplicates (technical repeats). To compare the two groups, two-tailed Student’s t-tests were applied on raw data using Microsoft Excel and p values below 0.05 (p <0.05) were considered significant. To compare more than two groups, ordinary one way or two way Analysis of Variance (ANOVA) was performed in combination with Bonferroni’s test using *GraphPad Prism 10*. Linear regression analyses and Pearson correlation coefficients (Pearson’s r) were determined using data analysis *ToolPak* in *Microsoft Excel*. All the results were represented as the mean of 3 individual experiments (n = 3) with standard error of the mean (± S.E.M).

## Results

### C328S mutant vimentin affects interaction with actin *in silico* and F-actin formation in cells

We selected MCF-7, a simple epithelial breast carcinoma cell line, which lacks endogenous vimentin [18], and transduced it with the full-length vimentin (wildtype, WT-VIM/pLPChygro-VIM) and the mutant vimentin (mutant, C328S-VIM/pLPChygro-C328S-VIM) retroviruses (**Table 1**). The expression of WT-VIM and C328S-VIM in MCF-7 cells was confirmed by qPCR (**Figure 1).**

**Figure 1:**
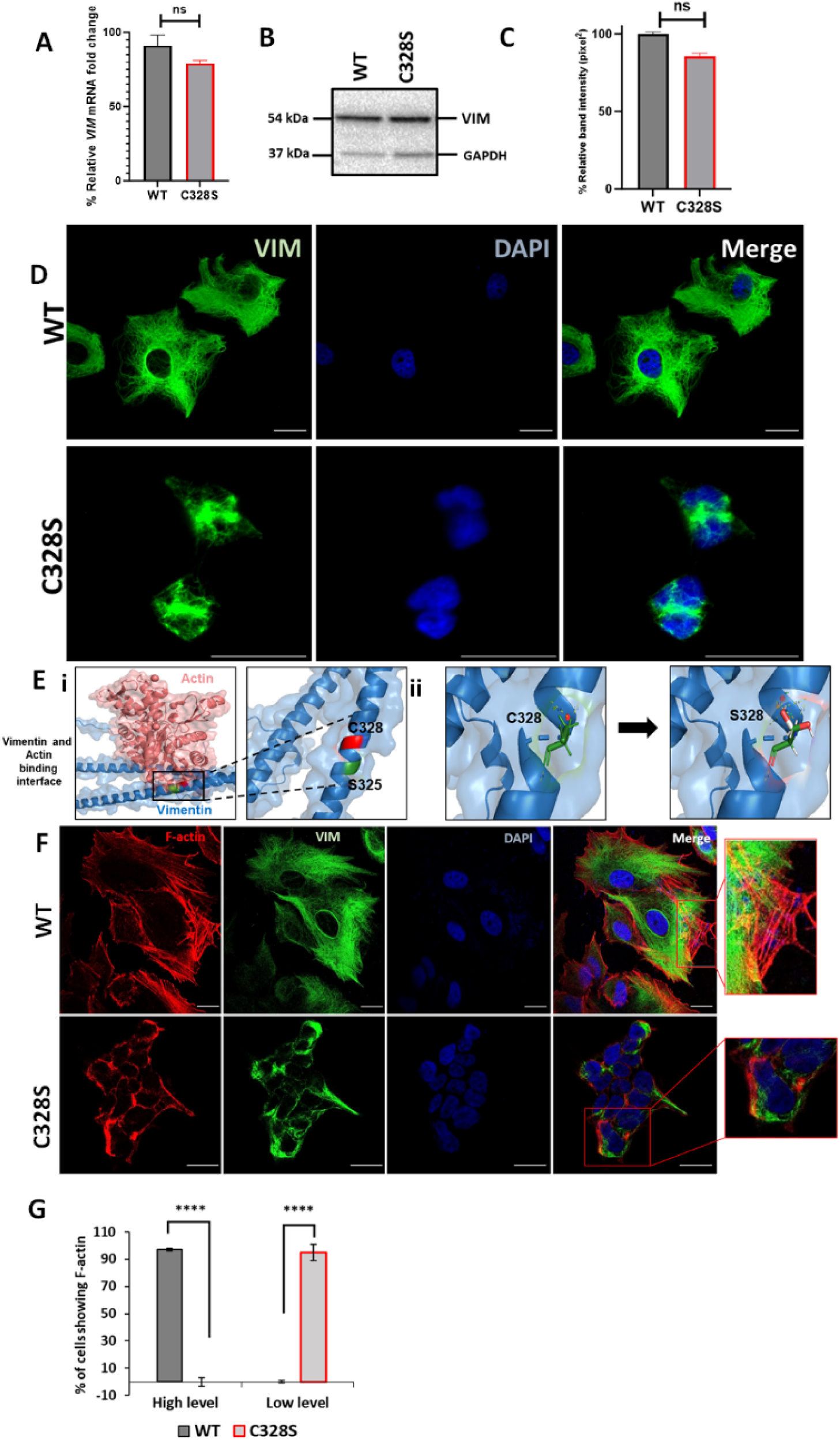
C328S mutant vimentin affects interaction with actin in silico and F-actin formation in cells. (A): Relative fold change of VIM mRNA in WT-VIM and C328S-VIM expressing MCF-7 cells normalised to POLR2A and YAP1. (B): Expression of WT-VIM and C328S-VIM in MCF-7 cells by western blotting. (C): Quantification of vimentin using ImageJ. (D): Immunostaining of MCF-7 cells expressing WT-VIM and C328S-VIM. Cells were immunostained with mouse anti-vimentin V9, AF-488 labelled goat anti-mouse showing green fluorescence. Nuclei were stained with DAPI in blue and the overlapping images are shown as Merge. Leica DM4000B Epi-fluorescence microscope was used for imaging (Scale bar = 20 µm; the scale bar in MCF-7C328S cells is much longer as these cells were reduced in size). (E): In silico modelling of actin binding at the interface where C328 and S325 residues are located. The solid rectangular area in the panel (i) is zoomed in the accompanying panel. The red colour indicates the C328 residue site whereas the green colour indicates the S325 residue site. (ii) In-silico modeling of C328 and S328 residue sites and rotamer conformations further zoomed in from the panel (i). The green colour indicates hydrogen bonds, the white colour indicates covalent bonds and the red colour indicates oxygen atom. (F): Immunostaining of MCF-7 cells expressing WT- and C328S-VIM. Cells were immunostained with AF568 Phalloidin (red colour) and anti-vimentin (green colour) antibody. Nuclei were in blue and the overlapping images are shown as Merge. Images were taken by Zeiss 880 laser scanning confocal microscope with Fast Airyscan and Multiphoton (inverted) system (scale bar = 20 µm). (G): Percentage of cells expressing F-actin in WT and C328 cells (see **Figure S3** for clarification of high and low expression). Student’s t-test was used to calculate p values using Microsoft Excel and are given by asterisks (****p□<□0.0001). Statistical analyses: n□=□3, error bars= ±SEM, ns= not significant, number of cells counted=200.

**A**) and western blotting (Figure 1B). The quantification of the relevant bands showed that the wildtype and mutant vimentin proteins were equally expressed in MCF-7 cells (Figure 1C). Immunostaining of the wildtype vimentin showed a fully extended vimentin network from perinuclear to peripheral cell boundaries whereas in cells transduced with the C328S mutant, the vimentin was more condensed around the perinuclear area (Figure 1D).

To confirm that C328S mutant vimentin was able to form filaments, A431 cells which are devoid of endogenous vimentin [18], were transduced with C328S-VIM. The immunostaining with mouse anti-vimentin V9 (referred as V9 from now on) followed by AF-488-labelled goat anti-mouse secondary antibody showed normal appearing polymerised IFs in A431 cells (**Figure S2A, compare a-c with d-f**). This shows that C328S-VIM retains its ability to polymerise into filament both in A431 and MCF-7 cells, however in MCF-7 these filaments were drastically reorganised affecting the cell shape (**Figure 1D**). Furthermore, to confirm that C328S-VIM does not disrupt pre-existing vimentin filaments (dominant negative), we transduced HFF-1 and MCF-7 cells expressing WT vimentin with either untagged C328S-VIM (**Figure S2B, g-l**) or AcGFP fused with C328S-VIM at its N-terminus (AcGFP-C328S-VIM) (**Figure S2C, m-r**). Cells were immunostained with V9 antibody. Our data showed that both untagged and AcGFP-C328S-VIM constructs were not dominant negative as they did not disturb the pre-existing filaments and appear to integrate into the pre-existing network. However, the limitation of using the V9 antibody was that it detected both endogenous vimentin and C328S-VIM constructs.

Next, to assess the impact of the C328S mutation on vimentin’s interaction with actin, in silico structural analysis was conducted for both the WT and mutant vimentin proteins. The Protein Data Bank (PDB) file corresponding to the wild-type vimentin (PDB ID: P08670) was obtained from the RCSB PDB database (www.rcsb.org). The specific mutation of interest, C328S, which replaces the cysteine residue at position 328 with a serine, was introduced into the WT structure and energy minimization steps were performed for best rotor fit to optimize the geometry of the mutant protein and resolve any steric clashes or unfavorable conformations resulting from the mutation. Subsequent structural simulation analyses were performed to evaluate the interactions between vimentin (both WT and C328S) and actin. These analyses focused on key binding sites, interaction energy, and potential structural changes caused by the C328S substitution. It should be noted that the thiol group of C328 enables specific hydrogen bonding and potential disulfide-mediated interactions with actin [33]. The computational results revealed that the C328S mutation induced alterations within the coiled-coil rod domain of vimentin, particularly in regions critical for actin binding. The substitution of cysteine with serine likely altered the confirmation of hydrogen bonds and hydrophobic contacts, thus weakening the overall binding affinity of the mutant vimentin for actin (**Figure 1E**).

To further test this, we investigated C328S mutation mediated structural changes in the cytoskeleton by staining these cells with Phalloidin that selectively labels F-actin. Quantification of stress fiber formation/F-actin staining pattern as described previously [34] confirmed that actin morphology and organisation was altered with low level F-actin/stress fibre staining with aggregates/fragments at the cortical margins of the cells, no stress fibres were observed in the centres of the cells in C328S-VIM expressing cells compared with high level F-actin/stress fibre staining in WT (**Figures 1F-G** and **Figure S3**). These data show that C328S-VIM when expressed in MCF-7 cells alters cell morphology through remodelling of actin cytoskeleton.

### C328S-VIM impacts cell adhesion, proliferation, migration by altering MCF-7 cell morphology

To compare the effect of C328S-VIM on the MCF-7 cell morphology (**Figure 2A**), cells were fixed and stained with CellMask^TM^ Deep Red dye and counterstained nuclei with DAPI (**Figure 2B**). Different cell morphological features, such as nuclear perimeter, cell diameter, nucleus/cell area, cell major axis, cell major axis angle, and cell minor axis, were analysed. C328S-VIM expressing MCF-7 cells showed more cellular projections when compared to WT and analysis of nuclear area, nuclear form factor, nuclear major axis, nuclear minor axis, cell area, cell diameter, cell perimeter, cell minor axis, cell major axis, and cell form factor were significantly (p<0.05) increased (**Figure 2C**), while the nucleus/cell area ratio and cell compactness were significantly decreased in cells expressing C328S-VIM in comparison to WT-VIM expressing cells (**Figure 2D**). These results indicate that expression of C328S-VIM significantly altered nuclear and cell perimeter as evident by more ruffled margins in MCF-7 cells. Next, we investigated the rate of proliferation between WT and C328S cells by colony formation (**Figure 2E, F**), MTT (**Figure 2G**) and CyQUANT proliferation assays (**Figure 2H**). The analyses showed a highly significant increase in cell proliferation and mitochondrial activity in C328S as compared to WT cells. Furthermore, C328S-VIM expressing MCF-7 cells showed reduced adhesion without substrate coating on the culture vessel as determined by CyQUANT cell adhesion assay (**Figure 2I**). However, when the tissue culture plates were coated with different substrates such as laminin, fibronectin and collagen, there was no significant difference in cell adhesion (**Figure 2J**). The chemotactic migration of the cells towards FCS through 8µm pore size culture inserts showed the number of cells migrating through the membrane was significantly higher (p<0.05) in C328S compared with the WT cells (**Figure 2K, L**). Similarly, the invasive capacity of the two cell lines was compared by coating the 8µm pore size culture inserts with Matrigel. The results indicated a significantly higher number of cells invading through Matrigel in C328S-VIM (p<0.05) compared with the WT-VIM (**Figure 2M, N**).

**Figure 2:**
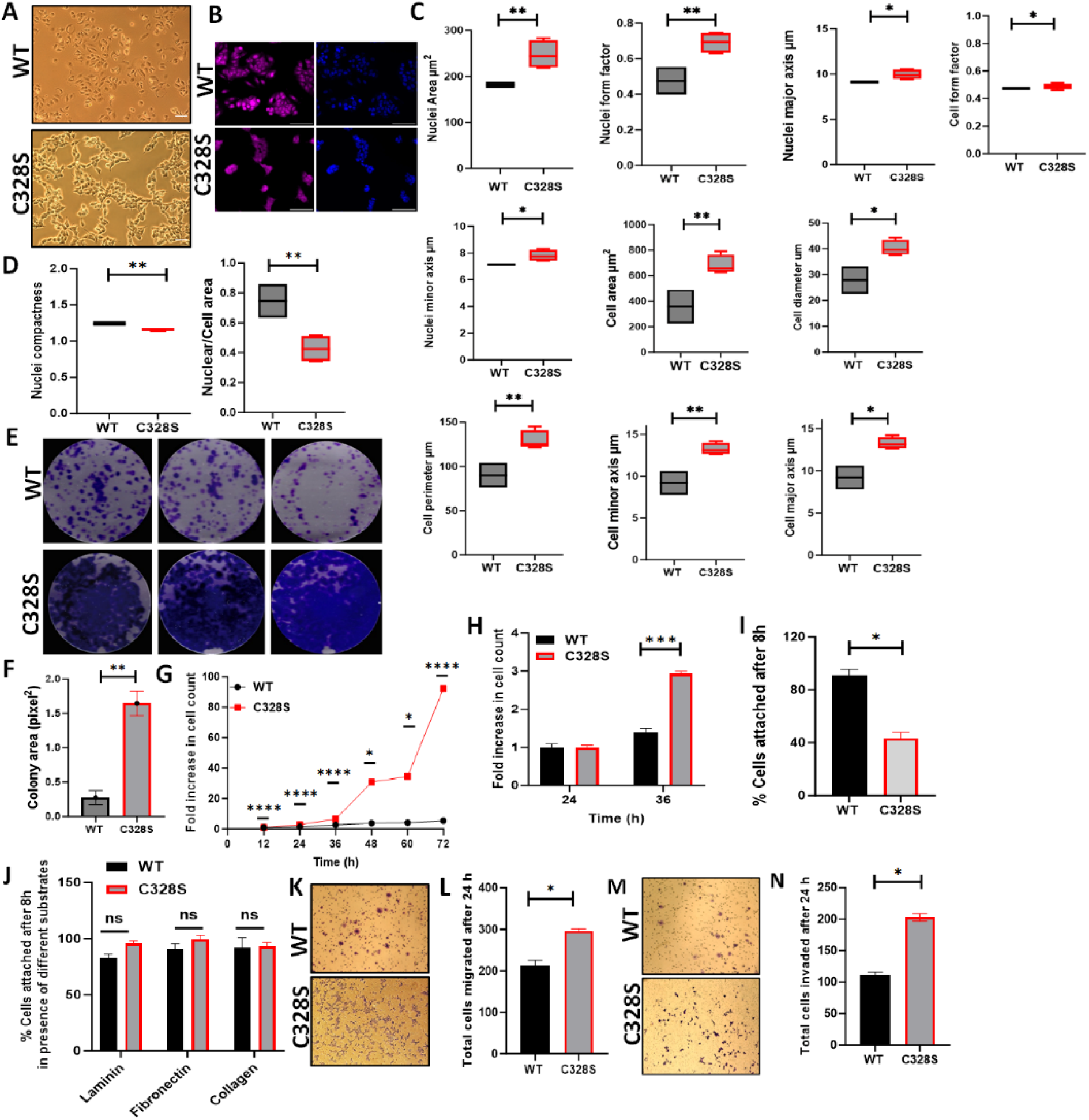
Effect of C328S-VIM on cell morphology, proliferation, adhesion and invasion. (A): Morphology of MCF-7 expressing WT-VIM and C328S-VIM in brightfield (scale bar = 100 µm). (B): Morphology of WT and C328S cells stained with CellmaskTM deep red dye. Images were captured by INCA 2200 and analysed by INCarta software (scale bar = 50 µm). (C): Differences in nuclear area, nuclei form factor, nuclear major axis, nuclear minor axis, cell area, cell diameter, cell perimeter, cell minor axis, cell major axis and cell form factor, between the two cell lines. (D): Significant reduction in nuclear compactness and nuclei/cell area between the two cell lines. Proliferation rate was compared between WT and C328S cells by (E, F) colony, (G) MTT, and (H) CyQUANT assays. (I): CyQUANT cell adhesion assay was performed to compare the cell adhesion between WT and C328S cells without substrate, and (J): with the addition of laminin, fibronectin and collagen, separately. (K): Chemotactic migration of the WT and C328S cells through 8.0 µm culture inserts. The cells were fixed and stained with 0.1% (w/v) crystal violet before imaging. (L): The cells on the outer surface of the inserts were counted and compared between WT and C328S. (M): Chemotactic invasion in WT and C328S cells through 8.0 µm culture inserts coated with Matrigel. The cells were fixed and stained with 0.1% (w/v) crystal violet. (N): Total number of cells invaded on the outer surface of the inserts were counted. Statistical analyses: n□=□3, Error bars= ± SEM, Student’s t-test was used to calculate p values using Microsoft Excel and are given by asterisks (*p□<□0.05, **p□<□0.01, ***p□<□0.001 and ****p□<□0.0001).

### Upregulation of EMT and cancer stemness related signatures by C328S-VIM in MCF-7

As C328S-VIM significantly increased the proliferation of MCF-7, reduced their adhesion capacity and increased migration and invasion towards FCS, we investigated the transcriptome profile of C328S-VIM vs WT-VIM expressing MCF-7 cells using RNA-Seq in order to understand the underlying mechanism for the changes observed. The analysis showed that a total 3421 out of 22645 genes were significantly upregulated whereas 3940 genes were downregulated (**Figure 3A**). Functional gene ontology (GO) analysis revealed that the significantly downregulated genes are involved in cell-cell adhesion, DNA packaging complex and keratinocyte differentiation (**Figure 3B**). The most upregulated cellular function was pattern specific processes, regionalisation and related to development (**Figure 3C**). The most upregulated gene was *XIST* (**Figure 3D),** a long noncoding oncogenic RNA (lncRNA) that is implicated in a large number of tumours (**Figure 3E)** [35]. The RNA-Seq data was validated using RT-qPCR and the regression analysis between two data sets showed significant Pearson correlation (R^2^ = 0.77, p=0.003, Pearson r=0.88; **Figure 3F**). The upregulated and downregulated DEGs are listed in **Table S4** and **Table S5**, respectively. **Table S6** and **Table S7** lists, respectively, the upregulated and downregulated lncRNAs. The widespread downregulation of keratin gene expression shown by RNA-Seq and qPCR was corroborated by immunostaining of K8 and K18 (**Figure 3G**) and western blotting (**Figure 3H**). The quantification of the relative band intensity showed significant downregulation in K8, K18, K19 and upregulation of TWIST1 and CDH2/N-cadherin in C328S-VIM expressing MCF-7 cells compared with the WT-VIM expressing cells (**Figure 3I**).

**Figure 3:**
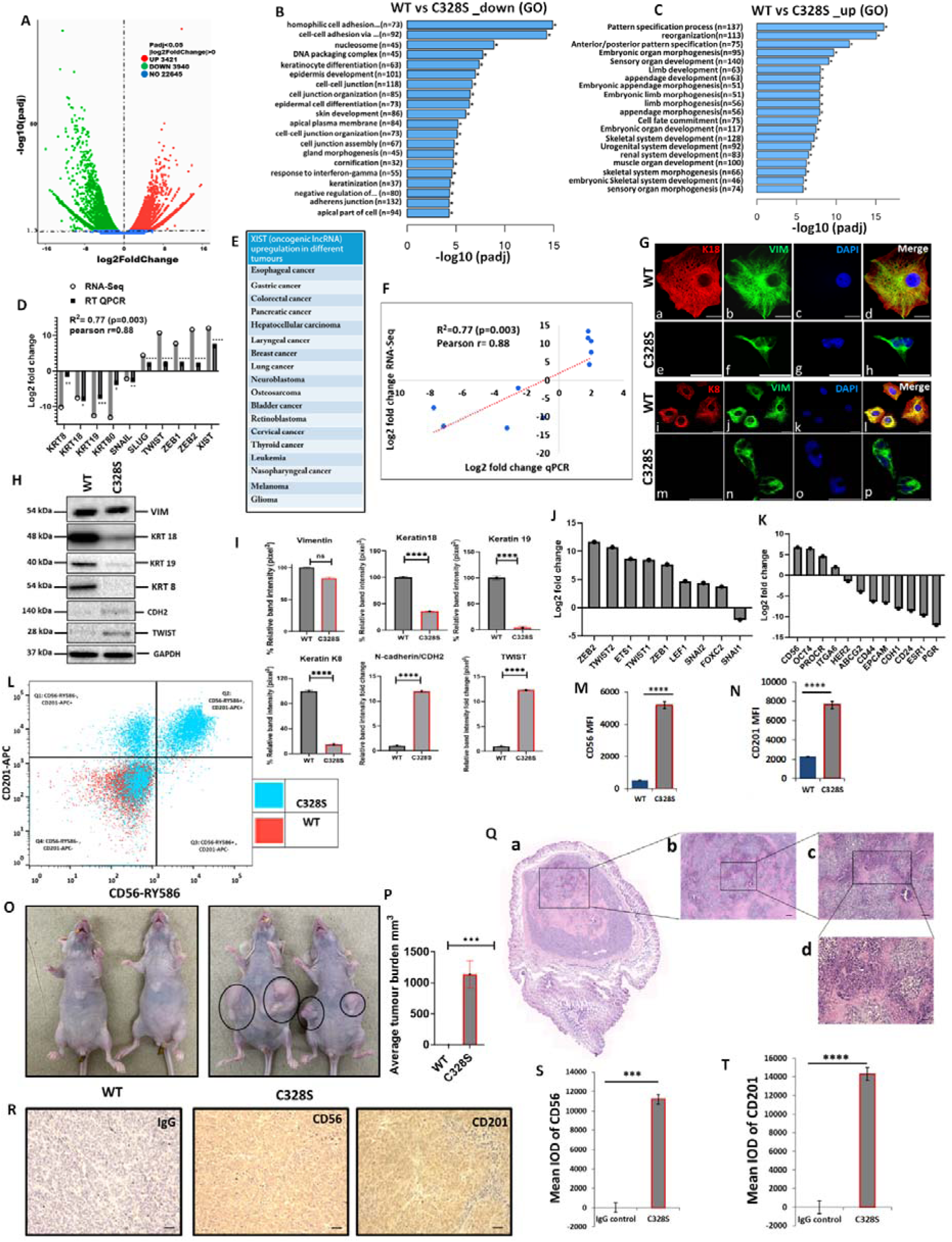
Transcriptomic insight into tumorigenic potential induced by C328S-VIM both *in vitro and in vivo*. (A): Volcano plot showing differentially expressed genes (DEGs) between WT and C328S cells. Gene Ontology (GO) showing overview of cellular functions, (B): downregulated and (C): upregulated by DEGs. (D): RNA seq analysis showing log2fold expression and validation by RT-qPCR for DEGs of interest. (E): *XIST*, the most upregulated gene, has been implicated in large number of solid tumours [35]. (F): Linear regression analysis of log2fold changes from RT-qPCR and RNA-Seq of DEGs. (G): Immunostaining of WT and C328S cells with V9, rabbit anti-K8 and rabbit anti-K18 using AF-488 (green) goat anti-mouse and AF-594 (red) goat anti-rabbit were used. Nuclei are in blue, overlapping images are shown as Merge. Leica DM4000B Epi-fluorescence microscope was used for imaging (scale bar = 20 µm). (H): VIM, K18, K19, K8, CDH2/N-cadherin and TWIST1 expression by western blotting in WT and C328S cells. Relevant bands were cropped from the original blots shown in Figure S1. (I): Quantification of the protein expression in panel H by ImageJ. (J): Relative log2fold changes in expression of EMT transcription factors, and (K): breast cancer stem cell markers in WT and C328S cells by RNA-Seq analysis. (L): Flow cytometry overlay dot plot of CD56-RY586 versus CD201-APC after gating on single and live cells for immunophenotype. (M): Comparison of mean fluorescence intensity (MFI) of CD56 in WT and C328S cells. (N): Comparison of MFI of CD201 in WT and C328S cells. (O): Transplantation of WT and C328 cells in nude mice without estrogen. (P): Average tumour burden after two weeks in nude mice injected with WT and C328S cells. (Q): H & E stained tumour sections scale bar= 50 µm. (R): Representative image from immunohistochemical staining of CD56 and CD201 in tumour sections as compared with IgG control, scale bar= 50 µm. (S): Quantification of CD56 and (T): CD201staining in tumour sections and control using ImageJ. Statistical analyses: n□=□3, Error bars= ± SEM, Student’s t-test to calculate p values using Microsoft Excel, and are given as asterisks (*p□<□0.05, **p□<□0.01, and ***p□<□0.001, ****p□<□0.0001).

The most upregulated and downregulated cellular functions in GO analyses are listed in **Figure S4** and **S5**, respectively. The KEGG pathway analysis is shown in **Figure S6**. The RNA-Seq analysis shows that multiple epithelial markers were downregulated, and mesenchymal markers were upregulated among DEGs indicating acquisition of EMT-like characteristics by C328S cells (**Figure S7**). Taken together, the data from RNA-Seq experiments begin to provide a mechanistic explanation for the structural and functional properties that we found to be associated with the C328S mutation in vimentin.

The RNA-Seq data showed upregulation of EMT transcription factors including *ZEB1, ZEB2, TWIST 1, TWIST 2, ETS1, LEF1, SNAI2*, and *FOXC2*, although *SNAI1* was downregulated (**Figure 3J**). The upregulated differentially expressed breast cancer stem cells markers included *CD56/ NCAM1, OCT4, PROCR/CD201, ITGA6*, and the downregulated were *ESR1, PGR, HER2/ERBB2, ABCG2, CD44, CD24, EPCAM* and *CDH1* (**Figure 3K**). Out of these presence or absence of three of them *ESR1* (estrogen receptor), *PGR* (progesterone receptor) and *HER2/ERBB2* (human epidermal growth factor receptor 2/HER2) makes a breast cancer triple positive or triple negative. Our results show that MCF-7 which are triple positive cells [36] became triple reduced by the expression of C328S-VIM. Collectively these results imply that C328S-VIM in MCF-7 induces EMT-like characteristics and increases cancer stemness.

To confirm the expression of breast cancer stem cell markers identified as being differentially expressed in RNA-Seq analysis, FACS analysis was performed. Cells were stained for surface antigens with RY586–conjugated antibody specific for CD56/NCAM1, APC-conjugated antibody specific for CD201/PROCR and the viability stain FVS575V was used for detecting the live cells. Gates were applied for cells (SSC-A:FSC-A), single cells (SSC-W: FSC-A) and live cells (FVS575V::L/d:FSC-A) for WT and C328S cells (**Figure S8**). Data analysis using *FlowJo v10* confirmed significant upregulation of breast cancer stem cell markers CD56/NCAM1 and CD201/PROCR in C328S-VIM expressing MCF-7 cells. MFI of CD56 and CD201 in C328S-VIM cells was significantly higher (p<0.0001) compared with WT-VIM cells (**Figure 3L, M, N**).

Next, to examine tumorigenic potential *in vivo*, we injected WT and C328S cells subcutaneously and bilaterally into the flanks of 6-week-old female athymic nu/nu mice. C328S cells were able to produce tumours without oestrogen however WT cells did not produce any tumour (**Figure 3O, P**). These results confirm the tumorigenic potential of C328S-VIM that is oestrogen independent. Furthermore to investigate breast cancer stem cell markers expression, routine H & E (**Figure 3Q**) and immunohistochemistry (**Figure 3R**) was performed on the tumour tissue sections for CD56 and CD201 expression, respectively. Normal mouse IgG was used as staining control. Tumour sections from C328S injected mice showed significantly higher IOD for CD56 (p<0.001) (**Figure 3S**) and CD201 (p<0.0001) (**Figure 3T**) as compared with IgG control. These results confirmed our *in-vitro* transcriptome and FACS analyses data.

To establish the specificity of tumour production by C328S-VIM we created another single substitution, Y117L, at the beginning of the rod domain and a double substitution containing C328S and Y117L in the same vimentin molecule. Y117L substitution has been reported to assembles into ULFs [14] and does not form long filaments. Interestingly transplantation of MCF-7Y117L-VIM did not induce tumour formation in nude mice whereas MCF-7DMT-VIM did produce tumours. **(Figure S8)**. These experiments suggest that tumour progession by C328S was specific and in DMT it was dominant over Y117L.

### shRNA mediated downregulation of mutant vimentin or *XIS*T in C328S-VIM expressing cells inhibits cancer potential

To determine whether the effects of C328S-VIM in MCF-7 can be reversed, we downregulated mutant vimentin or *XIST* in C328S-VIM cells by shRNA to more than 90 and 75%, respectively, as determined by RT-qPCR (p<0.01) (**Figure 4A and B)** and western blotting (p<0.001) **(Figure 4C and D; Figure S9)** compared with NTC. *XIST* RNA levels were reduced by 30% upon vimentin knockdown and VIM mRNA was downregulated by up to 20% (p<0.05) upon *XIST* knockdown (**Figure 4E and F)**. CyQUANT adhesion assay showed that adhesion capacity of cells treated with *VIM-*shRNA was significantly increased (p<0.001) (up to 54%±1.9) but not much with *XIST* shRNA when compared to NTC in C328S-VIM cells (**Figure 4G and H)**. Furthermore, cell proliferation (**Figure 4I and J)**, migration (**Figure 4K and L)** and invasion (**Figure 4M and N)** were also decreased but the results were significant (p<0.05) only for the invasion assay. Morphological analysis of MCF-7C328S-shVIM cells showed that several features (not all, see **Figure S10**) including changes in cell area (p<0.01), nuclear/cell area (p<0.01), cell diameter (p<0.01), cell perimeter (p<0.01), cell major axis (p<0.05) and cell minor axis (p<0.05) were significantly reversed (**Figure 4O)**. These results show that C328S-VIM is primarily responsible for the loss of adhesion and altered morphology of these cells. We also investigated the effect of mutant vimentin downregulation on breast cancer stem cell markers CD56/NCAM1 and CD201/PROCR in C328S-VIM expressing cells upon shRNA treatment. Flow cytometry analysis of CD56 and CD201 expression upon shRNA treatment in these cells showed insignificant differences (**Figure 4P, 4Q, Figure S11-12**) demonstrating that expression of cancer stem cell markers was irreversible.

**Figure 4:**
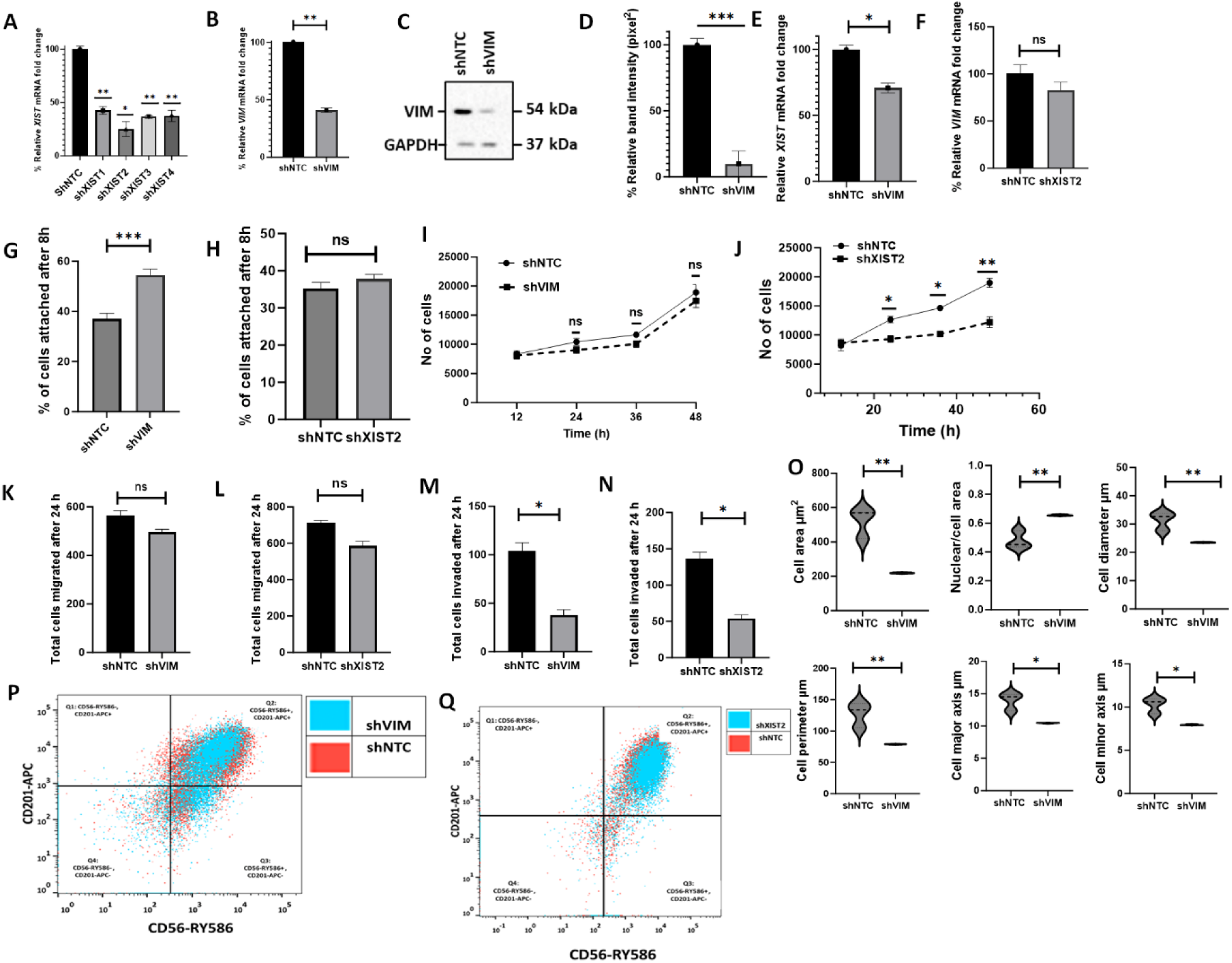
shRNA mediated downregulation of *XIST* in C328S-VIM reverts cell phenotype. (A): Downregulation of *XIST* in C328S cells by 4 different shRNAs (sh1-sh4) for *XIST* or NTC by RT-qPCR. sh*XIST*2 was the most potent (p < 0.05) compared with NTC. (B): VIM expression in MCF-7C328S_shVIM and MCF-7C328S_shNTC as determined by RT-qPCR. (C): Vimentin expression in MCF-7C328S_shVIM and MCF-7C328S_shNTC by western blotting. (D): Quantification of the protein expression in panel C by ImageJ. (E): *XIST* RNA in MCF-7 cells expressing C328S_shVIM and C328S_shNTC by RT-qPCR. (F): Relative VIM mRNA fold change (%) in MCF-7 cells expressing C328S_sh*XIST*2 and C328S_shNTC by RT-qPCR. Comparision of cell adhesion, (G): between MCF-7C328S_shVIM and MCF-7C328S_shNTC, and (H): between MCF-7C328S_sh*XIST*2 and MCF-7C328S_shNTC cells without substrate by CyQUANT assay. Comparision of cell proliferation, (I): between MCF-7C328S_shVIM and MCF-7C328S_shNTC, and (J): between MCF-7C328S_sh*XIST*2 and MCF-7C328S_shNTC by MTT assay. Comparision of chemotactic migration, (K): between MCF-7C328S_shVIM and MCF-7C328S_shNTC, and (L): between MCF-7C328S_sh*XIST*2 and MCF-7C328S_shNTC cells through 8.0 µm culture inserts. Comparision of chemotactic invasion, (M): between MCF-7C328S_shVIM and MCF-7C328S_shNTC, and (N): between MCF-7C328S_sh*XIST*2 and MCF-7C328S_shNTC cells through 8.0 µm Matrigel coated inserts. (O): Comparison of cell area, ratio of nuclei/cell area, cell diameter, cell perimeter, cell major axis, cell minor axis, between MCF-7C328S_shVIM and MCF-7C328S_shNTC. (P): Flow cytometry overlay dot plot of CD56-RY586 versus CD201-APC after gating on single and live cells for immunophenotyping of MCF-7C328S_shVIM and MCF-7C328S_shNTC cells. (Q): Flow cytometry overlay dot plot of CD56-RY586 versus CD201-APC after gating on single and live cells for immunophenotyping of MCF-7C328S_sh*XIST*2 and MCF-7C328S_shNTC cells. Statistical analyses: n□=□3, Error bars= ± SEM, Student’s t-test was used to calculate p values using Microsoft Excel when two groups were compared, one way ANOVA with Bonferroni test was applied using GraphPad Prism 10 when comparing more than two groups, (*p□<□0.05, **p□<□0.01 and ***P<0.001).

## Discussion

Vimentin is a key player in several pathophysiological processes such as cell migration, proliferation, adhesion, stress response, EMT and cancer metastasis [3]. It has a single cysteine residue at position 328 that is considered a preferred site for posttranslational modifications, and it is currently being intensely investigated due to its involvement in multiple cell functions, including filament assembly, cell polarity, elongation, and stress response to electrophiles and oxidants [17]. In spite of its likely importance in cellular physiology, the role of C328 in cancer progression and in determining the cancer stem cell phenotype has not been investigated before.

To investigate the significance of C328 in EMT and cancer, we made a mutant encoding a C328S substitution in vimentin (C328S-VIM) and transduced in MCF-7 cells, which are devoid of endogenous vimentin [18, 37, 38], and are widely used as a model to study EMT [19, 20]. Structurally, the two amino acids, cysteine and serine, are similar except an electronegative sulphur atom (atomic radius 0.88 A°) in cysteine was replaced by another electronegative oxygen atom (atomic radius 0.48 A°) in serine. Although both atoms are electronegative, oxygen is slightly more electronegative (EN=3.44) than sulphur (EN=2.58). These seemingly minor differences were unlikely to cause major structural perturbations, so we were surprised when *in silico* simulations demonstrated that C328S substitution was likely to affect interactions between actin and vimentin. These computational predictions were confirmed by our immunofluorescence analyses of F-actin in C328S cells. Whereas WT vimentin-expressing cells showed normal stress fibres with no fragments or aggregates, in contrast C328S-VIM cells expressed aggregated and fragmented F-actin which was limited to the cortical margins of the cells with no stress fibre formation. These observations highlight the critical interactions of the rod domain of vimentin at C328 with actin that has never been reported, although earlier studies have demonstrated direct binding of the tail domain of vimentin with actin [15]. It has recently been reported that C328 of vimentin modifies actin organisation in response to oxidants and electrophiles [33]. Our data support these findings and suggest that a local perturbation at C328 (in this case by C328S substitution) can disrupt actin organisation even when cells are not exposed to oxidants and electrophiles. These observations have widespread implications including cytoskeletal crosstalk that regulates cell behaviour especially during cancer development, progression and spread.

Our data show that the C328S mutation in vimentin can change the overall morphology of MCF-7 cells (**Figure 2**). Earlier studies have implicated vimentin filaments in the maintenance of cell shape and increasing vimentin levels are positively related to a mesenchymal elongated phenotype and EMT [39]. In contrast, we have recently shown that ectopic expression of the wildtype vimentin in MCF-7 can make them less elongated [18]. A possible explanation for these conflicting reports may be that cancer cells can express a wide spectrum of morphologies depending upon inducing factor and stage of EMT [40, 41]. As the C328S substitution in vimentin has increased the nuclear size and made the MCF-7 cells more rounded compared to WT vimentin, it is possible that the C328 residue may be an active player in vimentin-linked cell phenotypic changes since downregulation of the mutant vimentin significantly reversed the morphological changes (**Figure 4**).

We have previously shown that expression of WT vimentin in MCF-7 induces cell migration without affecting proliferation or cell adhesion [18]. Here we show that C328S-VIM enhances cell proliferation, migration, invasion and reduces cell adhesion, which are features closely associated with EMT mediated cancer metastasis. Some of these changes could be reversed by RNAi mediated downregulation of vimentin establishing a direct link between residue 328 of vimentin, or perhaps the surrounding region, in cancer progression, which is a novel finding. The increase in malignant characteristics of C328S cells imply that the original cysteine residue in vimentin acts as tumour suppressor. In addition, our RNA-Seq, RT-qPCR and western blot analyses showed upregulation of EMT-associated transcription factors (ZEB1, SNAI2, LEF1, ZEB2, TWIST1, TWIST2) and mesenchymal markers (CDH2, MMP2, FOXC2, FNDC1), as well as downregulation of epithelial markers (cytokeratins, CDH1, CLDN1, EPCAM), indicating possible EMT induction in C328S cells. There are 3 major keratins including K8 (type II), K18 and K19 (both type I) that are expressed in MCF-7 and since all of them were downregulated (**Figure 3**), a central mechanism involved in the expression of all the three keratins is likely to be affected by C328-VIM. One such mechanism is the involvement of transcription factor AP-1, which participates in the regulation of most keratin genes [42]. AP-1 is a complex of multiple different subunits [43] but our transcriptomic analysis suggests that none of these are affected by C328S-VIM (DEGs lists provided in supplementary material). It is therefore conceivable that C328S-VIM could suppress AP-1 activity, thereby switching off keratin gene expression. Further investigations are required to test this hypothesis.

Multiple breast cancer stem cell markers such as *OCT4* [44], *CD56* [45], *CD49f* [46], and *CD201/PROCR* [47] were also upregulated by C328-VIM, demonstrating increased stemness characteristics in MCF-7 cells as judged by the RNA-Seq analysis. Higher expression of the two stem cell markers, CD56 and CD201, was further validated by flow cytometry, and also observed *in vivo*. RNA-Seq analysis also showed that MCF-7, which is a triple positive (expression of ESR1, PGR and HER2) cell line, became triple reduced in the presence of the C328S-VIM. Breast oncologists broadly stratify breast cancers into triple positive, borderline and triple negative [48] which is indicative of their susceptibility to metastasise and their clinical course. Triple positive breast cancers have a relatively good prognosis, followed by borderline and triple negative lesions have by far the worst prognosis [48]. We have previously shown that wildtype vimentin does not affect expression of these receptors in MCF-7 [18]. In the present study, however, C328S-VIM makes MCF-7, a triple positive cell line [36], into triple reduced which has important clinical implications. It also suggests that a mechanism exists whereby disruption of molecular interactions between vimentin and actin caused by C328S, directly or indirectly, regulates expression of these receptors, cancer progression and metastasis.

The gene most upregulated by the C328S vimentin, as found in the RNA-Seq analysis, which is a long noncoding RNA *XIST* is known to induce proliferation, invasion and inhibit apoptosis in breast cancer [49]. It is also reported to be upregulated in multiple solid tumours [35, 50]. A recent study has unveiled its gatekeeping role in human mammary epithelium homeostasis especially differentiation aspect in human mammary stem cells (MaSCs) [51]. It therefore appears, based on our observations, that vimentin C328S mutation is inducing EMT-like changes via *XIST* upregulation, as downregulation of *XIST* in C328S cells, significantly reduced proliferation and invasion.

A highly significant and interesting observation was that presence of C328S-VIM in MCF-7, which are oestrogen-dependent tumorigenic cells [21], made these cells oestrogen-independent in nude mice that further supports our *in-vitro* data that C328S substitution had enhanced cancer stemness in MCF-7 cells. Expression of breast cancer stem cell markers CD56 and CD201 was evident in mouse tumours. These data suggest that the C328 in vimentin is an important regulator of cancer cell behaviour and that alterations in this region of the protein promotes EMT-like changes and may induce metastasis.

Recently, wildtype vimentin has been shown to increase malignant progression in lung cancer. However, our data suggest that wildtype vimentin is non-tumorigenic in breast cancer, inferring that effect of vimentin in cancer may be tissue specific, which has never been reported [52]. Previous studies have described the importance of C328 residue in filament assembly and organisation, stress response, aggresome formation, and lysosomal positioning only in SW13/cl.2 vimentin-deficient cells [16, 17]. However, the use of SW13/cl.2 cell model may not be appropriate to study the role of C328-VIM in EMT because these cells do not express any IFs. For a cancer cell to undergo EMT it must be an epithelial cancer cell expressing keratins. In that respect use of MCF-7 in this study is most approaprite because it does not express endogenous vimentin. As mutations in vimentin have been reported in patients around the area of C328 [1], our hypothesis that this mutation induces conformational changes that activate the complex processes of EMT in breast cancer cells, would have clinical implications.

## Conclusion

In summary, this study highlights that vimentin is no longer a mere marker of EMT and metastasis, but appears to be an active participant in cancer progression. The strong correlation between the single cysteine residue at position 328 in vimentin with actin organisation, *XIST* induction and hyperproliferation associated with breast oncogenesis is novel. This suggests that C328 in vimentin remodels actin cytoskeleton and protects against EMT and cancer growth via modulating lncRNA, *XIST.* Taken together, we propose that targeting vimentin via RNA interference should be considered a therapeutic strategy for breast cancer treatment.

## Declarations

### Ethical approval

Transplantation experiments were performed in this study according to animal welfare regulations and were approved by Augusta University under Institutional Animal Care and Use Committee (IACUC) protocol number 2015-0736.

### Authors’ contributions

SU, HT and AW conceived the idea, drafted the manuscript, and prepared the figures. SU, WAY performed the experiments and collected the data. MTT, WAY, HT, FG and AW revised and edited the manuscript. All authors have read and agreed to the submitted version of this manuscript.

### Availability of data and materials

All data generated including uncropped blot images to support the findings of this study are available within this manuscript and its supplementary information files.

## Acknowledgments

The authors are thankful to the Centre of Oral Immunobiology and Regenerative Medicine and the Blizard Institute for proving the research facilities necessary for this work. We are also thankful to Luke Gammon for help with the use of IN Cell Analyzer and Gary Warnes for FACS analysis. The authors also thank Professors Farida Fortune and Ian Mackenzie for useful discussion. We are also thankful to Mr Usman Baig for helping with figures.

