## Supplementary material for "A single cysteine residue in vimentin regulates long non-coding RNA *XIST* to suppress epithelial-mesenchymal transition and stemness in breast cancer": ATRIAL_SEPTUM_DEVELOPMENT(GO_0003283).html

Details for gene set ATRIAL\_SEPTUM\_DEVELOPMENT(GO:0003283)[GSEA]


|  || Dataset | fpkm.sample |
| Phenotype | sample.cls |
| Upregulated in class | C328SVIM |
| GeneSet | ATRIAL\_SEPTUM\_DEVELOPMENT(GO:0003283) |
| Enrichment Score (ES) | 0.89457643 |
| Normalized Enrichment Score (NES) | 1.4312274 |
| Nominal p-value | 0.0 |
| FDR q-value | 1.0 |
| FWER p-Value | 0.325 |
Table: GSEA Results Summary

  

Fig 1: Enrichment plot: ATRIAL\_SEPTUM\_DEVELOPMENT(GO:0003283)
  
Profile of the Running ES Score & Positions of GeneSet Members on the Rank Ordered List

  

| PROBE | DESCRIPTION   (from dataset) | GENE SYMBOL | GENE\_TITLE | RANK IN GENE LIST | RANK METRIC SCORE | RUNNING ES | CORE ENRICHMENT || 1 | ENSG00000183072 | NKX2-5 |  |  | 1494 | Infinity | -0.0253 | Yes |
| 2 | ENSG00000265107 | GJA5 |  |  | 1553 | Infinity | -0.0262 | Yes |
| 3 | ENSG00000128602 | SMO |  |  | 5833 | 1985.988 | 0.6037 | Yes |
| 4 | ENSG00000145362 | ANK2 |  |  | 5882 | 409.285 | 0.7477 | Yes |
| 5 | ENSG00000164532 | TBX20 |  |  | 5925 | 252.081 | 0.8361 | Yes |
| 6 | ENSG00000016082 | ISL1 |  |  | 5958 | 166.898 | 0.8946 | Yes |
| 7 | ENSG00000113971 | NPHP3 |  |  | 10660 | 2.610 | 0.8160 | No |
| 8 | ENSG00000136574 | GATA4 |  |  | 12258 | 1.997 | 0.7897 | No |
| 9 | ENSG00000179284 | DAND5 |  |  | 12524 | 1.915 | 0.7859 | No |
| 10 | ENSG00000135679 | MDM2 |  |  | 15941 | 1.240 | 0.7285 | No |
| 11 | ENSG00000204217 | BMPR2 |  |  | 17127 | 1.083 | 0.7089 | No |
| 12 | ENSG00000142871 | CYR61 |  |  | 17448 | 1.042 | 0.7038 | No |
| 13 | ENSG00000109685 | NSD2 |  |  | 18004 | 0.974 | 0.6948 | No |
| 14 | ENSG00000115170 | ACVR1 |  |  | 20567 | 0.736 | 0.6517 | No |
| 15 | ENSG00000124766 | SOX4 |  |  | 21256 | 0.665 | 0.6403 | No |
| 16 | ENSG00000134250 | NOTCH2 |  |  | 23322 | 0.493 | 0.6055 | No |
| 17 | ENSG00000135547 | HEY2 |  |  | 23743 | 0.461 | 0.5986 | No |
| 18 | ENSG00000179588 | ZFPM1 |  |  | 28202 | 0.102 | 0.5232 | No |
| 19 | ENSG00000092969 | TGFB2 |  |  | 28951 | 0.037 | 0.5106 | No |
| 20 | ENSG00000089225 | TBX5 |  |  | 31935 | 0.000 | 0.4601 | No |
Table: GSEA details
[plain text format]

  

Fig 2: ATRIAL\_SEPTUM\_DEVELOPMENT(GO:0003283)
  
Blue-Pink O' Gram in the Space of the Analyzed GeneSet

  

Fig 3: ATRIAL\_SEPTUM\_DEVELOPMENT(GO:0003283): Random ES distribution
  
Gene set null distribution of ES for
 **ATRIAL\_SEPTUM\_DEVELOPMENT(GO:0003283)**

  
