## Supplementary material for "A single cysteine residue in vimentin regulates long non-coding RNA *XIST* to suppress epithelial-mesenchymal transition and stemness in breast cancer": CELLULAR_RESPONSE_TO_INTERLEUKIN_12(GO_0071349).html

Details for gene set CELLULAR\_RESPONSE\_TO\_INTERLEUKIN\_12(GO:0071349)[GSEA]


|  || Dataset | fpkm.sample |
| Phenotype | sample.cls |
| Upregulated in class | C328SVIM |
| GeneSet | CELLULAR\_RESPONSE\_TO\_INTERLEUKIN\_12(GO:0071349) |
| Enrichment Score (ES) | 0.8929124 |
| Normalized Enrichment Score (NES) | 1.4470495 |
| Nominal p-value | 0.0 |
| FDR q-value | 1.0 |
| FWER p-Value | 0.325 |
Table: GSEA Results Summary

  

Fig 1: Enrichment plot: CELLULAR\_RESPONSE\_TO\_INTERLEUKIN\_12(GO:0071349)
  
Profile of the Running ES Score & Positions of GeneSet Members on the Rank Ordered List

  

| PROBE | DESCRIPTION   (from dataset) | GENE SYMBOL | GENE\_TITLE | RANK IN GENE LIST | RANK METRIC SCORE | RUNNING ES | CORE ENRICHMENT || 1 | ENSG00000244067 | GSTA2 |  |  | 2473 | Infinity | -0.0419 | Yes |
| 2 | ENSG00000113302 | IL12B |  |  | 4342 | Infinity | -0.0735 | Yes |
| 3 | ENSG00000285441 | SOD2 |  |  | 4837 | Infinity | -0.0818 | Yes |
| 4 | ENSG00000197632 | SERPINB2 |  |  | 5773 | Infinity | -0.0976 | Yes |
| 5 | ENSG00000147065 | MSN |  |  | 5823 | 15136.710 | 0.8897 | Yes |
| 6 | ENSG00000081985 | IL12RB2 |  |  | 6113 | 66.201 | 0.8891 | Yes |
| 7 | ENSG00000099810 | MTAP |  |  | 6128 | 61.949 | 0.8929 | Yes |
| 8 | ENSG00000168811 | IL12A |  |  | 6738 | 14.720 | 0.8836 | No |
| 9 | ENSG00000138378 | STAT4 |  |  | 7448 | 7.743 | 0.8721 | No |
| 10 | ENSG00000120438 | TCP1 |  |  | 10138 | 2.914 | 0.8268 | No |
| 11 | ENSG00000152795 | HNRNPDL |  |  | 11925 | 2.108 | 0.7967 | No |
| 12 | ENSG00000122566 | HNRNPA2B1 |  |  | 12376 | 1.964 | 0.7892 | No |
| 13 | ENSG00000116489 | CAPZA1 |  |  | 14453 | 1.486 | 0.7541 | No |
| 14 | ENSG00000070831 | CDC42 |  |  | 14664 | 1.457 | 0.7507 | No |
| 15 | ENSG00000148834 | GSTO1 |  |  | 14745 | 1.437 | 0.7494 | No |
| 16 | ENSG00000131876 | SNRPA1 |  |  | 14749 | 1.436 | 0.7495 | No |
| 17 | ENSG00000113368 | LMNB1 |  |  | 14801 | 1.427 | 0.7487 | No |
| 18 | ENSG00000142168 | SOD1 |  |  | 14948 | 1.400 | 0.7463 | No |
| 19 | ENSG00000006451 | RALA |  |  | 15266 | 1.342 | 0.7410 | No |
| 20 | ENSG00000174238 | PITPNA |  |  | 16755 | 1.132 | 0.7159 | No |
| 21 | ENSG00000150593 | PDCD4 |  |  | 16766 | 1.131 | 0.7158 | No |
| 22 | ENSG00000196262 | PPIA |  |  | 16939 | 1.112 | 0.7130 | No |
| 23 | ENSG00000127314 | RAP1B |  |  | 17393 | 1.049 | 0.7054 | No |
| 24 | ENSG00000182621 | PLCB1 |  |  | 18178 | 0.952 | 0.6922 | No |
| 25 | ENSG00000113013 | HSPA9 |  |  | 18525 | 0.914 | 0.6864 | No |
| 26 | ENSG00000162434 | JAK1 |  |  | 19242 | 0.836 | 0.6743 | No |
| 27 | ENSG00000143761 | ARF1 |  |  | 19660 | 0.807 | 0.6673 | No |
| 28 | ENSG00000105397 | TYK2 |  |  | 21073 | 0.682 | 0.6435 | No |
| 29 | ENSG00000177156 | TALDO1 |  |  | 21372 | 0.653 | 0.6385 | No |
| 30 | ENSG00000169813 | HNRNPF |  |  | 22036 | 0.595 | 0.6273 | No |
| 31 | ENSG00000089157 | RPLP0 |  |  | 23458 | 0.483 | 0.6033 | No |
| 32 | ENSG00000096968 | JAK2 |  |  | 23558 | 0.475 | 0.6016 | No |
| 33 | ENSG00000180370 | PAK2 |  |  | 24306 | 0.414 | 0.5890 | No |
| 34 | ENSG00000185624 | P4HB |  |  | 24505 | 0.402 | 0.5857 | No |
| 35 | ENSG00000110711 | AIP |  |  | 24785 | 0.379 | 0.5810 | No |
| 36 | ENSG00000183336 | BOLA2 |  |  | 24933 | 0.369 | 0.5785 | No |
| 37 | ENSG00000100911 | PSME2 |  |  | 25048 | 0.361 | 0.5766 | No |
| 38 | ENSG00000172757 | CFL1 |  |  | 25558 | 0.322 | 0.5680 | No |
| 39 | ENSG00000064666 | CNN2 |  |  | 26063 | 0.280 | 0.5595 | No |
| 40 | ENSG00000169627 | BOLA2B |  |  | 26235 | 0.269 | 0.5567 | No |
| 41 | ENSG00000182718 | ANXA2 |  |  | 27588 | 0.156 | 0.5338 | No |
| 42 | ENSG00000136167 | LCP1 |  |  | 28472 | 0.078 | 0.5188 | No |
| 43 | ENSG00000096996 | IL12RB1 |  |  | 31266 | 0.000 | 0.4716 | No |
| 44 | ENSG00000133742 | CA1 |  |  | 35559 | NaN | 0.3989 | No |
| 45 | ENSG00000240972 | MIF |  |  | 39917 | NaN | 0.3252 | No |
| 46 | ENSG00000136634 | IL10 |  |  | 48582 | NaN | 0.1786 | No |
| 47 | ENSG00000111537 | IFNG |  |  | 56672 | NaN | 0.0417 | No |
Table: GSEA details
[plain text format]

  

Fig 2: CELLULAR\_RESPONSE\_TO\_INTERLEUKIN\_12(GO:0071349)
  
Blue-Pink O' Gram in the Space of the Analyzed GeneSet

  

Fig 3: CELLULAR\_RESPONSE\_TO\_INTERLEUKIN\_12(GO:0071349): Random ES distribution
  
Gene set null distribution of ES for
 **CELLULAR\_RESPONSE\_TO\_INTERLEUKIN\_12(GO:0071349)**

  
