## Supplementary material for "A single cysteine residue in vimentin regulates long non-coding RNA *XIST* to suppress epithelial-mesenchymal transition and stemness in breast cancer": CEREBELLAR_CORTEX_MORPHOGENESIS(GO_0021696).html

Details for gene set CEREBELLAR\_CORTEX\_MORPHOGENESIS(GO:0021696)[GSEA]


|  || Dataset | fpkm.sample |
| Phenotype | sample.cls |
| Upregulated in class | C328SVIM |
| GeneSet | CEREBELLAR\_CORTEX\_MORPHOGENESIS(GO:0021696) |
| Enrichment Score (ES) | 0.8789515 |
| Normalized Enrichment Score (NES) | 1.4313596 |
| Nominal p-value | 0.0 |
| FDR q-value | 1.0 |
| FWER p-Value | 0.325 |
Table: GSEA Results Summary

  

Fig 1: Enrichment plot: CEREBELLAR\_CORTEX\_MORPHOGENESIS(GO:0021696)
  
Profile of the Running ES Score & Positions of GeneSet Members on the Rank Ordered List

  

| PROBE | DESCRIPTION   (from dataset) | GENE SYMBOL | GENE\_TITLE | RANK IN GENE LIST | RANK METRIC SCORE | RUNNING ES | CORE ENRICHMENT || 1 | ENSG00000152208 | GRID2 |  |  | 546 | Infinity | -0.0092 | Yes |
| 2 | ENSG00000117707 | PROX1 |  |  | 4531 | Infinity | -0.0766 | Yes |
| 3 | ENSG00000102924 | CBLN1 |  |  | 4642 | Infinity | -0.0785 | Yes |
| 4 | ENSG00000111783 | RFX4 |  |  | 5022 | Infinity | -0.0849 | Yes |
| 5 | ENSG00000128602 | SMO |  |  | 5833 | 1985.988 | 0.7884 | Yes |
| 6 | ENSG00000074047 | GLI2 |  |  | 5943 | 206.951 | 0.8790 | Yes |
| 7 | ENSG00000069667 | RORA |  |  | 7414 | 7.887 | 0.8576 | No |
| 8 | ENSG00000111087 | GLI1 |  |  | 7537 | 7.217 | 0.8588 | No |
| 9 | ENSG00000135919 | SERPINE2 |  |  | 7546 | 7.160 | 0.8618 | No |
| 10 | ENSG00000135049 | AGTPBP1 |  |  | 9481 | 3.416 | 0.8306 | No |
| 11 | ENSG00000273706 | LHX1 |  |  | 9951 | 3.035 | 0.8240 | No |
| 12 | ENSG00000184524 | CEND1 |  |  | 11496 | 2.276 | 0.7989 | No |
| 13 | ENSG00000118193 | KIF14 |  |  | 11858 | 2.135 | 0.7938 | No |
| 14 | ENSG00000165240 | ATP7A |  |  | 12090 | 2.050 | 0.7908 | No |
| 15 | ENSG00000138311 | ZNF365 |  |  | 13686 | 1.631 | 0.7645 | No |
| 16 | ENSG00000141837 | CACNA1A |  |  | 13932 | 1.585 | 0.7611 | No |
| 17 | ENSG00000179295 | PTPN11 |  |  | 15435 | 1.312 | 0.7363 | No |
| 18 | ENSG00000105887 | MTPN |  |  | 17738 | 1.007 | 0.6978 | No |
| 19 | ENSG00000253368 | TRNP1 |  |  | 18437 | 0.923 | 0.6864 | No |
| 20 | ENSG00000164885 | CDK5 |  |  | 20653 | 0.726 | 0.6492 | No |
| 21 | ENSG00000169032 | MAP2K1 |  |  | 20668 | 0.724 | 0.6493 | No |
| 22 | ENSG00000123815 | COQ8B |  |  | 21103 | 0.679 | 0.6423 | No |
| 23 | ENSG00000103657 | HERC1 |  |  | 23224 | 0.502 | 0.6066 | No |
| 24 | ENSG00000198728 | LDB1 |  |  | 23260 | 0.499 | 0.6063 | No |
| 25 | ENSG00000173898 | SPTBN2 |  |  | 24458 | 0.405 | 0.5862 | No |
| 26 | ENSG00000171798 | KNDC1 |  |  | 25241 | 0.348 | 0.5731 | No |
| 27 | ENSG00000079482 | OPHN1 |  |  | 25299 | 0.343 | 0.5723 | No |
| 28 | ENSG00000135472 | FAIM2 |  |  | 27391 | 0.171 | 0.5370 | No |
| 29 | ENSG00000154764 | WNT7A |  |  | 33251 | NaN | 0.4379 | No |
| 30 | ENSG00000089116 | LHX5 |  |  | 40770 | NaN | 0.3107 | No |
| 31 | ENSG00000198719 | DLL1 |  |  | 56299 | NaN | 0.0479 | No |
| 32 | ENSG00000179915 | NRXN1 |  |  | 58268 | NaN | 0.0146 | No |
| 33 | ENSG00000215474 | SKOR2 |  |  | 59082 | NaN | 0.0009 | No |
Table: GSEA details
[plain text format]

  

Fig 2: CEREBELLAR\_CORTEX\_MORPHOGENESIS(GO:0021696)
  
Blue-Pink O' Gram in the Space of the Analyzed GeneSet

  

Fig 3: CEREBELLAR\_CORTEX\_MORPHOGENESIS(GO:0021696): Random ES distribution
  
Gene set null distribution of ES for
 **CEREBELLAR\_CORTEX\_MORPHOGENESIS(GO:0021696)**

  
