## Supplementary material for "A single cysteine residue in vimentin regulates long non-coding RNA *XIST* to suppress epithelial-mesenchymal transition and stemness in breast cancer": CYTOSOLIC_TRANSPORT(GO_0016482).html

Details for gene set CYTOSOLIC\_TRANSPORT(GO:0016482)[GSEA]


|  || Dataset | fpkm.sample |
| Phenotype | sample.cls |
| Upregulated in class | C328SVIM |
| GeneSet | CYTOSOLIC\_TRANSPORT(GO:0016482) |
| Enrichment Score (ES) | 0.8871889 |
| Normalized Enrichment Score (NES) | 1.4577494 |
| Nominal p-value | 0.0 |
| FDR q-value | 1.0 |
| FWER p-Value | 0.141 |
Table: GSEA Results Summary

  

Fig 1: Enrichment plot: CYTOSOLIC\_TRANSPORT(GO:0016482)
  
Profile of the Running ES Score & Positions of GeneSet Members on the Rank Ordered List

  

| PROBE | DESCRIPTION   (from dataset) | GENE SYMBOL | GENE\_TITLE | RANK IN GENE LIST | RANK METRIC SCORE | RUNNING ES | CORE ENRICHMENT || 1 | ENSG00000147065 | MSN |  |  | 5823 | 15136.710 | 0.8872 | Yes |
| 2 | ENSG00000146966 | DENND2A |  |  | 6265 | 37.070 | 0.8821 | No |
| 3 | ENSG00000185345 | PRKN |  |  | 7293 | 8.647 | 0.8653 | No |
| 4 | ENSG00000123570 | RAB9B |  |  | 7409 | 7.916 | 0.8638 | No |
| 5 | ENSG00000188906 | LRRK2 |  |  | 7426 | 7.825 | 0.8641 | No |
| 6 | ENSG00000102805 | CLN5 |  |  | 8088 | 5.434 | 0.8532 | No |
| 7 | ENSG00000147127 | RAB41 |  |  | 8140 | 5.348 | 0.8527 | No |
| 8 | ENSG00000067208 | EVI5 |  |  | 8935 | 3.992 | 0.8395 | No |
| 9 | ENSG00000168172 | HOOK3 |  |  | 9332 | 3.574 | 0.8330 | No |
| 10 | ENSG00000147642 | SYBU |  |  | 9516 | 3.375 | 0.8302 | No |
| 11 | ENSG00000087053 | MTMR2 |  |  | 9520 | 3.368 | 0.8303 | No |
| 12 | ENSG00000078142 | PIK3C3 |  |  | 9875 | 3.078 | 0.8245 | No |
| 13 | ENSG00000134709 | HOOK1 |  |  | 9987 | 3.013 | 0.8228 | No |
| 14 | ENSG00000115020 | PIKFYVE |  |  | 10450 | 2.711 | 0.8152 | No |
| 15 | ENSG00000070371 | CLTCL1 |  |  | 10571 | 2.646 | 0.8133 | No |
| 16 | ENSG00000181404 | WASHC1 |  |  | 10622 | 2.625 | 0.8126 | No |
| 17 | ENSG00000135968 | GCC2 |  |  | 10828 | 2.532 | 0.8093 | No |
| 18 | ENSG00000123106 | CCDC91 |  |  | 11048 | 2.432 | 0.8058 | No |
| 19 | ENSG00000184014 | DENND5A |  |  | 11224 | 2.369 | 0.8030 | No |
| 20 | ENSG00000276600 | RAB7B |  |  | 11393 | 2.313 | 0.8003 | No |
| 21 | ENSG00000137710 | RDX |  |  | 11742 | 2.181 | 0.7945 | No |
| 22 | ENSG00000141252 | VPS53 |  |  | 11990 | 2.080 | 0.7905 | No |
| 23 | ENSG00000122958 | VPS26A |  |  | 12164 | 2.026 | 0.7877 | No |
| 24 | ENSG00000069329 | VPS35 |  |  | 12188 | 2.019 | 0.7874 | No |
| 25 | ENSG00000138071 | ACTR2 |  |  | 12327 | 1.977 | 0.7852 | No |
| 26 | ENSG00000143952 | VPS54 |  |  | 12435 | 1.945 | 0.7835 | No |
| 27 | ENSG00000153071 | DAB2 |  |  | 12594 | 1.898 | 0.7810 | No |
| 28 | ENSG00000129515 | SNX6 |  |  | 12682 | 1.874 | 0.7796 | No |
| 29 | ENSG00000083097 | DOPEY1 |  |  | 12799 | 1.845 | 0.7778 | No |
| 30 | ENSG00000185722 | ANKFY1 |  |  | 12912 | 1.807 | 0.7760 | No |
| 31 | ENSG00000205302 | SNX2 |  |  | 13079 | 1.767 | 0.7733 | No |
| 32 | ENSG00000197969 | VPS13A |  |  | 13144 | 1.749 | 0.7723 | No |
| 33 | ENSG00000089006 | SNX5 |  |  | 13150 | 1.748 | 0.7723 | No |
| 34 | ENSG00000172661 | WASHC2C |  |  | 13224 | 1.732 | 0.7712 | No |
| 35 | ENSG00000008294 | SPAG9 |  |  | 13524 | 1.655 | 0.7662 | No |
| 36 | ENSG00000111237 | VPS29 |  |  | 13525 | 1.655 | 0.7664 | No |
| 37 | ENSG00000049245 | VAMP3 |  |  | 13713 | 1.626 | 0.7633 | No |
| 38 | ENSG00000188529 | SRSF10 |  |  | 13826 | 1.605 | 0.7615 | No |
| 39 | ENSG00000120805 | ARL1 |  |  | 13971 | 1.576 | 0.7592 | No |
| 40 | ENSG00000254806 | SYS1-DBNDD2 |  |  | 14072 | 1.559 | 0.7576 | No |
| 41 | ENSG00000170310 | STX8 |  |  | 14439 | 1.489 | 0.7515 | No |
| 42 | ENSG00000102189 | EEA1 |  |  | 14683 | 1.453 | 0.7474 | No |
| 43 | ENSG00000175931 | UBE2O |  |  | 14717 | 1.444 | 0.7470 | No |
| 44 | ENSG00000082805 | ERC1 |  |  | 15073 | 1.377 | 0.7410 | No |
| 45 | ENSG00000123595 | RAB9A |  |  | 15124 | 1.367 | 0.7403 | No |
| 46 | ENSG00000105875 | WDR91 |  |  | 15203 | 1.353 | 0.7390 | No |
| 47 | ENSG00000154917 | RAB6B |  |  | 15360 | 1.327 | 0.7365 | No |
| 48 | ENSG00000085832 | EPS15 |  |  | 15436 | 1.312 | 0.7353 | No |
| 49 | ENSG00000036054 | TBC1D23 |  |  | 15490 | 1.305 | 0.7345 | No |
| 50 | ENSG00000108587 | GOSR1 |  |  | 15601 | 1.288 | 0.7327 | No |
| 51 | ENSG00000129003 | VPS13C |  |  | 16059 | 1.225 | 0.7250 | No |
| 52 | ENSG00000099290 | WASHC2A |  |  | 16155 | 1.209 | 0.7235 | No |
| 53 | ENSG00000138660 | AP1AR |  |  | 16164 | 1.208 | 0.7235 | No |
| 54 | ENSG00000107185 | RGP1 |  |  | 16266 | 1.191 | 0.7218 | No |
| 55 | ENSG00000177951 | BET1L |  |  | 16357 | 1.184 | 0.7204 | No |
| 56 | ENSG00000138246 | DNAJC13 |  |  | 16466 | 1.170 | 0.7186 | No |
| 57 | ENSG00000151532 | VTI1A |  |  | 16516 | 1.164 | 0.7179 | No |
| 58 | ENSG00000100030 | MAPK1 |  |  | 16571 | 1.158 | 0.7170 | No |
| 59 | ENSG00000051009 | FAM160A2 |  |  | 17031 | 1.099 | 0.7093 | No |
| 60 | ENSG00000126581 | BECN1 |  |  | 17082 | 1.090 | 0.7085 | No |
| 61 | ENSG00000112335 | SNX3 |  |  | 17188 | 1.076 | 0.7068 | No |
| 62 | ENSG00000070423 | RNF126 |  |  | 17190 | 1.076 | 0.7069 | No |
| 63 | ENSG00000107036 | RIC1 |  |  | 17328 | 1.057 | 0.7046 | No |
| 64 | ENSG00000204843 | DCTN1 |  |  | 17439 | 1.043 | 0.7028 | No |
| 65 | ENSG00000105186 | ANKRD27 |  |  | 17576 | 1.027 | 0.7006 | No |
| 66 | ENSG00000117280 | RAB29 |  |  | 17629 | 1.021 | 0.6998 | No |
| 67 | ENSG00000089177 | KIF16B |  |  | 17800 | 1.000 | 0.6970 | No |
| 68 | ENSG00000131381 | RBSN |  |  | 17807 | 0.999 | 0.6969 | No |
| 69 | ENSG00000104886 | PLEKHJ1 |  |  | 18026 | 0.972 | 0.6933 | No |
| 70 | ENSG00000132405 | TBC1D14 |  |  | 18468 | 0.920 | 0.6859 | No |
| 71 | ENSG00000106636 | YKT6 |  |  | 18536 | 0.912 | 0.6848 | No |
| 72 | ENSG00000151502 | VPS26B |  |  | 18673 | 0.895 | 0.6825 | No |
| 73 | ENSG00000013016 | EHD3 |  |  | 18676 | 0.894 | 0.6826 | No |
| 74 | ENSG00000100568 | VTI1B |  |  | 18762 | 0.886 | 0.6812 | No |
| 75 | ENSG00000152291 | TGOLN2 |  |  | 18824 | 0.879 | 0.6802 | No |
| 76 | ENSG00000108433 | GOSR2 |  |  | 19052 | 0.854 | 0.6764 | No |
| 77 | ENSG00000061987 | MON2 |  |  | 19069 | 0.853 | 0.6762 | No |
| 78 | ENSG00000169221 | TBC1D10B |  |  | 19192 | 0.841 | 0.6742 | No |
| 79 | ENSG00000175463 | TBC1D10C |  |  | 19646 | 0.809 | 0.6666 | No |
| 80 | ENSG00000159210 | SNF8 |  |  | 19701 | 0.803 | 0.6657 | No |
| 81 | ENSG00000107862 | GBF1 |  |  | 19817 | 0.791 | 0.6638 | No |
| 82 | ENSG00000176658 | MYO1D |  |  | 19993 | 0.778 | 0.6609 | No |
| 83 | ENSG00000137642 | SORL1 |  |  | 20346 | 0.742 | 0.6550 | No |
| 84 | ENSG00000115561 | CHMP3 |  |  | 20348 | 0.742 | 0.6550 | No |
| 85 | ENSG00000080371 | RAB21 |  |  | 20357 | 0.741 | 0.6549 | No |
| 86 | ENSG00000198324 | FAM109A |  |  | 20374 | 0.740 | 0.6547 | No |
| 87 | ENSG00000167716 | WDR81 |  |  | 20595 | 0.732 | 0.6510 | No |
| 88 | ENSG00000169032 | MAP2K1 |  |  | 20668 | 0.724 | 0.6498 | No |
| 89 | ENSG00000175582 | RAB6A |  |  | 20774 | 0.712 | 0.6481 | No |
| 90 | ENSG00000104915 | STX10 |  |  | 20977 | 0.691 | 0.6447 | No |
| 91 | ENSG00000162236 | STX5 |  |  | 21022 | 0.688 | 0.6440 | No |
| 92 | ENSG00000147164 | SNX12 |  |  | 21048 | 0.685 | 0.6436 | No |
| 93 | ENSG00000028528 | SNX1 |  |  | 21418 | 0.649 | 0.6374 | No |
| 94 | ENSG00000076321 | KLHL20 |  |  | 21458 | 0.645 | 0.6368 | No |
| 95 | ENSG00000262246 | CORO7 |  |  | 21559 | 0.635 | 0.6352 | No |
| 96 | ENSG00000119396 | RAB14 |  |  | 21645 | 0.628 | 0.6338 | No |
| 97 | ENSG00000103978 | TMEM87A |  |  | 21650 | 0.627 | 0.6337 | No |
| 98 | ENSG00000164715 | LMTK2 |  |  | 21981 | 0.599 | 0.6282 | No |
| 99 | ENSG00000126934 | MAP2K2 |  |  | 22444 | 0.562 | 0.6204 | No |
| 100 | ENSG00000144566 | RAB5A |  |  | 22605 | 0.555 | 0.6177 | No |
| 101 | ENSG00000280433 | FP565260.7 |  |  | 22902 | 0.528 | 0.6127 | No |
| 102 | ENSG00000104497 | SNX16 |  |  | 22925 | 0.526 | 0.6124 | No |
| 103 | ENSG00000197122 | SRC |  |  | 23479 | 0.482 | 0.6030 | No |
| 104 | ENSG00000092820 | EZR |  |  | 23520 | 0.479 | 0.6024 | No |
| 105 | ENSG00000166747 | AP1G1 |  |  | 23549 | 0.476 | 0.6019 | No |
| 106 | ENSG00000204070 | SYS1 |  |  | 23638 | 0.469 | 0.6005 | No |
| 107 | ENSG00000124222 | STX16 |  |  | 23782 | 0.457 | 0.5981 | No |
| 108 | ENSG00000075785 | RAB7A |  |  | 23825 | 0.454 | 0.5974 | No |
| 109 | ENSG00000076201 | PTPN23 |  |  | 24138 | 0.428 | 0.5921 | No |
| 110 | ENSG00000221838 | AP4M1 |  |  | 24456 | 0.405 | 0.5868 | No |
| 111 | ENSG00000141367 | CLTC |  |  | 24656 | 0.389 | 0.5835 | No |
| 112 | ENSG00000222014 | RAB6C |  |  | 24665 | 0.388 | 0.5833 | No |
| 113 | ENSG00000178950 | GAK |  |  | 24706 | 0.386 | 0.5827 | No |
| 114 | ENSG00000134243 | SORT1 |  |  | 24817 | 0.376 | 0.5808 | No |
| 115 | ENSG00000131374 | TBC1D5 |  |  | 24823 | 0.376 | 0.5808 | No |
| 116 | ENSG00000196961 | AP2A1 |  |  | 24869 | 0.372 | 0.5800 | No |
| 117 | ENSG00000185896 | LAMP1 |  |  | 24906 | 0.370 | 0.5795 | No |
| 118 | ENSG00000160218 | TRAPPC10 |  |  | 25083 | 0.359 | 0.5765 | No |
| 119 | ENSG00000102882 | MAPK3 |  |  | 25118 | 0.357 | 0.5760 | No |
| 120 | ENSG00000153214 | TMEM87B |  |  | 25182 | 0.353 | 0.5749 | No |
| 121 | ENSG00000141258 | SGSM2 |  |  | 25370 | 0.338 | 0.5718 | No |
| 122 | ENSG00000164292 | RHOBTB3 |  |  | 25547 | 0.323 | 0.5688 | No |
| 123 | ENSG00000149823 | VPS51 |  |  | 25886 | 0.295 | 0.5631 | No |
| 124 | ENSG00000106266 | SNX8 |  |  | 26043 | 0.281 | 0.5605 | No |
| 125 | ENSG00000101246 | ARFRP1 |  |  | 26064 | 0.280 | 0.5601 | No |
| 126 | ENSG00000099992 | TBC1D10A |  |  | 26238 | 0.268 | 0.5572 | No |
| 127 | ENSG00000135823 | STX6 |  |  | 26297 | 0.264 | 0.5563 | No |
| 128 | ENSG00000070540 | WIPI1 |  |  | 26350 | 0.259 | 0.5554 | No |
| 129 | ENSG00000104946 | TBC1D17 |  |  | 26630 | 0.235 | 0.5507 | No |
| 130 | ENSG00000166971 | AKTIP |  |  | 26721 | 0.228 | 0.5492 | No |
| 131 | ENSG00000213853 | EMP2 |  |  | 27502 | 0.161 | 0.5360 | No |
| 132 | ENSG00000106367 | AP1S1 |  |  | 27571 | 0.157 | 0.5348 | No |
| 133 | ENSG00000100711 | ZFYVE21 |  |  | 27755 | 0.140 | 0.5317 | No |
| 134 | ENSG00000095066 | HOOK2 |  |  | 28337 | 0.091 | 0.5219 | No |
| 135 | ENSG00000102879 | CORO1A |  |  | 28572 | 0.070 | 0.5179 | No |
| 136 | ENSG00000011347 | SYT7 |  |  | 28825 | 0.048 | 0.5136 | No |
| 137 | ENSG00000142197 | DOPEY2 |  |  | 28964 | 0.036 | 0.5113 | No |
| 138 | ENSG00000177096 | FAM109B |  |  | 29001 | 0.034 | 0.5107 | No |
| 139 | ENSG00000160695 | VPS11 |  |  | 35385 | NaN | 0.4025 | No |
| 140 | ENSG00000198848 | CES1 |  |  | 37903 | NaN | 0.3598 | No |
| 141 | ENSG00000254585 | MAGEL2 |  |  | 38829 | NaN | 0.3442 | No |
| 142 | ENSG00000223501 | VPS52 |  |  | 49118 | NaN | 0.1698 | No |
| 143 | ENSG00000204713 | TRIM27 |  |  | 49859 | NaN | 0.1572 | No |
| 144 | ENSG00000167705 | RILP |  |  | 52245 | NaN | 0.1168 | No |
| 145 | ENSG00000262633 | AC005670.2 |  |  | 53818 | NaN | 0.0901 | No |
Table: GSEA details
[plain text format]

  

Fig 2: CYTOSOLIC\_TRANSPORT(GO:0016482)
  
Blue-Pink O' Gram in the Space of the Analyzed GeneSet

  

Fig 3: CYTOSOLIC\_TRANSPORT(GO:0016482): Random ES distribution
  
Gene set null distribution of ES for
 **CYTOSOLIC\_TRANSPORT(GO:0016482)**

  
