## Supplementary material for "A single cysteine residue in vimentin regulates long non-coding RNA *XIST* to suppress epithelial-mesenchymal transition and stemness in breast cancer": ESTABLISHMENT_OR_MAINTENANCE_OF_EPITHELIAL_CELL_APICAL_BASAL_POLARITY(GO_0045197).html

Details for gene set ESTABLISHMENT\_OR\_MAINTENANCE\_OF\_EPITHELIAL\_CELL\_APICAL\_BASAL\_POLARITY(GO:0045197)[GSEA]


|  || Dataset | fpkm.sample |
| Phenotype | sample.cls |
| Upregulated in class | C328SVIM |
| GeneSet | ESTABLISHMENT\_OR\_MAINTENANCE\_OF\_EPITHELIAL\_CELL\_APICAL\_BASAL\_POLARITY(GO:0045197) |
| Enrichment Score (ES) | 0.8933289 |
| Normalized Enrichment Score (NES) | 1.4402307 |
| Nominal p-value | 0.0 |
| FDR q-value | 1.0 |
| FWER p-Value | 0.325 |
Table: GSEA Results Summary

  

Fig 1: Enrichment plot: ESTABLISHMENT\_OR\_MAINTENANCE\_OF\_EPITHELIAL\_CELL\_APICAL\_BASAL\_POLARITY(GO:0045197)
  
Profile of the Running ES Score & Positions of GeneSet Members on the Rank Ordered List

  

| PROBE | DESCRIPTION   (from dataset) | GENE SYMBOL | GENE\_TITLE | RANK IN GENE LIST | RANK METRIC SCORE | RUNNING ES | CORE ENRICHMENT || 1 | ENSG00000165556 | CDX2 |  |  | 725 | Infinity | -0.0123 | Yes |
| 2 | ENSG00000240720 | LRRD1 |  |  | 4671 | Infinity | -0.0790 | Yes |
| 3 | ENSG00000147065 | MSN |  |  | 5823 | 15136.710 | 0.8853 | Yes |
| 4 | ENSG00000103241 | FOXF1 |  |  | 5986 | 137.636 | 0.8915 | Yes |
| 5 | ENSG00000162738 | VANGL2 |  |  | 6121 | 63.248 | 0.8933 | Yes |
| 6 | ENSG00000114251 | WNT5A |  |  | 6473 | 22.250 | 0.8888 | No |
| 7 | ENSG00000132535 | DLG4 |  |  | 12455 | 1.934 | 0.7878 | No |
| 8 | ENSG00000148943 | LIN7C |  |  | 12611 | 1.893 | 0.7853 | No |
| 9 | ENSG00000067560 | RHOA |  |  | 14175 | 1.551 | 0.7589 | No |
| 10 | ENSG00000112851 | ERBIN |  |  | 14275 | 1.526 | 0.7573 | No |
| 11 | ENSG00000066933 | MYO9A |  |  | 14303 | 1.518 | 0.7570 | No |
| 12 | ENSG00000163558 | PRKCI |  |  | 14571 | 1.475 | 0.7526 | No |
| 13 | ENSG00000070831 | CDC42 |  |  | 14664 | 1.457 | 0.7511 | No |
| 14 | ENSG00000112655 | PTK7 |  |  | 14827 | 1.423 | 0.7484 | No |
| 15 | ENSG00000166333 | ILK |  |  | 15300 | 1.338 | 0.7405 | No |
| 16 | ENSG00000170248 | PDCD6IP |  |  | 15718 | 1.273 | 0.7336 | No |
| 17 | ENSG00000111052 | LIN7A |  |  | 16097 | 1.218 | 0.7273 | No |
| 18 | ENSG00000168374 | ARF4 |  |  | 17095 | 1.087 | 0.7105 | No |
| 19 | ENSG00000148204 | CRB2 |  |  | 17622 | 1.023 | 0.7016 | No |
| 20 | ENSG00000085741 | WNT11 |  |  | 17680 | 1.015 | 0.7007 | No |
| 21 | ENSG00000106689 | LHX2 |  |  | 18719 | 0.890 | 0.6832 | No |
| 22 | ENSG00000075711 | DLG1 |  |  | 19219 | 0.839 | 0.6748 | No |
| 23 | ENSG00000120509 | PDZD11 |  |  | 19638 | 0.809 | 0.6678 | No |
| 24 | ENSG00000168502 | MTCL1 |  |  | 21930 | 0.604 | 0.6291 | No |
| 25 | ENSG00000100092 | SH3BP1 |  |  | 22895 | 0.529 | 0.6128 | No |
| 26 | ENSG00000082458 | DLG3 |  |  | 22924 | 0.526 | 0.6124 | No |
| 27 | ENSG00000029534 | ANK1 |  |  | 23530 | 0.478 | 0.6022 | No |
| 28 | ENSG00000180900 | SCRIB |  |  | 24481 | 0.404 | 0.5861 | No |
| 29 | ENSG00000072518 | MARK2 |  |  | 25000 | 0.367 | 0.5774 | No |
| 30 | ENSG00000079482 | OPHN1 |  |  | 25299 | 0.343 | 0.5723 | No |
| 31 | ENSG00000076826 | CAMSAP3 |  |  | 26407 | 0.253 | 0.5536 | No |
| 32 | ENSG00000104863 | LIN7B |  |  | 27065 | 0.199 | 0.5425 | No |
| 33 | ENSG00000071127 | WDR1 |  |  | 27073 | 0.199 | 0.5424 | No |
| 34 | ENSG00000077454 | LRCH4 |  |  | 27150 | 0.190 | 0.5411 | No |
| 35 | ENSG00000109062 | SLC9A3R1 |  |  | 28790 | 0.051 | 0.5134 | No |
| 36 | ENSG00000181392 | SYNE4 |  |  | 29098 | 0.028 | 0.5082 | No |
| 37 | ENSG00000151208 | DLG5 |  |  | 30039 | 0.000 | 0.4923 | No |
| 38 | ENSG00000125878 | TCF15 |  |  | 58275 | NaN | 0.0146 | No |
Table: GSEA details
[plain text format]

  

Fig 2: ESTABLISHMENT\_OR\_MAINTENANCE\_OF\_EPITHELIAL\_CELL\_APICAL\_BASAL\_POLARITY(GO:0045197)
  
Blue-Pink O' Gram in the Space of the Analyzed GeneSet

  

Fig 3: ESTABLISHMENT\_OR\_MAINTENANCE\_OF\_EPITHELIAL\_CELL\_APICAL\_BASAL\_POLARITY(GO:0045197): Random ES distribution
  
Gene set null distribution of ES for
 **ESTABLISHMENT\_OR\_MAINTENANCE\_OF\_EPITHELIAL\_CELL\_APICAL\_BASAL\_POLARITY(GO:0045197)**

  
