## Supplementary material for "A single cysteine residue in vimentin regulates long non-coding RNA *XIST* to suppress epithelial-mesenchymal transition and stemness in breast cancer": gsea_report_for_C328SVIM_1625876782425.html

Report for C328SVIM 1625876782425 [GSEA]


| GS   follow link to MSigDB | GS DETAILS | SIZE | ES | NES | NOM p-val | FDR q-val | FWER p-val | RANK AT MAX | LEADING EDGE || 1 | CYTOSOLIC\_TRANSPORT(GO:0016482) | Details ... | 145 | 0.89 | 1.46 | 0.000 | 1.000 | 0.141 | 5823 | tags=1%, list=10%, signal=1% |
| 2 | VESICLE\_MEDIATED\_TRANSPORT\_BETWEEN\_ENDOSOMAL\_COMPARTMENTS(GO:0098927) | Details ... | 40 | 0.90 | 1.46 | 0.000 | 1.000 | 0.325 | 5823 | tags=3%, list=10%, signal=3% |
| 3 | EARLY\_ENDOSOME\_TO\_LATE\_ENDOSOME\_TRANSPORT(GO:0045022) | Details ... | 37 | 0.90 | 1.45 | 0.000 | 1.000 | 0.325 | 5823 | tags=3%, list=10%, signal=3% |
| 4 | NEGATIVE\_REGULATION\_OF\_INTERLEUKIN\_1\_PRODUCTION(GO:0032692) | Details ... | 27 | 0.90 | 1.45 | 0.000 | 1.000 | 0.325 | 5825 | tags=7%, list=10%, signal=8% |
| 5 | ESTABLISHMENT\_OR\_MAINTENANCE\_OF\_BIPOLAR\_CELL\_POLARITY(GO:0061245) | Details ... | 42 | 0.89 | 1.45 | 0.000 | 1.000 | 0.325 | 6121 | tags=12%, list=10%, signal=13% |
| 6 | ESTABLISHMENT\_OR\_MAINTENANCE\_OF\_APICAL\_BASAL\_CELL\_POLARITY(GO:0035088) | Details ... | 42 | 0.89 | 1.45 | 0.000 | 1.000 | 0.325 | 6121 | tags=12%, list=10%, signal=13% |
| 7 | CELLULAR\_RESPONSE\_TO\_INTERLEUKIN\_12(GO:0071349) | Details ... | 47 | 0.89 | 1.45 | 0.000 | 1.000 | 0.325 | 6128 | tags=15%, list=10%, signal=17% |
| 8 | INTERLEUKIN\_12\_MEDIATED\_SIGNALING\_PATHWAY(GO:0035722) | Details ... | 47 | 0.89 | 1.45 | 0.000 | 1.000 | 0.325 | 6128 | tags=15%, list=10%, signal=17% |
| 9 | RESPONSE\_TO\_INTERLEUKIN\_12(GO:0070671) | Details ... | 49 | 0.89 | 1.44 | 0.000 | 1.000 | 0.325 | 6128 | tags=16%, list=10%, signal=18% |
| 10 | NEGATIVE\_REGULATION\_OF\_INTERLEUKIN\_1\_BETA\_PRODUCTION(GO:0032691) | Details ... | 21 | 0.90 | 1.44 | 0.000 | 1.000 | 0.325 | 5825 | tags=10%, list=10%, signal=11% |
| 11 | REGULATION\_OF\_TUMOR\_NECROSIS\_FACTOR\_MEDIATED\_SIGNALING\_PATHWAY(GO:0010803) | Details ... | 61 | 0.89 | 1.44 | 0.000 | 1.000 | 0.325 | 5922 | tags=3%, list=10%, signal=4% |
| 12 | ESTABLISHMENT\_OR\_MAINTENANCE\_OF\_EPITHELIAL\_CELL\_APICAL\_BASAL\_POLARITY(GO:0045197) | Details ... | 38 | 0.89 | 1.44 | 0.000 | 1.000 | 0.325 | 6121 | tags=13%, list=10%, signal=15% |
| 13 | PROTEIN\_LOCALIZATION\_TO\_ENDOSOME(GO:0036010) | Details ... | 18 | 0.90 | 1.44 | 0.000 | 1.000 | 0.325 | 5823 | tags=6%, list=10%, signal=6% |
| 14 | INSULIN\_RECEPTOR\_BINDING(GO:0005158) | Details ... | 22 | 0.90 | 1.44 | 0.000 | 1.000 | 0.325 | 5824 | tags=5%, list=10%, signal=5% |
| 15 | CELLULAR\_RESPONSE\_TO\_INSULIN\_STIMULUS(GO:0032869) | Details ... | 200 | 0.89 | 1.44 | 0.000 | 1.000 | 0.325 | 5825 | tags=5%, list=10%, signal=6% |
| 16 | CEREBELLAR\_CORTEX\_MORPHOGENESIS(GO:0021696) | Details ... | 33 | 0.88 | 1.43 | 0.000 | 1.000 | 0.325 | 5943 | tags=18%, list=10%, signal=20% |
| 17 | ATRIAL\_SEPTUM\_DEVELOPMENT(GO:0003283) | Details ... | 20 | 0.89 | 1.43 | 0.000 | 1.000 | 0.325 | 5958 | tags=30%, list=10%, signal=33% |
| 18 | REGULATION\_OF\_CELLULAR\_PROTEIN\_CATABOLIC\_PROCESS(GO:1903362) | Details ... | 230 | 0.88 | 1.43 | 0.000 | 1.000 | 0.325 | 6154 | tags=6%, list=10%, signal=6% |
| 19 | MYELIN\_SHEATH(GO:0043209) | Details ... | 164 | 0.88 | 1.43 | 0.000 | 1.000 | 0.325 | 6303 | tags=11%, list=11%, signal=12% |
| 20 | PURINE\_NUCLEOTIDE\_CATABOLIC\_PROCESS(GO:0006195) | Details ... | 47 | 0.86 | 1.43 | 0.000 | 1.000 | 0.325 | 6423 | tags=19%, list=11%, signal=21% |
| 21 | MEMBRANE\_DOCKING(GO:0022406) |  | 170 | 0.88 | 1.43 | 0.000 | 1.000 | 0.325 | 5823 | tags=2%, list=10%, signal=2% |
| 22 | INSULIN\_RECEPTOR\_SIGNALING\_PATHWAY(GO:0008286) |  | 123 | 0.89 | 1.43 | 0.000 | 1.000 | 0.325 | 5824 | tags=4%, list=10%, signal=4% |
| 23 | CEREBELLAR\_CORTEX\_DEVELOPMENT(GO:0021695) |  | 52 | 0.87 | 1.43 | 0.000 | 1.000 | 0.325 | 5943 | tags=17%, list=10%, signal=19% |
| 24 | FICOLIN\_1\_RICH\_GRANULE\_LUMEN(GO:1904813) |  | 124 | 0.88 | 1.42 | 0.000 | 1.000 | 0.325 | 5825 | tags=2%, list=10%, signal=3% |
| 25 | CELLULAR\_RESPONSE\_TO\_PEPTIDE(GO:1901653) |  | 350 | 0.88 | 1.42 | 0.000 | 1.000 | 0.325 | 6005 | tags=7%, list=10%, signal=8% |
| 26 | TRANSFERASE\_ACTIVITY\_\_TRANSFERRING\_ALKYL\_OR\_ARYL\_(OTHER\_THAN\_METHYL)\_GROUPS(GO:0016765) |  | 57 | 0.89 | 1.42 | 0.000 | 1.000 | 0.325 | 5825 | tags=11%, list=10%, signal=12% |
| 27 | POSITIVE\_REGULATION\_OF\_CELLULAR\_PROTEIN\_CATABOLIC\_PROCESS(GO:1903364) |  | 136 | 0.88 | 1.42 | 0.000 | 1.000 | 0.325 | 6133 | tags=7%, list=10%, signal=7% |
| 28 | SIGNALING\_ADAPTOR\_ACTIVITY(GO:0035591) |  | 79 | 0.89 | 1.42 | 0.000 | 1.000 | 0.325 | 5824 | tags=9%, list=10%, signal=10% |
| 29 | CELLULAR\_RESPONSE\_TO\_PEPTIDE\_HORMONE\_STIMULUS(GO:0071375) |  | 307 | 0.88 | 1.42 | 0.000 | 1.000 | 0.325 | 6005 | tags=7%, list=10%, signal=8% |
| 30 | HOMEOSTASIS\_OF\_NUMBER\_OF\_CELLS\_WITHIN\_A\_TISSUE(GO:0048873) |  | 29 | 0.88 | 1.42 | 0.000 | 1.000 | 0.325 | 5916 | tags=17%, list=10%, signal=19% |
| 31 | POSITIVE\_REGULATION\_OF\_COFACTOR\_METABOLIC\_PROCESS(GO:0051194) |  | 23 | 0.88 | 1.42 | 0.000 | 1.000 | 0.325 | 6006 | tags=17%, list=10%, signal=19% |
| 32 | T\_CELL\_PROLIFERATION(GO:0042098) |  | 172 | 0.88 | 1.42 | 0.000 | 1.000 | 0.325 | 6178 | tags=8%, list=10%, signal=8% |
| 33 | POLYOL\_BIOSYNTHETIC\_PROCESS(GO:0046173) |  | 54 | 0.87 | 1.42 | 0.000 | 1.000 | 0.325 | 5955 | tags=15%, list=10%, signal=16% |
| 34 | FICOLIN\_1\_RICH\_GRANULE(GO:0101002) |  | 184 | 0.87 | 1.42 | 0.000 | 1.000 | 0.325 | 6057 | tags=3%, list=10%, signal=4% |
| 35 | FILOPODIUM(GO:0030175) |  | 95 | 0.89 | 1.42 | 0.000 | 1.000 | 0.325 | 6054 | tags=14%, list=10%, signal=15% |
| 36 | ESTABLISHMENT\_OF\_EPITHELIAL\_CELL\_POLARITY(GO:0090162) |  | 28 | 0.90 | 1.42 | 0.000 | 1.000 | 0.325 | 5986 | tags=7%, list=10%, signal=8% |
| 37 | REGULATION\_OF\_CYTOPLASMIC\_TRANSPORT(GO:1903649) |  | 21 | 0.90 | 1.42 | 0.000 | 1.000 | 0.325 | 5823 | tags=5%, list=10%, signal=5% |
| 38 | ESTABLISHMENT\_OF\_APICAL\_BASAL\_CELL\_POLARITY(GO:0035089) |  | 17 | 0.90 | 1.42 | 0.000 | 1.000 | 0.325 | 5986 | tags=12%, list=10%, signal=13% |
| 39 | HINDBRAIN\_MORPHOGENESIS(GO:0021575) |  | 44 | 0.88 | 1.42 | 0.000 | 1.000 | 0.325 | 5943 | tags=16%, list=10%, signal=18% |
| 40 | POSITIVE\_REGULATION\_OF\_PROTEIN\_CATABOLIC\_PROCESS(GO:0045732) |  | 205 | 0.88 | 1.42 | 0.000 | 1.000 | 0.325 | 6133 | tags=6%, list=10%, signal=7% |
| 41 | ALDITOL\_METABOLIC\_PROCESS(GO:0019400) |  | 21 | 0.89 | 1.42 | 0.000 | 1.000 | 0.325 | 5839 | tags=24%, list=10%, signal=26% |
| 42 | CEREBELLUM\_MORPHOGENESIS(GO:0021587) |  | 40 | 0.88 | 1.42 | 0.000 | 1.000 | 0.325 | 5943 | tags=18%, list=10%, signal=19% |
| 43 | NEGATIVE\_REGULATION\_OF\_FIBROBLAST\_PROLIFERATION(GO:0048147) |  | 33 | 0.89 | 1.42 | 0.000 | 1.000 | 0.325 | 6027 | tags=6%, list=10%, signal=7% |
| 44 | CORTICAL\_CYTOSKELETON\_ORGANIZATION(GO:0030865) |  | 35 | 0.87 | 1.41 | 0.000 | 1.000 | 0.325 | 5836 | tags=9%, list=10%, signal=10% |
| 45 | REGULATION\_OF\_PROTEIN\_CATABOLIC\_PROCESS(GO:0042176) |  | 352 | 0.87 | 1.41 | 0.000 | 1.000 | 0.325 | 6247 | tags=6%, list=11%, signal=6% |
| 46 | GMP\_METABOLIC\_PROCESS(GO:0046037) |  | 20 | 0.88 | 1.41 | 0.000 | 1.000 | 0.325 | 5849 | tags=5%, list=10%, signal=6% |
| 47 | REGULATION\_OF\_CELL\_SHAPE(GO:0008360) |  | 144 | 0.88 | 1.41 | 0.000 | 1.000 | 0.325 | 5877 | tags=4%, list=10%, signal=5% |
| 48 | ATRIAL\_SEPTUM\_MORPHOGENESIS(GO:0060413) |  | 16 | 0.90 | 1.41 | 0.000 | 1.000 | 0.325 | 5958 | tags=31%, list=10%, signal=35% |
| 49 | ESTABLISHMENT\_OF\_MONOPOLAR\_CELL\_POLARITY(GO:0061162) |  | 19 | 0.90 | 1.41 | 0.000 | 1.000 | 0.325 | 5986 | tags=11%, list=10%, signal=12% |
| 50 | POSITIVE\_REGULATION\_OF\_GLYCOLYTIC\_PROCESS(GO:0045821) |  | 19 | 0.88 | 1.41 | 0.000 | 1.000 | 0.325 | 6006 | tags=16%, list=10%, signal=18% |
| 51 | SH3\_SH2\_ADAPTOR\_ACTIVITY(GO:0005070) |  | 56 | 0.90 | 1.41 | 0.000 | 1.000 | 0.325 | 5824 | tags=13%, list=10%, signal=14% |
| 52 | MORPHOGENESIS\_OF\_A\_POLARIZED\_EPITHELIUM(GO:0001738) |  | 138 | 0.89 | 1.41 | 0.000 | 1.000 | 0.325 | 6121 | tags=7%, list=10%, signal=7% |
| 53 | RESPONSE\_TO\_INSULIN(GO:0032868) |  | 255 | 0.88 | 1.41 | 0.000 | 1.000 | 0.325 | 5825 | tags=5%, list=10%, signal=5% |
| 54 | POSITIVE\_REGULATION\_OF\_CELLULAR\_CATABOLIC\_PROCESS(GO:0031331) |  | 334 | 0.87 | 1.41 | 0.000 | 1.000 | 0.325 | 6212 | tags=6%, list=11%, signal=6% |
| 55 | DETECTION\_OF\_BIOTIC\_STIMULUS(GO:0009595) |  | 22 | 0.88 | 1.41 | 0.000 | 1.000 | 0.325 | 6838 | tags=27%, list=12%, signal=31% |
| 56 | NEGATIVE\_REGULATION\_OF\_STRESS\_ACTIVATED\_MAPK\_CASCADE(GO:0032873) |  | 47 | 0.89 | 1.41 | 0.000 | 1.000 | 0.325 | 6027 | tags=6%, list=10%, signal=7% |
| 57 | NEGATIVE\_REGULATION\_OF\_STRESS\_ACTIVATED\_PROTEIN\_KINASE\_SIGNALING\_CASCADE(GO:0070303) |  | 47 | 0.89 | 1.41 | 0.000 | 1.000 | 0.325 | 6027 | tags=6%, list=10%, signal=7% |
| 58 | MUSCLE\_CELL\_CELLULAR\_HOMEOSTASIS(GO:0046716) |  | 19 | 0.88 | 1.41 | 0.000 | 1.000 | 0.325 | 5856 | tags=21%, list=10%, signal=23% |
| 59 | NEGATIVE\_REGULATION\_OF\_MAPK\_CASCADE(GO:0043409) |  | 172 | 0.87 | 1.41 | 0.000 | 1.000 | 0.325 | 6047 | tags=7%, list=10%, signal=8% |
| 60 | GLUTATHIONE\_PEROXIDASE\_ACTIVITY(GO:0004602) |  | 19 | 0.90 | 1.41 | 0.000 | 1.000 | 0.325 | 5834 | tags=21%, list=10%, signal=23% |
| 61 | RESPONSE\_TO\_HYDROPEROXIDE(GO:0033194) |  | 19 | 0.89 | 1.41 | 0.000 | 1.000 | 0.325 | 6212 | tags=16%, list=11%, signal=18% |
| 62 | POSITIVE\_REGULATION\_OF\_CALCIUM\_ION\_TRANSMEMBRANE\_TRANSPORTER\_ACTIVITY(GO:1901021) |  | 33 | 0.87 | 1.41 | 0.000 | 1.000 | 0.325 | 5882 | tags=9%, list=10%, signal=10% |
| 63 | ESTABLISHMENT\_OR\_MAINTENANCE\_OF\_MONOPOLAR\_CELL\_POLARITY(GO:0061339) |  | 20 | 0.90 | 1.41 | 0.000 | 1.000 | 0.325 | 5986 | tags=10%, list=10%, signal=11% |
| 64 | NEGATIVE\_REGULATION\_OF\_MITOCHONDRION\_ORGANIZATION(GO:0010823) |  | 50 | 0.87 | 1.41 | 0.000 | 1.000 | 0.325 | 5834 | tags=4%, list=10%, signal=4% |
| 65 | REGULATION\_OF\_OXIDATIVE\_STRESS\_INDUCED\_INTRINSIC\_APOPTOTIC\_SIGNALING\_PATHWAY(GO:1902175) |  | 30 | 0.87 | 1.41 | 0.000 | 1.000 | 0.325 | 5834 | tags=7%, list=10%, signal=7% |
| 66 | PHOSPHOTRANSFERASE\_ACTIVITY\_\_PHOSPHATE\_GROUP\_AS\_ACCEPTOR(GO:0016776) |  | 42 | 0.85 | 1.41 | 0.000 | 1.000 | 0.325 | 5849 | tags=7%, list=10%, signal=8% |
| 67 | INTRINSIC\_APOPTOTIC\_SIGNALING\_PATHWAY\_IN\_RESPONSE\_TO\_OXIDATIVE\_STRESS(GO:0008631) |  | 45 | 0.87 | 1.41 | 0.000 | 1.000 | 0.325 | 5834 | tags=4%, list=10%, signal=5% |
| 68 | NEGATIVE\_REGULATION\_OF\_INFLAMMATORY\_RESPONSE(GO:0050728) |  | 120 | 0.89 | 1.41 | 0.000 | 1.000 | 0.325 | 6048 | tags=14%, list=10%, signal=16% |
| 69 | POSITIVE\_REGULATION\_OF\_CATABOLIC\_PROCESS(GO:0009896) |  | 386 | 0.87 | 1.41 | 0.000 | 1.000 | 0.325 | 6212 | tags=6%, list=11%, signal=7% |
| 70 | REGULATION\_OF\_CHEMOKINE\_PRODUCTION(GO:0032642) |  | 76 | 0.88 | 1.40 | 0.000 | 1.000 | 0.325 | 5904 | tags=12%, list=10%, signal=13% |
| 71 | NEGATIVE\_REGULATION\_OF\_RELEASE\_OF\_CYTOCHROME\_C\_FROM\_MITOCHONDRIA(GO:0090201) |  | 17 | 0.89 | 1.40 | 0.000 | 1.000 | 0.325 | 5834 | tags=12%, list=10%, signal=13% |
| 72 | REGULATION\_OF\_ORGANELLE\_ASSEMBLY(GO:1902115) |  | 162 | 0.88 | 1.40 | 0.000 | 1.000 | 0.325 | 5952 | tags=4%, list=10%, signal=5% |
| 73 | INTERACTION\_WITH\_SYMBIONT(GO:0051702) |  | 73 | 0.86 | 1.40 | 0.000 | 1.000 | 0.325 | 6381 | tags=14%, list=11%, signal=15% |
| 74 | PODOSOME\_ASSEMBLY(GO:0071800) |  | 20 | 0.90 | 1.40 | 0.000 | 1.000 | 0.325 | 6029 | tags=15%, list=10%, signal=17% |
| 75 | NEGATIVE\_REGULATION\_OF\_MAP\_KINASE\_ACTIVITY(GO:0043407) |  | 77 | 0.88 | 1.40 | 0.000 | 1.000 | 0.325 | 6047 | tags=8%, list=10%, signal=9% |
| 76 | LEUKOCYTE\_PROLIFERATION(GO:0070661) |  | 276 | 0.88 | 1.40 | 0.000 | 1.000 | 0.325 | 6178 | tags=11%, list=10%, signal=12% |
| 77 | ALCOHOL\_BIOSYNTHETIC\_PROCESS(GO:0046165) |  | 133 | 0.86 | 1.40 | 0.000 | 1.000 | 0.325 | 5955 | tags=9%, list=10%, signal=10% |
| 78 | CELLULAR\_RESPONSE\_TO\_OSMOTIC\_STRESS(GO:0071470) |  | 33 | 0.87 | 1.40 | 0.000 | 1.000 | 0.325 | 6424 | tags=15%, list=11%, signal=17% |
| 79 | CARDIAC\_ATRIUM\_DEVELOPMENT(GO:0003230) |  | 35 | 0.87 | 1.40 | 0.000 | 1.000 | 0.325 | 5958 | tags=20%, list=10%, signal=22% |
| 80 | POSITIVE\_REGULATION\_OF\_COENZYME\_METABOLIC\_PROCESS(GO:0051197) |  | 20 | 0.88 | 1.40 | 0.000 | 1.000 | 0.325 | 6006 | tags=15%, list=10%, signal=17% |
| 81 | BLOOD\_MICROPARTICLE(GO:0072562) |  | 189 | 0.88 | 1.40 | 0.000 | 1.000 | 0.325 | 6193 | tags=7%, list=10%, signal=8% |
| 82 | OXIDOREDUCTASE\_ACTIVITY\_\_ACTING\_ON\_PEROXIDE\_AS\_ACCEPTOR(GO:0016684) |  | 47 | 0.89 | 1.40 | 0.000 | 1.000 | 0.325 | 5834 | tags=15%, list=10%, signal=17% |
| 83 | NUCLEOBASE\_CONTAINING\_COMPOUND\_KINASE\_ACTIVITY(GO:0019205) |  | 50 | 0.84 | 1.40 | 0.000 | 1.000 | 0.325 | 5849 | tags=6%, list=10%, signal=7% |
| 84 | NEGATIVE\_REGULATION\_OF\_OXIDATIVE\_STRESS\_INDUCED\_INTRINSIC\_APOPTOTIC\_SIGNALING\_PATHWAY(GO:1902176) |  | 20 | 0.89 | 1.40 | 0.000 | 1.000 | 0.325 | 5834 | tags=10%, list=10%, signal=11% |
| 85 | POSITIVE\_REGULATION\_OF\_NUCLEOTIDE\_CATABOLIC\_PROCESS(GO:0030813) |  | 22 | 0.88 | 1.40 | 0.000 | 1.000 | 0.325 | 6006 | tags=14%, list=10%, signal=15% |
| 86 | PURINE\_CONTAINING\_COMPOUND\_CATABOLIC\_PROCESS(GO:0072523) |  | 54 | 0.86 | 1.40 | 0.000 | 1.000 | 0.325 | 6423 | tags=19%, list=11%, signal=21% |
| 87 | NEGATIVE\_REGULATION\_OF\_JNK\_CASCADE(GO:0046329) |  | 35 | 0.89 | 1.40 | 0.000 | 1.000 | 0.325 | 6027 | tags=9%, list=10%, signal=10% |
| 88 | NEGATIVE\_REGULATION\_OF\_JUN\_KINASE\_ACTIVITY(GO:0043508) |  | 16 | 0.89 | 1.40 | 0.000 | 1.000 | 0.325 | 6027 | tags=13%, list=10%, signal=14% |
| 89 | LONG\_CHAIN\_FATTY\_ACID\_METABOLIC\_PROCESS(GO:0001676) |  | 99 | 0.88 | 1.40 | 0.000 | 1.000 | 0.325 | 6048 | tags=12%, list=10%, signal=13% |
| 90 | LEUKOCYTE\_CELL\_CELL\_ADHESION(GO:0007159) |  | 323 | 0.87 | 1.40 | 0.000 | 1.000 | 0.325 | 6178 | tags=11%, list=10%, signal=12% |
| 91 | REGULATION\_OF\_EARLY\_ENDOSOME\_TO\_LATE\_ENDOSOME\_TRANSPORT(GO:2000641) |  | 17 | 0.90 | 1.40 | 0.000 | 1.000 | 0.325 | 5823 | tags=6%, list=10%, signal=7% |
| 92 | REGULATION\_OF\_RECEPTOR\_RECYCLING(GO:0001919) |  | 23 | 0.87 | 1.40 | 0.000 | 1.000 | 0.325 | 6259 | tags=17%, list=11%, signal=19% |
| 93 | CELLULAR\_OXIDANT\_DETOXIFICATION(GO:0098869) |  | 97 | 0.88 | 1.40 | 0.000 | 1.000 | 0.325 | 6560 | tags=14%, list=11%, signal=16% |
| 94 | NEGATIVE\_REGULATION\_OF\_MYELOID\_CELL\_DIFFERENTIATION(GO:0045638) |  | 86 | 0.88 | 1.40 | 0.000 | 1.000 | 0.325 | 6294 | tags=13%, list=11%, signal=14% |
| 95 | CELLULAR\_PROTEIN\_CATABOLIC\_PROCESS(GO:0044257) |  | 723 | 0.85 | 1.40 | 0.000 | 1.000 | 0.325 | 6270 | tags=4%, list=11%, signal=5% |
| 96 | CELLULAR\_RESPONSE\_TO\_HORMONE\_STIMULUS(GO:0032870) |  | 693 | 0.87 | 1.40 | 0.000 | 1.000 | 0.325 | 6381 | tags=10%, list=11%, signal=12% |
| 97 | NUCLEOTIDE\_KINASE\_ACTIVITY(GO:0019201) |  | 26 | 0.87 | 1.40 | 0.000 | 1.000 | 0.325 | 5849 | tags=8%, list=10%, signal=9% |
| 98 | POSITIVE\_REGULATION\_OF\_MULTICELLULAR\_ORGANISM\_GROWTH(GO:0040018) |  | 36 | 0.88 | 1.40 | 0.000 | 1.000 | 0.325 | 5936 | tags=14%, list=10%, signal=15% |
| 99 | NEGATIVE\_REGULATION\_OF\_TUMOR\_NECROSIS\_FACTOR\_MEDIATED\_SIGNALING\_PATHWAY(GO:0010804) |  | 21 | 0.90 | 1.40 | 0.000 | 1.000 | 0.325 | 5922 | tags=10%, list=10%, signal=11% |
| 100 | REGULATION\_OF\_ACUTE\_INFLAMMATORY\_RESPONSE(GO:0002673) |  | 148 | 0.89 | 1.40 | 0.000 | 1.000 | 0.325 | 5825 | tags=7%, list=10%, signal=8% |
| 101 | CELLULAR\_RESPONSE\_TO\_LIPOPOLYSACCHARIDE(GO:0071222) |  | 162 | 0.87 | 1.40 | 0.000 | 1.000 | 0.325 | 6247 | tags=11%, list=11%, signal=12% |
| 102 | CELLULAR\_DETOXIFICATION(GO:1990748) |  | 101 | 0.88 | 1.40 | 0.000 | 1.000 | 0.325 | 6560 | tags=14%, list=11%, signal=16% |
| 103 | CORTICAL\_ACTIN\_CYTOSKELETON\_ORGANIZATION(GO:0030866) |  | 30 | 0.87 | 1.40 | 0.000 | 1.000 | 0.325 | 5836 | tags=7%, list=10%, signal=7% |
| 104 | ANTIOXIDANT\_ACTIVITY(GO:0016209) |  | 84 | 0.89 | 1.40 | 0.000 | 1.000 | 0.325 | 5834 | tags=11%, list=10%, signal=12% |
| 105 | CELLULAR\_RESPONSE\_TO\_BIOTIC\_STIMULUS(GO:0071216) |  | 192 | 0.87 | 1.40 | 0.000 | 1.000 | 0.325 | 6530 | tags=11%, list=11%, signal=12% |
| 106 | ESTABLISHMENT\_OF\_CELL\_POLARITY(GO:0030010) |  | 122 | 0.88 | 1.40 | 0.000 | 1.000 | 0.325 | 6167 | tags=6%, list=10%, signal=6% |
| 107 | CELLULAR\_RESPONSE\_TO\_ORGANONITROGEN\_COMPOUND(GO:0071417) |  | 538 | 0.87 | 1.40 | 0.000 | 1.000 | 0.325 | 6256 | tags=9%, list=11%, signal=10% |
| 108 | REGULATION\_OF\_CALCIUM\_ION\_TRANSMEMBRANE\_TRANSPORTER\_ACTIVITY(GO:1901019) |  | 89 | 0.85 | 1.40 | 0.000 | 1.000 | 0.325 | 6148 | tags=11%, list=10%, signal=13% |
| 109 | TUMOR\_NECROSIS\_FACTOR\_SUPERFAMILY\_CYTOKINE\_PRODUCTION(GO:0071706) |  | 119 | 0.88 | 1.40 | 0.000 | 1.000 | 0.325 | 5978 | tags=11%, list=10%, signal=12% |
| 110 | LINOLEIC\_ACID\_METABOLIC\_PROCESS(GO:0043651) |  | 17 | 0.90 | 1.40 | 0.000 | 1.000 | 0.325 | 5825 | tags=24%, list=10%, signal=26% |
| 111 | REGULATION\_OF\_PODOSOME\_ASSEMBLY(GO:0071801) |  | 15 | 0.90 | 1.40 | 0.000 | 1.000 | 0.325 | 5823 | tags=13%, list=10%, signal=15% |
| 112 | LYMPHOCYTE\_PROLIFERATION(GO:0046651) |  | 259 | 0.87 | 1.40 | 0.000 | 1.000 | 0.325 | 6178 | tags=9%, list=10%, signal=10% |
| 113 | GLUTATHIONE\_TRANSFERASE\_ACTIVITY(GO:0004364) |  | 26 | 0.89 | 1.39 | 0.000 | 1.000 | 0.325 | 5825 | tags=15%, list=10%, signal=17% |
| 114 | VITAMIN\_BIOSYNTHETIC\_PROCESS(GO:0009110) |  | 23 | 0.86 | 1.39 | 0.000 | 1.000 | 0.325 | 6495 | tags=13%, list=11%, signal=15% |
| 115 | CELL\_COMMUNICATION\_BY\_ELECTRICAL\_COUPLING(GO:0010644) |  | 28 | 0.87 | 1.39 | 0.000 | 1.000 | 0.325 | 6507 | tags=29%, list=11%, signal=32% |
| 116 | LAMIN\_BINDING(GO:0005521) |  | 15 | 0.86 | 1.39 | 0.000 | 1.000 | 0.325 | 7184 | tags=20%, list=12%, signal=23% |
| 117 | CELLULAR\_RESPONSE\_TO\_AMINO\_ACID\_STIMULUS(GO:0071230) |  | 64 | 0.87 | 1.39 | 0.000 | 1.000 | 0.325 | 6662 | tags=17%, list=11%, signal=19% |
| 118 | MONONUCLEAR\_CELL\_PROLIFERATION(GO:0032943) |  | 261 | 0.87 | 1.39 | 0.000 | 1.000 | 0.325 | 6178 | tags=9%, list=10%, signal=10% |
| 119 | LEFT\_RIGHT\_PATTERN\_FORMATION(GO:0060972) |  | 21 | 0.88 | 1.39 | 0.000 | 1.000 | 0.325 | 5833 | tags=10%, list=10%, signal=11% |
| 120 | REGULATION\_OF\_TUMOR\_NECROSIS\_FACTOR\_SUPERFAMILY\_CYTOKINE\_PRODUCTION(GO:1903555) |  | 115 | 0.88 | 1.39 | 0.000 | 1.000 | 0.325 | 5978 | tags=10%, list=10%, signal=12% |
| 121 | ENDOTHELIUM\_DEVELOPMENT(GO:0003158) |  | 121 | 0.89 | 1.39 | 0.000 | 1.000 | 0.325 | 5959 | tags=15%, list=10%, signal=17% |
| 122 | REGULATION\_OF\_LIPID\_METABOLIC\_PROCESS(GO:0019216) |  | 365 | 0.86 | 1.39 | 0.000 | 1.000 | 0.325 | 6218 | tags=7%, list=11%, signal=8% |
| 123 | RESPONSE\_TO\_AMYLOID\_BETA(GO:1904645) |  | 27 | 0.87 | 1.39 | 0.000 | 1.000 | 0.325 | 6662 | tags=15%, list=11%, signal=17% |
| 124 | PEROXIDASE\_ACTIVITY(GO:0004601) |  | 44 | 0.89 | 1.39 | 0.000 | 1.000 | 0.325 | 5834 | tags=16%, list=10%, signal=18% |
| 125 | CELLULAR\_RESPONSE\_TO\_MOLECULE\_OF\_BACTERIAL\_ORIGIN(GO:0071219) |  | 170 | 0.87 | 1.39 | 0.000 | 1.000 | 0.325 | 6530 | tags=12%, list=11%, signal=14% |
| 126 | LYMPH\_NODE\_DEVELOPMENT(GO:0048535) |  | 16 | 0.87 | 1.39 | 0.000 | 1.000 | 0.325 | 5877 | tags=6%, list=10%, signal=7% |
| 127 | REGULATION\_OF\_TUMOR\_NECROSIS\_FACTOR\_PRODUCTION(GO:0032680) |  | 111 | 0.88 | 1.39 | 0.000 | 1.000 | 0.325 | 5978 | tags=10%, list=10%, signal=11% |
| 128 | CHEMOKINE\_PRODUCTION(GO:0032602) |  | 83 | 0.88 | 1.39 | 0.000 | 1.000 | 0.325 | 6495 | tags=16%, list=11%, signal=18% |
| 129 | UNSATURATED\_FATTY\_ACID\_METABOLIC\_PROCESS(GO:0033559) |  | 101 | 0.88 | 1.39 | 0.000 | 1.000 | 0.325 | 6048 | tags=11%, list=10%, signal=12% |
| 130 | CELLULAR\_RESPONSE\_TO\_AMYLOID\_BETA(GO:1904646) |  | 25 | 0.88 | 1.39 | 0.000 | 1.000 | 0.325 | 6413 | tags=8%, list=11%, signal=9% |
| 131 | REGULATION\_OF\_SUPEROXIDE\_METABOLIC\_PROCESS(GO:0090322) |  | 31 | 0.89 | 1.39 | 0.000 | 1.000 | 0.325 | 6169 | tags=6%, list=10%, signal=7% |
| 132 | ENDOCRINE\_PANCREAS\_DEVELOPMENT(GO:0031018) |  | 45 | 0.87 | 1.39 | 0.000 | 1.000 | 0.325 | 6473 | tags=24%, list=11%, signal=27% |
| 133 | TUMOR\_NECROSIS\_FACTOR\_PRODUCTION(GO:0032640) |  | 113 | 0.88 | 1.39 | 0.000 | 1.000 | 0.325 | 5978 | tags=10%, list=10%, signal=11% |
| 134 | REGULATION\_OF\_ACTION\_POTENTIAL(GO:0098900) |  | 52 | 0.87 | 1.39 | 0.000 | 1.000 | 0.325 | 6670 | tags=17%, list=11%, signal=19% |
| 135 | AMIDE\_BINDING(GO:0033218) |  | 299 | 0.85 | 1.39 | 0.000 | 1.000 | 0.325 | 7005 | tags=15%, list=12%, signal=17% |
| 136 | SULFUR\_AMINO\_ACID\_BIOSYNTHETIC\_PROCESS(GO:0000097) |  | 17 | 0.85 | 1.39 | 0.000 | 1.000 | 0.325 | 6628 | tags=24%, list=11%, signal=26% |
| 137 | PEPTIDE\_BINDING(GO:0042277) |  | 266 | 0.86 | 1.39 | 0.000 | 1.000 | 0.325 | 7005 | tags=16%, list=12%, signal=18% |
| 138 | CELLULAR\_RESPONSE\_TO\_EPIDERMAL\_GROWTH\_FACTOR\_STIMULUS(GO:0071364) |  | 41 | 0.89 | 1.39 | 0.000 | 1.000 | 0.325 | 5825 | tags=2%, list=10%, signal=3% |
| 139 | ENDOTHELIAL\_CELL\_DIFFERENTIATION(GO:0045446) |  | 104 | 0.89 | 1.39 | 0.000 | 1.000 | 0.325 | 5933 | tags=12%, list=10%, signal=13% |
| 140 | POLYOL\_METABOLIC\_PROCESS(GO:0019751) |  | 113 | 0.84 | 1.39 | 0.000 | 1.000 | 0.325 | 6153 | tags=12%, list=10%, signal=13% |
| 141 | CELL\_COMMUNICATION\_BY\_ELECTRICAL\_COUPLING\_INVOLVED\_IN\_CARDIAC\_CONDUCTION(GO:0086064) |  | 23 | 0.87 | 1.39 | 0.000 | 1.000 | 0.325 | 6507 | tags=30%, list=11%, signal=34% |
| 142 | CELLULAR\_RESPONSE\_TO\_EXTRACELLULAR\_STIMULUS(GO:0031668) |  | 235 | 0.86 | 1.39 | 0.000 | 1.000 | 0.325 | 6497 | tags=9%, list=11%, signal=10% |
| 143 | NEGATIVE\_REGULATION\_OF\_DOUBLE\_STRAND\_BREAK\_REPAIR(GO:2000780) |  | 23 | 0.87 | 1.39 | 0.000 | 1.000 | 0.325 | 5928 | tags=17%, list=10%, signal=19% |
| 144 | NEGATIVE\_REGULATION\_OF\_HEMOPOIESIS(GO:1903707) |  | 141 | 0.87 | 1.39 | 0.000 | 1.000 | 0.325 | 6294 | tags=13%, list=11%, signal=14% |
| 145 | RESPONSE\_TO\_EPIDERMAL\_GROWTH\_FACTOR(GO:0070849) |  | 45 | 0.89 | 1.39 | 0.000 | 1.000 | 0.325 | 5825 | tags=2%, list=10%, signal=2% |
| 146 | DETOXIFICATION(GO:0098754) |  | 106 | 0.88 | 1.39 | 0.000 | 1.000 | 0.325 | 6560 | tags=13%, list=11%, signal=15% |
| 147 | G\_PROTEIN\_BETA\_GAMMA\_SUBUNIT\_COMPLEX\_BINDING(GO:0031683) |  | 21 | 0.88 | 1.39 | 0.000 | 1.000 | 0.325 | 6471 | tags=14%, list=11%, signal=16% |
| 148 | REGULATION\_OF\_CYTOKINE\_MEDIATED\_SIGNALING\_PATHWAY(GO:0001959) |  | 169 | 0.87 | 1.39 | 0.000 | 1.000 | 0.325 | 5978 | tags=6%, list=10%, signal=7% |
| 149 | POSITIVE\_REGULATION\_OF\_ATP\_METABOLIC\_PROCESS(GO:1903580) |  | 33 | 0.86 | 1.39 | 0.000 | 1.000 | 0.325 | 6006 | tags=9%, list=10%, signal=10% |
| 150 | TRIGLYCERIDE\_METABOLIC\_PROCESS(GO:0006641) |  | 98 | 0.85 | 1.39 | 0.000 | 1.000 | 0.325 | 6622 | tags=19%, list=11%, signal=22% |
| 151 | REGULATION\_OF\_FATTY\_ACID\_BIOSYNTHETIC\_PROCESS(GO:0042304) |  | 41 | 0.83 | 1.39 | 0.000 | 1.000 | 0.325 | 7226 | tags=17%, list=12%, signal=19% |
| 152 | REGULATION\_OF\_RESPONSE\_TO\_CYTOKINE\_STIMULUS(GO:0060759) |  | 177 | 0.87 | 1.39 | 0.000 | 1.000 | 0.325 | 5978 | tags=6%, list=10%, signal=6% |
| 153 | ELECTRON\_TRANSFER\_ACTIVITY(GO:0009055) |  | 115 | 0.83 | 1.39 | 0.000 | 1.000 | 0.325 | 5920 | tags=5%, list=10%, signal=6% |
| 154 | INFLAMMATORY\_RESPONSE\_TO\_ANTIGENIC\_STIMULUS(GO:0002437) |  | 44 | 0.89 | 1.39 | 0.000 | 1.000 | 0.325 | 5834 | tags=7%, list=10%, signal=8% |
| 155 | NEUROMUSCULAR\_JUNCTION\_DEVELOPMENT(GO:0007528) |  | 42 | 0.88 | 1.39 | 0.000 | 1.000 | 0.325 | 6091 | tags=10%, list=10%, signal=11% |
| 156 | ESTABLISHMENT\_OR\_MAINTENANCE\_OF\_CELL\_POLARITY(GO:0007163) |  | 186 | 0.87 | 1.39 | 0.000 | 1.000 | 0.325 | 6374 | tags=8%, list=11%, signal=8% |
| 157 | NEGATIVE\_REGULATION\_OF\_ERK1\_AND\_ERK2\_CASCADE(GO:0070373) |  | 68 | 0.89 | 1.39 | 0.000 | 1.000 | 0.325 | 5856 | tags=4%, list=10%, signal=5% |
| 158 | POSITIVE\_REGULATION\_OF\_TRANSPORTER\_ACTIVITY(GO:0032411) |  | 95 | 0.85 | 1.39 | 0.000 | 1.000 | 0.325 | 7441 | tags=22%, list=13%, signal=25% |
| 159 | ANIMAL\_ORGAN\_REGENERATION(GO:0031100) |  | 79 | 0.88 | 1.39 | 0.000 | 1.000 | 0.325 | 5825 | tags=5%, list=10%, signal=6% |
| 160 | REGULATION\_OF\_VOLTAGE\_GATED\_CALCIUM\_CHANNEL\_ACTIVITY(GO:1901385) |  | 37 | 0.88 | 1.39 | 0.000 | 1.000 | 0.325 | 6148 | tags=16%, list=10%, signal=18% |
| 161 | CELL\_SUBSTRATE\_JUNCTION(GO:0030055) |  | 401 | 0.87 | 1.39 | 0.000 | 1.000 | 0.325 | 6626 | tags=9%, list=11%, signal=10% |
| 162 | NEUTRAL\_LIPID\_METABOLIC\_PROCESS(GO:0006638) |  | 115 | 0.85 | 1.39 | 0.000 | 1.000 | 0.325 | 6873 | tags=18%, list=12%, signal=21% |
| 163 | WIDE\_PORE\_CHANNEL\_ACTIVITY(GO:0022829) |  | 23 | 0.88 | 1.39 | 0.000 | 1.000 | 0.325 | 5959 | tags=22%, list=10%, signal=24% |
| 164 | POSITIVE\_REGULATION\_OF\_ION\_TRANSMEMBRANE\_TRANSPORTER\_ACTIVITY(GO:0032414) |  | 86 | 0.85 | 1.39 | 0.000 | 1.000 | 0.325 | 7441 | tags=23%, list=13%, signal=27% |
| 165 | FOCAL\_ADHESION(GO:0005925) |  | 394 | 0.87 | 1.39 | 0.000 | 1.000 | 0.325 | 6626 | tags=8%, list=11%, signal=9% |
| 166 | SUPEROXIDE\_METABOLIC\_PROCESS(GO:0006801) |  | 64 | 0.88 | 1.39 | 0.000 | 1.000 | 0.325 | 6247 | tags=11%, list=11%, signal=12% |
| 167 | CELL\_SUBSTRATE\_ADHERENS\_JUNCTION(GO:0005924) |  | 397 | 0.87 | 1.39 | 0.000 | 1.000 | 0.325 | 6626 | tags=9%, list=11%, signal=10% |
| 168 | INCLUSION\_BODY(GO:0016234) |  | 81 | 0.85 | 1.39 | 0.000 | 1.000 | 0.325 | 5894 | tags=7%, list=10%, signal=8% |
| 169 | RESPONSE\_TO\_PEPTIDE(GO:1901652) |  | 486 | 0.87 | 1.39 | 0.000 | 1.000 | 0.325 | 6256 | tags=9%, list=11%, signal=10% |
| 170 | NEGATIVE\_REGULATION\_OF\_TUMOR\_NECROSIS\_FACTOR\_SUPERFAMILY\_CYTOKINE\_PRODUCTION(GO:1903556) |  | 48 | 0.89 | 1.39 | 0.000 | 1.000 | 0.325 | 5904 | tags=13%, list=10%, signal=14% |
| 171 | PROTEIN\_CATABOLIC\_PROCESS(GO:0030163) |  | 870 | 0.84 | 1.38 | 0.000 | 1.000 | 0.325 | 6270 | tags=4%, list=11%, signal=5% |
| 172 | INORGANIC\_CATION\_IMPORT\_ACROSS\_PLASMA\_MEMBRANE(GO:0098659) |  | 49 | 0.86 | 1.38 | 0.000 | 1.000 | 0.325 | 6662 | tags=20%, list=11%, signal=23% |
| 173 | INORGANIC\_ION\_IMPORT\_ACROSS\_PLASMA\_MEMBRANE(GO:0099587) |  | 49 | 0.86 | 1.38 | 0.000 | 1.000 | 0.325 | 6662 | tags=20%, list=11%, signal=23% |
| 174 | CARDIAC\_ATRIUM\_MORPHOGENESIS(GO:0003209) |  | 31 | 0.87 | 1.38 | 0.000 | 1.000 | 0.325 | 6473 | tags=23%, list=11%, signal=25% |
| 175 | HYPEROSMOTIC\_RESPONSE(GO:0006972) |  | 23 | 0.88 | 1.38 | 0.000 | 1.000 | 0.325 | 6374 | tags=9%, list=11%, signal=10% |
| 176 | RESPONSE\_TO\_PEPTIDE\_HORMONE(GO:0043434) |  | 419 | 0.87 | 1.38 | 0.000 | 1.000 | 0.325 | 6256 | tags=8%, list=11%, signal=9% |
| 177 | TYPE\_B\_PANCREATIC\_CELL\_DIFFERENTIATION(GO:0003309) |  | 24 | 0.88 | 1.38 | 0.000 | 1.000 | 0.325 | 6473 | tags=33%, list=11%, signal=37% |
| 178 | NEGATIVE\_REGULATION\_OF\_DNA\_REPAIR(GO:0045738) |  | 25 | 0.86 | 1.38 | 0.000 | 1.000 | 0.325 | 5928 | tags=16%, list=10%, signal=18% |
| 179 | POSITIVE\_REGULATION\_OF\_ATP\_BIOSYNTHETIC\_PROCESS(GO:2001171) |  | 27 | 0.87 | 1.38 | 0.000 | 1.000 | 0.325 | 6006 | tags=11%, list=10%, signal=12% |
| 180 | REGULATION\_OF\_VENTRICULAR\_CARDIAC\_MUSCLE\_CELL\_MEMBRANE\_REPOLARIZATION(GO:0060307) |  | 24 | 0.88 | 1.38 | 0.000 | 1.000 | 0.325 | 6412 | tags=33%, list=11%, signal=37% |
| 181 | RENAL\_WATER\_HOMEOSTASIS(GO:0003091) |  | 36 | 0.88 | 1.38 | 0.000 | 1.000 | 0.325 | 6374 | tags=14%, list=11%, signal=16% |
| 182 | T\_CELL\_ACTIVATION(GO:0042110) |  | 448 | 0.86 | 1.38 | 0.000 | 1.000 | 0.325 | 6774 | tags=13%, list=11%, signal=14% |
| 183 | CELLULAR\_RESPONSE\_TO\_OXYGEN\_CONTAINING\_COMPOUND(GO:1901701) |  | 1036 | 0.87 | 1.38 | 0.000 | 1.000 | 0.325 | 6279 | tags=10%, list=11%, signal=11% |
| 184 | CELLULAR\_BIOGENIC\_AMINE\_BIOSYNTHETIC\_PROCESS(GO:0042401) |  | 22 | 0.85 | 1.38 | 0.000 | 1.000 | 0.325 | 7268 | tags=14%, list=12%, signal=16% |
| 185 | INTERLEUKIN\_1\_PRODUCTION(GO:0032612) |  | 83 | 0.88 | 1.38 | 0.000 | 1.000 | 0.325 | 6486 | tags=10%, list=11%, signal=11% |
| 186 | CELLULAR\_RESPONSE\_TO\_NITROGEN\_COMPOUND(GO:1901699) |  | 637 | 0.87 | 1.38 | 0.000 | 1.000 | 0.325 | 6560 | tags=10%, list=11%, signal=11% |
| 187 | WNT\_PROTEIN\_BINDING(GO:0017147) |  | 30 | 0.87 | 1.38 | 0.000 | 1.000 | 0.325 | 6677 | tags=27%, list=11%, signal=30% |
| 188 | POSITIVE\_REGULATION\_OF\_PROTEIN\_LOCALIZATION\_TO\_NUCLEUS(GO:1900182) |  | 67 | 0.86 | 1.38 | 0.000 | 1.000 | 0.325 | 6662 | tags=7%, list=11%, signal=8% |
| 189 | NEGATIVE\_REGULATION\_OF\_TUMOR\_NECROSIS\_FACTOR\_PRODUCTION(GO:0032720) |  | 46 | 0.89 | 1.38 | 0.000 | 1.000 | 0.325 | 5904 | tags=13%, list=10%, signal=14% |
| 190 | POSITIVE\_REGULATION\_OF\_G1\_S\_TRANSITION\_OF\_MITOTIC\_CELL\_CYCLE(GO:1900087) |  | 33 | 0.86 | 1.38 | 0.000 | 1.000 | 0.325 | 5852 | tags=9%, list=10%, signal=10% |
| 191 | BASEMENT\_MEMBRANE\_ORGANIZATION(GO:0071711) |  | 21 | 0.89 | 1.38 | 0.000 | 1.000 | 0.325 | 6186 | tags=19%, list=10%, signal=21% |
| 192 | NEGATIVE\_REGULATION\_OF\_DEFENSE\_RESPONSE(GO:0031348) |  | 177 | 0.87 | 1.38 | 0.000 | 1.000 | 0.325 | 6048 | tags=10%, list=10%, signal=11% |
| 193 | ENDOTHELIAL\_CELL\_DEVELOPMENT(GO:0001885) |  | 56 | 0.90 | 1.38 | 0.000 | 1.000 | 0.325 | 5856 | tags=9%, list=10%, signal=10% |
| 194 | NEGATIVE\_REGULATION\_OF\_LEUKOCYTE\_PROLIFERATION(GO:0070664) |  | 72 | 0.88 | 1.38 | 0.000 | 1.000 | 0.325 | 6154 | tags=6%, list=10%, signal=6% |
| 195 | RESPONSE\_TO\_ESTRADIOL(GO:0032355) |  | 128 | 0.88 | 1.38 | 0.000 | 1.000 | 0.325 | 6424 | tags=14%, list=11%, signal=16% |
| 196 | REGULATION\_OF\_INFLAMMATORY\_RESPONSE(GO:0050727) |  | 390 | 0.86 | 1.38 | 0.000 | 1.000 | 0.325 | 6520 | tags=12%, list=11%, signal=13% |
| 197 | CELLULAR\_RESPONSE\_TO\_KETONE(GO:1901655) |  | 86 | 0.89 | 1.38 | 0.000 | 1.000 | 0.325 | 6178 | tags=13%, list=10%, signal=14% |
| 198 | GLUTATHIONE\_METABOLIC\_PROCESS(GO:0006749) |  | 54 | 0.89 | 1.38 | 0.141 | 1.000 | 0.325 | 5834 | tags=7%, list=10%, signal=8% |
| 199 | RESPONSE\_TO\_REACTIVE\_OXYGEN\_SPECIES(GO:0000302) |  | 226 | 0.86 | 1.38 | 0.000 | 1.000 | 0.325 | 6507 | tags=8%, list=11%, signal=9% |
| 200 | POLARIZED\_EPITHELIAL\_CELL\_DIFFERENTIATION(GO:0030859) |  | 18 | 0.90 | 1.38 | 0.000 | 1.000 | 0.325 | 5986 | tags=11%, list=10%, signal=12% |
| 201 | CELLULAR\_RESPONSE\_TO\_LIPID(GO:0071396) |  | 572 | 0.87 | 1.38 | 0.000 | 1.000 | 0.325 | 6381 | tags=11%, list=11%, signal=13% |
| 202 | HYDROGEN\_PEROXIDE\_CATABOLIC\_PROCESS(GO:0042744) |  | 22 | 0.90 | 1.38 | 0.000 | 1.000 | 0.325 | 5894 | tags=18%, list=10%, signal=20% |
| 203 | ACUTE\_INFLAMMATORY\_RESPONSE(GO:0002526) |  | 212 | 0.87 | 1.38 | 0.000 | 1.000 | 0.325 | 6479 | tags=8%, list=11%, signal=10% |
| 204 | ENTEROENDOCRINE\_CELL\_DIFFERENTIATION(GO:0035883) |  | 26 | 0.88 | 1.38 | 0.000 | 1.000 | 0.325 | 6473 | tags=35%, list=11%, signal=39% |
| 205 | SKELETAL\_MUSCLE\_TISSUE\_REGENERATION(GO:0043403) |  | 34 | 0.88 | 1.38 | 0.000 | 1.000 | 0.325 | 6209 | tags=24%, list=10%, signal=26% |
| 206 | MODIFIED\_AMINO\_ACID\_BINDING(GO:0072341) |  | 69 | 0.89 | 1.38 | 0.000 | 1.000 | 0.325 | 6167 | tags=12%, list=10%, signal=13% |
| 207 | RIBONUCLEOSIDE\_CATABOLIC\_PROCESS(GO:0042454) |  | 20 | 0.83 | 1.38 | 0.000 | 1.000 | 0.325 | 6865 | tags=20%, list=12%, signal=23% |
| 208 | CELL\_REDOX\_HOMEOSTASIS(GO:0045454) |  | 73 | 0.85 | 1.38 | 0.000 | 1.000 | 0.325 | 6672 | tags=7%, list=11%, signal=8% |
| 209 | REGENERATION(GO:0031099) |  | 190 | 0.87 | 1.38 | 0.000 | 1.000 | 0.325 | 6292 | tags=13%, list=11%, signal=15% |
| 210 | DOUBLE\_STRANDED\_RNA\_BINDING(GO:0003725) |  | 67 | 0.90 | 1.38 | 0.000 | 1.000 | 0.325 | 5823 | tags=3%, list=10%, signal=3% |
| 211 | ACYLGLYCEROL\_METABOLIC\_PROCESS(GO:0006639) |  | 114 | 0.84 | 1.38 | 0.000 | 1.000 | 0.325 | 6873 | tags=18%, list=12%, signal=20% |
| 212 | MEMBRANE\_REPOLARIZATION(GO:0086009) |  | 48 | 0.87 | 1.38 | 0.000 | 1.000 | 0.325 | 6784 | tags=23%, list=11%, signal=26% |
| 213 | REGULATION\_OF\_URINE\_VOLUME(GO:0035809) |  | 21 | 0.88 | 1.38 | 0.000 | 1.000 | 0.325 | 6525 | tags=19%, list=11%, signal=21% |
| 214 | PROTEIN\_BINDING\_\_BRIDGING(GO:0030674) |  | 170 | 0.88 | 1.38 | 0.000 | 1.000 | 0.325 | 5949 | tags=8%, list=10%, signal=8% |
| 215 | REACTIVE\_OXYGEN\_SPECIES\_METABOLIC\_PROCESS(GO:0072593) |  | 248 | 0.87 | 1.38 | 0.000 | 1.000 | 0.325 | 6514 | tags=11%, list=11%, signal=12% |
| 216 | CARDIAC\_MUSCLE\_CELL\_CONTRACTION(GO:0086003) |  | 63 | 0.87 | 1.38 | 0.000 | 1.000 | 0.325 | 6569 | tags=24%, list=11%, signal=27% |
| 217 | POSITIVE\_REGULATION\_OF\_PROTEIN\_COMPLEX\_ASSEMBLY(GO:0031334) |  | 251 | 0.87 | 1.38 | 0.000 | 1.000 | 0.325 | 6115 | tags=7%, list=10%, signal=8% |
| 218 | NCRNA\_CATABOLIC\_PROCESS(GO:0034661) |  | 27 | 0.85 | 1.38 | 0.000 | 1.000 | 0.325 | 5858 | tags=7%, list=10%, signal=8% |
| 219 | REGULATION\_OF\_REACTIVE\_OXYGEN\_SPECIES\_METABOLIC\_PROCESS(GO:2000377) |  | 167 | 0.87 | 1.38 | 0.000 | 1.000 | 0.325 | 6465 | tags=11%, list=11%, signal=12% |
| 220 | ACTION\_POTENTIAL(GO:0001508) |  | 140 | 0.86 | 1.38 | 0.000 | 1.000 | 0.325 | 6844 | tags=24%, list=12%, signal=27% |
| 221 | IMPORT\_ACROSS\_PLASMA\_MEMBRANE(GO:0098739) |  | 67 | 0.86 | 1.38 | 0.000 | 1.000 | 0.325 | 7073 | tags=21%, list=12%, signal=24% |
| 222 | CARDIAC\_MUSCLE\_CELL\_ACTION\_POTENTIAL(GO:0086001) |  | 70 | 0.87 | 1.38 | 0.000 | 1.000 | 0.325 | 6670 | tags=26%, list=11%, signal=29% |
| 223 | VENTRICULAR\_CARDIAC\_MUSCLE\_CELL\_MEMBRANE\_REPOLARIZATION(GO:0099625) |  | 28 | 0.88 | 1.38 | 0.000 | 1.000 | 0.325 | 6569 | tags=32%, list=11%, signal=36% |
| 224 | ACTIN\_MEDIATED\_CELL\_CONTRACTION(GO:0070252) |  | 110 | 0.87 | 1.38 | 0.000 | 1.000 | 0.325 | 6583 | tags=26%, list=11%, signal=30% |
| 225 | REGULATION\_OF\_LEUKOCYTE\_PROLIFERATION(GO:0070663) |  | 210 | 0.86 | 1.38 | 0.000 | 1.000 | 0.325 | 6178 | tags=9%, list=10%, signal=10% |
| 226 | CELLULAR\_RESPONSE\_TO\_TOXIC\_SUBSTANCE(GO:0097237) |  | 231 | 0.87 | 1.38 | 0.000 | 1.000 | 0.325 | 6560 | tags=10%, list=11%, signal=11% |
| 227 | BRANCHING\_INVOLVED\_IN\_BLOOD\_VESSEL\_MORPHOGENESIS(GO:0001569) |  | 33 | 0.88 | 1.38 | 0.000 | 1.000 | 0.325 | 5925 | tags=6%, list=10%, signal=7% |
| 228 | RECEPTOR\_RECYCLING(GO:0001881) |  | 33 | 0.86 | 1.38 | 0.000 | 1.000 | 0.325 | 6259 | tags=15%, list=11%, signal=17% |
| 229 | REGULATION\_OF\_CARDIAC\_MUSCLE\_CELL\_MEMBRANE\_REPOLARIZATION(GO:0099623) |  | 29 | 0.88 | 1.38 | 0.000 | 1.000 | 0.325 | 6412 | tags=28%, list=11%, signal=31% |
| 230 | POSITIVE\_REGULATION\_OF\_ORGANELLE\_ASSEMBLY(GO:1902117) |  | 66 | 0.89 | 1.38 | 0.000 | 1.000 | 0.325 | 5952 | tags=8%, list=10%, signal=8% |
| 231 | MOLECULAR\_ADAPTOR\_ACTIVITY(GO:0060090) |  | 189 | 0.88 | 1.38 | 0.000 | 1.000 | 0.325 | 5949 | tags=7%, list=10%, signal=8% |
| 232 | HYDROGEN\_PEROXIDE\_METABOLIC\_PROCESS(GO:0042743) |  | 44 | 0.88 | 1.38 | 0.000 | 1.000 | 0.325 | 5894 | tags=11%, list=10%, signal=13% |
| 233 | CELLULAR\_RESPONSE\_TO\_GLUCOCORTICOID\_STIMULUS(GO:0071385) |  | 57 | 0.88 | 1.38 | 0.000 | 1.000 | 0.325 | 6155 | tags=11%, list=10%, signal=12% |
| 234 | NEGATIVE\_REGULATION\_OF\_CYTOKINE\_MEDIATED\_SIGNALING\_PATHWAY(GO:0001960) |  | 52 | 0.89 | 1.38 | 0.000 | 1.000 | 0.325 | 5922 | tags=6%, list=10%, signal=6% |
| 235 | INTERLEUKIN\_1\_BETA\_PRODUCTION(GO:0032611) |  | 69 | 0.88 | 1.38 | 0.000 | 1.000 | 0.325 | 6486 | tags=10%, list=11%, signal=11% |
| 236 | CELLULAR\_RESPONSE\_TO\_CORTICOSTEROID\_STIMULUS(GO:0071384) |  | 60 | 0.88 | 1.38 | 0.000 | 1.000 | 0.325 | 6155 | tags=10%, list=10%, signal=11% |
| 237 | ESTABLISHMENT\_OF\_ENDOTHELIAL\_BARRIER(GO:0061028) |  | 40 | 0.90 | 1.38 | 0.000 | 1.000 | 0.325 | 5823 | tags=5%, list=10%, signal=6% |
| 238 | RESPONSE\_TO\_MUSCLE\_ACTIVITY(GO:0014850) |  | 21 | 0.85 | 1.38 | 0.000 | 1.000 | 0.325 | 5861 | tags=19%, list=10%, signal=21% |
| 239 | TISSUE\_REGENERATION(GO:0042246) |  | 67 | 0.88 | 1.38 | 0.000 | 1.000 | 0.325 | 6292 | tags=18%, list=11%, signal=20% |
| 240 | CEREBELLUM\_DEVELOPMENT(GO:0021549) |  | 99 | 0.84 | 1.38 | 0.000 | 1.000 | 0.325 | 7546 | tags=24%, list=13%, signal=28% |
| 241 | LEUKOCYTE\_MIGRATION(GO:0050900) |  | 464 | 0.87 | 1.38 | 0.141 | 1.000 | 0.325 | 6473 | tags=11%, list=11%, signal=12% |
| 242 | PHOSPHATIDYLINOSITOL\_3\_KINASE\_BINDING(GO:0043548) |  | 29 | 0.89 | 1.38 | 0.000 | 1.000 | 0.325 | 5824 | tags=7%, list=10%, signal=8% |
| 243 | REGULATION\_OF\_LYMPHOCYTE\_MIGRATION(GO:2000401) |  | 47 | 0.90 | 1.38 | 0.000 | 1.000 | 0.325 | 5823 | tags=13%, list=10%, signal=14% |
| 244 | REGULATION\_OF\_LEUKOCYTE\_MIGRATION(GO:0002685) |  | 173 | 0.89 | 1.38 | 0.141 | 1.000 | 0.325 | 6011 | tags=13%, list=10%, signal=14% |
| 245 | T\_TUBULE(GO:0030315) |  | 44 | 0.87 | 1.38 | 0.000 | 1.000 | 0.325 | 6569 | tags=27%, list=11%, signal=31% |
| 246 | GLUTATHIONE\_DERIVATIVE\_BIOSYNTHETIC\_PROCESS(GO:1901687) |  | 21 | 0.89 | 1.37 | 0.000 | 1.000 | 0.325 | 5825 | tags=19%, list=10%, signal=21% |
| 247 | GLUTATHIONE\_DERIVATIVE\_METABOLIC\_PROCESS(GO:1901685) |  | 21 | 0.89 | 1.37 | 0.000 | 1.000 | 0.325 | 5825 | tags=19%, list=10%, signal=21% |
| 248 | RESPONSE\_TO\_MONOSACCHARIDE(GO:0034284) |  | 198 | 0.86 | 1.37 | 0.000 | 1.000 | 0.325 | 6578 | tags=12%, list=11%, signal=13% |
| 249 | INTERCALATED\_DISC(GO:0014704) |  | 52 | 0.87 | 1.37 | 0.000 | 1.000 | 0.325 | 6573 | tags=25%, list=11%, signal=28% |
| 250 | CARDIAC\_MUSCLE\_CELL\_ACTION\_POTENTIAL\_INVOLVED\_IN\_CONTRACTION(GO:0086002) |  | 53 | 0.87 | 1.37 | 0.000 | 1.000 | 0.325 | 6569 | tags=26%, list=11%, signal=30% |
| 251 | POSITIVE\_REGULATION\_OF\_PROTEIN\_IMPORT\_INTO\_NUCLEUS(GO:0042307) |  | 32 | 0.87 | 1.37 | 0.000 | 1.000 | 0.325 | 6642 | tags=13%, list=11%, signal=14% |
| 252 | SERINE\_FAMILY\_AMINO\_ACID\_BIOSYNTHETIC\_PROCESS(GO:0009070) |  | 19 | 0.86 | 1.37 | 0.000 | 1.000 | 0.325 | 6628 | tags=16%, list=11%, signal=18% |
| 253 | CILIARY\_TIP(GO:0097542) |  | 45 | 0.85 | 1.37 | 0.000 | 1.000 | 0.325 | 5943 | tags=4%, list=10%, signal=5% |
| 254 | RESPONSE\_TO\_CARBOHYDRATE(GO:0009743) |  | 223 | 0.86 | 1.37 | 0.000 | 1.000 | 0.325 | 6578 | tags=13%, list=11%, signal=14% |
| 255 | KINASE\_REGULATOR\_ACTIVITY(GO:0019207) |  | 215 | 0.87 | 1.37 | 0.000 | 1.000 | 0.325 | 6220 | tags=7%, list=11%, signal=8% |
| 256 | RESPONSE\_TO\_AMINO\_ACID(GO:0043200) |  | 111 | 0.88 | 1.37 | 0.000 | 1.000 | 0.325 | 5933 | tags=12%, list=10%, signal=13% |
| 257 | GUANOSINE\_CONTAINING\_COMPOUND\_METABOLIC\_PROCESS(GO:1901068) |  | 53 | 0.83 | 1.37 | 0.000 | 1.000 | 0.325 | 5849 | tags=4%, list=10%, signal=4% |
| 258 | SMOOTH\_MUSCLE\_CELL\_MIGRATION(GO:0014909) |  | 84 | 0.88 | 1.37 | 0.000 | 1.000 | 0.325 | 5889 | tags=11%, list=10%, signal=12% |
| 259 | CELLULAR\_RESPONSE\_TO\_ALCOHOL(GO:0097306) |  | 59 | 0.88 | 1.37 | 0.000 | 1.000 | 0.325 | 6178 | tags=12%, list=10%, signal=13% |
| 260 | CARBOXYPEPTIDASE\_ACTIVITY(GO:0004180) |  | 44 | 0.86 | 1.37 | 0.000 | 1.000 | 0.325 | 5838 | tags=11%, list=10%, signal=13% |
| 261 | CELL\_CELL\_RECOGNITION(GO:0009988) |  | 70 | 0.89 | 1.37 | 0.000 | 1.000 | 0.325 | 5998 | tags=10%, list=10%, signal=11% |
| 262 | REGULATION\_OF\_PROTEIN\_COMPLEX\_ASSEMBLY(GO:0043254) |  | 427 | 0.86 | 1.37 | 0.000 | 1.000 | 0.325 | 6240 | tags=6%, list=11%, signal=7% |
| 263 | DETECTION\_OF\_STIMULUS(GO:0051606) |  | 646 | 0.81 | 1.37 | 0.000 | 1.000 | 0.325 | 8258 | tags=12%, list=14%, signal=14% |
| 264 | RESPONSE\_TO\_TESTOSTERONE(GO:0033574) |  | 44 | 0.89 | 1.37 | 0.141 | 1.000 | 0.325 | 6182 | tags=16%, list=10%, signal=18% |
| 265 | NEGATIVE\_REGULATION\_OF\_PROTEIN\_SERINE\_THREONINE\_KINASE\_ACTIVITY(GO:0071901) |  | 140 | 0.88 | 1.37 | 0.141 | 1.000 | 0.325 | 6154 | tags=6%, list=10%, signal=6% |
| 266 | LYSOZYME\_ACTIVITY(GO:0003796) |  | 15 | 0.84 | 1.37 | 0.000 | 1.000 | 0.325 | 6109 | tags=13%, list=10%, signal=15% |
| 267 | COLLAGEN\_METABOLIC\_PROCESS(GO:0032963) |  | 124 | 0.87 | 1.37 | 0.000 | 1.000 | 0.325 | 6218 | tags=14%, list=11%, signal=15% |
| 268 | DORSAL\_VENTRAL\_AXIS\_SPECIFICATION(GO:0009950) |  | 16 | 0.86 | 1.37 | 0.000 | 1.000 | 0.325 | 6027 | tags=19%, list=10%, signal=21% |
| 269 | CELLULAR\_AMINO\_ACID\_BIOSYNTHETIC\_PROCESS(GO:0008652) |  | 83 | 0.83 | 1.37 | 0.000 | 1.000 | 0.325 | 6628 | tags=13%, list=11%, signal=15% |
| 270 | POSTSYNAPSE(GO:0098794) |  | 429 | 0.84 | 1.37 | 0.000 | 1.000 | 0.325 | 7487 | tags=22%, list=13%, signal=25% |
| 271 | MAINTENANCE\_OF\_PROTEIN\_LOCATION\_IN\_CELL(GO:0032507) |  | 68 | 0.85 | 1.37 | 0.000 | 1.000 | 0.325 | 6706 | tags=6%, list=11%, signal=7% |
| 272 | NEGATIVE\_REGULATION\_OF\_CELL\_AGING(GO:0090344) |  | 25 | 0.88 | 1.37 | 0.000 | 1.000 | 0.325 | 5916 | tags=16%, list=10%, signal=18% |
| 273 | LATERAL\_MESODERM\_DEVELOPMENT(GO:0048368) |  | 15 | 0.90 | 1.37 | 0.000 | 1.000 | 0.325 | 5986 | tags=27%, list=10%, signal=30% |
| 274 | CARDIAC\_MUSCLE\_CONTRACTION(GO:0060048) |  | 130 | 0.85 | 1.37 | 0.000 | 1.000 | 0.325 | 6832 | tags=18%, list=12%, signal=21% |
| 275 | XENOBIOTIC\_METABOLIC\_PROCESS(GO:0006805) |  | 115 | 0.88 | 1.37 | 0.000 | 1.000 | 0.325 | 6052 | tags=10%, list=10%, signal=11% |
| 276 | HISTONE\_DEACETYLASE\_BINDING(GO:0042826) |  | 109 | 0.83 | 1.37 | 0.000 | 1.000 | 0.325 | 7293 | tags=16%, list=12%, signal=18% |
| 277 | ACTIN\_BINDING(GO:0003779) |  | 408 | 0.87 | 1.37 | 0.000 | 1.000 | 0.325 | 6583 | tags=13%, list=11%, signal=15% |
| 278 | MEMBRANE\_DEPOLARIZATION\_DURING\_ACTION\_POTENTIAL(GO:0086010) |  | 48 | 0.87 | 1.37 | 0.000 | 1.000 | 0.325 | 6573 | tags=38%, list=11%, signal=42% |
| 279 | REGULATION\_OF\_SMOOTH\_MUSCLE\_CELL\_MIGRATION(GO:0014910) |  | 76 | 0.88 | 1.37 | 0.000 | 1.000 | 0.325 | 5825 | tags=11%, list=10%, signal=12% |
| 280 | NEGATIVE\_REGULATION\_OF\_I\_KAPPAB\_KINASE\_NF\_KAPPAB\_SIGNALING(GO:0043124) |  | 41 | 0.89 | 1.37 | 0.000 | 1.000 | 0.325 | 5825 | tags=5%, list=10%, signal=5% |
| 281 | NEGATIVE\_REGULATION\_OF\_RESPONSE\_TO\_CYTOKINE\_STIMULUS(GO:0060761) |  | 56 | 0.89 | 1.37 | 0.000 | 1.000 | 0.325 | 5922 | tags=5%, list=10%, signal=6% |
| 282 | RESPONSE\_TO\_HORMONE(GO:0009725) |  | 965 | 0.86 | 1.37 | 0.000 | 1.000 | 0.325 | 6381 | tags=11%, list=11%, signal=12% |
| 283 | CELL\_COMMUNICATION\_INVOLVED\_IN\_CARDIAC\_CONDUCTION(GO:0086065) |  | 54 | 0.87 | 1.37 | 0.000 | 1.000 | 0.325 | 6670 | tags=30%, list=11%, signal=33% |
| 284 | REGULATION\_OF\_CELL\_PROLIFERATION\_INVOLVED\_IN\_HEART\_MORPHOGENESIS(GO:2000136) |  | 18 | 0.88 | 1.37 | 0.000 | 1.000 | 0.325 | 5958 | tags=17%, list=10%, signal=19% |
| 285 | ELECTRON\_TRANSPORT\_CHAIN(GO:0022900) |  | 186 | 0.81 | 1.37 | 0.000 | 1.000 | 0.325 | 5920 | tags=4%, list=10%, signal=5% |
| 286 | REGULATION\_OF\_STEM\_CELL\_POPULATION\_MAINTENANCE(GO:2000036) |  | 27 | 0.89 | 1.37 | 0.000 | 1.000 | 0.325 | 6279 | tags=22%, list=11%, signal=25% |
| 287 | HETEROCHROMATIN\_ORGANIZATION(GO:0070828) |  | 16 | 0.85 | 1.37 | 0.000 | 1.000 | 0.325 | 6154 | tags=13%, list=10%, signal=14% |
| 288 | POSITIVE\_REGULATION\_OF\_SODIUM\_ION\_TRANSMEMBRANE\_TRANSPORT(GO:1902307) |  | 17 | 0.87 | 1.37 | 0.000 | 1.000 | 0.325 | 6432 | tags=24%, list=11%, signal=26% |
| 289 | CELL\_CELL\_CONTACT\_ZONE(GO:0044291) |  | 70 | 0.86 | 1.37 | 0.000 | 1.000 | 0.325 | 6573 | tags=21%, list=11%, signal=24% |
| 290 | PANCREAS\_DEVELOPMENT(GO:0031016) |  | 78 | 0.86 | 1.37 | 0.000 | 1.000 | 0.325 | 6585 | tags=26%, list=11%, signal=29% |
| 291 | REGULATION\_OF\_VASCULAR\_SMOOTH\_MUSCLE\_CELL\_PROLIFERATION(GO:1904705) |  | 57 | 0.89 | 1.37 | 0.000 | 1.000 | 0.325 | 5825 | tags=18%, list=10%, signal=19% |
| 292 | VASCULAR\_SMOOTH\_MUSCLE\_CELL\_PROLIFERATION(GO:1990874) |  | 57 | 0.89 | 1.37 | 0.000 | 1.000 | 0.325 | 5825 | tags=18%, list=10%, signal=19% |
| 293 | REGULATION\_OF\_CAMP\_MEDIATED\_SIGNALING(GO:0043949) |  | 43 | 0.86 | 1.37 | 0.000 | 1.000 | 0.325 | 6880 | tags=28%, list=12%, signal=32% |
| 294 | REGULATION\_OF\_CELL\_SIZE(GO:0008361) |  | 150 | 0.88 | 1.37 | 0.141 | 1.000 | 0.325 | 5994 | tags=8%, list=10%, signal=9% |
| 295 | MAINTENANCE\_OF\_LOCATION\_IN\_CELL(GO:0051651) |  | 87 | 0.84 | 1.37 | 0.000 | 1.000 | 0.325 | 6706 | tags=6%, list=11%, signal=6% |
| 296 | NEGATIVE\_REGULATION\_OF\_HISTONE\_ACETYLATION(GO:0035067) |  | 17 | 0.87 | 1.37 | 0.000 | 1.000 | 0.325 | 5904 | tags=12%, list=10%, signal=13% |
| 297 | REGULATION\_OF\_INTERLEUKIN\_1\_PRODUCTION(GO:0032652) |  | 73 | 0.88 | 1.37 | 0.000 | 1.000 | 0.325 | 6486 | tags=10%, list=11%, signal=11% |
| 298 | NEUTROPHIL\_DEGRANULATION(GO:0043312) |  | 480 | 0.84 | 1.37 | 0.000 | 1.000 | 0.325 | 6696 | tags=6%, list=11%, signal=7% |
| 299 | SECRETORY\_GRANULE\_LUMEN(GO:0034774) |  | 319 | 0.86 | 1.37 | 0.000 | 1.000 | 0.325 | 6479 | tags=8%, list=11%, signal=8% |
| 300 | NEUTROPHIL\_ACTIVATION\_INVOLVED\_IN\_IMMUNE\_RESPONSE(GO:0002283) |  | 483 | 0.84 | 1.37 | 0.000 | 1.000 | 0.325 | 6696 | tags=6%, list=11%, signal=7% |
| 301 | NEUTROPHIL\_MEDIATED\_IMMUNITY(GO:0002446) |  | 495 | 0.84 | 1.37 | 0.000 | 1.000 | 0.325 | 6696 | tags=6%, list=11%, signal=7% |
| 302 | POSITIVE\_REGULATION\_OF\_TRANSMEMBRANE\_TRANSPORT(GO:0034764) |  | 187 | 0.84 | 1.37 | 0.000 | 1.000 | 0.325 | 7441 | tags=19%, list=13%, signal=21% |
| 303 | BHLH\_TRANSCRIPTION\_FACTOR\_BINDING(GO:0043425) |  | 28 | 0.88 | 1.37 | 0.000 | 1.000 | 0.325 | 5958 | tags=14%, list=10%, signal=16% |
| 304 | CRANIOFACIAL\_SUTURE\_MORPHOGENESIS(GO:0097094) |  | 18 | 0.85 | 1.37 | 0.000 | 1.000 | 0.325 | 5904 | tags=11%, list=10%, signal=12% |
| 305 | APICAL\_PLASMA\_MEMBRANE(GO:0016324) |  | 298 | 0.87 | 1.37 | 0.141 | 1.000 | 0.325 | 6562 | tags=15%, list=11%, signal=17% |
| 306 | NEGATIVE\_REGULATION\_OF\_PROTEIN\_KINASE\_ACTIVITY(GO:0006469) |  | 260 | 0.86 | 1.37 | 0.000 | 1.000 | 0.325 | 6220 | tags=6%, list=11%, signal=7% |
| 307 | POSITIVE\_REGULATION\_OF\_PROTEIN\_IMPORT(GO:1904591) |  | 34 | 0.87 | 1.37 | 0.000 | 1.000 | 0.325 | 6642 | tags=12%, list=11%, signal=13% |
| 308 | MUSCLE\_CELL\_MIGRATION(GO:0014812) |  | 95 | 0.88 | 1.37 | 0.000 | 1.000 | 0.325 | 5889 | tags=11%, list=10%, signal=12% |
| 309 | NEUTROPHIL\_ACTIVATION(GO:0042119) |  | 493 | 0.84 | 1.37 | 0.000 | 1.000 | 0.325 | 6696 | tags=6%, list=11%, signal=7% |
| 310 | GRANULOCYTE\_ACTIVATION(GO:0036230) |  | 499 | 0.84 | 1.37 | 0.000 | 1.000 | 0.325 | 6696 | tags=6%, list=11%, signal=7% |
| 311 | CELLULAR\_MODIFIED\_AMINO\_ACID\_METABOLIC\_PROCESS(GO:0006575) |  | 182 | 0.88 | 1.37 | 0.000 | 1.000 | 0.325 | 5866 | tags=7%, list=10%, signal=8% |
| 312 | ALKALI\_METAL\_ION\_BINDING(GO:0031420) |  | 20 | 0.86 | 1.37 | 0.000 | 1.000 | 0.325 | 5913 | tags=25%, list=10%, signal=28% |
| 313 | CYTOPLASMIC\_VESICLE\_LUMEN(GO:0060205) |  | 336 | 0.86 | 1.37 | 0.000 | 1.000 | 0.325 | 6479 | tags=7%, list=11%, signal=8% |
| 314 | ALPHA\_AMINO\_ACID\_BIOSYNTHETIC\_PROCESS(GO:1901607) |  | 66 | 0.82 | 1.37 | 0.000 | 1.000 | 0.325 | 6628 | tags=14%, list=11%, signal=15% |
| 315 | REGULATION\_OF\_MITOTIC\_SPINDLE\_ORGANIZATION(GO:0060236) |  | 36 | 0.83 | 1.37 | 0.000 | 1.000 | 0.325 | 6626 | tags=6%, list=11%, signal=6% |
| 316 | POSITIVE\_REGULATION\_OF\_BRANCHING\_INVOLVED\_IN\_URETERIC\_BUD\_MORPHOGENESIS(GO:0090190) |  | 19 | 0.88 | 1.37 | 0.000 | 1.000 | 0.325 | 6279 | tags=16%, list=11%, signal=18% |
| 317 | CELLULAR\_RESPONSE\_TO\_ENDOGENOUS\_STIMULUS(GO:0071495) |  | 1344 | 0.86 | 1.37 | 0.000 | 1.000 | 0.325 | 6560 | tags=12%, list=11%, signal=13% |
| 318 | CELL\_DEATH\_IN\_RESPONSE\_TO\_OXIDATIVE\_STRESS(GO:0036473) |  | 90 | 0.85 | 1.37 | 0.000 | 1.000 | 0.325 | 6514 | tags=6%, list=11%, signal=6% |
| 319 | VESICLE\_LUMEN(GO:0031983) |  | 337 | 0.85 | 1.37 | 0.000 | 1.000 | 0.325 | 6479 | tags=7%, list=11%, signal=8% |
| 320 | STRUCTURAL\_MOLECULE\_ACTIVITY\_CONFERRING\_ELASTICITY(GO:0097493) |  | 17 | 0.89 | 1.37 | 0.000 | 1.000 | 0.325 | 6200 | tags=35%, list=10%, signal=39% |
| 321 | NEGATIVE\_REGULATION\_OF\_EXTRINSIC\_APOPTOTIC\_SIGNALING\_PATHWAY(GO:2001237) |  | 105 | 0.88 | 1.37 | 0.000 | 1.000 | 0.325 | 5871 | tags=8%, list=10%, signal=8% |
| 322 | COLLAGEN\_TRIMER(GO:0005581) |  | 98 | 0.86 | 1.37 | 0.000 | 1.000 | 0.325 | 7099 | tags=27%, list=12%, signal=30% |
| 323 | ERK1\_AND\_ERK2\_CASCADE(GO:0070371) |  | 293 | 0.86 | 1.37 | 0.000 | 1.000 | 0.325 | 6797 | tags=13%, list=11%, signal=14% |
| 324 | LYMPHOCYTE\_MIGRATION(GO:0072676) |  | 93 | 0.90 | 1.37 | 0.000 | 1.000 | 0.325 | 5823 | tags=15%, list=10%, signal=17% |
| 325 | REGULATION\_OF\_HEART\_RATE\_BY\_CARDIAC\_CONDUCTION(GO:0086091) |  | 36 | 0.87 | 1.37 | 0.000 | 1.000 | 0.325 | 6569 | tags=28%, list=11%, signal=31% |
| 326 | REGULATION\_OF\_PROTEIN\_IMPORT\_INTO\_NUCLEUS(GO:0042306) |  | 53 | 0.87 | 1.37 | 0.000 | 1.000 | 0.325 | 6642 | tags=13%, list=11%, signal=15% |
| 327 | REGULATION\_OF\_INTERLEUKIN\_1\_BETA\_PRODUCTION(GO:0032651) |  | 60 | 0.89 | 1.37 | 0.000 | 1.000 | 0.325 | 6486 | tags=10%, list=11%, signal=11% |
| 328 | TYPE\_B\_PANCREATIC\_CELL\_DEVELOPMENT(GO:0003323) |  | 19 | 0.88 | 1.37 | 0.000 | 1.000 | 0.325 | 6473 | tags=26%, list=11%, signal=30% |
| 329 | STRIATED\_MUSCLE\_CONTRACTION(GO:0006941) |  | 175 | 0.85 | 1.37 | 0.000 | 1.000 | 0.325 | 6832 | tags=20%, list=12%, signal=23% |
| 330 | CARDIAC\_MUSCLE\_CELL\_MEMBRANE\_REPOLARIZATION(GO:0099622) |  | 39 | 0.88 | 1.37 | 0.000 | 1.000 | 0.325 | 6569 | tags=23%, list=11%, signal=26% |
| 331 | POTASSIUM\_ION\_BINDING(GO:0030955) |  | 15 | 0.88 | 1.37 | 0.000 | 1.000 | 0.325 | 5913 | tags=13%, list=10%, signal=15% |
| 332 | REGULATION\_OF\_PROTEIN\_LOCALIZATION\_TO\_NUCLEUS(GO:1900180) |  | 106 | 0.85 | 1.37 | 0.000 | 1.000 | 0.325 | 6662 | tags=9%, list=11%, signal=11% |
| 333 | CELLULAR\_RESPONSE\_TO\_XENOBIOTIC\_STIMULUS(GO:0071466) |  | 167 | 0.87 | 1.37 | 0.000 | 1.000 | 0.325 | 6560 | tags=10%, list=11%, signal=11% |
| 334 | GLANDULAR\_EPITHELIAL\_CELL\_DEVELOPMENT(GO:0002068) |  | 24 | 0.88 | 1.37 | 0.000 | 1.000 | 0.325 | 6473 | tags=29%, list=11%, signal=33% |
| 335 | POSITIVE\_REGULATION\_OF\_CELL\_CYCLE\_G1\_S\_PHASE\_TRANSITION(GO:1902808) |  | 41 | 0.84 | 1.37 | 0.000 | 1.000 | 0.325 | 5852 | tags=7%, list=10%, signal=8% |
| 336 | ANCHORING\_JUNCTION(GO:0070161) |  | 527 | 0.86 | 1.36 | 0.000 | 1.000 | 0.325 | 6626 | tags=9%, list=11%, signal=10% |
| 337 | NEGATIVE\_REGULATION\_OF\_KINASE\_ACTIVITY(GO:0033673) |  | 279 | 0.86 | 1.36 | 0.000 | 1.000 | 0.325 | 6220 | tags=6%, list=11%, signal=7% |
| 338 | REGULATION\_OF\_ERK1\_AND\_ERK2\_CASCADE(GO:0070372) |  | 276 | 0.86 | 1.36 | 0.000 | 1.000 | 0.325 | 6797 | tags=13%, list=11%, signal=15% |
| 339 | INVADOPODIUM(GO:0071437) |  | 16 | 0.90 | 1.36 | 0.000 | 1.000 | 0.325 | 5877 | tags=13%, list=10%, signal=14% |
| 340 | PHOSPHATIDYLCHOLINE\_BIOSYNTHETIC\_PROCESS(GO:0006656) |  | 36 | 0.83 | 1.36 | 0.000 | 1.000 | 0.325 | 6084 | tags=14%, list=10%, signal=15% |
| 341 | ASPARTIC\_TYPE\_PEPTIDASE\_ACTIVITY(GO:0070001) |  | 27 | 0.87 | 1.36 | 0.000 | 1.000 | 0.325 | 5862 | tags=4%, list=10%, signal=4% |
| 342 | REGULATION\_OF\_PROTEIN\_KINASE\_A\_SIGNALING(GO:0010738) |  | 25 | 0.83 | 1.36 | 0.000 | 1.000 | 0.325 | 7441 | tags=28%, list=13%, signal=32% |
| 343 | MYELIN\_ASSEMBLY(GO:0032288) |  | 18 | 0.89 | 1.36 | 0.000 | 1.000 | 0.325 | 6004 | tags=17%, list=10%, signal=19% |
| 344 | CELLULAR\_RESPONSE\_TO\_OXIDATIVE\_STRESS(GO:0034599) |  | 283 | 0.86 | 1.36 | 0.000 | 1.000 | 0.325 | 6626 | tags=9%, list=11%, signal=10% |
| 345 | COFACTOR\_CATABOLIC\_PROCESS(GO:0051187) |  | 53 | 0.87 | 1.36 | 0.000 | 1.000 | 0.325 | 6438 | tags=9%, list=11%, signal=11% |
| 346 | RESPONSE\_TO\_KETONE(GO:1901654) |  | 186 | 0.88 | 1.36 | 0.000 | 1.000 | 0.325 | 6186 | tags=12%, list=10%, signal=13% |
| 347 | POSITIVE\_REGULATION\_OF\_CATION\_CHANNEL\_ACTIVITY(GO:2001259) |  | 58 | 0.85 | 1.36 | 0.000 | 1.000 | 0.325 | 7441 | tags=21%, list=13%, signal=24% |
| 348 | RIBONUCLEOSIDE\_METABOLIC\_PROCESS(GO:0009119) |  | 103 | 0.80 | 1.36 | 0.000 | 1.000 | 0.325 | 7635 | tags=12%, list=13%, signal=13% |
| 349 | REGULATION\_OF\_SPINDLE\_ORGANIZATION(GO:0090224) |  | 39 | 0.83 | 1.36 | 0.000 | 1.000 | 0.325 | 6626 | tags=5%, list=11%, signal=6% |
| 350 | PHOSPHATIDIC\_ACID\_METABOLIC\_PROCESS(GO:0046473) |  | 43 | 0.84 | 1.36 | 0.000 | 1.000 | 0.325 | 6622 | tags=14%, list=11%, signal=16% |
| 351 | CELL\_PROJECTION\_MEMBRANE(GO:0031253) |  | 317 | 0.87 | 1.36 | 0.000 | 1.000 | 0.325 | 6261 | tags=9%, list=11%, signal=10% |
| 352 | HYDROLASE\_ACTIVITY\_\_HYDROLYZING\_O\_GLYCOSYL\_COMPOUNDS(GO:0004553) |  | 106 | 0.82 | 1.36 | 0.000 | 1.000 | 0.325 | 6709 | tags=7%, list=11%, signal=7% |
| 353 | GLAND\_MORPHOGENESIS(GO:0022612) |  | 119 | 0.89 | 1.36 | 0.000 | 1.000 | 0.325 | 6182 | tags=14%, list=10%, signal=16% |
| 354 | MYOBLAST\_PROLIFERATION(GO:0051450) |  | 27 | 0.88 | 1.36 | 0.000 | 1.000 | 0.325 | 6209 | tags=22%, list=10%, signal=25% |
| 355 | ANGIOGENESIS\_INVOLVED\_IN\_WOUND\_HEALING(GO:0060055) |  | 23 | 0.88 | 1.36 | 0.000 | 1.000 | 0.325 | 6405 | tags=22%, list=11%, signal=24% |
| 356 | REGULATION\_OF\_CELLULAR\_COMPONENT\_SIZE(GO:0032535) |  | 329 | 0.87 | 1.36 | 0.141 | 1.000 | 0.325 | 6161 | tags=7%, list=10%, signal=8% |
| 357 | RESPONSE\_TO\_ORGANONITROGEN\_COMPOUND(GO:0010243) |  | 934 | 0.86 | 1.36 | 0.000 | 1.000 | 0.325 | 6578 | tags=11%, list=11%, signal=12% |
| 358 | CELLULAR\_RESPONSE\_TO\_ORGANIC\_CYCLIC\_COMPOUND(GO:0071407) |  | 581 | 0.87 | 1.36 | 0.000 | 1.000 | 0.325 | 6560 | tags=11%, list=11%, signal=12% |
| 359 | REGULATION\_OF\_MEMBRANE\_REPOLARIZATION(GO:0060306) |  | 37 | 0.87 | 1.36 | 0.000 | 1.000 | 0.325 | 6412 | tags=22%, list=11%, signal=24% |
| 360 | REGULATION\_OF\_FATTY\_ACID\_METABOLIC\_PROCESS(GO:0019217) |  | 85 | 0.82 | 1.36 | 0.000 | 1.000 | 0.325 | 7699 | tags=14%, list=13%, signal=16% |
| 361 | MUSCLE\_FIBER\_DEVELOPMENT(GO:0048747) |  | 56 | 0.87 | 1.36 | 0.000 | 1.000 | 0.325 | 6670 | tags=25%, list=11%, signal=28% |
| 362 | REGULATION\_OF\_RELEASE\_OF\_CYTOCHROME\_C\_FROM\_MITOCHONDRIA(GO:0090199) |  | 45 | 0.87 | 1.36 | 0.000 | 1.000 | 0.325 | 5834 | tags=4%, list=10%, signal=5% |
| 363 | REGULATION\_OF\_JUN\_KINASE\_ACTIVITY(GO:0043506) |  | 84 | 0.88 | 1.36 | 0.000 | 1.000 | 0.325 | 6121 | tags=10%, list=10%, signal=11% |
| 364 | CELLULAR\_RESPONSE\_TO\_PROSTAGLANDIN\_STIMULUS(GO:0071379) |  | 23 | 0.88 | 1.36 | 0.000 | 1.000 | 0.325 | 6178 | tags=26%, list=10%, signal=29% |
| 365 | T\_CELL\_MIGRATION(GO:0072678) |  | 50 | 0.90 | 1.36 | 0.000 | 1.000 | 0.325 | 5823 | tags=16%, list=10%, signal=18% |
| 366 | NEGATIVE\_REGULATION\_OF\_TRANSFERASE\_ACTIVITY(GO:0051348) |  | 305 | 0.86 | 1.36 | 0.000 | 1.000 | 0.325 | 6220 | tags=6%, list=11%, signal=7% |
| 367 | CELL\_SURFACE(GO:0009986) |  | 755 | 0.85 | 1.36 | 0.000 | 1.000 | 0.325 | 6821 | tags=14%, list=12%, signal=16% |
| 368 | NEGATIVE\_REGULATION\_OF\_SMOOTH\_MUSCLE\_CELL\_PROLIFERATION(GO:0048662) |  | 53 | 0.89 | 1.36 | 0.000 | 1.000 | 0.325 | 5825 | tags=4%, list=10%, signal=4% |
| 369 | PROTEIN\_LOCALIZATION\_TO\_CELL\_SURFACE(GO:0034394) |  | 53 | 0.85 | 1.36 | 0.000 | 1.000 | 0.325 | 7136 | tags=23%, list=12%, signal=26% |
| 370 | RESPONSE\_TO\_STEROL(GO:0036314) |  | 26 | 0.89 | 1.36 | 0.000 | 1.000 | 0.325 | 5833 | tags=8%, list=10%, signal=9% |
| 371 | MEMBRANE\_DEPOLARIZATION\_DURING\_CARDIAC\_MUSCLE\_CELL\_ACTION\_POTENTIAL(GO:0086012) |  | 22 | 0.88 | 1.36 | 0.000 | 1.000 | 0.325 | 6573 | tags=50%, list=11%, signal=56% |
| 372 | POSITIVE\_REGULATION\_OF\_REACTIVE\_OXYGEN\_SPECIES\_METABOLIC\_PROCESS(GO:2000379) |  | 88 | 0.88 | 1.36 | 0.000 | 1.000 | 0.325 | 6453 | tags=13%, list=11%, signal=14% |
| 373 | EPITHELIAL\_CELL\_DEVELOPMENT(GO:0002064) |  | 201 | 0.88 | 1.36 | 0.141 | 1.000 | 0.325 | 6473 | tags=11%, list=11%, signal=12% |
| 374 | ICOSANOID\_METABOLIC\_PROCESS(GO:0006690) |  | 106 | 0.86 | 1.36 | 0.000 | 1.000 | 0.325 | 6424 | tags=11%, list=11%, signal=13% |
| 375 | COLLAGEN\_CATABOLIC\_PROCESS(GO:0030574) |  | 68 | 0.89 | 1.36 | 0.000 | 1.000 | 0.325 | 6064 | tags=21%, list=10%, signal=23% |
| 376 | POSITIVE\_REGULATION\_OF\_ORGAN\_GROWTH(GO:0046622) |  | 60 | 0.86 | 1.36 | 0.000 | 1.000 | 0.325 | 5925 | tags=18%, list=10%, signal=20% |
| 377 | STRUCTURAL\_CONSTITUENT\_OF\_CYTOSKELETON(GO:0005200) |  | 109 | 0.89 | 1.36 | 0.141 | 1.000 | 0.325 | 5949 | tags=8%, list=10%, signal=9% |
| 378 | MIRNA\_METABOLIC\_PROCESS(GO:0010586) |  | 23 | 0.86 | 1.36 | 0.000 | 1.000 | 0.325 | 5858 | tags=9%, list=10%, signal=10% |
| 379 | CELLULAR\_RESPONSE\_TO\_PROSTAGLANDIN\_E\_STIMULUS(GO:0071380) |  | 18 | 0.88 | 1.36 | 0.000 | 1.000 | 0.325 | 6178 | tags=22%, list=10%, signal=25% |
| 380 | LYMPHOCYTE\_ACTIVATION(GO:0046649) |  | 680 | 0.85 | 1.36 | 0.000 | 1.000 | 0.325 | 6774 | tags=11%, list=11%, signal=12% |
| 381 | REGULATION\_OF\_PROTEIN\_IMPORT(GO:1904589) |  | 56 | 0.86 | 1.36 | 0.000 | 1.000 | 0.325 | 6642 | tags=13%, list=11%, signal=14% |
| 382 | NEURON\_PROJECTION\_MEMBRANE(GO:0032589) |  | 44 | 0.86 | 1.36 | 0.000 | 1.000 | 0.325 | 6818 | tags=25%, list=12%, signal=28% |
| 383 | APICAL\_PART\_OF\_CELL(GO:0045177) |  | 363 | 0.87 | 1.36 | 0.141 | 1.000 | 0.325 | 6784 | tags=16%, list=11%, signal=18% |
| 384 | CELL\_ADHESION\_MOLECULE\_BINDING(GO:0050839) |  | 468 | 0.87 | 1.36 | 0.141 | 1.000 | 0.325 | 6203 | tags=8%, list=10%, signal=9% |
| 385 | NEGATIVE\_REGULATION\_OF\_OXIDATIVE\_STRESS\_INDUCED\_CELL\_DEATH(GO:1903202) |  | 54 | 0.88 | 1.36 | 0.000 | 1.000 | 0.325 | 5834 | tags=6%, list=10%, signal=6% |
| 386 | NEGATIVE\_REGULATION\_OF\_CELLULAR\_RESPONSE\_TO\_OXIDATIVE\_STRESS(GO:1900408) |  | 54 | 0.88 | 1.36 | 0.000 | 1.000 | 0.325 | 5834 | tags=6%, list=10%, signal=6% |
| 387 | MAIN\_AXON(GO:0044304) |  | 66 | 0.85 | 1.36 | 0.000 | 1.000 | 0.325 | 7408 | tags=30%, list=13%, signal=35% |
| 388 | RESPONSE\_TO\_INTERLEUKIN\_6(GO:0070741) |  | 38 | 0.86 | 1.36 | 0.000 | 1.000 | 0.325 | 6211 | tags=16%, list=11%, signal=18% |
| 389 | NEGATIVE\_REGULATION\_OF\_CYTOKINE\_PRODUCTION(GO:0001818) |  | 245 | 0.86 | 1.36 | 0.000 | 1.000 | 0.325 | 6738 | tags=10%, list=11%, signal=11% |
| 390 | TRANSMEMBRANE\_RECEPTOR\_PROTEIN\_TYROSINE\_KINASE\_SIGNALING\_PATHWAY(GO:0007169) |  | 710 | 0.85 | 1.36 | 0.000 | 1.000 | 0.325 | 7103 | tags=12%, list=12%, signal=14% |
| 391 | MUSCLE\_CELL\_PROLIFERATION(GO:0033002) |  | 212 | 0.86 | 1.36 | 0.000 | 1.000 | 0.325 | 6964 | tags=15%, list=12%, signal=17% |
| 392 | ACTIN\_BASED\_CELL\_PROJECTION(GO:0098858) |  | 193 | 0.88 | 1.36 | 0.141 | 1.000 | 0.325 | 6733 | tags=18%, list=11%, signal=20% |
| 393 | SMOOTH\_MUSCLE\_CELL\_PROLIFERATION(GO:0048659) |  | 146 | 0.87 | 1.36 | 0.000 | 1.000 | 0.325 | 5950 | tags=10%, list=10%, signal=11% |
| 394 | GLIAL\_CELL\_DEVELOPMENT(GO:0021782) |  | 94 | 0.88 | 1.36 | 0.000 | 1.000 | 0.325 | 6421 | tags=12%, list=11%, signal=13% |
| 395 | CELLULAR\_RESPONSE\_TO\_GLUCOSE\_STARVATION(GO:0042149) |  | 37 | 0.83 | 1.36 | 0.000 | 1.000 | 0.325 | 7101 | tags=8%, list=12%, signal=9% |
| 396 | METENCEPHALON\_DEVELOPMENT(GO:0022037) |  | 108 | 0.84 | 1.36 | 0.000 | 1.000 | 0.325 | 7546 | tags=22%, list=13%, signal=25% |
| 397 | CELLULAR\_RESPONSE\_TO\_STEROID\_HORMONE\_STIMULUS(GO:0071383) |  | 261 | 0.86 | 1.36 | 0.000 | 1.000 | 0.325 | 6381 | tags=10%, list=11%, signal=11% |
| 398 | NEGATIVE\_REGULATION\_OF\_APOPTOTIC\_SIGNALING\_PATHWAY(GO:2001234) |  | 217 | 0.87 | 1.36 | 0.000 | 1.000 | 0.325 | 6662 | tags=9%, list=11%, signal=10% |
| 399 | FATTY\_ACID\_METABOLIC\_PROCESS(GO:0006631) |  | 360 | 0.86 | 1.36 | 0.000 | 1.000 | 0.325 | 6424 | tags=8%, list=11%, signal=8% |
| 400 | POSITIVE\_REGULATION\_OF\_ION\_TRANSMEMBRANE\_TRANSPORT(GO:0034767) |  | 134 | 0.85 | 1.36 | 0.000 | 1.000 | 0.325 | 7441 | tags=22%, list=13%, signal=25% |
| 401 | INTRINSIC\_COMPONENT\_OF\_SYNAPTIC\_VESICLE\_MEMBRANE(GO:0098563) |  | 22 | 0.85 | 1.36 | 0.000 | 1.000 | 0.325 | 6046 | tags=14%, list=10%, signal=15% |
| 402 | NUCLEOSIDE\_CATABOLIC\_PROCESS(GO:0009164) |  | 31 | 0.81 | 1.36 | 0.000 | 1.000 | 0.325 | 6865 | tags=19%, list=12%, signal=22% |
| 403 | ADHERENS\_JUNCTION(GO:0005912) |  | 510 | 0.86 | 1.36 | 0.000 | 1.000 | 0.325 | 6670 | tags=9%, list=11%, signal=10% |
| 404 | FILOPODIUM\_MEMBRANE(GO:0031527) |  | 17 | 0.88 | 1.36 | 0.000 | 1.000 | 0.325 | 5877 | tags=18%, list=10%, signal=20% |
| 405 | RESPONSE\_TO\_CHOLESTEROL(GO:0070723) |  | 22 | 0.89 | 1.36 | 0.000 | 1.000 | 0.325 | 5833 | tags=9%, list=10%, signal=10% |
| 406 | RETROGRADE\_TRANSPORT\_\_ENDOSOME\_TO\_PLASMA\_MEMBRANE(GO:1990126) |  | 20 | 0.83 | 1.36 | 0.000 | 1.000 | 0.325 | 5926 | tags=10%, list=10%, signal=11% |
| 407 | REGULATION\_OF\_CARDIAC\_MUSCLE\_CONTRACTION\_BY\_CALCIUM\_ION\_SIGNALING(GO:0010882) |  | 27 | 0.86 | 1.36 | 0.000 | 1.000 | 0.325 | 7008 | tags=22%, list=12%, signal=25% |
| 408 | EPIDERMIS\_MORPHOGENESIS(GO:0048730) |  | 29 | 0.89 | 1.36 | 0.000 | 1.000 | 0.325 | 5957 | tags=21%, list=10%, signal=23% |
| 409 | NEGATIVE\_REGULATION\_OF\_EXTRINSIC\_APOPTOTIC\_SIGNALING\_PATHWAY\_VIA\_DEATH\_DOMAIN\_RECEPTORS(GO:1902042) |  | 38 | 0.88 | 1.36 | 0.000 | 1.000 | 0.325 | 6212 | tags=16%, list=11%, signal=18% |
| 410 | RESPONSE\_TO\_XENOBIOTIC\_STIMULUS(GO:0009410) |  | 268 | 0.86 | 1.36 | 0.000 | 1.000 | 0.325 | 6560 | tags=12%, list=11%, signal=13% |
| 411 | ACTIN\_FILAMENT\_BASED\_MOVEMENT(GO:0030048) |  | 132 | 0.86 | 1.36 | 0.141 | 1.000 | 0.325 | 6583 | tags=23%, list=11%, signal=26% |
| 412 | REGULATION\_OF\_FIBROBLAST\_PROLIFERATION(GO:0048145) |  | 88 | 0.88 | 1.36 | 0.141 | 1.000 | 0.325 | 6617 | tags=9%, list=11%, signal=10% |
| 413 | PYRIMIDINE\_NUCLEOSIDE\_CATABOLIC\_PROCESS(GO:0046135) |  | 22 | 0.83 | 1.36 | 0.000 | 1.000 | 0.325 | 6865 | tags=27%, list=12%, signal=31% |
| 414 | PHOSPHATIDIC\_ACID\_BIOSYNTHETIC\_PROCESS(GO:0006654) |  | 42 | 0.84 | 1.36 | 0.000 | 1.000 | 0.325 | 6622 | tags=14%, list=11%, signal=16% |
| 415 | ASPARTIC\_TYPE\_ENDOPEPTIDASE\_ACTIVITY(GO:0004190) |  | 26 | 0.87 | 1.36 | 0.000 | 1.000 | 0.325 | 5862 | tags=4%, list=10%, signal=4% |
| 416 | GAP\_JUNCTION(GO:0005921) |  | 33 | 0.88 | 1.36 | 0.000 | 1.000 | 0.325 | 5959 | tags=21%, list=10%, signal=24% |
| 417 | REGULATION\_OF\_OXIDATIVE\_STRESS\_INDUCED\_CELL\_DEATH(GO:1903201) |  | 75 | 0.86 | 1.36 | 0.000 | 1.000 | 0.325 | 6514 | tags=7%, list=11%, signal=7% |
| 418 | POSITIVE\_REGULATION\_OF\_CATION\_TRANSMEMBRANE\_TRANSPORT(GO:1904064) |  | 124 | 0.85 | 1.36 | 0.000 | 0.998 | 0.325 | 7441 | tags=21%, list=13%, signal=24% |
| 419 | LEUKOCYTE\_DEGRANULATION(GO:0043299) |  | 527 | 0.83 | 1.36 | 0.000 | 0.998 | 0.325 | 6696 | tags=6%, list=11%, signal=6% |
| 420 | NEGATIVE\_REGULATION\_OF\_RESPONSE\_TO\_OXIDATIVE\_STRESS(GO:1902883) |  | 56 | 0.87 | 1.36 | 0.000 | 0.998 | 0.325 | 5834 | tags=5%, list=10%, signal=6% |
| 421 | NEGATIVE\_REGULATION\_OF\_CYSTEINE\_TYPE\_ENDOPEPTIDASE\_ACTIVITY(GO:2000117) |  | 111 | 0.87 | 1.36 | 0.141 | 0.996 | 0.325 | 6424 | tags=12%, list=11%, signal=13% |
| 422 | POSITIVE\_REGULATION\_OF\_SYNAPSE\_ASSEMBLY(GO:0051965) |  | 65 | 0.84 | 1.36 | 0.000 | 0.995 | 0.325 | 7405 | tags=29%, list=13%, signal=33% |
| 423 | REGULATION\_OF\_SMOOTH\_MUSCLE\_CELL\_PROLIFERATION(GO:0048660) |  | 143 | 0.87 | 1.36 | 0.000 | 0.993 | 0.325 | 5950 | tags=10%, list=10%, signal=11% |
| 424 | VASCULAR\_ENDOTHELIAL\_CELL\_PROLIFERATION(GO:0101023) |  | 40 | 0.87 | 1.36 | 0.000 | 0.992 | 0.325 | 7134 | tags=13%, list=12%, signal=14% |
| 425 | REGULATION\_OF\_VASCULAR\_ENDOTHELIAL\_CELL\_PROLIFERATION(GO:1905562) |  | 40 | 0.87 | 1.36 | 0.000 | 0.990 | 0.325 | 7134 | tags=13%, list=12%, signal=14% |
| 426 | MYELOID\_LEUKOCYTE\_MEDIATED\_IMMUNITY(GO:0002444) |  | 539 | 0.83 | 1.36 | 0.000 | 0.990 | 0.325 | 6696 | tags=6%, list=11%, signal=7% |
| 427 | CYSTEINE\_TYPE\_ENDOPEPTIDASE\_INHIBITOR\_ACTIVITY\_INVOLVED\_IN\_APOPTOTIC\_PROCESS(GO:0043027) |  | 25 | 0.85 | 1.36 | 0.000 | 0.988 | 0.325 | 6838 | tags=16%, list=12%, signal=18% |
| 428 | CELLULAR\_RESPONSE\_TO\_INTERLEUKIN\_6(GO:0071354) |  | 34 | 0.86 | 1.36 | 0.000 | 0.987 | 0.325 | 6048 | tags=12%, list=10%, signal=13% |
| 429 | POSTSYNAPTIC\_MEMBRANE(GO:0045211) |  | 229 | 0.83 | 1.36 | 0.000 | 0.986 | 0.325 | 7323 | tags=24%, list=12%, signal=28% |
| 430 | FIBROBLAST\_PROLIFERATION(GO:0048144) |  | 89 | 0.88 | 1.36 | 0.141 | 0.984 | 0.325 | 6617 | tags=9%, list=11%, signal=10% |
| 431 | VIRAL\_RNA\_GENOME\_REPLICATION(GO:0039694) |  | 24 | 0.83 | 1.36 | 0.000 | 0.983 | 0.325 | 6865 | tags=17%, list=12%, signal=19% |
| 432 | CYTOKINE\_MEDIATED\_SIGNALING\_PATHWAY(GO:0019221) |  | 789 | 0.86 | 1.36 | 0.141 | 0.981 | 0.325 | 6473 | tags=11%, list=11%, signal=12% |
| 433 | CENTRAL\_NERVOUS\_SYSTEM\_NEURON\_DIFFERENTIATION(GO:0021953) |  | 180 | 0.86 | 1.36 | 0.000 | 0.979 | 0.325 | 6777 | tags=23%, list=11%, signal=26% |
| 434 | REGULATION\_OF\_LIPOPOLYSACCHARIDE\_MEDIATED\_SIGNALING\_PATHWAY(GO:0031664) |  | 25 | 0.87 | 1.36 | 0.000 | 0.977 | 0.325 | 5858 | tags=8%, list=10%, signal=9% |
| 435 | POSITIVE\_REGULATION\_OF\_STEM\_CELL\_PROLIFERATION(GO:2000648) |  | 35 | 0.89 | 1.36 | 0.141 | 0.976 | 0.325 | 5892 | tags=20%, list=10%, signal=22% |
| 436 | NEGATIVE\_REGULATION\_OF\_CYSTEINE\_TYPE\_ENDOPEPTIDASE\_ACTIVITY\_INVOLVED\_IN\_APOPTOTIC\_PROCESS(GO:0043154) |  | 92 | 0.87 | 1.36 | 0.141 | 0.974 | 0.325 | 6424 | tags=13%, list=11%, signal=15% |
| 437 | MAINTENANCE\_OF\_PROTEIN\_LOCATION(GO:0045185) |  | 98 | 0.83 | 1.36 | 0.000 | 0.973 | 0.325 | 6706 | tags=8%, list=11%, signal=9% |
| 438 | RESPONSE\_TO\_CORTICOSTEROID(GO:0031960) |  | 158 | 0.87 | 1.36 | 0.000 | 0.972 | 0.325 | 6424 | tags=11%, list=11%, signal=12% |
| 439 | RESPONSE\_TO\_NITROGEN\_COMPOUND(GO:1901698) |  | 1060 | 0.86 | 1.36 | 0.000 | 0.971 | 0.325 | 6578 | tags=10%, list=11%, signal=12% |
| 440 | POSTSYNAPTIC\_DENSITY(GO:0014069) |  | 206 | 0.83 | 1.36 | 0.000 | 0.969 | 0.325 | 7995 | tags=21%, list=14%, signal=24% |
| 441 | POSTSYNAPTIC\_SPECIALIZATION(GO:0099572) |  | 207 | 0.83 | 1.36 | 0.000 | 0.967 | 0.325 | 7995 | tags=21%, list=14%, signal=24% |
| 442 | REGULATION\_OF\_CELLULAR\_CATABOLIC\_PROCESS(GO:0031329) |  | 807 | 0.84 | 1.36 | 0.000 | 0.966 | 0.325 | 6397 | tags=5%, list=11%, signal=5% |
| 443 | A\_BAND(GO:0031672) |  | 45 | 0.83 | 1.36 | 0.000 | 0.964 | 0.325 | 6605 | tags=31%, list=11%, signal=35% |
| 444 | ASYMMETRIC\_SYNAPSE(GO:0032279) |  | 211 | 0.83 | 1.36 | 0.000 | 0.962 | 0.325 | 7995 | tags=21%, list=14%, signal=25% |
| 445 | MYELOID\_CELL\_ACTIVATION\_INVOLVED\_IN\_IMMUNE\_RESPONSE(GO:0002275) |  | 534 | 0.83 | 1.36 | 0.000 | 0.961 | 0.325 | 6696 | tags=6%, list=11%, signal=7% |
| 446 | HISTONE\_PHOSPHORYLATION(GO:0016572) |  | 37 | 0.86 | 1.36 | 0.000 | 0.959 | 0.325 | 5904 | tags=14%, list=10%, signal=15% |
| 447 | EPITHELIAL\_CELL\_FATE\_COMMITMENT(GO:0072148) |  | 16 | 0.89 | 1.36 | 0.000 | 0.958 | 0.325 | 6058 | tags=31%, list=10%, signal=35% |
| 448 | RESPONSE\_TO\_TUMOR\_NECROSIS\_FACTOR(GO:0034612) |  | 307 | 0.86 | 1.36 | 0.141 | 0.956 | 0.325 | 6454 | tags=8%, list=11%, signal=9% |
| 449 | ANTIBIOTIC\_CATABOLIC\_PROCESS(GO:0017001) |  | 43 | 0.89 | 1.36 | 0.000 | 0.955 | 0.325 | 5920 | tags=14%, list=10%, signal=15% |
| 450 | NEGATIVE\_REGULATION\_OF\_TOR\_SIGNALING(GO:0032007) |  | 44 | 0.84 | 1.36 | 0.000 | 0.954 | 0.325 | 6256 | tags=11%, list=11%, signal=13% |
| 451 | NOREPINEPHRINE\_TRANSPORT(GO:0015874) |  | 22 | 0.88 | 1.36 | 0.000 | 0.954 | 0.325 | 5955 | tags=18%, list=10%, signal=20% |
| 452 | C21\_STEROID\_HORMONE\_BIOSYNTHETIC\_PROCESS(GO:0006700) |  | 25 | 0.87 | 1.36 | 0.000 | 0.953 | 0.325 | 5839 | tags=8%, list=10%, signal=9% |
| 453 | TRANSFERASE\_ACTIVITY\_\_TRANSFERRING\_NITROGENOUS\_GROUPS(GO:0016769) |  | 25 | 0.85 | 1.36 | 0.000 | 0.951 | 0.325 | 6145 | tags=12%, list=10%, signal=13% |
| 454 | SPECIFICATION\_OF\_SYMMETRY(GO:0009799) |  | 127 | 0.83 | 1.36 | 0.000 | 0.950 | 0.325 | 7344 | tags=26%, list=12%, signal=30% |
| 455 | AMINE\_BIOSYNTHETIC\_PROCESS(GO:0009309) |  | 23 | 0.85 | 1.36 | 0.000 | 0.950 | 0.325 | 7268 | tags=13%, list=12%, signal=15% |
| 456 | RESPONSE\_TO\_GLUCOCORTICOID(GO:0051384) |  | 142 | 0.87 | 1.36 | 0.000 | 0.949 | 0.325 | 6424 | tags=11%, list=11%, signal=13% |
| 457 | CELL\_RECOGNITION(GO:0008037) |  | 191 | 0.88 | 1.36 | 0.141 | 0.948 | 0.325 | 6000 | tags=10%, list=10%, signal=12% |
| 458 | REGULATION\_OF\_MAP\_KINASE\_ACTIVITY(GO:0043405) |  | 337 | 0.85 | 1.36 | 0.000 | 0.946 | 0.325 | 6928 | tags=14%, list=12%, signal=15% |
| 459 | SKELETAL\_MUSCLE\_FIBER\_DEVELOPMENT(GO:0048741) |  | 34 | 0.88 | 1.36 | 0.000 | 0.945 | 0.325 | 5834 | tags=18%, list=10%, signal=20% |
| 460 | REGULATION\_OF\_CELLULAR\_RESPONSE\_TO\_OXIDATIVE\_STRESS(GO:1900407) |  | 84 | 0.85 | 1.36 | 0.000 | 0.945 | 0.325 | 6514 | tags=7%, list=11%, signal=8% |
| 461 | RESPONSE\_TO\_ETHANOL(GO:0045471) |  | 138 | 0.87 | 1.36 | 0.141 | 0.944 | 0.325 | 5874 | tags=9%, list=10%, signal=10% |
| 462 | CENTRAL\_NERVOUS\_SYSTEM\_NEURON\_DEVELOPMENT(GO:0021954) |  | 73 | 0.86 | 1.36 | 0.000 | 0.942 | 0.325 | 7081 | tags=16%, list=12%, signal=19% |
| 463 | POSITIVE\_REGULATION\_OF\_PURINE\_NUCLEOTIDE\_BIOSYNTHETIC\_PROCESS(GO:1900373) |  | 67 | 0.83 | 1.36 | 0.000 | 0.941 | 0.325 | 7265 | tags=18%, list=12%, signal=20% |
| 464 | POSITIVE\_REGULATION\_OF\_NUCLEOTIDE\_BIOSYNTHETIC\_PROCESS(GO:0030810) |  | 67 | 0.83 | 1.36 | 0.000 | 0.939 | 0.325 | 7265 | tags=18%, list=12%, signal=20% |
| 465 | MODULATION\_OF\_TRANSCRIPTION\_IN\_OTHER\_ORGANISM\_INVOLVED\_IN\_SYMBIOTIC\_INTERACTION(GO:0052312) |  | 28 | 0.83 | 1.36 | 0.000 | 0.938 | 0.325 | 6381 | tags=7%, list=11%, signal=8% |
| 466 | HETEROTRIMERIC\_G\_PROTEIN\_COMPLEX(GO:0005834) |  | 38 | 0.86 | 1.36 | 0.000 | 0.937 | 0.325 | 6471 | tags=16%, list=11%, signal=18% |
| 467 | GTPASE\_COMPLEX(GO:1905360) |  | 38 | 0.86 | 1.36 | 0.000 | 0.935 | 0.325 | 6471 | tags=16%, list=11%, signal=18% |
| 468 | NEURON\_TO\_NEURON\_SYNAPSE(GO:0098984) |  | 214 | 0.83 | 1.36 | 0.000 | 0.934 | 0.325 | 7995 | tags=21%, list=14%, signal=24% |
| 469 | ARTERY\_MORPHOGENESIS(GO:0048844) |  | 73 | 0.84 | 1.36 | 0.000 | 0.933 | 0.325 | 6218 | tags=16%, list=11%, signal=18% |
| 470 | REGULATION\_OF\_OXIDATIVE\_PHOSPHORYLATION(GO:0002082) |  | 17 | 0.85 | 1.36 | 0.000 | 0.931 | 0.325 | 5894 | tags=6%, list=10%, signal=7% |
| 471 | SPLEEN\_DEVELOPMENT(GO:0048536) |  | 35 | 0.86 | 1.35 | 0.000 | 0.931 | 0.325 | 6216 | tags=11%, list=11%, signal=13% |
| 472 | RESPONSE\_TO\_NUTRIENT(GO:0007584) |  | 212 | 0.86 | 1.35 | 0.141 | 0.929 | 0.325 | 6706 | tags=15%, list=11%, signal=16% |
| 473 | GLAND\_DEVELOPMENT(GO:0048732) |  | 435 | 0.87 | 1.35 | 0.141 | 0.929 | 0.325 | 6617 | tags=17%, list=11%, signal=19% |
| 474 | NEGATIVE\_REGULATION\_OF\_HISTONE\_MODIFICATION(GO:0031057) |  | 39 | 0.84 | 1.35 | 0.000 | 0.928 | 0.325 | 5904 | tags=5%, list=10%, signal=6% |
| 475 | CELLULAR\_RESPONSE\_TO\_TUMOR\_NECROSIS\_FACTOR(GO:0071356) |  | 289 | 0.87 | 1.35 | 0.141 | 0.926 | 0.325 | 6454 | tags=9%, list=11%, signal=10% |
| 476 | CELL\_CHEMOTAXIS(GO:0060326) |  | 267 | 0.87 | 1.35 | 0.141 | 0.926 | 0.325 | 6738 | tags=18%, list=11%, signal=20% |
| 477 | MEMBRANE\_DEPOLARIZATION(GO:0051899) |  | 100 | 0.86 | 1.35 | 0.000 | 0.927 | 0.325 | 6573 | tags=26%, list=11%, signal=29% |
| 478 | REGULATION\_OF\_ANATOMICAL\_STRUCTURE\_SIZE(GO:0090066) |  | 466 | 0.86 | 1.35 | 0.141 | 0.925 | 0.325 | 6525 | tags=11%, list=11%, signal=12% |
| 479 | DETERMINATION\_OF\_BILATERAL\_SYMMETRY(GO:0009855) |  | 126 | 0.83 | 1.35 | 0.000 | 0.923 | 0.325 | 7344 | tags=26%, list=12%, signal=30% |
| 480 | NEGATIVE\_REGULATION\_OF\_CHEMOKINE\_PRODUCTION(GO:0032682) |  | 19 | 0.90 | 1.35 | 0.000 | 0.922 | 0.325 | 5825 | tags=5%, list=10%, signal=6% |
| 481 | PYRIMIDINE\_RIBONUCLEOSIDE\_METABOLIC\_PROCESS(GO:0046131) |  | 34 | 0.80 | 1.35 | 0.000 | 0.921 | 0.325 | 6865 | tags=15%, list=12%, signal=17% |
| 482 | NEGATIVE\_REGULATION\_OF\_RESPONSE\_TO\_EXTERNAL\_STIMULUS(GO:0032102) |  | 287 | 0.86 | 1.35 | 0.000 | 0.919 | 0.325 | 7159 | tags=18%, list=12%, signal=21% |
| 483 | CYCLIN\_DEPENDENT\_PROTEIN\_KINASE\_HOLOENZYME\_COMPLEX(GO:0000307) |  | 40 | 0.83 | 1.35 | 0.000 | 0.918 | 0.325 | 6745 | tags=8%, list=11%, signal=8% |
| 484 | RESPONSE\_TO\_SALT\_STRESS(GO:0009651) |  | 24 | 0.88 | 1.35 | 0.141 | 0.918 | 0.325 | 6374 | tags=17%, list=11%, signal=19% |
| 485 | DNA\_BINDING\_\_BENDING(GO:0008301) |  | 18 | 0.82 | 1.35 | 0.000 | 0.917 | 0.325 | 7260 | tags=22%, list=12%, signal=25% |
| 486 | PROTEIN\_KINASE\_BINDING(GO:0019901) |  | 586 | 0.86 | 1.35 | 0.141 | 0.916 | 0.325 | 6218 | tags=6%, list=11%, signal=6% |
| 487 | TUMOR\_NECROSIS\_FACTOR\_MEDIATED\_SIGNALING\_PATHWAY(GO:0033209) |  | 174 | 0.87 | 1.35 | 0.141 | 0.916 | 0.325 | 6454 | tags=6%, list=11%, signal=7% |
| 488 | HYDROLASE\_ACTIVITY\_\_ACTING\_ON\_GLYCOSYL\_BONDS(GO:0016798) |  | 132 | 0.81 | 1.35 | 0.000 | 0.914 | 0.325 | 6709 | tags=8%, list=11%, signal=9% |
| 489 | BRAIN\_MORPHOGENESIS(GO:0048854) |  | 35 | 0.87 | 1.35 | 0.000 | 0.913 | 0.325 | 6473 | tags=23%, list=11%, signal=26% |
| 490 | SULFUR\_COMPOUND\_METABOLIC\_PROCESS(GO:0006790) |  | 360 | 0.86 | 1.35 | 0.000 | 0.911 | 0.325 | 6683 | tags=11%, list=11%, signal=13% |
| 491 | DETERMINATION\_OF\_LEFT\_RIGHT\_SYMMETRY(GO:0007368) |  | 117 | 0.83 | 1.35 | 0.000 | 0.911 | 0.325 | 7344 | tags=26%, list=12%, signal=29% |
| 492 | REGULATION\_OF\_CALCIUM\_ION\_TRANSMEMBRANE\_TRANSPORT(GO:1903169) |  | 146 | 0.84 | 1.35 | 0.000 | 0.910 | 0.325 | 6662 | tags=14%, list=11%, signal=15% |
| 493 | NEGATIVE\_REGULATION\_OF\_CELLULAR\_SENESCENCE(GO:2000773) |  | 19 | 0.88 | 1.35 | 0.000 | 0.911 | 0.325 | 5916 | tags=21%, list=10%, signal=23% |
| 494 | NUCLEUS\_LOCALIZATION(GO:0051647) |  | 24 | 0.85 | 1.35 | 0.000 | 0.910 | 0.325 | 5856 | tags=17%, list=10%, signal=18% |
| 495 | MODULATION\_BY\_HOST\_OF\_SYMBIONT\_TRANSCRIPTION(GO:0052472) |  | 27 | 0.83 | 1.35 | 0.000 | 0.909 | 0.325 | 6381 | tags=7%, list=11%, signal=8% |
| 496 | MODULATION\_BY\_HOST\_OF\_VIRAL\_TRANSCRIPTION(GO:0043921) |  | 27 | 0.83 | 1.35 | 0.000 | 0.907 | 0.325 | 6381 | tags=7%, list=11%, signal=8% |
| 497 | ATRIOVENTRICULAR\_VALVE\_DEVELOPMENT(GO:0003171) |  | 24 | 0.85 | 1.35 | 0.000 | 0.907 | 0.325 | 6282 | tags=21%, list=11%, signal=23% |
| 498 | PEPTIDE\_HORMONE\_BINDING(GO:0017046) |  | 43 | 0.83 | 1.35 | 0.000 | 0.906 | 0.325 | 6186 | tags=23%, list=10%, signal=26% |
| 499 | REGULATION\_OF\_RESPONSE\_TO\_OXIDATIVE\_STRESS(GO:1902882) |  | 94 | 0.85 | 1.35 | 0.000 | 0.904 | 0.325 | 6514 | tags=7%, list=11%, signal=8% |
| 500 | MYOTUBE\_DIFFERENTIATION(GO:0014902) |  | 112 | 0.86 | 1.35 | 0.000 | 0.903 | 0.325 | 7134 | tags=17%, list=12%, signal=19% |
| 501 | INNER\_EAR\_DEVELOPMENT(GO:0048839) |  | 188 | 0.84 | 1.35 | 0.000 | 0.904 | 0.325 | 7396 | tags=24%, list=13%, signal=28% |
| 502 | MYOTUBE\_CELL\_DEVELOPMENT(GO:0014904) |  | 38 | 0.89 | 1.35 | 0.000 | 0.902 | 0.325 | 5856 | tags=18%, list=10%, signal=20% |
| 503 | M\_BAND(GO:0031430) |  | 32 | 0.84 | 1.35 | 0.000 | 0.903 | 0.325 | 8083 | tags=41%, list=14%, signal=47% |
| 504 | POSITIVE\_REGULATION\_OF\_CELLULAR\_COMPONENT\_BIOGENESIS(GO:0044089) |  | 502 | 0.85 | 1.35 | 0.000 | 0.902 | 0.325 | 7102 | tags=11%, list=12%, signal=13% |
| 505 | RESPONSE\_TO\_ENDOGENOUS\_STIMULUS(GO:0009719) |  | 1594 | 0.85 | 1.35 | 0.000 | 0.902 | 0.325 | 6578 | tags=12%, list=11%, signal=13% |
| 506 | EXTRINSIC\_APOPTOTIC\_SIGNALING\_PATHWAY\_VIA\_DEATH\_DOMAIN\_RECEPTORS(GO:0008625) |  | 84 | 0.86 | 1.35 | 0.000 | 0.901 | 0.325 | 6212 | tags=11%, list=11%, signal=12% |
| 507 | BASOLATERAL\_PLASMA\_MEMBRANE(GO:0016323) |  | 210 | 0.87 | 1.35 | 0.141 | 0.900 | 0.325 | 6602 | tags=17%, list=11%, signal=19% |
| 508 | CELL\_PROLIFERATION\_INVOLVED\_IN\_HEART\_MORPHOGENESIS(GO:0061323) |  | 19 | 0.86 | 1.35 | 0.000 | 0.898 | 0.325 | 5958 | tags=16%, list=10%, signal=18% |
| 509 | PURINE\_RIBONUCLEOSIDE\_METABOLIC\_PROCESS(GO:0046128) |  | 75 | 0.81 | 1.35 | 0.000 | 0.899 | 0.325 | 6423 | tags=5%, list=11%, signal=6% |
| 510 | POSITIVE\_REGULATION\_OF\_CELLULAR\_PROTEIN\_LOCALIZATION(GO:1903829) |  | 347 | 0.87 | 1.35 | 0.141 | 0.898 | 0.325 | 6179 | tags=4%, list=10%, signal=4% |
| 511 | POSITIVE\_REGULATION\_OF\_INTRACELLULAR\_TRANSPORT(GO:0032388) |  | 235 | 0.88 | 1.35 | 0.141 | 0.896 | 0.325 | 5938 | tags=5%, list=10%, signal=6% |
| 512 | COPPER\_ION\_BINDING(GO:0005507) |  | 57 | 0.84 | 1.35 | 0.000 | 0.897 | 0.325 | 7225 | tags=18%, list=12%, signal=20% |
| 513 | POSITIVE\_REGULATION\_OF\_PURINE\_NUCLEOTIDE\_METABOLIC\_PROCESS(GO:1900544) |  | 90 | 0.82 | 1.35 | 0.000 | 0.896 | 0.325 | 7265 | tags=19%, list=12%, signal=22% |
| 514 | POSITIVE\_REGULATION\_OF\_NUCLEOTIDE\_METABOLIC\_PROCESS(GO:0045981) |  | 90 | 0.82 | 1.35 | 0.000 | 0.894 | 0.325 | 7265 | tags=19%, list=12%, signal=22% |
| 515 | PROTEIN\_STABILIZATION(GO:0050821) |  | 158 | 0.82 | 1.35 | 0.000 | 0.893 | 0.325 | 6626 | tags=4%, list=11%, signal=4% |
| 516 | SARCOPLASMIC\_RETICULUM\_CALCIUM\_ION\_TRANSPORT(GO:0070296) |  | 39 | 0.85 | 1.35 | 0.000 | 0.891 | 0.325 | 7544 | tags=26%, list=13%, signal=29% |
| 517 | RESPONSE\_TO\_ANGIOTENSIN(GO:1990776) |  | 27 | 0.84 | 1.35 | 0.000 | 0.891 | 0.325 | 6424 | tags=19%, list=11%, signal=21% |
| 518 | RESPONSE\_TO\_THYROID\_HORMONE(GO:0097066) |  | 26 | 0.87 | 1.35 | 0.000 | 0.890 | 0.325 | 5839 | tags=4%, list=10%, signal=4% |
| 519 | MEMBRANE\_REPOLARIZATION\_DURING\_ACTION\_POTENTIAL(GO:0086011) |  | 29 | 0.88 | 1.35 | 0.000 | 0.891 | 0.325 | 5835 | tags=7%, list=10%, signal=8% |
| 520 | RESPONSE\_TO\_VITAMIN(GO:0033273) |  | 91 | 0.88 | 1.35 | 0.141 | 0.890 | 0.325 | 6495 | tags=18%, list=11%, signal=20% |
| 521 | SPECTRIN\_BINDING(GO:0030507) |  | 26 | 0.82 | 1.35 | 0.000 | 0.889 | 0.325 | 7847 | tags=27%, list=13%, signal=31% |
| 522 | REGULATION\_OF\_VESICLE\_MEDIATED\_TRANSPORT(GO:0060627) |  | 476 | 0.86 | 1.35 | 0.141 | 0.887 | 0.325 | 6283 | tags=7%, list=11%, signal=7% |
| 523 | REGULATION\_OF\_BEHAVIOR(GO:0050795) |  | 67 | 0.83 | 1.35 | 0.000 | 0.886 | 0.325 | 7134 | tags=22%, list=12%, signal=25% |
| 524 | ANATOMICAL\_STRUCTURE\_ARRANGEMENT(GO:0048532) |  | 17 | 0.88 | 1.35 | 0.000 | 0.885 | 0.325 | 6279 | tags=29%, list=11%, signal=33% |
| 525 | REGULATION\_OF\_ATPASE\_ACTIVITY(GO:0043462) |  | 73 | 0.82 | 1.35 | 0.000 | 0.884 | 0.325 | 7008 | tags=11%, list=12%, signal=12% |
| 526 | E\_BOX\_BINDING(GO:0070888) |  | 37 | 0.87 | 1.35 | 0.141 | 0.883 | 0.325 | 6537 | tags=30%, list=11%, signal=33% |
| 527 | CILIUM\_OR\_FLAGELLUM\_DEPENDENT\_CELL\_MOTILITY(GO:0001539) |  | 30 | 0.83 | 1.35 | 0.000 | 0.881 | 0.325 | 7628 | tags=37%, list=13%, signal=42% |
| 528 | REGULATION\_OF\_CHEMOTAXIS(GO:0050920) |  | 183 | 0.87 | 1.35 | 0.141 | 0.881 | 0.325 | 6738 | tags=17%, list=11%, signal=20% |
| 529 | CELL\_CORTEX\_REGION(GO:0099738) |  | 25 | 0.86 | 1.35 | 0.000 | 0.880 | 0.325 | 6581 | tags=8%, list=11%, signal=9% |
| 530 | POSITIVE\_REGULATION\_OF\_NEUROBLAST\_PROLIFERATION(GO:0002052) |  | 20 | 0.89 | 1.35 | 0.000 | 0.879 | 0.325 | 5892 | tags=25%, list=10%, signal=28% |
| 531 | REGULATION\_OF\_EXTRINSIC\_APOPTOTIC\_SIGNALING\_PATHWAY(GO:2001236) |  | 157 | 0.87 | 1.35 | 0.141 | 0.878 | 0.325 | 6212 | tags=8%, list=11%, signal=9% |
| 532 | REGULATION\_OF\_CATABOLIC\_PROCESS(GO:0009894) |  | 901 | 0.84 | 1.35 | 0.000 | 0.877 | 0.325 | 6397 | tags=5%, list=11%, signal=6% |
| 533 | RESPONSE\_TO\_ALCOHOL(GO:0097305) |  | 200 | 0.87 | 1.35 | 0.141 | 0.876 | 0.325 | 6202 | tags=10%, list=10%, signal=11% |
| 534 | AORTA\_MORPHOGENESIS(GO:0035909) |  | 35 | 0.85 | 1.35 | 0.000 | 0.875 | 0.325 | 6218 | tags=14%, list=11%, signal=16% |
| 535 | PURINE\_NUCLEOSIDE\_METABOLIC\_PROCESS(GO:0042278) |  | 78 | 0.80 | 1.35 | 0.000 | 0.874 | 0.325 | 6423 | tags=5%, list=11%, signal=6% |
| 536 | HAIR\_FOLLICLE\_MORPHOGENESIS(GO:0031069) |  | 26 | 0.89 | 1.35 | 0.000 | 0.873 | 0.325 | 5957 | tags=23%, list=10%, signal=26% |
| 537 | REGULATION\_OF\_HEART\_RATE(GO:0002027) |  | 99 | 0.86 | 1.35 | 0.000 | 0.873 | 0.325 | 6573 | tags=23%, list=11%, signal=26% |
| 538 | HISTONE\_KINASE\_ACTIVITY(GO:0035173) |  | 17 | 0.85 | 1.35 | 0.000 | 0.872 | 0.325 | 5861 | tags=12%, list=10%, signal=13% |
| 539 | RESPONSE\_TO\_ACID\_CHEMICAL(GO:0001101) |  | 335 | 0.87 | 1.35 | 0.141 | 0.871 | 0.325 | 6527 | tags=14%, list=11%, signal=16% |
| 540 | MORPHOGENESIS\_OF\_AN\_EPITHELIUM(GO:0002009) |  | 537 | 0.86 | 1.35 | 0.141 | 0.870 | 0.325 | 6495 | tags=12%, list=11%, signal=14% |
| 541 | RELEASE\_OF\_SEQUESTERED\_CALCIUM\_ION\_INTO\_CYTOSOL\_BY\_ENDOPLASMIC\_RETICULUM(GO:1903514) |  | 35 | 0.85 | 1.35 | 0.000 | 0.871 | 0.325 | 7544 | tags=26%, list=13%, signal=29% |
| 542 | NEGATIVE\_REGULATION\_OF\_PROTEIN\_ACETYLATION(GO:1901984) |  | 23 | 0.85 | 1.35 | 0.000 | 0.870 | 0.325 | 6648 | tags=22%, list=11%, signal=24% |
| 543 | COFACTOR\_METABOLIC\_PROCESS(GO:0051186) |  | 563 | 0.85 | 1.35 | 0.000 | 0.869 | 0.325 | 6562 | tags=6%, list=11%, signal=7% |
| 544 | REGULATION\_OF\_CATION\_CHANNEL\_ACTIVITY(GO:2001257) |  | 156 | 0.84 | 1.35 | 0.000 | 0.869 | 0.325 | 7979 | tags=25%, list=13%, signal=29% |
| 545 | POSITIVE\_REGULATION\_OF\_CALCIUM\_ION\_TRANSMEMBRANE\_TRANSPORT(GO:1904427) |  | 65 | 0.86 | 1.35 | 0.000 | 0.868 | 0.325 | 6560 | tags=15%, list=11%, signal=17% |
| 546 | RELEASE\_OF\_SEQUESTERED\_CALCIUM\_ION\_INTO\_CYTOSOL\_BY\_SARCOPLASMIC\_RETICULUM(GO:0014808) |  | 34 | 0.85 | 1.35 | 0.000 | 0.867 | 0.325 | 7544 | tags=26%, list=13%, signal=30% |
| 547 | SYNAPTIC\_MEMBRANE(GO:0097060) |  | 301 | 0.82 | 1.35 | 0.000 | 0.866 | 0.325 | 8040 | tags=28%, list=14%, signal=32% |
| 548 | LIPOPOLYSACCHARIDE\_MEDIATED\_SIGNALING\_PATHWAY(GO:0031663) |  | 56 | 0.85 | 1.35 | 0.000 | 0.865 | 0.325 | 5858 | tags=4%, list=10%, signal=4% |
| 549 | CELLULAR\_RESPONSE\_TO\_STEROL(GO:0036315) |  | 17 | 0.89 | 1.35 | 0.000 | 0.864 | 0.325 | 5833 | tags=12%, list=10%, signal=13% |
| 550 | SUPEROXIDE\_ANION\_GENERATION(GO:0042554) |  | 29 | 0.90 | 1.35 | 0.000 | 0.863 | 0.325 | 5825 | tags=7%, list=10%, signal=8% |
| 551 | REGULATION\_OF\_AMINO\_ACID\_TRANSPORT(GO:0051955) |  | 30 | 0.84 | 1.35 | 0.000 | 0.864 | 0.325 | 5912 | tags=17%, list=10%, signal=19% |
| 552 | LYMPH\_VESSEL\_MORPHOGENESIS(GO:0036303) |  | 18 | 0.84 | 1.35 | 0.000 | 0.863 | 0.325 | 7260 | tags=39%, list=12%, signal=44% |
| 553 | RESPONSE\_TO\_ANTIBIOTIC(GO:0046677) |  | 337 | 0.86 | 1.35 | 0.141 | 0.863 | 0.325 | 6507 | tags=10%, list=11%, signal=11% |
| 554 | REGULATION\_OF\_HAIR\_CYCLE(GO:0042634) |  | 29 | 0.88 | 1.35 | 0.000 | 0.863 | 0.325 | 6473 | tags=10%, list=11%, signal=12% |
| 555 | RESPONSE\_TO\_ORGANIC\_CYCLIC\_COMPOUND(GO:0014070) |  | 952 | 0.85 | 1.35 | 0.000 | 0.862 | 0.325 | 6560 | tags=12%, list=11%, signal=13% |
| 556 | STEM\_CELL\_POPULATION\_MAINTENANCE(GO:0019827) |  | 158 | 0.84 | 1.35 | 0.000 | 0.861 | 0.325 | 6279 | tags=12%, list=11%, signal=13% |
| 557 | SULFUR\_COMPOUND\_BIOSYNTHETIC\_PROCESS(GO:0044272) |  | 184 | 0.86 | 1.35 | 0.000 | 0.860 | 0.325 | 6683 | tags=14%, list=11%, signal=15% |
| 558 | RESPONSE\_TO\_BACTERIUM(GO:0009617) |  | 585 | 0.85 | 1.35 | 0.141 | 0.859 | 0.325 | 6838 | tags=10%, list=12%, signal=11% |
| 559 | NEGATIVE\_REGULATION\_OF\_CELL\_MIGRATION(GO:0030336) |  | 293 | 0.86 | 1.35 | 0.141 | 0.858 | 0.325 | 6027 | tags=8%, list=10%, signal=9% |
| 560 | PSEUDOPODIUM(GO:0031143) |  | 17 | 0.90 | 1.35 | 0.141 | 0.857 | 0.325 | 5938 | tags=18%, list=10%, signal=20% |
| 561 | NEGATIVE\_REGULATION\_OF\_PEPTIDYL\_LYSINE\_ACETYLATION(GO:2000757) |  | 20 | 0.86 | 1.35 | 0.000 | 0.856 | 0.325 | 6648 | tags=15%, list=11%, signal=17% |
| 562 | SULFUR\_AMINO\_ACID\_METABOLIC\_PROCESS(GO:0000096) |  | 39 | 0.81 | 1.35 | 0.000 | 0.855 | 0.325 | 6628 | tags=13%, list=11%, signal=14% |
| 563 | REGULATION\_OF\_HEART\_MORPHOGENESIS(GO:2000826) |  | 36 | 0.88 | 1.35 | 0.000 | 0.855 | 0.325 | 5958 | tags=14%, list=10%, signal=15% |
| 564 | CARDIAC\_RIGHT\_VENTRICLE\_MORPHOGENESIS(GO:0003215) |  | 19 | 0.88 | 1.35 | 0.000 | 0.854 | 0.325 | 5958 | tags=32%, list=10%, signal=35% |
| 565 | ATRIOVENTRICULAR\_VALVE\_MORPHOGENESIS(GO:0003181) |  | 21 | 0.85 | 1.35 | 0.000 | 0.854 | 0.325 | 6282 | tags=14%, list=11%, signal=16% |
| 566 | REGULATION\_OF\_INTRACELLULAR\_TRANSPORT(GO:0032386) |  | 392 | 0.86 | 1.35 | 0.141 | 0.853 | 0.325 | 6259 | tags=6%, list=11%, signal=7% |
| 567 | MAINTENANCE\_OF\_CELL\_NUMBER(GO:0098727) |  | 160 | 0.84 | 1.35 | 0.000 | 0.852 | 0.325 | 6279 | tags=12%, list=11%, signal=13% |
| 568 | NEGATIVE\_REGULATION\_OF\_SMOOTH\_MUSCLE\_CELL\_MIGRATION(GO:0014912) |  | 31 | 0.90 | 1.35 | 0.000 | 0.851 | 0.325 | 5825 | tags=6%, list=10%, signal=7% |
| 569 | NEURON\_SPINE(GO:0044309) |  | 140 | 0.83 | 1.35 | 0.000 | 0.851 | 0.325 | 7962 | tags=23%, list=13%, signal=26% |
| 570 | CELLULAR\_RESPONSE\_TO\_CYTOKINE\_STIMULUS(GO:0071345) |  | 1072 | 0.85 | 1.35 | 0.141 | 0.850 | 0.325 | 6738 | tags=11%, list=11%, signal=13% |
| 571 | CALCIUM\_ION\_IMPORT(GO:0070509) |  | 71 | 0.86 | 1.35 | 0.141 | 0.849 | 0.325 | 6662 | tags=14%, list=11%, signal=16% |
| 572 | GLYCOSYL\_COMPOUND\_METABOLIC\_PROCESS(GO:1901657) |  | 151 | 0.83 | 1.35 | 0.000 | 0.848 | 0.325 | 6555 | tags=8%, list=11%, signal=9% |
| 573 | ORGANIC\_HYDROXY\_COMPOUND\_BIOSYNTHETIC\_PROCESS(GO:1901617) |  | 234 | 0.83 | 1.35 | 0.000 | 0.847 | 0.325 | 7393 | tags=17%, list=13%, signal=19% |
| 574 | REGULATION\_OF\_CYTOKINE\_PRODUCTION(GO:0001817) |  | 625 | 0.83 | 1.35 | 0.000 | 0.847 | 0.325 | 7488 | tags=15%, list=13%, signal=17% |
| 575 | EPITHELIAL\_CELL\_DIFFERENTIATION(GO:0030855) |  | 753 | 0.87 | 1.35 | 0.141 | 0.846 | 0.325 | 6473 | tags=11%, list=11%, signal=12% |
| 576 | ENZYME\_LINKED\_RECEPTOR\_PROTEIN\_SIGNALING\_PATHWAY(GO:0007167) |  | 1026 | 0.84 | 1.35 | 0.000 | 0.845 | 0.325 | 7263 | tags=14%, list=12%, signal=16% |
| 577 | METANEPHRIC\_NEPHRON\_MORPHOGENESIS(GO:0072273) |  | 26 | 0.88 | 1.35 | 0.000 | 0.844 | 0.325 | 6279 | tags=27%, list=11%, signal=30% |
| 578 | 3'\_5'\_CYCLIC\_AMP\_PHOSPHODIESTERASE\_ACTIVITY(GO:0004115) |  | 16 | 0.82 | 1.35 | 0.000 | 0.843 | 0.325 | 7935 | tags=56%, list=13%, signal=65% |
| 579 | REGULATION\_OF\_PROTEIN\_STABILITY(GO:0031647) |  | 253 | 0.81 | 1.35 | 0.000 | 0.842 | 0.325 | 6626 | tags=5%, list=11%, signal=6% |
| 580 | RESPONSE\_TO\_OSMOTIC\_STRESS(GO:0006970) |  | 72 | 0.85 | 1.35 | 0.000 | 0.841 | 0.325 | 6424 | tags=11%, list=11%, signal=12% |
| 581 | CONVERGENT\_EXTENSION(GO:0060026) |  | 15 | 0.82 | 1.35 | 0.000 | 0.840 | 0.325 | 7792 | tags=33%, list=13%, signal=38% |
| 582 | CELLULAR\_RESPONSE\_TO\_ACID\_CHEMICAL(GO:0071229) |  | 203 | 0.86 | 1.35 | 0.000 | 0.839 | 0.325 | 6662 | tags=16%, list=11%, signal=18% |
| 583 | INNER\_EAR\_MORPHOGENESIS(GO:0042472) |  | 100 | 0.86 | 1.35 | 0.000 | 0.837 | 0.325 | 6473 | tags=23%, list=11%, signal=26% |
| 584 | DORSAL\_VENTRAL\_PATTERN\_FORMATION(GO:0009953) |  | 97 | 0.87 | 1.35 | 0.000 | 0.837 | 0.325 | 6817 | tags=31%, list=12%, signal=35% |
| 585 | MICROVILLUS\_MEMBRANE(GO:0031528) |  | 23 | 0.90 | 1.35 | 0.000 | 0.837 | 0.325 | 5877 | tags=9%, list=10%, signal=10% |
| 586 | CELL\_ACTIVATION(GO:0001775) |  | 1368 | 0.85 | 1.35 | 0.141 | 0.837 | 0.325 | 6774 | tags=11%, list=11%, signal=12% |
| 587 | EXOPEPTIDASE\_ACTIVITY(GO:0008238) |  | 110 | 0.80 | 1.35 | 0.000 | 0.836 | 0.325 | 6596 | tags=8%, list=11%, signal=9% |
| 588 | RESPONSE\_TO\_CYTOKINE(GO:0034097) |  | 1156 | 0.85 | 1.35 | 0.141 | 0.836 | 0.325 | 6738 | tags=11%, list=11%, signal=13% |
| 589 | CAVEOLA(GO:0005901) |  | 78 | 0.86 | 1.35 | 0.000 | 0.835 | 0.325 | 6573 | tags=21%, list=11%, signal=23% |
| 590 | REGULATION\_OF\_MESENCHYMAL\_CELL\_PROLIFERATION(GO:0010464) |  | 36 | 0.88 | 1.35 | 0.000 | 0.834 | 0.325 | 6473 | tags=22%, list=11%, signal=25% |
| 591 | QUINONE\_METABOLIC\_PROCESS(GO:1901661) |  | 30 | 0.88 | 1.35 | 0.000 | 0.834 | 0.325 | 5839 | tags=3%, list=10%, signal=4% |
| 592 | NEGATIVE\_REGULATION\_OF\_NOTCH\_SIGNALING\_PATHWAY(GO:0045746) |  | 41 | 0.88 | 1.34 | 0.000 | 0.833 | 0.325 | 6155 | tags=5%, list=10%, signal=5% |
| 593 | EXTRINSIC\_APOPTOTIC\_SIGNALING\_PATHWAY(GO:0097191) |  | 223 | 0.86 | 1.34 | 0.141 | 0.833 | 0.325 | 6738 | tags=9%, list=11%, signal=11% |
| 594 | ORGANONITROGEN\_COMPOUND\_CATABOLIC\_PROCESS(GO:1901565) |  | 1223 | 0.82 | 1.34 | 0.000 | 0.832 | 0.325 | 6688 | tags=7%, list=11%, signal=7% |
| 595 | NEGATIVE\_REGULATION\_OF\_CATALYTIC\_ACTIVITY(GO:0043086) |  | 818 | 0.84 | 1.34 | 0.000 | 0.831 | 0.325 | 6695 | tags=10%, list=11%, signal=11% |
| 596 | HEART\_CONTRACTION(GO:0060047) |  | 274 | 0.85 | 1.34 | 0.141 | 0.830 | 0.325 | 7015 | tags=22%, list=12%, signal=25% |
| 597 | HEART\_PROCESS(GO:0003015) |  | 277 | 0.85 | 1.34 | 0.141 | 0.829 | 0.325 | 7015 | tags=22%, list=12%, signal=24% |
| 598 | NUCLEOSOME\_BINDING(GO:0031491) |  | 66 | 0.83 | 1.34 | 0.000 | 0.828 | 0.325 | 6422 | tags=5%, list=11%, signal=5% |
| 599 | POSITIVE\_REGULATION\_OF\_INTERFERON\_GAMMA\_PRODUCTION(GO:0032729) |  | 62 | 0.83 | 1.34 | 0.000 | 0.827 | 0.325 | 7258 | tags=23%, list=12%, signal=26% |
| 600 | TISSUE\_MORPHOGENESIS(GO:0048729) |  | 645 | 0.86 | 1.34 | 0.141 | 0.826 | 0.325 | 6535 | tags=13%, list=11%, signal=15% |
| 601 | NEGATIVE\_REGULATION\_OF\_CHEMOTAXIS(GO:0050922) |  | 41 | 0.89 | 1.34 | 0.141 | 0.826 | 0.325 | 6000 | tags=17%, list=10%, signal=19% |
| 602 | REGULATION\_OF\_CELL\_MIGRATION(GO:0030334) |  | 825 | 0.86 | 1.34 | 0.141 | 0.825 | 0.325 | 6578 | tags=12%, list=11%, signal=13% |
| 603 | REGULATION\_OF\_RELEASE\_OF\_SEQUESTERED\_CALCIUM\_ION\_INTO\_CYTOSOL\_BY\_SARCOPLASMIC\_RETICULUM(GO:0010880) |  | 30 | 0.85 | 1.34 | 0.000 | 0.824 | 0.325 | 7544 | tags=30%, list=13%, signal=34% |
| 604 | FATTY\_ACID\_DERIVATIVE\_METABOLIC\_PROCESS(GO:1901568) |  | 157 | 0.84 | 1.34 | 0.000 | 0.823 | 0.325 | 6424 | tags=9%, list=11%, signal=10% |
| 605 | SIGNALING\_RECEPTOR\_BINDING(GO:0005102) |  | 1553 | 0.85 | 1.34 | 0.141 | 0.822 | 0.325 | 6857 | tags=13%, list=12%, signal=14% |
| 606 | RESPONSE\_TO\_LIPID(GO:0033993) |  | 910 | 0.86 | 1.34 | 0.141 | 0.821 | 0.325 | 6535 | tags=13%, list=11%, signal=14% |
| 607 | REGULATION\_OF\_CELL\_MORPHOGENESIS(GO:0022604) |  | 439 | 0.86 | 1.34 | 0.141 | 0.820 | 0.325 | 7169 | tags=14%, list=12%, signal=16% |
| 608 | CELLULAR\_MACROMOLECULE\_CATABOLIC\_PROCESS(GO:0044265) |  | 1100 | 0.83 | 1.34 | 0.000 | 0.819 | 0.325 | 6626 | tags=4%, list=11%, signal=5% |
| 609 | INTERLEUKIN\_6\_PRODUCTION(GO:0032635) |  | 123 | 0.83 | 1.34 | 0.000 | 0.818 | 0.325 | 7256 | tags=18%, list=12%, signal=20% |
| 610 | REGULATION\_OF\_EXTRINSIC\_APOPTOTIC\_SIGNALING\_PATHWAY\_VIA\_DEATH\_DOMAIN\_RECEPTORS(GO:1902041) |  | 59 | 0.87 | 1.34 | 0.000 | 0.817 | 0.325 | 6212 | tags=14%, list=11%, signal=15% |
| 611 | DENDRITIC\_SPINE(GO:0043197) |  | 137 | 0.83 | 1.34 | 0.000 | 0.816 | 0.325 | 7962 | tags=23%, list=13%, signal=26% |
| 612 | RESPONSE\_TO\_OXIDATIVE\_STRESS(GO:0006979) |  | 435 | 0.85 | 1.34 | 0.000 | 0.815 | 0.325 | 6626 | tags=9%, list=11%, signal=10% |
| 613 | MICROVILLUS(GO:0005902) |  | 84 | 0.89 | 1.34 | 0.141 | 0.815 | 0.325 | 6115 | tags=11%, list=10%, signal=12% |
| 614 | PEROXISOME\_PROLIFERATOR\_ACTIVATED\_RECEPTOR\_SIGNALING\_PATHWAY(GO:0035357) |  | 20 | 0.87 | 1.34 | 0.000 | 0.815 | 0.325 | 6048 | tags=15%, list=10%, signal=17% |
| 615 | TRANSAMINASE\_ACTIVITY(GO:0008483) |  | 23 | 0.86 | 1.34 | 0.000 | 0.816 | 0.325 | 6145 | tags=13%, list=10%, signal=15% |
| 616 | ALDO\_KETO\_REDUCTASE\_(NADP)\_ACTIVITY(GO:0004033) |  | 25 | 0.88 | 1.34 | 0.000 | 0.815 | 0.325 | 5839 | tags=4%, list=10%, signal=4% |
| 617 | REGULATION\_OF\_RYANODINE\_SENSITIVE\_CALCIUM\_RELEASE\_CHANNEL\_ACTIVITY(GO:0060314) |  | 26 | 0.83 | 1.34 | 0.000 | 0.814 | 0.325 | 6560 | tags=15%, list=11%, signal=17% |
| 618 | HEART\_LOOPING(GO:0001947) |  | 61 | 0.85 | 1.34 | 0.000 | 0.813 | 0.325 | 6585 | tags=18%, list=11%, signal=20% |
| 619 | STEROID\_BIOSYNTHETIC\_PROCESS(GO:0006694) |  | 185 | 0.82 | 1.34 | 0.000 | 0.812 | 0.325 | 7634 | tags=15%, list=13%, signal=17% |
| 620 | POLY(A)\_BINDING(GO:0008143) |  | 16 | 0.82 | 1.34 | 0.000 | 0.811 | 0.325 | 5932 | tags=6%, list=10%, signal=7% |
| 621 | NEGATIVE\_REGULATION\_OF\_CELLULAR\_COMPONENT\_MOVEMENT(GO:0051271) |  | 326 | 0.85 | 1.34 | 0.141 | 0.811 | 0.325 | 6578 | tags=10%, list=11%, signal=11% |
| 622 | DETERMINATION\_OF\_HEART\_LEFT\_RIGHT\_ASYMMETRY(GO:0061371) |  | 66 | 0.85 | 1.34 | 0.000 | 0.810 | 0.325 | 6585 | tags=20%, list=11%, signal=22% |
| 623 | NEGATIVE\_REGULATION\_OF\_CELL\_MOTILITY(GO:2000146) |  | 309 | 0.85 | 1.34 | 0.141 | 0.809 | 0.325 | 6027 | tags=8%, list=10%, signal=9% |
| 624 | NEGATIVE\_REGULATION\_OF\_INTRACELLULAR\_STEROID\_HORMONE\_RECEPTOR\_SIGNALING\_PATHWAY(GO:0033144) |  | 34 | 0.84 | 1.34 | 0.000 | 0.809 | 0.325 | 6027 | tags=18%, list=10%, signal=20% |
| 625 | RESPONSE\_TO\_OXYGEN\_CONTAINING\_COMPOUND(GO:1901700) |  | 1563 | 0.86 | 1.34 | 0.141 | 0.808 | 0.325 | 6578 | tags=12%, list=11%, signal=13% |
| 626 | RESPONSE\_TO\_LIPOPOLYSACCHARIDE(GO:0032496) |  | 313 | 0.86 | 1.34 | 0.141 | 0.807 | 0.325 | 6486 | tags=12%, list=11%, signal=13% |
| 627 | TRANSCRIPTIONAL\_ACTIVATOR\_ACTIVITY\_\_RNA\_POLYMERASE\_II\_DISTAL\_ENHANCER\_SEQUENCE\_SPECIFIC\_DNA\_BINDING(GO:0001205) |  | 30 | 0.83 | 1.34 | 0.000 | 0.807 | 0.325 | 7834 | tags=30%, list=13%, signal=35% |
| 628 | NEGATIVE\_REGULATION\_OF\_LOCOMOTION(GO:0040013) |  | 339 | 0.85 | 1.34 | 0.141 | 0.807 | 0.325 | 6578 | tags=10%, list=11%, signal=11% |
| 629 | CHONDROCYTE\_PROLIFERATION(GO:0035988) |  | 18 | 0.86 | 1.34 | 0.000 | 0.806 | 0.325 | 6831 | tags=28%, list=12%, signal=31% |
| 630 | REGULATION\_OF\_I\_KAPPAB\_KINASE\_NF\_KAPPAB\_SIGNALING(GO:0043122) |  | 213 | 0.87 | 1.34 | 0.141 | 0.807 | 0.325 | 6706 | tags=10%, list=11%, signal=11% |
| 631 | INTEGRAL\_COMPONENT\_OF\_SYNAPTIC\_VESICLE\_MEMBRANE(GO:0030285) |  | 15 | 0.86 | 1.34 | 0.000 | 0.806 | 0.325 | 6046 | tags=20%, list=10%, signal=22% |
| 632 | GLIOGENESIS(GO:0042063) |  | 262 | 0.87 | 1.34 | 0.141 | 0.805 | 0.325 | 6442 | tags=16%, list=11%, signal=18% |
| 633 | NEGATIVE\_REGULATION\_OF\_PROTEIN\_PHOSPHORYLATION(GO:0001933) |  | 431 | 0.84 | 1.34 | 0.141 | 0.804 | 0.325 | 6695 | tags=8%, list=11%, signal=9% |
| 634 | GLIAL\_CELL\_ACTIVATION(GO:0061900) |  | 31 | 0.87 | 1.34 | 0.000 | 0.804 | 0.325 | 6413 | tags=19%, list=11%, signal=22% |
| 635 | LEUKOCYTE\_ACTIVATION\_INVOLVED\_IN\_IMMUNE\_RESPONSE(GO:0002366) |  | 679 | 0.82 | 1.34 | 0.000 | 0.803 | 0.325 | 6696 | tags=8%, list=11%, signal=9% |
| 636 | LEUKOCYTE\_ACTIVATION(GO:0045321) |  | 1214 | 0.84 | 1.34 | 0.141 | 0.802 | 0.325 | 6738 | tags=10%, list=11%, signal=11% |
| 637 | CORNIFIED\_ENVELOPE(GO:0001533) |  | 63 | 0.90 | 1.34 | 0.141 | 0.801 | 0.325 | 5827 | tags=5%, list=10%, signal=5% |
| 638 | KINASE\_BINDING(GO:0019900) |  | 665 | 0.86 | 1.34 | 0.141 | 0.800 | 0.325 | 6218 | tags=6%, list=11%, signal=6% |
| 639 | CILIUM\_DEPENDENT\_CELL\_MOTILITY(GO:0060285) |  | 28 | 0.82 | 1.34 | 0.000 | 0.800 | 0.325 | 7628 | tags=32%, list=13%, signal=37% |
| 640 | REGULATION\_OF\_PROTEASOMAL\_PROTEIN\_CATABOLIC\_PROCESS(GO:0061136) |  | 171 | 0.81 | 1.34 | 0.000 | 0.799 | 0.325 | 6133 | tags=4%, list=10%, signal=5% |
| 641 | RESPONSE\_TO\_MOLECULE\_OF\_BACTERIAL\_ORIGIN(GO:0002237) |  | 330 | 0.86 | 1.34 | 0.141 | 0.798 | 0.325 | 6530 | tags=12%, list=11%, signal=13% |
| 642 | BLOOD\_VESSEL\_MORPHOGENESIS(GO:0048514) |  | 653 | 0.85 | 1.34 | 0.141 | 0.798 | 0.325 | 7414 | tags=21%, list=13%, signal=24% |
| 643 | REGULATION\_OF\_JAK\_STAT\_CASCADE(GO:0046425) |  | 153 | 0.83 | 1.34 | 0.141 | 0.797 | 0.325 | 7449 | tags=21%, list=13%, signal=24% |
| 644 | CELL\_ACTIVATION\_INVOLVED\_IN\_IMMUNE\_RESPONSE(GO:0002263) |  | 683 | 0.82 | 1.34 | 0.000 | 0.796 | 0.325 | 6696 | tags=8%, list=11%, signal=9% |
| 645 | MESENCHYME\_MORPHOGENESIS(GO:0072132) |  | 47 | 0.84 | 1.34 | 0.000 | 0.795 | 0.325 | 8177 | tags=45%, list=14%, signal=52% |
| 646 | REGULATION\_OF\_CELL\_MOTILITY(GO:2000145) |  | 877 | 0.85 | 1.34 | 0.141 | 0.795 | 0.325 | 6578 | tags=12%, list=11%, signal=13% |
| 647 | PLASMA\_MEMBRANE\_RAFT(GO:0044853) |  | 98 | 0.85 | 1.34 | 0.000 | 0.794 | 0.325 | 7426 | tags=24%, list=13%, signal=28% |
| 648 | MAMMARY\_GLAND\_EPITHELIUM\_DEVELOPMENT(GO:0061180) |  | 70 | 0.88 | 1.34 | 0.141 | 0.795 | 0.325 | 6473 | tags=17%, list=11%, signal=19% |
| 649 | REGULATION\_OF\_DOUBLE\_STRAND\_BREAK\_REPAIR\_VIA\_HOMOLOGOUS\_RECOMBINATION(GO:0010569) |  | 31 | 0.81 | 1.34 | 0.000 | 0.794 | 0.325 | 6893 | tags=10%, list=12%, signal=11% |
| 650 | ANTIBIOTIC\_METABOLIC\_PROCESS(GO:0016999) |  | 136 | 0.86 | 1.34 | 0.000 | 0.793 | 0.325 | 6612 | tags=10%, list=11%, signal=11% |
| 651 | EMBRYONIC\_HEART\_TUBE\_MORPHOGENESIS(GO:0003143) |  | 67 | 0.85 | 1.34 | 0.000 | 0.792 | 0.325 | 7134 | tags=21%, list=12%, signal=24% |
| 652 | NEGATIVE\_REGULATION\_OF\_IMMUNE\_SYSTEM\_PROCESS(GO:0002683) |  | 412 | 0.83 | 1.34 | 0.000 | 0.792 | 0.325 | 7170 | tags=13%, list=12%, signal=15% |
| 653 | LIPID\_METABOLIC\_PROCESS(GO:0006629) |  | 1424 | 0.84 | 1.34 | 0.000 | 0.791 | 0.325 | 6688 | tags=11%, list=11%, signal=12% |
| 654 | CELL\_CELL\_SIGNALING\_INVOLVED\_IN\_CARDIAC\_CONDUCTION(GO:0086019) |  | 31 | 0.88 | 1.34 | 0.000 | 0.791 | 0.325 | 6670 | tags=42%, list=11%, signal=47% |
| 655 | I\_KAPPAB\_KINASE\_NF\_KAPPAB\_SIGNALING(GO:0007249) |  | 244 | 0.86 | 1.34 | 0.141 | 0.790 | 0.325 | 6706 | tags=9%, list=11%, signal=10% |
| 656 | CELLULAR\_RESPONSE\_TO\_EXTERNAL\_STIMULUS(GO:0071496) |  | 305 | 0.85 | 1.34 | 0.141 | 0.789 | 0.325 | 6497 | tags=9%, list=11%, signal=10% |
| 657 | HORMONE\_BIOSYNTHETIC\_PROCESS(GO:0042446) |  | 84 | 0.84 | 1.34 | 0.000 | 0.788 | 0.325 | 5839 | tags=8%, list=10%, signal=9% |
| 658 | REGULATION\_OF\_STAT\_CASCADE(GO:1904892) |  | 155 | 0.83 | 1.34 | 0.141 | 0.787 | 0.325 | 7449 | tags=21%, list=13%, signal=24% |
| 659 | CYTOKINE\_SECRETION\_INVOLVED\_IN\_IMMUNE\_RESPONSE(GO:0002374) |  | 17 | 0.83 | 1.34 | 0.000 | 0.786 | 0.325 | 7233 | tags=24%, list=12%, signal=27% |
| 660 | NEGATIVE\_REGULATION\_OF\_CHROMATIN\_ORGANIZATION(GO:1905268) |  | 48 | 0.82 | 1.34 | 0.000 | 0.787 | 0.325 | 5904 | tags=6%, list=10%, signal=7% |
| 661 | SKELETAL\_MUSCLE\_ORGAN\_DEVELOPMENT(GO:0060538) |  | 175 | 0.85 | 1.34 | 0.000 | 0.786 | 0.325 | 6918 | tags=19%, list=12%, signal=22% |
| 662 | REGULATION\_OF\_CELLULAR\_PROTEIN\_LOCALIZATION(GO:1903827) |  | 512 | 0.86 | 1.34 | 0.141 | 0.785 | 0.325 | 6662 | tags=6%, list=11%, signal=7% |
| 663 | EMBRYONIC\_HEART\_TUBE\_DEVELOPMENT(GO:0035050) |  | 77 | 0.85 | 1.34 | 0.000 | 0.785 | 0.325 | 7343 | tags=23%, list=12%, signal=27% |
| 664 | GLIAL\_CELL\_DIFFERENTIATION(GO:0010001) |  | 194 | 0.87 | 1.34 | 0.000 | 0.784 | 0.325 | 6442 | tags=16%, list=11%, signal=18% |
| 665 | REGULATION\_OF\_CARDIAC\_MUSCLE\_CELL\_ACTION\_POTENTIAL(GO:0098901) |  | 32 | 0.85 | 1.34 | 0.000 | 0.783 | 0.325 | 7008 | tags=22%, list=12%, signal=25% |
| 666 | LAMELLIPODIUM\_MEMBRANE(GO:0031258) |  | 22 | 0.85 | 1.34 | 0.000 | 0.782 | 0.325 | 6248 | tags=9%, list=11%, signal=10% |
| 667 | POSITIVE\_REGULATION\_OF\_ORGANELLE\_ORGANIZATION(GO:0010638) |  | 632 | 0.84 | 1.34 | 0.000 | 0.781 | 0.325 | 6218 | tags=6%, list=11%, signal=6% |
| 668 | REGULATION\_OF\_CARDIAC\_MUSCLE\_CONTRACTION\_BY\_REGULATION\_OF\_THE\_RELEASE\_OF\_SEQUESTERED\_CALCIUM\_ION(GO:0010881) |  | 22 | 0.86 | 1.34 | 0.000 | 0.780 | 0.325 | 7008 | tags=27%, list=12%, signal=31% |
| 669 | REGULATION\_OF\_CATECHOLAMINE\_SECRETION(GO:0050433) |  | 48 | 0.86 | 1.34 | 0.000 | 0.780 | 0.325 | 6248 | tags=25%, list=11%, signal=28% |
| 670 | STRIATED\_MUSCLE\_TISSUE\_DEVELOPMENT(GO:0014706) |  | 378 | 0.84 | 1.34 | 0.000 | 0.780 | 0.325 | 6918 | tags=19%, list=12%, signal=22% |
| 671 | DRUG\_CATABOLIC\_PROCESS(GO:0042737) |  | 117 | 0.85 | 1.34 | 0.000 | 0.779 | 0.325 | 6555 | tags=10%, list=11%, signal=12% |
| 672 | REGULATION\_OF\_STRIATED\_MUSCLE\_CONTRACTION(GO:0006942) |  | 94 | 0.82 | 1.34 | 0.000 | 0.779 | 0.325 | 7979 | tags=22%, list=13%, signal=26% |
| 673 | REGULATION\_OF\_SYNAPTIC\_VESICLE\_RECYCLING(GO:1903421) |  | 17 | 0.84 | 1.34 | 0.000 | 0.779 | 0.325 | 7426 | tags=29%, list=13%, signal=34% |
| 674 | CILIUM\_MOVEMENT\_INVOLVED\_IN\_CELL\_MOTILITY(GO:0060294) |  | 18 | 0.83 | 1.34 | 0.000 | 0.778 | 0.325 | 7628 | tags=39%, list=13%, signal=45% |
| 675 | REGULATION\_OF\_LOCOMOTION(GO:0040012) |  | 947 | 0.85 | 1.34 | 0.141 | 0.777 | 0.325 | 6578 | tags=12%, list=11%, signal=13% |
| 676 | CATECHOLAMINE\_TRANSPORT(GO:0051937) |  | 64 | 0.85 | 1.34 | 0.000 | 0.777 | 0.325 | 7298 | tags=25%, list=12%, signal=28% |
| 677 | REGULATION\_OF\_CARDIAC\_MUSCLE\_CONTRACTION(GO:0055117) |  | 79 | 0.82 | 1.34 | 0.000 | 0.776 | 0.325 | 7979 | tags=22%, list=13%, signal=25% |
| 678 | INFLAMMATORY\_RESPONSE(GO:0006954) |  | 754 | 0.84 | 1.34 | 0.141 | 0.775 | 0.325 | 7238 | tags=15%, list=12%, signal=17% |
| 679 | CELLULAR\_RESPONSE\_TO\_VASCULAR\_ENDOTHELIAL\_GROWTH\_FACTOR\_STIMULUS(GO:0035924) |  | 55 | 0.85 | 1.34 | 0.000 | 0.775 | 0.325 | 7754 | tags=33%, list=13%, signal=38% |
| 680 | RESPONSE\_TO\_COPPER\_ION(GO:0046688) |  | 28 | 0.84 | 1.34 | 0.000 | 0.774 | 0.325 | 6374 | tags=11%, list=11%, signal=12% |
| 681 | NEGATIVE\_REGULATION\_OF\_DNA\_BINDING(GO:0043392) |  | 51 | 0.87 | 1.34 | 0.000 | 0.775 | 0.325 | 6381 | tags=14%, list=11%, signal=15% |
| 682 | MAMMARY\_GLAND\_DEVELOPMENT(GO:0030879) |  | 141 | 0.87 | 1.34 | 0.141 | 0.774 | 0.325 | 6473 | tags=14%, list=11%, signal=16% |
| 683 | MUSCLE\_TISSUE\_DEVELOPMENT(GO:0060537) |  | 395 | 0.84 | 1.34 | 0.000 | 0.773 | 0.325 | 7408 | tags=21%, list=13%, signal=24% |
| 684 | RESPONSE\_TO\_TUMOR\_CELL(GO:0002347) |  | 18 | 0.84 | 1.34 | 0.000 | 0.773 | 0.325 | 6738 | tags=33%, list=11%, signal=38% |
| 685 | MONOCARBOXYLIC\_ACID\_METABOLIC\_PROCESS(GO:0032787) |  | 604 | 0.84 | 1.34 | 0.000 | 0.772 | 0.325 | 6797 | tags=10%, list=11%, signal=11% |
| 686 | SECRETORY\_GRANULE(GO:0030141) |  | 824 | 0.83 | 1.34 | 0.000 | 0.771 | 0.325 | 7546 | tags=12%, list=13%, signal=14% |
| 687 | CELL\_FATE\_DETERMINATION(GO:0001709) |  | 42 | 0.88 | 1.34 | 0.000 | 0.771 | 0.325 | 6297 | tags=33%, list=11%, signal=37% |
| 688 | NEGATIVE\_REGULATION\_OF\_EXOCYTOSIS(GO:0045920) |  | 30 | 0.84 | 1.34 | 0.000 | 0.770 | 0.325 | 7457 | tags=20%, list=13%, signal=23% |
| 689 | NUCLEOSIDE\_METABOLIC\_PROCESS(GO:0009116) |  | 128 | 0.79 | 1.34 | 0.000 | 0.769 | 0.325 | 7635 | tags=11%, list=13%, signal=13% |
| 690 | REGULATION\_OF\_INTERLEUKIN\_6\_PRODUCTION(GO:0032675) |  | 114 | 0.83 | 1.34 | 0.000 | 0.769 | 0.325 | 7256 | tags=18%, list=12%, signal=20% |
| 691 | CELL\_FATE\_COMMITMENT\_INVOLVED\_IN\_FORMATION\_OF\_PRIMARY\_GERM\_LAYER(GO:0060795) |  | 35 | 0.85 | 1.34 | 0.000 | 0.768 | 0.325 | 6871 | tags=17%, list=12%, signal=19% |
| 692 | CELL\_CELL\_ADHESION(GO:0098609) |  | 768 | 0.86 | 1.34 | 0.141 | 0.767 | 0.325 | 6758 | tags=14%, list=11%, signal=15% |
| 693 | RETINA\_VASCULATURE\_DEVELOPMENT\_IN\_CAMERA\_TYPE\_EYE(GO:0061298) |  | 16 | 0.89 | 1.34 | 0.000 | 0.766 | 0.325 | 6218 | tags=25%, list=11%, signal=28% |
| 694 | POSITIVE\_REGULATION\_OF\_EPITHELIAL\_TO\_MESENCHYMAL\_TRANSITION(GO:0010718) |  | 48 | 0.84 | 1.34 | 0.000 | 0.766 | 0.325 | 6381 | tags=15%, list=11%, signal=16% |
| 695 | INOSITOL\_PHOSPHATE\_BIOSYNTHETIC\_PROCESS(GO:0032958) |  | 27 | 0.87 | 1.34 | 0.000 | 0.766 | 0.325 | 5955 | tags=22%, list=10%, signal=25% |
| 696 | ORGANIC\_HYDROXY\_COMPOUND\_CATABOLIC\_PROCESS(GO:1901616) |  | 63 | 0.85 | 1.34 | 0.141 | 0.766 | 0.325 | 5920 | tags=10%, list=10%, signal=11% |
| 697 | CYTOKINE\_PRODUCTION(GO:0001816) |  | 691 | 0.83 | 1.34 | 0.141 | 0.766 | 0.325 | 7488 | tags=15%, list=13%, signal=17% |
| 698 | METANEPHRIC\_NEPHRON\_DEVELOPMENT(GO:0072210) |  | 41 | 0.87 | 1.34 | 0.141 | 0.765 | 0.325 | 6831 | tags=24%, list=12%, signal=28% |
| 699 | SKELETAL\_MUSCLE\_TISSUE\_DEVELOPMENT(GO:0007519) |  | 166 | 0.86 | 1.34 | 0.000 | 0.764 | 0.325 | 6918 | tags=20%, list=12%, signal=22% |
| 700 | MONOAMINE\_TRANSPORT(GO:0015844) |  | 74 | 0.84 | 1.34 | 0.000 | 0.764 | 0.325 | 7298 | tags=24%, list=12%, signal=28% |
| 701 | REGULATION\_OF\_RESPONSE\_TO\_EXTERNAL\_STIMULUS(GO:0032101) |  | 782 | 0.84 | 1.34 | 0.141 | 0.765 | 0.325 | 7546 | tags=17%, list=13%, signal=19% |
| 702 | ANGIOGENESIS(GO:0001525) |  | 564 | 0.85 | 1.34 | 0.141 | 0.765 | 0.325 | 7414 | tags=21%, list=13%, signal=23% |
| 703 | KERATINOCYTE\_DIFFERENTIATION(GO:0030216) |  | 298 | 0.87 | 1.34 | 0.141 | 0.764 | 0.325 | 6473 | tags=8%, list=11%, signal=9% |
| 704 | REGULATION\_OF\_JNK\_CASCADE(GO:0046328) |  | 167 | 0.87 | 1.34 | 0.141 | 0.763 | 0.325 | 6797 | tags=11%, list=11%, signal=12% |
| 705 | MUSCLE\_CONTRACTION(GO:0006936) |  | 354 | 0.83 | 1.34 | 0.000 | 0.763 | 0.325 | 7979 | tags=26%, list=13%, signal=30% |
| 706 | REGULATION\_OF\_CELLULAR\_COMPONENT\_MOVEMENT(GO:0051270) |  | 957 | 0.85 | 1.34 | 0.141 | 0.762 | 0.325 | 6681 | tags=12%, list=11%, signal=14% |
| 707 | REGULATION\_OF\_PROTEIN\_SERINE\_THREONINE\_KINASE\_ACTIVITY(GO:0071900) |  | 500 | 0.84 | 1.34 | 0.141 | 0.761 | 0.325 | 6940 | tags=11%, list=12%, signal=13% |
| 708 | PYRIMIDINE\_CONTAINING\_COMPOUND\_CATABOLIC\_PROCESS(GO:0072529) |  | 34 | 0.81 | 1.34 | 0.000 | 0.760 | 0.325 | 6865 | tags=18%, list=12%, signal=20% |
| 709 | CYTOSOLIC\_CALCIUM\_ION\_TRANSPORT(GO:0060401) |  | 161 | 0.84 | 1.34 | 0.000 | 0.760 | 0.325 | 6706 | tags=15%, list=11%, signal=17% |
| 710 | METANEPHROS\_MORPHOGENESIS(GO:0003338) |  | 33 | 0.88 | 1.34 | 0.141 | 0.759 | 0.325 | 6279 | tags=21%, list=11%, signal=24% |
| 711 | VASCULAR\_PROCESS\_IN\_CIRCULATORY\_SYSTEM(GO:0003018) |  | 169 | 0.84 | 1.34 | 0.000 | 0.758 | 0.325 | 7937 | tags=27%, list=13%, signal=31% |
| 712 | TISSUE\_HOMEOSTASIS(GO:0001894) |  | 207 | 0.84 | 1.34 | 0.141 | 0.757 | 0.325 | 7504 | tags=20%, list=13%, signal=23% |
| 713 | CELL\_MIGRATION(GO:0016477) |  | 1435 | 0.85 | 1.34 | 0.141 | 0.757 | 0.325 | 7134 | tags=14%, list=12%, signal=16% |
| 714 | REGULATION\_OF\_CELLULAR\_RESPIRATION(GO:0043457) |  | 24 | 0.84 | 1.34 | 0.000 | 0.756 | 0.325 | 6247 | tags=13%, list=11%, signal=14% |
| 715 | VASCULATURE\_DEVELOPMENT(GO:0001944) |  | 758 | 0.85 | 1.34 | 0.141 | 0.755 | 0.325 | 7427 | tags=21%, list=13%, signal=23% |
| 716 | SYNAPSE\_PART(GO:0044456) |  | 705 | 0.82 | 1.34 | 0.000 | 0.755 | 0.325 | 7487 | tags=20%, list=13%, signal=23% |
| 717 | CELL\_CELL\_JUNCTION\_ASSEMBLY(GO:0007043) |  | 95 | 0.86 | 1.34 | 0.141 | 0.754 | 0.325 | 6495 | tags=11%, list=11%, signal=12% |
| 718 | REGULATION\_OF\_T\_CELL\_DIFFERENTIATION\_IN\_THYMUS(GO:0033081) |  | 25 | 0.87 | 1.34 | 0.000 | 0.753 | 0.325 | 6154 | tags=16%, list=10%, signal=18% |
| 719 | GLYCOSYL\_COMPOUND\_CATABOLIC\_PROCESS(GO:1901658) |  | 38 | 0.79 | 1.34 | 0.000 | 0.753 | 0.325 | 6865 | tags=16%, list=12%, signal=18% |
| 720 | REGULATION\_OF\_STRESS\_ACTIVATED\_MAPK\_CASCADE(GO:0032872) |  | 214 | 0.86 | 1.34 | 0.141 | 0.753 | 0.325 | 6797 | tags=11%, list=11%, signal=13% |
| 721 | ENDOCYTIC\_VESICLE\_MEMBRANE(GO:0030666) |  | 155 | 0.83 | 1.34 | 0.000 | 0.752 | 0.325 | 6912 | tags=14%, list=12%, signal=15% |
| 722 | CARDIOVASCULAR\_SYSTEM\_DEVELOPMENT(GO:0072358) |  | 766 | 0.85 | 1.34 | 0.141 | 0.752 | 0.325 | 7427 | tags=21%, list=13%, signal=23% |
| 723 | COMPLEX\_OF\_COLLAGEN\_TRIMERS(GO:0098644) |  | 20 | 0.89 | 1.34 | 0.000 | 0.751 | 0.325 | 5973 | tags=20%, list=10%, signal=22% |
| 724 | REGULATION\_OF\_NEUROBLAST\_PROLIFERATION(GO:1902692) |  | 27 | 0.89 | 1.34 | 0.000 | 0.750 | 0.325 | 5953 | tags=22%, list=10%, signal=25% |
| 725 | MESODERM\_MORPHOGENESIS(GO:0048332) |  | 78 | 0.84 | 1.34 | 0.000 | 0.749 | 0.325 | 7358 | tags=24%, list=12%, signal=28% |
| 726 | FATTY\_ACID\_LIGASE\_ACTIVITY(GO:0015645) |  | 21 | 0.83 | 1.34 | 0.000 | 0.749 | 0.325 | 6782 | tags=14%, list=11%, signal=16% |
| 727 | NEGATIVE\_REGULATION\_OF\_AMINE\_TRANSPORT(GO:0051953) |  | 23 | 0.87 | 1.34 | 0.000 | 0.749 | 0.325 | 5955 | tags=17%, list=10%, signal=19% |
| 728 | REGULATION\_OF\_AMINE\_TRANSPORT(GO:0051952) |  | 79 | 0.83 | 1.34 | 0.000 | 0.749 | 0.325 | 7435 | tags=24%, list=13%, signal=27% |
| 729 | REGULATION\_OF\_STRESS\_ACTIVATED\_PROTEIN\_KINASE\_SIGNALING\_CASCADE(GO:0070302) |  | 215 | 0.86 | 1.34 | 0.141 | 0.748 | 0.325 | 6797 | tags=11%, list=11%, signal=13% |
| 730 | POSITIVE\_REGULATION\_OF\_TRANSPORT(GO:0051050) |  | 942 | 0.85 | 1.34 | 0.141 | 0.747 | 0.325 | 6797 | tags=12%, list=11%, signal=14% |
| 731 | REGULATION\_OF\_DEFENSE\_RESPONSE(GO:0031347) |  | 789 | 0.83 | 1.34 | 0.141 | 0.747 | 0.325 | 7256 | tags=12%, list=12%, signal=14% |
| 732 | REGULATION\_OF\_TUBE\_DIAMETER(GO:0035296) |  | 135 | 0.84 | 1.34 | 0.000 | 0.746 | 0.325 | 7845 | tags=27%, list=13%, signal=31% |
| 733 | REGULATION\_OF\_BLOOD\_VESSEL\_DIAMETER(GO:0097746) |  | 135 | 0.84 | 1.34 | 0.000 | 0.745 | 0.325 | 7845 | tags=27%, list=13%, signal=31% |
| 734 | NEURON\_FATE\_COMMITMENT(GO:0048663) |  | 70 | 0.88 | 1.34 | 0.000 | 0.744 | 0.325 | 6338 | tags=40%, list=11%, signal=45% |
| 735 | POSITIVE\_REGULATION\_OF\_PROTEIN\_LOCALIZATION\_TO\_CELL\_PERIPHERY(GO:1904377) |  | 60 | 0.84 | 1.34 | 0.000 | 0.744 | 0.325 | 6642 | tags=10%, list=11%, signal=11% |
| 736 | RETINAL\_GANGLION\_CELL\_AXON\_GUIDANCE(GO:0031290) |  | 20 | 0.84 | 1.34 | 0.000 | 0.744 | 0.325 | 7211 | tags=40%, list=12%, signal=46% |
| 737 | MESODERMAL\_CELL\_DIFFERENTIATION(GO:0048333) |  | 35 | 0.87 | 1.34 | 0.000 | 0.743 | 0.325 | 6871 | tags=26%, list=12%, signal=29% |
| 738 | NEGATIVE\_REGULATION\_OF\_INTRINSIC\_APOPTOTIC\_SIGNALING\_PATHWAY(GO:2001243) |  | 97 | 0.84 | 1.34 | 0.141 | 0.742 | 0.325 | 6495 | tags=7%, list=11%, signal=8% |
| 739 | NEGATIVE\_REGULATION\_OF\_PHOSPHORYLATION(GO:0042326) |  | 467 | 0.84 | 1.34 | 0.141 | 0.742 | 0.325 | 6695 | tags=8%, list=11%, signal=9% |
| 740 | GAP\_JUNCTION\_CHANNEL\_ACTIVITY(GO:0005243) |  | 16 | 0.89 | 1.34 | 0.000 | 0.741 | 0.325 | 5959 | tags=25%, list=10%, signal=28% |
| 741 | REGULATION\_OF\_PROTEOLYSIS\_INVOLVED\_IN\_CELLULAR\_PROTEIN\_CATABOLIC\_PROCESS(GO:1903050) |  | 200 | 0.80 | 1.34 | 0.000 | 0.741 | 0.325 | 6154 | tags=5%, list=10%, signal=5% |
| 742 | REGULATION\_OF\_HEART\_CONTRACTION(GO:0008016) |  | 245 | 0.84 | 1.34 | 0.141 | 0.740 | 0.325 | 7015 | tags=22%, list=12%, signal=25% |
| 743 | NEURONAL\_ACTION\_POTENTIAL(GO:0019228) |  | 34 | 0.85 | 1.34 | 0.000 | 0.739 | 0.325 | 6844 | tags=32%, list=12%, signal=37% |
| 744 | REGULATION\_OF\_NOTCH\_SIGNALING\_PATHWAY(GO:0008593) |  | 93 | 0.85 | 1.34 | 0.000 | 0.739 | 0.325 | 6155 | tags=3%, list=10%, signal=4% |
| 745 | NEUROBLAST\_PROLIFERATION(GO:0007405) |  | 54 | 0.87 | 1.34 | 0.000 | 0.738 | 0.325 | 6381 | tags=17%, list=11%, signal=19% |
| 746 | STRESS\_ACTIVATED\_PROTEIN\_KINASE\_SIGNALING\_CASCADE(GO:0031098) |  | 294 | 0.86 | 1.34 | 0.141 | 0.738 | 0.325 | 6832 | tags=10%, list=12%, signal=11% |
| 747 | COLUMNAR\_CUBOIDAL\_EPITHELIAL\_CELL\_DEVELOPMENT(GO:0002066) |  | 57 | 0.86 | 1.34 | 0.141 | 0.737 | 0.325 | 6473 | tags=18%, list=11%, signal=20% |
| 748 | FOREBRAIN\_GENERATION\_OF\_NEURONS(GO:0021872) |  | 67 | 0.86 | 1.34 | 0.000 | 0.736 | 0.325 | 6473 | tags=21%, list=11%, signal=23% |
| 749 | DESMOSOME(GO:0030057) |  | 25 | 0.89 | 1.34 | 0.141 | 0.736 | 0.325 | 6011 | tags=8%, list=10%, signal=9% |
| 750 | NEGATIVE\_REGULATION\_OF\_EXTRINSIC\_APOPTOTIC\_SIGNALING\_PATHWAY\_IN\_ABSENCE\_OF\_LIGAND(GO:2001240) |  | 36 | 0.82 | 1.34 | 0.000 | 0.735 | 0.325 | 7322 | tags=19%, list=12%, signal=22% |
| 751 | NEGATIVE\_REGULATION\_OF\_SIGNAL\_TRANSDUCTION\_IN\_ABSENCE\_OF\_LIGAND(GO:1901099) |  | 36 | 0.82 | 1.34 | 0.000 | 0.734 | 0.325 | 7322 | tags=19%, list=12%, signal=22% |
| 752 | REGULATION\_OF\_MYELOID\_CELL\_DIFFERENTIATION(GO:0045637) |  | 246 | 0.84 | 1.34 | 0.000 | 0.734 | 0.325 | 6706 | tags=12%, list=11%, signal=13% |
| 753 | PERINUCLEAR\_REGION\_OF\_CYTOPLASM(GO:0048471) |  | 673 | 0.85 | 1.34 | 0.141 | 0.733 | 0.325 | 6626 | tags=9%, list=11%, signal=10% |
| 754 | CELLULAR\_RESPONSE\_TO\_ANGIOTENSIN(GO:1904385) |  | 24 | 0.85 | 1.34 | 0.000 | 0.733 | 0.325 | 6005 | tags=17%, list=10%, signal=19% |
| 755 | BLOOD\_VESSEL\_DEVELOPMENT(GO:0001568) |  | 733 | 0.85 | 1.34 | 0.141 | 0.732 | 0.325 | 7427 | tags=20%, list=13%, signal=23% |
| 756 | NEGATIVE\_REGULATION\_OF\_OSTEOCLAST\_DIFFERENTIATION(GO:0045671) |  | 26 | 0.81 | 1.33 | 0.000 | 0.732 | 0.325 | 7504 | tags=27%, list=13%, signal=31% |
| 757 | MORPHOGENESIS\_OF\_A\_BRANCHING\_EPITHELIUM(GO:0061138) |  | 183 | 0.87 | 1.33 | 0.141 | 0.731 | 0.325 | 6495 | tags=20%, list=11%, signal=23% |
| 758 | JNK\_CASCADE(GO:0007254) |  | 199 | 0.86 | 1.33 | 0.141 | 0.731 | 0.325 | 6797 | tags=10%, list=11%, signal=11% |
| 759 | MESODERM\_FORMATION(GO:0001707) |  | 76 | 0.85 | 1.33 | 0.000 | 0.730 | 0.325 | 7358 | tags=25%, list=12%, signal=29% |
| 760 | CALCIUM\_ION\_TRANSMEMBRANE\_TRANSPORT\_VIA\_HIGH\_VOLTAGE\_GATED\_CALCIUM\_CHANNEL(GO:0061577) |  | 15 | 0.88 | 1.33 | 0.000 | 0.729 | 0.325 | 6332 | tags=13%, list=11%, signal=15% |
| 761 | MULTICELLULAR\_ORGANISMAL\_SIGNALING(GO:0035637) |  | 199 | 0.84 | 1.33 | 0.141 | 0.729 | 0.325 | 6844 | tags=25%, list=12%, signal=28% |
| 762 | CELL\_CELL\_JUNCTION(GO:0005911) |  | 427 | 0.85 | 1.33 | 0.141 | 0.728 | 0.325 | 7055 | tags=14%, list=12%, signal=16% |
| 763 | CRANIAL\_NERVE\_MORPHOGENESIS(GO:0021602) |  | 30 | 0.88 | 1.33 | 0.000 | 0.727 | 0.325 | 6733 | tags=37%, list=11%, signal=41% |
| 764 | ARACHIDONIC\_ACID\_METABOLIC\_PROCESS(GO:0019369) |  | 54 | 0.84 | 1.33 | 0.000 | 0.727 | 0.325 | 6424 | tags=17%, list=11%, signal=19% |
| 765 | RENAL\_SYSTEM\_PROCESS(GO:0003014) |  | 115 | 0.84 | 1.33 | 0.141 | 0.726 | 0.325 | 7038 | tags=18%, list=12%, signal=21% |
| 766 | LEUKOCYTE\_MEDIATED\_IMMUNITY(GO:0002443) |  | 833 | 0.81 | 1.33 | 0.000 | 0.725 | 0.325 | 7515 | tags=8%, list=13%, signal=9% |
| 767 | CATECHOLAMINE\_SECRETION(GO:0050432) |  | 51 | 0.86 | 1.33 | 0.000 | 0.725 | 0.325 | 6248 | tags=24%, list=11%, signal=26% |
| 768 | CADHERIN\_BINDING\_INVOLVED\_IN\_CELL\_CELL\_ADHESION(GO:0098641) |  | 18 | 0.89 | 1.33 | 0.141 | 0.725 | 0.325 | 5845 | tags=6%, list=10%, signal=6% |
| 769 | NEGATIVE\_REGULATION\_OF\_PEPTIDYL\_SERINE\_PHOSPHORYLATION(GO:0033137) |  | 26 | 0.86 | 1.33 | 0.000 | 0.724 | 0.325 | 5856 | tags=8%, list=10%, signal=9% |
| 770 | RESPONSE\_TO\_BIOTIC\_STIMULUS(GO:0009607) |  | 937 | 0.84 | 1.33 | 0.141 | 0.724 | 0.325 | 7265 | tags=11%, list=12%, signal=13% |
| 771 | REGULATION\_OF\_TUBE\_SIZE(GO:0035150) |  | 144 | 0.84 | 1.33 | 0.000 | 0.723 | 0.325 | 7937 | tags=28%, list=13%, signal=32% |
| 772 | REGULATION\_OF\_BLOOD\_VESSEL\_SIZE(GO:0050880) |  | 143 | 0.84 | 1.33 | 0.000 | 0.723 | 0.325 | 7937 | tags=28%, list=13%, signal=32% |
| 773 | REGULATION\_OF\_IMMUNE\_SYSTEM\_PROCESS(GO:0002682) |  | 1541 | 0.83 | 1.33 | 0.000 | 0.723 | 0.325 | 7260 | tags=11%, list=12%, signal=13% |
| 774 | REGULATION\_OF\_SULFUR\_METABOLIC\_PROCESS(GO:0042762) |  | 20 | 0.85 | 1.33 | 0.000 | 0.723 | 0.325 | 5894 | tags=5%, list=10%, signal=6% |
| 775 | REGULATION\_OF\_MICROTUBULE\_POLYMERIZATION(GO:0031113) |  | 45 | 0.80 | 1.33 | 0.000 | 0.722 | 0.325 | 8112 | tags=18%, list=14%, signal=21% |
| 776 | RESPONSE\_TO\_STEROID\_HORMONE(GO:0048545) |  | 396 | 0.84 | 1.33 | 0.141 | 0.722 | 0.325 | 7217 | tags=13%, list=12%, signal=14% |
| 777 | RESPONSE\_TO\_AUDITORY\_STIMULUS(GO:0010996) |  | 23 | 0.85 | 1.33 | 0.000 | 0.721 | 0.325 | 7249 | tags=26%, list=12%, signal=30% |
| 778 | POSITIVE\_REGULATION\_OF\_INTERLEUKIN\_6\_PRODUCTION(GO:0032755) |  | 75 | 0.85 | 1.33 | 0.000 | 0.721 | 0.325 | 6955 | tags=19%, list=12%, signal=21% |
| 779 | PROTEIN\_LOCALIZATION\_TO\_ORGANELLE(GO:0033365) |  | 819 | 0.83 | 1.33 | 0.000 | 0.720 | 0.325 | 6933 | tags=5%, list=12%, signal=6% |
| 780 | NEGATIVE\_REGULATION\_OF\_BINDING(GO:0051100) |  | 158 | 0.84 | 1.33 | 0.000 | 0.720 | 0.325 | 6381 | tags=10%, list=11%, signal=11% |
| 781 | POSITIVE\_REGULATION\_OF\_JAK\_STAT\_CASCADE(GO:0046427) |  | 77 | 0.85 | 1.33 | 0.141 | 0.720 | 0.325 | 7170 | tags=18%, list=12%, signal=21% |
| 782 | CALCIUM\_CHANNEL\_ACTIVITY(GO:0005262) |  | 116 | 0.83 | 1.33 | 0.000 | 0.719 | 0.325 | 7881 | tags=26%, list=13%, signal=30% |
| 783 | PHOSPHATIDYLSERINE\_METABOLIC\_PROCESS(GO:0006658) |  | 30 | 0.87 | 1.33 | 0.000 | 0.719 | 0.325 | 5866 | tags=13%, list=10%, signal=15% |
| 784 | PRESYNAPSE\_ASSEMBLY(GO:0099054) |  | 19 | 0.85 | 1.33 | 0.000 | 0.718 | 0.325 | 7373 | tags=32%, list=12%, signal=36% |
| 785 | OXIDOREDUCTASE\_ACTIVITY(GO:0016491) |  | 755 | 0.83 | 1.33 | 0.000 | 0.718 | 0.325 | 7054 | tags=11%, list=12%, signal=12% |
| 786 | RESPONSE\_TO\_NUTRIENT\_LEVELS(GO:0031667) |  | 459 | 0.84 | 1.33 | 0.141 | 0.717 | 0.325 | 6731 | tags=13%, list=11%, signal=14% |
| 787 | FILAMENTOUS\_ACTIN(GO:0031941) |  | 31 | 0.85 | 1.33 | 0.141 | 0.716 | 0.325 | 6561 | tags=16%, list=11%, signal=18% |
| 788 | MUSCLE\_SYSTEM\_PROCESS(GO:0003012) |  | 447 | 0.82 | 1.33 | 0.000 | 0.716 | 0.325 | 7979 | tags=23%, list=13%, signal=26% |
| 789 | POSITIVE\_REGULATION\_OF\_CELL\_MATRIX\_ADHESION(GO:0001954) |  | 46 | 0.87 | 1.33 | 0.000 | 0.715 | 0.325 | 6027 | tags=11%, list=10%, signal=12% |
| 790 | FOREBRAIN\_NEURON\_DIFFERENTIATION(GO:0021879) |  | 56 | 0.87 | 1.33 | 0.000 | 0.715 | 0.325 | 6473 | tags=21%, list=11%, signal=24% |
| 791 | EAR\_MORPHOGENESIS(GO:0042471) |  | 120 | 0.85 | 1.33 | 0.000 | 0.714 | 0.325 | 6473 | tags=22%, list=11%, signal=24% |
| 792 | CELL\_MOTILITY(GO:0048870) |  | 1579 | 0.85 | 1.33 | 0.141 | 0.714 | 0.325 | 7169 | tags=14%, list=12%, signal=16% |
| 793 | LOCALIZATION\_OF\_CELL(GO:0051674) |  | 1579 | 0.85 | 1.33 | 0.141 | 0.713 | 0.325 | 7169 | tags=14%, list=12%, signal=16% |
| 794 | CALCIUM\_ION\_TRANSMEMBRANE\_IMPORT\_INTO\_CYTOSOL(GO:0097553) |  | 129 | 0.85 | 1.33 | 0.000 | 0.712 | 0.325 | 6662 | tags=15%, list=11%, signal=17% |
| 795 | PEPTIDOGLYCAN\_MURALYTIC\_ACTIVITY(GO:0061783) |  | 19 | 0.81 | 1.33 | 0.000 | 0.712 | 0.325 | 10529 | tags=21%, list=18%, signal=26% |
| 796 | SULFUR\_COMPOUND\_BINDING(GO:1901681) |  | 229 | 0.86 | 1.33 | 0.141 | 0.711 | 0.325 | 7008 | tags=22%, list=12%, signal=25% |
| 797 | NEGATIVE\_REGULATION\_OF\_MOLECULAR\_FUNCTION(GO:0044092) |  | 1116 | 0.83 | 1.33 | 0.000 | 0.711 | 0.325 | 6695 | tags=10%, list=11%, signal=12% |
| 798 | CELLULAR\_RESPONSE\_TO\_FATTY\_ACID(GO:0071398) |  | 56 | 0.86 | 1.33 | 0.000 | 0.710 | 0.325 | 6178 | tags=13%, list=10%, signal=14% |
| 799 | PROTEIN\_SERINE\_THREONINE\_TYROSINE\_KINASE\_ACTIVITY(GO:0004712) |  | 41 | 0.82 | 1.33 | 0.000 | 0.709 | 0.325 | 7426 | tags=12%, list=13%, signal=14% |
| 800 | PLATELET\_ALPHA\_GRANULE\_MEMBRANE(GO:0031092) |  | 16 | 0.84 | 1.33 | 0.000 | 0.709 | 0.325 | 7921 | tags=38%, list=13%, signal=43% |
| 801 | REGULATION\_OF\_SUPEROXIDE\_ANION\_GENERATION(GO:0032928) |  | 18 | 0.90 | 1.33 | 0.000 | 0.709 | 0.325 | 5825 | tags=6%, list=10%, signal=6% |
| 802 | HINDBRAIN\_DEVELOPMENT(GO:0030902) |  | 151 | 0.83 | 1.33 | 0.000 | 0.708 | 0.325 | 8261 | tags=28%, list=14%, signal=32% |
| 803 | ENDOCRINE\_SYSTEM\_DEVELOPMENT(GO:0035270) |  | 128 | 0.85 | 1.33 | 0.000 | 0.708 | 0.325 | 6617 | tags=27%, list=11%, signal=31% |
| 804 | GLIAL\_CELL\_MIGRATION(GO:0008347) |  | 47 | 0.88 | 1.33 | 0.141 | 0.707 | 0.325 | 6328 | tags=21%, list=11%, signal=24% |
| 805 | REGULATION\_OF\_TOR\_SIGNALING(GO:0032006) |  | 92 | 0.80 | 1.33 | 0.000 | 0.706 | 0.325 | 6256 | tags=8%, list=11%, signal=8% |
| 806 | SKIN\_DEVELOPMENT(GO:0043588) |  | 411 | 0.86 | 1.33 | 0.141 | 0.706 | 0.325 | 6473 | tags=9%, list=11%, signal=9% |
| 807 | NEGATIVE\_REGULATION\_OF\_SYNAPTIC\_TRANSMISSION(GO:0050805) |  | 74 | 0.85 | 1.33 | 0.000 | 0.705 | 0.325 | 7426 | tags=27%, list=13%, signal=31% |
| 808 | PROTEIN\_KINASE\_COMPLEX(GO:1902911) |  | 97 | 0.80 | 1.33 | 0.000 | 0.705 | 0.325 | 6745 | tags=7%, list=11%, signal=8% |
| 809 | NEGATIVE\_REGULATION\_OF\_PEPTIDASE\_ACTIVITY(GO:0010466) |  | 262 | 0.84 | 1.33 | 0.141 | 0.704 | 0.325 | 7238 | tags=16%, list=12%, signal=18% |
| 810 | HIPPOCAMPUS\_DEVELOPMENT(GO:0021766) |  | 78 | 0.86 | 1.33 | 0.000 | 0.704 | 0.325 | 6381 | tags=15%, list=11%, signal=17% |
| 811 | CHEMOTAXIS(GO:0006935) |  | 582 | 0.85 | 1.33 | 0.141 | 0.703 | 0.325 | 6989 | tags=18%, list=12%, signal=20% |
| 812 | SEMI\_LUNAR\_VALVE\_DEVELOPMENT(GO:1905314) |  | 16 | 0.88 | 1.33 | 0.000 | 0.703 | 0.325 | 6155 | tags=25%, list=10%, signal=28% |
| 813 | TAXIS(GO:0042330) |  | 583 | 0.85 | 1.33 | 0.141 | 0.703 | 0.325 | 6989 | tags=18%, list=12%, signal=20% |
| 814 | CALCIUM\_ION\_TRANSMEMBRANE\_TRANSPORTER\_ACTIVITY(GO:0015085) |  | 137 | 0.83 | 1.33 | 0.000 | 0.703 | 0.325 | 7881 | tags=26%, list=13%, signal=29% |
| 815 | MACROMOLECULE\_CATABOLIC\_PROCESS(GO:0009057) |  | 1327 | 0.82 | 1.33 | 0.000 | 0.702 | 0.325 | 6696 | tags=5%, list=11%, signal=6% |
| 816 | POSITIVE\_REGULATION\_OF\_STAT\_CASCADE(GO:1904894) |  | 79 | 0.85 | 1.33 | 0.141 | 0.702 | 0.325 | 7170 | tags=18%, list=12%, signal=20% |
| 817 | CONNEXIN\_COMPLEX(GO:0005922) |  | 22 | 0.89 | 1.33 | 0.000 | 0.701 | 0.325 | 5959 | tags=32%, list=10%, signal=35% |
| 818 | POSITIVE\_REGULATION\_OF\_NUCLEOCYTOPLASMIC\_TRANSPORT(GO:0046824) |  | 56 | 0.86 | 1.33 | 0.141 | 0.701 | 0.325 | 6642 | tags=7%, list=11%, signal=8% |
| 819 | VOLTAGE\_GATED\_CALCIUM\_CHANNEL\_ACTIVITY(GO:0005245) |  | 41 | 0.87 | 1.33 | 0.000 | 0.701 | 0.325 | 6332 | tags=17%, list=11%, signal=19% |
| 820 | CARDIAC\_CONDUCTION(GO:0061337) |  | 143 | 0.85 | 1.33 | 0.141 | 0.700 | 0.325 | 6670 | tags=23%, list=11%, signal=26% |
| 821 | STRESS\_ACTIVATED\_MAPK\_CASCADE(GO:0051403) |  | 266 | 0.85 | 1.33 | 0.141 | 0.699 | 0.325 | 6797 | tags=9%, list=11%, signal=11% |
| 822 | CARDIAC\_SEPTUM\_DEVELOPMENT(GO:0003279) |  | 101 | 0.85 | 1.33 | 0.000 | 0.699 | 0.325 | 6527 | tags=17%, list=11%, signal=19% |
| 823 | REGULATION\_OF\_ACTIN\_FILAMENT\_BASED\_MOVEMENT(GO:1903115) |  | 45 | 0.85 | 1.33 | 0.000 | 0.698 | 0.325 | 7008 | tags=22%, list=12%, signal=25% |
| 824 | NEGATIVE\_REGULATION\_OF\_IMMUNE\_RESPONSE(GO:0050777) |  | 127 | 0.85 | 1.33 | 0.141 | 0.697 | 0.325 | 6685 | tags=12%, list=11%, signal=13% |
| 825 | RENAL\_SYSTEM\_DEVELOPMENT(GO:0072001) |  | 281 | 0.86 | 1.33 | 0.141 | 0.697 | 0.325 | 7426 | tags=25%, list=13%, signal=29% |
| 826 | PEPTIDYL\_CYSTEINE\_MODIFICATION(GO:0018198) |  | 29 | 0.86 | 1.33 | 0.000 | 0.697 | 0.325 | 6247 | tags=7%, list=11%, signal=8% |
| 827 | ADULT\_BEHAVIOR(GO:0030534) |  | 146 | 0.84 | 1.33 | 0.000 | 0.697 | 0.325 | 7298 | tags=24%, list=12%, signal=27% |
| 828 | PROTEIN\_CONTAINING\_COMPLEX\_BINDING(GO:0044877) |  | 907 | 0.84 | 1.33 | 0.141 | 0.696 | 0.325 | 6894 | tags=12%, list=12%, signal=13% |
| 829 | NODE\_OF\_RANVIER(GO:0033268) |  | 15 | 0.84 | 1.33 | 0.000 | 0.695 | 0.325 | 7932 | tags=40%, list=13%, signal=46% |
| 830 | AXOLEMMA(GO:0030673) |  | 18 | 0.88 | 1.33 | 0.000 | 0.695 | 0.325 | 6000 | tags=28%, list=10%, signal=31% |
| 831 | NEGATIVE\_REGULATION\_OF\_VIRAL\_GENOME\_REPLICATION(GO:0045071) |  | 53 | 0.85 | 1.33 | 0.141 | 0.695 | 0.325 | 7293 | tags=21%, list=12%, signal=24% |
| 832 | BLOOD\_VESSEL\_ENDOTHELIAL\_CELL\_MIGRATION(GO:0043534) |  | 167 | 0.84 | 1.33 | 0.000 | 0.695 | 0.325 | 7267 | tags=14%, list=12%, signal=16% |
| 833 | SECRETORY\_VESICLE(GO:0099503) |  | 940 | 0.82 | 1.33 | 0.000 | 0.694 | 0.325 | 7546 | tags=13%, list=13%, signal=15% |
| 834 | MESENCHYMAL\_CELL\_PROLIFERATION(GO:0010463) |  | 47 | 0.87 | 1.33 | 0.000 | 0.694 | 0.325 | 6918 | tags=30%, list=12%, signal=34% |
| 835 | REGULATION\_OF\_PROTEIN\_LOCALIZATION(GO:0032880) |  | 958 | 0.84 | 1.33 | 0.141 | 0.693 | 0.325 | 6702 | tags=9%, list=11%, signal=10% |
| 836 | WATER\_HOMEOSTASIS(GO:0030104) |  | 68 | 0.87 | 1.33 | 0.141 | 0.693 | 0.325 | 6374 | tags=13%, list=11%, signal=15% |
| 837 | REGULATION\_OF\_BLOOD\_CIRCULATION(GO:1903522) |  | 294 | 0.84 | 1.33 | 0.141 | 0.693 | 0.325 | 7015 | tags=22%, list=12%, signal=25% |
| 838 | CALCIUM\_CHANNEL\_COMPLEX(GO:0034704) |  | 68 | 0.84 | 1.33 | 0.000 | 0.692 | 0.325 | 6332 | tags=16%, list=11%, signal=18% |
| 839 | PHOTORECEPTOR\_CELL\_DEVELOPMENT(GO:0042461) |  | 42 | 0.86 | 1.33 | 0.141 | 0.692 | 0.325 | 5892 | tags=17%, list=10%, signal=18% |
| 840 | ACTIN\_FILAMENT(GO:0005884) |  | 91 | 0.83 | 1.33 | 0.141 | 0.692 | 0.325 | 6785 | tags=18%, list=11%, signal=20% |
| 841 | VASCULOGENESIS(GO:0001570) |  | 77 | 0.87 | 1.33 | 0.141 | 0.691 | 0.325 | 6527 | tags=23%, list=11%, signal=26% |
| 842 | ARTERY\_DEVELOPMENT(GO:0060840) |  | 93 | 0.83 | 1.33 | 0.000 | 0.692 | 0.325 | 7343 | tags=17%, list=12%, signal=20% |
| 843 | COA\_LIGASE\_ACTIVITY(GO:0016405) |  | 26 | 0.82 | 1.33 | 0.000 | 0.691 | 0.325 | 6782 | tags=12%, list=11%, signal=13% |
| 844 | REGULATION\_OF\_EXTRACELLULAR\_MATRIX\_DISASSEMBLY(GO:0010715) |  | 15 | 0.87 | 1.33 | 0.000 | 0.690 | 0.325 | 6229 | tags=20%, list=11%, signal=22% |
| 845 | RESPONSE\_TO\_EXTERNAL\_BIOTIC\_STIMULUS(GO:0043207) |  | 891 | 0.84 | 1.33 | 0.141 | 0.690 | 0.325 | 7265 | tags=11%, list=12%, signal=12% |
| 846 | CELL\_JUNCTION(GO:0030054) |  | 1238 | 0.85 | 1.33 | 0.141 | 0.689 | 0.325 | 6818 | tags=13%, list=12%, signal=14% |
| 847 | HOMOPHILIC\_CELL\_ADHESION\_VIA\_PLASMA\_MEMBRANE\_ADHESION\_MOLECULES(GO:0007156) |  | 157 | 0.87 | 1.33 | 0.141 | 0.689 | 0.325 | 6726 | tags=16%, list=11%, signal=18% |
| 848 | SENSORY\_PERCEPTION\_OF\_SOUND(GO:0007605) |  | 148 | 0.85 | 1.33 | 0.141 | 0.688 | 0.325 | 6957 | tags=18%, list=12%, signal=20% |
| 849 | MORPHOGENESIS\_OF\_A\_BRANCHING\_STRUCTURE(GO:0001763) |  | 196 | 0.86 | 1.33 | 0.141 | 0.688 | 0.325 | 6495 | tags=20%, list=11%, signal=23% |
| 850 | RESPONSE\_TO\_OTHER\_ORGANISM(GO:0051707) |  | 889 | 0.84 | 1.33 | 0.141 | 0.687 | 0.325 | 7265 | tags=11%, list=12%, signal=13% |
| 851 | REGULATION\_OF\_CELL\_AGING(GO:0090342) |  | 52 | 0.85 | 1.33 | 0.141 | 0.686 | 0.325 | 6685 | tags=12%, list=11%, signal=13% |
| 852 | REGULATION\_OF\_ANATOMICAL\_STRUCTURE\_MORPHOGENESIS(GO:0022603) |  | 1096 | 0.84 | 1.33 | 0.141 | 0.686 | 0.325 | 7169 | tags=15%, list=12%, signal=17% |
| 853 | EAR\_DEVELOPMENT(GO:0043583) |  | 215 | 0.84 | 1.33 | 0.000 | 0.685 | 0.325 | 7837 | tags=27%, list=13%, signal=30% |
| 854 | SPINAL\_CORD\_PATTERNING(GO:0021511) |  | 24 | 0.88 | 1.33 | 0.000 | 0.685 | 0.325 | 6058 | tags=33%, list=10%, signal=37% |
| 855 | C21\_STEROID\_HORMONE\_METABOLIC\_PROCESS(GO:0008207) |  | 36 | 0.87 | 1.33 | 0.000 | 0.684 | 0.325 | 5839 | tags=11%, list=10%, signal=12% |
| 856 | SKIN\_EPIDERMIS\_DEVELOPMENT(GO:0098773) |  | 89 | 0.85 | 1.33 | 0.000 | 0.684 | 0.325 | 6724 | tags=15%, list=11%, signal=16% |
| 857 | CALCIUM\_ION\_TRANSMEMBRANE\_TRANSPORT(GO:0070588) |  | 311 | 0.82 | 1.33 | 0.000 | 0.683 | 0.325 | 8046 | tags=20%, list=14%, signal=23% |
| 858 | ASTROCYTE\_DIFFERENTIATION(GO:0048708) |  | 68 | 0.86 | 1.33 | 0.141 | 0.683 | 0.325 | 6442 | tags=19%, list=11%, signal=21% |
| 859 | T\_HELPER\_1\_CELL\_DIFFERENTIATION(GO:0045063) |  | 17 | 0.84 | 1.33 | 0.000 | 0.682 | 0.325 | 6381 | tags=29%, list=11%, signal=33% |
| 860 | UROGENITAL\_SYSTEM\_DEVELOPMENT(GO:0001655) |  | 317 | 0.85 | 1.33 | 0.141 | 0.682 | 0.325 | 7426 | tags=25%, list=13%, signal=28% |
| 861 | NEGATIVE\_REGULATION\_OF\_INTRACELLULAR\_SIGNAL\_TRANSDUCTION(GO:1902532) |  | 527 | 0.84 | 1.33 | 0.141 | 0.682 | 0.325 | 6695 | tags=9%, list=11%, signal=10% |
| 862 | INTERLEUKIN\_1\_MEDIATED\_SIGNALING\_PATHWAY(GO:0070498) |  | 101 | 0.81 | 1.33 | 0.000 | 0.681 | 0.325 | 5846 | tags=1%, list=10%, signal=1% |
| 863 | REGULATION\_OF\_TRANSCRIPTION\_INVOLVED\_IN\_CELL\_FATE\_COMMITMENT(GO:0060850) |  | 17 | 0.86 | 1.33 | 0.000 | 0.681 | 0.325 | 7668 | tags=41%, list=13%, signal=47% |
| 864 | GROWTH\_FACTOR\_BINDING(GO:0019838) |  | 130 | 0.86 | 1.33 | 0.141 | 0.681 | 0.325 | 6731 | tags=12%, list=11%, signal=14% |
| 865 | PERICARDIUM\_DEVELOPMENT(GO:0060039) |  | 20 | 0.83 | 1.33 | 0.000 | 0.681 | 0.325 | 8261 | tags=40%, list=14%, signal=46% |
| 866 | COMPACT\_MYELIN(GO:0043218) |  | 15 | 0.89 | 1.33 | 0.000 | 0.680 | 0.325 | 6011 | tags=13%, list=10%, signal=15% |
| 867 | REGULATION\_OF\_MEMBRANE\_POTENTIAL(GO:0042391) |  | 411 | 0.83 | 1.33 | 0.000 | 0.680 | 0.325 | 7963 | tags=25%, list=13%, signal=29% |
| 868 | HAIR\_CYCLE(GO:0042633) |  | 111 | 0.84 | 1.33 | 0.141 | 0.680 | 0.325 | 6779 | tags=14%, list=11%, signal=15% |
| 869 | MOLTING\_CYCLE(GO:0042303) |  | 111 | 0.84 | 1.33 | 0.141 | 0.679 | 0.325 | 6779 | tags=14%, list=11%, signal=15% |
| 870 | OLIGODENDROCYTE\_DIFFERENTIATION(GO:0048709) |  | 86 | 0.88 | 1.33 | 0.000 | 0.678 | 0.325 | 6442 | tags=20%, list=11%, signal=22% |
| 871 | CALCIUM\_ION\_TRANSPORT\_INTO\_CYTOSOL(GO:0060402) |  | 147 | 0.84 | 1.33 | 0.000 | 0.678 | 0.325 | 6706 | tags=16%, list=11%, signal=18% |
| 872 | LEADING\_EDGE\_MEMBRANE(GO:0031256) |  | 151 | 0.82 | 1.33 | 0.000 | 0.678 | 0.325 | 6455 | tags=9%, list=11%, signal=10% |
| 873 | REGULATION\_OF\_EXTRACELLULAR\_MATRIX\_ORGANIZATION(GO:1903053) |  | 33 | 0.82 | 1.33 | 0.000 | 0.678 | 0.325 | 7640 | tags=18%, list=13%, signal=21% |
| 874 | POSITIVE\_REGULATION\_OF\_LIPID\_CATABOLIC\_PROCESS(GO:0050996) |  | 22 | 0.86 | 1.33 | 0.000 | 0.677 | 0.325 | 5904 | tags=18%, list=10%, signal=20% |
| 875 | INOSITOL\_PHOSPHATE\_METABOLIC\_PROCESS(GO:0043647) |  | 67 | 0.81 | 1.33 | 0.000 | 0.677 | 0.325 | 6991 | tags=15%, list=12%, signal=17% |
| 876 | POSITIVE\_REGULATION\_OF\_MACROPHAGE\_DERIVED\_FOAM\_CELL\_DIFFERENTIATION(GO:0010744) |  | 16 | 0.85 | 1.33 | 0.000 | 0.676 | 0.325 | 6005 | tags=25%, list=10%, signal=28% |
| 877 | LYMPHANGIOGENESIS(GO:0001946) |  | 15 | 0.84 | 1.33 | 0.000 | 0.676 | 0.325 | 7260 | tags=47%, list=12%, signal=53% |
| 878 | BRANCHING\_MORPHOGENESIS\_OF\_AN\_EPITHELIAL\_TUBE(GO:0048754) |  | 152 | 0.87 | 1.33 | 0.141 | 0.676 | 0.325 | 6473 | tags=18%, list=11%, signal=20% |
| 879 | CELLULAR\_CALCIUM\_ION\_HOMEOSTASIS(GO:0006874) |  | 413 | 0.82 | 1.33 | 0.000 | 0.675 | 0.325 | 7544 | tags=19%, list=13%, signal=21% |
| 880 | RESPONSE\_TO\_EXTRACELLULAR\_STIMULUS(GO:0009991) |  | 488 | 0.83 | 1.33 | 0.141 | 0.675 | 0.325 | 7172 | tags=14%, list=12%, signal=16% |
| 881 | VOLTAGE\_GATED\_SODIUM\_CHANNEL\_ACTIVITY(GO:0005248) |  | 23 | 0.87 | 1.33 | 0.000 | 0.674 | 0.325 | 6844 | tags=43%, list=12%, signal=49% |
| 882 | NEGATIVE\_REGULATION\_OF\_ENDOPEPTIDASE\_ACTIVITY(GO:0010951) |  | 251 | 0.84 | 1.33 | 0.141 | 0.674 | 0.325 | 7238 | tags=15%, list=12%, signal=17% |
| 883 | PERIPHERAL\_NERVOUS\_SYSTEM\_DEVELOPMENT(GO:0007422) |  | 72 | 0.86 | 1.33 | 0.000 | 0.673 | 0.325 | 7046 | tags=22%, list=12%, signal=25% |
| 884 | CARDIAC\_SEPTUM\_MORPHOGENESIS(GO:0060411) |  | 68 | 0.86 | 1.33 | 0.000 | 0.673 | 0.325 | 6527 | tags=21%, list=11%, signal=23% |
| 885 | RESPONSE\_TO\_UV(GO:0009411) |  | 136 | 0.79 | 1.33 | 0.000 | 0.672 | 0.325 | 8427 | tags=13%, list=14%, signal=15% |
| 886 | POSITIVE\_REGULATION\_OF\_SMOOTH\_MUSCLE\_CELL\_PROLIFERATION(GO:0048661) |  | 88 | 0.85 | 1.33 | 0.141 | 0.672 | 0.325 | 6514 | tags=17%, list=11%, signal=19% |
| 887 | REGULATION\_OF\_TRANSPOSITION(GO:0010528) |  | 19 | 0.86 | 1.33 | 0.141 | 0.671 | 0.325 | 6865 | tags=32%, list=12%, signal=36% |
| 888 | NEGATIVE\_REGULATION\_OF\_TRANSPOSITION(GO:0010529) |  | 19 | 0.86 | 1.33 | 0.141 | 0.671 | 0.325 | 6865 | tags=32%, list=12%, signal=36% |
| 889 | VASODILATION(GO:0042311) |  | 33 | 0.87 | 1.33 | 0.000 | 0.670 | 0.325 | 6015 | tags=15%, list=10%, signal=17% |
| 890 | TOR\_SIGNALING(GO:0031929) |  | 108 | 0.81 | 1.33 | 0.000 | 0.670 | 0.325 | 6359 | tags=8%, list=11%, signal=9% |
| 891 | CYTOSKELETAL\_PROTEIN\_BINDING(GO:0008092) |  | 883 | 0.84 | 1.33 | 0.141 | 0.669 | 0.325 | 7256 | tags=14%, list=12%, signal=16% |
| 892 | NEGATIVE\_REGULATION\_OF\_APOPTOTIC\_PROCESS(GO:0043066) |  | 874 | 0.84 | 1.33 | 0.141 | 0.669 | 0.325 | 6871 | tags=12%, list=12%, signal=14% |
| 893 | ADULT\_LOCOMOTORY\_BEHAVIOR(GO:0008344) |  | 83 | 0.86 | 1.33 | 0.000 | 0.669 | 0.325 | 6240 | tags=23%, list=11%, signal=26% |
| 894 | INTERLEUKIN\_6\_SECRETION(GO:0072604) |  | 34 | 0.83 | 1.33 | 0.141 | 0.669 | 0.325 | 7256 | tags=15%, list=12%, signal=17% |
| 895 | MICROVILLUS\_ORGANIZATION(GO:0032528) |  | 24 | 0.85 | 1.33 | 0.000 | 0.669 | 0.325 | 6763 | tags=21%, list=11%, signal=24% |
| 896 | SIGNAL\_TRANSDUCER\_ACTIVITY(GO:0004871) |  | 1696 | 0.84 | 1.33 | 0.141 | 0.668 | 0.325 | 7556 | tags=16%, list=13%, signal=17% |
| 897 | REGULATED\_EXOCYTOSIS(GO:0045055) |  | 736 | 0.82 | 1.33 | 0.000 | 0.668 | 0.325 | 7515 | tags=10%, list=13%, signal=12% |
| 898 | RESPONSE\_TO\_STIMULUS\_INVOLVED\_IN\_REGULATION\_OF\_MUSCLE\_ADAPTATION(GO:0014874) |  | 16 | 0.87 | 1.33 | 0.000 | 0.668 | 0.325 | 6314 | tags=13%, list=11%, signal=14% |
| 899 | CELLULAR\_RESPONSE\_TO\_GROWTH\_FACTOR\_STIMULUS(GO:0071363) |  | 672 | 0.83 | 1.33 | 0.000 | 0.667 | 0.325 | 7396 | tags=17%, list=13%, signal=19% |
| 900 | REGULATION\_OF\_APOPTOTIC\_SIGNALING\_PATHWAY(GO:2001233) |  | 388 | 0.85 | 1.33 | 0.141 | 0.667 | 0.325 | 6662 | tags=6%, list=11%, signal=7% |
| 901 | NEGATIVE\_REGULATION\_OF\_PROGRAMMED\_CELL\_DEATH(GO:0043069) |  | 889 | 0.84 | 1.33 | 0.141 | 0.666 | 0.325 | 6957 | tags=13%, list=12%, signal=14% |
| 902 | POSITIVE\_REGULATION\_OF\_DNA\_BINDING(GO:0043388) |  | 56 | 0.82 | 1.33 | 0.000 | 0.666 | 0.325 | 7313 | tags=23%, list=12%, signal=26% |
| 903 | REGULATION\_OF\_TRANSMEMBRANE\_TRANSPORTER\_ACTIVITY(GO:0022898) |  | 234 | 0.84 | 1.33 | 0.000 | 0.665 | 0.325 | 7979 | tags=26%, list=13%, signal=31% |
| 904 | GLUCURONOSYLTRANSFERASE\_ACTIVITY(GO:0015020) |  | 35 | 0.82 | 1.33 | 0.000 | 0.665 | 0.325 | 6557 | tags=17%, list=11%, signal=19% |
| 905 | POSITIVE\_REGULATION\_OF\_BLOOD\_VESSEL\_DIAMETER(GO:0097755) |  | 59 | 0.85 | 1.33 | 0.000 | 0.664 | 0.325 | 7543 | tags=24%, list=13%, signal=27% |
| 906 | MITOCHONDRIAL\_GENOME\_MAINTENANCE(GO:0000002) |  | 28 | 0.79 | 1.33 | 0.000 | 0.664 | 0.325 | 5916 | tags=4%, list=10%, signal=4% |
| 907 | PHENOL\_CONTAINING\_COMPOUND\_METABOLIC\_PROCESS(GO:0018958) |  | 95 | 0.85 | 1.33 | 0.141 | 0.664 | 0.325 | 7393 | tags=23%, list=13%, signal=26% |
| 908 | MESENCHYMAL\_CELL\_DEVELOPMENT(GO:0014031) |  | 65 | 0.86 | 1.33 | 0.141 | 0.663 | 0.325 | 6871 | tags=25%, list=12%, signal=28% |
| 909 | PROTEIN\_LIPID\_COMPLEX\_REMODELING(GO:0034368) |  | 28 | 0.85 | 1.33 | 0.000 | 0.664 | 0.325 | 6362 | tags=36%, list=11%, signal=40% |
| 910 | PLASMA\_LIPOPROTEIN\_PARTICLE\_REMODELING(GO:0034369) |  | 28 | 0.85 | 1.33 | 0.000 | 0.663 | 0.325 | 6362 | tags=36%, list=11%, signal=40% |
| 911 | PROTEIN\_CONTAINING\_COMPLEX\_REMODELING(GO:0034367) |  | 28 | 0.85 | 1.33 | 0.000 | 0.662 | 0.325 | 6362 | tags=36%, list=11%, signal=40% |
| 912 | HAIR\_FOLLICLE\_DEVELOPMENT(GO:0001942) |  | 87 | 0.85 | 1.33 | 0.000 | 0.662 | 0.325 | 6724 | tags=15%, list=11%, signal=17% |
| 913 | REGULATION\_OF\_TRANSPORTER\_ACTIVITY(GO:0032409) |  | 248 | 0.84 | 1.33 | 0.000 | 0.662 | 0.325 | 7979 | tags=26%, list=13%, signal=30% |
| 914 | REGULATION\_OF\_NOREPINEPHRINE\_SECRETION(GO:0014061) |  | 17 | 0.86 | 1.33 | 0.000 | 0.661 | 0.325 | 5955 | tags=18%, list=10%, signal=20% |
| 915 | CELL\_CELL\_ADHESION\_VIA\_PLASMA\_MEMBRANE\_ADHESION\_MOLECULES(GO:0098742) |  | 239 | 0.86 | 1.33 | 0.141 | 0.661 | 0.325 | 6726 | tags=18%, list=11%, signal=21% |
| 916 | REGULATION\_OF\_CELLULAR\_LOCALIZATION(GO:0060341) |  | 812 | 0.84 | 1.33 | 0.141 | 0.661 | 0.325 | 6809 | tags=9%, list=12%, signal=10% |
| 917 | CELL\_FATE\_COMMITMENT(GO:0045165) |  | 256 | 0.85 | 1.33 | 0.141 | 0.660 | 0.325 | 6871 | tags=26%, list=12%, signal=29% |
| 918 | ASTROCYTE\_DEVELOPMENT(GO:0014002) |  | 31 | 0.88 | 1.33 | 0.000 | 0.660 | 0.325 | 6303 | tags=16%, list=11%, signal=18% |
| 919 | VENTRAL\_SPINAL\_CORD\_INTERNEURON\_DIFFERENTIATION(GO:0021514) |  | 17 | 0.86 | 1.33 | 0.000 | 0.659 | 0.325 | 7006 | tags=41%, list=12%, signal=47% |
| 920 | ANTERIOR\_POSTERIOR\_PATTERN\_SPECIFICATION(GO:0009952) |  | 204 | 0.85 | 1.33 | 0.000 | 0.659 | 0.325 | 6871 | tags=28%, list=12%, signal=32% |
| 921 | NEGATIVE\_REGULATION\_OF\_PROTEOLYSIS(GO:0045861) |  | 354 | 0.83 | 1.33 | 0.141 | 0.659 | 0.325 | 7238 | tags=14%, list=12%, signal=16% |
| 922 | PURINE\_RIBONUCLEOSIDE\_BISPHOSPHATE\_METABOLIC\_PROCESS(GO:0034035) |  | 25 | 0.85 | 1.33 | 0.000 | 0.658 | 0.325 | 7287 | tags=20%, list=12%, signal=23% |
| 923 | 3'\_PHOSPHOADENOSINE\_5'\_PHOSPHOSULFATE\_METABOLIC\_PROCESS(GO:0050427) |  | 25 | 0.85 | 1.33 | 0.000 | 0.658 | 0.325 | 7287 | tags=20%, list=12%, signal=23% |
| 924 | LOCOMOTION(GO:0040011) |  | 1815 | 0.84 | 1.33 | 0.141 | 0.657 | 0.325 | 7169 | tags=15%, list=12%, signal=16% |
| 925 | MULTICELLULAR\_ORGANISMAL\_HOMEOSTASIS(GO:0048871) |  | 330 | 0.84 | 1.33 | 0.141 | 0.657 | 0.325 | 7504 | tags=18%, list=13%, signal=21% |
| 926 | PROTEIN\_SELF\_ASSOCIATION(GO:0043621) |  | 56 | 0.81 | 1.33 | 0.000 | 0.657 | 0.325 | 7721 | tags=14%, list=13%, signal=16% |
| 927 | LONG\_CHAIN\_FATTY\_ACID\_TRANSPORT(GO:0015909) |  | 65 | 0.85 | 1.33 | 0.141 | 0.656 | 0.325 | 5866 | tags=9%, list=10%, signal=10% |
| 928 | MULTICELLULAR\_ORGANISMAL\_WATER\_HOMEOSTASIS(GO:0050891) |  | 63 | 0.87 | 1.33 | 0.141 | 0.656 | 0.325 | 6374 | tags=13%, list=11%, signal=14% |
| 929 | NEURAL\_CREST\_CELL\_DEVELOPMENT(GO:0014032) |  | 61 | 0.86 | 1.33 | 0.141 | 0.656 | 0.325 | 6871 | tags=26%, list=12%, signal=30% |
| 930 | NEURAL\_CREST\_CELL\_MIGRATION(GO:0001755) |  | 37 | 0.88 | 1.33 | 0.000 | 0.655 | 0.325 | 6381 | tags=30%, list=11%, signal=33% |
| 931 | CRANIAL\_NERVE\_DEVELOPMENT(GO:0021545) |  | 50 | 0.87 | 1.33 | 0.000 | 0.655 | 0.325 | 6733 | tags=36%, list=11%, signal=41% |
| 932 | ATRIAL\_CARDIAC\_MUSCLE\_CELL\_TO\_AV\_NODE\_CELL\_COMMUNICATION(GO:0086066) |  | 20 | 0.88 | 1.33 | 0.141 | 0.655 | 0.325 | 6569 | tags=45%, list=11%, signal=51% |
| 933 | ATRIAL\_CARDIAC\_MUSCLE\_CELL\_TO\_AV\_NODE\_CELL\_SIGNALING(GO:0086026) |  | 20 | 0.88 | 1.33 | 0.141 | 0.654 | 0.325 | 6569 | tags=45%, list=11%, signal=51% |
| 934 | ATRIAL\_CARDIAC\_MUSCLE\_CELL\_ACTION\_POTENTIAL(GO:0086014) |  | 20 | 0.88 | 1.33 | 0.141 | 0.654 | 0.325 | 6569 | tags=45%, list=11%, signal=51% |
| 935 | LIMBIC\_SYSTEM\_DEVELOPMENT(GO:0021761) |  | 105 | 0.85 | 1.33 | 0.000 | 0.653 | 0.325 | 6381 | tags=18%, list=11%, signal=20% |
| 936 | POSITIVE\_REGULATION\_OF\_NEUROTRANSMITTER\_TRANSPORT(GO:0051590) |  | 29 | 0.84 | 1.33 | 0.000 | 0.653 | 0.325 | 7298 | tags=28%, list=12%, signal=31% |
| 937 | SYNTAXIN\_1\_BINDING(GO:0017075) |  | 20 | 0.81 | 1.33 | 0.000 | 0.652 | 0.325 | 7426 | tags=25%, list=13%, signal=29% |
| 938 | NEGATIVE\_REGULATION\_OF\_HYDROLASE\_ACTIVITY(GO:0051346) |  | 449 | 0.83 | 1.33 | 0.141 | 0.652 | 0.325 | 7546 | tags=16%, list=13%, signal=18% |
| 939 | REGULATION\_OF\_DNA\_BINDING(GO:0051101) |  | 115 | 0.84 | 1.33 | 0.000 | 0.651 | 0.325 | 7313 | tags=17%, list=12%, signal=20% |
| 940 | STEREOCILIUM\_BUNDLE(GO:0032421) |  | 52 | 0.86 | 1.33 | 0.141 | 0.651 | 0.325 | 6502 | tags=19%, list=11%, signal=22% |
| 941 | PLASMA\_MEMBRANE\_ORGANIZATION(GO:0007009) |  | 77 | 0.85 | 1.33 | 0.141 | 0.651 | 0.325 | 5969 | tags=12%, list=10%, signal=13% |
| 942 | REGULATION\_OF\_CELLULAR\_RESPONSE\_TO\_VASCULAR\_ENDOTHELIAL\_GROWTH\_FACTOR\_STIMULUS(GO:1902547) |  | 19 | 0.88 | 1.33 | 0.000 | 0.651 | 0.325 | 5929 | tags=21%, list=10%, signal=23% |
| 943 | CELL\_FATE\_SPECIFICATION(GO:0001708) |  | 84 | 0.87 | 1.33 | 0.141 | 0.650 | 0.325 | 6831 | tags=35%, list=12%, signal=39% |
| 944 | RESPONSE\_TO\_GROWTH\_FACTOR(GO:0070848) |  | 697 | 0.83 | 1.33 | 0.000 | 0.650 | 0.325 | 7396 | tags=17%, list=13%, signal=19% |
| 945 | RESPONSE\_TO\_PROSTAGLANDIN(GO:0034694) |  | 33 | 0.87 | 1.33 | 0.000 | 0.650 | 0.325 | 6527 | tags=21%, list=11%, signal=24% |
| 946 | NEGATIVE\_REGULATION\_OF\_PHOSPHATE\_METABOLIC\_PROCESS(GO:0045936) |  | 604 | 0.83 | 1.33 | 0.141 | 0.649 | 0.325 | 6531 | tags=8%, list=11%, signal=9% |
| 947 | NEGATIVE\_REGULATION\_OF\_PHOSPHORUS\_METABOLIC\_PROCESS(GO:0010563) |  | 605 | 0.83 | 1.33 | 0.141 | 0.649 | 0.325 | 6531 | tags=8%, list=11%, signal=9% |
| 948 | RELEASE\_OF\_CYTOCHROME\_C\_FROM\_MITOCHONDRIA(GO:0001836) |  | 57 | 0.86 | 1.33 | 0.141 | 0.649 | 0.325 | 5834 | tags=5%, list=10%, signal=6% |
| 949 | POSITIVE\_REGULATION\_OF\_MITOTIC\_CELL\_CYCLE\_PHASE\_TRANSITION(GO:1901992) |  | 68 | 0.79 | 1.33 | 0.000 | 0.649 | 0.325 | 5852 | tags=6%, list=10%, signal=7% |
| 950 | REGULATION\_OF\_INFLAMMATORY\_RESPONSE\_TO\_ANTIGENIC\_STIMULUS(GO:0002861) |  | 21 | 0.90 | 1.33 | 0.000 | 0.648 | 0.325 | 5834 | tags=10%, list=10%, signal=11% |
| 951 | JAK\_STAT\_CASCADE(GO:0007259) |  | 173 | 0.82 | 1.33 | 0.141 | 0.648 | 0.325 | 7449 | tags=19%, list=13%, signal=22% |
| 952 | CELLULAR\_RESPONSE\_TO\_ORGANIC\_SUBSTANCE(GO:0071310) |  | 2623 | 0.84 | 1.33 | 0.141 | 0.647 | 0.325 | 6628 | tags=11%, list=11%, signal=12% |
| 953 | REGULATION\_OF\_CATION\_TRANSMEMBRANE\_TRANSPORT(GO:1904062) |  | 299 | 0.83 | 1.33 | 0.000 | 0.647 | 0.325 | 7979 | tags=24%, list=13%, signal=27% |
| 954 | INTRACELLULAR\_CHLORIDE\_CHANNEL\_ACTIVITY(GO:0061778) |  | 16 | 0.88 | 1.33 | 0.000 | 0.647 | 0.325 | 5969 | tags=19%, list=10%, signal=21% |
| 955 | INTRACELLULAR\_CALCIUM\_ACTIVATED\_CHLORIDE\_CHANNEL\_ACTIVITY(GO:0005229) |  | 16 | 0.88 | 1.33 | 0.000 | 0.646 | 0.325 | 5969 | tags=19%, list=10%, signal=21% |
| 956 | NUCLEAR\_OUTER\_MEMBRANE(GO:0005640) |  | 25 | 0.85 | 1.33 | 0.000 | 0.646 | 0.325 | 7184 | tags=20%, list=12%, signal=23% |
| 957 | STEREOCILIUM(GO:0032420) |  | 44 | 0.86 | 1.33 | 0.141 | 0.645 | 0.325 | 6502 | tags=20%, list=11%, signal=23% |
| 958 | MOLTING\_CYCLE\_PROCESS(GO:0022404) |  | 89 | 0.85 | 1.33 | 0.000 | 0.645 | 0.325 | 6724 | tags=15%, list=11%, signal=16% |
| 959 | HAIR\_CYCLE\_PROCESS(GO:0022405) |  | 89 | 0.85 | 1.33 | 0.000 | 0.644 | 0.325 | 6724 | tags=15%, list=11%, signal=16% |
| 960 | MESENCHYMAL\_CELL\_DIFFERENTIATION(GO:0048762) |  | 193 | 0.83 | 1.33 | 0.000 | 0.644 | 0.325 | 7360 | tags=21%, list=12%, signal=24% |
| 961 | SPINAL\_CORD\_DEVELOPMENT(GO:0021510) |  | 103 | 0.86 | 1.33 | 0.000 | 0.643 | 0.325 | 6209 | tags=26%, list=10%, signal=29% |
| 962 | SENSORY\_PERCEPTION\_OF\_MECHANICAL\_STIMULUS(GO:0050954) |  | 164 | 0.85 | 1.33 | 0.141 | 0.643 | 0.325 | 6957 | tags=20%, list=12%, signal=23% |
| 963 | MESENCHYME\_DEVELOPMENT(GO:0060485) |  | 246 | 0.83 | 1.32 | 0.000 | 0.643 | 0.325 | 7360 | tags=24%, list=12%, signal=27% |
| 964 | CELL\_JUNCTION\_ASSEMBLY(GO:0034329) |  | 205 | 0.85 | 1.32 | 0.141 | 0.643 | 0.325 | 6495 | tags=8%, list=11%, signal=9% |
| 965 | CATECHOLAMINE\_METABOLIC\_PROCESS(GO:0006584) |  | 52 | 0.86 | 1.32 | 0.141 | 0.643 | 0.325 | 6441 | tags=13%, list=11%, signal=15% |
| 966 | CATECHOL\_CONTAINING\_COMPOUND\_METABOLIC\_PROCESS(GO:0009712) |  | 52 | 0.86 | 1.32 | 0.141 | 0.642 | 0.325 | 6441 | tags=13%, list=11%, signal=15% |
| 967 | METANEPHROS\_DEVELOPMENT(GO:0001656) |  | 89 | 0.86 | 1.32 | 0.141 | 0.642 | 0.325 | 7382 | tags=24%, list=12%, signal=27% |
| 968 | REGULATION\_OF\_CORTICOSTEROID\_HORMONE\_SECRETION(GO:2000846) |  | 16 | 0.82 | 1.32 | 0.000 | 0.641 | 0.325 | 8485 | tags=31%, list=14%, signal=36% |
| 969 | REGULATION\_OF\_ION\_TRANSMEMBRANE\_TRANSPORTER\_ACTIVITY(GO:0032412) |  | 228 | 0.83 | 1.32 | 0.000 | 0.641 | 0.325 | 7979 | tags=26%, list=13%, signal=30% |
| 970 | REGULATION\_OF\_MULTICELLULAR\_ORGANISM\_GROWTH(GO:0040014) |  | 72 | 0.86 | 1.32 | 0.141 | 0.641 | 0.325 | 5936 | tags=8%, list=10%, signal=9% |
| 971 | STEM\_CELL\_DEVELOPMENT(GO:0048864) |  | 64 | 0.86 | 1.32 | 0.141 | 0.640 | 0.325 | 6871 | tags=25%, list=12%, signal=28% |
| 972 | RNA\_POLYMERASE\_II\_ACTIVATING\_TRANSCRIPTION\_FACTOR\_BINDING(GO:0001102) |  | 39 | 0.81 | 1.32 | 0.000 | 0.641 | 0.325 | 7356 | tags=18%, list=12%, signal=20% |
| 973 | ANION\_HOMEOSTASIS(GO:0055081) |  | 44 | 0.82 | 1.32 | 0.000 | 0.640 | 0.325 | 8109 | tags=23%, list=14%, signal=26% |
| 974 | AGING(GO:0007568) |  | 306 | 0.83 | 1.32 | 0.000 | 0.640 | 0.325 | 6731 | tags=11%, list=11%, signal=12% |
| 975 | STAT\_CASCADE(GO:0097696) |  | 175 | 0.82 | 1.32 | 0.141 | 0.640 | 0.325 | 7449 | tags=19%, list=13%, signal=22% |
| 976 | EPITHELIUM\_DEVELOPMENT(GO:0060429) |  | 1261 | 0.85 | 1.32 | 0.141 | 0.640 | 0.325 | 6779 | tags=13%, list=11%, signal=14% |
| 977 | OUTFLOW\_TRACT\_SEPTUM\_MORPHOGENESIS(GO:0003148) |  | 25 | 0.86 | 1.32 | 0.000 | 0.639 | 0.325 | 7343 | tags=32%, list=12%, signal=37% |
| 978 | REGULATION\_OF\_CALCIUM\_ION\_TRANSPORT(GO:0051924) |  | 233 | 0.84 | 1.32 | 0.000 | 0.639 | 0.325 | 6706 | tags=16%, list=11%, signal=18% |
| 979 | REGULATION\_OF\_HAIR\_FOLLICLE\_DEVELOPMENT(GO:0051797) |  | 21 | 0.88 | 1.32 | 0.000 | 0.638 | 0.325 | 6473 | tags=14%, list=11%, signal=16% |
| 980 | MYOSIN\_BINDING(GO:0017022) |  | 60 | 0.85 | 1.32 | 0.141 | 0.638 | 0.325 | 7176 | tags=23%, list=12%, signal=27% |
| 981 | RESPONSE\_TO\_TOXIC\_SUBSTANCE(GO:0009636) |  | 499 | 0.84 | 1.32 | 0.141 | 0.638 | 0.325 | 6560 | tags=11%, list=11%, signal=12% |
| 982 | RESPONSE\_TO\_MUSCLE\_STRETCH(GO:0035994) |  | 19 | 0.85 | 1.32 | 0.141 | 0.637 | 0.325 | 7937 | tags=32%, list=13%, signal=36% |
| 983 | CENTRAL\_NERVOUS\_SYSTEM\_DEVELOPMENT(GO:0007417) |  | 933 | 0.84 | 1.32 | 0.141 | 0.637 | 0.325 | 6841 | tags=17%, list=12%, signal=19% |
| 984 | ENDOCARDIAL\_CUSHION\_MORPHOGENESIS(GO:0003203) |  | 29 | 0.84 | 1.32 | 0.000 | 0.637 | 0.325 | 8177 | tags=41%, list=14%, signal=48% |
| 985 | VOLTAGE\_GATED\_CALCIUM\_CHANNEL\_COMPLEX(GO:0005891) |  | 45 | 0.86 | 1.32 | 0.000 | 0.636 | 0.325 | 6332 | tags=18%, list=11%, signal=20% |
| 986 | NEGATIVE\_REGULATION\_OF\_PROTEIN\_MODIFICATION\_PROCESS(GO:0031400) |  | 597 | 0.83 | 1.32 | 0.000 | 0.636 | 0.325 | 6695 | tags=8%, list=11%, signal=9% |
| 987 | VENOUS\_BLOOD\_VESSEL\_DEVELOPMENT(GO:0060841) |  | 16 | 0.85 | 1.32 | 0.000 | 0.635 | 0.325 | 7374 | tags=44%, list=12%, signal=50% |
| 988 | CELL\_ADHESION\_MEDIATOR\_ACTIVITY(GO:0098631) |  | 39 | 0.85 | 1.32 | 0.141 | 0.635 | 0.325 | 6670 | tags=8%, list=11%, signal=9% |
| 989 | TRANSCRIPTION\_FACTOR\_ACTIVITY\_\_RNA\_POLYMERASE\_II\_DISTAL\_ENHANCER\_SEQUENCE\_SPECIFIC\_BINDING(GO:0003705) |  | 99 | 0.81 | 1.32 | 0.000 | 0.635 | 0.325 | 7834 | tags=32%, list=13%, signal=37% |
| 990 | SPINAL\_CORD\_DORSAL\_VENTRAL\_PATTERNING(GO:0021513) |  | 22 | 0.88 | 1.32 | 0.000 | 0.635 | 0.325 | 6058 | tags=36%, list=10%, signal=40% |
| 991 | ENDOPLASMIC\_RETICULUM\_LUMEN(GO:0005788) |  | 297 | 0.84 | 1.32 | 0.000 | 0.634 | 0.325 | 7347 | tags=18%, list=12%, signal=20% |
| 992 | ALCOHOL\_METABOLIC\_PROCESS(GO:0006066) |  | 317 | 0.81 | 1.32 | 0.000 | 0.634 | 0.325 | 7358 | tags=13%, list=12%, signal=15% |
| 993 | NITRIC\_OXIDE\_SYNTHASE\_BIOSYNTHETIC\_PROCESS(GO:0051767) |  | 19 | 0.90 | 1.32 | 0.000 | 0.634 | 0.325 | 5825 | tags=5%, list=10%, signal=6% |
| 994 | REGULATION\_OF\_NITRIC\_OXIDE\_SYNTHASE\_BIOSYNTHETIC\_PROCESS(GO:0051769) |  | 19 | 0.90 | 1.32 | 0.000 | 0.633 | 0.325 | 5825 | tags=5%, list=10%, signal=6% |
| 995 | GLANDULAR\_EPITHELIAL\_CELL\_DIFFERENTIATION(GO:0002067) |  | 46 | 0.87 | 1.32 | 0.141 | 0.633 | 0.325 | 6473 | tags=28%, list=11%, signal=32% |
| 996 | POSITIVE\_REGULATION\_OF\_CYCLIN\_DEPENDENT\_PROTEIN\_KINASE\_ACTIVITY(GO:1904031) |  | 37 | 0.85 | 1.32 | 0.000 | 0.632 | 0.325 | 5852 | tags=5%, list=10%, signal=6% |
| 997 | TYPE\_2\_IMMUNE\_RESPONSE(GO:0042092) |  | 37 | 0.84 | 1.32 | 0.141 | 0.632 | 0.325 | 7435 | tags=30%, list=13%, signal=34% |
| 998 | POSITIVE\_REGULATION\_OF\_CYTOSOLIC\_CALCIUM\_ION\_CONCENTRATION(GO:0007204) |  | 283 | 0.83 | 1.32 | 0.141 | 0.632 | 0.325 | 7544 | tags=22%, list=13%, signal=25% |
| 999 | SARCOPLASM(GO:0016528) |  | 71 | 0.84 | 1.32 | 0.141 | 0.632 | 0.325 | 7754 | tags=24%, list=13%, signal=28% |
| 1000 | EXTRACELLULAR\_STRUCTURE\_ORGANIZATION(GO:0043062) |  | 389 | 0.84 | 1.32 | 0.141 | 0.631 | 0.325 | 7346 | tags=24%, list=12%, signal=27% |
| 1001 | CELL\_CELL\_ADHESION\_MEDIATOR\_ACTIVITY(GO:0098632) |  | 30 | 0.86 | 1.32 | 0.141 | 0.631 | 0.325 | 5845 | tags=3%, list=10%, signal=4% |
| 1002 | EXOCYTIC\_VESICLE\_MEMBRANE(GO:0099501) |  | 82 | 0.83 | 1.32 | 0.141 | 0.631 | 0.325 | 7924 | tags=22%, list=13%, signal=25% |
| 1003 | SYNAPTIC\_VESICLE\_MEMBRANE(GO:0030672) |  | 82 | 0.83 | 1.32 | 0.141 | 0.630 | 0.325 | 7924 | tags=22%, list=13%, signal=25% |
| 1004 | THIOESTER\_BIOSYNTHETIC\_PROCESS(GO:0035384) |  | 51 | 0.80 | 1.32 | 0.000 | 0.630 | 0.325 | 6042 | tags=6%, list=10%, signal=7% |
| 1005 | ACYL\_COA\_BIOSYNTHETIC\_PROCESS(GO:0071616) |  | 51 | 0.80 | 1.32 | 0.000 | 0.630 | 0.325 | 6042 | tags=6%, list=10%, signal=7% |
| 1006 | EPITHELIAL\_CELL\_DIFFERENTIATION\_INVOLVED\_IN\_KIDNEY\_DEVELOPMENT(GO:0035850) |  | 44 | 0.86 | 1.32 | 0.141 | 0.630 | 0.325 | 7134 | tags=25%, list=12%, signal=28% |
| 1007 | REGULATION\_OF\_STEM\_CELL\_PROLIFERATION(GO:0072091) |  | 62 | 0.88 | 1.32 | 0.141 | 0.629 | 0.325 | 6614 | tags=21%, list=11%, signal=24% |
| 1008 | COPULATION(GO:0007620) |  | 20 | 0.86 | 1.32 | 0.000 | 0.629 | 0.325 | 5956 | tags=25%, list=10%, signal=28% |
| 1009 | STRIATED\_MUSCLE\_CELL\_DIFFERENTIATION(GO:0051146) |  | 277 | 0.84 | 1.32 | 0.141 | 0.628 | 0.325 | 6670 | tags=17%, list=11%, signal=19% |
| 1010 | CEREBRAL\_CORTEX\_DEVELOPMENT(GO:0021987) |  | 114 | 0.87 | 1.32 | 0.141 | 0.628 | 0.325 | 6338 | tags=16%, list=11%, signal=18% |
| 1011 | AMMONIUM\_ION\_METABOLIC\_PROCESS(GO:0097164) |  | 195 | 0.83 | 1.32 | 0.141 | 0.628 | 0.325 | 6393 | tags=11%, list=11%, signal=12% |
| 1012 | CELLULAR\_DIVALENT\_INORGANIC\_CATION\_HOMEOSTASIS(GO:0072503) |  | 438 | 0.82 | 1.32 | 0.141 | 0.627 | 0.325 | 7544 | tags=19%, list=13%, signal=21% |
| 1013 | NEGATIVE\_REGULATION\_OF\_VASCULAR\_ENDOTHELIAL\_CELL\_PROLIFERATION(GO:1905563) |  | 20 | 0.88 | 1.32 | 0.000 | 0.627 | 0.325 | 7134 | tags=10%, list=12%, signal=11% |
| 1014 | RESPONSE\_TO\_INTERLEUKIN\_1(GO:0070555) |  | 196 | 0.81 | 1.32 | 0.000 | 0.627 | 0.325 | 6211 | tags=8%, list=11%, signal=9% |
| 1015 | MESODERM\_DEVELOPMENT(GO:0007498) |  | 132 | 0.85 | 1.32 | 0.141 | 0.627 | 0.325 | 7360 | tags=23%, list=12%, signal=26% |
| 1016 | COSTAMERE(GO:0043034) |  | 19 | 0.88 | 1.32 | 0.141 | 0.627 | 0.325 | 6605 | tags=21%, list=11%, signal=24% |
| 1017 | POSITIVE\_REGULATION\_OF\_FATTY\_ACID\_METABOLIC\_PROCESS(GO:0045923) |  | 32 | 0.82 | 1.32 | 0.000 | 0.627 | 0.325 | 6424 | tags=13%, list=11%, signal=14% |
| 1018 | CHEMOKINE\_BINDING(GO:0019956) |  | 24 | 0.85 | 1.32 | 0.141 | 0.627 | 0.325 | 5877 | tags=17%, list=10%, signal=18% |
| 1019 | NEURON\_APOPTOTIC\_PROCESS(GO:0051402) |  | 219 | 0.82 | 1.32 | 0.000 | 0.626 | 0.325 | 7639 | tags=20%, list=13%, signal=23% |
| 1020 | MAST\_CELL\_GRANULE(GO:0042629) |  | 22 | 0.87 | 1.32 | 0.141 | 0.626 | 0.325 | 5839 | tags=9%, list=10%, signal=10% |
| 1021 | NEGATIVE\_REGULATION\_OF\_VASCULAR\_SMOOTH\_MUSCLE\_CELL\_PROLIFERATION(GO:1904706) |  | 25 | 0.90 | 1.32 | 0.000 | 0.626 | 0.325 | 5825 | tags=4%, list=10%, signal=4% |
| 1022 | PHOSPHATIDYLSERINE\_ACYL\_CHAIN\_REMODELING(GO:0036150) |  | 21 | 0.88 | 1.32 | 0.000 | 0.626 | 0.325 | 5866 | tags=19%, list=10%, signal=21% |
| 1023 | POSITIVE\_REGULATION\_OF\_POTASSIUM\_ION\_TRANSMEMBRANE\_TRANSPORT(GO:1901381) |  | 36 | 0.83 | 1.32 | 0.000 | 0.625 | 0.325 | 9543 | tags=42%, list=16%, signal=50% |
| 1024 | MUSCLE\_ORGAN\_DEVELOPMENT(GO:0007517) |  | 395 | 0.83 | 1.32 | 0.000 | 0.625 | 0.325 | 7449 | tags=21%, list=13%, signal=23% |
| 1025 | REGULATION\_OF\_SYNAPSE\_ASSEMBLY(GO:0051963) |  | 86 | 0.85 | 1.32 | 0.141 | 0.624 | 0.325 | 7442 | tags=29%, list=13%, signal=33% |
| 1026 | POSITIVE\_REGULATION\_OF\_SODIUM\_ION\_TRANSPORT(GO:0010765) |  | 31 | 0.86 | 1.32 | 0.141 | 0.624 | 0.325 | 6432 | tags=29%, list=11%, signal=33% |
| 1027 | MOVEMENT\_OF\_CELL\_OR\_SUBCELLULAR\_COMPONENT(GO:0006928) |  | 2042 | 0.84 | 1.32 | 0.141 | 0.624 | 0.325 | 7169 | tags=15%, list=12%, signal=17% |
| 1028 | CLATHRIN\_COAT\_ASSEMBLY(GO:0048268) |  | 17 | 0.81 | 1.32 | 0.000 | 0.624 | 0.325 | 6847 | tags=12%, list=12%, signal=13% |
| 1029 | EXTRACELLULAR\_MATRIX\_ORGANIZATION(GO:0030198) |  | 336 | 0.85 | 1.32 | 0.141 | 0.624 | 0.325 | 7346 | tags=24%, list=12%, signal=27% |
| 1030 | VENTRAL\_SPINAL\_CORD\_DEVELOPMENT(GO:0021517) |  | 46 | 0.87 | 1.32 | 0.000 | 0.623 | 0.325 | 7149 | tags=41%, list=12%, signal=47% |
| 1031 | NEGATIVE\_REGULATION\_OF\_REGULATED\_SECRETORY\_PATHWAY(GO:1903306) |  | 22 | 0.81 | 1.32 | 0.000 | 0.623 | 0.325 | 6126 | tags=14%, list=10%, signal=15% |
| 1032 | PLASMA\_MEMBRANE\_REGION(GO:0098590) |  | 1049 | 0.84 | 1.32 | 0.141 | 0.623 | 0.325 | 7267 | tags=17%, list=12%, signal=19% |
| 1033 | VASCULAR\_ENDOTHELIAL\_GROWTH\_FACTOR\_SIGNALING\_PATHWAY(GO:0038084) |  | 35 | 0.86 | 1.32 | 0.000 | 0.622 | 0.325 | 7754 | tags=43%, list=13%, signal=49% |
| 1034 | AXIS\_SPECIFICATION(GO:0009798) |  | 89 | 0.85 | 1.32 | 0.141 | 0.622 | 0.325 | 7373 | tags=26%, list=12%, signal=29% |
| 1035 | NEGATIVE\_REGULATION\_OF\_TRANSPORTER\_ACTIVITY(GO:0032410) |  | 83 | 0.81 | 1.32 | 0.000 | 0.622 | 0.325 | 7979 | tags=23%, list=13%, signal=26% |
| 1036 | PYRIMIDINE\_NUCLEOSIDE\_METABOLIC\_PROCESS(GO:0006213) |  | 50 | 0.77 | 1.32 | 0.000 | 0.622 | 0.325 | 6865 | tags=16%, list=12%, signal=18% |
| 1037 | RESPONSE\_TO\_HYDROGEN\_PEROXIDE(GO:0042542) |  | 137 | 0.83 | 1.32 | 0.000 | 0.621 | 0.325 | 7544 | tags=12%, list=13%, signal=13% |
| 1038 | REGULATION\_OF\_RENAL\_SYSTEM\_PROCESS(GO:0098801) |  | 34 | 0.86 | 1.32 | 0.141 | 0.622 | 0.325 | 6525 | tags=29%, list=11%, signal=33% |
| 1039 | POSITIVE\_REGULATION\_OF\_SUPRAMOLECULAR\_FIBER\_ORGANIZATION(GO:1902905) |  | 190 | 0.82 | 1.32 | 0.141 | 0.621 | 0.325 | 6799 | tags=10%, list=11%, signal=11% |
| 1040 | GROOMING\_BEHAVIOR(GO:0007625) |  | 20 | 0.87 | 1.32 | 0.000 | 0.621 | 0.325 | 5983 | tags=15%, list=10%, signal=17% |
| 1041 | REGULATION\_OF\_ION\_TRANSMEMBRANE\_TRANSPORT(GO:0034765) |  | 439 | 0.82 | 1.32 | 0.141 | 0.621 | 0.325 | 7979 | tags=24%, list=13%, signal=28% |
| 1042 | CELL\_DIFFERENTIATION\_INVOLVED\_IN\_KIDNEY\_DEVELOPMENT(GO:0061005) |  | 54 | 0.86 | 1.32 | 0.141 | 0.621 | 0.325 | 7360 | tags=26%, list=12%, signal=30% |
| 1043 | LONG\_TERM\_MEMORY(GO:0007616) |  | 35 | 0.86 | 1.32 | 0.000 | 0.620 | 0.325 | 5852 | tags=11%, list=10%, signal=13% |
| 1044 | SULFUR\_COMPOUND\_CATABOLIC\_PROCESS(GO:0044273) |  | 48 | 0.80 | 1.32 | 0.000 | 0.620 | 0.325 | 7148 | tags=17%, list=12%, signal=19% |
| 1045 | MAMMARY\_GLAND\_EPITHELIAL\_CELL\_PROLIFERATION(GO:0033598) |  | 28 | 0.88 | 1.32 | 0.141 | 0.619 | 0.325 | 6473 | tags=14%, list=11%, signal=16% |
| 1046 | MAINTENANCE\_OF\_LOCATION(GO:0051235) |  | 306 | 0.80 | 1.32 | 0.000 | 0.619 | 0.325 | 8046 | tags=15%, list=14%, signal=17% |
| 1047 | RESPONSE\_TO\_WATER(GO:0009415) |  | 15 | 0.89 | 1.32 | 0.141 | 0.619 | 0.325 | 5856 | tags=20%, list=10%, signal=22% |
| 1048 | POSITIVE\_REGULATION\_OF\_MORPHOGENESIS\_OF\_AN\_EPITHELIUM(GO:1905332) |  | 33 | 0.88 | 1.32 | 0.141 | 0.619 | 0.325 | 6279 | tags=12%, list=11%, signal=14% |
| 1049 | PLATELET\_DERIVED\_GROWTH\_FACTOR\_RECEPTOR\_SIGNALING\_PATHWAY(GO:0048008) |  | 52 | 0.84 | 1.32 | 0.000 | 0.619 | 0.325 | 6617 | tags=10%, list=11%, signal=11% |
| 1050 | REGULATION\_OF\_STEROID\_HORMONE\_SECRETION(GO:2000831) |  | 19 | 0.80 | 1.32 | 0.000 | 0.618 | 0.325 | 8485 | tags=26%, list=14%, signal=31% |
| 1051 | PHOSPHOLIPID\_CATABOLIC\_PROCESS(GO:0009395) |  | 38 | 0.86 | 1.32 | 0.000 | 0.618 | 0.325 | 6393 | tags=18%, list=11%, signal=21% |
| 1052 | REGULATION\_OF\_CYTOSOLIC\_CALCIUM\_ION\_CONCENTRATION(GO:0051480) |  | 316 | 0.83 | 1.32 | 0.141 | 0.618 | 0.325 | 7544 | tags=21%, list=13%, signal=24% |
| 1053 | POSITIVE\_REGULATION\_OF\_POTASSIUM\_ION\_TRANSMEMBRANE\_TRANSPORTER\_ACTIVITY(GO:1901018) |  | 24 | 0.85 | 1.32 | 0.000 | 0.618 | 0.325 | 7249 | tags=25%, list=12%, signal=28% |
| 1054 | OLIGODENDROCYTE\_DEVELOPMENT(GO:0014003) |  | 39 | 0.89 | 1.32 | 0.141 | 0.617 | 0.325 | 6058 | tags=8%, list=10%, signal=9% |
| 1055 | POSITIVE\_REGULATION\_OF\_MESENCHYMAL\_CELL\_PROLIFERATION(GO:0002053) |  | 28 | 0.88 | 1.32 | 0.000 | 0.617 | 0.325 | 6473 | tags=25%, list=11%, signal=28% |
| 1056 | NEURAL\_CREST\_CELL\_DIFFERENTIATION(GO:0014033) |  | 69 | 0.86 | 1.32 | 0.141 | 0.617 | 0.325 | 7313 | tags=30%, list=12%, signal=35% |
| 1057 | ALCOHOL\_DEHYDROGENASE\_(NADP+)\_ACTIVITY(GO:0008106) |  | 15 | 0.89 | 1.32 | 0.000 | 0.616 | 0.325 | 5839 | tags=7%, list=10%, signal=7% |
| 1058 | POSITIVE\_REGULATION\_OF\_NEURAL\_PRECURSOR\_CELL\_PROLIFERATION(GO:2000179) |  | 46 | 0.88 | 1.32 | 0.141 | 0.616 | 0.325 | 6441 | tags=22%, list=11%, signal=24% |
| 1059 | NEURON\_PROJECTION\_TERMINUS(GO:0044306) |  | 131 | 0.83 | 1.32 | 0.000 | 0.615 | 0.325 | 7426 | tags=21%, list=13%, signal=24% |
| 1060 | MYELOID\_LEUKOCYTE\_ACTIVATION(GO:0002274) |  | 620 | 0.83 | 1.32 | 0.141 | 0.615 | 0.325 | 6696 | tags=8%, list=11%, signal=9% |
| 1061 | SARCOPLASMIC\_RETICULUM(GO:0016529) |  | 62 | 0.84 | 1.32 | 0.141 | 0.615 | 0.325 | 7549 | tags=19%, list=13%, signal=22% |
| 1062 | ANIMAL\_ORGAN\_MORPHOGENESIS(GO:0009887) |  | 1005 | 0.85 | 1.32 | 0.141 | 0.615 | 0.325 | 7155 | tags=18%, list=12%, signal=21% |
| 1063 | APOPTOTIC\_SIGNALING\_PATHWAY(GO:0097190) |  | 595 | 0.84 | 1.32 | 0.141 | 0.615 | 0.325 | 6662 | tags=7%, list=11%, signal=8% |
| 1064 | LYMPH\_VESSEL\_DEVELOPMENT(GO:0001945) |  | 24 | 0.84 | 1.32 | 0.000 | 0.614 | 0.325 | 7374 | tags=38%, list=12%, signal=43% |
| 1065 | VOLTAGE\_GATED\_CHANNEL\_ACTIVITY(GO:0022832) |  | 201 | 0.82 | 1.32 | 0.141 | 0.614 | 0.325 | 7979 | tags=25%, list=13%, signal=29% |
| 1066 | VOLTAGE\_GATED\_ION\_CHANNEL\_ACTIVITY(GO:0005244) |  | 201 | 0.82 | 1.32 | 0.141 | 0.613 | 0.325 | 7979 | tags=25%, list=13%, signal=29% |
| 1067 | POSITIVE\_REGULATION\_OF\_CALCIUM\_ION\_TRANSPORT(GO:0051928) |  | 111 | 0.85 | 1.32 | 0.141 | 0.613 | 0.325 | 6706 | tags=16%, list=11%, signal=18% |
| 1068 | STRUCTURAL\_MOLECULE\_ACTIVITY(GO:0005198) |  | 730 | 0.86 | 1.32 | 0.141 | 0.613 | 0.325 | 7015 | tags=11%, list=12%, signal=12% |
| 1069 | WNT\_ACTIVATED\_RECEPTOR\_ACTIVITY(GO:0042813) |  | 20 | 0.81 | 1.32 | 0.000 | 0.613 | 0.325 | 8109 | tags=35%, list=14%, signal=41% |
| 1070 | SARCOLEMMA(GO:0042383) |  | 121 | 0.85 | 1.32 | 0.141 | 0.612 | 0.325 | 7068 | tags=22%, list=12%, signal=25% |
| 1071 | MEMBRANE\_REGION(GO:0098589) |  | 311 | 0.84 | 1.32 | 0.141 | 0.612 | 0.325 | 7149 | tags=18%, list=12%, signal=20% |
| 1072 | HOMEOSTASIS\_OF\_NUMBER\_OF\_CELLS(GO:0048872) |  | 240 | 0.81 | 1.32 | 0.000 | 0.611 | 0.325 | 7258 | tags=15%, list=12%, signal=17% |
| 1073 | NEGATIVE\_REGULATION\_OF\_CELL\_DEATH(GO:0060548) |  | 969 | 0.84 | 1.32 | 0.141 | 0.611 | 0.325 | 6970 | tags=12%, list=12%, signal=14% |
| 1074 | REGULATION\_OF\_PEPTIDYL\_SERINE\_PHOSPHORYLATION(GO:0033135) |  | 136 | 0.79 | 1.32 | 0.000 | 0.611 | 0.325 | 8112 | tags=20%, list=14%, signal=23% |
| 1075 | GAMMA\_AMINOBUTYRIC\_ACID\_SIGNALING\_PATHWAY(GO:0007214) |  | 25 | 0.86 | 1.32 | 0.000 | 0.610 | 0.325 | 7049 | tags=24%, list=12%, signal=27% |
| 1076 | COFACTOR\_BINDING(GO:0048037) |  | 495 | 0.82 | 1.32 | 0.000 | 0.610 | 0.325 | 7265 | tags=10%, list=12%, signal=11% |
| 1077 | RENAL\_TUBULE\_DEVELOPMENT(GO:0061326) |  | 94 | 0.87 | 1.32 | 0.141 | 0.610 | 0.325 | 6282 | tags=18%, list=11%, signal=20% |
| 1078 | MEMBRANE\_RAFT(GO:0045121) |  | 300 | 0.84 | 1.32 | 0.141 | 0.609 | 0.325 | 7149 | tags=19%, list=12%, signal=21% |
| 1079 | MEMBRANE\_MICRODOMAIN(GO:0098857) |  | 300 | 0.84 | 1.32 | 0.141 | 0.609 | 0.325 | 7149 | tags=19%, list=12%, signal=21% |
| 1080 | TUBE\_MORPHOGENESIS(GO:0035239) |  | 889 | 0.84 | 1.32 | 0.141 | 0.609 | 0.325 | 7414 | tags=20%, list=13%, signal=23% |
| 1081 | PHOSPHATIDYLETHANOLAMINE\_ACYL\_CHAIN\_REMODELING(GO:0036152) |  | 26 | 0.88 | 1.32 | 0.141 | 0.608 | 0.325 | 5866 | tags=15%, list=10%, signal=17% |
| 1082 | MUSCLE\_CELL\_DEVELOPMENT(GO:0055001) |  | 174 | 0.85 | 1.32 | 0.141 | 0.608 | 0.325 | 6670 | tags=20%, list=11%, signal=23% |
| 1083 | CELL\_SURFACE\_RECEPTOR\_SIGNALING\_PATHWAY(GO:0007166) |  | 2976 | 0.84 | 1.32 | 0.141 | 0.607 | 0.325 | 7167 | tags=14%, list=12%, signal=15% |
| 1084 | PHOSPHATIDYLCHOLINE\_METABOLIC\_PROCESS(GO:0046470) |  | 80 | 0.84 | 1.32 | 0.141 | 0.607 | 0.325 | 6393 | tags=16%, list=11%, signal=18% |
| 1085 | TRANSPOSITION(GO:0032196) |  | 23 | 0.84 | 1.32 | 0.141 | 0.607 | 0.325 | 6865 | tags=30%, list=12%, signal=34% |
| 1086 | NEGATIVE\_REGULATION\_OF\_VIRAL\_LIFE\_CYCLE(GO:1903901) |  | 77 | 0.83 | 1.32 | 0.141 | 0.607 | 0.325 | 7293 | tags=16%, list=12%, signal=18% |
| 1087 | POSITIVE\_REGULATION\_OF\_VASCULAR\_ENDOTHELIAL\_CELL\_PROLIFERATION(GO:1905564) |  | 20 | 0.86 | 1.32 | 0.000 | 0.606 | 0.325 | 6958 | tags=15%, list=12%, signal=17% |
| 1088 | REGULATION\_OF\_VASCULAR\_ENDOTHELIAL\_GROWTH\_FACTOR\_SIGNALING\_PATHWAY(GO:1900746) |  | 18 | 0.88 | 1.32 | 0.000 | 0.606 | 0.325 | 5929 | tags=22%, list=10%, signal=25% |
| 1089 | AMELOGENESIS(GO:0097186) |  | 23 | 0.84 | 1.32 | 0.000 | 0.605 | 0.325 | 6068 | tags=13%, list=10%, signal=15% |
| 1090 | LYSOPHOSPHOLIPASE\_ACTIVITY(GO:0004622) |  | 21 | 0.86 | 1.32 | 0.141 | 0.605 | 0.325 | 6949 | tags=19%, list=12%, signal=22% |
| 1091 | STRIATED\_MUSCLE\_CELL\_DEVELOPMENT(GO:0055002) |  | 159 | 0.85 | 1.32 | 0.141 | 0.605 | 0.325 | 6670 | tags=21%, list=11%, signal=23% |
| 1092 | REGULATION\_OF\_CARDIAC\_MUSCLE\_CELL\_CONTRACTION(GO:0086004) |  | 40 | 0.83 | 1.32 | 0.000 | 0.605 | 0.325 | 7008 | tags=20%, list=12%, signal=23% |
| 1093 | NEGATIVE\_REGULATION\_OF\_CHONDROCYTE\_DIFFERENTIATION(GO:0032331) |  | 19 | 0.84 | 1.32 | 0.000 | 0.604 | 0.325 | 6635 | tags=32%, list=11%, signal=36% |
| 1094 | REGULATION\_OF\_CELLULAR\_SENESCENCE(GO:2000772) |  | 44 | 0.85 | 1.32 | 0.141 | 0.604 | 0.325 | 6685 | tags=14%, list=11%, signal=15% |
| 1095 | CELL\_MORPHOGENESIS(GO:0000902) |  | 951 | 0.84 | 1.32 | 0.141 | 0.603 | 0.325 | 7211 | tags=17%, list=12%, signal=19% |
| 1096 | ARACHIDONIC\_ACID\_SECRETION(GO:0050482) |  | 31 | 0.87 | 1.32 | 0.141 | 0.603 | 0.325 | 5866 | tags=10%, list=10%, signal=11% |
| 1097 | ARACHIDONATE\_TRANSPORT(GO:1903963) |  | 31 | 0.87 | 1.32 | 0.141 | 0.603 | 0.325 | 5866 | tags=10%, list=10%, signal=11% |
| 1098 | RESPONSE\_TO\_GAMMA\_RADIATION(GO:0010332) |  | 55 | 0.86 | 1.32 | 0.141 | 0.602 | 0.325 | 5834 | tags=4%, list=10%, signal=4% |
| 1099 | PRIMARY\_ALCOHOL\_CATABOLIC\_PROCESS(GO:0034310) |  | 15 | 0.89 | 1.32 | 0.141 | 0.602 | 0.325 | 5920 | tags=13%, list=10%, signal=15% |
| 1100 | DOPAMINE\_SECRETION(GO:0014046) |  | 28 | 0.87 | 1.32 | 0.000 | 0.602 | 0.325 | 7293 | tags=36%, list=12%, signal=41% |
| 1101 | REGULATION\_OF\_DOPAMINE\_SECRETION(GO:0014059) |  | 28 | 0.87 | 1.32 | 0.000 | 0.601 | 0.325 | 7293 | tags=36%, list=12%, signal=41% |
| 1102 | NEURAL\_PRECURSOR\_CELL\_PROLIFERATION(GO:0061351) |  | 138 | 0.86 | 1.32 | 0.141 | 0.601 | 0.325 | 6817 | tags=20%, list=12%, signal=23% |
| 1103 | GRANULOCYTE\_MACROPHAGE\_COLONY\_STIMULATING\_FACTOR\_PRODUCTION(GO:0032604) |  | 16 | 0.86 | 1.32 | 0.000 | 0.601 | 0.325 | 5958 | tags=25%, list=10%, signal=28% |
| 1104 | EPIDERMAL\_CELL\_DIFFERENTIATION(GO:0009913) |  | 351 | 0.85 | 1.32 | 0.141 | 0.600 | 0.325 | 7313 | tags=10%, list=12%, signal=11% |
| 1105 | NEGATIVE\_REGULATION\_OF\_MICROTUBULE\_POLYMERIZATION\_OR\_DEPOLYMERIZATION(GO:0031111) |  | 30 | 0.82 | 1.32 | 0.000 | 0.600 | 0.325 | 6040 | tags=7%, list=10%, signal=7% |
| 1106 | POSITIVE\_REGULATION\_OF\_CYCLIN\_DEPENDENT\_PROTEIN\_SERINE\_THREONINE\_KINASE\_ACTIVITY(GO:0045737) |  | 32 | 0.85 | 1.32 | 0.000 | 0.600 | 0.325 | 5852 | tags=6%, list=10%, signal=7% |
| 1107 | PHARYNGEAL\_SYSTEM\_DEVELOPMENT(GO:0060037) |  | 25 | 0.85 | 1.32 | 0.000 | 0.600 | 0.325 | 7913 | tags=36%, list=13%, signal=42% |
| 1108 | OXIDOREDUCTASE\_ACTIVITY\_\_ACTING\_ON\_SINGLE\_DONORS\_WITH\_INCORPORATION\_OF\_MOLECULAR\_OXYGEN\_\_INCORPORATION\_OF\_TWO\_ATOMS\_OF\_OXYGEN(GO:0016702) |  | 27 | 0.85 | 1.32 | 0.000 | 0.599 | 0.325 | 6465 | tags=19%, list=11%, signal=21% |
| 1109 | OXIDOREDUCTASE\_ACTIVITY\_\_ACTING\_ON\_SINGLE\_DONORS\_WITH\_INCORPORATION\_OF\_MOLECULAR\_OXYGEN(GO:0016701) |  | 28 | 0.84 | 1.32 | 0.000 | 0.599 | 0.325 | 6465 | tags=18%, list=11%, signal=20% |
| 1110 | FILOPODIUM\_TIP(GO:0032433) |  | 17 | 0.85 | 1.32 | 0.000 | 0.598 | 0.325 | 6733 | tags=24%, list=11%, signal=27% |
| 1111 | REGULATION\_OF\_ALCOHOL\_BIOSYNTHETIC\_PROCESS(GO:1902930) |  | 64 | 0.81 | 1.32 | 0.000 | 0.598 | 0.325 | 5955 | tags=9%, list=10%, signal=10% |
| 1112 | RESPONSE\_TO\_PROSTAGLANDIN\_E(GO:0034695) |  | 26 | 0.87 | 1.32 | 0.000 | 0.598 | 0.325 | 6527 | tags=19%, list=11%, signal=22% |
| 1113 | REGULATION\_OF\_MAMMARY\_GLAND\_EPITHELIAL\_CELL\_PROLIFERATION(GO:0033599) |  | 18 | 0.89 | 1.32 | 0.000 | 0.598 | 0.325 | 6294 | tags=11%, list=11%, signal=12% |
| 1114 | TUBE\_DEVELOPMENT(GO:0035295) |  | 1066 | 0.84 | 1.32 | 0.141 | 0.597 | 0.325 | 7435 | tags=20%, list=13%, signal=23% |
| 1115 | RESPONSE\_TO\_ACTIVITY(GO:0014823) |  | 61 | 0.82 | 1.32 | 0.141 | 0.597 | 0.325 | 7217 | tags=20%, list=12%, signal=22% |
| 1116 | CHEMICAL\_HOMEOSTASIS(GO:0048878) |  | 1074 | 0.83 | 1.32 | 0.141 | 0.597 | 0.325 | 7517 | tags=17%, list=13%, signal=19% |
| 1117 | BLOOD\_CIRCULATION(GO:0008015) |  | 526 | 0.83 | 1.32 | 0.141 | 0.597 | 0.325 | 7981 | tags=25%, list=13%, signal=28% |
| 1118 | BASEMENT\_MEMBRANE(GO:0005604) |  | 91 | 0.87 | 1.32 | 0.141 | 0.597 | 0.325 | 7010 | tags=25%, list=12%, signal=29% |
| 1119 | GLYCOSPHINGOLIPID\_BIOSYNTHETIC\_PROCESS(GO:0006688) |  | 24 | 0.83 | 1.32 | 0.000 | 0.596 | 0.325 | 8052 | tags=38%, list=14%, signal=43% |
| 1120 | DIGESTIVE\_SYSTEM\_DEVELOPMENT(GO:0055123) |  | 143 | 0.85 | 1.32 | 0.141 | 0.596 | 0.325 | 7042 | tags=29%, list=12%, signal=32% |
| 1121 | EXTRACELLULAR\_MATRIX\_COMPONENT(GO:0044420) |  | 117 | 0.87 | 1.32 | 0.141 | 0.596 | 0.325 | 7169 | tags=27%, list=12%, signal=31% |
| 1122 | PEPTIDYL\_ASPARAGINE\_MODIFICATION(GO:0018196) |  | 46 | 0.80 | 1.32 | 0.000 | 0.596 | 0.325 | 6848 | tags=15%, list=12%, signal=17% |
| 1123 | CIRCULATORY\_SYSTEM\_PROCESS(GO:0003013) |  | 530 | 0.83 | 1.32 | 0.141 | 0.595 | 0.325 | 7981 | tags=25%, list=13%, signal=28% |
| 1124 | PROTEIN\_LOCALIZATION\_TO\_CELL\_PERIPHERY(GO:1990778) |  | 266 | 0.83 | 1.32 | 0.141 | 0.595 | 0.325 | 7205 | tags=11%, list=12%, signal=12% |
| 1125 | REGULATION\_OF\_GRANULOCYTE\_MACROPHAGE\_COLONY\_STIMULATING\_FACTOR\_PRODUCTION(GO:0032645) |  | 15 | 0.86 | 1.32 | 0.000 | 0.594 | 0.325 | 5958 | tags=27%, list=10%, signal=30% |
| 1126 | SYNAPSE\_ORGANIZATION(GO:0050808) |  | 299 | 0.85 | 1.32 | 0.141 | 0.594 | 0.325 | 7487 | tags=18%, list=13%, signal=21% |
| 1127 | NEGATIVE\_REGULATION\_OF\_MULTICELLULAR\_ORGANISMAL\_PROCESS(GO:0051241) |  | 1164 | 0.84 | 1.32 | 0.141 | 0.594 | 0.325 | 7546 | tags=18%, list=13%, signal=20% |
| 1128 | FORMATION\_OF\_PRIMARY\_GERM\_LAYER(GO:0001704) |  | 127 | 0.83 | 1.32 | 0.000 | 0.593 | 0.325 | 7358 | tags=21%, list=12%, signal=24% |
| 1129 | REGULATION\_OF\_NEUTROPHIL\_MIGRATION(GO:1902622) |  | 39 | 0.86 | 1.32 | 0.141 | 0.593 | 0.325 | 7081 | tags=23%, list=12%, signal=26% |
| 1130 | ADENYLATE\_CYCLASE\_INHIBITING\_G\_PROTEIN\_COUPLED\_RECEPTOR\_SIGNALING\_PATHWAY(GO:0007193) |  | 88 | 0.85 | 1.32 | 0.141 | 0.593 | 0.325 | 7479 | tags=20%, list=13%, signal=23% |
| 1131 | REGULATION\_OF\_CYSTEINE\_TYPE\_ENDOPEPTIDASE\_ACTIVITY(GO:2000116) |  | 242 | 0.83 | 1.32 | 0.141 | 0.593 | 0.325 | 7211 | tags=10%, list=12%, signal=12% |
| 1132 | PROTEIN\_N\_LINKED\_GLYCOSYLATION\_VIA\_ASPARAGINE(GO:0018279) |  | 45 | 0.80 | 1.32 | 0.000 | 0.592 | 0.325 | 6848 | tags=16%, list=12%, signal=18% |
| 1133 | CALCIUM\_ION\_HOMEOSTASIS(GO:0055074) |  | 424 | 0.82 | 1.32 | 0.141 | 0.592 | 0.325 | 7544 | tags=19%, list=13%, signal=21% |
| 1134 | MYELINATION(GO:0042552) |  | 111 | 0.85 | 1.32 | 0.141 | 0.592 | 0.325 | 6572 | tags=15%, list=11%, signal=17% |
| 1135 | REGULATION\_OF\_TRANSMEMBRANE\_TRANSPORT(GO:0034762) |  | 525 | 0.81 | 1.32 | 0.141 | 0.591 | 0.325 | 7979 | tags=22%, list=13%, signal=25% |
| 1136 | RESPONSE\_TO\_GLUCAGON(GO:0033762) |  | 53 | 0.80 | 1.32 | 0.000 | 0.591 | 0.325 | 7785 | tags=19%, list=13%, signal=22% |
| 1137 | NEGATIVE\_REGULATION\_OF\_MUSCLE\_CONTRACTION(GO:0045932) |  | 26 | 0.82 | 1.32 | 0.000 | 0.591 | 0.325 | 8279 | tags=35%, list=14%, signal=40% |
| 1138 | MULTICELLULAR\_ORGANISMAL\_MOVEMENT(GO:0050879) |  | 50 | 0.83 | 1.32 | 0.000 | 0.591 | 0.325 | 6583 | tags=14%, list=11%, signal=16% |
| 1139 | MUSCULOSKELETAL\_MOVEMENT(GO:0050881) |  | 50 | 0.83 | 1.32 | 0.000 | 0.590 | 0.325 | 6583 | tags=14%, list=11%, signal=16% |
| 1140 | REGULATION\_OF\_MAPK\_CASCADE(GO:0043408) |  | 723 | 0.83 | 1.32 | 0.141 | 0.590 | 0.325 | 7754 | tags=17%, list=13%, signal=19% |
| 1141 | NEUROMUSCULAR\_JUNCTION(GO:0031594) |  | 56 | 0.81 | 1.32 | 0.000 | 0.589 | 0.325 | 8462 | tags=36%, list=14%, signal=42% |
| 1142 | NOREPINEPHRINE\_SECRETION(GO:0048243) |  | 18 | 0.85 | 1.32 | 0.000 | 0.589 | 0.325 | 5955 | tags=17%, list=10%, signal=19% |
| 1143 | CELLULAR\_KETONE\_METABOLIC\_PROCESS(GO:0042180) |  | 235 | 0.80 | 1.32 | 0.000 | 0.589 | 0.325 | 7226 | tags=9%, list=12%, signal=10% |
| 1144 | EPIDERMIS\_DEVELOPMENT(GO:0008544) |  | 457 | 0.85 | 1.32 | 0.141 | 0.589 | 0.325 | 6473 | tags=9%, list=11%, signal=10% |
| 1145 | NEGATIVE\_REGULATION\_OF\_NF\_KAPPAB\_TRANSCRIPTION\_FACTOR\_ACTIVITY(GO:0032088) |  | 79 | 0.80 | 1.32 | 0.000 | 0.588 | 0.325 | 6154 | tags=8%, list=10%, signal=8% |
| 1146 | REGULATION\_OF\_CYSTEINE\_TYPE\_ENDOPEPTIDASE\_ACTIVITY\_INVOLVED\_IN\_APOPTOTIC\_PROCESS(GO:0043281) |  | 209 | 0.83 | 1.32 | 0.141 | 0.588 | 0.325 | 7211 | tags=11%, list=12%, signal=12% |
| 1147 | COLUMNAR\_CUBOIDAL\_EPITHELIAL\_CELL\_DIFFERENTIATION(GO:0002065) |  | 111 | 0.84 | 1.32 | 0.141 | 0.588 | 0.325 | 6473 | tags=19%, list=11%, signal=21% |
| 1148 | GLYCOSYLCERAMIDE\_METABOLIC\_PROCESS(GO:0006677) |  | 17 | 0.80 | 1.32 | 0.000 | 0.588 | 0.325 | 8052 | tags=29%, list=14%, signal=34% |
| 1149 | CELLULAR\_RESPONSE\_TO\_CHEMICAL\_STIMULUS(GO:0070887) |  | 3126 | 0.84 | 1.32 | 0.141 | 0.588 | 0.325 | 6738 | tags=12%, list=11%, signal=12% |
| 1150 | VENTRICULAR\_CARDIAC\_MUSCLE\_CELL\_ACTION\_POTENTIAL(GO:0086005) |  | 35 | 0.83 | 1.32 | 0.000 | 0.587 | 0.325 | 9054 | tags=37%, list=15%, signal=44% |
| 1151 | ODONTOGENESIS\_OF\_DENTIN\_CONTAINING\_TOOTH(GO:0042475) |  | 85 | 0.85 | 1.32 | 0.000 | 0.587 | 0.325 | 6617 | tags=19%, list=11%, signal=21% |
| 1152 | REGULATION\_OF\_POTASSIUM\_ION\_TRANSMEMBRANE\_TRANSPORTER\_ACTIVITY(GO:1901016) |  | 59 | 0.81 | 1.32 | 0.000 | 0.587 | 0.325 | 7937 | tags=24%, list=13%, signal=27% |
| 1153 | KINESIN\_BINDING(GO:0019894) |  | 41 | 0.80 | 1.32 | 0.000 | 0.587 | 0.325 | 7973 | tags=20%, list=13%, signal=23% |
| 1154 | NEURONAL\_STEM\_CELL\_POPULATION\_MAINTENANCE(GO:0097150) |  | 22 | 0.85 | 1.32 | 0.000 | 0.586 | 0.325 | 5941 | tags=18%, list=10%, signal=20% |
| 1155 | LAMELLIPODIUM(GO:0030027) |  | 187 | 0.80 | 1.32 | 0.000 | 0.586 | 0.325 | 8112 | tags=15%, list=14%, signal=17% |
| 1156 | POTASSIUM\_ION\_ANTIPORTER\_ACTIVITY(GO:0022821) |  | 16 | 0.86 | 1.32 | 0.000 | 0.586 | 0.325 | 7073 | tags=25%, list=12%, signal=28% |
| 1157 | RETINA\_DEVELOPMENT\_IN\_CAMERA\_TYPE\_EYE(GO:0060041) |  | 140 | 0.86 | 1.32 | 0.141 | 0.585 | 0.325 | 7046 | tags=24%, list=12%, signal=27% |
| 1158 | ENSHEATHMENT\_OF\_NEURONS(GO:0007272) |  | 113 | 0.85 | 1.32 | 0.141 | 0.585 | 0.325 | 6572 | tags=15%, list=11%, signal=17% |
| 1159 | AXON\_ENSHEATHMENT(GO:0008366) |  | 113 | 0.85 | 1.32 | 0.141 | 0.585 | 0.325 | 6572 | tags=15%, list=11%, signal=17% |
| 1160 | AXONEME\_PART(GO:0044447) |  | 34 | 0.82 | 1.32 | 0.000 | 0.585 | 0.325 | 8261 | tags=32%, list=14%, signal=38% |
| 1161 | PROTEIN\_IMPORT\_INTO\_NUCLEUS(GO:0006606) |  | 142 | 0.82 | 1.32 | 0.000 | 0.585 | 0.325 | 7005 | tags=11%, list=12%, signal=12% |
| 1162 | TELENCEPHALON\_DEVELOPMENT(GO:0021537) |  | 246 | 0.85 | 1.32 | 0.141 | 0.585 | 0.325 | 6614 | tags=18%, list=11%, signal=20% |
| 1163 | COCHLEA\_MORPHOGENESIS(GO:0090103) |  | 25 | 0.88 | 1.32 | 0.000 | 0.584 | 0.325 | 6473 | tags=44%, list=11%, signal=49% |
| 1164 | KIDNEY\_MORPHOGENESIS(GO:0060993) |  | 92 | 0.86 | 1.32 | 0.141 | 0.584 | 0.325 | 7426 | tags=27%, list=13%, signal=31% |
| 1165 | EXOCYTOSIS(GO:0006887) |  | 849 | 0.81 | 1.32 | 0.000 | 0.583 | 0.325 | 7515 | tags=10%, list=13%, signal=12% |
| 1166 | SH3\_DOMAIN\_BINDING(GO:0017124) |  | 124 | 0.82 | 1.32 | 0.141 | 0.583 | 0.325 | 7963 | tags=15%, list=13%, signal=18% |
| 1167 | RENAL\_TUBULE\_MORPHOGENESIS(GO:0061333) |  | 77 | 0.88 | 1.32 | 0.141 | 0.583 | 0.325 | 6282 | tags=21%, list=11%, signal=23% |
| 1168 | POSITIVE\_REGULATION\_OF\_VASCULAR\_ENDOTHELIAL\_GROWTH\_FACTOR\_RECEPTOR\_SIGNALING\_PATHWAY(GO:0030949) |  | 19 | 0.85 | 1.32 | 0.000 | 0.582 | 0.325 | 5908 | tags=21%, list=10%, signal=23% |
| 1169 | GTP\_DEPENDENT\_PROTEIN\_BINDING(GO:0030742) |  | 22 | 0.82 | 1.32 | 0.000 | 0.582 | 0.325 | 8333 | tags=27%, list=14%, signal=32% |
| 1170 | CELL\_PROLIFERATION\_IN\_FOREBRAIN(GO:0021846) |  | 27 | 0.84 | 1.32 | 0.000 | 0.582 | 0.325 | 6614 | tags=26%, list=11%, signal=29% |
| 1171 | SODIUM\_ION\_TRANSMEMBRANE\_TRANSPORT(GO:0035725) |  | 137 | 0.84 | 1.32 | 0.141 | 0.582 | 0.325 | 7360 | tags=24%, list=12%, signal=27% |
| 1172 | KERATAN\_SULFATE\_METABOLIC\_PROCESS(GO:0042339) |  | 33 | 0.84 | 1.32 | 0.000 | 0.582 | 0.325 | 6985 | tags=24%, list=12%, signal=27% |
| 1173 | RENAL\_VESICLE\_MORPHOGENESIS(GO:0072077) |  | 19 | 0.88 | 1.32 | 0.000 | 0.582 | 0.325 | 6279 | tags=26%, list=11%, signal=29% |
| 1174 | REGULATION\_OF\_NEURON\_APOPTOTIC\_PROCESS(GO:0043523) |  | 195 | 0.82 | 1.32 | 0.000 | 0.582 | 0.325 | 7583 | tags=21%, list=13%, signal=24% |
| 1175 | REGULATION\_OF\_LIPASE\_ACTIVITY(GO:0060191) |  | 85 | 0.83 | 1.32 | 0.141 | 0.582 | 0.325 | 6928 | tags=25%, list=12%, signal=28% |
| 1176 | INTRACELLULAR\_LIPID\_TRANSPORT(GO:0032365) |  | 31 | 0.83 | 1.32 | 0.000 | 0.581 | 0.325 | 7172 | tags=13%, list=12%, signal=15% |
| 1177 | CELL\_ADHESION(GO:0007155) |  | 1330 | 0.84 | 1.32 | 0.141 | 0.581 | 0.325 | 7275 | tags=16%, list=12%, signal=18% |
| 1178 | ORGAN\_GROWTH(GO:0035265) |  | 174 | 0.84 | 1.32 | 0.141 | 0.581 | 0.325 | 7583 | tags=24%, list=13%, signal=27% |
| 1179 | EMBRYONIC\_CAMERA\_TYPE\_EYE\_MORPHOGENESIS(GO:0048596) |  | 26 | 0.88 | 1.32 | 0.141 | 0.580 | 0.325 | 6614 | tags=31%, list=11%, signal=35% |
| 1180 | PHAGOSOME\_MATURATION(GO:0090382) |  | 41 | 0.83 | 1.32 | 0.000 | 0.580 | 0.325 | 6008 | tags=7%, list=10%, signal=8% |
| 1181 | NEGATIVE\_REGULATION\_OF\_SMOOTH\_MUSCLE\_CONTRACTION(GO:0045986) |  | 16 | 0.84 | 1.32 | 0.000 | 0.580 | 0.325 | 8279 | tags=56%, list=14%, signal=65% |
| 1182 | METANEPHRIC\_RENAL\_VESICLE\_MORPHOGENESIS(GO:0072283) |  | 16 | 0.88 | 1.32 | 0.000 | 0.579 | 0.325 | 6279 | tags=25%, list=11%, signal=28% |
| 1183 | POSITIVE\_REGULATION\_OF\_ION\_TRANSPORT(GO:0043270) |  | 244 | 0.83 | 1.32 | 0.141 | 0.579 | 0.325 | 7516 | tags=21%, list=13%, signal=24% |
| 1184 | ODONTOGENESIS(GO:0042476) |  | 123 | 0.85 | 1.32 | 0.000 | 0.579 | 0.325 | 6617 | tags=19%, list=11%, signal=21% |
| 1185 | HEART\_VALVE\_DEVELOPMENT(GO:0003170) |  | 40 | 0.84 | 1.32 | 0.000 | 0.578 | 0.325 | 7800 | tags=23%, list=13%, signal=26% |
| 1186 | NEGATIVE\_REGULATION\_OF\_CELLULAR\_PROTEIN\_METABOLIC\_PROCESS(GO:0032269) |  | 1284 | 0.82 | 1.32 | 0.141 | 0.578 | 0.325 | 7170 | tags=10%, list=12%, signal=11% |
| 1187 | PALLIUM\_DEVELOPMENT(GO:0021543) |  | 169 | 0.86 | 1.32 | 0.141 | 0.578 | 0.325 | 6381 | tags=15%, list=11%, signal=17% |
| 1188 | POSITIVE\_REGULATION\_OF\_TOR\_SIGNALING(GO:0032008) |  | 36 | 0.79 | 1.32 | 0.000 | 0.578 | 0.325 | 5916 | tags=3%, list=10%, signal=3% |
| 1189 | NEGATIVE\_REGULATION\_OF\_PROTEIN\_METABOLIC\_PROCESS(GO:0051248) |  | 1341 | 0.82 | 1.32 | 0.141 | 0.578 | 0.325 | 7170 | tags=10%, list=12%, signal=11% |
| 1190 | NEGATIVE\_REGULATION\_OF\_CATION\_TRANSMEMBRANE\_TRANSPORT(GO:1904063) |  | 85 | 0.79 | 1.32 | 0.000 | 0.577 | 0.325 | 7979 | tags=21%, list=13%, signal=24% |
| 1191 | NEGATIVE\_REGULATION\_OF\_TISSUE\_REMODELING(GO:0034104) |  | 20 | 0.83 | 1.32 | 0.000 | 0.577 | 0.325 | 7504 | tags=25%, list=13%, signal=29% |
| 1192 | PURINE\_RIBONUCLEOTIDE\_CATABOLIC\_PROCESS(GO:0009154) |  | 34 | 0.79 | 1.32 | 0.000 | 0.577 | 0.325 | 9361 | tags=38%, list=16%, signal=45% |
| 1193 | NEUTROPHIL\_MIGRATION(GO:1990266) |  | 101 | 0.85 | 1.32 | 0.141 | 0.576 | 0.325 | 7081 | tags=18%, list=12%, signal=20% |
| 1194 | NUCLEOTIDE\_CATABOLIC\_PROCESS(GO:0009166) |  | 168 | 0.82 | 1.32 | 0.141 | 0.576 | 0.325 | 6696 | tags=8%, list=11%, signal=9% |
| 1195 | BIOLOGICAL\_ADHESION(GO:0022610) |  | 1338 | 0.84 | 1.32 | 0.141 | 0.576 | 0.325 | 7275 | tags=16%, list=12%, signal=18% |
| 1196 | PRESYNAPSE\_ORGANIZATION(GO:0099172) |  | 21 | 0.84 | 1.32 | 0.000 | 0.576 | 0.325 | 7373 | tags=29%, list=12%, signal=33% |
| 1197 | REGULATION\_OF\_EXCRETION(GO:0044062) |  | 24 | 0.85 | 1.32 | 0.141 | 0.575 | 0.325 | 6525 | tags=25%, list=11%, signal=28% |
| 1198 | CELL\_PROLIFERATION(GO:0008283) |  | 2082 | 0.84 | 1.32 | 0.141 | 0.575 | 0.325 | 6970 | tags=13%, list=12%, signal=15% |
| 1199 | CARDIAC\_CHAMBER\_DEVELOPMENT(GO:0003205) |  | 160 | 0.84 | 1.32 | 0.141 | 0.575 | 0.325 | 7374 | tags=22%, list=12%, signal=25% |
| 1200 | REGULATION\_OF\_SYNAPTIC\_VESICLE\_CYCLE(GO:0098693) |  | 45 | 0.80 | 1.32 | 0.000 | 0.575 | 0.325 | 9227 | tags=31%, list=16%, signal=37% |
| 1201 | CELL\_DIFFERENTIATION\_INVOLVED\_IN\_METANEPHROS\_DEVELOPMENT(GO:0072202) |  | 26 | 0.87 | 1.32 | 0.141 | 0.574 | 0.325 | 6279 | tags=23%, list=11%, signal=26% |
| 1202 | NEGATIVE\_REGULATION\_OF\_CARTILAGE\_DEVELOPMENT(GO:0061037) |  | 25 | 0.84 | 1.32 | 0.000 | 0.574 | 0.325 | 7913 | tags=36%, list=13%, signal=42% |
| 1203 | NEURON\_MIGRATION(GO:0001764) |  | 147 | 0.84 | 1.32 | 0.141 | 0.574 | 0.325 | 7134 | tags=18%, list=12%, signal=21% |
| 1204 | CELLULAR\_COMPONENT\_MORPHOGENESIS(GO:0032989) |  | 1049 | 0.84 | 1.32 | 0.141 | 0.574 | 0.325 | 7169 | tags=17%, list=12%, signal=19% |
| 1205 | ARP2\_3\_COMPLEX\_MEDIATED\_ACTIN\_NUCLEATION(GO:0034314) |  | 35 | 0.79 | 1.32 | 0.000 | 0.573 | 0.325 | 5927 | tags=6%, list=10%, signal=6% |
| 1206 | POSITIVE\_REGULATION\_OF\_OXIDOREDUCTASE\_ACTIVITY(GO:0051353) |  | 46 | 0.82 | 1.31 | 0.141 | 0.573 | 0.325 | 6005 | tags=7%, list=10%, signal=7% |
| 1207 | KIDNEY\_DEVELOPMENT(GO:0001822) |  | 263 | 0.84 | 1.31 | 0.141 | 0.573 | 0.325 | 7604 | tags=26%, list=13%, signal=30% |
| 1208 | EXTRACELLULAR\_MATRIX\_STRUCTURAL\_CONSTITUENT(GO:0005201) |  | 76 | 0.87 | 1.31 | 0.000 | 0.573 | 0.325 | 7222 | tags=32%, list=12%, signal=36% |
| 1209 | POSITIVE\_REGULATION\_OF\_INTERLEUKIN\_6\_SECRETION(GO:2000778) |  | 24 | 0.85 | 1.31 | 0.000 | 0.572 | 0.325 | 6486 | tags=13%, list=11%, signal=14% |
| 1210 | REGULATION\_OF\_LONG\_TERM\_NEURONAL\_SYNAPTIC\_PLASTICITY(GO:0048169) |  | 25 | 0.82 | 1.31 | 0.000 | 0.572 | 0.325 | 7721 | tags=12%, list=13%, signal=14% |
| 1211 | EPITHELIAL\_TUBE\_MORPHOGENESIS(GO:0060562) |  | 323 | 0.85 | 1.31 | 0.141 | 0.572 | 0.325 | 7382 | tags=19%, list=12%, signal=21% |
| 1212 | MITOCHONDRIAL\_RESPIRATORY\_CHAIN\_COMPLEX\_I(GO:0005747) |  | 52 | 0.78 | 1.31 | 0.000 | 0.572 | 0.325 | 5894 | tags=6%, list=10%, signal=6% |
| 1213 | RESPIRATORY\_CHAIN\_COMPLEX\_I(GO:0045271) |  | 52 | 0.78 | 1.31 | 0.000 | 0.572 | 0.325 | 5894 | tags=6%, list=10%, signal=6% |
| 1214 | NADH\_DEHYDROGENASE\_COMPLEX(GO:0030964) |  | 52 | 0.78 | 1.31 | 0.000 | 0.571 | 0.325 | 5894 | tags=6%, list=10%, signal=6% |
| 1215 | POSITIVE\_REGULATION\_OF\_POTASSIUM\_ION\_TRANSPORT(GO:0043268) |  | 43 | 0.83 | 1.31 | 0.141 | 0.571 | 0.325 | 9543 | tags=37%, list=16%, signal=44% |
| 1216 | NEUTROPHIL\_HOMEOSTASIS(GO:0001780) |  | 16 | 0.83 | 1.31 | 0.000 | 0.571 | 0.325 | 7545 | tags=25%, list=13%, signal=29% |
| 1217 | CARDIAC\_CHAMBER\_MORPHOGENESIS(GO:0003206) |  | 123 | 0.85 | 1.31 | 0.141 | 0.571 | 0.325 | 7374 | tags=25%, list=12%, signal=29% |
| 1218 | FOREBRAIN\_NEURON\_DEVELOPMENT(GO:0021884) |  | 30 | 0.87 | 1.31 | 0.000 | 0.570 | 0.325 | 6473 | tags=17%, list=11%, signal=19% |
| 1219 | GASTRULATION(GO:0007369) |  | 187 | 0.81 | 1.31 | 0.000 | 0.570 | 0.325 | 7360 | tags=19%, list=12%, signal=22% |
| 1220 | MUSCLE\_CELL\_DIFFERENTIATION(GO:0042692) |  | 364 | 0.83 | 1.31 | 0.141 | 0.570 | 0.325 | 7414 | tags=20%, list=13%, signal=22% |
| 1221 | REGULATION\_OF\_EPITHELIAL\_CELL\_PROLIFERATION(GO:0050678) |  | 355 | 0.84 | 1.31 | 0.141 | 0.570 | 0.325 | 7217 | tags=16%, list=12%, signal=18% |
| 1222 | COLLAGEN\_FIBRIL\_ORGANIZATION(GO:0030199) |  | 43 | 0.84 | 1.31 | 0.000 | 0.569 | 0.325 | 7322 | tags=26%, list=12%, signal=29% |
| 1223 | CAMERA\_TYPE\_EYE\_DEVELOPMENT(GO:0043010) |  | 298 | 0.84 | 1.31 | 0.141 | 0.569 | 0.325 | 7046 | tags=20%, list=12%, signal=22% |
| 1224 | SALIVARY\_GLAND\_MORPHOGENESIS(GO:0007435) |  | 33 | 0.84 | 1.31 | 0.141 | 0.569 | 0.325 | 7343 | tags=18%, list=12%, signal=21% |
| 1225 | REGULATION\_OF\_PROTEIN\_KINASE\_ACTIVITY(GO:0045859) |  | 787 | 0.81 | 1.31 | 0.141 | 0.569 | 0.325 | 6940 | tags=10%, list=12%, signal=11% |
| 1226 | EPITHELIAL\_TO\_MESENCHYMAL\_TRANSITION(GO:0001837) |  | 136 | 0.81 | 1.31 | 0.000 | 0.568 | 0.325 | 8177 | tags=21%, list=14%, signal=25% |
| 1227 | NEPHRON\_DEVELOPMENT(GO:0072006) |  | 134 | 0.85 | 1.31 | 0.141 | 0.568 | 0.325 | 7382 | tags=26%, list=12%, signal=30% |
| 1228 | VOLTAGE\_GATED\_CATION\_CHANNEL\_ACTIVITY(GO:0022843) |  | 145 | 0.83 | 1.31 | 0.141 | 0.568 | 0.325 | 7881 | tags=25%, list=13%, signal=29% |
| 1229 | CELLULAR\_RESPONSE\_TO\_INTERLEUKIN\_1(GO:0071347) |  | 173 | 0.80 | 1.31 | 0.000 | 0.568 | 0.325 | 6048 | tags=6%, list=10%, signal=7% |
| 1230 | RESPONSE\_TO\_ISCHEMIA(GO:0002931) |  | 37 | 0.82 | 1.31 | 0.000 | 0.568 | 0.325 | 7134 | tags=19%, list=12%, signal=22% |
| 1231 | PROTEIN\_LOCALIZATION\_TO\_PLASMA\_MEMBRANE(GO:0072659) |  | 237 | 0.83 | 1.31 | 0.141 | 0.568 | 0.325 | 6763 | tags=9%, list=11%, signal=10% |
| 1232 | COCHLEA\_DEVELOPMENT(GO:0090102) |  | 48 | 0.86 | 1.31 | 0.000 | 0.568 | 0.325 | 6473 | tags=25%, list=11%, signal=28% |
| 1233 | POSITIVE\_REGULATION\_OF\_ALCOHOL\_BIOSYNTHETIC\_PROCESS(GO:1902932) |  | 21 | 0.89 | 1.31 | 0.000 | 0.568 | 0.325 | 5955 | tags=24%, list=10%, signal=26% |
| 1234 | MONOSACCHARIDE\_BIOSYNTHETIC\_PROCESS(GO:0046364) |  | 88 | 0.84 | 1.31 | 0.141 | 0.568 | 0.325 | 6696 | tags=7%, list=11%, signal=8% |
| 1235 | OXIDOREDUCTASE\_ACTIVITY\_\_ACTING\_ON\_THE\_ALDEHYDE\_OR\_OXO\_GROUP\_OF\_DONORS(GO:0016903) |  | 46 | 0.86 | 1.31 | 0.141 | 0.567 | 0.325 | 6438 | tags=13%, list=11%, signal=15% |
| 1236 | LATERAL\_PLASMA\_MEMBRANE(GO:0016328) |  | 60 | 0.86 | 1.31 | 0.141 | 0.567 | 0.325 | 6784 | tags=17%, list=11%, signal=19% |
| 1237 | MULTIVESICULAR\_BODY(GO:0005771) |  | 39 | 0.83 | 1.31 | 0.000 | 0.567 | 0.325 | 5959 | tags=13%, list=10%, signal=14% |
| 1238 | EMBRYONIC\_ORGAN\_MORPHOGENESIS(GO:0048562) |  | 291 | 0.84 | 1.31 | 0.141 | 0.567 | 0.325 | 7134 | tags=24%, list=12%, signal=28% |
| 1239 | HEXOSE\_BIOSYNTHETIC\_PROCESS(GO:0019319) |  | 82 | 0.84 | 1.31 | 0.141 | 0.567 | 0.325 | 6696 | tags=7%, list=11%, signal=8% |
| 1240 | R\_SMAD\_BINDING(GO:0070412) |  | 23 | 0.82 | 1.31 | 0.000 | 0.566 | 0.325 | 5892 | tags=9%, list=10%, signal=10% |
| 1241 | NERVE\_DEVELOPMENT(GO:0021675) |  | 74 | 0.85 | 1.31 | 0.000 | 0.566 | 0.325 | 7115 | tags=28%, list=12%, signal=32% |
| 1242 | AMMONIUM\_TRANSMEMBRANE\_TRANSPORTER\_ACTIVITY(GO:0008519) |  | 29 | 0.85 | 1.31 | 0.000 | 0.566 | 0.325 | 7268 | tags=28%, list=12%, signal=31% |
| 1243 | HINDLIMB\_MORPHOGENESIS(GO:0035137) |  | 37 | 0.86 | 1.31 | 0.141 | 0.565 | 0.325 | 7360 | tags=41%, list=12%, signal=46% |
| 1244 | REGULATION\_OF\_RENAL\_SODIUM\_EXCRETION(GO:0035813) |  | 21 | 0.85 | 1.31 | 0.000 | 0.565 | 0.325 | 6525 | tags=29%, list=11%, signal=32% |
| 1245 | AMINE\_TRANSPORT(GO:0015837) |  | 86 | 0.83 | 1.31 | 0.141 | 0.565 | 0.325 | 7435 | tags=22%, list=13%, signal=25% |
| 1246 | PROTEIN\_DESTABILIZATION(GO:0031648) |  | 44 | 0.79 | 1.31 | 0.000 | 0.565 | 0.325 | 6154 | tags=5%, list=10%, signal=5% |
| 1247 | PHOSPHOLIPASE\_A2\_ACTIVITY(GO:0004623) |  | 37 | 0.85 | 1.31 | 0.141 | 0.565 | 0.325 | 6622 | tags=16%, list=11%, signal=18% |
| 1248 | CIRCULATORY\_SYSTEM\_DEVELOPMENT(GO:0072359) |  | 1094 | 0.83 | 1.31 | 0.141 | 0.564 | 0.325 | 7427 | tags=19%, list=13%, signal=21% |
| 1249 | ENDOCARDIAL\_CUSHION\_DEVELOPMENT(GO:0003197) |  | 39 | 0.84 | 1.31 | 0.141 | 0.564 | 0.325 | 8177 | tags=33%, list=14%, signal=39% |
| 1250 | SENSORY\_PERCEPTION(GO:0007600) |  | 925 | 0.82 | 1.31 | 0.141 | 0.564 | 0.325 | 8119 | tags=17%, list=14%, signal=19% |
| 1251 | RETINA\_HOMEOSTASIS(GO:0001895) |  | 67 | 0.83 | 1.31 | 0.141 | 0.564 | 0.325 | 7467 | tags=19%, list=13%, signal=22% |
| 1252 | ESTABLISHMENT\_OF\_SPINDLE\_LOCALIZATION(GO:0051293) |  | 34 | 0.81 | 1.31 | 0.000 | 0.563 | 0.325 | 6115 | tags=12%, list=10%, signal=13% |
| 1253 | FAT\_CELL\_DIFFERENTIATION(GO:0045444) |  | 211 | 0.81 | 1.31 | 0.000 | 0.563 | 0.325 | 7884 | tags=15%, list=13%, signal=17% |
| 1254 | CALCIUM\_DEPENDENT\_CELL\_CELL\_ADHESION\_VIA\_PLASMA\_MEMBRANE\_CELL\_ADHESION\_MOLECULES(GO:0016339) |  | 31 | 0.87 | 1.31 | 0.141 | 0.563 | 0.325 | 6502 | tags=19%, list=11%, signal=22% |
| 1255 | CARDIAC\_MUSCLE\_TISSUE\_DEVELOPMENT(GO:0048738) |  | 218 | 0.82 | 1.31 | 0.000 | 0.563 | 0.325 | 8122 | tags=25%, list=14%, signal=29% |
| 1256 | STEROID\_METABOLIC\_PROCESS(GO:0008202) |  | 311 | 0.81 | 1.31 | 0.000 | 0.562 | 0.325 | 7491 | tags=16%, list=13%, signal=19% |
| 1257 | REGULATION\_OF\_INOSITOL\_PHOSPHATE\_BIOSYNTHETIC\_PROCESS(GO:0010919) |  | 15 | 0.89 | 1.31 | 0.000 | 0.562 | 0.325 | 5955 | tags=33%, list=10%, signal=37% |
| 1258 | CELLULAR\_HORMONE\_METABOLIC\_PROCESS(GO:0034754) |  | 115 | 0.83 | 1.31 | 0.141 | 0.562 | 0.325 | 7255 | tags=16%, list=12%, signal=18% |
| 1259 | REGULATION\_OF\_NUCLEOCYTOPLASMIC\_TRANSPORT(GO:0046822) |  | 102 | 0.84 | 1.31 | 0.141 | 0.562 | 0.325 | 6642 | tags=7%, list=11%, signal=8% |
| 1260 | REGULATION\_OF\_SODIUM\_ION\_TRANSMEMBRANE\_TRANSPORTER\_ACTIVITY(GO:2000649) |  | 47 | 0.86 | 1.31 | 0.141 | 0.562 | 0.325 | 7360 | tags=32%, list=12%, signal=36% |
| 1261 | INTERLEUKIN\_8\_SECRETION(GO:0072606) |  | 28 | 0.81 | 1.31 | 0.000 | 0.562 | 0.325 | 6530 | tags=18%, list=11%, signal=20% |
| 1262 | CAMP\_CATABOLIC\_PROCESS(GO:0006198) |  | 19 | 0.78 | 1.31 | 0.000 | 0.562 | 0.325 | 7935 | tags=42%, list=13%, signal=49% |
| 1263 | GLUTAMATE\_SECRETION(GO:0014047) |  | 44 | 0.82 | 1.31 | 0.000 | 0.561 | 0.325 | 7873 | tags=25%, list=13%, signal=29% |
| 1264 | POSITIVE\_REGULATION\_OF\_CYTOSOLIC\_CALCIUM\_ION\_CONCENTRATION\_INVOLVED\_IN\_PHOSPHOLIPASE\_C\_ACTIVATING\_G\_PROTEIN\_COUPLED\_SIGNALING\_PATHWAY(GO:0051482) |  | 28 | 0.85 | 1.31 | 0.141 | 0.561 | 0.325 | 7479 | tags=32%, list=13%, signal=37% |
| 1265 | SENSORY\_ORGAN\_DEVELOPMENT(GO:0007423) |  | 530 | 0.83 | 1.31 | 0.141 | 0.561 | 0.325 | 7920 | tags=26%, list=13%, signal=29% |
| 1266 | PLASMA\_MEMBRANE\_BOUNDED\_CELL\_PROJECTION\_CYTOPLASM(GO:0032838) |  | 180 | 0.80 | 1.31 | 0.000 | 0.562 | 0.325 | 8290 | tags=22%, list=14%, signal=26% |
| 1267 | TEMPERATURE\_HOMEOSTASIS(GO:0001659) |  | 40 | 0.86 | 1.31 | 0.141 | 0.561 | 0.325 | 6424 | tags=13%, list=11%, signal=14% |
| 1268 | CELL\_CORTEX\_PART(GO:0044448) |  | 138 | 0.84 | 1.31 | 0.141 | 0.561 | 0.325 | 6581 | tags=12%, list=11%, signal=13% |
| 1269 | REGULATION\_OF\_INTRINSIC\_APOPTOTIC\_SIGNALING\_PATHWAY(GO:2001242) |  | 160 | 0.82 | 1.31 | 0.141 | 0.561 | 0.325 | 6514 | tags=6%, list=11%, signal=6% |
| 1270 | REGULATION\_OF\_VIRAL\_TRANSCRIPTION(GO:0046782) |  | 62 | 0.79 | 1.31 | 0.000 | 0.561 | 0.325 | 6381 | tags=5%, list=11%, signal=5% |
| 1271 | SALIVARY\_GLAND\_DEVELOPMENT(GO:0007431) |  | 37 | 0.84 | 1.31 | 0.141 | 0.560 | 0.325 | 7343 | tags=19%, list=12%, signal=22% |
| 1272 | REGULATION\_OF\_ORGAN\_GROWTH(GO:0046620) |  | 106 | 0.84 | 1.31 | 0.141 | 0.560 | 0.325 | 7583 | tags=24%, list=13%, signal=27% |
| 1273 | REGULATION\_OF\_LIPID\_CATABOLIC\_PROCESS(GO:0050994) |  | 51 | 0.82 | 1.31 | 0.000 | 0.560 | 0.325 | 6197 | tags=10%, list=10%, signal=11% |
| 1274 | SENSORY\_PERCEPTION\_OF\_SMELL(GO:0007608) |  | 419 | 0.78 | 1.31 | 0.141 | 0.560 | 0.325 | 7761 | tags=7%, list=13%, signal=8% |
| 1275 | POSITIVE\_REGULATION\_OF\_CHEMOKINE\_PRODUCTION(GO:0032722) |  | 56 | 0.80 | 1.31 | 0.000 | 0.560 | 0.325 | 6473 | tags=18%, list=11%, signal=20% |
| 1276 | CELL\_JUNCTION\_ORGANIZATION(GO:0034330) |  | 268 | 0.84 | 1.31 | 0.141 | 0.559 | 0.325 | 6796 | tags=9%, list=11%, signal=10% |
| 1277 | MESENCHYMAL\_TO\_EPITHELIAL\_TRANSITION(GO:0060231) |  | 20 | 0.88 | 1.31 | 0.141 | 0.559 | 0.325 | 6279 | tags=25%, list=11%, signal=28% |
| 1278 | POSITIVE\_REGULATION\_OF\_KIDNEY\_DEVELOPMENT(GO:0090184) |  | 39 | 0.88 | 1.31 | 0.141 | 0.559 | 0.325 | 6279 | tags=13%, list=11%, signal=14% |
| 1279 | POSITIVE\_REGULATION\_OF\_NERVOUS\_SYSTEM\_DEVELOPMENT(GO:0051962) |  | 490 | 0.83 | 1.31 | 0.141 | 0.559 | 0.325 | 7583 | tags=20%, list=13%, signal=23% |
| 1280 | LACTATE\_METABOLIC\_PROCESS(GO:0006089) |  | 15 | 0.82 | 1.31 | 0.000 | 0.559 | 0.325 | 6164 | tags=13%, list=10%, signal=15% |
| 1281 | CARDIAC\_MUSCLE\_CELL\_PROLIFERATION(GO:0060038) |  | 65 | 0.81 | 1.31 | 0.000 | 0.559 | 0.325 | 8380 | tags=26%, list=14%, signal=30% |
| 1282 | SULFOTRANSFERASE\_ACTIVITY(GO:0008146) |  | 55 | 0.83 | 1.31 | 0.000 | 0.559 | 0.325 | 8052 | tags=33%, list=14%, signal=38% |
| 1283 | CELLULAR\_SENESCENCE(GO:0090398) |  | 70 | 0.83 | 1.31 | 0.141 | 0.559 | 0.325 | 6685 | tags=13%, list=11%, signal=14% |
| 1284 | IMMUNE\_EFFECTOR\_PROCESS(GO:0002252) |  | 1204 | 0.80 | 1.31 | 0.000 | 0.558 | 0.325 | 7669 | tags=11%, list=13%, signal=12% |
| 1285 | POSITIVE\_REGULATION\_OF\_SMALL\_MOLECULE\_METABOLIC\_PROCESS(GO:0062013) |  | 170 | 0.82 | 1.31 | 0.141 | 0.558 | 0.325 | 7272 | tags=18%, list=12%, signal=21% |
| 1286 | FORELIMB\_MORPHOGENESIS(GO:0035136) |  | 40 | 0.86 | 1.31 | 0.141 | 0.559 | 0.325 | 7427 | tags=43%, list=13%, signal=49% |
| 1287 | REGULATION\_OF\_SYSTEM\_PROCESS(GO:0044057) |  | 537 | 0.81 | 1.31 | 0.141 | 0.558 | 0.325 | 7979 | tags=24%, list=13%, signal=28% |
| 1288 | SKELETAL\_MUSCLE\_CONTRACTION(GO:0003009) |  | 39 | 0.85 | 1.31 | 0.000 | 0.558 | 0.325 | 6583 | tags=18%, list=11%, signal=20% |
| 1289 | SUBSTRATE\_DEPENDENT\_CELL\_MIGRATION(GO:0006929) |  | 27 | 0.81 | 1.31 | 0.000 | 0.558 | 0.325 | 7898 | tags=26%, list=13%, signal=30% |
| 1290 | CELLULAR\_LIPID\_METABOLIC\_PROCESS(GO:0044255) |  | 1092 | 0.82 | 1.31 | 0.141 | 0.558 | 0.325 | 7358 | tags=13%, list=12%, signal=15% |
| 1291 | BRAIN\_DEVELOPMENT(GO:0007420) |  | 708 | 0.83 | 1.31 | 0.141 | 0.557 | 0.325 | 7668 | tags=20%, list=13%, signal=23% |
| 1292 | ORGANOPHOSPHATE\_CATABOLIC\_PROCESS(GO:0046434) |  | 225 | 0.82 | 1.31 | 0.141 | 0.558 | 0.325 | 6423 | tags=9%, list=11%, signal=10% |
| 1293 | ESTABLISHMENT\_OF\_MITOTIC\_SPINDLE\_ORIENTATION(GO:0000132) |  | 20 | 0.82 | 1.31 | 0.000 | 0.557 | 0.325 | 5892 | tags=5%, list=10%, signal=6% |
| 1294 | C2H2\_ZINC\_FINGER\_DOMAIN\_BINDING(GO:0070742) |  | 15 | 0.86 | 1.31 | 0.000 | 0.557 | 0.325 | 7024 | tags=27%, list=12%, signal=30% |
| 1295 | PEPTIDYL\_SERINE\_PHOSPHORYLATION(GO:0018105) |  | 293 | 0.79 | 1.31 | 0.000 | 0.557 | 0.325 | 8112 | tags=17%, list=14%, signal=19% |
| 1296 | NEGATIVE\_REGULATION\_OF\_WNT\_SIGNALING\_PATHWAY(GO:0030178) |  | 203 | 0.80 | 1.31 | 0.000 | 0.556 | 0.325 | 7913 | tags=17%, list=13%, signal=19% |
| 1297 | NEGATIVE\_REGULATION\_OF\_CELL\_DIFFERENTIATION(GO:0045596) |  | 679 | 0.83 | 1.31 | 0.141 | 0.556 | 0.325 | 7461 | tags=18%, list=13%, signal=20% |
| 1298 | DIVALENT\_INORGANIC\_CATION\_HOMEOSTASIS(GO:0072507) |  | 456 | 0.81 | 1.31 | 0.141 | 0.556 | 0.325 | 7544 | tags=19%, list=13%, signal=21% |
| 1299 | AMEBOIDAL\_TYPE\_CELL\_MIGRATION(GO:0001667) |  | 413 | 0.83 | 1.31 | 0.141 | 0.556 | 0.325 | 7396 | tags=17%, list=13%, signal=19% |
| 1300 | CELL\_CELL\_JUNCTION\_ORGANIZATION(GO:0045216) |  | 234 | 0.84 | 1.31 | 0.141 | 0.556 | 0.325 | 6796 | tags=10%, list=11%, signal=11% |
| 1301 | REFLEX(GO:0060004) |  | 20 | 0.87 | 1.31 | 0.000 | 0.555 | 0.325 | 6045 | tags=20%, list=10%, signal=22% |
| 1302 | NEGATIVE\_REGULATION\_OF\_VIRAL\_TRANSCRIPTION(GO:0032897) |  | 23 | 0.86 | 1.31 | 0.000 | 0.555 | 0.325 | 5890 | tags=4%, list=10%, signal=5% |
| 1303 | CELLULAR\_METAL\_ION\_HOMEOSTASIS(GO:0006875) |  | 528 | 0.80 | 1.31 | 0.000 | 0.555 | 0.325 | 7544 | tags=17%, list=13%, signal=19% |
| 1304 | CORTICAL\_CYTOSKELETON(GO:0030863) |  | 89 | 0.85 | 1.31 | 0.141 | 0.554 | 0.325 | 6581 | tags=16%, list=11%, signal=18% |
| 1305 | CYSTEINE\_TYPE\_ENDOPEPTIDASE\_REGULATOR\_ACTIVITY\_INVOLVED\_IN\_APOPTOTIC\_PROCESS(GO:0043028) |  | 42 | 0.80 | 1.31 | 0.000 | 0.554 | 0.325 | 7639 | tags=14%, list=13%, signal=16% |
| 1306 | TISSUE\_DEVELOPMENT(GO:0009888) |  | 1954 | 0.84 | 1.31 | 0.141 | 0.554 | 0.325 | 7162 | tags=16%, list=12%, signal=17% |
| 1307 | PROTEIN\_IMPORT(GO:0017038) |  | 194 | 0.80 | 1.31 | 0.000 | 0.554 | 0.325 | 7005 | tags=8%, list=12%, signal=9% |
| 1308 | CELL\_PROJECTION\_PART(GO:0044463) |  | 1318 | 0.83 | 1.31 | 0.141 | 0.554 | 0.325 | 7386 | tags=14%, list=12%, signal=15% |
| 1309 | PLASMA\_MEMBRANE\_BOUNDED\_CELL\_PROJECTION\_PART(GO:0120038) |  | 1318 | 0.83 | 1.31 | 0.141 | 0.553 | 0.325 | 7386 | tags=14%, list=12%, signal=15% |
| 1310 | ACTOMYOSIN\_STRUCTURE\_ORGANIZATION(GO:0031032) |  | 182 | 0.84 | 1.31 | 0.141 | 0.553 | 0.325 | 7898 | tags=23%, list=13%, signal=27% |
| 1311 | NUCLEOSIDE\_BISPHOSPHATE\_METABOLIC\_PROCESS(GO:0033865) |  | 40 | 0.82 | 1.31 | 0.000 | 0.553 | 0.325 | 7287 | tags=13%, list=12%, signal=14% |
| 1312 | PURINE\_NUCLEOSIDE\_BISPHOSPHATE\_METABOLIC\_PROCESS(GO:0034032) |  | 40 | 0.82 | 1.31 | 0.000 | 0.553 | 0.325 | 7287 | tags=13%, list=12%, signal=14% |
| 1313 | RIBONUCLEOSIDE\_BISPHOSPHATE\_METABOLIC\_PROCESS(GO:0033875) |  | 40 | 0.82 | 1.31 | 0.000 | 0.553 | 0.325 | 7287 | tags=13%, list=12%, signal=14% |
| 1314 | POSTSYNAPTIC\_MEMBRANE\_ORGANIZATION(GO:0001941) |  | 35 | 0.79 | 1.31 | 0.000 | 0.553 | 0.325 | 7442 | tags=20%, list=13%, signal=23% |
| 1315 | KIDNEY\_EPITHELIUM\_DEVELOPMENT(GO:0072073) |  | 138 | 0.85 | 1.31 | 0.141 | 0.552 | 0.325 | 7396 | tags=26%, list=13%, signal=30% |
| 1316 | POSITIVE\_REGULATION\_OF\_CELLULAR\_COMPONENT\_ORGANIZATION(GO:0051130) |  | 1203 | 0.83 | 1.31 | 0.141 | 0.552 | 0.325 | 7154 | tags=11%, list=12%, signal=13% |
| 1317 | NEGATIVE\_REGULATION\_OF\_UBIQUITIN\_PROTEIN\_TRANSFERASE\_ACTIVITY(GO:0051444) |  | 15 | 0.81 | 1.31 | 0.000 | 0.552 | 0.325 | 6154 | tags=13%, list=10%, signal=15% |
| 1318 | POSITIVE\_REGULATION\_OF\_CELL\_AGING(GO:0090343) |  | 19 | 0.87 | 1.31 | 0.141 | 0.552 | 0.325 | 6685 | tags=16%, list=11%, signal=18% |
| 1319 | HEART\_VALVE\_MORPHOGENESIS(GO:0003179) |  | 36 | 0.84 | 1.31 | 0.000 | 0.551 | 0.325 | 7134 | tags=17%, list=12%, signal=19% |
| 1320 | RIBONUCLEOTIDE\_CATABOLIC\_PROCESS(GO:0009261) |  | 35 | 0.78 | 1.31 | 0.000 | 0.551 | 0.325 | 9361 | tags=37%, list=16%, signal=44% |
| 1321 | RENAL\_VESICLE\_DEVELOPMENT(GO:0072087) |  | 20 | 0.88 | 1.31 | 0.000 | 0.551 | 0.325 | 6279 | tags=25%, list=11%, signal=28% |
| 1322 | REGULATION\_OF\_MUSCLE\_CONTRACTION(GO:0006937) |  | 167 | 0.81 | 1.31 | 0.000 | 0.551 | 0.325 | 8363 | tags=26%, list=14%, signal=30% |
| 1323 | REGULATION\_OF\_HISTONE\_ACETYLATION(GO:0035065) |  | 53 | 0.82 | 1.31 | 0.000 | 0.551 | 0.325 | 7029 | tags=9%, list=12%, signal=11% |
| 1324 | CELLULAR\_CARBOHYDRATE\_BIOSYNTHETIC\_PROCESS(GO:0034637) |  | 78 | 0.84 | 1.31 | 0.141 | 0.551 | 0.325 | 6006 | tags=9%, list=10%, signal=10% |
| 1325 | SOMITE\_DEVELOPMENT(GO:0061053) |  | 85 | 0.84 | 1.31 | 0.141 | 0.551 | 0.325 | 7913 | tags=28%, list=13%, signal=33% |
| 1326 | NEGATIVE\_REGULATION\_OF\_CELL\_PROLIFERATION(GO:0008285) |  | 725 | 0.82 | 1.31 | 0.141 | 0.551 | 0.325 | 7792 | tags=14%, list=13%, signal=16% |
| 1327 | NEPHRON\_EPITHELIUM\_DEVELOPMENT(GO:0072009) |  | 107 | 0.85 | 1.31 | 0.141 | 0.551 | 0.325 | 7382 | tags=26%, list=12%, signal=30% |
| 1328 | RENAL\_SODIUM\_EXCRETION(GO:0035812) |  | 22 | 0.84 | 1.31 | 0.000 | 0.550 | 0.325 | 6525 | tags=27%, list=11%, signal=31% |
| 1329 | RESPONSE\_TO\_DRUG(GO:0042493) |  | 962 | 0.82 | 1.31 | 0.141 | 0.550 | 0.325 | 7360 | tags=15%, list=12%, signal=16% |
| 1330 | POSITIVE\_REGULATION\_OF\_CARBOHYDRATE\_METABOLIC\_PROCESS(GO:0045913) |  | 71 | 0.84 | 1.31 | 0.141 | 0.550 | 0.325 | 6006 | tags=11%, list=10%, signal=13% |
| 1331 | CYTOSKELETON\_ORGANIZATION(GO:0007010) |  | 1222 | 0.83 | 1.31 | 0.141 | 0.550 | 0.325 | 7202 | tags=12%, list=12%, signal=13% |
| 1332 | FOREBRAIN\_CELL\_MIGRATION(GO:0021885) |  | 62 | 0.86 | 1.31 | 0.141 | 0.550 | 0.325 | 6338 | tags=18%, list=11%, signal=20% |
| 1333 | CAMP\_BINDING(GO:0030552) |  | 23 | 0.80 | 1.31 | 0.000 | 0.549 | 0.325 | 8582 | tags=35%, list=15%, signal=41% |
| 1334 | REGULATION\_OF\_NEUTROPHIL\_CHEMOTAXIS(GO:0090022) |  | 35 | 0.86 | 1.31 | 0.141 | 0.549 | 0.325 | 7081 | tags=26%, list=12%, signal=29% |
| 1335 | REGULATION\_OF\_CELLULAR\_COMPONENT\_BIOGENESIS(GO:0044087) |  | 871 | 0.83 | 1.31 | 0.141 | 0.549 | 0.325 | 7154 | tags=10%, list=12%, signal=11% |
| 1336 | RNA\_POLYMERASE\_II\_TRANSCRIPTION\_FACTOR\_ACTIVITY\_\_SEQUENCE\_SPECIFIC\_DNA\_BINDING(GO:0000981) |  | 1098 | 0.83 | 1.31 | 0.141 | 0.549 | 0.325 | 7385 | tags=21%, list=12%, signal=24% |
| 1337 | STEM\_CELL\_PROLIFERATION(GO:0072089) |  | 112 | 0.86 | 1.31 | 0.141 | 0.549 | 0.325 | 6831 | tags=21%, list=12%, signal=23% |
| 1338 | BETA\_TUBULIN\_BINDING(GO:0048487) |  | 37 | 0.80 | 1.31 | 0.000 | 0.548 | 0.325 | 7345 | tags=14%, list=12%, signal=15% |
| 1339 | EPITHELIAL\_CELL\_PROLIFERATION(GO:0050673) |  | 409 | 0.83 | 1.31 | 0.141 | 0.548 | 0.325 | 7217 | tags=16%, list=12%, signal=18% |
| 1340 | SYNAPTIC\_VESICLE(GO:0008021) |  | 155 | 0.82 | 1.31 | 0.141 | 0.548 | 0.325 | 6702 | tags=17%, list=11%, signal=19% |
| 1341 | OXOACID\_METABOLIC\_PROCESS(GO:0043436) |  | 1140 | 0.81 | 1.31 | 0.000 | 0.548 | 0.325 | 7287 | tags=12%, list=12%, signal=13% |
| 1342 | TRANSMEMBRANE\_RECEPTOR\_PROTEIN\_SERINE\_THREONINE\_KINASE\_ACTIVITY(GO:0004675) |  | 17 | 0.80 | 1.31 | 0.000 | 0.547 | 0.325 | 6578 | tags=24%, list=11%, signal=26% |
| 1343 | HEAD\_DEVELOPMENT(GO:0060322) |  | 751 | 0.83 | 1.31 | 0.141 | 0.547 | 0.325 | 7668 | tags=20%, list=13%, signal=23% |
| 1344 | PDZ\_DOMAIN\_BINDING(GO:0030165) |  | 89 | 0.79 | 1.31 | 0.000 | 0.547 | 0.325 | 8287 | tags=21%, list=14%, signal=25% |
| 1345 | HYDROLASE\_ACTIVITY\_\_ACTING\_ON\_CARBON\_NITROGEN\_(BUT\_NOT\_PEPTIDE)\_BONDS\_\_IN\_CYCLIC\_AMIDINES(GO:0016814) |  | 36 | 0.81 | 1.31 | 0.000 | 0.547 | 0.325 | 8202 | tags=22%, list=14%, signal=26% |
| 1346 | LIPOXYGENASE\_PATHWAY(GO:0019372) |  | 16 | 0.89 | 1.31 | 0.141 | 0.547 | 0.325 | 6424 | tags=31%, list=11%, signal=35% |
| 1347 | STRIATED\_MUSCLE\_CELL\_PROLIFERATION(GO:0014855) |  | 78 | 0.81 | 1.31 | 0.000 | 0.546 | 0.325 | 8380 | tags=27%, list=14%, signal=31% |
| 1348 | NEURON\_DEATH(GO:0070997) |  | 318 | 0.80 | 1.31 | 0.000 | 0.546 | 0.325 | 7963 | tags=18%, list=13%, signal=21% |
| 1349 | GLIAL\_CELL\_PROJECTION(GO:0097386) |  | 18 | 0.89 | 1.31 | 0.141 | 0.546 | 0.325 | 5856 | tags=17%, list=10%, signal=18% |
| 1350 | ACTIN\_MYOSIN\_FILAMENT\_SLIDING(GO:0033275) |  | 39 | 0.85 | 1.31 | 0.000 | 0.546 | 0.325 | 7015 | tags=36%, list=12%, signal=41% |
| 1351 | MUSCLE\_FILAMENT\_SLIDING(GO:0030049) |  | 39 | 0.85 | 1.31 | 0.000 | 0.545 | 0.325 | 7015 | tags=36%, list=12%, signal=41% |
| 1352 | MAPK\_CASCADE(GO:0000165) |  | 914 | 0.82 | 1.31 | 0.141 | 0.545 | 0.325 | 7754 | tags=16%, list=13%, signal=18% |
| 1353 | REGULATION\_OF\_EXTRINSIC\_APOPTOTIC\_SIGNALING\_PATHWAY\_IN\_ABSENCE\_OF\_LIGAND(GO:2001239) |  | 48 | 0.80 | 1.31 | 0.000 | 0.545 | 0.325 | 7322 | tags=15%, list=12%, signal=17% |
| 1354 | CARBOXYLIC\_ACID\_METABOLIC\_PROCESS(GO:0019752) |  | 1051 | 0.81 | 1.31 | 0.000 | 0.544 | 0.325 | 7270 | tags=11%, list=12%, signal=12% |
| 1355 | PEPTIDE\_METABOLIC\_PROCESS(GO:0006518) |  | 1073 | 0.81 | 1.31 | 0.000 | 0.544 | 0.325 | 6695 | tags=5%, list=11%, signal=6% |
| 1356 | DOPAMINE\_TRANSPORT(GO:0015872) |  | 40 | 0.86 | 1.31 | 0.000 | 0.544 | 0.325 | 7298 | tags=33%, list=12%, signal=37% |
| 1357 | REGULATION\_OF\_KINASE\_ACTIVITY(GO:0043549) |  | 849 | 0.81 | 1.31 | 0.141 | 0.544 | 0.325 | 7727 | tags=12%, list=13%, signal=14% |
| 1358 | CELLULAR\_RESPONSE\_TO\_ANTIBIOTIC(GO:0071236) |  | 144 | 0.81 | 1.31 | 0.141 | 0.544 | 0.325 | 7435 | tags=11%, list=13%, signal=13% |
| 1359 | POSITIVE\_REGULATION\_OF\_NEUROGENESIS(GO:0050769) |  | 427 | 0.83 | 1.31 | 0.141 | 0.544 | 0.325 | 7668 | tags=20%, list=13%, signal=23% |
| 1360 | NEUROTRANSMITTER\_BIOSYNTHETIC\_PROCESS(GO:0042136) |  | 95 | 0.82 | 1.31 | 0.141 | 0.544 | 0.325 | 7873 | tags=19%, list=13%, signal=22% |
| 1361 | CATECHOL\_CONTAINING\_COMPOUND\_BIOSYNTHETIC\_PROCESS(GO:0009713) |  | 20 | 0.88 | 1.31 | 0.141 | 0.544 | 0.325 | 6441 | tags=25%, list=11%, signal=28% |
| 1362 | CATECHOLAMINE\_BIOSYNTHETIC\_PROCESS(GO:0042423) |  | 20 | 0.88 | 1.31 | 0.141 | 0.544 | 0.325 | 6441 | tags=25%, list=11%, signal=28% |
| 1363 | POSITIVE\_REGULATION\_OF\_COAGULATION(GO:0050820) |  | 27 | 0.85 | 1.31 | 0.141 | 0.543 | 0.325 | 7128 | tags=19%, list=12%, signal=21% |
| 1364 | IN\_UTERO\_EMBRYONIC\_DEVELOPMENT(GO:0001701) |  | 322 | 0.84 | 1.31 | 0.141 | 0.543 | 0.325 | 6618 | tags=10%, list=11%, signal=11% |
| 1365 | CARDIAC\_EPITHELIAL\_TO\_MESENCHYMAL\_TRANSITION(GO:0060317) |  | 32 | 0.83 | 1.31 | 0.000 | 0.543 | 0.325 | 8177 | tags=34%, list=14%, signal=40% |
| 1366 | ENDOCARDIAL\_CUSHION\_FORMATION(GO:0003272) |  | 21 | 0.83 | 1.31 | 0.000 | 0.543 | 0.325 | 8177 | tags=43%, list=14%, signal=50% |
| 1367 | ORGANIC\_ACID\_METABOLIC\_PROCESS(GO:0006082) |  | 1156 | 0.81 | 1.31 | 0.000 | 0.542 | 0.325 | 7287 | tags=12%, list=12%, signal=13% |
| 1368 | CILIARY\_MEMBRANE(GO:0060170) |  | 84 | 0.85 | 1.31 | 0.141 | 0.542 | 0.325 | 7149 | tags=14%, list=12%, signal=16% |
| 1369 | CHOLESTEROL\_BIOSYNTHETIC\_PROCESS(GO:0006695) |  | 67 | 0.79 | 1.31 | 0.000 | 0.542 | 0.325 | 5861 | tags=4%, list=10%, signal=5% |
| 1370 | REGIONALIZATION(GO:0003002) |  | 333 | 0.85 | 1.31 | 0.141 | 0.542 | 0.325 | 7385 | tags=29%, list=12%, signal=33% |
| 1371 | RESPONSE\_TO\_ORGANIC\_SUBSTANCE(GO:0010033) |  | 3188 | 0.84 | 1.31 | 0.141 | 0.541 | 0.325 | 6628 | tags=11%, list=11%, signal=12% |
| 1372 | PASSIVE\_TRANSMEMBRANE\_TRANSPORTER\_ACTIVITY(GO:0022803) |  | 469 | 0.82 | 1.31 | 0.141 | 0.541 | 0.325 | 7979 | tags=26%, list=13%, signal=30% |
| 1373 | CHANNEL\_ACTIVITY(GO:0015267) |  | 468 | 0.82 | 1.31 | 0.141 | 0.541 | 0.325 | 7979 | tags=26%, list=13%, signal=30% |
| 1374 | CALCIUM\_ION\_TRANSPORT(GO:0006816) |  | 415 | 0.81 | 1.31 | 0.141 | 0.541 | 0.325 | 7979 | tags=20%, list=13%, signal=24% |
| 1375 | EMBRYONIC\_MORPHOGENESIS(GO:0048598) |  | 591 | 0.83 | 1.31 | 0.141 | 0.540 | 0.325 | 8003 | tags=25%, list=14%, signal=29% |
| 1376 | FRUCTOSE\_METABOLIC\_PROCESS(GO:0006000) |  | 16 | 0.90 | 1.31 | 0.141 | 0.540 | 0.325 | 5839 | tags=19%, list=10%, signal=21% |
| 1377 | ENDOTHELIAL\_CELL\_MIGRATION(GO:0043542) |  | 247 | 0.82 | 1.31 | 0.141 | 0.540 | 0.325 | 7698 | tags=17%, list=13%, signal=19% |
| 1378 | REGULATION\_OF\_CARDIAC\_MUSCLE\_CELL\_PROLIFERATION(GO:0060043) |  | 55 | 0.81 | 1.31 | 0.000 | 0.540 | 0.325 | 8380 | tags=27%, list=14%, signal=32% |
| 1379 | DIGESTIVE\_TRACT\_DEVELOPMENT(GO:0048565) |  | 130 | 0.84 | 1.31 | 0.141 | 0.540 | 0.325 | 7537 | tags=31%, list=13%, signal=35% |
| 1380 | REGULATION\_OF\_ANGIOGENESIS(GO:0045765) |  | 364 | 0.82 | 1.31 | 0.141 | 0.539 | 0.325 | 7267 | tags=19%, list=12%, signal=21% |
| 1381 | REGULATION\_OF\_GRANULOCYTE\_CHEMOTAXIS(GO:0071622) |  | 47 | 0.86 | 1.31 | 0.141 | 0.539 | 0.325 | 7081 | tags=21%, list=12%, signal=24% |
| 1382 | OSTEOBLAST\_DIFFERENTIATION(GO:0001649) |  | 206 | 0.82 | 1.31 | 0.141 | 0.539 | 0.325 | 7346 | tags=18%, list=12%, signal=20% |
| 1383 | POSITIVE\_REGULATION\_OF\_HEMOSTASIS(GO:1900048) |  | 26 | 0.86 | 1.31 | 0.141 | 0.539 | 0.325 | 7128 | tags=19%, list=12%, signal=22% |
| 1384 | POSITIVE\_REGULATION\_OF\_BLOOD\_COAGULATION(GO:0030194) |  | 26 | 0.86 | 1.31 | 0.141 | 0.538 | 0.325 | 7128 | tags=19%, list=12%, signal=22% |
| 1385 | POSITIVE\_REGULATION\_OF\_MESONEPHROS\_DEVELOPMENT(GO:0061213) |  | 22 | 0.88 | 1.31 | 0.141 | 0.538 | 0.325 | 6279 | tags=18%, list=11%, signal=20% |
| 1386 | PATTERN\_SPECIFICATION\_PROCESS(GO:0007389) |  | 423 | 0.84 | 1.31 | 0.141 | 0.538 | 0.325 | 7385 | tags=27%, list=12%, signal=31% |
| 1387 | REGULATION\_OF\_EPITHELIAL\_TO\_MESENCHYMAL\_TRANSITION(GO:0010717) |  | 87 | 0.82 | 1.31 | 0.000 | 0.538 | 0.325 | 7913 | tags=16%, list=13%, signal=19% |
| 1388 | EXOCYTIC\_VESICLE(GO:0070382) |  | 169 | 0.82 | 1.31 | 0.141 | 0.538 | 0.325 | 6702 | tags=15%, list=11%, signal=17% |
| 1389 | MATERNAL\_PROCESS\_INVOLVED\_IN\_FEMALE\_PREGNANCY(GO:0060135) |  | 61 | 0.84 | 1.31 | 0.141 | 0.538 | 0.325 | 7508 | tags=13%, list=13%, signal=15% |
| 1390 | REGULATION\_OF\_PEPTIDYL\_LYSINE\_ACETYLATION(GO:2000756) |  | 57 | 0.81 | 1.31 | 0.000 | 0.537 | 0.325 | 7029 | tags=11%, list=12%, signal=12% |
| 1391 | ANATOMICAL\_STRUCTURE\_FORMATION\_INVOLVED\_IN\_MORPHOGENESIS(GO:0048646) |  | 1130 | 0.83 | 1.31 | 0.141 | 0.537 | 0.325 | 7414 | tags=19%, list=13%, signal=21% |
| 1392 | EYE\_DEVELOPMENT(GO:0001654) |  | 346 | 0.83 | 1.31 | 0.141 | 0.537 | 0.325 | 7545 | tags=22%, list=13%, signal=25% |
| 1393 | STEROL\_BIOSYNTHETIC\_PROCESS(GO:0016126) |  | 72 | 0.78 | 1.31 | 0.000 | 0.537 | 0.325 | 5861 | tags=4%, list=10%, signal=5% |
| 1394 | POSITIVE\_REGULATION\_OF\_SMOOTHENED\_SIGNALING\_PATHWAY(GO:0045880) |  | 32 | 0.87 | 1.31 | 0.000 | 0.537 | 0.325 | 7059 | tags=34%, list=12%, signal=39% |
| 1395 | SECONDARY\_ALCOHOL\_BIOSYNTHETIC\_PROCESS(GO:1902653) |  | 68 | 0.78 | 1.31 | 0.000 | 0.537 | 0.325 | 5861 | tags=4%, list=10%, signal=5% |
| 1396 | ORGANIC\_HYDROXY\_COMPOUND\_METABOLIC\_PROCESS(GO:1901615) |  | 495 | 0.81 | 1.31 | 0.141 | 0.536 | 0.325 | 7393 | tags=15%, list=13%, signal=17% |
| 1397 | REGULATION\_OF\_HEMOPOIESIS(GO:1903706) |  | 454 | 0.81 | 1.31 | 0.000 | 0.536 | 0.325 | 7134 | tags=11%, list=12%, signal=12% |
| 1398 | REGULATION\_OF\_CELL\_PROLIFERATION(GO:0042127) |  | 1669 | 0.83 | 1.31 | 0.141 | 0.536 | 0.325 | 7169 | tags=14%, list=12%, signal=16% |
| 1399 | SIGNAL\_TRANSDUCTION\_BY\_PROTEIN\_PHOSPHORYLATION(GO:0023014) |  | 945 | 0.82 | 1.31 | 0.141 | 0.536 | 0.325 | 7754 | tags=16%, list=13%, signal=18% |
| 1400 | POSITIVE\_REGULATION\_OF\_SIGNAL\_TRANSDUCTION(GO:0009967) |  | 1554 | 0.83 | 1.31 | 0.141 | 0.536 | 0.325 | 7143 | tags=12%, list=12%, signal=14% |
| 1401 | ION\_CHANNEL\_ACTIVITY(GO:0005216) |  | 428 | 0.82 | 1.31 | 0.141 | 0.535 | 0.325 | 7979 | tags=26%, list=13%, signal=30% |
| 1402 | ANATOMICAL\_STRUCTURE\_HOMEOSTASIS(GO:0060249) |  | 387 | 0.81 | 1.31 | 0.141 | 0.536 | 0.325 | 7547 | tags=13%, list=13%, signal=14% |
| 1403 | RELAXATION\_OF\_MUSCLE(GO:0090075) |  | 32 | 0.80 | 1.31 | 0.000 | 0.535 | 0.325 | 8279 | tags=28%, list=14%, signal=33% |
| 1404 | PEPTIDYL\_SERINE\_MODIFICATION(GO:0018209) |  | 312 | 0.79 | 1.31 | 0.000 | 0.535 | 0.325 | 8112 | tags=18%, list=14%, signal=21% |
| 1405 | NEGATIVE\_REGULATION\_OF\_NEURON\_DIFFERENTIATION(GO:0045665) |  | 200 | 0.83 | 1.31 | 0.141 | 0.535 | 0.325 | 7461 | tags=21%, list=13%, signal=24% |
| 1406 | SUBSTRATE\_SPECIFIC\_CHANNEL\_ACTIVITY(GO:0022838) |  | 439 | 0.82 | 1.31 | 0.141 | 0.535 | 0.325 | 7979 | tags=26%, list=13%, signal=30% |
| 1407 | MESONEPHROS\_DEVELOPMENT(GO:0001823) |  | 102 | 0.85 | 1.31 | 0.141 | 0.535 | 0.325 | 7396 | tags=28%, list=13%, signal=32% |
| 1408 | LACTATION(GO:0007595) |  | 46 | 0.83 | 1.31 | 0.000 | 0.535 | 0.325 | 6958 | tags=13%, list=12%, signal=15% |
| 1409 | REGULATION\_OF\_CHONDROCYTE\_DIFFERENTIATION(GO:0032330) |  | 47 | 0.83 | 1.31 | 0.000 | 0.534 | 0.325 | 7781 | tags=40%, list=13%, signal=47% |
| 1410 | NEUTROPHIL\_CHEMOTAXIS(GO:0030593) |  | 90 | 0.85 | 1.31 | 0.141 | 0.534 | 0.325 | 7081 | tags=19%, list=12%, signal=21% |
| 1411 | PROTEIN\_TRANSPORT\_WITHIN\_LIPID\_BILAYER(GO:0032594) |  | 22 | 0.81 | 1.31 | 0.000 | 0.534 | 0.325 | 7558 | tags=14%, list=13%, signal=16% |
| 1412 | INTERMEDIATE\_FILAMENT(GO:0005882) |  | 200 | 0.85 | 1.31 | 0.141 | 0.534 | 0.325 | 5959 | tags=9%, list=10%, signal=10% |
| 1413 | APOPTOTIC\_PROCESS\_INVOLVED\_IN\_DEVELOPMENT(GO:1902742) |  | 34 | 0.83 | 1.31 | 0.000 | 0.534 | 0.325 | 7260 | tags=32%, list=12%, signal=37% |
| 1414 | MESONEPHRIC\_TUBULE\_DEVELOPMENT(GO:0072164) |  | 98 | 0.85 | 1.31 | 0.141 | 0.534 | 0.325 | 7396 | tags=29%, list=13%, signal=33% |
| 1415 | MESONEPHRIC\_EPITHELIUM\_DEVELOPMENT(GO:0072163) |  | 98 | 0.85 | 1.31 | 0.141 | 0.534 | 0.325 | 7396 | tags=29%, list=13%, signal=33% |
| 1416 | MONOCYTE\_CHEMOTACTIC\_PROTEIN\_1\_PRODUCTION(GO:0071605) |  | 18 | 0.89 | 1.31 | 0.000 | 0.534 | 0.325 | 5904 | tags=11%, list=10%, signal=12% |
| 1417 | REGULATION\_OF\_MONOCYTE\_CHEMOTACTIC\_PROTEIN\_1\_PRODUCTION(GO:0071637) |  | 18 | 0.89 | 1.31 | 0.000 | 0.534 | 0.325 | 5904 | tags=11%, list=10%, signal=12% |
| 1418 | CELL\_DEVELOPMENT(GO:0048468) |  | 2013 | 0.84 | 1.31 | 0.141 | 0.533 | 0.325 | 7217 | tags=16%, list=12%, signal=17% |
| 1419 | POSITIVE\_REGULATION\_OF\_EXTRACELLULAR\_MATRIX\_ORGANIZATION(GO:1903055) |  | 18 | 0.84 | 1.31 | 0.000 | 0.533 | 0.325 | 7640 | tags=22%, list=13%, signal=26% |
| 1420 | SEQUENCE\_SPECIFIC\_DNA\_BINDING(GO:0043565) |  | 1067 | 0.83 | 1.31 | 0.141 | 0.533 | 0.325 | 7358 | tags=19%, list=12%, signal=22% |
| 1421 | IMMUNE\_SYSTEM\_PROCESS(GO:0002376) |  | 3028 | 0.82 | 1.31 | 0.141 | 0.533 | 0.325 | 7492 | tags=12%, list=13%, signal=13% |
| 1422 | DENDRITE\_EXTENSION(GO:0097484) |  | 31 | 0.80 | 1.31 | 0.000 | 0.532 | 0.325 | 8949 | tags=45%, list=15%, signal=53% |
| 1423 | REGULATION\_OF\_DOUBLE\_STRAND\_BREAK\_REPAIR\_VIA\_NONHOMOLOGOUS\_END\_JOINING(GO:2001032) |  | 15 | 0.85 | 1.31 | 0.000 | 0.532 | 0.325 | 6910 | tags=13%, list=12%, signal=15% |
| 1424 | WOUND\_HEALING\_\_SPREADING\_OF\_CELLS(GO:0044319) |  | 31 | 0.84 | 1.31 | 0.000 | 0.532 | 0.325 | 5877 | tags=13%, list=10%, signal=14% |
| 1425 | EPIBOLY\_INVOLVED\_IN\_WOUND\_HEALING(GO:0090505) |  | 31 | 0.84 | 1.31 | 0.000 | 0.532 | 0.325 | 5877 | tags=13%, list=10%, signal=14% |
| 1426 | BONE\_MORPHOGENESIS(GO:0060349) |  | 90 | 0.81 | 1.31 | 0.000 | 0.532 | 0.325 | 8445 | tags=28%, list=14%, signal=32% |
| 1427 | REGULATION\_OF\_LEUKOCYTE\_CHEMOTAXIS(GO:0002688) |  | 107 | 0.84 | 1.31 | 0.141 | 0.532 | 0.325 | 7081 | tags=18%, list=12%, signal=20% |
| 1428 | CORTICOSTEROID\_HORMONE\_SECRETION(GO:0035930) |  | 17 | 0.80 | 1.31 | 0.000 | 0.531 | 0.325 | 8485 | tags=29%, list=14%, signal=34% |
| 1429 | MUSCLE\_STRUCTURE\_DEVELOPMENT(GO:0061061) |  | 642 | 0.83 | 1.31 | 0.141 | 0.531 | 0.325 | 7449 | tags=20%, list=13%, signal=23% |
| 1430 | NEGATIVE\_REGULATION\_OF\_CELL\_COMMUNICATION(GO:0010648) |  | 1333 | 0.82 | 1.31 | 0.141 | 0.531 | 0.325 | 7670 | tags=14%, list=13%, signal=16% |
| 1431 | CARDIAC\_MUSCLE\_TISSUE\_GROWTH(GO:0055017) |  | 95 | 0.80 | 1.31 | 0.000 | 0.531 | 0.325 | 8380 | tags=25%, list=14%, signal=29% |
| 1432 | PHENOL\_CONTAINING\_COMPOUND\_BIOSYNTHETIC\_PROCESS(GO:0046189) |  | 39 | 0.86 | 1.31 | 0.141 | 0.531 | 0.325 | 6923 | tags=33%, list=12%, signal=38% |
| 1433 | NEGATIVE\_REGULATION\_OF\_SIGNALING(GO:0023057) |  | 1337 | 0.82 | 1.31 | 0.141 | 0.531 | 0.325 | 7670 | tags=14%, list=13%, signal=16% |
| 1434 | AXON\_GUIDANCE(GO:0007411) |  | 233 | 0.83 | 1.31 | 0.141 | 0.531 | 0.325 | 7362 | tags=22%, list=12%, signal=25% |
| 1435 | REGULATION\_OF\_PROTEIN\_KINASE\_C\_SIGNALING(GO:0090036) |  | 17 | 0.85 | 1.31 | 0.000 | 0.531 | 0.325 | 6473 | tags=29%, list=11%, signal=33% |
| 1436 | RESPONSE\_TO\_X\_RAY(GO:0010165) |  | 35 | 0.85 | 1.31 | 0.141 | 0.530 | 0.325 | 6027 | tags=9%, list=10%, signal=10% |
| 1437 | POSITIVE\_REGULATION\_OF\_CELL\_DEVELOPMENT(GO:0010720) |  | 502 | 0.82 | 1.31 | 0.141 | 0.530 | 0.325 | 7293 | tags=18%, list=12%, signal=20% |
| 1438 | REGULATION\_OF\_PHOSPHOLIPASE\_ACTIVITY(GO:0010517) |  | 59 | 0.83 | 1.31 | 0.141 | 0.530 | 0.325 | 6928 | tags=29%, list=12%, signal=33% |
| 1439 | LIPID\_HOMEOSTASIS(GO:0055088) |  | 112 | 0.79 | 1.31 | 0.000 | 0.530 | 0.325 | 7699 | tags=15%, list=13%, signal=17% |
| 1440 | IMPORT\_INTO\_NUCLEUS(GO:0051170) |  | 163 | 0.81 | 1.30 | 0.000 | 0.530 | 0.325 | 7005 | tags=10%, list=12%, signal=11% |
| 1441 | RAL\_GTPASE\_BINDING(GO:0017160) |  | 16 | 0.85 | 1.30 | 0.000 | 0.530 | 0.325 | 6478 | tags=13%, list=11%, signal=14% |
| 1442 | ESTABLISHMENT\_OF\_MITOCHONDRION\_LOCALIZATION(GO:0051654) |  | 21 | 0.80 | 1.30 | 0.000 | 0.529 | 0.325 | 6798 | tags=19%, list=11%, signal=22% |
| 1443 | EYE\_PHOTORECEPTOR\_CELL\_DEVELOPMENT(GO:0042462) |  | 33 | 0.84 | 1.30 | 0.141 | 0.529 | 0.325 | 8035 | tags=30%, list=14%, signal=35% |
| 1444 | REGULATION\_OF\_NEURAL\_PRECURSOR\_CELL\_PROLIFERATION(GO:2000177) |  | 80 | 0.87 | 1.30 | 0.141 | 0.529 | 0.325 | 6614 | tags=21%, list=11%, signal=24% |
| 1445 | CYCLIC\_NUCLEOTIDE\_BINDING(GO:0030551) |  | 37 | 0.79 | 1.30 | 0.000 | 0.529 | 0.325 | 7591 | tags=38%, list=13%, signal=43% |
| 1446 | POSITIVE\_REGULATION\_OF\_PROTEIN\_KINASE\_B\_SIGNALING(GO:0051897) |  | 160 | 0.83 | 1.30 | 0.141 | 0.529 | 0.325 | 7056 | tags=17%, list=12%, signal=19% |
| 1447 | REGULATION\_OF\_MUSCLE\_ADAPTATION(GO:0043502) |  | 94 | 0.80 | 1.30 | 0.000 | 0.529 | 0.325 | 8328 | tags=13%, list=14%, signal=15% |
| 1448 | SKELETAL\_SYSTEM\_MORPHOGENESIS(GO:0048705) |  | 212 | 0.83 | 1.30 | 0.141 | 0.528 | 0.325 | 7360 | tags=25%, list=12%, signal=28% |
| 1449 | MICROVILLUS\_ASSEMBLY(GO:0030033) |  | 16 | 0.85 | 1.30 | 0.000 | 0.529 | 0.325 | 6763 | tags=19%, list=11%, signal=21% |
| 1450 | NEGATIVE\_REGULATION\_OF\_SIGNAL\_TRANSDUCTION(GO:0009968) |  | 1231 | 0.82 | 1.30 | 0.141 | 0.528 | 0.325 | 7670 | tags=14%, list=13%, signal=16% |
| 1451 | GRANULOCYTE\_MIGRATION(GO:0097530) |  | 121 | 0.84 | 1.30 | 0.141 | 0.528 | 0.325 | 7081 | tags=16%, list=12%, signal=18% |
| 1452 | CARDIOCYTE\_DIFFERENTIATION(GO:0035051) |  | 148 | 0.80 | 1.30 | 0.000 | 0.528 | 0.325 | 8112 | tags=23%, list=14%, signal=27% |
| 1453 | POSITIVE\_REGULATION\_OF\_CELL\_PROLIFERATION(GO:0008284) |  | 911 | 0.82 | 1.30 | 0.141 | 0.528 | 0.325 | 7134 | tags=16%, list=12%, signal=18% |
| 1454 | NEGATIVE\_REGULATION\_OF\_VIRAL\_PROCESS(GO:0048525) |  | 90 | 0.82 | 1.30 | 0.141 | 0.528 | 0.325 | 7293 | tags=14%, list=12%, signal=16% |
| 1455 | REGULATION\_OF\_METAL\_ION\_TRANSPORT(GO:0010959) |  | 364 | 0.82 | 1.30 | 0.141 | 0.528 | 0.325 | 7546 | tags=21%, list=13%, signal=24% |
| 1456 | URETERIC\_BUD\_DEVELOPMENT(GO:0001657) |  | 97 | 0.85 | 1.30 | 0.141 | 0.527 | 0.325 | 7396 | tags=29%, list=13%, signal=33% |
| 1457 | FOREBRAIN\_DEVELOPMENT(GO:0030900) |  | 378 | 0.83 | 1.30 | 0.141 | 0.527 | 0.325 | 7081 | tags=19%, list=12%, signal=22% |
| 1458 | NEGATIVE\_REGULATION\_OF\_PLATELET\_DERIVED\_GROWTH\_FACTOR\_RECEPTOR\_SIGNALING\_PATHWAY(GO:0010642) |  | 17 | 0.86 | 1.30 | 0.141 | 0.527 | 0.325 | 6617 | tags=18%, list=11%, signal=20% |
| 1459 | REGULATION\_OF\_TRANSPORT(GO:0051049) |  | 1803 | 0.82 | 1.30 | 0.141 | 0.527 | 0.325 | 7558 | tags=16%, list=13%, signal=17% |
| 1460 | REGULATION\_OF\_PLATELET\_DERIVED\_GROWTH\_FACTOR\_RECEPTOR\_SIGNALING\_PATHWAY(GO:0010640) |  | 25 | 0.85 | 1.30 | 0.141 | 0.527 | 0.325 | 6617 | tags=12%, list=11%, signal=14% |
| 1461 | CALCIUM\_ION\_BINDING(GO:0005509) |  | 708 | 0.82 | 1.30 | 0.141 | 0.526 | 0.325 | 7374 | tags=16%, list=12%, signal=18% |
| 1462 | REGULATION\_OF\_DEVELOPMENTAL\_PIGMENTATION(GO:0048070) |  | 17 | 0.78 | 1.30 | 0.000 | 0.526 | 0.325 | 6928 | tags=18%, list=12%, signal=20% |
| 1463 | HEART\_FORMATION(GO:0060914) |  | 25 | 0.85 | 1.30 | 0.000 | 0.526 | 0.325 | 5958 | tags=12%, list=10%, signal=13% |
| 1464 | NEURON\_PROJECTION\_GUIDANCE(GO:0097485) |  | 235 | 0.83 | 1.30 | 0.141 | 0.526 | 0.325 | 7362 | tags=22%, list=12%, signal=25% |
| 1465 | POSITIVE\_REGULATION\_OF\_PHOSPHATIDYLINOSITOL\_3\_KINASE\_SIGNALING(GO:0014068) |  | 68 | 0.82 | 1.30 | 0.141 | 0.526 | 0.325 | 7046 | tags=21%, list=12%, signal=23% |
| 1466 | POSITIVE\_REGULATION\_OF\_DNA\_BINDING\_TRANSCRIPTION\_FACTOR\_ACTIVITY(GO:0051091) |  | 246 | 0.81 | 1.30 | 0.141 | 0.525 | 0.325 | 6928 | tags=12%, list=12%, signal=14% |
| 1467 | TRANSMISSION\_OF\_NERVE\_IMPULSE(GO:0019226) |  | 70 | 0.81 | 1.30 | 0.141 | 0.525 | 0.325 | 7932 | tags=31%, list=13%, signal=36% |
| 1468 | PHOSPHOLIPASE\_BINDING(GO:0043274) |  | 15 | 0.85 | 1.30 | 0.000 | 0.525 | 0.325 | 5894 | tags=13%, list=10%, signal=15% |
| 1469 | POSITIVE\_REGULATION\_OF\_NEURON\_DIFFERENTIATION(GO:0045666) |  | 339 | 0.82 | 1.30 | 0.141 | 0.525 | 0.325 | 7727 | tags=21%, list=13%, signal=24% |
| 1470 | POSITIVE\_REGULATION\_OF\_EPITHELIAL\_CELL\_PROLIFERATION(GO:0050679) |  | 192 | 0.84 | 1.30 | 0.141 | 0.525 | 0.325 | 6965 | tags=17%, list=12%, signal=19% |
| 1471 | MYELOID\_LEUKOCYTE\_MIGRATION(GO:0097529) |  | 177 | 0.83 | 1.30 | 0.141 | 0.524 | 0.325 | 7081 | tags=17%, list=12%, signal=19% |
| 1472 | CELL\_DIFFERENTIATION\_IN\_SPINAL\_CORD(GO:0021515) |  | 53 | 0.86 | 1.30 | 0.000 | 0.524 | 0.325 | 7149 | tags=40%, list=12%, signal=45% |
| 1473 | REGULATION\_OF\_EPIDERMIS\_DEVELOPMENT(GO:0045682) |  | 79 | 0.85 | 1.30 | 0.141 | 0.524 | 0.325 | 7260 | tags=13%, list=12%, signal=14% |
| 1474 | CENTRAL\_NERVOUS\_SYSTEM\_PROJECTION\_NEURON\_AXONOGENESIS(GO:0021952) |  | 24 | 0.83 | 1.30 | 0.000 | 0.524 | 0.325 | 7081 | tags=21%, list=12%, signal=24% |
| 1475 | CEREBRAL\_CORTEX\_RADIAL\_GLIA\_GUIDED\_MIGRATION(GO:0021801) |  | 23 | 0.89 | 1.30 | 0.141 | 0.524 | 0.325 | 6328 | tags=17%, list=11%, signal=19% |
| 1476 | TELENCEPHALON\_GLIAL\_CELL\_MIGRATION(GO:0022030) |  | 23 | 0.89 | 1.30 | 0.141 | 0.523 | 0.325 | 6328 | tags=17%, list=11%, signal=19% |
| 1477 | REPLACEMENT\_BONE\_MORPHOGENESIS(GO:0061971) |  | 51 | 0.82 | 1.30 | 0.000 | 0.524 | 0.325 | 7322 | tags=25%, list=12%, signal=29% |
| 1478 | ENDOCHONDRAL\_BONE\_MORPHOGENESIS(GO:0060350) |  | 51 | 0.82 | 1.30 | 0.000 | 0.523 | 0.325 | 7322 | tags=25%, list=12%, signal=29% |
| 1479 | NEURON\_RECOGNITION(GO:0008038) |  | 34 | 0.88 | 1.30 | 0.141 | 0.523 | 0.325 | 6000 | tags=15%, list=10%, signal=16% |
| 1480 | ESTABLISHMENT\_OF\_SPINDLE\_ORIENTATION(GO:0051294) |  | 26 | 0.81 | 1.30 | 0.000 | 0.523 | 0.325 | 5892 | tags=12%, list=10%, signal=13% |
| 1481 | AORTA\_DEVELOPMENT(GO:0035904) |  | 54 | 0.81 | 1.30 | 0.000 | 0.523 | 0.325 | 7343 | tags=13%, list=12%, signal=15% |
| 1482 | OXIDOREDUCTASE\_ACTIVITY\_\_ACTING\_ON\_THE\_CH\_OH\_GROUP\_OF\_DONORS\_\_NAD\_OR\_NADP\_AS\_ACCEPTOR(GO:0016616) |  | 120 | 0.82 | 1.30 | 0.141 | 0.523 | 0.325 | 7408 | tags=13%, list=13%, signal=14% |
| 1483 | MUSCLE\_ORGAN\_MORPHOGENESIS(GO:0048644) |  | 84 | 0.80 | 1.30 | 0.000 | 0.523 | 0.325 | 8328 | tags=33%, list=14%, signal=39% |
| 1484 | PENTOSE\_METABOLIC\_PROCESS(GO:0019321) |  | 15 | 0.82 | 1.30 | 0.000 | 0.523 | 0.325 | 5961 | tags=7%, list=10%, signal=7% |
| 1485 | APOPTOTIC\_MITOCHONDRIAL\_CHANGES(GO:0008637) |  | 127 | 0.83 | 1.30 | 0.141 | 0.523 | 0.325 | 6154 | tags=4%, list=10%, signal=4% |
| 1486 | DEFENSE\_RESPONSE(GO:0006952) |  | 1676 | 0.81 | 1.30 | 0.141 | 0.523 | 0.325 | 7488 | tags=12%, list=13%, signal=14% |
| 1487 | MESODERMAL\_CELL\_FATE\_COMMITMENT(GO:0001710) |  | 19 | 0.87 | 1.30 | 0.000 | 0.523 | 0.325 | 6871 | tags=26%, list=12%, signal=30% |
| 1488 | NUCLEOSIDE\_PHOSPHATE\_CATABOLIC\_PROCESS(GO:1901292) |  | 174 | 0.82 | 1.30 | 0.141 | 0.522 | 0.325 | 6696 | tags=8%, list=11%, signal=9% |
| 1489 | MITOCHONDRIAL\_ELECTRON\_TRANSPORT\_\_NADH\_TO\_UBIQUINONE(GO:0006120) |  | 53 | 0.77 | 1.30 | 0.000 | 0.522 | 0.325 | 5894 | tags=6%, list=10%, signal=6% |
| 1490 | GRANULOCYTE\_CHEMOTAXIS(GO:0071621) |  | 107 | 0.85 | 1.30 | 0.141 | 0.522 | 0.325 | 7081 | tags=17%, list=12%, signal=19% |
| 1491 | REGULATION\_OF\_VASCULATURE\_DEVELOPMENT(GO:1901342) |  | 394 | 0.81 | 1.30 | 0.141 | 0.522 | 0.325 | 7267 | tags=19%, list=12%, signal=22% |
| 1492 | TELENCEPHALON\_CELL\_MIGRATION(GO:0022029) |  | 59 | 0.87 | 1.30 | 0.141 | 0.522 | 0.325 | 6338 | tags=17%, list=11%, signal=19% |
| 1493 | PYRIDINE\_CONTAINING\_COMPOUND\_BIOSYNTHETIC\_PROCESS(GO:0072525) |  | 137 | 0.81 | 1.30 | 0.141 | 0.522 | 0.325 | 6696 | tags=6%, list=11%, signal=7% |
| 1494 | POSITIVE\_REGULATION\_OF\_FAT\_CELL\_DIFFERENTIATION(GO:0045600) |  | 60 | 0.80 | 1.30 | 0.000 | 0.522 | 0.325 | 7792 | tags=20%, list=13%, signal=23% |
| 1495 | REGULATION\_OF\_RESPONSE\_TO\_DRUG(GO:2001023) |  | 89 | 0.82 | 1.30 | 0.141 | 0.522 | 0.325 | 7435 | tags=18%, list=13%, signal=21% |
| 1496 | ICOSANOID\_SECRETION(GO:0032309) |  | 45 | 0.85 | 1.30 | 0.141 | 0.522 | 0.325 | 7298 | tags=18%, list=12%, signal=20% |
| 1497 | PLASMA\_MEMBRANE\_PROTEIN\_COMPLEX(GO:0098797) |  | 553 | 0.82 | 1.30 | 0.141 | 0.521 | 0.325 | 7963 | tags=23%, list=13%, signal=26% |
| 1498 | REGULATION\_OF\_SODIUM\_ION\_TRANSMEMBRANE\_TRANSPORT(GO:1902305) |  | 56 | 0.85 | 1.30 | 0.141 | 0.521 | 0.325 | 6608 | tags=27%, list=11%, signal=30% |
| 1499 | POSITIVE\_REGULATION\_OF\_CELL\_COMMUNICATION(GO:0010647) |  | 1696 | 0.82 | 1.30 | 0.141 | 0.521 | 0.325 | 7175 | tags=13%, list=12%, signal=14% |
| 1500 | VESICLE\_MEDIATED\_TRANSPORT(GO:0016192) |  | 1994 | 0.82 | 1.30 | 0.141 | 0.521 | 0.325 | 7467 | tags=9%, list=13%, signal=10% |
| 1501 | ACTIN\_FILAMENT\_BASED\_PROCESS(GO:0030029) |  | 702 | 0.82 | 1.30 | 0.141 | 0.521 | 0.325 | 7180 | tags=16%, list=12%, signal=18% |
| 1502 | AMMONIUM\_TRANSPORT(GO:0015696) |  | 94 | 0.81 | 1.30 | 0.141 | 0.520 | 0.325 | 7298 | tags=22%, list=12%, signal=25% |
| 1503 | HISTONE\_ACETYLTRANSFERASE\_BINDING(GO:0035035) |  | 27 | 0.82 | 1.30 | 0.000 | 0.520 | 0.325 | 6584 | tags=11%, list=11%, signal=12% |
| 1504 | REGULATION\_OF\_PEPTIDASE\_ACTIVITY(GO:0052547) |  | 432 | 0.81 | 1.30 | 0.141 | 0.520 | 0.325 | 7238 | tags=12%, list=12%, signal=14% |
| 1505 | POSITIVE\_REGULATION\_OF\_SIGNALING(GO:0023056) |  | 1702 | 0.82 | 1.30 | 0.141 | 0.520 | 0.325 | 7175 | tags=13%, list=12%, signal=14% |
| 1506 | NEURAL\_TUBE\_PATTERNING(GO:0021532) |  | 41 | 0.86 | 1.30 | 0.000 | 0.520 | 0.325 | 6714 | tags=17%, list=11%, signal=19% |
| 1507 | ENDODERM\_FORMATION(GO:0001706) |  | 55 | 0.83 | 1.30 | 0.000 | 0.520 | 0.325 | 6355 | tags=16%, list=11%, signal=18% |
| 1508 | GLYCOSIDE\_METABOLIC\_PROCESS(GO:0016137) |  | 17 | 0.89 | 1.30 | 0.141 | 0.520 | 0.325 | 5839 | tags=6%, list=10%, signal=7% |
| 1509 | NUCLEAR\_TRANSCRIBED\_MRNA\_POLY(A)\_TAIL\_SHORTENING(GO:0000289) |  | 40 | 0.84 | 1.30 | 0.141 | 0.520 | 0.325 | 6508 | tags=8%, list=11%, signal=8% |
| 1510 | RESPONSE\_TO\_INACTIVITY(GO:0014854) |  | 15 | 0.87 | 1.30 | 0.000 | 0.520 | 0.325 | 6314 | tags=13%, list=11%, signal=15% |
| 1511 | STEM\_CELL\_DIFFERENTIATION(GO:0048863) |  | 261 | 0.82 | 1.30 | 0.141 | 0.520 | 0.325 | 7360 | tags=16%, list=12%, signal=19% |
| 1512 | REGULATION\_OF\_PROTEIN\_KINASE\_B\_SIGNALING(GO:0051896) |  | 221 | 0.82 | 1.30 | 0.141 | 0.519 | 0.325 | 7754 | tags=17%, list=13%, signal=19% |
| 1513 | SECOND\_MESSENGER\_MEDIATED\_SIGNALING(GO:0019932) |  | 330 | 0.80 | 1.30 | 0.141 | 0.519 | 0.325 | 7544 | tags=21%, list=13%, signal=24% |
| 1514 | FATTY\_ACID\_TRANSPORT(GO:0015908) |  | 93 | 0.83 | 1.30 | 0.141 | 0.519 | 0.325 | 7699 | tags=17%, list=13%, signal=20% |
| 1515 | POSITIVE\_REGULATION\_OF\_GENE\_EXPRESSION(GO:0010628) |  | 1866 | 0.82 | 1.30 | 0.000 | 0.519 | 0.325 | 7450 | tags=14%, list=13%, signal=15% |
| 1516 | DYNEIN\_COMPLEX\_BINDING(GO:0070840) |  | 24 | 0.81 | 1.30 | 0.000 | 0.519 | 0.325 | 7023 | tags=13%, list=12%, signal=14% |
| 1517 | REGULATION\_OF\_ORGAN\_MORPHOGENESIS(GO:2000027) |  | 251 | 0.82 | 1.30 | 0.141 | 0.518 | 0.325 | 7537 | tags=15%, list=13%, signal=17% |
| 1518 | REGULATION\_OF\_CALCIUM\_ION\_TRANSPORT\_INTO\_CYTOSOL(GO:0010522) |  | 99 | 0.83 | 1.30 | 0.000 | 0.518 | 0.325 | 8198 | tags=27%, list=14%, signal=32% |
| 1519 | ESTABLISHMENT\_OF\_PLANAR\_POLARITY\_INVOLVED\_IN\_NEURAL\_TUBE\_CLOSURE(GO:0090177) |  | 15 | 0.84 | 1.30 | 0.000 | 0.518 | 0.325 | 6473 | tags=20%, list=11%, signal=22% |
| 1520 | REGULATION\_OF\_ION\_TRANSPORT(GO:0043269) |  | 620 | 0.82 | 1.30 | 0.141 | 0.518 | 0.325 | 8109 | tags=25%, list=14%, signal=28% |
| 1521 | REGULATION\_OF\_RELEASE\_OF\_SEQUESTERED\_CALCIUM\_ION\_INTO\_CYTOSOL(GO:0051279) |  | 78 | 0.83 | 1.30 | 0.000 | 0.518 | 0.325 | 7544 | tags=22%, list=13%, signal=25% |
| 1522 | EXOCRINE\_SYSTEM\_DEVELOPMENT(GO:0035272) |  | 49 | 0.84 | 1.30 | 0.141 | 0.518 | 0.325 | 7343 | tags=24%, list=12%, signal=28% |
| 1523 | NEGATIVE\_REGULATION\_OF\_B\_CELL\_PROLIFERATION(GO:0030889) |  | 17 | 0.78 | 1.30 | 0.000 | 0.518 | 0.325 | 10693 | tags=29%, list=18%, signal=36% |
| 1524 | REGULATION\_OF\_NEURON\_DEATH(GO:1901214) |  | 284 | 0.80 | 1.30 | 0.000 | 0.518 | 0.325 | 7963 | tags=19%, list=13%, signal=21% |
| 1525 | NEGATIVE\_REGULATION\_OF\_RESPONSE\_TO\_STIMULUS(GO:0048585) |  | 1546 | 0.82 | 1.30 | 0.141 | 0.518 | 0.325 | 7670 | tags=14%, list=13%, signal=16% |
| 1526 | EMBRYONIC\_FORELIMB\_MORPHOGENESIS(GO:0035115) |  | 32 | 0.86 | 1.30 | 0.141 | 0.518 | 0.325 | 6395 | tags=31%, list=11%, signal=35% |
| 1527 | POSITIVE\_REGULATION\_OF\_DEVELOPMENTAL\_GROWTH(GO:0048639) |  | 180 | 0.82 | 1.30 | 0.141 | 0.518 | 0.325 | 7537 | tags=19%, list=13%, signal=22% |
| 1528 | REGULATION\_OF\_SMOOTHENED\_SIGNALING\_PATHWAY(GO:0008589) |  | 76 | 0.85 | 1.30 | 0.000 | 0.517 | 0.325 | 7546 | tags=30%, list=13%, signal=35% |
| 1529 | NEPHRON\_MORPHOGENESIS(GO:0072028) |  | 77 | 0.86 | 1.30 | 0.141 | 0.517 | 0.325 | 7382 | tags=29%, list=12%, signal=33% |
| 1530 | MOTOR\_NEURON\_AXON\_GUIDANCE(GO:0008045) |  | 23 | 0.83 | 1.30 | 0.000 | 0.517 | 0.325 | 8305 | tags=35%, list=14%, signal=40% |
| 1531 | NEGATIVE\_REGULATION\_OF\_G\_PROTEIN\_COUPLED\_RECEPTOR\_PROTEIN\_SIGNALING\_PATHWAY(GO:0045744) |  | 47 | 0.78 | 1.30 | 0.000 | 0.517 | 0.325 | 7441 | tags=23%, list=13%, signal=27% |
| 1532 | HOMEOSTATIC\_PROCESS(GO:0042592) |  | 1683 | 0.82 | 1.30 | 0.141 | 0.517 | 0.325 | 7547 | tags=15%, list=13%, signal=17% |
| 1533 | POSITIVE\_REGULATION\_OF\_ANGIOGENESIS(GO:0045766) |  | 193 | 0.82 | 1.30 | 0.141 | 0.517 | 0.325 | 7267 | tags=21%, list=12%, signal=24% |
| 1534 | POLY\_PURINE\_TRACT\_BINDING(GO:0070717) |  | 23 | 0.79 | 1.30 | 0.000 | 0.517 | 0.325 | 5932 | tags=4%, list=10%, signal=5% |
| 1535 | REGULATION\_OF\_POTASSIUM\_ION\_TRANSMEMBRANE\_TRANSPORT(GO:1901379) |  | 81 | 0.80 | 1.30 | 0.000 | 0.516 | 0.325 | 9608 | tags=30%, list=16%, signal=35% |
| 1536 | POSITIVE\_REGULATION\_OF\_INTERLEUKIN\_12\_PRODUCTION(GO:0032735) |  | 32 | 0.83 | 1.30 | 0.141 | 0.516 | 0.325 | 6178 | tags=16%, list=10%, signal=17% |
| 1537 | NEPHRON\_TUBULE\_MORPHOGENESIS(GO:0072078) |  | 73 | 0.86 | 1.30 | 0.141 | 0.516 | 0.325 | 7382 | tags=29%, list=12%, signal=33% |
| 1538 | FATTY\_ACID\_DERIVATIVE\_TRANSPORT(GO:1901571) |  | 49 | 0.85 | 1.30 | 0.141 | 0.516 | 0.325 | 7298 | tags=16%, list=12%, signal=19% |
| 1539 | ICOSANOID\_TRANSPORT(GO:0071715) |  | 49 | 0.85 | 1.30 | 0.141 | 0.516 | 0.325 | 7298 | tags=16%, list=12%, signal=19% |
| 1540 | NEURON\_PROJECTION\_ARBORIZATION(GO:0140058) |  | 16 | 0.81 | 1.30 | 0.000 | 0.515 | 0.325 | 7426 | tags=31%, list=13%, signal=36% |
| 1541 | EMBRYO\_DEVELOPMENT\_ENDING\_IN\_BIRTH\_OR\_EGG\_HATCHING(GO:0009792) |  | 591 | 0.83 | 1.30 | 0.141 | 0.515 | 0.325 | 7374 | tags=18%, list=12%, signal=20% |
| 1542 | REGULATION\_OF\_CARTILAGE\_DEVELOPMENT(GO:0061035) |  | 65 | 0.82 | 1.30 | 0.000 | 0.515 | 0.325 | 7913 | tags=34%, list=13%, signal=39% |
| 1543 | CELLULAR\_COMPONENT\_ASSEMBLY\_INVOLVED\_IN\_MORPHOGENESIS(GO:0010927) |  | 104 | 0.84 | 1.30 | 0.141 | 0.515 | 0.325 | 6617 | tags=22%, list=11%, signal=25% |
| 1544 | ACID\_SECRETION(GO:0046717) |  | 122 | 0.83 | 1.30 | 0.141 | 0.515 | 0.325 | 7397 | tags=20%, list=13%, signal=23% |
| 1545 | CHORDATE\_EMBRYONIC\_DEVELOPMENT(GO:0043009) |  | 572 | 0.83 | 1.30 | 0.141 | 0.515 | 0.325 | 7374 | tags=18%, list=12%, signal=20% |
| 1546 | NERVOUS\_SYSTEM\_PROCESS(GO:0050877) |  | 1281 | 0.81 | 1.30 | 0.141 | 0.515 | 0.325 | 7962 | tags=18%, list=13%, signal=20% |
| 1547 | HEART\_GROWTH(GO:0060419) |  | 102 | 0.80 | 1.30 | 0.141 | 0.514 | 0.325 | 8380 | tags=25%, list=14%, signal=30% |
| 1548 | PROTEINACEOUS\_EXTRACELLULAR\_MATRIX(GO:0005578) |  | 363 | 0.84 | 1.30 | 0.141 | 0.514 | 0.325 | 7322 | tags=23%, list=12%, signal=26% |
| 1549 | AMINO\_ACID\_TRANSPORT(GO:0006865) |  | 137 | 0.78 | 1.30 | 0.000 | 0.514 | 0.325 | 8074 | tags=23%, list=14%, signal=27% |
| 1550 | CELLULAR\_CATION\_HOMEOSTASIS(GO:0030003) |  | 609 | 0.79 | 1.30 | 0.141 | 0.514 | 0.325 | 7544 | tags=18%, list=13%, signal=20% |
| 1551 | LEUKOCYTE\_CHEMOTAXIS(GO:0030595) |  | 201 | 0.83 | 1.30 | 0.141 | 0.514 | 0.325 | 7081 | tags=19%, list=12%, signal=21% |
| 1552 | MUSCLE\_TISSUE\_MORPHOGENESIS(GO:0060415) |  | 77 | 0.81 | 1.30 | 0.000 | 0.514 | 0.325 | 8328 | tags=34%, list=14%, signal=39% |
| 1553 | PROTEIN\_KINASE\_A\_SIGNALING(GO:0010737) |  | 35 | 0.79 | 1.30 | 0.000 | 0.514 | 0.325 | 8485 | tags=29%, list=14%, signal=33% |
| 1554 | REGULATION\_OF\_VASCULAR\_ENDOTHELIAL\_GROWTH\_FACTOR\_RECEPTOR\_SIGNALING\_PATHWAY(GO:0030947) |  | 32 | 0.83 | 1.30 | 0.000 | 0.514 | 0.325 | 5908 | tags=13%, list=10%, signal=14% |
| 1555 | NEGATIVE\_REGULATION\_OF\_DEVELOPMENTAL\_PROCESS(GO:0051093) |  | 967 | 0.82 | 1.30 | 0.141 | 0.513 | 0.325 | 7461 | tags=17%, list=13%, signal=19% |
| 1556 | ICOSANOID\_BIOSYNTHETIC\_PROCESS(GO:0046456) |  | 46 | 0.83 | 1.30 | 0.141 | 0.513 | 0.325 | 7682 | tags=17%, list=13%, signal=20% |
| 1557 | REGULATION\_OF\_ORGANELLE\_ORGANIZATION(GO:0033043) |  | 1220 | 0.82 | 1.30 | 0.141 | 0.513 | 0.325 | 6654 | tags=7%, list=11%, signal=7% |
| 1558 | ACTIN\_MONOMER\_BINDING(GO:0003785) |  | 28 | 0.79 | 1.30 | 0.000 | 0.513 | 0.325 | 8787 | tags=25%, list=15%, signal=29% |
| 1559 | ACID\_THIOL\_LIGASE\_ACTIVITY(GO:0016878) |  | 30 | 0.79 | 1.30 | 0.000 | 0.513 | 0.325 | 6782 | tags=10%, list=11%, signal=11% |
| 1560 | SMAD\_BINDING(GO:0046332) |  | 75 | 0.78 | 1.30 | 0.000 | 0.513 | 0.325 | 7907 | tags=13%, list=13%, signal=15% |
| 1561 | COLLAGEN\_BIOSYNTHETIC\_PROCESS(GO:0032964) |  | 47 | 0.80 | 1.30 | 0.000 | 0.513 | 0.325 | 7640 | tags=15%, list=13%, signal=17% |
| 1562 | KERATAN\_SULFATE\_BIOSYNTHETIC\_PROCESS(GO:0018146) |  | 28 | 0.84 | 1.30 | 0.000 | 0.513 | 0.325 | 6985 | tags=29%, list=12%, signal=32% |
| 1563 | NEGATIVE\_REGULATION\_OF\_ENDOTHELIAL\_CELL\_PROLIFERATION(GO:0001937) |  | 68 | 0.82 | 1.30 | 0.141 | 0.513 | 0.325 | 7754 | tags=16%, list=13%, signal=19% |
| 1564 | NEPHRON\_TUBULE\_DEVELOPMENT(GO:0072080) |  | 91 | 0.85 | 1.30 | 0.141 | 0.512 | 0.325 | 7382 | tags=25%, list=12%, signal=29% |
| 1565 | EMBRYONIC\_EYE\_MORPHOGENESIS(GO:0048048) |  | 35 | 0.86 | 1.30 | 0.141 | 0.513 | 0.325 | 6614 | tags=29%, list=11%, signal=32% |
| 1566 | WATER\_SOLUBLE\_VITAMIN\_METABOLIC\_PROCESS(GO:0006767) |  | 87 | 0.80 | 1.30 | 0.000 | 0.513 | 0.325 | 6438 | tags=11%, list=11%, signal=13% |
| 1567 | REGULATION\_OF\_BONE\_REMODELING(GO:0046850) |  | 46 | 0.81 | 1.30 | 0.000 | 0.512 | 0.325 | 8241 | tags=28%, list=14%, signal=33% |
| 1568 | POSITIVE\_REGULATION\_OF\_PHOSPHOPROTEIN\_PHOSPHATASE\_ACTIVITY(GO:0032516) |  | 17 | 0.79 | 1.30 | 0.000 | 0.512 | 0.325 | 9543 | tags=29%, list=16%, signal=35% |
| 1569 | TUMOR\_NECROSIS\_FACTOR\_ACTIVATED\_RECEPTOR\_ACTIVITY(GO:0005031) |  | 24 | 0.87 | 1.30 | 0.141 | 0.512 | 0.325 | 5851 | tags=8%, list=10%, signal=9% |
| 1570 | DEATH\_RECEPTOR\_ACTIVITY(GO:0005035) |  | 24 | 0.87 | 1.30 | 0.141 | 0.512 | 0.325 | 5851 | tags=8%, list=10%, signal=9% |
| 1571 | NEUROTRANSMITTER\_TRANSPORTER\_ACTIVITY(GO:0005326) |  | 55 | 0.82 | 1.30 | 0.000 | 0.512 | 0.325 | 6374 | tags=25%, list=11%, signal=29% |
| 1572 | CELLULAR\_ION\_HOMEOSTASIS(GO:0006873) |  | 624 | 0.79 | 1.30 | 0.141 | 0.511 | 0.325 | 7727 | tags=18%, list=13%, signal=21% |
| 1573 | SYNAPSE(GO:0045202) |  | 846 | 0.81 | 1.30 | 0.141 | 0.512 | 0.325 | 7995 | tags=22%, list=14%, signal=25% |
| 1574 | PURINE\_NUCLEOTIDE\_METABOLIC\_PROCESS(GO:0006163) |  | 553 | 0.78 | 1.30 | 0.000 | 0.511 | 0.325 | 7559 | tags=11%, list=13%, signal=13% |
| 1575 | NEGATIVE\_REGULATION\_OF\_ANGIOGENESIS(GO:0016525) |  | 167 | 0.81 | 1.30 | 0.000 | 0.511 | 0.325 | 7260 | tags=14%, list=12%, signal=16% |
| 1576 | RESPONSE\_TO\_MAGNESIUM\_ION(GO:0032026) |  | 18 | 0.86 | 1.30 | 0.141 | 0.511 | 0.325 | 7217 | tags=22%, list=12%, signal=25% |
| 1577 | REGULATION\_OF\_ORGANIC\_ACID\_TRANSPORT(GO:0032890) |  | 51 | 0.83 | 1.30 | 0.141 | 0.511 | 0.325 | 7516 | tags=20%, list=13%, signal=22% |
| 1578 | EPIBOLY(GO:0090504) |  | 32 | 0.84 | 1.30 | 0.000 | 0.511 | 0.325 | 5877 | tags=13%, list=10%, signal=14% |
| 1579 | NEPHRON\_EPITHELIUM\_MORPHOGENESIS(GO:0072088) |  | 75 | 0.86 | 1.30 | 0.141 | 0.511 | 0.325 | 7382 | tags=28%, list=12%, signal=32% |
| 1580 | DIVALENT\_METAL\_ION\_TRANSPORT(GO:0070838) |  | 461 | 0.80 | 1.30 | 0.141 | 0.510 | 0.325 | 7979 | tags=20%, list=13%, signal=23% |
| 1581 | CELL\_SURFACE\_RECEPTOR\_SIGNALING\_PATHWAY\_INVOLVED\_IN\_CELL\_CELL\_SIGNALING(GO:1905114) |  | 605 | 0.80 | 1.30 | 0.000 | 0.510 | 0.325 | 7635 | tags=18%, list=13%, signal=20% |
| 1582 | CAMP\_MEDIATED\_SIGNALING(GO:0019933) |  | 137 | 0.82 | 1.30 | 0.000 | 0.510 | 0.325 | 7441 | tags=20%, list=13%, signal=23% |
| 1583 | REGULATION\_OF\_SODIUM\_ION\_TRANSPORT(GO:0002028) |  | 83 | 0.84 | 1.30 | 0.141 | 0.510 | 0.325 | 7546 | tags=29%, list=13%, signal=33% |
| 1584 | G\_PROTEIN\_COUPLED\_RECEPTOR\_ACTIVITY(GO:0004930) |  | 844 | 0.81 | 1.30 | 0.141 | 0.510 | 0.325 | 7792 | tags=14%, list=13%, signal=16% |
| 1585 | SYNAPSE\_ASSEMBLY(GO:0007416) |  | 149 | 0.83 | 1.30 | 0.141 | 0.509 | 0.325 | 7442 | tags=23%, list=13%, signal=26% |
| 1586 | PHOTORECEPTOR\_OUTER\_SEGMENT(GO:0001750) |  | 80 | 0.80 | 1.30 | 0.000 | 0.509 | 0.325 | 7947 | tags=20%, list=13%, signal=23% |
| 1587 | PROSTATE\_GLAND\_MORPHOGENESIS(GO:0060512) |  | 28 | 0.85 | 1.30 | 0.141 | 0.509 | 0.325 | 6473 | tags=25%, list=11%, signal=28% |
| 1588 | DIVALENT\_INORGANIC\_CATION\_TRANSPORT(GO:0072511) |  | 465 | 0.80 | 1.30 | 0.141 | 0.509 | 0.325 | 7979 | tags=20%, list=13%, signal=23% |
| 1589 | NEUROGENESIS(GO:0022008) |  | 1523 | 0.83 | 1.30 | 0.141 | 0.509 | 0.325 | 7461 | tags=19%, list=13%, signal=21% |
| 1590 | SMOOTHENED\_SIGNALING\_PATHWAY(GO:0007224) |  | 132 | 0.83 | 1.30 | 0.000 | 0.509 | 0.325 | 7546 | tags=23%, list=13%, signal=27% |
| 1591 | SIGNAL\_TRANSDUCTION\_INVOLVED\_IN\_REGULATION\_OF\_GENE\_EXPRESSION(GO:0023019) |  | 20 | 0.85 | 1.30 | 0.000 | 0.509 | 0.325 | 8177 | tags=50%, list=14%, signal=58% |
| 1592 | MRNA\_5'\_UTR\_BINDING(GO:0048027) |  | 23 | 0.79 | 1.30 | 0.000 | 0.509 | 0.325 | 6397 | tags=13%, list=11%, signal=15% |
| 1593 | VENTRAL\_SPINAL\_CORD\_INTERNEURON\_FATE\_COMMITMENT(GO:0060579) |  | 15 | 0.87 | 1.30 | 0.000 | 0.509 | 0.325 | 7006 | tags=47%, list=12%, signal=53% |
| 1594 | CELL\_FATE\_COMMITMENT\_INVOLVED\_IN\_PATTERN\_SPECIFICATION(GO:0060581) |  | 15 | 0.87 | 1.30 | 0.000 | 0.508 | 0.325 | 7006 | tags=47%, list=12%, signal=53% |
| 1595 | LUNG\_ALVEOLUS\_DEVELOPMENT(GO:0048286) |  | 39 | 0.82 | 1.30 | 0.141 | 0.508 | 0.325 | 6458 | tags=13%, list=11%, signal=14% |
| 1596 | REGULATION\_OF\_CELLULAR\_RESPONSE\_TO\_STRESS(GO:0080135) |  | 665 | 0.83 | 1.30 | 0.141 | 0.508 | 0.325 | 6957 | tags=9%, list=12%, signal=10% |
| 1597 | POSITIVE\_REGULATION\_OF\_CELLULAR\_SENESCENCE(GO:2000774) |  | 17 | 0.87 | 1.30 | 0.141 | 0.508 | 0.325 | 6685 | tags=18%, list=11%, signal=20% |
| 1598 | EMBRYO\_DEVELOPMENT(GO:0009790) |  | 966 | 0.82 | 1.30 | 0.141 | 0.508 | 0.325 | 7639 | tags=20%, list=13%, signal=22% |
| 1599 | MEMBRANE\_ASSEMBLY(GO:0071709) |  | 27 | 0.83 | 1.30 | 0.000 | 0.508 | 0.325 | 8937 | tags=37%, list=15%, signal=44% |
| 1600 | REGULATION\_OF\_RECEPTOR\_BINDING(GO:1900120) |  | 23 | 0.83 | 1.30 | 0.141 | 0.508 | 0.325 | 7975 | tags=39%, list=13%, signal=45% |
| 1601 | BILE\_ACID\_BIOSYNTHETIC\_PROCESS(GO:0006699) |  | 36 | 0.82 | 1.30 | 0.000 | 0.508 | 0.325 | 7491 | tags=28%, list=13%, signal=32% |
| 1602 | NEGATIVE\_REGULATION\_OF\_TRANSLATIONAL\_INITIATION(GO:0045947) |  | 19 | 0.84 | 1.30 | 0.000 | 0.508 | 0.325 | 5902 | tags=5%, list=10%, signal=6% |
| 1603 | PURINE\_NUCLEOSIDE\_DIPHOSPHATE\_METABOLIC\_PROCESS(GO:0009135) |  | 125 | 0.82 | 1.30 | 0.141 | 0.508 | 0.325 | 6006 | tags=5%, list=10%, signal=5% |
| 1604 | PURINE\_RIBONUCLEOSIDE\_DIPHOSPHATE\_METABOLIC\_PROCESS(GO:0009179) |  | 125 | 0.82 | 1.30 | 0.141 | 0.508 | 0.325 | 6006 | tags=5%, list=10%, signal=5% |
| 1605 | RELEASE\_OF\_SEQUESTERED\_CALCIUM\_ION\_INTO\_CYTOSOL(GO:0051209) |  | 113 | 0.82 | 1.30 | 0.000 | 0.508 | 0.325 | 8198 | tags=22%, list=14%, signal=26% |
| 1606 | NEGATIVE\_REGULATION\_OF\_SEQUESTERING\_OF\_CALCIUM\_ION(GO:0051283) |  | 113 | 0.82 | 1.30 | 0.000 | 0.508 | 0.325 | 8198 | tags=22%, list=14%, signal=26% |
| 1607 | RESPONSE\_TO\_CHEMICAL(GO:0042221) |  | 4568 | 0.83 | 1.30 | 0.141 | 0.508 | 0.325 | 7172 | tags=12%, list=12%, signal=13% |
| 1608 | URETERIC\_BUD\_MORPHOGENESIS(GO:0060675) |  | 64 | 0.86 | 1.30 | 0.141 | 0.508 | 0.325 | 6456 | tags=22%, list=11%, signal=25% |
| 1609 | REGULATION\_OF\_BICELLULAR\_TIGHT\_JUNCTION\_ASSEMBLY(GO:2000810) |  | 20 | 0.83 | 1.30 | 0.000 | 0.507 | 0.325 | 6796 | tags=15%, list=11%, signal=17% |
| 1610 | CORONARY\_VASCULATURE\_MORPHOGENESIS(GO:0060977) |  | 17 | 0.83 | 1.30 | 0.000 | 0.507 | 0.325 | 6527 | tags=24%, list=11%, signal=26% |
| 1611 | REGULATION\_OF\_SERINE\_TYPE\_ENDOPEPTIDASE\_ACTIVITY(GO:1900003) |  | 15 | 0.84 | 1.30 | 0.000 | 0.507 | 0.325 | 8252 | tags=40%, list=14%, signal=46% |
| 1612 | REGULATION\_OF\_SERINE\_TYPE\_PEPTIDASE\_ACTIVITY(GO:1902571) |  | 15 | 0.84 | 1.30 | 0.000 | 0.507 | 0.325 | 8252 | tags=40%, list=14%, signal=46% |
| 1613 | ADULT\_HEART\_DEVELOPMENT(GO:0007512) |  | 15 | 0.84 | 1.30 | 0.000 | 0.507 | 0.325 | 7898 | tags=33%, list=13%, signal=38% |
| 1614 | REGULATION\_OF\_CELLULAR\_RESPONSE\_TO\_GROWTH\_FACTOR\_STIMULUS(GO:0090287) |  | 273 | 0.81 | 1.30 | 0.141 | 0.506 | 0.325 | 8177 | tags=22%, list=14%, signal=25% |
| 1615 | PLASMA\_MEMBRANE\_BOUNDED\_CELL\_PROJECTION\_MORPHOGENESIS(GO:0120039) |  | 622 | 0.82 | 1.30 | 0.141 | 0.506 | 0.325 | 7461 | tags=20%, list=13%, signal=23% |
| 1616 | CYSTEINE\_TYPE\_ENDOPEPTIDASE\_INHIBITOR\_ACTIVITY(GO:0004869) |  | 57 | 0.82 | 1.30 | 0.141 | 0.506 | 0.325 | 6838 | tags=11%, list=12%, signal=12% |
| 1617 | TRABECULA\_FORMATION(GO:0060343) |  | 26 | 0.83 | 1.30 | 0.000 | 0.506 | 0.325 | 6527 | tags=38%, list=11%, signal=43% |
| 1618 | CLUSTER\_OF\_ACTIN\_BASED\_CELL\_PROJECTIONS(GO:0098862) |  | 149 | 0.81 | 1.30 | 0.141 | 0.506 | 0.325 | 8119 | tags=19%, list=14%, signal=23% |
| 1619 | POSITIVE\_REGULATION\_OF\_CELL\_MIGRATION\_INVOLVED\_IN\_SPROUTING\_ANGIOGENESIS(GO:0090050) |  | 33 | 0.84 | 1.30 | 0.000 | 0.506 | 0.325 | 7698 | tags=30%, list=13%, signal=35% |
| 1620 | CELL\_PROJECTION\_MORPHOGENESIS(GO:0048858) |  | 625 | 0.82 | 1.30 | 0.141 | 0.506 | 0.325 | 7461 | tags=20%, list=13%, signal=23% |
| 1621 | EMBRYONIC\_ORGAN\_DEVELOPMENT(GO:0048568) |  | 428 | 0.83 | 1.30 | 0.141 | 0.506 | 0.325 | 7792 | tags=24%, list=13%, signal=28% |
| 1622 | REGULATION\_OF\_SYSTEMIC\_ARTERIAL\_BLOOD\_PRESSURE\_BY\_RENIN\_ANGIOTENSIN(GO:0003081) |  | 26 | 0.79 | 1.30 | 0.000 | 0.506 | 0.325 | 7711 | tags=31%, list=13%, signal=35% |
| 1623 | POSITIVE\_REGULATION\_OF\_CILIUM\_ASSEMBLY(GO:0045724) |  | 20 | 0.77 | 1.30 | 0.000 | 0.506 | 0.325 | 8273 | tags=20%, list=14%, signal=23% |
| 1624 | SPINDLE\_LOCALIZATION(GO:0051653) |  | 39 | 0.79 | 1.30 | 0.000 | 0.506 | 0.325 | 6115 | tags=10%, list=10%, signal=11% |
| 1625 | REGULATION\_OF\_MACROPHAGE\_ACTIVATION(GO:0043030) |  | 40 | 0.84 | 1.30 | 0.141 | 0.506 | 0.325 | 7547 | tags=30%, list=13%, signal=34% |
| 1626 | ASYMMETRIC\_CELL\_DIVISION(GO:0008356) |  | 19 | 0.85 | 1.30 | 0.000 | 0.506 | 0.325 | 6227 | tags=21%, list=11%, signal=24% |
| 1627 | REGULATION\_OF\_ENDOPEPTIDASE\_ACTIVITY(GO:0052548) |  | 407 | 0.81 | 1.30 | 0.141 | 0.505 | 0.325 | 7238 | tags=12%, list=12%, signal=13% |
| 1628 | REGULATION\_OF\_MORPHOGENESIS\_OF\_AN\_EPITHELIUM(GO:1905330) |  | 181 | 0.82 | 1.30 | 0.141 | 0.505 | 0.325 | 6984 | tags=11%, list=12%, signal=12% |
| 1629 | REGULATION\_OF\_SEQUESTERING\_OF\_CALCIUM\_ION(GO:0051282) |  | 114 | 0.82 | 1.30 | 0.000 | 0.505 | 0.325 | 8198 | tags=22%, list=14%, signal=25% |
| 1630 | SECRETION\_BY\_CELL(GO:0032940) |  | 1435 | 0.80 | 1.30 | 0.141 | 0.505 | 0.325 | 7659 | tags=14%, list=13%, signal=16% |
| 1631 | ORGANELLE\_ASSEMBLY(GO:0070925) |  | 777 | 0.82 | 1.30 | 0.141 | 0.505 | 0.325 | 7177 | tags=9%, list=12%, signal=10% |
| 1632 | NEGATIVE\_REGULATION\_OF\_OXIDOREDUCTASE\_ACTIVITY(GO:0051354) |  | 29 | 0.82 | 1.30 | 0.000 | 0.505 | 0.325 | 8627 | tags=34%, list=15%, signal=40% |
| 1633 | PHOSPHATIDYLCHOLINE\_ACYL\_CHAIN\_REMODELING(GO:0036151) |  | 27 | 0.86 | 1.30 | 0.141 | 0.505 | 0.325 | 5866 | tags=15%, list=10%, signal=16% |
| 1634 | RESPONSE\_TO\_EXTERNAL\_STIMULUS(GO:0009605) |  | 2277 | 0.82 | 1.30 | 0.141 | 0.504 | 0.325 | 7727 | tags=16%, list=13%, signal=18% |
| 1635 | AXON\_TERMINUS(GO:0043679) |  | 115 | 0.81 | 1.30 | 0.000 | 0.504 | 0.325 | 7969 | tags=23%, list=13%, signal=26% |
| 1636 | REGULATION\_OF\_BINDING(GO:0051098) |  | 349 | 0.79 | 1.30 | 0.000 | 0.504 | 0.325 | 7995 | tags=16%, list=14%, signal=18% |
| 1637 | FATTY\_ACID\_TRANSMEMBRANE\_TRANSPORT(GO:1902001) |  | 16 | 0.88 | 1.30 | 0.141 | 0.504 | 0.325 | 5861 | tags=6%, list=10%, signal=7% |
| 1638 | POSITIVE\_REGULATION\_OF\_PEPTIDYL\_SERINE\_PHOSPHORYLATION(GO:0033138) |  | 104 | 0.77 | 1.30 | 0.000 | 0.504 | 0.325 | 9145 | tags=25%, list=15%, signal=30% |
| 1639 | VASOCONSTRICTION(GO:0042310) |  | 79 | 0.80 | 1.30 | 0.141 | 0.503 | 0.325 | 7845 | tags=29%, list=13%, signal=34% |
| 1640 | GATED\_CHANNEL\_ACTIVITY(GO:0022836) |  | 337 | 0.81 | 1.30 | 0.141 | 0.503 | 0.325 | 7979 | tags=26%, list=13%, signal=29% |
| 1641 | ETHANOL\_METABOLIC\_PROCESS(GO:0006067) |  | 21 | 0.87 | 1.30 | 0.141 | 0.503 | 0.325 | 5920 | tags=19%, list=10%, signal=21% |
| 1642 | PURINE\_NUCLEOTIDE\_TRANSPORT(GO:0015865) |  | 15 | 0.82 | 1.30 | 0.000 | 0.503 | 0.325 | 5959 | tags=7%, list=10%, signal=7% |
| 1643 | CARDIAC\_MUSCLE\_TISSUE\_MORPHOGENESIS(GO:0055008) |  | 64 | 0.79 | 1.30 | 0.000 | 0.503 | 0.325 | 8328 | tags=34%, list=14%, signal=40% |
| 1644 | ESTABLISHMENT\_OF\_MITOCHONDRION\_LOCALIZATION\_\_MICROTUBULE\_MEDIATED(GO:0034643) |  | 19 | 0.80 | 1.30 | 0.000 | 0.503 | 0.325 | 6798 | tags=21%, list=11%, signal=24% |
| 1645 | MITOCHONDRION\_TRANSPORT\_ALONG\_MICROTUBULE(GO:0047497) |  | 19 | 0.80 | 1.30 | 0.000 | 0.502 | 0.325 | 6798 | tags=21%, list=11%, signal=24% |
| 1646 | NUCLEOSIDE\_DIPHOSPHATE\_METABOLIC\_PROCESS(GO:0009132) |  | 146 | 0.81 | 1.30 | 0.141 | 0.502 | 0.325 | 6006 | tags=5%, list=10%, signal=5% |
| 1647 | OXIDOREDUCTASE\_ACTIVITY\_\_ACTING\_ON\_CH\_OH\_GROUP\_OF\_DONORS(GO:0016614) |  | 138 | 0.81 | 1.30 | 0.141 | 0.502 | 0.325 | 7408 | tags=12%, list=13%, signal=14% |
| 1648 | CELL\_WALL\_MACROMOLECULE\_METABOLIC\_PROCESS(GO:0044036) |  | 17 | 0.80 | 1.30 | 0.000 | 0.502 | 0.325 | 6109 | tags=12%, list=10%, signal=13% |
| 1649 | CELL\_WALL\_ORGANIZATION\_OR\_BIOGENESIS(GO:0071554) |  | 17 | 0.80 | 1.30 | 0.000 | 0.502 | 0.325 | 6109 | tags=12%, list=10%, signal=13% |
| 1650 | POSITIVE\_REGULATION\_OF\_VASCULATURE\_DEVELOPMENT(GO:1904018) |  | 215 | 0.82 | 1.30 | 0.141 | 0.502 | 0.325 | 7267 | tags=21%, list=12%, signal=24% |
| 1651 | NEGATIVE\_REGULATION\_OF\_BLOOD\_VESSEL\_DIAMETER(GO:0097756) |  | 87 | 0.80 | 1.30 | 0.141 | 0.502 | 0.325 | 7845 | tags=28%, list=13%, signal=32% |
| 1652 | HETEROPHILIC\_CELL\_CELL\_ADHESION\_VIA\_PLASMA\_MEMBRANE\_CELL\_ADHESION\_MOLECULES(GO:0007157) |  | 51 | 0.82 | 1.30 | 0.141 | 0.501 | 0.325 | 6670 | tags=35%, list=11%, signal=40% |
| 1653 | FACE\_MORPHOGENESIS(GO:0060325) |  | 31 | 0.82 | 1.30 | 0.000 | 0.502 | 0.325 | 8177 | tags=29%, list=14%, signal=34% |
| 1654 | NEURON\_PROJECTION\_MORPHOGENESIS(GO:0048812) |  | 608 | 0.82 | 1.30 | 0.141 | 0.501 | 0.325 | 7461 | tags=20%, list=13%, signal=23% |
| 1655 | PRESYNAPTIC\_ACTIVE\_ZONE(GO:0048786) |  | 33 | 0.80 | 1.30 | 0.000 | 0.501 | 0.325 | 8247 | tags=30%, list=14%, signal=35% |
| 1656 | EMBRYONIC\_CAMERA\_TYPE\_EYE\_DEVELOPMENT(GO:0031076) |  | 38 | 0.86 | 1.30 | 0.141 | 0.501 | 0.325 | 6614 | tags=24%, list=11%, signal=27% |
| 1657 | DOPAMINE\_METABOLIC\_PROCESS(GO:0042417) |  | 36 | 0.85 | 1.30 | 0.141 | 0.501 | 0.325 | 5894 | tags=8%, list=10%, signal=9% |
| 1658 | CELLULAR\_AMIDE\_METABOLIC\_PROCESS(GO:0043603) |  | 1239 | 0.80 | 1.30 | 0.000 | 0.501 | 0.325 | 6695 | tags=6%, list=11%, signal=7% |
| 1659 | REGULATION\_OF\_CELLULAR\_KETONE\_METABOLIC\_PROCESS(GO:0010565) |  | 168 | 0.78 | 1.30 | 0.000 | 0.500 | 0.325 | 7699 | tags=12%, list=13%, signal=14% |
| 1660 | LYSOSOME\_ORGANIZATION(GO:0007040) |  | 56 | 0.78 | 1.30 | 0.000 | 0.500 | 0.325 | 6008 | tags=7%, list=10%, signal=8% |
| 1661 | LYTIC\_VACUOLE\_ORGANIZATION(GO:0080171) |  | 56 | 0.78 | 1.30 | 0.000 | 0.500 | 0.325 | 6008 | tags=7%, list=10%, signal=8% |
| 1662 | SIALYLATION(GO:0097503) |  | 22 | 0.86 | 1.30 | 0.000 | 0.500 | 0.325 | 6848 | tags=36%, list=12%, signal=41% |
| 1663 | SIALYLTRANSFERASE\_ACTIVITY(GO:0008373) |  | 22 | 0.86 | 1.30 | 0.000 | 0.500 | 0.325 | 6848 | tags=36%, list=12%, signal=41% |
| 1664 | REGULATION\_OF\_CELLULAR\_RESPONSE\_TO\_DRUG(GO:2001038) |  | 36 | 0.83 | 1.30 | 0.141 | 0.500 | 0.325 | 5890 | tags=6%, list=10%, signal=6% |
| 1665 | MITOTIC\_G2\_DNA\_DAMAGE\_CHECKPOINT(GO:0007095) |  | 23 | 0.79 | 1.30 | 0.000 | 0.500 | 0.325 | 7547 | tags=13%, list=13%, signal=15% |
| 1666 | PHOSPHATIDYLINOSITOL\_ACYL\_CHAIN\_REMODELING(GO:0036149) |  | 16 | 0.89 | 1.30 | 0.141 | 0.499 | 0.325 | 5866 | tags=25%, list=10%, signal=28% |
| 1667 | EXTRACELLULAR\_MATRIX(GO:0031012) |  | 466 | 0.84 | 1.30 | 0.141 | 0.499 | 0.325 | 7322 | tags=20%, list=12%, signal=23% |
| 1668 | TRANSLATION\_REGULATOR\_ACTIVITY(GO:0045182) |  | 48 | 0.78 | 1.30 | 0.000 | 0.499 | 0.325 | 9105 | tags=19%, list=15%, signal=22% |
| 1669 | CATION:CATION\_ANTIPORTER\_ACTIVITY(GO:0015491) |  | 30 | 0.83 | 1.30 | 0.000 | 0.499 | 0.325 | 7809 | tags=30%, list=13%, signal=35% |
| 1670 | RIBONUCLEOSIDE\_DIPHOSPHATE\_METABOLIC\_PROCESS(GO:0009185) |  | 127 | 0.81 | 1.30 | 0.141 | 0.499 | 0.325 | 6006 | tags=5%, list=10%, signal=5% |
| 1671 | CALCIUM\_DEPENDENT\_PHOSPHOLIPID\_BINDING(GO:0005544) |  | 47 | 0.83 | 1.30 | 0.141 | 0.499 | 0.325 | 6702 | tags=15%, list=11%, signal=17% |
| 1672 | SEQUESTERING\_OF\_CALCIUM\_ION(GO:0051208) |  | 117 | 0.82 | 1.30 | 0.000 | 0.499 | 0.325 | 8198 | tags=21%, list=14%, signal=25% |
| 1673 | MAMMARY\_GLAND\_DUCT\_MORPHOGENESIS(GO:0060603) |  | 32 | 0.84 | 1.30 | 0.141 | 0.498 | 0.325 | 7149 | tags=16%, list=12%, signal=18% |
| 1674 | SECONDARY\_METABOLIC\_PROCESS(GO:0019748) |  | 62 | 0.85 | 1.29 | 0.141 | 0.498 | 0.325 | 7172 | tags=19%, list=12%, signal=22% |
| 1675 | REGULATION\_OF\_MORPHOGENESIS\_OF\_A\_BRANCHING\_STRUCTURE(GO:0060688) |  | 55 | 0.86 | 1.29 | 0.141 | 0.498 | 0.325 | 6918 | tags=25%, list=12%, signal=29% |
| 1676 | POSITIVE\_REGULATION\_OF\_CELL\_CYCLE\_PHASE\_TRANSITION(GO:1901989) |  | 81 | 0.77 | 1.29 | 0.000 | 0.498 | 0.325 | 8638 | tags=14%, list=15%, signal=16% |
| 1677 | REGULATION\_OF\_INTERFERON\_GAMMA\_PRODUCTION(GO:0032649) |  | 91 | 0.81 | 1.29 | 0.141 | 0.498 | 0.325 | 7258 | tags=19%, list=12%, signal=21% |
| 1678 | MESONEPHRIC\_TUBULE\_MORPHOGENESIS(GO:0072171) |  | 65 | 0.86 | 1.29 | 0.141 | 0.498 | 0.325 | 6456 | tags=22%, list=11%, signal=24% |
| 1679 | INTERLEUKIN\_6\_MEDIATED\_SIGNALING\_PATHWAY(GO:0070102) |  | 18 | 0.86 | 1.29 | 0.000 | 0.498 | 0.325 | 5858 | tags=6%, list=10%, signal=6% |
| 1680 | REGULATION\_OF\_DEVELOPMENTAL\_PROCESS(GO:0050793) |  | 2534 | 0.83 | 1.29 | 0.141 | 0.498 | 0.325 | 7489 | tags=16%, list=13%, signal=17% |
| 1681 | NUCLEOBASE\_CONTAINING\_SMALL\_MOLECULE\_BIOSYNTHETIC\_PROCESS(GO:0034404) |  | 217 | 0.80 | 1.29 | 0.141 | 0.498 | 0.325 | 6696 | tags=8%, list=11%, signal=9% |
| 1682 | SULFATE\_TRANSMEMBRANE\_TRANSPORT(GO:1902358) |  | 15 | 0.80 | 1.29 | 0.000 | 0.498 | 0.325 | 9957 | tags=53%, list=17%, signal=64% |
| 1683 | SULFATE\_TRANSMEMBRANE\_TRANSPORTER\_ACTIVITY(GO:0015116) |  | 15 | 0.80 | 1.29 | 0.000 | 0.497 | 0.325 | 9957 | tags=53%, list=17%, signal=64% |
| 1684 | MEMORY(GO:0007613) |  | 108 | 0.82 | 1.29 | 0.141 | 0.497 | 0.325 | 6970 | tags=15%, list=12%, signal=17% |
| 1685 | PURINE\_CONTAINING\_COMPOUND\_METABOLIC\_PROCESS(GO:0072521) |  | 588 | 0.78 | 1.29 | 0.000 | 0.497 | 0.325 | 7559 | tags=12%, list=13%, signal=13% |
| 1686 | NEGATIVE\_REGULATION\_OF\_SECRETION\_BY\_CELL(GO:1903531) |  | 181 | 0.78 | 1.29 | 0.141 | 0.497 | 0.325 | 7640 | tags=16%, list=13%, signal=18% |
| 1687 | NEGATIVE\_REGULATION\_OF\_EPIDERMIS\_DEVELOPMENT(GO:0045683) |  | 17 | 0.89 | 1.29 | 0.141 | 0.497 | 0.325 | 5833 | tags=6%, list=10%, signal=7% |
| 1688 | DNA\_DAMAGE\_RESPONSE\_\_DETECTION\_OF\_DNA\_DAMAGE(GO:0042769) |  | 39 | 0.76 | 1.29 | 0.000 | 0.497 | 0.325 | 8258 | tags=15%, list=14%, signal=18% |
| 1689 | REGULATION\_OF\_PROTEIN\_ACETYLATION(GO:1901983) |  | 68 | 0.80 | 1.29 | 0.000 | 0.497 | 0.325 | 7029 | tags=12%, list=12%, signal=13% |
| 1690 | CYCLIC\_NUCLEOTIDE\_CATABOLIC\_PROCESS(GO:0009214) |  | 22 | 0.76 | 1.29 | 0.000 | 0.496 | 0.325 | 9361 | tags=45%, list=16%, signal=54% |
| 1691 | ANATOMICAL\_STRUCTURE\_MORPHOGENESIS(GO:0009653) |  | 2611 | 0.83 | 1.29 | 0.141 | 0.496 | 0.325 | 7547 | tags=18%, list=13%, signal=20% |
| 1692 | CELL\_PART\_MORPHOGENESIS(GO:0032990) |  | 644 | 0.82 | 1.29 | 0.141 | 0.496 | 0.325 | 7461 | tags=19%, list=13%, signal=22% |
| 1693 | SYSTEM\_PROCESS(GO:0003008) |  | 2045 | 0.81 | 1.29 | 0.141 | 0.496 | 0.325 | 7981 | tags=19%, list=13%, signal=22% |
| 1694 | REGULATION\_OF\_DNA\_BINDING\_TRANSCRIPTION\_FACTOR\_ACTIVITY(GO:0051090) |  | 404 | 0.79 | 1.29 | 0.141 | 0.496 | 0.325 | 7158 | tags=12%, list=12%, signal=14% |
| 1695 | SODIUM\_ION\_TRANSPORT(GO:0006814) |  | 222 | 0.82 | 1.29 | 0.141 | 0.496 | 0.325 | 7642 | tags=26%, list=13%, signal=29% |
| 1696 | POSITIVE\_REGULATION\_OF\_INFLAMMATORY\_RESPONSE(GO:0050729) |  | 117 | 0.79 | 1.29 | 0.141 | 0.496 | 0.325 | 6473 | tags=15%, list=11%, signal=17% |
| 1697 | PROTEIN\_KINASE\_B\_SIGNALING(GO:0043491) |  | 255 | 0.81 | 1.29 | 0.141 | 0.495 | 0.325 | 7754 | tags=16%, list=13%, signal=19% |
| 1698 | 3'\_5'\_CYCLIC\_NUCLEOTIDE\_PHOSPHODIESTERASE\_ACTIVITY(GO:0004114) |  | 27 | 0.77 | 1.29 | 0.000 | 0.495 | 0.325 | 7935 | tags=44%, list=13%, signal=51% |
| 1699 | I\_BAND(GO:0031674) |  | 137 | 0.82 | 1.29 | 0.141 | 0.495 | 0.325 | 7739 | tags=26%, list=13%, signal=30% |
| 1700 | SCHWANN\_CELL\_DEVELOPMENT(GO:0014044) |  | 27 | 0.81 | 1.29 | 0.141 | 0.495 | 0.325 | 6421 | tags=7%, list=11%, signal=8% |
| 1701 | ANIMAL\_ORGAN\_DEVELOPMENT(GO:0048513) |  | 3433 | 0.84 | 1.29 | 0.141 | 0.495 | 0.325 | 7180 | tags=15%, list=12%, signal=16% |
| 1702 | MICROTUBULE\_CYTOSKELETON\_ORGANIZATION\_INVOLVED\_IN\_MITOSIS(GO:1902850) |  | 121 | 0.77 | 1.29 | 0.000 | 0.495 | 0.325 | 7983 | tags=10%, list=13%, signal=11% |
| 1703 | ENZYME\_REGULATOR\_ACTIVITY(GO:0030234) |  | 1026 | 0.80 | 1.29 | 0.141 | 0.495 | 0.325 | 7467 | tags=11%, list=13%, signal=12% |
| 1704 | REGULATION\_OF\_MUSCLE\_SYSTEM\_PROCESS(GO:0090257) |  | 241 | 0.79 | 1.29 | 0.000 | 0.495 | 0.325 | 8363 | tags=21%, list=14%, signal=25% |
| 1705 | CELL\_MIGRATION\_INVOLVED\_IN\_HEART\_DEVELOPMENT(GO:0060973) |  | 19 | 0.86 | 1.29 | 0.000 | 0.495 | 0.325 | 7969 | tags=42%, list=13%, signal=49% |
| 1706 | REGULATION\_OF\_EPITHELIAL\_CELL\_DIFFERENTIATION(GO:0030856) |  | 145 | 0.84 | 1.29 | 0.141 | 0.494 | 0.325 | 7360 | tags=12%, list=12%, signal=13% |
| 1707 | NEUROTROPHIN\_TRK\_RECEPTOR\_SIGNALING\_PATHWAY(GO:0048011) |  | 26 | 0.78 | 1.29 | 0.000 | 0.495 | 0.325 | 8823 | tags=27%, list=15%, signal=32% |
| 1708 | DNA\_BINDING\_TRANSCRIPTION\_FACTOR\_ACTIVITY(GO:0003700) |  | 1432 | 0.82 | 1.29 | 0.141 | 0.495 | 0.325 | 7414 | tags=19%, list=13%, signal=21% |
| 1709 | SECRETION(GO:0046903) |  | 1569 | 0.80 | 1.29 | 0.141 | 0.494 | 0.325 | 7659 | tags=15%, list=13%, signal=16% |
| 1710 | ESTABLISHMENT\_OF\_MITOTIC\_SPINDLE\_LOCALIZATION(GO:0040001) |  | 24 | 0.81 | 1.29 | 0.000 | 0.494 | 0.325 | 5892 | tags=4%, list=10%, signal=5% |
| 1711 | RENAL\_SYSTEM\_PROCESS\_INVOLVED\_IN\_REGULATION\_OF\_SYSTEMIC\_ARTERIAL\_BLOOD\_PRESSURE(GO:0003071) |  | 24 | 0.86 | 1.29 | 0.141 | 0.494 | 0.325 | 6525 | tags=29%, list=11%, signal=33% |
| 1712 | CHLORIDE\_CHANNEL\_ACTIVITY(GO:0005254) |  | 81 | 0.82 | 1.29 | 0.141 | 0.494 | 0.325 | 7287 | tags=22%, list=12%, signal=25% |
| 1713 | POSITIVE\_REGULATION\_OF\_CELLULAR\_PROTEIN\_METABOLIC\_PROCESS(GO:0032270) |  | 1502 | 0.81 | 1.29 | 0.141 | 0.494 | 0.325 | 7754 | tags=13%, list=13%, signal=15% |
| 1714 | REGULATION\_OF\_BLOOD\_COAGULATION(GO:0030193) |  | 78 | 0.83 | 1.29 | 0.141 | 0.494 | 0.325 | 7546 | tags=24%, list=13%, signal=28% |
| 1715 | REGULATION\_OF\_HEMOSTASIS(GO:1900046) |  | 78 | 0.83 | 1.29 | 0.141 | 0.494 | 0.325 | 7546 | tags=24%, list=13%, signal=28% |
| 1716 | REGULATION\_OF\_PHOSPHATIDYLINOSITOL\_3\_KINASE\_ACTIVITY(GO:0043551) |  | 46 | 0.82 | 1.29 | 0.141 | 0.494 | 0.325 | 6928 | tags=20%, list=12%, signal=22% |
| 1717 | NEGATIVE\_REGULATION\_OF\_BLOOD\_VESSEL\_MORPHOGENESIS(GO:2000181) |  | 170 | 0.81 | 1.29 | 0.141 | 0.494 | 0.325 | 7754 | tags=15%, list=13%, signal=18% |
| 1718 | ENDOCHONDRAL\_OSSIFICATION(GO:0001958) |  | 26 | 0.84 | 1.29 | 0.000 | 0.494 | 0.325 | 7322 | tags=31%, list=12%, signal=35% |
| 1719 | REPLACEMENT\_OSSIFICATION(GO:0036075) |  | 26 | 0.84 | 1.29 | 0.000 | 0.494 | 0.325 | 7322 | tags=31%, list=12%, signal=35% |
| 1720 | SENSORY\_ORGAN\_MORPHOGENESIS(GO:0090596) |  | 258 | 0.83 | 1.29 | 0.141 | 0.493 | 0.325 | 8177 | tags=29%, list=14%, signal=33% |
| 1721 | NEGATIVE\_REGULATION\_OF\_NERVOUS\_SYSTEM\_DEVELOPMENT(GO:0051961) |  | 274 | 0.82 | 1.29 | 0.141 | 0.493 | 0.325 | 7461 | tags=20%, list=13%, signal=23% |
| 1722 | ION\_GATED\_CHANNEL\_ACTIVITY(GO:0022839) |  | 334 | 0.81 | 1.29 | 0.141 | 0.493 | 0.325 | 7979 | tags=25%, list=13%, signal=29% |
| 1723 | REGULATION\_OF\_MICROTUBULE\_POLYMERIZATION\_OR\_DEPOLYMERIZATION(GO:0031110) |  | 69 | 0.77 | 1.29 | 0.000 | 0.493 | 0.325 | 8112 | tags=16%, list=14%, signal=18% |
| 1724 | MELANOCYTE\_DIFFERENTIATION(GO:0030318) |  | 24 | 0.80 | 1.29 | 0.000 | 0.493 | 0.325 | 7134 | tags=29%, list=12%, signal=33% |
| 1725 | PERIPHERAL\_NERVOUS\_SYSTEM\_AXON\_ENSHEATHMENT(GO:0032292) |  | 24 | 0.82 | 1.29 | 0.141 | 0.493 | 0.325 | 6421 | tags=8%, list=11%, signal=9% |
| 1726 | MYELINATION\_IN\_PERIPHERAL\_NERVOUS\_SYSTEM(GO:0022011) |  | 24 | 0.82 | 1.29 | 0.141 | 0.492 | 0.325 | 6421 | tags=8%, list=11%, signal=9% |
| 1727 | ACTIN\_FILAMENT\_BINDING(GO:0051015) |  | 164 | 0.82 | 1.29 | 0.141 | 0.492 | 0.325 | 8328 | tags=24%, list=14%, signal=28% |
| 1728 | COGNITION(GO:0050890) |  | 277 | 0.80 | 1.29 | 0.000 | 0.492 | 0.325 | 7962 | tags=20%, list=13%, signal=23% |
| 1729 | PROTEIN\_LOCALIZATION\_TO\_NUCLEUS(GO:0034504) |  | 251 | 0.78 | 1.29 | 0.000 | 0.492 | 0.325 | 7184 | tags=9%, list=12%, signal=10% |
| 1730 | SERINE\_FAMILY\_AMINO\_ACID\_METABOLIC\_PROCESS(GO:0009069) |  | 44 | 0.84 | 1.29 | 0.141 | 0.492 | 0.325 | 6628 | tags=14%, list=11%, signal=15% |
| 1731 | Z\_DISC(GO:0030018) |  | 122 | 0.83 | 1.29 | 0.141 | 0.492 | 0.325 | 7180 | tags=23%, list=12%, signal=26% |
| 1732 | REGULATION\_OF\_KIDNEY\_DEVELOPMENT(GO:0090183) |  | 53 | 0.86 | 1.29 | 0.141 | 0.492 | 0.325 | 6279 | tags=11%, list=11%, signal=13% |
| 1733 | MYOSIN\_FILAMENT(GO:0032982) |  | 22 | 0.83 | 1.29 | 0.000 | 0.492 | 0.325 | 7898 | tags=55%, list=13%, signal=63% |
| 1734 | NOTCH\_SIGNALING\_PATHWAY(GO:0007219) |  | 181 | 0.82 | 1.29 | 0.141 | 0.492 | 0.325 | 7367 | tags=9%, list=12%, signal=11% |
| 1735 | MATING(GO:0007618) |  | 38 | 0.83 | 1.29 | 0.141 | 0.491 | 0.325 | 8185 | tags=26%, list=14%, signal=31% |
| 1736 | PROMOTER\_SPECIFIC\_CHROMATIN\_BINDING(GO:1990841) |  | 39 | 0.79 | 1.29 | 0.000 | 0.491 | 0.325 | 7260 | tags=13%, list=12%, signal=15% |
| 1737 | CENTRAL\_NERVOUS\_SYSTEM\_NEURON\_AXONOGENESIS(GO:0021955) |  | 32 | 0.80 | 1.29 | 0.000 | 0.491 | 0.325 | 7081 | tags=16%, list=12%, signal=18% |
| 1738 | AMINO\_ACID\_IMPORT\_ACROSS\_PLASMA\_MEMBRANE(GO:0089718) |  | 16 | 0.81 | 1.29 | 0.141 | 0.491 | 0.325 | 7352 | tags=25%, list=12%, signal=29% |
| 1739 | METALLOENDOPEPTIDASE\_ACTIVITY(GO:0004222) |  | 114 | 0.78 | 1.29 | 0.000 | 0.491 | 0.325 | 7067 | tags=17%, list=12%, signal=19% |
| 1740 | NEGATIVE\_REGULATION\_OF\_MACROMOLECULE\_BIOSYNTHETIC\_PROCESS(GO:0010558) |  | 1669 | 0.82 | 1.29 | 0.141 | 0.491 | 0.325 | 6618 | tags=10%, list=11%, signal=11% |
| 1741 | CATABOLIC\_PROCESS(GO:0009056) |  | 2509 | 0.80 | 1.29 | 0.000 | 0.491 | 0.325 | 6696 | tags=7%, list=11%, signal=7% |
| 1742 | NEGATIVE\_REGULATION\_OF\_NEUROGENESIS(GO:0050768) |  | 255 | 0.82 | 1.29 | 0.141 | 0.491 | 0.325 | 7461 | tags=20%, list=13%, signal=23% |
| 1743 | REGULATION\_OF\_LOCALIZATION(GO:0032879) |  | 2653 | 0.82 | 1.29 | 0.141 | 0.491 | 0.325 | 7546 | tags=15%, list=13%, signal=17% |
| 1744 | PROTEIN\_SERINE\_THREONINE\_KINASE\_ACTIVITY(GO:0004674) |  | 457 | 0.76 | 1.29 | 0.000 | 0.490 | 0.325 | 8112 | tags=15%, list=14%, signal=17% |
| 1745 | REGULATION\_OF\_COAGULATION(GO:0050818) |  | 85 | 0.83 | 1.29 | 0.141 | 0.490 | 0.325 | 7546 | tags=24%, list=13%, signal=27% |
| 1746 | MICROGLIAL\_CELL\_ACTIVATION(GO:0001774) |  | 25 | 0.83 | 1.29 | 0.141 | 0.490 | 0.325 | 8894 | tags=40%, list=15%, signal=47% |
| 1747 | CATION\_CHANNEL\_COMPLEX(GO:0034703) |  | 219 | 0.82 | 1.29 | 0.141 | 0.490 | 0.325 | 7963 | tags=28%, list=13%, signal=33% |
| 1748 | POSITIVE\_REGULATION\_OF\_SPROUTING\_ANGIOGENESIS(GO:1903672) |  | 59 | 0.83 | 1.29 | 0.000 | 0.490 | 0.325 | 7698 | tags=22%, list=13%, signal=25% |
| 1749 | CARBOHYDRATE\_BIOSYNTHETIC\_PROCESS(GO:0016051) |  | 197 | 0.81 | 1.29 | 0.141 | 0.490 | 0.325 | 6848 | tags=11%, list=12%, signal=12% |
| 1750 | CARDIAC\_MUSCLE\_CELL\_DEVELOPMENT(GO:0055013) |  | 72 | 0.80 | 1.29 | 0.000 | 0.490 | 0.325 | 8112 | tags=26%, list=14%, signal=31% |
| 1751 | POSITIVE\_REGULATION\_OF\_LIPID\_METABOLIC\_PROCESS(GO:0045834) |  | 124 | 0.81 | 1.29 | 0.141 | 0.489 | 0.325 | 7281 | tags=19%, list=12%, signal=21% |
| 1752 | PLASMA\_MEMBRANE\_BOUNDED\_CELL\_PROJECTION(GO:0120025) |  | 1894 | 0.82 | 1.29 | 0.141 | 0.489 | 0.325 | 7995 | tags=18%, list=14%, signal=20% |
| 1753 | MONOOXYGENASE\_ACTIVITY(GO:0004497) |  | 97 | 0.80 | 1.29 | 0.141 | 0.489 | 0.325 | 7967 | tags=25%, list=13%, signal=29% |
| 1754 | CELL\_CELL\_ADHERENS\_JUNCTION(GO:0005913) |  | 92 | 0.83 | 1.29 | 0.141 | 0.489 | 0.325 | 7155 | tags=14%, list=12%, signal=16% |
| 1755 | MITOCHONDRIAL\_TRANSMEMBRANE\_TRANSPORT(GO:1990542) |  | 93 | 0.75 | 1.29 | 0.000 | 0.489 | 0.325 | 5861 | tags=2%, list=10%, signal=2% |
| 1756 | ION\_CHANNEL\_BINDING(GO:0044325) |  | 119 | 0.78 | 1.29 | 0.000 | 0.489 | 0.325 | 8452 | tags=24%, list=14%, signal=28% |
| 1757 | REGULATION\_OF\_APOPTOTIC\_PROCESS(GO:0042981) |  | 1528 | 0.82 | 1.29 | 0.141 | 0.489 | 0.325 | 7260 | tags=12%, list=12%, signal=13% |
| 1758 | POSITIVE\_REGULATION\_OF\_PHOSPHOLIPID\_METABOLIC\_PROCESS(GO:1903727) |  | 43 | 0.82 | 1.29 | 0.141 | 0.489 | 0.325 | 8046 | tags=28%, list=14%, signal=32% |
| 1759 | MODIFICATION\_BY\_HOST\_OF\_SYMBIONT\_MORPHOLOGY\_OR\_PHYSIOLOGY(GO:0051851) |  | 69 | 0.77 | 1.29 | 0.000 | 0.488 | 0.325 | 7435 | tags=16%, list=13%, signal=18% |
| 1760 | ESTABLISHMENT\_OF\_PROTEIN\_LOCALIZATION\_TO\_PLASMA\_MEMBRANE(GO:0061951) |  | 42 | 0.82 | 1.29 | 0.141 | 0.488 | 0.325 | 5926 | tags=7%, list=10%, signal=8% |
| 1761 | CARDIAC\_CELL\_DEVELOPMENT(GO:0055006) |  | 78 | 0.80 | 1.29 | 0.000 | 0.488 | 0.325 | 8112 | tags=26%, list=14%, signal=30% |
| 1762 | POSITIVE\_REGULATION\_OF\_CARDIAC\_MUSCLE\_CELL\_PROLIFERATION(GO:0060045) |  | 36 | 0.81 | 1.29 | 0.000 | 0.488 | 0.325 | 8122 | tags=25%, list=14%, signal=29% |
| 1763 | POSITIVE\_REGULATION\_OF\_PROTEIN\_METABOLIC\_PROCESS(GO:0051247) |  | 1592 | 0.81 | 1.29 | 0.141 | 0.488 | 0.325 | 7754 | tags=13%, list=13%, signal=15% |
| 1764 | PHOTORECEPTOR\_CELL\_DIFFERENTIATION(GO:0046530) |  | 59 | 0.84 | 1.29 | 0.141 | 0.488 | 0.325 | 8035 | tags=27%, list=14%, signal=31% |
| 1765 | REGULATION\_OF\_NERVOUS\_SYSTEM\_DEVELOPMENT(GO:0051960) |  | 810 | 0.82 | 1.29 | 0.141 | 0.487 | 0.325 | 7546 | tags=19%, list=13%, signal=21% |
| 1766 | EMBRYONIC\_SKELETAL\_SYSTEM\_DEVELOPMENT(GO:0048706) |  | 123 | 0.85 | 1.29 | 0.141 | 0.487 | 0.325 | 7360 | tags=30%, list=12%, signal=34% |
| 1767 | NEGATIVE\_REGULATION\_OF\_BONE\_REMODELING(GO:0046851) |  | 15 | 0.86 | 1.29 | 0.000 | 0.487 | 0.325 | 7504 | tags=33%, list=13%, signal=38% |
| 1768 | 9+0\_NON\_MOTILE\_CILIUM(GO:0097731) |  | 103 | 0.78 | 1.29 | 0.000 | 0.487 | 0.325 | 9717 | tags=30%, list=16%, signal=36% |
| 1769 | PHOTORECEPTOR\_CELL\_CILIUM(GO:0097733) |  | 103 | 0.78 | 1.29 | 0.000 | 0.487 | 0.325 | 9717 | tags=30%, list=16%, signal=36% |
| 1770 | RESPONSE\_TO\_IONIZING\_RADIATION(GO:0010212) |  | 150 | 0.83 | 1.29 | 0.141 | 0.487 | 0.325 | 6027 | tags=5%, list=10%, signal=5% |
| 1771 | FIBROBLAST\_GROWTH\_FACTOR\_RECEPTOR\_BINDING(GO:0005104) |  | 27 | 0.82 | 1.29 | 0.000 | 0.487 | 0.325 | 7969 | tags=41%, list=13%, signal=47% |
| 1772 | STEROID\_HORMONE\_SECRETION(GO:0035929) |  | 22 | 0.78 | 1.29 | 0.000 | 0.487 | 0.325 | 8485 | tags=23%, list=14%, signal=27% |
| 1773 | POSITIVE\_REGULATION\_OF\_RESPONSE\_TO\_STIMULUS(GO:0048584) |  | 2278 | 0.82 | 1.29 | 0.141 | 0.486 | 0.325 | 7669 | tags=14%, list=13%, signal=15% |
| 1774 | AXONAL\_TRANSPORT(GO:0098930) |  | 40 | 0.79 | 1.29 | 0.000 | 0.486 | 0.325 | 6047 | tags=10%, list=10%, signal=11% |
| 1775 | REGULATION\_OF\_SYNAPSE\_ORGANIZATION(GO:0050807) |  | 158 | 0.81 | 1.29 | 0.141 | 0.486 | 0.325 | 7712 | tags=24%, list=13%, signal=28% |
| 1776 | SKELETAL\_SYSTEM\_DEVELOPMENT(GO:0001501) |  | 491 | 0.81 | 1.29 | 0.141 | 0.486 | 0.325 | 7837 | tags=26%, list=13%, signal=29% |
| 1777 | REGULATION\_OF\_PROGRAMMED\_CELL\_DEATH(GO:0043067) |  | 1544 | 0.81 | 1.29 | 0.141 | 0.486 | 0.325 | 7260 | tags=12%, list=12%, signal=13% |
| 1778 | TRANSMEMBRANE\_RECEPTOR\_PROTEIN\_TYROSINE\_KINASE\_ACTIVITY(GO:0004714) |  | 64 | 0.84 | 1.29 | 0.141 | 0.486 | 0.325 | 7837 | tags=36%, list=13%, signal=41% |
| 1779 | ESTABLISHMENT\_OF\_PLANAR\_POLARITY\_OF\_EMBRYONIC\_EPITHELIUM(GO:0042249) |  | 17 | 0.83 | 1.29 | 0.000 | 0.486 | 0.325 | 6473 | tags=24%, list=11%, signal=26% |
| 1780 | PLASMA\_MEMBRANE\_PART(GO:0044459) |  | 2732 | 0.82 | 1.29 | 0.141 | 0.486 | 0.325 | 7492 | tags=18%, list=13%, signal=19% |
| 1781 | REGULATION\_OF\_POSTSYNAPTIC\_MEMBRANE\_POTENTIAL(GO:0060078) |  | 122 | 0.79 | 1.29 | 0.000 | 0.486 | 0.325 | 7963 | tags=30%, list=13%, signal=34% |
| 1782 | RNA\_POLYMERASE\_II\_DISTAL\_ENHANCER\_SEQUENCE\_SPECIFIC\_DNA\_BINDING(GO:0000980) |  | 71 | 0.78 | 1.29 | 0.000 | 0.485 | 0.325 | 7834 | tags=24%, list=13%, signal=28% |
| 1783 | REGULATION\_OF\_CARDIAC\_MUSCLE\_TISSUE\_GROWTH(GO:0055021) |  | 69 | 0.80 | 1.29 | 0.141 | 0.485 | 0.325 | 8380 | tags=26%, list=14%, signal=30% |
| 1784 | ION\_CHANNEL\_COMPLEX(GO:0034702) |  | 295 | 0.81 | 1.29 | 0.141 | 0.485 | 0.325 | 8420 | tags=29%, list=14%, signal=33% |
| 1785 | CYTOKINE\_BINDING(GO:0019955) |  | 101 | 0.80 | 1.29 | 0.141 | 0.485 | 0.325 | 7921 | tags=24%, list=13%, signal=27% |
| 1786 | ACTIN\_FILAMENT\_BUNDLE(GO:0032432) |  | 59 | 0.80 | 1.29 | 0.000 | 0.485 | 0.325 | 7898 | tags=20%, list=13%, signal=23% |
| 1787 | CELL\_PROJECTION(GO:0042995) |  | 1963 | 0.82 | 1.29 | 0.141 | 0.485 | 0.325 | 7995 | tags=18%, list=14%, signal=20% |
| 1788 | PARAXIAL\_MESODERM\_DEVELOPMENT(GO:0048339) |  | 19 | 0.81 | 1.29 | 0.000 | 0.485 | 0.325 | 8673 | tags=42%, list=15%, signal=49% |
| 1789 | NEURAL\_NUCLEUS\_DEVELOPMENT(GO:0048857) |  | 65 | 0.79 | 1.29 | 0.000 | 0.485 | 0.325 | 7818 | tags=18%, list=13%, signal=21% |
| 1790 | APPENDAGE\_MORPHOGENESIS(GO:0035107) |  | 148 | 0.84 | 1.29 | 0.141 | 0.485 | 0.325 | 7427 | tags=34%, list=13%, signal=39% |
| 1791 | LIMB\_MORPHOGENESIS(GO:0035108) |  | 148 | 0.84 | 1.29 | 0.141 | 0.484 | 0.325 | 7427 | tags=34%, list=13%, signal=39% |
| 1792 | IRON\_ION\_IMPORT(GO:0097286) |  | 15 | 0.80 | 1.29 | 0.141 | 0.484 | 0.325 | 7467 | tags=13%, list=13%, signal=15% |
| 1793 | PHOSPHATIDYLGLYCEROL\_METABOLIC\_PROCESS(GO:0046471) |  | 32 | 0.85 | 1.29 | 0.141 | 0.484 | 0.325 | 5866 | tags=9%, list=10%, signal=10% |
| 1794 | REGULATION\_OF\_PROTON\_TRANSPORT(GO:0010155) |  | 16 | 0.84 | 1.29 | 0.000 | 0.484 | 0.325 | 7298 | tags=19%, list=12%, signal=21% |
| 1795 | SMALL\_MOLECULE\_BIOSYNTHETIC\_PROCESS(GO:0044283) |  | 744 | 0.79 | 1.29 | 0.141 | 0.484 | 0.325 | 7821 | tags=12%, list=13%, signal=14% |
| 1796 | REGULATION\_OF\_PROTEIN\_HOMODIMERIZATION\_ACTIVITY(GO:0043496) |  | 21 | 0.80 | 1.29 | 0.000 | 0.484 | 0.325 | 7134 | tags=19%, list=12%, signal=22% |
| 1797 | REGULATION\_OF\_MESONEPHROS\_DEVELOPMENT(GO:0061217) |  | 26 | 0.88 | 1.29 | 0.141 | 0.484 | 0.325 | 6279 | tags=15%, list=11%, signal=17% |
| 1798 | POSITIVE\_REGULATION\_OF\_PHOSPHATIDYLINOSITOL\_3\_KINASE\_ACTIVITY(GO:0043552) |  | 30 | 0.83 | 1.29 | 0.141 | 0.483 | 0.325 | 6928 | tags=23%, list=12%, signal=26% |
| 1799 | KERATINIZATION(GO:0031424) |  | 224 | 0.89 | 1.29 | 0.141 | 0.484 | 0.325 | 6275 | tags=8%, list=11%, signal=9% |
| 1800 | CALCIUM\_MEDIATED\_SIGNALING(GO:0019722) |  | 169 | 0.79 | 1.29 | 0.141 | 0.483 | 0.325 | 7979 | tags=21%, list=13%, signal=24% |
| 1801 | REGULATION\_OF\_NEUROGENESIS(GO:0050767) |  | 723 | 0.82 | 1.29 | 0.141 | 0.483 | 0.325 | 7546 | tags=19%, list=13%, signal=21% |
| 1802 | NEGATIVE\_REGULATION\_OF\_STEM\_CELL\_PROLIFERATION(GO:2000647) |  | 15 | 0.87 | 1.29 | 0.141 | 0.483 | 0.325 | 6531 | tags=33%, list=11%, signal=37% |
| 1803 | NUCLEOTIDE\_TRANSPORT(GO:0006862) |  | 20 | 0.79 | 1.29 | 0.000 | 0.483 | 0.325 | 5959 | tags=5%, list=10%, signal=6% |
| 1804 | LOCOMOTORY\_BEHAVIOR(GO:0007626) |  | 200 | 0.82 | 1.29 | 0.141 | 0.483 | 0.325 | 7479 | tags=22%, list=13%, signal=25% |
| 1805 | SOLUTE:SODIUM\_SYMPORTER\_ACTIVITY(GO:0015370) |  | 77 | 0.79 | 1.29 | 0.000 | 0.483 | 0.325 | 9451 | tags=38%, list=16%, signal=45% |
| 1806 | UTERUS\_DEVELOPMENT(GO:0060065) |  | 20 | 0.83 | 1.29 | 0.141 | 0.483 | 0.325 | 6473 | tags=15%, list=11%, signal=17% |
| 1807 | FERROUS\_IRON\_BINDING(GO:0008198) |  | 21 | 0.86 | 1.29 | 0.141 | 0.483 | 0.325 | 5894 | tags=10%, list=10%, signal=11% |
| 1808 | LONG\_CHAIN\_FATTY\_ACYL\_COA\_BIOSYNTHETIC\_PROCESS(GO:0035338) |  | 18 | 0.82 | 1.29 | 0.000 | 0.482 | 0.325 | 6042 | tags=11%, list=10%, signal=12% |
| 1809 | NEGATIVE\_REGULATION\_OF\_TRANSCRIPTION\_BY\_RNA\_POLYMERASE\_II(GO:0000122) |  | 799 | 0.82 | 1.29 | 0.141 | 0.482 | 0.325 | 7364 | tags=15%, list=12%, signal=17% |
| 1810 | REGULATION\_OF\_PROTEIN\_LOCALIZATION\_TO\_CELL\_SURFACE(GO:2000008) |  | 34 | 0.80 | 1.29 | 0.000 | 0.482 | 0.325 | 7136 | tags=18%, list=12%, signal=20% |
| 1811 | SOLUTE:CATION\_ANTIPORTER\_ACTIVITY(GO:0015298) |  | 33 | 0.82 | 1.29 | 0.000 | 0.482 | 0.325 | 7809 | tags=27%, list=13%, signal=31% |
| 1812 | DIENCEPHALON\_DEVELOPMENT(GO:0021536) |  | 76 | 0.86 | 1.29 | 0.141 | 0.482 | 0.325 | 6818 | tags=33%, list=12%, signal=37% |
| 1813 | GLYCOPROTEIN\_COMPLEX(GO:0090665) |  | 21 | 0.87 | 1.29 | 0.141 | 0.482 | 0.325 | 5856 | tags=14%, list=10%, signal=16% |
| 1814 | DYSTROPHIN\_ASSOCIATED\_GLYCOPROTEIN\_COMPLEX(GO:0016010) |  | 21 | 0.87 | 1.29 | 0.141 | 0.481 | 0.325 | 5856 | tags=14%, list=10%, signal=16% |
| 1815 | REGULATION\_OF\_TRANSFERASE\_ACTIVITY(GO:0051338) |  | 950 | 0.80 | 1.29 | 0.141 | 0.481 | 0.325 | 7727 | tags=11%, list=13%, signal=13% |
| 1816 | NEGATIVE\_REGULATION\_OF\_GROWTH\_OF\_SYMBIONT\_INVOLVED\_IN\_INTERACTION\_WITH\_HOST(GO:0044146) |  | 15 | 0.86 | 1.29 | 0.000 | 0.481 | 0.325 | 6114 | tags=27%, list=10%, signal=30% |
| 1817 | NEGATIVE\_REGULATION\_OF\_GROWTH\_OF\_SYMBIONT\_IN\_HOST(GO:0044130) |  | 15 | 0.86 | 1.29 | 0.000 | 0.481 | 0.325 | 6114 | tags=27%, list=10%, signal=30% |
| 1818 | NEGATIVE\_REGULATION\_OF\_EPITHELIAL\_CELL\_DIFFERENTIATION(GO:0030857) |  | 45 | 0.87 | 1.29 | 0.141 | 0.481 | 0.325 | 6211 | tags=7%, list=11%, signal=7% |
| 1819 | NEGATIVE\_REGULATION\_OF\_CARDIAC\_MUSCLE\_CELL\_PROLIFERATION(GO:0060044) |  | 20 | 0.83 | 1.29 | 0.000 | 0.481 | 0.325 | 8380 | tags=20%, list=14%, signal=23% |
| 1820 | REGULATION\_OF\_FATTY\_ACID\_BETA\_OXIDATION(GO:0031998) |  | 16 | 0.86 | 1.29 | 0.000 | 0.481 | 0.325 | 5904 | tags=13%, list=10%, signal=14% |
| 1821 | POSITIVE\_REGULATION\_OF\_PHOSPHOLIPASE\_ACTIVITY(GO:0010518) |  | 49 | 0.82 | 1.29 | 0.141 | 0.481 | 0.325 | 6928 | tags=29%, list=12%, signal=32% |
| 1822 | NEURAL\_TUBE\_DEVELOPMENT(GO:0021915) |  | 162 | 0.83 | 1.29 | 0.141 | 0.481 | 0.325 | 7362 | tags=17%, list=12%, signal=20% |
| 1823 | GABA\_RECEPTOR\_COMPLEX(GO:1902710) |  | 18 | 0.88 | 1.29 | 0.000 | 0.481 | 0.325 | 7049 | tags=33%, list=12%, signal=38% |
| 1824 | EXCITATORY\_POSTSYNAPTIC\_POTENTIAL(GO:0060079) |  | 110 | 0.79 | 1.29 | 0.000 | 0.481 | 0.325 | 7963 | tags=30%, list=13%, signal=35% |
| 1825 | STRUCTURAL\_CONSTITUENT\_OF\_MUSCLE(GO:0008307) |  | 46 | 0.86 | 1.29 | 0.141 | 0.480 | 0.325 | 6540 | tags=26%, list=11%, signal=29% |
| 1826 | NUCLEOSOMAL\_DNA\_BINDING(GO:0031492) |  | 46 | 0.80 | 1.29 | 0.000 | 0.480 | 0.325 | 6422 | tags=4%, list=11%, signal=5% |
| 1827 | DEFENSE\_RESPONSE\_TO\_PROTOZOAN(GO:0042832) |  | 20 | 0.81 | 1.29 | 0.141 | 0.481 | 0.325 | 7435 | tags=35%, list=13%, signal=40% |
| 1828 | POSITIVE\_REGULATION\_OF\_CELL\_DIFFERENTIATION(GO:0045597) |  | 922 | 0.81 | 1.29 | 0.141 | 0.481 | 0.325 | 7668 | tags=18%, list=13%, signal=20% |
| 1829 | REGULATION\_OF\_CELLULAR\_PROTEIN\_METABOLIC\_PROCESS(GO:0032268) |  | 2722 | 0.81 | 1.29 | 0.141 | 0.480 | 0.325 | 7238 | tags=10%, list=12%, signal=11% |
| 1830 | METAL\_ION\_HOMEOSTASIS(GO:0055065) |  | 595 | 0.80 | 1.29 | 0.141 | 0.480 | 0.325 | 7544 | tags=17%, list=13%, signal=19% |
| 1831 | POTASSIUM\_ION\_IMPORT(GO:0010107) |  | 28 | 0.78 | 1.29 | 0.000 | 0.480 | 0.325 | 9364 | tags=39%, list=16%, signal=47% |
| 1832 | STEROL\_TRANSPORTER\_ACTIVITY(GO:0015248) |  | 30 | 0.82 | 1.29 | 0.000 | 0.480 | 0.325 | 7172 | tags=27%, list=12%, signal=30% |
| 1833 | T\_CELL\_DIFFERENTIATION\_IN\_THYMUS(GO:0033077) |  | 67 | 0.81 | 1.29 | 0.141 | 0.480 | 0.325 | 7451 | tags=12%, list=13%, signal=14% |
| 1834 | INTERFERON\_GAMMA\_PRODUCTION(GO:0032609) |  | 102 | 0.81 | 1.29 | 0.141 | 0.480 | 0.325 | 7258 | tags=20%, list=12%, signal=22% |
| 1835 | PHOSPHOLIPASE\_A2\_ACTIVITY\_CONSUMING\_1\_2\_DIOLEOYLPHOSPHATIDYLETHANOLAMINE)(GO:0102568) |  | 22 | 0.87 | 1.29 | 0.141 | 0.480 | 0.325 | 5866 | tags=18%, list=10%, signal=20% |
| 1836 | PHOSPHOLIPASE\_A2\_ACTIVITY\_(CONSUMING\_1\_2\_DIPALMITOYLPHOSPHATIDYLCHOLINE)(GO:0102567) |  | 22 | 0.87 | 1.29 | 0.141 | 0.479 | 0.325 | 5866 | tags=18%, list=10%, signal=20% |
| 1837 | TRANSMEMBRANE\_RECEPTOR\_PROTEIN\_KINASE\_ACTIVITY(GO:0019199) |  | 81 | 0.82 | 1.29 | 0.141 | 0.479 | 0.325 | 7837 | tags=33%, list=13%, signal=38% |
| 1838 | HEART\_MORPHOGENESIS(GO:0003007) |  | 239 | 0.82 | 1.29 | 0.141 | 0.479 | 0.325 | 7374 | tags=20%, list=12%, signal=23% |
| 1839 | MAMMARY\_GLAND\_EPITHELIAL\_CELL\_DIFFERENTIATION(GO:0060644) |  | 17 | 0.89 | 1.29 | 0.141 | 0.479 | 0.325 | 6294 | tags=29%, list=11%, signal=33% |
| 1840 | CELL\_CORTEX(GO:0005938) |  | 249 | 0.82 | 1.29 | 0.141 | 0.479 | 0.325 | 6894 | tags=12%, list=12%, signal=14% |
| 1841 | ACIDIC\_AMINO\_ACID\_TRANSPORT(GO:0015800) |  | 61 | 0.79 | 1.29 | 0.000 | 0.479 | 0.325 | 7873 | tags=20%, list=13%, signal=23% |
| 1842 | OSSIFICATION(GO:0001503) |  | 372 | 0.80 | 1.29 | 0.141 | 0.479 | 0.325 | 8046 | tags=23%, list=14%, signal=27% |
| 1843 | TRANSMEMBRANE\_TRANSPORTER\_COMPLEX(GO:1902495) |  | 321 | 0.80 | 1.29 | 0.141 | 0.479 | 0.325 | 8420 | tags=29%, list=14%, signal=34% |
| 1844 | PIGMENT\_CELL\_DIFFERENTIATION(GO:0050931) |  | 33 | 0.77 | 1.29 | 0.000 | 0.478 | 0.325 | 7134 | tags=24%, list=12%, signal=28% |
| 1845 | ENDODERM\_DEVELOPMENT(GO:0007492) |  | 80 | 0.81 | 1.29 | 0.000 | 0.478 | 0.325 | 7343 | tags=16%, list=12%, signal=19% |
| 1846 | NEGATIVE\_REGULATION\_OF\_BIOSYNTHETIC\_PROCESS(GO:0009890) |  | 1768 | 0.81 | 1.29 | 0.141 | 0.478 | 0.325 | 6618 | tags=10%, list=11%, signal=11% |
| 1847 | LIPOSACCHARIDE\_METABOLIC\_PROCESS(GO:1903509) |  | 124 | 0.77 | 1.29 | 0.000 | 0.478 | 0.325 | 8560 | tags=20%, list=14%, signal=24% |
| 1848 | NEUROTRANSMITTER\_UPTAKE(GO:0001504) |  | 30 | 0.85 | 1.29 | 0.141 | 0.478 | 0.325 | 7352 | tags=30%, list=12%, signal=34% |
| 1849 | POSTSYNAPSE\_ORGANIZATION(GO:0099173) |  | 99 | 0.76 | 1.29 | 0.000 | 0.478 | 0.325 | 7962 | tags=19%, list=13%, signal=22% |
| 1850 | MONOCARBOXYLIC\_ACID\_TRANSPORT(GO:0015718) |  | 164 | 0.81 | 1.29 | 0.141 | 0.478 | 0.325 | 7699 | tags=18%, list=13%, signal=21% |
| 1851 | EXTRACELLULAR\_EXOSOME(GO:0070062) |  | 2784 | 0.82 | 1.29 | 0.141 | 0.478 | 0.325 | 7128 | tags=9%, list=12%, signal=10% |
| 1852 | MULTICELLULAR\_ORGANISM\_GROWTH(GO:0035264) |  | 149 | 0.82 | 1.29 | 0.141 | 0.478 | 0.325 | 7547 | tags=13%, list=13%, signal=15% |
| 1853 | CALMODULIN\_BINDING(GO:0005516) |  | 194 | 0.80 | 1.29 | 0.141 | 0.478 | 0.325 | 8417 | tags=27%, list=14%, signal=32% |
| 1854 | PHOSPHATIDYLGLYCEROL\_ACYL\_CHAIN\_REMODELING(GO:0036148) |  | 18 | 0.87 | 1.29 | 0.141 | 0.477 | 0.325 | 5866 | tags=17%, list=10%, signal=18% |
| 1855 | REGULATION\_OF\_SUPRAMOLECULAR\_FIBER\_ORGANIZATION(GO:1902903) |  | 320 | 0.80 | 1.29 | 0.141 | 0.477 | 0.325 | 7973 | tags=13%, list=13%, signal=15% |
| 1856 | REGULATION\_OF\_SYNAPSE\_STRUCTURE\_OR\_ACTIVITY(GO:0050803) |  | 163 | 0.81 | 1.29 | 0.141 | 0.477 | 0.325 | 7712 | tags=23%, list=13%, signal=27% |
| 1857 | CYTOPLASMIC\_VESICLE\_PART(GO:0044433) |  | 1438 | 0.80 | 1.29 | 0.141 | 0.477 | 0.325 | 6738 | tags=9%, list=11%, signal=10% |
| 1858 | EMBRYONIC\_LIMB\_MORPHOGENESIS(GO:0030326) |  | 128 | 0.84 | 1.29 | 0.141 | 0.477 | 0.325 | 7427 | tags=35%, list=13%, signal=40% |
| 1859 | EMBRYONIC\_APPENDAGE\_MORPHOGENESIS(GO:0035113) |  | 128 | 0.84 | 1.29 | 0.141 | 0.477 | 0.325 | 7427 | tags=35%, list=13%, signal=40% |
| 1860 | MULTI\_MULTICELLULAR\_ORGANISM\_PROCESS(GO:0044706) |  | 224 | 0.81 | 1.29 | 0.141 | 0.477 | 0.325 | 7787 | tags=17%, list=13%, signal=19% |
| 1861 | EXTRACELLULAR\_VESICLE(GO:1903561) |  | 2804 | 0.82 | 1.29 | 0.141 | 0.477 | 0.325 | 7128 | tags=9%, list=12%, signal=10% |
| 1862 | PRESYNAPSE(GO:0098793) |  | 358 | 0.79 | 1.29 | 0.141 | 0.477 | 0.325 | 8040 | tags=21%, list=14%, signal=24% |
| 1863 | EXTRACELLULAR\_ORGANELLE(GO:0043230) |  | 2806 | 0.82 | 1.29 | 0.141 | 0.477 | 0.325 | 7128 | tags=9%, list=12%, signal=10% |
| 1864 | BRANCHING\_INVOLVED\_IN\_URETERIC\_BUD\_MORPHOGENESIS(GO:0001658) |  | 59 | 0.87 | 1.29 | 0.141 | 0.477 | 0.325 | 6456 | tags=24%, list=11%, signal=27% |
| 1865 | NEGATIVE\_REGULATION\_OF\_RESPONSE\_TO\_DNA\_DAMAGE\_STIMULUS(GO:2001021) |  | 69 | 0.82 | 1.29 | 0.141 | 0.477 | 0.325 | 6893 | tags=10%, list=12%, signal=11% |
| 1866 | APPENDAGE\_DEVELOPMENT(GO:0048736) |  | 175 | 0.83 | 1.29 | 0.141 | 0.477 | 0.325 | 7427 | tags=31%, list=13%, signal=35% |
| 1867 | LIMB\_DEVELOPMENT(GO:0060173) |  | 175 | 0.83 | 1.29 | 0.141 | 0.477 | 0.325 | 7427 | tags=31%, list=13%, signal=35% |
| 1868 | EMBRYONIC\_SKELETAL\_SYSTEM\_MORPHOGENESIS(GO:0048704) |  | 93 | 0.85 | 1.29 | 0.141 | 0.477 | 0.325 | 7360 | tags=32%, list=12%, signal=37% |
| 1869 | ADENYLATE\_CYCLASE\_MODULATING\_G\_PROTEIN\_COUPLED\_RECEPTOR\_SIGNALING\_PATHWAY(GO:0007188) |  | 189 | 0.82 | 1.29 | 0.141 | 0.476 | 0.325 | 7479 | tags=17%, list=13%, signal=19% |
| 1870 | SODIUM\_CHANNEL\_REGULATOR\_ACTIVITY(GO:0017080) |  | 36 | 0.84 | 1.29 | 0.141 | 0.476 | 0.325 | 6608 | tags=33%, list=11%, signal=38% |
| 1871 | ORGANIC\_SUBSTANCE\_CATABOLIC\_PROCESS(GO:1901575) |  | 2063 | 0.80 | 1.29 | 0.000 | 0.476 | 0.325 | 6696 | tags=7%, list=11%, signal=7% |
| 1872 | FATTY\_ACID\_DERIVATIVE\_BIOSYNTHETIC\_PROCESS(GO:1901570) |  | 89 | 0.79 | 1.29 | 0.141 | 0.476 | 0.325 | 7682 | tags=13%, list=13%, signal=15% |
| 1873 | POSITIVE\_REGULATION\_OF\_CELL\_SUBSTRATE\_ADHESION(GO:0010811) |  | 108 | 0.81 | 1.29 | 0.141 | 0.476 | 0.325 | 8070 | tags=19%, list=14%, signal=22% |
| 1874 | REGULATION\_OF\_DOUBLE\_STRAND\_BREAK\_REPAIR(GO:2000779) |  | 58 | 0.83 | 1.29 | 0.141 | 0.476 | 0.325 | 6910 | tags=12%, list=12%, signal=14% |
| 1875 | CELLULAR\_CATABOLIC\_PROCESS(GO:0044248) |  | 2210 | 0.79 | 1.29 | 0.000 | 0.476 | 0.325 | 6628 | tags=6%, list=11%, signal=7% |
| 1876 | NEGATIVE\_REGULATION\_OF\_JAK\_STAT\_CASCADE(GO:0046426) |  | 52 | 0.80 | 1.29 | 0.141 | 0.476 | 0.325 | 7969 | tags=23%, list=13%, signal=27% |
| 1877 | NEGATIVE\_REGULATION\_OF\_STAT\_CASCADE(GO:1904893) |  | 52 | 0.80 | 1.29 | 0.141 | 0.475 | 0.325 | 7969 | tags=23%, list=13%, signal=27% |
| 1878 | DEVELOPMENTAL\_GROWTH(GO:0048589) |  | 612 | 0.81 | 1.29 | 0.141 | 0.476 | 0.325 | 7583 | tags=19%, list=13%, signal=22% |
| 1879 | RESPONSE\_TO\_FATTY\_ACID(GO:0070542) |  | 86 | 0.81 | 1.29 | 0.000 | 0.475 | 0.325 | 7699 | tags=15%, list=13%, signal=17% |
| 1880 | TRANSPORT\_VESICLE\_MEMBRANE(GO:0030658) |  | 181 | 0.78 | 1.29 | 0.141 | 0.475 | 0.325 | 8063 | tags=17%, list=14%, signal=19% |
| 1881 | ANION\_CHANNEL\_ACTIVITY(GO:0005253) |  | 95 | 0.80 | 1.29 | 0.141 | 0.475 | 0.325 | 8281 | tags=23%, list=14%, signal=27% |
| 1882 | EPITHELIAL\_CELL\_MATURATION(GO:0002070) |  | 16 | 0.87 | 1.29 | 0.141 | 0.475 | 0.325 | 6614 | tags=38%, list=11%, signal=42% |
| 1883 | REGULATION\_OF\_REACTIVE\_OXYGEN\_SPECIES\_BIOSYNTHETIC\_PROCESS(GO:1903426) |  | 80 | 0.80 | 1.29 | 0.000 | 0.475 | 0.325 | 6685 | tags=15%, list=11%, signal=17% |
| 1884 | GLYCOLIPID\_METABOLIC\_PROCESS(GO:0006664) |  | 123 | 0.77 | 1.29 | 0.000 | 0.475 | 0.325 | 8560 | tags=20%, list=14%, signal=23% |
| 1885 | INORGANIC\_CATION\_TRANSMEMBRANE\_TRANSPORT(GO:0098662) |  | 741 | 0.79 | 1.29 | 0.141 | 0.475 | 0.325 | 7979 | tags=20%, list=13%, signal=23% |
| 1886 | NEURON\_DIFFERENTIATION(GO:0030182) |  | 1293 | 0.82 | 1.29 | 0.141 | 0.475 | 0.325 | 7975 | tags=21%, list=13%, signal=24% |
| 1887 | ION\_ANTIPORTER\_ACTIVITY(GO:0099516) |  | 60 | 0.79 | 1.29 | 0.000 | 0.475 | 0.325 | 9293 | tags=35%, list=16%, signal=41% |
| 1888 | ENDODERMAL\_CELL\_DIFFERENTIATION(GO:0035987) |  | 46 | 0.82 | 1.29 | 0.141 | 0.474 | 0.325 | 7099 | tags=17%, list=12%, signal=20% |
| 1889 | CATION\_CHANNEL\_ACTIVITY(GO:0005261) |  | 318 | 0.81 | 1.29 | 0.141 | 0.474 | 0.325 | 7963 | tags=26%, list=13%, signal=30% |
| 1890 | REGULATION\_OF\_LAMELLIPODIUM\_ORGANIZATION(GO:1902743) |  | 45 | 0.81 | 1.29 | 0.000 | 0.474 | 0.325 | 7081 | tags=13%, list=12%, signal=15% |
| 1891 | TRANSCRIPTIONAL\_REPRESSOR\_ACTIVITY\_\_RNA\_POLYMERASE\_II\_TRANSCRIPTION\_REGULATORY\_REGION\_SEQUENCE\_SPECIFIC\_DNA\_BINDING(GO:0001227) |  | 225 | 0.82 | 1.29 | 0.141 | 0.474 | 0.325 | 7364 | tags=23%, list=12%, signal=26% |
| 1892 | HYPOTHALAMUS\_DEVELOPMENT(GO:0021854) |  | 22 | 0.86 | 1.29 | 0.000 | 0.474 | 0.325 | 7313 | tags=41%, list=12%, signal=47% |
| 1893 | NEGATIVE\_REGULATION\_OF\_MYOBLAST\_DIFFERENTIATION(GO:0045662) |  | 24 | 0.83 | 1.29 | 0.000 | 0.474 | 0.325 | 6271 | tags=21%, list=11%, signal=23% |
| 1894 | AXONOGENESIS(GO:0007409) |  | 444 | 0.81 | 1.29 | 0.141 | 0.474 | 0.325 | 7461 | tags=20%, list=13%, signal=23% |
| 1895 | NEGATIVE\_REGULATION\_OF\_RESPONSE\_TO\_DRUG(GO:2001024) |  | 34 | 0.83 | 1.29 | 0.141 | 0.474 | 0.325 | 7426 | tags=12%, list=13%, signal=13% |
| 1896 | MYELOID\_CELL\_DIFFERENTIATION(GO:0030099) |  | 397 | 0.81 | 1.29 | 0.141 | 0.474 | 0.493 | 7159 | tags=13%, list=12%, signal=15% |
| 1897 | MICROTUBULE\_BINDING(GO:0008017) |  | 263 | 0.77 | 1.29 | 0.000 | 0.473 | 0.493 | 7973 | tags=18%, list=13%, signal=21% |
| 1898 | MITOTIC\_SPINDLE\_ORGANIZATION(GO:0007052) |  | 102 | 0.76 | 1.29 | 0.000 | 0.473 | 0.493 | 7948 | tags=10%, list=13%, signal=11% |
| 1899 | REGULATION\_OF\_CELL\_DEVELOPMENT(GO:0060284) |  | 850 | 0.81 | 1.29 | 0.141 | 0.473 | 0.493 | 7670 | tags=19%, list=13%, signal=21% |
| 1900 | NEUROTROPHIN\_SIGNALING\_PATHWAY(GO:0038179) |  | 34 | 0.76 | 1.29 | 0.000 | 0.473 | 0.493 | 8937 | tags=26%, list=15%, signal=31% |
| 1901 | REGULATION\_OF\_MEMBRANE\_DEPOLARIZATION(GO:0003254) |  | 46 | 0.80 | 1.29 | 0.141 | 0.473 | 0.493 | 7907 | tags=30%, list=13%, signal=35% |
| 1902 | AXO\_DENDRITIC\_TRANSPORT(GO:0008088) |  | 48 | 0.78 | 1.29 | 0.000 | 0.473 | 0.493 | 6071 | tags=10%, list=10%, signal=12% |
| 1903 | POSITIVE\_REGULATION\_OF\_TUMOR\_NECROSIS\_FACTOR\_SUPERFAMILY\_CYTOKINE\_PRODUCTION(GO:1903557) |  | 69 | 0.80 | 1.29 | 0.141 | 0.473 | 0.493 | 7461 | tags=16%, list=13%, signal=18% |
| 1904 | NERVOUS\_SYSTEM\_DEVELOPMENT(GO:0007399) |  | 2270 | 0.82 | 1.29 | 0.141 | 0.473 | 0.493 | 7547 | tags=18%, list=13%, signal=20% |
| 1905 | MODULATION\_OF\_EXCITATORY\_POSTSYNAPTIC\_POTENTIAL(GO:0098815) |  | 38 | 0.81 | 1.29 | 0.000 | 0.473 | 0.493 | 7962 | tags=34%, list=13%, signal=40% |
| 1906 | CELL\_AGING(GO:0007569) |  | 111 | 0.80 | 1.29 | 0.141 | 0.473 | 0.493 | 6685 | tags=10%, list=11%, signal=11% |
| 1907 | NEGATIVE\_REGULATION\_OF\_PROTEIN\_LOCALIZATION\_TO\_NUCLEUS(GO:1900181) |  | 25 | 0.83 | 1.29 | 0.000 | 0.473 | 0.493 | 7632 | tags=24%, list=13%, signal=28% |
| 1908 | OUTFLOW\_TRACT\_MORPHOGENESIS(GO:0003151) |  | 73 | 0.84 | 1.29 | 0.141 | 0.472 | 0.493 | 7343 | tags=25%, list=12%, signal=28% |
| 1909 | REGULATION\_OF\_NEURON\_DIFFERENTIATION(GO:0045664) |  | 588 | 0.81 | 1.29 | 0.141 | 0.472 | 0.493 | 7668 | tags=20%, list=13%, signal=22% |
| 1910 | NEUROMUSCULAR\_PROCESS(GO:0050905) |  | 103 | 0.79 | 1.29 | 0.000 | 0.472 | 0.493 | 7962 | tags=24%, list=13%, signal=28% |
| 1911 | CATION\_TRANSMEMBRANE\_TRANSPORT(GO:0098655) |  | 834 | 0.79 | 1.29 | 0.141 | 0.472 | 0.493 | 8217 | tags=21%, list=14%, signal=24% |
| 1912 | LIPID\_DROPLET(GO:0005811) |  | 74 | 0.81 | 1.29 | 0.141 | 0.472 | 0.493 | 7162 | tags=12%, list=12%, signal=14% |
| 1913 | POSITIVE\_REGULATION\_OF\_LIPID\_KINASE\_ACTIVITY(GO:0090218) |  | 33 | 0.83 | 1.29 | 0.141 | 0.472 | 0.493 | 6928 | tags=21%, list=12%, signal=24% |
| 1914 | RESPONSE\_TO\_FLUID\_SHEAR\_STRESS(GO:0034405) |  | 37 | 0.81 | 1.29 | 0.000 | 0.472 | 0.493 | 8241 | tags=22%, list=14%, signal=25% |
| 1915 | POSITIVE\_REGULATION\_OF\_TRANSCRIPTION\_FROM\_RNA\_POLYMERASE\_II\_PROMOTER\_INVOLVED\_IN\_CELLULAR\_RESPONSE\_TO\_CHEMICAL\_STIMULUS(GO:1901522) |  | 23 | 0.80 | 1.29 | 0.000 | 0.472 | 0.493 | 5935 | tags=9%, list=10%, signal=10% |
| 1916 | SODIUM\_CHANNEL\_COMPLEX(GO:0034706) |  | 21 | 0.86 | 1.29 | 0.141 | 0.471 | 0.493 | 6844 | tags=48%, list=12%, signal=54% |
| 1917 | SERINE\_THREONINE\_PROTEIN\_KINASE\_COMPLEX(GO:1902554) |  | 83 | 0.77 | 1.28 | 0.000 | 0.471 | 0.493 | 7154 | tags=6%, list=12%, signal=7% |
| 1918 | CEREBRAL\_CORTEX\_CELL\_MIGRATION(GO:0021795) |  | 45 | 0.87 | 1.28 | 0.141 | 0.471 | 0.493 | 6338 | tags=18%, list=11%, signal=20% |
| 1919 | REGULATION\_OF\_MULTICELLULAR\_ORGANISMAL\_PROCESS(GO:0051239) |  | 2954 | 0.82 | 1.28 | 0.141 | 0.471 | 0.493 | 7546 | tags=18%, list=13%, signal=19% |
| 1920 | TRANSPORTER\_COMPLEX(GO:1990351) |  | 328 | 0.80 | 1.28 | 0.141 | 0.471 | 0.493 | 8420 | tags=28%, list=14%, signal=33% |
| 1921 | SECRETION\_BY\_TISSUE(GO:0032941) |  | 38 | 0.82 | 1.28 | 0.141 | 0.471 | 0.493 | 7038 | tags=26%, list=12%, signal=30% |
| 1922 | NEGATIVE\_REGULATION\_OF\_VASCULATURE\_DEVELOPMENT(GO:1901343) |  | 177 | 0.81 | 1.28 | 0.141 | 0.471 | 0.493 | 7754 | tags=15%, list=13%, signal=18% |
| 1923 | TRANSFERASE\_ACTIVITY\_\_TRANSFERRING\_SULFUR\_CONTAINING\_GROUPS(GO:0016782) |  | 75 | 0.80 | 1.28 | 0.141 | 0.471 | 0.493 | 8052 | tags=25%, list=14%, signal=29% |
| 1924 | POSITIVE\_REGULATION\_OF\_TUMOR\_NECROSIS\_FACTOR\_PRODUCTION(GO:0032760) |  | 66 | 0.80 | 1.28 | 0.141 | 0.471 | 0.493 | 7461 | tags=15%, list=13%, signal=17% |
| 1925 | NEGATIVE\_REGULATION\_OF\_ION\_TRANSPORT(GO:0043271) |  | 141 | 0.80 | 1.28 | 0.141 | 0.470 | 0.493 | 8109 | tags=23%, list=14%, signal=26% |
| 1926 | POSITIVE\_REGULATION\_OF\_MULTICELLULAR\_ORGANISMAL\_PROCESS(GO:0051240) |  | 1607 | 0.80 | 1.28 | 0.141 | 0.470 | 0.493 | 7995 | tags=19%, list=14%, signal=22% |
| 1927 | RNA\_POLYMERASE\_II\_REGULATORY\_REGION\_DNA\_BINDING(GO:0001012) |  | 638 | 0.81 | 1.28 | 0.141 | 0.470 | 0.493 | 8038 | tags=22%, list=14%, signal=25% |
| 1928 | POSITIVE\_REGULATION\_OF\_DEVELOPMENTAL\_PROCESS(GO:0051094) |  | 1326 | 0.81 | 1.28 | 0.141 | 0.470 | 0.493 | 7546 | tags=18%, list=13%, signal=20% |
| 1929 | G\_PROTEIN\_COUPLED\_RECEPTOR\_SIGNALING\_PATHWAY(GO:0007186) |  | 1349 | 0.79 | 1.28 | 0.141 | 0.470 | 0.493 | 8279 | tags=16%, list=14%, signal=18% |
| 1930 | REGULATION\_OF\_CELL\_DEATH(GO:0010941) |  | 1660 | 0.81 | 1.28 | 0.141 | 0.470 | 0.493 | 7260 | tags=12%, list=12%, signal=13% |
| 1931 | CHONDROCYTE\_DIFFERENTIATION(GO:0002062) |  | 99 | 0.81 | 1.28 | 0.141 | 0.470 | 0.493 | 7407 | tags=26%, list=13%, signal=30% |
| 1932 | PROSTATE\_GLAND\_EPITHELIUM\_MORPHOGENESIS(GO:0060740) |  | 26 | 0.85 | 1.28 | 0.141 | 0.470 | 0.493 | 6473 | tags=27%, list=11%, signal=30% |
| 1933 | POSITIVE\_REGULATION\_OF\_INTERLEUKIN\_1\_BETA\_PRODUCTION(GO:0032731) |  | 35 | 0.83 | 1.28 | 0.000 | 0.469 | 0.493 | 6473 | tags=9%, list=11%, signal=10% |
| 1934 | NEGATIVE\_REGULATION\_OF\_SECRETION(GO:0051048) |  | 206 | 0.78 | 1.28 | 0.141 | 0.469 | 0.493 | 7640 | tags=18%, list=13%, signal=21% |
| 1935 | PARASYMPATHETIC\_NERVOUS\_SYSTEM\_DEVELOPMENT(GO:0048486) |  | 18 | 0.84 | 1.28 | 0.000 | 0.469 | 0.493 | 6068 | tags=11%, list=10%, signal=12% |
| 1936 | NEGATIVE\_REGULATION\_OF\_NEURAL\_PRECURSOR\_CELL\_PROLIFERATION(GO:2000178) |  | 20 | 0.88 | 1.28 | 0.141 | 0.469 | 0.493 | 6473 | tags=25%, list=11%, signal=28% |
| 1937 | CYCLIC\_NUCLEOTIDE\_MEDIATED\_SIGNALING(GO:0019935) |  | 148 | 0.81 | 1.28 | 0.141 | 0.469 | 0.493 | 7543 | tags=21%, list=13%, signal=24% |
| 1938 | RNA\_POLYMERASE\_II\_REGULATORY\_REGION\_SEQUENCE\_SPECIFIC\_DNA\_BINDING(GO:0000977) |  | 637 | 0.81 | 1.28 | 0.141 | 0.469 | 0.493 | 8038 | tags=22%, list=14%, signal=26% |
| 1939 | IMMUNE\_RESPONSE(GO:0006955) |  | 2157 | 0.80 | 1.28 | 0.141 | 0.469 | 0.493 | 7669 | tags=12%, list=13%, signal=13% |
| 1940 | MORPHOGENESIS\_OF\_AN\_EPITHELIAL\_SHEET(GO:0002011) |  | 54 | 0.82 | 1.28 | 0.141 | 0.469 | 0.493 | 6473 | tags=15%, list=11%, signal=17% |
| 1941 | TERMINAL\_BOUTON(GO:0043195) |  | 64 | 0.82 | 1.28 | 0.141 | 0.469 | 0.493 | 7963 | tags=20%, list=13%, signal=23% |
| 1942 | PROGRAMMED\_CELL\_DEATH(GO:0012501) |  | 2050 | 0.82 | 1.28 | 0.141 | 0.469 | 0.493 | 7217 | tags=11%, list=12%, signal=12% |
| 1943 | TRANSLATION\_INITIATION\_FACTOR\_BINDING(GO:0031369) |  | 33 | 0.79 | 1.28 | 0.000 | 0.468 | 0.493 | 7256 | tags=9%, list=12%, signal=10% |
| 1944 | POSITIVE\_REGULATION\_OF\_LIPASE\_ACTIVITY(GO:0060193) |  | 62 | 0.81 | 1.28 | 0.141 | 0.468 | 0.493 | 6928 | tags=23%, list=12%, signal=26% |
| 1945 | TISSUE\_MIGRATION(GO:0090130) |  | 330 | 0.81 | 1.28 | 0.141 | 0.468 | 0.493 | 7698 | tags=17%, list=13%, signal=19% |
| 1946 | CELL\_DIFFERENTIATION(GO:0030154) |  | 4072 | 0.83 | 1.28 | 0.141 | 0.468 | 0.493 | 7547 | tags=16%, list=13%, signal=18% |
| 1947 | REGULATION\_OF\_HOMOTYPIC\_CELL\_CELL\_ADHESION(GO:0034110) |  | 24 | 0.85 | 1.28 | 0.000 | 0.468 | 0.493 | 7546 | tags=29%, list=13%, signal=33% |
| 1948 | RESPONSE\_TO\_PROTOZOAN(GO:0001562) |  | 21 | 0.81 | 1.28 | 0.141 | 0.468 | 0.493 | 7435 | tags=33%, list=13%, signal=38% |
| 1949 | APOPTOTIC\_PROCESS(GO:0006915) |  | 1919 | 0.81 | 1.28 | 0.141 | 0.468 | 0.493 | 7217 | tags=11%, list=12%, signal=12% |
| 1950 | POSITIVE\_REGULATION\_OF\_BEHAVIOR(GO:0048520) |  | 23 | 0.82 | 1.28 | 0.000 | 0.468 | 0.493 | 7134 | tags=30%, list=12%, signal=35% |
| 1951 | HEART\_DEVELOPMENT(GO:0007507) |  | 539 | 0.81 | 1.28 | 0.141 | 0.468 | 0.493 | 7969 | tags=20%, list=13%, signal=23% |
| 1952 | FEMALE\_PREGNANCY(GO:0007565) |  | 192 | 0.81 | 1.28 | 0.141 | 0.467 | 0.493 | 7730 | tags=14%, list=13%, signal=16% |
| 1953 | REGULATION\_OF\_TISSUE\_REMODELING(GO:0034103) |  | 86 | 0.78 | 1.28 | 0.141 | 0.467 | 0.493 | 8241 | tags=20%, list=14%, signal=23% |
| 1954 | CHLORIDE\_TRANSMEMBRANE\_TRANSPORTER\_ACTIVITY(GO:0015108) |  | 93 | 0.81 | 1.28 | 0.141 | 0.467 | 0.493 | 8596 | tags=25%, list=15%, signal=29% |
| 1955 | RNA\_POLYMERASE\_II\_TRANSCRIPTION\_COACTIVATOR\_ACTIVITY(GO:0001105) |  | 43 | 0.83 | 1.28 | 0.141 | 0.467 | 0.493 | 6421 | tags=19%, list=11%, signal=21% |
| 1956 | POSITIVE\_REGULATION\_OF\_PROTEIN\_DEPOLYMERIZATION(GO:1901881) |  | 19 | 0.83 | 1.28 | 0.141 | 0.467 | 0.493 | 5938 | tags=11%, list=10%, signal=12% |
| 1957 | EPITHELIAL\_CELL\_MIGRATION(GO:0010631) |  | 321 | 0.80 | 1.28 | 0.141 | 0.467 | 0.493 | 7698 | tags=16%, list=13%, signal=18% |
| 1958 | CELL\_LEADING\_EDGE(GO:0031252) |  | 373 | 0.80 | 1.28 | 0.141 | 0.467 | 0.493 | 8249 | tags=14%, list=14%, signal=16% |
| 1959 | REGULATION\_OF\_HEART\_GROWTH(GO:0060420) |  | 75 | 0.79 | 1.28 | 0.141 | 0.467 | 0.493 | 8380 | tags=27%, list=14%, signal=31% |
| 1960 | NEUROTRANSMITTER\_METABOLIC\_PROCESS(GO:0042133) |  | 142 | 0.79 | 1.28 | 0.141 | 0.467 | 0.493 | 8382 | tags=20%, list=14%, signal=24% |
| 1961 | REGULATION\_OF\_SIGNAL\_TRANSDUCTION(GO:0009966) |  | 3163 | 0.82 | 1.28 | 0.141 | 0.467 | 0.493 | 7170 | tags=13%, list=12%, signal=14% |
| 1962 | RECEPTOR\_METABOLIC\_PROCESS(GO:0043112) |  | 167 | 0.80 | 1.28 | 0.141 | 0.467 | 0.493 | 7154 | tags=12%, list=12%, signal=14% |
| 1963 | PLASMINOGEN\_ACTIVATION(GO:0031639) |  | 23 | 0.85 | 1.28 | 0.141 | 0.467 | 0.493 | 7916 | tags=35%, list=13%, signal=40% |
| 1964 | SPINAL\_CORD\_MOTOR\_NEURON\_DIFFERENTIATION(GO:0021522) |  | 33 | 0.87 | 1.28 | 0.000 | 0.466 | 0.493 | 6760 | tags=45%, list=11%, signal=51% |
| 1965 | LEARNING\_OR\_MEMORY(GO:0007611) |  | 240 | 0.79 | 1.28 | 0.141 | 0.466 | 0.493 | 7962 | tags=21%, list=13%, signal=24% |
| 1966 | NEGATIVE\_REGULATION\_OF\_POTASSIUM\_ION\_TRANSPORT(GO:0043267) |  | 36 | 0.79 | 1.28 | 0.000 | 0.466 | 0.493 | 7697 | tags=22%, list=13%, signal=26% |
| 1967 | WOUND\_HEALING(GO:0042060) |  | 555 | 0.79 | 1.28 | 0.141 | 0.466 | 0.493 | 7546 | tags=16%, list=13%, signal=18% |
| 1968 | CEREBRAL\_CORTEX\_RADIALLY\_ORIENTED\_CELL\_MIGRATION(GO:0021799) |  | 30 | 0.88 | 1.28 | 0.141 | 0.466 | 0.493 | 6328 | tags=20%, list=11%, signal=22% |
| 1969 | MEMBRANE\_BIOGENESIS(GO:0044091) |  | 32 | 0.81 | 1.28 | 0.000 | 0.466 | 0.493 | 8937 | tags=31%, list=15%, signal=37% |
| 1970 | GABA\_RECEPTOR\_ACTIVITY(GO:0016917) |  | 22 | 0.86 | 1.28 | 0.000 | 0.466 | 0.493 | 7049 | tags=27%, list=12%, signal=31% |
| 1971 | REGULATION\_OF\_PROTEIN\_METABOLIC\_PROCESS(GO:0051246) |  | 2955 | 0.81 | 1.28 | 0.141 | 0.466 | 0.493 | 7669 | tags=12%, list=13%, signal=13% |
| 1972 | PHOSPHATIDYLINOSITOL\_3\_KINASE\_SIGNALING(GO:0014065) |  | 130 | 0.79 | 1.28 | 0.141 | 0.465 | 0.493 | 7046 | tags=16%, list=12%, signal=18% |
| 1973 | MRNA\_3'\_UTR\_BINDING(GO:0003730) |  | 68 | 0.77 | 1.28 | 0.000 | 0.465 | 0.493 | 6397 | tags=7%, list=11%, signal=8% |
| 1974 | SYNAPTIC\_TRANSMISSION\_\_DOPAMINERGIC(GO:0001963) |  | 32 | 0.86 | 1.28 | 0.141 | 0.465 | 0.493 | 7298 | tags=22%, list=12%, signal=25% |
| 1975 | VESICLE\_MEDIATED\_TRANSPORT\_TO\_THE\_PLASMA\_MEMBRANE(GO:0098876) |  | 72 | 0.79 | 1.28 | 0.000 | 0.465 | 0.493 | 6066 | tags=6%, list=10%, signal=6% |
| 1976 | PEPTIDASE\_INHIBITOR\_ACTIVITY(GO:0030414) |  | 179 | 0.80 | 1.28 | 0.141 | 0.465 | 0.493 | 8070 | tags=20%, list=14%, signal=23% |
| 1977 | CELLULAR\_HOMEOSTASIS(GO:0019725) |  | 893 | 0.78 | 1.28 | 0.141 | 0.465 | 0.493 | 7544 | tags=15%, list=13%, signal=17% |
| 1978 | DISORDERED\_DOMAIN\_SPECIFIC\_BINDING(GO:0097718) |  | 35 | 0.77 | 1.28 | 0.000 | 0.465 | 0.493 | 6662 | tags=17%, list=11%, signal=19% |
| 1979 | INORGANIC\_ION\_TRANSMEMBRANE\_TRANSPORT(GO:0098660) |  | 824 | 0.79 | 1.28 | 0.141 | 0.465 | 0.493 | 7979 | tags=20%, list=13%, signal=23% |
| 1980 | CHEMICAL\_SYNAPTIC\_TRANSMISSION\_\_POSTSYNAPTIC(GO:0099565) |  | 116 | 0.78 | 1.28 | 0.000 | 0.465 | 0.493 | 7963 | tags=29%, list=13%, signal=34% |
| 1981 | CELLULAR\_RESPONSE\_TO\_IONIZING\_RADIATION(GO:0071479) |  | 68 | 0.83 | 1.28 | 0.141 | 0.465 | 0.493 | 6910 | tags=9%, list=12%, signal=10% |
| 1982 | ENZYME\_BINDING(GO:0019899) |  | 2202 | 0.82 | 1.28 | 0.141 | 0.465 | 0.493 | 6940 | tags=8%, list=12%, signal=9% |
| 1983 | REGULATION\_OF\_PROTEIN\_PHOSPHORYLATION(GO:0001932) |  | 1388 | 0.80 | 1.28 | 0.141 | 0.465 | 0.493 | 7702 | tags=14%, list=13%, signal=16% |
| 1984 | ALPHA\_TUBULIN\_BINDING(GO:0043014) |  | 29 | 0.79 | 1.28 | 0.000 | 0.464 | 0.493 | 7345 | tags=14%, list=12%, signal=16% |
| 1985 | NEGATIVE\_CHEMOTAXIS(GO:0050919) |  | 34 | 0.84 | 1.28 | 0.141 | 0.464 | 0.493 | 7969 | tags=32%, list=13%, signal=37% |
| 1986 | HEART\_TRABECULA\_FORMATION(GO:0060347) |  | 15 | 0.83 | 1.28 | 0.000 | 0.464 | 0.493 | 6527 | tags=40%, list=11%, signal=45% |
| 1987 | NEGATIVE\_REGULATION\_OF\_PHOSPHATASE\_ACTIVITY(GO:0010923) |  | 97 | 0.77 | 1.28 | 0.000 | 0.464 | 0.493 | 7544 | tags=13%, list=13%, signal=15% |
| 1988 | TRANSCRIPTION\_FACTOR\_ACTIVITY\_\_RNA\_POLYMERASE\_II\_CORE\_PROMOTER\_SEQUENCE\_SPECIFIC\_DNA\_BINDING(GO:0000983) |  | 23 | 0.76 | 1.28 | 0.000 | 0.464 | 0.493 | 7800 | tags=30%, list=13%, signal=35% |
| 1989 | METAL\_ION\_TRANSMEMBRANE\_TRANSPORTER\_ACTIVITY(GO:0046873) |  | 460 | 0.79 | 1.28 | 0.141 | 0.464 | 0.493 | 9496 | tags=32%, list=16%, signal=37% |
| 1990 | NEGATIVE\_REGULATION\_OF\_CELL\_DEVELOPMENT(GO:0010721) |  | 295 | 0.82 | 1.28 | 0.141 | 0.464 | 0.493 | 8570 | tags=25%, list=14%, signal=29% |
| 1991 | CARTILAGE\_DEVELOPMENT(GO:0051216) |  | 186 | 0.81 | 1.28 | 0.141 | 0.464 | 0.493 | 8003 | tags=31%, list=14%, signal=35% |
| 1992 | GENERATION\_OF\_NEURONS(GO:0048699) |  | 1428 | 0.81 | 1.28 | 0.141 | 0.464 | 0.493 | 8003 | tags=21%, list=14%, signal=23% |
| 1993 | PROSTATE\_GLAND\_DEVELOPMENT(GO:0030850) |  | 48 | 0.82 | 1.28 | 0.141 | 0.464 | 0.493 | 7913 | tags=27%, list=13%, signal=31% |
| 1994 | SINGLE\_FERTILIZATION(GO:0007338) |  | 145 | 0.81 | 1.28 | 0.141 | 0.463 | 0.493 | 8643 | tags=23%, list=15%, signal=27% |
| 1995 | RESPONSE\_TO\_WOUNDING(GO:0009611) |  | 667 | 0.79 | 1.28 | 0.141 | 0.463 | 0.493 | 7546 | tags=16%, list=13%, signal=18% |
| 1996 | SULFATE\_TRANSPORT(GO:0008272) |  | 16 | 0.79 | 1.28 | 0.000 | 0.463 | 0.493 | 9957 | tags=50%, list=17%, signal=60% |
| 1997 | REGULATION\_OF\_BIOLOGICAL\_QUALITY(GO:0065008) |  | 3666 | 0.81 | 1.28 | 0.141 | 0.463 | 0.493 | 7745 | tags=15%, list=13%, signal=17% |
| 1998 | CHLORIDE\_TRANSPORT(GO:0006821) |  | 105 | 0.81 | 1.28 | 0.141 | 0.463 | 0.493 | 8596 | tags=28%, list=15%, signal=32% |
| 1999 | POSITIVE\_REGULATION\_OF\_WOUND\_HEALING(GO:0090303) |  | 57 | 0.82 | 1.28 | 0.141 | 0.463 | 0.493 | 7128 | tags=16%, list=12%, signal=18% |
| 2000 | INORGANIC\_MOLECULAR\_ENTITY\_TRANSMEMBRANE\_TRANSPORTER\_ACTIVITY(GO:0015318) |  | 856 | 0.79 | 1.28 | 0.141 | 0.463 | 0.493 | 8432 | tags=24%, list=14%, signal=28% |
| 2001 | TRANSCRIPTIONAL\_ACTIVATOR\_ACTIVITY\_\_RNA\_POLYMERASE\_II\_TRANSCRIPTION\_REGULATORY\_REGION\_SEQUENCE\_SPECIFIC\_DNA\_BINDING(GO:0001228) |  | 413 | 0.81 | 1.28 | 0.141 | 0.463 | 0.493 | 8038 | tags=24%, list=14%, signal=27% |
| 2002 | PHAGOCYTOSIS\_\_RECOGNITION(GO:0006910) |  | 66 | 0.77 | 1.28 | 0.000 | 0.462 | 0.493 | 6886 | tags=6%, list=12%, signal=7% |
| 2003 | GLYCEROLIPID\_METABOLIC\_PROCESS(GO:0046486) |  | 447 | 0.78 | 1.28 | 0.141 | 0.462 | 0.493 | 7699 | tags=15%, list=13%, signal=18% |
| 2004 | BASAL\_TRANSCRIPTION\_MACHINERY\_BINDING(GO:0001098) |  | 57 | 0.84 | 1.28 | 0.141 | 0.462 | 0.493 | 6006 | tags=7%, list=10%, signal=8% |
| 2005 | BASAL\_RNA\_POLYMERASE\_II\_TRANSCRIPTION\_MACHINERY\_BINDING(GO:0001099) |  | 57 | 0.84 | 1.28 | 0.141 | 0.462 | 0.493 | 6006 | tags=7%, list=10%, signal=8% |
| 2006 | ION\_TRANSMEMBRANE\_TRANSPORT(GO:0034220) |  | 1110 | 0.79 | 1.28 | 0.141 | 0.462 | 0.493 | 8066 | tags=21%, list=14%, signal=23% |
| 2007 | ZYMOGEN\_ACTIVATION(GO:0031638) |  | 44 | 0.82 | 1.28 | 0.141 | 0.462 | 0.493 | 7916 | tags=20%, list=13%, signal=24% |
| 2008 | CELL\_MORPHOGENESIS\_INVOLVED\_IN\_NEURON\_DIFFERENTIATION(GO:0048667) |  | 548 | 0.81 | 1.28 | 0.141 | 0.462 | 0.493 | 7461 | tags=19%, list=13%, signal=22% |
| 2009 | EPITHELIUM\_MIGRATION(GO:0090132) |  | 324 | 0.80 | 1.28 | 0.141 | 0.462 | 0.493 | 7698 | tags=16%, list=13%, signal=18% |
| 2010 | EXTRACELLULAR\_SPACE(GO:0005615) |  | 3964 | 0.82 | 1.28 | 0.141 | 0.461 | 0.493 | 7467 | tags=12%, list=13%, signal=13% |
| 2011 | BRANCHING\_INVOLVED\_IN\_SALIVARY\_GLAND\_MORPHOGENESIS(GO:0060445) |  | 22 | 0.84 | 1.28 | 0.141 | 0.461 | 0.493 | 7343 | tags=23%, list=12%, signal=26% |
| 2012 | ION\_HOMEOSTASIS(GO:0050801) |  | 751 | 0.79 | 1.28 | 0.141 | 0.461 | 0.493 | 7727 | tags=19%, list=13%, signal=21% |
| 2013 | PHOSPHOLIPASE\_C\_ACTIVATING\_G\_PROTEIN\_COUPLED\_RECEPTOR\_SIGNALING\_PATHWAY(GO:0007200) |  | 113 | 0.81 | 1.28 | 0.141 | 0.461 | 0.493 | 7479 | tags=21%, list=13%, signal=24% |
| 2014 | REGULATION\_OF\_INTERLEUKIN\_8\_SECRETION(GO:2000482) |  | 24 | 0.79 | 1.28 | 0.000 | 0.461 | 0.493 | 6530 | tags=17%, list=11%, signal=19% |
| 2015 | DORSAL\_VENTRAL\_NEURAL\_TUBE\_PATTERNING(GO:0021904) |  | 25 | 0.87 | 1.28 | 0.141 | 0.461 | 0.493 | 6714 | tags=20%, list=11%, signal=23% |
| 2016 | ORGANIC\_HYDROXY\_COMPOUND\_TRANSMEMBRANE\_TRANSPORTER\_ACTIVITY(GO:1901618) |  | 47 | 0.81 | 1.28 | 0.141 | 0.461 | 0.493 | 6374 | tags=26%, list=11%, signal=29% |
| 2017 | TRANSMEMBRANE\_SIGNALING\_RECEPTOR\_ACTIVITY(GO:0004888) |  | 1242 | 0.80 | 1.28 | 0.141 | 0.461 | 0.493 | 7981 | tags=17%, list=13%, signal=20% |
| 2018 | REGULATION\_OF\_PHOSPHATIDYLINOSITOL\_3\_KINASE\_SIGNALING(GO:0014066) |  | 100 | 0.81 | 1.28 | 0.141 | 0.461 | 0.493 | 7046 | tags=17%, list=12%, signal=19% |
| 2019 | REGULATION\_OF\_ANION\_TRANSPORT(GO:0044070) |  | 84 | 0.81 | 1.28 | 0.141 | 0.461 | 0.493 | 8452 | tags=25%, list=14%, signal=29% |
| 2020 | MITOTIC\_G2\_M\_TRANSITION\_CHECKPOINT(GO:0044818) |  | 31 | 0.77 | 1.28 | 0.000 | 0.461 | 0.493 | 10550 | tags=39%, list=18%, signal=47% |
| 2021 | SODIUM\_ION\_TRANSMEMBRANE\_TRANSPORTER\_ACTIVITY(GO:0015081) |  | 160 | 0.81 | 1.28 | 0.141 | 0.461 | 0.493 | 7352 | tags=29%, list=12%, signal=33% |
| 2022 | INNERVATION(GO:0060384) |  | 25 | 0.83 | 1.28 | 0.000 | 0.460 | 0.493 | 7789 | tags=32%, list=13%, signal=37% |
| 2023 | REGULATION\_OF\_SYNAPTIC\_TRANSMISSION\_\_DOPAMINERGIC(GO:0032225) |  | 17 | 0.87 | 1.28 | 0.141 | 0.460 | 0.493 | 7298 | tags=29%, list=12%, signal=34% |
| 2024 | REGULATION\_OF\_MULTICELLULAR\_ORGANISMAL\_DEVELOPMENT(GO:2000026) |  | 1996 | 0.81 | 1.28 | 0.141 | 0.460 | 0.493 | 7546 | tags=17%, list=13%, signal=19% |
| 2025 | SERINE\_TYPE\_PEPTIDASE\_ACTIVITY(GO:0008236) |  | 280 | 0.82 | 1.28 | 0.141 | 0.460 | 0.493 | 6810 | tags=13%, list=12%, signal=15% |
| 2026 | NEGATIVE\_REGULATION\_OF\_SIGNAL\_TRANSDUCTION\_BY\_P53\_CLASS\_MEDIATOR(GO:1901797) |  | 30 | 0.81 | 1.28 | 0.000 | 0.460 | 0.493 | 7939 | tags=13%, list=13%, signal=15% |
| 2027 | BODY\_MORPHOGENESIS(GO:0010171) |  | 50 | 0.79 | 1.28 | 0.000 | 0.460 | 0.493 | 8177 | tags=26%, list=14%, signal=30% |
| 2028 | RESPONSE\_TO\_COCAINE(GO:0042220) |  | 52 | 0.82 | 1.28 | 0.000 | 0.460 | 0.493 | 7298 | tags=21%, list=12%, signal=24% |
| 2029 | RESPONSE\_TO\_VITAMIN\_D(GO:0033280) |  | 36 | 0.80 | 1.28 | 0.141 | 0.460 | 0.493 | 6937 | tags=28%, list=12%, signal=31% |
| 2030 | SERINE\_HYDROLASE\_ACTIVITY(GO:0017171) |  | 285 | 0.82 | 1.28 | 0.141 | 0.460 | 0.493 | 6810 | tags=13%, list=12%, signal=15% |
| 2031 | CELLULAR\_DEVELOPMENTAL\_PROCESS(GO:0048869) |  | 4247 | 0.83 | 1.28 | 0.141 | 0.460 | 0.493 | 7547 | tags=16%, list=13%, signal=17% |
| 2032 | CYTOKINE\_SECRETION(GO:0050663) |  | 199 | 0.77 | 1.28 | 0.141 | 0.459 | 0.493 | 7984 | tags=17%, list=14%, signal=19% |
| 2033 | HETEROCHROMATIN(GO:0000792) |  | 73 | 0.75 | 1.28 | 0.000 | 0.459 | 0.493 | 6154 | tags=7%, list=10%, signal=8% |
| 2034 | CATION\_HOMEOSTASIS(GO:0055080) |  | 678 | 0.79 | 1.28 | 0.141 | 0.459 | 0.493 | 7544 | tags=18%, list=13%, signal=20% |
| 2035 | MONOCYTE\_CHEMOTAXIS(GO:0002548) |  | 62 | 0.83 | 1.28 | 0.141 | 0.459 | 0.493 | 7081 | tags=16%, list=12%, signal=18% |
| 2036 | DICARBOXYLIC\_ACID\_TRANSPORT(GO:0006835) |  | 89 | 0.77 | 1.28 | 0.000 | 0.459 | 0.493 | 7873 | tags=22%, list=13%, signal=26% |
| 2037 | RESPONSE\_TO\_INORGANIC\_SUBSTANCE(GO:0010035) |  | 520 | 0.80 | 1.28 | 0.141 | 0.459 | 0.493 | 7696 | tags=13%, list=13%, signal=15% |
| 2038 | PYRIDINE\_CONTAINING\_COMPOUND\_METABOLIC\_PROCESS(GO:0072524) |  | 180 | 0.79 | 1.28 | 0.141 | 0.459 | 0.493 | 6696 | tags=7%, list=11%, signal=8% |
| 2039 | INTEGRIN\_BINDING(GO:0005178) |  | 118 | 0.81 | 1.28 | 0.141 | 0.459 | 0.493 | 6857 | tags=18%, list=12%, signal=20% |
| 2040 | NEGATIVE\_REGULATION\_OF\_MONOOXYGENASE\_ACTIVITY(GO:0032769) |  | 15 | 0.86 | 1.28 | 0.000 | 0.459 | 0.493 | 5894 | tags=33%, list=10%, signal=37% |
| 2041 | BASE\_EXCISION\_REPAIR(GO:0006284) |  | 44 | 0.77 | 1.28 | 0.000 | 0.458 | 0.493 | 5890 | tags=2%, list=10%, signal=3% |
| 2042 | MODULATION\_OF\_CHEMICAL\_SYNAPTIC\_TRANSMISSION(GO:0050804) |  | 316 | 0.79 | 1.28 | 0.141 | 0.458 | 0.493 | 8241 | tags=22%, list=14%, signal=26% |
| 2043 | REGULATION\_OF\_TRANS\_SYNAPTIC\_SIGNALING(GO:0099177) |  | 316 | 0.79 | 1.28 | 0.141 | 0.458 | 0.493 | 8241 | tags=22%, list=14%, signal=26% |
| 2044 | HORMONE\_METABOLIC\_PROCESS(GO:0042445) |  | 219 | 0.79 | 1.28 | 0.141 | 0.458 | 0.493 | 8013 | tags=18%, list=14%, signal=21% |
| 2045 | CELL\_DEATH(GO:0008219) |  | 2178 | 0.82 | 1.28 | 0.141 | 0.458 | 0.493 | 7217 | tags=11%, list=12%, signal=12% |
| 2046 | CELL\_CELL\_ADHESION\_MEDIATED\_BY\_CADHERIN(GO:0044331) |  | 16 | 0.83 | 1.28 | 0.141 | 0.458 | 0.493 | 8673 | tags=31%, list=15%, signal=37% |
| 2047 | HEAD\_MORPHOGENESIS(GO:0060323) |  | 37 | 0.80 | 1.28 | 0.141 | 0.458 | 0.493 | 8177 | tags=27%, list=14%, signal=31% |
| 2048 | PHOTORECEPTOR\_CELL\_MAINTENANCE(GO:0045494) |  | 34 | 0.84 | 1.28 | 0.141 | 0.458 | 0.493 | 7255 | tags=29%, list=12%, signal=34% |
| 2049 | REGULATION\_OF\_LONG\_TERM\_SYNAPTIC\_POTENTIATION(GO:1900271) |  | 31 | 0.79 | 1.28 | 0.000 | 0.458 | 0.493 | 7738 | tags=13%, list=13%, signal=15% |
| 2050 | NEGATIVE\_REGULATION\_OF\_POTASSIUM\_ION\_TRANSMEMBRANE\_TRANSPORTER\_ACTIVITY(GO:1901017) |  | 25 | 0.81 | 1.28 | 0.000 | 0.458 | 0.493 | 7697 | tags=24%, list=13%, signal=28% |
| 2051 | ORGANIC\_HYDROXY\_COMPOUND\_TRANSPORT(GO:0015850) |  | 230 | 0.80 | 1.28 | 0.141 | 0.457 | 0.493 | 7358 | tags=21%, list=12%, signal=24% |
| 2052 | REGULATION\_OF\_LIPID\_KINASE\_ACTIVITY(GO:0043550) |  | 54 | 0.80 | 1.28 | 0.141 | 0.457 | 0.493 | 6928 | tags=17%, list=12%, signal=19% |
| 2053 | RENAL\_SYSTEM\_PROCESS\_INVOLVED\_IN\_REGULATION\_OF\_BLOOD\_VOLUME(GO:0001977) |  | 16 | 0.86 | 1.28 | 0.141 | 0.457 | 0.493 | 6525 | tags=31%, list=11%, signal=35% |
| 2054 | BEHAVIORAL\_RESPONSE\_TO\_COCAINE(GO:0048148) |  | 21 | 0.86 | 1.28 | 0.141 | 0.457 | 0.493 | 5894 | tags=19%, list=10%, signal=21% |
| 2055 | AMINO\_ACID\_IMPORT(GO:0043090) |  | 19 | 0.80 | 1.28 | 0.141 | 0.457 | 0.493 | 7352 | tags=26%, list=12%, signal=30% |
| 2056 | CELLULAR\_RESPONSE\_TO\_MONOAMINE\_STIMULUS(GO:0071868) |  | 35 | 0.82 | 1.28 | 0.000 | 0.457 | 0.493 | 7426 | tags=20%, list=13%, signal=23% |
| 2057 | CELLULAR\_RESPONSE\_TO\_CATECHOLAMINE\_STIMULUS(GO:0071870) |  | 35 | 0.82 | 1.28 | 0.000 | 0.457 | 0.493 | 7426 | tags=20%, list=13%, signal=23% |
| 2058 | NEGATIVE\_REGULATION\_OF\_POTASSIUM\_ION\_TRANSMEMBRANE\_TRANSPORT(GO:1901380) |  | 28 | 0.81 | 1.28 | 0.000 | 0.457 | 0.493 | 7697 | tags=25%, list=13%, signal=29% |
| 2059 | POSITIVE\_REGULATION\_OF\_INTRACELLULAR\_PROTEIN\_TRANSPORT(GO:0090316) |  | 190 | 0.82 | 1.28 | 0.141 | 0.456 | 0.493 | 6793 | tags=8%, list=11%, signal=9% |
| 2060 | REGULATION\_OF\_RESPONSE\_TO\_STRESS(GO:0080134) |  | 1467 | 0.80 | 1.28 | 0.141 | 0.456 | 0.493 | 7727 | tags=13%, list=13%, signal=15% |
| 2061 | FEMALE\_GONAD\_DEVELOPMENT(GO:0008585) |  | 95 | 0.80 | 1.28 | 0.141 | 0.456 | 0.493 | 7547 | tags=21%, list=13%, signal=24% |
| 2062 | BODY\_FLUID\_SECRETION(GO:0007589) |  | 92 | 0.81 | 1.28 | 0.141 | 0.456 | 0.493 | 7038 | tags=16%, list=12%, signal=18% |
| 2063 | DRUG\_METABOLIC\_PROCESS(GO:0017144) |  | 794 | 0.78 | 1.28 | 0.141 | 0.456 | 0.493 | 7935 | tags=12%, list=13%, signal=14% |
| 2064 | REGULATION\_OF\_PROTEIN\_LOCALIZATION\_TO\_CELL\_PERIPHERY(GO:1904375) |  | 96 | 0.80 | 1.28 | 0.141 | 0.456 | 0.493 | 7407 | tags=10%, list=13%, signal=12% |
| 2065 | AXON\_DEVELOPMENT(GO:0061564) |  | 482 | 0.81 | 1.28 | 0.141 | 0.456 | 0.493 | 7975 | tags=22%, list=13%, signal=26% |
| 2066 | DEVELOPMENT\_OF\_PRIMARY\_FEMALE\_SEXUAL\_CHARACTERISTICS(GO:0046545) |  | 100 | 0.80 | 1.28 | 0.141 | 0.456 | 0.493 | 7547 | tags=23%, list=13%, signal=26% |
| 2067 | G\_PROTEIN\_COUPLED\_RECEPTOR\_SIGNALING\_PATHWAY\_\_COUPLED\_TO\_CYCLIC\_NUCLEOTIDE\_SECOND\_MESSENGER(GO:0007187) |  | 213 | 0.82 | 1.28 | 0.141 | 0.456 | 0.493 | 7479 | tags=18%, list=13%, signal=21% |
| 2068 | REGULATION\_OF\_MICROTUBULE\_CYTOSKELETON\_ORGANIZATION(GO:0070507) |  | 167 | 0.75 | 1.28 | 0.000 | 0.456 | 0.493 | 8262 | tags=11%, list=14%, signal=12% |
| 2069 | RENAL\_FILTRATION(GO:0097205) |  | 21 | 0.86 | 1.28 | 0.141 | 0.456 | 0.493 | 6525 | tags=33%, list=11%, signal=37% |
| 2070 | REGULATION\_OF\_VASOCONSTRICTION(GO:0019229) |  | 62 | 0.80 | 1.28 | 0.141 | 0.456 | 0.493 | 7845 | tags=29%, list=13%, signal=33% |
| 2071 | CYTOPLASMIC\_REGION(GO:0099568) |  | 422 | 0.81 | 1.28 | 0.141 | 0.455 | 0.493 | 8290 | tags=19%, list=14%, signal=22% |
| 2072 | METAL\_ION\_TRANSPORT(GO:0030001) |  | 875 | 0.79 | 1.28 | 0.141 | 0.455 | 0.493 | 8217 | tags=22%, list=14%, signal=25% |
| 2073 | CYTOPLASMIC\_VESICLE\_MEMBRANE(GO:0030659) |  | 729 | 0.79 | 1.28 | 0.141 | 0.455 | 0.493 | 7940 | tags=13%, list=13%, signal=15% |
| 2074 | REGULATION\_OF\_POTASSIUM\_ION\_TRANSPORT(GO:0043266) |  | 100 | 0.79 | 1.28 | 0.141 | 0.455 | 0.493 | 9608 | tags=25%, list=16%, signal=30% |
| 2075 | PRODUCTION\_OF\_MIRNAS\_INVOLVED\_IN\_GENE\_SILENCING\_BY\_MIRNA(GO:0035196) |  | 41 | 0.84 | 1.28 | 0.141 | 0.455 | 0.493 | 5858 | tags=5%, list=10%, signal=5% |
| 2076 | REGULATION\_OF\_CELL\_COMMUNICATION(GO:0010646) |  | 3463 | 0.82 | 1.28 | 0.141 | 0.455 | 0.493 | 7175 | tags=13%, list=12%, signal=14% |
| 2077 | RECEPTOR\_LOCALIZATION\_TO\_SYNAPSE(GO:0097120) |  | 26 | 0.83 | 1.28 | 0.000 | 0.455 | 0.493 | 7530 | tags=19%, list=13%, signal=22% |
| 2078 | CYCLIC\_NUCLEOTIDE\_PHOSPHODIESTERASE\_ACTIVITY(GO:0004112) |  | 28 | 0.75 | 1.28 | 0.000 | 0.455 | 0.493 | 7935 | tags=43%, list=13%, signal=49% |
| 2079 | POSITIVE\_REGULATION\_OF\_INTERLEUKIN\_8\_SECRETION(GO:2000484) |  | 15 | 0.81 | 1.28 | 0.000 | 0.455 | 0.493 | 6530 | tags=20%, list=11%, signal=22% |
| 2080 | ENDOPLASMIC\_RETICULUM\_PART(GO:0044432) |  | 1296 | 0.78 | 1.28 | 0.141 | 0.455 | 0.493 | 7564 | tags=12%, list=13%, signal=13% |
| 2081 | CELLULAR\_BIOGENIC\_AMINE\_METABOLIC\_PROCESS(GO:0006576) |  | 53 | 0.80 | 1.28 | 0.141 | 0.455 | 0.493 | 7830 | tags=11%, list=13%, signal=13% |
| 2082 | CHEMICAL\_SYNAPTIC\_TRANSMISSION(GO:0007268) |  | 612 | 0.80 | 1.28 | 0.141 | 0.455 | 0.493 | 7981 | tags=23%, list=13%, signal=26% |
| 2083 | ANTEROGRADE\_TRANS\_SYNAPTIC\_SIGNALING(GO:0098916) |  | 612 | 0.80 | 1.28 | 0.141 | 0.455 | 0.493 | 7981 | tags=23%, list=13%, signal=26% |
| 2084 | POSITIVE\_REGULATION\_OF\_CHONDROCYTE\_DIFFERENTIATION(GO:0032332) |  | 19 | 0.83 | 1.28 | 0.000 | 0.455 | 0.493 | 7407 | tags=42%, list=13%, signal=48% |
| 2085 | ENDOCYTIC\_VESICLE(GO:0030139) |  | 282 | 0.80 | 1.28 | 0.141 | 0.454 | 0.493 | 7467 | tags=12%, list=13%, signal=13% |
| 2086 | DOPAMINE\_RECEPTOR\_SIGNALING\_PATHWAY(GO:0007212) |  | 46 | 0.81 | 1.28 | 0.000 | 0.454 | 0.493 | 7426 | tags=13%, list=13%, signal=15% |
| 2087 | CARDIAC\_MUSCLE\_CELL\_DIFFERENTIATION(GO:0055007) |  | 118 | 0.77 | 1.28 | 0.000 | 0.454 | 0.493 | 8112 | tags=25%, list=14%, signal=28% |
| 2088 | REGULATION\_OF\_FATTY\_ACID\_OXIDATION(GO:0046320) |  | 28 | 0.81 | 1.28 | 0.000 | 0.454 | 0.493 | 8474 | tags=18%, list=14%, signal=21% |
| 2089 | TRANS\_SYNAPTIC\_SIGNALING(GO:0099537) |  | 614 | 0.80 | 1.28 | 0.141 | 0.454 | 0.493 | 7981 | tags=22%, list=13%, signal=26% |
| 2090 | SYNAPTIC\_SIGNALING(GO:0099536) |  | 614 | 0.80 | 1.28 | 0.141 | 0.454 | 0.493 | 7981 | tags=22%, list=13%, signal=26% |
| 2091 | PHOSPHOLIPASE\_ACTIVITY(GO:0004620) |  | 103 | 0.79 | 1.28 | 0.141 | 0.454 | 0.493 | 6949 | tags=16%, list=12%, signal=18% |
| 2092 | EXTRACELLULAR\_REGION\_PART(GO:0044421) |  | 4181 | 0.82 | 1.28 | 0.141 | 0.454 | 0.493 | 7467 | tags=13%, list=13%, signal=13% |
| 2093 | REGULATION\_OF\_ENDOTHELIAL\_CELL\_DIFFERENTIATION(GO:0045601) |  | 40 | 0.79 | 1.28 | 0.000 | 0.454 | 0.493 | 7754 | tags=10%, list=13%, signal=12% |
| 2094 | TRANSCRIPTION\_REGULATOR\_ACTIVITY(GO:0140110) |  | 1782 | 0.80 | 1.28 | 0.141 | 0.453 | 0.493 | 7414 | tags=16%, list=13%, signal=18% |
| 2095 | TRANSCRIPTION\_REGULATORY\_REGION\_SEQUENCE\_SPECIFIC\_DNA\_BINDING(GO:0000976) |  | 716 | 0.80 | 1.28 | 0.141 | 0.453 | 0.493 | 8038 | tags=21%, list=14%, signal=24% |
| 2096 | PROXIMAL\_DISTAL\_PATTERN\_FORMATION(GO:0009954) |  | 32 | 0.86 | 1.28 | 0.000 | 0.453 | 0.493 | 7668 | tags=47%, list=13%, signal=54% |
| 2097 | CELLULAR\_PROTEIN\_LOCALIZATION(GO:0034613) |  | 1659 | 0.81 | 1.28 | 0.141 | 0.453 | 0.493 | 7184 | tags=7%, list=12%, signal=7% |
| 2098 | CORNIFICATION(GO:0070268) |  | 112 | 0.89 | 1.28 | 0.141 | 0.453 | 0.493 | 6275 | tags=7%, list=11%, signal=8% |
| 2099 | DYNEIN\_LIGHT\_INTERMEDIATE\_CHAIN\_BINDING(GO:0051959) |  | 29 | 0.78 | 1.28 | 0.000 | 0.453 | 0.493 | 8569 | tags=24%, list=14%, signal=28% |
| 2100 | PREGANGLIONIC\_PARASYMPATHETIC\_FIBER\_DEVELOPMENT(GO:0021783) |  | 16 | 0.84 | 1.28 | 0.000 | 0.453 | 0.493 | 6068 | tags=13%, list=10%, signal=14% |
| 2101 | REGULATION\_OF\_WOUND\_HEALING(GO:0061041) |  | 135 | 0.81 | 1.28 | 0.141 | 0.453 | 0.493 | 7546 | tags=21%, list=13%, signal=24% |
| 2102 | REGULATION\_OF\_GROWTH\_OF\_SYMBIONT\_IN\_HOST(GO:0044126) |  | 16 | 0.84 | 1.28 | 0.141 | 0.453 | 0.493 | 6114 | tags=25%, list=10%, signal=28% |
| 2103 | MODULATION\_OF\_GROWTH\_OF\_SYMBIONT\_INVOLVED\_IN\_INTERACTION\_WITH\_HOST(GO:0044144) |  | 16 | 0.84 | 1.28 | 0.141 | 0.453 | 0.493 | 6114 | tags=25%, list=10%, signal=28% |
| 2104 | POSITIVE\_REGULATION\_OF\_PHOSPHOLIPASE\_C\_ACTIVITY(GO:0010863) |  | 33 | 0.83 | 1.28 | 0.141 | 0.453 | 0.493 | 6928 | tags=39%, list=12%, signal=45% |
| 2105 | SARCOMERE(GO:0030017) |  | 195 | 0.81 | 1.28 | 0.141 | 0.452 | 0.493 | 8328 | tags=30%, list=14%, signal=35% |
| 2106 | REGULATION\_OF\_SIGNALING(GO:0023051) |  | 3514 | 0.81 | 1.28 | 0.141 | 0.452 | 0.493 | 7758 | tags=15%, list=13%, signal=17% |
| 2107 | METANEPHRIC\_GLOMERULUS\_DEVELOPMENT(GO:0072224) |  | 16 | 0.82 | 1.28 | 0.141 | 0.452 | 0.493 | 8107 | tags=44%, list=14%, signal=51% |
| 2108 | POSITIVE\_REGULATION\_OF\_CARTILAGE\_DEVELOPMENT(GO:0061036) |  | 30 | 0.81 | 1.28 | 0.000 | 0.452 | 0.493 | 7407 | tags=30%, list=13%, signal=34% |
| 2109 | POSITIVE\_REGULATION\_OF\_TYPE\_2\_IMMUNE\_RESPONSE(GO:0002830) |  | 15 | 0.83 | 1.28 | 0.141 | 0.452 | 0.493 | 6178 | tags=27%, list=10%, signal=30% |
| 2110 | PLATELET\_ALPHA\_GRANULE(GO:0031091) |  | 91 | 0.79 | 1.28 | 0.141 | 0.452 | 0.493 | 7921 | tags=19%, list=13%, signal=22% |
| 2111 | PEPTIDASE\_REGULATOR\_ACTIVITY(GO:0061134) |  | 216 | 0.78 | 1.28 | 0.141 | 0.452 | 0.493 | 8070 | tags=18%, list=14%, signal=21% |
| 2112 | AXON\_REGENERATION(GO:0031103) |  | 43 | 0.79 | 1.28 | 0.000 | 0.452 | 0.493 | 8743 | tags=30%, list=15%, signal=35% |
| 2113 | INORGANIC\_ION\_HOMEOSTASIS(GO:0098771) |  | 693 | 0.79 | 1.28 | 0.141 | 0.451 | 0.493 | 7727 | tags=18%, list=13%, signal=21% |
| 2114 | CELLULAR\_CHEMICAL\_HOMEOSTASIS(GO:0055082) |  | 753 | 0.77 | 1.28 | 0.141 | 0.451 | 0.493 | 7745 | tags=17%, list=13%, signal=19% |
| 2115 | MIDDLE\_EAR\_MORPHOGENESIS(GO:0042474) |  | 20 | 0.84 | 1.28 | 0.141 | 0.451 | 0.493 | 8177 | tags=45%, list=14%, signal=52% |
| 2116 | CLATHRIN\_BINDING(GO:0030276) |  | 63 | 0.81 | 1.28 | 0.141 | 0.451 | 0.493 | 7426 | tags=21%, list=13%, signal=24% |
| 2117 | CELLULAR\_MACROMOLECULE\_LOCALIZATION(GO:0070727) |  | 1670 | 0.81 | 1.28 | 0.141 | 0.451 | 0.493 | 7184 | tags=7%, list=12%, signal=7% |
| 2118 | SIGNALING\_RECEPTOR\_ACTIVITY(GO:0038023) |  | 1356 | 0.80 | 1.28 | 0.141 | 0.451 | 0.493 | 7981 | tags=18%, list=13%, signal=20% |
| 2119 | MOLECULAR\_TRANSDUCER\_ACTIVITY(GO:0060089) |  | 1356 | 0.80 | 1.28 | 0.141 | 0.451 | 0.493 | 7981 | tags=18%, list=13%, signal=20% |
| 2120 | REGULATION\_OF\_DEVELOPMENTAL\_GROWTH(GO:0048638) |  | 319 | 0.81 | 1.28 | 0.141 | 0.451 | 0.493 | 7583 | tags=20%, list=13%, signal=23% |
| 2121 | CILIARY\_PART(GO:0044441) |  | 424 | 0.78 | 1.28 | 0.000 | 0.451 | 0.493 | 8933 | tags=23%, list=15%, signal=27% |
| 2122 | RESPONSE\_TO\_LIGHT\_STIMULUS(GO:0009416) |  | 304 | 0.77 | 1.28 | 0.000 | 0.451 | 0.493 | 8237 | tags=13%, list=14%, signal=15% |
| 2123 | POSITIVE\_REGULATION\_OF\_CGMP\_METABOLIC\_PROCESS(GO:0030825) |  | 21 | 0.79 | 1.28 | 0.000 | 0.451 | 0.493 | 8279 | tags=33%, list=14%, signal=39% |
| 2124 | AXON\_PART(GO:0033267) |  | 352 | 0.80 | 1.28 | 0.141 | 0.450 | 0.493 | 8030 | tags=20%, list=14%, signal=23% |
| 2125 | NEGATIVE\_REGULATION\_OF\_EPITHELIAL\_TO\_MESENCHYMAL\_TRANSITION(GO:0010719) |  | 30 | 0.80 | 1.28 | 0.141 | 0.450 | 0.493 | 7913 | tags=17%, list=13%, signal=19% |
| 2126 | POSITIVE\_REGULATION\_OF\_TRANSCRIPTION\_BY\_RNA\_POLYMERASE\_II(GO:0045944) |  | 1114 | 0.79 | 1.28 | 0.141 | 0.450 | 0.493 | 8066 | tags=18%, list=14%, signal=21% |
| 2127 | REGULATION\_OF\_CIRCADIAN\_RHYTHM(GO:0042752) |  | 98 | 0.76 | 1.28 | 0.000 | 0.450 | 0.493 | 7414 | tags=13%, list=13%, signal=15% |
| 2128 | REGULATION\_OF\_MACROPHAGE\_DERIVED\_FOAM\_CELL\_DIFFERENTIATION(GO:0010743) |  | 29 | 0.79 | 1.28 | 0.000 | 0.450 | 0.493 | 8474 | tags=24%, list=14%, signal=28% |
| 2129 | REGULATION\_OF\_NEURONAL\_SYNAPTIC\_PLASTICITY(GO:0048168) |  | 52 | 0.78 | 1.28 | 0.000 | 0.450 | 0.493 | 7906 | tags=13%, list=13%, signal=16% |
| 2130 | BEHAVIOR(GO:0007610) |  | 573 | 0.80 | 1.28 | 0.141 | 0.450 | 0.493 | 7981 | tags=24%, list=13%, signal=27% |
| 2131 | REGULATION\_OF\_MITOCHONDRION\_ORGANIZATION(GO:0010821) |  | 222 | 0.80 | 1.28 | 0.141 | 0.450 | 0.493 | 5977 | tags=4%, list=10%, signal=4% |
| 2132 | CONTRACTILE\_FIBER(GO:0043292) |  | 229 | 0.81 | 1.28 | 0.141 | 0.450 | 0.493 | 8328 | tags=28%, list=14%, signal=32% |
| 2133 | CHRONIC\_INFLAMMATORY\_RESPONSE(GO:0002544) |  | 19 | 0.86 | 1.28 | 0.141 | 0.450 | 0.493 | 5959 | tags=11%, list=10%, signal=12% |
| 2134 | REGULATION\_OF\_VASCULAR\_PERMEABILITY(GO:0043114) |  | 34 | 0.78 | 1.27 | 0.000 | 0.450 | 0.659 | 7081 | tags=15%, list=12%, signal=17% |
| 2135 | NEGATIVE\_REGULATION\_OF\_CARDIAC\_MUSCLE\_TISSUE\_GROWTH(GO:0055022) |  | 26 | 0.82 | 1.27 | 0.000 | 0.450 | 0.659 | 8380 | tags=23%, list=14%, signal=27% |
| 2136 | NEGATIVE\_REGULATION\_OF\_HEART\_GROWTH(GO:0061117) |  | 26 | 0.82 | 1.27 | 0.000 | 0.449 | 0.659 | 8380 | tags=23%, list=14%, signal=27% |
| 2137 | AU\_RICH\_ELEMENT\_BINDING(GO:0017091) |  | 25 | 0.79 | 1.27 | 0.000 | 0.449 | 0.659 | 5932 | tags=4%, list=10%, signal=4% |
| 2138 | NEGATIVE\_REGULATION\_OF\_PROTEIN\_CATABOLIC\_PROCESS(GO:0042177) |  | 115 | 0.75 | 1.27 | 0.000 | 0.449 | 0.659 | 7546 | tags=11%, list=13%, signal=13% |
| 2139 | ACTIVATION\_OF\_CYSTEINE\_TYPE\_ENDOPEPTIDASE\_ACTIVITY\_INVOLVED\_IN\_APOPTOTIC\_PROCESS(GO:0006919) |  | 84 | 0.77 | 1.27 | 0.000 | 0.449 | 0.659 | 7754 | tags=12%, list=13%, signal=14% |
| 2140 | POSITIVE\_REGULATION\_OF\_CELLULAR\_METABOLIC\_PROCESS(GO:0031325) |  | 3018 | 0.81 | 1.27 | 0.141 | 0.449 | 0.659 | 7450 | tags=13%, list=13%, signal=14% |
| 2141 | REGULATION\_OF\_RESPONSE\_TO\_BIOTIC\_STIMULUS(GO:0002831) |  | 126 | 0.78 | 1.27 | 0.141 | 0.449 | 0.659 | 7663 | tags=16%, list=13%, signal=18% |
| 2142 | ACTIN\_CYTOSKELETON(GO:0015629) |  | 467 | 0.80 | 1.27 | 0.141 | 0.449 | 0.659 | 7955 | tags=18%, list=13%, signal=21% |
| 2143 | VESICLE\_MEMBRANE(GO:0012506) |  | 747 | 0.78 | 1.27 | 0.141 | 0.449 | 0.659 | 7940 | tags=13%, list=13%, signal=15% |
| 2144 | TRANSCRIPTIONAL\_ACTIVATOR\_ACTIVITY\_\_RNA\_POLYMERASE\_II\_TRANSCRIPTION\_FACTOR\_BINDING(GO:0001190) |  | 60 | 0.81 | 1.27 | 0.141 | 0.449 | 0.659 | 6421 | tags=15%, list=11%, signal=17% |
| 2145 | POSITIVE\_REGULATION\_OF\_GROWTH(GO:0045927) |  | 261 | 0.80 | 1.27 | 0.141 | 0.449 | 0.659 | 7537 | tags=16%, list=13%, signal=18% |
| 2146 | OLFACTORY\_LOBE\_DEVELOPMENT(GO:0021988) |  | 35 | 0.82 | 1.27 | 0.000 | 0.449 | 0.659 | 7668 | tags=31%, list=13%, signal=36% |
| 2147 | EXTRACELLULAR\_REGION(GO:0005576) |  | 4937 | 0.82 | 1.27 | 0.141 | 0.449 | 0.659 | 7467 | tags=13%, list=13%, signal=14% |
| 2148 | NEURON\_MATURATION(GO:0042551) |  | 40 | 0.77 | 1.27 | 0.000 | 0.449 | 0.659 | 8447 | tags=25%, list=14%, signal=29% |
| 2149 | VISUAL\_BEHAVIOR(GO:0007632) |  | 51 | 0.81 | 1.27 | 0.141 | 0.449 | 0.659 | 7102 | tags=14%, list=12%, signal=16% |
| 2150 | CELL\_CELL\_SIGNALING(GO:0007267) |  | 1601 | 0.80 | 1.27 | 0.141 | 0.448 | 0.659 | 7995 | tags=19%, list=14%, signal=22% |
| 2151 | AXON\_CYTOPLASM(GO:1904115) |  | 43 | 0.77 | 1.27 | 0.000 | 0.448 | 0.659 | 6047 | tags=9%, list=10%, signal=10% |
| 2152 | POSITIVE\_REGULATION\_OF\_STEROID\_METABOLIC\_PROCESS(GO:0045940) |  | 23 | 0.83 | 1.27 | 0.000 | 0.448 | 0.659 | 7217 | tags=30%, list=12%, signal=35% |
| 2153 | NEGATIVE\_REGULATION\_OF\_MULTI\_ORGANISM\_PROCESS(GO:0043901) |  | 170 | 0.79 | 1.27 | 0.141 | 0.448 | 0.659 | 7777 | tags=16%, list=13%, signal=19% |
| 2154 | REGULATION\_OF\_RESPONSE\_TO\_WOUNDING(GO:1903034) |  | 162 | 0.80 | 1.27 | 0.141 | 0.448 | 0.659 | 7975 | tags=22%, list=13%, signal=26% |
| 2155 | CATION\_TRANSPORT(GO:0006812) |  | 1115 | 0.78 | 1.27 | 0.141 | 0.448 | 0.659 | 8074 | tags=21%, list=14%, signal=24% |
| 2156 | LIGAND\_DEPENDENT\_NUCLEAR\_RECEPTOR\_BINDING(GO:0016922) |  | 21 | 0.80 | 1.27 | 0.000 | 0.448 | 0.659 | 5958 | tags=10%, list=10%, signal=11% |
| 2157 | PRESYNAPTIC\_MEMBRANE(GO:0042734) |  | 73 | 0.78 | 1.27 | 0.000 | 0.448 | 0.659 | 9105 | tags=32%, list=15%, signal=37% |
| 2158 | GROWTH(GO:0040007) |  | 960 | 0.80 | 1.27 | 0.141 | 0.448 | 0.659 | 7583 | tags=17%, list=13%, signal=19% |
| 2159 | NEGATIVE\_REGULATION\_OF\_NITROGEN\_COMPOUND\_METABOLIC\_PROCESS(GO:0051172) |  | 2548 | 0.80 | 1.27 | 0.141 | 0.448 | 0.659 | 7268 | tags=11%, list=12%, signal=12% |
| 2160 | G\_PROTEIN\_COUPLED\_PURINERGIC\_RECEPTOR\_SIGNALING\_PATHWAY(GO:0035588) |  | 25 | 0.80 | 1.27 | 0.000 | 0.448 | 0.659 | 5955 | tags=12%, list=10%, signal=13% |
| 2161 | MYOFIBRIL(GO:0030016) |  | 217 | 0.80 | 1.27 | 0.141 | 0.448 | 0.659 | 8328 | tags=29%, list=14%, signal=33% |
| 2162 | GLOMERULUS\_VASCULATURE\_DEVELOPMENT(GO:0072012) |  | 22 | 0.84 | 1.27 | 0.141 | 0.447 | 0.659 | 8107 | tags=41%, list=14%, signal=47% |
| 2163 | PROTEIN\_LOCALIZATION\_TO\_SYNAPSE(GO:0035418) |  | 35 | 0.80 | 1.27 | 0.141 | 0.448 | 0.659 | 8247 | tags=17%, list=14%, signal=20% |
| 2164 | BLOOD\_VESSEL\_REMODELING(GO:0001974) |  | 46 | 0.77 | 1.27 | 0.000 | 0.447 | 0.659 | 7343 | tags=17%, list=12%, signal=20% |
| 2165 | REGULATION\_OF\_GONAD\_DEVELOPMENT(GO:1905939) |  | 19 | 0.84 | 1.27 | 0.141 | 0.447 | 0.659 | 6344 | tags=26%, list=11%, signal=29% |
| 2166 | REGULATION\_OF\_WNT\_SIGNALING\_PATHWAY(GO:0030111) |  | 340 | 0.77 | 1.27 | 0.000 | 0.447 | 0.659 | 8122 | tags=17%, list=14%, signal=20% |
| 2167 | PROTEIN\_LIPID\_COMPLEX\_SUBUNIT\_ORGANIZATION(GO:0071825) |  | 47 | 0.77 | 1.27 | 0.000 | 0.447 | 0.659 | 7659 | tags=26%, list=13%, signal=29% |
| 2168 | GLOMERULAR\_FILTRATION(GO:0003094) |  | 20 | 0.86 | 1.27 | 0.141 | 0.447 | 0.659 | 6525 | tags=35%, list=11%, signal=39% |
| 2169 | NEGATIVE\_REGULATION\_OF\_EPITHELIAL\_CELL\_PROLIFERATION(GO:0050680) |  | 155 | 0.80 | 1.27 | 0.141 | 0.447 | 0.659 | 8058 | tags=17%, list=14%, signal=20% |
| 2170 | ORGANELLE\_OUTER\_MEMBRANE(GO:0031968) |  | 206 | 0.77 | 1.27 | 0.000 | 0.447 | 0.659 | 7435 | tags=9%, list=13%, signal=11% |
| 2171 | POSITIVE\_REGULATION\_OF\_BLOOD\_VESSEL\_ENDOTHELIAL\_CELL\_MIGRATION(GO:0043536) |  | 74 | 0.80 | 1.27 | 0.141 | 0.447 | 0.659 | 8046 | tags=20%, list=14%, signal=23% |
| 2172 | CALCIUM\_CHANNEL\_REGULATOR\_ACTIVITY(GO:0005246) |  | 36 | 0.77 | 1.27 | 0.000 | 0.447 | 0.659 | 10556 | tags=39%, list=18%, signal=47% |
| 2173 | FERTILIZATION(GO:0009566) |  | 177 | 0.79 | 1.27 | 0.141 | 0.447 | 0.659 | 8659 | tags=23%, list=15%, signal=26% |
| 2174 | REACTIVE\_OXYGEN\_SPECIES\_BIOSYNTHETIC\_PROCESS(GO:1903409) |  | 100 | 0.78 | 1.27 | 0.141 | 0.447 | 0.659 | 7873 | tags=17%, list=13%, signal=20% |
| 2175 | RENAL\_SYSTEM\_VASCULATURE\_DEVELOPMENT(GO:0061437) |  | 24 | 0.83 | 1.27 | 0.141 | 0.447 | 0.659 | 8107 | tags=38%, list=14%, signal=43% |
| 2176 | KIDNEY\_VASCULATURE\_DEVELOPMENT(GO:0061440) |  | 24 | 0.83 | 1.27 | 0.141 | 0.447 | 0.659 | 8107 | tags=38%, list=14%, signal=43% |
| 2177 | SODIUM\_ION\_HOMEOSTASIS(GO:0055078) |  | 44 | 0.82 | 1.27 | 0.141 | 0.447 | 0.659 | 6525 | tags=18%, list=11%, signal=20% |
| 2178 | BMP\_SIGNALING\_PATHWAY(GO:0030509) |  | 159 | 0.79 | 1.27 | 0.141 | 0.446 | 0.659 | 8177 | tags=27%, list=14%, signal=31% |
| 2179 | REGULATION\_OF\_DOPAMINE\_METABOLIC\_PROCESS(GO:0042053) |  | 19 | 0.84 | 1.27 | 0.141 | 0.446 | 0.659 | 7935 | tags=21%, list=13%, signal=24% |
| 2180 | REGULATION\_OF\_CATECHOLAMINE\_METABOLIC\_PROCESS(GO:0042069) |  | 19 | 0.84 | 1.27 | 0.141 | 0.446 | 0.659 | 7935 | tags=21%, list=13%, signal=24% |
| 2181 | G2\_DNA\_DAMAGE\_CHECKPOINT(GO:0031572) |  | 35 | 0.77 | 1.27 | 0.000 | 0.446 | 0.659 | 10943 | tags=43%, list=19%, signal=53% |
| 2182 | SENSORY\_PERCEPTION\_OF\_LIGHT\_STIMULUS(GO:0050953) |  | 210 | 0.80 | 1.27 | 0.141 | 0.446 | 0.659 | 8182 | tags=29%, list=14%, signal=33% |
| 2183 | REGULATION\_OF\_INTRACELLULAR\_SIGNAL\_TRANSDUCTION(GO:1902531) |  | 1850 | 0.80 | 1.27 | 0.141 | 0.446 | 0.659 | 7732 | tags=14%, list=13%, signal=16% |
| 2184 | VISUAL\_PERCEPTION(GO:0007601) |  | 207 | 0.80 | 1.27 | 0.141 | 0.446 | 0.659 | 8182 | tags=29%, list=14%, signal=33% |
| 2185 | SUPRAMOLECULAR\_POLYMER(GO:0099081) |  | 919 | 0.81 | 1.27 | 0.141 | 0.446 | 0.659 | 7739 | tags=16%, list=13%, signal=19% |
| 2186 | CONNECTIVE\_TISSUE\_DEVELOPMENT(GO:0061448) |  | 242 | 0.79 | 1.27 | 0.141 | 0.446 | 0.659 | 8003 | tags=28%, list=14%, signal=32% |
| 2187 | MICROTUBULE\_POLYMERIZATION(GO:0046785) |  | 71 | 0.74 | 1.27 | 0.000 | 0.446 | 0.659 | 8972 | tags=17%, list=15%, signal=20% |
| 2188 | NEGATIVE\_REGULATION\_OF\_MUSCLE\_ORGAN\_DEVELOPMENT(GO:0048635) |  | 55 | 0.81 | 1.27 | 0.000 | 0.446 | 0.659 | 8774 | tags=29%, list=15%, signal=34% |
| 2189 | REGULATION\_OF\_POSTSYNAPSE\_ORGANIZATION(GO:0099175) |  | 57 | 0.77 | 1.27 | 0.000 | 0.446 | 0.659 | 7487 | tags=21%, list=13%, signal=24% |
| 2190 | SUPRAMOLECULAR\_COMPLEX(GO:0099080) |  | 920 | 0.81 | 1.27 | 0.141 | 0.446 | 0.659 | 7739 | tags=16%, list=13%, signal=19% |
| 2191 | MATERNAL\_PLACENTA\_DEVELOPMENT(GO:0001893) |  | 33 | 0.81 | 1.27 | 0.000 | 0.446 | 0.659 | 7508 | tags=15%, list=13%, signal=17% |
| 2192 | HEMATOPOIETIC\_OR\_LYMPHOID\_ORGAN\_DEVELOPMENT(GO:0048534) |  | 896 | 0.79 | 1.27 | 0.141 | 0.445 | 0.659 | 7583 | tags=15%, list=13%, signal=17% |
| 2193 | SYSTEM\_DEVELOPMENT(GO:0048731) |  | 4732 | 0.83 | 1.27 | 0.141 | 0.445 | 0.659 | 7548 | tags=16%, list=13%, signal=17% |
| 2194 | REGULATION\_OF\_CELL\_MIGRATION\_INVOLVED\_IN\_SPROUTING\_ANGIOGENESIS(GO:0090049) |  | 75 | 0.80 | 1.27 | 0.141 | 0.445 | 0.659 | 9016 | tags=24%, list=15%, signal=28% |
| 2195 | SEQUENCE\_SPECIFIC\_DOUBLE\_STRANDED\_DNA\_BINDING(GO:1990837) |  | 753 | 0.80 | 1.27 | 0.141 | 0.445 | 0.659 | 8038 | tags=20%, list=14%, signal=23% |
| 2196 | POSITIVE\_REGULATION\_OF\_MACROAUTOPHAGY(GO:0016239) |  | 60 | 0.79 | 1.27 | 0.141 | 0.445 | 0.659 | 5861 | tags=5%, list=10%, signal=6% |
| 2197 | NEGATIVE\_REGULATION\_OF\_STRIATED\_MUSCLE\_TISSUE\_DEVELOPMENT(GO:0045843) |  | 54 | 0.81 | 1.27 | 0.000 | 0.445 | 0.659 | 8774 | tags=30%, list=15%, signal=35% |
| 2198 | SMOOTH\_MUSCLE\_TISSUE\_DEVELOPMENT(GO:0048745) |  | 22 | 0.82 | 1.27 | 0.141 | 0.445 | 0.659 | 7360 | tags=18%, list=12%, signal=21% |
| 2199 | SYNAPTIC\_VESICLE\_ENDOCYTOSIS(GO:0048488) |  | 35 | 0.82 | 1.27 | 0.141 | 0.445 | 0.659 | 7721 | tags=26%, list=13%, signal=30% |
| 2200 | CONTRACTILE\_FIBER\_PART(GO:0044449) |  | 212 | 0.80 | 1.27 | 0.141 | 0.445 | 0.659 | 8328 | tags=28%, list=14%, signal=33% |
| 2201 | NEGATIVE\_REGULATION\_OF\_PROTEIN\_POLYMERIZATION(GO:0032272) |  | 64 | 0.76 | 1.27 | 0.000 | 0.445 | 0.659 | 7859 | tags=13%, list=13%, signal=14% |
| 2202 | MITOCHONDRION\_LOCALIZATION(GO:0051646) |  | 41 | 0.74 | 1.27 | 0.000 | 0.445 | 0.659 | 7426 | tags=20%, list=13%, signal=22% |
| 2203 | ION\_TRANSMEMBRANE\_TRANSPORTER\_ACTIVITY(GO:0015075) |  | 908 | 0.79 | 1.27 | 0.141 | 0.445 | 0.659 | 8432 | tags=24%, list=14%, signal=27% |
| 2204 | PLASMA\_LIPOPROTEIN\_PARTICLE\_ORGANIZATION(GO:0071827) |  | 44 | 0.77 | 1.27 | 0.000 | 0.445 | 0.659 | 7659 | tags=25%, list=13%, signal=29% |
| 2205 | REGULATION\_OF\_VIRAL\_GENOME\_REPLICATION(GO:0045069) |  | 87 | 0.80 | 1.27 | 0.141 | 0.445 | 0.659 | 7293 | tags=13%, list=12%, signal=14% |
| 2206 | REGULATION\_OF\_PHOSPHORYLATION(GO:0042325) |  | 1510 | 0.80 | 1.27 | 0.141 | 0.445 | 0.659 | 7702 | tags=14%, list=13%, signal=15% |
| 2207 | NOTCH\_BINDING(GO:0005112) |  | 21 | 0.81 | 1.27 | 0.141 | 0.445 | 0.659 | 6846 | tags=10%, list=12%, signal=11% |
| 2208 | RNA\_BINDING(GO:0003723) |  | 1845 | 0.78 | 1.27 | 0.000 | 0.445 | 0.659 | 6528 | tags=4%, list=11%, signal=4% |
| 2209 | THYROID\_GLAND\_DEVELOPMENT(GO:0030878) |  | 26 | 0.83 | 1.27 | 0.000 | 0.444 | 0.659 | 7343 | tags=35%, list=12%, signal=40% |
| 2210 | PHOSPHATIDYLINOSITOL\_3\_4\_5\_TRISPHOSPHATE\_BINDING(GO:0005547) |  | 37 | 0.77 | 1.27 | 0.000 | 0.444 | 0.659 | 9358 | tags=22%, list=16%, signal=26% |
| 2211 | MONONUCLEAR\_CELL\_MIGRATION(GO:0071674) |  | 79 | 0.81 | 1.27 | 0.141 | 0.444 | 0.659 | 7634 | tags=18%, list=13%, signal=20% |
| 2212 | REGULATION\_OF\_ANIMAL\_ORGAN\_FORMATION(GO:0003156) |  | 34 | 0.85 | 1.27 | 0.141 | 0.444 | 0.659 | 6473 | tags=21%, list=11%, signal=23% |
| 2213 | OUTER\_MEMBRANE(GO:0019867) |  | 209 | 0.77 | 1.27 | 0.000 | 0.444 | 0.659 | 7435 | tags=9%, list=13%, signal=10% |
| 2214 | MONOVALENT\_INORGANIC\_CATION\_TRANSPORT(GO:0015672) |  | 526 | 0.78 | 1.27 | 0.141 | 0.444 | 0.659 | 8627 | tags=23%, list=15%, signal=27% |
| 2215 | ACTIN\_CYTOSKELETON\_ORGANIZATION(GO:0030036) |  | 609 | 0.80 | 1.27 | 0.141 | 0.444 | 0.659 | 7976 | tags=17%, list=13%, signal=20% |
| 2216 | NUCLEOTIDE\_METABOLIC\_PROCESS(GO:0009117) |  | 685 | 0.76 | 1.27 | 0.000 | 0.444 | 0.659 | 7559 | tags=10%, list=13%, signal=12% |
| 2217 | RESPONSE\_TO\_CATECHOLAMINE(GO:0071869) |  | 38 | 0.82 | 1.27 | 0.141 | 0.444 | 0.659 | 7426 | tags=18%, list=13%, signal=21% |
| 2218 | RESPONSE\_TO\_MONOAMINE(GO:0071867) |  | 38 | 0.82 | 1.27 | 0.141 | 0.444 | 0.659 | 7426 | tags=18%, list=13%, signal=21% |
| 2219 | EXTRINSIC\_APOPTOTIC\_SIGNALING\_PATHWAY\_IN\_ABSENCE\_OF\_LIGAND(GO:0097192) |  | 73 | 0.78 | 1.27 | 0.000 | 0.444 | 0.659 | 9857 | tags=21%, list=17%, signal=25% |
| 2220 | SIGNAL\_TRANSDUCTION\_IN\_ABSENCE\_OF\_LIGAND(GO:0038034) |  | 73 | 0.78 | 1.27 | 0.000 | 0.444 | 0.659 | 9857 | tags=21%, list=17%, signal=25% |
| 2221 | DETECTION\_OF\_MECHANICAL\_STIMULUS\_INVOLVED\_IN\_SENSORY\_PERCEPTION(GO:0050974) |  | 27 | 0.80 | 1.27 | 0.000 | 0.444 | 0.659 | 9354 | tags=48%, list=16%, signal=57% |
| 2222 | NEGATIVE\_REGULATION\_OF\_NEUROTRANSMITTER\_TRANSPORT(GO:0051589) |  | 20 | 0.83 | 1.27 | 0.000 | 0.444 | 0.659 | 7293 | tags=30%, list=12%, signal=34% |
| 2223 | FEMALE\_SEX\_DIFFERENTIATION(GO:0046660) |  | 115 | 0.79 | 1.27 | 0.141 | 0.443 | 0.659 | 7547 | tags=23%, list=13%, signal=26% |
| 2224 | REGULATION\_OF\_BODY\_FLUID\_LEVELS(GO:0050878) |  | 505 | 0.79 | 1.27 | 0.141 | 0.443 | 0.659 | 7546 | tags=15%, list=13%, signal=18% |
| 2225 | ANTIPORTER\_ACTIVITY(GO:0015297) |  | 80 | 0.77 | 1.27 | 0.000 | 0.443 | 0.659 | 10298 | tags=36%, list=17%, signal=44% |
| 2226 | SPERM\_MOTILITY(GO:0097722) |  | 87 | 0.77 | 1.27 | 0.141 | 0.443 | 0.828 | 10309 | tags=34%, list=17%, signal=42% |
| 2227 | NEGATIVE\_REGULATION\_OF\_WOUND\_HEALING(GO:0061045) |  | 70 | 0.83 | 1.27 | 0.141 | 0.443 | 0.828 | 7916 | tags=27%, list=13%, signal=31% |
| 2228 | REGULATION\_OF\_CELL\_DIFFERENTIATION(GO:0045595) |  | 1750 | 0.80 | 1.27 | 0.141 | 0.443 | 0.828 | 7546 | tags=16%, list=13%, signal=18% |
| 2229 | NEGATIVE\_REGULATION\_OF\_B\_CELL\_ACTIVATION(GO:0050869) |  | 31 | 0.77 | 1.27 | 0.141 | 0.443 | 0.828 | 10693 | tags=23%, list=18%, signal=28% |
| 2230 | TRANSCRIPTIONAL\_ACTIVATOR\_ACTIVITY\_\_RNA\_POLYMERASE\_II\_PROXIMAL\_PROMOTER\_SEQUENCE\_SPECIFIC\_DNA\_BINDING(GO:0001077) |  | 275 | 0.80 | 1.27 | 0.141 | 0.443 | 0.828 | 7358 | tags=20%, list=12%, signal=23% |
| 2231 | RESPONSE\_TO\_MECHANICAL\_STIMULUS(GO:0009612) |  | 213 | 0.80 | 1.27 | 0.141 | 0.443 | 0.828 | 7652 | tags=21%, list=13%, signal=24% |
| 2232 | INTERMEDIATE\_FILAMENT\_CYTOSKELETON(GO:0045111) |  | 240 | 0.82 | 1.27 | 0.141 | 0.443 | 0.828 | 5959 | tags=8%, list=10%, signal=9% |
| 2233 | NEGATIVE\_REGULATION\_OF\_STEROL\_TRANSPORT(GO:0032372) |  | 20 | 0.85 | 1.27 | 0.141 | 0.443 | 0.828 | 6197 | tags=25%, list=10%, signal=28% |
| 2234 | NEGATIVE\_REGULATION\_OF\_CHOLESTEROL\_TRANSPORT(GO:0032375) |  | 20 | 0.85 | 1.27 | 0.141 | 0.443 | 0.828 | 6197 | tags=25%, list=10%, signal=28% |
| 2235 | BEHAVIORAL\_FEAR\_RESPONSE(GO:0001662) |  | 34 | 0.81 | 1.27 | 0.141 | 0.443 | 0.828 | 7298 | tags=21%, list=12%, signal=23% |
| 2236 | BEHAVIORAL\_DEFENSE\_RESPONSE(GO:0002209) |  | 34 | 0.81 | 1.27 | 0.141 | 0.442 | 0.828 | 7298 | tags=21%, list=12%, signal=23% |
| 2237 | NEGATIVE\_REGULATION\_OF\_STRESS\_FIBER\_ASSEMBLY(GO:0051497) |  | 22 | 0.84 | 1.27 | 0.141 | 0.442 | 0.828 | 5903 | tags=18%, list=10%, signal=20% |
| 2238 | NEURON\_FATE\_SPECIFICATION(GO:0048665) |  | 34 | 0.88 | 1.27 | 0.141 | 0.442 | 0.828 | 6058 | tags=53%, list=10%, signal=59% |
| 2239 | VOCALIZATION\_BEHAVIOR(GO:0071625) |  | 18 | 0.86 | 1.27 | 0.141 | 0.442 | 0.828 | 7962 | tags=44%, list=13%, signal=51% |
| 2240 | CELLULAR\_CARBOHYDRATE\_METABOLIC\_PROCESS(GO:0044262) |  | 250 | 0.78 | 1.27 | 0.141 | 0.442 | 0.828 | 7414 | tags=11%, list=13%, signal=13% |
| 2241 | NEGATIVE\_REGULATION\_OF\_RESPONSE\_TO\_WOUNDING(GO:1903035) |  | 83 | 0.82 | 1.27 | 0.141 | 0.442 | 0.828 | 7975 | tags=28%, list=13%, signal=32% |
| 2242 | POSITIVE\_REGULATION\_OF\_METABOLIC\_PROCESS(GO:0009893) |  | 3268 | 0.81 | 1.27 | 0.141 | 0.442 | 0.828 | 7756 | tags=14%, list=13%, signal=15% |
| 2243 | POSITIVE\_REGULATION\_OF\_CELLULAR\_CARBOHYDRATE\_METABOLIC\_PROCESS(GO:0010676) |  | 54 | 0.81 | 1.27 | 0.141 | 0.442 | 0.828 | 7272 | tags=13%, list=12%, signal=15% |
| 2244 | ROOF\_OF\_MOUTH\_DEVELOPMENT(GO:0060021) |  | 88 | 0.80 | 1.27 | 0.141 | 0.442 | 0.828 | 8177 | tags=31%, list=14%, signal=36% |
| 2245 | BILE\_ACID\_METABOLIC\_PROCESS(GO:0008206) |  | 47 | 0.81 | 1.27 | 0.000 | 0.441 | 0.828 | 7491 | tags=26%, list=13%, signal=29% |
| 2246 | EYE\_PHOTORECEPTOR\_CELL\_DIFFERENTIATION(GO:0001754) |  | 45 | 0.82 | 1.27 | 0.141 | 0.441 | 0.828 | 8035 | tags=29%, list=14%, signal=33% |
| 2247 | REGULATION\_OF\_PLATELET\_ACTIVATION(GO:0010543) |  | 31 | 0.85 | 1.27 | 0.141 | 0.441 | 0.828 | 7546 | tags=29%, list=13%, signal=33% |
| 2248 | REGULATION\_OF\_CALCIUM\_MEDIATED\_SIGNALING(GO:0050848) |  | 79 | 0.80 | 1.27 | 0.141 | 0.441 | 0.828 | 7544 | tags=22%, list=13%, signal=25% |
| 2249 | BONE\_DEVELOPMENT(GO:0060348) |  | 186 | 0.78 | 1.27 | 0.000 | 0.441 | 0.828 | 8445 | tags=24%, list=14%, signal=28% |
| 2250 | POSITIVE\_REGULATION\_OF\_INSULIN\_SECRETION(GO:0032024) |  | 71 | 0.76 | 1.27 | 0.000 | 0.441 | 0.828 | 5959 | tags=14%, list=10%, signal=16% |
| 2251 | VASCULAR\_ENDOTHELIAL\_GROWTH\_FACTOR\_PRODUCTION(GO:0010573) |  | 34 | 0.82 | 1.27 | 0.141 | 0.441 | 0.828 | 7414 | tags=24%, list=13%, signal=27% |
| 2252 | FOREBRAIN\_REGIONALIZATION(GO:0021871) |  | 25 | 0.84 | 1.27 | 0.141 | 0.441 | 0.828 | 7343 | tags=28%, list=12%, signal=32% |
| 2253 | NEGATIVE\_REGULATION\_OF\_CELLULAR\_METABOLIC\_PROCESS(GO:0031324) |  | 2696 | 0.80 | 1.27 | 0.141 | 0.441 | 0.828 | 7260 | tags=11%, list=12%, signal=12% |
| 2254 | REGULATION\_OF\_RESPONSE\_TO\_STIMULUS(GO:0048583) |  | 4146 | 0.81 | 1.27 | 0.141 | 0.441 | 0.828 | 7732 | tags=15%, list=13%, signal=16% |
| 2255 | SMALL\_MOLECULE\_METABOLIC\_PROCESS(GO:0044281) |  | 2144 | 0.79 | 1.27 | 0.141 | 0.441 | 0.828 | 7516 | tags=12%, list=13%, signal=13% |
| 2256 | REGULATION\_OF\_SYSTEMIC\_ARTERIAL\_BLOOD\_PRESSURE\_BY\_HORMONE(GO:0001990) |  | 40 | 0.80 | 1.27 | 0.141 | 0.441 | 0.828 | 7711 | tags=28%, list=13%, signal=32% |
| 2257 | INTEGRAL\_COMPONENT\_OF\_MITOCHONDRIAL\_OUTER\_MEMBRANE(GO:0031307) |  | 23 | 0.76 | 1.27 | 0.000 | 0.440 | 0.828 | 5977 | tags=4%, list=10%, signal=5% |
| 2258 | CARBOXYLIC\_ACID\_TRANSPORT(GO:0046942) |  | 315 | 0.79 | 1.27 | 0.141 | 0.440 | 0.828 | 8074 | tags=21%, list=14%, signal=24% |
| 2259 | ORGANIC\_ACID\_TRANSPORT(GO:0015849) |  | 315 | 0.79 | 1.27 | 0.141 | 0.440 | 0.828 | 8074 | tags=21%, list=14%, signal=24% |
| 2260 | INWARD\_RECTIFIER\_POTASSIUM\_CHANNEL\_ACTIVITY(GO:0005242) |  | 21 | 0.82 | 1.27 | 0.141 | 0.440 | 0.828 | 9364 | tags=48%, list=16%, signal=57% |
| 2261 | LONG\_TERM\_SYNAPTIC\_POTENTIATION(GO:0060291) |  | 67 | 0.81 | 1.27 | 0.141 | 0.440 | 0.828 | 7738 | tags=21%, list=13%, signal=24% |
| 2262 | CELLULAR\_RESPONSE\_TO\_VITAMIN\_D(GO:0071305) |  | 20 | 0.79 | 1.27 | 0.000 | 0.440 | 0.828 | 6495 | tags=20%, list=11%, signal=22% |
| 2263 | FIBRINOLYSIS(GO:0042730) |  | 27 | 0.85 | 1.27 | 0.141 | 0.440 | 0.828 | 7916 | tags=30%, list=13%, signal=34% |
| 2264 | NEGATIVE\_REGULATION\_OF\_HEMOSTASIS(GO:1900047) |  | 51 | 0.83 | 1.27 | 0.141 | 0.440 | 0.828 | 7916 | tags=29%, list=13%, signal=34% |
| 2265 | NEGATIVE\_REGULATION\_OF\_BLOOD\_COAGULATION(GO:0030195) |  | 51 | 0.83 | 1.27 | 0.141 | 0.440 | 0.828 | 7916 | tags=29%, list=13%, signal=34% |
| 2266 | POSITIVE\_REGULATION\_OF\_CYTOKINE\_SECRETION(GO:0050715) |  | 117 | 0.77 | 1.27 | 0.141 | 0.440 | 0.828 | 7640 | tags=16%, list=13%, signal=19% |
| 2267 | NEGATIVE\_REGULATION\_OF\_INTRACELLULAR\_PROTEIN\_TRANSPORT(GO:0090317) |  | 39 | 0.80 | 1.27 | 0.000 | 0.440 | 0.828 | 7426 | tags=13%, list=13%, signal=15% |
| 2268 | MACROPHAGE\_ACTIVATION(GO:0042116) |  | 66 | 0.80 | 1.27 | 0.141 | 0.440 | 0.828 | 8894 | tags=24%, list=15%, signal=29% |
| 2269 | NEUROTRANSMITTER:SODIUM\_SYMPORTER\_ACTIVITY(GO:0005328) |  | 28 | 0.83 | 1.27 | 0.141 | 0.440 | 0.828 | 8074 | tags=39%, list=14%, signal=45% |
| 2270 | NEGATIVE\_REGULATION\_OF\_TRANSCRIPTION\_\_DNA\_TEMPLATED(GO:0045892) |  | 1172 | 0.80 | 1.27 | 0.141 | 0.440 | 0.828 | 7364 | tags=13%, list=12%, signal=15% |
| 2271 | NEGATIVE\_REGULATION\_OF\_SYNAPSE\_ORGANIZATION(GO:1905809) |  | 22 | 0.82 | 1.27 | 0.000 | 0.440 | 0.828 | 8237 | tags=45%, list=14%, signal=53% |
| 2272 | SECRETORY\_GRANULE\_MEMBRANE(GO:0030667) |  | 296 | 0.76 | 1.27 | 0.000 | 0.440 | 0.828 | 7932 | tags=13%, list=13%, signal=15% |
| 2273 | POSITIVE\_REGULATION\_OF\_STRIATED\_MUSCLE\_TISSUE\_DEVELOPMENT(GO:0045844) |  | 83 | 0.79 | 1.27 | 0.141 | 0.440 | 0.828 | 8774 | tags=24%, list=15%, signal=28% |
| 2274 | POSITIVE\_REGULATION\_OF\_MUSCLE\_ORGAN\_DEVELOPMENT(GO:0048636) |  | 83 | 0.79 | 1.27 | 0.141 | 0.440 | 0.828 | 8774 | tags=24%, list=15%, signal=28% |
| 2275 | MAMMARY\_GLAND\_MORPHOGENESIS(GO:0060443) |  | 46 | 0.82 | 1.27 | 0.141 | 0.439 | 0.828 | 7435 | tags=13%, list=13%, signal=15% |
| 2276 | MUCOPOLYSACCHARIDE\_METABOLIC\_PROCESS(GO:1903510) |  | 113 | 0.79 | 1.27 | 0.141 | 0.439 | 0.828 | 9031 | tags=34%, list=15%, signal=40% |
| 2277 | MECHANORECEPTOR\_DIFFERENTIATION(GO:0042490) |  | 65 | 0.76 | 1.27 | 0.000 | 0.439 | 0.828 | 9354 | tags=28%, list=16%, signal=33% |
| 2278 | CELLULAR\_RESPONSE\_TO\_CALCIUM\_ION(GO:0071277) |  | 61 | 0.78 | 1.27 | 0.141 | 0.439 | 0.828 | 7134 | tags=15%, list=12%, signal=17% |
| 2279 | REGULATORY\_REGION\_NUCLEIC\_ACID\_BINDING(GO:0001067) |  | 850 | 0.80 | 1.27 | 0.141 | 0.439 | 0.828 | 8038 | tags=19%, list=14%, signal=22% |
| 2280 | TRANSCRIPTION\_REGULATORY\_REGION\_DNA\_BINDING(GO:0044212) |  | 849 | 0.80 | 1.27 | 0.141 | 0.439 | 0.828 | 8038 | tags=19%, list=14%, signal=22% |
| 2281 | MAMMARY\_GLAND\_LOBULE\_DEVELOPMENT(GO:0061377) |  | 20 | 0.85 | 1.27 | 0.141 | 0.439 | 0.828 | 6923 | tags=25%, list=12%, signal=28% |
| 2282 | MAMMARY\_GLAND\_ALVEOLUS\_DEVELOPMENT(GO:0060749) |  | 20 | 0.85 | 1.27 | 0.141 | 0.439 | 0.828 | 6923 | tags=25%, list=12%, signal=28% |
| 2283 | CILIUM\_MOVEMENT(GO:0003341) |  | 69 | 0.76 | 1.27 | 0.000 | 0.439 | 0.828 | 8933 | tags=35%, list=15%, signal=41% |
| 2284 | REGULATION\_OF\_GENE\_EXPRESSION\_\_EPIGENETIC(GO:0040029) |  | 496 | 0.78 | 1.27 | 0.141 | 0.439 | 0.828 | 5858 | tags=6%, list=10%, signal=6% |
| 2285 | NEUROTRANSMITTER\_REUPTAKE(GO:0098810) |  | 17 | 0.86 | 1.27 | 0.141 | 0.439 | 0.828 | 7298 | tags=29%, list=12%, signal=34% |
| 2286 | ENDOPLASMIC\_RETICULUM(GO:0005783) |  | 1818 | 0.78 | 1.27 | 0.141 | 0.439 | 0.828 | 7807 | tags=12%, list=13%, signal=13% |
| 2287 | TRANSMEMBRANE\_TRANSPORTER\_ACTIVITY(GO:0022857) |  | 1078 | 0.79 | 1.27 | 0.141 | 0.439 | 0.828 | 8432 | tags=23%, list=14%, signal=26% |
| 2288 | LUNG\_EPITHELIUM\_DEVELOPMENT(GO:0060428) |  | 35 | 0.81 | 1.27 | 0.141 | 0.439 | 0.828 | 6338 | tags=11%, list=11%, signal=13% |
| 2289 | OTIC\_VESICLE\_DEVELOPMENT(GO:0071599) |  | 15 | 0.85 | 1.27 | 0.141 | 0.439 | 0.828 | 7515 | tags=33%, list=13%, signal=38% |
| 2290 | NAD\_BIOSYNTHESIS\_VIA\_NICOTINAMIDE\_RIBOSIDE\_SALVAGE\_PATHWAY(GO:0034356) |  | 15 | 0.81 | 1.27 | 0.141 | 0.438 | 0.828 | 6424 | tags=13%, list=11%, signal=15% |
| 2291 | OLFACTORY\_BULB\_DEVELOPMENT(GO:0021772) |  | 34 | 0.82 | 1.27 | 0.000 | 0.438 | 0.828 | 7668 | tags=32%, list=13%, signal=37% |
| 2292 | PATTERN\_RECOGNITION\_RECEPTOR\_ACTIVITY(GO:0038187) |  | 17 | 0.77 | 1.27 | 0.000 | 0.438 | 0.828 | 9039 | tags=12%, list=15%, signal=14% |
| 2293 | SIGNALING\_PATTERN\_RECOGNITION\_RECEPTOR\_ACTIVITY(GO:0008329) |  | 17 | 0.77 | 1.27 | 0.000 | 0.438 | 0.828 | 9039 | tags=12%, list=15%, signal=14% |
| 2294 | POSITIVE\_REGULATION\_OF\_RELEASE\_OF\_SEQUESTERED\_CALCIUM\_ION\_INTO\_CYTOSOL(GO:0051281) |  | 38 | 0.82 | 1.27 | 0.141 | 0.438 | 0.828 | 8627 | tags=29%, list=15%, signal=34% |
| 2295 | ACTIN\_NUCLEATION(GO:0045010) |  | 49 | 0.78 | 1.27 | 0.141 | 0.438 | 0.828 | 7737 | tags=14%, list=13%, signal=16% |
| 2296 | CELL\_MORPHOGENESIS\_INVOLVED\_IN\_DIFFERENTIATION(GO:0000904) |  | 680 | 0.80 | 1.27 | 0.141 | 0.438 | 0.828 | 7975 | tags=20%, list=13%, signal=23% |
| 2297 | REGULATION\_OF\_PHOSPHOLIPID\_METABOLIC\_PROCESS(GO:1903725) |  | 74 | 0.79 | 1.27 | 0.141 | 0.438 | 0.828 | 8046 | tags=22%, list=14%, signal=25% |
| 2298 | SYMPORTER\_ACTIVITY(GO:0015293) |  | 147 | 0.77 | 1.27 | 0.141 | 0.438 | 0.828 | 8716 | tags=27%, list=15%, signal=32% |
| 2299 | POSITIVE\_REGULATION\_OF\_INTRACELLULAR\_SIGNAL\_TRANSDUCTION(GO:1902533) |  | 989 | 0.79 | 1.27 | 0.141 | 0.438 | 0.828 | 7754 | tags=15%, list=13%, signal=17% |
| 2300 | POSITIVE\_REGULATION\_OF\_MUSCLE\_TISSUE\_DEVELOPMENT(GO:1901863) |  | 84 | 0.79 | 1.27 | 0.141 | 0.438 | 0.828 | 8774 | tags=24%, list=15%, signal=28% |
| 2301 | REGULATION\_OF\_DNA\_REPAIR(GO:0006282) |  | 99 | 0.82 | 1.27 | 0.141 | 0.438 | 0.828 | 6957 | tags=10%, list=12%, signal=11% |
| 2302 | BLASTODERM\_SEGMENTATION(GO:0007350) |  | 18 | 0.83 | 1.27 | 0.141 | 0.438 | 0.828 | 6473 | tags=28%, list=11%, signal=31% |
| 2303 | POSITIVE\_REGULATION\_OF\_RESPONSE\_TO\_WOUNDING(GO:1903036) |  | 67 | 0.80 | 1.27 | 0.141 | 0.437 | 0.828 | 7544 | tags=18%, list=13%, signal=21% |
| 2304 | POSITIVE\_REGULATION\_OF\_NEURON\_DEATH(GO:1901216) |  | 80 | 0.79 | 1.27 | 0.000 | 0.437 | 0.828 | 7963 | tags=16%, list=13%, signal=19% |
| 2305 | REGULATION\_OF\_MICROTUBULE\_BASED\_PROCESS(GO:0032886) |  | 195 | 0.74 | 1.27 | 0.000 | 0.437 | 0.828 | 8972 | tags=15%, list=15%, signal=18% |
| 2306 | REGULATION\_OF\_PHOSPHOLIPASE\_C\_ACTIVITY(GO:1900274) |  | 35 | 0.83 | 1.27 | 0.141 | 0.437 | 0.828 | 6928 | tags=37%, list=12%, signal=42% |
| 2307 | L\_AMINO\_ACID\_TRANSMEMBRANE\_TRANSPORTER\_ACTIVITY(GO:0015179) |  | 53 | 0.76 | 1.27 | 0.000 | 0.437 | 0.828 | 8897 | tags=23%, list=15%, signal=27% |
| 2308 | NEGATIVE\_REGULATION\_OF\_NUCLEIC\_ACID\_TEMPLATED\_TRANSCRIPTION(GO:1903507) |  | 1215 | 0.80 | 1.27 | 0.141 | 0.437 | 0.828 | 7404 | tags=13%, list=13%, signal=15% |
| 2309 | PROTEIN\_DOMAIN\_SPECIFIC\_BINDING(GO:0019904) |  | 706 | 0.78 | 1.27 | 0.141 | 0.437 | 0.828 | 7963 | tags=13%, list=13%, signal=15% |
| 2310 | PROSTAGLANDIN\_BIOSYNTHETIC\_PROCESS(GO:0001516) |  | 23 | 0.80 | 1.27 | 0.141 | 0.437 | 0.828 | 7682 | tags=30%, list=13%, signal=35% |
| 2311 | PROSTANOID\_BIOSYNTHETIC\_PROCESS(GO:0046457) |  | 23 | 0.80 | 1.27 | 0.141 | 0.437 | 0.828 | 7682 | tags=30%, list=13%, signal=35% |
| 2312 | T\_HELPER\_1\_TYPE\_IMMUNE\_RESPONSE(GO:0042088) |  | 40 | 0.80 | 1.27 | 0.000 | 0.437 | 0.828 | 6738 | tags=28%, list=11%, signal=31% |
| 2313 | UBIQUITIN\_PROTEIN\_LIGASE\_ACTIVITY(GO:0061630) |  | 221 | 0.75 | 1.27 | 0.000 | 0.437 | 0.828 | 8341 | tags=13%, list=14%, signal=15% |
| 2314 | NEGATIVE\_REGULATION\_OF\_COAGULATION(GO:0050819) |  | 57 | 0.83 | 1.27 | 0.141 | 0.437 | 0.828 | 7916 | tags=28%, list=13%, signal=32% |
| 2315 | L\_ALPHA\_AMINO\_ACID\_TRANSMEMBRANE\_TRANSPORT(GO:1902475) |  | 43 | 0.77 | 1.27 | 0.000 | 0.436 | 0.828 | 9357 | tags=23%, list=16%, signal=28% |
| 2316 | IMMUNE\_SYSTEM\_DEVELOPMENT(GO:0002520) |  | 937 | 0.79 | 1.27 | 0.141 | 0.436 | 0.828 | 7583 | tags=15%, list=13%, signal=17% |
| 2317 | REGULATION\_OF\_NEUROTRANSMITTER\_LEVELS(GO:0001505) |  | 303 | 0.77 | 1.27 | 0.141 | 0.436 | 0.828 | 8554 | tags=23%, list=14%, signal=26% |
| 2318 | NEGATIVE\_REGULATION\_OF\_ENDOTHELIAL\_CELL\_APOPTOTIC\_PROCESS(GO:2000352) |  | 31 | 0.82 | 1.27 | 0.141 | 0.436 | 0.828 | 6186 | tags=19%, list=10%, signal=22% |
| 2319 | NEGATIVE\_REGULATION\_OF\_RNA\_BIOSYNTHETIC\_PROCESS(GO:1902679) |  | 1217 | 0.80 | 1.27 | 0.141 | 0.436 | 0.828 | 7404 | tags=13%, list=13%, signal=15% |
| 2320 | PHAGOCYTIC\_VESICLE\_MEMBRANE(GO:0030670) |  | 70 | 0.78 | 1.27 | 0.141 | 0.436 | 0.828 | 7409 | tags=13%, list=13%, signal=15% |
| 2321 | NEGATIVE\_REGULATION\_OF\_DEPHOSPHORYLATION(GO:0035305) |  | 103 | 0.76 | 1.27 | 0.000 | 0.436 | 0.828 | 7544 | tags=14%, list=13%, signal=16% |
| 2322 | REGULATION\_OF\_MONOOXYGENASE\_ACTIVITY(GO:0032768) |  | 57 | 0.77 | 1.27 | 0.141 | 0.436 | 0.828 | 6453 | tags=11%, list=11%, signal=12% |
| 2323 | NEGATIVE\_REGULATION\_OF\_TRANSMEMBRANE\_RECEPTOR\_PROTEIN\_SERINE\_THREONINE\_KINASE\_SIGNALING\_PATHWAY(GO:0090101) |  | 116 | 0.77 | 1.27 | 0.000 | 0.436 | 0.828 | 7396 | tags=18%, list=13%, signal=21% |
| 2324 | CELLULAR\_RESPONSE\_TO\_PH(GO:0071467) |  | 22 | 0.80 | 1.27 | 0.141 | 0.436 | 0.828 | 8066 | tags=36%, list=14%, signal=42% |
| 2325 | REGULATION\_OF\_UBIQUITIN\_PROTEIN\_LIGASE\_ACTIVITY(GO:1904666) |  | 21 | 0.75 | 1.27 | 0.000 | 0.436 | 0.828 | 10469 | tags=33%, list=18%, signal=40% |
| 2326 | INORGANIC\_CATION\_TRANSMEMBRANE\_TRANSPORTER\_ACTIVITY(GO:0022890) |  | 611 | 0.77 | 1.27 | 0.141 | 0.435 | 0.828 | 9628 | tags=28%, list=16%, signal=34% |
| 2327 | NEGATIVE\_REGULATION\_OF\_INTRACELLULAR\_TRANSPORT(GO:0032387) |  | 61 | 0.78 | 1.27 | 0.000 | 0.435 | 0.828 | 7426 | tags=15%, list=13%, signal=17% |
| 2328 | FACE\_DEVELOPMENT(GO:0060324) |  | 49 | 0.81 | 1.27 | 0.141 | 0.435 | 0.828 | 8177 | tags=24%, list=14%, signal=28% |
| 2329 | NEURON\_DEVELOPMENT(GO:0048666) |  | 1045 | 0.80 | 1.27 | 0.141 | 0.435 | 0.828 | 8119 | tags=20%, list=14%, signal=22% |
| 2330 | DSRNA\_FRAGMENTATION(GO:0031050) |  | 44 | 0.83 | 1.27 | 0.141 | 0.435 | 0.828 | 5858 | tags=5%, list=10%, signal=5% |
| 2331 | PRODUCTION\_OF\_SMALL\_RNA\_INVOLVED\_IN\_GENE\_SILENCING\_BY\_RNA(GO:0070918) |  | 44 | 0.83 | 1.27 | 0.141 | 0.435 | 0.828 | 5858 | tags=5%, list=10%, signal=5% |
| 2332 | PROTEIN\_KINASE\_ACTIVITY(GO:0004672) |  | 674 | 0.76 | 1.27 | 0.000 | 0.435 | 0.828 | 8115 | tags=18%, list=14%, signal=20% |
| 2333 | AMINO\_ACID:SODIUM\_SYMPORTER\_ACTIVITY(GO:0005283) |  | 18 | 0.81 | 1.27 | 0.141 | 0.435 | 0.828 | 7352 | tags=44%, list=12%, signal=51% |
| 2334 | ION\_TRANSPORT(GO:0006811) |  | 1620 | 0.79 | 1.27 | 0.141 | 0.435 | 0.828 | 8485 | tags=21%, list=14%, signal=24% |
| 2335 | CORTICAL\_ACTIN\_CYTOSKELETON(GO:0030864) |  | 63 | 0.82 | 1.27 | 0.141 | 0.435 | 0.828 | 6581 | tags=16%, list=11%, signal=18% |
| 2336 | CATION\_TRANSMEMBRANE\_TRANSPORTER\_ACTIVITY(GO:0008324) |  | 672 | 0.77 | 1.27 | 0.141 | 0.435 | 0.828 | 8420 | tags=24%, list=14%, signal=27% |
| 2337 | CAMERA\_TYPE\_EYE\_MORPHOGENESIS(GO:0048593) |  | 110 | 0.81 | 1.27 | 0.141 | 0.435 | 0.828 | 8122 | tags=26%, list=14%, signal=31% |
| 2338 | ACTIVATING\_TRANSCRIPTION\_FACTOR\_BINDING(GO:0033613) |  | 67 | 0.76 | 1.27 | 0.000 | 0.435 | 0.828 | 7356 | tags=13%, list=12%, signal=15% |
| 2339 | GLYCOSAMINOGLYCAN\_BIOSYNTHETIC\_PROCESS(GO:0006024) |  | 106 | 0.79 | 1.27 | 0.141 | 0.435 | 0.828 | 9277 | tags=30%, list=16%, signal=36% |
| 2340 | ANIMAL\_ORGAN\_FORMATION(GO:0048645) |  | 61 | 0.83 | 1.27 | 0.141 | 0.435 | 0.828 | 6473 | tags=16%, list=11%, signal=18% |
| 2341 | REGULATION\_OF\_ENDOTHELIAL\_CELL\_PROLIFERATION(GO:0001936) |  | 161 | 0.80 | 1.27 | 0.141 | 0.435 | 0.828 | 8122 | tags=20%, list=14%, signal=24% |
| 2342 | BONE\_MINERALIZATION(GO:0030282) |  | 100 | 0.78 | 1.27 | 0.141 | 0.434 | 0.828 | 7837 | tags=26%, list=13%, signal=30% |
| 2343 | SEGMENT\_SPECIFICATION(GO:0007379) |  | 17 | 0.82 | 1.27 | 0.000 | 0.434 | 0.828 | 7846 | tags=35%, list=13%, signal=41% |
| 2344 | CELLULAR\_RESPONSE\_TO\_BMP\_STIMULUS(GO:0071773) |  | 171 | 0.78 | 1.27 | 0.141 | 0.434 | 0.828 | 8177 | tags=27%, list=14%, signal=31% |
| 2345 | RESPONSE\_TO\_BMP(GO:0071772) |  | 171 | 0.78 | 1.27 | 0.141 | 0.434 | 0.828 | 8177 | tags=27%, list=14%, signal=31% |
| 2346 | FLAGELLATED\_SPERM\_MOTILITY(GO:0030317) |  | 84 | 0.77 | 1.27 | 0.141 | 0.434 | 0.828 | 10309 | tags=33%, list=17%, signal=40% |
| 2347 | DETECTION\_OF\_EXTERNAL\_BIOTIC\_STIMULUS(GO:0098581) |  | 19 | 0.76 | 1.27 | 0.000 | 0.434 | 0.828 | 9039 | tags=26%, list=15%, signal=31% |
| 2348 | PURINE\_NUCLEOSIDE\_MONOPHOSPHATE\_METABOLIC\_PROCESS(GO:0009126) |  | 330 | 0.76 | 1.27 | 0.000 | 0.434 | 0.828 | 6696 | tags=6%, list=11%, signal=7% |
| 2349 | PHOSPHOPROTEIN\_BINDING(GO:0051219) |  | 76 | 0.81 | 1.27 | 0.141 | 0.434 | 0.828 | 7947 | tags=14%, list=13%, signal=17% |
| 2350 | NEURON\_PROJECTION\_REGENERATION(GO:0031102) |  | 54 | 0.78 | 1.27 | 0.000 | 0.434 | 0.828 | 8743 | tags=31%, list=15%, signal=37% |
| 2351 | LAMELLIPODIUM\_MORPHOGENESIS(GO:0072673) |  | 17 | 0.85 | 1.27 | 0.000 | 0.434 | 0.828 | 6799 | tags=18%, list=11%, signal=20% |
| 2352 | SYNAPTIC\_VESICLE\_RECYCLING(GO:0036465) |  | 46 | 0.80 | 1.27 | 0.141 | 0.434 | 0.828 | 8763 | tags=24%, list=15%, signal=28% |
| 2353 | FEAR\_RESPONSE(GO:0042596) |  | 36 | 0.80 | 1.26 | 0.141 | 0.434 | 0.828 | 7298 | tags=19%, list=12%, signal=22% |
| 2354 | AMINOGLYCAN\_BIOSYNTHETIC\_PROCESS(GO:0006023) |  | 107 | 0.79 | 1.26 | 0.141 | 0.434 | 0.828 | 9277 | tags=30%, list=16%, signal=35% |
| 2355 | NUCLEOSIDE\_PHOSPHATE\_METABOLIC\_PROCESS(GO:0006753) |  | 691 | 0.76 | 1.26 | 0.141 | 0.433 | 0.828 | 7769 | tags=11%, list=13%, signal=12% |
| 2356 | POSITIVE\_REGULATION\_OF\_NITROGEN\_COMPOUND\_METABOLIC\_PROCESS(GO:0051173) |  | 2905 | 0.80 | 1.26 | 0.141 | 0.433 | 0.828 | 7756 | tags=15%, list=13%, signal=16% |
| 2357 | OLIGOSACCHARIDE\_METABOLIC\_PROCESS(GO:0009311) |  | 56 | 0.80 | 1.26 | 0.141 | 0.433 | 0.828 | 6848 | tags=18%, list=12%, signal=20% |
| 2358 | VITAMIN\_METABOLIC\_PROCESS(GO:0006766) |  | 133 | 0.77 | 1.26 | 0.141 | 0.433 | 0.828 | 8743 | tags=18%, list=15%, signal=21% |
| 2359 | VENTRICULAR\_CARDIAC\_MUSCLE\_CELL\_DIFFERENTIATION(GO:0055012) |  | 18 | 0.78 | 1.26 | 0.000 | 0.433 | 0.828 | 8774 | tags=39%, list=15%, signal=46% |
| 2360 | REGULATION\_OF\_GLUTAMATE\_SECRETION(GO:0014048) |  | 16 | 0.87 | 1.26 | 0.000 | 0.433 | 0.828 | 5894 | tags=19%, list=10%, signal=21% |
| 2361 | REGULATION\_OF\_ATP\_METABOLIC\_PROCESS(GO:1903578) |  | 89 | 0.80 | 1.26 | 0.141 | 0.433 | 0.828 | 6006 | tags=4%, list=10%, signal=5% |
| 2362 | GLYCOLIPID\_BIOSYNTHETIC\_PROCESS(GO:0009247) |  | 67 | 0.74 | 1.26 | 0.000 | 0.433 | 0.828 | 9651 | tags=27%, list=16%, signal=32% |
| 2363 | POSITIVE\_REGULATION\_OF\_CGMP\_BIOSYNTHETIC\_PROCESS(GO:0030828) |  | 19 | 0.78 | 1.26 | 0.000 | 0.433 | 0.828 | 8279 | tags=32%, list=14%, signal=37% |
| 2364 | REGULATION\_OF\_BRANCHING\_INVOLVED\_IN\_URETERIC\_BUD\_MORPHOGENESIS(GO:0090189) |  | 23 | 0.88 | 1.26 | 0.141 | 0.433 | 0.828 | 6279 | tags=13%, list=11%, signal=15% |
| 2365 | NEURON\_PROJECTION(GO:0043005) |  | 1139 | 0.79 | 1.26 | 0.141 | 0.433 | 0.828 | 7975 | tags=19%, list=13%, signal=21% |
| 2366 | REGULATION\_OF\_INTRACELLULAR\_PROTEIN\_TRANSPORT(GO:0033157) |  | 260 | 0.80 | 1.26 | 0.141 | 0.433 | 0.828 | 6793 | tags=7%, list=11%, signal=8% |
| 2367 | CARTILAGE\_DEVELOPMENT\_INVOLVED\_IN\_ENDOCHONDRAL\_BONE\_MORPHOGENESIS(GO:0060351) |  | 26 | 0.81 | 1.26 | 0.000 | 0.433 | 0.828 | 7322 | tags=23%, list=12%, signal=26% |
| 2368 | TRANSPORT\_VESICLE(GO:0030133) |  | 355 | 0.77 | 1.26 | 0.141 | 0.432 | 0.828 | 8063 | tags=16%, list=14%, signal=18% |
| 2369 | DECIDUALIZATION(GO:0046697) |  | 22 | 0.84 | 1.26 | 0.000 | 0.432 | 0.828 | 6937 | tags=18%, list=12%, signal=21% |
| 2370 | PHOSPHATIDYLINOSITOL\_MEDIATED\_SIGNALING(GO:0048015) |  | 157 | 0.78 | 1.26 | 0.141 | 0.432 | 0.828 | 7046 | tags=15%, list=12%, signal=17% |
| 2371 | CYTOPLASMIC\_VESICLE(GO:0031410) |  | 2220 | 0.79 | 1.26 | 0.141 | 0.432 | 0.828 | 7441 | tags=12%, list=13%, signal=13% |
| 2372 | NEGATIVE\_REGULATION\_OF\_MACROMOLECULE\_METABOLIC\_PROCESS(GO:0010605) |  | 2811 | 0.80 | 1.26 | 0.141 | 0.432 | 0.828 | 7268 | tags=10%, list=12%, signal=11% |
| 2373 | NEURON\_PART(GO:0097458) |  | 1442 | 0.80 | 1.26 | 0.141 | 0.432 | 0.828 | 7995 | tags=19%, list=14%, signal=21% |
| 2374 | NEGATIVE\_REGULATION\_OF\_OSTEOBLAST\_DIFFERENTIATION(GO:0045668) |  | 43 | 0.83 | 1.26 | 0.141 | 0.432 | 0.828 | 6211 | tags=26%, list=11%, signal=29% |
| 2375 | REGULATION\_OF\_PLATELET\_AGGREGATION(GO:0090330) |  | 17 | 0.86 | 1.26 | 0.000 | 0.432 | 0.828 | 7546 | tags=35%, list=13%, signal=40% |
| 2376 | SYMPATHETIC\_NERVOUS\_SYSTEM\_DEVELOPMENT(GO:0048485) |  | 21 | 0.84 | 1.26 | 0.141 | 0.432 | 0.828 | 6965 | tags=24%, list=12%, signal=27% |
| 2377 | NEGATIVE\_REGULATION\_OF\_BMP\_SIGNALING\_PATHWAY(GO:0030514) |  | 51 | 0.79 | 1.26 | 0.000 | 0.432 | 0.828 | 7913 | tags=29%, list=13%, signal=34% |
| 2378 | AMINO\_ACID\_TRANSMEMBRANE\_TRANSPORT(GO:0003333) |  | 74 | 0.77 | 1.26 | 0.000 | 0.432 | 0.828 | 9451 | tags=30%, list=16%, signal=35% |
| 2379 | REGULATION\_OF\_CELL\_MATRIX\_ADHESION(GO:0001952) |  | 106 | 0.80 | 1.26 | 0.141 | 0.432 | 0.828 | 7372 | tags=12%, list=12%, signal=14% |
| 2380 | VACUOLE(GO:0005773) |  | 759 | 0.77 | 1.26 | 0.141 | 0.431 | 0.828 | 7441 | tags=9%, list=13%, signal=10% |
| 2381 | CARBOHYDRATE\_METABOLIC\_PROCESS(GO:0005975) |  | 610 | 0.78 | 1.26 | 0.141 | 0.431 | 0.828 | 7293 | tags=12%, list=12%, signal=13% |
| 2382 | POTASSIUM\_ION\_TRANSPORT(GO:0006813) |  | 246 | 0.78 | 1.26 | 0.141 | 0.431 | 0.828 | 9703 | tags=28%, list=16%, signal=33% |
| 2383 | INTRACELLULAR\_VESICLE(GO:0097708) |  | 2223 | 0.79 | 1.26 | 0.141 | 0.431 | 0.828 | 7441 | tags=12%, list=13%, signal=13% |
| 2384 | AMINOGLYCAN\_METABOLIC\_PROCESS(GO:0006022) |  | 164 | 0.78 | 1.26 | 0.000 | 0.431 | 0.828 | 9277 | tags=32%, list=16%, signal=38% |
| 2385 | RECEPTOR\_COMPLEX(GO:0043235) |  | 327 | 0.77 | 1.26 | 0.141 | 0.431 | 0.828 | 9000 | tags=27%, list=15%, signal=31% |
| 2386 | EYE\_MORPHOGENESIS(GO:0048592) |  | 147 | 0.80 | 1.26 | 0.141 | 0.431 | 0.828 | 8490 | tags=28%, list=14%, signal=32% |
| 2387 | SWI\_SNF\_SUPERFAMILY\_TYPE\_COMPLEX(GO:0070603) |  | 74 | 0.75 | 1.26 | 0.000 | 0.431 | 0.828 | 5860 | tags=5%, list=10%, signal=6% |
| 2388 | POSITIVE\_REGULATION\_OF\_CARDIAC\_MUSCLE\_TISSUE\_GROWTH(GO:0055023) |  | 44 | 0.79 | 1.26 | 0.141 | 0.431 | 0.828 | 8122 | tags=23%, list=14%, signal=26% |
| 2389 | HSP70\_PROTEIN\_BINDING(GO:0030544) |  | 38 | 0.77 | 1.26 | 0.000 | 0.431 | 0.828 | 5894 | tags=11%, list=10%, signal=12% |
| 2390 | PURINE\_RIBONUCLEOSIDE\_MONOPHOSPHATE\_METABOLIC\_PROCESS(GO:0009167) |  | 329 | 0.76 | 1.26 | 0.000 | 0.430 | 0.828 | 6696 | tags=6%, list=11%, signal=6% |
| 2391 | CILIUM(GO:0005929) |  | 559 | 0.76 | 1.26 | 0.000 | 0.430 | 0.828 | 9484 | tags=26%, list=16%, signal=30% |
| 2392 | TRANSMEMBRANE\_TRANSPORT(GO:0055085) |  | 1564 | 0.78 | 1.26 | 0.141 | 0.430 | 0.828 | 8596 | tags=21%, list=15%, signal=23% |
| 2393 | ORGANIC\_CATION\_TRANSMEMBRANE\_TRANSPORTER\_ACTIVITY(GO:0015101) |  | 23 | 0.80 | 1.26 | 0.000 | 0.430 | 0.828 | 7268 | tags=17%, list=12%, signal=20% |
| 2394 | TUBULIN\_BINDING(GO:0015631) |  | 348 | 0.75 | 1.26 | 0.000 | 0.430 | 0.828 | 10334 | tags=29%, list=17%, signal=35% |
| 2395 | REGULATION\_OF\_BONE\_MINERALIZATION(GO:0030500) |  | 67 | 0.79 | 1.26 | 0.141 | 0.430 | 0.828 | 7441 | tags=28%, list=13%, signal=32% |
| 2396 | GLYCOSAMINOGLYCAN\_METABOLIC\_PROCESS(GO:0030203) |  | 157 | 0.78 | 1.26 | 0.000 | 0.430 | 0.828 | 9277 | tags=31%, list=16%, signal=37% |
| 2397 | NEGATIVE\_REGULATION\_OF\_NUCLEOCYTOPLASMIC\_TRANSPORT(GO:0046823) |  | 20 | 0.83 | 1.26 | 0.141 | 0.430 | 0.828 | 6541 | tags=15%, list=11%, signal=17% |
| 2398 | NEGATIVE\_REGULATION\_OF\_PRODUCTION\_OF\_MOLECULAR\_MEDIATOR\_OF\_IMMUNE\_RESPONSE(GO:0002701) |  | 32 | 0.79 | 1.26 | 0.141 | 0.430 | 0.828 | 10458 | tags=44%, list=18%, signal=53% |
| 2399 | REGULATION\_OF\_SUBSTRATE\_ADHESION\_DEPENDENT\_CELL\_SPREADING(GO:1900024) |  | 49 | 0.81 | 1.26 | 0.141 | 0.430 | 0.828 | 6531 | tags=12%, list=11%, signal=14% |
| 2400 | INTRINSIC\_COMPONENT\_OF\_MITOCHONDRIAL\_OUTER\_MEMBRANE(GO:0031306) |  | 24 | 0.75 | 1.26 | 0.000 | 0.430 | 0.828 | 5977 | tags=4%, list=10%, signal=5% |
| 2401 | BONE\_GROWTH(GO:0098868) |  | 24 | 0.82 | 1.26 | 0.000 | 0.430 | 0.828 | 6375 | tags=33%, list=11%, signal=37% |
| 2402 | MIDBODY(GO:0030496) |  | 167 | 0.74 | 1.26 | 0.000 | 0.430 | 0.828 | 9404 | tags=15%, list=16%, signal=18% |
| 2403 | NEGATIVE\_REGULATION\_OF\_ION\_TRANSMEMBRANE\_TRANSPORT(GO:0034766) |  | 96 | 0.77 | 1.26 | 0.141 | 0.430 | 0.828 | 7979 | tags=19%, list=13%, signal=22% |
| 2404 | CHROMOSOME\_CONDENSATION(GO:0030261) |  | 33 | 0.77 | 1.26 | 0.000 | 0.430 | 0.828 | 10719 | tags=33%, list=18%, signal=41% |
| 2405 | STEROL\_BINDING(GO:0032934) |  | 48 | 0.81 | 1.26 | 0.141 | 0.430 | 0.828 | 7721 | tags=27%, list=13%, signal=31% |
| 2406 | INTRINSIC\_APOPTOTIC\_SIGNALING\_PATHWAY(GO:0097193) |  | 290 | 0.78 | 1.26 | 0.141 | 0.430 | 0.828 | 7783 | tags=9%, list=13%, signal=10% |
| 2407 | REGULATION\_OF\_PHOSPHATE\_METABOLIC\_PROCESS(GO:0019220) |  | 1748 | 0.79 | 1.26 | 0.141 | 0.430 | 0.828 | 7702 | tags=14%, list=13%, signal=15% |
| 2408 | CELLULAR\_RESPONSE\_TO\_DRUG(GO:0035690) |  | 340 | 0.77 | 1.26 | 0.141 | 0.430 | 0.828 | 7745 | tags=14%, list=13%, signal=15% |
| 2409 | REGULATION\_OF\_PROTEOLYSIS(GO:0030162) |  | 794 | 0.78 | 1.26 | 0.141 | 0.430 | 0.828 | 7853 | tags=12%, list=13%, signal=13% |
| 2410 | LIPASE\_ACTIVITY(GO:0016298) |  | 128 | 0.78 | 1.26 | 0.141 | 0.429 | 0.828 | 6949 | tags=15%, list=12%, signal=17% |
| 2411 | POSITIVE\_REGULATION\_OF\_BINDING(GO:0051099) |  | 172 | 0.75 | 1.26 | 0.000 | 0.429 | 0.828 | 8784 | tags=20%, list=15%, signal=24% |
| 2412 | G\_PROTEIN\_COUPLED\_PURINERGIC\_NUCLEOTIDE\_RECEPTOR\_SIGNALING\_PATHWAY(GO:0035589) |  | 15 | 0.86 | 1.26 | 0.000 | 0.429 | 0.828 | 5955 | tags=20%, list=10%, signal=22% |
| 2413 | G\_PROTEIN\_COUPLED\_PURINERGIC\_NUCLEOTIDE\_RECEPTOR\_ACTIVITY(GO:0045028) |  | 15 | 0.86 | 1.26 | 0.000 | 0.429 | 0.828 | 5955 | tags=20%, list=10%, signal=22% |
| 2414 | G\_PROTEIN\_COUPLED\_NUCLEOTIDE\_RECEPTOR\_ACTIVITY(GO:0001608) |  | 15 | 0.86 | 1.26 | 0.000 | 0.429 | 0.828 | 5955 | tags=20%, list=10%, signal=22% |
| 2415 | REGULATION\_OF\_SPROUTING\_ANGIOGENESIS(GO:1903670) |  | 130 | 0.78 | 1.26 | 0.141 | 0.429 | 0.828 | 7698 | tags=15%, list=13%, signal=18% |
| 2416 | PROTEIN\_PHOSPHORYLATION(GO:0006468) |  | 1953 | 0.79 | 1.26 | 0.141 | 0.429 | 0.828 | 7973 | tags=15%, list=13%, signal=16% |
| 2417 | CARBOHYDRATE\_DERIVATIVE\_CATABOLIC\_PROCESS(GO:1901136) |  | 183 | 0.74 | 1.26 | 0.000 | 0.429 | 0.828 | 9382 | tags=23%, list=16%, signal=28% |
| 2418 | REGULATION\_OF\_OSTEOBLAST\_DIFFERENTIATION(GO:0045667) |  | 114 | 0.80 | 1.26 | 0.141 | 0.429 | 0.828 | 8046 | tags=22%, list=14%, signal=25% |
| 2419 | AXON(GO:0030424) |  | 528 | 0.80 | 1.26 | 0.141 | 0.429 | 0.828 | 7969 | tags=20%, list=13%, signal=22% |
| 2420 | SENSORY\_PERCEPTION\_OF\_PAIN(GO:0019233) |  | 95 | 0.82 | 1.26 | 0.141 | 0.429 | 0.828 | 7873 | tags=34%, list=13%, signal=39% |
| 2421 | INOSITOL\_LIPID\_MEDIATED\_SIGNALING(GO:0048017) |  | 160 | 0.77 | 1.26 | 0.141 | 0.429 | 0.828 | 7046 | tags=14%, list=12%, signal=16% |
| 2422 | NEUROTRANSMITTER\_TRANSPORT(GO:0006836) |  | 240 | 0.78 | 1.26 | 0.141 | 0.428 | 0.828 | 8249 | tags=25%, list=14%, signal=29% |
| 2423 | INNER\_EAR\_RECEPTOR\_CELL\_DIFFERENTIATION(GO:0060113) |  | 60 | 0.76 | 1.26 | 0.000 | 0.428 | 0.828 | 9354 | tags=27%, list=16%, signal=32% |
| 2424 | REGULATION\_OF\_CELLULAR\_COMPONENT\_ORGANIZATION(GO:0051128) |  | 2360 | 0.80 | 1.26 | 0.141 | 0.428 | 0.828 | 7975 | tags=13%, list=13%, signal=14% |
| 2425 | NUCLEOBASE\_CONTAINING\_SMALL\_MOLECULE\_METABOLIC\_PROCESS(GO:0055086) |  | 776 | 0.76 | 1.26 | 0.141 | 0.428 | 0.828 | 7769 | tags=11%, list=13%, signal=12% |
| 2426 | DIGESTIVE\_TRACT\_MORPHOGENESIS(GO:0048546) |  | 49 | 0.81 | 1.26 | 0.141 | 0.428 | 0.828 | 8107 | tags=35%, list=14%, signal=40% |
| 2427 | REGULATION\_OF\_PHOSPHORUS\_METABOLIC\_PROCESS(GO:0051174) |  | 1762 | 0.79 | 1.26 | 0.141 | 0.428 | 0.828 | 7702 | tags=14%, list=13%, signal=15% |
| 2428 | REGULATION\_OF\_LIPID\_BIOSYNTHETIC\_PROCESS(GO:0046890) |  | 168 | 0.76 | 1.26 | 0.000 | 0.428 | 0.828 | 7699 | tags=13%, list=13%, signal=15% |
| 2429 | RESPONSE\_TO\_IRON\_ION(GO:0010039) |  | 32 | 0.82 | 1.26 | 0.141 | 0.428 | 0.828 | 7986 | tags=22%, list=14%, signal=25% |
| 2430 | POSITIVE\_REGULATION\_OF\_RNA\_BIOSYNTHETIC\_PROCESS(GO:1902680) |  | 1440 | 0.78 | 1.26 | 0.141 | 0.428 | 0.828 | 8066 | tags=17%, list=14%, signal=20% |
| 2431 | POTASSIUM\_ION\_TRANSMEMBRANE\_TRANSPORT(GO:0071805) |  | 209 | 0.77 | 1.26 | 0.141 | 0.428 | 0.828 | 9703 | tags=29%, list=16%, signal=35% |
| 2432 | CELLULAR\_POTASSIUM\_ION\_TRANSPORT(GO:0071804) |  | 209 | 0.77 | 1.26 | 0.141 | 0.428 | 0.828 | 9703 | tags=29%, list=16%, signal=35% |
| 2433 | CELL\_MIGRATION\_INVOLVED\_IN\_SPROUTING\_ANGIOGENESIS(GO:0002042) |  | 86 | 0.79 | 1.26 | 0.141 | 0.427 | 0.828 | 9016 | tags=22%, list=15%, signal=26% |
| 2434 | INTRASPECIES\_INTERACTION\_BETWEEN\_ORGANISMS(GO:0051703) |  | 52 | 0.81 | 1.26 | 0.141 | 0.427 | 0.828 | 7962 | tags=27%, list=13%, signal=31% |
| 2435 | SOCIAL\_BEHAVIOR(GO:0035176) |  | 52 | 0.81 | 1.26 | 0.141 | 0.427 | 0.828 | 7962 | tags=27%, list=13%, signal=31% |
| 2436 | NON\_MOTILE\_CILIUM(GO:0097730) |  | 130 | 0.77 | 1.26 | 0.141 | 0.427 | 0.828 | 9717 | tags=28%, list=16%, signal=34% |
| 2437 | UDP\_GALACTOSYLTRANSFERASE\_ACTIVITY(GO:0035250) |  | 30 | 0.82 | 1.26 | 0.141 | 0.427 | 0.828 | 6683 | tags=20%, list=11%, signal=23% |
| 2438 | VIRAL\_GENOME\_REPLICATION(GO:0019079) |  | 114 | 0.78 | 1.26 | 0.141 | 0.427 | 0.828 | 7293 | tags=10%, list=12%, signal=11% |
| 2439 | GLYCEROPHOSPHOLIPID\_CATABOLIC\_PROCESS(GO:0046475) |  | 19 | 0.78 | 1.26 | 0.000 | 0.427 | 0.828 | 6393 | tags=16%, list=11%, signal=18% |
| 2440 | POSITIVE\_REGULATION\_OF\_TRANSCRIPTION\_\_DNA\_TEMPLATED(GO:0045893) |  | 1439 | 0.78 | 1.26 | 0.141 | 0.427 | 0.828 | 8066 | tags=17%, list=14%, signal=20% |
| 2441 | POSITIVE\_REGULATION\_OF\_NUCLEIC\_ACID\_TEMPLATED\_TRANSCRIPTION(GO:1903508) |  | 1439 | 0.78 | 1.26 | 0.141 | 0.427 | 0.828 | 8066 | tags=17%, list=14%, signal=20% |
| 2442 | MOLECULAR\_FUNCTION\_REGULATOR(GO:0098772) |  | 1822 | 0.78 | 1.26 | 0.141 | 0.427 | 0.828 | 7876 | tags=16%, list=13%, signal=17% |
| 2443 | REGULATION\_OF\_NEUROTRANSMITTER\_UPTAKE(GO:0051580) |  | 16 | 0.84 | 1.26 | 0.141 | 0.427 | 0.828 | 7298 | tags=25%, list=12%, signal=29% |
| 2444 | CARBOXYLIC\_ACID\_TRANSMEMBRANE\_TRANSPORT(GO:1905039) |  | 115 | 0.78 | 1.26 | 0.141 | 0.427 | 0.828 | 9451 | tags=25%, list=16%, signal=30% |
| 2445 | ORGANIC\_ACID\_TRANSMEMBRANE\_TRANSPORT(GO:1903825) |  | 115 | 0.78 | 1.26 | 0.141 | 0.427 | 0.828 | 9451 | tags=25%, list=16%, signal=30% |
| 2446 | UBIQUITIN\_LIKE\_PROTEIN\_LIGASE\_ACTIVITY(GO:0061659) |  | 226 | 0.75 | 1.26 | 0.000 | 0.426 | 0.828 | 8341 | tags=12%, list=14%, signal=14% |
| 2447 | NEGATIVE\_REGULATION\_OF\_ORGAN\_GROWTH(GO:0046621) |  | 35 | 0.82 | 1.26 | 0.141 | 0.426 | 0.828 | 8380 | tags=20%, list=14%, signal=23% |
| 2448 | INTRACELLULAR\_TRANSPORT(GO:0046907) |  | 1660 | 0.80 | 1.26 | 0.141 | 0.426 | 0.828 | 6933 | tags=5%, list=12%, signal=6% |
| 2449 | GLUCOSAMINE\_CONTAINING\_COMPOUND\_METABOLIC\_PROCESS(GO:1901071) |  | 23 | 0.75 | 1.26 | 0.000 | 0.426 | 0.828 | 9382 | tags=43%, list=16%, signal=52% |
| 2450 | UBIQUITIN\_CONJUGATING\_ENZYME\_BINDING(GO:0031624) |  | 34 | 0.78 | 1.26 | 0.000 | 0.426 | 0.828 | 8490 | tags=24%, list=14%, signal=27% |
| 2451 | ENDOTHELIAL\_CELL\_PROLIFERATION(GO:0001935) |  | 176 | 0.79 | 1.26 | 0.141 | 0.426 | 0.828 | 8122 | tags=20%, list=14%, signal=24% |
| 2452 | POSITIVE\_REGULATION\_OF\_MACROMOLECULE\_METABOLIC\_PROCESS(GO:0010604) |  | 3051 | 0.80 | 1.26 | 0.141 | 0.426 | 0.828 | 7846 | tags=15%, list=13%, signal=16% |
| 2453 | AMINO\_ACID\_TRANSMEMBRANE\_TRANSPORTER\_ACTIVITY(GO:0015171) |  | 77 | 0.76 | 1.26 | 0.000 | 0.426 | 0.828 | 9451 | tags=32%, list=16%, signal=39% |
| 2454 | SOLUTE:CATION\_SYMPORTER\_ACTIVITY(GO:0015294) |  | 111 | 0.76 | 1.26 | 0.000 | 0.426 | 0.828 | 9451 | tags=33%, list=16%, signal=40% |
| 2455 | RIBONUCLEOSIDE\_MONOPHOSPHATE\_METABOLIC\_PROCESS(GO:0009161) |  | 341 | 0.75 | 1.26 | 0.000 | 0.426 | 0.828 | 6696 | tags=6%, list=11%, signal=6% |
| 2456 | CELLULAR\_EXTRAVASATION(GO:0045123) |  | 50 | 0.77 | 1.26 | 0.141 | 0.426 | 0.828 | 8784 | tags=22%, list=15%, signal=26% |
| 2457 | PROTEIN\_CONTAINING\_COMPLEX\_ASSEMBLY(GO:0065003) |  | 1822 | 0.79 | 1.26 | 0.141 | 0.426 | 0.828 | 7659 | tags=10%, list=13%, signal=11% |
| 2458 | EMBRYONIC\_HINDLIMB\_MORPHOGENESIS(GO:0035116) |  | 29 | 0.84 | 1.26 | 0.141 | 0.425 | 0.828 | 8261 | tags=45%, list=14%, signal=52% |
| 2459 | PROXIMAL\_PROMOTER\_SEQUENCE\_SPECIFIC\_DNA\_BINDING(GO:0000987) |  | 435 | 0.79 | 1.26 | 0.141 | 0.426 | 0.828 | 8033 | tags=21%, list=14%, signal=24% |
| 2460 | REGULATION\_OF\_IMMUNOGLOBULIN\_SECRETION(GO:0051023) |  | 16 | 0.81 | 1.26 | 0.000 | 0.425 | 0.828 | 6178 | tags=25%, list=10%, signal=28% |
| 2461 | POSITIVE\_REGULATION\_OF\_PROTEIN\_TRANSPORT(GO:0051222) |  | 426 | 0.79 | 1.26 | 0.141 | 0.425 | 0.828 | 7652 | tags=13%, list=13%, signal=14% |
| 2462 | RNA\_POLYMERASE\_II\_PROXIMAL\_PROMOTER\_SEQUENCE\_SPECIFIC\_DNA\_BINDING(GO:0000978) |  | 421 | 0.79 | 1.26 | 0.141 | 0.425 | 0.828 | 8524 | tags=23%, list=14%, signal=26% |
| 2463 | L\_AMINO\_ACID\_TRANSPORT(GO:0015807) |  | 63 | 0.76 | 1.26 | 0.000 | 0.425 | 0.828 | 9357 | tags=24%, list=16%, signal=28% |
| 2464 | NEGATIVE\_REGULATION\_OF\_DNA\_BINDING\_TRANSCRIPTION\_FACTOR\_ACTIVITY(GO:0043433) |  | 157 | 0.77 | 1.26 | 0.141 | 0.425 | 0.828 | 6178 | tags=11%, list=10%, signal=12% |
| 2465 | RECEPTOR\_INTERNALIZATION(GO:0031623) |  | 87 | 0.79 | 1.26 | 0.141 | 0.425 | 0.828 | 7154 | tags=16%, list=12%, signal=18% |
| 2466 | REGULATION\_OF\_CELL\_SUBSTRATE\_ADHESION(GO:0010810) |  | 192 | 0.80 | 1.26 | 0.141 | 0.425 | 0.828 | 7372 | tags=15%, list=12%, signal=17% |
| 2467 | REGULATION\_OF\_CARDIAC\_CONDUCTION(GO:1903779) |  | 75 | 0.79 | 1.26 | 0.141 | 0.425 | 0.828 | 8198 | tags=32%, list=14%, signal=37% |
| 2468 | REGULATION\_OF\_STRIATED\_MUSCLE\_TISSUE\_DEVELOPMENT(GO:0016202) |  | 143 | 0.78 | 1.26 | 0.141 | 0.425 | 0.828 | 8774 | tags=27%, list=15%, signal=32% |
| 2469 | NEURON\_PROJECTION\_DEVELOPMENT(GO:0031175) |  | 919 | 0.79 | 1.26 | 0.141 | 0.425 | 0.828 | 7975 | tags=19%, list=13%, signal=22% |
| 2470 | HYALURONIC\_ACID\_BINDING(GO:0005540) |  | 27 | 0.84 | 1.26 | 0.000 | 0.425 | 0.828 | 8242 | tags=33%, list=14%, signal=39% |
| 2471 | LIPID\_TRANSPORT(GO:0006869) |  | 347 | 0.78 | 1.26 | 0.141 | 0.425 | 0.828 | 7699 | tags=15%, list=13%, signal=17% |
| 2472 | NEGATIVE\_REGULATION\_OF\_MUSCLE\_TISSUE\_DEVELOPMENT(GO:1901862) |  | 56 | 0.81 | 1.26 | 0.141 | 0.425 | 0.828 | 8774 | tags=29%, list=15%, signal=34% |
| 2473 | RESPONSE\_TO\_ORGANOPHOSPHORUS(GO:0046683) |  | 134 | 0.78 | 1.26 | 0.141 | 0.425 | 0.828 | 7172 | tags=19%, list=12%, signal=22% |
| 2474 | TRABECULA\_MORPHOGENESIS(GO:0061383) |  | 50 | 0.76 | 1.26 | 0.000 | 0.425 | 0.828 | 9633 | tags=36%, list=16%, signal=43% |
| 2475 | SUPRAMOLECULAR\_FIBER\_ORGANIZATION(GO:0097435) |  | 597 | 0.79 | 1.26 | 0.141 | 0.425 | 0.828 | 7973 | tags=17%, list=13%, signal=19% |
| 2476 | CYTOSKELETON(GO:0005856) |  | 2099 | 0.80 | 1.26 | 0.141 | 0.425 | 0.828 | 7955 | tags=14%, list=13%, signal=15% |
| 2477 | REGULATION\_OF\_GROWTH(GO:0040008) |  | 657 | 0.79 | 1.26 | 0.141 | 0.425 | 0.828 | 7583 | tags=16%, list=13%, signal=18% |
| 2478 | POSITIVE\_REGULATION\_OF\_CALCIUM\_ION\_TRANSPORT\_INTO\_CYTOSOL(GO:0010524) |  | 52 | 0.81 | 1.26 | 0.141 | 0.425 | 0.828 | 7347 | tags=25%, list=12%, signal=29% |
| 2479 | CD4\_POSITIVE\_\_ALPHA\_BETA\_T\_CELL\_DIFFERENTIATION(GO:0043367) |  | 64 | 0.78 | 1.26 | 0.141 | 0.425 | 0.828 | 9411 | tags=30%, list=16%, signal=35% |
| 2480 | EMBRYONIC\_SKELETAL\_JOINT\_DEVELOPMENT(GO:0072498) |  | 15 | 0.82 | 1.26 | 0.000 | 0.425 | 0.828 | 7913 | tags=33%, list=13%, signal=38% |
| 2481 | CELL\_CELL\_SIGNALING\_BY\_WNT(GO:0198738) |  | 498 | 0.77 | 1.26 | 0.141 | 0.424 | 0.828 | 8801 | tags=19%, list=15%, signal=22% |
| 2482 | REGULATION\_OF\_MUSCLE\_ORGAN\_DEVELOPMENT(GO:0048634) |  | 147 | 0.78 | 1.26 | 0.141 | 0.424 | 0.828 | 8774 | tags=27%, list=15%, signal=32% |
| 2483 | WNT\_SIGNALING\_PATHWAY(GO:0016055) |  | 496 | 0.77 | 1.26 | 0.141 | 0.424 | 0.828 | 8801 | tags=19%, list=15%, signal=22% |
| 2484 | POSITIVE\_REGULATION\_OF\_I\_KAPPAB\_KINASE\_NF\_KAPPAB\_SIGNALING(GO:0043123) |  | 170 | 0.80 | 1.26 | 0.141 | 0.424 | 0.828 | 7293 | tags=11%, list=12%, signal=12% |
| 2485 | PRESYNAPTIC\_PROCESS\_INVOLVED\_IN\_CHEMICAL\_SYNAPTIC\_TRANSMISSION(GO:0099531) |  | 150 | 0.78 | 1.26 | 0.141 | 0.424 | 0.828 | 8249 | tags=24%, list=14%, signal=28% |
| 2486 | REGULATION\_OF\_MUSCLE\_TISSUE\_DEVELOPMENT(GO:1901861) |  | 145 | 0.78 | 1.26 | 0.141 | 0.424 | 0.828 | 8774 | tags=27%, list=15%, signal=32% |
| 2487 | NEGATIVE\_REGULATION\_OF\_METABOLIC\_PROCESS(GO:0009892) |  | 3033 | 0.79 | 1.26 | 0.141 | 0.424 | 0.828 | 7268 | tags=10%, list=12%, signal=11% |
| 2488 | REGULATION\_OF\_VASCULAR\_ENDOTHELIAL\_GROWTH\_FACTOR\_PRODUCTION(GO:0010574) |  | 32 | 0.78 | 1.26 | 0.000 | 0.424 | 0.828 | 8279 | tags=25%, list=14%, signal=29% |
| 2489 | REGULATION\_OF\_NEUROTRANSMITTER\_TRANSPORT(GO:0051588) |  | 88 | 0.80 | 1.26 | 0.141 | 0.424 | 0.828 | 8040 | tags=27%, list=14%, signal=32% |
| 2490 | BICARBONATE\_TRANSPORT(GO:0015701) |  | 44 | 0.76 | 1.26 | 0.000 | 0.424 | 0.828 | 9957 | tags=41%, list=17%, signal=49% |
| 2491 | CELL\_MATRIX\_ADHESION(GO:0007160) |  | 211 | 0.79 | 1.26 | 0.141 | 0.424 | 0.828 | 7397 | tags=15%, list=13%, signal=17% |
| 2492 | REGULATION\_OF\_TRANSCRIPTION\_BY\_RNA\_POLYMERASE\_II(GO:0006357) |  | 2210 | 0.79 | 1.26 | 0.141 | 0.424 | 0.828 | 7450 | tags=16%, list=13%, signal=17% |
| 2493 | NEGATIVE\_REGULATION\_OF\_RNA\_METABOLIC\_PROCESS(GO:0051253) |  | 1297 | 0.79 | 1.26 | 0.141 | 0.424 | 0.828 | 7404 | tags=13%, list=13%, signal=14% |
| 2494 | IMPORT\_INTO\_CELL(GO:0098657) |  | 819 | 0.77 | 1.26 | 0.141 | 0.424 | 0.828 | 8112 | tags=13%, list=14%, signal=15% |
| 2495 | DOUBLE\_STRANDED\_DNA\_BINDING(GO:0003690) |  | 836 | 0.79 | 1.26 | 0.141 | 0.424 | 0.828 | 8038 | tags=19%, list=14%, signal=21% |
| 2496 | CARBON\_OXYGEN\_LYASE\_ACTIVITY(GO:0016835) |  | 70 | 0.75 | 1.26 | 0.000 | 0.424 | 0.828 | 8925 | tags=17%, list=15%, signal=20% |
| 2497 | VESICLE(GO:0031982) |  | 4292 | 0.81 | 1.26 | 0.141 | 0.424 | 0.828 | 7174 | tags=11%, list=12%, signal=11% |
| 2498 | REGULATION\_OF\_BIOMINERAL\_TISSUE\_DEVELOPMENT(GO:0070167) |  | 76 | 0.79 | 1.26 | 0.141 | 0.424 | 0.828 | 7441 | tags=26%, list=13%, signal=30% |
| 2499 | REGULATION\_OF\_HORMONE\_LEVELS(GO:0010817) |  | 513 | 0.76 | 1.26 | 0.141 | 0.424 | 0.828 | 8987 | tags=20%, list=15%, signal=23% |
| 2500 | VASCULAR\_ENDOTHELIAL\_GROWTH\_FACTOR\_RECEPTOR\_SIGNALING\_PATHWAY(GO:0048010) |  | 97 | 0.75 | 1.26 | 0.000 | 0.424 | 0.828 | 8046 | tags=16%, list=14%, signal=19% |
| 2501 | NUCLEOSIDE\_MONOPHOSPHATE\_METABOLIC\_PROCESS(GO:0009123) |  | 355 | 0.75 | 1.26 | 0.000 | 0.423 | 0.828 | 6696 | tags=6%, list=11%, signal=6% |
| 2502 | RESPIRATORY\_SYSTEM\_DEVELOPMENT(GO:0060541) |  | 196 | 0.79 | 1.26 | 0.141 | 0.423 | 0.828 | 8823 | tags=25%, list=15%, signal=29% |
| 2503 | PROTEOLYSIS\_INVOLVED\_IN\_CELLULAR\_PROTEIN\_CATABOLIC\_PROCESS(GO:0051603) |  | 671 | 0.74 | 1.26 | 0.000 | 0.423 | 0.828 | 7795 | tags=7%, list=13%, signal=8% |
| 2504 | THYMUS\_DEVELOPMENT(GO:0048538) |  | 46 | 0.80 | 1.26 | 0.141 | 0.423 | 0.828 | 6068 | tags=15%, list=10%, signal=17% |
| 2505 | NEGATIVE\_REGULATION\_OF\_MYELOID\_LEUKOCYTE\_DIFFERENTIATION(GO:0002762) |  | 45 | 0.77 | 1.26 | 0.141 | 0.423 | 0.828 | 7504 | tags=18%, list=13%, signal=20% |
| 2506 | COLLAGEN\_BINDING(GO:0005518) |  | 63 | 0.82 | 1.26 | 0.141 | 0.423 | 0.828 | 8732 | tags=37%, list=15%, signal=43% |
| 2507 | NEURONAL\_CELL\_BODY(GO:0043025) |  | 453 | 0.79 | 1.26 | 0.141 | 0.423 | 0.828 | 7963 | tags=20%, list=13%, signal=23% |
| 2508 | GLOMERULUS\_DEVELOPMENT(GO:0032835) |  | 59 | 0.79 | 1.26 | 0.141 | 0.423 | 0.828 | 8107 | tags=32%, list=14%, signal=37% |
| 2509 | RESPONSE\_TO\_CAMP(GO:0051591) |  | 99 | 0.78 | 1.26 | 0.141 | 0.423 | 0.828 | 7172 | tags=17%, list=12%, signal=20% |
| 2510 | CARBOXYLIC\_ACID\_BIOSYNTHETIC\_PROCESS(GO:0046394) |  | 415 | 0.77 | 1.26 | 0.141 | 0.423 | 0.828 | 7358 | tags=12%, list=12%, signal=13% |
| 2511 | LUNG\_DEVELOPMENT(GO:0030324) |  | 169 | 0.79 | 1.26 | 0.141 | 0.423 | 0.828 | 8823 | tags=25%, list=15%, signal=29% |
| 2512 | ORGANIC\_ACID\_BIOSYNTHETIC\_PROCESS(GO:0016053) |  | 416 | 0.77 | 1.26 | 0.141 | 0.423 | 0.828 | 7358 | tags=12%, list=12%, signal=13% |
| 2513 | RESPONSE\_TO\_EPINEPHRINE(GO:0071871) |  | 16 | 0.85 | 1.26 | 0.141 | 0.423 | 0.828 | 7172 | tags=25%, list=12%, signal=28% |
| 2514 | EMBRYONIC\_CRANIAL\_SKELETON\_MORPHOGENESIS(GO:0048701) |  | 46 | 0.83 | 1.26 | 0.141 | 0.423 | 0.828 | 7134 | tags=22%, list=12%, signal=25% |
| 2515 | PHOSPHATIDYLINOSITOL\_4\_5\_BISPHOSPHATE\_BINDING(GO:0005546) |  | 63 | 0.77 | 1.26 | 0.141 | 0.422 | 0.828 | 8363 | tags=17%, list=14%, signal=20% |
| 2516 | ESTABLISHMENT\_OF\_LOCALIZATION\_IN\_CELL(GO:0051649) |  | 2026 | 0.79 | 1.26 | 0.141 | 0.422 | 0.828 | 7298 | tags=8%, list=12%, signal=8% |
| 2517 | INTERLEUKIN\_13\_PRODUCTION(GO:0032616) |  | 22 | 0.81 | 1.26 | 0.141 | 0.422 | 0.828 | 6685 | tags=32%, list=11%, signal=36% |
| 2518 | SYNCYTIUM\_FORMATION\_BY\_PLASMA\_MEMBRANE\_FUSION(GO:0000768) |  | 52 | 0.79 | 1.26 | 0.141 | 0.422 | 0.828 | 9145 | tags=29%, list=15%, signal=34% |
| 2519 | GLYCEROPHOSPHOLIPID\_BIOSYNTHETIC\_PROCESS(GO:0046474) |  | 246 | 0.74 | 1.26 | 0.000 | 0.422 | 0.828 | 9669 | tags=18%, list=16%, signal=22% |
| 2520 | RELAXATION\_OF\_CARDIAC\_MUSCLE(GO:0055119) |  | 17 | 0.77 | 1.26 | 0.000 | 0.422 | 0.828 | 6695 | tags=24%, list=11%, signal=27% |
| 2521 | EXCITATORY\_SYNAPSE(GO:0060076) |  | 29 | 0.80 | 1.26 | 0.141 | 0.422 | 0.828 | 7962 | tags=31%, list=13%, signal=36% |
| 2522 | CELLULAR\_RESPONSE\_TO\_VITAMIN(GO:0071295) |  | 29 | 0.76 | 1.26 | 0.000 | 0.422 | 0.828 | 6495 | tags=21%, list=11%, signal=23% |
| 2523 | DRUG\_TRANSPORT(GO:0015893) |  | 196 | 0.78 | 1.26 | 0.141 | 0.422 | 0.828 | 8485 | tags=24%, list=14%, signal=28% |
| 2524 | SPROUTING\_ANGIOGENESIS(GO:0002040) |  | 166 | 0.79 | 1.26 | 0.141 | 0.422 | 0.828 | 7698 | tags=19%, list=13%, signal=22% |
| 2525 | NOTOCHORD\_DEVELOPMENT(GO:0030903) |  | 18 | 0.83 | 1.26 | 0.000 | 0.422 | 0.828 | 7913 | tags=39%, list=13%, signal=45% |
| 2526 | POSITIVE\_REGULATION\_OF\_RNA\_METABOLIC\_PROCESS(GO:0051254) |  | 1513 | 0.78 | 1.26 | 0.141 | 0.422 | 0.828 | 8066 | tags=17%, list=14%, signal=19% |
| 2527 | SULFUR\_COMPOUND\_TRANSMEMBRANE\_TRANSPORTER\_ACTIVITY(GO:1901682) |  | 32 | 0.73 | 1.26 | 0.000 | 0.422 | 0.828 | 10186 | tags=38%, list=17%, signal=45% |
| 2528 | CELL\_SUBSTRATE\_ADHESION(GO:0031589) |  | 320 | 0.79 | 1.26 | 0.141 | 0.422 | 0.828 | 8119 | tags=18%, list=14%, signal=20% |
| 2529 | REGULATION\_OF\_SMOOTH\_MUSCLE\_CELL\_DIFFERENTIATION(GO:0051150) |  | 42 | 0.81 | 1.26 | 0.141 | 0.422 | 0.828 | 6928 | tags=17%, list=12%, signal=19% |
| 2530 | REGULATION\_OF\_CATALYTIC\_ACTIVITY(GO:0050790) |  | 2253 | 0.78 | 1.26 | 0.141 | 0.422 | 0.828 | 7921 | tags=13%, list=13%, signal=14% |
| 2531 | LYSOSOME(GO:0005764) |  | 663 | 0.76 | 1.26 | 0.141 | 0.422 | 0.828 | 7441 | tags=9%, list=13%, signal=10% |
| 2532 | LYTIC\_VACUOLE(GO:0000323) |  | 663 | 0.76 | 1.26 | 0.141 | 0.422 | 0.828 | 7441 | tags=9%, list=13%, signal=10% |
| 2533 | OSTEOBLAST\_DEVELOPMENT(GO:0002076) |  | 18 | 0.85 | 1.26 | 0.141 | 0.422 | 0.828 | 5988 | tags=17%, list=10%, signal=19% |
| 2534 | G\_PROTEIN\_COUPLED\_RECEPTOR\_BINDING(GO:0001664) |  | 265 | 0.79 | 1.26 | 0.141 | 0.421 | 0.828 | 8763 | tags=22%, list=15%, signal=26% |
| 2535 | POSITIVE\_REGULATION\_OF\_CELL\_ADHESION(GO:0045785) |  | 381 | 0.78 | 1.26 | 0.141 | 0.421 | 0.828 | 8112 | tags=16%, list=14%, signal=19% |
| 2536 | REGULATION\_OF\_AXON\_GUIDANCE(GO:1902667) |  | 22 | 0.83 | 1.26 | 0.141 | 0.421 | 0.828 | 7362 | tags=36%, list=12%, signal=42% |
| 2537 | RESPONSE\_TO\_INTERFERON\_GAMMA(GO:0034341) |  | 187 | 0.77 | 1.26 | 0.141 | 0.421 | 0.828 | 8063 | tags=13%, list=14%, signal=15% |
| 2538 | TRANSCRIPTION\_FACTOR\_ACTIVITY\_\_RNA\_POLYMERASE\_II\_PROXIMAL\_PROMOTER\_SEQUENCE\_SPECIFIC\_DNA\_BINDING(GO:0000982) |  | 402 | 0.79 | 1.26 | 0.141 | 0.421 | 0.828 | 8524 | tags=25%, list=14%, signal=29% |
| 2539 | INTEGRATOR\_COMPLEX(GO:0032039) |  | 16 | 0.77 | 1.26 | 0.000 | 0.421 | 0.828 | 6006 | tags=6%, list=10%, signal=7% |
| 2540 | ANKYRIN\_BINDING(GO:0030506) |  | 19 | 0.83 | 1.26 | 0.141 | 0.421 | 0.828 | 8590 | tags=37%, list=15%, signal=43% |
| 2541 | POSITIVE\_REGULATION\_OF\_TRANSCRIPTION\_REGULATORY\_REGION\_DNA\_BINDING(GO:2000679) |  | 23 | 0.85 | 1.26 | 0.141 | 0.421 | 0.828 | 7260 | tags=35%, list=12%, signal=40% |
| 2542 | CARDIAC\_VENTRICLE\_MORPHOGENESIS(GO:0003208) |  | 72 | 0.81 | 1.26 | 0.141 | 0.421 | 0.828 | 7913 | tags=28%, list=13%, signal=32% |
| 2543 | CRANIAL\_SKELETAL\_SYSTEM\_DEVELOPMENT(GO:1904888) |  | 66 | 0.82 | 1.26 | 0.141 | 0.421 | 0.828 | 7787 | tags=21%, list=13%, signal=24% |
| 2544 | AXONEME(GO:0005930) |  | 112 | 0.76 | 1.26 | 0.000 | 0.421 | 0.828 | 8933 | tags=32%, list=15%, signal=38% |
| 2545 | CELLULAR\_RESPONSE\_TO\_ENVIRONMENTAL\_STIMULUS(GO:0104004) |  | 306 | 0.79 | 1.26 | 0.141 | 0.421 | 0.828 | 7937 | tags=12%, list=13%, signal=14% |
| 2546 | CELLULAR\_RESPONSE\_TO\_ABIOTIC\_STIMULUS(GO:0071214) |  | 306 | 0.79 | 1.26 | 0.141 | 0.421 | 0.828 | 7937 | tags=12%, list=13%, signal=14% |
| 2547 | NEGATIVE\_REGULATION\_OF\_CELLULAR\_MACROMOLECULE\_BIOSYNTHETIC\_PROCESS(GO:2000113) |  | 1602 | 0.79 | 1.25 | 0.141 | 0.421 | 0.828 | 7364 | tags=11%, list=12%, signal=13% |
| 2548 | CHOLESTEROL\_BINDING(GO:0015485) |  | 43 | 0.81 | 1.25 | 0.141 | 0.420 | 0.828 | 7721 | tags=26%, list=13%, signal=29% |
| 2549 | PURINERGIC\_NUCLEOTIDE\_RECEPTOR\_SIGNALING\_PATHWAY(GO:0035590) |  | 24 | 0.84 | 1.25 | 0.000 | 0.420 | 0.828 | 5955 | tags=21%, list=10%, signal=23% |
| 2550 | NEGATIVE\_REGULATION\_OF\_AUTOPHAGY(GO:0010507) |  | 75 | 0.74 | 1.25 | 0.000 | 0.420 | 0.828 | 6375 | tags=7%, list=11%, signal=7% |
| 2551 | RESPIRATORY\_TUBE\_DEVELOPMENT(GO:0030323) |  | 173 | 0.79 | 1.25 | 0.141 | 0.420 | 0.828 | 8823 | tags=25%, list=15%, signal=29% |
| 2552 | REGULATION\_OF\_SYSTEMIC\_ARTERIAL\_BLOOD\_PRESSURE\_MEDIATED\_BY\_A\_CHEMICAL\_SIGNAL(GO:0003044) |  | 50 | 0.78 | 1.25 | 0.141 | 0.420 | 0.828 | 8969 | tags=30%, list=15%, signal=35% |
| 2553 | NEURON\_PROJECTION\_EXTENSION\_INVOLVED\_IN\_NEURON\_PROJECTION\_GUIDANCE(GO:1902284) |  | 21 | 0.83 | 1.25 | 0.141 | 0.420 | 0.828 | 7362 | tags=33%, list=12%, signal=38% |
| 2554 | AXON\_EXTENSION\_INVOLVED\_IN\_AXON\_GUIDANCE(GO:0048846) |  | 21 | 0.83 | 1.25 | 0.141 | 0.420 | 0.828 | 7362 | tags=33%, list=12%, signal=38% |
| 2555 | CELLULAR\_RESPONSE\_TO\_DSRNA(GO:0071359) |  | 66 | 0.81 | 1.25 | 0.141 | 0.420 | 0.828 | 6255 | tags=5%, list=11%, signal=5% |
| 2556 | REGULATION\_OF\_COENZYME\_METABOLIC\_PROCESS(GO:0051196) |  | 88 | 0.80 | 1.25 | 0.141 | 0.420 | 0.828 | 6006 | tags=5%, list=10%, signal=5% |
| 2557 | CILIARY\_PLASM(GO:0097014) |  | 113 | 0.76 | 1.25 | 0.000 | 0.420 | 0.828 | 11087 | tags=43%, list=19%, signal=53% |
| 2558 | HEPARAN\_SULFATE\_SULFOTRANSFERASE\_ACTIVITY(GO:0034483) |  | 16 | 0.78 | 1.25 | 0.000 | 0.420 | 0.828 | 11047 | tags=44%, list=19%, signal=54% |
| 2559 | LIPID\_LOCALIZATION(GO:0010876) |  | 380 | 0.78 | 1.25 | 0.141 | 0.420 | 0.828 | 7699 | tags=15%, list=13%, signal=17% |
| 2560 | MULTI\_ORGANISM\_BEHAVIOR(GO:0051705) |  | 68 | 0.79 | 1.25 | 0.141 | 0.420 | 0.828 | 7962 | tags=29%, list=13%, signal=34% |
| 2561 | HMG\_BOX\_DOMAIN\_BINDING(GO:0071837) |  | 18 | 0.88 | 1.25 | 0.141 | 0.420 | 0.828 | 6162 | tags=56%, list=10%, signal=62% |
| 2562 | GOLGI\_CISTERNA(GO:0031985) |  | 93 | 0.78 | 1.25 | 0.141 | 0.419 | 0.828 | 7121 | tags=12%, list=12%, signal=13% |
| 2563 | INTERFERON\_GAMMA\_SECRETION(GO:0072643) |  | 17 | 0.82 | 1.25 | 0.141 | 0.419 | 0.828 | 6892 | tags=12%, list=12%, signal=13% |
| 2564 | SOMATODENDRITIC\_COMPARTMENT(GO:0036477) |  | 717 | 0.78 | 1.25 | 0.141 | 0.419 | 0.828 | 7963 | tags=19%, list=13%, signal=21% |
| 2565 | REGULATION\_OF\_SMOOTH\_MUSCLE\_CONTRACTION(GO:0006940) |  | 65 | 0.78 | 1.25 | 0.141 | 0.419 | 0.828 | 9682 | tags=35%, list=16%, signal=42% |
| 2566 | SMOOTH\_MUSCLE\_CELL\_DIFFERENTIATION(GO:0051145) |  | 67 | 0.79 | 1.25 | 0.141 | 0.419 | 0.828 | 7313 | tags=19%, list=12%, signal=22% |
| 2567 | CHOLESTEROL\_METABOLIC\_PROCESS(GO:0008203) |  | 133 | 0.76 | 1.25 | 0.000 | 0.419 | 0.828 | 7358 | tags=14%, list=12%, signal=16% |
| 2568 | GDP\_BINDING(GO:0019003) |  | 59 | 0.83 | 1.25 | 0.141 | 0.419 | 0.828 | 5850 | tags=3%, list=10%, signal=4% |
| 2569 | NEURON\_PROJECTION\_FASCICULATION(GO:0106030) |  | 21 | 0.82 | 1.25 | 0.141 | 0.419 | 0.828 | 5994 | tags=10%, list=10%, signal=11% |
| 2570 | AXONAL\_FASCICULATION(GO:0007413) |  | 21 | 0.82 | 1.25 | 0.141 | 0.419 | 0.828 | 5994 | tags=10%, list=10%, signal=11% |
| 2571 | CILIARY\_BASE(GO:0097546) |  | 29 | 0.75 | 1.25 | 0.000 | 0.419 | 0.828 | 10468 | tags=31%, list=18%, signal=38% |
| 2572 | POSITIVE\_REGULATION\_OF\_SMOOTH\_MUSCLE\_CELL\_APOPTOTIC\_PROCESS(GO:0034393) |  | 16 | 0.77 | 1.25 | 0.000 | 0.419 | 0.828 | 6738 | tags=19%, list=11%, signal=21% |
| 2573 | AMINO\_ACID:CATION\_SYMPORTER\_ACTIVITY(GO:0005416) |  | 22 | 0.80 | 1.25 | 0.141 | 0.419 | 0.828 | 7352 | tags=41%, list=12%, signal=47% |
| 2574 | LENS\_DEVELOPMENT\_IN\_CAMERA\_TYPE\_EYE(GO:0002088) |  | 70 | 0.80 | 1.25 | 0.141 | 0.419 | 0.828 | 8380 | tags=26%, list=14%, signal=30% |
| 2575 | SIGNAL\_RELEASE\_FROM\_SYNAPSE(GO:0099643) |  | 140 | 0.77 | 1.25 | 0.141 | 0.419 | 0.828 | 9227 | tags=28%, list=16%, signal=33% |
| 2576 | NEUROTRANSMITTER\_SECRETION(GO:0007269) |  | 140 | 0.77 | 1.25 | 0.141 | 0.419 | 0.828 | 9227 | tags=28%, list=16%, signal=33% |
| 2577 | POSITIVE\_REGULATION\_OF\_VASOCONSTRICTION(GO:0045907) |  | 35 | 0.80 | 1.25 | 0.141 | 0.419 | 0.828 | 7682 | tags=34%, list=13%, signal=39% |
| 2578 | CATALYTIC\_ACTIVITY\_\_ACTING\_ON\_A\_GLYCOPROTEIN(GO:0140103) |  | 23 | 0.75 | 1.25 | 0.000 | 0.419 | 0.828 | 7412 | tags=22%, list=13%, signal=25% |
| 2579 | REGULATION\_OF\_CARBOHYDRATE\_BIOSYNTHETIC\_PROCESS(GO:0043255) |  | 79 | 0.80 | 1.25 | 0.141 | 0.419 | 0.828 | 6375 | tags=11%, list=11%, signal=13% |
| 2580 | PLATELET\_AGGREGATION(GO:0070527) |  | 57 | 0.79 | 1.25 | 0.141 | 0.419 | 0.828 | 7921 | tags=19%, list=13%, signal=22% |
| 2581 | POSITIVE\_REGULATION\_OF\_MAPK\_CASCADE(GO:0043410) |  | 512 | 0.78 | 1.25 | 0.141 | 0.419 | 0.828 | 7920 | tags=19%, list=13%, signal=21% |
| 2582 | ATPASE\_COMPLEX(GO:1904949) |  | 99 | 0.74 | 1.25 | 0.000 | 0.419 | 0.828 | 6432 | tags=12%, list=11%, signal=14% |
| 2583 | INORGANIC\_ANION\_TRANSMEMBRANE\_TRANSPORTER\_ACTIVITY(GO:0015103) |  | 155 | 0.78 | 1.25 | 0.141 | 0.419 | 0.828 | 8596 | tags=25%, list=15%, signal=29% |
| 2584 | POSITIVE\_REGULATION\_OF\_ENDOCYTOSIS(GO:0045807) |  | 127 | 0.76 | 1.25 | 0.141 | 0.419 | 0.828 | 9368 | tags=21%, list=16%, signal=25% |
| 2585 | POSITIVE\_REGULATION\_OF\_INTERLEUKIN\_1\_PRODUCTION(GO:0032732) |  | 42 | 0.81 | 1.25 | 0.141 | 0.418 | 0.828 | 6473 | tags=10%, list=11%, signal=11% |
| 2586 | REGULATION\_OF\_PROTEIN\_MODIFICATION\_PROCESS(GO:0031399) |  | 1763 | 0.78 | 1.25 | 0.141 | 0.418 | 0.828 | 7969 | tags=13%, list=13%, signal=15% |
| 2587 | SEGMENTATION(GO:0035282) |  | 95 | 0.79 | 1.25 | 0.141 | 0.418 | 0.828 | 7846 | tags=27%, list=13%, signal=32% |
| 2588 | POSITIVE\_REGULATION\_OF\_HEART\_GROWTH(GO:0060421) |  | 48 | 0.79 | 1.25 | 0.141 | 0.418 | 0.828 | 8122 | tags=25%, list=14%, signal=29% |
| 2589 | AMINE\_METABOLIC\_PROCESS(GO:0009308) |  | 137 | 0.78 | 1.25 | 0.141 | 0.418 | 0.828 | 7935 | tags=10%, list=13%, signal=12% |
| 2590 | N\_ACETYLGLUCOSAMINE\_METABOLIC\_PROCESS(GO:0006044) |  | 17 | 0.76 | 1.25 | 0.000 | 0.418 | 0.828 | 9382 | tags=41%, list=16%, signal=49% |
| 2591 | REGULATION\_OF\_CARDIAC\_MUSCLE\_TISSUE\_DEVELOPMENT(GO:0055024) |  | 90 | 0.78 | 1.25 | 0.141 | 0.418 | 0.828 | 8774 | tags=26%, list=15%, signal=30% |
| 2592 | PITUITARY\_GLAND\_DEVELOPMENT(GO:0021983) |  | 43 | 0.84 | 1.25 | 0.141 | 0.418 | 0.828 | 8397 | tags=49%, list=14%, signal=57% |
| 2593 | TRANSPORTER\_ACTIVITY(GO:0005215) |  | 1266 | 0.77 | 1.25 | 0.141 | 0.418 | 0.828 | 8596 | tags=22%, list=15%, signal=25% |
| 2594 | SYNCYTIUM\_FORMATION(GO:0006949) |  | 53 | 0.79 | 1.25 | 0.141 | 0.418 | 0.828 | 9145 | tags=28%, list=15%, signal=33% |
| 2595 | ACTIVATION\_OF\_PROTEIN\_KINASE\_B\_ACTIVITY(GO:0032148) |  | 34 | 0.80 | 1.25 | 0.141 | 0.418 | 0.828 | 6473 | tags=15%, list=11%, signal=17% |
| 2596 | REGULATION\_OF\_COFACTOR\_METABOLIC\_PROCESS(GO:0051193) |  | 102 | 0.79 | 1.25 | 0.141 | 0.418 | 0.828 | 6006 | tags=4%, list=10%, signal=4% |
| 2597 | REGULATION\_OF\_BLOOD\_VESSEL\_ENDOTHELIAL\_CELL\_PROLIFERATION\_INVOLVED\_IN\_SPROUTING\_ANGIOGENESIS(GO:1903587) |  | 41 | 0.83 | 1.25 | 0.141 | 0.418 | 0.828 | 9016 | tags=20%, list=15%, signal=23% |
| 2598 | THYMOCYTE\_APOPTOTIC\_PROCESS(GO:0070242) |  | 17 | 0.78 | 1.25 | 0.000 | 0.418 | 0.828 | 6473 | tags=12%, list=11%, signal=13% |
| 2599 | EXTRINSIC\_COMPONENT\_OF\_CYTOPLASMIC\_SIDE\_OF\_PLASMA\_MEMBRANE(GO:0031234) |  | 112 | 0.79 | 1.25 | 0.141 | 0.418 | 0.828 | 7951 | tags=13%, list=13%, signal=15% |
| 2600 | POSITIVE\_REGULATION\_OF\_PHOSPHATASE\_ACTIVITY(GO:0010922) |  | 30 | 0.75 | 1.25 | 0.000 | 0.418 | 0.828 | 9543 | tags=30%, list=16%, signal=36% |
| 2601 | AMYLOID\_FIBRIL\_FORMATION(GO:1990000) |  | 16 | 0.76 | 1.25 | 0.000 | 0.418 | 0.828 | 6154 | tags=13%, list=10%, signal=14% |
| 2602 | POSITIVE\_REGULATION\_OF\_SYNAPTIC\_TRANSMISSION(GO:0050806) |  | 133 | 0.79 | 1.25 | 0.141 | 0.418 | 0.828 | 7962 | tags=20%, list=13%, signal=23% |
| 2603 | DENTATE\_GYRUS\_DEVELOPMENT(GO:0021542) |  | 17 | 0.89 | 1.25 | 0.141 | 0.418 | 0.828 | 6381 | tags=41%, list=11%, signal=46% |
| 2604 | HIGH\_DENSITY\_LIPOPROTEIN\_PARTICLE\_REMODELING(GO:0034375) |  | 17 | 0.83 | 1.25 | 0.000 | 0.418 | 0.828 | 6362 | tags=29%, list=11%, signal=33% |
| 2605 | SENSORY\_PERCEPTION\_OF\_CHEMICAL\_STIMULUS(GO:0007606) |  | 491 | 0.75 | 1.25 | 0.141 | 0.418 | 0.828 | 9891 | tags=9%, list=17%, signal=10% |
| 2606 | HEMOPOIESIS(GO:0030097) |  | 857 | 0.78 | 1.25 | 0.141 | 0.418 | 0.828 | 7583 | tags=14%, list=13%, signal=16% |
| 2607 | CARDIOBLAST\_DIFFERENTIATION(GO:0010002) |  | 18 | 0.79 | 1.25 | 0.141 | 0.418 | 0.828 | 5958 | tags=22%, list=10%, signal=25% |
| 2608 | POSITIVE\_REGULATION\_OF\_REPRODUCTIVE\_PROCESS(GO:2000243) |  | 57 | 0.80 | 1.25 | 0.141 | 0.418 | 0.828 | 8276 | tags=28%, list=14%, signal=33% |
| 2609 | MULTI\_ORGANISM\_PROCESS(GO:0051704) |  | 2470 | 0.78 | 1.25 | 0.141 | 0.418 | 0.828 | 7796 | tags=13%, list=13%, signal=14% |
| 2610 | NEGATIVE\_REGULATION\_OF\_CELLULAR\_BIOSYNTHETIC\_PROCESS(GO:0031327) |  | 1748 | 0.78 | 1.25 | 0.141 | 0.417 | 0.828 | 7404 | tags=11%, list=13%, signal=13% |
| 2611 | PYRIMIDINE\_CONTAINING\_COMPOUND\_METABOLIC\_PROCESS(GO:0072527) |  | 81 | 0.73 | 1.25 | 0.000 | 0.417 | 0.828 | 8641 | tags=19%, list=15%, signal=22% |
| 2612 | PHOSPHATIDIC\_ACID\_BINDING(GO:0070300) |  | 18 | 0.76 | 1.25 | 0.000 | 0.418 | 0.828 | 7738 | tags=17%, list=13%, signal=19% |
| 2613 | CELLULAR\_AMINE\_METABOLIC\_PROCESS(GO:0044106) |  | 128 | 0.77 | 1.25 | 0.141 | 0.417 | 0.828 | 7935 | tags=9%, list=13%, signal=10% |
| 2614 | REGULATION\_OF\_TRANSCRIPTION\_REGULATORY\_REGION\_DNA\_BINDING(GO:2000677) |  | 48 | 0.82 | 1.25 | 0.141 | 0.417 | 0.828 | 8946 | tags=35%, list=15%, signal=42% |
| 2615 | POSITIVE\_REGULATION\_OF\_ESTABLISHMENT\_OF\_PROTEIN\_LOCALIZATION(GO:1904951) |  | 461 | 0.78 | 1.25 | 0.141 | 0.417 | 0.828 | 7652 | tags=12%, list=13%, signal=14% |
| 2616 | BIOMINERAL\_TISSUE\_DEVELOPMENT(GO:0031214) |  | 133 | 0.77 | 1.25 | 0.141 | 0.417 | 0.828 | 7441 | tags=23%, list=13%, signal=27% |
| 2617 | REGULATION\_OF\_HYDROLASE\_ACTIVITY(GO:0051336) |  | 1254 | 0.77 | 1.25 | 0.141 | 0.417 | 0.828 | 7906 | tags=14%, list=13%, signal=15% |
| 2618 | SYNTAXIN\_BINDING(GO:0019905) |  | 81 | 0.78 | 1.25 | 0.141 | 0.417 | 0.828 | 7426 | tags=17%, list=13%, signal=20% |
| 2619 | INORGANIC\_ANION\_TRANSPORT(GO:0015698) |  | 177 | 0.78 | 1.25 | 0.141 | 0.417 | 0.828 | 8596 | tags=27%, list=15%, signal=32% |
| 2620 | REGULATION\_OF\_MOLECULAR\_FUNCTION(GO:0065009) |  | 3327 | 0.78 | 1.25 | 0.141 | 0.417 | 0.828 | 7979 | tags=15%, list=13%, signal=16% |
| 2621 | REGULATION\_OF\_INTERLEUKIN\_13\_PRODUCTION(GO:0032656) |  | 20 | 0.81 | 1.25 | 0.141 | 0.417 | 0.828 | 6685 | tags=35%, list=11%, signal=39% |
| 2622 | EMBRYONIC\_DIGIT\_MORPHOGENESIS(GO:0042733) |  | 64 | 0.81 | 1.25 | 0.141 | 0.417 | 0.828 | 8177 | tags=27%, list=14%, signal=31% |
| 2623 | GALACTOSYLTRANSFERASE\_ACTIVITY(GO:0008378) |  | 35 | 0.80 | 1.25 | 0.141 | 0.417 | 0.828 | 7412 | tags=20%, list=13%, signal=23% |
| 2624 | NEPHRIC\_DUCT\_DEVELOPMENT(GO:0072176) |  | 15 | 0.82 | 1.25 | 0.141 | 0.417 | 0.828 | 7360 | tags=27%, list=12%, signal=30% |
| 2625 | LEARNING(GO:0007612) |  | 139 | 0.77 | 1.25 | 0.141 | 0.417 | 0.828 | 7962 | tags=21%, list=13%, signal=24% |
| 2626 | IRON\_ION\_BINDING(GO:0005506) |  | 164 | 0.78 | 1.25 | 0.141 | 0.417 | 0.828 | 8511 | tags=21%, list=14%, signal=25% |
| 2627 | OVULATION\_CYCLE\_PROCESS(GO:0022602) |  | 44 | 0.79 | 1.25 | 0.141 | 0.417 | 0.828 | 7081 | tags=25%, list=12%, signal=28% |
| 2628 | NEGATIVE\_REGULATION\_OF\_CYTOKINE\_PRODUCTION\_INVOLVED\_IN\_IMMUNE\_RESPONSE(GO:0002719) |  | 23 | 0.80 | 1.25 | 0.141 | 0.417 | 0.828 | 10458 | tags=43%, list=18%, signal=53% |
| 2629 | POSITIVE\_REGULATION\_OF\_BIOSYNTHETIC\_PROCESS(GO:0009891) |  | 1829 | 0.77 | 1.25 | 0.141 | 0.417 | 0.828 | 8066 | tags=16%, list=14%, signal=18% |
| 2630 | POSITIVE\_REGULATION\_OF\_AUTOPHAGY(GO:0010508) |  | 103 | 0.75 | 1.25 | 0.000 | 0.417 | 0.828 | 7441 | tags=7%, list=13%, signal=8% |
| 2631 | PURINE\_RIBONUCLEOTIDE\_METABOLIC\_PROCESS(GO:0009150) |  | 537 | 0.75 | 1.25 | 0.141 | 0.417 | 0.828 | 7635 | tags=11%, list=13%, signal=13% |
| 2632 | CD4\_POSITIVE\_\_ALPHA\_BETA\_T\_CELL\_ACTIVATION(GO:0035710) |  | 73 | 0.77 | 1.25 | 0.141 | 0.417 | 0.828 | 7488 | tags=26%, list=13%, signal=30% |
| 2633 | ORGANONITROGEN\_COMPOUND\_BIOSYNTHETIC\_PROCESS(GO:1901566) |  | 2150 | 0.77 | 1.25 | 0.141 | 0.417 | 0.828 | 7472 | tags=10%, list=13%, signal=11% |
| 2634 | NEGATIVE\_REGULATION\_OF\_CANONICAL\_WNT\_SIGNALING\_PATHWAY(GO:0090090) |  | 172 | 0.76 | 1.25 | 0.000 | 0.416 | 0.828 | 8109 | tags=16%, list=14%, signal=18% |
| 2635 | GLYCEROLIPID\_BIOSYNTHETIC\_PROCESS(GO:0045017) |  | 274 | 0.73 | 1.25 | 0.000 | 0.416 | 0.828 | 9669 | tags=19%, list=16%, signal=23% |
| 2636 | CYCLASE\_ACTIVITY(GO:0009975) |  | 22 | 0.79 | 1.25 | 0.000 | 0.416 | 0.828 | 9489 | tags=41%, list=16%, signal=49% |
| 2637 | LIPID\_BIOSYNTHETIC\_PROCESS(GO:0008610) |  | 691 | 0.76 | 1.25 | 0.141 | 0.416 | 0.828 | 8607 | tags=16%, list=15%, signal=19% |
| 2638 | INTRACELLULAR\_SIGNAL\_TRANSDUCTION(GO:0035556) |  | 2761 | 0.79 | 1.25 | 0.141 | 0.416 | 0.828 | 7783 | tags=14%, list=13%, signal=15% |
| 2639 | POSITIVE\_REGULATION\_OF\_CYTOKINE\_PRODUCTION(GO:0001819) |  | 410 | 0.75 | 1.25 | 0.141 | 0.416 | 0.828 | 7663 | tags=17%, list=13%, signal=19% |
| 2640 | REGULATION\_OF\_VENTRICULAR\_CARDIAC\_MUSCLE\_CELL\_ACTION\_POTENTIAL(GO:0098911) |  | 18 | 0.80 | 1.25 | 0.141 | 0.416 | 0.828 | 9054 | tags=22%, list=15%, signal=26% |
| 2641 | MICROTUBULE\_POLYMERIZATION\_OR\_DEPOLYMERIZATION(GO:0031109) |  | 101 | 0.72 | 1.25 | 0.000 | 0.416 | 0.828 | 8972 | tags=19%, list=15%, signal=22% |
| 2642 | CELL\_MIGRATION\_INVOLVED\_IN\_GASTRULATION(GO:0042074) |  | 15 | 0.78 | 1.25 | 0.000 | 0.416 | 0.828 | 7343 | tags=33%, list=12%, signal=38% |
| 2643 | OSTEOCLAST\_DIFFERENTIATION(GO:0030316) |  | 92 | 0.76 | 1.25 | 0.141 | 0.416 | 0.828 | 10798 | tags=32%, list=18%, signal=39% |
| 2644 | MITOCHONDRIAL\_MEMBRANE\_PART(GO:0044455) |  | 226 | 0.72 | 1.25 | 0.000 | 0.416 | 0.828 | 5977 | tags=4%, list=10%, signal=4% |
| 2645 | MAGNESIUM\_ION\_BINDING(GO:0000287) |  | 201 | 0.76 | 1.25 | 0.000 | 0.416 | 0.828 | 8235 | tags=11%, list=14%, signal=13% |
| 2646 | SARCOMERE\_ORGANIZATION(GO:0045214) |  | 45 | 0.81 | 1.25 | 0.141 | 0.416 | 0.828 | 8232 | tags=36%, list=14%, signal=41% |
| 2647 | REGULATION\_OF\_CYTOKINE\_BIOSYNTHETIC\_PROCESS(GO:0042035) |  | 96 | 0.79 | 1.25 | 0.141 | 0.416 | 0.828 | 7461 | tags=20%, list=13%, signal=23% |
| 2648 | NEGATIVE\_REGULATION\_OF\_NEURON\_APOPTOTIC\_PROCESS(GO:0043524) |  | 138 | 0.76 | 1.25 | 0.141 | 0.416 | 0.828 | 9445 | tags=28%, list=16%, signal=34% |
| 2649 | ENDOSOME\_LUMEN(GO:0031904) |  | 34 | 0.74 | 1.25 | 0.000 | 0.416 | 0.828 | 6738 | tags=15%, list=11%, signal=17% |
| 2650 | POSITIVE\_REGULATION\_OF\_CELLULAR\_BIOSYNTHETIC\_PROCESS(GO:0031328) |  | 1799 | 0.77 | 1.25 | 0.141 | 0.416 | 0.828 | 8066 | tags=16%, list=14%, signal=18% |
| 2651 | TRANSCRIPTION\_FACTOR\_ACTIVITY\_\_DIRECT\_LIGAND\_REGULATED\_SEQUENCE\_SPECIFIC\_DNA\_BINDING(GO:0098531) |  | 49 | 0.79 | 1.25 | 0.141 | 0.416 | 0.828 | 9711 | tags=31%, list=16%, signal=37% |
| 2652 | NUCLEAR\_RECEPTOR\_ACTIVITY(GO:0004879) |  | 49 | 0.79 | 1.25 | 0.141 | 0.416 | 0.828 | 9711 | tags=31%, list=16%, signal=37% |
| 2653 | RETINOIC\_ACID\_RECEPTOR\_SIGNALING\_PATHWAY(GO:0048384) |  | 30 | 0.79 | 1.25 | 0.141 | 0.416 | 0.828 | 6278 | tags=23%, list=11%, signal=26% |
| 2654 | MYOFIBRIL\_ASSEMBLY(GO:0030239) |  | 66 | 0.80 | 1.25 | 0.141 | 0.416 | 0.828 | 8232 | tags=41%, list=14%, signal=47% |
| 2655 | POSITIVE\_REGULATION\_OF\_T\_CELL\_DIFFERENTIATION(GO:0045582) |  | 73 | 0.78 | 1.25 | 0.141 | 0.415 | 0.828 | 9368 | tags=25%, list=16%, signal=29% |
| 2656 | REGULATION\_OF\_CYTOKINE\_SECRETION(GO:0050707) |  | 174 | 0.75 | 1.25 | 0.141 | 0.415 | 0.828 | 7984 | tags=16%, list=14%, signal=18% |
| 2657 | GLUCOSE\_TRANSMEMBRANE\_TRANSPORTER\_ACTIVITY(GO:0005355) |  | 21 | 0.78 | 1.25 | 0.141 | 0.415 | 0.828 | 8716 | tags=43%, list=15%, signal=50% |
| 2658 | IMMUNOGLOBULIN\_BINDING(GO:0019865) |  | 21 | 0.82 | 1.25 | 0.141 | 0.415 | 0.828 | 7330 | tags=19%, list=12%, signal=22% |
| 2659 | BICARBONATE\_TRANSMEMBRANE\_TRANSPORTER\_ACTIVITY(GO:0015106) |  | 23 | 0.77 | 1.25 | 0.000 | 0.415 | 0.828 | 9957 | tags=48%, list=17%, signal=57% |
| 2660 | EXTRINSIC\_COMPONENT\_OF\_PLASMA\_MEMBRANE(GO:0019897) |  | 155 | 0.79 | 1.25 | 0.141 | 0.415 | 0.828 | 7951 | tags=15%, list=13%, signal=17% |
| 2661 | NEGATIVE\_REGULATION\_OF\_TRANSPORT(GO:0051051) |  | 459 | 0.76 | 1.25 | 0.141 | 0.415 | 0.828 | 8109 | tags=17%, list=14%, signal=20% |
| 2662 | RESPONSE\_TO\_RADIATION(GO:0009314) |  | 441 | 0.78 | 1.25 | 0.141 | 0.415 | 0.828 | 7276 | tags=10%, list=12%, signal=11% |
| 2663 | KINASE\_ACTIVITY(GO:0016301) |  | 914 | 0.75 | 1.25 | 0.000 | 0.415 | 0.828 | 8312 | tags=16%, list=14%, signal=19% |
| 2664 | CELL\_BODY(GO:0044297) |  | 518 | 0.78 | 1.25 | 0.141 | 0.415 | 0.828 | 8771 | tags=22%, list=15%, signal=26% |
| 2665 | PHYSIOLOGICAL\_MUSCLE\_HYPERTROPHY(GO:0003298) |  | 31 | 0.78 | 1.25 | 0.141 | 0.415 | 0.828 | 8112 | tags=16%, list=14%, signal=19% |
| 2666 | PHYSIOLOGICAL\_CARDIAC\_MUSCLE\_HYPERTROPHY(GO:0003301) |  | 31 | 0.78 | 1.25 | 0.141 | 0.415 | 0.828 | 8112 | tags=16%, list=14%, signal=19% |
| 2667 | CELL\_GROWTH\_INVOLVED\_IN\_CARDIAC\_MUSCLE\_CELL\_DEVELOPMENT(GO:0061049) |  | 31 | 0.78 | 1.25 | 0.141 | 0.415 | 0.828 | 8112 | tags=16%, list=14%, signal=19% |
| 2668 | RETINOID\_X\_RECEPTOR\_BINDING(GO:0046965) |  | 16 | 0.80 | 1.25 | 0.000 | 0.415 | 0.828 | 6051 | tags=6%, list=10%, signal=7% |
| 2669 | LIGASE\_ACTIVITY\_\_FORMING\_CARBON\_SULFUR\_BONDS(GO:0016877) |  | 40 | 0.75 | 1.25 | 0.000 | 0.415 | 0.828 | 11022 | tags=28%, list=19%, signal=34% |
| 2670 | LONG\_CHAIN\_FATTY\_ACYL\_COA\_METABOLIC\_PROCESS(GO:0035336) |  | 24 | 0.80 | 1.25 | 0.141 | 0.415 | 0.828 | 6042 | tags=8%, list=10%, signal=9% |
| 2671 | MITOCHONDRIAL\_OUTER\_MEMBRANE(GO:0005741) |  | 183 | 0.75 | 1.25 | 0.000 | 0.414 | 0.828 | 7435 | tags=8%, list=13%, signal=9% |
| 2672 | ACYLGLYCEROL\_O\_ACYLTRANSFERASE\_ACTIVITY(GO:0016411) |  | 28 | 0.78 | 1.25 | 0.000 | 0.414 | 0.828 | 6622 | tags=11%, list=11%, signal=12% |
| 2673 | GOLGI\_TO\_PLASMA\_MEMBRANE\_PROTEIN\_TRANSPORT(GO:0043001) |  | 32 | 0.80 | 1.25 | 0.141 | 0.414 | 0.828 | 5891 | tags=3%, list=10%, signal=3% |
| 2674 | T\_HELPER\_CELL\_DIFFERENTIATION(GO:0042093) |  | 52 | 0.77 | 1.25 | 0.141 | 0.414 | 0.828 | 7414 | tags=21%, list=13%, signal=24% |
| 2675 | PURINERGIC\_RECEPTOR\_SIGNALING\_PATHWAY(GO:0035587) |  | 34 | 0.78 | 1.25 | 0.000 | 0.414 | 0.828 | 5955 | tags=15%, list=10%, signal=16% |
| 2676 | PROTEASOMAL\_PROTEIN\_CATABOLIC\_PROCESS(GO:0010498) |  | 439 | 0.73 | 1.25 | 0.000 | 0.414 | 0.828 | 7795 | tags=6%, list=13%, signal=7% |
| 2677 | INTERLEUKIN\_12\_PRODUCTION(GO:0032615) |  | 53 | 0.78 | 1.25 | 0.141 | 0.414 | 0.828 | 7170 | tags=19%, list=12%, signal=21% |
| 2678 | TRACHEA\_DEVELOPMENT(GO:0060438) |  | 19 | 0.84 | 1.25 | 0.141 | 0.414 | 0.828 | 6395 | tags=21%, list=11%, signal=24% |
| 2679 | NEGATIVE\_REGULATION\_OF\_NUCLEOBASE\_CONTAINING\_COMPOUND\_METABOLIC\_PROCESS(GO:0045934) |  | 1446 | 0.78 | 1.25 | 0.141 | 0.414 | 0.828 | 7404 | tags=12%, list=13%, signal=13% |
| 2680 | PROTEIN\_N\_LINKED\_GLYCOSYLATION(GO:0006487) |  | 87 | 0.73 | 1.25 | 0.000 | 0.414 | 0.828 | 10532 | tags=23%, list=18%, signal=28% |
| 2681 | FATTY\_ACYL\_COA\_BIOSYNTHETIC\_PROCESS(GO:0046949) |  | 31 | 0.76 | 1.25 | 0.000 | 0.414 | 0.828 | 8607 | tags=16%, list=15%, signal=19% |
| 2682 | NEGATIVE\_REGULATION\_OF\_ATP\_METABOLIC\_PROCESS(GO:1903579) |  | 21 | 0.84 | 1.25 | 0.141 | 0.414 | 0.828 | 5894 | tags=5%, list=10%, signal=5% |
| 2683 | RESPONSE\_TO\_RETINOIC\_ACID(GO:0032526) |  | 104 | 0.79 | 1.25 | 0.141 | 0.414 | 0.828 | 8122 | tags=25%, list=14%, signal=29% |
| 2684 | ENDOCHONDRAL\_BONE\_GROWTH(GO:0003416) |  | 22 | 0.81 | 1.25 | 0.141 | 0.414 | 0.828 | 6229 | tags=32%, list=11%, signal=36% |
| 2685 | SCHWANN\_CELL\_DIFFERENTIATION(GO:0014037) |  | 35 | 0.78 | 1.25 | 0.141 | 0.414 | 0.828 | 6850 | tags=9%, list=12%, signal=10% |
| 2686 | PROTEIN\_CONTAINING\_COMPLEX\_SUBUNIT\_ORGANIZATION(GO:0043933) |  | 2112 | 0.78 | 1.25 | 0.141 | 0.414 | 0.828 | 7659 | tags=10%, list=13%, signal=11% |
| 2687 | SODIUM\_CHANNEL\_ACTIVITY(GO:0005272) |  | 45 | 0.82 | 1.25 | 0.141 | 0.414 | 0.828 | 6844 | tags=24%, list=12%, signal=28% |
| 2688 | NEGATIVE\_REGULATION\_OF\_CELLULAR\_AMIDE\_METABOLIC\_PROCESS(GO:0034249) |  | 428 | 0.75 | 1.25 | 0.141 | 0.414 | 0.828 | 6397 | tags=7%, list=11%, signal=7% |
| 2689 | POLY\_PYRIMIDINE\_TRACT\_BINDING(GO:0008187) |  | 24 | 0.75 | 1.25 | 0.000 | 0.414 | 0.828 | 9997 | tags=25%, list=17%, signal=30% |
| 2690 | DENDRITIC\_TREE(GO:0097447) |  | 519 | 0.78 | 1.25 | 0.141 | 0.414 | 0.828 | 7963 | tags=18%, list=13%, signal=20% |
| 2691 | ACTIN\_FILAMENT\_BUNDLE\_ORGANIZATION(GO:0061572) |  | 143 | 0.78 | 1.25 | 0.141 | 0.414 | 0.828 | 8112 | tags=18%, list=14%, signal=21% |
| 2692 | DENDRITE(GO:0030425) |  | 518 | 0.78 | 1.25 | 0.141 | 0.414 | 0.828 | 7963 | tags=18%, list=13%, signal=20% |
| 2693 | ANION\_TRANSPORT(GO:0006820) |  | 598 | 0.77 | 1.25 | 0.141 | 0.414 | 0.828 | 8474 | tags=21%, list=14%, signal=24% |
| 2694 | SECONDARY\_ACTIVE\_TRANSMEMBRANE\_TRANSPORTER\_ACTIVITY(GO:0015291) |  | 238 | 0.75 | 1.25 | 0.141 | 0.414 | 0.828 | 10298 | tags=33%, list=17%, signal=40% |
| 2695 | EPHRIN\_RECEPTOR\_ACTIVITY(GO:0005003) |  | 19 | 0.82 | 1.25 | 0.141 | 0.414 | 0.828 | 7211 | tags=32%, list=12%, signal=36% |
| 2696 | POSITIVE\_REGULATION\_OF\_RECEPTOR\_INTERNALIZATION(GO:0002092) |  | 25 | 0.78 | 1.25 | 0.000 | 0.414 | 0.828 | 9105 | tags=32%, list=15%, signal=38% |
| 2697 | GROWTH\_OF\_SYMBIONT\_IN\_HOST(GO:0044117) |  | 19 | 0.79 | 1.25 | 0.141 | 0.414 | 0.828 | 9039 | tags=26%, list=15%, signal=31% |
| 2698 | GROWTH\_INVOLVED\_IN\_SYMBIOTIC\_INTERACTION(GO:0044110) |  | 19 | 0.79 | 1.25 | 0.141 | 0.414 | 0.828 | 9039 | tags=26%, list=15%, signal=31% |
| 2699 | GROWTH\_OF\_SYMBIONT\_INVOLVED\_IN\_INTERACTION\_WITH\_HOST(GO:0044116) |  | 19 | 0.79 | 1.25 | 0.141 | 0.414 | 0.828 | 9039 | tags=26%, list=15%, signal=31% |
| 2700 | ALCOHOL\_CATABOLIC\_PROCESS(GO:0046164) |  | 46 | 0.78 | 1.25 | 0.141 | 0.414 | 0.828 | 5920 | tags=9%, list=10%, signal=10% |
| 2701 | CHANNEL\_REGULATOR\_ACTIVITY(GO:0016247) |  | 133 | 0.78 | 1.25 | 0.141 | 0.414 | 0.828 | 7977 | tags=28%, list=13%, signal=32% |
| 2702 | HOMOTYPIC\_CELL\_CELL\_ADHESION(GO:0034109) |  | 76 | 0.78 | 1.25 | 0.141 | 0.414 | 0.828 | 9054 | tags=25%, list=15%, signal=29% |
| 2703 | NEGATIVE\_REGULATION\_OF\_PURINE\_NUCLEOTIDE\_METABOLIC\_PROCESS(GO:1900543) |  | 56 | 0.79 | 1.25 | 0.141 | 0.414 | 0.828 | 8474 | tags=11%, list=14%, signal=12% |
| 2704 | AXONEME\_ASSEMBLY(GO:0035082) |  | 58 | 0.77 | 1.25 | 0.141 | 0.414 | 0.828 | 8290 | tags=24%, list=14%, signal=28% |
| 2705 | TRANSCRIPTION\_BY\_RNA\_POLYMERASE\_II(GO:0006366) |  | 2401 | 0.78 | 1.25 | 0.141 | 0.414 | 0.828 | 7450 | tags=15%, list=13%, signal=16% |
| 2706 | GABA\_A\_RECEPTOR\_ACTIVITY(GO:0004890) |  | 19 | 0.88 | 1.25 | 0.000 | 0.414 | 0.828 | 7049 | tags=32%, list=12%, signal=36% |
| 2707 | PROTEIN\_LOCALIZATION(GO:0008104) |  | 2578 | 0.79 | 1.25 | 0.141 | 0.413 | 0.828 | 7583 | tags=10%, list=13%, signal=10% |
| 2708 | ADP\_BINDING(GO:0043531) |  | 36 | 0.75 | 1.25 | 0.000 | 0.413 | 0.828 | 9341 | tags=25%, list=16%, signal=30% |
| 2709 | POSITIVE\_REGULATION\_OF\_LIPID\_BIOSYNTHETIC\_PROCESS(GO:0046889) |  | 65 | 0.75 | 1.25 | 0.000 | 0.413 | 0.828 | 9326 | tags=22%, list=16%, signal=26% |
| 2710 | RESPONSE\_TO\_PURINE\_CONTAINING\_COMPOUND(GO:0014074) |  | 147 | 0.76 | 1.25 | 0.141 | 0.413 | 0.828 | 7873 | tags=20%, list=13%, signal=23% |
| 2711 | RIBONUCLEOTIDE\_METABOLIC\_PROCESS(GO:0009259) |  | 552 | 0.74 | 1.25 | 0.141 | 0.413 | 0.828 | 7635 | tags=11%, list=13%, signal=13% |
| 2712 | HEXOSE\_TRANSMEMBRANE\_TRANSPORTER\_ACTIVITY(GO:0015149) |  | 22 | 0.77 | 1.25 | 0.141 | 0.413 | 0.828 | 8716 | tags=41%, list=15%, signal=48% |
| 2713 | NEGATIVE\_REGULATION\_OF\_DEVELOPMENTAL\_GROWTH(GO:0048640) |  | 89 | 0.81 | 1.25 | 0.141 | 0.413 | 0.828 | 8673 | tags=28%, list=15%, signal=33% |
| 2714 | PROTEIN\_TYROSINE\_KINASE\_BINDING(GO:1990782) |  | 70 | 0.74 | 1.25 | 0.000 | 0.413 | 0.828 | 10041 | tags=23%, list=17%, signal=27% |
| 2715 | ALDEHYDE\_DEHYDROGENASE\_(NAD)\_ACTIVITY(GO:0004029) |  | 16 | 0.84 | 1.25 | 0.000 | 0.413 | 0.828 | 6438 | tags=25%, list=11%, signal=28% |
| 2716 | POSITIVE\_REGULATION\_OF\_MACROMOLECULE\_BIOSYNTHETIC\_PROCESS(GO:0010557) |  | 1691 | 0.77 | 1.25 | 0.141 | 0.413 | 0.828 | 8066 | tags=16%, list=14%, signal=18% |
| 2717 | ARGININE\_METABOLIC\_PROCESS(GO:0006525) |  | 21 | 0.76 | 1.25 | 0.000 | 0.413 | 0.828 | 10309 | tags=48%, list=17%, signal=58% |
| 2718 | ZINC\_ION\_BINDING(GO:0008270) |  | 814 | 0.77 | 1.25 | 0.141 | 0.413 | 0.828 | 7955 | tags=14%, list=13%, signal=16% |
| 2719 | SECONDARY\_ALCOHOL\_METABOLIC\_PROCESS(GO:1902652) |  | 138 | 0.75 | 1.25 | 0.000 | 0.413 | 0.828 | 7358 | tags=14%, list=12%, signal=16% |
| 2720 | DRUG\_BINDING(GO:0008144) |  | 1777 | 0.77 | 1.25 | 0.141 | 0.413 | 0.828 | 7675 | tags=12%, list=13%, signal=14% |
| 2721 | SOMITOGENESIS(GO:0001756) |  | 66 | 0.78 | 1.25 | 0.000 | 0.413 | 0.828 | 7837 | tags=30%, list=13%, signal=35% |
| 2722 | REGULATION\_OF\_GLYCOLYTIC\_PROCESS(GO:0006110) |  | 67 | 0.79 | 1.24 | 0.141 | 0.413 | 0.828 | 6006 | tags=4%, list=10%, signal=5% |
| 2723 | REGULATION\_OF\_VIRAL\_LIFE\_CYCLE(GO:1903900) |  | 135 | 0.77 | 1.24 | 0.141 | 0.413 | 0.828 | 7293 | tags=10%, list=12%, signal=12% |
| 2724 | BONE\_REMODELING(GO:0046849) |  | 83 | 0.76 | 1.24 | 0.141 | 0.413 | 0.828 | 8241 | tags=25%, list=14%, signal=29% |
| 2725 | GOLGI\_TO\_PLASMA\_MEMBRANE\_TRANSPORT(GO:0006893) |  | 47 | 0.78 | 1.24 | 0.141 | 0.413 | 0.828 | 7457 | tags=9%, list=13%, signal=10% |
| 2726 | CELLULAR\_LIPID\_CATABOLIC\_PROCESS(GO:0044242) |  | 198 | 0.75 | 1.24 | 0.000 | 0.413 | 0.828 | 8139 | tags=13%, list=14%, signal=15% |
| 2727 | REGULATION\_OF\_CELL\_FATE\_COMMITMENT(GO:0010453) |  | 35 | 0.81 | 1.24 | 0.141 | 0.413 | 0.828 | 6209 | tags=17%, list=10%, signal=19% |
| 2728 | MORPHOGENESIS\_OF\_AN\_EPITHELIAL\_BUD(GO:0060572) |  | 17 | 0.83 | 1.24 | 0.141 | 0.412 | 0.828 | 6473 | tags=18%, list=11%, signal=20% |
| 2729 | POLY(U)\_RNA\_BINDING(GO:0008266) |  | 20 | 0.77 | 1.24 | 0.000 | 0.412 | 0.828 | 9639 | tags=25%, list=16%, signal=30% |
| 2730 | CAMP\_METABOLIC\_PROCESS(GO:0046058) |  | 104 | 0.73 | 1.24 | 0.000 | 0.412 | 0.828 | 7935 | tags=22%, list=13%, signal=25% |
| 2731 | NEGATIVE\_REGULATION\_OF\_CELLULAR\_PROTEIN\_LOCALIZATION(GO:1903828) |  | 103 | 0.75 | 1.24 | 0.000 | 0.412 | 0.828 | 7973 | tags=16%, list=13%, signal=18% |
| 2732 | ATP\_DEPENDENT\_MICROTUBULE\_MOTOR\_ACTIVITY\_\_MINUS\_END\_DIRECTED(GO:0008569) |  | 18 | 0.76 | 1.24 | 0.000 | 0.412 | 0.828 | 8776 | tags=44%, list=15%, signal=52% |
| 2733 | RESPONSE\_TO\_AXON\_INJURY(GO:0048678) |  | 66 | 0.76 | 1.24 | 0.141 | 0.412 | 0.828 | 8743 | tags=30%, list=15%, signal=36% |
| 2734 | DENDRITE\_MEMBRANE(GO:0032590) |  | 24 | 0.78 | 1.24 | 0.000 | 0.412 | 0.828 | 8210 | tags=38%, list=14%, signal=44% |
| 2735 | AMYLOID\_PRECURSOR\_PROTEIN\_METABOLIC\_PROCESS(GO:0042982) |  | 46 | 0.75 | 1.24 | 0.141 | 0.412 | 0.828 | 7068 | tags=7%, list=12%, signal=7% |
| 2736 | POSITIVE\_REGULATION\_OF\_SECRETION\_BY\_CELL(GO:1903532) |  | 364 | 0.75 | 1.24 | 0.141 | 0.412 | 0.828 | 7937 | tags=15%, list=13%, signal=17% |
| 2737 | SECONDARY\_METABOLITE\_BIOSYNTHETIC\_PROCESS(GO:0044550) |  | 25 | 0.80 | 1.24 | 0.141 | 0.412 | 0.828 | 8714 | tags=44%, list=15%, signal=52% |
| 2738 | RIBOSE\_PHOSPHATE\_METABOLIC\_PROCESS(GO:0019693) |  | 567 | 0.74 | 1.24 | 0.141 | 0.412 | 0.828 | 7635 | tags=11%, list=13%, signal=12% |
| 2739 | ALPHA\_ACTININ\_BINDING(GO:0051393) |  | 33 | 0.78 | 1.24 | 0.000 | 0.412 | 0.828 | 7652 | tags=27%, list=13%, signal=31% |
| 2740 | HEPARIN\_BINDING(GO:0008201) |  | 156 | 0.80 | 1.24 | 0.141 | 0.412 | 0.828 | 8073 | tags=31%, list=14%, signal=36% |
| 2741 | POSITIVE\_REGULATION\_OF\_ADHERENS\_JUNCTION\_ORGANIZATION(GO:1903393) |  | 26 | 0.79 | 1.24 | 0.141 | 0.412 | 0.828 | 6027 | tags=12%, list=10%, signal=13% |
| 2742 | POSITIVE\_REGULATION\_OF\_CYCLIC\_NUCLEOTIDE\_METABOLIC\_PROCESS(GO:0030801) |  | 57 | 0.74 | 1.24 | 0.000 | 0.412 | 0.828 | 8279 | tags=26%, list=14%, signal=31% |
| 2743 | NEGATIVE\_REGULATION\_OF\_OSSIFICATION(GO:0030279) |  | 72 | 0.80 | 1.24 | 0.141 | 0.412 | 0.828 | 7373 | tags=28%, list=12%, signal=32% |
| 2744 | PEPTIDYL\_THREONINE\_PHOSPHORYLATION(GO:0018107) |  | 120 | 0.73 | 1.24 | 0.000 | 0.412 | 0.828 | 8046 | tags=13%, list=14%, signal=15% |
| 2745 | VESICLE\_FUSION(GO:0006906) |  | 160 | 0.76 | 1.24 | 0.141 | 0.412 | 0.828 | 9144 | tags=15%, list=15%, signal=18% |
| 2746 | GABA\_A\_RECEPTOR\_COMPLEX(GO:1902711) |  | 17 | 0.88 | 1.24 | 0.000 | 0.412 | 0.828 | 7049 | tags=35%, list=12%, signal=40% |
| 2747 | NUCLEAR\_TRANSCRIBED\_MRNA\_CATABOLIC\_PROCESS\_\_DEADENYLATION\_DEPENDENT\_DECAY(GO:0000288) |  | 80 | 0.77 | 1.24 | 0.141 | 0.412 | 0.828 | 6508 | tags=5%, list=11%, signal=6% |
| 2748 | REGULATION\_OF\_ACTIN\_FILAMENT\_BASED\_PROCESS(GO:0032970) |  | 349 | 0.77 | 1.24 | 0.141 | 0.412 | 0.828 | 7962 | tags=15%, list=13%, signal=17% |
| 2749 | NEGATIVE\_REGULATION\_OF\_CARDIAC\_MUSCLE\_TISSUE\_DEVELOPMENT(GO:0055026) |  | 37 | 0.79 | 1.24 | 0.141 | 0.411 | 0.828 | 8380 | tags=24%, list=14%, signal=28% |
| 2750 | DETECTION\_OF\_STIMULUS\_INVOLVED\_IN\_SENSORY\_PERCEPTION(GO:0050906) |  | 484 | 0.71 | 1.24 | 0.141 | 0.411 | 0.828 | 9774 | tags=10%, list=17%, signal=12% |
| 2751 | CHONDROITIN\_SULFATE\_PROTEOGLYCAN\_BIOSYNTHETIC\_PROCESS(GO:0050650) |  | 30 | 0.76 | 1.24 | 0.000 | 0.411 | 0.828 | 9569 | tags=50%, list=16%, signal=60% |
| 2752 | MACROMOLECULE\_LOCALIZATION(GO:0033036) |  | 2907 | 0.79 | 1.24 | 0.141 | 0.411 | 0.828 | 7583 | tags=10%, list=13%, signal=11% |
| 2753 | SIDE\_OF\_MEMBRANE(GO:0098552) |  | 491 | 0.76 | 1.24 | 0.141 | 0.411 | 0.828 | 8063 | tags=15%, list=14%, signal=17% |
| 2754 | GLUCOSE\_HOMEOSTASIS(GO:0042593) |  | 219 | 0.75 | 1.24 | 0.141 | 0.411 | 0.828 | 8987 | tags=17%, list=15%, signal=20% |
| 2755 | CARBOHYDRATE\_HOMEOSTASIS(GO:0033500) |  | 219 | 0.75 | 1.24 | 0.141 | 0.411 | 0.828 | 8987 | tags=17%, list=15%, signal=20% |
| 2756 | ENDOCYTIC\_RECYCLING(GO:0032456) |  | 33 | 0.81 | 1.24 | 0.141 | 0.411 | 0.828 | 6259 | tags=6%, list=11%, signal=7% |
| 2757 | REGULATION\_OF\_NUCLEOTIDE\_CATABOLIC\_PROCESS(GO:0030811) |  | 71 | 0.78 | 1.24 | 0.141 | 0.411 | 0.828 | 6006 | tags=4%, list=10%, signal=5% |
| 2758 | 1\_ACYLGLYCEROL\_3\_PHOSPHATE\_O\_ACYLTRANSFERASE\_ACTIVITY(GO:0003841) |  | 19 | 0.79 | 1.24 | 0.000 | 0.411 | 0.828 | 5931 | tags=5%, list=10%, signal=6% |
| 2759 | POSITIVE\_REGULATION\_OF\_FOCAL\_ADHESION\_ASSEMBLY(GO:0051894) |  | 24 | 0.80 | 1.24 | 0.141 | 0.411 | 0.828 | 6027 | tags=13%, list=10%, signal=14% |
| 2760 | STEM\_CELL\_DIVISION(GO:0017145) |  | 42 | 0.79 | 1.24 | 0.141 | 0.411 | 0.828 | 8953 | tags=33%, list=15%, signal=39% |
| 2761 | POSITIVE\_REGULATION\_OF\_CELL\_CYCLE(GO:0045787) |  | 368 | 0.77 | 1.24 | 0.141 | 0.411 | 0.828 | 7640 | tags=8%, list=13%, signal=9% |
| 2762 | PEPTIDASE\_ACTIVITY\_\_ACTING\_ON\_L\_AMINO\_ACID\_PEPTIDES(GO:0070011) |  | 701 | 0.76 | 1.24 | 0.141 | 0.411 | 0.828 | 8285 | tags=13%, list=14%, signal=15% |
| 2763 | MYOBLAST\_FUSION(GO:0007520) |  | 39 | 0.78 | 1.24 | 0.141 | 0.411 | 0.828 | 10390 | tags=26%, list=18%, signal=31% |
| 2764 | ORGANELLE\_MEMBRANE\_FUSION(GO:0090174) |  | 165 | 0.75 | 1.24 | 0.141 | 0.411 | 0.828 | 9144 | tags=15%, list=15%, signal=17% |
| 2765 | REGULATION\_OF\_INTERLEUKIN\_12\_PRODUCTION(GO:0032655) |  | 51 | 0.78 | 1.24 | 0.141 | 0.411 | 0.828 | 7170 | tags=20%, list=12%, signal=22% |
| 2766 | REGULATION\_OF\_CARBOHYDRATE\_CATABOLIC\_PROCESS(GO:0043470) |  | 76 | 0.78 | 1.24 | 0.141 | 0.411 | 0.828 | 6006 | tags=4%, list=10%, signal=4% |
| 2767 | REGULATION\_OF\_PURINE\_NUCLEOTIDE\_METABOLIC\_PROCESS(GO:1900542) |  | 191 | 0.77 | 1.24 | 0.141 | 0.410 | 0.828 | 8541 | tags=16%, list=14%, signal=18% |
| 2768 | NEGATIVE\_REGULATION\_OF\_PROTEIN\_COMPLEX\_ASSEMBLY(GO:0031333) |  | 128 | 0.75 | 1.24 | 0.141 | 0.410 | 0.828 | 7859 | tags=10%, list=13%, signal=12% |
| 2769 | GLYCOSAMINOGLYCAN\_BINDING(GO:0005539) |  | 215 | 0.79 | 1.24 | 0.141 | 0.410 | 0.828 | 8242 | tags=28%, list=14%, signal=33% |
| 2770 | POSITIVE\_REGULATION\_OF\_PROTEIN\_SERINE\_THREONINE\_KINASE\_ACTIVITY(GO:0071902) |  | 349 | 0.75 | 1.24 | 0.141 | 0.410 | 0.828 | 7837 | tags=15%, list=13%, signal=17% |
| 2771 | PHOSPHORUS\_OXYGEN\_LYASE\_ACTIVITY(GO:0016849) |  | 22 | 0.79 | 1.24 | 0.000 | 0.410 | 0.828 | 7265 | tags=36%, list=12%, signal=41% |
| 2772 | TRANSCRIPTION\_FACTOR\_BINDING(GO:0008134) |  | 598 | 0.75 | 1.24 | 0.000 | 0.410 | 0.828 | 7907 | tags=12%, list=13%, signal=14% |
| 2773 | STEROL\_METABOLIC\_PROCESS(GO:0016125) |  | 151 | 0.75 | 1.24 | 0.000 | 0.410 | 0.828 | 8511 | tags=17%, list=14%, signal=19% |
| 2774 | DEVELOPMENTAL\_PIGMENTATION(GO:0048066) |  | 45 | 0.76 | 1.24 | 0.141 | 0.410 | 0.828 | 7134 | tags=27%, list=12%, signal=30% |
| 2775 | MULTICELLULAR\_ORGANISMAL\_RESPONSE\_TO\_STRESS(GO:0033555) |  | 68 | 0.78 | 1.24 | 0.141 | 0.410 | 0.828 | 7873 | tags=25%, list=13%, signal=29% |
| 2776 | REGULATION\_OF\_GENE\_EXPRESSION(GO:0010468) |  | 4846 | 0.79 | 1.24 | 0.141 | 0.410 | 0.828 | 7456 | tags=12%, list=13%, signal=12% |
| 2777 | MORPHOGENESIS\_OF\_AN\_EPITHELIAL\_FOLD(GO:0060571) |  | 27 | 0.82 | 1.24 | 0.141 | 0.410 | 0.828 | 8122 | tags=33%, list=14%, signal=39% |
| 2778 | REGULATION\_OF\_AXON\_EXTENSION\_INVOLVED\_IN\_AXON\_GUIDANCE(GO:0048841) |  | 16 | 0.82 | 1.24 | 0.141 | 0.410 | 0.828 | 8673 | tags=31%, list=15%, signal=37% |
| 2779 | PROTEIN\_AUTOPHOSPHORYLATION(GO:0046777) |  | 235 | 0.74 | 1.24 | 0.000 | 0.410 | 0.828 | 8112 | tags=16%, list=14%, signal=19% |
| 2780 | REGULATION\_OF\_HISTONE\_MODIFICATION(GO:0031056) |  | 135 | 0.74 | 1.24 | 0.000 | 0.410 | 0.828 | 8774 | tags=11%, list=15%, signal=13% |
| 2781 | OXIDATION\_REDUCTION\_PROCESS(GO:0055114) |  | 1005 | 0.75 | 1.24 | 0.141 | 0.410 | 0.828 | 8607 | tags=14%, list=15%, signal=16% |
| 2782 | STEROID\_BINDING(GO:0005496) |  | 86 | 0.78 | 1.24 | 0.141 | 0.410 | 0.828 | 7721 | tags=20%, list=13%, signal=23% |
| 2783 | NEURON\_PROJECTION\_CYTOPLASM(GO:0120111) |  | 68 | 0.74 | 1.24 | 0.000 | 0.409 | 0.828 | 7426 | tags=13%, list=13%, signal=15% |
| 2784 | ORGANIC\_ANION\_TRANSPORT(GO:0015711) |  | 463 | 0.77 | 1.24 | 0.141 | 0.409 | 0.828 | 8474 | tags=21%, list=14%, signal=24% |
| 2785 | T\_CELL\_DIFFERENTIATION(GO:0030217) |  | 224 | 0.76 | 1.24 | 0.141 | 0.409 | 0.828 | 7488 | tags=16%, list=13%, signal=18% |
| 2786 | POSITIVE\_REGULATION\_OF\_NUCLEOBASE\_CONTAINING\_COMPOUND\_METABOLIC\_PROCESS(GO:0045935) |  | 1734 | 0.77 | 1.24 | 0.141 | 0.410 | 0.828 | 8066 | tags=16%, list=14%, signal=18% |
| 2787 | NEGATIVE\_REGULATION\_OF\_GTPASE\_ACTIVITY(GO:0034260) |  | 47 | 0.76 | 1.24 | 0.141 | 0.409 | 0.828 | 7426 | tags=15%, list=13%, signal=17% |
| 2788 | POSITIVE\_REGULATION\_OF\_ENDOTHELIAL\_CELL\_MIGRATION(GO:0010595) |  | 114 | 0.78 | 1.24 | 0.141 | 0.409 | 0.828 | 8046 | tags=22%, list=14%, signal=25% |
| 2789 | REGULATION\_OF\_SKELETAL\_MUSCLE\_TISSUE\_DEVELOPMENT(GO:0048641) |  | 53 | 0.79 | 1.24 | 0.141 | 0.409 | 0.828 | 8774 | tags=32%, list=15%, signal=38% |
| 2790 | ACTIN\_FILAMENT\_BUNDLE\_ASSEMBLY(GO:0051017) |  | 139 | 0.78 | 1.24 | 0.141 | 0.409 | 0.828 | 8112 | tags=17%, list=14%, signal=19% |
| 2791 | VENTRICULAR\_CARDIAC\_MUSCLE\_TISSUE\_MORPHOGENESIS(GO:0055010) |  | 48 | 0.76 | 1.24 | 0.000 | 0.409 | 0.828 | 8328 | tags=31%, list=14%, signal=36% |
| 2792 | TRANSFERASE\_ACTIVITY\_\_TRANSFERRING\_PHOSPHORUS\_CONTAINING\_GROUPS(GO:0016772) |  | 1074 | 0.74 | 1.24 | 0.000 | 0.409 | 0.828 | 8272 | tags=15%, list=14%, signal=17% |
| 2793 | BLOOD\_VESSEL\_ENDOTHELIAL\_CELL\_PROLIFERATION\_INVOLVED\_IN\_SPROUTING\_ANGIOGENESIS(GO:0002043) |  | 47 | 0.82 | 1.24 | 0.141 | 0.409 | 0.828 | 9016 | tags=21%, list=15%, signal=25% |
| 2794 | G\_PROTEIN\_COUPLED\_PEPTIDE\_RECEPTOR\_ACTIVITY(GO:0008528) |  | 132 | 0.79 | 1.24 | 0.141 | 0.409 | 0.828 | 9434 | tags=31%, list=16%, signal=37% |
| 2795 | CARBOXYLIC\_ESTER\_HYDROLASE\_ACTIVITY(GO:0052689) |  | 144 | 0.78 | 1.24 | 0.141 | 0.409 | 0.828 | 7270 | tags=15%, list=12%, signal=17% |
| 2796 | DNA\_GEOMETRIC\_CHANGE(GO:0032392) |  | 84 | 0.72 | 1.24 | 0.000 | 0.409 | 0.828 | 5860 | tags=1%, list=10%, signal=1% |
| 2797 | STEROL\_TRANSPORT(GO:0015918) |  | 92 | 0.78 | 1.24 | 0.141 | 0.409 | 0.828 | 7358 | tags=16%, list=12%, signal=19% |
| 2798 | HISTONE\_ACETYLATION(GO:0016573) |  | 150 | 0.74 | 1.24 | 0.000 | 0.409 | 0.828 | 7358 | tags=7%, list=12%, signal=8% |
| 2799 | LUNG\_MORPHOGENESIS(GO:0060425) |  | 52 | 0.81 | 1.24 | 0.141 | 0.409 | 0.828 | 8122 | tags=27%, list=14%, signal=31% |
| 2800 | POSITIVE\_REGULATION\_OF\_VASCULAR\_ENDOTHELIAL\_GROWTH\_FACTOR\_PRODUCTION(GO:0010575) |  | 27 | 0.78 | 1.24 | 0.000 | 0.409 | 0.828 | 8279 | tags=30%, list=14%, signal=34% |
| 2801 | NEGATIVE\_REGULATION\_OF\_ORGANELLE\_ORGANIZATION(GO:0010639) |  | 350 | 0.77 | 1.24 | 0.141 | 0.409 | 0.828 | 6126 | tags=7%, list=10%, signal=7% |
| 2802 | REGULATION\_OF\_CELL\_CELL\_ADHESION(GO:0022407) |  | 374 | 0.75 | 1.24 | 0.141 | 0.409 | 0.828 | 9023 | tags=18%, list=15%, signal=22% |
| 2803 | MODIFICATION\_OF\_MORPHOLOGY\_OR\_PHYSIOLOGY\_OF\_OTHER\_ORGANISM\_INVOLVED\_IN\_SYMBIOTIC\_INTERACTION(GO:0051817) |  | 107 | 0.73 | 1.24 | 0.000 | 0.408 | 0.828 | 7980 | tags=13%, list=13%, signal=15% |
| 2804 | MONOSACCHARIDE\_METABOLIC\_PROCESS(GO:0005996) |  | 279 | 0.78 | 1.24 | 0.141 | 0.408 | 0.828 | 7414 | tags=9%, list=13%, signal=11% |
| 2805 | MOTILE\_CILIUM(GO:0031514) |  | 154 | 0.76 | 1.24 | 0.141 | 0.408 | 0.828 | 10544 | tags=37%, list=18%, signal=45% |
| 2806 | PHOSPHOLIPID\_BINDING(GO:0005543) |  | 384 | 0.75 | 1.24 | 0.141 | 0.408 | 0.828 | 8363 | tags=17%, list=14%, signal=20% |
| 2807 | GLUTAMINE\_FAMILY\_AMINO\_ACID\_BIOSYNTHETIC\_PROCESS(GO:0009084) |  | 20 | 0.75 | 1.24 | 0.000 | 0.408 | 0.828 | 6020 | tags=10%, list=10%, signal=11% |
| 2808 | REGULATION\_OF\_GENERATION\_OF\_PRECURSOR\_METABOLITES\_AND\_ENERGY(GO:0043467) |  | 123 | 0.78 | 1.24 | 0.141 | 0.408 | 0.828 | 6247 | tags=7%, list=11%, signal=7% |
| 2809 | DNA\_BINDING(GO:0003677) |  | 2487 | 0.78 | 1.24 | 0.141 | 0.408 | 0.828 | 7631 | tags=13%, list=13%, signal=14% |
| 2810 | ACTIVATED\_T\_CELL\_PROLIFERATION(GO:0050798) |  | 43 | 0.72 | 1.24 | 0.000 | 0.408 | 0.828 | 9023 | tags=23%, list=15%, signal=27% |
| 2811 | INTEGRAL\_COMPONENT\_OF\_PLASMA\_MEMBRANE(GO:0005887) |  | 1604 | 0.78 | 1.24 | 0.141 | 0.408 | 0.828 | 9472 | tags=26%, list=16%, signal=31% |
| 2812 | ORGANOPHOSPHATE\_METABOLIC\_PROCESS(GO:0019637) |  | 1201 | 0.75 | 1.24 | 0.141 | 0.408 | 0.828 | 7937 | tags=12%, list=13%, signal=14% |
| 2813 | CORONARY\_VASCULATURE\_DEVELOPMENT(GO:0060976) |  | 50 | 0.74 | 1.24 | 0.000 | 0.408 | 0.828 | 7898 | tags=18%, list=13%, signal=21% |
| 2814 | NEGATIVE\_REGULATION\_OF\_T\_CELL\_APOPTOTIC\_PROCESS(GO:0070233) |  | 18 | 0.80 | 1.24 | 0.000 | 0.408 | 0.828 | 6685 | tags=17%, list=11%, signal=19% |
| 2815 | INSULIN\_SECRETION(GO:0030073) |  | 206 | 0.74 | 1.24 | 0.141 | 0.408 | 0.828 | 10124 | tags=23%, list=17%, signal=28% |
| 2816 | PHOSPHOLIPID\_BIOSYNTHETIC\_PROCESS(GO:0008654) |  | 286 | 0.72 | 1.24 | 0.000 | 0.408 | 0.828 | 9669 | tags=18%, list=16%, signal=21% |
| 2817 | RESPONSE\_TO\_ANESTHETIC(GO:0072347) |  | 76 | 0.79 | 1.24 | 0.141 | 0.408 | 0.828 | 7298 | tags=17%, list=12%, signal=19% |
| 2818 | TRANSCRIPTION\_FACTOR\_ACTIVITY\_\_RNA\_POLYMERASE\_II\_TRANSCRIPTION\_FACTOR\_BINDING(GO:0001076) |  | 157 | 0.78 | 1.24 | 0.141 | 0.408 | 0.828 | 6559 | tags=10%, list=11%, signal=11% |
| 2819 | RESPIRATORY\_CHAIN\_COMPLEX(GO:0098803) |  | 89 | 0.72 | 1.24 | 0.000 | 0.408 | 0.828 | 5894 | tags=6%, list=10%, signal=6% |
| 2820 | GLOMERULAR\_EPITHELIUM\_DEVELOPMENT(GO:0072010) |  | 20 | 0.76 | 1.24 | 0.141 | 0.408 | 0.828 | 8937 | tags=35%, list=15%, signal=41% |
| 2821 | NEGATIVE\_REGULATION\_OF\_TRANSMEMBRANE\_TRANSPORT(GO:0034763) |  | 119 | 0.75 | 1.24 | 0.141 | 0.408 | 0.828 | 7979 | tags=17%, list=13%, signal=19% |
| 2822 | ATP\_DEPENDENT\_CHROMATIN\_REMODELING(GO:0043044) |  | 82 | 0.75 | 1.24 | 0.000 | 0.408 | 0.828 | 10645 | tags=22%, list=18%, signal=27% |
| 2823 | RESPONSE\_TO\_PH(GO:0009268) |  | 41 | 0.76 | 1.24 | 0.000 | 0.407 | 0.828 | 11630 | tags=44%, list=20%, signal=55% |
| 2824 | PEPTIDE\_RECEPTOR\_ACTIVITY(GO:0001653) |  | 133 | 0.79 | 1.24 | 0.141 | 0.407 | 0.828 | 9434 | tags=31%, list=16%, signal=37% |
| 2825 | RNA\_POLYMERASE\_II\_CORE\_PROMOTER\_SEQUENCE\_SPECIFIC\_DNA\_BINDING(GO:0000979) |  | 31 | 0.81 | 1.24 | 0.141 | 0.407 | 0.828 | 6271 | tags=6%, list=11%, signal=7% |
| 2826 | INTERNAL\_PEPTIDYL\_LYSINE\_ACETYLATION(GO:0018393) |  | 154 | 0.73 | 1.24 | 0.000 | 0.407 | 0.828 | 7358 | tags=8%, list=12%, signal=9% |
| 2827 | LEUKOCYTE\_HOMEOSTASIS(GO:0001776) |  | 85 | 0.71 | 1.24 | 0.000 | 0.407 | 0.828 | 7545 | tags=14%, list=13%, signal=16% |
| 2828 | POSITIVE\_REGULATION\_OF\_SECRETION(GO:0051047) |  | 390 | 0.75 | 1.24 | 0.141 | 0.407 | 0.828 | 7937 | tags=16%, list=13%, signal=18% |
| 2829 | PEPTIDYL\_TYROSINE\_DEPHOSPHORYLATION(GO:0035335) |  | 101 | 0.75 | 1.24 | 0.000 | 0.407 | 0.828 | 9557 | tags=20%, list=16%, signal=24% |
| 2830 | REGULATION\_OF\_CARBOHYDRATE\_METABOLIC\_PROCESS(GO:0006109) |  | 175 | 0.76 | 1.24 | 0.141 | 0.407 | 0.828 | 8594 | tags=14%, list=15%, signal=16% |
| 2831 | CELLULAR\_RESPONSE\_TO\_RETINOIC\_ACID(GO:0071300) |  | 69 | 0.78 | 1.24 | 0.141 | 0.407 | 0.828 | 8122 | tags=26%, list=14%, signal=30% |
| 2832 | ANION\_TRANSMEMBRANE\_TRANSPORT(GO:0098656) |  | 251 | 0.76 | 1.24 | 0.141 | 0.407 | 0.828 | 9451 | tags=25%, list=16%, signal=29% |
| 2833 | TRANSITION\_METAL\_ION\_BINDING(GO:0046914) |  | 1079 | 0.76 | 1.24 | 0.141 | 0.407 | 0.828 | 7967 | tags=15%, list=13%, signal=17% |
| 2834 | PEPTIDASE\_ACTIVITY(GO:0008233) |  | 729 | 0.75 | 1.24 | 0.141 | 0.407 | 0.828 | 7956 | tags=12%, list=13%, signal=14% |
| 2835 | ALCOHOL\_BINDING(GO:0043178) |  | 79 | 0.78 | 1.24 | 0.141 | 0.407 | 0.828 | 8521 | tags=29%, list=14%, signal=34% |
| 2836 | BINDING\_OF\_SPERM\_TO\_ZONA\_PELLUCIDA(GO:0007339) |  | 38 | 0.74 | 1.24 | 0.000 | 0.407 | 0.828 | 12193 | tags=32%, list=21%, signal=40% |
| 2837 | NEGATIVE\_REGULATION\_OF\_NUCLEOTIDE\_METABOLIC\_PROCESS(GO:0045980) |  | 57 | 0.78 | 1.24 | 0.141 | 0.407 | 0.828 | 8474 | tags=11%, list=14%, signal=12% |
| 2838 | PROTEIN\_TYROSINE\_PHOSPHATASE\_ACTIVITY(GO:0004725) |  | 103 | 0.74 | 1.24 | 0.000 | 0.407 | 0.828 | 9557 | tags=19%, list=16%, signal=23% |
| 2839 | SULFUR\_COMPOUND\_TRANSPORT(GO:0072348) |  | 36 | 0.74 | 1.24 | 0.000 | 0.407 | 0.828 | 10186 | tags=39%, list=17%, signal=47% |
| 2840 | SOMATIC\_STEM\_CELL\_DIVISION(GO:0048103) |  | 25 | 0.82 | 1.24 | 0.000 | 0.407 | 0.828 | 8953 | tags=44%, list=15%, signal=52% |
| 2841 | PHOSPHATIDYLINOSITOL\_PHOSPHATE\_BINDING(GO:1901981) |  | 138 | 0.74 | 1.24 | 0.000 | 0.407 | 0.828 | 9358 | tags=17%, list=16%, signal=21% |
| 2842 | CD4\_POSITIVE\_\_ALPHA\_BETA\_T\_CELL\_DIFFERENTIATION\_INVOLVED\_IN\_IMMUNE\_RESPONSE(GO:0002294) |  | 54 | 0.76 | 1.24 | 0.141 | 0.407 | 0.828 | 7414 | tags=20%, list=13%, signal=23% |
| 2843 | PEPTIDYL\_THREONINE\_MODIFICATION(GO:0018210) |  | 127 | 0.73 | 1.24 | 0.000 | 0.407 | 0.828 | 7632 | tags=13%, list=13%, signal=14% |
| 2844 | LAMELLIPODIUM\_ORGANIZATION(GO:0097581) |  | 80 | 0.79 | 1.24 | 0.141 | 0.407 | 0.828 | 8605 | tags=20%, list=15%, signal=23% |
| 2845 | ORGANELLE\_MEMBRANE\_CONTACT\_SITE(GO:0044232) |  | 17 | 0.76 | 1.24 | 0.000 | 0.407 | 0.828 | 9101 | tags=35%, list=15%, signal=42% |
| 2846 | TRANSCRIPTIONAL\_REPRESSOR\_ACTIVITY\_\_RNA\_POLYMERASE\_II\_PROXIMAL\_PROMOTER\_SEQUENCE\_SPECIFIC\_DNA\_BINDING(GO:0001078) |  | 134 | 0.78 | 1.24 | 0.141 | 0.407 | 0.828 | 8647 | tags=29%, list=15%, signal=34% |
| 2847 | POSITIVE\_REGULATION\_OF\_CYSTEINE\_TYPE\_ENDOPEPTIDASE\_ACTIVITY\_INVOLVED\_IN\_APOPTOTIC\_PROCESS(GO:0043280) |  | 121 | 0.73 | 1.24 | 0.000 | 0.407 | 0.828 | 7754 | tags=12%, list=13%, signal=13% |
| 2848 | DOPAMINERGIC\_NEURON\_DIFFERENTIATION(GO:0071542) |  | 36 | 0.81 | 1.24 | 0.141 | 0.407 | 0.828 | 8673 | tags=42%, list=15%, signal=49% |
| 2849 | PHOSPHOTRANSFERASE\_ACTIVITY\_\_ALCOHOL\_GROUP\_AS\_ACCEPTOR(GO:0016773) |  | 808 | 0.74 | 1.24 | 0.141 | 0.407 | 0.828 | 8115 | tags=16%, list=14%, signal=19% |
| 2850 | ENHANCER\_BINDING(GO:0035326) |  | 106 | 0.76 | 1.24 | 0.141 | 0.406 | 0.828 | 7385 | tags=20%, list=12%, signal=23% |
| 2851 | O\_ACYLTRANSFERASE\_ACTIVITY(GO:0008374) |  | 49 | 0.74 | 1.24 | 0.000 | 0.406 | 0.828 | 11950 | tags=31%, list=20%, signal=38% |
| 2852 | MUCOSAL\_IMMUNE\_RESPONSE(GO:0002385) |  | 31 | 0.79 | 1.24 | 0.141 | 0.406 | 0.828 | 10916 | tags=23%, list=18%, signal=28% |
| 2853 | GROWTH\_CONE\_PART(GO:0090723) |  | 16 | 0.75 | 1.24 | 0.000 | 0.406 | 0.828 | 10556 | tags=38%, list=18%, signal=46% |
| 2854 | PODOSOME(GO:0002102) |  | 30 | 0.75 | 1.24 | 0.000 | 0.407 | 0.828 | 7787 | tags=10%, list=13%, signal=12% |
| 2855 | NUCLEOBASE\_CONTAINING\_SMALL\_MOLECULE\_CATABOLIC\_PROCESS(GO:0034656) |  | 47 | 0.77 | 1.24 | 0.141 | 0.406 | 0.828 | 6865 | tags=13%, list=12%, signal=14% |
| 2856 | MICROTUBULE\_CYTOSKELETON\_ORGANIZATION(GO:0000226) |  | 485 | 0.73 | 1.24 | 0.000 | 0.406 | 0.828 | 10026 | tags=23%, list=17%, signal=28% |
| 2857 | POSITIVE\_REGULATION\_OF\_DENDRITE\_EXTENSION(GO:1903861) |  | 18 | 0.75 | 1.24 | 0.000 | 0.407 | 0.828 | 8949 | tags=39%, list=15%, signal=46% |
| 2858 | REGULATION\_OF\_EXECUTION\_PHASE\_OF\_APOPTOSIS(GO:1900117) |  | 24 | 0.77 | 1.24 | 0.000 | 0.406 | 0.828 | 10461 | tags=25%, list=18%, signal=30% |
| 2859 | MITOCHONDRIAL\_CALCIUM\_ION\_HOMEOSTASIS(GO:0051560) |  | 25 | 0.74 | 1.24 | 0.000 | 0.406 | 0.828 | 6198 | tags=8%, list=10%, signal=9% |
| 2860 | CELLULAR\_LOCALIZATION(GO:0051641) |  | 2664 | 0.78 | 1.24 | 0.141 | 0.406 | 0.828 | 7588 | tags=9%, list=13%, signal=9% |
| 2861 | INTRINSIC\_COMPONENT\_OF\_PLASMA\_MEMBRANE(GO:0031226) |  | 1666 | 0.78 | 1.24 | 0.141 | 0.406 | 0.828 | 9472 | tags=26%, list=16%, signal=30% |
| 2862 | POSITIVE\_REGULATION\_OF\_SKELETAL\_MUSCLE\_TISSUE\_DEVELOPMENT(GO:0048643) |  | 28 | 0.77 | 1.24 | 0.141 | 0.406 | 0.828 | 8673 | tags=29%, list=15%, signal=33% |
| 2863 | INTERNAL\_PROTEIN\_AMINO\_ACID\_ACETYLATION(GO:0006475) |  | 160 | 0.73 | 1.24 | 0.000 | 0.406 | 0.828 | 7358 | tags=9%, list=12%, signal=10% |
| 2864 | NEGATIVE\_REGULATION\_OF\_CELLULAR\_PROCESS(GO:0048523) |  | 4730 | 0.79 | 1.24 | 0.141 | 0.406 | 0.828 | 7472 | tags=13%, list=13%, signal=13% |
| 2865 | REGULATION\_OF\_CANONICAL\_WNT\_SIGNALING\_PATHWAY(GO:0060828) |  | 262 | 0.74 | 1.24 | 0.000 | 0.406 | 0.828 | 8771 | tags=17%, list=15%, signal=20% |
| 2866 | MEMBRANE\_PROTEIN\_COMPLEX(GO:0098796) |  | 1155 | 0.76 | 1.24 | 0.141 | 0.406 | 0.828 | 7979 | tags=14%, list=13%, signal=16% |
| 2867 | MITOCHONDRION(GO:0005739) |  | 1744 | 0.75 | 1.24 | 0.141 | 0.406 | 0.828 | 6841 | tags=6%, list=12%, signal=6% |
| 2868 | REGULATION\_OF\_PURINE\_NUCLEOTIDE\_BIOSYNTHETIC\_PROCESS(GO:1900371) |  | 148 | 0.76 | 1.24 | 0.141 | 0.406 | 0.828 | 8541 | tags=16%, list=14%, signal=18% |
| 2869 | LIGAND\_DEPENDENT\_NUCLEAR\_RECEPTOR\_TRANSCRIPTION\_COACTIVATOR\_ACTIVITY(GO:0030374) |  | 51 | 0.74 | 1.24 | 0.000 | 0.406 | 0.828 | 6759 | tags=10%, list=11%, signal=11% |
| 2870 | PROTEIN\_C\_TERMINUS\_BINDING(GO:0008022) |  | 190 | 0.72 | 1.24 | 0.000 | 0.406 | 0.828 | 8763 | tags=13%, list=15%, signal=15% |
| 2871 | PROTEIN\_KINASE\_A\_BINDING(GO:0051018) |  | 49 | 0.74 | 1.24 | 0.000 | 0.406 | 0.828 | 8605 | tags=29%, list=15%, signal=33% |
| 2872 | PLATELET\_ACTIVATION(GO:0030168) |  | 160 | 0.77 | 1.24 | 0.141 | 0.406 | 0.828 | 7921 | tags=19%, list=13%, signal=22% |
| 2873 | ENHANCER\_SEQUENCE\_SPECIFIC\_DNA\_BINDING(GO:0001158) |  | 92 | 0.78 | 1.24 | 0.141 | 0.406 | 0.828 | 7385 | tags=22%, list=12%, signal=25% |
| 2874 | REGULATION\_OF\_OSTEOCLAST\_DIFFERENTIATION(GO:0045670) |  | 65 | 0.76 | 1.24 | 0.141 | 0.406 | 0.828 | 10798 | tags=34%, list=18%, signal=41% |
| 2875 | CELL\_PROLIFERATION\_INVOLVED\_IN\_KIDNEY\_DEVELOPMENT(GO:0072111) |  | 19 | 0.84 | 1.24 | 0.141 | 0.406 | 0.828 | 7360 | tags=26%, list=12%, signal=30% |
| 2876 | UBIQUITIN\_LIKE\_PROTEIN\_CONJUGATING\_ENZYME\_BINDING(GO:0044390) |  | 40 | 0.75 | 1.24 | 0.000 | 0.406 | 0.828 | 8490 | tags=20%, list=14%, signal=23% |
| 2877 | PROTEIN\_KINASE\_C\_SIGNALING(GO:0070528) |  | 31 | 0.83 | 1.24 | 0.141 | 0.406 | 0.828 | 6473 | tags=23%, list=11%, signal=25% |
| 2878 | MELANIN\_BIOSYNTHETIC\_PROCESS(GO:0042438) |  | 17 | 0.81 | 1.24 | 0.141 | 0.406 | 0.828 | 8714 | tags=53%, list=15%, signal=62% |
| 2879 | REGULATION\_OF\_NUCLEOTIDE\_BIOSYNTHETIC\_PROCESS(GO:0030808) |  | 149 | 0.76 | 1.24 | 0.141 | 0.406 | 0.828 | 8541 | tags=15%, list=14%, signal=18% |
| 2880 | NEGATIVE\_REGULATION\_OF\_GENE\_EXPRESSION(GO:0010629) |  | 1982 | 0.77 | 1.24 | 0.141 | 0.406 | 0.828 | 7450 | tags=11%, list=13%, signal=12% |
| 2881 | INNATE\_IMMUNE\_RESPONSE\_IN\_MUCOSA(GO:0002227) |  | 21 | 0.79 | 1.24 | 0.141 | 0.406 | 0.828 | 10916 | tags=19%, list=18%, signal=23% |
| 2882 | POSITIVE\_REGULATION\_OF\_LYMPHOCYTE\_DIFFERENTIATION(GO:0045621) |  | 84 | 0.76 | 1.24 | 0.141 | 0.406 | 0.828 | 9368 | tags=25%, list=16%, signal=30% |
| 2883 | LYSOSOME\_LOCALIZATION(GO:0032418) |  | 68 | 0.76 | 1.24 | 0.141 | 0.405 | 0.828 | 9722 | tags=19%, list=16%, signal=23% |
| 2884 | REGULATION\_OF\_SKELETAL\_MUSCLE\_CELL\_DIFFERENTIATION(GO:2001014) |  | 20 | 0.77 | 1.24 | 0.000 | 0.405 | 0.828 | 9208 | tags=45%, list=16%, signal=53% |
| 2885 | POSITIVE\_REGULATION\_OF\_PROTEIN\_TARGETING\_TO\_MITOCHONDRION(GO:1903955) |  | 82 | 0.76 | 1.24 | 0.141 | 0.406 | 0.828 | 7422 | tags=9%, list=13%, signal=10% |
| 2886 | HEXOSE\_METABOLIC\_PROCESS(GO:0019318) |  | 235 | 0.77 | 1.24 | 0.141 | 0.405 | 0.828 | 7414 | tags=9%, list=13%, signal=10% |
| 2887 | MULTI\_ORGANISM\_REPRODUCTIVE\_PROCESS(GO:0044703) |  | 987 | 0.76 | 1.24 | 0.141 | 0.405 | 0.828 | 8690 | tags=19%, list=15%, signal=22% |
| 2888 | REGULATION\_OF\_LYMPHOCYTE\_DIFFERENTIATION(GO:0045619) |  | 155 | 0.77 | 1.24 | 0.141 | 0.405 | 0.828 | 9411 | tags=21%, list=16%, signal=25% |
| 2889 | SIGNAL\_RELEASE(GO:0023061) |  | 434 | 0.75 | 1.24 | 0.141 | 0.405 | 0.828 | 10131 | tags=25%, list=17%, signal=30% |
| 2890 | PHOSPHATIDYLINOSITOL\_BISPHOSPHATE\_BINDING(GO:1902936) |  | 92 | 0.75 | 1.24 | 0.141 | 0.405 | 0.828 | 8363 | tags=17%, list=14%, signal=20% |
| 2891 | RENAL\_ABSORPTION(GO:0070293) |  | 16 | 0.81 | 1.24 | 0.141 | 0.405 | 0.828 | 8363 | tags=31%, list=14%, signal=36% |
| 2892 | DICARBOXYLIC\_ACID\_TRANSMEMBRANE\_TRANSPORTER\_ACTIVITY(GO:0005310) |  | 39 | 0.73 | 1.24 | 0.000 | 0.405 | 0.828 | 11402 | tags=46%, list=19%, signal=57% |
| 2893 | NEGATIVE\_REGULATION\_OF\_IMMUNE\_EFFECTOR\_PROCESS(GO:0002698) |  | 106 | 0.74 | 1.24 | 0.141 | 0.405 | 0.828 | 10458 | tags=25%, list=18%, signal=30% |
| 2894 | DETECTION\_OF\_MECHANICAL\_STIMULUS(GO:0050982) |  | 45 | 0.76 | 1.23 | 0.000 | 0.405 | 0.828 | 10033 | tags=49%, list=17%, signal=59% |
| 2895 | PROTEIN\_TYROSINE\_KINASE\_ACTIVITY(GO:0004713) |  | 184 | 0.75 | 1.23 | 0.141 | 0.405 | 0.828 | 8365 | tags=26%, list=14%, signal=30% |
| 2896 | POSITIVE\_REGULATION\_OF\_CELL\_CELL\_ADHESION(GO:0022409) |  | 245 | 0.75 | 1.23 | 0.141 | 0.405 | 0.828 | 9023 | tags=19%, list=15%, signal=22% |
| 2897 | REGULATION\_OF\_ATP\_BIOSYNTHETIC\_PROCESS(GO:2001169) |  | 75 | 0.78 | 1.23 | 0.141 | 0.405 | 0.828 | 6006 | tags=4%, list=10%, signal=4% |
| 2898 | REGULATION\_OF\_EPITHELIAL\_TUBE\_FORMATION(GO:1905276) |  | 18 | 0.82 | 1.23 | 0.141 | 0.405 | 0.828 | 6473 | tags=17%, list=11%, signal=19% |
| 2899 | TRIGLYCERIDE\_CATABOLIC\_PROCESS(GO:0019433) |  | 33 | 0.76 | 1.23 | 0.000 | 0.405 | 0.828 | 11563 | tags=36%, list=20%, signal=45% |
| 2900 | RNA\_POLYMERASE\_II\_TRANSCRIPTION\_FACTOR\_BINDING(GO:0001085) |  | 119 | 0.78 | 1.23 | 0.141 | 0.405 | 0.828 | 5958 | tags=8%, list=10%, signal=8% |
| 2901 | NEGATIVE\_REGULATION\_OF\_ACTIN\_FILAMENT\_BUNDLE\_ASSEMBLY(GO:0032232) |  | 25 | 0.81 | 1.23 | 0.141 | 0.405 | 0.828 | 7962 | tags=24%, list=13%, signal=28% |
| 2902 | PEPTIDYL\_LYSINE\_ACETYLATION(GO:0018394) |  | 160 | 0.73 | 1.23 | 0.000 | 0.405 | 0.828 | 7358 | tags=8%, list=12%, signal=9% |
| 2903 | FATTY\_ACID\_BIOSYNTHETIC\_PROCESS(GO:0006633) |  | 133 | 0.76 | 1.23 | 0.141 | 0.405 | 0.828 | 8043 | tags=15%, list=14%, signal=17% |
| 2904 | B\_CELL\_PROLIFERATION(GO:0042100) |  | 92 | 0.74 | 1.23 | 0.141 | 0.405 | 0.828 | 10693 | tags=36%, list=18%, signal=44% |
| 2905 | APOPTOTIC\_PROCESS\_INVOLVED\_IN\_MORPHOGENESIS(GO:0060561) |  | 25 | 0.78 | 1.23 | 0.000 | 0.404 | 0.828 | 7260 | tags=24%, list=12%, signal=27% |
| 2906 | REGULATION\_OF\_DNA\_DAMAGE\_RESPONSE\_\_SIGNAL\_TRANSDUCTION\_BY\_P53\_CLASS\_MEDIATOR(GO:0043516) |  | 30 | 0.80 | 1.23 | 0.000 | 0.405 | 0.828 | 8177 | tags=20%, list=14%, signal=23% |
| 2907 | RESPIRATORY\_SYSTEM\_PROCESS(GO:0003016) |  | 30 | 0.73 | 1.23 | 0.000 | 0.405 | 0.828 | 11201 | tags=40%, list=19%, signal=49% |
| 2908 | TRANSMEMBRANE\_RECEPTOR\_PROTEIN\_SERINE\_THREONINE\_KINASE\_SIGNALING\_PATHWAY(GO:0007178) |  | 334 | 0.75 | 1.23 | 0.141 | 0.405 | 0.828 | 8177 | tags=20%, list=14%, signal=23% |
| 2909 | LYSOPHOSPHOLIPID\_ACYLTRANSFERASE\_ACTIVITY(GO:0071617) |  | 21 | 0.78 | 1.23 | 0.000 | 0.405 | 0.828 | 9965 | tags=33%, list=17%, signal=40% |
| 2910 | LYSOPHOSPHATIDIC\_ACID\_ACYLTRANSFERASE\_ACTIVITY(GO:0042171) |  | 21 | 0.78 | 1.23 | 0.000 | 0.405 | 0.828 | 9965 | tags=33%, list=17%, signal=40% |
| 2911 | REGULATION\_OF\_MACROMOLECULE\_BIOSYNTHETIC\_PROCESS(GO:0010556) |  | 4440 | 0.78 | 1.23 | 0.141 | 0.405 | 0.828 | 7450 | tags=12%, list=13%, signal=12% |
| 2912 | POSITIVE\_REGULATION\_OF\_TYROSINE\_PHOSPHORYLATION\_OF\_STAT\_PROTEIN(GO:0042531) |  | 64 | 0.77 | 1.23 | 0.141 | 0.404 | 0.828 | 10643 | tags=30%, list=18%, signal=36% |
| 2913 | CELLULAR\_COMPONENT\_ASSEMBLY(GO:0022607) |  | 2843 | 0.78 | 1.23 | 0.141 | 0.404 | 0.828 | 7973 | tags=12%, list=13%, signal=14% |
| 2914 | SYNAPTIC\_VESICLE\_CYCLE(GO:0099504) |  | 124 | 0.76 | 1.23 | 0.141 | 0.404 | 0.828 | 9395 | tags=26%, list=16%, signal=31% |
| 2915 | STRIATUM\_DEVELOPMENT(GO:0021756) |  | 17 | 0.86 | 1.23 | 0.000 | 0.404 | 0.828 | 7426 | tags=29%, list=13%, signal=34% |
| 2916 | REGULATION\_OF\_NUCLEOTIDE\_METABOLIC\_PROCESS(GO:0006140) |  | 198 | 0.76 | 1.23 | 0.141 | 0.404 | 0.828 | 8541 | tags=15%, list=14%, signal=18% |
| 2917 | CELLULAR\_COMPONENT\_MAINTENANCE(GO:0043954) |  | 40 | 0.74 | 1.23 | 0.000 | 0.404 | 0.828 | 7487 | tags=10%, list=13%, signal=11% |
| 2918 | POSITIVE\_REGULATION\_OF\_MOLECULAR\_FUNCTION(GO:0044093) |  | 1704 | 0.76 | 1.23 | 0.141 | 0.404 | 0.828 | 7979 | tags=14%, list=13%, signal=15% |
| 2919 | REGULATION\_OF\_T\_CELL\_DIFFERENTIATION(GO:0045580) |  | 129 | 0.77 | 1.23 | 0.141 | 0.404 | 0.828 | 7488 | tags=16%, list=13%, signal=19% |
| 2920 | POSITIVE\_REGULATION\_OF\_LYMPHOCYTE\_CHEMOTAXIS(GO:0140131) |  | 15 | 0.77 | 1.23 | 0.000 | 0.404 | 0.828 | 6473 | tags=20%, list=11%, signal=22% |
| 2921 | NEGATIVE\_REGULATION\_OF\_CYTOKINE\_SECRETION(GO:0050710) |  | 58 | 0.74 | 1.23 | 0.141 | 0.404 | 0.828 | 9648 | tags=19%, list=16%, signal=23% |
| 2922 | PATTERN\_BINDING(GO:0001871) |  | 22 | 0.74 | 1.23 | 0.141 | 0.404 | 0.828 | 11717 | tags=41%, list=20%, signal=51% |
| 2923 | POLYSACCHARIDE\_BINDING(GO:0030247) |  | 22 | 0.74 | 1.23 | 0.141 | 0.404 | 0.828 | 11717 | tags=41%, list=20%, signal=51% |
| 2924 | NEGATIVE\_REGULATION\_OF\_SUPRAMOLECULAR\_FIBER\_ORGANIZATION(GO:1902904) |  | 122 | 0.76 | 1.23 | 0.141 | 0.404 | 0.828 | 7973 | tags=15%, list=13%, signal=17% |
| 2925 | POSITIVE\_REGULATION\_OF\_EXCITATORY\_POSTSYNAPTIC\_POTENTIAL(GO:2000463) |  | 24 | 0.78 | 1.23 | 0.141 | 0.404 | 0.828 | 7962 | tags=33%, list=13%, signal=39% |
| 2926 | ADRENAL\_GLAND\_DEVELOPMENT(GO:0030325) |  | 21 | 0.78 | 1.23 | 0.141 | 0.404 | 0.828 | 6617 | tags=33%, list=11%, signal=38% |
| 2927 | RESPONSE\_TO\_AMMONIUM\_ION(GO:0060359) |  | 127 | 0.74 | 1.23 | 0.000 | 0.404 | 0.828 | 7671 | tags=15%, list=13%, signal=17% |
| 2928 | TRANSFERASE\_ACTIVITY(GO:0016740) |  | 2504 | 0.76 | 1.23 | 0.141 | 0.404 | 0.828 | 8341 | tags=13%, list=14%, signal=15% |
| 2929 | DNA\_DEPENDENT\_ATPASE\_ACTIVITY(GO:0008094) |  | 85 | 0.72 | 1.23 | 0.000 | 0.404 | 0.828 | 11599 | tags=29%, list=20%, signal=37% |
| 2930 | MODIFICATION\_OF\_MORPHOLOGY\_OR\_PHYSIOLOGY\_OF\_OTHER\_ORGANISM(GO:0035821) |  | 151 | 0.72 | 1.23 | 0.000 | 0.404 | 0.828 | 8121 | tags=11%, list=14%, signal=13% |
| 2931 | EXTRACELLULAR\_MATRIX\_DISASSEMBLY(GO:0022617) |  | 92 | 0.78 | 1.23 | 0.141 | 0.403 | 0.828 | 8165 | tags=20%, list=14%, signal=23% |
| 2932 | GLUTAMATE\_METABOLIC\_PROCESS(GO:0006536) |  | 31 | 0.74 | 1.23 | 0.000 | 0.403 | 0.828 | 10534 | tags=32%, list=18%, signal=39% |
| 2933 | RESPONSE\_TO\_DSRNA(GO:0043331) |  | 98 | 0.77 | 1.23 | 0.141 | 0.403 | 0.828 | 7158 | tags=11%, list=12%, signal=13% |
| 2934 | NEUROPEPTIDE\_HORMONE\_ACTIVITY(GO:0005184) |  | 28 | 0.82 | 1.23 | 0.141 | 0.403 | 0.828 | 8485 | tags=29%, list=14%, signal=33% |
| 2935 | REGULATION\_OF\_CGMP\_BIOSYNTHETIC\_PROCESS(GO:0030826) |  | 22 | 0.76 | 1.23 | 0.000 | 0.403 | 0.828 | 8279 | tags=27%, list=14%, signal=32% |
| 2936 | HOMOLOGOUS\_CHROMOSOME\_SEGREGATION(GO:0045143) |  | 60 | 0.78 | 1.23 | 0.141 | 0.403 | 0.828 | 9155 | tags=37%, list=15%, signal=43% |
| 2937 | STRESS\_FIBER(GO:0001725) |  | 53 | 0.76 | 1.23 | 0.141 | 0.403 | 0.828 | 8594 | tags=19%, list=15%, signal=22% |
| 2938 | CONTRACTILE\_ACTIN\_FILAMENT\_BUNDLE(GO:0097517) |  | 53 | 0.76 | 1.23 | 0.141 | 0.403 | 0.828 | 8594 | tags=19%, list=15%, signal=22% |
| 2939 | SYNAPSIS(GO:0007129) |  | 47 | 0.78 | 1.23 | 0.141 | 0.403 | 0.828 | 9155 | tags=36%, list=15%, signal=43% |
| 2940 | REGULATION\_OF\_SPINDLE\_CHECKPOINT(GO:0090231) |  | 15 | 0.74 | 1.23 | 0.000 | 0.403 | 0.828 | 6039 | tags=7%, list=10%, signal=7% |
| 2941 | REGULATION\_OF\_MULTI\_ORGANISM\_PROCESS(GO:0043900) |  | 387 | 0.75 | 1.23 | 0.141 | 0.403 | 0.828 | 7777 | tags=13%, list=13%, signal=15% |
| 2942 | REGULATION\_OF\_PROTEIN\_TRANSPORT(GO:0051223) |  | 687 | 0.76 | 1.23 | 0.141 | 0.403 | 0.828 | 7652 | tags=13%, list=13%, signal=14% |
| 2943 | NEGATIVE\_REGULATION\_OF\_NEURON\_DEATH(GO:1901215) |  | 189 | 0.73 | 1.23 | 0.141 | 0.403 | 0.828 | 9445 | tags=25%, list=16%, signal=29% |
| 2944 | CELLULAR\_RESPONSE\_TO\_METAL\_ION(GO:0071248) |  | 156 | 0.76 | 1.23 | 0.141 | 0.403 | 0.828 | 8349 | tags=17%, list=14%, signal=20% |
| 2945 | TISSUE\_REMODELING(GO:0048771) |  | 170 | 0.74 | 1.23 | 0.141 | 0.403 | 0.828 | 10332 | tags=27%, list=17%, signal=33% |
| 2946 | ORGANIC\_ACID:SODIUM\_SYMPORTER\_ACTIVITY(GO:0005343) |  | 31 | 0.79 | 1.23 | 0.141 | 0.403 | 0.828 | 9376 | tags=48%, list=16%, signal=57% |
| 2947 | ALPHA\_BETA\_T\_CELL\_DIFFERENTIATION\_INVOLVED\_IN\_IMMUNE\_RESPONSE(GO:0002293) |  | 55 | 0.76 | 1.23 | 0.141 | 0.403 | 0.828 | 7414 | tags=20%, list=13%, signal=23% |
| 2948 | ALPHA\_BETA\_T\_CELL\_ACTIVATION\_INVOLVED\_IN\_IMMUNE\_RESPONSE(GO:0002287) |  | 55 | 0.76 | 1.23 | 0.141 | 0.403 | 0.828 | 7414 | tags=20%, list=13%, signal=23% |
| 2949 | SPHINGOLIPID\_BINDING(GO:0046625) |  | 17 | 0.75 | 1.23 | 0.000 | 0.403 | 0.828 | 10643 | tags=35%, list=18%, signal=43% |
| 2950 | ANATOMICAL\_STRUCTURE\_MATURATION(GO:0071695) |  | 152 | 0.75 | 1.23 | 0.141 | 0.403 | 0.828 | 8892 | tags=26%, list=15%, signal=31% |
| 2951 | PHOTORECEPTOR\_ACTIVITY(GO:0009881) |  | 15 | 0.79 | 1.23 | 0.000 | 0.403 | 0.828 | 7933 | tags=27%, list=13%, signal=31% |
| 2952 | RECEPTOR\_CLUSTERING(GO:0043113) |  | 51 | 0.73 | 1.23 | 0.000 | 0.403 | 0.828 | 7963 | tags=18%, list=13%, signal=20% |
| 2953 | OVARIAN\_FOLLICLE\_DEVELOPMENT(GO:0001541) |  | 55 | 0.78 | 1.23 | 0.141 | 0.403 | 0.828 | 7547 | tags=13%, list=13%, signal=15% |
| 2954 | REGULATION\_OF\_T\_CELL\_APOPTOTIC\_PROCESS(GO:0070232) |  | 34 | 0.77 | 1.23 | 0.141 | 0.403 | 0.828 | 6685 | tags=12%, list=11%, signal=13% |
| 2955 | EMBRYONIC\_DIGESTIVE\_TRACT\_DEVELOPMENT(GO:0048566) |  | 35 | 0.84 | 1.23 | 0.141 | 0.402 | 0.828 | 7407 | tags=46%, list=13%, signal=52% |
| 2956 | MRNA\_STABILIZATION(GO:0048255) |  | 37 | 0.78 | 1.23 | 0.000 | 0.402 | 0.828 | 5906 | tags=3%, list=10%, signal=3% |
| 2957 | DNA\_DEALKYLATION(GO:0035510) |  | 29 | 0.81 | 1.23 | 0.141 | 0.402 | 0.828 | 7450 | tags=14%, list=13%, signal=16% |
| 2958 | DORSAL\_SPINAL\_CORD\_DEVELOPMENT(GO:0021516) |  | 22 | 0.83 | 1.23 | 0.141 | 0.402 | 0.828 | 8673 | tags=55%, list=15%, signal=64% |
| 2959 | REGULATION\_OF\_OSSIFICATION(GO:0030278) |  | 186 | 0.77 | 1.23 | 0.141 | 0.402 | 0.828 | 8046 | tags=24%, list=14%, signal=27% |
| 2960 | REGULATION\_OF\_OXIDOREDUCTASE\_ACTIVITY(GO:0051341) |  | 90 | 0.75 | 1.23 | 0.141 | 0.402 | 0.828 | 8755 | tags=17%, list=15%, signal=20% |
| 2961 | PHOSPHORYLATION(GO:0016310) |  | 2397 | 0.77 | 1.23 | 0.141 | 0.402 | 0.828 | 7973 | tags=13%, list=13%, signal=15% |
| 2962 | RESPONSE\_TO\_ABIOTIC\_STIMULUS(GO:0009628) |  | 1191 | 0.77 | 1.23 | 0.141 | 0.402 | 0.828 | 8241 | tags=14%, list=14%, signal=16% |
| 2963 | DNA\_DEMETHYLATION(GO:0080111) |  | 23 | 0.82 | 1.23 | 0.141 | 0.402 | 0.828 | 7450 | tags=17%, list=13%, signal=20% |
| 2964 | CYTOKINE\_METABOLIC\_PROCESS(GO:0042107) |  | 107 | 0.78 | 1.23 | 0.141 | 0.402 | 0.828 | 7461 | tags=20%, list=13%, signal=22% |
| 2965 | NEGATIVE\_REGULATION\_OF\_CHROMOSOME\_ORGANIZATION(GO:2001251) |  | 125 | 0.72 | 1.23 | 0.000 | 0.402 | 0.828 | 11042 | tags=22%, list=19%, signal=27% |
| 2966 | POTASSIUM\_ION\_TRANSMEMBRANE\_TRANSPORTER\_ACTIVITY(GO:0015079) |  | 168 | 0.76 | 1.23 | 0.141 | 0.402 | 0.828 | 9703 | tags=32%, list=16%, signal=38% |
| 2967 | NEUROMUSCULAR\_PROCESS\_CONTROLLING\_POSTURE(GO:0050884) |  | 16 | 0.77 | 1.23 | 0.000 | 0.402 | 0.828 | 11201 | tags=38%, list=19%, signal=46% |
| 2968 | REGULATION\_OF\_AMPA\_RECEPTOR\_ACTIVITY(GO:2000311) |  | 21 | 0.80 | 1.23 | 0.141 | 0.402 | 0.828 | 7962 | tags=38%, list=13%, signal=44% |
| 2969 | MONOVALENT\_INORGANIC\_CATION\_TRANSMEMBRANE\_TRANSPORTER\_ACTIVITY(GO:0015077) |  | 398 | 0.75 | 1.23 | 0.141 | 0.402 | 0.828 | 9554 | tags=28%, list=16%, signal=33% |
| 2970 | CGMP\_BIOSYNTHETIC\_PROCESS(GO:0006182) |  | 41 | 0.75 | 1.23 | 0.000 | 0.402 | 0.828 | 10245 | tags=34%, list=17%, signal=41% |
| 2971 | REGULATION\_OF\_OSTEOBLAST\_PROLIFERATION(GO:0033688) |  | 23 | 0.76 | 1.23 | 0.000 | 0.402 | 0.828 | 8946 | tags=22%, list=15%, signal=26% |
| 2972 | OXIDOREDUCTASE\_ACTIVITY\_\_ACTING\_ON\_THE\_ALDEHYDE\_OR\_OXO\_GROUP\_OF\_DONORS\_\_NAD\_OR\_NADP\_AS\_ACCEPTOR(GO:0016620) |  | 36 | 0.80 | 1.23 | 0.141 | 0.402 | 0.828 | 6438 | tags=14%, list=11%, signal=16% |
| 2973 | NUCLEIC\_ACID\_BINDING(GO:0003676) |  | 4300 | 0.78 | 1.23 | 0.141 | 0.402 | 0.828 | 7385 | tags=10%, list=12%, signal=10% |
| 2974 | SKELETAL\_MUSCLE\_CELL\_DIFFERENTIATION(GO:0035914) |  | 70 | 0.77 | 1.23 | 0.141 | 0.402 | 0.828 | 9208 | tags=30%, list=16%, signal=35% |
| 2975 | REGULATION\_OF\_BIOSYNTHETIC\_PROCESS(GO:0009889) |  | 4676 | 0.78 | 1.23 | 0.141 | 0.402 | 0.828 | 7517 | tags=12%, list=13%, signal=13% |
| 2976 | NICOTINAMIDE\_NUCLEOTIDE\_BIOSYNTHETIC\_PROCESS(GO:0019359) |  | 134 | 0.75 | 1.23 | 0.141 | 0.402 | 0.828 | 6696 | tags=5%, list=11%, signal=6% |
| 2977 | PYRIDINE\_NUCLEOTIDE\_BIOSYNTHETIC\_PROCESS(GO:0019363) |  | 134 | 0.75 | 1.23 | 0.141 | 0.402 | 0.828 | 6696 | tags=5%, list=11%, signal=6% |
| 2978 | REGULATION\_OF\_G\_PROTEIN\_COUPLED\_RECEPTOR\_PROTEIN\_SIGNALING\_PATHWAY(GO:0008277) |  | 142 | 0.73 | 1.23 | 0.000 | 0.402 | 0.828 | 10309 | tags=28%, list=17%, signal=34% |
| 2979 | AMMONIUM\_ION\_BINDING(GO:0070405) |  | 72 | 0.77 | 1.23 | 0.141 | 0.402 | 0.828 | 7830 | tags=28%, list=13%, signal=32% |
| 2980 | EMBRYO\_IMPLANTATION(GO:0007566) |  | 53 | 0.78 | 1.23 | 0.000 | 0.402 | 0.828 | 7727 | tags=25%, list=13%, signal=28% |
| 2981 | PLASMA\_MEMBRANE\_BOUNDED\_CELL\_PROJECTION\_ORGANIZATION(GO:0120036) |  | 1410 | 0.77 | 1.23 | 0.141 | 0.402 | 0.828 | 8953 | tags=22%, list=15%, signal=25% |
| 2982 | STEROID\_HORMONE\_RECEPTOR\_ACTIVITY(GO:0003707) |  | 59 | 0.78 | 1.23 | 0.141 | 0.402 | 0.828 | 9711 | tags=36%, list=16%, signal=43% |
| 2983 | BILE\_ACID\_TRANSMEMBRANE\_TRANSPORTER\_ACTIVITY(GO:0015125) |  | 20 | 0.79 | 1.23 | 0.141 | 0.402 | 0.828 | 9376 | tags=30%, list=16%, signal=36% |
| 2984 | CYTOKINE\_BIOSYNTHETIC\_PROCESS(GO:0042089) |  | 106 | 0.78 | 1.23 | 0.141 | 0.402 | 0.828 | 7461 | tags=20%, list=13%, signal=23% |
| 2985 | ENDOPEPTIDASE\_ACTIVITY(GO:0004175) |  | 486 | 0.75 | 1.23 | 0.141 | 0.402 | 0.828 | 7916 | tags=13%, list=13%, signal=15% |
| 2986 | CELLULAR\_RESPONSE\_TO\_STRESS(GO:0033554) |  | 1933 | 0.77 | 1.23 | 0.141 | 0.402 | 0.828 | 7640 | tags=8%, list=13%, signal=9% |
| 2987 | SYNAPTIC\_VESICLE\_LOCALIZATION(GO:0097479) |  | 117 | 0.76 | 1.23 | 0.141 | 0.402 | 0.828 | 9227 | tags=25%, list=16%, signal=29% |
| 2988 | DENDRITE\_CYTOPLASM(GO:0032839) |  | 25 | 0.74 | 1.23 | 0.000 | 0.402 | 0.828 | 8210 | tags=24%, list=14%, signal=28% |
| 2989 | CELL\_PROJECTION\_ORGANIZATION(GO:0030030) |  | 1438 | 0.77 | 1.23 | 0.141 | 0.402 | 0.828 | 8953 | tags=22%, list=15%, signal=25% |
| 2990 | POSITIVE\_REGULATION\_OF\_DNA\_TEMPLATED\_TRANSCRIPTION\_\_INITIATION(GO:2000144) |  | 29 | 0.80 | 1.23 | 0.141 | 0.402 | 0.828 | 6114 | tags=14%, list=10%, signal=15% |
| 2991 | RESPONSE\_TO\_CAFFEINE(GO:0031000) |  | 15 | 0.76 | 1.23 | 0.000 | 0.402 | 0.828 | 9628 | tags=47%, list=16%, signal=56% |
| 2992 | RESPONSE\_TO\_DIURETIC(GO:0036270) |  | 15 | 0.76 | 1.23 | 0.000 | 0.402 | 0.828 | 9628 | tags=47%, list=16%, signal=56% |
| 2993 | POSITIVE\_REGULATION\_OF\_RESPONSE\_TO\_DNA\_DAMAGE\_STIMULUS(GO:2001022) |  | 78 | 0.79 | 1.23 | 0.141 | 0.402 | 0.828 | 7101 | tags=8%, list=12%, signal=9% |
| 2994 | VOLTAGE\_GATED\_POTASSIUM\_CHANNEL\_ACTIVITY(GO:0005249) |  | 97 | 0.75 | 1.23 | 0.141 | 0.402 | 0.828 | 9703 | tags=36%, list=16%, signal=43% |
| 2995 | ANDROGEN\_RECEPTOR\_BINDING(GO:0050681) |  | 43 | 0.74 | 1.23 | 0.000 | 0.401 | 0.828 | 5918 | tags=5%, list=10%, signal=5% |
| 2996 | REGULATION\_OF\_PEPTIDE\_TRANSPORT(GO:0090087) |  | 718 | 0.76 | 1.23 | 0.141 | 0.401 | 0.828 | 7652 | tags=13%, list=13%, signal=15% |
| 2997 | REGULATION\_OF\_BLOOD\_VESSEL\_ENDOTHELIAL\_CELL\_MIGRATION(GO:0043535) |  | 147 | 0.76 | 1.23 | 0.141 | 0.401 | 0.828 | 10901 | tags=23%, list=18%, signal=28% |
| 2998 | POSITIVE\_REGULATION\_OF\_VOLTAGE\_GATED\_POTASSIUM\_CHANNEL\_ACTIVITY(GO:1903818) |  | 15 | 0.81 | 1.23 | 0.141 | 0.401 | 0.828 | 9543 | tags=40%, list=16%, signal=48% |
| 2999 | TRANSCRIPTION\_FACTOR\_COMPLEX(GO:0005667) |  | 323 | 0.73 | 1.23 | 0.000 | 0.401 | 0.828 | 8380 | tags=17%, list=14%, signal=20% |
| 3000 | AUTOPHAGOSOME\_MEMBRANE(GO:0000421) |  | 31 | 0.73 | 1.23 | 0.000 | 0.401 | 0.828 | 9096 | tags=23%, list=15%, signal=27% |
| 3001 | NAD\_BINDING(GO:0051287) |  | 57 | 0.75 | 1.23 | 0.000 | 0.401 | 0.828 | 6975 | tags=7%, list=12%, signal=8% |
| 3002 | NEGATIVE\_REGULATION\_OF\_PROTEIN\_TRANSPORT(GO:0051224) |  | 164 | 0.73 | 1.23 | 0.141 | 0.401 | 0.828 | 9686 | tags=20%, list=16%, signal=23% |
| 3003 | REGULATION\_OF\_SYSTEMIC\_ARTERIAL\_BLOOD\_PRESSURE(GO:0003073) |  | 92 | 0.77 | 1.23 | 0.141 | 0.401 | 0.828 | 8969 | tags=27%, list=15%, signal=32% |
| 3004 | MACROPHAGE\_DERIVED\_FOAM\_CELL\_DIFFERENTIATION(GO:0010742) |  | 35 | 0.74 | 1.23 | 0.000 | 0.402 | 0.828 | 8474 | tags=20%, list=14%, signal=23% |
| 3005 | FOAM\_CELL\_DIFFERENTIATION(GO:0090077) |  | 35 | 0.74 | 1.23 | 0.000 | 0.401 | 0.828 | 8474 | tags=20%, list=14%, signal=23% |
| 3006 | NEGATIVE\_REGULATION\_OF\_LYMPHOCYTE\_DIFFERENTIATION(GO:0045620) |  | 46 | 0.75 | 1.23 | 0.141 | 0.401 | 0.828 | 6178 | tags=11%, list=10%, signal=12% |
| 3007 | SNARE\_BINDING(GO:0000149) |  | 119 | 0.75 | 1.23 | 0.141 | 0.401 | 0.828 | 7426 | tags=13%, list=13%, signal=14% |
| 3008 | TRANSCRIPTION\_COREPRESSOR\_ACTIVITY(GO:0003714) |  | 223 | 0.73 | 1.23 | 0.000 | 0.401 | 0.828 | 7401 | tags=12%, list=13%, signal=13% |
| 3009 | INTEGRIN\_COMPLEX(GO:0008305) |  | 33 | 0.76 | 1.23 | 0.141 | 0.401 | 0.828 | 8865 | tags=27%, list=15%, signal=32% |
| 3010 | REGULATION\_OF\_CELL\_GROWTH(GO:0001558) |  | 385 | 0.77 | 1.23 | 0.141 | 0.401 | 0.828 | 7975 | tags=16%, list=13%, signal=18% |
| 3011 | PROTEIN\_LOCALIZATION\_TO\_MEMBRANE(GO:0072657) |  | 541 | 0.76 | 1.23 | 0.141 | 0.401 | 0.828 | 7963 | tags=10%, list=13%, signal=11% |
| 3012 | CARDIAC\_VENTRICLE\_DEVELOPMENT(GO:0003231) |  | 121 | 0.77 | 1.23 | 0.141 | 0.401 | 0.828 | 7913 | tags=21%, list=13%, signal=25% |
| 3013 | RESPONSE\_TO\_STARVATION(GO:0042594) |  | 174 | 0.74 | 1.23 | 0.141 | 0.402 | 0.828 | 10519 | tags=21%, list=18%, signal=26% |
| 3014 | SEMAPHORIN\_PLEXIN\_SIGNALING\_PATHWAY(GO:0071526) |  | 26 | 0.79 | 1.23 | 0.141 | 0.401 | 0.828 | 9233 | tags=23%, list=16%, signal=27% |
| 3015 | ACYLGLYCEROL\_CATABOLIC\_PROCESS(GO:0046464) |  | 39 | 0.75 | 1.23 | 0.000 | 0.401 | 0.828 | 11563 | tags=36%, list=20%, signal=45% |
| 3016 | NEUTRAL\_LIPID\_CATABOLIC\_PROCESS(GO:0046461) |  | 39 | 0.75 | 1.23 | 0.000 | 0.401 | 0.828 | 11563 | tags=36%, list=20%, signal=45% |
| 3017 | REGULATION\_OF\_CHROMATIN\_ORGANIZATION(GO:1902275) |  | 162 | 0.72 | 1.23 | 0.000 | 0.401 | 0.828 | 10543 | tags=19%, list=18%, signal=22% |
| 3018 | PURINERGIC\_RECEPTOR\_ACTIVITY(GO:0035586) |  | 27 | 0.80 | 1.23 | 0.000 | 0.401 | 0.828 | 5955 | tags=19%, list=10%, signal=21% |
| 3019 | OXIDOREDUCTASE\_ACTIVITY\_\_ACTING\_ON\_PAIRED\_DONORS\_\_WITH\_INCORPORATION\_OR\_REDUCTION\_OF\_MOLECULAR\_OXYGEN(GO:0016705) |  | 155 | 0.76 | 1.23 | 0.141 | 0.401 | 0.828 | 11264 | tags=34%, list=19%, signal=42% |
| 3020 | RNA\_POLYMERASE\_II\_TRANSCRIPTION\_FACTOR\_COMPLEX(GO:0090575) |  | 139 | 0.74 | 1.23 | 0.000 | 0.401 | 0.828 | 7257 | tags=9%, list=12%, signal=10% |
| 3021 | RNA\_STABILIZATION(GO:0043489) |  | 41 | 0.76 | 1.23 | 0.000 | 0.401 | 0.828 | 5906 | tags=2%, list=10%, signal=3% |
| 3022 | NUCLEOTIDYLTRANSFERASE\_ACTIVITY(GO:0016779) |  | 140 | 0.75 | 1.23 | 0.141 | 0.401 | 0.828 | 6761 | tags=5%, list=11%, signal=6% |
| 3023 | POSITIVE\_REGULATION\_OF\_DNA\_REPAIR(GO:0045739) |  | 48 | 0.79 | 1.23 | 0.141 | 0.401 | 0.828 | 6957 | tags=8%, list=12%, signal=9% |
| 3024 | MITOCHONDRIAL\_RESPIRATORY\_CHAIN(GO:0005746) |  | 95 | 0.71 | 1.23 | 0.000 | 0.401 | 0.828 | 5894 | tags=5%, list=10%, signal=6% |
| 3025 | PEPTIDE\_HORMONE\_PROCESSING(GO:0016486) |  | 37 | 0.73 | 1.23 | 0.000 | 0.401 | 0.828 | 8969 | tags=35%, list=15%, signal=41% |
| 3026 | POSITIVE\_REGULATION\_OF\_CARDIAC\_MUSCLE\_TISSUE\_DEVELOPMENT(GO:0055025) |  | 57 | 0.77 | 1.23 | 0.141 | 0.401 | 0.828 | 8774 | tags=23%, list=15%, signal=27% |
| 3027 | GLOMERULAR\_VISCERAL\_EPITHELIAL\_CELL\_DIFFERENTIATION(GO:0072112) |  | 17 | 0.76 | 1.23 | 0.141 | 0.401 | 0.828 | 8937 | tags=29%, list=15%, signal=35% |
| 3028 | RENAL\_FILTRATION\_CELL\_DIFFERENTIATION(GO:0061318) |  | 17 | 0.76 | 1.23 | 0.141 | 0.401 | 0.828 | 8937 | tags=29%, list=15%, signal=35% |
| 3029 | REGULATION\_OF\_ENDOTHELIAL\_CELL\_APOPTOTIC\_PROCESS(GO:2000351) |  | 50 | 0.77 | 1.23 | 0.141 | 0.401 | 0.828 | 8627 | tags=18%, list=15%, signal=21% |
| 3030 | CHAPERONE\_BINDING(GO:0051087) |  | 93 | 0.71 | 1.23 | 0.000 | 0.401 | 0.828 | 12107 | tags=28%, list=20%, signal=35% |
| 3031 | ORGANELLE\_FUSION(GO:0048284) |  | 190 | 0.74 | 1.23 | 0.141 | 0.401 | 0.828 | 9144 | tags=14%, list=15%, signal=17% |
| 3032 | REGULATION\_OF\_CELLULAR\_CARBOHYDRATE\_METABOLIC\_PROCESS(GO:0010675) |  | 120 | 0.76 | 1.23 | 0.141 | 0.401 | 0.828 | 7821 | tags=13%, list=13%, signal=14% |
| 3033 | POSITIVE\_REGULATION\_OF\_PHOSPHORYLATION(GO:0042327) |  | 997 | 0.76 | 1.23 | 0.141 | 0.401 | 0.828 | 7754 | tags=15%, list=13%, signal=17% |
| 3034 | SUBPALLIUM\_DEVELOPMENT(GO:0021544) |  | 23 | 0.85 | 1.23 | 0.000 | 0.401 | 0.828 | 7668 | tags=30%, list=13%, signal=35% |
| 3035 | CHLORIDE\_CHANNEL\_COMPLEX(GO:0034707) |  | 49 | 0.77 | 1.23 | 0.141 | 0.401 | 0.828 | 8596 | tags=27%, list=15%, signal=31% |
| 3036 | INTERFERON\_GAMMA\_MEDIATED\_SIGNALING\_PATHWAY(GO:0060333) |  | 84 | 0.76 | 1.23 | 0.141 | 0.401 | 0.828 | 8063 | tags=12%, list=14%, signal=14% |
| 3037 | CHROMATIN\_BINDING(GO:0003682) |  | 513 | 0.76 | 1.23 | 0.141 | 0.401 | 0.828 | 7583 | tags=12%, list=13%, signal=13% |
| 3038 | MULTICELLULAR\_ORGANISM\_REPRODUCTION(GO:0032504) |  | 820 | 0.75 | 1.23 | 0.141 | 0.400 | 0.828 | 8690 | tags=20%, list=15%, signal=23% |
| 3039 | TRIGLYCERIDE\_LIPASE\_ACTIVITY(GO:0004806) |  | 20 | 0.76 | 1.23 | 0.000 | 0.400 | 0.828 | 11563 | tags=35%, list=20%, signal=43% |
| 3040 | REGULATION\_OF\_RESPIRATORY\_SYSTEM\_PROCESS(GO:0044065) |  | 16 | 0.74 | 1.23 | 0.000 | 0.400 | 0.828 | 11201 | tags=44%, list=19%, signal=54% |
| 3041 | OMEGA\_PEPTIDASE\_ACTIVITY(GO:0008242) |  | 15 | 0.78 | 1.23 | 0.000 | 0.400 | 0.828 | 9578 | tags=20%, list=16%, signal=24% |
| 3042 | NEGATIVE\_REGULATION\_OF\_ION\_TRANSMEMBRANE\_TRANSPORTER\_ACTIVITY(GO:0032413) |  | 75 | 0.73 | 1.23 | 0.141 | 0.400 | 0.828 | 9472 | tags=25%, list=16%, signal=30% |
| 3043 | PROTEIN\_FOLDING(GO:0006457) |  | 229 | 0.71 | 1.23 | 0.000 | 0.400 | 0.828 | 11101 | tags=23%, list=19%, signal=28% |
| 3044 | REGULATION\_OF\_B\_CELL\_PROLIFERATION(GO:0030888) |  | 60 | 0.74 | 1.23 | 0.141 | 0.400 | 0.828 | 10693 | tags=37%, list=18%, signal=45% |
| 3045 | MICROTUBULE\_BASED\_PROCESS(GO:0007017) |  | 728 | 0.72 | 1.23 | 0.000 | 0.400 | 0.828 | 10064 | tags=24%, list=17%, signal=29% |
| 3046 | ENDOPEPTIDASE\_INHIBITOR\_ACTIVITY(GO:0004866) |  | 171 | 0.75 | 1.23 | 0.141 | 0.400 | 0.828 | 8070 | tags=19%, list=14%, signal=22% |
| 3047 | REGULATION\_OF\_ENDOCRINE\_PROCESS(GO:0044060) |  | 45 | 0.74 | 1.23 | 0.141 | 0.400 | 0.828 | 8490 | tags=24%, list=14%, signal=29% |
| 3048 | CARBOHYDRATE\_DERIVATIVE\_METABOLIC\_PROCESS(GO:1901135) |  | 1264 | 0.74 | 1.23 | 0.141 | 0.400 | 0.828 | 8696 | tags=15%, list=15%, signal=17% |
| 3049 | CYTOPLASMIC\_SIDE\_OF\_PLASMA\_MEMBRANE(GO:0009898) |  | 180 | 0.76 | 1.23 | 0.141 | 0.400 | 0.828 | 7951 | tags=11%, list=13%, signal=12% |
| 3050 | CORECEPTOR\_ACTIVITY(GO:0015026) |  | 41 | 0.79 | 1.23 | 0.141 | 0.400 | 0.828 | 7975 | tags=32%, list=13%, signal=37% |
| 3051 | REGULATION\_OF\_PROTEIN\_TARGETING\_TO\_MITOCHONDRION(GO:1903214) |  | 92 | 0.75 | 1.23 | 0.141 | 0.400 | 0.828 | 7426 | tags=10%, list=13%, signal=11% |
| 3052 | ATP\_GENERATION\_FROM\_ADP(GO:0006757) |  | 103 | 0.75 | 1.23 | 0.141 | 0.400 | 0.828 | 6696 | tags=5%, list=11%, signal=5% |
| 3053 | METALLOPEPTIDASE\_ACTIVITY(GO:0008237) |  | 188 | 0.72 | 1.23 | 0.000 | 0.400 | 0.828 | 11757 | tags=30%, list=20%, signal=37% |
| 3054 | REGULATION\_OF\_CELL\_ACTIVATION(GO:0050865) |  | 555 | 0.75 | 1.23 | 0.141 | 0.400 | 0.828 | 9198 | tags=19%, list=16%, signal=22% |
| 3055 | REGULATION\_OF\_T\_HELPER\_CELL\_DIFFERENTIATION(GO:0045622) |  | 30 | 0.75 | 1.23 | 0.141 | 0.400 | 0.828 | 9411 | tags=30%, list=16%, signal=36% |
| 3056 | NEUROPEPTIDE\_BINDING(GO:0042923) |  | 23 | 0.81 | 1.22 | 0.141 | 0.400 | 0.828 | 9434 | tags=39%, list=16%, signal=47% |
| 3057 | POSITIVE\_REGULATION\_OF\_DENDRITIC\_SPINE\_MORPHOGENESIS(GO:0061003) |  | 17 | 0.74 | 1.22 | 0.000 | 0.400 | 0.828 | 8450 | tags=24%, list=14%, signal=27% |
| 3058 | CYTOSOL(GO:0005829) |  | 4956 | 0.80 | 1.22 | 0.141 | 0.400 | 0.828 | 7175 | tags=7%, list=12%, signal=7% |
| 3059 | GLYCEROLIPID\_CATABOLIC\_PROCESS(GO:0046503) |  | 56 | 0.74 | 1.22 | 0.000 | 0.400 | 0.828 | 11717 | tags=32%, list=20%, signal=40% |
| 3060 | POSITIVE\_REGULATION\_OF\_BLOOD\_VESSEL\_ENDOTHELIAL\_CELL\_PROLIFERATION\_INVOLVED\_IN\_SPROUTING\_ANGIOGENESIS(GO:1903589) |  | 17 | 0.84 | 1.22 | 0.141 | 0.400 | 0.828 | 9016 | tags=29%, list=15%, signal=35% |
| 3061 | NEGATIVE\_REGULATION\_OF\_STEM\_CELL\_DIFFERENTIATION(GO:2000737) |  | 20 | 0.75 | 1.22 | 0.000 | 0.400 | 0.828 | 8673 | tags=20%, list=15%, signal=23% |
| 3062 | REGULATION\_OF\_TRANSCRIPTION\_\_DNA\_TEMPLATED(GO:0006355) |  | 3751 | 0.77 | 1.22 | 0.141 | 0.400 | 0.828 | 7927 | tags=14%, list=13%, signal=15% |
| 3063 | GLYCOLYTIC\_PROCESS(GO:0006096) |  | 102 | 0.75 | 1.22 | 0.141 | 0.400 | 0.828 | 6696 | tags=5%, list=11%, signal=6% |
| 3064 | REGULATION\_OF\_NUCLEIC\_ACID\_TEMPLATED\_TRANSCRIPTION(GO:1903506) |  | 3772 | 0.77 | 1.22 | 0.141 | 0.400 | 0.828 | 7927 | tags=14%, list=13%, signal=15% |
| 3065 | REGULATION\_OF\_RNA\_BIOSYNTHETIC\_PROCESS(GO:2001141) |  | 3780 | 0.77 | 1.22 | 0.141 | 0.400 | 0.828 | 7927 | tags=14%, list=13%, signal=15% |
| 3066 | PROTEIN\_LOCALIZATION\_TO\_CILIUM(GO:0061512) |  | 38 | 0.74 | 1.22 | 0.000 | 0.399 | 0.828 | 8306 | tags=29%, list=14%, signal=34% |
| 3067 | TUBE\_FORMATION(GO:0035148) |  | 149 | 0.77 | 1.22 | 0.141 | 0.399 | 0.828 | 7975 | tags=16%, list=13%, signal=19% |
| 3068 | CARDIAC\_MYOFIBRIL\_ASSEMBLY(GO:0055003) |  | 17 | 0.75 | 1.22 | 0.000 | 0.399 | 0.828 | 10033 | tags=53%, list=17%, signal=64% |
| 3069 | NEGATIVE\_REGULATION\_OF\_ESTABLISHMENT\_OF\_PROTEIN\_LOCALIZATION(GO:1904950) |  | 168 | 0.73 | 1.22 | 0.141 | 0.399 | 0.828 | 9686 | tags=20%, list=16%, signal=23% |
| 3070 | PHOSPHATE\_CONTAINING\_COMPOUND\_METABOLIC\_PROCESS(GO:0006796) |  | 3294 | 0.77 | 1.22 | 0.141 | 0.399 | 0.828 | 7973 | tags=13%, list=13%, signal=14% |
| 3071 | MULTICELLULAR\_ORGANISMAL\_REPRODUCTIVE\_PROCESS(GO:0048609) |  | 807 | 0.75 | 1.22 | 0.141 | 0.399 | 0.828 | 8690 | tags=20%, list=15%, signal=23% |
| 3072 | POSITIVE\_REGULATION\_OF\_CYSTEINE\_TYPE\_ENDOPEPTIDASE\_ACTIVITY(GO:2001056) |  | 136 | 0.73 | 1.22 | 0.141 | 0.399 | 0.828 | 7754 | tags=11%, list=13%, signal=13% |
| 3073 | SUBSTRATE\_ADHESION\_DEPENDENT\_CELL\_SPREADING(GO:0034446) |  | 84 | 0.76 | 1.22 | 0.141 | 0.399 | 0.828 | 8726 | tags=20%, list=15%, signal=24% |
| 3074 | OVULATION\_CYCLE(GO:0042698) |  | 67 | 0.76 | 1.22 | 0.141 | 0.399 | 0.828 | 7356 | tags=25%, list=12%, signal=29% |
| 3075 | PHOSPHOLIPID\_METABOLIC\_PROCESS(GO:0006644) |  | 470 | 0.74 | 1.22 | 0.141 | 0.399 | 0.828 | 9096 | tags=18%, list=15%, signal=21% |
| 3076 | HEPARAN\_SULFATE\_PROTEOGLYCAN\_BINDING(GO:0043395) |  | 19 | 0.81 | 1.22 | 0.141 | 0.399 | 0.828 | 6968 | tags=32%, list=12%, signal=36% |
| 3077 | REGULATION\_OF\_FAT\_CELL\_DIFFERENTIATION(GO:0045598) |  | 120 | 0.74 | 1.22 | 0.141 | 0.399 | 0.828 | 10811 | tags=26%, list=18%, signal=32% |
| 3078 | EATING\_BEHAVIOR(GO:0042755) |  | 33 | 0.81 | 1.22 | 0.141 | 0.399 | 0.828 | 6240 | tags=12%, list=11%, signal=14% |
| 3079 | ENDOPEPTIDASE\_REGULATOR\_ACTIVITY(GO:0061135) |  | 177 | 0.75 | 1.22 | 0.141 | 0.399 | 0.828 | 8070 | tags=18%, list=14%, signal=21% |
| 3080 | GLOMERULAR\_EPITHELIAL\_CELL\_DIFFERENTIATION(GO:0072311) |  | 18 | 0.75 | 1.22 | 0.141 | 0.399 | 0.828 | 8937 | tags=28%, list=15%, signal=33% |
| 3081 | CALMODULIN\_DEPENDENT\_PROTEIN\_KINASE\_ACTIVITY(GO:0004683) |  | 28 | 0.76 | 1.22 | 0.000 | 0.399 | 0.828 | 6212 | tags=14%, list=11%, signal=16% |
| 3082 | REGULATION\_OF\_SYNAPTIC\_PLASTICITY(GO:0048167) |  | 162 | 0.75 | 1.22 | 0.141 | 0.399 | 0.828 | 9453 | tags=23%, list=16%, signal=28% |
| 3083 | ENERGY\_HOMEOSTASIS(GO:0097009) |  | 18 | 0.79 | 1.22 | 0.141 | 0.399 | 0.828 | 10112 | tags=39%, list=17%, signal=47% |
| 3084 | PROTEIN\_COMPLEX\_INVOLVED\_IN\_CELL\_ADHESION(GO:0098636) |  | 36 | 0.76 | 1.22 | 0.141 | 0.399 | 0.828 | 8865 | tags=28%, list=15%, signal=33% |
| 3085 | T\_CELL\_DIFFERENTIATION\_INVOLVED\_IN\_IMMUNE\_RESPONSE(GO:0002292) |  | 61 | 0.75 | 1.22 | 0.141 | 0.399 | 0.828 | 7414 | tags=18%, list=13%, signal=21% |
| 3086 | CELL\_GROWTH(GO:0016049) |  | 475 | 0.76 | 1.22 | 0.141 | 0.399 | 0.828 | 8820 | tags=18%, list=15%, signal=21% |
| 3087 | REGULATION\_OF\_TRANSMEMBRANE\_RECEPTOR\_PROTEIN\_SERINE\_THREONINE\_KINASE\_SIGNALING\_PATHWAY(GO:0090092) |  | 225 | 0.75 | 1.22 | 0.141 | 0.399 | 0.828 | 8177 | tags=21%, list=14%, signal=25% |
| 3088 | POSITIVE\_REGULATION\_OF\_PROTEIN\_SECRETION(GO:0050714) |  | 240 | 0.74 | 1.22 | 0.141 | 0.399 | 0.828 | 7898 | tags=16%, list=13%, signal=18% |
| 3089 | AUTONOMIC\_NERVOUS\_SYSTEM\_DEVELOPMENT(GO:0048483) |  | 42 | 0.80 | 1.22 | 0.141 | 0.399 | 0.828 | 9208 | tags=31%, list=16%, signal=37% |
| 3090 | NITRIC\_OXIDE\_BIOSYNTHETIC\_PROCESS(GO:0006809) |  | 65 | 0.75 | 1.22 | 0.141 | 0.399 | 0.828 | 7873 | tags=20%, list=13%, signal=23% |
| 3091 | PEPTIDE\_HORMONE\_SECRETION(GO:0030072) |  | 247 | 0.73 | 1.22 | 0.141 | 0.399 | 0.828 | 10131 | tags=23%, list=17%, signal=28% |
| 3092 | LYSOSOMAL\_LUMEN(GO:0043202) |  | 94 | 0.74 | 1.22 | 0.000 | 0.399 | 0.828 | 7148 | tags=13%, list=12%, signal=14% |
| 3093 | AROMATIC\_COMPOUND\_CATABOLIC\_PROCESS(GO:0019439) |  | 680 | 0.75 | 1.22 | 0.141 | 0.399 | 0.828 | 6696 | tags=6%, list=11%, signal=7% |
| 3094 | GLYCOSPHINGOLIPID\_METABOLIC\_PROCESS(GO:0006687) |  | 73 | 0.74 | 1.22 | 0.141 | 0.399 | 0.828 | 8411 | tags=19%, list=14%, signal=22% |
| 3095 | HORMONE\_SECRETION(GO:0046879) |  | 311 | 0.73 | 1.22 | 0.141 | 0.399 | 0.828 | 10131 | tags=24%, list=17%, signal=29% |
| 3096 | IRON\_ION\_HOMEOSTASIS(GO:0055072) |  | 86 | 0.72 | 1.22 | 0.000 | 0.399 | 0.828 | 9361 | tags=17%, list=16%, signal=21% |
| 3097 | REGULATION\_OF\_SECRETION(GO:0051046) |  | 732 | 0.74 | 1.22 | 0.141 | 0.398 | 0.828 | 10131 | tags=23%, list=17%, signal=27% |
| 3098 | ENAMEL\_MINERALIZATION(GO:0070166) |  | 16 | 0.77 | 1.22 | 0.000 | 0.399 | 0.828 | 9405 | tags=25%, list=16%, signal=30% |
| 3099 | VENTRICULAR\_CARDIAC\_MUSCLE\_TISSUE\_DEVELOPMENT(GO:0003229) |  | 54 | 0.74 | 1.22 | 0.000 | 0.399 | 0.828 | 12990 | tags=46%, list=22%, signal=59% |
| 3100 | PHOSPHATIDYLCHOLINE\_BINDING(GO:0031210) |  | 23 | 0.73 | 1.22 | 0.000 | 0.399 | 0.828 | 6362 | tags=26%, list=11%, signal=29% |
| 3101 | QUATERNARY\_AMMONIUM\_GROUP\_BINDING(GO:0050997) |  | 23 | 0.73 | 1.22 | 0.000 | 0.398 | 0.828 | 6362 | tags=26%, list=11%, signal=29% |
| 3102 | PURINERGIC\_NUCLEOTIDE\_RECEPTOR\_ACTIVITY(GO:0001614) |  | 23 | 0.84 | 1.22 | 0.000 | 0.398 | 0.828 | 5955 | tags=22%, list=10%, signal=24% |
| 3103 | NUCLEOTIDE\_RECEPTOR\_ACTIVITY(GO:0016502) |  | 23 | 0.84 | 1.22 | 0.000 | 0.398 | 0.828 | 5955 | tags=22%, list=10%, signal=24% |
| 3104 | REGULATION\_OF\_NEUROTRANSMITTER\_SECRETION(GO:0046928) |  | 55 | 0.77 | 1.22 | 0.141 | 0.398 | 0.828 | 8040 | tags=25%, list=14%, signal=29% |
| 3105 | REGULATION\_OF\_ESTABLISHMENT\_OF\_PROTEIN\_LOCALIZATION(GO:0070201) |  | 730 | 0.75 | 1.22 | 0.141 | 0.398 | 0.828 | 7652 | tags=12%, list=13%, signal=14% |
| 3106 | NUCLEAR\_OUTER\_MEMBRANE\_ENDOPLASMIC\_RETICULUM\_MEMBRANE\_NETWORK(GO:0042175) |  | 1046 | 0.73 | 1.22 | 0.141 | 0.398 | 0.828 | 8013 | tags=11%, list=14%, signal=13% |
| 3107 | PHOSPHORUS\_METABOLIC\_PROCESS(GO:0006793) |  | 3383 | 0.76 | 1.22 | 0.141 | 0.398 | 0.828 | 7973 | tags=13%, list=13%, signal=14% |
| 3108 | PROTEIN\_DNA\_COMPLEX(GO:0032993) |  | 202 | 0.75 | 1.22 | 0.141 | 0.398 | 0.828 | 6564 | tags=7%, list=11%, signal=8% |
| 3109 | CD4\_POSITIVE\_\_ALPHA\_BETA\_T\_CELL\_CYTOKINE\_PRODUCTION(GO:0035743) |  | 18 | 0.78 | 1.22 | 0.141 | 0.398 | 0.828 | 7435 | tags=44%, list=13%, signal=51% |
| 3110 | MELANIN\_METABOLIC\_PROCESS(GO:0006582) |  | 18 | 0.80 | 1.22 | 0.141 | 0.398 | 0.828 | 8714 | tags=50%, list=15%, signal=59% |
| 3111 | DOUBLE\_STRAND\_BREAK\_REPAIR\_VIA\_HOMOLOGOUS\_RECOMBINATION(GO:0000724) |  | 111 | 0.71 | 1.22 | 0.000 | 0.398 | 0.828 | 13120 | tags=43%, list=22%, signal=55% |
| 3112 | DRUG\_TRANSMEMBRANE\_TRANSPORTER\_ACTIVITY(GO:0015238) |  | 95 | 0.78 | 1.22 | 0.141 | 0.398 | 0.828 | 8066 | tags=25%, list=14%, signal=29% |
| 3113 | HORMONE\_TRANSPORT(GO:0009914) |  | 323 | 0.73 | 1.22 | 0.141 | 0.398 | 0.828 | 10131 | tags=24%, list=17%, signal=29% |
| 3114 | DRUG\_TRANSMEMBRANE\_TRANSPORT(GO:0006855) |  | 76 | 0.76 | 1.22 | 0.141 | 0.398 | 0.828 | 8066 | tags=28%, list=14%, signal=32% |
| 3115 | GTP\_BINDING(GO:0005525) |  | 384 | 0.76 | 1.22 | 0.141 | 0.398 | 0.828 | 7575 | tags=12%, list=13%, signal=13% |
| 3116 | REGULATION\_OF\_ANDROGEN\_RECEPTOR\_SIGNALING\_PATHWAY(GO:0060765) |  | 26 | 0.71 | 1.22 | 0.000 | 0.398 | 0.828 | 12594 | tags=38%, list=21%, signal=49% |
| 3117 | INORGANIC\_ANION\_EXCHANGER\_ACTIVITY(GO:0005452) |  | 23 | 0.74 | 1.22 | 0.000 | 0.398 | 0.828 | 11359 | tags=43%, list=19%, signal=54% |
| 3118 | HEART\_TRABECULA\_MORPHOGENESIS(GO:0061384) |  | 35 | 0.72 | 1.22 | 0.000 | 0.398 | 0.828 | 12860 | tags=49%, list=22%, signal=62% |
| 3119 | REGULATION\_OF\_CYTOSKELETON\_ORGANIZATION(GO:0051493) |  | 476 | 0.75 | 1.22 | 0.141 | 0.398 | 0.828 | 7973 | tags=13%, list=13%, signal=15% |
| 3120 | OSTEOBLAST\_PROLIFERATION(GO:0033687) |  | 26 | 0.74 | 1.22 | 0.141 | 0.398 | 0.828 | 8946 | tags=19%, list=15%, signal=23% |
| 3121 | ORGANIC\_CYCLIC\_COMPOUND\_CATABOLIC\_PROCESS(GO:1901361) |  | 711 | 0.75 | 1.22 | 0.141 | 0.398 | 0.828 | 8602 | tags=11%, list=15%, signal=12% |
| 3122 | ENDOMEMBRANE\_SYSTEM\_ORGANIZATION(GO:0010256) |  | 397 | 0.74 | 1.22 | 0.141 | 0.398 | 0.828 | 8541 | tags=12%, list=14%, signal=13% |
| 3123 | WHOLE\_MEMBRANE(GO:0098805) |  | 1595 | 0.76 | 1.22 | 0.141 | 0.398 | 0.828 | 7492 | tags=12%, list=13%, signal=13% |
| 3124 | PURINE\_NUCLEOSIDE\_BIOSYNTHETIC\_PROCESS(GO:0042451) |  | 24 | 0.70 | 1.22 | 0.000 | 0.398 | 0.828 | 13707 | tags=46%, list=23%, signal=60% |
| 3125 | PURINE\_RIBONUCLEOSIDE\_BIOSYNTHETIC\_PROCESS(GO:0046129) |  | 24 | 0.70 | 1.22 | 0.000 | 0.398 | 0.828 | 13707 | tags=46%, list=23%, signal=60% |
| 3126 | MIDBRAIN\_DOPAMINERGIC\_NEURON\_DIFFERENTIATION(GO:1904948) |  | 16 | 0.82 | 1.22 | 0.141 | 0.398 | 0.828 | 8673 | tags=31%, list=15%, signal=37% |
| 3127 | LYMPHOCYTE\_DIFFERENTIATION(GO:0030098) |  | 329 | 0.74 | 1.22 | 0.141 | 0.398 | 0.828 | 7547 | tags=18%, list=13%, signal=20% |
| 3128 | POSITIVE\_REGULATION\_OF\_ENDOTHELIAL\_CELL\_PROLIFERATION(GO:0001938) |  | 101 | 0.77 | 1.22 | 0.141 | 0.398 | 0.828 | 9255 | tags=25%, list=16%, signal=29% |
| 3129 | REGULATION\_OF\_CELL\_ADHESION(GO:0030155) |  | 633 | 0.75 | 1.22 | 0.141 | 0.398 | 0.828 | 8112 | tags=16%, list=14%, signal=19% |
| 3130 | POSITIVE\_REGULATION\_OF\_ALPHA\_BETA\_T\_CELL\_PROLIFERATION(GO:0046641) |  | 17 | 0.77 | 1.22 | 0.000 | 0.398 | 0.828 | 9023 | tags=29%, list=15%, signal=35% |
| 3131 | PROTEOGLYCAN\_BIOSYNTHETIC\_PROCESS(GO:0030166) |  | 58 | 0.72 | 1.22 | 0.000 | 0.398 | 0.828 | 9569 | tags=38%, list=16%, signal=45% |
| 3132 | RHYTHMIC\_PROCESS(GO:0048511) |  | 280 | 0.75 | 1.22 | 0.141 | 0.397 | 0.828 | 7676 | tags=16%, list=13%, signal=18% |
| 3133 | REGULATION\_OF\_DNA\_TEMPLATED\_TRANSCRIPTION\_\_INITIATION(GO:2000142) |  | 38 | 0.79 | 1.22 | 0.141 | 0.397 | 0.828 | 6114 | tags=13%, list=10%, signal=15% |
| 3134 | TOOTH\_MINERALIZATION(GO:0034505) |  | 23 | 0.76 | 1.22 | 0.000 | 0.397 | 0.828 | 9837 | tags=26%, list=17%, signal=31% |
| 3135 | ENDOMEMBRANE\_SYSTEM(GO:0012505) |  | 4363 | 0.77 | 1.22 | 0.141 | 0.397 | 0.828 | 7980 | tags=12%, list=13%, signal=13% |
| 3136 | DIOXYGENASE\_ACTIVITY(GO:0051213) |  | 92 | 0.76 | 1.22 | 0.141 | 0.397 | 0.828 | 8451 | tags=16%, list=14%, signal=19% |
| 3137 | POSITIVE\_REGULATION\_OF\_PROTEIN\_PHOSPHORYLATION(GO:0001934) |  | 954 | 0.75 | 1.22 | 0.141 | 0.397 | 0.828 | 7921 | tags=16%, list=13%, signal=18% |
| 3138 | GUANYL\_RIBONUCLEOTIDE\_BINDING(GO:0032561) |  | 403 | 0.75 | 1.22 | 0.141 | 0.397 | 0.828 | 7575 | tags=13%, list=13%, signal=15% |
| 3139 | POSITIVE\_REGULATION\_OF\_MUSCLE\_CELL\_DIFFERENTIATION(GO:0051149) |  | 102 | 0.74 | 1.22 | 0.141 | 0.397 | 0.828 | 9543 | tags=22%, list=16%, signal=26% |
| 3140 | POSITIVE\_REGULATION\_OF\_PHOSPHORUS\_METABOLIC\_PROCESS(GO:0010562) |  | 1089 | 0.75 | 1.22 | 0.141 | 0.397 | 0.828 | 7754 | tags=16%, list=13%, signal=18% |
| 3141 | POSITIVE\_REGULATION\_OF\_PHOSPHATE\_METABOLIC\_PROCESS(GO:0045937) |  | 1089 | 0.75 | 1.22 | 0.141 | 0.397 | 0.828 | 7754 | tags=16%, list=13%, signal=18% |
| 3142 | THIOESTER\_METABOLIC\_PROCESS(GO:0035383) |  | 98 | 0.74 | 1.22 | 0.141 | 0.397 | 0.828 | 8607 | tags=11%, list=15%, signal=13% |
| 3143 | ACYL\_COA\_METABOLIC\_PROCESS(GO:0006637) |  | 98 | 0.74 | 1.22 | 0.141 | 0.397 | 0.828 | 8607 | tags=11%, list=15%, signal=13% |
| 3144 | DENDRITE\_MORPHOGENESIS(GO:0048813) |  | 128 | 0.76 | 1.22 | 0.141 | 0.397 | 0.828 | 9191 | tags=21%, list=16%, signal=25% |
| 3145 | NEGATIVE\_REGULATION\_OF\_PROTEIN\_SECRETION(GO:0050709) |  | 116 | 0.73 | 1.22 | 0.141 | 0.397 | 0.828 | 10004 | tags=21%, list=17%, signal=25% |
| 3146 | COFACTOR\_BIOSYNTHETIC\_PROCESS(GO:0051188) |  | 302 | 0.75 | 1.22 | 0.141 | 0.397 | 0.828 | 6696 | tags=4%, list=11%, signal=4% |
| 3147 | REGULATION\_OF\_MUSCLE\_CELL\_DIFFERENTIATION(GO:0051147) |  | 169 | 0.74 | 1.22 | 0.141 | 0.397 | 0.828 | 9268 | tags=21%, list=16%, signal=24% |
| 3148 | POSITIVE\_REGULATION\_OF\_EPITHELIAL\_CELL\_DIFFERENTIATION(GO:0030858) |  | 62 | 0.79 | 1.22 | 0.141 | 0.397 | 0.828 | 8744 | tags=23%, list=15%, signal=26% |
| 3149 | FATTY\_ACID\_BINDING(GO:0005504) |  | 28 | 0.78 | 1.22 | 0.141 | 0.397 | 0.828 | 9096 | tags=32%, list=15%, signal=38% |
| 3150 | NEGATIVE\_REGULATION\_OF\_AXON\_EXTENSION(GO:0030517) |  | 24 | 0.78 | 1.22 | 0.141 | 0.397 | 0.828 | 8673 | tags=21%, list=15%, signal=24% |
| 3151 | ATP\_DEPENDENT\_MICROTUBULE\_MOTOR\_ACTIVITY(GO:1990939) |  | 46 | 0.72 | 1.22 | 0.000 | 0.397 | 0.828 | 13157 | tags=59%, list=22%, signal=75% |
| 3152 | PHOSPHATIDYLINOSITOL\_BINDING(GO:0035091) |  | 222 | 0.72 | 1.22 | 0.141 | 0.397 | 0.828 | 9358 | tags=17%, list=16%, signal=20% |
| 3153 | GUANYL\_NUCLEOTIDE\_BINDING(GO:0019001) |  | 404 | 0.75 | 1.22 | 0.141 | 0.397 | 0.828 | 7575 | tags=13%, list=13%, signal=15% |
| 3154 | NEGATIVE\_REGULATION\_OF\_TRANSLATION(GO:0017148) |  | 412 | 0.73 | 1.22 | 0.141 | 0.397 | 0.828 | 6397 | tags=6%, list=11%, signal=7% |
| 3155 | NEGATIVE\_REGULATION\_OF\_PEPTIDE\_SECRETION(GO:0002792) |  | 122 | 0.72 | 1.22 | 0.141 | 0.397 | 0.828 | 10004 | tags=20%, list=17%, signal=25% |
| 3156 | PHAGOSOME\_ACIDIFICATION(GO:0090383) |  | 27 | 0.75 | 1.22 | 0.141 | 0.397 | 0.828 | 6008 | tags=7%, list=10%, signal=8% |
| 3157 | RECOMBINATIONAL\_REPAIR(GO:0000725) |  | 112 | 0.71 | 1.22 | 0.000 | 0.396 | 0.828 | 13120 | tags=43%, list=22%, signal=55% |
| 3158 | RAC\_GTPASE\_BINDING(GO:0048365) |  | 64 | 0.74 | 1.22 | 0.141 | 0.396 | 0.828 | 8953 | tags=14%, list=15%, signal=17% |
| 3159 | REGULATION\_OF\_PHOSPHATASE\_ACTIVITY(GO:0010921) |  | 168 | 0.72 | 1.22 | 0.141 | 0.396 | 0.828 | 7544 | tags=12%, list=13%, signal=14% |
| 3160 | ASPARTATE\_FAMILY\_AMINO\_ACID\_BIOSYNTHETIC\_PROCESS(GO:0009067) |  | 22 | 0.72 | 1.22 | 0.000 | 0.396 | 0.828 | 11214 | tags=41%, list=19%, signal=50% |
| 3161 | METANEPHRIC\_MESENCHYME\_DEVELOPMENT(GO:0072075) |  | 15 | 0.79 | 1.22 | 0.141 | 0.396 | 0.828 | 9208 | tags=40%, list=16%, signal=47% |
| 3162 | PROTEIN\_HYDROXYLATION(GO:0018126) |  | 23 | 0.73 | 1.22 | 0.000 | 0.396 | 0.828 | 11786 | tags=35%, list=20%, signal=43% |
| 3163 | NUCLEAR\_CHROMATIN(GO:0000790) |  | 347 | 0.75 | 1.22 | 0.141 | 0.396 | 0.828 | 6774 | tags=8%, list=11%, signal=9% |
| 3164 | REGULATION\_OF\_SIGNAL\_TRANSDUCTION\_BY\_P53\_CLASS\_MEDIATOR(GO:1901796) |  | 177 | 0.71 | 1.22 | 0.000 | 0.396 | 0.828 | 10674 | tags=23%, list=18%, signal=27% |
| 3165 | PYRUVATE\_BIOSYNTHETIC\_PROCESS(GO:0042866) |  | 106 | 0.75 | 1.22 | 0.141 | 0.396 | 0.828 | 6696 | tags=5%, list=11%, signal=5% |
| 3166 | NEUROPEPTIDE\_RECEPTOR\_BINDING(GO:0071855) |  | 27 | 0.80 | 1.22 | 0.141 | 0.396 | 0.828 | 9985 | tags=48%, list=17%, signal=58% |
| 3167 | ADP\_METABOLIC\_PROCESS(GO:0046031) |  | 109 | 0.75 | 1.22 | 0.141 | 0.396 | 0.828 | 8541 | tags=11%, list=14%, signal=13% |
| 3168 | REGULATION\_OF\_BMP\_SIGNALING\_PATHWAY(GO:0030510) |  | 91 | 0.75 | 1.22 | 0.141 | 0.396 | 0.828 | 8177 | tags=24%, list=14%, signal=28% |
| 3169 | NUCLEIC\_ACID\_TEMPLATED\_TRANSCRIPTION(GO:0097659) |  | 3950 | 0.77 | 1.22 | 0.141 | 0.396 | 0.828 | 8072 | tags=14%, list=14%, signal=15% |
| 3170 | RESPONSE\_TO\_FOLLICLE\_STIMULATING\_HORMONE(GO:0032354) |  | 18 | 0.77 | 1.22 | 0.000 | 0.396 | 0.828 | 7172 | tags=22%, list=12%, signal=25% |
| 3171 | TRANSCRIPTION\_\_DNA\_TEMPLATED(GO:0006351) |  | 3935 | 0.77 | 1.22 | 0.141 | 0.396 | 0.828 | 8072 | tags=14%, list=14%, signal=15% |
| 3172 | PHAGOCYTOSIS(GO:0006909) |  | 326 | 0.72 | 1.22 | 0.141 | 0.396 | 0.828 | 8112 | tags=9%, list=14%, signal=10% |
| 3173 | CELLULAR\_COMPONENT\_BIOGENESIS(GO:0044085) |  | 3115 | 0.77 | 1.22 | 0.141 | 0.396 | 0.828 | 7973 | tags=11%, list=13%, signal=12% |
| 3174 | PRIMARY\_ALCOHOL\_METABOLIC\_PROCESS(GO:0034308) |  | 59 | 0.77 | 1.22 | 0.141 | 0.396 | 0.828 | 7255 | tags=14%, list=12%, signal=15% |
| 3175 | REGULATION\_OF\_DENDRITE\_EXTENSION(GO:1903859) |  | 21 | 0.74 | 1.22 | 0.000 | 0.396 | 0.828 | 8949 | tags=43%, list=15%, signal=50% |
| 3176 | REGULATION\_OF\_NON\_CANONICAL\_WNT\_SIGNALING\_PATHWAY(GO:2000050) |  | 21 | 0.81 | 1.22 | 0.141 | 0.396 | 0.828 | 8572 | tags=43%, list=14%, signal=50% |
| 3177 | RNA\_BIOSYNTHETIC\_PROCESS(GO:0032774) |  | 3966 | 0.77 | 1.22 | 0.141 | 0.396 | 0.828 | 8072 | tags=14%, list=14%, signal=15% |
| 3178 | LIPID\_BINDING(GO:0008289) |  | 674 | 0.74 | 1.22 | 0.141 | 0.396 | 0.828 | 9144 | tags=19%, list=15%, signal=22% |
| 3179 | CYTOPLASMIC\_MICROTUBULE\_ORGANIZATION(GO:0031122) |  | 43 | 0.71 | 1.22 | 0.000 | 0.396 | 0.828 | 7647 | tags=14%, list=13%, signal=16% |
| 3180 | PURINE\_RIBONUCLEOSIDE\_BINDING(GO:0032550) |  | 388 | 0.75 | 1.22 | 0.141 | 0.396 | 0.828 | 7575 | tags=12%, list=13%, signal=13% |
| 3181 | NITRIC\_OXIDE\_MEDIATED\_SIGNAL\_TRANSDUCTION(GO:0007263) |  | 25 | 0.79 | 1.22 | 0.141 | 0.396 | 0.828 | 10309 | tags=44%, list=17%, signal=53% |
| 3182 | CHONDROITIN\_SULFATE\_BIOSYNTHETIC\_PROCESS(GO:0030206) |  | 25 | 0.74 | 1.22 | 0.000 | 0.396 | 0.828 | 9277 | tags=44%, list=16%, signal=52% |
| 3183 | REGULATION\_OF\_MYOBLAST\_DIFFERENTIATION(GO:0045661) |  | 50 | 0.77 | 1.22 | 0.141 | 0.396 | 0.828 | 9268 | tags=32%, list=16%, signal=38% |
| 3184 | REGULATION\_OF\_CILIUM\_ASSEMBLY(GO:1902017) |  | 43 | 0.71 | 1.22 | 0.000 | 0.396 | 0.828 | 9765 | tags=26%, list=17%, signal=31% |
| 3185 | POSITIVE\_REGULATION\_OF\_INTERLEUKIN\_2\_PRODUCTION(GO:0032743) |  | 30 | 0.74 | 1.22 | 0.000 | 0.396 | 0.828 | 7488 | tags=30%, list=13%, signal=34% |
| 3186 | RECEPTOR\_MEDIATED\_ENDOCYTOSIS(GO:0006898) |  | 323 | 0.74 | 1.22 | 0.141 | 0.396 | 0.828 | 8937 | tags=17%, list=15%, signal=20% |
| 3187 | NITRIC\_OXIDE\_METABOLIC\_PROCESS(GO:0046209) |  | 69 | 0.75 | 1.22 | 0.141 | 0.396 | 0.828 | 7873 | tags=19%, list=13%, signal=22% |
| 3188 | NEUROTRANSMITTER\_RECEPTOR\_ACTIVITY(GO:0030594) |  | 107 | 0.75 | 1.22 | 0.141 | 0.396 | 0.828 | 9798 | tags=28%, list=17%, signal=34% |
| 3189 | NOSE\_DEVELOPMENT(GO:0043584) |  | 16 | 0.80 | 1.22 | 0.000 | 0.396 | 0.828 | 5935 | tags=13%, list=10%, signal=14% |
| 3190 | REGULATION\_OF\_METANEPHROS\_DEVELOPMENT(GO:0072215) |  | 23 | 0.80 | 1.22 | 0.141 | 0.396 | 0.828 | 8339 | tags=22%, list=14%, signal=25% |
| 3191 | POSITIVE\_REGULATION\_OF\_PEPTIDYL\_LYSINE\_ACETYLATION(GO:2000758) |  | 29 | 0.76 | 1.22 | 0.141 | 0.395 | 0.828 | 7029 | tags=10%, list=12%, signal=12% |
| 3192 | REGULATION\_OF\_CYTOKINE\_PRODUCTION\_INVOLVED\_IN\_IMMUNE\_RESPONSE(GO:0002718) |  | 65 | 0.74 | 1.22 | 0.141 | 0.395 | 0.828 | 10793 | tags=28%, list=18%, signal=34% |
| 3193 | POSITIVE\_REGULATION\_OF\_CD4\_POSITIVE\_\_ALPHA\_BETA\_T\_CELL\_DIFFERENTIATION(GO:0043372) |  | 25 | 0.74 | 1.22 | 0.141 | 0.395 | 0.828 | 9368 | tags=36%, list=16%, signal=43% |
| 3194 | ENDOPLASMIC\_RETICULUM\_MEMBRANE(GO:0005789) |  | 1025 | 0.73 | 1.22 | 0.141 | 0.395 | 0.828 | 8013 | tags=11%, list=14%, signal=13% |
| 3195 | CGMP\_METABOLIC\_PROCESS(GO:0046068) |  | 53 | 0.77 | 1.22 | 0.141 | 0.395 | 0.828 | 10245 | tags=40%, list=17%, signal=48% |
| 3196 | POSITIVE\_REGULATION\_OF\_HISTONE\_ACETYLATION(GO:0035066) |  | 28 | 0.76 | 1.22 | 0.141 | 0.395 | 0.828 | 7029 | tags=11%, list=12%, signal=12% |
| 3197 | REGULATION\_OF\_INTRACELLULAR\_STEROID\_HORMONE\_RECEPTOR\_SIGNALING\_PATHWAY(GO:0033143) |  | 78 | 0.77 | 1.22 | 0.141 | 0.395 | 0.828 | 8112 | tags=18%, list=14%, signal=21% |
| 3198 | PURINE\_NUCLEOSIDE\_BINDING(GO:0001883) |  | 391 | 0.75 | 1.22 | 0.141 | 0.395 | 0.828 | 7575 | tags=12%, list=13%, signal=13% |
| 3199 | TRANSFERASE\_COMPLEX\_\_TRANSFERRING\_PHOSPHORUS\_CONTAINING\_GROUPS(GO:0061695) |  | 267 | 0.70 | 1.22 | 0.000 | 0.395 | 0.828 | 10725 | tags=17%, list=18%, signal=21% |
| 3200 | POSITIVE\_REGULATION\_OF\_CELL\_JUNCTION\_ASSEMBLY(GO:1901890) |  | 31 | 0.77 | 1.22 | 0.141 | 0.395 | 0.828 | 7711 | tags=16%, list=13%, signal=19% |
| 3201 | ENDOPLASMIC\_RETICULUM\_SUBCOMPARTMENT(GO:0098827) |  | 1029 | 0.73 | 1.22 | 0.141 | 0.395 | 0.828 | 8013 | tags=11%, list=14%, signal=13% |
| 3202 | OXIDOREDUCTASE\_ACTIVITY\_\_OXIDIZING\_METAL\_IONS(GO:0016722) |  | 15 | 0.78 | 1.22 | 0.141 | 0.395 | 0.828 | 11214 | tags=40%, list=19%, signal=49% |
| 3203 | DEFINITIVE\_HEMOPOIESIS(GO:0060216) |  | 20 | 0.75 | 1.22 | 0.000 | 0.395 | 0.828 | 12195 | tags=55%, list=21%, signal=69% |
| 3204 | CHROMATIN(GO:0000785) |  | 520 | 0.74 | 1.22 | 0.141 | 0.395 | 0.828 | 6774 | tags=7%, list=11%, signal=8% |
| 3205 | PROTEIN\_HETERODIMERIZATION\_ACTIVITY(GO:0046982) |  | 528 | 0.73 | 1.22 | 0.141 | 0.395 | 0.828 | 9484 | tags=18%, list=16%, signal=21% |
| 3206 | GENERATION\_OF\_PRECURSOR\_METABOLITES\_AND\_ENERGY(GO:0006091) |  | 513 | 0.72 | 1.21 | 0.141 | 0.395 | 0.828 | 7821 | tags=7%, list=13%, signal=8% |
| 3207 | REGULATION\_OF\_RNA\_METABOLIC\_PROCESS(GO:0051252) |  | 3977 | 0.77 | 1.21 | 0.141 | 0.395 | 0.828 | 7957 | tags=13%, list=13%, signal=15% |
| 3208 | PLATELET\_DERIVED\_GROWTH\_FACTOR\_RECEPTOR\_BINDING(GO:0005161) |  | 15 | 0.80 | 1.21 | 0.141 | 0.395 | 0.828 | 10643 | tags=47%, list=18%, signal=57% |
| 3209 | POSITIVE\_CHEMOTAXIS(GO:0050918) |  | 60 | 0.78 | 1.21 | 0.141 | 0.395 | 0.828 | 7343 | tags=18%, list=12%, signal=21% |
| 3210 | CHLORIDE\_TRANSMEMBRANE\_TRANSPORT(GO:1902476) |  | 85 | 0.75 | 1.21 | 0.141 | 0.395 | 0.828 | 8596 | tags=25%, list=15%, signal=29% |
| 3211 | POSITIVE\_REGULATION\_OF\_T\_HELPER\_CELL\_DIFFERENTIATION(GO:0045624) |  | 19 | 0.77 | 1.21 | 0.141 | 0.395 | 0.828 | 9023 | tags=32%, list=15%, signal=37% |
| 3212 | CELL\_MIGRATION\_IN\_HINDBRAIN(GO:0021535) |  | 15 | 0.76 | 1.21 | 0.000 | 0.395 | 0.828 | 11902 | tags=53%, list=20%, signal=67% |
| 3213 | CADHERIN\_BINDING(GO:0045296) |  | 304 | 0.75 | 1.21 | 0.141 | 0.395 | 0.828 | 6203 | tags=2%, list=10%, signal=2% |
| 3214 | ACTIN\_FILAMENT\_ORGANIZATION(GO:0007015) |  | 364 | 0.75 | 1.21 | 0.141 | 0.395 | 0.828 | 8787 | tags=17%, list=15%, signal=20% |
| 3215 | PURINE\_CONTAINING\_COMPOUND\_SALVAGE(GO:0043101) |  | 16 | 0.72 | 1.21 | 0.000 | 0.395 | 0.828 | 9191 | tags=19%, list=16%, signal=22% |
| 3216 | DISTAL\_AXON(GO:0150034) |  | 262 | 0.76 | 1.21 | 0.141 | 0.395 | 0.828 | 10143 | tags=27%, list=17%, signal=32% |
| 3217 | BICELLULAR\_TIGHT\_JUNCTION\_ASSEMBLY(GO:0070830) |  | 51 | 0.76 | 1.21 | 0.141 | 0.395 | 0.828 | 6796 | tags=8%, list=11%, signal=9% |
| 3218 | GONAD\_DEVELOPMENT(GO:0008406) |  | 213 | 0.75 | 1.21 | 0.141 | 0.395 | 0.828 | 8613 | tags=20%, list=15%, signal=24% |
| 3219 | DEVELOPMENT\_OF\_PRIMARY\_SEXUAL\_CHARACTERISTICS(GO:0045137) |  | 219 | 0.76 | 1.21 | 0.141 | 0.395 | 0.828 | 8613 | tags=21%, list=15%, signal=24% |
| 3220 | POSITIVE\_REGULATION\_OF\_PEPTIDE\_SECRETION(GO:0002793) |  | 261 | 0.73 | 1.21 | 0.141 | 0.395 | 0.828 | 7898 | tags=16%, list=13%, signal=19% |
| 3221 | CORE\_PROMOTER\_BINDING(GO:0001047) |  | 133 | 0.76 | 1.21 | 0.141 | 0.395 | 0.828 | 6441 | tags=8%, list=11%, signal=8% |
| 3222 | ION\_CHANNEL\_REGULATOR\_ACTIVITY(GO:0099106) |  | 94 | 0.72 | 1.21 | 0.141 | 0.395 | 0.828 | 9554 | tags=28%, list=16%, signal=33% |
| 3223 | REGULATION\_OF\_MYOBLAST\_PROLIFERATION(GO:2000291) |  | 23 | 0.75 | 1.21 | 0.000 | 0.395 | 0.828 | 12440 | tags=39%, list=21%, signal=50% |
| 3224 | REGULATION\_OF\_RESPONSE\_TO\_DNA\_DAMAGE\_STIMULUS(GO:2001020) |  | 184 | 0.77 | 1.21 | 0.141 | 0.394 | 0.828 | 7101 | tags=7%, list=12%, signal=8% |
| 3225 | PROTEIN\_SECRETION(GO:0009306) |  | 525 | 0.72 | 1.21 | 0.141 | 0.394 | 0.828 | 10552 | tags=22%, list=18%, signal=27% |
| 3226 | MIDBRAIN\_DEVELOPMENT(GO:0030901) |  | 94 | 0.74 | 1.21 | 0.141 | 0.394 | 0.828 | 8969 | tags=27%, list=15%, signal=31% |
| 3227 | NEUROEPITHELIAL\_CELL\_DIFFERENTIATION(GO:0060563) |  | 54 | 0.76 | 1.21 | 0.141 | 0.394 | 0.828 | 9354 | tags=26%, list=16%, signal=31% |
| 3228 | NUCLEOSIDE\_DIPHOSPHATE\_PHOSPHORYLATION(GO:0006165) |  | 120 | 0.74 | 1.21 | 0.141 | 0.394 | 0.828 | 6696 | tags=6%, list=11%, signal=7% |
| 3229 | NEGATIVE\_REGULATION\_OF\_EPITHELIAL\_CELL\_APOPTOTIC\_PROCESS(GO:1904036) |  | 42 | 0.77 | 1.21 | 0.141 | 0.394 | 0.828 | 6186 | tags=19%, list=10%, signal=21% |
| 3230 | REGULATION\_OF\_VASCULAR\_SMOOTH\_MUSCLE\_CELL\_DIFFERENTIATION(GO:1905063) |  | 20 | 0.79 | 1.21 | 0.141 | 0.394 | 0.828 | 8774 | tags=30%, list=15%, signal=35% |
| 3231 | RIBONUCLEOSIDE\_BINDING(GO:0032549) |  | 392 | 0.75 | 1.21 | 0.141 | 0.394 | 0.828 | 7575 | tags=11%, list=13%, signal=13% |
| 3232 | NUCLEAR\_MEMBRANE(GO:0031965) |  | 302 | 0.71 | 1.21 | 0.000 | 0.394 | 0.828 | 9315 | tags=14%, list=16%, signal=16% |
| 3233 | PYRUVATE\_METABOLIC\_PROCESS(GO:0006090) |  | 138 | 0.74 | 1.21 | 0.141 | 0.394 | 0.828 | 6696 | tags=7%, list=11%, signal=8% |
| 3234 | REGULATION\_OF\_HORMONE\_SECRETION(GO:0046883) |  | 265 | 0.73 | 1.21 | 0.141 | 0.394 | 0.828 | 10131 | tags=25%, list=17%, signal=30% |
| 3235 | REGULATION\_OF\_ALPHA\_BETA\_T\_CELL\_PROLIFERATION(GO:0046640) |  | 21 | 0.75 | 1.21 | 0.000 | 0.394 | 0.828 | 9023 | tags=29%, list=15%, signal=34% |
| 3236 | T\_HELPER\_17\_TYPE\_IMMUNE\_RESPONSE(GO:0072538) |  | 22 | 0.77 | 1.21 | 0.141 | 0.394 | 0.828 | 9023 | tags=27%, list=15%, signal=32% |
| 3237 | NUCLEOBASE\_CONTAINING\_COMPOUND\_CATABOLIC\_PROCESS(GO:0034655) |  | 621 | 0.74 | 1.21 | 0.141 | 0.394 | 0.828 | 8602 | tags=10%, list=15%, signal=12% |
| 3238 | MAP\_KINASE\_KINASE\_ACTIVITY(GO:0004708) |  | 15 | 0.76 | 1.21 | 0.000 | 0.394 | 0.828 | 7426 | tags=20%, list=13%, signal=23% |
| 3239 | ANION\_TRANSMEMBRANE\_TRANSPORTER\_ACTIVITY(GO:0008509) |  | 337 | 0.74 | 1.21 | 0.141 | 0.394 | 0.828 | 9669 | tags=26%, list=16%, signal=31% |
| 3240 | CELLULAR\_RESPONSE\_TO\_HYPOXIA(GO:0071456) |  | 202 | 0.72 | 1.21 | 0.141 | 0.394 | 0.828 | 7727 | tags=9%, list=13%, signal=10% |
| 3241 | SCAFFOLD\_PROTEIN\_BINDING(GO:0097110) |  | 59 | 0.76 | 1.21 | 0.141 | 0.394 | 0.828 | 8363 | tags=17%, list=14%, signal=20% |
| 3242 | PROSTAGLANDIN\_METABOLIC\_PROCESS(GO:0006693) |  | 34 | 0.75 | 1.21 | 0.141 | 0.394 | 0.828 | 7682 | tags=21%, list=13%, signal=24% |
| 3243 | PROSTANOID\_METABOLIC\_PROCESS(GO:0006692) |  | 34 | 0.75 | 1.21 | 0.141 | 0.393 | 0.828 | 7682 | tags=21%, list=13%, signal=24% |
| 3244 | MALE\_MEIOTIC\_NUCLEAR\_DIVISION(GO:0007140) |  | 47 | 0.76 | 1.21 | 0.141 | 0.393 | 0.828 | 8276 | tags=21%, list=14%, signal=25% |
| 3245 | REGULATION\_OF\_VIRAL\_PROCESS(GO:0050792) |  | 190 | 0.74 | 1.21 | 0.141 | 0.393 | 0.828 | 7293 | tags=9%, list=12%, signal=10% |
| 3246 | MONOSACCHARIDE\_TRANSMEMBRANE\_TRANSPORTER\_ACTIVITY(GO:0015145) |  | 24 | 0.75 | 1.21 | 0.141 | 0.393 | 0.828 | 8716 | tags=38%, list=15%, signal=44% |
| 3247 | GROWTH\_PLATE\_CARTILAGE\_DEVELOPMENT(GO:0003417) |  | 15 | 0.80 | 1.21 | 0.141 | 0.393 | 0.828 | 6051 | tags=20%, list=10%, signal=22% |
| 3248 | PHAGOCYTIC\_VESICLE(GO:0045335) |  | 121 | 0.72 | 1.21 | 0.141 | 0.393 | 0.828 | 8121 | tags=10%, list=14%, signal=11% |
| 3249 | COENZYME\_BIOSYNTHETIC\_PROCESS(GO:0009108) |  | 244 | 0.73 | 1.21 | 0.141 | 0.393 | 0.828 | 6696 | tags=4%, list=11%, signal=5% |
| 3250 | ENZYME\_INHIBITOR\_ACTIVITY(GO:0004857) |  | 397 | 0.73 | 1.21 | 0.141 | 0.393 | 0.828 | 8070 | tags=16%, list=14%, signal=19% |
| 3251 | REGULATION\_OF\_SECRETION\_BY\_CELL(GO:1903530) |  | 678 | 0.73 | 1.21 | 0.141 | 0.393 | 0.828 | 10131 | tags=22%, list=17%, signal=26% |
| 3252 | POSITIVE\_REGULATION\_OF\_EPITHELIAL\_CELL\_MIGRATION(GO:0010634) |  | 154 | 0.75 | 1.21 | 0.141 | 0.393 | 0.828 | 8046 | tags=19%, list=14%, signal=22% |
| 3253 | ROUGH\_ENDOPLASMIC\_RETICULUM(GO:0005791) |  | 84 | 0.72 | 1.21 | 0.000 | 0.393 | 0.828 | 7659 | tags=12%, list=13%, signal=14% |
| 3254 | FATTY\_ACID\_BETA\_OXIDATION(GO:0006635) |  | 72 | 0.72 | 1.21 | 0.000 | 0.393 | 0.828 | 10056 | tags=17%, list=17%, signal=20% |
| 3255 | HIGH\_DENSITY\_LIPOPROTEIN\_PARTICLE(GO:0034364) |  | 27 | 0.76 | 1.21 | 0.000 | 0.393 | 0.828 | 6362 | tags=19%, list=11%, signal=21% |
| 3256 | MICROTUBULE\_BASED\_MOVEMENT(GO:0007018) |  | 307 | 0.70 | 1.21 | 0.000 | 0.393 | 0.828 | 10334 | tags=29%, list=17%, signal=35% |
| 3257 | CHROMATIN\_DNA\_BINDING(GO:0031490) |  | 104 | 0.75 | 1.21 | 0.141 | 0.393 | 0.828 | 9479 | tags=17%, list=16%, signal=21% |
| 3258 | SUPRAMOLECULAR\_FIBER(GO:0099512) |  | 912 | 0.76 | 1.21 | 0.141 | 0.393 | 0.828 | 8823 | tags=20%, list=15%, signal=23% |
| 3259 | ORGANIC\_CYCLIC\_COMPOUND\_BIOSYNTHETIC\_PROCESS(GO:1901362) |  | 4720 | 0.76 | 1.21 | 0.141 | 0.393 | 0.828 | 8072 | tags=13%, list=14%, signal=14% |
| 3260 | POSITIVE\_REGULATION\_OF\_DNA\_REPLICATION(GO:0045740) |  | 55 | 0.74 | 1.21 | 0.141 | 0.393 | 0.828 | 11102 | tags=31%, list=19%, signal=38% |
| 3261 | INNER\_DYNEIN\_ARM\_ASSEMBLY(GO:0036159) |  | 16 | 0.73 | 1.21 | 0.000 | 0.393 | 0.828 | 14289 | tags=63%, list=24%, signal=82% |
| 3262 | DELAYED\_RECTIFIER\_POTASSIUM\_CHANNEL\_ACTIVITY(GO:0005251) |  | 21 | 0.79 | 1.21 | 0.141 | 0.393 | 0.828 | 9472 | tags=52%, list=16%, signal=62% |
| 3263 | REGULATION\_OF\_MITOTIC\_CELL\_CYCLE(GO:0007346) |  | 662 | 0.74 | 1.21 | 0.141 | 0.393 | 0.828 | 10601 | tags=19%, list=18%, signal=23% |
| 3264 | HISTONE\_MODIFICATION(GO:0016570) |  | 451 | 0.72 | 1.21 | 0.000 | 0.393 | 0.828 | 8046 | tags=8%, list=14%, signal=9% |
| 3265 | POSITIVE\_REGULATION\_OF\_PROTEIN\_KINASE\_ACTIVITY(GO:0045860) |  | 516 | 0.72 | 1.21 | 0.141 | 0.393 | 0.828 | 7921 | tags=14%, list=13%, signal=16% |
| 3266 | ACTOMYOSIN(GO:0042641) |  | 65 | 0.73 | 1.21 | 0.141 | 0.393 | 0.828 | 8594 | tags=18%, list=15%, signal=22% |
| 3267 | REGULATION\_OF\_DENDRITE\_MORPHOGENESIS(GO:0048814) |  | 76 | 0.73 | 1.21 | 0.141 | 0.393 | 0.828 | 9607 | tags=24%, list=16%, signal=28% |
| 3268 | REGULATION\_OF\_MICROTUBULE\_DEPOLYMERIZATION(GO:0031114) |  | 21 | 0.75 | 1.21 | 0.141 | 0.393 | 0.828 | 11641 | tags=38%, list=20%, signal=47% |
| 3269 | REGULATION\_OF\_DNA\_RECOMBINATION(GO:0000018) |  | 73 | 0.75 | 1.21 | 0.141 | 0.393 | 0.828 | 7022 | tags=12%, list=12%, signal=14% |
| 3270 | MEMBRANE\_REPOLARIZATION\_DURING\_VENTRICULAR\_CARDIAC\_MUSCLE\_CELL\_ACTION\_POTENTIAL(GO:0098915) |  | 16 | 0.76 | 1.21 | 0.141 | 0.393 | 0.828 | 9472 | tags=25%, list=16%, signal=30% |
| 3271 | ENDOTHELIAL\_CELL\_APOPTOTIC\_PROCESS(GO:0072577) |  | 56 | 0.76 | 1.21 | 0.141 | 0.393 | 0.828 | 8627 | tags=16%, list=15%, signal=19% |
| 3272 | CHONDROITIN\_SULFATE\_PROTEOGLYCAN\_METABOLIC\_PROCESS(GO:0050654) |  | 43 | 0.72 | 1.21 | 0.000 | 0.392 | 0.828 | 9569 | tags=44%, list=16%, signal=53% |
| 3273 | CYTOPLASMIC\_SIDE\_OF\_MEMBRANE(GO:0098562) |  | 204 | 0.75 | 1.21 | 0.141 | 0.392 | 0.828 | 9411 | tags=15%, list=16%, signal=17% |
| 3274 | REGULATION\_OF\_CELLULAR\_MACROMOLECULE\_BIOSYNTHETIC\_PROCESS(GO:2000112) |  | 4347 | 0.76 | 1.21 | 0.141 | 0.392 | 0.828 | 7957 | tags=13%, list=13%, signal=14% |
| 3275 | RESPONSE\_TO\_STRESS(GO:0006950) |  | 3890 | 0.76 | 1.21 | 0.141 | 0.392 | 0.828 | 7975 | tags=12%, list=13%, signal=13% |
| 3276 | EXTRACELLULAR\_MATRIX\_BINDING(GO:0050840) |  | 53 | 0.76 | 1.21 | 0.141 | 0.392 | 0.828 | 8865 | tags=25%, list=15%, signal=29% |
| 3277 | PROSTAGLANDIN\_TRANSPORT(GO:0015732) |  | 16 | 0.81 | 1.21 | 0.141 | 0.392 | 0.828 | 6964 | tags=25%, list=12%, signal=28% |
| 3278 | SPECIFICATION\_OF\_ANIMAL\_ORGAN\_IDENTITY(GO:0010092) |  | 30 | 0.79 | 1.21 | 0.141 | 0.392 | 0.828 | 9208 | tags=30%, list=16%, signal=36% |
| 3279 | ACETYLGALACTOSAMINYLTRANSFERASE\_ACTIVITY(GO:0008376) |  | 34 | 0.74 | 1.21 | 0.141 | 0.392 | 0.828 | 7053 | tags=24%, list=12%, signal=27% |
| 3280 | CYTOKINE\_PRODUCTION\_INVOLVED\_IN\_IMMUNE\_RESPONSE(GO:0002367) |  | 84 | 0.72 | 1.21 | 0.141 | 0.392 | 0.828 | 10793 | tags=30%, list=18%, signal=36% |
| 3281 | CHOLESTEROL\_EFFLUX(GO:0033344) |  | 48 | 0.75 | 1.21 | 0.141 | 0.392 | 0.828 | 11522 | tags=29%, list=19%, signal=36% |
| 3282 | PLATELET\_DENSE\_GRANULE(GO:0042827) |  | 21 | 0.74 | 1.21 | 0.141 | 0.392 | 0.828 | 7060 | tags=29%, list=12%, signal=32% |
| 3283 | POSITIVE\_REGULATION\_OF\_CELL\_ACTIVATION(GO:0050867) |  | 361 | 0.73 | 1.21 | 0.141 | 0.392 | 0.828 | 9411 | tags=19%, list=16%, signal=22% |
| 3284 | CELLULAR\_RESPONSE\_TO\_STARVATION(GO:0009267) |  | 137 | 0.73 | 1.21 | 0.141 | 0.392 | 0.828 | 8210 | tags=12%, list=14%, signal=14% |
| 3285 | ADRENERGIC\_RECEPTOR\_BINDING(GO:0031690) |  | 19 | 0.73 | 1.21 | 0.000 | 0.392 | 0.828 | 12549 | tags=32%, list=21%, signal=40% |
| 3286 | TROPOMYOSIN\_BINDING(GO:0005523) |  | 15 | 0.79 | 1.21 | 0.141 | 0.392 | 0.828 | 7737 | tags=40%, list=13%, signal=46% |
| 3287 | AMINOGLYCAN\_CATABOLIC\_PROCESS(GO:0006026) |  | 66 | 0.73 | 1.21 | 0.141 | 0.392 | 0.828 | 11868 | tags=36%, list=20%, signal=45% |
| 3288 | ORGAN\_OR\_TISSUE\_SPECIFIC\_IMMUNE\_RESPONSE(GO:0002251) |  | 33 | 0.76 | 1.21 | 0.141 | 0.392 | 0.828 | 10916 | tags=21%, list=18%, signal=26% |
| 3289 | POSITIVE\_REGULATION\_OF\_STEM\_CELL\_DIFFERENTIATION(GO:2000738) |  | 18 | 0.79 | 1.21 | 0.141 | 0.392 | 0.828 | 7260 | tags=28%, list=12%, signal=32% |
| 3290 | AROMATIC\_COMPOUND\_BIOSYNTHETIC\_PROCESS(GO:0019438) |  | 4574 | 0.76 | 1.21 | 0.141 | 0.392 | 0.828 | 8072 | tags=13%, list=14%, signal=14% |
| 3291 | REGULATION\_OF\_PROTEIN\_POLYMERIZATION(GO:0032271) |  | 195 | 0.72 | 1.21 | 0.141 | 0.392 | 0.828 | 8112 | tags=12%, list=14%, signal=14% |
| 3292 | REGULATION\_OF\_SYMBIOSIS\_\_ENCOMPASSING\_MUTUALISM\_THROUGH\_PARASITISM(GO:0043903) |  | 218 | 0.74 | 1.21 | 0.141 | 0.392 | 0.828 | 7435 | tags=10%, list=13%, signal=11% |
| 3293 | CELLULAR\_NITROGEN\_COMPOUND\_CATABOLIC\_PROCESS(GO:0044270) |  | 668 | 0.74 | 1.21 | 0.141 | 0.392 | 0.828 | 8602 | tags=10%, list=15%, signal=12% |
| 3294 | ALPHA\_AMINO\_ACID\_METABOLIC\_PROCESS(GO:1901605) |  | 224 | 0.75 | 1.21 | 0.141 | 0.392 | 0.828 | 8755 | tags=18%, list=15%, signal=21% |
| 3295 | TONGUE\_DEVELOPMENT(GO:0043586) |  | 21 | 0.78 | 1.21 | 0.000 | 0.392 | 0.828 | 11988 | tags=67%, list=20%, signal=84% |
| 3296 | NEGATIVE\_REGULATION\_OF\_CYTOSKELETON\_ORGANIZATION(GO:0051494) |  | 129 | 0.75 | 1.21 | 0.141 | 0.392 | 0.828 | 7973 | tags=15%, list=13%, signal=17% |
| 3297 | REGULATION\_OF\_INTERLEUKIN\_6\_BIOSYNTHETIC\_PROCESS(GO:0045408) |  | 15 | 0.77 | 1.21 | 0.000 | 0.392 | 0.828 | 6530 | tags=20%, list=11%, signal=22% |
| 3298 | CYTOPLASMIC\_STRESS\_GRANULE(GO:0010494) |  | 44 | 0.74 | 1.21 | 0.000 | 0.392 | 0.828 | 5906 | tags=2%, list=10%, signal=3% |
| 3299 | GOLGI\_STACK(GO:0005795) |  | 125 | 0.74 | 1.21 | 0.141 | 0.392 | 0.828 | 7121 | tags=9%, list=12%, signal=10% |
| 3300 | NUCLEOSIDE\_BINDING(GO:0001882) |  | 399 | 0.75 | 1.21 | 0.141 | 0.392 | 0.828 | 7575 | tags=11%, list=13%, signal=13% |
| 3301 | COVALENT\_CHROMATIN\_MODIFICATION(GO:0016569) |  | 464 | 0.72 | 1.21 | 0.000 | 0.392 | 0.828 | 8046 | tags=8%, list=14%, signal=9% |
| 3302 | NEGATIVE\_REGULATION\_OF\_CELL\_CELL\_ADHESION(GO:0022408) |  | 157 | 0.73 | 1.21 | 0.141 | 0.392 | 0.828 | 9446 | tags=17%, list=16%, signal=20% |
| 3303 | REPLICATION\_FORK(GO:0005657) |  | 66 | 0.70 | 1.21 | 0.000 | 0.391 | 0.828 | 13312 | tags=52%, list=23%, signal=66% |
| 3304 | PEPTIDYL\_TYROSINE\_PHOSPHORYLATION(GO:0018108) |  | 378 | 0.73 | 1.21 | 0.141 | 0.392 | 0.828 | 9522 | tags=23%, list=16%, signal=27% |
| 3305 | GLYCOSAMINOGLYCAN\_CATABOLIC\_PROCESS(GO:0006027) |  | 61 | 0.73 | 1.21 | 0.141 | 0.391 | 0.828 | 11868 | tags=34%, list=20%, signal=43% |
| 3306 | GTPASE\_REGULATOR\_ACTIVITY(GO:0030695) |  | 314 | 0.74 | 1.21 | 0.141 | 0.391 | 0.828 | 8030 | tags=13%, list=14%, signal=15% |
| 3307 | DEVELOPMENTAL\_INDUCTION(GO:0031128) |  | 30 | 0.79 | 1.21 | 0.141 | 0.391 | 0.828 | 9208 | tags=37%, list=16%, signal=43% |
| 3308 | HETEROCYCLE\_CATABOLIC\_PROCESS(GO:0046700) |  | 667 | 0.74 | 1.21 | 0.141 | 0.391 | 0.828 | 8650 | tags=10%, list=15%, signal=12% |
| 3309 | NEGATIVE\_REGULATION\_OF\_LEUKOCYTE\_DIFFERENTIATION(GO:1902106) |  | 91 | 0.73 | 1.21 | 0.141 | 0.391 | 0.828 | 10798 | tags=24%, list=18%, signal=30% |
| 3310 | POSITIVE\_REGULATION\_OF\_CELL\_MOTILITY(GO:2000147) |  | 488 | 0.75 | 1.21 | 0.141 | 0.391 | 0.828 | 8195 | tags=17%, list=14%, signal=20% |
| 3311 | REGULATION\_OF\_CELL\_MORPHOGENESIS\_INVOLVED\_IN\_DIFFERENTIATION(GO:0010769) |  | 259 | 0.75 | 1.21 | 0.141 | 0.391 | 0.828 | 7459 | tags=18%, list=13%, signal=21% |
| 3312 | SPERM\_EGG\_RECOGNITION(GO:0035036) |  | 51 | 0.72 | 1.21 | 0.000 | 0.391 | 0.828 | 10928 | tags=29%, list=18%, signal=36% |
| 3313 | MORPHOGENESIS\_OF\_EMBRYONIC\_EPITHELIUM(GO:0016331) |  | 153 | 0.76 | 1.21 | 0.141 | 0.391 | 0.828 | 8122 | tags=16%, list=14%, signal=19% |
| 3314 | REGULATION\_OF\_CELLULAR\_BIOSYNTHETIC\_PROCESS(GO:0031326) |  | 4607 | 0.76 | 1.21 | 0.141 | 0.391 | 0.828 | 7957 | tags=13%, list=13%, signal=14% |
| 3315 | LIPID\_TRANSPORTER\_ACTIVITY(GO:0005319) |  | 140 | 0.75 | 1.21 | 0.141 | 0.391 | 0.828 | 7699 | tags=18%, list=13%, signal=20% |
| 3316 | REGULATION\_OF\_INSULIN\_SECRETION(GO:0050796) |  | 176 | 0.71 | 1.21 | 0.141 | 0.391 | 0.828 | 10124 | tags=23%, list=17%, signal=27% |
| 3317 | REGULATION\_OF\_EXOCYTOSIS(GO:0017157) |  | 171 | 0.73 | 1.21 | 0.141 | 0.391 | 0.828 | 9227 | tags=20%, list=16%, signal=23% |
| 3318 | HETEROCYCLE\_BIOSYNTHETIC\_PROCESS(GO:0018130) |  | 4568 | 0.76 | 1.21 | 0.141 | 0.391 | 0.828 | 8072 | tags=13%, list=14%, signal=14% |
| 3319 | COMPLEMENT\_ACTIVATION\_\_CLASSICAL\_PATHWAY(GO:0006958) |  | 133 | 0.70 | 1.21 | 0.141 | 0.391 | 0.828 | 11910 | tags=8%, list=20%, signal=9% |
| 3320 | CANONICAL\_WNT\_SIGNALING\_PATHWAY(GO:0060070) |  | 313 | 0.73 | 1.21 | 0.141 | 0.391 | 0.828 | 8771 | tags=17%, list=15%, signal=20% |
| 3321 | NUCLEOBASE\_CONTAINING\_COMPOUND\_BIOSYNTHETIC\_PROCESS(GO:0034654) |  | 4504 | 0.76 | 1.21 | 0.141 | 0.391 | 0.828 | 8072 | tags=13%, list=14%, signal=14% |
| 3322 | NEGATIVE\_REGULATION\_OF\_CELLULAR\_COMPONENT\_ORGANIZATION(GO:0051129) |  | 640 | 0.74 | 1.21 | 0.141 | 0.391 | 0.828 | 9316 | tags=17%, list=16%, signal=20% |
| 3323 | MANNOSE\_BINDING(GO:0005537) |  | 18 | 0.74 | 1.21 | 0.000 | 0.391 | 0.828 | 10008 | tags=39%, list=17%, signal=47% |
| 3324 | MICROTUBULE\_BUNDLE\_FORMATION(GO:0001578) |  | 83 | 0.73 | 1.21 | 0.141 | 0.391 | 0.828 | 8933 | tags=22%, list=15%, signal=26% |
| 3325 | ALPHA\_BETA\_T\_CELL\_PROLIFERATION(GO:0046633) |  | 24 | 0.73 | 1.21 | 0.000 | 0.391 | 0.828 | 9023 | tags=25%, list=15%, signal=29% |
| 3326 | ANION\_BINDING(GO:0043168) |  | 2778 | 0.75 | 1.21 | 0.141 | 0.391 | 0.828 | 8258 | tags=15%, list=14%, signal=17% |
| 3327 | REGULATION\_OF\_PLASMA\_LIPOPROTEIN\_PARTICLE\_LEVELS(GO:0097006) |  | 88 | 0.72 | 1.21 | 0.000 | 0.391 | 0.828 | 10458 | tags=22%, list=18%, signal=26% |
| 3328 | REGULATION\_OF\_LIPOPROTEIN\_PARTICLE\_CLEARANCE(GO:0010984) |  | 21 | 0.76 | 1.21 | 0.141 | 0.391 | 0.828 | 6197 | tags=10%, list=10%, signal=11% |
| 3329 | NEGATIVE\_REGULATION\_OF\_FATTY\_ACID\_BIOSYNTHETIC\_PROCESS(GO:0045717) |  | 16 | 0.73 | 1.21 | 0.000 | 0.391 | 0.828 | 13039 | tags=50%, list=22%, signal=64% |
| 3330 | REGULATION\_OF\_CAMP\_DEPENDENT\_PROTEIN\_KINASE\_ACTIVITY(GO:2000479) |  | 16 | 0.75 | 1.21 | 0.000 | 0.391 | 0.828 | 10468 | tags=31%, list=18%, signal=38% |
| 3331 | RNA\_3'\_END\_PROCESSING(GO:0031123) |  | 122 | 0.69 | 1.21 | 0.000 | 0.391 | 0.828 | 14342 | tags=40%, list=24%, signal=53% |
| 3332 | NEGATIVE\_REGULATION\_OF\_LIPASE\_ACTIVITY(GO:0060192) |  | 16 | 0.76 | 1.21 | 0.141 | 0.391 | 0.828 | 8252 | tags=31%, list=14%, signal=36% |
| 3333 | NEURON\_CELL\_CELL\_ADHESION(GO:0007158) |  | 15 | 0.84 | 1.21 | 0.141 | 0.391 | 0.828 | 7102 | tags=27%, list=12%, signal=30% |
| 3334 | ATPASE\_BINDING(GO:0051117) |  | 79 | 0.75 | 1.21 | 0.141 | 0.391 | 0.828 | 7164 | tags=8%, list=12%, signal=9% |
| 3335 | POSITIVE\_REGULATION\_OF\_ENDOPEPTIDASE\_ACTIVITY(GO:0010950) |  | 163 | 0.71 | 1.21 | 0.141 | 0.391 | 0.828 | 7754 | tags=9%, list=13%, signal=11% |
| 3336 | TYROSINE\_PHOSPHORYLATION\_OF\_STAT\_PROTEIN(GO:0007260) |  | 76 | 0.74 | 1.21 | 0.141 | 0.391 | 0.828 | 9454 | tags=25%, list=16%, signal=30% |
| 3337 | POSITIVE\_REGULATION\_OF\_PROTEIN\_ACETYLATION(GO:1901985) |  | 37 | 0.75 | 1.21 | 0.141 | 0.391 | 0.828 | 10856 | tags=27%, list=18%, signal=33% |
| 3338 | REGULATION\_OF\_NMDA\_RECEPTOR\_ACTIVITY(GO:2000310) |  | 19 | 0.78 | 1.21 | 0.000 | 0.391 | 0.828 | 7906 | tags=32%, list=13%, signal=36% |
| 3339 | ORGANELLE\_ORGANIZATION(GO:0006996) |  | 3750 | 0.76 | 1.21 | 0.141 | 0.391 | 0.828 | 8009 | tags=10%, list=14%, signal=11% |
| 3340 | PEROXISOMAL\_MEMBRANE(GO:0005778) |  | 57 | 0.71 | 1.21 | 0.000 | 0.391 | 0.828 | 12418 | tags=37%, list=21%, signal=47% |
| 3341 | MICROBODY\_MEMBRANE(GO:0031903) |  | 57 | 0.71 | 1.21 | 0.000 | 0.391 | 0.828 | 12418 | tags=37%, list=21%, signal=47% |
| 3342 | GLUTAMINE\_FAMILY\_AMINO\_ACID\_METABOLIC\_PROCESS(GO:0009064) |  | 76 | 0.71 | 1.21 | 0.000 | 0.391 | 0.828 | 12333 | tags=41%, list=21%, signal=51% |
| 3343 | REGULATION\_OF\_MYELINATION(GO:0031641) |  | 34 | 0.70 | 1.21 | 0.000 | 0.391 | 0.828 | 10295 | tags=29%, list=17%, signal=36% |
| 3344 | REGULATION\_OF\_CYCLIN\_DEPENDENT\_PROTEIN\_SERINE\_THREONINE\_KINASE\_ACTIVITY(GO:0000079) |  | 84 | 0.76 | 1.21 | 0.141 | 0.391 | 0.828 | 7092 | tags=7%, list=12%, signal=8% |
| 3345 | RESPIRATORY\_CHAIN(GO:0070469) |  | 103 | 0.69 | 1.21 | 0.000 | 0.391 | 0.828 | 5894 | tags=5%, list=10%, signal=5% |
| 3346 | PEPTIDE\_SECRETION(GO:0002790) |  | 558 | 0.72 | 1.21 | 0.141 | 0.391 | 0.828 | 10136 | tags=21%, list=17%, signal=25% |
| 3347 | RESPONSE\_TO\_LEUKEMIA\_INHIBITORY\_FACTOR(GO:1990823) |  | 94 | 0.74 | 1.21 | 0.141 | 0.391 | 0.828 | 8246 | tags=19%, list=14%, signal=22% |
| 3348 | CELLULAR\_RESPONSE\_TO\_LEUKEMIA\_INHIBITORY\_FACTOR(GO:1990830) |  | 94 | 0.74 | 1.21 | 0.141 | 0.391 | 0.828 | 8246 | tags=19%, list=14%, signal=22% |
| 3349 | CARBOHYDRATE\_TRANSMEMBRANE\_TRANSPORTER\_ACTIVITY(GO:0015144) |  | 44 | 0.71 | 1.21 | 0.141 | 0.390 | 0.828 | 9386 | tags=34%, list=16%, signal=40% |
| 3350 | NEGATIVE\_REGULATION\_OF\_NITRIC\_OXIDE\_BIOSYNTHETIC\_PROCESS(GO:0045019) |  | 16 | 0.77 | 1.21 | 0.000 | 0.391 | 0.828 | 10671 | tags=31%, list=18%, signal=38% |
| 3351 | NEGATIVE\_REGULATION\_OF\_NITRIC\_OXIDE\_METABOLIC\_PROCESS(GO:1904406) |  | 16 | 0.77 | 1.21 | 0.000 | 0.390 | 0.828 | 10671 | tags=31%, list=18%, signal=38% |
| 3352 | REGULATION\_OF\_CYCLIN\_DEPENDENT\_PROTEIN\_KINASE\_ACTIVITY(GO:1904029) |  | 90 | 0.76 | 1.21 | 0.141 | 0.390 | 0.828 | 10303 | tags=22%, list=17%, signal=27% |
| 3353 | NUCLEAR\_TRANSCRIPTION\_FACTOR\_COMPLEX(GO:0044798) |  | 165 | 0.72 | 1.21 | 0.000 | 0.390 | 0.828 | 7257 | tags=8%, list=12%, signal=9% |
| 3354 | PHOSPHORIC\_DIESTER\_HYDROLASE\_ACTIVITY(GO:0008081) |  | 94 | 0.70 | 1.21 | 0.000 | 0.390 | 0.828 | 12699 | tags=41%, list=21%, signal=53% |
| 3355 | POSITIVE\_REGULATION\_OF\_HORMONE\_SECRETION(GO:0046887) |  | 129 | 0.73 | 1.21 | 0.141 | 0.390 | 0.828 | 10131 | tags=23%, list=17%, signal=28% |
| 3356 | TRIGLYCERIDE\_RICH\_PLASMA\_LIPOPROTEIN\_PARTICLE(GO:0034385) |  | 20 | 0.73 | 1.21 | 0.141 | 0.390 | 0.828 | 6197 | tags=15%, list=10%, signal=17% |
| 3357 | VERY\_LOW\_DENSITY\_LIPOPROTEIN\_PARTICLE(GO:0034361) |  | 20 | 0.73 | 1.21 | 0.141 | 0.390 | 0.828 | 6197 | tags=15%, list=10%, signal=17% |
| 3358 | NUCLEOTIDE\_PHOSPHORYLATION(GO:0046939) |  | 126 | 0.73 | 1.21 | 0.141 | 0.390 | 0.828 | 8541 | tags=10%, list=14%, signal=12% |
| 3359 | CHONDROITIN\_SULFATE\_METABOLIC\_PROCESS(GO:0030204) |  | 41 | 0.71 | 1.21 | 0.000 | 0.390 | 0.828 | 9569 | tags=41%, list=16%, signal=49% |
| 3360 | POSITIVE\_REGULATION\_OF\_RAS\_PROTEIN\_SIGNAL\_TRANSDUCTION(GO:0046579) |  | 50 | 0.75 | 1.21 | 0.141 | 0.390 | 0.828 | 7906 | tags=30%, list=13%, signal=35% |
| 3361 | POSITIVE\_REGULATION\_OF\_KINASE\_ACTIVITY(GO:0033674) |  | 551 | 0.72 | 1.21 | 0.141 | 0.390 | 0.828 | 9289 | tags=17%, list=16%, signal=20% |
| 3362 | L\_ASCORBIC\_ACID\_BINDING(GO:0031418) |  | 20 | 0.76 | 1.21 | 0.141 | 0.390 | 1.000 | 11786 | tags=45%, list=20%, signal=56% |
| 3363 | REGULATION\_OF\_NITRIC\_OXIDE\_BIOSYNTHETIC\_PROCESS(GO:0045428) |  | 54 | 0.75 | 1.21 | 0.141 | 0.390 | 1.000 | 8755 | tags=20%, list=15%, signal=24% |
| 3364 | 9+2\_MOTILE\_CILIUM(GO:0097729) |  | 87 | 0.75 | 1.21 | 0.141 | 0.390 | 1.000 | 11087 | tags=40%, list=19%, signal=49% |
| 3365 | SPERM\_FLAGELLUM(GO:0036126) |  | 87 | 0.75 | 1.21 | 0.141 | 0.390 | 1.000 | 11087 | tags=40%, list=19%, signal=49% |
| 3366 | MICROTUBULE\_DEPOLYMERIZATION(GO:0007019) |  | 34 | 0.72 | 1.21 | 0.000 | 0.390 | 1.000 | 11641 | tags=47%, list=20%, signal=59% |
| 3367 | CELL\_SURFACE\_RECEPTOR\_SIGNALING\_PATHWAY\_INVOLVED\_IN\_HEART\_DEVELOPMENT(GO:0061311) |  | 28 | 0.78 | 1.21 | 0.141 | 0.390 | 1.000 | 8673 | tags=29%, list=15%, signal=33% |
| 3368 | NEGATIVE\_REGULATION\_OF\_LIPID\_CATABOLIC\_PROCESS(GO:0050995) |  | 22 | 0.73 | 1.21 | 0.141 | 0.390 | 1.000 | 11397 | tags=23%, list=19%, signal=28% |
| 3369 | REGULATION\_OF\_TYROSINE\_PHOSPHORYLATION\_OF\_STAT\_PROTEIN(GO:0042509) |  | 74 | 0.74 | 1.21 | 0.141 | 0.390 | 1.000 | 10643 | tags=26%, list=18%, signal=31% |
| 3370 | CELLULAR\_RESPONSE\_TO\_INORGANIC\_SUBSTANCE(GO:0071241) |  | 180 | 0.74 | 1.21 | 0.141 | 0.390 | 1.000 | 9405 | tags=19%, list=16%, signal=23% |
| 3371 | PYRIDINE\_NUCLEOTIDE\_METABOLIC\_PROCESS(GO:0019362) |  | 175 | 0.73 | 1.21 | 0.141 | 0.390 | 1.000 | 8602 | tags=11%, list=15%, signal=13% |
| 3372 | NICOTINAMIDE\_NUCLEOTIDE\_METABOLIC\_PROCESS(GO:0046496) |  | 175 | 0.73 | 1.21 | 0.141 | 0.390 | 1.000 | 8602 | tags=11%, list=15%, signal=13% |
| 3373 | RESPONSE\_TO\_ALKALOID(GO:0043279) |  | 108 | 0.76 | 1.21 | 0.141 | 0.390 | 1.000 | 7873 | tags=19%, list=13%, signal=21% |
| 3374 | METAL\_ION\_BINDING(GO:0046872) |  | 4142 | 0.76 | 1.21 | 0.141 | 0.390 | 1.000 | 8390 | tags=15%, list=14%, signal=16% |
| 3375 | CYCLIC\_PURINE\_NUCLEOTIDE\_METABOLIC\_PROCESS(GO:0052652) |  | 97 | 0.71 | 1.21 | 0.000 | 0.390 | 1.000 | 8279 | tags=22%, list=14%, signal=25% |
| 3376 | CELLULAR\_RESPONSE\_TO\_RADIATION(GO:0071478) |  | 174 | 0.75 | 1.20 | 0.141 | 0.390 | 1.000 | 8427 | tags=13%, list=14%, signal=15% |
| 3377 | ALPHA\_BETA\_T\_CELL\_DIFFERENTIATION(GO:0046632) |  | 92 | 0.73 | 1.20 | 0.141 | 0.390 | 1.000 | 9411 | tags=25%, list=16%, signal=30% |
| 3378 | MEMBRANE\_FUSION(GO:0061025) |  | 243 | 0.72 | 1.20 | 0.141 | 0.390 | 1.000 | 9144 | tags=15%, list=15%, signal=18% |
| 3379 | PEPTIDYL\_TYROSINE\_MODIFICATION(GO:0018212) |  | 381 | 0.73 | 1.20 | 0.141 | 0.390 | 1.000 | 9522 | tags=23%, list=16%, signal=27% |
| 3380 | REGULATION\_OF\_LEUKOCYTE\_ACTIVATION(GO:0002694) |  | 518 | 0.73 | 1.20 | 0.141 | 0.390 | 1.000 | 9198 | tags=19%, list=16%, signal=22% |
| 3381 | POST\_ANAL\_TAIL\_MORPHOGENESIS(GO:0036342) |  | 17 | 0.78 | 1.20 | 0.141 | 0.390 | 1.000 | 10461 | tags=53%, list=18%, signal=64% |
| 3382 | PROTEIN\_K63\_LINKED\_DEUBIQUITINATION(GO:0070536) |  | 29 | 0.71 | 1.20 | 0.000 | 0.390 | 1.000 | 8931 | tags=21%, list=15%, signal=24% |
| 3383 | INTERLEUKIN\_1\_BETA\_SECRETION(GO:0050702) |  | 45 | 0.75 | 1.20 | 0.141 | 0.390 | 1.000 | 6838 | tags=11%, list=12%, signal=13% |
| 3384 | REGULATION\_OF\_NEURON\_PROJECTION\_DEVELOPMENT(GO:0010975) |  | 435 | 0.75 | 1.20 | 0.141 | 0.390 | 1.000 | 9607 | tags=25%, list=16%, signal=30% |
| 3385 | EXTRACELLULAR\_LIGAND\_GATED\_ION\_CHANNEL\_ACTIVITY(GO:0005230) |  | 76 | 0.75 | 1.20 | 0.141 | 0.390 | 1.000 | 9607 | tags=36%, list=16%, signal=42% |
| 3386 | POSITIVE\_REGULATION\_OF\_OLIGODENDROCYTE\_DIFFERENTIATION(GO:0048714) |  | 17 | 0.78 | 1.20 | 0.141 | 0.390 | 1.000 | 9664 | tags=29%, list=16%, signal=35% |
| 3387 | NEGATIVE\_REGULATION\_OF\_PEPTIDE\_HORMONE\_SECRETION(GO:0090278) |  | 40 | 0.73 | 1.20 | 0.141 | 0.390 | 1.000 | 7544 | tags=15%, list=13%, signal=17% |
| 3388 | REGULATION\_OF\_NUCLEOBASE\_CONTAINING\_COMPOUND\_METABOLIC\_PROCESS(GO:0019219) |  | 4311 | 0.76 | 1.20 | 0.141 | 0.390 | 1.000 | 7957 | tags=13%, list=13%, signal=14% |
| 3389 | INTRACELLULAR\_RECEPTOR\_SIGNALING\_PATHWAY(GO:0030522) |  | 290 | 0.73 | 1.20 | 0.141 | 0.390 | 1.000 | 8112 | tags=14%, list=14%, signal=16% |
| 3390 | MONOCARBOXYLIC\_ACID\_BIOSYNTHETIC\_PROCESS(GO:0072330) |  | 296 | 0.73 | 1.20 | 0.141 | 0.390 | 1.000 | 8607 | tags=16%, list=15%, signal=18% |
| 3391 | CELLULAR\_RESPONSE\_TO\_DECREASED\_OXYGEN\_LEVELS(GO:0036294) |  | 209 | 0.71 | 1.20 | 0.141 | 0.390 | 1.000 | 7727 | tags=9%, list=13%, signal=10% |
| 3392 | INTERLEUKIN\_6\_BIOSYNTHETIC\_PROCESS(GO:0042226) |  | 17 | 0.74 | 1.20 | 0.000 | 0.389 | 1.000 | 6530 | tags=18%, list=11%, signal=20% |
| 3393 | REGULATION\_OF\_GLIOGENESIS(GO:0014013) |  | 104 | 0.75 | 1.20 | 0.141 | 0.389 | 1.000 | 7913 | tags=23%, list=13%, signal=27% |
| 3394 | POSITIVE\_REGULATION\_OF\_BLOOD\_CIRCULATION(GO:1903524) |  | 74 | 0.75 | 1.20 | 0.141 | 0.389 | 1.000 | 8882 | tags=31%, list=15%, signal=37% |
| 3395 | GLYCEROPHOSPHOLIPID\_METABOLIC\_PROCESS(GO:0006650) |  | 361 | 0.72 | 1.20 | 0.141 | 0.389 | 1.000 | 9669 | tags=20%, list=16%, signal=23% |
| 3396 | REGULATION\_OF\_INTERLEUKIN\_4\_PRODUCTION(GO:0032673) |  | 27 | 0.78 | 1.20 | 0.141 | 0.389 | 1.000 | 7488 | tags=30%, list=13%, signal=34% |
| 3397 | RUFFLE\_MEMBRANE(GO:0032587) |  | 88 | 0.73 | 1.20 | 0.141 | 0.389 | 1.000 | 8214 | tags=7%, list=14%, signal=8% |
| 3398 | MITOCHONDRION\_ORGANIZATION(GO:0007005) |  | 571 | 0.72 | 1.20 | 0.141 | 0.389 | 1.000 | 7782 | tags=6%, list=13%, signal=7% |
| 3399 | EPITHELIAL\_CELL\_CELL\_ADHESION(GO:0090136) |  | 16 | 0.78 | 1.20 | 0.141 | 0.389 | 1.000 | 10860 | tags=31%, list=18%, signal=38% |
| 3400 | FATTY\_ACYL\_COA\_METABOLIC\_PROCESS(GO:0035337) |  | 39 | 0.74 | 1.20 | 0.141 | 0.389 | 1.000 | 9812 | tags=18%, list=17%, signal=22% |
| 3401 | REACTIVE\_NITROGEN\_SPECIES\_METABOLIC\_PROCESS(GO:2001057) |  | 72 | 0.74 | 1.20 | 0.141 | 0.389 | 1.000 | 7873 | tags=18%, list=13%, signal=21% |
| 3402 | POSITIVE\_REGULATION\_OF\_LOCOMOTION(GO:0040017) |  | 518 | 0.74 | 1.20 | 0.141 | 0.389 | 1.000 | 8195 | tags=17%, list=14%, signal=20% |
| 3403 | POSITIVE\_REGULATION\_OF\_GLYCOGEN\_BIOSYNTHETIC\_PROCESS(GO:0045725) |  | 16 | 0.80 | 1.20 | 0.141 | 0.389 | 1.000 | 8594 | tags=25%, list=15%, signal=29% |
| 3404 | CATION\_BINDING(GO:0043169) |  | 4232 | 0.76 | 1.20 | 0.141 | 0.389 | 1.000 | 8390 | tags=15%, list=14%, signal=16% |
| 3405 | SMOOTH\_MUSCLE\_CONTRACTION(GO:0006939) |  | 108 | 0.73 | 1.20 | 0.141 | 0.389 | 1.000 | 9682 | tags=32%, list=16%, signal=39% |
| 3406 | EXTRINSIC\_COMPONENT\_OF\_MEMBRANE(GO:0019898) |  | 268 | 0.74 | 1.20 | 0.141 | 0.389 | 1.000 | 7951 | tags=12%, list=13%, signal=14% |
| 3407 | VITAMIN\_TRANSMEMBRANE\_TRANSPORTER\_ACTIVITY(GO:0090482) |  | 26 | 0.79 | 1.20 | 0.141 | 0.389 | 1.000 | 6269 | tags=19%, list=11%, signal=22% |
| 3408 | PROTEIN\_ACETYLATION(GO:0006473) |  | 191 | 0.71 | 1.20 | 0.000 | 0.389 | 1.000 | 10656 | tags=20%, list=18%, signal=25% |
| 3409 | NEGATIVE\_REGULATION\_OF\_INSULIN\_SECRETION(GO:0046676) |  | 35 | 0.74 | 1.20 | 0.141 | 0.389 | 1.000 | 7544 | tags=14%, list=13%, signal=16% |
| 3410 | POSITIVE\_REGULATION\_OF\_PROTEIN\_COMPLEX\_DISASSEMBLY(GO:0043243) |  | 35 | 0.76 | 1.20 | 0.141 | 0.389 | 1.000 | 8232 | tags=11%, list=14%, signal=13% |
| 3411 | POSITIVE\_REGULATION\_OF\_BIOMINERAL\_TISSUE\_DEVELOPMENT(GO:0070169) |  | 41 | 0.73 | 1.20 | 0.141 | 0.389 | 1.000 | 10980 | tags=37%, list=19%, signal=45% |
| 3412 | IRON\_ION\_TRANSMEMBRANE\_TRANSPORT(GO:0034755) |  | 16 | 0.75 | 1.20 | 0.000 | 0.389 | 1.000 | 6066 | tags=13%, list=10%, signal=14% |
| 3413 | MITOCHONDRIAL\_ATP\_SYNTHESIS\_COUPLED\_ELECTRON\_TRANSPORT(GO:0042775) |  | 98 | 0.69 | 1.20 | 0.000 | 0.389 | 1.000 | 5894 | tags=4%, list=10%, signal=5% |
| 3414 | OVULATION(GO:0030728) |  | 22 | 0.74 | 1.20 | 0.141 | 0.389 | 1.000 | 6424 | tags=18%, list=11%, signal=20% |
| 3415 | NEGATIVE\_REGULATION\_OF\_SMALL\_MOLECULE\_METABOLIC\_PROCESS(GO:0062014) |  | 121 | 0.74 | 1.20 | 0.141 | 0.389 | 1.000 | 8474 | tags=16%, list=14%, signal=18% |
| 3416 | ACYLGLYCEROL\_HOMEOSTASIS(GO:0055090) |  | 32 | 0.73 | 1.20 | 0.141 | 0.389 | 1.000 | 10458 | tags=25%, list=18%, signal=30% |
| 3417 | TRIGLYCERIDE\_HOMEOSTASIS(GO:0070328) |  | 32 | 0.73 | 1.20 | 0.141 | 0.389 | 1.000 | 10458 | tags=25%, list=18%, signal=30% |
| 3418 | GAMMA\_TUBULIN\_BINDING(GO:0043015) |  | 28 | 0.69 | 1.20 | 0.000 | 0.389 | 1.000 | 12528 | tags=50%, list=21%, signal=63% |
| 3419 | REGULATION\_OF\_PROTEIN\_SECRETION(GO:0050708) |  | 429 | 0.72 | 1.20 | 0.141 | 0.389 | 1.000 | 10786 | tags=23%, list=18%, signal=28% |
| 3420 | REGULATION\_OF\_MAST\_CELL\_ACTIVATION(GO:0033003) |  | 39 | 0.74 | 1.20 | 0.141 | 0.389 | 1.000 | 9198 | tags=28%, list=16%, signal=33% |
| 3421 | POSITIVE\_REGULATION\_OF\_OSSIFICATION(GO:0045778) |  | 86 | 0.74 | 1.20 | 0.141 | 0.389 | 1.000 | 8046 | tags=20%, list=14%, signal=23% |
| 3422 | ANCHORED\_COMPONENT\_OF\_MEMBRANE(GO:0031225) |  | 156 | 0.77 | 1.20 | 0.141 | 0.389 | 1.000 | 7975 | tags=22%, list=13%, signal=26% |
| 3423 | L\_GLUTAMATE\_TRANSMEMBRANE\_TRANSPORT(GO:0015813) |  | 22 | 0.74 | 1.20 | 0.141 | 0.389 | 1.000 | 7352 | tags=18%, list=12%, signal=21% |
| 3424 | EPITHELIAL\_TUBE\_BRANCHING\_INVOLVED\_IN\_LUNG\_MORPHOGENESIS(GO:0060441) |  | 29 | 0.80 | 1.20 | 0.141 | 0.389 | 1.000 | 6817 | tags=24%, list=12%, signal=27% |
| 3425 | POST\_GOLGI\_VESICLE\_MEDIATED\_TRANSPORT(GO:0006892) |  | 92 | 0.72 | 1.20 | 0.141 | 0.389 | 1.000 | 7457 | tags=4%, list=13%, signal=5% |
| 3426 | EXCRETION(GO:0007588) |  | 66 | 0.76 | 1.20 | 0.141 | 0.389 | 1.000 | 10857 | tags=35%, list=18%, signal=43% |
| 3427 | T\_HELPER\_17\_CELL\_DIFFERENTIATION(GO:0072539) |  | 20 | 0.77 | 1.20 | 0.141 | 0.389 | 1.000 | 9023 | tags=25%, list=15%, signal=29% |
| 3428 | MYOBLAST\_DIFFERENTIATION(GO:0045445) |  | 82 | 0.75 | 1.20 | 0.141 | 0.389 | 1.000 | 9633 | tags=26%, list=16%, signal=31% |
| 3429 | POSITIVE\_REGULATION\_OF\_NITRIC\_OXIDE\_METABOLIC\_PROCESS(GO:1904407) |  | 38 | 0.74 | 1.20 | 0.141 | 0.389 | 1.000 | 8755 | tags=21%, list=15%, signal=25% |
| 3430 | POSITIVE\_REGULATION\_OF\_NITRIC\_OXIDE\_BIOSYNTHETIC\_PROCESS(GO:0045429) |  | 38 | 0.74 | 1.20 | 0.141 | 0.389 | 1.000 | 8755 | tags=21%, list=15%, signal=25% |
| 3431 | SEXUAL\_REPRODUCTION(GO:0019953) |  | 816 | 0.73 | 1.20 | 0.141 | 0.389 | 1.000 | 8740 | tags=20%, list=15%, signal=23% |
| 3432 | ACTININ\_BINDING(GO:0042805) |  | 42 | 0.74 | 1.20 | 0.141 | 0.389 | 1.000 | 7652 | tags=24%, list=13%, signal=27% |
| 3433 | POSITIVE\_REGULATION\_OF\_CYCLIC\_NUCLEOTIDE\_BIOSYNTHETIC\_PROCESS(GO:0030804) |  | 40 | 0.71 | 1.20 | 0.000 | 0.389 | 1.000 | 8279 | tags=25%, list=14%, signal=29% |
| 3434 | REGULATION\_OF\_FIBROBLAST\_APOPTOTIC\_PROCESS(GO:2000269) |  | 17 | 0.76 | 1.20 | 0.141 | 0.389 | 1.000 | 6027 | tags=6%, list=10%, signal=7% |
| 3435 | REGULATION\_OF\_MAST\_CELL\_DEGRANULATION(GO:0043304) |  | 29 | 0.76 | 1.20 | 0.141 | 0.389 | 1.000 | 9198 | tags=24%, list=16%, signal=29% |
| 3436 | REGULATION\_OF\_MAST\_CELL\_ACTIVATION\_INVOLVED\_IN\_IMMUNE\_RESPONSE(GO:0033006) |  | 29 | 0.76 | 1.20 | 0.141 | 0.388 | 1.000 | 9198 | tags=24%, list=16%, signal=29% |
| 3437 | EXECUTION\_PHASE\_OF\_APOPTOSIS(GO:0097194) |  | 80 | 0.70 | 1.20 | 0.000 | 0.388 | 1.000 | 10939 | tags=25%, list=18%, signal=31% |
| 3438 | REGULATION\_OF\_CYCLIC\_NUCLEOTIDE\_BIOSYNTHETIC\_PROCESS(GO:0030802) |  | 73 | 0.71 | 1.20 | 0.000 | 0.388 | 1.000 | 8279 | tags=19%, list=14%, signal=22% |
| 3439 | CELLULAR\_RESPONSE\_TO\_FIBROBLAST\_GROWTH\_FACTOR\_STIMULUS(GO:0044344) |  | 145 | 0.74 | 1.20 | 0.141 | 0.388 | 1.000 | 8479 | tags=19%, list=14%, signal=22% |
| 3440 | CYCLIC\_NUCLEOTIDE\_METABOLIC\_PROCESS(GO:0009187) |  | 145 | 0.73 | 1.20 | 0.141 | 0.388 | 1.000 | 8279 | tags=25%, list=14%, signal=29% |
| 3441 | RESPONSE\_TO\_FIBROBLAST\_GROWTH\_FACTOR(GO:0071774) |  | 152 | 0.75 | 1.20 | 0.141 | 0.388 | 1.000 | 8479 | tags=20%, list=14%, signal=23% |
| 3442 | INNER\_EAR\_RECEPTOR\_CELL\_STEREOCILIUM\_ORGANIZATION(GO:0060122) |  | 31 | 0.73 | 1.20 | 0.000 | 0.388 | 1.000 | 9354 | tags=35%, list=16%, signal=42% |
| 3443 | NEGATIVE\_REGULATION\_OF\_GROWTH(GO:0045926) |  | 238 | 0.76 | 1.20 | 0.141 | 0.388 | 1.000 | 8744 | tags=21%, list=15%, signal=25% |
| 3444 | FEEDING\_BEHAVIOR(GO:0007631) |  | 105 | 0.75 | 1.20 | 0.141 | 0.388 | 1.000 | 8714 | tags=29%, list=15%, signal=33% |
| 3445 | NATURAL\_KILLER\_CELL\_ACTIVATION(GO:0030101) |  | 73 | 0.71 | 1.20 | 0.141 | 0.388 | 1.000 | 9266 | tags=26%, list=16%, signal=31% |
| 3446 | NEUTRAL\_AMINO\_ACID\_TRANSPORT(GO:0015804) |  | 39 | 0.74 | 1.20 | 0.000 | 0.388 | 1.000 | 8432 | tags=23%, list=14%, signal=27% |
| 3447 | RESPONSE\_TO\_PLATELET\_DERIVED\_GROWTH\_FACTOR(GO:0036119) |  | 22 | 0.70 | 1.20 | 0.000 | 0.388 | 1.000 | 12241 | tags=36%, list=21%, signal=46% |
| 3448 | POSITIVE\_REGULATION\_OF\_CELLULAR\_COMPONENT\_MOVEMENT(GO:0051272) |  | 502 | 0.74 | 1.20 | 0.141 | 0.388 | 1.000 | 8195 | tags=17%, list=14%, signal=19% |
| 3449 | POSITIVE\_REGULATION\_OF\_NEURON\_PROJECTION\_DEVELOPMENT(GO:0010976) |  | 253 | 0.74 | 1.20 | 0.141 | 0.388 | 1.000 | 9248 | tags=25%, list=16%, signal=29% |
| 3450 | UBIQUITIN\_PROTEIN\_TRANSFERASE\_ACTIVITY(GO:0004842) |  | 443 | 0.71 | 1.20 | 0.000 | 0.388 | 1.000 | 10958 | tags=21%, list=19%, signal=26% |
| 3451 | POSITIVE\_REGULATION\_OF\_LEUKOCYTE\_APOPTOTIC\_PROCESS(GO:2000108) |  | 29 | 0.70 | 1.20 | 0.000 | 0.388 | 1.000 | 13291 | tags=28%, list=22%, signal=36% |
| 3452 | T\_CELL\_COSTIMULATION(GO:0031295) |  | 68 | 0.73 | 1.20 | 0.141 | 0.388 | 1.000 | 10643 | tags=25%, list=18%, signal=30% |
| 3453 | POSITIVE\_REGULATION\_OF\_PEPTIDASE\_ACTIVITY(GO:0010952) |  | 180 | 0.70 | 1.20 | 0.141 | 0.388 | 1.000 | 8673 | tags=11%, list=15%, signal=13% |
| 3454 | T\_CELL\_CYTOKINE\_PRODUCTION(GO:0002369) |  | 34 | 0.72 | 1.20 | 0.141 | 0.388 | 1.000 | 7435 | tags=29%, list=13%, signal=34% |
| 3455 | POSITIVE\_REGULATION\_OF\_CD4\_POSITIVE\_\_ALPHA\_BETA\_T\_CELL\_ACTIVATION(GO:2000516) |  | 29 | 0.72 | 1.20 | 0.141 | 0.388 | 1.000 | 9368 | tags=38%, list=16%, signal=45% |
| 3456 | REGULATION\_OF\_ACTIN\_CYTOSKELETON\_ORGANIZATION(GO:0032956) |  | 305 | 0.74 | 1.20 | 0.141 | 0.388 | 1.000 | 8232 | tags=15%, list=14%, signal=17% |
| 3457 | REGULATION\_OF\_BLOOD\_PRESSURE(GO:0008217) |  | 181 | 0.74 | 1.20 | 0.141 | 0.388 | 1.000 | 10488 | tags=31%, list=18%, signal=37% |
| 3458 | SOLUTE:PROTON\_ANTIPORTER\_ACTIVITY(GO:0015299) |  | 19 | 0.75 | 1.20 | 0.141 | 0.388 | 1.000 | 10139 | tags=42%, list=17%, signal=51% |
| 3459 | NEUROPEPTIDE\_RECEPTOR\_ACTIVITY(GO:0008188) |  | 48 | 0.79 | 1.20 | 0.141 | 0.388 | 1.000 | 10588 | tags=33%, list=18%, signal=41% |
| 3460 | RNA\_POLYMERASE\_II\_TRANSCRIPTION\_COFACTOR\_ACTIVITY(GO:0001104) |  | 100 | 0.75 | 1.20 | 0.141 | 0.388 | 1.000 | 6559 | tags=12%, list=11%, signal=13% |
| 3461 | NEGATIVE\_REGULATION\_OF\_DNA\_METABOLIC\_PROCESS(GO:0051053) |  | 132 | 0.75 | 1.20 | 0.141 | 0.388 | 1.000 | 7945 | tags=9%, list=13%, signal=10% |
| 3462 | RECYCLING\_ENDOSOME(GO:0055037) |  | 158 | 0.75 | 1.20 | 0.141 | 0.388 | 1.000 | 7467 | tags=14%, list=13%, signal=16% |
| 3463 | TRANSCRIPTION\_INITIATION\_FROM\_RNA\_POLYMERASE\_II\_PROMOTER(GO:0006367) |  | 200 | 0.73 | 1.20 | 0.141 | 0.387 | 1.000 | 7414 | tags=12%, list=13%, signal=14% |
| 3464 | PLASMA\_LIPOPROTEIN\_PARTICLE(GO:0034358) |  | 40 | 0.72 | 1.20 | 0.141 | 0.387 | 1.000 | 6362 | tags=18%, list=11%, signal=20% |
| 3465 | LIPOPROTEIN\_PARTICLE(GO:1990777) |  | 40 | 0.72 | 1.20 | 0.141 | 0.387 | 1.000 | 6362 | tags=18%, list=11%, signal=20% |
| 3466 | GUANYLATE\_CYCLASE\_COMPLEX\_\_SOLUBLE(GO:0008074) |  | 17 | 0.78 | 1.20 | 0.000 | 0.387 | 1.000 | 7265 | tags=29%, list=12%, signal=34% |
| 3467 | GUANYLATE\_CYCLASE\_ACTIVITY(GO:0004383) |  | 17 | 0.78 | 1.20 | 0.000 | 0.387 | 1.000 | 7265 | tags=29%, list=12%, signal=34% |
| 3468 | NEGATIVE\_REGULATION\_OF\_AXONOGENESIS(GO:0050771) |  | 52 | 0.75 | 1.20 | 0.141 | 0.387 | 1.000 | 8673 | tags=31%, list=15%, signal=36% |
| 3469 | ORGANIC\_ACID\_BINDING(GO:0043177) |  | 192 | 0.74 | 1.20 | 0.141 | 0.387 | 1.000 | 8383 | tags=22%, list=14%, signal=25% |
| 3470 | INORGANIC\_ANION\_TRANSMEMBRANE\_TRANSPORT(GO:0098661) |  | 102 | 0.74 | 1.20 | 0.141 | 0.387 | 1.000 | 9293 | tags=25%, list=16%, signal=30% |
| 3471 | REGULATION\_OF\_LEUKOCYTE\_DIFFERENTIATION(GO:1902105) |  | 256 | 0.73 | 1.20 | 0.141 | 0.387 | 1.000 | 9411 | tags=21%, list=16%, signal=25% |
| 3472 | ENDOCYTOSIS(GO:0006897) |  | 744 | 0.73 | 1.20 | 0.141 | 0.387 | 1.000 | 8825 | tags=13%, list=15%, signal=16% |
| 3473 | ESTABLISHMENT\_OF\_PROTEIN\_LOCALIZATION\_TO\_ORGANELLE(GO:0072594) |  | 515 | 0.73 | 1.20 | 0.141 | 0.387 | 1.000 | 7426 | tags=6%, list=13%, signal=6% |
| 3474 | ANION:ANION\_ANTIPORTER\_ACTIVITY(GO:0015301) |  | 30 | 0.73 | 1.20 | 0.141 | 0.387 | 1.000 | 11402 | tags=43%, list=19%, signal=54% |
| 3475 | RIBOSOME(GO:0005840) |  | 255 | 0.70 | 1.20 | 0.000 | 0.387 | 1.000 | 6146 | tags=4%, list=10%, signal=4% |
| 3476 | CELLULAR\_RESPONSE\_TO\_OXYGEN\_LEVELS(GO:0071453) |  | 225 | 0.71 | 1.20 | 0.141 | 0.387 | 1.000 | 7727 | tags=9%, list=13%, signal=11% |
| 3477 | REPRODUCTIVE\_PROCESS(GO:0022414) |  | 1405 | 0.74 | 1.20 | 0.141 | 0.387 | 1.000 | 10839 | tags=27%, list=18%, signal=32% |
| 3478 | REPRODUCTION(GO:0000003) |  | 1406 | 0.74 | 1.20 | 0.141 | 0.387 | 1.000 | 10839 | tags=27%, list=18%, signal=32% |
| 3479 | REGULATION\_OF\_ENDOTHELIAL\_CELL\_MIGRATION(GO:0010594) |  | 208 | 0.74 | 1.20 | 0.141 | 0.387 | 1.000 | 9016 | tags=21%, list=15%, signal=25% |
| 3480 | DYNEIN\_LIGHT\_CHAIN\_BINDING(GO:0045503) |  | 24 | 0.73 | 1.20 | 0.000 | 0.387 | 1.000 | 8569 | tags=38%, list=14%, signal=44% |
| 3481 | SPINDLE\_ORGANIZATION(GO:0007051) |  | 158 | 0.72 | 1.20 | 0.141 | 0.387 | 1.000 | 11524 | tags=27%, list=19%, signal=33% |
| 3482 | CELLULAR\_RESPONSE\_TO\_VIRUS(GO:0098586) |  | 48 | 0.72 | 1.20 | 0.141 | 0.387 | 1.000 | 11567 | tags=27%, list=20%, signal=34% |
| 3483 | REGULATION\_OF\_SYSTEMIC\_ARTERIAL\_BLOOD\_PRESSURE\_BY\_CIRCULATORY\_RENIN\_ANGIOTENSIN(GO:0001991) |  | 18 | 0.72 | 1.20 | 0.000 | 0.387 | 1.000 | 13678 | tags=56%, list=23%, signal=72% |
| 3484 | TOR\_COMPLEX(GO:0038201) |  | 15 | 0.75 | 1.20 | 0.000 | 0.387 | 1.000 | 6359 | tags=13%, list=11%, signal=15% |
| 3485 | CELLULAR\_RESPONSE\_TO\_NUTRIENT(GO:0031670) |  | 61 | 0.70 | 1.20 | 0.141 | 0.387 | 1.000 | 11906 | tags=31%, list=20%, signal=39% |
| 3486 | SEX\_DIFFERENTIATION(GO:0007548) |  | 265 | 0.74 | 1.20 | 0.141 | 0.387 | 1.000 | 8613 | tags=22%, list=15%, signal=26% |
| 3487 | PROTEOLYSIS(GO:0006508) |  | 1852 | 0.73 | 1.20 | 0.141 | 0.387 | 1.000 | 8285 | tags=12%, list=14%, signal=13% |
| 3488 | BLOOD\_COAGULATION(GO:0007596) |  | 347 | 0.72 | 1.20 | 0.141 | 0.387 | 1.000 | 7921 | tags=15%, list=13%, signal=18% |
| 3489 | CARBOXYLIC\_ACID\_BINDING(GO:0031406) |  | 190 | 0.74 | 1.20 | 0.141 | 0.387 | 1.000 | 8383 | tags=22%, list=14%, signal=25% |
| 3490 | ATP\_BIOSYNTHETIC\_PROCESS(GO:0006754) |  | 145 | 0.73 | 1.20 | 0.141 | 0.387 | 1.000 | 8541 | tags=9%, list=14%, signal=10% |
| 3491 | TRANSMEMBRANE\_RECEPTOR\_PROTEIN\_TYROSINE\_PHOSPHATASE\_ACTIVITY(GO:0005001) |  | 15 | 0.76 | 1.20 | 0.141 | 0.387 | 1.000 | 7155 | tags=27%, list=12%, signal=30% |
| 3492 | TRANSMEMBRANE\_RECEPTOR\_PROTEIN\_PHOSPHATASE\_ACTIVITY(GO:0019198) |  | 15 | 0.76 | 1.20 | 0.141 | 0.386 | 1.000 | 7155 | tags=27%, list=12%, signal=30% |
| 3493 | GTPASE\_ACTIVITY(GO:0003924) |  | 290 | 0.74 | 1.20 | 0.141 | 0.386 | 1.000 | 8409 | tags=11%, list=14%, signal=13% |
| 3494 | CELLULAR\_RESPONSE\_TO\_NUTRIENT\_LEVELS(GO:0031669) |  | 207 | 0.71 | 1.20 | 0.141 | 0.386 | 1.000 | 12070 | tags=28%, list=20%, signal=35% |
| 3495 | CATION\_TRANSPORTING\_ATPASE\_COMPLEX(GO:0090533) |  | 15 | 0.77 | 1.20 | 0.000 | 0.386 | 1.000 | 6432 | tags=40%, list=11%, signal=45% |
| 3496 | REGULATION\_OF\_SENSORY\_PERCEPTION\_OF\_PAIN(GO:0051930) |  | 34 | 0.78 | 1.20 | 0.141 | 0.386 | 1.000 | 9682 | tags=32%, list=16%, signal=39% |
| 3497 | REGULATION\_OF\_SENSORY\_PERCEPTION(GO:0051931) |  | 34 | 0.78 | 1.20 | 0.141 | 0.386 | 1.000 | 9682 | tags=32%, list=16%, signal=39% |
| 3498 | VIRUS\_RECEPTOR\_ACTIVITY(GO:0001618) |  | 75 | 0.74 | 1.20 | 0.141 | 0.386 | 1.000 | 10550 | tags=27%, list=18%, signal=32% |
| 3499 | HIJACKED\_MOLECULAR\_FUNCTION(GO:0104005) |  | 75 | 0.74 | 1.20 | 0.141 | 0.386 | 1.000 | 10550 | tags=27%, list=18%, signal=32% |
| 3500 | ATP\_SYNTHESIS\_COUPLED\_ELECTRON\_TRANSPORT(GO:0042773) |  | 99 | 0.68 | 1.20 | 0.000 | 0.386 | 1.000 | 5894 | tags=4%, list=10%, signal=4% |
| 3501 | GTPASE\_ACTIVATOR\_ACTIVITY(GO:0005096) |  | 285 | 0.73 | 1.20 | 0.141 | 0.386 | 1.000 | 8030 | tags=13%, list=14%, signal=15% |
| 3502 | TRANSEPITHELIAL\_TRANSPORT(GO:0070633) |  | 18 | 0.75 | 1.20 | 0.141 | 0.386 | 1.000 | 11397 | tags=39%, list=19%, signal=48% |
| 3503 | DNA\_TEMPLATED\_TRANSCRIPTION\_\_INITIATION(GO:0006352) |  | 263 | 0.73 | 1.20 | 0.141 | 0.386 | 1.000 | 7414 | tags=10%, list=13%, signal=12% |
| 3504 | PROTEIN\_K48\_LINKED\_DEUBIQUITINATION(GO:0071108) |  | 30 | 0.70 | 1.20 | 0.000 | 0.386 | 1.000 | 11513 | tags=30%, list=19%, signal=37% |
| 3505 | BEHAVIORAL\_RESPONSE\_TO\_ETHANOL(GO:0048149) |  | 15 | 0.75 | 1.20 | 0.141 | 0.386 | 1.000 | 11807 | tags=40%, list=20%, signal=50% |
| 3506 | COAGULATION(GO:0050817) |  | 354 | 0.72 | 1.20 | 0.141 | 0.386 | 1.000 | 7921 | tags=15%, list=13%, signal=17% |
| 3507 | CILIARY\_BASAL\_BODY(GO:0036064) |  | 109 | 0.68 | 1.20 | 0.000 | 0.386 | 1.000 | 15119 | tags=52%, list=26%, signal=70% |
| 3508 | FIBROBLAST\_APOPTOTIC\_PROCESS(GO:0044346) |  | 20 | 0.74 | 1.20 | 0.141 | 0.386 | 1.000 | 6027 | tags=5%, list=10%, signal=6% |
| 3509 | PEPTIDYL\_AMINO\_ACID\_MODIFICATION(GO:0018193) |  | 1256 | 0.72 | 1.20 | 0.141 | 0.386 | 1.000 | 10483 | tags=21%, list=18%, signal=25% |
| 3510 | REGULATION\_OF\_PLASMA\_MEMBRANE\_BOUNDED\_CELL\_PROJECTION\_ORGANIZATION(GO:0120035) |  | 586 | 0.74 | 1.20 | 0.141 | 0.386 | 1.000 | 9316 | tags=23%, list=16%, signal=27% |
| 3511 | REGULATION\_OF\_CELL\_PROJECTION\_ORGANIZATION(GO:0031344) |  | 596 | 0.74 | 1.20 | 0.141 | 0.386 | 1.000 | 9316 | tags=23%, list=16%, signal=27% |
| 3512 | INTERLEUKIN\_1\_SECRETION(GO:0050701) |  | 51 | 0.73 | 1.20 | 0.141 | 0.386 | 1.000 | 6838 | tags=10%, list=12%, signal=11% |
| 3513 | REGULATION\_OF\_TRANSLATIONAL\_INITIATION(GO:0006446) |  | 76 | 0.71 | 1.20 | 0.000 | 0.386 | 1.000 | 5902 | tags=3%, list=10%, signal=3% |
| 3514 | UBIQUITIN\_LIKE\_PROTEIN\_TRANSFERASE\_ACTIVITY(GO:0019787) |  | 465 | 0.70 | 1.20 | 0.000 | 0.386 | 1.000 | 11101 | tags=22%, list=19%, signal=26% |
| 3515 | NUCLEAR\_SPECK(GO:0016607) |  | 385 | 0.70 | 1.20 | 0.000 | 0.386 | 1.000 | 8237 | tags=10%, list=14%, signal=12% |
| 3516 | POSITIVE\_REGULATION\_OF\_GLYCOGEN\_METABOLIC\_PROCESS(GO:0070875) |  | 18 | 0.78 | 1.20 | 0.141 | 0.386 | 1.000 | 8594 | tags=22%, list=15%, signal=26% |
| 3517 | REGULATION\_OF\_RECEPTOR\_INTERNALIZATION(GO:0002090) |  | 39 | 0.74 | 1.20 | 0.141 | 0.386 | 1.000 | 9520 | tags=28%, list=16%, signal=34% |
| 3518 | LIPID\_CATABOLIC\_PROCESS(GO:0016042) |  | 311 | 0.71 | 1.20 | 0.141 | 0.386 | 1.000 | 8216 | tags=13%, list=14%, signal=14% |
| 3519 | SNARE\_COMPLEX\_ASSEMBLY(GO:0035493) |  | 17 | 0.70 | 1.20 | 0.000 | 0.386 | 1.000 | 7457 | tags=12%, list=13%, signal=13% |
| 3520 | ORGANOPHOSPHATE\_BIOSYNTHETIC\_PROCESS(GO:0090407) |  | 697 | 0.71 | 1.20 | 0.141 | 0.386 | 1.000 | 10458 | tags=18%, list=18%, signal=22% |
| 3521 | HEAT\_SHOCK\_PROTEIN\_BINDING(GO:0031072) |  | 109 | 0.70 | 1.20 | 0.000 | 0.386 | 1.000 | 11054 | tags=23%, list=19%, signal=28% |
| 3522 | NEGATIVE\_REGULATION\_OF\_LEUKOCYTE\_MEDIATED\_IMMUNITY(GO:0002704) |  | 43 | 0.73 | 1.20 | 0.141 | 0.386 | 1.000 | 10239 | tags=21%, list=17%, signal=25% |
| 3523 | ORGANIC\_CATION\_TRANSPORT(GO:0015695) |  | 40 | 0.75 | 1.19 | 0.141 | 0.386 | 1.000 | 7809 | tags=15%, list=13%, signal=17% |
| 3524 | NEGATIVE\_REGULATION\_OF\_KIDNEY\_DEVELOPMENT(GO:0090185) |  | 17 | 0.81 | 1.19 | 0.141 | 0.386 | 1.000 | 9208 | tags=29%, list=16%, signal=35% |
| 3525 | PROTEIN\_PROCESSING(GO:0016485) |  | 314 | 0.72 | 1.19 | 0.141 | 0.386 | 1.000 | 8263 | tags=17%, list=14%, signal=20% |
| 3526 | HEMOSTASIS(GO:0007599) |  | 352 | 0.72 | 1.19 | 0.141 | 0.386 | 1.000 | 7921 | tags=15%, list=13%, signal=18% |
| 3527 | EARLY\_ENDOSOME\_MEMBRANE(GO:0031901) |  | 134 | 0.71 | 1.19 | 0.141 | 0.386 | 1.000 | 10713 | tags=25%, list=18%, signal=31% |
| 3528 | REPRODUCTIVE\_STRUCTURE\_DEVELOPMENT(GO:0048608) |  | 428 | 0.74 | 1.19 | 0.141 | 0.386 | 1.000 | 8490 | tags=19%, list=14%, signal=22% |
| 3529 | POLYMERIC\_CYTOSKELETAL\_FIBER(GO:0099513) |  | 693 | 0.74 | 1.19 | 0.141 | 0.386 | 1.000 | 9344 | tags=18%, list=16%, signal=22% |
| 3530 | COMPLEMENT\_BINDING(GO:0001848) |  | 22 | 0.76 | 1.19 | 0.141 | 0.386 | 1.000 | 10025 | tags=27%, list=17%, signal=33% |
| 3531 | NEGATIVE\_REGULATION\_OF\_REACTIVE\_OXYGEN\_SPECIES\_BIOSYNTHETIC\_PROCESS(GO:1903427) |  | 28 | 0.71 | 1.19 | 0.000 | 0.386 | 1.000 | 12925 | tags=32%, list=22%, signal=41% |
| 3532 | CONDENSED\_NUCLEAR\_CHROMOSOME\_\_CENTROMERIC\_REGION(GO:0000780) |  | 21 | 0.71 | 1.19 | 0.000 | 0.386 | 1.000 | 15072 | tags=62%, list=25%, signal=83% |
| 3533 | STRUCTURAL\_CONSTITUENT\_OF\_EYE\_LENS(GO:0005212) |  | 21 | 0.72 | 1.19 | 0.000 | 0.385 | 1.000 | 14138 | tags=52%, list=24%, signal=69% |
| 3534 | DOUBLE\_STRAND\_BREAK\_REPAIR(GO:0006302) |  | 217 | 0.75 | 1.19 | 0.141 | 0.385 | 1.000 | 8385 | tags=11%, list=14%, signal=12% |
| 3535 | ADHERENS\_JUNCTION\_ORGANIZATION(GO:0034332) |  | 124 | 0.75 | 1.19 | 0.141 | 0.385 | 1.000 | 7136 | tags=10%, list=12%, signal=11% |
| 3536 | REGULATION\_OF\_PEPTIDE\_SECRETION(GO:0002791) |  | 458 | 0.71 | 1.19 | 0.141 | 0.386 | 1.000 | 10786 | tags=23%, list=18%, signal=28% |
| 3537 | PLATELET\_ALPHA\_GRANULE\_LUMEN(GO:0031093) |  | 68 | 0.74 | 1.19 | 0.141 | 0.386 | 1.000 | 7238 | tags=15%, list=12%, signal=17% |
| 3538 | PROTEIN\_SUMOYLATION(GO:0016925) |  | 94 | 0.71 | 1.19 | 0.000 | 0.386 | 1.000 | 6154 | tags=6%, list=10%, signal=7% |
| 3539 | ENDOPLASMIC\_RETICULUM\_TO\_CYTOSOL\_TRANSPORT(GO:1903513) |  | 28 | 0.71 | 1.19 | 0.000 | 0.386 | 1.000 | 5999 | tags=4%, list=10%, signal=4% |
| 3540 | RETROGRADE\_PROTEIN\_TRANSPORT\_\_ER\_TO\_CYTOSOL(GO:0030970) |  | 28 | 0.71 | 1.19 | 0.000 | 0.386 | 1.000 | 5999 | tags=4%, list=10%, signal=4% |
| 3541 | POSITIVE\_REGULATION\_OF\_LEUKOCYTE\_MEDIATED\_CYTOTOXICITY(GO:0001912) |  | 33 | 0.69 | 1.19 | 0.000 | 0.385 | 1.000 | 14247 | tags=55%, list=24%, signal=72% |
| 3542 | NEGATIVE\_REGULATION\_OF\_MICROTUBULE\_DEPOLYMERIZATION(GO:0007026) |  | 19 | 0.75 | 1.19 | 0.141 | 0.385 | 1.000 | 11641 | tags=37%, list=20%, signal=46% |
| 3543 | NEGATIVE\_REGULATION\_OF\_DNA\_RECOMBINATION(GO:0045910) |  | 24 | 0.78 | 1.19 | 0.141 | 0.385 | 1.000 | 6893 | tags=8%, list=12%, signal=9% |
| 3544 | COMPLEMENT\_ACTIVATION(GO:0006956) |  | 166 | 0.70 | 1.19 | 0.141 | 0.385 | 1.000 | 8285 | tags=11%, list=14%, signal=13% |
| 3545 | NEURAL\_TUBE\_FORMATION(GO:0001841) |  | 106 | 0.75 | 1.19 | 0.141 | 0.385 | 1.000 | 7975 | tags=16%, list=13%, signal=19% |
| 3546 | GLUTAMATE\_RECEPTOR\_ACTIVITY(GO:0008066) |  | 27 | 0.78 | 1.19 | 0.141 | 0.385 | 1.000 | 9000 | tags=37%, list=15%, signal=44% |
| 3547 | POSITIVE\_REGULATION\_OF\_PEPTIDYL\_THREONINE\_PHOSPHORYLATION(GO:0010800) |  | 30 | 0.70 | 1.19 | 0.000 | 0.385 | 1.000 | 7233 | tags=13%, list=12%, signal=15% |
| 3548 | ANIMAL\_ORGAN\_MATURATION(GO:0048799) |  | 23 | 0.78 | 1.19 | 0.141 | 0.385 | 1.000 | 10811 | tags=39%, list=18%, signal=48% |
| 3549 | ADENYLATE\_CYCLASE\_ACTIVATING\_G\_PROTEIN\_COUPLED\_RECEPTOR\_SIGNALING\_PATHWAY(GO:0007189) |  | 92 | 0.74 | 1.19 | 0.141 | 0.385 | 1.000 | 10901 | tags=24%, list=18%, signal=29% |
| 3550 | MALE\_GONAD\_DEVELOPMENT(GO:0008584) |  | 136 | 0.74 | 1.19 | 0.141 | 0.385 | 1.000 | 8613 | tags=20%, list=15%, signal=23% |
| 3551 | DNA\_CATABOLIC\_PROCESS\_\_ENDONUCLEOLYTIC(GO:0000737) |  | 26 | 0.69 | 1.19 | 0.000 | 0.385 | 1.000 | 14434 | tags=54%, list=24%, signal=71% |
| 3552 | INTRACELLULAR\_NON\_MEMBRANE\_BOUNDED\_ORGANELLE(GO:0043232) |  | 4113 | 0.76 | 1.19 | 0.141 | 0.385 | 1.000 | 8092 | tags=11%, list=14%, signal=12% |
| 3553 | NON\_MEMBRANE\_BOUNDED\_ORGANELLE(GO:0043228) |  | 4113 | 0.76 | 1.19 | 0.141 | 0.385 | 1.000 | 8092 | tags=11%, list=14%, signal=12% |
| 3554 | REGULATION\_OF\_CIRCADIAN\_SLEEP\_WAKE\_CYCLE(GO:0042749) |  | 21 | 0.73 | 1.19 | 0.000 | 0.385 | 1.000 | 6525 | tags=33%, list=11%, signal=37% |
| 3555 | BONE\_MATURATION(GO:0070977) |  | 19 | 0.74 | 1.19 | 0.000 | 0.385 | 1.000 | 10811 | tags=37%, list=18%, signal=45% |
| 3556 | CARBON\_NITROGEN\_LYASE\_ACTIVITY(GO:0016840) |  | 15 | 0.71 | 1.19 | 0.000 | 0.385 | 1.000 | 10786 | tags=27%, list=18%, signal=33% |
| 3557 | CIRCADIAN\_SLEEP\_WAKE\_CYCLE\_PROCESS(GO:0022410) |  | 25 | 0.73 | 1.19 | 0.000 | 0.385 | 1.000 | 7172 | tags=32%, list=12%, signal=36% |
| 3558 | RESPONSE\_TO\_ACIDIC\_PH(GO:0010447) |  | 22 | 0.72 | 1.19 | 0.000 | 0.385 | 1.000 | 11211 | tags=45%, list=19%, signal=56% |
| 3559 | REGULATION\_OF\_CIRCADIAN\_SLEEP\_WAKE\_CYCLE\_\_SLEEP(GO:0045187) |  | 19 | 0.73 | 1.19 | 0.000 | 0.385 | 1.000 | 6525 | tags=32%, list=11%, signal=35% |
| 3560 | REGULATION\_OF\_PROTEIN\_EXIT\_FROM\_ENDOPLASMIC\_RETICULUM(GO:0070861) |  | 22 | 0.79 | 1.19 | 0.141 | 0.385 | 1.000 | 6179 | tags=14%, list=10%, signal=15% |
| 3561 | ALPHA\_BETA\_T\_CELL\_ACTIVATION(GO:0046631) |  | 116 | 0.72 | 1.19 | 0.141 | 0.385 | 1.000 | 9411 | tags=23%, list=16%, signal=28% |
| 3562 | REPRODUCTIVE\_SYSTEM\_DEVELOPMENT(GO:0061458) |  | 432 | 0.74 | 1.19 | 0.141 | 0.385 | 1.000 | 8490 | tags=19%, list=14%, signal=22% |
| 3563 | HEPARAN\_SULFATE\_PROTEOGLYCAN\_BIOSYNTHETIC\_PROCESS(GO:0015012) |  | 24 | 0.73 | 1.19 | 0.000 | 0.385 | 1.000 | 12257 | tags=46%, list=21%, signal=58% |
| 3564 | CELL\_MATURATION(GO:0048469) |  | 165 | 0.72 | 1.19 | 0.141 | 0.385 | 1.000 | 10295 | tags=26%, list=17%, signal=31% |
| 3565 | ESTABLISHMENT\_OF\_SYNAPTIC\_VESICLE\_LOCALIZATION(GO:0097480) |  | 109 | 0.73 | 1.19 | 0.141 | 0.385 | 1.000 | 9227 | tags=24%, list=16%, signal=28% |
| 3566 | SYNAPTIC\_VESICLE\_TRANSPORT(GO:0048489) |  | 109 | 0.73 | 1.19 | 0.141 | 0.385 | 1.000 | 9227 | tags=24%, list=16%, signal=28% |
| 3567 | POSITIVE\_REGULATION\_OF\_CYTOKINE\_PRODUCTION\_INVOLVED\_IN\_INFLAMMATORY\_RESPONSE(GO:1900017) |  | 16 | 0.75 | 1.19 | 0.141 | 0.385 | 1.000 | 6413 | tags=19%, list=11%, signal=21% |
| 3568 | CERAMIDE\_BIOSYNTHETIC\_PROCESS(GO:0046513) |  | 54 | 0.72 | 1.19 | 0.141 | 0.385 | 1.000 | 8876 | tags=22%, list=15%, signal=26% |
| 3569 | REGULATION\_OF\_CYTOLYSIS(GO:0042268) |  | 16 | 0.75 | 1.19 | 0.000 | 0.385 | 1.000 | 11397 | tags=31%, list=19%, signal=39% |
| 3570 | BOUNDING\_MEMBRANE\_OF\_ORGANELLE(GO:0098588) |  | 1998 | 0.73 | 1.19 | 0.141 | 0.384 | 1.000 | 7940 | tags=12%, list=13%, signal=13% |
| 3571 | LYMPHOCYTE\_COSTIMULATION(GO:0031294) |  | 69 | 0.72 | 1.19 | 0.141 | 0.384 | 1.000 | 10643 | tags=25%, list=18%, signal=30% |
| 3572 | PHOSPHATIDYLINOSITOL\_4\_5\_BISPHOSPHATE\_3\_KINASE\_ACTIVITY(GO:0046934) |  | 68 | 0.74 | 1.19 | 0.141 | 0.384 | 1.000 | 7656 | tags=32%, list=13%, signal=37% |
| 3573 | REGULATION\_OF\_GLUCOSE\_IMPORT(GO:0046324) |  | 68 | 0.68 | 1.19 | 0.141 | 0.384 | 1.000 | 10571 | tags=18%, list=18%, signal=21% |
| 3574 | REGULATION\_OF\_FIBROBLAST\_MIGRATION(GO:0010762) |  | 32 | 0.76 | 1.19 | 0.141 | 0.384 | 1.000 | 8112 | tags=22%, list=14%, signal=25% |
| 3575 | EMBRYONIC\_DIGESTIVE\_TRACT\_MORPHOGENESIS(GO:0048557) |  | 19 | 0.81 | 1.19 | 0.141 | 0.384 | 1.000 | 8107 | tags=42%, list=14%, signal=49% |
| 3576 | POSITIVE\_REGULATION\_OF\_CELL\_MIGRATION(GO:0030335) |  | 472 | 0.73 | 1.19 | 0.141 | 0.384 | 1.000 | 8195 | tags=17%, list=14%, signal=20% |
| 3577 | EXTERNAL\_SIDE\_OF\_PLASMA\_MEMBRANE(GO:0009897) |  | 269 | 0.71 | 1.19 | 0.141 | 0.384 | 1.000 | 9121 | tags=20%, list=15%, signal=24% |
| 3578 | CIRCADIAN\_RHYTHM(GO:0007623) |  | 196 | 0.73 | 1.19 | 0.141 | 0.384 | 1.000 | 7659 | tags=13%, list=13%, signal=15% |
| 3579 | REGULATION\_OF\_TYPE\_2\_IMMUNE\_RESPONSE(GO:0002828) |  | 30 | 0.75 | 1.19 | 0.141 | 0.384 | 1.000 | 7435 | tags=27%, list=13%, signal=30% |
| 3580 | G\_PROTEIN\_ALPHA\_SUBUNIT\_BINDING(GO:0001965) |  | 19 | 0.71 | 1.19 | 0.000 | 0.384 | 1.000 | 10220 | tags=42%, list=17%, signal=51% |
| 3581 | RHO\_PROTEIN\_SIGNAL\_TRANSDUCTION(GO:0007266) |  | 190 | 0.73 | 1.19 | 0.141 | 0.384 | 1.000 | 9584 | tags=19%, list=16%, signal=23% |
| 3582 | DEVELOPMENT\_OF\_PRIMARY\_MALE\_SEXUAL\_CHARACTERISTICS(GO:0046546) |  | 137 | 0.74 | 1.19 | 0.141 | 0.384 | 1.000 | 8613 | tags=20%, list=15%, signal=23% |
| 3583 | POSITIVE\_REGULATION\_OF\_COLLAGEN\_BIOSYNTHETIC\_PROCESS(GO:0032967) |  | 24 | 0.73 | 1.19 | 0.141 | 0.384 | 1.000 | 11254 | tags=33%, list=19%, signal=41% |
| 3584 | POSITIVE\_REGULATION\_OF\_COLLAGEN\_METABOLIC\_PROCESS(GO:0010714) |  | 24 | 0.73 | 1.19 | 0.141 | 0.384 | 1.000 | 11254 | tags=33%, list=19%, signal=41% |
| 3585 | SERINE\_TYPE\_ENDOPEPTIDASE\_ACTIVITY(GO:0004252) |  | 253 | 0.76 | 1.19 | 0.141 | 0.384 | 1.000 | 8392 | tags=17%, list=14%, signal=19% |
| 3586 | PEPTIDYL\_PROLINE\_MODIFICATION(GO:0018208) |  | 59 | 0.69 | 1.19 | 0.000 | 0.384 | 1.000 | 11786 | tags=36%, list=20%, signal=44% |
| 3587 | ALPHA\_AMINO\_ACID\_CATABOLIC\_PROCESS(GO:1901606) |  | 103 | 0.74 | 1.19 | 0.141 | 0.384 | 1.000 | 10534 | tags=26%, list=18%, signal=32% |
| 3588 | SCAVENGER\_RECEPTOR\_ACTIVITY(GO:0005044) |  | 53 | 0.74 | 1.19 | 0.141 | 0.384 | 1.000 | 11589 | tags=32%, list=20%, signal=40% |
| 3589 | ORGAN\_INDUCTION(GO:0001759) |  | 19 | 0.78 | 1.19 | 0.141 | 0.384 | 1.000 | 10839 | tags=42%, list=18%, signal=52% |
| 3590 | PROTEIN\_POLYUBIQUITINATION(GO:0000209) |  | 295 | 0.71 | 1.19 | 0.141 | 0.384 | 1.000 | 7795 | tags=6%, list=13%, signal=7% |
| 3591 | GLUCOSE\_TRANSMEMBRANE\_TRANSPORT(GO:1904659) |  | 111 | 0.70 | 1.19 | 0.141 | 0.384 | 1.000 | 9042 | tags=22%, list=15%, signal=25% |
| 3592 | CHROMATIN\_REMODELING(GO:0006338) |  | 165 | 0.74 | 1.19 | 0.141 | 0.384 | 1.000 | 7522 | tags=7%, list=13%, signal=8% |
| 3593 | CELLULAR\_RESPONSE\_TO\_PLATELET\_DERIVED\_GROWTH\_FACTOR\_STIMULUS(GO:0036120) |  | 21 | 0.69 | 1.19 | 0.000 | 0.384 | 1.000 | 12241 | tags=33%, list=21%, signal=42% |
| 3594 | POSITIVE\_REGULATION\_OF\_BONE\_MINERALIZATION(GO:0030501) |  | 36 | 0.73 | 1.19 | 0.141 | 0.383 | 1.000 | 10861 | tags=39%, list=18%, signal=48% |
| 3595 | REGULATION\_OF\_ALTERNATIVE\_MRNA\_SPLICING\_\_VIA\_SPLICEOSOME(GO:0000381) |  | 42 | 0.71 | 1.19 | 0.000 | 0.383 | 1.000 | 13512 | tags=55%, list=23%, signal=71% |
| 3596 | MONOVALENT\_INORGANIC\_CATION\_HOMEOSTASIS(GO:0055067) |  | 148 | 0.73 | 1.19 | 0.141 | 0.383 | 1.000 | 9957 | tags=26%, list=17%, signal=31% |
| 3597 | REGULATION\_OF\_MYELOID\_LEUKOCYTE\_MEDIATED\_IMMUNITY(GO:0002886) |  | 44 | 0.74 | 1.19 | 0.141 | 0.383 | 1.000 | 9198 | tags=23%, list=16%, signal=27% |
| 3598 | MEMBRANE\_LIPID\_METABOLIC\_PROCESS(GO:0006643) |  | 218 | 0.72 | 1.19 | 0.141 | 0.383 | 1.000 | 8560 | tags=17%, list=14%, signal=19% |
| 3599 | TUBE\_CLOSURE(GO:0060606) |  | 92 | 0.75 | 1.19 | 0.141 | 0.383 | 1.000 | 7975 | tags=16%, list=13%, signal=19% |
| 3600 | ANCHORED\_COMPONENT\_OF\_PLASMA\_MEMBRANE(GO:0046658) |  | 46 | 0.78 | 1.19 | 0.141 | 0.383 | 1.000 | 7658 | tags=24%, list=13%, signal=27% |
| 3601 | CARDIAC\_MUSCLE\_HYPERTROPHY(GO:0003300) |  | 96 | 0.70 | 1.19 | 0.000 | 0.383 | 1.000 | 13771 | tags=33%, list=23%, signal=43% |
| 3602 | NEURAL\_TUBE\_CLOSURE(GO:0001843) |  | 91 | 0.75 | 1.19 | 0.141 | 0.383 | 1.000 | 7975 | tags=16%, list=13%, signal=19% |
| 3603 | GROWTH\_CONE(GO:0030426) |  | 157 | 0.75 | 1.19 | 0.141 | 0.383 | 1.000 | 10556 | tags=29%, list=18%, signal=36% |
| 3604 | NEGATIVE\_REGULATION\_OF\_PHOSPHOPROTEIN\_PHOSPHATASE\_ACTIVITY(GO:0032515) |  | 53 | 0.70 | 1.19 | 0.000 | 0.383 | 1.000 | 7544 | tags=13%, list=13%, signal=15% |
| 3605 | TYPE\_B\_PANCREATIC\_CELL\_PROLIFERATION(GO:0044342) |  | 19 | 0.77 | 1.19 | 0.141 | 0.383 | 1.000 | 10505 | tags=26%, list=18%, signal=32% |
| 3606 | PRIMARY\_NEURAL\_TUBE\_FORMATION(GO:0014020) |  | 98 | 0.75 | 1.19 | 0.141 | 0.383 | 1.000 | 7975 | tags=16%, list=13%, signal=19% |
| 3607 | CARBOHYDRATE\_TRANSMEMBRANE\_TRANSPORT(GO:0034219) |  | 122 | 0.70 | 1.19 | 0.141 | 0.383 | 1.000 | 9571 | tags=23%, list=16%, signal=27% |
| 3608 | POSITIVE\_REGULATION\_OF\_MITOTIC\_CELL\_CYCLE(GO:0045931) |  | 142 | 0.72 | 1.19 | 0.141 | 0.383 | 1.000 | 11362 | tags=24%, list=19%, signal=30% |
| 3609 | NUCLEOSIDE\_TRIPHOSPHATASE\_REGULATOR\_ACTIVITY(GO:0060589) |  | 350 | 0.72 | 1.19 | 0.141 | 0.383 | 1.000 | 10220 | tags=19%, list=17%, signal=23% |
| 3610 | REGULATION\_OF\_AXONOGENESIS(GO:0050770) |  | 157 | 0.74 | 1.19 | 0.141 | 0.383 | 1.000 | 8673 | tags=22%, list=15%, signal=26% |
| 3611 | LENS\_MORPHOGENESIS\_IN\_CAMERA\_TYPE\_EYE(GO:0002089) |  | 21 | 0.79 | 1.19 | 0.141 | 0.383 | 1.000 | 8380 | tags=29%, list=14%, signal=33% |
| 3612 | REGULATION\_OF\_T\_CELL\_CYTOKINE\_PRODUCTION(GO:0002724) |  | 23 | 0.73 | 1.19 | 0.141 | 0.383 | 1.000 | 10014 | tags=26%, list=17%, signal=31% |
| 3613 | POSITIVE\_REGULATION\_OF\_PEPTIDE\_HORMONE\_SECRETION(GO:0090277) |  | 95 | 0.72 | 1.19 | 0.141 | 0.383 | 1.000 | 10131 | tags=24%, list=17%, signal=29% |
| 3614 | CYSTEINE\_TYPE\_ENDOPEPTIDASE\_ACTIVITY(GO:0004197) |  | 87 | 0.69 | 1.19 | 0.000 | 0.383 | 1.000 | 13863 | tags=37%, list=23%, signal=48% |
| 3615 | REGULATION\_OF\_B\_CELL\_DIFFERENTIATION(GO:0045577) |  | 26 | 0.74 | 1.19 | 0.141 | 0.383 | 1.000 | 9368 | tags=19%, list=16%, signal=23% |
| 3616 | RESPONSE\_TO\_NERVE\_GROWTH\_FACTOR(GO:1990089) |  | 49 | 0.69 | 1.19 | 0.000 | 0.383 | 1.000 | 8937 | tags=16%, list=15%, signal=19% |
| 3617 | POSITIVE\_REGULATION\_OF\_LEUKOCYTE\_DIFFERENTIATION(GO:1902107) |  | 134 | 0.72 | 1.19 | 0.141 | 0.383 | 1.000 | 9368 | tags=25%, list=16%, signal=30% |
| 3618 | POSITIVE\_REGULATION\_OF\_PROTEIN\_DEPHOSPHORYLATION(GO:0035307) |  | 37 | 0.70 | 1.19 | 0.000 | 0.383 | 1.000 | 9543 | tags=16%, list=16%, signal=19% |
| 3619 | POSITIVE\_REGULATION\_OF\_MEIOTIC\_CELL\_CYCLE(GO:0051446) |  | 17 | 0.77 | 1.19 | 0.141 | 0.383 | 1.000 | 10543 | tags=41%, list=18%, signal=50% |
| 3620 | NAD+\_KINASE\_ACTIVITY(GO:0003951) |  | 17 | 0.70 | 1.19 | 0.000 | 0.383 | 1.000 | 8235 | tags=29%, list=14%, signal=34% |
| 3621 | REGULATION\_OF\_DEPHOSPHORYLATION(GO:0035303) |  | 196 | 0.70 | 1.19 | 0.141 | 0.383 | 1.000 | 10905 | tags=20%, list=18%, signal=24% |
| 3622 | REGULATION\_OF\_REPRODUCTIVE\_PROCESS(GO:2000241) |  | 141 | 0.73 | 1.19 | 0.141 | 0.383 | 1.000 | 9128 | tags=27%, list=15%, signal=32% |
| 3623 | NEGATIVE\_REGULATION\_OF\_CELL\_GROWTH(GO:0030308) |  | 160 | 0.74 | 1.19 | 0.141 | 0.383 | 1.000 | 8744 | tags=22%, list=15%, signal=26% |
| 3624 | PLASMA\_MEMBRANE\_RECEPTOR\_COMPLEX(GO:0098802) |  | 178 | 0.71 | 1.19 | 0.141 | 0.383 | 1.000 | 10811 | tags=30%, list=18%, signal=36% |
| 3625 | EXTRACELLULAR\_MATRIX\_ASSEMBLY(GO:0085029) |  | 32 | 0.70 | 1.19 | 0.000 | 0.383 | 1.000 | 12645 | tags=41%, list=21%, signal=52% |
| 3626 | MEMBRANE\_ORGANIZATION(GO:0061024) |  | 917 | 0.73 | 1.19 | 0.141 | 0.382 | 1.000 | 9006 | tags=12%, list=15%, signal=14% |
| 3627 | ACTIVE\_TRANSMEMBRANE\_TRANSPORTER\_ACTIVITY(GO:0022804) |  | 366 | 0.71 | 1.19 | 0.141 | 0.382 | 1.000 | 10330 | tags=30%, list=17%, signal=37% |
| 3628 | G\_PROTEIN\_COUPLED\_SEROTONIN\_RECEPTOR\_SIGNALING\_PATHWAY(GO:0098664) |  | 22 | 0.72 | 1.19 | 0.141 | 0.383 | 1.000 | 14753 | tags=36%, list=25%, signal=48% |
| 3629 | SEROTONIN\_RECEPTOR\_ACTIVITY(GO:0099589) |  | 22 | 0.72 | 1.19 | 0.141 | 0.382 | 1.000 | 14753 | tags=36%, list=25%, signal=48% |
| 3630 | SEROTONIN\_RECEPTOR\_SIGNALING\_PATHWAY(GO:0007210) |  | 22 | 0.72 | 1.19 | 0.141 | 0.382 | 1.000 | 14753 | tags=36%, list=25%, signal=48% |
| 3631 | G\_PROTEIN\_COUPLED\_SEROTONIN\_RECEPTOR\_ACTIVITY(GO:0004993) |  | 22 | 0.72 | 1.19 | 0.141 | 0.382 | 1.000 | 14753 | tags=36%, list=25%, signal=48% |
| 3632 | SUGAR\_TRANSMEMBRANE\_TRANSPORTER\_ACTIVITY(GO:0051119) |  | 28 | 0.73 | 1.19 | 0.141 | 0.382 | 1.000 | 8716 | tags=36%, list=15%, signal=42% |
| 3633 | EMBRYONIC\_EPITHELIAL\_TUBE\_FORMATION(GO:0001838) |  | 127 | 0.74 | 1.19 | 0.141 | 0.382 | 1.000 | 7975 | tags=16%, list=13%, signal=18% |
| 3634 | PROTEIN\_TARGETING\_TO\_MITOCHONDRION(GO:0006626) |  | 149 | 0.70 | 1.19 | 0.141 | 0.382 | 1.000 | 7426 | tags=7%, list=13%, signal=8% |
| 3635 | DEMETHYLATION(GO:0070988) |  | 66 | 0.74 | 1.19 | 0.141 | 0.382 | 1.000 | 7450 | tags=9%, list=13%, signal=10% |
| 3636 | STRIATED\_MUSCLE\_HYPERTROPHY(GO:0014897) |  | 98 | 0.70 | 1.19 | 0.141 | 0.382 | 1.000 | 13771 | tags=33%, list=23%, signal=42% |
| 3637 | TRANSFERASE\_ACTIVITY\_\_TRANSFERRING\_GLYCOSYL\_GROUPS(GO:0016757) |  | 289 | 0.71 | 1.19 | 0.141 | 0.382 | 1.000 | 9318 | tags=21%, list=16%, signal=25% |
| 3638 | MONOSACCHARIDE\_BINDING(GO:0048029) |  | 67 | 0.74 | 1.19 | 0.141 | 0.382 | 1.000 | 8383 | tags=24%, list=14%, signal=28% |
| 3639 | REGULATION\_OF\_MEMBRANE\_LIPID\_DISTRIBUTION(GO:0097035) |  | 40 | 0.75 | 1.19 | 0.141 | 0.382 | 1.000 | 8507 | tags=18%, list=14%, signal=20% |
| 3640 | PURINE\_NUCLEOSIDE\_TRIPHOSPHATE\_BIOSYNTHETIC\_PROCESS(GO:0009145) |  | 158 | 0.72 | 1.19 | 0.141 | 0.382 | 1.000 | 6696 | tags=5%, list=11%, signal=6% |
| 3641 | REGULATION\_OF\_RESPIRATORY\_GASEOUS\_EXCHANGE(GO:0043576) |  | 23 | 0.74 | 1.19 | 0.141 | 0.382 | 1.000 | 11201 | tags=39%, list=19%, signal=48% |
| 3642 | PROTEOGLYCAN\_BINDING(GO:0043394) |  | 37 | 0.76 | 1.19 | 0.141 | 0.382 | 1.000 | 7081 | tags=24%, list=12%, signal=28% |
| 3643 | ATP\_DEPENDENT\_MICROTUBULE\_MOTOR\_ACTIVITY\_\_PLUS\_END\_DIRECTED(GO:0008574) |  | 27 | 0.72 | 1.19 | 0.000 | 0.382 | 1.000 | 10334 | tags=44%, list=17%, signal=54% |
| 3644 | REGULATION\_OF\_PEPTIDE\_HORMONE\_SECRETION(GO:0090276) |  | 207 | 0.71 | 1.19 | 0.141 | 0.382 | 1.000 | 10131 | tags=24%, list=17%, signal=29% |
| 3645 | HISTONE\_MONOUBIQUITINATION(GO:0010390) |  | 29 | 0.70 | 1.19 | 0.000 | 0.382 | 1.000 | 10434 | tags=21%, list=18%, signal=25% |
| 3646 | CELLULAR\_AMINO\_ACID\_CATABOLIC\_PROCESS(GO:0009063) |  | 118 | 0.74 | 1.19 | 0.141 | 0.382 | 1.000 | 10534 | tags=25%, list=18%, signal=31% |
| 3647 | REGULATION\_OF\_INTERLEUKIN\_2\_BIOSYNTHETIC\_PROCESS(GO:0045076) |  | 18 | 0.78 | 1.19 | 0.141 | 0.382 | 1.000 | 7355 | tags=39%, list=12%, signal=44% |
| 3648 | SITE\_OF\_POLARIZED\_GROWTH(GO:0030427) |  | 162 | 0.75 | 1.19 | 0.141 | 0.381 | 1.000 | 10556 | tags=28%, list=18%, signal=34% |
| 3649 | VASCULAR\_SMOOTH\_MUSCLE\_CELL\_DIFFERENTIATION(GO:0035886) |  | 31 | 0.75 | 1.19 | 0.141 | 0.381 | 1.000 | 8774 | tags=26%, list=15%, signal=30% |
| 3650 | POSITIVE\_REGULATION\_OF\_RHO\_PROTEIN\_SIGNAL\_TRANSDUCTION(GO:0035025) |  | 26 | 0.78 | 1.19 | 0.141 | 0.381 | 1.000 | 6572 | tags=19%, list=11%, signal=22% |
| 3651 | NUCLEOSIDE\_SALVAGE(GO:0043174) |  | 17 | 0.70 | 1.19 | 0.000 | 0.381 | 1.000 | 9191 | tags=24%, list=16%, signal=28% |
| 3652 | RESPONSE\_TO\_INTERLEUKIN\_4(GO:0070670) |  | 34 | 0.74 | 1.19 | 0.141 | 0.381 | 1.000 | 7435 | tags=21%, list=13%, signal=24% |
| 3653 | BETA\_1\_3\_GALACTOSYLTRANSFERASE\_ACTIVITY(GO:0048531) |  | 16 | 0.76 | 1.19 | 0.141 | 0.381 | 1.000 | 9569 | tags=25%, list=16%, signal=30% |
| 3654 | REGULATION\_OF\_MEIOTIC\_NUCLEAR\_DIVISION(GO:0040020) |  | 30 | 0.69 | 1.19 | 0.141 | 0.381 | 1.000 | 11130 | tags=47%, list=19%, signal=57% |
| 3655 | OSSIFICATION\_INVOLVED\_IN\_BONE\_MATURATION(GO:0043931) |  | 17 | 0.74 | 1.19 | 0.000 | 0.382 | 1.000 | 10811 | tags=41%, list=18%, signal=50% |
| 3656 | MUSCLE\_HYPERTROPHY(GO:0014896) |  | 100 | 0.70 | 1.19 | 0.141 | 0.382 | 1.000 | 13771 | tags=33%, list=23%, signal=43% |
| 3657 | GOLGI\_APPARATUS(GO:0005794) |  | 1486 | 0.73 | 1.19 | 0.141 | 0.381 | 1.000 | 9585 | tags=16%, list=16%, signal=19% |
| 3658 | PHOTORECEPTOR\_INNER\_SEGMENT(GO:0001917) |  | 43 | 0.73 | 1.19 | 0.141 | 0.382 | 1.000 | 11588 | tags=44%, list=20%, signal=55% |
| 3659 | PURINE\_RIBONUCLEOSIDE\_TRIPHOSPHATE\_BIOSYNTHETIC\_PROCESS(GO:0009206) |  | 157 | 0.72 | 1.19 | 0.141 | 0.381 | 1.000 | 6696 | tags=5%, list=11%, signal=6% |
| 3660 | NEGATIVE\_REGULATION\_OF\_DNA\_BIOSYNTHETIC\_PROCESS(GO:2000279) |  | 36 | 0.69 | 1.19 | 0.000 | 0.381 | 1.000 | 14468 | tags=44%, list=24%, signal=59% |
| 3661 | FATTY\_ACID\_OXIDATION(GO:0019395) |  | 99 | 0.69 | 1.19 | 0.000 | 0.381 | 1.000 | 10056 | tags=15%, list=17%, signal=18% |
| 3662 | CIRCADIAN\_SLEEP\_WAKE\_CYCLE\_\_SLEEP(GO:0050802) |  | 23 | 0.73 | 1.19 | 0.000 | 0.381 | 1.000 | 7172 | tags=30%, list=12%, signal=35% |
| 3663 | APOPTOTIC\_NUCLEAR\_CHANGES(GO:0030262) |  | 30 | 0.69 | 1.19 | 0.000 | 0.381 | 1.000 | 15560 | tags=60%, list=26%, signal=81% |
| 3664 | POSITIVE\_REGULATION\_OF\_PROTEIN\_MODIFICATION\_PROCESS(GO:0031401) |  | 1151 | 0.73 | 1.19 | 0.141 | 0.381 | 1.000 | 9543 | tags=19%, list=16%, signal=22% |
| 3665 | MUSCLE\_CELL\_FATE\_COMMITMENT(GO:0042693) |  | 19 | 0.80 | 1.19 | 0.141 | 0.381 | 1.000 | 8673 | tags=37%, list=15%, signal=43% |
| 3666 | MITOCHONDRIAL\_MATRIX(GO:0005759) |  | 454 | 0.69 | 1.19 | 0.000 | 0.381 | 1.000 | 7671 | tags=5%, list=13%, signal=6% |
| 3667 | ENDOCRINE\_PROCESS(GO:0050886) |  | 86 | 0.72 | 1.19 | 0.141 | 0.381 | 1.000 | 8969 | tags=23%, list=15%, signal=27% |
| 3668 | POSITIVE\_REGULATION\_OF\_INTERLEUKIN\_10\_PRODUCTION(GO:0032733) |  | 30 | 0.71 | 1.19 | 0.141 | 0.381 | 1.000 | 9023 | tags=40%, list=15%, signal=47% |
| 3669 | REGULATION\_OF\_STRESS\_FIBER\_ASSEMBLY(GO:0051492) |  | 79 | 0.74 | 1.19 | 0.141 | 0.381 | 1.000 | 8112 | tags=14%, list=14%, signal=16% |
| 3670 | OXIDOREDUCTION\_COENZYME\_METABOLIC\_PROCESS(GO:0006733) |  | 191 | 0.72 | 1.19 | 0.141 | 0.381 | 1.000 | 8867 | tags=12%, list=15%, signal=14% |
| 3671 | NEGATIVE\_REGULATION\_OF\_RECEPTOR\_MEDIATED\_ENDOCYTOSIS(GO:0048261) |  | 18 | 0.72 | 1.19 | 0.000 | 0.381 | 1.000 | 7256 | tags=22%, list=12%, signal=25% |
| 3672 | POSITIVE\_REGULATION\_OF\_CELL\_PROJECTION\_ORGANIZATION(GO:0031346) |  | 344 | 0.73 | 1.19 | 0.141 | 0.381 | 1.000 | 9311 | tags=22%, list=16%, signal=26% |
| 3673 | REGULATION\_OF\_ERAD\_PATHWAY(GO:1904292) |  | 27 | 0.71 | 1.19 | 0.000 | 0.381 | 1.000 | 5999 | tags=7%, list=10%, signal=8% |
| 3674 | POSITIVE\_REGULATION\_OF\_RECEPTOR\_MEDIATED\_ENDOCYTOSIS(GO:0048260) |  | 50 | 0.72 | 1.19 | 0.141 | 0.381 | 1.000 | 12594 | tags=36%, list=21%, signal=46% |
| 3675 | RECYCLING\_ENDOSOME\_MEMBRANE(GO:0055038) |  | 68 | 0.75 | 1.19 | 0.141 | 0.381 | 1.000 | 9119 | tags=19%, list=15%, signal=23% |
| 3676 | POSITIVE\_REGULATION\_OF\_INTERLEUKIN\_4\_PRODUCTION(GO:0032753) |  | 21 | 0.78 | 1.19 | 0.141 | 0.381 | 1.000 | 6564 | tags=29%, list=11%, signal=32% |
| 3677 | RESPONSE\_TO\_VIRUS(GO:0009615) |  | 308 | 0.71 | 1.19 | 0.141 | 0.381 | 1.000 | 7663 | tags=11%, list=13%, signal=13% |
| 3678 | EXPORT\_ACROSS\_PLASMA\_MEMBRANE(GO:0140115) |  | 18 | 0.74 | 1.19 | 0.141 | 0.381 | 1.000 | 8363 | tags=39%, list=14%, signal=45% |
| 3679 | EARLY\_ENDOSOME(GO:0005769) |  | 330 | 0.73 | 1.19 | 0.141 | 0.381 | 1.000 | 7467 | tags=13%, list=13%, signal=15% |
| 3680 | VESICLE\_MEDIATED\_TRANSPORT\_IN\_SYNAPSE(GO:0099003) |  | 128 | 0.72 | 1.19 | 0.141 | 0.381 | 1.000 | 9395 | tags=23%, list=16%, signal=28% |
| 3681 | REGULATION\_OF\_LEUKOCYTE\_DEGRANULATION(GO:0043300) |  | 42 | 0.73 | 1.19 | 0.141 | 0.381 | 1.000 | 9198 | tags=17%, list=16%, signal=20% |
| 3682 | SPERM\_PART(GO:0097223) |  | 198 | 0.71 | 1.19 | 0.141 | 0.381 | 1.000 | 11087 | tags=28%, list=19%, signal=34% |
| 3683 | REGULATION\_OF\_CELL\_CYCLE(GO:0051726) |  | 1181 | 0.72 | 1.19 | 0.141 | 0.381 | 1.000 | 10717 | tags=19%, list=18%, signal=23% |
| 3684 | POST\_EMBRYONIC\_DEVELOPMENT(GO:0009791) |  | 95 | 0.74 | 1.18 | 0.141 | 0.381 | 1.000 | 8035 | tags=18%, list=14%, signal=21% |
| 3685 | AXONEMAL\_DYNEIN\_COMPLEX(GO:0005858) |  | 18 | 0.73 | 1.18 | 0.141 | 0.381 | 1.000 | 8044 | tags=22%, list=14%, signal=26% |
| 3686 | PROTEOGLYCAN\_METABOLIC\_PROCESS(GO:0006029) |  | 84 | 0.69 | 1.18 | 0.000 | 0.381 | 1.000 | 12257 | tags=44%, list=21%, signal=55% |
| 3687 | POSITIVE\_REGULATION\_OF\_DEPHOSPHORYLATION(GO:0035306) |  | 50 | 0.69 | 1.18 | 0.000 | 0.381 | 1.000 | 11536 | tags=28%, list=20%, signal=35% |
| 3688 | LOCALIZATION\_WITHIN\_MEMBRANE(GO:0051668) |  | 107 | 0.75 | 1.18 | 0.141 | 0.381 | 1.000 | 8937 | tags=19%, list=15%, signal=22% |
| 3689 | DENDRITE\_DEVELOPMENT(GO:0016358) |  | 211 | 0.73 | 1.18 | 0.141 | 0.381 | 1.000 | 9722 | tags=21%, list=16%, signal=25% |
| 3690 | PERIKARYON(GO:0043204) |  | 119 | 0.72 | 1.18 | 0.141 | 0.381 | 1.000 | 9231 | tags=24%, list=16%, signal=28% |
| 3691 | SPERMATID\_DEVELOPMENT(GO:0007286) |  | 139 | 0.72 | 1.18 | 0.141 | 0.381 | 1.000 | 10143 | tags=24%, list=17%, signal=29% |
| 3692 | MEIOTIC\_CHROMOSOME\_SEGREGATION(GO:0045132) |  | 90 | 0.73 | 1.18 | 0.141 | 0.381 | 1.000 | 12327 | tags=49%, list=21%, signal=62% |
| 3693 | NEGATIVE\_REGULATION\_OF\_FATTY\_ACID\_METABOLIC\_PROCESS(GO:0045922) |  | 25 | 0.72 | 1.18 | 0.141 | 0.381 | 1.000 | 13039 | tags=40%, list=22%, signal=51% |
| 3694 | REGULATION\_OF\_GTPASE\_ACTIVITY(GO:0043087) |  | 482 | 0.71 | 1.18 | 0.141 | 0.380 | 1.000 | 9584 | tags=17%, list=16%, signal=20% |
| 3695 | CHOLESTEROL\_TRANSPORTER\_ACTIVITY(GO:0017127) |  | 19 | 0.76 | 1.18 | 0.141 | 0.380 | 1.000 | 7172 | tags=37%, list=12%, signal=42% |
| 3696 | CELLULAR\_RESPONSE\_TO\_GONADOTROPIN\_STIMULUS(GO:0071371) |  | 19 | 0.73 | 1.18 | 0.000 | 0.380 | 1.000 | 7172 | tags=21%, list=12%, signal=24% |
| 3697 | DETECTION\_OF\_EXTERNAL\_STIMULUS(GO:0009581) |  | 132 | 0.71 | 1.18 | 0.141 | 0.380 | 1.000 | 12044 | tags=37%, list=20%, signal=47% |
| 3698 | CELLULAR\_RESPONSE\_TO\_INTERFERON\_GAMMA(GO:0071346) |  | 167 | 0.71 | 1.18 | 0.141 | 0.380 | 1.000 | 8063 | tags=13%, list=14%, signal=15% |
| 3699 | CELL\_CYCLE\_ARREST(GO:0007050) |  | 248 | 0.73 | 1.18 | 0.141 | 0.380 | 1.000 | 10012 | tags=17%, list=17%, signal=20% |
| 3700 | REGULATION\_OF\_CALCIUM\_ION\_IMPORT(GO:0090279) |  | 37 | 0.77 | 1.18 | 0.141 | 0.380 | 1.000 | 6662 | tags=16%, list=11%, signal=18% |
| 3701 | NEGATIVE\_REGULATION\_OF\_PROTEIN\_PROCESSING(GO:0010955) |  | 42 | 0.74 | 1.18 | 0.141 | 0.380 | 1.000 | 11568 | tags=33%, list=20%, signal=41% |
| 3702 | NEGATIVE\_REGULATION\_OF\_PROTEIN\_MATURATION(GO:1903318) |  | 42 | 0.74 | 1.18 | 0.141 | 0.380 | 1.000 | 11568 | tags=33%, list=20%, signal=41% |
| 3703 | REGULATION\_OF\_ACTIN\_FILAMENT\_ORGANIZATION(GO:0110053) |  | 241 | 0.72 | 1.18 | 0.141 | 0.380 | 1.000 | 8605 | tags=14%, list=15%, signal=16% |
| 3704 | NEGATIVE\_REGULATION\_OF\_CD4\_POSITIVE\_\_ALPHA\_BETA\_T\_CELL\_DIFFERENTIATION(GO:0043371) |  | 15 | 0.71 | 1.18 | 0.141 | 0.380 | 1.000 | 9411 | tags=27%, list=16%, signal=32% |
| 3705 | APOPTOTIC\_DNA\_FRAGMENTATION(GO:0006309) |  | 21 | 0.71 | 1.18 | 0.000 | 0.380 | 1.000 | 14434 | tags=62%, list=24%, signal=82% |
| 3706 | BASAL\_LAMINA(GO:0005605) |  | 21 | 0.77 | 1.18 | 0.141 | 0.380 | 1.000 | 11288 | tags=48%, list=19%, signal=59% |
| 3707 | REGULATION\_OF\_PROTEIN\_TARGETING(GO:1903533) |  | 126 | 0.72 | 1.18 | 0.141 | 0.380 | 1.000 | 7426 | tags=9%, list=13%, signal=10% |
| 3708 | NEGATIVE\_REGULATION\_OF\_PLATELET\_ACTIVATION(GO:0010544) |  | 18 | 0.80 | 1.18 | 0.141 | 0.380 | 1.000 | 7546 | tags=28%, list=13%, signal=32% |
| 3709 | PLASMA\_LIPOPROTEIN\_PARTICLE\_CLEARANCE(GO:0034381) |  | 56 | 0.71 | 1.18 | 0.141 | 0.380 | 1.000 | 12095 | tags=21%, list=20%, signal=27% |
| 3710 | CHROMOSOME\_ORGANIZATION\_INVOLVED\_IN\_MEIOTIC\_CELL\_CYCLE(GO:0070192) |  | 70 | 0.74 | 1.18 | 0.141 | 0.380 | 1.000 | 11597 | tags=44%, list=20%, signal=55% |
| 3711 | PROTEIN\_ACYLATION(GO:0043543) |  | 229 | 0.69 | 1.18 | 0.000 | 0.380 | 1.000 | 10862 | tags=21%, list=18%, signal=26% |
| 3712 | CELL\_ADHESION\_MEDIATED\_BY\_INTEGRIN(GO:0033627) |  | 56 | 0.71 | 1.18 | 0.141 | 0.380 | 1.000 | 10643 | tags=29%, list=18%, signal=35% |
| 3713 | RETINOIC\_ACID\_METABOLIC\_PROCESS(GO:0042573) |  | 23 | 0.78 | 1.18 | 0.141 | 0.380 | 1.000 | 11950 | tags=43%, list=20%, signal=54% |
| 3714 | REGULATION\_OF\_LIPOPROTEIN\_LIPASE\_ACTIVITY(GO:0051004) |  | 22 | 0.74 | 1.18 | 0.141 | 0.380 | 1.000 | 6197 | tags=18%, list=10%, signal=20% |
| 3715 | STEROID\_HORMONE\_MEDIATED\_SIGNALING\_PATHWAY(GO:0043401) |  | 196 | 0.72 | 1.18 | 0.141 | 0.380 | 1.000 | 8112 | tags=16%, list=14%, signal=18% |
| 3716 | LIGAND\_GATED\_CHANNEL\_ACTIVITY(GO:0022834) |  | 140 | 0.73 | 1.18 | 0.141 | 0.380 | 1.000 | 9703 | tags=32%, list=16%, signal=38% |
| 3717 | LIGAND\_GATED\_ION\_CHANNEL\_ACTIVITY(GO:0015276) |  | 140 | 0.73 | 1.18 | 0.141 | 0.380 | 1.000 | 9703 | tags=32%, list=16%, signal=38% |
| 3718 | REGULATION\_OF\_INTRACELLULAR\_PH(GO:0051453) |  | 91 | 0.70 | 1.18 | 0.141 | 0.380 | 1.000 | 9957 | tags=29%, list=17%, signal=34% |
| 3719 | POSITIVE\_REGULATION\_OF\_ACTIN\_FILAMENT\_POLYMERIZATION(GO:0030838) |  | 93 | 0.71 | 1.18 | 0.141 | 0.380 | 1.000 | 7951 | tags=12%, list=13%, signal=14% |
| 3720 | IONOTROPIC\_GLUTAMATE\_RECEPTOR\_ACTIVITY(GO:0004970) |  | 19 | 0.79 | 1.18 | 0.141 | 0.380 | 1.000 | 9000 | tags=37%, list=15%, signal=43% |
| 3721 | REGULATION\_OF\_ADENYLATE\_CYCLASE\_ACTIVITY(GO:0045761) |  | 37 | 0.70 | 1.18 | 0.141 | 0.380 | 1.000 | 7472 | tags=14%, list=13%, signal=15% |
| 3722 | COPPER\_ION\_TRANSPORT(GO:0006825) |  | 20 | 0.73 | 1.18 | 0.141 | 0.380 | 1.000 | 6066 | tags=10%, list=10%, signal=11% |
| 3723 | SPERMATID\_DIFFERENTIATION(GO:0048515) |  | 144 | 0.72 | 1.18 | 0.141 | 0.380 | 1.000 | 10143 | tags=24%, list=17%, signal=29% |
| 3724 | POSITIVE\_REGULATION\_OF\_MYOTUBE\_DIFFERENTIATION(GO:0010831) |  | 38 | 0.74 | 1.18 | 0.141 | 0.380 | 1.000 | 9121 | tags=21%, list=15%, signal=25% |
| 3725 | CYSTEINE\_TYPE\_ENDOPEPTIDASE\_ACTIVATOR\_ACTIVITY\_INVOLVED\_IN\_APOPTOTIC\_PROCESS(GO:0008656) |  | 16 | 0.73 | 1.18 | 0.000 | 0.380 | 1.000 | 12564 | tags=44%, list=21%, signal=56% |
| 3726 | PHOSPHATIDYLINOSITOL\_METABOLIC\_PROCESS(GO:0046488) |  | 221 | 0.71 | 1.18 | 0.141 | 0.380 | 1.000 | 9651 | tags=21%, list=16%, signal=25% |
| 3727 | MYOSIN\_II\_COMPLEX(GO:0016460) |  | 33 | 0.74 | 1.18 | 0.000 | 0.380 | 1.000 | 7898 | tags=39%, list=13%, signal=45% |
| 3728 | REGULATION\_OF\_ACTIN\_FILAMENT\_BUNDLE\_ASSEMBLY(GO:0032231) |  | 91 | 0.73 | 1.18 | 0.141 | 0.380 | 1.000 | 8112 | tags=15%, list=14%, signal=18% |
| 3729 | SOMATIC\_STEM\_CELL\_POPULATION\_MAINTENANCE(GO:0035019) |  | 74 | 0.73 | 1.18 | 0.141 | 0.380 | 1.000 | 7913 | tags=18%, list=13%, signal=20% |
| 3730 | CATALYTIC\_ACTIVITY\_\_ACTING\_ON\_A\_PROTEIN(GO:0140096) |  | 2423 | 0.71 | 1.18 | 0.141 | 0.380 | 1.000 | 10967 | tags=22%, list=19%, signal=26% |
| 3731 | IMMUNOGLOBULIN\_SECRETION(GO:0048305) |  | 19 | 0.74 | 1.18 | 0.141 | 0.380 | 1.000 | 6178 | tags=21%, list=10%, signal=24% |
| 3732 | PHOSPHATIDYLINOSITOL\_BISPHOSPHATE\_KINASE\_ACTIVITY(GO:0052813) |  | 71 | 0.73 | 1.18 | 0.141 | 0.380 | 1.000 | 7656 | tags=31%, list=13%, signal=36% |
| 3733 | POSITIVE\_REGULATION\_OF\_T\_CELL\_ACTIVATION(GO:0050870) |  | 201 | 0.70 | 1.18 | 0.141 | 0.380 | 1.000 | 10712 | tags=21%, list=18%, signal=26% |
| 3734 | POSITIVE\_REGULATION\_OF\_GTPASE\_ACTIVITY(GO:0043547) |  | 410 | 0.71 | 1.18 | 0.141 | 0.379 | 1.000 | 8030 | tags=13%, list=14%, signal=15% |
| 3735 | REGULATION\_OF\_MACROAUTOPHAGY(GO:0016241) |  | 164 | 0.68 | 1.18 | 0.141 | 0.379 | 1.000 | 7441 | tags=5%, list=13%, signal=6% |
| 3736 | CELLULAR\_PROTEIN\_MODIFICATION\_PROCESS(GO:0006464) |  | 4205 | 0.74 | 1.18 | 0.141 | 0.379 | 1.000 | 8123 | tags=13%, list=14%, signal=13% |
| 3737 | PROTEIN\_MODIFICATION\_PROCESS(GO:0036211) |  | 4205 | 0.74 | 1.18 | 0.141 | 0.379 | 1.000 | 8123 | tags=13%, list=14%, signal=13% |
| 3738 | CELLULAR\_RESPONSE\_TO\_NERVE\_GROWTH\_FACTOR\_STIMULUS(GO:1990090) |  | 46 | 0.68 | 1.18 | 0.000 | 0.379 | 1.000 | 8937 | tags=15%, list=15%, signal=18% |
| 3739 | CELLULAR\_PROCESS\_INVOLVED\_IN\_REPRODUCTION\_IN\_MULTICELLULAR\_ORGANISM(GO:0022412) |  | 334 | 0.71 | 1.18 | 0.141 | 0.379 | 1.000 | 10998 | tags=27%, list=19%, signal=33% |
| 3740 | CATECHOLAMINE\_BINDING(GO:1901338) |  | 20 | 0.76 | 1.18 | 0.141 | 0.379 | 1.000 | 7441 | tags=10%, list=13%, signal=11% |
| 3741 | GOLGI\_CISTERNA\_MEMBRANE(GO:0032580) |  | 73 | 0.72 | 1.18 | 0.141 | 0.379 | 1.000 | 9585 | tags=21%, list=16%, signal=24% |
| 3742 | ORGANIC\_ACID\_TRANSMEMBRANE\_TRANSPORTER\_ACTIVITY(GO:0005342) |  | 156 | 0.72 | 1.18 | 0.141 | 0.379 | 1.000 | 11768 | tags=36%, list=20%, signal=45% |
| 3743 | CARBOXYLIC\_ACID\_TRANSMEMBRANE\_TRANSPORTER\_ACTIVITY(GO:0046943) |  | 156 | 0.72 | 1.18 | 0.141 | 0.379 | 1.000 | 11768 | tags=36%, list=20%, signal=45% |
| 3744 | PROTEIN\_POLYMERIZATION(GO:0051258) |  | 259 | 0.69 | 1.18 | 0.141 | 0.379 | 1.000 | 9368 | tags=16%, list=16%, signal=19% |
| 3745 | REGULATION\_OF\_CGMP\_METABOLIC\_PROCESS(GO:0030823) |  | 28 | 0.76 | 1.18 | 0.141 | 0.379 | 1.000 | 8279 | tags=29%, list=14%, signal=33% |
| 3746 | METHIONINE\_METABOLIC\_PROCESS(GO:0006555) |  | 20 | 0.70 | 1.18 | 0.000 | 0.379 | 1.000 | 13127 | tags=40%, list=22%, signal=51% |
| 3747 | RIBONUCLEOSIDE\_TRIPHOSPHATE\_BIOSYNTHETIC\_PROCESS(GO:0009201) |  | 163 | 0.71 | 1.18 | 0.141 | 0.379 | 1.000 | 6696 | tags=5%, list=11%, signal=6% |
| 3748 | RETINOID\_METABOLIC\_PROCESS(GO:0001523) |  | 89 | 0.73 | 1.18 | 0.141 | 0.379 | 1.000 | 11950 | tags=31%, list=20%, signal=39% |
| 3749 | INTERLEUKIN\_17\_PRODUCTION(GO:0032620) |  | 26 | 0.75 | 1.18 | 0.141 | 0.379 | 1.000 | 9368 | tags=27%, list=16%, signal=32% |
| 3750 | B\_CELL\_DIFFERENTIATION(GO:0030183) |  | 121 | 0.69 | 1.18 | 0.141 | 0.379 | 1.000 | 11130 | tags=27%, list=19%, signal=34% |
| 3751 | NUCLEOCYTOPLASMIC\_TRANSPORT(GO:0006913) |  | 346 | 0.73 | 1.18 | 0.141 | 0.379 | 1.000 | 7005 | tags=6%, list=12%, signal=7% |
| 3752 | ORGANELLE\_SUBCOMPARTMENT(GO:0031984) |  | 1613 | 0.71 | 1.18 | 0.141 | 0.379 | 1.000 | 9677 | tags=15%, list=16%, signal=18% |
| 3753 | LIPID\_OXIDATION(GO:0034440) |  | 101 | 0.69 | 1.18 | 0.000 | 0.379 | 1.000 | 10056 | tags=16%, list=17%, signal=19% |
| 3754 | EPITHELIAL\_TUBE\_FORMATION(GO:0072175) |  | 135 | 0.74 | 1.18 | 0.141 | 0.379 | 1.000 | 7975 | tags=17%, list=13%, signal=20% |
| 3755 | POTASSIUM\_CHANNEL\_ACTIVITY(GO:0005267) |  | 133 | 0.73 | 1.18 | 0.141 | 0.379 | 1.000 | 9703 | tags=32%, list=16%, signal=39% |
| 3756 | VACUOLE\_ORGANIZATION(GO:0007033) |  | 152 | 0.69 | 1.18 | 0.141 | 0.379 | 1.000 | 11630 | tags=22%, list=20%, signal=27% |
| 3757 | POSITIVE\_REGULATION\_OF\_OSTEOBLAST\_DIFFERENTIATION(GO:0045669) |  | 60 | 0.74 | 1.18 | 0.141 | 0.379 | 1.000 | 8046 | tags=13%, list=14%, signal=15% |
| 3758 | MITOCHONDRIAL\_CALCIUM\_ION\_TRANSMEMBRANE\_TRANSPORT(GO:0006851) |  | 25 | 0.70 | 1.18 | 0.000 | 0.379 | 1.000 | 6323 | tags=8%, list=11%, signal=9% |
| 3759 | HEXOSE\_TRANSMEMBRANE\_TRANSPORT(GO:0008645) |  | 117 | 0.69 | 1.18 | 0.141 | 0.378 | 1.000 | 9042 | tags=21%, list=15%, signal=25% |
| 3760 | CYTOKINE\_RECEPTOR\_ACTIVITY(GO:0004896) |  | 93 | 0.73 | 1.18 | 0.141 | 0.378 | 1.000 | 11737 | tags=32%, list=20%, signal=40% |
| 3761 | DEVELOPMENTAL\_PROCESS\_INVOLVED\_IN\_REPRODUCTION(GO:0003006) |  | 657 | 0.73 | 1.18 | 0.141 | 0.378 | 1.000 | 10998 | tags=27%, list=19%, signal=33% |
| 3762 | T\_CELL\_ACTIVATION\_INVOLVED\_IN\_IMMUNE\_RESPONSE(GO:0002286) |  | 95 | 0.71 | 1.18 | 0.141 | 0.378 | 1.000 | 7414 | tags=20%, list=13%, signal=23% |
| 3763 | UDP\_GLYCOSYLTRANSFERASE\_ACTIVITY(GO:0008194) |  | 145 | 0.69 | 1.18 | 0.141 | 0.378 | 1.000 | 9901 | tags=23%, list=17%, signal=28% |
| 3764 | REGULATION\_OF\_ACTOMYOSIN\_STRUCTURE\_ORGANIZATION(GO:0110020) |  | 89 | 0.74 | 1.18 | 0.141 | 0.378 | 1.000 | 8112 | tags=16%, list=14%, signal=18% |
| 3765 | POSITIVE\_REGULATION\_OF\_BONE\_REMODELING(GO:0046852) |  | 16 | 0.73 | 1.18 | 0.141 | 0.378 | 1.000 | 10901 | tags=44%, list=18%, signal=54% |
| 3766 | POSITIVE\_REGULATION\_OF\_BONE\_RESORPTION(GO:0045780) |  | 16 | 0.73 | 1.18 | 0.141 | 0.378 | 1.000 | 10901 | tags=44%, list=18%, signal=54% |
| 3767 | REGULATION\_OF\_INTERLEUKIN\_17\_PRODUCTION(GO:0032660) |  | 23 | 0.77 | 1.18 | 0.141 | 0.378 | 1.000 | 9368 | tags=30%, list=16%, signal=36% |
| 3768 | BILE\_ACID\_AND\_BILE\_SALT\_TRANSPORT(GO:0015721) |  | 33 | 0.75 | 1.18 | 0.141 | 0.378 | 1.000 | 9376 | tags=24%, list=16%, signal=29% |
| 3769 | REGULATION\_OF\_SMALL\_MOLECULE\_METABOLIC\_PROCESS(GO:0062012) |  | 484 | 0.71 | 1.18 | 0.141 | 0.378 | 1.000 | 8594 | tags=14%, list=15%, signal=17% |
| 3770 | POSITIVE\_REGULATION\_OF\_NEURON\_APOPTOTIC\_PROCESS(GO:0043525) |  | 49 | 0.72 | 1.18 | 0.141 | 0.378 | 1.000 | 7963 | tags=16%, list=13%, signal=19% |
| 3771 | REGULATION\_OF\_CELLULAR\_AMINE\_METABOLIC\_PROCESS(GO:0033238) |  | 82 | 0.72 | 1.18 | 0.141 | 0.378 | 1.000 | 7935 | tags=6%, list=13%, signal=7% |
| 3772 | RESPONSE\_TO\_PAIN(GO:0048265) |  | 29 | 0.76 | 1.18 | 0.141 | 0.378 | 1.000 | 7873 | tags=31%, list=13%, signal=36% |
| 3773 | LOW\_DENSITY\_LIPOPROTEIN\_PARTICLE\_CLEARANCE(GO:0034383) |  | 34 | 0.70 | 1.18 | 0.141 | 0.378 | 1.000 | 12095 | tags=21%, list=20%, signal=26% |
| 3774 | REGULATION\_OF\_PH(GO:0006885) |  | 102 | 0.70 | 1.18 | 0.141 | 0.378 | 1.000 | 9957 | tags=28%, list=17%, signal=34% |
| 3775 | REGULATION\_OF\_OLIGODENDROCYTE\_DIFFERENTIATION(GO:0048713) |  | 32 | 0.74 | 1.18 | 0.141 | 0.378 | 1.000 | 9664 | tags=22%, list=16%, signal=26% |
| 3776 | POSITIVE\_REGULATION\_OF\_MAP\_KINASE\_ACTIVITY(GO:0043406) |  | 262 | 0.71 | 1.18 | 0.141 | 0.378 | 1.000 | 8279 | tags=19%, list=14%, signal=22% |
| 3777 | REGULATION\_OF\_PROTEIN\_DEPOLYMERIZATION(GO:1901879) |  | 73 | 0.72 | 1.18 | 0.141 | 0.378 | 1.000 | 11742 | tags=29%, list=20%, signal=36% |
| 3778 | REGULATION\_OF\_CD4\_POSITIVE\_\_ALPHA\_BETA\_T\_CELL\_DIFFERENTIATION(GO:0043370) |  | 37 | 0.73 | 1.18 | 0.141 | 0.378 | 1.000 | 9411 | tags=32%, list=16%, signal=39% |
| 3779 | CELL\_DIVISION(GO:0051301) |  | 591 | 0.72 | 1.18 | 0.141 | 0.378 | 1.000 | 10939 | tags=25%, list=18%, signal=30% |
| 3780 | NEGATIVE\_REGULATION\_OF\_DNA\_DEPENDENT\_DNA\_REPLICATION(GO:2000104) |  | 21 | 0.71 | 1.18 | 0.000 | 0.378 | 1.000 | 13120 | tags=71%, list=22%, signal=92% |
| 3781 | EXTRACELLULARLY\_GLUTAMATE\_GATED\_ION\_CHANNEL\_ACTIVITY(GO:0005234) |  | 19 | 0.78 | 1.18 | 0.141 | 0.378 | 1.000 | 9000 | tags=37%, list=15%, signal=43% |
| 3782 | MICROTUBULE\_MOTOR\_ACTIVITY(GO:0003777) |  | 126 | 0.68 | 1.18 | 0.000 | 0.378 | 1.000 | 13157 | tags=43%, list=22%, signal=55% |
| 3783 | SERTOLI\_CELL\_DEVELOPMENT(GO:0060009) |  | 15 | 0.73 | 1.18 | 0.141 | 0.378 | 1.000 | 10743 | tags=33%, list=18%, signal=41% |
| 3784 | NUCLEAR\_TRANSPORT(GO:0051169) |  | 350 | 0.72 | 1.18 | 0.141 | 0.378 | 1.000 | 7005 | tags=6%, list=12%, signal=7% |
| 3785 | NEPHRON\_TUBULE\_FORMATION(GO:0072079) |  | 19 | 0.80 | 1.18 | 0.141 | 0.378 | 1.000 | 9208 | tags=26%, list=16%, signal=31% |
| 3786 | REGULATION\_OF\_LIPID\_TRANSPORT(GO:0032368) |  | 109 | 0.71 | 1.18 | 0.141 | 0.378 | 1.000 | 12117 | tags=30%, list=20%, signal=38% |
| 3787 | REGULATION\_OF\_GLUCOSE\_TRANSMEMBRANE\_TRANSPORT(GO:0010827) |  | 81 | 0.67 | 1.18 | 0.141 | 0.378 | 1.000 | 10571 | tags=19%, list=18%, signal=23% |
| 3788 | NEGATIVE\_REGULATION\_OF\_CD4\_POSITIVE\_\_ALPHA\_BETA\_T\_CELL\_ACTIVATION(GO:2000515) |  | 20 | 0.70 | 1.18 | 0.000 | 0.378 | 1.000 | 6685 | tags=25%, list=11%, signal=28% |
| 3789 | NEGATIVE\_REGULATION\_OF\_DIGESTIVE\_SYSTEM\_PROCESS(GO:0060457) |  | 17 | 0.76 | 1.18 | 0.141 | 0.378 | 1.000 | 6987 | tags=35%, list=12%, signal=40% |
| 3790 | MALE\_SEX\_DIFFERENTIATION(GO:0046661) |  | 158 | 0.73 | 1.18 | 0.141 | 0.378 | 1.000 | 8613 | tags=21%, list=15%, signal=24% |
| 3791 | PEPTIDASE\_ACTIVATOR\_ACTIVITY\_INVOLVED\_IN\_APOPTOTIC\_PROCESS(GO:0016505) |  | 19 | 0.73 | 1.18 | 0.000 | 0.378 | 1.000 | 12564 | tags=42%, list=21%, signal=53% |
| 3792 | REGULATION\_OF\_ACTIN\_FILAMENT\_DEPOLYMERIZATION(GO:0030834) |  | 49 | 0.72 | 1.18 | 0.141 | 0.378 | 1.000 | 8232 | tags=16%, list=14%, signal=19% |
| 3793 | MITOTIC\_CELL\_CYCLE(GO:0000278) |  | 1003 | 0.71 | 1.18 | 0.141 | 0.378 | 1.000 | 10952 | tags=22%, list=19%, signal=26% |
| 3794 | LAMELLIPODIUM\_ASSEMBLY(GO:0030032) |  | 62 | 0.74 | 1.18 | 0.141 | 0.378 | 1.000 | 9336 | tags=23%, list=16%, signal=27% |
| 3795 | POSITIVE\_REGULATION\_OF\_CREB\_TRANSCRIPTION\_FACTOR\_ACTIVITY(GO:0032793) |  | 15 | 0.74 | 1.18 | 0.141 | 0.378 | 1.000 | 8046 | tags=20%, list=14%, signal=23% |
| 3796 | THIOL\_DEPENDENT\_UBIQUITIN\_SPECIFIC\_PROTEASE\_ACTIVITY(GO:0004843) |  | 85 | 0.69 | 1.18 | 0.000 | 0.378 | 1.000 | 9353 | tags=15%, list=16%, signal=18% |
| 3797 | DYNEIN\_INTERMEDIATE\_CHAIN\_BINDING(GO:0045505) |  | 27 | 0.71 | 1.18 | 0.000 | 0.378 | 1.000 | 8569 | tags=26%, list=14%, signal=30% |
| 3798 | ERYTHROCYTE\_DEVELOPMENT(GO:0048821) |  | 30 | 0.70 | 1.18 | 0.141 | 0.378 | 1.000 | 10768 | tags=43%, list=18%, signal=53% |
| 3799 | KIDNEY\_MESENCHYME\_DEVELOPMENT(GO:0072074) |  | 19 | 0.76 | 1.18 | 0.141 | 0.378 | 1.000 | 9208 | tags=37%, list=16%, signal=44% |
| 3800 | HELICASE\_ACTIVITY(GO:0004386) |  | 153 | 0.68 | 1.18 | 0.000 | 0.378 | 1.000 | 13756 | tags=37%, list=23%, signal=48% |
| 3801 | TRANSCRIPTION\_COFACTOR\_ACTIVITY(GO:0003712) |  | 541 | 0.70 | 1.18 | 0.141 | 0.378 | 1.000 | 10905 | tags=21%, list=18%, signal=25% |
| 3802 | NEGATIVE\_REGULATION\_OF\_SMOOTHENED\_SIGNALING\_PATHWAY(GO:0045879) |  | 30 | 0.76 | 1.18 | 0.141 | 0.378 | 1.000 | 12402 | tags=50%, list=21%, signal=63% |
| 3803 | POSITIVE\_REGULATION\_OF\_HYDROLASE\_ACTIVITY(GO:0051345) |  | 745 | 0.71 | 1.18 | 0.141 | 0.378 | 1.000 | 8046 | tags=13%, list=14%, signal=15% |
| 3804 | CONNECTIVE\_TISSUE\_REPLACEMENT(GO:0097709) |  | 18 | 0.72 | 1.18 | 0.000 | 0.378 | 1.000 | 6486 | tags=6%, list=11%, signal=6% |
| 3805 | SPERMATOGENESIS(GO:0007283) |  | 514 | 0.71 | 1.18 | 0.141 | 0.378 | 1.000 | 10832 | tags=26%, list=18%, signal=31% |
| 3806 | ACTIVATION\_OF\_PHOSPHOLIPASE\_C\_ACTIVITY(GO:0007202) |  | 24 | 0.72 | 1.18 | 0.141 | 0.378 | 1.000 | 13992 | tags=58%, list=24%, signal=76% |
| 3807 | POSITIVE\_REGULATION\_OF\_SMOOTH\_MUSCLE\_CELL\_DIFFERENTIATION(GO:0051152) |  | 20 | 0.73 | 1.18 | 0.141 | 0.378 | 1.000 | 11362 | tags=25%, list=19%, signal=31% |
| 3808 | PROTON\_TRANSMEMBRANE\_TRANSPORT(GO:1902600) |  | 142 | 0.69 | 1.18 | 0.141 | 0.378 | 1.000 | 10330 | tags=21%, list=17%, signal=26% |
| 3809 | RESPONSE\_TO\_HYPOXIA(GO:0001666) |  | 349 | 0.72 | 1.18 | 0.141 | 0.378 | 1.000 | 9042 | tags=15%, list=15%, signal=18% |
| 3810 | RECEPTOR\_TYROSINE\_KINASE\_BINDING(GO:0030971) |  | 57 | 0.68 | 1.18 | 0.141 | 0.378 | 1.000 | 10041 | tags=21%, list=17%, signal=25% |
| 3811 | NEGATIVE\_REGULATION\_OF\_PROTEIN\_BINDING(GO:0032091) |  | 99 | 0.71 | 1.18 | 0.141 | 0.378 | 1.000 | 9311 | tags=20%, list=16%, signal=24% |
| 3812 | 1\_PHOSPHATIDYLINOSITOL\_3\_KINASE\_ACTIVITY(GO:0016303) |  | 43 | 0.67 | 1.18 | 0.141 | 0.378 | 1.000 | 7547 | tags=26%, list=13%, signal=29% |
| 3813 | COENZYME\_METABOLIC\_PROCESS(GO:0006732) |  | 377 | 0.70 | 1.18 | 0.141 | 0.378 | 1.000 | 8607 | tags=10%, list=15%, signal=11% |
| 3814 | RESPONSE\_TO\_HEXOSE(GO:0009746) |  | 193 | 0.71 | 1.18 | 0.141 | 0.378 | 1.000 | 11480 | tags=28%, list=19%, signal=35% |
| 3815 | HORMONE\_MEDIATED\_SIGNALING\_PATHWAY(GO:0009755) |  | 235 | 0.71 | 1.18 | 0.141 | 0.378 | 1.000 | 10768 | tags=24%, list=18%, signal=30% |
| 3816 | REGULATION\_OF\_G1\_S\_TRANSITION\_OF\_MITOTIC\_CELL\_CYCLE(GO:2000045) |  | 169 | 0.72 | 1.18 | 0.141 | 0.378 | 1.000 | 7640 | tags=8%, list=13%, signal=9% |
| 3817 | REGULATION\_OF\_CELLULAR\_PH(GO:0030641) |  | 95 | 0.69 | 1.18 | 0.141 | 0.378 | 1.000 | 9957 | tags=28%, list=17%, signal=34% |
| 3818 | PEPTIDYL\_LYSINE\_MODIFICATION(GO:0018205) |  | 390 | 0.68 | 1.18 | 0.000 | 0.377 | 1.000 | 13833 | tags=34%, list=23%, signal=44% |
| 3819 | PROTEIN\_KINASE\_C\_ACTIVATING\_G\_PROTEIN\_COUPLED\_RECEPTOR\_SIGNALING\_PATHWAY(GO:0007205) |  | 29 | 0.71 | 1.18 | 0.141 | 0.377 | 1.000 | 7271 | tags=28%, list=12%, signal=31% |
| 3820 | REGULATION\_OF\_DNA\_METABOLIC\_PROCESS(GO:0051052) |  | 397 | 0.73 | 1.18 | 0.141 | 0.377 | 1.000 | 7945 | tags=11%, list=13%, signal=12% |
| 3821 | REGULATION\_OF\_T\_CELL\_ACTIVATION(GO:0050863) |  | 308 | 0.70 | 1.18 | 0.141 | 0.377 | 1.000 | 8351 | tags=16%, list=14%, signal=19% |
| 3822 | POSITIVE\_REGULATION\_OF\_LEUKOCYTE\_CELL\_CELL\_ADHESION(GO:1903039) |  | 212 | 0.70 | 1.18 | 0.141 | 0.377 | 1.000 | 10712 | tags=22%, list=18%, signal=26% |
| 3823 | ANTERIOR\_POSTERIOR\_AXIS\_SPECIFICATION(GO:0009948) |  | 51 | 0.73 | 1.18 | 0.141 | 0.377 | 1.000 | 8733 | tags=31%, list=15%, signal=37% |
| 3824 | GAS\_TRANSPORT(GO:0015669) |  | 22 | 0.76 | 1.18 | 0.141 | 0.377 | 1.000 | 10857 | tags=41%, list=18%, signal=50% |
| 3825 | GENE\_SILENCING\_BY\_MIRNA(GO:0035195) |  | 320 | 0.70 | 1.18 | 0.141 | 0.377 | 1.000 | 5858 | tags=6%, list=10%, signal=7% |
| 3826 | RESPONSE\_TO\_GLUCOSE(GO:0009749) |  | 188 | 0.70 | 1.18 | 0.141 | 0.377 | 1.000 | 11480 | tags=28%, list=19%, signal=34% |
| 3827 | CHLORIDE\_CHANNEL\_REGULATOR\_ACTIVITY(GO:0017081) |  | 16 | 0.74 | 1.18 | 0.141 | 0.377 | 1.000 | 9050 | tags=38%, list=15%, signal=44% |
| 3828 | REGULATION\_OF\_CYCLASE\_ACTIVITY(GO:0031279) |  | 53 | 0.70 | 1.18 | 0.141 | 0.377 | 1.000 | 8279 | tags=13%, list=14%, signal=15% |
| 3829 | RNA\_METABOLIC\_PROCESS(GO:0016070) |  | 4837 | 0.74 | 1.18 | 0.141 | 0.377 | 1.000 | 8683 | tags=14%, list=15%, signal=15% |
| 3830 | POSITIVE\_REGULATION\_OF\_CELL\_KILLING(GO:0031343) |  | 42 | 0.68 | 1.18 | 0.000 | 0.377 | 1.000 | 14247 | tags=48%, list=24%, signal=63% |
| 3831 | CONDENSED\_CHROMOSOME\_\_CENTROMERIC\_REGION(GO:0000779) |  | 117 | 0.68 | 1.18 | 0.000 | 0.377 | 1.000 | 14323 | tags=58%, list=24%, signal=77% |
| 3832 | PROTEIN\_TRANSPORTER\_ACTIVITY(GO:0008565) |  | 82 | 0.66 | 1.18 | 0.000 | 0.377 | 1.000 | 16328 | tags=56%, list=28%, signal=77% |
| 3833 | SMALL\_MOLECULE\_CATABOLIC\_PROCESS(GO:0044282) |  | 417 | 0.71 | 1.18 | 0.141 | 0.377 | 1.000 | 10549 | tags=20%, list=18%, signal=25% |
| 3834 | PROTEIN\_LIPID\_COMPLEX(GO:0032994) |  | 42 | 0.70 | 1.17 | 0.141 | 0.377 | 1.000 | 7068 | tags=19%, list=12%, signal=22% |
| 3835 | ANTEROGRADE\_AXONAL\_TRANSPORT(GO:0008089) |  | 32 | 0.70 | 1.17 | 0.000 | 0.377 | 1.000 | 15340 | tags=59%, list=26%, signal=80% |
| 3836 | RESPONSE\_TO\_HYPEROXIA(GO:0055093) |  | 21 | 0.73 | 1.17 | 0.000 | 0.377 | 1.000 | 9746 | tags=29%, list=16%, signal=34% |
| 3837 | REGULATION\_OF\_TRANSCRIPTION\_INVOLVED\_IN\_G1\_S\_TRANSITION\_OF\_MITOTIC\_CELL\_CYCLE(GO:0000083) |  | 28 | 0.69 | 1.17 | 0.000 | 0.377 | 1.000 | 16141 | tags=68%, list=27%, signal=93% |
| 3838 | GASTRULATION\_WITH\_MOUTH\_FORMING\_SECOND(GO:0001702) |  | 27 | 0.68 | 1.17 | 0.000 | 0.377 | 1.000 | 13004 | tags=41%, list=22%, signal=52% |
| 3839 | INTERLEUKIN\_8\_PRODUCTION(GO:0032637) |  | 71 | 0.70 | 1.17 | 0.141 | 0.377 | 1.000 | 6626 | tags=13%, list=11%, signal=14% |
| 3840 | TERPENOID\_METABOLIC\_PROCESS(GO:0006721) |  | 106 | 0.72 | 1.17 | 0.141 | 0.377 | 1.000 | 11550 | tags=32%, list=20%, signal=40% |
| 3841 | REGULATION\_OF\_DNA\_REPLICATION(GO:0006275) |  | 119 | 0.70 | 1.17 | 0.141 | 0.377 | 1.000 | 14016 | tags=45%, list=24%, signal=59% |
| 3842 | CELL\_SUBSTRATE\_ADHERENS\_JUNCTION\_ASSEMBLY(GO:0007045) |  | 73 | 0.73 | 1.17 | 0.141 | 0.377 | 1.000 | 7136 | tags=8%, list=12%, signal=9% |
| 3843 | FOCAL\_ADHESION\_ASSEMBLY(GO:0048041) |  | 73 | 0.73 | 1.17 | 0.141 | 0.377 | 1.000 | 7136 | tags=8%, list=12%, signal=9% |
| 3844 | STRESS\_FIBER\_ASSEMBLY(GO:0043149) |  | 91 | 0.73 | 1.17 | 0.141 | 0.377 | 1.000 | 8594 | tags=14%, list=15%, signal=17% |
| 3845 | CONTRACTILE\_ACTIN\_FILAMENT\_BUNDLE\_ASSEMBLY(GO:0030038) |  | 91 | 0.73 | 1.17 | 0.141 | 0.377 | 1.000 | 8594 | tags=14%, list=15%, signal=17% |
| 3846 | NEGATIVE\_REGULATION\_OF\_G2\_M\_TRANSITION\_OF\_MITOTIC\_CELL\_CYCLE(GO:0010972) |  | 98 | 0.69 | 1.17 | 0.141 | 0.377 | 1.000 | 12431 | tags=24%, list=21%, signal=31% |
| 3847 | INTRACELLULAR\_CHOLESTEROL\_TRANSPORT(GO:0032367) |  | 18 | 0.74 | 1.17 | 0.141 | 0.377 | 1.000 | 11715 | tags=39%, list=20%, signal=48% |
| 3848 | INTRACELLULAR\_STEROL\_TRANSPORT(GO:0032366) |  | 18 | 0.74 | 1.17 | 0.141 | 0.377 | 1.000 | 11715 | tags=39%, list=20%, signal=48% |
| 3849 | REGULATION\_OF\_IMMUNE\_RESPONSE(GO:0050776) |  | 1022 | 0.70 | 1.17 | 0.141 | 0.377 | 1.000 | 9411 | tags=14%, list=16%, signal=17% |
| 3850 | REGULATION\_OF\_NUCLEAR\_DIVISION(GO:0051783) |  | 179 | 0.69 | 1.17 | 0.141 | 0.377 | 1.000 | 11409 | tags=35%, list=19%, signal=43% |
| 3851 | LIPASE\_INHIBITOR\_ACTIVITY(GO:0055102) |  | 16 | 0.73 | 1.17 | 0.141 | 0.377 | 1.000 | 11260 | tags=38%, list=19%, signal=46% |
| 3852 | FATTY\_ACID\_CATABOLIC\_PROCESS(GO:0009062) |  | 102 | 0.69 | 1.17 | 0.000 | 0.377 | 1.000 | 10056 | tags=14%, list=17%, signal=17% |
| 3853 | AMINO\_SUGAR\_METABOLIC\_PROCESS(GO:0006040) |  | 38 | 0.69 | 1.17 | 0.000 | 0.377 | 1.000 | 9382 | tags=32%, list=16%, signal=38% |
| 3854 | POSITIVE\_REGULATION\_OF\_HEMOPOIESIS(GO:1903708) |  | 174 | 0.70 | 1.17 | 0.141 | 0.377 | 1.000 | 9368 | tags=25%, list=16%, signal=29% |
| 3855 | REGULATION\_OF\_LYMPHOCYTE\_CHEMOTAXIS(GO:1901623) |  | 20 | 0.74 | 1.17 | 0.141 | 0.377 | 1.000 | 6473 | tags=20%, list=11%, signal=22% |
| 3856 | REGULATION\_OF\_MITOTIC\_NUCLEAR\_DIVISION(GO:0007088) |  | 155 | 0.69 | 1.17 | 0.141 | 0.376 | 1.000 | 11409 | tags=34%, list=19%, signal=41% |
| 3857 | T\_CELL\_APOPTOTIC\_PROCESS(GO:0070231) |  | 48 | 0.71 | 1.17 | 0.141 | 0.376 | 1.000 | 6685 | tags=8%, list=11%, signal=9% |
| 3858 | MITOCHONDRIAL\_PART(GO:0044429) |  | 1031 | 0.68 | 1.17 | 0.000 | 0.376 | 1.000 | 10782 | tags=15%, list=18%, signal=18% |
| 3859 | PHOSPHATIDYLINOSITOL\_3\_KINASE\_ACTIVITY(GO:0035004) |  | 76 | 0.72 | 1.17 | 0.141 | 0.376 | 1.000 | 10131 | tags=34%, list=17%, signal=41% |
| 3860 | CARBOHYDRATE\_BINDING(GO:0030246) |  | 275 | 0.72 | 1.17 | 0.141 | 0.376 | 1.000 | 11786 | tags=29%, list=20%, signal=36% |
| 3861 | ADHERENS\_JUNCTION\_ASSEMBLY(GO:0034333) |  | 83 | 0.73 | 1.17 | 0.141 | 0.376 | 1.000 | 7136 | tags=8%, list=12%, signal=10% |
| 3862 | NUCLEOLUS(GO:0005730) |  | 941 | 0.70 | 1.17 | 0.141 | 0.376 | 1.000 | 10795 | tags=18%, list=18%, signal=22% |
| 3863 | POSITIVE\_REGULATION\_OF\_GLIOGENESIS(GO:0014015) |  | 57 | 0.72 | 1.17 | 0.141 | 0.376 | 1.000 | 10811 | tags=32%, list=18%, signal=39% |
| 3864 | MONOSACCHARIDE\_TRANSMEMBRANE\_TRANSPORT(GO:0015749) |  | 119 | 0.68 | 1.17 | 0.141 | 0.376 | 1.000 | 9042 | tags=21%, list=15%, signal=25% |
| 3865 | POSITIVE\_REGULATION\_OF\_INTERLEUKIN\_1\_SECRETION(GO:0050716) |  | 29 | 0.73 | 1.17 | 0.141 | 0.376 | 1.000 | 6473 | tags=7%, list=11%, signal=8% |
| 3866 | PROTEIN\_HOMODIMERIZATION\_ACTIVITY(GO:0042803) |  | 797 | 0.71 | 1.17 | 0.141 | 0.376 | 1.000 | 9857 | tags=21%, list=17%, signal=25% |
| 3867 | STEROID\_HORMONE\_RECEPTOR\_BINDING(GO:0035258) |  | 90 | 0.71 | 1.17 | 0.141 | 0.376 | 1.000 | 8490 | tags=10%, list=14%, signal=12% |
| 3868 | COBALAMIN\_METABOLIC\_PROCESS(GO:0009235) |  | 22 | 0.67 | 1.17 | 0.000 | 0.376 | 1.000 | 18441 | tags=77%, list=31%, signal=112% |
| 3869 | DITERPENOID\_METABOLIC\_PROCESS(GO:0016101) |  | 96 | 0.72 | 1.17 | 0.141 | 0.376 | 1.000 | 11550 | tags=31%, list=20%, signal=39% |
| 3870 | REGULATION\_OF\_BONE\_RESORPTION(GO:0045124) |  | 38 | 0.72 | 1.17 | 0.141 | 0.376 | 1.000 | 10901 | tags=34%, list=18%, signal=42% |
| 3871 | REGULATION\_OF\_LYASE\_ACTIVITY(GO:0051339) |  | 54 | 0.69 | 1.17 | 0.141 | 0.376 | 1.000 | 11214 | tags=19%, list=19%, signal=23% |
| 3872 | MALE\_GAMETE\_GENERATION(GO:0048232) |  | 532 | 0.71 | 1.17 | 0.141 | 0.376 | 1.000 | 11004 | tags=27%, list=19%, signal=32% |
| 3873 | NEGATIVE\_REGULATION\_OF\_LYMPHOCYTE\_APOPTOTIC\_PROCESS(GO:0070229) |  | 29 | 0.74 | 1.17 | 0.141 | 0.376 | 1.000 | 9411 | tags=17%, list=16%, signal=20% |
| 3874 | NEGATIVE\_REGULATION\_OF\_LEUKOCYTE\_ACTIVATION(GO:0002695) |  | 151 | 0.70 | 1.17 | 0.141 | 0.376 | 1.000 | 10712 | tags=25%, list=18%, signal=30% |
| 3875 | DENDRITIC\_SPINE\_DEVELOPMENT(GO:0060996) |  | 86 | 0.67 | 1.17 | 0.141 | 0.376 | 1.000 | 12455 | tags=30%, list=21%, signal=38% |
| 3876 | REGULATION\_OF\_ENDOCYTOSIS(GO:0030100) |  | 228 | 0.72 | 1.17 | 0.141 | 0.376 | 1.000 | 9520 | tags=16%, list=16%, signal=19% |
| 3877 | HEPARAN\_SULFATE\_PROTEOGLYCAN\_METABOLIC\_PROCESS(GO:0030201) |  | 30 | 0.73 | 1.17 | 0.141 | 0.376 | 1.000 | 12257 | tags=43%, list=21%, signal=55% |
| 3878 | TRANSCRIPTION\_FACTOR\_ACTIVITY\_\_TRANSCRIPTION\_FACTOR\_BINDING(GO:0000989) |  | 607 | 0.70 | 1.17 | 0.141 | 0.376 | 1.000 | 8774 | tags=13%, list=15%, signal=15% |
| 3879 | REGULATION\_OF\_T\_HELPER\_1\_TYPE\_IMMUNE\_RESPONSE(GO:0002825) |  | 22 | 0.74 | 1.17 | 0.000 | 0.376 | 1.000 | 9411 | tags=36%, list=16%, signal=43% |
| 3880 | DNA\_CATABOLIC\_PROCESS(GO:0006308) |  | 35 | 0.67 | 1.17 | 0.000 | 0.376 | 1.000 | 15700 | tags=51%, list=27%, signal=70% |
| 3881 | NUCLEOSIDE\_TRIPHOSPHATE\_BIOSYNTHETIC\_PROCESS(GO:0009142) |  | 174 | 0.70 | 1.17 | 0.141 | 0.376 | 1.000 | 6696 | tags=5%, list=11%, signal=6% |
| 3882 | PROTEIN\_ACTIVATION\_CASCADE(GO:0072376) |  | 189 | 0.69 | 1.17 | 0.141 | 0.376 | 1.000 | 11960 | tags=16%, list=20%, signal=20% |
| 3883 | VACUOLAR\_LUMEN(GO:0005775) |  | 170 | 0.69 | 1.17 | 0.141 | 0.376 | 1.000 | 12327 | tags=24%, list=21%, signal=30% |
| 3884 | NEUROMUSCULAR\_PROCESS\_CONTROLLING\_BALANCE(GO:0050885) |  | 53 | 0.67 | 1.17 | 0.000 | 0.376 | 1.000 | 13386 | tags=40%, list=23%, signal=51% |
| 3885 | REGULATION\_OF\_RECEPTOR\_MEDIATED\_ENDOCYTOSIS(GO:0048259) |  | 80 | 0.70 | 1.17 | 0.141 | 0.376 | 1.000 | 12594 | tags=33%, list=21%, signal=41% |
| 3886 | ORGANIC\_ACID\_CATABOLIC\_PROCESS(GO:0016054) |  | 260 | 0.71 | 1.17 | 0.141 | 0.376 | 1.000 | 10534 | tags=22%, list=18%, signal=26% |
| 3887 | CARBOXYLIC\_ACID\_CATABOLIC\_PROCESS(GO:0046395) |  | 260 | 0.71 | 1.17 | 0.141 | 0.376 | 1.000 | 10534 | tags=22%, list=18%, signal=26% |
| 3888 | REGULATION\_OF\_CELL\_ADHESION\_MEDIATED\_BY\_INTEGRIN(GO:0033628) |  | 40 | 0.72 | 1.17 | 0.141 | 0.376 | 1.000 | 10643 | tags=25%, list=18%, signal=30% |
| 3889 | INTERLEUKIN\_4\_PRODUCTION(GO:0032633) |  | 33 | 0.73 | 1.17 | 0.141 | 0.376 | 1.000 | 12217 | tags=36%, list=21%, signal=46% |
| 3890 | NECROTIC\_CELL\_DEATH(GO:0070265) |  | 52 | 0.69 | 1.17 | 0.141 | 0.376 | 1.000 | 12202 | tags=19%, list=21%, signal=24% |
| 3891 | REGULATION\_OF\_MYOTUBE\_DIFFERENTIATION(GO:0010830) |  | 63 | 0.72 | 1.17 | 0.141 | 0.376 | 1.000 | 9121 | tags=21%, list=15%, signal=24% |
| 3892 | POSITIVE\_REGULATION\_OF\_GLUCOSE\_IMPORT(GO:0046326) |  | 44 | 0.68 | 1.17 | 0.141 | 0.376 | 1.000 | 8594 | tags=14%, list=15%, signal=16% |
| 3893 | POSITIVE\_REGULATION\_OF\_RESPONSE\_TO\_EXTERNAL\_STIMULUS(GO:0032103) |  | 283 | 0.72 | 1.17 | 0.141 | 0.376 | 1.000 | 9446 | tags=19%, list=16%, signal=23% |
| 3894 | TRANSCRIPTION\_FACTOR\_ACTIVITY\_\_PROTEIN\_BINDING(GO:0000988) |  | 611 | 0.70 | 1.17 | 0.141 | 0.376 | 1.000 | 10674 | tags=19%, list=18%, signal=23% |
| 3895 | APICAL\_JUNCTION\_ASSEMBLY(GO:0043297) |  | 60 | 0.73 | 1.17 | 0.141 | 0.376 | 1.000 | 6796 | tags=8%, list=11%, signal=9% |
| 3896 | SERINE\_TYPE\_ENDOPEPTIDASE\_INHIBITOR\_ACTIVITY(GO:0004867) |  | 94 | 0.73 | 1.17 | 0.141 | 0.376 | 1.000 | 9448 | tags=26%, list=16%, signal=30% |
| 3897 | DEVELOPMENTAL\_MATURATION(GO:0021700) |  | 257 | 0.70 | 1.17 | 0.141 | 0.376 | 1.000 | 10347 | tags=27%, list=17%, signal=32% |
| 3898 | RESPONSE\_TO\_DECREASED\_OXYGEN\_LEVELS(GO:0036293) |  | 357 | 0.72 | 1.17 | 0.141 | 0.376 | 1.000 | 9042 | tags=15%, list=15%, signal=17% |
| 3899 | MACROMOLECULE\_MODIFICATION(GO:0043412) |  | 4395 | 0.73 | 1.17 | 0.141 | 0.376 | 1.000 | 8341 | tags=13%, list=14%, signal=14% |
| 3900 | RETINOIC\_ACID\_RECEPTOR\_BINDING(GO:0042974) |  | 24 | 0.70 | 1.17 | 0.000 | 0.376 | 1.000 | 12545 | tags=25%, list=21%, signal=32% |
| 3901 | REGULATION\_OF\_NEURON\_MIGRATION(GO:2001222) |  | 35 | 0.73 | 1.17 | 0.141 | 0.376 | 1.000 | 9681 | tags=23%, list=16%, signal=27% |
| 3902 | CARBOHYDRATE\_TRANSPORT(GO:0008643) |  | 151 | 0.68 | 1.17 | 0.141 | 0.376 | 1.000 | 9571 | tags=23%, list=16%, signal=27% |
| 3903 | CYCLIN\_BINDING(GO:0030332) |  | 32 | 0.71 | 1.17 | 0.141 | 0.376 | 1.000 | 12984 | tags=44%, list=22%, signal=56% |
| 3904 | HISTONE\_BINDING(GO:0042393) |  | 190 | 0.68 | 1.17 | 0.000 | 0.376 | 1.000 | 13760 | tags=36%, list=23%, signal=46% |
| 3905 | REGULATION\_OF\_ANION\_TRANSMEMBRANE\_TRANSPORT(GO:1903959) |  | 23 | 0.73 | 1.17 | 0.141 | 0.376 | 1.000 | 8452 | tags=17%, list=14%, signal=20% |
| 3906 | HORMONE\_BINDING(GO:0042562) |  | 96 | 0.72 | 1.17 | 0.141 | 0.376 | 1.000 | 6708 | tags=19%, list=11%, signal=21% |
| 3907 | PROTEIN\_N\_TERMINUS\_BINDING(GO:0047485) |  | 108 | 0.67 | 1.17 | 0.000 | 0.376 | 1.000 | 13572 | tags=33%, list=23%, signal=43% |
| 3908 | EPITHELIAL\_TO\_MESENCHYMAL\_TRANSITION\_INVOLVED\_IN\_ENDOCARDIAL\_CUSHION\_FORMATION(GO:0003198) |  | 16 | 0.73 | 1.17 | 0.141 | 0.376 | 1.000 | 12990 | tags=56%, list=22%, signal=72% |
| 3909 | TUMOR\_NECROSIS\_FACTOR\_RECEPTOR\_BINDING(GO:0005164) |  | 32 | 0.70 | 1.17 | 0.141 | 0.376 | 1.000 | 10751 | tags=22%, list=18%, signal=27% |
| 3910 | PROTEIN\_LIPID\_COMPLEX\_ASSEMBLY(GO:0065005) |  | 30 | 0.69 | 1.17 | 0.000 | 0.376 | 1.000 | 13100 | tags=30%, list=22%, signal=39% |
| 3911 | ADAPTIVE\_IMMUNE\_RESPONSE(GO:0002250) |  | 547 | 0.69 | 1.17 | 0.141 | 0.376 | 1.000 | 11438 | tags=18%, list=19%, signal=23% |
| 3912 | TRANSCRIPTION\_COACTIVATOR\_ACTIVITY(GO:0003713) |  | 305 | 0.70 | 1.17 | 0.141 | 0.376 | 1.000 | 10485 | tags=20%, list=18%, signal=25% |
| 3913 | CELLULAR\_RESPONSE\_TO\_GLUCAGON\_STIMULUS(GO:0071377) |  | 41 | 0.70 | 1.17 | 0.141 | 0.376 | 1.000 | 10468 | tags=34%, list=18%, signal=41% |
| 3914 | POSITIVE\_REGULATION\_OF\_ACUTE\_INFLAMMATORY\_RESPONSE(GO:0002675) |  | 29 | 0.73 | 1.17 | 0.141 | 0.376 | 1.000 | 11372 | tags=28%, list=19%, signal=34% |
| 3915 | HYDROLASE\_ACTIVITY\_\_ACTING\_ON\_CARBON\_NITROGEN\_(BUT\_NOT\_PEPTIDE)\_BONDS(GO:0016810) |  | 149 | 0.72 | 1.17 | 0.141 | 0.376 | 1.000 | 11227 | tags=27%, list=19%, signal=33% |
| 3916 | NEGATIVE\_REGULATION\_OF\_MRNA\_CATABOLIC\_PROCESS(GO:1902373) |  | 44 | 0.76 | 1.17 | 0.141 | 0.376 | 1.000 | 5906 | tags=2%, list=10%, signal=3% |
| 3917 | ACTIVATION\_OF\_NF\_KAPPAB\_INDUCING\_KINASE\_ACTIVITY(GO:0007250) |  | 18 | 0.69 | 1.17 | 0.141 | 0.376 | 1.000 | 15981 | tags=50%, list=27%, signal=68% |
| 3918 | CYCLIC\_NUCLEOTIDE\_BIOSYNTHETIC\_PROCESS(GO:0009190) |  | 98 | 0.71 | 1.17 | 0.141 | 0.376 | 1.000 | 8279 | tags=21%, list=14%, signal=25% |
| 3919 | NEGATIVE\_REGULATION\_OF\_PROTEIN\_DEPHOSPHORYLATION(GO:0035308) |  | 57 | 0.68 | 1.17 | 0.000 | 0.376 | 1.000 | 10702 | tags=23%, list=18%, signal=28% |
| 3920 | POSITIVE\_REGULATION\_OF\_PROTEASOMAL\_UBIQUITIN\_DEPENDENT\_PROTEIN\_CATABOLIC\_PROCESS(GO:0032436) |  | 81 | 0.69 | 1.17 | 0.141 | 0.376 | 1.000 | 7795 | tags=12%, list=13%, signal=14% |
| 3921 | PHOSPHOLIPID\_TRANSPORTER\_ACTIVITY(GO:0005548) |  | 56 | 0.73 | 1.17 | 0.141 | 0.376 | 1.000 | 7675 | tags=16%, list=13%, signal=18% |
| 3922 | WNT\_SIGNALING\_PATHWAY\_\_PLANAR\_CELL\_POLARITY\_PATHWAY(GO:0060071) |  | 110 | 0.69 | 1.17 | 0.141 | 0.376 | 1.000 | 12594 | tags=27%, list=21%, signal=35% |
| 3923 | MICROTUBULE\_BASED\_TRANSPORT(GO:0099111) |  | 133 | 0.67 | 1.17 | 0.000 | 0.376 | 1.000 | 15406 | tags=49%, list=26%, signal=66% |
| 3924 | TRANSPORT\_ALONG\_MICROTUBULE(GO:0010970) |  | 133 | 0.67 | 1.17 | 0.000 | 0.375 | 1.000 | 15406 | tags=49%, list=26%, signal=66% |
| 3925 | INTERCELLULAR\_BRIDGE(GO:0045171) |  | 58 | 0.68 | 1.17 | 0.000 | 0.375 | 1.000 | 13819 | tags=40%, list=23%, signal=52% |
| 3926 | ENDOCRINE\_HORMONE\_SECRETION(GO:0060986) |  | 49 | 0.71 | 1.17 | 0.141 | 0.375 | 1.000 | 13419 | tags=37%, list=23%, signal=47% |
| 3927 | ATP\_METABOLIC\_PROCESS(GO:0046034) |  | 286 | 0.69 | 1.17 | 0.141 | 0.375 | 1.000 | 6696 | tags=6%, list=11%, signal=6% |
| 3928 | POSITIVE\_REGULATION\_OF\_TRANSFERASE\_ACTIVITY(GO:0051347) |  | 627 | 0.70 | 1.17 | 0.141 | 0.375 | 1.000 | 10829 | tags=21%, list=18%, signal=25% |
| 3929 | POSITIVE\_REGULATION\_OF\_EXTRINSIC\_APOPTOTIC\_SIGNALING\_PATHWAY\_VIA\_DEATH\_DOMAIN\_RECEPTORS(GO:1902043) |  | 17 | 0.75 | 1.17 | 0.141 | 0.375 | 1.000 | 7101 | tags=18%, list=12%, signal=20% |
| 3930 | EMBRYONIC\_PATTERN\_SPECIFICATION(GO:0009880) |  | 63 | 0.73 | 1.17 | 0.141 | 0.375 | 1.000 | 11411 | tags=37%, list=19%, signal=45% |
| 3931 | POSITIVE\_REGULATION\_OF\_CAMP\_METABOLIC\_PROCESS(GO:0030816) |  | 36 | 0.70 | 1.17 | 0.141 | 0.375 | 1.000 | 6671 | tags=22%, list=11%, signal=25% |
| 3932 | OXIDOREDUCTASE\_ACTIVITY\_\_ACTING\_ON\_PAIRED\_DONORS\_\_WITH\_INCORPORATION\_OR\_REDUCTION\_OF\_MOLECULAR\_OXYGEN\_\_REDUCED\_FLAVIN\_OR\_FLAVOPROTEIN\_AS\_ONE\_DONOR\_\_AND\_INCORPORATION\_OF\_ONE\_ATOM\_OF\_OXYGEN(GO:0016712) |  | 27 | 0.71 | 1.17 | 0.141 | 0.375 | 1.000 | 11149 | tags=30%, list=19%, signal=36% |
| 3933 | INTRAMOLECULAR\_OXIDOREDUCTASE\_ACTIVITY(GO:0016860) |  | 49 | 0.67 | 1.17 | 0.000 | 0.375 | 1.000 | 16824 | tags=59%, list=28%, signal=83% |
| 3934 | GLUCOSE\_IMPORT(GO:0046323) |  | 81 | 0.68 | 1.17 | 0.141 | 0.375 | 1.000 | 8801 | tags=17%, list=15%, signal=20% |
| 3935 | POSITIVE\_REGULATION\_OF\_ERK1\_AND\_ERK2\_CASCADE(GO:0070374) |  | 197 | 0.71 | 1.17 | 0.141 | 0.375 | 1.000 | 7727 | tags=17%, list=13%, signal=20% |
| 3936 | ADIPOSE\_TISSUE\_DEVELOPMENT(GO:0060612) |  | 37 | 0.69 | 1.17 | 0.141 | 0.376 | 1.000 | 13967 | tags=38%, list=24%, signal=50% |
| 3937 | DIACYLGLYCEROL\_METABOLIC\_PROCESS(GO:0046339) |  | 16 | 0.76 | 1.17 | 0.000 | 0.376 | 1.000 | 9255 | tags=31%, list=16%, signal=37% |
| 3938 | CELLULAR\_MONOVALENT\_INORGANIC\_CATION\_HOMEOSTASIS(GO:0030004) |  | 111 | 0.68 | 1.17 | 0.141 | 0.376 | 1.000 | 9957 | tags=27%, list=17%, signal=32% |
| 3939 | HEME\_BINDING(GO:0020037) |  | 136 | 0.69 | 1.17 | 0.141 | 0.376 | 1.000 | 11264 | tags=29%, list=19%, signal=36% |
| 3940 | INTEGRAL\_COMPONENT\_OF\_ORGANELLE\_MEMBRANE(GO:0031301) |  | 179 | 0.67 | 1.17 | 0.000 | 0.376 | 1.000 | 13102 | tags=29%, list=22%, signal=37% |
| 3941 | REGULATION\_OF\_MYELOID\_LEUKOCYTE\_DIFFERENTIATION(GO:0002761) |  | 114 | 0.70 | 1.17 | 0.141 | 0.376 | 1.000 | 11089 | tags=29%, list=19%, signal=36% |
| 3942 | ATPASE\_DEPENDENT\_TRANSMEMBRANE\_TRANSPORT\_COMPLEX(GO:0098533) |  | 24 | 0.70 | 1.17 | 0.141 | 0.376 | 1.000 | 12215 | tags=46%, list=21%, signal=58% |
| 3943 | INTERLEUKIN\_10\_PRODUCTION(GO:0032613) |  | 46 | 0.69 | 1.17 | 0.141 | 0.375 | 1.000 | 15466 | tags=48%, list=26%, signal=65% |
| 3944 | SUBSTANTIA\_NIGRA\_DEVELOPMENT(GO:0021762) |  | 44 | 0.69 | 1.17 | 0.141 | 0.375 | 1.000 | 7818 | tags=16%, list=13%, signal=18% |
| 3945 | NUCLEAR\_CHROMOSOME(GO:0000228) |  | 576 | 0.70 | 1.17 | 0.141 | 0.375 | 1.000 | 11810 | tags=27%, list=20%, signal=33% |
| 3946 | REGULATION\_OF\_EPITHELIAL\_CELL\_MIGRATION(GO:0010632) |  | 266 | 0.71 | 1.17 | 0.141 | 0.375 | 1.000 | 9016 | tags=19%, list=15%, signal=23% |
| 3947 | LEUKOCYTE\_DIFFERENTIATION(GO:0002521) |  | 486 | 0.72 | 1.17 | 0.141 | 0.375 | 1.000 | 7967 | tags=18%, list=13%, signal=21% |
| 3948 | NEGATIVE\_REGULATION\_OF\_CELLULAR\_RESPONSE\_TO\_GROWTH\_FACTOR\_STIMULUS(GO:0090288) |  | 149 | 0.71 | 1.17 | 0.141 | 0.375 | 1.000 | 7913 | tags=17%, list=13%, signal=20% |
| 3949 | PURINE\_RIBONUCLEOSIDE\_MONOPHOSPHATE\_BIOSYNTHETIC\_PROCESS(GO:0009168) |  | 169 | 0.70 | 1.17 | 0.141 | 0.375 | 1.000 | 8541 | tags=9%, list=14%, signal=11% |
| 3950 | PURINE\_NUCLEOSIDE\_MONOPHOSPHATE\_BIOSYNTHETIC\_PROCESS(GO:0009127) |  | 169 | 0.70 | 1.17 | 0.141 | 0.375 | 1.000 | 8541 | tags=9%, list=14%, signal=11% |
| 3951 | MAST\_CELL\_ACTIVATION\_INVOLVED\_IN\_IMMUNE\_RESPONSE(GO:0002279) |  | 45 | 0.71 | 1.17 | 0.141 | 0.375 | 1.000 | 10643 | tags=27%, list=18%, signal=32% |
| 3952 | MAST\_CELL\_DEGRANULATION(GO:0043303) |  | 45 | 0.71 | 1.17 | 0.141 | 0.375 | 1.000 | 10643 | tags=27%, list=18%, signal=32% |
| 3953 | REGULATION\_OF\_CD4\_POSITIVE\_\_ALPHA\_BETA\_T\_CELL\_ACTIVATION(GO:2000514) |  | 45 | 0.71 | 1.17 | 0.141 | 0.375 | 1.000 | 9411 | tags=33%, list=16%, signal=40% |
| 3954 | POSITIVE\_REGULATION\_OF\_INTERLEUKIN\_1\_BETA\_SECRETION(GO:0050718) |  | 26 | 0.74 | 1.17 | 0.141 | 0.375 | 1.000 | 6473 | tags=8%, list=11%, signal=9% |
| 3955 | POSITIVE\_REGULATION\_OF\_REACTIVE\_OXYGEN\_SPECIES\_BIOSYNTHETIC\_PROCESS(GO:1903428) |  | 49 | 0.71 | 1.17 | 0.141 | 0.375 | 1.000 | 8755 | tags=16%, list=15%, signal=19% |
| 3956 | DOUBLE\_STRAND\_BREAK\_REPAIR\_VIA\_NONHOMOLOGOUS\_END\_JOINING(GO:0006303) |  | 76 | 0.72 | 1.17 | 0.141 | 0.375 | 1.000 | 9900 | tags=13%, list=17%, signal=16% |
| 3957 | PROTEIN\_HOMOTETRAMERIZATION(GO:0051289) |  | 80 | 0.72 | 1.17 | 0.141 | 0.375 | 1.000 | 11915 | tags=30%, list=20%, signal=38% |
| 3958 | RESPONSE\_TO\_OXYGEN\_LEVELS(GO:0070482) |  | 380 | 0.72 | 1.17 | 0.141 | 0.375 | 1.000 | 10294 | tags=19%, list=17%, signal=23% |
| 3959 | NEGATIVE\_REGULATION\_OF\_MITOTIC\_NUCLEAR\_DIVISION(GO:0045839) |  | 48 | 0.67 | 1.17 | 0.000 | 0.375 | 1.000 | 15525 | tags=56%, list=26%, signal=76% |
| 3960 | POSITIVE\_REGULATION\_OF\_ESTABLISHMENT\_OF\_PROTEIN\_LOCALIZATION\_TO\_MITOCHONDRION(GO:1903749) |  | 112 | 0.73 | 1.17 | 0.141 | 0.375 | 1.000 | 7583 | tags=8%, list=13%, signal=9% |
| 3961 | NEGATIVE\_REGULATION\_OF\_GLYCOPROTEIN\_METABOLIC\_PROCESS(GO:1903019) |  | 15 | 0.70 | 1.17 | 0.000 | 0.375 | 1.000 | 12547 | tags=27%, list=21%, signal=34% |
| 3962 | BASAL\_PART\_OF\_CELL(GO:0045178) |  | 45 | 0.73 | 1.17 | 0.141 | 0.375 | 1.000 | 11790 | tags=38%, list=20%, signal=47% |
| 3963 | SMAD\_PROTEIN\_SIGNAL\_TRANSDUCTION(GO:0060395) |  | 69 | 0.70 | 1.17 | 0.141 | 0.375 | 1.000 | 13279 | tags=45%, list=22%, signal=58% |
| 3964 | VENTRICULAR\_SEPTUM\_MORPHOGENESIS(GO:0060412) |  | 39 | 0.74 | 1.17 | 0.141 | 0.375 | 1.000 | 7913 | tags=28%, list=13%, signal=33% |
| 3965 | REGULATION\_OF\_ESTABLISHMENT\_OF\_PLANAR\_POLARITY(GO:0090175) |  | 112 | 0.69 | 1.17 | 0.141 | 0.375 | 1.000 | 12594 | tags=27%, list=21%, signal=34% |
| 3966 | NON\_MOTILE\_CILIUM\_ASSEMBLY(GO:1905515) |  | 52 | 0.68 | 1.16 | 0.000 | 0.375 | 1.000 | 13376 | tags=52%, list=23%, signal=67% |
| 3967 | ACTIN\_FILAMENT\_DEPOLYMERIZATION(GO:0030042) |  | 53 | 0.71 | 1.16 | 0.141 | 0.375 | 1.000 | 8232 | tags=15%, list=14%, signal=18% |
| 3968 | POSITIVE\_REGULATION\_OF\_MACROPHAGE\_ACTIVATION(GO:0043032) |  | 23 | 0.75 | 1.16 | 0.141 | 0.375 | 1.000 | 8627 | tags=30%, list=15%, signal=36% |
| 3969 | INFLAMMATORY\_CELL\_APOPTOTIC\_PROCESS(GO:0006925) |  | 20 | 0.72 | 1.16 | 0.000 | 0.375 | 1.000 | 10262 | tags=20%, list=17%, signal=24% |
| 3970 | FIBROBLAST\_MIGRATION(GO:0010761) |  | 42 | 0.72 | 1.16 | 0.141 | 0.375 | 1.000 | 8112 | tags=17%, list=14%, signal=19% |
| 3971 | CELLULAR\_COMPONENT\_DISASSEMBLY\_INVOLVED\_IN\_EXECUTION\_PHASE\_OF\_APOPTOSIS(GO:0006921) |  | 32 | 0.69 | 1.16 | 0.141 | 0.375 | 1.000 | 15560 | tags=56%, list=26%, signal=76% |
| 3972 | REGULATION\_OF\_DELAYED\_RECTIFIER\_POTASSIUM\_CHANNEL\_ACTIVITY(GO:1902259) |  | 20 | 0.69 | 1.16 | 0.000 | 0.375 | 1.000 | 16616 | tags=55%, list=28%, signal=76% |
| 3973 | MAST\_CELL\_MEDIATED\_IMMUNITY(GO:0002448) |  | 47 | 0.71 | 1.16 | 0.141 | 0.375 | 1.000 | 10643 | tags=26%, list=18%, signal=31% |
| 3974 | PHOTORECEPTOR\_CONNECTING\_CILIUM(GO:0032391) |  | 34 | 0.68 | 1.16 | 0.000 | 0.375 | 1.000 | 13479 | tags=53%, list=23%, signal=69% |
| 3975 | OXIDOREDUCTASE\_COMPLEX(GO:1990204) |  | 114 | 0.65 | 1.16 | 0.000 | 0.375 | 1.000 | 9653 | tags=11%, list=16%, signal=13% |
| 3976 | NEGATIVE\_REGULATION\_OF\_VIRAL\_ENTRY\_INTO\_HOST\_CELL(GO:0046597) |  | 19 | 0.73 | 1.16 | 0.141 | 0.375 | 1.000 | 6283 | tags=5%, list=11%, signal=6% |
| 3977 | REGULATION\_OF\_INTERLEUKIN\_10\_PRODUCTION(GO:0032653) |  | 44 | 0.69 | 1.16 | 0.141 | 0.375 | 1.000 | 15466 | tags=48%, list=26%, signal=65% |
| 3978 | REGULATION\_OF\_STEROL\_TRANSPORT(GO:0032371) |  | 50 | 0.72 | 1.16 | 0.141 | 0.375 | 1.000 | 11906 | tags=32%, list=20%, signal=40% |
| 3979 | REGULATION\_OF\_CHOLESTEROL\_TRANSPORT(GO:0032374) |  | 50 | 0.72 | 1.16 | 0.141 | 0.375 | 1.000 | 11906 | tags=32%, list=20%, signal=40% |
| 3980 | REGULATION\_OF\_STEROID\_METABOLIC\_PROCESS(GO:0019218) |  | 108 | 0.68 | 1.16 | 0.141 | 0.375 | 1.000 | 11640 | tags=28%, list=20%, signal=35% |
| 3981 | POSITIVE\_REGULATION\_OF\_B\_CELL\_PROLIFERATION(GO:0030890) |  | 40 | 0.70 | 1.16 | 0.141 | 0.375 | 1.000 | 10136 | tags=40%, list=17%, signal=48% |
| 3982 | NEGATIVE\_REGULATION\_OF\_MITOTIC\_SISTER\_CHROMATID\_SEGREGATION(GO:0033048) |  | 38 | 0.67 | 1.16 | 0.000 | 0.375 | 1.000 | 15943 | tags=61%, list=27%, signal=83% |
| 3983 | CALCIUM\_INDEPENDENT\_CELL\_CELL\_ADHESION\_VIA\_PLASMA\_MEMBRANE\_CELL\_ADHESION\_MOLECULES(GO:0016338) |  | 23 | 0.77 | 1.16 | 0.141 | 0.375 | 1.000 | 6320 | tags=22%, list=11%, signal=24% |
| 3984 | UBIQUITIN\_PROTEIN\_TRANSFERASE\_REGULATOR\_ACTIVITY(GO:0055106) |  | 18 | 0.70 | 1.16 | 0.000 | 0.375 | 1.000 | 10469 | tags=33%, list=18%, signal=40% |
| 3985 | MEIOSIS\_I(GO:0007127) |  | 114 | 0.69 | 1.16 | 0.141 | 0.375 | 1.000 | 11304 | tags=34%, list=19%, signal=42% |
| 3986 | PROTEIN\_DIMERIZATION\_ACTIVITY(GO:0046983) |  | 1296 | 0.71 | 1.16 | 0.141 | 0.375 | 1.000 | 10556 | tags=22%, list=18%, signal=26% |
| 3987 | CALCIUM\_ION\_REGULATED\_EXOCYTOSIS\_OF\_NEUROTRANSMITTER(GO:0048791) |  | 30 | 0.73 | 1.16 | 0.141 | 0.375 | 1.000 | 9144 | tags=40%, list=15%, signal=47% |
| 3988 | NEGATIVE\_REGULATION\_OF\_CELL\_CYCLE\_G2\_M\_PHASE\_TRANSITION(GO:1902750) |  | 110 | 0.68 | 1.16 | 0.141 | 0.375 | 1.000 | 13482 | tags=31%, list=23%, signal=40% |
| 3989 | REGULATION\_OF\_GLUTAMATE\_RECEPTOR\_SIGNALING\_PATHWAY(GO:1900449) |  | 39 | 0.73 | 1.16 | 0.141 | 0.375 | 1.000 | 9608 | tags=33%, list=16%, signal=40% |
| 3990 | REGULATION\_OF\_AUTOPHAGY(GO:0010506) |  | 304 | 0.67 | 1.16 | 0.141 | 0.375 | 1.000 | 9745 | tags=10%, list=16%, signal=12% |
| 3991 | RESPONSE\_TO\_ATP(GO:0033198) |  | 31 | 0.71 | 1.16 | 0.000 | 0.375 | 1.000 | 7873 | tags=29%, list=13%, signal=33% |
| 3992 | TRANSITION\_METAL\_ION\_HOMEOSTASIS(GO:0055076) |  | 130 | 0.68 | 1.16 | 0.141 | 0.374 | 1.000 | 9726 | tags=20%, list=16%, signal=24% |
| 3993 | CELLULAR\_RESPONSE\_TO\_MECHANICAL\_STIMULUS(GO:0071260) |  | 78 | 0.72 | 1.16 | 0.141 | 0.374 | 1.000 | 9087 | tags=14%, list=15%, signal=17% |
| 3994 | REGULATION\_OF\_CELL\_CYCLE\_G1\_S\_PHASE\_TRANSITION(GO:1902806) |  | 186 | 0.71 | 1.16 | 0.141 | 0.374 | 1.000 | 10637 | tags=17%, list=18%, signal=20% |
| 3995 | NEUROPEPTIDE\_SIGNALING\_PATHWAY(GO:0007218) |  | 101 | 0.73 | 1.16 | 0.141 | 0.374 | 1.000 | 11744 | tags=39%, list=20%, signal=48% |
| 3996 | NEGATIVE\_REGULATION\_OF\_CELL\_ACTIVATION(GO:0050866) |  | 170 | 0.71 | 1.16 | 0.141 | 0.375 | 1.000 | 10712 | tags=25%, list=18%, signal=30% |
| 3997 | NEGATIVE\_REGULATION\_OF\_CELL\_MORPHOGENESIS\_INVOLVED\_IN\_DIFFERENTIATION(GO:0010771) |  | 79 | 0.72 | 1.16 | 0.141 | 0.375 | 1.000 | 9137 | tags=30%, list=15%, signal=36% |
| 3998 | GLUTAMINE\_FAMILY\_AMINO\_ACID\_CATABOLIC\_PROCESS(GO:0009065) |  | 29 | 0.70 | 1.16 | 0.141 | 0.375 | 1.000 | 11311 | tags=38%, list=19%, signal=47% |
| 3999 | NEGATIVE\_REGULATION\_OF\_RESPONSE\_TO\_ENDOPLASMIC\_RETICULUM\_STRESS(GO:1903573) |  | 43 | 0.68 | 1.16 | 0.000 | 0.375 | 1.000 | 7426 | tags=12%, list=13%, signal=13% |
| 4000 | LABYRINTHINE\_LAYER\_BLOOD\_VESSEL\_DEVELOPMENT(GO:0060716) |  | 20 | 0.70 | 1.16 | 0.141 | 0.374 | 1.000 | 13159 | tags=35%, list=22%, signal=45% |
| 4001 | REGULATION\_OF\_ACTIN\_POLYMERIZATION\_OR\_DEPOLYMERIZATION(GO:0008064) |  | 165 | 0.70 | 1.16 | 0.141 | 0.374 | 1.000 | 8232 | tags=11%, list=14%, signal=13% |
| 4002 | PURINE\_RIBONUCLEOTIDE\_BIOSYNTHETIC\_PROCESS(GO:0009152) |  | 290 | 0.69 | 1.16 | 0.141 | 0.374 | 1.000 | 8541 | tags=13%, list=14%, signal=16% |
| 4003 | POSITIVE\_REGULATION\_OF\_SMALL\_GTPASE\_MEDIATED\_SIGNAL\_TRANSDUCTION(GO:0051057) |  | 58 | 0.72 | 1.16 | 0.141 | 0.374 | 1.000 | 7920 | tags=29%, list=13%, signal=34% |
| 4004 | POSITIVE\_REGULATION\_OF\_CYTOSKELETON\_ORGANIZATION(GO:0051495) |  | 212 | 0.69 | 1.16 | 0.141 | 0.374 | 1.000 | 8232 | tags=14%, list=14%, signal=16% |
| 4005 | REGULATION\_OF\_PEPTIDYL\_TYROSINE\_PHOSPHORYLATION(GO:0050730) |  | 233 | 0.70 | 1.16 | 0.141 | 0.374 | 1.000 | 10131 | tags=21%, list=17%, signal=25% |
| 4006 | REGULATION\_OF\_REGULATED\_SECRETORY\_PATHWAY(GO:1903305) |  | 111 | 0.69 | 1.16 | 0.141 | 0.374 | 1.000 | 9227 | tags=22%, list=16%, signal=26% |
| 4007 | CYTOSKELETON\_DEPENDENT\_CYTOKINESIS(GO:0061640) |  | 66 | 0.67 | 1.16 | 0.000 | 0.374 | 1.000 | 16021 | tags=53%, list=27%, signal=73% |
| 4008 | REGULATION\_OF\_NEUROTRANSMITTER\_RECEPTOR\_ACTIVITY(GO:0099601) |  | 45 | 0.73 | 1.16 | 0.141 | 0.374 | 1.000 | 9608 | tags=31%, list=16%, signal=37% |
| 4009 | REGULATION\_OF\_GLYCOGEN\_BIOSYNTHETIC\_PROCESS(GO:0005979) |  | 27 | 0.74 | 1.16 | 0.141 | 0.374 | 1.000 | 8594 | tags=15%, list=15%, signal=17% |
| 4010 | REGULATION\_OF\_GLUCAN\_BIOSYNTHETIC\_PROCESS(GO:0010962) |  | 27 | 0.74 | 1.16 | 0.141 | 0.374 | 1.000 | 8594 | tags=15%, list=15%, signal=17% |
| 4011 | POSITIVE\_REGULATION\_OF\_CANONICAL\_WNT\_SIGNALING\_PATHWAY(GO:0090263) |  | 137 | 0.68 | 1.16 | 0.141 | 0.374 | 1.000 | 12668 | tags=28%, list=21%, signal=36% |
| 4012 | POTASSIUM\_ION\_HOMEOSTASIS(GO:0055075) |  | 17 | 0.74 | 1.16 | 0.141 | 0.374 | 1.000 | 11400 | tags=35%, list=19%, signal=44% |
| 4013 | POSITIVE\_REGULATION\_OF\_WNT\_SIGNALING\_PATHWAY(GO:0030177) |  | 171 | 0.68 | 1.16 | 0.141 | 0.374 | 1.000 | 12668 | tags=32%, list=21%, signal=41% |
| 4014 | POSITIVE\_REGULATION\_OF\_MUSCLE\_CELL\_APOPTOTIC\_PROCESS(GO:0010661) |  | 32 | 0.70 | 1.16 | 0.000 | 0.374 | 1.000 | 6816 | tags=19%, list=12%, signal=21% |
| 4015 | REGULATION\_OF\_LYMPHOCYTE\_ACTIVATION(GO:0051249) |  | 451 | 0.69 | 1.16 | 0.141 | 0.374 | 1.000 | 9411 | tags=17%, list=16%, signal=20% |
| 4016 | POSITIVE\_REGULATION\_OF\_PEPTIDYL\_TYROSINE\_PHOSPHORYLATION(GO:0050731) |  | 174 | 0.71 | 1.16 | 0.141 | 0.374 | 1.000 | 9784 | tags=22%, list=17%, signal=27% |
| 4017 | NEGATIVE\_REGULATION\_OF\_METAPHASE\_ANAPHASE\_TRANSITION\_OF\_CELL\_CYCLE(GO:1902100) |  | 34 | 0.67 | 1.16 | 0.000 | 0.374 | 1.000 | 15943 | tags=62%, list=27%, signal=85% |
| 4018 | NEGATIVE\_REGULATION\_OF\_MITOTIC\_METAPHASE\_ANAPHASE\_TRANSITION(GO:0045841) |  | 33 | 0.67 | 1.16 | 0.000 | 0.374 | 1.000 | 15943 | tags=61%, list=27%, signal=83% |
| 4019 | CARBOHYDRATE\_DERIVATIVE\_BIOSYNTHETIC\_PROCESS(GO:1901137) |  | 773 | 0.69 | 1.16 | 0.141 | 0.374 | 1.000 | 11522 | tags=24%, list=19%, signal=30% |
| 4020 | REGULATION\_OF\_CELL\_JUNCTION\_ASSEMBLY(GO:1901888) |  | 81 | 0.72 | 1.16 | 0.141 | 0.374 | 1.000 | 7711 | tags=11%, list=13%, signal=13% |
| 4021 | RESPONSE\_TO\_PROGESTERONE(GO:0032570) |  | 44 | 0.72 | 1.16 | 0.141 | 0.374 | 1.000 | 9111 | tags=14%, list=15%, signal=16% |
| 4022 | KILLING\_OF\_CELLS\_OF\_OTHER\_ORGANISM(GO:0031640) |  | 59 | 0.69 | 1.16 | 0.141 | 0.374 | 1.000 | 9039 | tags=14%, list=15%, signal=16% |
| 4023 | DISRUPTION\_OF\_CELLS\_OF\_OTHER\_ORGANISM(GO:0044364) |  | 59 | 0.69 | 1.16 | 0.141 | 0.374 | 1.000 | 9039 | tags=14%, list=15%, signal=16% |
| 4024 | POSITIVE\_REGULATION\_OF\_DENDRITIC\_SPINE\_DEVELOPMENT(GO:0060999) |  | 39 | 0.68 | 1.16 | 0.000 | 0.374 | 1.000 | 12327 | tags=36%, list=21%, signal=45% |
| 4025 | REGULATION\_OF\_LYMPHOCYTE\_APOPTOTIC\_PROCESS(GO:0070228) |  | 54 | 0.71 | 1.16 | 0.141 | 0.374 | 1.000 | 13741 | tags=28%, list=23%, signal=36% |
| 4026 | PROTEIN\_MATURATION(GO:0051604) |  | 374 | 0.69 | 1.16 | 0.141 | 0.374 | 1.000 | 11570 | tags=23%, list=20%, signal=28% |
| 4027 | MONOCARBOXYLIC\_ACID\_CATABOLIC\_PROCESS(GO:0072329) |  | 124 | 0.67 | 1.16 | 0.000 | 0.374 | 1.000 | 12718 | tags=25%, list=22%, signal=32% |
| 4028 | POSITIVE\_REGULATION\_OF\_LIPID\_TRANSPORT(GO:0032370) |  | 54 | 0.71 | 1.16 | 0.141 | 0.374 | 1.000 | 12117 | tags=33%, list=20%, signal=42% |
| 4029 | CYSTEINE\_TYPE\_PEPTIDASE\_ACTIVITY(GO:0008234) |  | 206 | 0.67 | 1.16 | 0.000 | 0.374 | 1.000 | 12385 | tags=26%, list=21%, signal=33% |
| 4030 | POSITIVE\_REGULATION\_OF\_CYCLASE\_ACTIVITY(GO:0031281) |  | 21 | 0.68 | 1.16 | 0.000 | 0.374 | 1.000 | 14425 | tags=33%, list=24%, signal=44% |
| 4031 | GAMETE\_GENERATION(GO:0007276) |  | 665 | 0.70 | 1.16 | 0.141 | 0.374 | 1.000 | 13524 | tags=35%, list=23%, signal=45% |
| 4032 | DNA\_RECOMBINATION(GO:0006310) |  | 257 | 0.67 | 1.16 | 0.141 | 0.374 | 1.000 | 13260 | tags=40%, list=22%, signal=51% |
| 4033 | RESPONSE\_TO\_MURAMYL\_DIPEPTIDE(GO:0032495) |  | 16 | 0.72 | 1.16 | 0.141 | 0.374 | 1.000 | 6199 | tags=6%, list=10%, signal=7% |
| 4034 | DEVELOPMENTAL\_CELL\_GROWTH(GO:0048588) |  | 206 | 0.71 | 1.16 | 0.141 | 0.374 | 1.000 | 8949 | tags=24%, list=15%, signal=28% |
| 4035 | REGULATION\_OF\_DIGESTIVE\_SYSTEM\_PROCESS(GO:0044058) |  | 40 | 0.70 | 1.16 | 0.141 | 0.374 | 1.000 | 10458 | tags=35%, list=18%, signal=42% |
| 4036 | PHOSPHATIDYLINOSITOL\_PHOSPHOLIPASE\_C\_ACTIVITY(GO:0004435) |  | 27 | 0.69 | 1.16 | 0.141 | 0.374 | 1.000 | 9042 | tags=26%, list=15%, signal=31% |
| 4037 | VITAMIN\_TRANSPORT(GO:0051180) |  | 37 | 0.74 | 1.16 | 0.141 | 0.374 | 1.000 | 11359 | tags=35%, list=19%, signal=43% |
| 4038 | MAST\_CELL\_ACTIVATION(GO:0045576) |  | 57 | 0.70 | 1.16 | 0.141 | 0.374 | 1.000 | 10643 | tags=26%, list=18%, signal=32% |
| 4039 | REGULATION\_OF\_PROTEIN\_PROCESSING(GO:0070613) |  | 166 | 0.69 | 1.16 | 0.141 | 0.374 | 1.000 | 13179 | tags=25%, list=22%, signal=32% |
| 4040 | CILIARY\_TRANSITION\_ZONE(GO:0035869) |  | 60 | 0.67 | 1.16 | 0.000 | 0.374 | 1.000 | 13542 | tags=58%, list=23%, signal=76% |
| 4041 | MYELOID\_CELL\_DEVELOPMENT(GO:0061515) |  | 65 | 0.70 | 1.16 | 0.141 | 0.374 | 1.000 | 10768 | tags=32%, list=18%, signal=39% |
| 4042 | NUCLEOTIDASE\_ACTIVITY(GO:0008252) |  | 16 | 0.68 | 1.16 | 0.000 | 0.374 | 1.000 | 14787 | tags=56%, list=25%, signal=75% |
| 4043 | NEGATIVE\_REGULATION\_OF\_CHROMOSOME\_SEPARATION(GO:1905819) |  | 36 | 0.67 | 1.16 | 0.000 | 0.374 | 1.000 | 15943 | tags=61%, list=27%, signal=84% |
| 4044 | NEGATIVE\_REGULATION\_OF\_MITOTIC\_SISTER\_CHROMATID\_SEPARATION(GO:2000816) |  | 35 | 0.67 | 1.16 | 0.000 | 0.374 | 1.000 | 15943 | tags=60%, list=27%, signal=82% |
| 4045 | POSITIVE\_REGULATION\_OF\_CALCIUM\_ION\_IMPORT(GO:0090280) |  | 19 | 0.77 | 1.16 | 0.141 | 0.374 | 1.000 | 9496 | tags=11%, list=16%, signal=13% |
| 4046 | REGULATION\_OF\_PROTEIN\_BINDING(GO:0043393) |  | 206 | 0.69 | 1.16 | 0.141 | 0.373 | 1.000 | 9311 | tags=19%, list=16%, signal=22% |
| 4047 | POSTTRANSCRIPTIONAL\_GENE\_SILENCING\_BY\_RNA(GO:0035194) |  | 326 | 0.69 | 1.16 | 0.141 | 0.373 | 1.000 | 5858 | tags=6%, list=10%, signal=6% |
| 4048 | AXIS\_ELONGATION(GO:0003401) |  | 31 | 0.77 | 1.16 | 0.141 | 0.373 | 1.000 | 9208 | tags=35%, list=16%, signal=42% |
| 4049 | PURINE\_NUCLEOTIDE\_BIOSYNTHETIC\_PROCESS(GO:0006164) |  | 294 | 0.68 | 1.16 | 0.141 | 0.373 | 1.000 | 11574 | tags=22%, list=20%, signal=27% |
| 4050 | REGULATION\_OF\_PROTEIN\_MATURATION(GO:1903317) |  | 168 | 0.69 | 1.16 | 0.141 | 0.373 | 1.000 | 13179 | tags=25%, list=22%, signal=32% |
| 4051 | SPHINGOLIPID\_BIOSYNTHETIC\_PROCESS(GO:0030148) |  | 98 | 0.70 | 1.16 | 0.141 | 0.373 | 1.000 | 8128 | tags=17%, list=14%, signal=20% |
| 4052 | POSITIVE\_REGULATION\_OF\_CIRCADIAN\_RHYTHM(GO:0042753) |  | 17 | 0.73 | 1.16 | 0.000 | 0.373 | 1.000 | 13103 | tags=59%, list=22%, signal=76% |
| 4053 | PURINE\_CONTAINING\_COMPOUND\_BIOSYNTHETIC\_PROCESS(GO:0072522) |  | 306 | 0.68 | 1.16 | 0.141 | 0.373 | 1.000 | 11574 | tags=22%, list=20%, signal=27% |
| 4054 | POSITIVE\_REGULATION\_OF\_GLUCOSE\_TRANSMEMBRANE\_TRANSPORT(GO:0010828) |  | 50 | 0.67 | 1.16 | 0.141 | 0.373 | 1.000 | 8594 | tags=14%, list=15%, signal=16% |
| 4055 | INTERLEUKIN\_2\_BIOSYNTHETIC\_PROCESS(GO:0042094) |  | 21 | 0.73 | 1.16 | 0.141 | 0.373 | 1.000 | 9574 | tags=38%, list=16%, signal=45% |
| 4056 | REGULATION\_OF\_ACTIN\_FILAMENT\_LENGTH(GO:0030832) |  | 166 | 0.70 | 1.16 | 0.141 | 0.373 | 1.000 | 8232 | tags=11%, list=14%, signal=13% |
| 4057 | INTRINSIC\_COMPONENT\_OF\_ORGANELLE\_MEMBRANE(GO:0031300) |  | 193 | 0.67 | 1.16 | 0.141 | 0.373 | 1.000 | 13102 | tags=28%, list=22%, signal=36% |
| 4058 | REGULATION\_OF\_CHROMOSOME\_ORGANIZATION(GO:0033044) |  | 310 | 0.66 | 1.16 | 0.000 | 0.373 | 1.000 | 14015 | tags=36%, list=24%, signal=47% |
| 4059 | SERTOLI\_CELL\_DIFFERENTIATION(GO:0060008) |  | 21 | 0.71 | 1.16 | 0.141 | 0.373 | 1.000 | 11480 | tags=38%, list=19%, signal=47% |
| 4060 | FEMALE\_MEIOTIC\_NUCLEAR\_DIVISION(GO:0007143) |  | 28 | 0.73 | 1.16 | 0.141 | 0.373 | 1.000 | 9128 | tags=18%, list=15%, signal=21% |
| 4061 | CELLULAR\_AMINO\_ACID\_METABOLIC\_PROCESS(GO:0006520) |  | 379 | 0.70 | 1.16 | 0.141 | 0.373 | 1.000 | 12125 | tags=27%, list=21%, signal=34% |
| 4062 | POSITIVE\_REGULATION\_OF\_INTERLEUKIN\_8\_PRODUCTION(GO:0032757) |  | 46 | 0.69 | 1.16 | 0.141 | 0.373 | 1.000 | 6626 | tags=13%, list=11%, signal=15% |
| 4063 | NON\_CANONICAL\_WNT\_SIGNALING\_PATHWAY(GO:0035567) |  | 152 | 0.68 | 1.16 | 0.141 | 0.373 | 1.000 | 12594 | tags=29%, list=21%, signal=37% |
| 4064 | POSITIVE\_REGULATION\_OF\_GLIAL\_CELL\_DIFFERENTIATION(GO:0045687) |  | 37 | 0.72 | 1.16 | 0.141 | 0.373 | 1.000 | 11343 | tags=32%, list=19%, signal=40% |
| 4065 | MEIOSIS\_I\_CELL\_CYCLE\_PROCESS(GO:0061982) |  | 116 | 0.69 | 1.16 | 0.141 | 0.373 | 1.000 | 11304 | tags=34%, list=19%, signal=41% |
| 4066 | CORE\_PROMOTER\_SEQUENCE\_SPECIFIC\_DNA\_BINDING(GO:0001046) |  | 82 | 0.72 | 1.16 | 0.141 | 0.373 | 1.000 | 7800 | tags=10%, list=13%, signal=11% |
| 4067 | CHROMATIN\_ORGANIZATION(GO:0006325) |  | 779 | 0.70 | 1.16 | 0.141 | 0.373 | 1.000 | 10856 | tags=18%, list=18%, signal=22% |
| 4068 | SODIUM\_INDEPENDENT\_ORGANIC\_ANION\_TRANSMEMBRANE\_TRANSPORTER\_ACTIVITY(GO:0015347) |  | 25 | 0.73 | 1.16 | 0.141 | 0.373 | 1.000 | 11359 | tags=40%, list=19%, signal=49% |
| 4069 | REGULATION\_OF\_CELLULAR\_AMIDE\_METABOLIC\_PROCESS(GO:0034248) |  | 651 | 0.68 | 1.16 | 0.141 | 0.373 | 1.000 | 8746 | tags=8%, list=15%, signal=9% |
| 4070 | OXIDOREDUCTASE\_ACTIVITY\_\_ACTING\_ON\_THE\_CH\_NH\_GROUP\_OF\_DONORS\_\_NAD\_OR\_NADP\_AS\_ACCEPTOR(GO:0016646) |  | 18 | 0.70 | 1.16 | 0.141 | 0.373 | 1.000 | 14192 | tags=50%, list=24%, signal=66% |
| 4071 | CHROMOSOMAL\_PART(GO:0044427) |  | 884 | 0.70 | 1.16 | 0.141 | 0.373 | 1.000 | 13260 | tags=33%, list=22%, signal=43% |
| 4072 | POSTTRANSCRIPTIONAL\_GENE\_SILENCING(GO:0016441) |  | 327 | 0.69 | 1.16 | 0.141 | 0.373 | 1.000 | 5858 | tags=6%, list=10%, signal=6% |
| 4073 | B\_CELL\_ACTIVATION(GO:0042113) |  | 279 | 0.68 | 1.16 | 0.141 | 0.373 | 1.000 | 10861 | tags=23%, list=18%, signal=28% |
| 4074 | NEGATIVE\_REGULATION\_OF\_CELL\_ADHESION(GO:0007162) |  | 247 | 0.71 | 1.16 | 0.141 | 0.373 | 1.000 | 10000 | tags=19%, list=17%, signal=22% |
| 4075 | ACTIN\_POLYMERIZATION\_OR\_DEPOLYMERIZATION(GO:0008154) |  | 190 | 0.69 | 1.16 | 0.141 | 0.373 | 1.000 | 9686 | tags=14%, list=16%, signal=17% |
| 4076 | ORGANIC\_ANION\_TRANSMEMBRANE\_TRANSPORTER\_ACTIVITY(GO:0008514) |  | 226 | 0.71 | 1.16 | 0.141 | 0.373 | 1.000 | 11579 | tags=33%, list=20%, signal=41% |
| 4077 | CHOLESTEROL\_TRANSPORT(GO:0030301) |  | 85 | 0.71 | 1.16 | 0.141 | 0.373 | 1.000 | 11906 | tags=28%, list=20%, signal=35% |
| 4078 | PROTEIN\_PHOSPHATASE\_1\_BINDING(GO:0008157) |  | 19 | 0.70 | 1.16 | 0.000 | 0.373 | 1.000 | 11571 | tags=42%, list=20%, signal=52% |
| 4079 | REGULATION\_OF\_B\_CELL\_ACTIVATION(GO:0050864) |  | 161 | 0.69 | 1.16 | 0.141 | 0.373 | 1.000 | 11130 | tags=19%, list=19%, signal=24% |
| 4080 | NEGATIVE\_REGULATION\_OF\_RNA\_CATABOLIC\_PROCESS(GO:1902369) |  | 50 | 0.74 | 1.16 | 0.141 | 0.373 | 1.000 | 5906 | tags=2%, list=10%, signal=2% |
| 4081 | CELLULAR\_RESPONSE\_TO\_NITRIC\_OXIDE(GO:0071732) |  | 16 | 0.69 | 1.16 | 0.000 | 0.373 | 1.000 | 15560 | tags=56%, list=26%, signal=76% |
| 4082 | ANTIBACTERIAL\_HUMORAL\_RESPONSE(GO:0019731) |  | 43 | 0.72 | 1.16 | 0.141 | 0.373 | 1.000 | 10916 | tags=14%, list=18%, signal=17% |
| 4083 | RIBONUCLEOSIDE\_MONOPHOSPHATE\_BIOSYNTHETIC\_PROCESS(GO:0009156) |  | 181 | 0.69 | 1.16 | 0.141 | 0.373 | 1.000 | 8541 | tags=9%, list=14%, signal=10% |
| 4084 | DENDRITIC\_SPINE\_MORPHOGENESIS(GO:0060997) |  | 49 | 0.67 | 1.16 | 0.141 | 0.373 | 1.000 | 15406 | tags=45%, list=26%, signal=61% |
| 4085 | CELL\_CYCLE(GO:0007049) |  | 1831 | 0.70 | 1.16 | 0.141 | 0.373 | 1.000 | 13128 | tags=33%, list=22%, signal=41% |
| 4086 | REGULATION\_OF\_SYNAPTIC\_TRANSMISSION\_\_GLUTAMATERGIC(GO:0051966) |  | 60 | 0.71 | 1.16 | 0.141 | 0.373 | 1.000 | 8759 | tags=27%, list=15%, signal=31% |
| 4087 | REGULATION\_OF\_COLLATERAL\_SPROUTING(GO:0048670) |  | 20 | 0.75 | 1.16 | 0.141 | 0.373 | 1.000 | 8673 | tags=40%, list=15%, signal=47% |
| 4088 | NON\_RECOMBINATIONAL\_REPAIR(GO:0000726) |  | 83 | 0.71 | 1.16 | 0.141 | 0.373 | 1.000 | 10674 | tags=17%, list=18%, signal=21% |
| 4089 | NUCLEAR\_HETEROCHROMATIN(GO:0005720) |  | 35 | 0.67 | 1.16 | 0.000 | 0.373 | 1.000 | 13983 | tags=43%, list=24%, signal=56% |
| 4090 | CELL\_BODY\_MEMBRANE(GO:0044298) |  | 21 | 0.70 | 1.16 | 0.141 | 0.373 | 1.000 | 12773 | tags=33%, list=22%, signal=43% |
| 4091 | NEURONAL\_CELL\_BODY\_MEMBRANE(GO:0032809) |  | 21 | 0.70 | 1.16 | 0.141 | 0.373 | 1.000 | 12773 | tags=33%, list=22%, signal=43% |
| 4092 | REGULATION\_OF\_TRANSFORMING\_GROWTH\_FACTOR\_BETA\_RECEPTOR\_SIGNALING\_PATHWAY(GO:0017015) |  | 108 | 0.71 | 1.16 | 0.141 | 0.373 | 1.000 | 7159 | tags=12%, list=12%, signal=14% |
| 4093 | CYTOLYSIS(GO:0019835) |  | 43 | 0.69 | 1.16 | 0.141 | 0.373 | 1.000 | 11397 | tags=23%, list=19%, signal=29% |
| 4094 | POSITIVE\_REGULATION\_OF\_CYTOKINE\_BIOSYNTHETIC\_PROCESS(GO:0042108) |  | 58 | 0.72 | 1.16 | 0.141 | 0.373 | 1.000 | 9016 | tags=26%, list=15%, signal=30% |
| 4095 | HETEROTYPIC\_CELL\_CELL\_ADHESION(GO:0034113) |  | 50 | 0.70 | 1.16 | 0.141 | 0.373 | 1.000 | 10458 | tags=26%, list=18%, signal=32% |
| 4096 | REGULATION\_OF\_COLLAGEN\_BIOSYNTHETIC\_PROCESS(GO:0032965) |  | 39 | 0.68 | 1.16 | 0.141 | 0.373 | 1.000 | 11254 | tags=23%, list=19%, signal=28% |
| 4097 | VASCULAR\_SMOOTH\_MUSCLE\_CONTRACTION(GO:0014829) |  | 23 | 0.73 | 1.15 | 0.141 | 0.373 | 1.000 | 14175 | tags=61%, list=24%, signal=80% |
| 4098 | ORGANIC\_SUBSTANCE\_TRANSPORT(GO:0071702) |  | 2626 | 0.71 | 1.15 | 0.141 | 0.373 | 1.000 | 9686 | tags=16%, list=16%, signal=19% |
| 4099 | POSITIVE\_REGULATION\_OF\_LYASE\_ACTIVITY(GO:0051349) |  | 21 | 0.69 | 1.15 | 0.000 | 0.373 | 1.000 | 11031 | tags=24%, list=19%, signal=29% |
| 4100 | T\_CELL\_HOMEOSTASIS(GO:0043029) |  | 39 | 0.67 | 1.15 | 0.141 | 0.373 | 1.000 | 11567 | tags=21%, list=20%, signal=25% |
| 4101 | ESTROGEN\_RECEPTOR\_BINDING(GO:0030331) |  | 40 | 0.72 | 1.15 | 0.141 | 0.373 | 1.000 | 12546 | tags=35%, list=21%, signal=44% |
| 4102 | REGULATION\_OF\_LIPID\_LOCALIZATION(GO:1905952) |  | 137 | 0.69 | 1.15 | 0.141 | 0.373 | 1.000 | 13676 | tags=30%, list=23%, signal=39% |
| 4103 | LYTIC\_VACUOLE\_MEMBRANE(GO:0098852) |  | 346 | 0.69 | 1.15 | 0.141 | 0.373 | 1.000 | 6602 | tags=5%, list=11%, signal=6% |
| 4104 | LYSOSOMAL\_MEMBRANE(GO:0005765) |  | 346 | 0.69 | 1.15 | 0.141 | 0.373 | 1.000 | 6602 | tags=5%, list=11%, signal=6% |
| 4105 | MOTOR\_ACTIVITY(GO:0003774) |  | 143 | 0.67 | 1.15 | 0.141 | 0.373 | 1.000 | 13157 | tags=41%, list=22%, signal=52% |
| 4106 | PEROXISOMAL\_PART(GO:0044439) |  | 91 | 0.68 | 1.15 | 0.141 | 0.373 | 1.000 | 13455 | tags=36%, list=23%, signal=47% |
| 4107 | MICROBODY\_PART(GO:0044438) |  | 91 | 0.68 | 1.15 | 0.141 | 0.373 | 1.000 | 13455 | tags=36%, list=23%, signal=47% |
| 4108 | POSITIVE\_REGULATION\_OF\_PATHWAY\_RESTRICTED\_SMAD\_PROTEIN\_PHOSPHORYLATION(GO:0010862) |  | 48 | 0.70 | 1.15 | 0.141 | 0.372 | 1.000 | 13279 | tags=44%, list=22%, signal=56% |
| 4109 | CYTOSKELETAL\_PART(GO:0044430) |  | 1592 | 0.71 | 1.15 | 0.141 | 0.372 | 1.000 | 10939 | tags=24%, list=18%, signal=29% |
| 4110 | REGULATION\_OF\_GLYCOGEN\_METABOLIC\_PROCESS(GO:0070873) |  | 32 | 0.73 | 1.15 | 0.141 | 0.372 | 1.000 | 8594 | tags=16%, list=15%, signal=18% |
| 4111 | REGULATION\_OF\_MRNA\_SPLICING\_\_VIA\_SPLICEOSOME(GO:0048024) |  | 76 | 0.67 | 1.15 | 0.000 | 0.372 | 1.000 | 17646 | tags=74%, list=30%, signal=105% |
| 4112 | REGULATION\_OF\_ESTABLISHMENT\_OF\_PROTEIN\_LOCALIZATION\_TO\_MITOCHONDRION(GO:1903747) |  | 125 | 0.72 | 1.15 | 0.141 | 0.372 | 1.000 | 7583 | tags=9%, list=13%, signal=10% |
| 4113 | RIBOSE\_PHOSPHATE\_BIOSYNTHETIC\_PROCESS(GO:0046390) |  | 306 | 0.68 | 1.15 | 0.141 | 0.372 | 1.000 | 11574 | tags=21%, list=20%, signal=26% |
| 4114 | NEGATIVE\_REGULATION\_OF\_TORC1\_SIGNALING(GO:1904262) |  | 15 | 0.71 | 1.15 | 0.141 | 0.372 | 1.000 | 11929 | tags=33%, list=20%, signal=42% |
| 4115 | REGULATION\_OF\_POLYSACCHARIDE\_BIOSYNTHETIC\_PROCESS(GO:0032885) |  | 34 | 0.73 | 1.15 | 0.141 | 0.372 | 1.000 | 8594 | tags=15%, list=15%, signal=17% |
| 4116 | REGULATION\_OF\_DENDRITIC\_SPINE\_MORPHOGENESIS(GO:0061001) |  | 36 | 0.69 | 1.15 | 0.141 | 0.372 | 1.000 | 8450 | tags=22%, list=14%, signal=26% |
| 4117 | PROTEIN\_LOCALIZATION\_TO\_KINETOCHORE(GO:0034501) |  | 18 | 0.70 | 1.15 | 0.000 | 0.372 | 1.000 | 14424 | tags=78%, list=24%, signal=103% |
| 4118 | CELLULAR\_RESPONSE\_TO\_CAMP(GO:0071320) |  | 53 | 0.68 | 1.15 | 0.141 | 0.372 | 1.000 | 9633 | tags=28%, list=16%, signal=34% |
| 4119 | POSITIVE\_REGULATION\_OF\_UBIQUITIN\_DEPENDENT\_PROTEIN\_CATABOLIC\_PROCESS(GO:2000060) |  | 94 | 0.68 | 1.15 | 0.141 | 0.372 | 1.000 | 7795 | tags=12%, list=13%, signal=13% |
| 4120 | POSITIVE\_REGULATION\_OF\_DENDRITE\_MORPHOGENESIS(GO:0050775) |  | 32 | 0.67 | 1.15 | 0.000 | 0.372 | 1.000 | 8450 | tags=16%, list=14%, signal=18% |
| 4121 | TETRAPYRROLE\_BINDING(GO:0046906) |  | 146 | 0.68 | 1.15 | 0.141 | 0.372 | 1.000 | 11264 | tags=30%, list=19%, signal=37% |
| 4122 | MYELOID\_LEUKOCYTE\_DIFFERENTIATION(GO:0002573) |  | 194 | 0.71 | 1.15 | 0.141 | 0.372 | 1.000 | 10901 | tags=27%, list=18%, signal=33% |
| 4123 | REGULATION\_OF\_EPITHELIAL\_CELL\_APOPTOTIC\_PROCESS(GO:1904035) |  | 81 | 0.71 | 1.15 | 0.141 | 0.372 | 1.000 | 9016 | tags=16%, list=15%, signal=19% |
| 4124 | NEGATIVE\_REGULATION\_OF\_NUCLEAR\_DIVISION(GO:0051784) |  | 59 | 0.66 | 1.15 | 0.000 | 0.372 | 1.000 | 15968 | tags=59%, list=27%, signal=81% |
| 4125 | TRANS\_GOLGI\_NETWORK\_TRANSPORT\_VESICLE(GO:0030140) |  | 29 | 0.68 | 1.15 | 0.141 | 0.372 | 1.000 | 8452 | tags=17%, list=14%, signal=20% |
| 4126 | VACUOLAR\_PART(GO:0044437) |  | 555 | 0.68 | 1.15 | 0.141 | 0.372 | 1.000 | 12215 | tags=20%, list=21%, signal=25% |
| 4127 | POSITIVE\_REGULATION\_OF\_SEQUESTERING\_OF\_CALCIUM\_ION(GO:0051284) |  | 16 | 0.71 | 1.15 | 0.141 | 0.372 | 1.000 | 10116 | tags=38%, list=17%, signal=45% |
| 4128 | DNA\_METHYLATION\_OR\_DEMETHYLATION(GO:0044728) |  | 88 | 0.72 | 1.15 | 0.141 | 0.372 | 1.000 | 8202 | tags=11%, list=14%, signal=13% |
| 4129 | POSITIVE\_REGULATION\_OF\_MITOCHONDRION\_ORGANIZATION(GO:0010822) |  | 166 | 0.71 | 1.15 | 0.141 | 0.372 | 1.000 | 7782 | tags=8%, list=13%, signal=10% |
| 4130 | RIBONUCLEOTIDE\_BIOSYNTHETIC\_PROCESS(GO:0009260) |  | 303 | 0.68 | 1.15 | 0.141 | 0.372 | 1.000 | 11574 | tags=21%, list=20%, signal=26% |
| 4131 | VIRAL\_LIFE\_CYCLE(GO:0019058) |  | 314 | 0.69 | 1.15 | 0.141 | 0.372 | 1.000 | 8541 | tags=12%, list=14%, signal=14% |
| 4132 | MITOTIC\_SPINDLE\_CHECKPOINT(GO:0071174) |  | 32 | 0.67 | 1.15 | 0.000 | 0.372 | 1.000 | 14793 | tags=53%, list=25%, signal=71% |
| 4133 | SPINDLE\_ASSEMBLY\_CHECKPOINT(GO:0071173) |  | 32 | 0.67 | 1.15 | 0.000 | 0.372 | 1.000 | 14793 | tags=53%, list=25%, signal=71% |
| 4134 | MITOTIC\_SPINDLE\_ASSEMBLY\_CHECKPOINT(GO:0007094) |  | 32 | 0.67 | 1.15 | 0.000 | 0.372 | 1.000 | 14793 | tags=53%, list=25%, signal=71% |
| 4135 | RETINAL\_BINDING(GO:0016918) |  | 16 | 0.75 | 1.15 | 0.141 | 0.372 | 1.000 | 8521 | tags=38%, list=14%, signal=44% |
| 4136 | REGULATION\_OF\_INTERLEUKIN\_1\_BETA\_SECRETION(GO:0050706) |  | 39 | 0.72 | 1.15 | 0.141 | 0.372 | 1.000 | 12242 | tags=18%, list=21%, signal=23% |
| 4137 | HUMORAL\_IMMUNE\_RESPONSE\_MEDIATED\_BY\_CIRCULATING\_IMMUNOGLOBULIN(GO:0002455) |  | 144 | 0.69 | 1.15 | 0.141 | 0.372 | 1.000 | 11910 | tags=8%, list=20%, signal=10% |
| 4138 | METANEPHRIC\_EPITHELIUM\_DEVELOPMENT(GO:0072207) |  | 27 | 0.74 | 1.15 | 0.141 | 0.372 | 1.000 | 13927 | tags=41%, list=24%, signal=53% |
| 4139 | DEVELOPMENTAL\_GROWTH\_INVOLVED\_IN\_MORPHOGENESIS(GO:0060560) |  | 214 | 0.71 | 1.15 | 0.141 | 0.372 | 1.000 | 11111 | tags=31%, list=19%, signal=38% |
| 4140 | MEIOTIC\_NUCLEAR\_DIVISION(GO:0140013) |  | 172 | 0.68 | 1.15 | 0.141 | 0.372 | 1.000 | 13476 | tags=42%, list=23%, signal=55% |
| 4141 | ACETYLCHOLINE\_BINDING(GO:0042166) |  | 17 | 0.72 | 1.15 | 0.141 | 0.372 | 1.000 | 12401 | tags=53%, list=21%, signal=67% |
| 4142 | NEGATIVE\_REGULATION\_OF\_MYOTUBE\_DIFFERENTIATION(GO:0010832) |  | 17 | 0.72 | 1.15 | 0.141 | 0.372 | 1.000 | 8774 | tags=24%, list=15%, signal=28% |
| 4143 | REGULATION\_OF\_CYCLIC\_NUCLEOTIDE\_METABOLIC\_PROCESS(GO:0030799) |  | 102 | 0.69 | 1.15 | 0.141 | 0.372 | 1.000 | 8279 | tags=20%, list=14%, signal=23% |
| 4144 | NUCLEAR\_CHROMOSOME\_PART(GO:0044454) |  | 540 | 0.69 | 1.15 | 0.141 | 0.372 | 1.000 | 12135 | tags=26%, list=21%, signal=33% |
| 4145 | POSITIVE\_REGULATION\_OF\_CELL\_CYCLE\_PROCESS(GO:0090068) |  | 269 | 0.70 | 1.15 | 0.141 | 0.372 | 1.000 | 10898 | tags=20%, list=18%, signal=24% |
| 4146 | SINGLE\_STRANDED\_RNA\_BINDING(GO:0003727) |  | 79 | 0.67 | 1.15 | 0.000 | 0.372 | 1.000 | 15297 | tags=47%, list=26%, signal=63% |
| 4147 | PROLINE\_RICH\_REGION\_BINDING(GO:0070064) |  | 19 | 0.70 | 1.15 | 0.000 | 0.372 | 1.000 | 11939 | tags=32%, list=20%, signal=40% |
| 4148 | CHEMOKINE\_SECRETION(GO:0090195) |  | 16 | 0.72 | 1.15 | 0.141 | 0.372 | 1.000 | 6258 | tags=38%, list=11%, signal=42% |
| 4149 | POSITIVE\_REGULATION\_OF\_PROTEIN\_POLYMERIZATION(GO:0032273) |  | 123 | 0.68 | 1.15 | 0.141 | 0.372 | 1.000 | 9686 | tags=15%, list=16%, signal=18% |
| 4150 | SPINDLE\_CHECKPOINT(GO:0031577) |  | 35 | 0.67 | 1.15 | 0.000 | 0.371 | 1.000 | 14793 | tags=54%, list=25%, signal=72% |
| 4151 | PLATELET\_DEGRANULATION(GO:0002576) |  | 126 | 0.69 | 1.15 | 0.141 | 0.371 | 1.000 | 10860 | tags=25%, list=18%, signal=31% |
| 4152 | AZOLE\_TRANSPORT(GO:0045117) |  | 21 | 0.71 | 1.15 | 0.141 | 0.371 | 1.000 | 13990 | tags=52%, list=24%, signal=69% |
| 4153 | DETECTION\_OF\_ABIOTIC\_STIMULUS(GO:0009582) |  | 135 | 0.69 | 1.15 | 0.141 | 0.372 | 1.000 | 12044 | tags=36%, list=20%, signal=45% |
| 4154 | KINESIN\_COMPLEX(GO:0005871) |  | 55 | 0.67 | 1.15 | 0.000 | 0.372 | 1.000 | 13386 | tags=45%, list=23%, signal=59% |
| 4155 | AXONEMAL\_DYNEIN\_COMPLEX\_ASSEMBLY(GO:0070286) |  | 32 | 0.68 | 1.15 | 0.141 | 0.372 | 1.000 | 14289 | tags=53%, list=24%, signal=70% |
| 4156 | BETA\_CATENIN\_BINDING(GO:0008013) |  | 83 | 0.73 | 1.15 | 0.141 | 0.372 | 1.000 | 7414 | tags=17%, list=13%, signal=19% |
| 4157 | UNSATURATED\_FATTY\_ACID\_BIOSYNTHETIC\_PROCESS(GO:0006636) |  | 43 | 0.70 | 1.15 | 0.141 | 0.372 | 1.000 | 8876 | tags=23%, list=15%, signal=27% |
| 4158 | MITOTIC\_CELL\_CYCLE\_PROCESS(GO:1903047) |  | 837 | 0.69 | 1.15 | 0.141 | 0.372 | 1.000 | 12586 | tags=30%, list=21%, signal=37% |
| 4159 | METANEPHRIC\_TUBULE\_DEVELOPMENT(GO:0072170) |  | 23 | 0.74 | 1.15 | 0.141 | 0.372 | 1.000 | 13927 | tags=43%, list=24%, signal=57% |
| 4160 | FAT\_SOLUBLE\_VITAMIN\_METABOLIC\_PROCESS(GO:0006775) |  | 44 | 0.69 | 1.15 | 0.141 | 0.372 | 1.000 | 13642 | tags=43%, list=23%, signal=56% |
| 4161 | RESPIRATORY\_ELECTRON\_TRANSPORT\_CHAIN(GO:0022904) |  | 117 | 0.65 | 1.15 | 0.000 | 0.372 | 1.000 | 5894 | tags=3%, list=10%, signal=4% |
| 4162 | REGULATION\_OF\_CELLULAR\_RESPONSE\_TO\_TRANSFORMING\_GROWTH\_FACTOR\_BETA\_STIMULUS(GO:1903844) |  | 110 | 0.70 | 1.15 | 0.141 | 0.372 | 1.000 | 7159 | tags=12%, list=12%, signal=13% |
| 4163 | METANEPHRIC\_NEPHRON\_EPITHELIUM\_DEVELOPMENT(GO:0072243) |  | 23 | 0.74 | 1.15 | 0.141 | 0.372 | 1.000 | 13927 | tags=43%, list=24%, signal=57% |
| 4164 | GLUTAMATE\_RECEPTOR\_SIGNALING\_PATHWAY(GO:0007215) |  | 81 | 0.70 | 1.15 | 0.141 | 0.372 | 1.000 | 9608 | tags=30%, list=16%, signal=35% |
| 4165 | RAS\_GUANYL\_NUCLEOTIDE\_EXCHANGE\_FACTOR\_ACTIVITY(GO:0005088) |  | 257 | 0.70 | 1.15 | 0.141 | 0.372 | 1.000 | 12000 | tags=31%, list=20%, signal=39% |
| 4166 | ACTIVIN\_RECEPTOR\_SIGNALING\_PATHWAY(GO:0032924) |  | 39 | 0.70 | 1.15 | 0.141 | 0.372 | 1.000 | 12702 | tags=41%, list=21%, signal=52% |
| 4167 | PROTEIN\_CHROMOPHORE\_LINKAGE(GO:0018298) |  | 15 | 0.74 | 1.15 | 0.141 | 0.371 | 1.000 | 7933 | tags=20%, list=13%, signal=23% |
| 4168 | REGULATION\_OF\_PRODUCTION\_OF\_MOLECULAR\_MEDIATOR\_OF\_IMMUNE\_RESPONSE(GO:0002700) |  | 114 | 0.69 | 1.15 | 0.141 | 0.371 | 1.000 | 11130 | tags=26%, list=19%, signal=32% |
| 4169 | CHROMOSOME(GO:0005694) |  | 1013 | 0.69 | 1.15 | 0.141 | 0.372 | 1.000 | 13260 | tags=34%, list=22%, signal=43% |
| 4170 | CONDENSED\_CHROMOSOME\_KINETOCHORE(GO:0000777) |  | 104 | 0.67 | 1.15 | 0.000 | 0.372 | 1.000 | 14323 | tags=58%, list=24%, signal=76% |
| 4171 | MITOTIC\_CYTOKINESIS(GO:0000281) |  | 52 | 0.67 | 1.15 | 0.000 | 0.372 | 1.000 | 16021 | tags=58%, list=27%, signal=79% |
| 4172 | HYDROLASE\_ACTIVITY(GO:0016787) |  | 2679 | 0.70 | 1.15 | 0.141 | 0.372 | 1.000 | 12125 | tags=25%, list=21%, signal=30% |
| 4173 | PHOSPHOPROTEIN\_PHOSPHATASE\_ACTIVITY(GO:0004721) |  | 192 | 0.67 | 1.15 | 0.141 | 0.372 | 1.000 | 10781 | tags=18%, list=18%, signal=22% |
| 4174 | MITOCHONDRIAL\_TRANSPORT(GO:0006839) |  | 326 | 0.69 | 1.15 | 0.141 | 0.372 | 1.000 | 7583 | tags=7%, list=13%, signal=8% |
| 4175 | RESPIRATORY\_GASEOUS\_EXCHANGE(GO:0007585) |  | 66 | 0.69 | 1.15 | 0.141 | 0.372 | 1.000 | 11650 | tags=36%, list=20%, signal=45% |
| 4176 | LYMPHOCYTE\_HOMEOSTASIS(GO:0002260) |  | 62 | 0.66 | 1.15 | 0.141 | 0.372 | 1.000 | 14695 | tags=34%, list=25%, signal=45% |
| 4177 | PROTEIN\_MANNOSYLATION(GO:0035268) |  | 20 | 0.65 | 1.15 | 0.000 | 0.372 | 1.000 | 15816 | tags=50%, list=27%, signal=68% |
| 4178 | GLUCOSYLTRANSFERASE\_ACTIVITY(GO:0046527) |  | 16 | 0.70 | 1.15 | 0.000 | 0.372 | 1.000 | 15330 | tags=75%, list=26%, signal=101% |
| 4179 | IONOTROPIC\_GLUTAMATE\_RECEPTOR\_SIGNALING\_PATHWAY(GO:0035235) |  | 25 | 0.73 | 1.15 | 0.141 | 0.372 | 1.000 | 9000 | tags=28%, list=15%, signal=33% |
| 4180 | NEGATIVE\_REGULATION\_OF\_CHROMOSOME\_SEGREGATION(GO:0051985) |  | 41 | 0.66 | 1.15 | 0.000 | 0.372 | 1.000 | 16227 | tags=61%, list=27%, signal=84% |
| 4181 | NEGATIVE\_REGULATION\_OF\_SISTER\_CHROMATID\_SEGREGATION(GO:0033046) |  | 40 | 0.66 | 1.15 | 0.000 | 0.372 | 1.000 | 16227 | tags=60%, list=27%, signal=83% |
| 4182 | UBIQUITINYL\_HYDROLASE\_ACTIVITY(GO:0101005) |  | 130 | 0.66 | 1.15 | 0.000 | 0.372 | 1.000 | 12385 | tags=25%, list=21%, signal=32% |
| 4183 | PROTEIN\_GLYCOSYLATION(GO:0006486) |  | 258 | 0.69 | 1.15 | 0.141 | 0.372 | 1.000 | 11789 | tags=25%, list=20%, signal=31% |
| 4184 | MACROMOLECULE\_GLYCOSYLATION(GO:0043413) |  | 258 | 0.69 | 1.15 | 0.141 | 0.372 | 1.000 | 11789 | tags=25%, list=20%, signal=31% |
| 4185 | LABYRINTHINE\_LAYER\_DEVELOPMENT(GO:0060711) |  | 47 | 0.72 | 1.15 | 0.141 | 0.372 | 1.000 | 9659 | tags=26%, list=16%, signal=30% |
| 4186 | NUCLEOBASE\_CONTAINING\_COMPOUND\_TRANSPORT(GO:0015931) |  | 235 | 0.65 | 1.15 | 0.000 | 0.372 | 1.000 | 17041 | tags=57%, list=29%, signal=80% |
| 4187 | NUCLEOSIDE\_MONOPHOSPHATE\_BIOSYNTHETIC\_PROCESS(GO:0009124) |  | 190 | 0.69 | 1.15 | 0.141 | 0.372 | 1.000 | 9272 | tags=11%, list=16%, signal=12% |
| 4188 | TRANSLATION\_REGULATOR\_ACTIVITY\_\_NUCLEIC\_ACID\_BINDING(GO:0090079) |  | 24 | 0.69 | 1.15 | 0.000 | 0.371 | 1.000 | 8668 | tags=17%, list=15%, signal=20% |
| 4189 | POSITIVE\_REGULATION\_OF\_SMOOTH\_MUSCLE\_CONTRACTION(GO:0045987) |  | 34 | 0.71 | 1.15 | 0.141 | 0.371 | 1.000 | 9682 | tags=26%, list=16%, signal=32% |
| 4190 | REGULATION\_OF\_EPITHELIAL\_CELL\_DIFFERENTIATION\_INVOLVED\_IN\_KIDNEY\_DEVELOPMENT(GO:2000696) |  | 17 | 0.76 | 1.15 | 0.141 | 0.371 | 1.000 | 9951 | tags=29%, list=17%, signal=35% |
| 4191 | CARGO\_RECEPTOR\_ACTIVITY(GO:0038024) |  | 83 | 0.71 | 1.15 | 0.141 | 0.371 | 1.000 | 12594 | tags=30%, list=21%, signal=38% |
| 4192 | POSITIVE\_REGULATION\_OF\_CATALYTIC\_ACTIVITY(GO:0043085) |  | 1373 | 0.69 | 1.15 | 0.141 | 0.371 | 1.000 | 11372 | tags=21%, list=19%, signal=26% |
| 4193 | ORGANELLE\_TRANSPORT\_ALONG\_MICROTUBULE(GO:0072384) |  | 70 | 0.67 | 1.15 | 0.000 | 0.371 | 1.000 | 15406 | tags=47%, list=26%, signal=64% |
| 4194 | REGULATION\_OF\_LEUKOCYTE\_CELL\_CELL\_ADHESION(GO:1903037) |  | 290 | 0.67 | 1.15 | 0.141 | 0.371 | 1.000 | 10712 | tags=21%, list=18%, signal=26% |
| 4195 | ORGANELLE\_MEMBRANE(GO:0031090) |  | 2909 | 0.70 | 1.15 | 0.141 | 0.371 | 1.000 | 9677 | tags=15%, list=16%, signal=17% |
| 4196 | DETECTION\_OF\_LIGHT\_STIMULUS\_INVOLVED\_IN\_VISUAL\_PERCEPTION(GO:0050908) |  | 17 | 0.73 | 1.15 | 0.141 | 0.371 | 1.000 | 9703 | tags=18%, list=16%, signal=21% |
| 4197 | DETECTION\_OF\_LIGHT\_STIMULUS\_INVOLVED\_IN\_SENSORY\_PERCEPTION(GO:0050962) |  | 17 | 0.73 | 1.15 | 0.141 | 0.371 | 1.000 | 9703 | tags=18%, list=16%, signal=21% |
| 4198 | REGULATION\_OF\_METAPHASE\_ANAPHASE\_TRANSITION\_OF\_CELL\_CYCLE(GO:1902099) |  | 49 | 0.65 | 1.15 | 0.000 | 0.371 | 1.000 | 15525 | tags=59%, list=26%, signal=80% |
| 4199 | REGULATION\_OF\_MITOTIC\_METAPHASE\_ANAPHASE\_TRANSITION(GO:0030071) |  | 48 | 0.65 | 1.15 | 0.000 | 0.371 | 1.000 | 15525 | tags=58%, list=26%, signal=79% |
| 4200 | ENZYME\_ACTIVATOR\_ACTIVITY(GO:0008047) |  | 514 | 0.68 | 1.15 | 0.141 | 0.371 | 1.000 | 11609 | tags=22%, list=20%, signal=27% |
| 4201 | RESPONSE\_TO\_TRANSFORMING\_GROWTH\_FACTOR\_BETA(GO:0071559) |  | 232 | 0.69 | 1.15 | 0.141 | 0.371 | 1.000 | 12996 | tags=31%, list=22%, signal=40% |
| 4202 | PROTEIN\_DEPOLYMERIZATION(GO:0051261) |  | 98 | 0.70 | 1.15 | 0.141 | 0.371 | 1.000 | 11742 | tags=32%, list=20%, signal=39% |
| 4203 | POSITIVE\_REGULATION\_OF\_ALPHA\_BETA\_T\_CELL\_ACTIVATION(GO:0046635) |  | 50 | 0.69 | 1.15 | 0.141 | 0.371 | 1.000 | 14175 | tags=46%, list=24%, signal=60% |
| 4204 | MICROTUBULE\_ASSOCIATED\_COMPLEX(GO:0005875) |  | 150 | 0.66 | 1.15 | 0.000 | 0.371 | 1.000 | 13902 | tags=42%, list=24%, signal=55% |
| 4205 | CELLULAR\_RESPONSE\_TO\_TRANSFORMING\_GROWTH\_FACTOR\_BETA\_STIMULUS(GO:0071560) |  | 228 | 0.69 | 1.15 | 0.141 | 0.371 | 1.000 | 12996 | tags=32%, list=22%, signal=40% |
| 4206 | RAS\_GTPASE\_BINDING(GO:0017016) |  | 526 | 0.69 | 1.15 | 0.141 | 0.371 | 1.000 | 12119 | tags=27%, list=20%, signal=34% |
| 4207 | ESTABLISHMENT\_OF\_PLANAR\_POLARITY(GO:0001736) |  | 123 | 0.68 | 1.15 | 0.141 | 0.371 | 1.000 | 12594 | tags=27%, list=21%, signal=34% |
| 4208 | ESTABLISHMENT\_OF\_TISSUE\_POLARITY(GO:0007164) |  | 123 | 0.68 | 1.15 | 0.141 | 0.371 | 1.000 | 12594 | tags=27%, list=21%, signal=34% |
| 4209 | IDENTICAL\_PROTEIN\_BINDING(GO:0042802) |  | 1641 | 0.71 | 1.15 | 0.141 | 0.371 | 1.000 | 11131 | tags=24%, list=19%, signal=28% |
| 4210 | MACROAUTOPHAGY(GO:0016236) |  | 285 | 0.65 | 1.15 | 0.141 | 0.371 | 1.000 | 13150 | tags=23%, list=22%, signal=30% |
| 4211 | CENTROSOME(GO:0005813) |  | 488 | 0.69 | 1.15 | 0.141 | 0.371 | 1.000 | 12577 | tags=37%, list=21%, signal=47% |
| 4212 | LIGAND\_GATED\_CATION\_CHANNEL\_ACTIVITY(GO:0099094) |  | 103 | 0.72 | 1.15 | 0.141 | 0.371 | 1.000 | 9703 | tags=32%, list=16%, signal=38% |
| 4213 | GERM\_CELL\_DEVELOPMENT(GO:0007281) |  | 245 | 0.69 | 1.15 | 0.141 | 0.371 | 1.000 | 11229 | tags=28%, list=19%, signal=35% |
| 4214 | CEREBELLAR\_PURKINJE\_CELL\_LAYER\_MORPHOGENESIS(GO:0021692) |  | 15 | 0.67 | 1.15 | 0.000 | 0.371 | 1.000 | 13932 | tags=47%, list=24%, signal=61% |
| 4215 | CYSTEINE\_TYPE\_ENDOPEPTIDASE\_ACTIVITY\_INVOLVED\_IN\_APOPTOTIC\_PROCESS(GO:0097153) |  | 15 | 0.74 | 1.15 | 0.000 | 0.371 | 1.000 | 6187 | tags=7%, list=10%, signal=7% |
| 4216 | RESPONSE\_TO\_INCREASED\_OXYGEN\_LEVELS(GO:0036296) |  | 28 | 0.68 | 1.15 | 0.141 | 0.371 | 1.000 | 9746 | tags=21%, list=16%, signal=26% |
| 4217 | GLYCOSYLATION(GO:0070085) |  | 269 | 0.68 | 1.15 | 0.141 | 0.371 | 1.000 | 11789 | tags=25%, list=20%, signal=31% |
| 4218 | POSITIVE\_REGULATION\_OF\_FATTY\_ACID\_TRANSPORT(GO:2000193) |  | 16 | 0.74 | 1.15 | 0.141 | 0.371 | 1.000 | 11372 | tags=31%, list=19%, signal=39% |
| 4219 | FK506\_BINDING(GO:0005528) |  | 16 | 0.67 | 1.15 | 0.000 | 0.371 | 1.000 | 14311 | tags=50%, list=24%, signal=66% |
| 4220 | MACROLIDE\_BINDING(GO:0005527) |  | 16 | 0.67 | 1.15 | 0.000 | 0.371 | 1.000 | 14311 | tags=50%, list=24%, signal=66% |
| 4221 | POSITIVE\_REGULATION\_OF\_PROTEOLYSIS(GO:0045862) |  | 340 | 0.66 | 1.15 | 0.141 | 0.371 | 1.000 | 15082 | tags=33%, list=26%, signal=44% |
| 4222 | REGULATION\_OF\_MITOCHONDRIAL\_MEMBRANE\_POTENTIAL(GO:0051881) |  | 65 | 0.68 | 1.15 | 0.141 | 0.371 | 1.000 | 7907 | tags=15%, list=13%, signal=18% |
| 4223 | POSITIVE\_REGULATION\_OF\_CELL\_CYCLE\_ARREST(GO:0071158) |  | 88 | 0.71 | 1.15 | 0.141 | 0.371 | 1.000 | 7640 | tags=9%, list=13%, signal=10% |
| 4224 | GLYCOPROTEIN\_BIOSYNTHETIC\_PROCESS(GO:0009101) |  | 331 | 0.69 | 1.15 | 0.141 | 0.371 | 1.000 | 12346 | tags=29%, list=21%, signal=37% |
| 4225 | GAMMA\_TUBULIN\_COMPLEX(GO:0000930) |  | 22 | 0.65 | 1.15 | 0.000 | 0.371 | 1.000 | 15351 | tags=50%, list=26%, signal=68% |
| 4226 | POSITIVE\_REGULATION\_OF\_NATURAL\_KILLER\_CELL\_ACTIVATION(GO:0032816) |  | 16 | 0.66 | 1.15 | 0.141 | 0.371 | 1.000 | 16662 | tags=56%, list=28%, signal=78% |
| 4227 | MEMBRANE\_REPOLARIZATION\_DURING\_CARDIAC\_MUSCLE\_CELL\_ACTION\_POTENTIAL(GO:0086013) |  | 26 | 0.70 | 1.15 | 0.141 | 0.371 | 1.000 | 10556 | tags=27%, list=18%, signal=33% |
| 4228 | REGULATION\_OF\_CHROMOSOME\_SEPARATION(GO:1905818) |  | 56 | 0.65 | 1.15 | 0.000 | 0.371 | 1.000 | 15525 | tags=59%, list=26%, signal=80% |
| 4229 | REGULATION\_OF\_CELL\_CYCLE\_PROCESS(GO:0010564) |  | 740 | 0.69 | 1.15 | 0.141 | 0.371 | 1.000 | 12586 | tags=29%, list=21%, signal=36% |
| 4230 | REGULATION\_OF\_CELL\_DIVISION(GO:0051302) |  | 156 | 0.71 | 1.15 | 0.141 | 0.371 | 1.000 | 12392 | tags=29%, list=21%, signal=36% |
| 4231 | RESPONSE\_TO\_MANGANESE\_ION(GO:0010042) |  | 17 | 0.73 | 1.15 | 0.141 | 0.371 | 1.000 | 7435 | tags=35%, list=13%, signal=40% |
| 4232 | RETINA\_MORPHOGENESIS\_IN\_CAMERA\_TYPE\_EYE(GO:0060042) |  | 49 | 0.74 | 1.15 | 0.141 | 0.371 | 1.000 | 10712 | tags=39%, list=18%, signal=47% |
| 4233 | CARBOHYDRATE\_CATABOLIC\_PROCESS(GO:0016052) |  | 181 | 0.69 | 1.15 | 0.141 | 0.371 | 1.000 | 11506 | tags=20%, list=19%, signal=25% |
| 4234 | DNA\_CONFORMATION\_CHANGE(GO:0071103) |  | 288 | 0.70 | 1.15 | 0.141 | 0.371 | 1.000 | 10856 | tags=19%, list=18%, signal=23% |
| 4235 | TRANS\_GOLGI\_NETWORK\_MEMBRANE(GO:0032588) |  | 83 | 0.68 | 1.15 | 0.141 | 0.371 | 1.000 | 7297 | tags=7%, list=12%, signal=8% |
| 4236 | REGULATION\_OF\_COLLAGEN\_METABOLIC\_PROCESS(GO:0010712) |  | 43 | 0.67 | 1.15 | 0.141 | 0.371 | 1.000 | 14441 | tags=30%, list=24%, signal=40% |
| 4237 | REGULATION\_OF\_MITOTIC\_SISTER\_CHROMATID\_SEPARATION(GO:0010965) |  | 52 | 0.65 | 1.15 | 0.000 | 0.371 | 1.000 | 15525 | tags=58%, list=26%, signal=78% |
| 4238 | MONOCARBOXYLIC\_ACID\_BINDING(GO:0033293) |  | 59 | 0.71 | 1.15 | 0.141 | 0.371 | 1.000 | 11950 | tags=27%, list=20%, signal=34% |
| 4239 | HAIR\_CELL\_DIFFERENTIATION(GO:0035315) |  | 42 | 0.68 | 1.15 | 0.141 | 0.371 | 1.000 | 9354 | tags=24%, list=16%, signal=28% |
| 4240 | POSITIVE\_REGULATION\_OF\_LYMPHOCYTE\_ACTIVATION(GO:0051251) |  | 313 | 0.67 | 1.15 | 0.141 | 0.371 | 1.000 | 12242 | tags=22%, list=21%, signal=28% |
| 4241 | HEAT\_GENERATION(GO:0031649) |  | 20 | 0.75 | 1.15 | 0.141 | 0.371 | 1.000 | 11372 | tags=40%, list=19%, signal=50% |
| 4242 | CELLULAR\_RESPONSE\_TO\_ESTRADIOL\_STIMULUS(GO:0071392) |  | 36 | 0.72 | 1.14 | 0.141 | 0.371 | 1.000 | 12241 | tags=39%, list=21%, signal=49% |
| 4243 | ADAPTIVE\_IMMUNE\_RESPONSE\_BASED\_ON\_SOMATIC\_RECOMBINATION\_OF\_IMMUNE\_RECEPTORS\_BUILT\_FROM\_IMMUNOGLOBULIN\_SUPERFAMILY\_DOMAINS(GO:0002460) |  | 317 | 0.67 | 1.14 | 0.141 | 0.371 | 1.000 | 10239 | tags=17%, list=17%, signal=20% |
| 4244 | METAPHASE\_ANAPHASE\_TRANSITION\_OF\_CELL\_CYCLE(GO:0044784) |  | 51 | 0.65 | 1.14 | 0.000 | 0.371 | 1.000 | 15525 | tags=59%, list=26%, signal=80% |
| 4245 | METAPHASE\_ANAPHASE\_TRANSITION\_OF\_MITOTIC\_CELL\_CYCLE(GO:0007091) |  | 50 | 0.65 | 1.14 | 0.000 | 0.371 | 1.000 | 15525 | tags=58%, list=26%, signal=79% |
| 4246 | POSITIVE\_REGULATION\_OF\_T\_CELL\_MEDIATED\_IMMUNITY(GO:0002711) |  | 31 | 0.68 | 1.14 | 0.141 | 0.371 | 1.000 | 15082 | tags=42%, list=26%, signal=56% |
| 4247 | SMALL\_GTPASE\_BINDING(GO:0031267) |  | 541 | 0.69 | 1.14 | 0.141 | 0.371 | 1.000 | 12119 | tags=27%, list=20%, signal=33% |
| 4248 | RECEPTOR\_LIGAND\_ACTIVITY(GO:0048018) |  | 459 | 0.70 | 1.14 | 0.141 | 0.371 | 1.000 | 10951 | tags=28%, list=19%, signal=34% |
| 4249 | FRIZZLED\_BINDING(GO:0005109) |  | 35 | 0.74 | 1.14 | 0.141 | 0.370 | 1.000 | 11918 | tags=37%, list=20%, signal=46% |
| 4250 | PLASMA\_LIPOPROTEIN\_PARTICLE\_ASSEMBLY(GO:0034377) |  | 27 | 0.67 | 1.14 | 0.000 | 0.370 | 1.000 | 11522 | tags=22%, list=19%, signal=28% |
| 4251 | POSITIVE\_REGULATION\_OF\_PROGRAMMED\_CELL\_DEATH(GO:0043068) |  | 626 | 0.70 | 1.14 | 0.141 | 0.370 | 1.000 | 10967 | tags=19%, list=19%, signal=24% |
| 4252 | DENDRITIC\_SPINE\_ORGANIZATION(GO:0097061) |  | 68 | 0.66 | 1.14 | 0.141 | 0.370 | 1.000 | 15406 | tags=43%, list=26%, signal=58% |
| 4253 | MUSCLE\_MYOSIN\_COMPLEX(GO:0005859) |  | 27 | 0.72 | 1.14 | 0.166 | 0.370 | 1.000 | 7652 | tags=41%, list=13%, signal=47% |
| 4254 | THIOL\_DEPENDENT\_UBIQUITINYL\_HYDROLASE\_ACTIVITY(GO:0036459) |  | 128 | 0.66 | 1.14 | 0.000 | 0.370 | 1.000 | 12385 | tags=24%, list=21%, signal=31% |
| 4255 | PROTEIN\_DNA\_COMPLEX\_ASSEMBLY(GO:0065004) |  | 247 | 0.70 | 1.14 | 0.141 | 0.370 | 1.000 | 12160 | tags=26%, list=21%, signal=32% |
| 4256 | NADP\_BINDING(GO:0050661) |  | 47 | 0.67 | 1.14 | 0.141 | 0.370 | 1.000 | 11214 | tags=32%, list=19%, signal=39% |
| 4257 | POSITIVE\_REGULATION\_OF\_APOPTOTIC\_PROCESS(GO:0043065) |  | 620 | 0.70 | 1.14 | 0.141 | 0.370 | 1.000 | 10967 | tags=19%, list=19%, signal=23% |
| 4258 | PROTEIN\_EXIT\_FROM\_ENDOPLASMIC\_RETICULUM(GO:0032527) |  | 40 | 0.73 | 1.14 | 0.141 | 0.370 | 1.000 | 6573 | tags=10%, list=11%, signal=11% |
| 4259 | PHOSPHATIDYLSERINE\_BINDING(GO:0001786) |  | 38 | 0.69 | 1.14 | 0.141 | 0.370 | 1.000 | 7056 | tags=18%, list=12%, signal=21% |
| 4260 | MONOCARBOXYLIC\_ACID\_TRANSMEMBRANE\_TRANSPORTER\_ACTIVITY(GO:0008028) |  | 57 | 0.70 | 1.14 | 0.141 | 0.370 | 1.000 | 13242 | tags=40%, list=22%, signal=52% |
| 4261 | ACETYLGLUCOSAMINYLTRANSFERASE\_ACTIVITY(GO:0008375) |  | 51 | 0.66 | 1.14 | 0.141 | 0.370 | 1.000 | 14353 | tags=41%, list=24%, signal=54% |
| 4262 | PROTEIN\_LOCALIZATION\_TO\_MICROTUBULE\_CYTOSKELETON(GO:0072698) |  | 43 | 0.66 | 1.14 | 0.000 | 0.370 | 1.000 | 15989 | tags=60%, list=27%, signal=83% |
| 4263 | MATING\_BEHAVIOR(GO:0007617) |  | 21 | 0.71 | 1.14 | 0.141 | 0.370 | 1.000 | 8185 | tags=29%, list=14%, signal=33% |
| 4264 | T\_HELPER\_CELL\_LINEAGE\_COMMITMENT(GO:0002295) |  | 15 | 0.73 | 1.14 | 0.141 | 0.370 | 1.000 | 9023 | tags=27%, list=15%, signal=31% |
| 4265 | NEGATIVE\_REGULATION\_OF\_LEUKOCYTE\_APOPTOTIC\_PROCESS(GO:2000107) |  | 46 | 0.71 | 1.14 | 0.141 | 0.370 | 1.000 | 9411 | tags=15%, list=16%, signal=18% |
| 4266 | INTRACELLULAR\_STEROID\_HORMONE\_RECEPTOR\_SIGNALING\_PATHWAY(GO:0030518) |  | 143 | 0.70 | 1.14 | 0.141 | 0.370 | 1.000 | 8112 | tags=12%, list=14%, signal=14% |
| 4267 | LIPID\_MODIFICATION(GO:0030258) |  | 279 | 0.68 | 1.14 | 0.141 | 0.370 | 1.000 | 10669 | tags=24%, list=18%, signal=30% |
| 4268 | REGULATION\_OF\_NATURAL\_KILLER\_CELL\_ACTIVATION(GO:0032814) |  | 26 | 0.66 | 1.14 | 0.141 | 0.370 | 1.000 | 13757 | tags=38%, list=23%, signal=50% |
| 4269 | RESPONSE\_TO\_LITHIUM\_ION(GO:0010226) |  | 23 | 0.71 | 1.14 | 0.141 | 0.370 | 1.000 | 8349 | tags=17%, list=14%, signal=20% |
| 4270 | CELL\_CYCLE\_PROCESS(GO:0022402) |  | 1336 | 0.69 | 1.14 | 0.141 | 0.370 | 1.000 | 12586 | tags=30%, list=21%, signal=38% |
| 4271 | POSITIVE\_REGULATION\_OF\_OXIDATIVE\_STRESS\_INDUCED\_CELL\_DEATH(GO:1903209) |  | 17 | 0.69 | 1.14 | 0.000 | 0.370 | 1.000 | 6514 | tags=12%, list=11%, signal=13% |
| 4272 | CENTROSOME\_SEPARATION(GO:0051299) |  | 17 | 0.68 | 1.14 | 0.000 | 0.370 | 1.000 | 17160 | tags=94%, list=29%, signal=133% |
| 4273 | NEGATIVE\_REGULATION\_OF\_NEURON\_PROJECTION\_DEVELOPMENT(GO:0010977) |  | 131 | 0.70 | 1.14 | 0.141 | 0.370 | 1.000 | 9454 | tags=28%, list=16%, signal=34% |
| 4274 | REGULATION\_OF\_MYELOID\_CELL\_APOPTOTIC\_PROCESS(GO:0033032) |  | 28 | 0.68 | 1.14 | 0.000 | 0.370 | 1.000 | 10262 | tags=21%, list=17%, signal=26% |
| 4275 | LYASE\_ACTIVITY(GO:0016829) |  | 187 | 0.69 | 1.14 | 0.141 | 0.370 | 1.000 | 10855 | tags=25%, list=18%, signal=31% |
| 4276 | REGULATION\_OF\_PROTEIN\_ACTIVATION\_CASCADE(GO:2000257) |  | 106 | 0.67 | 1.14 | 0.141 | 0.370 | 1.000 | 15756 | tags=19%, list=27%, signal=26% |
| 4277 | VOLTAGE\_GATED\_POTASSIUM\_CHANNEL\_COMPLEX(GO:0008076) |  | 90 | 0.71 | 1.14 | 0.141 | 0.370 | 1.000 | 13305 | tags=42%, list=22%, signal=54% |
| 4278 | NEGATIVE\_REGULATION\_OF\_HORMONE\_SECRETION(GO:0046888) |  | 61 | 0.69 | 1.14 | 0.141 | 0.370 | 1.000 | 7544 | tags=15%, list=13%, signal=17% |
| 4279 | REGULATION\_OF\_POLYSACCHARIDE\_METABOLIC\_PROCESS(GO:0032881) |  | 39 | 0.71 | 1.14 | 0.141 | 0.370 | 1.000 | 8594 | tags=15%, list=15%, signal=18% |
| 4280 | TRANSLATION\_REPRESSOR\_ACTIVITY(GO:0030371) |  | 25 | 0.67 | 1.14 | 0.000 | 0.370 | 1.000 | 9105 | tags=24%, list=15%, signal=28% |
| 4281 | POSITIVE\_REGULATION\_OF\_INTRINSIC\_APOPTOTIC\_SIGNALING\_PATHWAY(GO:2001244) |  | 57 | 0.69 | 1.14 | 0.141 | 0.370 | 1.000 | 13155 | tags=25%, list=22%, signal=32% |
| 4282 | CHROMATIN\_ASSEMBLY(GO:0031497) |  | 162 | 0.69 | 1.14 | 0.141 | 0.370 | 1.000 | 10856 | tags=17%, list=18%, signal=20% |
| 4283 | TRANSFERASE\_ACTIVITY\_\_TRANSFERRING\_HEXOSYL\_GROUPS(GO:0016758) |  | 209 | 0.68 | 1.14 | 0.141 | 0.370 | 1.000 | 9901 | tags=22%, list=17%, signal=27% |
| 4284 | MITOTIC\_SISTER\_CHROMATID\_SEPARATION(GO:0051306) |  | 54 | 0.65 | 1.14 | 0.000 | 0.370 | 1.000 | 17976 | tags=70%, list=30%, signal=101% |
| 4285 | POSITIVE\_REGULATION\_OF\_ALPHA\_BETA\_T\_CELL\_DIFFERENTIATION(GO:0046638) |  | 39 | 0.69 | 1.14 | 0.141 | 0.370 | 1.000 | 14175 | tags=49%, list=24%, signal=64% |
| 4286 | O\_GLYCAN\_PROCESSING(GO:0016266) |  | 56 | 0.70 | 1.14 | 0.141 | 0.370 | 1.000 | 13170 | tags=36%, list=22%, signal=46% |
| 4287 | ISOPRENOID\_BIOSYNTHETIC\_PROCESS(GO:0008299) |  | 27 | 0.69 | 1.14 | 0.141 | 0.370 | 1.000 | 14587 | tags=44%, list=25%, signal=59% |
| 4288 | ACTIVATION\_OF\_ADENYLATE\_CYCLASE\_ACTIVITY(GO:0007190) |  | 36 | 0.71 | 1.14 | 0.141 | 0.370 | 1.000 | 9042 | tags=28%, list=15%, signal=33% |
| 4289 | POSITIVE\_REGULATION\_OF\_EPIDERMAL\_CELL\_DIFFERENTIATION(GO:0045606) |  | 23 | 0.74 | 1.14 | 0.141 | 0.370 | 1.000 | 10768 | tags=39%, list=18%, signal=48% |
| 4290 | CELL\_SUBSTRATE\_JUNCTION\_ASSEMBLY(GO:0007044) |  | 90 | 0.71 | 1.14 | 0.141 | 0.370 | 1.000 | 8865 | tags=11%, list=15%, signal=13% |
| 4291 | CELLULAR\_RESPONSE\_TO\_EXOGENOUS\_DSRNA(GO:0071360) |  | 16 | 0.72 | 1.14 | 0.141 | 0.370 | 1.000 | 6255 | tags=6%, list=11%, signal=7% |
| 4292 | PURINE\_RIBONUCLEOSIDE\_TRIPHOSPHATE\_METABOLIC\_PROCESS(GO:0009205) |  | 317 | 0.67 | 1.14 | 0.141 | 0.370 | 1.000 | 11589 | tags=17%, list=20%, signal=21% |
| 4293 | CYTOKINE\_ACTIVITY(GO:0005125) |  | 221 | 0.69 | 1.14 | 0.141 | 0.370 | 1.000 | 10951 | tags=28%, list=19%, signal=34% |
| 4294 | REGULATION\_OF\_UBIQUITIN\_PROTEIN\_TRANSFERASE\_ACTIVITY(GO:0051438) |  | 54 | 0.65 | 1.14 | 0.000 | 0.370 | 1.000 | 14185 | tags=39%, list=24%, signal=51% |
| 4295 | VACUOLAR\_MEMBRANE(GO:0005774) |  | 397 | 0.68 | 1.14 | 0.141 | 0.370 | 1.000 | 9641 | tags=12%, list=16%, signal=14% |
| 4296 | POSITIVE\_REGULATION\_OF\_TRANSMEMBRANE\_RECEPTOR\_PROTEIN\_SERINE\_THREONINE\_KINASE\_SIGNALING\_PATHWAY(GO:0090100) |  | 102 | 0.70 | 1.14 | 0.141 | 0.370 | 1.000 | 13476 | tags=39%, list=23%, signal=51% |
| 4297 | POTASSIUM\_CHANNEL\_COMPLEX(GO:0034705) |  | 97 | 0.71 | 1.14 | 0.141 | 0.370 | 1.000 | 13305 | tags=43%, list=22%, signal=56% |
| 4298 | RESPONSE\_TO\_CALCIUM\_ION(GO:0051592) |  | 125 | 0.70 | 1.14 | 0.141 | 0.370 | 1.000 | 8198 | tags=14%, list=14%, signal=17% |
| 4299 | PROCESS\_UTILIZING\_AUTOPHAGIC\_MECHANISM(GO:0061919) |  | 478 | 0.65 | 1.14 | 0.141 | 0.370 | 1.000 | 13150 | tags=22%, list=22%, signal=28% |
| 4300 | AUTOPHAGY(GO:0006914) |  | 478 | 0.65 | 1.14 | 0.141 | 0.370 | 1.000 | 13150 | tags=22%, list=22%, signal=28% |
| 4301 | PROTEIN\_PHOSPHATASE\_BINDING(GO:0019903) |  | 120 | 0.69 | 1.14 | 0.141 | 0.370 | 1.000 | 10352 | tags=18%, list=18%, signal=21% |
| 4302 | ION\_CHANNEL\_INHIBITOR\_ACTIVITY(GO:0008200) |  | 36 | 0.67 | 1.14 | 0.141 | 0.370 | 1.000 | 9409 | tags=22%, list=16%, signal=26% |
| 4303 | INNER\_EAR\_RECEPTOR\_CELL\_DEVELOPMENT(GO:0060119) |  | 45 | 0.68 | 1.14 | 0.141 | 0.369 | 1.000 | 9354 | tags=24%, list=16%, signal=29% |
| 4304 | KINETOCHORE(GO:0000776) |  | 135 | 0.65 | 1.14 | 0.000 | 0.369 | 1.000 | 15387 | tags=60%, list=26%, signal=81% |
| 4305 | NEGATIVE\_REGULATION\_OF\_LIPID\_BIOSYNTHETIC\_PROCESS(GO:0051055) |  | 49 | 0.69 | 1.14 | 0.141 | 0.369 | 1.000 | 13039 | tags=37%, list=22%, signal=47% |
| 4306 | FOLIC\_ACID\_METABOLIC\_PROCESS(GO:0046655) |  | 18 | 0.66 | 1.14 | 0.000 | 0.370 | 1.000 | 14644 | tags=56%, list=25%, signal=74% |
| 4307 | RESPONSE\_TO\_METAL\_ION(GO:0010038) |  | 334 | 0.69 | 1.14 | 0.141 | 0.370 | 1.000 | 8349 | tags=16%, list=14%, signal=18% |
| 4308 | RESPONSE\_TO\_VITAMIN\_A(GO:0033189) |  | 19 | 0.70 | 1.14 | 0.000 | 0.370 | 1.000 | 12258 | tags=42%, list=21%, signal=53% |
| 4309 | RIBONUCLEOSIDE\_TRIPHOSPHATE\_METABOLIC\_PROCESS(GO:0009199) |  | 323 | 0.66 | 1.14 | 0.141 | 0.370 | 1.000 | 11589 | tags=17%, list=20%, signal=21% |
| 4310 | INHIBITORY\_SYNAPSE(GO:0060077) |  | 16 | 0.72 | 1.14 | 0.141 | 0.370 | 1.000 | 13572 | tags=44%, list=23%, signal=57% |
| 4311 | CARBOHYDRATE\_DERIVATIVE\_BINDING(GO:0097367) |  | 2224 | 0.70 | 1.14 | 0.141 | 0.370 | 1.000 | 12246 | tags=29%, list=21%, signal=35% |
| 4312 | MITOTIC\_CENTROSOME\_SEPARATION(GO:0007100) |  | 16 | 0.68 | 1.14 | 0.000 | 0.369 | 1.000 | 17160 | tags=94%, list=29%, signal=132% |
| 4313 | REGULATION\_OF\_BONE\_DEVELOPMENT(GO:1903010) |  | 22 | 0.71 | 1.14 | 0.141 | 0.369 | 1.000 | 12925 | tags=41%, list=22%, signal=52% |
| 4314 | NEGATIVE\_REGULATION\_OF\_TRANSCRIPTION\_REGULATORY\_REGION\_DNA\_BINDING(GO:2000678) |  | 21 | 0.73 | 1.14 | 0.141 | 0.369 | 1.000 | 10116 | tags=33%, list=17%, signal=40% |
| 4315 | REGULATION\_OF\_HUMORAL\_IMMUNE\_RESPONSE(GO:0002920) |  | 123 | 0.69 | 1.14 | 0.141 | 0.369 | 1.000 | 7853 | tags=12%, list=13%, signal=14% |
| 4316 | REGULATION\_OF\_LEUKOCYTE\_APOPTOTIC\_PROCESS(GO:2000106) |  | 82 | 0.70 | 1.14 | 0.141 | 0.369 | 1.000 | 13741 | tags=28%, list=23%, signal=36% |
| 4317 | POSITIVE\_REGULATION\_OF\_DNA\_METABOLIC\_PROCESS(GO:0051054) |  | 220 | 0.70 | 1.14 | 0.141 | 0.369 | 1.000 | 11130 | tags=25%, list=19%, signal=31% |
| 4318 | NEGATIVE\_REGULATION\_OF\_INTERLEUKIN\_12\_PRODUCTION(GO:0032695) |  | 15 | 0.71 | 1.14 | 0.141 | 0.369 | 1.000 | 13873 | tags=53%, list=23%, signal=70% |
| 4319 | REGULATION\_OF\_INTERLEUKIN\_8\_PRODUCTION(GO:0032677) |  | 64 | 0.68 | 1.14 | 0.141 | 0.369 | 1.000 | 6626 | tags=13%, list=11%, signal=14% |
| 4320 | PEPTIDE\_HORMONE\_RECEPTOR\_BINDING(GO:0051428) |  | 19 | 0.70 | 1.14 | 0.000 | 0.369 | 1.000 | 9985 | tags=26%, list=17%, signal=32% |
| 4321 | REGULATION\_OF\_NEUROLOGICAL\_SYSTEM\_PROCESS(GO:0031644) |  | 71 | 0.67 | 1.14 | 0.141 | 0.369 | 1.000 | 9682 | tags=30%, list=16%, signal=35% |
| 4322 | HORMONE\_RECEPTOR\_BINDING(GO:0051427) |  | 173 | 0.67 | 1.14 | 0.141 | 0.369 | 1.000 | 12688 | tags=25%, list=21%, signal=32% |
| 4323 | POSITIVE\_REGULATION\_OF\_CELL\_DEATH(GO:0010942) |  | 674 | 0.69 | 1.14 | 0.141 | 0.369 | 1.000 | 10967 | tags=19%, list=19%, signal=23% |
| 4324 | T\_CELL\_MEDIATED\_IMMUNITY(GO:0002456) |  | 78 | 0.67 | 1.14 | 0.141 | 0.369 | 1.000 | 15529 | tags=37%, list=26%, signal=50% |
| 4325 | HORMONE\_ACTIVITY(GO:0005179) |  | 123 | 0.70 | 1.14 | 0.141 | 0.369 | 1.000 | 10430 | tags=28%, list=18%, signal=34% |
| 4326 | T\_CELL\_CHEMOTAXIS(GO:0010818) |  | 22 | 0.73 | 1.14 | 0.141 | 0.369 | 1.000 | 6473 | tags=23%, list=11%, signal=26% |
| 4327 | NEURON\_PROJECTION\_EXTENSION(GO:1990138) |  | 147 | 0.70 | 1.14 | 0.141 | 0.369 | 1.000 | 11339 | tags=31%, list=19%, signal=38% |
| 4328 | MITOCHONDRIAL\_INNER\_MEMBRANE(GO:0005743) |  | 505 | 0.65 | 1.14 | 0.000 | 0.369 | 1.000 | 17291 | tags=48%, list=29%, signal=67% |
| 4329 | HYDROLASE\_ACTIVITY\_\_HYDROLYZING\_N\_GLYCOSYL\_COMPOUNDS(GO:0016799) |  | 23 | 0.66 | 1.14 | 0.000 | 0.369 | 1.000 | 8925 | tags=17%, list=15%, signal=20% |
| 4330 | REGULATION\_OF\_GLIAL\_CELL\_DIFFERENTIATION(GO:0045685) |  | 61 | 0.71 | 1.14 | 0.141 | 0.369 | 1.000 | 11343 | tags=31%, list=19%, signal=39% |
| 4331 | POSITIVE\_REGULATION\_OF\_HISTONE\_MODIFICATION(GO:0031058) |  | 84 | 0.68 | 1.14 | 0.141 | 0.369 | 1.000 | 10856 | tags=21%, list=18%, signal=26% |
| 4332 | FOUR\_WAY\_JUNCTION\_DNA\_BINDING(GO:0000400) |  | 16 | 0.69 | 1.14 | 0.000 | 0.369 | 1.000 | 17317 | tags=88%, list=29%, signal=124% |
| 4333 | METANEPHRIC\_NEPHRON\_TUBULE\_DEVELOPMENT(GO:0072234) |  | 20 | 0.74 | 1.14 | 0.141 | 0.369 | 1.000 | 13927 | tags=50%, list=24%, signal=65% |
| 4334 | ADULT\_WALKING\_BEHAVIOR(GO:0007628) |  | 30 | 0.72 | 1.14 | 0.141 | 0.369 | 1.000 | 13179 | tags=60%, list=22%, signal=77% |
| 4335 | ORGANELLE\_LOCALIZATION(GO:0051640) |  | 627 | 0.70 | 1.14 | 0.141 | 0.369 | 1.000 | 10763 | tags=22%, list=18%, signal=27% |
| 4336 | PURINE\_NUCLEOSIDE\_TRIPHOSPHATE\_METABOLIC\_PROCESS(GO:0009144) |  | 324 | 0.66 | 1.14 | 0.141 | 0.369 | 1.000 | 11589 | tags=17%, list=20%, signal=21% |
| 4337 | INTEGRIN\_MEDIATED\_SIGNALING\_PATHWAY(GO:0007229) |  | 102 | 0.66 | 1.14 | 0.141 | 0.369 | 1.000 | 15437 | tags=37%, list=26%, signal=50% |
| 4338 | POSITIVE\_REGULATION\_OF\_CHOLESTEROL\_TRANSPORT(GO:0032376) |  | 19 | 0.73 | 1.14 | 0.141 | 0.369 | 1.000 | 11906 | tags=37%, list=20%, signal=46% |
| 4339 | POSITIVE\_REGULATION\_OF\_STEROL\_TRANSPORT(GO:0032373) |  | 19 | 0.73 | 1.14 | 0.141 | 0.369 | 1.000 | 11906 | tags=37%, list=20%, signal=46% |
| 4340 | CELLULAR\_RESPONSE\_TO\_FLUID\_SHEAR\_STRESS(GO:0071498) |  | 21 | 0.68 | 1.14 | 0.000 | 0.369 | 1.000 | 8241 | tags=24%, list=14%, signal=28% |
| 4341 | SOMATIC\_CELL\_DNA\_RECOMBINATION(GO:0016444) |  | 56 | 0.71 | 1.14 | 0.141 | 0.369 | 1.000 | 10412 | tags=34%, list=18%, signal=41% |
| 4342 | SOMATIC\_DIVERSIFICATION\_OF\_IMMUNE\_RECEPTORS\_VIA\_GERMLINE\_RECOMBINATION\_WITHIN\_A\_SINGLE\_LOCUS(GO:0002562) |  | 56 | 0.71 | 1.14 | 0.141 | 0.369 | 1.000 | 10412 | tags=34%, list=18%, signal=41% |
| 4343 | ACETYLCHOLINE\_RECEPTOR\_ACTIVITY(GO:0015464) |  | 18 | 0.68 | 1.14 | 0.141 | 0.369 | 1.000 | 12401 | tags=39%, list=21%, signal=49% |
| 4344 | UBIQUITIN\_LIKE\_PROTEIN\_SPECIFIC\_PROTEASE\_ACTIVITY(GO:0019783) |  | 137 | 0.65 | 1.14 | 0.000 | 0.369 | 1.000 | 18524 | tags=55%, list=31%, signal=81% |
| 4345 | RIBONUCLEOSIDE\_BIOSYNTHETIC\_PROCESS(GO:0042455) |  | 41 | 0.64 | 1.14 | 0.000 | 0.369 | 1.000 | 17206 | tags=56%, list=29%, signal=79% |
| 4346 | PROTEIN\_LOCALIZATION\_TO\_CYTOSKELETON(GO:0044380) |  | 47 | 0.65 | 1.14 | 0.000 | 0.369 | 1.000 | 15989 | tags=57%, list=27%, signal=79% |
| 4347 | ORGANELLE\_INNER\_MEMBRANE(GO:0019866) |  | 562 | 0.65 | 1.14 | 0.000 | 0.369 | 1.000 | 17291 | tags=48%, list=29%, signal=68% |
| 4348 | POSITIVE\_REGULATION\_OF\_STRESS\_ACTIVATED\_MAPK\_CASCADE(GO:0032874) |  | 149 | 0.70 | 1.14 | 0.141 | 0.369 | 1.000 | 8112 | tags=16%, list=14%, signal=19% |
| 4349 | OXIDOREDUCTASE\_ACTIVITY\_\_ACTING\_ON\_PAIRED\_DONORS\_\_WITH\_INCORPORATION\_OR\_REDUCTION\_OF\_MOLECULAR\_OXYGEN\_\_NAD(P)H\_AS\_ONE\_DONOR\_\_AND\_INCORPORATION\_OF\_ONE\_ATOM\_OF\_OXYGEN(GO:0016709) |  | 34 | 0.67 | 1.14 | 0.141 | 0.369 | 1.000 | 11264 | tags=32%, list=19%, signal=40% |
| 4350 | INTERSPECIES\_INTERACTION\_BETWEEN\_ORGANISMS(GO:0044419) |  | 882 | 0.70 | 1.14 | 0.141 | 0.369 | 1.000 | 8349 | tags=8%, list=14%, signal=9% |
| 4351 | ESTABLISHMENT\_OF\_RNA\_LOCALIZATION(GO:0051236) |  | 195 | 0.65 | 1.14 | 0.000 | 0.369 | 1.000 | 17552 | tags=66%, list=30%, signal=94% |
| 4352 | POSITIVE\_REGULATION\_OF\_SYNAPTIC\_TRANSMISSION\_\_GLUTAMATERGIC(GO:0051968) |  | 22 | 0.76 | 1.14 | 0.141 | 0.369 | 1.000 | 7837 | tags=27%, list=13%, signal=31% |
| 4353 | REGULATION\_OF\_COMPLEMENT\_ACTIVATION(GO:0030449) |  | 105 | 0.66 | 1.14 | 0.141 | 0.369 | 1.000 | 15756 | tags=18%, list=27%, signal=25% |
| 4354 | POSITIVE\_REGULATION\_OF\_GLUCOSE\_METABOLIC\_PROCESS(GO:0010907) |  | 35 | 0.70 | 1.14 | 0.141 | 0.369 | 1.000 | 9405 | tags=20%, list=16%, signal=24% |
| 4355 | NEGATIVE\_REGULATION\_OF\_REACTIVE\_OXYGEN\_SPECIES\_METABOLIC\_PROCESS(GO:2000378) |  | 54 | 0.67 | 1.14 | 0.141 | 0.369 | 1.000 | 12925 | tags=24%, list=22%, signal=31% |
| 4356 | NUCLEAR\_DIVISION(GO:0000280) |  | 396 | 0.66 | 1.14 | 0.141 | 0.369 | 1.000 | 17160 | tags=58%, list=29%, signal=81% |
| 4357 | ASPARTATE\_FAMILY\_AMINO\_ACID\_CATABOLIC\_PROCESS(GO:0009068) |  | 19 | 0.68 | 1.14 | 0.000 | 0.369 | 1.000 | 10534 | tags=37%, list=18%, signal=45% |
| 4358 | WATER\_TRANSPORT(GO:0006833) |  | 20 | 0.74 | 1.14 | 0.141 | 0.369 | 1.000 | 10857 | tags=40%, list=18%, signal=49% |
| 4359 | INHIBITORY\_POSTSYNAPTIC\_POTENTIAL(GO:0060080) |  | 15 | 0.72 | 1.14 | 0.141 | 0.369 | 1.000 | 7298 | tags=40%, list=12%, signal=46% |
| 4360 | MITOTIC\_NUCLEAR\_DIVISION(GO:0140014) |  | 254 | 0.65 | 1.14 | 0.141 | 0.369 | 1.000 | 17160 | tags=60%, list=29%, signal=84% |
| 4361 | LENS\_FIBER\_CELL\_DIFFERENTIATION(GO:0070306) |  | 29 | 0.70 | 1.14 | 0.141 | 0.369 | 1.000 | 11402 | tags=34%, list=19%, signal=43% |
| 4362 | PEPTIDASE\_ACTIVATOR\_ACTIVITY(GO:0016504) |  | 38 | 0.67 | 1.14 | 0.141 | 0.369 | 1.000 | 13760 | tags=34%, list=23%, signal=45% |
| 4363 | REGULATION\_OF\_MITOTIC\_SISTER\_CHROMATID\_SEGREGATION(GO:0033047) |  | 62 | 0.65 | 1.14 | 0.000 | 0.369 | 1.000 | 19207 | tags=79%, list=32%, signal=117% |
| 4364 | GOLGI\_VESICLE\_TRANSPORT(GO:0048193) |  | 342 | 0.65 | 1.14 | 0.141 | 0.369 | 1.000 | 13159 | tags=23%, list=22%, signal=30% |
| 4365 | PEROXISOME(GO:0005777) |  | 130 | 0.65 | 1.14 | 0.141 | 0.369 | 1.000 | 14137 | tags=38%, list=24%, signal=49% |
| 4366 | MICROBODY(GO:0042579) |  | 130 | 0.65 | 1.14 | 0.141 | 0.369 | 1.000 | 14137 | tags=38%, list=24%, signal=49% |
| 4367 | ENDOCYTIC\_VESICLE\_LUMEN(GO:0071682) |  | 19 | 0.66 | 1.14 | 0.141 | 0.369 | 1.000 | 13552 | tags=37%, list=23%, signal=48% |
| 4368 | DEFENSE\_RESPONSE\_TO\_VIRUS(GO:0051607) |  | 222 | 0.68 | 1.14 | 0.141 | 0.369 | 1.000 | 11578 | tags=21%, list=20%, signal=26% |
| 4369 | MICROTUBULE\_ORGANIZING\_CENTER(GO:0005815) |  | 690 | 0.67 | 1.14 | 0.141 | 0.369 | 1.000 | 12577 | tags=35%, list=21%, signal=44% |
| 4370 | NEURON\_PROJECTION\_ORGANIZATION(GO:0106027) |  | 71 | 0.65 | 1.14 | 0.141 | 0.369 | 1.000 | 15406 | tags=42%, list=26%, signal=57% |
| 4371 | NUCLEIC\_ACID\_TRANSPORT(GO:0050657) |  | 192 | 0.65 | 1.14 | 0.000 | 0.369 | 1.000 | 17552 | tags=66%, list=30%, signal=93% |
| 4372 | RNA\_TRANSPORT(GO:0050658) |  | 192 | 0.65 | 1.14 | 0.000 | 0.369 | 1.000 | 17552 | tags=66%, list=30%, signal=93% |
| 4373 | BENZENE\_CONTAINING\_COMPOUND\_METABOLIC\_PROCESS(GO:0042537) |  | 22 | 0.74 | 1.14 | 0.141 | 0.369 | 1.000 | 10534 | tags=27%, list=18%, signal=33% |
| 4374 | NUCLEAR\_HORMONE\_RECEPTOR\_BINDING(GO:0035257) |  | 149 | 0.68 | 1.14 | 0.141 | 0.368 | 1.000 | 12688 | tags=26%, list=21%, signal=32% |
| 4375 | REGULATION\_OF\_TUMOR\_NECROSIS\_FACTOR\_SECRETION(GO:1904467) |  | 19 | 0.74 | 1.14 | 0.141 | 0.368 | 1.000 | 7426 | tags=21%, list=13%, signal=24% |
| 4376 | NITROGEN\_COMPOUND\_TRANSPORT(GO:0071705) |  | 2221 | 0.69 | 1.14 | 0.141 | 0.368 | 1.000 | 10183 | tags=16%, list=17%, signal=19% |
| 4377 | REGULATION\_OF\_INTERLEUKIN\_1\_SECRETION(GO:0050704) |  | 44 | 0.70 | 1.14 | 0.141 | 0.368 | 1.000 | 12242 | tags=16%, list=21%, signal=20% |
| 4378 | NEGATIVE\_REGULATION\_OF\_LIPID\_TRANSPORT(GO:0032369) |  | 36 | 0.70 | 1.14 | 0.141 | 0.368 | 1.000 | 7921 | tags=19%, list=13%, signal=22% |
| 4379 | NUCLEOTIDE\_BIOSYNTHETIC\_PROCESS(GO:0009165) |  | 368 | 0.67 | 1.14 | 0.141 | 0.368 | 1.000 | 11391 | tags=20%, list=19%, signal=24% |
| 4380 | POSITIVE\_REGULATION\_OF\_LEUKOCYTE\_ACTIVATION(GO:0002696) |  | 349 | 0.67 | 1.14 | 0.141 | 0.368 | 1.000 | 11130 | tags=21%, list=19%, signal=25% |
| 4381 | SYMBIONT\_PROCESS(GO:0044403) |  | 841 | 0.70 | 1.14 | 0.141 | 0.369 | 1.000 | 8349 | tags=8%, list=14%, signal=10% |
| 4382 | MITOCHONDRIAL\_MEMBRANE(GO:0031966) |  | 713 | 0.65 | 1.14 | 0.141 | 0.368 | 1.000 | 13538 | tags=25%, list=23%, signal=32% |
| 4383 | NEGATIVE\_REGULATION\_OF\_DENDRITE\_MORPHOGENESIS(GO:0050774) |  | 15 | 0.70 | 1.14 | 0.141 | 0.368 | 1.000 | 9137 | tags=40%, list=15%, signal=47% |
| 4384 | BASAL\_PLASMA\_MEMBRANE(GO:0009925) |  | 30 | 0.71 | 1.14 | 0.141 | 0.368 | 1.000 | 14275 | tags=50%, list=24%, signal=66% |
| 4385 | RIBONUCLEOTIDE\_BINDING(GO:0032553) |  | 1925 | 0.69 | 1.14 | 0.141 | 0.368 | 1.000 | 13355 | tags=33%, list=23%, signal=41% |
| 4386 | PURINE\_RIBONUCLEOTIDE\_BINDING(GO:0032555) |  | 1910 | 0.69 | 1.14 | 0.141 | 0.368 | 1.000 | 13355 | tags=33%, list=23%, signal=41% |
| 4387 | NUCLEOSIDE\_PHOSPHATE\_BIOSYNTHETIC\_PROCESS(GO:1901293) |  | 370 | 0.67 | 1.14 | 0.141 | 0.368 | 1.000 | 11391 | tags=20%, list=19%, signal=24% |
| 4388 | OCCLUDING\_JUNCTION(GO:0070160) |  | 125 | 0.69 | 1.14 | 0.141 | 0.368 | 1.000 | 8340 | tags=14%, list=14%, signal=16% |
| 4389 | KINETOCHORE\_ORGANIZATION(GO:0051383) |  | 21 | 0.66 | 1.14 | 0.000 | 0.368 | 1.000 | 15072 | tags=67%, list=25%, signal=89% |
| 4390 | OLIGOSACCHARIDE\_BIOSYNTHETIC\_PROCESS(GO:0009312) |  | 19 | 0.73 | 1.14 | 0.141 | 0.368 | 1.000 | 10433 | tags=26%, list=18%, signal=32% |
| 4391 | PHOSPHATIDYLINOSITOL\_PHOSPHORYLATION(GO:0046854) |  | 100 | 0.69 | 1.14 | 0.141 | 0.368 | 1.000 | 7805 | tags=26%, list=13%, signal=30% |
| 4392 | PURINE\_NUCLEOTIDE\_BINDING(GO:0017076) |  | 1922 | 0.69 | 1.14 | 0.141 | 0.368 | 1.000 | 13355 | tags=33%, list=23%, signal=41% |
| 4393 | REGULATION\_OF\_CHOLESTEROL\_EFFLUX(GO:0010874) |  | 28 | 0.73 | 1.13 | 0.141 | 0.368 | 1.000 | 11225 | tags=21%, list=19%, signal=26% |
| 4394 | AMINO\_ACID\_BINDING(GO:0016597) |  | 59 | 0.70 | 1.13 | 0.141 | 0.368 | 1.000 | 12031 | tags=37%, list=20%, signal=47% |
| 4395 | MRNA\_TRANSPORT(GO:0051028) |  | 146 | 0.65 | 1.13 | 0.000 | 0.368 | 1.000 | 17526 | tags=66%, list=30%, signal=94% |
| 4396 | POSITIVE\_REGULATION\_OF\_STRIATED\_MUSCLE\_CELL\_DIFFERENTIATION(GO:0051155) |  | 69 | 0.68 | 1.13 | 0.141 | 0.368 | 1.000 | 9543 | tags=23%, list=16%, signal=28% |
| 4397 | SMALL\_MOLECULE\_BINDING(GO:0036094) |  | 2503 | 0.69 | 1.13 | 0.141 | 0.368 | 1.000 | 12337 | tags=28%, list=21%, signal=34% |
| 4398 | SLEEP(GO:0030431) |  | 34 | 0.69 | 1.13 | 0.141 | 0.368 | 1.000 | 7172 | tags=24%, list=12%, signal=27% |
| 4399 | REGULATION\_OF\_CELL\_CYCLE\_CHECKPOINT(GO:1901976) |  | 33 | 0.67 | 1.13 | 0.000 | 0.368 | 1.000 | 15985 | tags=61%, list=27%, signal=83% |
| 4400 | POSITIVE\_REGULATION\_OF\_EPIDERMIS\_DEVELOPMENT(GO:0045684) |  | 36 | 0.73 | 1.13 | 0.141 | 0.368 | 1.000 | 11868 | tags=36%, list=20%, signal=45% |
| 4401 | PURINE\_RIBONUCLEOSIDE\_TRIPHOSPHATE\_BINDING(GO:0035639) |  | 1867 | 0.69 | 1.13 | 0.141 | 0.368 | 1.000 | 13355 | tags=33%, list=23%, signal=41% |
| 4402 | REGULATION\_OF\_CELL\_CYCLE\_ARREST(GO:0071156) |  | 113 | 0.70 | 1.13 | 0.141 | 0.368 | 1.000 | 7640 | tags=10%, list=13%, signal=11% |
| 4403 | NUCLEOSIDE\_TRIPHOSPHATE\_METABOLIC\_PROCESS(GO:0009141) |  | 343 | 0.66 | 1.13 | 0.141 | 0.368 | 1.000 | 11589 | tags=17%, list=20%, signal=21% |
| 4404 | RECEPTOR\_REGULATOR\_ACTIVITY(GO:0030545) |  | 483 | 0.69 | 1.13 | 0.141 | 0.368 | 1.000 | 10951 | tags=28%, list=19%, signal=34% |
| 4405 | SYNAPTIC\_TRANSMISSION\_\_GABAERGIC(GO:0051932) |  | 38 | 0.65 | 1.13 | 0.000 | 0.368 | 1.000 | 14929 | tags=45%, list=25%, signal=60% |
| 4406 | NEGATIVE\_REGULATION\_OF\_CATION\_CHANNEL\_ACTIVITY(GO:2001258) |  | 51 | 0.65 | 1.13 | 0.141 | 0.368 | 1.000 | 15565 | tags=43%, list=26%, signal=58% |
| 4407 | PROTEIN\_DEPHOSPHORYLATION(GO:0006470) |  | 314 | 0.66 | 1.13 | 0.141 | 0.368 | 1.000 | 10888 | tags=18%, list=18%, signal=22% |
| 4408 | REGULATION\_OF\_CAMP\_BIOSYNTHETIC\_PROCESS(GO:0030817) |  | 52 | 0.65 | 1.13 | 0.141 | 0.368 | 1.000 | 7472 | tags=15%, list=13%, signal=18% |
| 4409 | POSITIVE\_REGULATION\_OF\_PROTEASOMAL\_PROTEIN\_CATABOLIC\_PROCESS(GO:1901800) |  | 102 | 0.66 | 1.13 | 0.141 | 0.368 | 1.000 | 14983 | tags=37%, list=25%, signal=50% |
| 4410 | REGULATION\_OF\_PROTEIN\_COMPLEX\_DISASSEMBLY(GO:0043244) |  | 99 | 0.69 | 1.13 | 0.141 | 0.368 | 1.000 | 11742 | tags=26%, list=20%, signal=33% |
| 4411 | G\_PROTEIN\_COUPLED\_NEUROTRANSMITTER\_RECEPTOR\_ACTIVITY(GO:0099528) |  | 36 | 0.70 | 1.13 | 0.141 | 0.368 | 1.000 | 12916 | tags=31%, list=22%, signal=39% |
| 4412 | LATE\_ENDOSOME\_MEMBRANE(GO:0031902) |  | 126 | 0.66 | 1.13 | 0.141 | 0.368 | 1.000 | 10450 | tags=17%, list=18%, signal=21% |
| 4413 | G1\_S\_TRANSITION\_OF\_MITOTIC\_CELL\_CYCLE(GO:0000082) |  | 268 | 0.69 | 1.13 | 0.141 | 0.368 | 1.000 | 13095 | tags=32%, list=22%, signal=41% |
| 4414 | PHOSPHOLIPASE\_C\_ACTIVITY(GO:0004629) |  | 30 | 0.66 | 1.13 | 0.141 | 0.368 | 1.000 | 9042 | tags=23%, list=15%, signal=28% |
| 4415 | ESTROGEN\_METABOLIC\_PROCESS(GO:0008210) |  | 24 | 0.72 | 1.13 | 0.141 | 0.368 | 1.000 | 11544 | tags=33%, list=20%, signal=41% |
| 4416 | REGULATION\_OF\_TRANSLATION(GO:0006417) |  | 607 | 0.66 | 1.13 | 0.141 | 0.368 | 1.000 | 14716 | tags=25%, list=25%, signal=33% |
| 4417 | INTRACELLULAR\_LIGAND\_GATED\_ION\_CHANNEL\_ACTIVITY(GO:0005217) |  | 28 | 0.69 | 1.13 | 0.141 | 0.368 | 1.000 | 9703 | tags=32%, list=16%, signal=38% |
| 4418 | MONOVALENT\_INORGANIC\_ANION\_HOMEOSTASIS(GO:0055083) |  | 19 | 0.68 | 1.13 | 0.141 | 0.368 | 1.000 | 13932 | tags=47%, list=24%, signal=62% |
| 4419 | POSITIVE\_REGULATION\_OF\_STRESS\_ACTIVATED\_PROTEIN\_KINASE\_SIGNALING\_CASCADE(GO:0070304) |  | 150 | 0.70 | 1.13 | 0.141 | 0.368 | 1.000 | 8112 | tags=16%, list=14%, signal=18% |
| 4420 | ACTIVATION\_OF\_PROTEIN\_KINASE\_ACTIVITY(GO:0032147) |  | 321 | 0.67 | 1.13 | 0.141 | 0.368 | 1.000 | 10811 | tags=22%, list=18%, signal=27% |
| 4421 | ADENYL\_RIBONUCLEOTIDE\_BINDING(GO:0032559) |  | 1553 | 0.67 | 1.13 | 0.141 | 0.368 | 1.000 | 13347 | tags=34%, list=23%, signal=42% |
| 4422 | CELLULAR\_RESPONSE\_TO\_ZINC\_ION(GO:0071294) |  | 18 | 0.67 | 1.13 | 0.000 | 0.368 | 1.000 | 15462 | tags=39%, list=26%, signal=53% |
| 4423 | REGULATION\_OF\_DNA\_DEPENDENT\_DNA\_REPLICATION(GO:0090329) |  | 49 | 0.65 | 1.13 | 0.000 | 0.368 | 1.000 | 15720 | tags=65%, list=27%, signal=89% |
| 4424 | POST\_TRANSLATIONAL\_PROTEIN\_MODIFICATION(GO:0043687) |  | 441 | 0.66 | 1.13 | 0.141 | 0.368 | 1.000 | 12454 | tags=27%, list=21%, signal=34% |
| 4425 | POSITIVE\_REGULATION\_OF\_FIBROBLAST\_PROLIFERATION(GO:0048146) |  | 54 | 0.71 | 1.13 | 0.141 | 0.368 | 1.000 | 12241 | tags=33%, list=21%, signal=42% |
| 4426 | DETECTION\_OF\_CHEMICAL\_STIMULUS(GO:0009593) |  | 467 | 0.61 | 1.13 | 0.141 | 0.368 | 1.000 | 13901 | tags=9%, list=24%, signal=12% |
| 4427 | WALKING\_BEHAVIOR(GO:0090659) |  | 31 | 0.72 | 1.13 | 0.141 | 0.368 | 1.000 | 13179 | tags=58%, list=22%, signal=75% |
| 4428 | MACROPHAGE\_DIFFERENTIATION(GO:0030225) |  | 40 | 0.66 | 1.13 | 0.141 | 0.368 | 1.000 | 12402 | tags=30%, list=21%, signal=38% |
| 4429 | REGULATION\_OF\_SIGNALING\_RECEPTOR\_ACTIVITY(GO:0010469) |  | 544 | 0.68 | 1.13 | 0.141 | 0.368 | 1.000 | 10951 | tags=27%, list=19%, signal=33% |
| 4430 | RESPONSE\_TO\_GONADOTROPIN(GO:0034698) |  | 31 | 0.71 | 1.13 | 0.141 | 0.368 | 1.000 | 7172 | tags=16%, list=12%, signal=18% |
| 4431 | ADENYL\_NUCLEOTIDE\_BINDING(GO:0030554) |  | 1563 | 0.67 | 1.13 | 0.141 | 0.368 | 1.000 | 13347 | tags=34%, list=23%, signal=42% |
| 4432 | NEGATIVE\_REGULATION\_OF\_CATABOLIC\_PROCESS(GO:0009895) |  | 272 | 0.68 | 1.13 | 0.141 | 0.368 | 1.000 | 10469 | tags=15%, list=18%, signal=18% |
| 4433 | REGULATION\_OF\_SYNAPTIC\_TRANSMISSION\_\_GABAERGIC(GO:0032228) |  | 32 | 0.66 | 1.13 | 0.000 | 0.368 | 1.000 | 14929 | tags=47%, list=25%, signal=63% |
| 4434 | CARBOHYDRATE\_DERIVATIVE\_TRANSPORT(GO:1901264) |  | 70 | 0.69 | 1.13 | 0.141 | 0.368 | 1.000 | 13707 | tags=27%, list=23%, signal=35% |
| 4435 | ATP\_BINDING(GO:0005524) |  | 1518 | 0.67 | 1.13 | 0.141 | 0.368 | 1.000 | 13347 | tags=33%, list=23%, signal=42% |
| 4436 | NEGATIVE\_REGULATION\_OF\_TOLL\_LIKE\_RECEPTOR\_SIGNALING\_PATHWAY(GO:0034122) |  | 36 | 0.68 | 1.13 | 0.141 | 0.368 | 1.000 | 13678 | tags=39%, list=23%, signal=51% |
| 4437 | MRNA\_SPLICE\_SITE\_SELECTION(GO:0006376) |  | 29 | 0.69 | 1.13 | 0.000 | 0.368 | 1.000 | 17789 | tags=90%, list=30%, signal=128% |
| 4438 | MICROTUBULE\_CYTOSKELETON(GO:0015630) |  | 1134 | 0.67 | 1.13 | 0.141 | 0.368 | 1.000 | 13530 | tags=37%, list=23%, signal=47% |
| 4439 | LAMELLAR\_BODY(GO:0042599) |  | 19 | 0.71 | 1.13 | 0.141 | 0.368 | 1.000 | 6275 | tags=26%, list=11%, signal=29% |
| 4440 | BLOOD\_COAGULATION\_\_INTRINSIC\_PATHWAY(GO:0007597) |  | 19 | 0.67 | 1.13 | 0.141 | 0.368 | 1.000 | 15756 | tags=53%, list=27%, signal=72% |
| 4441 | REGULATION\_OF\_SISTER\_CHROMATID\_SEGREGATION(GO:0033045) |  | 73 | 0.64 | 1.13 | 0.000 | 0.368 | 1.000 | 19305 | tags=81%, list=33%, signal=120% |
| 4442 | MYOSIN\_COMPLEX(GO:0016459) |  | 72 | 0.67 | 1.13 | 0.141 | 0.368 | 1.000 | 10126 | tags=29%, list=17%, signal=35% |
| 4443 | STRUCTURAL\_CONSTITUENT\_OF\_NUCLEAR\_PORE(GO:0017056) |  | 28 | 0.67 | 1.13 | 0.000 | 0.368 | 1.000 | 18239 | tags=82%, list=31%, signal=119% |
| 4444 | MRNA\_MODIFICATION(GO:0016556) |  | 19 | 0.67 | 1.13 | 0.000 | 0.368 | 1.000 | 18912 | tags=89%, list=32%, signal=131% |
| 4445 | DEFENSE\_RESPONSE\_TO\_OTHER\_ORGANISM(GO:0098542) |  | 520 | 0.67 | 1.13 | 0.141 | 0.368 | 1.000 | 11578 | tags=17%, list=20%, signal=21% |
| 4446 | CATALYTIC\_COMPLEX(GO:1902494) |  | 1392 | 0.65 | 1.13 | 0.000 | 0.368 | 1.000 | 13813 | tags=31%, list=23%, signal=39% |
| 4447 | NEGATIVE\_REGULATION\_OF\_RESPONSE\_TO\_BIOTIC\_STIMULUS(GO:0002832) |  | 44 | 0.66 | 1.13 | 0.000 | 0.368 | 1.000 | 10185 | tags=20%, list=17%, signal=25% |
| 4448 | G2\_M\_TRANSITION\_OF\_MITOTIC\_CELL\_CYCLE(GO:0000086) |  | 254 | 0.65 | 1.13 | 0.000 | 0.368 | 1.000 | 16931 | tags=53%, list=29%, signal=74% |
| 4449 | RNA\_LOCALIZATION(GO:0006403) |  | 230 | 0.65 | 1.13 | 0.000 | 0.368 | 1.000 | 17552 | tags=66%, list=30%, signal=94% |
| 4450 | NEGATIVE\_REGULATION\_OF\_BONE\_MINERALIZATION(GO:0030502) |  | 16 | 0.71 | 1.13 | 0.141 | 0.368 | 1.000 | 9837 | tags=44%, list=17%, signal=52% |
| 4451 | PROTEIN\_TETRAMERIZATION(GO:0051262) |  | 153 | 0.68 | 1.13 | 0.141 | 0.368 | 1.000 | 12248 | tags=27%, list=21%, signal=34% |
| 4452 | DNA\_METABOLIC\_PROCESS(GO:0006259) |  | 1002 | 0.68 | 1.13 | 0.141 | 0.368 | 1.000 | 13434 | tags=35%, list=23%, signal=44% |
| 4453 | NEGATIVE\_REGULATION\_OF\_T\_CELL\_DIFFERENTIATION(GO:0045581) |  | 38 | 0.68 | 1.13 | 0.141 | 0.368 | 1.000 | 9411 | tags=16%, list=16%, signal=19% |
| 4454 | GLYCOPROTEIN\_METABOLIC\_PROCESS(GO:0009100) |  | 403 | 0.67 | 1.13 | 0.141 | 0.368 | 1.000 | 12346 | tags=28%, list=21%, signal=35% |
| 4455 | BICELLULAR\_TIGHT\_JUNCTION(GO:0005923) |  | 122 | 0.69 | 1.13 | 0.141 | 0.368 | 1.000 | 8340 | tags=14%, list=14%, signal=16% |
| 4456 | REGULATION\_OF\_G2\_M\_TRANSITION\_OF\_MITOTIC\_CELL\_CYCLE(GO:0010389) |  | 200 | 0.65 | 1.13 | 0.000 | 0.368 | 1.000 | 16830 | tags=52%, list=28%, signal=73% |
| 4457 | POSITIVE\_REGULATION\_OF\_TISSUE\_REMODELING(GO:0034105) |  | 43 | 0.67 | 1.13 | 0.141 | 0.368 | 1.000 | 11164 | tags=23%, list=19%, signal=29% |
| 4458 | PHOSPHATIDYLINOSITOL\_3\_PHOSPHATE\_BIOSYNTHETIC\_PROCESS(GO:0036092) |  | 50 | 0.64 | 1.13 | 0.141 | 0.368 | 1.000 | 16324 | tags=46%, list=28%, signal=63% |
| 4459 | PLASMA\_MEMBRANE\_BOUNDED\_CELL\_PROJECTION\_ASSEMBLY(GO:0120031) |  | 532 | 0.69 | 1.13 | 0.141 | 0.368 | 1.000 | 13564 | tags=40%, list=23%, signal=51% |
| 4460 | PROTEIN\_DNA\_COMPLEX\_SUBUNIT\_ORGANIZATION(GO:0071824) |  | 277 | 0.69 | 1.13 | 0.141 | 0.368 | 1.000 | 12766 | tags=27%, list=22%, signal=35% |
| 4461 | NAD\_METABOLIC\_PROCESS(GO:0019674) |  | 73 | 0.66 | 1.13 | 0.141 | 0.368 | 1.000 | 6654 | tags=8%, list=11%, signal=9% |
| 4462 | NEGATIVE\_REGULATION\_OF\_MORPHOGENESIS\_OF\_AN\_EPITHELIUM(GO:1905331) |  | 19 | 0.74 | 1.13 | 0.141 | 0.367 | 1.000 | 12645 | tags=37%, list=21%, signal=47% |
| 4463 | NEGATIVE\_REGULATION\_OF\_INTERFERON\_GAMMA\_PRODUCTION(GO:0032689) |  | 30 | 0.71 | 1.13 | 0.141 | 0.367 | 1.000 | 12217 | tags=30%, list=21%, signal=38% |
| 4464 | ORGANELLE\_FISSION(GO:0048285) |  | 439 | 0.65 | 1.13 | 0.141 | 0.367 | 1.000 | 17160 | tags=57%, list=29%, signal=79% |
| 4465 | NUCLEOSIDE\_BIOSYNTHETIC\_PROCESS(GO:0009163) |  | 48 | 0.63 | 1.13 | 0.000 | 0.367 | 1.000 | 17206 | tags=56%, list=29%, signal=79% |
| 4466 | INTRACILIARY\_TRANSPORT\_INVOLVED\_IN\_CILIUM\_ASSEMBLY(GO:0035735) |  | 40 | 0.66 | 1.13 | 0.000 | 0.367 | 1.000 | 15522 | tags=63%, list=26%, signal=85% |
| 4467 | CAMP\_BIOSYNTHETIC\_PROCESS(GO:0006171) |  | 66 | 0.65 | 1.13 | 0.141 | 0.367 | 1.000 | 7472 | tags=17%, list=13%, signal=19% |
| 4468 | ORGANOPHOSPHATE\_ESTER\_TRANSPORT(GO:0015748) |  | 90 | 0.69 | 1.13 | 0.141 | 0.367 | 1.000 | 8886 | tags=12%, list=15%, signal=14% |
| 4469 | MITOCHONDRIAL\_ENVELOPE(GO:0005740) |  | 755 | 0.65 | 1.13 | 0.141 | 0.367 | 1.000 | 13132 | tags=23%, list=22%, signal=30% |
| 4470 | NEGATIVE\_REGULATION\_OF\_CELLULAR\_CATABOLIC\_PROCESS(GO:0031330) |  | 231 | 0.69 | 1.13 | 0.141 | 0.367 | 1.000 | 6375 | tags=5%, list=11%, signal=6% |
| 4471 | PHOSPHATIDYLINOSITOL\_KINASE\_ACTIVITY(GO:0052742) |  | 50 | 0.64 | 1.13 | 0.141 | 0.367 | 1.000 | 17092 | tags=46%, list=29%, signal=65% |
| 4472 | REGULATION\_OF\_MICROTUBULE\_BASED\_MOVEMENT(GO:0060632) |  | 24 | 0.67 | 1.13 | 0.000 | 0.367 | 1.000 | 15770 | tags=67%, list=27%, signal=91% |
| 4473 | NUCLEOTIDE\_BINDING(GO:0000166) |  | 2157 | 0.69 | 1.13 | 0.141 | 0.367 | 1.000 | 13355 | tags=32%, list=23%, signal=40% |
| 4474 | NUCLEOSIDE\_PHOSPHATE\_BINDING(GO:1901265) |  | 2158 | 0.69 | 1.13 | 0.141 | 0.367 | 1.000 | 13355 | tags=32%, list=23%, signal=40% |
| 4475 | LYMPHOCYTE\_APOPTOTIC\_PROCESS(GO:0070227) |  | 69 | 0.68 | 1.13 | 0.141 | 0.367 | 1.000 | 13741 | tags=25%, list=23%, signal=32% |
| 4476 | ASPARTATE\_FAMILY\_AMINO\_ACID\_METABOLIC\_PROCESS(GO:0009066) |  | 54 | 0.66 | 1.13 | 0.000 | 0.367 | 1.000 | 11214 | tags=30%, list=19%, signal=37% |
| 4477 | CHANNEL\_INHIBITOR\_ACTIVITY(GO:0016248) |  | 37 | 0.66 | 1.13 | 0.141 | 0.367 | 1.000 | 9409 | tags=22%, list=16%, signal=26% |
| 4478 | DEFENSE\_RESPONSE\_TO\_BACTERIUM(GO:0042742) |  | 294 | 0.67 | 1.13 | 0.141 | 0.367 | 1.000 | 13053 | tags=17%, list=22%, signal=21% |
| 4479 | NEGATIVE\_REGULATION\_OF\_LEUKOCYTE\_MIGRATION(GO:0002686) |  | 37 | 0.68 | 1.13 | 0.141 | 0.367 | 1.000 | 9016 | tags=24%, list=15%, signal=29% |
| 4480 | GROWTH\_FACTOR\_RECEPTOR\_BINDING(GO:0070851) |  | 136 | 0.67 | 1.13 | 0.141 | 0.367 | 1.000 | 11343 | tags=29%, list=19%, signal=35% |
| 4481 | ANTIMICROBIAL\_HUMORAL\_IMMUNE\_RESPONSE\_MEDIATED\_BY\_ANTIMICROBIAL\_PEPTIDE(GO:0061844) |  | 57 | 0.69 | 1.13 | 0.141 | 0.367 | 1.000 | 10916 | tags=16%, list=18%, signal=19% |
| 4482 | CIRCADIAN\_SLEEP\_WAKE\_CYCLE(GO:0042745) |  | 27 | 0.72 | 1.13 | 0.141 | 0.367 | 1.000 | 7172 | tags=30%, list=12%, signal=34% |
| 4483 | SULFATION(GO:0051923) |  | 16 | 0.69 | 1.13 | 0.141 | 0.367 | 1.000 | 15691 | tags=50%, list=27%, signal=68% |
| 4484 | DERMATAN\_SULFATE\_PROTEOGLYCAN\_BIOSYNTHETIC\_PROCESS(GO:0050651) |  | 15 | 0.68 | 1.13 | 0.000 | 0.367 | 1.000 | 11459 | tags=47%, list=19%, signal=58% |
| 4485 | CYTOSKELETON\_DEPENDENT\_INTRACELLULAR\_TRANSPORT(GO:0030705) |  | 156 | 0.69 | 1.13 | 0.141 | 0.367 | 1.000 | 10185 | tags=20%, list=17%, signal=24% |
| 4486 | REGULATION\_OF\_PHOSPHOPROTEIN\_PHOSPHATASE\_ACTIVITY(GO:0043666) |  | 110 | 0.66 | 1.13 | 0.141 | 0.367 | 1.000 | 15304 | tags=38%, list=26%, signal=51% |
| 4487 | CELL\_CYCLE\_G1\_S\_PHASE\_TRANSITION(GO:0044843) |  | 286 | 0.68 | 1.13 | 0.141 | 0.367 | 1.000 | 13095 | tags=32%, list=22%, signal=41% |
| 4488 | CELL\_PROJECTION\_ASSEMBLY(GO:0030031) |  | 538 | 0.69 | 1.13 | 0.141 | 0.367 | 1.000 | 13564 | tags=40%, list=23%, signal=51% |
| 4489 | PROTEIN\_HOMOOLIGOMERIZATION(GO:0051260) |  | 323 | 0.68 | 1.13 | 0.141 | 0.367 | 1.000 | 13156 | tags=33%, list=22%, signal=42% |
| 4490 | REGULATION\_OF\_DENDRITIC\_SPINE\_DEVELOPMENT(GO:0060998) |  | 64 | 0.65 | 1.13 | 0.141 | 0.367 | 1.000 | 12327 | tags=33%, list=21%, signal=41% |
| 4491 | PROTEIN\_MODIFICATION\_BY\_SMALL\_PROTEIN\_CONJUGATION(GO:0032446) |  | 914 | 0.67 | 1.13 | 0.141 | 0.367 | 1.000 | 11101 | tags=18%, list=19%, signal=22% |
| 4492 | CD4\_POSITIVE\_\_ALPHA\_BETA\_T\_CELL\_LINEAGE\_COMMITMENT(GO:0043373) |  | 16 | 0.71 | 1.13 | 0.307 | 0.367 | 1.000 | 9023 | tags=25%, list=15%, signal=29% |
| 4493 | KINETOCHORE\_ASSEMBLY(GO:0051382) |  | 17 | 0.66 | 1.13 | 0.000 | 0.367 | 1.000 | 15072 | tags=65%, list=25%, signal=87% |
| 4494 | TUMOR\_NECROSIS\_FACTOR\_SECRETION(GO:1990774) |  | 20 | 0.72 | 1.13 | 0.141 | 0.367 | 1.000 | 7426 | tags=20%, list=13%, signal=23% |
| 4495 | TRANSFORMING\_GROWTH\_FACTOR\_BETA\_RECEPTOR\_BINDING(GO:0005160) |  | 50 | 0.68 | 1.13 | 0.141 | 0.367 | 1.000 | 13279 | tags=36%, list=22%, signal=46% |
| 4496 | DNA\_DIRECTED\_DNA\_POLYMERASE\_ACTIVITY(GO:0003887) |  | 28 | 0.64 | 1.13 | 0.000 | 0.367 | 1.000 | 20209 | tags=86%, list=34%, signal=130% |
| 4497 | B\_CELL\_HOMEOSTASIS(GO:0001782) |  | 28 | 0.63 | 1.13 | 0.000 | 0.367 | 1.000 | 15381 | tags=46%, list=26%, signal=63% |
| 4498 | RNA\_STEM\_LOOP\_BINDING(GO:0035613) |  | 15 | 0.65 | 1.13 | 0.000 | 0.367 | 1.000 | 18535 | tags=80%, list=31%, signal=116% |
| 4499 | PROTEIN\_DISULFIDE\_OXIDOREDUCTASE\_ACTIVITY(GO:0015035) |  | 25 | 0.65 | 1.13 | 0.000 | 0.367 | 1.000 | 17916 | tags=64%, list=30%, signal=92% |
| 4500 | NEGATIVE\_REGULATION\_OF\_FIBROBLAST\_GROWTH\_FACTOR\_RECEPTOR\_SIGNALING\_PATHWAY(GO:0040037) |  | 19 | 0.75 | 1.13 | 0.141 | 0.367 | 1.000 | 13062 | tags=32%, list=22%, signal=41% |
| 4501 | VIRAL\_ENTRY\_INTO\_HOST\_CELL(GO:0046718) |  | 120 | 0.68 | 1.13 | 0.141 | 0.367 | 1.000 | 7921 | tags=14%, list=13%, signal=16% |
| 4502 | POSITIVE\_REGULATION\_OF\_CHROMATIN\_ORGANIZATION(GO:1905269) |  | 91 | 0.67 | 1.13 | 0.141 | 0.367 | 1.000 | 10856 | tags=22%, list=18%, signal=27% |
| 4503 | SMOOTH\_MUSCLE\_CELL\_APOPTOTIC\_PROCESS(GO:0034390) |  | 30 | 0.71 | 1.13 | 0.141 | 0.367 | 1.000 | 13825 | tags=30%, list=23%, signal=39% |
| 4504 | REGULATION\_OF\_SMOOTH\_MUSCLE\_CELL\_APOPTOTIC\_PROCESS(GO:0034391) |  | 30 | 0.71 | 1.13 | 0.141 | 0.367 | 1.000 | 13825 | tags=30%, list=23%, signal=39% |
| 4505 | NEGATIVE\_REGULATION\_OF\_CELL\_PROJECTION\_ORGANIZATION(GO:0031345) |  | 161 | 0.69 | 1.13 | 0.141 | 0.367 | 1.000 | 9919 | tags=29%, list=17%, signal=35% |
| 4506 | PROTEIN\_NEDDYLATION(GO:0045116) |  | 16 | 0.69 | 1.13 | 0.000 | 0.367 | 1.000 | 12933 | tags=44%, list=22%, signal=56% |
| 4507 | RETINOL\_METABOLIC\_PROCESS(GO:0042572) |  | 31 | 0.68 | 1.13 | 0.141 | 0.367 | 1.000 | 12814 | tags=32%, list=22%, signal=41% |
| 4508 | NEGATIVE\_REGULATION\_OF\_PROTEIN\_UBIQUITINATION(GO:0031397) |  | 66 | 0.65 | 1.13 | 0.141 | 0.367 | 1.000 | 14185 | tags=32%, list=24%, signal=42% |
| 4509 | CELLULAR\_RESPONSE\_TO\_ALKALOID(GO:0071312) |  | 35 | 0.72 | 1.13 | 0.141 | 0.367 | 1.000 | 12241 | tags=43%, list=21%, signal=54% |
| 4510 | REGULATION\_OF\_CELL\_CYCLE\_G2\_M\_PHASE\_TRANSITION(GO:1902749) |  | 218 | 0.65 | 1.13 | 0.000 | 0.367 | 1.000 | 16830 | tags=53%, list=28%, signal=74% |
| 4511 | CELLULAR\_RESPONSE\_TO\_INTERLEUKIN\_4(GO:0071353) |  | 31 | 0.70 | 1.13 | 0.141 | 0.367 | 1.000 | 9411 | tags=23%, list=16%, signal=27% |
| 4512 | LIPID\_PHOSPHORYLATION(GO:0046834) |  | 112 | 0.68 | 1.13 | 0.141 | 0.366 | 1.000 | 10616 | tags=30%, list=18%, signal=37% |
| 4513 | NUCLEAR\_ENVELOPE(GO:0005635) |  | 447 | 0.68 | 1.13 | 0.141 | 0.367 | 1.000 | 14181 | tags=36%, list=24%, signal=46% |
| 4514 | DERMATAN\_SULFATE\_PROTEOGLYCAN\_METABOLIC\_PROCESS(GO:0050655) |  | 16 | 0.68 | 1.13 | 0.141 | 0.367 | 1.000 | 11459 | tags=44%, list=19%, signal=54% |
| 4515 | CONDENSED\_CHROMOSOME(GO:0000793) |  | 218 | 0.66 | 1.13 | 0.141 | 0.367 | 1.000 | 15552 | tags=56%, list=26%, signal=75% |
| 4516 | POSITIVE\_REGULATION\_OF\_OSTEOCLAST\_DIFFERENTIATION(GO:0045672) |  | 24 | 0.70 | 1.13 | 0.141 | 0.367 | 1.000 | 15462 | tags=50%, list=26%, signal=68% |
| 4517 | POSITIVE\_REGULATION\_OF\_ACTIN\_FILAMENT\_BUNDLE\_ASSEMBLY(GO:0032233) |  | 57 | 0.67 | 1.13 | 0.141 | 0.366 | 1.000 | 11474 | tags=23%, list=19%, signal=28% |
| 4518 | MICROTUBULE\_ORGANIZING\_CENTER\_PART(GO:0044450) |  | 167 | 0.66 | 1.13 | 0.141 | 0.366 | 1.000 | 15192 | tags=54%, list=26%, signal=72% |
| 4519 | POSITIVE\_REGULATION\_OF\_JUN\_KINASE\_ACTIVITY(GO:0043507) |  | 69 | 0.70 | 1.13 | 0.141 | 0.366 | 1.000 | 8112 | tags=17%, list=14%, signal=20% |
| 4520 | PHOSPHATASE\_COMPLEX(GO:1903293) |  | 47 | 0.65 | 1.13 | 0.000 | 0.366 | 1.000 | 6984 | tags=4%, list=12%, signal=5% |
| 4521 | PROTEIN\_SERINE\_THREONINE\_PHOSPHATASE\_COMPLEX(GO:0008287) |  | 47 | 0.65 | 1.13 | 0.000 | 0.366 | 1.000 | 6984 | tags=4%, list=12%, signal=5% |
| 4522 | CHROMATIN\_ASSEMBLY\_OR\_DISASSEMBLY(GO:0006333) |  | 187 | 0.68 | 1.13 | 0.141 | 0.366 | 1.000 | 13380 | tags=26%, list=23%, signal=34% |
| 4523 | INTRACILIARY\_TRANSPORT\_PARTICLE\_B(GO:0030992) |  | 21 | 0.68 | 1.13 | 0.000 | 0.366 | 1.000 | 15522 | tags=71%, list=26%, signal=97% |
| 4524 | REGULATION\_OF\_CAMP\_METABOLIC\_PROCESS(GO:0030814) |  | 76 | 0.64 | 1.13 | 0.141 | 0.366 | 1.000 | 7472 | tags=16%, list=13%, signal=18% |
| 4525 | LIGASE\_ACTIVITY(GO:0016874) |  | 172 | 0.64 | 1.13 | 0.000 | 0.366 | 1.000 | 13149 | tags=35%, list=22%, signal=45% |
| 4526 | DEPHOSPHORYLATION(GO:0016311) |  | 475 | 0.67 | 1.13 | 0.141 | 0.366 | 1.000 | 11680 | tags=22%, list=20%, signal=27% |
| 4527 | SIGNAL\_TRANSDUCTION\_BY\_P53\_CLASS\_MEDIATOR(GO:0072331) |  | 271 | 0.68 | 1.13 | 0.141 | 0.366 | 1.000 | 10674 | tags=20%, list=18%, signal=24% |
| 4528 | CELL\_CYCLE\_G2\_M\_PHASE\_TRANSITION(GO:0044839) |  | 270 | 0.65 | 1.13 | 0.000 | 0.366 | 1.000 | 16931 | tags=53%, list=29%, signal=73% |
| 4529 | PRODUCTION\_OF\_MOLECULAR\_MEDIATOR\_OF\_IMMUNE\_RESPONSE(GO:0002440) |  | 230 | 0.66 | 1.13 | 0.141 | 0.366 | 1.000 | 14181 | tags=27%, list=24%, signal=36% |
| 4530 | POSITIVE\_REGULATION\_OF\_ANION\_TRANSPORT(GO:1903793) |  | 44 | 0.67 | 1.13 | 0.141 | 0.366 | 1.000 | 13419 | tags=39%, list=23%, signal=50% |
| 4531 | EPITHELIAL\_CELL\_APOPTOTIC\_PROCESS(GO:1904019) |  | 103 | 0.69 | 1.13 | 0.141 | 0.366 | 1.000 | 9016 | tags=15%, list=15%, signal=17% |
| 4532 | POSITIVE\_REGULATION\_OF\_DEFENSE\_RESPONSE(GO:0031349) |  | 452 | 0.67 | 1.13 | 0.141 | 0.366 | 1.000 | 14441 | tags=29%, list=24%, signal=38% |
| 4533 | REGULATION\_OF\_EXIT\_FROM\_MITOSIS(GO:0007096) |  | 16 | 0.67 | 1.13 | 0.000 | 0.366 | 1.000 | 12001 | tags=50%, list=20%, signal=63% |
| 4534 | FIBRILLAR\_CENTER(GO:0001650) |  | 135 | 0.68 | 1.13 | 0.141 | 0.366 | 1.000 | 14820 | tags=42%, list=25%, signal=56% |
| 4535 | GTPASE\_BINDING(GO:0051020) |  | 641 | 0.67 | 1.13 | 0.141 | 0.366 | 1.000 | 12119 | tags=26%, list=20%, signal=32% |
| 4536 | DETECTION\_OF\_TEMPERATURE\_STIMULUS\_INVOLVED\_IN\_SENSORY\_PERCEPTION(GO:0050961) |  | 15 | 0.72 | 1.13 | 0.141 | 0.366 | 1.000 | 7873 | tags=33%, list=13%, signal=38% |
| 4537 | DETECTION\_OF\_TEMPERATURE\_STIMULUS\_INVOLVED\_IN\_SENSORY\_PERCEPTION\_OF\_PAIN(GO:0050965) |  | 15 | 0.72 | 1.13 | 0.141 | 0.366 | 1.000 | 7873 | tags=33%, list=13%, signal=38% |
| 4538 | NEGATIVE\_REGULATION\_OF\_GLIAL\_CELL\_DIFFERENTIATION(GO:0045686) |  | 23 | 0.73 | 1.13 | 0.141 | 0.366 | 1.000 | 7913 | tags=30%, list=13%, signal=35% |
| 4539 | POSITIVE\_REGULATION\_OF\_CYTOKINE\_PRODUCTION\_INVOLVED\_IN\_IMMUNE\_RESPONSE(GO:0002720) |  | 36 | 0.67 | 1.13 | 0.141 | 0.366 | 1.000 | 13741 | tags=33%, list=23%, signal=43% |
| 4540 | NEGATIVE\_ADAPTATION\_OF\_SIGNALING\_PATHWAY(GO:0022401) |  | 20 | 0.68 | 1.13 | 0.000 | 0.366 | 1.000 | 13268 | tags=50%, list=22%, signal=64% |
| 4541 | DESENSITIZATION\_OF\_G\_PROTEIN\_COUPLED\_RECEPTOR\_PROTEIN\_SIGNALING\_PATHWAY(GO:0002029) |  | 20 | 0.68 | 1.13 | 0.000 | 0.366 | 1.000 | 13268 | tags=50%, list=22%, signal=64% |
| 4542 | NEGATIVE\_REGULATION\_OF\_CELL\_DIVISION(GO:0051782) |  | 17 | 0.70 | 1.13 | 0.141 | 0.366 | 1.000 | 10637 | tags=35%, list=18%, signal=43% |
| 4543 | MODULATION\_BY\_HOST\_OF\_VIRAL\_PROCESS(GO:0044788) |  | 24 | 0.65 | 1.13 | 0.000 | 0.366 | 1.000 | 11583 | tags=38%, list=20%, signal=47% |
| 4544 | REGULATION\_OF\_PEPTIDYL\_SERINE\_PHOSPHORYLATION\_OF\_STAT\_PROTEIN(GO:0033139) |  | 23 | 0.68 | 1.13 | 0.141 | 0.366 | 1.000 | 13297 | tags=39%, list=22%, signal=50% |
| 4545 | REGULATION\_OF\_T\_CELL\_MEDIATED\_IMMUNITY(GO:0002709) |  | 50 | 0.67 | 1.12 | 0.141 | 0.366 | 1.000 | 15529 | tags=38%, list=26%, signal=51% |
| 4546 | NEGATIVE\_REGULATION\_OF\_STEROID\_METABOLIC\_PROCESS(GO:0045939) |  | 25 | 0.67 | 1.12 | 0.141 | 0.366 | 1.000 | 14948 | tags=48%, list=25%, signal=64% |
| 4547 | NEGATIVE\_REGULATION\_OF\_STEROID\_BIOSYNTHETIC\_PROCESS(GO:0010894) |  | 23 | 0.67 | 1.12 | 0.141 | 0.366 | 1.000 | 14948 | tags=52%, list=25%, signal=70% |
| 4548 | MYELOID\_CELL\_APOPTOTIC\_PROCESS(GO:0033028) |  | 32 | 0.66 | 1.12 | 0.141 | 0.366 | 1.000 | 10262 | tags=19%, list=17%, signal=23% |
| 4549 | REGULATION\_OF\_LEUKOCYTE\_MEDIATED\_CYTOTOXICITY(GO:0001910) |  | 54 | 0.65 | 1.12 | 0.141 | 0.366 | 1.000 | 16562 | tags=50%, list=28%, signal=69% |
| 4550 | GOLGI\_ASSOCIATED\_VESICLE(GO:0005798) |  | 156 | 0.64 | 1.12 | 0.141 | 0.366 | 1.000 | 13667 | tags=30%, list=23%, signal=39% |
| 4551 | GLYCOSYL\_COMPOUND\_BIOSYNTHETIC\_PROCESS(GO:1901659) |  | 50 | 0.62 | 1.12 | 0.000 | 0.366 | 1.000 | 17206 | tags=56%, list=29%, signal=79% |
| 4552 | INTRACILIARY\_TRANSPORT\_PARTICLE(GO:0030990) |  | 33 | 0.66 | 1.12 | 0.000 | 0.366 | 1.000 | 15522 | tags=67%, list=26%, signal=90% |
| 4553 | REGULATION\_OF\_MUSCLE\_CELL\_APOPTOTIC\_PROCESS(GO:0010660) |  | 84 | 0.64 | 1.12 | 0.141 | 0.366 | 1.000 | 14472 | tags=27%, list=24%, signal=36% |
| 4554 | RHO\_GTPASE\_BINDING(GO:0017048) |  | 177 | 0.68 | 1.12 | 0.141 | 0.366 | 1.000 | 14031 | tags=35%, list=24%, signal=46% |
| 4555 | MULTICELLULAR\_ORGANISM\_AGING(GO:0010259) |  | 29 | 0.71 | 1.12 | 0.141 | 0.366 | 1.000 | 11480 | tags=45%, list=19%, signal=56% |
| 4556 | CENTRIOLAR\_SATELLITE(GO:0034451) |  | 26 | 0.71 | 1.12 | 0.141 | 0.366 | 1.000 | 14028 | tags=65%, list=24%, signal=86% |
| 4557 | POSITIVE\_REGULATION\_OF\_STRESS\_FIBER\_ASSEMBLY(GO:0051496) |  | 48 | 0.68 | 1.12 | 0.141 | 0.366 | 1.000 | 11474 | tags=23%, list=19%, signal=28% |
| 4558 | COLLATERAL\_SPROUTING(GO:0048668) |  | 26 | 0.71 | 1.12 | 0.141 | 0.366 | 1.000 | 8673 | tags=35%, list=15%, signal=41% |
| 4559 | ISOPRENOID\_METABOLIC\_PROCESS(GO:0006720) |  | 127 | 0.68 | 1.12 | 0.141 | 0.366 | 1.000 | 11550 | tags=28%, list=20%, signal=35% |
| 4560 | PHOSPHORIC\_ESTER\_HYDROLASE\_ACTIVITY(GO:0042578) |  | 381 | 0.66 | 1.12 | 0.141 | 0.366 | 1.000 | 12794 | tags=29%, list=22%, signal=37% |
| 4561 | REGULATION\_OF\_ACTIN\_FILAMENT\_POLYMERIZATION(GO:0030833) |  | 146 | 0.66 | 1.12 | 0.141 | 0.366 | 1.000 | 7951 | tags=10%, list=13%, signal=12% |
| 4562 | RAN\_GTPASE\_BINDING(GO:0008536) |  | 34 | 0.64 | 1.12 | 0.000 | 0.366 | 1.000 | 16538 | tags=74%, list=28%, signal=102% |
| 4563 | SMALL\_GTPASE\_MEDIATED\_SIGNAL\_TRANSDUCTION(GO:0007264) |  | 497 | 0.68 | 1.12 | 0.141 | 0.366 | 1.000 | 13969 | tags=33%, list=24%, signal=43% |
| 4564 | RECEPTOR\_BIOSYNTHETIC\_PROCESS(GO:0032800) |  | 24 | 0.68 | 1.12 | 0.141 | 0.366 | 1.000 | 10604 | tags=29%, list=18%, signal=36% |
| 4565 | ACTIN\_FILAMENT\_POLYMERIZATION(GO:0030041) |  | 161 | 0.65 | 1.12 | 0.141 | 0.366 | 1.000 | 9686 | tags=14%, list=16%, signal=16% |
| 4566 | ESTABLISHMENT\_OF\_PROTEIN\_LOCALIZATION\_TO\_TELOMERE(GO:0070200) |  | 15 | 0.65 | 1.12 | 0.000 | 0.366 | 1.000 | 19773 | tags=93%, list=33%, signal=140% |
| 4567 | RESPONSE\_TO\_DIETARY\_EXCESS(GO:0002021) |  | 22 | 0.70 | 1.12 | 0.141 | 0.366 | 1.000 | 8818 | tags=27%, list=15%, signal=32% |
| 4568 | NEGATIVE\_REGULATION\_OF\_GLIOGENESIS(GO:0014014) |  | 39 | 0.72 | 1.12 | 0.141 | 0.366 | 1.000 | 7913 | tags=23%, list=13%, signal=27% |
| 4569 | OXIDOREDUCTASE\_ACTIVITY\_\_ACTING\_ON\_THE\_CH\_NH2\_GROUP\_OF\_DONORS(GO:0016638) |  | 21 | 0.71 | 1.12 | 0.141 | 0.366 | 1.000 | 8602 | tags=33%, list=15%, signal=39% |
| 4570 | REGULATION\_OF\_RESPONSE\_TO\_NUTRIENT\_LEVELS(GO:0032107) |  | 24 | 0.66 | 1.12 | 0.000 | 0.366 | 1.000 | 18462 | tags=63%, list=31%, signal=91% |
| 4571 | REGULATION\_OF\_RESPONSE\_TO\_EXTRACELLULAR\_STIMULUS(GO:0032104) |  | 24 | 0.66 | 1.12 | 0.000 | 0.366 | 1.000 | 18462 | tags=63%, list=31%, signal=91% |
| 4572 | MEMBRANE\_LIPID\_BIOSYNTHETIC\_PROCESS(GO:0046467) |  | 141 | 0.67 | 1.12 | 0.141 | 0.366 | 1.000 | 9651 | tags=19%, list=16%, signal=23% |
| 4573 | INTRINSIC\_COMPONENT\_OF\_GOLGI\_MEMBRANE(GO:0031228) |  | 51 | 0.64 | 1.12 | 0.141 | 0.366 | 1.000 | 18337 | tags=61%, list=31%, signal=88% |
| 4574 | TRANSMITTER\_GATED\_ION\_CHANNEL\_ACTIVITY(GO:0022824) |  | 50 | 0.69 | 1.12 | 0.141 | 0.366 | 1.000 | 14022 | tags=46%, list=24%, signal=60% |
| 4575 | TRANSMITTER\_GATED\_CHANNEL\_ACTIVITY(GO:0022835) |  | 50 | 0.69 | 1.12 | 0.141 | 0.366 | 1.000 | 14022 | tags=46%, list=24%, signal=60% |
| 4576 | ATPASE\_ACTIVITY\_\_COUPLED(GO:0042623) |  | 362 | 0.67 | 1.12 | 0.141 | 0.366 | 1.000 | 13323 | tags=37%, list=23%, signal=48% |
| 4577 | CHROMOSOME\_\_CENTROMERIC\_REGION(GO:0000775) |  | 194 | 0.64 | 1.12 | 0.000 | 0.366 | 1.000 | 17739 | tags=68%, list=30%, signal=96% |
| 4578 | GENE\_SILENCING\_BY\_RNA(GO:0031047) |  | 350 | 0.67 | 1.12 | 0.141 | 0.366 | 1.000 | 5858 | tags=6%, list=10%, signal=6% |
| 4579 | PHOSPHOLIPID\_TRANSLOCATING\_ATPASE\_ACTIVITY(GO:0004012) |  | 20 | 0.69 | 1.12 | 0.141 | 0.366 | 1.000 | 13125 | tags=30%, list=22%, signal=39% |
| 4580 | NUCLEAR\_CHROMOSOME\_SEGREGATION(GO:0098813) |  | 304 | 0.64 | 1.12 | 0.000 | 0.366 | 1.000 | 17170 | tags=64%, list=29%, signal=89% |
| 4581 | ER\_TO\_GOLGI\_VESICLE\_MEDIATED\_TRANSPORT(GO:0006888) |  | 184 | 0.64 | 1.12 | 0.141 | 0.366 | 1.000 | 17553 | tags=47%, list=30%, signal=67% |
| 4582 | CELLULAR\_RESPONSE\_TO\_CARBOHYDRATE\_STIMULUS(GO:0071322) |  | 136 | 0.66 | 1.12 | 0.141 | 0.366 | 1.000 | 11397 | tags=26%, list=19%, signal=32% |
| 4583 | POSITIVE\_REGULATION\_OF\_PROTEOLYSIS\_INVOLVED\_IN\_CELLULAR\_PROTEIN\_CATABOLIC\_PROCESS(GO:1903052) |  | 117 | 0.65 | 1.12 | 0.141 | 0.366 | 1.000 | 14983 | tags=35%, list=25%, signal=47% |
| 4584 | POSITIVE\_REGULATION\_OF\_PLASMA\_MEMBRANE\_BOUNDED\_CELL\_PROJECTION\_ASSEMBLY(GO:0120034) |  | 89 | 0.68 | 1.12 | 0.141 | 0.366 | 1.000 | 10939 | tags=24%, list=18%, signal=29% |
| 4585 | GUANYL\_NUCLEOTIDE\_EXCHANGE\_FACTOR\_ACTIVITY(GO:0005085) |  | 334 | 0.67 | 1.12 | 0.141 | 0.366 | 1.000 | 12000 | tags=29%, list=20%, signal=36% |
| 4586 | DNA\_SECONDARY\_STRUCTURE\_BINDING(GO:0000217) |  | 23 | 0.68 | 1.12 | 0.000 | 0.365 | 1.000 | 17317 | tags=87%, list=29%, signal=123% |
| 4587 | EXORIBONUCLEASE\_ACTIVITY(GO:0004532) |  | 35 | 0.65 | 1.12 | 0.000 | 0.365 | 1.000 | 17651 | tags=77%, list=30%, signal=110% |
| 4588 | MUSCLE\_CELL\_APOPTOTIC\_PROCESS(GO:0010657) |  | 89 | 0.64 | 1.12 | 0.141 | 0.366 | 1.000 | 14472 | tags=28%, list=24%, signal=37% |
| 4589 | PROTEIN\_UBIQUITINATION(GO:0016567) |  | 825 | 0.66 | 1.12 | 0.141 | 0.366 | 1.000 | 11094 | tags=18%, list=19%, signal=22% |
| 4590 | CATALYTIC\_ACTIVITY\_\_ACTING\_ON\_DNA(GO:0140097) |  | 189 | 0.69 | 1.12 | 0.141 | 0.366 | 1.000 | 13412 | tags=33%, list=23%, signal=42% |
| 4591 | MITOTIC\_SPINDLE\_ASSEMBLY(GO:0090307) |  | 59 | 0.63 | 1.12 | 0.000 | 0.366 | 1.000 | 15695 | tags=59%, list=27%, signal=81% |
| 4592 | NEGATIVE\_REGULATION\_OF\_SYSTEMIC\_ARTERIAL\_BLOOD\_PRESSURE(GO:0003085) |  | 17 | 0.70 | 1.12 | 0.141 | 0.366 | 1.000 | 17432 | tags=53%, list=29%, signal=75% |
| 4593 | OXIDOREDUCTASE\_ACTIVITY\_\_ACTING\_ON\_THE\_CH\_NH2\_GROUP\_OF\_DONORS\_\_OXYGEN\_AS\_ACCEPTOR(GO:0016641) |  | 16 | 0.71 | 1.12 | 0.141 | 0.366 | 1.000 | 7225 | tags=25%, list=12%, signal=28% |
| 4594 | STARTLE\_RESPONSE(GO:0001964) |  | 27 | 0.68 | 1.12 | 0.141 | 0.366 | 1.000 | 14441 | tags=44%, list=24%, signal=59% |
| 4595 | CELL\_DIFFERENTIATION\_IN\_HINDBRAIN(GO:0021533) |  | 22 | 0.65 | 1.12 | 0.141 | 0.366 | 1.000 | 13932 | tags=45%, list=24%, signal=59% |
| 4596 | LEUKOCYTE\_APOPTOTIC\_PROCESS(GO:0071887) |  | 100 | 0.68 | 1.12 | 0.141 | 0.365 | 1.000 | 13741 | tags=25%, list=23%, signal=33% |
| 4597 | EXORIBONUCLEASE\_ACTIVITY\_\_PRODUCING\_5'\_PHOSPHOMONOESTERS(GO:0016896) |  | 33 | 0.66 | 1.12 | 0.000 | 0.365 | 1.000 | 17651 | tags=82%, list=30%, signal=117% |
| 4598 | PHASIC\_SMOOTH\_MUSCLE\_CONTRACTION(GO:0014821) |  | 21 | 0.70 | 1.12 | 0.141 | 0.365 | 1.000 | 6952 | tags=24%, list=12%, signal=27% |
| 4599 | HISTONE\_UBIQUITINATION(GO:0016574) |  | 45 | 0.64 | 1.12 | 0.141 | 0.365 | 1.000 | 10434 | tags=18%, list=18%, signal=22% |
| 4600 | POSITIVE\_REGULATION\_OF\_JNK\_CASCADE(GO:0046330) |  | 122 | 0.69 | 1.12 | 0.141 | 0.366 | 1.000 | 12564 | tags=28%, list=21%, signal=35% |
| 4601 | SEH1\_ASSOCIATED\_COMPLEX(GO:0035859) |  | 16 | 0.67 | 1.12 | 0.000 | 0.366 | 1.000 | 14928 | tags=38%, list=25%, signal=50% |
| 4602 | PROTEIN\_KINASE\_C\_ACTIVITY(GO:0004697) |  | 16 | 0.68 | 1.12 | 0.000 | 0.366 | 1.000 | 11049 | tags=44%, list=19%, signal=54% |
| 4603 | REGULATION\_OF\_MITOTIC\_CELL\_CYCLE\_PHASE\_TRANSITION(GO:1901990) |  | 412 | 0.67 | 1.12 | 0.141 | 0.366 | 1.000 | 13421 | tags=31%, list=23%, signal=40% |
| 4604 | CENTRIOLE(GO:0005814) |  | 123 | 0.66 | 1.12 | 0.141 | 0.366 | 1.000 | 14176 | tags=48%, list=24%, signal=63% |
| 4605 | NEGATIVE\_REGULATION\_OF\_LYMPHOCYTE\_ACTIVATION(GO:0051250) |  | 129 | 0.67 | 1.12 | 0.141 | 0.366 | 1.000 | 10712 | tags=22%, list=18%, signal=26% |
| 4606 | PROTEIN\_PHOSPHATASE\_REGULATOR\_ACTIVITY(GO:0019888) |  | 79 | 0.66 | 1.12 | 0.141 | 0.365 | 1.000 | 6922 | tags=10%, list=12%, signal=11% |
| 4607 | PYROPHOSPHATASE\_ACTIVITY(GO:0016462) |  | 876 | 0.68 | 1.12 | 0.141 | 0.365 | 1.000 | 13355 | tags=33%, list=23%, signal=42% |
| 4608 | NEGATIVE\_REGULATION\_OF\_ALPHA\_BETA\_T\_CELL\_DIFFERENTIATION(GO:0046639) |  | 18 | 0.70 | 1.12 | 0.141 | 0.365 | 1.000 | 13828 | tags=44%, list=23%, signal=58% |
| 4609 | NUCLEOTIDE\_BINDING\_DOMAIN\_\_LEUCINE\_RICH\_REPEAT\_CONTAINING\_RECEPTOR\_SIGNALING\_PATHWAY(GO:0035872) |  | 38 | 0.63 | 1.12 | 0.000 | 0.365 | 1.000 | 17085 | tags=58%, list=29%, signal=81% |
| 4610 | NEGATIVE\_REGULATION\_OF\_VASCULAR\_ASSOCIATED\_SMOOTH\_MUSCLE\_CELL\_MIGRATION(GO:1904753) |  | 15 | 0.81 | 1.12 | 0.000 | 0.365 | 1.000 | 8774 | tags=27%, list=15%, signal=31% |
| 4611 | HYDROLASE\_ACTIVITY\_\_ACTING\_ON\_ACID\_ANHYDRIDES\_\_IN\_PHOSPHORUS\_CONTAINING\_ANHYDRIDES(GO:0016818) |  | 879 | 0.68 | 1.12 | 0.141 | 0.365 | 1.000 | 13355 | tags=33%, list=23%, signal=42% |
| 4612 | HYDROLASE\_ACTIVITY\_\_ACTING\_ON\_ACID\_ANHYDRIDES(GO:0016817) |  | 879 | 0.68 | 1.12 | 0.141 | 0.365 | 1.000 | 13355 | tags=33%, list=23%, signal=42% |
| 4613 | CELLULAR\_RESPONSE\_TO\_UV(GO:0034644) |  | 76 | 0.64 | 1.12 | 0.000 | 0.365 | 1.000 | 17367 | tags=62%, list=29%, signal=87% |
| 4614 | INNER\_MITOCHONDRIAL\_MEMBRANE\_PROTEIN\_COMPLEX(GO:0098800) |  | 144 | 0.63 | 1.12 | 0.141 | 0.365 | 1.000 | 5894 | tags=4%, list=10%, signal=5% |
| 4615 | REGULATION\_OF\_ALPHA\_BETA\_T\_CELL\_DIFFERENTIATION(GO:0046637) |  | 53 | 0.68 | 1.12 | 0.141 | 0.365 | 1.000 | 14175 | tags=43%, list=24%, signal=57% |
| 4616 | ENDOSOME(GO:0005768) |  | 869 | 0.67 | 1.12 | 0.141 | 0.365 | 1.000 | 9637 | tags=14%, list=16%, signal=17% |
| 4617 | DNA\_MODIFICATION(GO:0006304) |  | 110 | 0.69 | 1.12 | 0.141 | 0.365 | 1.000 | 8202 | tags=12%, list=14%, signal=14% |
| 4618 | FIBROBLAST\_GROWTH\_FACTOR\_BINDING(GO:0017134) |  | 23 | 0.70 | 1.12 | 0.141 | 0.365 | 1.000 | 11519 | tags=30%, list=19%, signal=38% |
| 4619 | REGULATION\_OF\_T\_CELL\_PROLIFERATION(GO:0042129) |  | 147 | 0.64 | 1.12 | 0.141 | 0.365 | 1.000 | 16684 | tags=37%, list=28%, signal=52% |
| 4620 | DNA\_PACKAGING(GO:0006323) |  | 206 | 0.68 | 1.12 | 0.141 | 0.365 | 1.000 | 13828 | tags=30%, list=23%, signal=39% |
| 4621 | APICAL\_JUNCTION\_COMPLEX(GO:0043296) |  | 141 | 0.68 | 1.12 | 0.141 | 0.365 | 1.000 | 8937 | tags=15%, list=15%, signal=18% |
| 4622 | GOLGI\_SUBCOMPARTMENT(GO:0098791) |  | 795 | 0.66 | 1.12 | 0.141 | 0.365 | 1.000 | 11641 | tags=21%, list=20%, signal=25% |
| 4623 | NEGATIVE\_REGULATION\_OF\_CALCIUM\_ION\_TRANSPORT\_INTO\_CYTOSOL(GO:0010523) |  | 20 | 0.68 | 1.12 | 0.141 | 0.365 | 1.000 | 10245 | tags=40%, list=17%, signal=48% |
| 4624 | NEGATIVE\_REGULATION\_OF\_PEPTIDYL\_TYROSINE\_PHOSPHORYLATION(GO:0050732) |  | 47 | 0.68 | 1.12 | 0.141 | 0.365 | 1.000 | 7170 | tags=9%, list=12%, signal=10% |
| 4625 | POSITIVE\_REGULATION\_OF\_RESPONSE\_TO\_OXIDATIVE\_STRESS(GO:1902884) |  | 21 | 0.66 | 1.12 | 0.141 | 0.365 | 1.000 | 14948 | tags=33%, list=25%, signal=45% |
| 4626 | RNA\_PHOSPHODIESTER\_BOND\_HYDROLYSIS\_\_EXONUCLEOLYTIC(GO:0090503) |  | 36 | 0.65 | 1.12 | 0.000 | 0.365 | 1.000 | 17651 | tags=78%, list=30%, signal=111% |
| 4627 | SOMATIC\_DIVERSIFICATION\_OF\_IMMUNE\_RECEPTORS(GO:0002200) |  | 62 | 0.69 | 1.12 | 0.141 | 0.365 | 1.000 | 10412 | tags=31%, list=18%, signal=37% |
| 4628 | REGULATION\_OF\_PROTEIN\_DEPHOSPHORYLATION(GO:0035304) |  | 128 | 0.65 | 1.12 | 0.141 | 0.365 | 1.000 | 15304 | tags=37%, list=26%, signal=49% |
| 4629 | NUCLEOTIDE\_BINDING\_OLIGOMERIZATION\_DOMAIN\_CONTAINING\_SIGNALING\_PATHWAY(GO:0070423) |  | 37 | 0.63 | 1.12 | 0.000 | 0.365 | 1.000 | 17085 | tags=57%, list=29%, signal=80% |
| 4630 | CELLULAR\_PROTEIN\_CONTAINING\_COMPLEX\_ASSEMBLY(GO:0034622) |  | 1075 | 0.66 | 1.12 | 0.141 | 0.365 | 1.000 | 15578 | tags=40%, list=26%, signal=54% |
| 4631 | POSITIVE\_REGULATION\_OF\_CYTOKINESIS(GO:0032467) |  | 36 | 0.66 | 1.12 | 0.000 | 0.365 | 1.000 | 15403 | tags=44%, list=26%, signal=60% |
| 4632 | POSITIVE\_REGULATION\_OF\_FATTY\_ACID\_BIOSYNTHETIC\_PROCESS(GO:0045723) |  | 16 | 0.65 | 1.12 | 0.000 | 0.365 | 1.000 | 17139 | tags=56%, list=29%, signal=79% |
| 4633 | SPERM\_PRINCIPAL\_PIECE(GO:0097228) |  | 24 | 0.72 | 1.12 | 0.141 | 0.365 | 1.000 | 13081 | tags=63%, list=22%, signal=80% |
| 4634 | LABYRINTHINE\_LAYER\_MORPHOGENESIS(GO:0060713) |  | 20 | 0.73 | 1.12 | 0.141 | 0.365 | 1.000 | 7634 | tags=20%, list=13%, signal=23% |
| 4635 | PATHWAY\_RESTRICTED\_SMAD\_PROTEIN\_PHOSPHORYLATION(GO:0060389) |  | 64 | 0.67 | 1.12 | 0.141 | 0.365 | 1.000 | 13279 | tags=38%, list=22%, signal=48% |
| 4636 | CATALYTIC\_ACTIVITY\_\_ACTING\_ON\_RNA(GO:0140098) |  | 382 | 0.63 | 1.12 | 0.000 | 0.365 | 1.000 | 18471 | tags=60%, list=31%, signal=87% |
| 4637 | GLUCAN\_BIOSYNTHETIC\_PROCESS(GO:0009250) |  | 51 | 0.68 | 1.12 | 0.141 | 0.365 | 1.000 | 8951 | tags=18%, list=15%, signal=21% |
| 4638 | GLYCOGEN\_BIOSYNTHETIC\_PROCESS(GO:0005978) |  | 51 | 0.68 | 1.12 | 0.141 | 0.365 | 1.000 | 8951 | tags=18%, list=15%, signal=21% |
| 4639 | HYDROLASE\_ACTIVITY\_\_ACTING\_ON\_ESTER\_BONDS(GO:0016788) |  | 779 | 0.66 | 1.12 | 0.141 | 0.365 | 1.000 | 13097 | tags=29%, list=22%, signal=37% |
| 4640 | REGULATION\_OF\_PROTEASOMAL\_UBIQUITIN\_DEPENDENT\_PROTEIN\_CATABOLIC\_PROCESS(GO:0032434) |  | 118 | 0.65 | 1.12 | 0.141 | 0.365 | 1.000 | 15098 | tags=37%, list=26%, signal=50% |
| 4641 | REPLISOME(GO:0030894) |  | 28 | 0.64 | 1.12 | 0.000 | 0.365 | 1.000 | 17064 | tags=75%, list=29%, signal=105% |
| 4642 | POSITIVE\_REGULATION\_OF\_HISTONE\_METHYLATION(GO:0031062) |  | 36 | 0.65 | 1.12 | 0.000 | 0.365 | 1.000 | 16613 | tags=53%, list=28%, signal=73% |
| 4643 | 3'\_5'\_EXORIBONUCLEASE\_ACTIVITY(GO:0000175) |  | 31 | 0.66 | 1.12 | 0.000 | 0.365 | 1.000 | 17651 | tags=81%, list=30%, signal=115% |
| 4644 | TRANSFERASE\_COMPLEX(GO:1990234) |  | 779 | 0.64 | 1.12 | 0.141 | 0.365 | 1.000 | 17881 | tags=56%, list=30%, signal=79% |
| 4645 | POSITIVE\_REGULATION\_OF\_T\_CELL\_PROLIFERATION(GO:0042102) |  | 93 | 0.65 | 1.12 | 0.141 | 0.365 | 1.000 | 16684 | tags=38%, list=28%, signal=52% |
| 4646 | CELLULAR\_MONOVALENT\_INORGANIC\_ANION\_HOMEOSTASIS(GO:0030320) |  | 15 | 0.69 | 1.12 | 0.141 | 0.365 | 1.000 | 13932 | tags=53%, list=24%, signal=70% |
| 4647 | CELLULAR\_ANION\_HOMEOSTASIS(GO:0030002) |  | 15 | 0.69 | 1.12 | 0.141 | 0.365 | 1.000 | 13932 | tags=53%, list=24%, signal=70% |
| 4648 | REGULATION\_OF\_MITOTIC\_SPINDLE\_ASSEMBLY(GO:1901673) |  | 19 | 0.65 | 1.12 | 0.000 | 0.365 | 1.000 | 15444 | tags=53%, list=26%, signal=71% |
| 4649 | REGULATION\_OF\_RESPONSE\_TO\_FOOD(GO:0032095) |  | 18 | 0.66 | 1.12 | 0.000 | 0.365 | 1.000 | 18462 | tags=61%, list=31%, signal=89% |
| 4650 | NEGATIVE\_REGULATION\_OF\_PROTEIN\_OLIGOMERIZATION(GO:0032460) |  | 17 | 0.69 | 1.12 | 0.141 | 0.365 | 1.000 | 6240 | tags=6%, list=11%, signal=7% |
| 4651 | POSITIVE\_REGULATION\_OF\_ICOSANOID\_SECRETION(GO:0032305) |  | 15 | 0.71 | 1.12 | 0.141 | 0.365 | 1.000 | 11372 | tags=27%, list=19%, signal=33% |
| 4652 | GOLGI\_APPARATUS\_PART(GO:0044431) |  | 910 | 0.65 | 1.12 | 0.141 | 0.365 | 1.000 | 11641 | tags=21%, list=20%, signal=26% |
| 4653 | PROTEIN\_HOMOTRIMERIZATION(GO:0070207) |  | 28 | 0.67 | 1.12 | 0.141 | 0.365 | 1.000 | 16434 | tags=57%, list=28%, signal=79% |
| 4654 | POSITIVE\_REGULATION\_OF\_FILOPODIUM\_ASSEMBLY(GO:0051491) |  | 30 | 0.69 | 1.12 | 0.141 | 0.365 | 1.000 | 16153 | tags=57%, list=27%, signal=78% |
| 4655 | NUCLEAR\_REPLICATION\_FORK(GO:0043596) |  | 40 | 0.64 | 1.12 | 0.000 | 0.365 | 1.000 | 17066 | tags=75%, list=29%, signal=105% |
| 4656 | POSITIVE\_REGULATION\_OF\_LIPID\_LOCALIZATION(GO:1905954) |  | 74 | 0.67 | 1.12 | 0.141 | 0.365 | 1.000 | 12613 | tags=27%, list=21%, signal=34% |
| 4657 | DNA\_REPAIR(GO:0006281) |  | 536 | 0.67 | 1.12 | 0.141 | 0.365 | 1.000 | 14200 | tags=39%, list=24%, signal=51% |
| 4658 | CHROMOSOME\_ORGANIZATION(GO:0051276) |  | 1220 | 0.67 | 1.12 | 0.141 | 0.365 | 1.000 | 14441 | tags=37%, list=24%, signal=48% |
| 4659 | REGULATION\_OF\_ALPHA\_BETA\_T\_CELL\_ACTIVATION(GO:0046634) |  | 73 | 0.67 | 1.12 | 0.141 | 0.365 | 1.000 | 14175 | tags=40%, list=24%, signal=52% |
| 4660 | SODIUM\_INDEPENDENT\_ORGANIC\_ANION\_TRANSPORT(GO:0043252) |  | 29 | 0.70 | 1.12 | 0.141 | 0.365 | 1.000 | 11359 | tags=38%, list=19%, signal=47% |
| 4661 | SYNAPTIC\_TRANSMISSION\_\_CHOLINERGIC(GO:0007271) |  | 35 | 0.68 | 1.12 | 0.141 | 0.365 | 1.000 | 13932 | tags=43%, list=24%, signal=56% |
| 4662 | VESICLE\_ORGANIZATION(GO:0016050) |  | 385 | 0.65 | 1.12 | 0.141 | 0.365 | 1.000 | 10763 | tags=16%, list=18%, signal=20% |
| 4663 | MAP\_KINASE\_KINASE\_KINASE\_ACTIVITY(GO:0004709) |  | 26 | 0.64 | 1.12 | 0.141 | 0.365 | 1.000 | 14456 | tags=38%, list=24%, signal=51% |
| 4664 | NUCLEOSIDE\_TRIPHOSPHATASE\_ACTIVITY(GO:0017111) |  | 834 | 0.68 | 1.12 | 0.141 | 0.365 | 1.000 | 13434 | tags=33%, list=23%, signal=42% |
| 4665 | HAIR\_FOLLICLE\_MATURATION(GO:0048820) |  | 16 | 0.71 | 1.12 | 0.141 | 0.365 | 1.000 | 9633 | tags=25%, list=16%, signal=30% |
| 4666 | SMALL\_SUBUNIT\_PROCESSOME(GO:0032040) |  | 36 | 0.62 | 1.12 | 0.000 | 0.365 | 1.000 | 19119 | tags=78%, list=32%, signal=115% |
| 4667 | POSITIVE\_REGULATION\_OF\_CHROMOSOME\_ORGANIZATION(GO:2001252) |  | 159 | 0.64 | 1.12 | 0.141 | 0.365 | 1.000 | 16693 | tags=52%, list=28%, signal=73% |
| 4668 | TRIVALENT\_INORGANIC\_CATION\_TRANSPORT(GO:0072512) |  | 39 | 0.66 | 1.12 | 0.141 | 0.365 | 1.000 | 7467 | tags=8%, list=13%, signal=9% |
| 4669 | FERRIC\_IRON\_TRANSPORT(GO:0015682) |  | 39 | 0.66 | 1.12 | 0.141 | 0.365 | 1.000 | 7467 | tags=8%, list=13%, signal=9% |
| 4670 | REGULATION\_OF\_CHROMOSOME\_SEGREGATION(GO:0051983) |  | 96 | 0.64 | 1.12 | 0.000 | 0.365 | 1.000 | 19305 | tags=80%, list=33%, signal=119% |
| 4671 | PROTEIN\_AUTOPROCESSING(GO:0016540) |  | 15 | 0.71 | 1.12 | 0.141 | 0.365 | 1.000 | 11480 | tags=40%, list=19%, signal=50% |
| 4672 | OUTER\_MITOCHONDRIAL\_MEMBRANE\_PROTEIN\_COMPLEX(GO:0098799) |  | 17 | 0.64 | 1.12 | 0.000 | 0.365 | 1.000 | 18967 | tags=76%, list=32%, signal=113% |
| 4673 | PROTEIN\_CONTAINING\_COMPLEX\_LOCALIZATION(GO:0031503) |  | 234 | 0.64 | 1.12 | 0.000 | 0.365 | 1.000 | 17443 | tags=59%, list=29%, signal=83% |
| 4674 | CYTOKINESIS(GO:0000910) |  | 134 | 0.63 | 1.12 | 0.000 | 0.365 | 1.000 | 17160 | tags=49%, list=29%, signal=69% |
| 4675 | MATURE\_B\_CELL\_DIFFERENTIATION(GO:0002335) |  | 22 | 0.68 | 1.12 | 0.141 | 0.365 | 1.000 | 13707 | tags=36%, list=23%, signal=47% |
| 4676 | NUCLEAR\_LUMEN(GO:0031981) |  | 4076 | 0.69 | 1.12 | 0.141 | 0.364 | 1.000 | 13192 | tags=30%, list=22%, signal=35% |
| 4677 | GOLGI\_MEMBRANE(GO:0000139) |  | 706 | 0.65 | 1.12 | 0.141 | 0.364 | 1.000 | 11640 | tags=21%, list=20%, signal=26% |
| 4678 | PROTEIN\_TRANSPORT(GO:0015031) |  | 1869 | 0.68 | 1.12 | 0.141 | 0.364 | 1.000 | 11143 | tags=18%, list=19%, signal=21% |
| 4679 | REGULATION\_OF\_UBIQUITIN\_DEPENDENT\_PROTEIN\_CATABOLIC\_PROCESS(GO:2000058) |  | 143 | 0.64 | 1.12 | 0.141 | 0.364 | 1.000 | 15098 | tags=36%, list=26%, signal=48% |
| 4680 | VISUAL\_LEARNING(GO:0008542) |  | 46 | 0.68 | 1.12 | 0.141 | 0.364 | 1.000 | 15752 | tags=46%, list=27%, signal=62% |
| 4681 | NEGATIVE\_REGULATION\_OF\_ALPHA\_BETA\_T\_CELL\_ACTIVATION(GO:0046636) |  | 27 | 0.69 | 1.12 | 0.141 | 0.364 | 1.000 | 14700 | tags=48%, list=25%, signal=64% |
| 4682 | REGULATION\_OF\_CALCIUM\_ION\_DEPENDENT\_EXOCYTOSIS(GO:0017158) |  | 66 | 0.66 | 1.12 | 0.141 | 0.364 | 1.000 | 9227 | tags=26%, list=16%, signal=30% |
| 4683 | MICROTUBULE(GO:0005874) |  | 403 | 0.65 | 1.12 | 0.141 | 0.364 | 1.000 | 13746 | tags=39%, list=23%, signal=51% |
| 4684 | REGULATION\_OF\_PEPTIDYL\_THREONINE\_PHOSPHORYLATION(GO:0010799) |  | 42 | 0.65 | 1.12 | 0.141 | 0.364 | 1.000 | 8823 | tags=14%, list=15%, signal=17% |
| 4685 | NEGATIVE\_REGULATION\_OF\_EPIDERMAL\_GROWTH\_FACTOR\_RECEPTOR\_SIGNALING\_PATHWAY(GO:0042059) |  | 44 | 0.64 | 1.12 | 0.141 | 0.364 | 1.000 | 17190 | tags=43%, list=29%, signal=61% |
| 4686 | AMIDE\_TRANSPORT(GO:0042886) |  | 1928 | 0.68 | 1.12 | 0.141 | 0.364 | 1.000 | 10876 | tags=17%, list=18%, signal=20% |
| 4687 | FATTY\_ACYL\_COA\_BINDING(GO:0000062) |  | 17 | 0.66 | 1.11 | 0.000 | 0.364 | 1.000 | 11715 | tags=35%, list=20%, signal=44% |
| 4688 | REGULATION\_OF\_MRNA\_PROCESSING(GO:0050684) |  | 110 | 0.64 | 1.11 | 0.000 | 0.364 | 1.000 | 18552 | tags=69%, list=31%, signal=100% |
| 4689 | REGULATION\_OF\_TORC1\_SIGNALING(GO:1903432) |  | 29 | 0.66 | 1.11 | 0.141 | 0.364 | 1.000 | 12070 | tags=34%, list=20%, signal=43% |
| 4690 | PEPTIDE\_TRANSPORT(GO:0015833) |  | 1907 | 0.68 | 1.11 | 0.141 | 0.364 | 1.000 | 11143 | tags=18%, list=19%, signal=21% |
| 4691 | CHROMOSOME\_SEGREGATION(GO:0007059) |  | 350 | 0.64 | 1.11 | 0.000 | 0.364 | 1.000 | 17317 | tags=63%, list=29%, signal=89% |
| 4692 | CILIUM\_ORGANIZATION(GO:0044782) |  | 362 | 0.68 | 1.11 | 0.141 | 0.364 | 1.000 | 13542 | tags=44%, list=23%, signal=57% |
| 4693 | NUCLEAR\_BODY(GO:0016604) |  | 744 | 0.68 | 1.11 | 0.141 | 0.364 | 1.000 | 13169 | tags=33%, list=22%, signal=41% |
| 4694 | INNATE\_IMMUNE\_RESPONSE(GO:0045087) |  | 943 | 0.66 | 1.11 | 0.141 | 0.364 | 1.000 | 12564 | tags=19%, list=21%, signal=23% |
| 4695 | RAS\_PROTEIN\_SIGNAL\_TRANSDUCTION(GO:0007265) |  | 354 | 0.67 | 1.11 | 0.141 | 0.364 | 1.000 | 14031 | tags=32%, list=24%, signal=42% |
| 4696 | CILIUM\_ASSEMBLY(GO:0060271) |  | 351 | 0.68 | 1.11 | 0.141 | 0.364 | 1.000 | 13542 | tags=45%, list=23%, signal=58% |
| 4697 | RNA\_DESTABILIZATION(GO:0050779) |  | 24 | 0.66 | 1.11 | 0.141 | 0.364 | 1.000 | 16375 | tags=63%, list=28%, signal=86% |
| 4698 | DNA\_POLYMERASE\_ACTIVITY(GO:0034061) |  | 35 | 0.63 | 1.11 | 0.000 | 0.364 | 1.000 | 20209 | tags=86%, list=34%, signal=130% |
| 4699 | AROMATASE\_ACTIVITY(GO:0070330) |  | 23 | 0.67 | 1.11 | 0.141 | 0.364 | 1.000 | 11149 | tags=26%, list=19%, signal=32% |
| 4700 | PHOSPHOTRANSFERASE\_ACTIVITY\_\_NITROGENOUS\_GROUP\_AS\_ACCEPTOR(GO:0016775) |  | 15 | 0.68 | 1.11 | 0.141 | 0.364 | 1.000 | 17118 | tags=60%, list=29%, signal=84% |
| 4701 | NUCLEAR\_INNER\_MEMBRANE(GO:0005637) |  | 57 | 0.64 | 1.11 | 0.000 | 0.364 | 1.000 | 16128 | tags=54%, list=27%, signal=75% |
| 4702 | SUMO\_TRANSFERASE\_ACTIVITY(GO:0019789) |  | 19 | 0.66 | 1.11 | 0.000 | 0.364 | 1.000 | 15941 | tags=47%, list=27%, signal=65% |
| 4703 | HEMATOPOIETIC\_PROGENITOR\_CELL\_DIFFERENTIATION(GO:0002244) |  | 197 | 0.65 | 1.11 | 0.141 | 0.364 | 1.000 | 15258 | tags=32%, list=26%, signal=43% |
| 4704 | CHONDROCYTE\_DEVELOPMENT(GO:0002063) |  | 26 | 0.67 | 1.11 | 0.141 | 0.364 | 1.000 | 11139 | tags=27%, list=19%, signal=33% |
| 4705 | POLYPEPTIDE\_N\_ACETYLGALACTOSAMINYLTRANSFERASE\_ACTIVITY(GO:0004653) |  | 19 | 0.70 | 1.11 | 0.141 | 0.364 | 1.000 | 15062 | tags=63%, list=25%, signal=85% |
| 4706 | POSTTRANSCRIPTIONAL\_REGULATION\_OF\_GENE\_EXPRESSION(GO:0010608) |  | 742 | 0.65 | 1.11 | 0.141 | 0.364 | 1.000 | 16211 | tags=33%, list=27%, signal=45% |
| 4707 | REGULATION\_OF\_PATHWAY\_RESTRICTED\_SMAD\_PROTEIN\_PHOSPHORYLATION(GO:0060393) |  | 61 | 0.67 | 1.11 | 0.141 | 0.364 | 1.000 | 13279 | tags=38%, list=22%, signal=49% |
| 4708 | ENDOSOME\_MEMBRANE(GO:0010008) |  | 447 | 0.67 | 1.11 | 0.141 | 0.364 | 1.000 | 10793 | tags=18%, list=18%, signal=22% |
| 4709 | REGULATION\_OF\_MRNA\_STABILITY(GO:0043488) |  | 160 | 0.65 | 1.11 | 0.141 | 0.364 | 1.000 | 16931 | tags=47%, list=29%, signal=66% |
| 4710 | ERROR\_PRONE\_TRANSLESION\_SYNTHESIS(GO:0042276) |  | 20 | 0.64 | 1.11 | 0.000 | 0.364 | 1.000 | 19834 | tags=90%, list=34%, signal=135% |
| 4711 | RNA\_POLYMERASE\_II\_CORE\_BINDING(GO:0000993) |  | 43 | 0.67 | 1.11 | 0.000 | 0.365 | 1.000 | 10656 | tags=16%, list=18%, signal=20% |
| 4712 | INTRACILIARY\_TRANSPORT(GO:0042073) |  | 50 | 0.64 | 1.11 | 0.000 | 0.364 | 1.000 | 15522 | tags=60%, list=26%, signal=81% |
| 4713 | REGULATION\_OF\_RESPONSE\_TO\_REACTIVE\_OXYGEN\_SPECIES(GO:1901031) |  | 45 | 0.67 | 1.11 | 0.141 | 0.365 | 1.000 | 14492 | tags=29%, list=25%, signal=38% |
| 4714 | POSITIVE\_REGULATION\_OF\_HEART\_CONTRACTION(GO:0045823) |  | 39 | 0.68 | 1.11 | 0.141 | 0.365 | 1.000 | 11254 | tags=36%, list=19%, signal=44% |
| 4715 | PROTEIN\_LOCALIZATION\_TO\_ENDOPLASMIC\_RETICULUM(GO:0070972) |  | 126 | 0.63 | 1.11 | 0.141 | 0.365 | 1.000 | 17054 | tags=44%, list=29%, signal=61% |
| 4716 | POSITIVE\_REGULATION\_OF\_T\_CELL\_APOPTOTIC\_PROCESS(GO:0070234) |  | 15 | 0.68 | 1.11 | 0.141 | 0.365 | 1.000 | 13000 | tags=20%, list=22%, signal=26% |
| 4717 | CAMERA\_TYPE\_EYE\_PHOTORECEPTOR\_CELL\_DIFFERENTIATION(GO:0060219) |  | 18 | 0.72 | 1.11 | 0.141 | 0.365 | 1.000 | 10712 | tags=39%, list=18%, signal=47% |
| 4718 | BITTER\_TASTE\_RECEPTOR\_ACTIVITY(GO:0033038) |  | 20 | 0.66 | 1.11 | 0.168 | 0.365 | 1.000 | 9774 | tags=15%, list=17%, signal=18% |
| 4719 | OXIDOREDUCTASE\_ACTIVITY\_\_ACTING\_ON\_THE\_CH\_NH\_GROUP\_OF\_DONORS(GO:0016645) |  | 28 | 0.69 | 1.11 | 0.141 | 0.365 | 1.000 | 14864 | tags=46%, list=25%, signal=62% |
| 4720 | ACUTE\_PHASE\_RESPONSE(GO:0006953) |  | 49 | 0.68 | 1.11 | 0.141 | 0.365 | 1.000 | 14996 | tags=29%, list=25%, signal=38% |
| 4721 | REGULATION\_OF\_RNA\_STABILITY(GO:0043487) |  | 168 | 0.65 | 1.11 | 0.141 | 0.365 | 1.000 | 17026 | tags=48%, list=29%, signal=67% |
| 4722 | PROTEIN\_MONOUBIQUITINATION(GO:0006513) |  | 64 | 0.63 | 1.11 | 0.000 | 0.365 | 1.000 | 16251 | tags=50%, list=27%, signal=69% |
| 4723 | IONOTROPIC\_GLUTAMATE\_RECEPTOR\_COMPLEX(GO:0008328) |  | 47 | 0.69 | 1.11 | 0.141 | 0.365 | 1.000 | 10305 | tags=34%, list=17%, signal=41% |
| 4724 | NEUROTRANSMITTER\_BINDING(GO:0042165) |  | 52 | 0.66 | 1.11 | 0.141 | 0.365 | 1.000 | 14855 | tags=52%, list=25%, signal=69% |
| 4725 | NEGATIVE\_REGULATION\_OF\_BLOOD\_CIRCULATION(GO:1903523) |  | 34 | 0.68 | 1.11 | 0.141 | 0.364 | 1.000 | 12699 | tags=35%, list=21%, signal=45% |
| 4726 | NEGATIVE\_REGULATION\_OF\_RELEASE\_OF\_SEQUESTERED\_CALCIUM\_ION\_INTO\_CYTOSOL(GO:0051280) |  | 15 | 0.68 | 1.11 | 0.141 | 0.364 | 1.000 | 10116 | tags=33%, list=17%, signal=40% |
| 4727 | POSITIVE\_REGULATION\_OF\_CELLULAR\_RESPONSE\_TO\_OXIDATIVE\_STRESS(GO:1900409) |  | 19 | 0.67 | 1.11 | 0.141 | 0.364 | 1.000 | 6514 | tags=11%, list=11%, signal=12% |
| 4728 | NUCLEOLAR\_PART(GO:0044452) |  | 182 | 0.65 | 1.11 | 0.141 | 0.364 | 1.000 | 14820 | tags=41%, list=25%, signal=54% |
| 4729 | OSTEOCLAST\_DEVELOPMENT(GO:0036035) |  | 17 | 0.72 | 1.11 | 0.141 | 0.364 | 1.000 | 13386 | tags=35%, list=23%, signal=46% |
| 4730 | MITOTIC\_SISTER\_CHROMATID\_SEGREGATION(GO:0000070) |  | 146 | 0.63 | 1.11 | 0.000 | 0.364 | 1.000 | 19621 | tags=78%, list=33%, signal=117% |
| 4731 | CELL\_CYCLE\_DNA\_REPLICATION(GO:0044786) |  | 59 | 0.64 | 1.11 | 0.000 | 0.364 | 1.000 | 17484 | tags=75%, list=30%, signal=106% |
| 4732 | ASSOCIATIVE\_LEARNING(GO:0008306) |  | 72 | 0.66 | 1.11 | 0.141 | 0.364 | 1.000 | 15752 | tags=43%, list=27%, signal=59% |
| 4733 | INTEGRAL\_COMPONENT\_OF\_MITOCHONDRIAL\_MEMBRANE(GO:0032592) |  | 72 | 0.64 | 1.11 | 0.000 | 0.364 | 1.000 | 18095 | tags=60%, list=31%, signal=86% |
| 4734 | SYNAPTIC\_TRANSMISSION\_\_GLUTAMATERGIC(GO:0035249) |  | 79 | 0.68 | 1.11 | 0.141 | 0.364 | 1.000 | 8759 | tags=25%, list=15%, signal=30% |
| 4735 | RUFFLE(GO:0001726) |  | 163 | 0.67 | 1.11 | 0.141 | 0.364 | 1.000 | 13195 | tags=22%, list=22%, signal=28% |
| 4736 | EPITHELIAL\_CILIUM\_MOVEMENT(GO:0003351) |  | 22 | 0.64 | 1.11 | 0.000 | 0.364 | 1.000 | 13286 | tags=41%, list=22%, signal=53% |
| 4737 | ESTABLISHMENT\_OF\_PROTEIN\_LOCALIZATION\_TO\_MITOCHONDRION(GO:0072655) |  | 189 | 0.67 | 1.11 | 0.141 | 0.364 | 1.000 | 12293 | tags=22%, list=21%, signal=27% |
| 4738 | AXONAL\_GROWTH\_CONE(GO:0044295) |  | 24 | 0.69 | 1.11 | 0.141 | 0.364 | 1.000 | 11853 | tags=33%, list=20%, signal=42% |
| 4739 | PROTEIN\_LOCALIZATION\_TO\_CHROMOSOME\_\_CENTROMERIC\_REGION(GO:0071459) |  | 22 | 0.66 | 1.11 | 0.000 | 0.364 | 1.000 | 15072 | tags=73%, list=25%, signal=98% |
| 4740 | PLACENTA\_DEVELOPMENT(GO:0001890) |  | 152 | 0.68 | 1.11 | 0.141 | 0.364 | 1.000 | 13707 | tags=32%, list=23%, signal=41% |
| 4741 | SISTER\_CHROMATID\_SEGREGATION(GO:0000819) |  | 234 | 0.63 | 1.11 | 0.000 | 0.364 | 1.000 | 19843 | tags=81%, list=34%, signal=122% |
| 4742 | PROTEIN\_COMPLEX\_OLIGOMERIZATION(GO:0051259) |  | 533 | 0.67 | 1.11 | 0.141 | 0.364 | 1.000 | 13169 | tags=30%, list=22%, signal=39% |
| 4743 | NEGATIVE\_REGULATION\_OF\_ADAPTIVE\_IMMUNE\_RESPONSE\_BASED\_ON\_SOMATIC\_RECOMBINATION\_OF\_IMMUNE\_RECEPTORS\_BUILT\_FROM\_IMMUNOGLOBULIN\_SUPERFAMILY\_DOMAINS(GO:0002823) |  | 33 | 0.68 | 1.11 | 0.141 | 0.364 | 1.000 | 15529 | tags=36%, list=26%, signal=49% |
| 4744 | TRNA\_TRANSPORT(GO:0051031) |  | 34 | 0.64 | 1.11 | 0.000 | 0.364 | 1.000 | 18197 | tags=82%, list=31%, signal=119% |
| 4745 | NUCLEAR\_PART(GO:0044428) |  | 4430 | 0.69 | 1.11 | 0.141 | 0.364 | 1.000 | 13552 | tags=31%, list=23%, signal=37% |
| 4746 | VENTRICULAR\_SEPTUM\_DEVELOPMENT(GO:0003281) |  | 66 | 0.68 | 1.11 | 0.141 | 0.364 | 1.000 | 12990 | tags=36%, list=22%, signal=47% |
| 4747 | MITOTIC\_CELL\_CYCLE\_PHASE\_TRANSITION(GO:0044772) |  | 555 | 0.66 | 1.11 | 0.141 | 0.364 | 1.000 | 13522 | tags=34%, list=23%, signal=44% |
| 4748 | VENTRICULAR\_SYSTEM\_DEVELOPMENT(GO:0021591) |  | 26 | 0.65 | 1.11 | 0.000 | 0.364 | 1.000 | 17987 | tags=65%, list=30%, signal=94% |
| 4749 | HISTONE\_H4\_ACETYLATION(GO:0043967) |  | 62 | 0.64 | 1.11 | 0.000 | 0.364 | 1.000 | 19832 | tags=76%, list=34%, signal=114% |
| 4750 | AMPA\_GLUTAMATE\_RECEPTOR\_COMPLEX(GO:0032281) |  | 28 | 0.70 | 1.11 | 0.141 | 0.364 | 1.000 | 9440 | tags=32%, list=16%, signal=38% |
| 4751 | NUCLEAR\_REPLISOME(GO:0043601) |  | 26 | 0.63 | 1.11 | 0.000 | 0.364 | 1.000 | 17064 | tags=73%, list=29%, signal=103% |
| 4752 | GOLGI\_LUMEN(GO:0005796) |  | 100 | 0.69 | 1.11 | 0.141 | 0.364 | 1.000 | 11459 | tags=24%, list=19%, signal=30% |
| 4753 | REGULATION\_OF\_CELL\_KILLING(GO:0031341) |  | 67 | 0.64 | 1.11 | 0.141 | 0.364 | 1.000 | 14247 | tags=42%, list=24%, signal=55% |
| 4754 | REGULATION\_OF\_CELL\_CYCLE\_PHASE\_TRANSITION(GO:1901987) |  | 446 | 0.67 | 1.11 | 0.141 | 0.364 | 1.000 | 13110 | tags=30%, list=22%, signal=38% |
| 4755 | INTEGRAL\_COMPONENT\_OF\_GOLGI\_MEMBRANE(GO:0030173) |  | 46 | 0.64 | 1.11 | 0.141 | 0.364 | 1.000 | 18337 | tags=57%, list=31%, signal=82% |
| 4756 | MRNA\_BINDING(GO:0003729) |  | 377 | 0.67 | 1.11 | 0.141 | 0.364 | 1.000 | 15159 | tags=29%, list=26%, signal=38% |
| 4757 | CELLULAR\_RESPONSE\_TO\_LIGHT\_STIMULUS(GO:0071482) |  | 114 | 0.63 | 1.11 | 0.000 | 0.364 | 1.000 | 17568 | tags=60%, list=30%, signal=85% |
| 4758 | REGULATION\_OF\_CHROMATIN\_BINDING(GO:0035561) |  | 22 | 0.64 | 1.11 | 0.141 | 0.364 | 1.000 | 16227 | tags=50%, list=27%, signal=69% |
| 4759 | NEGATIVE\_REGULATION\_OF\_PROTEIN\_MODIFICATION\_BY\_SMALL\_PROTEIN\_CONJUGATION\_OR\_REMOVAL(GO:1903321) |  | 77 | 0.64 | 1.11 | 0.141 | 0.364 | 1.000 | 14185 | tags=32%, list=24%, signal=43% |
| 4760 | TERMINATION\_OF\_RNA\_POLYMERASE\_I\_TRANSCRIPTION(GO:0006363) |  | 30 | 0.63 | 1.11 | 0.000 | 0.364 | 1.000 | 20344 | tags=80%, list=34%, signal=122% |
| 4761 | RNA\_POLYMERASE\_BINDING(GO:0070063) |  | 66 | 0.65 | 1.11 | 0.000 | 0.364 | 1.000 | 15530 | tags=41%, list=26%, signal=55% |
| 4762 | REGULATION\_OF\_INTRACELLULAR\_ESTROGEN\_RECEPTOR\_SIGNALING\_PATHWAY(GO:0033146) |  | 40 | 0.71 | 1.11 | 0.141 | 0.364 | 1.000 | 8112 | tags=18%, list=14%, signal=20% |
| 4763 | INTERLEUKIN\_5\_PRODUCTION(GO:0032634) |  | 22 | 0.70 | 1.11 | 0.141 | 0.364 | 1.000 | 14363 | tags=36%, list=24%, signal=48% |
| 4764 | NEUROTRANSMITTER\_RECEPTOR\_COMPLEX(GO:0098878) |  | 49 | 0.69 | 1.11 | 0.141 | 0.364 | 1.000 | 10305 | tags=37%, list=17%, signal=44% |
| 4765 | REGULATION\_OF\_DENDRITE\_DEVELOPMENT(GO:0050773) |  | 130 | 0.68 | 1.11 | 0.141 | 0.364 | 1.000 | 9607 | tags=22%, list=16%, signal=26% |
| 4766 | PROTEIN\_LOCALIZATION\_TO\_MITOCHONDRION(GO:0070585) |  | 191 | 0.67 | 1.11 | 0.141 | 0.364 | 1.000 | 12293 | tags=21%, list=21%, signal=27% |
| 4767 | ACTIVATION\_OF\_MAPK\_ACTIVITY(GO:0000187) |  | 143 | 0.66 | 1.11 | 0.141 | 0.364 | 1.000 | 10811 | tags=24%, list=18%, signal=30% |
| 4768 | NEGATIVE\_REGULATION\_OF\_T\_CELL\_PROLIFERATION(GO:0042130) |  | 54 | 0.66 | 1.11 | 0.141 | 0.364 | 1.000 | 15466 | tags=37%, list=26%, signal=50% |
| 4769 | REGULATION\_OF\_INTERLEUKIN\_5\_PRODUCTION(GO:0032674) |  | 21 | 0.70 | 1.11 | 0.141 | 0.364 | 1.000 | 14363 | tags=38%, list=24%, signal=50% |
| 4770 | TUMOR\_NECROSIS\_FACTOR\_BIOSYNTHETIC\_PROCESS(GO:0042533) |  | 18 | 0.70 | 1.11 | 0.141 | 0.364 | 1.000 | 7461 | tags=11%, list=13%, signal=13% |
| 4771 | REGULATION\_OF\_TUMOR\_NECROSIS\_FACTOR\_BIOSYNTHETIC\_PROCESS(GO:0042534) |  | 18 | 0.70 | 1.11 | 0.141 | 0.364 | 1.000 | 7461 | tags=11%, list=13%, signal=13% |
| 4772 | OXIDATIVE\_PHOSPHORYLATION(GO:0006119) |  | 136 | 0.62 | 1.11 | 0.141 | 0.364 | 1.000 | 19377 | tags=59%, list=33%, signal=87% |
| 4773 | POSITIVE\_REGULATION\_OF\_BMP\_SIGNALING\_PATHWAY(GO:0030513) |  | 34 | 0.70 | 1.11 | 0.141 | 0.364 | 1.000 | 12258 | tags=29%, list=21%, signal=37% |
| 4774 | POSITIVE\_REGULATION\_OF\_MRNA\_SPLICING\_\_VIA\_SPLICEOSOME(GO:0048026) |  | 17 | 0.65 | 1.11 | 0.000 | 0.364 | 1.000 | 18535 | tags=76%, list=31%, signal=111% |
| 4775 | NCRNA\_PROCESSING(GO:0034470) |  | 407 | 0.62 | 1.11 | 0.000 | 0.364 | 1.000 | 19172 | tags=67%, list=32%, signal=98% |
| 4776 | ESTABLISHMENT\_OF\_PROTEIN\_LOCALIZATION(GO:0045184) |  | 1969 | 0.67 | 1.11 | 0.141 | 0.364 | 1.000 | 12700 | tags=23%, list=21%, signal=29% |
| 4777 | AUTOPHAGOSOME(GO:0005776) |  | 89 | 0.62 | 1.11 | 0.141 | 0.364 | 1.000 | 11317 | tags=21%, list=19%, signal=26% |
| 4778 | MEIOTIC\_CELL\_CYCLE\_PROCESS(GO:1903046) |  | 186 | 0.66 | 1.11 | 0.141 | 0.364 | 1.000 | 14321 | tags=44%, list=24%, signal=58% |
| 4779 | INTESTINAL\_EPITHELIAL\_CELL\_DIFFERENTIATION(GO:0060575) |  | 20 | 0.70 | 1.11 | 0.141 | 0.364 | 1.000 | 12444 | tags=40%, list=21%, signal=51% |
| 4780 | POSITIVE\_REGULATION\_OF\_RNA\_SPLICING(GO:0033120) |  | 31 | 0.66 | 1.11 | 0.000 | 0.364 | 1.000 | 13647 | tags=39%, list=23%, signal=50% |
| 4781 | PROTEIN\_MODIFICATION\_BY\_SMALL\_PROTEIN\_CONJUGATION\_OR\_REMOVAL(GO:0070647) |  | 1123 | 0.66 | 1.11 | 0.141 | 0.364 | 1.000 | 15185 | tags=36%, list=26%, signal=47% |
| 4782 | PROTEIN\_KINASE\_A\_REGULATORY\_SUBUNIT\_BINDING(GO:0034237) |  | 19 | 0.65 | 1.11 | 0.000 | 0.364 | 1.000 | 13269 | tags=42%, list=22%, signal=54% |
| 4783 | CELLULAR\_RESPONSE\_TO\_DEXAMETHASONE\_STIMULUS(GO:0071549) |  | 30 | 0.66 | 1.11 | 0.141 | 0.364 | 1.000 | 10131 | tags=23%, list=17%, signal=28% |
| 4784 | DNA\_REPLICATION(GO:0006260) |  | 295 | 0.65 | 1.11 | 0.141 | 0.364 | 1.000 | 16570 | tags=57%, list=28%, signal=79% |
| 4785 | NUCLEAR\_LOCALIZATION\_SEQUENCE\_BINDING(GO:0008139) |  | 27 | 0.65 | 1.11 | 0.000 | 0.364 | 1.000 | 17378 | tags=67%, list=29%, signal=94% |
| 4786 | POSITIVE\_REGULATION\_OF\_DENDRITE\_DEVELOPMENT(GO:1900006) |  | 66 | 0.63 | 1.11 | 0.141 | 0.364 | 1.000 | 16153 | tags=42%, list=27%, signal=58% |
| 4787 | ANTERIOR\_POSTERIOR\_AXIS\_SPECIFICATION\_\_EMBRYO(GO:0008595) |  | 15 | 0.71 | 1.11 | 0.141 | 0.364 | 1.000 | 11411 | tags=40%, list=19%, signal=50% |
| 4788 | TRIPARTITE\_REGIONAL\_SUBDIVISION(GO:0007351) |  | 15 | 0.71 | 1.11 | 0.141 | 0.364 | 1.000 | 11411 | tags=40%, list=19%, signal=50% |
| 4789 | PROTEIN\_LOCALIZATION\_TO\_CHROMOSOME\_\_TELOMERIC\_REGION(GO:0070198) |  | 27 | 0.64 | 1.11 | 0.000 | 0.364 | 1.000 | 20841 | tags=96%, list=35%, signal=149% |
| 4790 | REGULATION\_OF\_APPETITE(GO:0032098) |  | 22 | 0.65 | 1.11 | 0.169 | 0.364 | 1.000 | 18462 | tags=59%, list=31%, signal=86% |
| 4791 | LATE\_ENDOSOME(GO:0005770) |  | 235 | 0.65 | 1.11 | 0.141 | 0.364 | 1.000 | 13564 | tags=29%, list=23%, signal=38% |
| 4792 | PROTEIN\_AUTOUBIQUITINATION(GO:0051865) |  | 59 | 0.64 | 1.11 | 0.141 | 0.364 | 1.000 | 16700 | tags=53%, list=28%, signal=73% |
| 4793 | SISTER\_CHROMATID\_COHESION(GO:0007062) |  | 134 | 0.63 | 1.11 | 0.000 | 0.364 | 1.000 | 19843 | tags=87%, list=34%, signal=130% |
| 4794 | COFACTOR\_TRANSMEMBRANE\_TRANSPORTER\_ACTIVITY(GO:0051184) |  | 20 | 0.65 | 1.11 | 0.141 | 0.364 | 1.000 | 15294 | tags=55%, list=26%, signal=74% |
| 4795 | RESPONSE\_TO\_AMPHETAMINE(GO:0001975) |  | 26 | 0.70 | 1.11 | 0.141 | 0.364 | 1.000 | 12828 | tags=42%, list=22%, signal=54% |
| 4796 | G\_PROTEIN\_COUPLED\_AMINE\_RECEPTOR\_ACTIVITY(GO:0008227) |  | 48 | 0.67 | 1.11 | 0.141 | 0.364 | 1.000 | 9156 | tags=21%, list=15%, signal=25% |
| 4797 | MITOTIC\_DNA\_DAMAGE\_CHECKPOINT(GO:0044773) |  | 107 | 0.68 | 1.11 | 0.141 | 0.364 | 1.000 | 13110 | tags=31%, list=22%, signal=40% |
| 4798 | NEGATIVE\_REGULATION\_OF\_PROTEIN\_DEPOLYMERIZATION(GO:1901880) |  | 58 | 0.64 | 1.11 | 0.141 | 0.364 | 1.000 | 17265 | tags=48%, list=29%, signal=68% |
| 4799 | ISOMERASE\_ACTIVITY(GO:0016853) |  | 155 | 0.62 | 1.11 | 0.141 | 0.364 | 1.000 | 18912 | tags=59%, list=32%, signal=86% |
| 4800 | AXON\_ENSHEATHMENT\_IN\_CENTRAL\_NERVOUS\_SYSTEM(GO:0032291) |  | 17 | 0.69 | 1.11 | 0.141 | 0.364 | 1.000 | 13179 | tags=41%, list=22%, signal=53% |
| 4801 | CENTRAL\_NERVOUS\_SYSTEM\_MYELINATION(GO:0022010) |  | 17 | 0.69 | 1.11 | 0.141 | 0.364 | 1.000 | 13179 | tags=41%, list=22%, signal=53% |
| 4802 | TETRAHYDROFOLATE\_METABOLIC\_PROCESS(GO:0046653) |  | 19 | 0.63 | 1.11 | 0.000 | 0.364 | 1.000 | 14192 | tags=42%, list=24%, signal=55% |
| 4803 | TUMOR\_NECROSIS\_FACTOR\_RECEPTOR\_SUPERFAMILY\_BINDING(GO:0032813) |  | 46 | 0.66 | 1.11 | 0.141 | 0.364 | 1.000 | 11164 | tags=26%, list=19%, signal=32% |
| 4804 | CENTROSOME\_CYCLE(GO:0007098) |  | 121 | 0.62 | 1.11 | 0.000 | 0.364 | 1.000 | 21117 | tags=84%, list=36%, signal=131% |
| 4805 | RETINOID\_BINDING(GO:0005501) |  | 34 | 0.71 | 1.11 | 0.141 | 0.364 | 1.000 | 11950 | tags=32%, list=20%, signal=41% |
| 4806 | DNA\_DEPENDENT\_DNA\_REPLICATION\_MAINTENANCE\_OF\_FIDELITY(GO:0045005) |  | 35 | 0.62 | 1.11 | 0.000 | 0.364 | 1.000 | 17539 | tags=69%, list=30%, signal=97% |
| 4807 | INTRINSIC\_COMPONENT\_OF\_MITOCHONDRIAL\_MEMBRANE(GO:0098573) |  | 73 | 0.63 | 1.11 | 0.000 | 0.364 | 1.000 | 19384 | tags=67%, list=33%, signal=100% |
| 4808 | STRIATED\_MUSCLE\_THIN\_FILAMENT(GO:0005865) |  | 21 | 0.68 | 1.11 | 0.141 | 0.364 | 1.000 | 13148 | tags=48%, list=22%, signal=61% |
| 4809 | NEGATIVE\_REGULATION\_OF\_CALCIUM\_ION\_TRANSPORT(GO:0051926) |  | 65 | 0.66 | 1.11 | 0.141 | 0.364 | 1.000 | 10556 | tags=26%, list=18%, signal=32% |
| 4810 | ADAPTATION\_OF\_SIGNALING\_PATHWAY(GO:0023058) |  | 22 | 0.66 | 1.11 | 0.000 | 0.364 | 1.000 | 14399 | tags=55%, list=24%, signal=72% |
| 4811 | NEGATIVE\_REGULATION\_OF\_BLOOD\_PRESSURE(GO:0045776) |  | 45 | 0.69 | 1.11 | 0.141 | 0.364 | 1.000 | 10245 | tags=24%, list=17%, signal=30% |
| 4812 | NUCLEAR\_TRANSCRIBED\_MRNA\_CATABOLIC\_PROCESS(GO:0000956) |  | 215 | 0.64 | 1.11 | 0.141 | 0.364 | 1.000 | 17054 | tags=46%, list=29%, signal=64% |
| 4813 | NUCLEAR\_DNA\_REPLICATION(GO:0033260) |  | 50 | 0.63 | 1.11 | 0.000 | 0.364 | 1.000 | 17484 | tags=72%, list=30%, signal=102% |
| 4814 | MYELOID\_LEUKOCYTE\_CYTOKINE\_PRODUCTION(GO:0061082) |  | 22 | 0.66 | 1.11 | 0.141 | 0.364 | 1.000 | 14181 | tags=36%, list=24%, signal=48% |
| 4815 | POSITIVE\_REGULATION\_OF\_AXONOGENESIS(GO:0050772) |  | 77 | 0.68 | 1.11 | 0.141 | 0.364 | 1.000 | 9919 | tags=26%, list=17%, signal=31% |
| 4816 | CELLULAR\_RESPONSE\_TO\_REACTIVE\_NITROGEN\_SPECIES(GO:1902170) |  | 19 | 0.66 | 1.11 | 0.000 | 0.364 | 1.000 | 15560 | tags=47%, list=26%, signal=64% |
| 4817 | DNA\_DAMAGE\_RESPONSE\_\_SIGNAL\_TRANSDUCTION\_BY\_P53\_CLASS\_MEDIATOR(GO:0030330) |  | 110 | 0.67 | 1.11 | 0.141 | 0.364 | 1.000 | 8177 | tags=9%, list=14%, signal=11% |
| 4818 | NEGATIVE\_REGULATION\_OF\_VOLTAGE\_GATED\_POTASSIUM\_CHANNEL\_ACTIVITY(GO:1903817) |  | 17 | 0.68 | 1.11 | 0.172 | 0.363 | 1.000 | 14718 | tags=41%, list=25%, signal=55% |
| 4819 | MITOTIC\_DNA\_INTEGRITY\_CHECKPOINT(GO:0044774) |  | 113 | 0.68 | 1.11 | 0.141 | 0.363 | 1.000 | 13110 | tags=32%, list=22%, signal=41% |
| 4820 | NEGATIVE\_REGULATION\_OF\_ANION\_TRANSPORT(GO:1903792) |  | 24 | 0.69 | 1.11 | 0.141 | 0.363 | 1.000 | 11264 | tags=29%, list=19%, signal=36% |
| 4821 | GROWTH\_FACTOR\_ACTIVITY(GO:0008083) |  | 161 | 0.68 | 1.11 | 0.141 | 0.363 | 1.000 | 11459 | tags=32%, list=19%, signal=39% |
| 4822 | TRANSCRIPTION\_FACTOR\_TFIID\_COMPLEX(GO:0005669) |  | 37 | 0.63 | 1.11 | 0.000 | 0.364 | 1.000 | 19664 | tags=78%, list=33%, signal=117% |
| 4823 | POSITIVE\_REGULATION\_OF\_TELOMERE\_MAINTENANCE\_VIA\_TELOMERASE(GO:0032212) |  | 34 | 0.64 | 1.11 | 0.000 | 0.363 | 1.000 | 16850 | tags=76%, list=28%, signal=107% |
| 4824 | REGULATION\_OF\_CYTOKINE\_PRODUCTION\_INVOLVED\_IN\_INFLAMMATORY\_RESPONSE(GO:1900015) |  | 34 | 0.66 | 1.10 | 0.141 | 0.364 | 1.000 | 6413 | tags=18%, list=11%, signal=20% |
| 4825 | REGULATION\_OF\_IMMUNE\_EFFECTOR\_PROCESS(GO:0002697) |  | 398 | 0.64 | 1.10 | 0.141 | 0.364 | 1.000 | 15529 | tags=29%, list=26%, signal=40% |
| 4826 | POLYSACCHARIDE\_BIOSYNTHETIC\_PROCESS(GO:0000271) |  | 72 | 0.67 | 1.10 | 0.141 | 0.364 | 1.000 | 9277 | tags=18%, list=16%, signal=21% |
| 4827 | PROTEIN\_REFOLDING(GO:0042026) |  | 19 | 0.66 | 1.10 | 0.141 | 0.364 | 1.000 | 18589 | tags=68%, list=31%, signal=100% |
| 4828 | DNA\_STRAND\_ELONGATION(GO:0022616) |  | 25 | 0.63 | 1.10 | 0.000 | 0.364 | 1.000 | 21373 | tags=96%, list=36%, signal=150% |
| 4829 | COPII\_COATED\_ER\_TO\_GOLGI\_TRANSPORT\_VESICLE(GO:0030134) |  | 76 | 0.62 | 1.10 | 0.141 | 0.363 | 1.000 | 19349 | tags=58%, list=33%, signal=86% |
| 4830 | MICROTUBULE\_ORGANIZING\_CENTER\_ORGANIZATION(GO:0031023) |  | 130 | 0.62 | 1.10 | 0.000 | 0.363 | 1.000 | 21117 | tags=83%, list=36%, signal=129% |
| 4831 | NEGATIVE\_REGULATION\_OF\_CELLULAR\_PROTEIN\_CATABOLIC\_PROCESS(GO:1903363) |  | 72 | 0.63 | 1.10 | 0.141 | 0.363 | 1.000 | 18043 | tags=54%, list=31%, signal=78% |
| 4832 | ATPASE\_ACTIVITY\_\_COUPLED\_TO\_TRANSMEMBRANE\_MOVEMENT\_OF\_IONS\_\_PHOSPHORYLATIVE\_MECHANISM(GO:0015662) |  | 37 | 0.65 | 1.10 | 0.141 | 0.363 | 1.000 | 16848 | tags=54%, list=28%, signal=76% |
| 4833 | ACTIVATION\_OF\_MAPKK\_ACTIVITY(GO:0000186) |  | 54 | 0.64 | 1.10 | 0.141 | 0.363 | 1.000 | 7702 | tags=19%, list=13%, signal=21% |
| 4834 | TRANSCRIPTION\_ELONGATION\_FROM\_RNA\_POLYMERASE\_I\_PROMOTER(GO:0006362) |  | 30 | 0.63 | 1.10 | 0.000 | 0.363 | 1.000 | 20344 | tags=80%, list=34%, signal=122% |
| 4835 | NEGATIVE\_REGULATION\_OF\_EPITHELIAL\_CELL\_MIGRATION(GO:0010633) |  | 108 | 0.68 | 1.10 | 0.141 | 0.363 | 1.000 | 14175 | tags=29%, list=24%, signal=38% |
| 4836 | ENVELOPE(GO:0031975) |  | 1182 | 0.65 | 1.10 | 0.141 | 0.363 | 1.000 | 13695 | tags=29%, list=23%, signal=36% |
| 4837 | ORGANELLE\_ENVELOPE(GO:0031967) |  | 1182 | 0.65 | 1.10 | 0.141 | 0.363 | 1.000 | 13695 | tags=29%, list=23%, signal=36% |
| 4838 | BRANCHED\_CHAIN\_AMINO\_ACID\_CATABOLIC\_PROCESS(GO:0009083) |  | 21 | 0.66 | 1.10 | 0.000 | 0.363 | 1.000 | 17081 | tags=57%, list=29%, signal=80% |
| 4839 | POSITIVE\_REGULATION\_OF\_TELOMERE\_MAINTENANCE\_VIA\_TELOMERE\_LENGTHENING(GO:1904358) |  | 36 | 0.64 | 1.10 | 0.000 | 0.363 | 1.000 | 16850 | tags=75%, list=28%, signal=105% |
| 4840 | EXCITATORY\_EXTRACELLULAR\_LIGAND\_GATED\_ION\_CHANNEL\_ACTIVITY(GO:0005231) |  | 50 | 0.67 | 1.10 | 0.141 | 0.363 | 1.000 | 13742 | tags=50%, list=23%, signal=65% |
| 4841 | NUCLEIC\_ACID\_PHOSPHODIESTER\_BOND\_HYDROLYSIS(GO:0090305) |  | 290 | 0.62 | 1.10 | 0.000 | 0.363 | 1.000 | 18087 | tags=62%, list=31%, signal=89% |
| 4842 | CHEMOATTRACTANT\_ACTIVITY(GO:0042056) |  | 31 | 0.70 | 1.10 | 0.141 | 0.363 | 1.000 | 14434 | tags=35%, list=24%, signal=47% |
| 4843 | MYELOID\_CELL\_HOMEOSTASIS(GO:0002262) |  | 142 | 0.65 | 1.10 | 0.141 | 0.363 | 1.000 | 10998 | tags=25%, list=19%, signal=30% |
| 4844 | FIBRONECTIN\_BINDING(GO:0001968) |  | 25 | 0.68 | 1.10 | 0.141 | 0.363 | 1.000 | 8013 | tags=20%, list=14%, signal=23% |
| 4845 | SENSORY\_PERCEPTION\_OF\_BITTER\_TASTE(GO:0050913) |  | 41 | 0.65 | 1.10 | 0.141 | 0.363 | 1.000 | 9774 | tags=12%, list=17%, signal=15% |
| 4846 | CHROMOSOMAL\_REGION(GO:0098687) |  | 345 | 0.62 | 1.10 | 0.000 | 0.363 | 1.000 | 19369 | tags=70%, list=33%, signal=103% |
| 4847 | NUCLEOTIDE\_EXCISION\_REPAIR\_\_DNA\_INCISION\_\_5'\_TO\_LESION(GO:0006296) |  | 37 | 0.63 | 1.10 | 0.000 | 0.363 | 1.000 | 19834 | tags=81%, list=34%, signal=122% |
| 4848 | MYOSIN\_FILAMENT\_ORGANIZATION(GO:0031033) |  | 15 | 0.68 | 1.10 | 0.166 | 0.363 | 1.000 | 7652 | tags=40%, list=13%, signal=46% |
| 4849 | POSITIVE\_REGULATION\_OF\_NIK\_NF\_KAPPAB\_SIGNALING(GO:1901224) |  | 44 | 0.62 | 1.10 | 0.141 | 0.363 | 1.000 | 19641 | tags=66%, list=33%, signal=99% |
| 4850 | NEGATIVE\_REGULATION\_OF\_LYMPHOCYTE\_PROLIFERATION(GO:0050672) |  | 68 | 0.66 | 1.10 | 0.141 | 0.363 | 1.000 | 15466 | tags=35%, list=26%, signal=48% |
| 4851 | NEGATIVE\_REGULATION\_OF\_MONONUCLEAR\_CELL\_PROLIFERATION(GO:0032945) |  | 68 | 0.66 | 1.10 | 0.141 | 0.363 | 1.000 | 15466 | tags=35%, list=26%, signal=48% |
| 4852 | CELLULAR\_METABOLIC\_COMPOUND\_SALVAGE(GO:0043094) |  | 36 | 0.63 | 1.10 | 0.141 | 0.363 | 1.000 | 14931 | tags=36%, list=25%, signal=48% |
| 4853 | REGULATION\_OF\_EMBRYONIC\_DEVELOPMENT(GO:0045995) |  | 125 | 0.67 | 1.10 | 0.141 | 0.363 | 1.000 | 10839 | tags=26%, list=18%, signal=32% |
| 4854 | NEGATIVE\_REGULATION\_OF\_PROTEOLYSIS\_INVOLVED\_IN\_CELLULAR\_PROTEIN\_CATABOLIC\_PROCESS(GO:1903051) |  | 61 | 0.65 | 1.10 | 0.141 | 0.363 | 1.000 | 18043 | tags=57%, list=31%, signal=82% |
| 4855 | DETECTION\_OF\_TEMPERATURE\_STIMULUS(GO:0016048) |  | 19 | 0.70 | 1.10 | 0.141 | 0.363 | 1.000 | 14001 | tags=47%, list=24%, signal=62% |
| 4856 | RECEPTOR\_SERINE\_THREONINE\_KINASE\_BINDING(GO:0033612) |  | 15 | 0.70 | 1.10 | 0.141 | 0.363 | 1.000 | 10951 | tags=40%, list=19%, signal=49% |
| 4857 | CD4\_POSITIVE\_OR\_CD8\_POSITIVE\_\_ALPHA\_BETA\_T\_CELL\_LINEAGE\_COMMITMENT(GO:0043369) |  | 20 | 0.67 | 1.10 | 0.141 | 0.363 | 1.000 | 9023 | tags=25%, list=15%, signal=29% |
| 4858 | REGULATION\_OF\_SMALL\_GTPASE\_MEDIATED\_SIGNAL\_TRANSDUCTION(GO:0051056) |  | 323 | 0.66 | 1.10 | 0.141 | 0.363 | 1.000 | 14699 | tags=36%, list=25%, signal=47% |
| 4859 | REGULATION\_OF\_NIK\_NF\_KAPPAB\_SIGNALING(GO:1901222) |  | 62 | 0.62 | 1.10 | 0.141 | 0.363 | 1.000 | 16766 | tags=50%, list=28%, signal=70% |
| 4860 | ESTABLISHMENT\_OF\_ORGANELLE\_LOCALIZATION(GO:0051656) |  | 435 | 0.67 | 1.10 | 0.141 | 0.363 | 1.000 | 11063 | tags=23%, list=19%, signal=28% |
| 4861 | BLOOD\_COAGULATION\_\_FIBRIN\_CLOT\_FORMATION(GO:0072378) |  | 27 | 0.68 | 1.10 | 0.141 | 0.363 | 1.000 | 11960 | tags=37%, list=20%, signal=46% |
| 4862 | RESPONSE\_TO\_BRONCHODILATOR(GO:0097366) |  | 45 | 0.69 | 1.10 | 0.141 | 0.363 | 1.000 | 15560 | tags=49%, list=26%, signal=66% |
| 4863 | METALLOEXOPEPTIDASE\_ACTIVITY(GO:0008235) |  | 58 | 0.63 | 1.10 | 0.141 | 0.363 | 1.000 | 18396 | tags=60%, list=31%, signal=88% |
| 4864 | NEURAL\_RETINA\_DEVELOPMENT(GO:0003407) |  | 60 | 0.70 | 1.10 | 0.141 | 0.363 | 1.000 | 12010 | tags=40%, list=20%, signal=50% |
| 4865 | BRUSH\_BORDER\_MEMBRANE(GO:0031526) |  | 53 | 0.66 | 1.10 | 0.141 | 0.363 | 1.000 | 12090 | tags=32%, list=20%, signal=40% |
| 4866 | PHOSPHATASE\_BINDING(GO:0019902) |  | 165 | 0.66 | 1.10 | 0.141 | 0.363 | 1.000 | 11777 | tags=20%, list=20%, signal=25% |
| 4867 | CELLULAR\_RESPIRATION(GO:0045333) |  | 188 | 0.62 | 1.10 | 0.141 | 0.363 | 1.000 | 20227 | tags=65%, list=34%, signal=98% |
| 4868 | BIOACTIVE\_LIPID\_RECEPTOR\_ACTIVITY(GO:0045125) |  | 15 | 0.71 | 1.10 | 0.141 | 0.363 | 1.000 | 13333 | tags=33%, list=23%, signal=43% |
| 4869 | ENDOSOMAL\_PART(GO:0044440) |  | 492 | 0.66 | 1.10 | 0.141 | 0.363 | 1.000 | 10450 | tags=17%, list=18%, signal=21% |
| 4870 | REGULATION\_OF\_GENE\_EXPRESSION\_BY\_GENETIC\_IMPRINTING(GO:0006349) |  | 18 | 0.63 | 1.10 | 0.000 | 0.363 | 1.000 | 21613 | tags=89%, list=37%, signal=140% |
| 4871 | BONE\_RESORPTION(GO:0045453) |  | 59 | 0.66 | 1.10 | 0.141 | 0.363 | 1.000 | 10901 | tags=32%, list=18%, signal=39% |
| 4872 | NUCLEOPLASM(GO:0005654) |  | 3470 | 0.68 | 1.10 | 0.141 | 0.363 | 1.000 | 13576 | tags=31%, list=23%, signal=38% |
| 4873 | REGULATION\_OF\_EXTENT\_OF\_CELL\_GROWTH(GO:0061387) |  | 91 | 0.68 | 1.10 | 0.141 | 0.363 | 1.000 | 11339 | tags=27%, list=19%, signal=34% |
| 4874 | REGULATION\_OF\_INTERLEUKIN\_2\_PRODUCTION(GO:0032663) |  | 51 | 0.67 | 1.10 | 0.141 | 0.363 | 1.000 | 15869 | tags=47%, list=27%, signal=64% |
| 4875 | INTRACELLULAR\_PROTEIN\_TRANSPORT(GO:0006886) |  | 972 | 0.67 | 1.10 | 0.141 | 0.363 | 1.000 | 9047 | tags=9%, list=15%, signal=10% |
| 4876 | NEGATIVE\_REGULATION\_OF\_RHO\_PROTEIN\_SIGNAL\_TRANSDUCTION(GO:0035024) |  | 19 | 0.67 | 1.10 | 0.141 | 0.363 | 1.000 | 15403 | tags=53%, list=26%, signal=71% |
| 4877 | ATTACHMENT\_OF\_SPINDLE\_MICROTUBULES\_TO\_KINETOCHORE(GO:0008608) |  | 33 | 0.64 | 1.10 | 0.000 | 0.363 | 1.000 | 14934 | tags=67%, list=25%, signal=89% |
| 4878 | NEGATIVE\_REGULATION\_OF\_LEUKOCYTE\_CELL\_CELL\_ADHESION(GO:1903038) |  | 111 | 0.65 | 1.10 | 0.141 | 0.363 | 1.000 | 15466 | tags=36%, list=26%, signal=49% |
| 4879 | NEGATIVE\_REGULATION\_OF\_PROTEIN\_COMPLEX\_DISASSEMBLY(GO:0043242) |  | 65 | 0.63 | 1.10 | 0.141 | 0.363 | 1.000 | 17265 | tags=46%, list=29%, signal=65% |
| 4880 | CELLULAR\_RESPONSE\_TO\_AMINO\_ACID\_STARVATION(GO:0034198) |  | 45 | 0.62 | 1.10 | 0.000 | 0.362 | 1.000 | 17567 | tags=60%, list=30%, signal=85% |
| 4881 | EXONUCLEASE\_ACTIVITY(GO:0004527) |  | 80 | 0.62 | 1.10 | 0.000 | 0.362 | 1.000 | 21004 | tags=83%, list=36%, signal=128% |
| 4882 | REGULATION\_OF\_SYNAPTIC\_VESICLE\_EXOCYTOSIS(GO:2000300) |  | 26 | 0.64 | 1.10 | 0.000 | 0.362 | 1.000 | 17682 | tags=69%, list=30%, signal=99% |
| 4883 | ACETYLATION\_DEPENDENT\_PROTEIN\_BINDING(GO:0140033) |  | 18 | 0.64 | 1.10 | 0.000 | 0.362 | 1.000 | 13760 | tags=39%, list=23%, signal=51% |
| 4884 | LYSINE\_ACETYLATED\_HISTONE\_BINDING(GO:0070577) |  | 18 | 0.64 | 1.10 | 0.000 | 0.362 | 1.000 | 13760 | tags=39%, list=23%, signal=51% |
| 4885 | MOTILE\_CILIUM\_ASSEMBLY(GO:0044458) |  | 23 | 0.64 | 1.10 | 0.000 | 0.362 | 1.000 | 18091 | tags=70%, list=31%, signal=100% |
| 4886 | METALLOCARBOXYPEPTIDASE\_ACTIVITY(GO:0004181) |  | 27 | 0.65 | 1.10 | 0.141 | 0.362 | 1.000 | 18368 | tags=67%, list=31%, signal=97% |
| 4887 | STEROID\_HYDROXYLASE\_ACTIVITY(GO:0008395) |  | 35 | 0.64 | 1.10 | 0.141 | 0.362 | 1.000 | 13642 | tags=37%, list=23%, signal=48% |
| 4888 | NEGATIVE\_REGULATION\_OF\_ADAPTIVE\_IMMUNE\_RESPONSE(GO:0002820) |  | 36 | 0.68 | 1.10 | 0.141 | 0.362 | 1.000 | 15529 | tags=33%, list=26%, signal=45% |
| 4889 | HYALURONAN\_METABOLIC\_PROCESS(GO:0030212) |  | 36 | 0.67 | 1.10 | 0.141 | 0.362 | 1.000 | 9021 | tags=28%, list=15%, signal=33% |
| 4890 | RNA\_POLYMERASE\_CORE\_ENZYME\_BINDING(GO:0043175) |  | 50 | 0.65 | 1.10 | 0.000 | 0.362 | 1.000 | 15530 | tags=34%, list=26%, signal=46% |
| 4891 | GLUCOCORTICOID\_METABOLIC\_PROCESS(GO:0008211) |  | 24 | 0.68 | 1.10 | 0.141 | 0.362 | 1.000 | 10811 | tags=38%, list=18%, signal=46% |
| 4892 | DYNEIN\_COMPLEX(GO:0030286) |  | 49 | 0.65 | 1.10 | 0.141 | 0.362 | 1.000 | 14289 | tags=45%, list=24%, signal=59% |
| 4893 | POSITIVE\_REGULATION\_BY\_HOST\_OF\_VIRAL\_TRANSCRIPTION(GO:0043923) |  | 17 | 0.65 | 1.10 | 0.000 | 0.362 | 1.000 | 18343 | tags=71%, list=31%, signal=102% |
| 4894 | REGULATION\_OF\_STRIATED\_MUSCLE\_CELL\_DIFFERENTIATION(GO:0051153) |  | 111 | 0.65 | 1.10 | 0.141 | 0.362 | 1.000 | 9543 | tags=22%, list=16%, signal=26% |
| 4895 | TRNA\_EXPORT\_FROM\_NUCLEUS(GO:0006409) |  | 32 | 0.64 | 1.10 | 0.000 | 0.362 | 1.000 | 20253 | tags=91%, list=34%, signal=138% |
| 4896 | TRNA\_CONTAINING\_RIBONUCLEOPROTEIN\_COMPLEX\_EXPORT\_FROM\_NUCLEUS(GO:0071431) |  | 32 | 0.64 | 1.10 | 0.000 | 0.362 | 1.000 | 20253 | tags=91%, list=34%, signal=138% |
| 4897 | VIRAL\_GENE\_EXPRESSION(GO:0019080) |  | 188 | 0.63 | 1.10 | 0.141 | 0.362 | 1.000 | 18212 | tags=53%, list=31%, signal=76% |
| 4898 | VIRAL\_TRANSCRIPTION(GO:0019083) |  | 174 | 0.63 | 1.10 | 0.141 | 0.362 | 1.000 | 17228 | tags=45%, list=29%, signal=63% |
| 4899 | ATPASE\_ACTIVITY(GO:0016887) |  | 443 | 0.65 | 1.10 | 0.141 | 0.362 | 1.000 | 15400 | tags=45%, list=26%, signal=61% |
| 4900 | NEGATIVE\_REGULATION\_OF\_CARDIAC\_MUSCLE\_HYPERTROPHY(GO:0010614) |  | 28 | 0.67 | 1.10 | 0.141 | 0.362 | 1.000 | 13557 | tags=25%, list=23%, signal=32% |
| 4901 | POSITIVE\_REGULATION\_OF\_VIRAL\_TRANSCRIPTION(GO:0050434) |  | 40 | 0.62 | 1.10 | 0.141 | 0.362 | 1.000 | 15831 | tags=40%, list=27%, signal=55% |
| 4902 | POSITIVE\_REGULATION\_OF\_HEART\_RATE(GO:0010460) |  | 25 | 0.67 | 1.10 | 0.141 | 0.362 | 1.000 | 17517 | tags=52%, list=30%, signal=74% |
| 4903 | POSITIVE\_REGULATION\_OF\_NUCLEAR\_DIVISION(GO:0051785) |  | 61 | 0.65 | 1.10 | 0.141 | 0.362 | 1.000 | 17160 | tags=54%, list=29%, signal=76% |
| 4904 | NEGATIVE\_REGULATION\_OF\_DNA\_REPLICATION(GO:0008156) |  | 38 | 0.64 | 1.10 | 0.000 | 0.362 | 1.000 | 18905 | tags=71%, list=32%, signal=104% |
| 4905 | NCRNA\_EXPORT\_FROM\_NUCLEUS(GO:0097064) |  | 36 | 0.64 | 1.10 | 0.000 | 0.362 | 1.000 | 20253 | tags=92%, list=34%, signal=139% |
| 4906 | LIPOPROTEIN\_PARTICLE\_BINDING(GO:0071813) |  | 24 | 0.68 | 1.10 | 0.141 | 0.362 | 1.000 | 6255 | tags=17%, list=11%, signal=19% |
| 4907 | PROTEIN\_LIPID\_COMPLEX\_BINDING(GO:0071814) |  | 24 | 0.68 | 1.10 | 0.141 | 0.362 | 1.000 | 6255 | tags=17%, list=11%, signal=19% |
| 4908 | NEGATIVE\_REGULATION\_OF\_LIPID\_METABOLIC\_PROCESS(GO:0045833) |  | 82 | 0.66 | 1.10 | 0.141 | 0.362 | 1.000 | 13039 | tags=29%, list=22%, signal=37% |
| 4909 | NUCLEOSIDE\_BISPHOSPHATE\_BIOSYNTHETIC\_PROCESS(GO:0033866) |  | 16 | 0.63 | 1.10 | 0.141 | 0.362 | 1.000 | 18244 | tags=69%, list=31%, signal=99% |
| 4910 | PURINE\_NUCLEOSIDE\_BISPHOSPHATE\_BIOSYNTHETIC\_PROCESS(GO:0034033) |  | 16 | 0.63 | 1.10 | 0.141 | 0.362 | 1.000 | 18244 | tags=69%, list=31%, signal=99% |
| 4911 | RIBONUCLEOSIDE\_BISPHOSPHATE\_BIOSYNTHETIC\_PROCESS(GO:0034030) |  | 16 | 0.63 | 1.10 | 0.141 | 0.362 | 1.000 | 18244 | tags=69%, list=31%, signal=99% |
| 4912 | MALE\_MEIOSIS\_I(GO:0007141) |  | 23 | 0.63 | 1.10 | 0.000 | 0.362 | 1.000 | 17719 | tags=61%, list=30%, signal=87% |
| 4913 | MITOTIC\_CELL\_CYCLE\_CHECKPOINT(GO:0007093) |  | 168 | 0.67 | 1.10 | 0.141 | 0.362 | 1.000 | 16045 | tags=50%, list=27%, signal=68% |
| 4914 | MICROFILAMENT\_MOTOR\_ACTIVITY(GO:0000146) |  | 22 | 0.66 | 1.10 | 0.141 | 0.362 | 1.000 | 7898 | tags=32%, list=13%, signal=37% |
| 4915 | UNFOLDED\_PROTEIN\_BINDING(GO:0051082) |  | 109 | 0.62 | 1.10 | 0.000 | 0.362 | 1.000 | 18646 | tags=61%, list=32%, signal=90% |
| 4916 | POSITIVE\_REGULATION\_OF\_MUSCLE\_CONTRACTION(GO:0045933) |  | 52 | 0.68 | 1.10 | 0.141 | 0.362 | 1.000 | 9682 | tags=25%, list=16%, signal=30% |
| 4917 | TRANSFORMING\_GROWTH\_FACTOR\_BETA\_RECEPTOR\_SIGNALING\_PATHWAY(GO:0007179) |  | 181 | 0.65 | 1.10 | 0.141 | 0.362 | 1.000 | 14781 | tags=35%, list=25%, signal=47% |
| 4918 | NEGATIVE\_REGULATION\_OF\_T\_CELL\_ACTIVATION(GO:0050868) |  | 99 | 0.65 | 1.10 | 0.141 | 0.362 | 1.000 | 15466 | tags=36%, list=26%, signal=49% |
| 4919 | DNA\_DEPENDENT\_DNA\_REPLICATION(GO:0006261) |  | 149 | 0.62 | 1.10 | 0.000 | 0.362 | 1.000 | 17539 | tags=68%, list=30%, signal=96% |
| 4920 | INTERSTRAND\_CROSS\_LINK\_REPAIR(GO:0036297) |  | 51 | 0.62 | 1.10 | 0.000 | 0.362 | 1.000 | 20681 | tags=80%, list=35%, signal=124% |
| 4921 | GENITALIA\_DEVELOPMENT(GO:0048806) |  | 46 | 0.69 | 1.10 | 0.141 | 0.362 | 1.000 | 9951 | tags=33%, list=17%, signal=39% |
| 4922 | MHC\_PROTEIN\_BINDING(GO:0042287) |  | 30 | 0.66 | 1.10 | 0.141 | 0.363 | 1.000 | 8325 | tags=27%, list=14%, signal=31% |
| 4923 | REGULATION\_OF\_ODONTOGENESIS(GO:0042481) |  | 26 | 0.70 | 1.10 | 0.141 | 0.363 | 1.000 | 10980 | tags=35%, list=19%, signal=42% |
| 4924 | REGULATION\_OF\_THE\_FORCE\_OF\_HEART\_CONTRACTION(GO:0002026) |  | 27 | 0.67 | 1.10 | 0.141 | 0.362 | 1.000 | 10309 | tags=26%, list=17%, signal=31% |
| 4925 | NATURAL\_KILLER\_CELL\_DIFFERENTIATION(GO:0001779) |  | 19 | 0.62 | 1.10 | 0.000 | 0.363 | 1.000 | 20690 | tags=68%, list=35%, signal=105% |
| 4926 | AXON\_EXTENSION(GO:0048675) |  | 105 | 0.67 | 1.10 | 0.141 | 0.363 | 1.000 | 11339 | tags=27%, list=19%, signal=33% |
| 4927 | NEGATIVE\_REGULATION\_OF\_ERBB\_SIGNALING\_PATHWAY(GO:1901185) |  | 56 | 0.64 | 1.10 | 0.141 | 0.363 | 1.000 | 14684 | tags=32%, list=25%, signal=43% |
| 4928 | RACEMASE\_AND\_EPIMERASE\_ACTIVITY(GO:0016854) |  | 16 | 0.65 | 1.10 | 0.141 | 0.363 | 1.000 | 18771 | tags=63%, list=32%, signal=92% |
| 4929 | REGULATION\_OF\_KERATINOCYTE\_PROLIFERATION(GO:0010837) |  | 29 | 0.70 | 1.10 | 0.141 | 0.363 | 1.000 | 14978 | tags=41%, list=25%, signal=55% |
| 4930 | POSITIVE\_REGULATION\_OF\_DNA\_DEPENDENT\_DNA\_REPLICATION(GO:2000105) |  | 15 | 0.64 | 1.10 | 0.000 | 0.363 | 1.000 | 15720 | tags=53%, list=27%, signal=73% |
| 4931 | RETINA\_LAYER\_FORMATION(GO:0010842) |  | 22 | 0.70 | 1.10 | 0.141 | 0.363 | 1.000 | 13338 | tags=50%, list=23%, signal=65% |
| 4932 | PROTEIN\_PEPTIDYL\_PROLYL\_ISOMERIZATION(GO:0000413) |  | 44 | 0.63 | 1.10 | 0.000 | 0.363 | 1.000 | 17939 | tags=55%, list=30%, signal=78% |
| 4933 | PEPTIDYL\_PROLYL\_CIS\_TRANS\_ISOMERASE\_ACTIVITY(GO:0003755) |  | 44 | 0.63 | 1.10 | 0.000 | 0.363 | 1.000 | 17939 | tags=55%, list=30%, signal=78% |
| 4934 | AMYLOID\_BETA\_BINDING(GO:0001540) |  | 56 | 0.64 | 1.10 | 0.141 | 0.363 | 1.000 | 7441 | tags=14%, list=13%, signal=16% |
| 4935 | REGULATION\_OF\_VIRAL\_INDUCED\_CYTOPLASMIC\_PATTERN\_RECOGNITION\_RECEPTOR\_SIGNALING\_PATHWAY(GO:0039531) |  | 20 | 0.64 | 1.10 | 0.141 | 0.363 | 1.000 | 15128 | tags=40%, list=26%, signal=54% |
| 4936 | MITOTIC\_NUCLEAR\_ENVELOPE\_DISASSEMBLY(GO:0007077) |  | 43 | 0.64 | 1.10 | 0.000 | 0.363 | 1.000 | 20557 | tags=91%, list=35%, signal=139% |
| 4937 | CELLULAR\_DEFENSE\_RESPONSE(GO:0006968) |  | 56 | 0.63 | 1.10 | 0.141 | 0.363 | 1.000 | 13741 | tags=30%, list=23%, signal=40% |
| 4938 | FOLIC\_ACID\_CONTAINING\_COMPOUND\_METABOLIC\_PROCESS(GO:0006760) |  | 28 | 0.63 | 1.10 | 0.141 | 0.363 | 1.000 | 14644 | tags=46%, list=25%, signal=62% |
| 4939 | SITE\_OF\_DNA\_DAMAGE(GO:0090734) |  | 37 | 0.64 | 1.10 | 0.000 | 0.363 | 1.000 | 16139 | tags=62%, list=27%, signal=85% |
| 4940 | MICROTUBULE\_END(GO:1990752) |  | 27 | 0.64 | 1.10 | 0.000 | 0.363 | 1.000 | 17976 | tags=63%, list=30%, signal=90% |
| 4941 | CARBOXY\_TERMINAL\_DOMAIN\_PROTEIN\_KINASE\_COMPLEX(GO:0032806) |  | 24 | 0.65 | 1.10 | 0.000 | 0.363 | 1.000 | 19635 | tags=83%, list=33%, signal=125% |
| 4942 | PROTEIN\_TRIMERIZATION(GO:0070206) |  | 50 | 0.65 | 1.10 | 0.141 | 0.363 | 1.000 | 17593 | tags=54%, list=30%, signal=77% |
| 4943 | PHOSPHATASE\_ACTIVITY(GO:0016791) |  | 284 | 0.64 | 1.10 | 0.141 | 0.363 | 1.000 | 12794 | tags=25%, list=22%, signal=32% |
| 4944 | EPHRIN\_RECEPTOR\_SIGNALING\_PATHWAY(GO:0048013) |  | 88 | 0.63 | 1.10 | 0.141 | 0.363 | 1.000 | 18659 | tags=55%, list=32%, signal=80% |
| 4945 | VASCULAR\_ASSOCIATED\_SMOOTH\_MUSCLE\_CELL\_APOPTOTIC\_PROCESS(GO:1905288) |  | 15 | 0.67 | 1.10 | 0.000 | 0.363 | 1.000 | 19010 | tags=40%, list=32%, signal=59% |
| 4946 | REGULATION\_OF\_VASCULAR\_ASSOCIATED\_SMOOTH\_MUSCLE\_CELL\_APOPTOTIC\_PROCESS(GO:1905459) |  | 15 | 0.67 | 1.10 | 0.000 | 0.363 | 1.000 | 19010 | tags=40%, list=32%, signal=59% |
| 4947 | REGULATION\_OF\_TRANSLATIONAL\_ELONGATION(GO:0006448) |  | 17 | 0.62 | 1.10 | 0.000 | 0.363 | 1.000 | 21293 | tags=88%, list=36%, signal=138% |
| 4948 | CENTRIOLE\_REPLICATION(GO:0007099) |  | 30 | 0.65 | 1.10 | 0.141 | 0.363 | 1.000 | 20378 | tags=90%, list=34%, signal=137% |
| 4949 | CUL3\_RING\_UBIQUITIN\_LIGASE\_COMPLEX(GO:0031463) |  | 71 | 0.63 | 1.10 | 0.141 | 0.363 | 1.000 | 15403 | tags=42%, list=26%, signal=57% |
| 4950 | MEIOTIC\_CELL\_CYCLE(GO:0051321) |  | 243 | 0.65 | 1.10 | 0.141 | 0.363 | 1.000 | 15525 | tags=48%, list=26%, signal=64% |
| 4951 | T\_CELL\_RECEPTOR\_COMPLEX(GO:0042101) |  | 19 | 0.66 | 1.10 | 0.141 | 0.363 | 1.000 | 7103 | tags=16%, list=12%, signal=18% |
| 4952 | REGULATION\_OF\_TOLL\_LIKE\_RECEPTOR\_SIGNALING\_PATHWAY(GO:0034121) |  | 60 | 0.66 | 1.10 | 0.141 | 0.363 | 1.000 | 13828 | tags=32%, list=23%, signal=41% |
| 4953 | NUCLEAR\_ENVELOPE\_DISASSEMBLY(GO:0051081) |  | 47 | 0.64 | 1.10 | 0.000 | 0.363 | 1.000 | 20557 | tags=91%, list=35%, signal=140% |
| 4954 | MEMBRANE\_DISASSEMBLY(GO:0030397) |  | 47 | 0.64 | 1.10 | 0.000 | 0.363 | 1.000 | 20557 | tags=91%, list=35%, signal=140% |
| 4955 | DNA\_STRAND\_ELONGATION\_INVOLVED\_IN\_DNA\_REPLICATION(GO:0006271) |  | 18 | 0.65 | 1.10 | 0.000 | 0.362 | 1.000 | 19692 | tags=94%, list=33%, signal=142% |
| 4956 | GASTRIC\_ACID\_SECRETION(GO:0001696) |  | 17 | 0.67 | 1.10 | 0.141 | 0.362 | 1.000 | 9682 | tags=35%, list=16%, signal=42% |
| 4957 | RNA\_EXPORT\_FROM\_NUCLEUS(GO:0006405) |  | 142 | 0.62 | 1.10 | 0.000 | 0.362 | 1.000 | 20302 | tags=82%, list=34%, signal=124% |
| 4958 | AUTOPHAGOSOME\_MATURATION(GO:0097352) |  | 36 | 0.64 | 1.10 | 0.141 | 0.362 | 1.000 | 11317 | tags=25%, list=19%, signal=31% |
| 4959 | NEGATIVE\_REGULATION\_OF\_ANOIKIS(GO:2000811) |  | 16 | 0.67 | 1.10 | 0.141 | 0.362 | 1.000 | 9784 | tags=19%, list=17%, signal=22% |
| 4960 | RESPONSE\_TO\_ESTROGEN(GO:0043627) |  | 76 | 0.69 | 1.10 | 0.141 | 0.362 | 1.000 | 11951 | tags=34%, list=20%, signal=43% |
| 4961 | ENDONUCLEASE\_ACTIVITY(GO:0004519) |  | 137 | 0.62 | 1.10 | 0.000 | 0.362 | 1.000 | 17983 | tags=56%, list=30%, signal=81% |
| 4962 | EXTRACELLULAR\_NEGATIVE\_REGULATION\_OF\_SIGNAL\_TRANSDUCTION(GO:1900116) |  | 17 | 0.69 | 1.10 | 0.141 | 0.362 | 1.000 | 10430 | tags=29%, list=18%, signal=36% |
| 4963 | EXTRACELLULAR\_REGULATION\_OF\_SIGNAL\_TRANSDUCTION(GO:1900115) |  | 17 | 0.69 | 1.10 | 0.141 | 0.362 | 1.000 | 10430 | tags=29%, list=18%, signal=36% |
| 4964 | NUCLEOTIDE\_EXCISION\_REPAIR\_\_DNA\_GAP\_FILLING(GO:0006297) |  | 23 | 0.62 | 1.10 | 0.000 | 0.362 | 1.000 | 19834 | tags=83%, list=34%, signal=124% |
| 4965 | RESPONSE\_TO\_FOOD(GO:0032094) |  | 39 | 0.65 | 1.10 | 0.141 | 0.362 | 1.000 | 18462 | tags=56%, list=31%, signal=82% |
| 4966 | CENTROSOME\_DUPLICATION(GO:0051298) |  | 64 | 0.62 | 1.10 | 0.141 | 0.362 | 1.000 | 21117 | tags=81%, list=36%, signal=126% |
| 4967 | ADRENERGIC\_RECEPTOR\_SIGNALING\_PATHWAY(GO:0071875) |  | 27 | 0.64 | 1.10 | 0.141 | 0.362 | 1.000 | 10887 | tags=22%, list=18%, signal=27% |
| 4968 | NUCLEOTIDE\_EXCISION\_REPAIR\_\_DNA\_INCISION\_\_3'\_TO\_LESION(GO:0006295) |  | 22 | 0.66 | 1.10 | 0.000 | 0.362 | 1.000 | 18883 | tags=82%, list=32%, signal=120% |
| 4969 | NEGATIVE\_REGULATION\_OF\_MUSCLE\_HYPERTROPHY(GO:0014741) |  | 30 | 0.66 | 1.10 | 0.141 | 0.362 | 1.000 | 13557 | tags=27%, list=23%, signal=35% |
| 4970 | CELL\_CYCLE\_PHASE\_TRANSITION(GO:0044770) |  | 593 | 0.66 | 1.09 | 0.141 | 0.362 | 1.000 | 13522 | tags=34%, list=23%, signal=43% |
| 4971 | REMOVAL\_OF\_SUPEROXIDE\_RADICALS(GO:0019430) |  | 25 | 0.66 | 1.09 | 0.141 | 0.362 | 1.000 | 15716 | tags=40%, list=27%, signal=54% |
| 4972 | POSITIVE\_REGULATION\_OF\_TRANSFORMING\_GROWTH\_FACTOR\_BETA\_PRODUCTION(GO:0071636) |  | 17 | 0.67 | 1.09 | 0.141 | 0.362 | 1.000 | 15462 | tags=47%, list=26%, signal=64% |
| 4973 | NUCLEOTIDE\_EXCISION\_REPAIR\_\_PREINCISION\_COMPLEX\_STABILIZATION(GO:0006293) |  | 21 | 0.66 | 1.09 | 0.000 | 0.362 | 1.000 | 18883 | tags=86%, list=32%, signal=126% |
| 4974 | POSITIVE\_T\_CELL\_SELECTION(GO:0043368) |  | 31 | 0.65 | 1.09 | 0.141 | 0.362 | 1.000 | 9023 | tags=19%, list=15%, signal=23% |
| 4975 | BROWN\_FAT\_CELL\_DIFFERENTIATION(GO:0050873) |  | 39 | 0.68 | 1.09 | 0.141 | 0.362 | 1.000 | 10262 | tags=26%, list=17%, signal=31% |
| 4976 | POSITIVE\_REGULATION\_OF\_RESPONSE\_TO\_BIOTIC\_STIMULUS(GO:0002833) |  | 40 | 0.63 | 1.09 | 0.141 | 0.362 | 1.000 | 15744 | tags=43%, list=27%, signal=58% |
| 4977 | MITOTIC\_SPINDLE\_POLE(GO:0097431) |  | 26 | 0.65 | 1.09 | 0.000 | 0.362 | 1.000 | 15088 | tags=62%, list=26%, signal=83% |
| 4978 | POSITIVE\_REGULATION\_OF\_ORGANIC\_ACID\_TRANSPORT(GO:0032892) |  | 31 | 0.67 | 1.09 | 0.141 | 0.362 | 1.000 | 17522 | tags=55%, list=30%, signal=78% |
| 4979 | SENSORY\_PERCEPTION\_OF\_TASTE(GO:0050909) |  | 65 | 0.68 | 1.09 | 0.141 | 0.362 | 1.000 | 11683 | tags=18%, list=20%, signal=23% |
| 4980 | CYTOKINE\_PRODUCTION\_INVOLVED\_IN\_INFLAMMATORY\_RESPONSE(GO:0002534) |  | 37 | 0.65 | 1.09 | 0.141 | 0.362 | 1.000 | 14390 | tags=35%, list=24%, signal=46% |
| 4981 | VESICLE\_LOCALIZATION(GO:0051648) |  | 246 | 0.67 | 1.09 | 0.141 | 0.362 | 1.000 | 10763 | tags=21%, list=18%, signal=25% |
| 4982 | PROTEIN\_LOCALIZATION\_TO\_CENTROSOME(GO:0071539) |  | 27 | 0.64 | 1.09 | 0.000 | 0.362 | 1.000 | 19782 | tags=89%, list=33%, signal=134% |
| 4983 | NEGATIVE\_REGULATION\_OF\_CELL\_CYCLE(GO:0045786) |  | 625 | 0.66 | 1.09 | 0.141 | 0.362 | 1.000 | 13315 | tags=30%, list=23%, signal=38% |
| 4984 | GOLGI\_ASSOCIATED\_VESICLE\_MEMBRANE(GO:0030660) |  | 105 | 0.62 | 1.09 | 0.141 | 0.362 | 1.000 | 19349 | tags=52%, list=33%, signal=78% |
| 4985 | REGULATION\_OF\_LEUKOCYTE\_MEDIATED\_IMMUNITY(GO:0002703) |  | 162 | 0.64 | 1.09 | 0.141 | 0.362 | 1.000 | 15869 | tags=37%, list=27%, signal=50% |
| 4986 | REGULATION\_OF\_NITRIC\_OXIDE\_SYNTHASE\_ACTIVITY(GO:0050999) |  | 46 | 0.62 | 1.09 | 0.141 | 0.362 | 1.000 | 16715 | tags=48%, list=28%, signal=67% |
| 4987 | EXONUCLEOLYTIC\_NUCLEAR\_TRANSCRIBED\_MRNA\_CATABOLIC\_PROCESS\_INVOLVED\_IN\_DEADENYLATION\_DEPENDENT\_DECAY(GO:0043928) |  | 33 | 0.61 | 1.09 | 0.000 | 0.362 | 1.000 | 21067 | tags=82%, list=36%, signal=127% |
| 4988 | EMBRYONIC\_AXIS\_SPECIFICATION(GO:0000578) |  | 35 | 0.68 | 1.09 | 0.141 | 0.362 | 1.000 | 13668 | tags=43%, list=23%, signal=56% |
| 4989 | MAGNESIUM\_ION\_TRANSPORT(GO:0015693) |  | 18 | 0.65 | 1.09 | 0.000 | 0.362 | 1.000 | 18680 | tags=78%, list=32%, signal=114% |
| 4990 | POSITIVE\_REGULATION\_OF\_TELOMERE\_MAINTENANCE(GO:0032206) |  | 51 | 0.62 | 1.09 | 0.000 | 0.362 | 1.000 | 20903 | tags=84%, list=35%, signal=130% |
| 4991 | CEREBRAL\_CORTEX\_NEURON\_DIFFERENTIATION(GO:0021895) |  | 24 | 0.68 | 1.09 | 0.141 | 0.362 | 1.000 | 13386 | tags=46%, list=23%, signal=59% |
| 4992 | AGGRESOME(GO:0016235) |  | 42 | 0.63 | 1.09 | 0.000 | 0.362 | 1.000 | 19749 | tags=69%, list=33%, signal=104% |
| 4993 | PERICENTRIOLAR\_MATERIAL(GO:0000242) |  | 20 | 0.66 | 1.09 | 0.000 | 0.362 | 1.000 | 19263 | tags=90%, list=33%, signal=133% |
| 4994 | REGULATION\_OF\_RECEPTOR\_BIOSYNTHETIC\_PROCESS(GO:0010869) |  | 20 | 0.67 | 1.09 | 0.141 | 0.362 | 1.000 | 10604 | tags=30%, list=18%, signal=37% |
| 4995 | REPLICATION\_FORK\_PROCESSING(GO:0031297) |  | 30 | 0.61 | 1.09 | 0.000 | 0.362 | 1.000 | 15764 | tags=60%, list=27%, signal=82% |
| 4996 | NEGATIVE\_REGULATION\_OF\_LEUKOCYTE\_CHEMOTAXIS(GO:0002689) |  | 15 | 0.69 | 1.09 | 0.141 | 0.362 | 1.000 | 7081 | tags=20%, list=12%, signal=23% |
| 4997 | OTHER\_ORGANISM\_CELL(GO:0044216) |  | 74 | 0.63 | 1.09 | 0.000 | 0.362 | 1.000 | 18624 | tags=66%, list=31%, signal=97% |
| 4998 | OTHER\_ORGANISM\_PART(GO:0044217) |  | 74 | 0.63 | 1.09 | 0.000 | 0.362 | 1.000 | 18624 | tags=66%, list=31%, signal=97% |
| 4999 | OTHER\_ORGANISM(GO:0044215) |  | 74 | 0.63 | 1.09 | 0.000 | 0.362 | 1.000 | 18624 | tags=66%, list=31%, signal=97% |
| 5000 | RESPONSE\_TO\_AMINE(GO:0014075) |  | 39 | 0.67 | 1.09 | 0.141 | 0.362 | 1.000 | 13132 | tags=41%, list=22%, signal=53% |
| 5001 | NUCLEASE\_ACTIVITY(GO:0004518) |  | 218 | 0.61 | 1.09 | 0.000 | 0.362 | 1.000 | 18099 | tags=60%, list=31%, signal=86% |
| 5002 | HOST\_CELL(GO:0043657) |  | 68 | 0.63 | 1.09 | 0.000 | 0.362 | 1.000 | 18624 | tags=72%, list=31%, signal=105% |
| 5003 | HOST(GO:0018995) |  | 68 | 0.63 | 1.09 | 0.000 | 0.362 | 1.000 | 18624 | tags=72%, list=31%, signal=105% |
| 5004 | DNA\_REPLICATION\_INITIATION(GO:0006270) |  | 43 | 0.61 | 1.09 | 0.000 | 0.362 | 1.000 | 22441 | tags=95%, list=38%, signal=154% |
| 5005 | INTRINSIC\_COMPONENT\_OF\_THE\_CYTOPLASMIC\_SIDE\_OF\_THE\_PLASMA\_MEMBRANE(GO:0031235) |  | 16 | 0.63 | 1.09 | 0.141 | 0.362 | 1.000 | 16021 | tags=50%, list=27%, signal=69% |
| 5006 | AUDITORY\_RECEPTOR\_CELL\_DEVELOPMENT(GO:0060117) |  | 19 | 0.63 | 1.09 | 0.141 | 0.362 | 1.000 | 18101 | tags=63%, list=31%, signal=91% |
| 5007 | SITE\_OF\_DOUBLE\_STRAND\_BREAK(GO:0035861) |  | 34 | 0.63 | 1.09 | 0.000 | 0.362 | 1.000 | 16139 | tags=62%, list=27%, signal=85% |
| 5008 | OXIDOREDUCTASE\_ACTIVITY\_\_ACTING\_ON\_PAIRED\_DONORS\_\_WITH\_INCORPORATION\_OR\_REDUCTION\_OF\_MOLECULAR\_OXYGEN\_\_2\_OXOGLUTARATE\_AS\_ONE\_DONOR\_\_AND\_INCORPORATION\_OF\_ONE\_ATOM\_EACH\_OF\_OXYGEN\_INTO\_BOTH\_DONORS(GO:0016706) |  | 45 | 0.68 | 1.09 | 0.141 | 0.362 | 1.000 | 14421 | tags=47%, list=24%, signal=62% |
| 5009 | DNA\_TEMPLATED\_TRANSCRIPTION\_\_TERMINATION(GO:0006353) |  | 107 | 0.62 | 1.09 | 0.000 | 0.362 | 1.000 | 20344 | tags=84%, list=34%, signal=128% |
| 5010 | HIPPO\_SIGNALING(GO:0035329) |  | 37 | 0.68 | 1.09 | 0.141 | 0.362 | 1.000 | 13943 | tags=46%, list=24%, signal=60% |
| 5011 | SENSORY\_PERCEPTION\_OF\_TEMPERATURE\_STIMULUS(GO:0050951) |  | 21 | 0.69 | 1.09 | 0.141 | 0.362 | 1.000 | 14001 | tags=48%, list=24%, signal=62% |
| 5012 | TRANSFORMING\_GROWTH\_FACTOR\_BETA\_PRODUCTION(GO:0071604) |  | 28 | 0.65 | 1.09 | 0.141 | 0.362 | 1.000 | 15462 | tags=46%, list=26%, signal=63% |
| 5013 | NUCLEOCYTOPLASMIC\_CARRIER\_ACTIVITY(GO:0140142) |  | 19 | 0.66 | 1.09 | 0.000 | 0.362 | 1.000 | 17173 | tags=79%, list=29%, signal=111% |
| 5014 | SIGNAL\_SEQUENCE\_BINDING(GO:0005048) |  | 46 | 0.63 | 1.09 | 0.000 | 0.362 | 1.000 | 18116 | tags=65%, list=31%, signal=94% |
| 5015 | NEGATIVE\_REGULATION\_OF\_PROTEASOMAL\_PROTEIN\_CATABOLIC\_PROCESS(GO:1901799) |  | 47 | 0.64 | 1.09 | 0.141 | 0.362 | 1.000 | 18043 | tags=57%, list=31%, signal=83% |
| 5016 | FICOLIN\_1\_RICH\_GRANULE\_MEMBRANE(GO:0101003) |  | 60 | 0.63 | 1.09 | 0.141 | 0.362 | 1.000 | 15074 | tags=25%, list=25%, signal=34% |
| 5017 | CILIARY\_BASAL\_BODY\_PLASMA\_MEMBRANE\_DOCKING(GO:0097711) |  | 95 | 0.62 | 1.09 | 0.000 | 0.362 | 1.000 | 19532 | tags=80%, list=33%, signal=119% |
| 5018 | BLASTOCYST\_GROWTH(GO:0001832) |  | 17 | 0.65 | 1.09 | 0.141 | 0.362 | 1.000 | 20722 | tags=94%, list=35%, signal=145% |
| 5019 | REGULATION\_OF\_G0\_TO\_G1\_TRANSITION(GO:0070316) |  | 44 | 0.63 | 1.09 | 0.000 | 0.362 | 1.000 | 19453 | tags=75%, list=33%, signal=112% |
| 5020 | GLYCINE\_BINDING(GO:0016594) |  | 15 | 0.67 | 1.09 | 0.141 | 0.362 | 1.000 | 16842 | tags=67%, list=28%, signal=93% |
| 5021 | POSITIVE\_REGULATION\_OF\_MONONUCLEAR\_CELL\_PROLIFERATION(GO:0032946) |  | 132 | 0.64 | 1.09 | 0.141 | 0.362 | 1.000 | 16684 | tags=38%, list=28%, signal=53% |
| 5022 | CYTOPLASMIC\_MICROTUBULE(GO:0005881) |  | 58 | 0.66 | 1.09 | 0.141 | 0.362 | 1.000 | 13741 | tags=38%, list=23%, signal=49% |
| 5023 | PURINE\_NUCLEOBASE\_METABOLIC\_PROCESS(GO:0006144) |  | 21 | 0.67 | 1.09 | 0.141 | 0.362 | 1.000 | 14615 | tags=62%, list=25%, signal=82% |
| 5024 | MITOCHONDRIAL\_PROTEIN\_COMPLEX(GO:0098798) |  | 268 | 0.61 | 1.09 | 0.141 | 0.362 | 1.000 | 20294 | tags=69%, list=34%, signal=105% |
| 5025 | GENETIC\_IMPRINTING(GO:0071514) |  | 26 | 0.61 | 1.09 | 0.141 | 0.362 | 1.000 | 21613 | tags=77%, list=37%, signal=121% |
| 5026 | PROTEASOME\_MEDIATED\_UBIQUITIN\_DEPENDENT\_PROTEIN\_CATABOLIC\_PROCESS(GO:0043161) |  | 390 | 0.61 | 1.09 | 0.141 | 0.362 | 1.000 | 17876 | tags=48%, list=30%, signal=69% |
| 5027 | AMINOPEPTIDASE\_ACTIVITY(GO:0004177) |  | 44 | 0.61 | 1.09 | 0.141 | 0.362 | 1.000 | 19837 | tags=64%, list=34%, signal=96% |
| 5028 | HISTONE\_MRNA\_METABOLIC\_PROCESS(GO:0008334) |  | 24 | 0.65 | 1.09 | 0.000 | 0.362 | 1.000 | 18871 | tags=88%, list=32%, signal=128% |
| 5029 | LIPID\_TRANSLOCATION(GO:0034204) |  | 25 | 0.68 | 1.09 | 0.141 | 0.362 | 1.000 | 13125 | tags=32%, list=22%, signal=41% |
| 5030 | SPECIFIC\_GRANULE\_LUMEN(GO:0035580) |  | 61 | 0.65 | 1.09 | 0.141 | 0.362 | 1.000 | 13765 | tags=25%, list=23%, signal=32% |
| 5031 | POSITIVE\_REGULATION\_OF\_LYMPHOCYTE\_PROLIFERATION(GO:0050671) |  | 131 | 0.63 | 1.09 | 0.141 | 0.362 | 1.000 | 16684 | tags=37%, list=28%, signal=52% |
| 5032 | ANTIGEN\_PROCESSING\_AND\_PRESENTATION(GO:0019882) |  | 225 | 0.64 | 1.09 | 0.141 | 0.362 | 1.000 | 13472 | tags=24%, list=23%, signal=31% |
| 5033 | NCRNA\_METABOLIC\_PROCESS(GO:0034660) |  | 584 | 0.62 | 1.09 | 0.141 | 0.362 | 1.000 | 19271 | tags=67%, list=33%, signal=98% |
| 5034 | PROTEIN\_LOCALIZATION\_TO\_MICROTUBULE\_ORGANIZING\_CENTER(GO:1905508) |  | 28 | 0.64 | 1.09 | 0.000 | 0.362 | 1.000 | 19782 | tags=86%, list=33%, signal=129% |
| 5035 | NONRIBOSOMAL\_PEPTIDE\_BIOSYNTHETIC\_PROCESS(GO:0019184) |  | 17 | 0.67 | 1.09 | 0.141 | 0.362 | 1.000 | 10576 | tags=35%, list=18%, signal=43% |
| 5036 | CARDIAC\_MUSCLE\_ADAPTATION(GO:0014887) |  | 31 | 0.64 | 1.09 | 0.141 | 0.362 | 1.000 | 18262 | tags=39%, list=31%, signal=56% |
| 5037 | MUSCLE\_HYPERTROPHY\_IN\_RESPONSE\_TO\_STRESS(GO:0003299) |  | 31 | 0.64 | 1.09 | 0.141 | 0.361 | 1.000 | 18262 | tags=39%, list=31%, signal=56% |
| 5038 | CARDIAC\_MUSCLE\_HYPERTROPHY\_IN\_RESPONSE\_TO\_STRESS(GO:0014898) |  | 31 | 0.64 | 1.09 | 0.141 | 0.361 | 1.000 | 18262 | tags=39%, list=31%, signal=56% |
| 5039 | MITOCHONDRIAL\_RNA\_METABOLIC\_PROCESS(GO:0000959) |  | 40 | 0.63 | 1.09 | 0.000 | 0.361 | 1.000 | 20597 | tags=93%, list=35%, signal=142% |
| 5040 | TELOMERE\_MAINTENANCE\_VIA\_SEMI\_CONSERVATIVE\_REPLICATION(GO:0032201) |  | 27 | 0.64 | 1.09 | 0.000 | 0.361 | 1.000 | 20209 | tags=89%, list=34%, signal=135% |
| 5041 | ACETYL\_COA\_BIOSYNTHETIC\_PROCESS(GO:0006085) |  | 20 | 0.64 | 1.09 | 0.000 | 0.361 | 1.000 | 20175 | tags=85%, list=34%, signal=129% |
| 5042 | REGULATION\_OF\_GLIAL\_CELL\_PROLIFERATION(GO:0060251) |  | 28 | 0.67 | 1.09 | 0.141 | 0.361 | 1.000 | 8554 | tags=14%, list=14%, signal=17% |
| 5043 | REGULATION\_OF\_TELOMERE\_MAINTENANCE\_VIA\_TELOMERASE(GO:0032210) |  | 53 | 0.62 | 1.09 | 0.000 | 0.361 | 1.000 | 20903 | tags=87%, list=35%, signal=134% |
| 5044 | ACROSOMAL\_VESICLE(GO:0001669) |  | 117 | 0.65 | 1.09 | 0.141 | 0.361 | 1.000 | 15889 | tags=33%, list=27%, signal=45% |
| 5045 | MICROTUBULE\_NUCLEATION(GO:0007020) |  | 26 | 0.62 | 1.09 | 0.141 | 0.361 | 1.000 | 21933 | tags=88%, list=37%, signal=141% |
| 5046 | PRE\_MRNA\_BINDING(GO:0036002) |  | 37 | 0.63 | 1.09 | 0.000 | 0.361 | 1.000 | 20302 | tags=84%, list=34%, signal=128% |
| 5047 | CYCLIN\_DEPENDENT\_PROTEIN\_SERINE\_THREONINE\_KINASE\_REGULATOR\_ACTIVITY(GO:0016538) |  | 29 | 0.64 | 1.09 | 0.000 | 0.361 | 1.000 | 19048 | tags=69%, list=32%, signal=102% |
| 5048 | ETHER\_METABOLIC\_PROCESS(GO:0018904) |  | 17 | 0.62 | 1.09 | 0.141 | 0.361 | 1.000 | 18032 | tags=59%, list=30%, signal=85% |
| 5049 | REGULATION\_OF\_CENTROSOME\_CYCLE(GO:0046605) |  | 55 | 0.62 | 1.09 | 0.141 | 0.361 | 1.000 | 21117 | tags=82%, list=36%, signal=127% |
| 5050 | REGULATION\_OF\_TELOMERE\_MAINTENANCE(GO:0032204) |  | 79 | 0.62 | 1.09 | 0.000 | 0.361 | 1.000 | 20903 | tags=85%, list=35%, signal=131% |
| 5051 | REGULATION\_OF\_TRANSFORMING\_GROWTH\_FACTOR\_BETA\_PRODUCTION(GO:0071634) |  | 27 | 0.66 | 1.09 | 0.141 | 0.361 | 1.000 | 15462 | tags=48%, list=26%, signal=65% |
| 5052 | TELOMERE\_MAINTENANCE\_VIA\_TELOMERASE(GO:0007004) |  | 67 | 0.62 | 1.09 | 0.000 | 0.361 | 1.000 | 20903 | tags=87%, list=35%, signal=134% |
| 5053 | CIS\_TRANS\_ISOMERASE\_ACTIVITY(GO:0016859) |  | 46 | 0.62 | 1.09 | 0.000 | 0.361 | 1.000 | 17939 | tags=52%, list=30%, signal=75% |
| 5054 | RIBONUCLEOPROTEIN\_COMPLEX\_EXPORT\_FROM\_NUCLEUS(GO:0071426) |  | 128 | 0.62 | 1.09 | 0.000 | 0.361 | 1.000 | 20302 | tags=81%, list=34%, signal=123% |
| 5055 | NUCLEOTIDE\_EXCISION\_REPAIR\_\_DNA\_INCISION(GO:0033683) |  | 40 | 0.62 | 1.09 | 0.000 | 0.361 | 1.000 | 19834 | tags=75%, list=34%, signal=113% |
| 5056 | KERATINOCYTE\_PROLIFERATION(GO:0043616) |  | 39 | 0.69 | 1.09 | 0.141 | 0.361 | 1.000 | 11480 | tags=28%, list=19%, signal=35% |
| 5057 | POSITIVE\_REGULATION\_OF\_LEUKOCYTE\_PROLIFERATION(GO:0070665) |  | 138 | 0.63 | 1.09 | 0.141 | 0.361 | 1.000 | 16684 | tags=38%, list=28%, signal=53% |
| 5058 | DNA\_DIRECTED\_RNA\_POLYMERASE\_II\_\_HOLOENZYME(GO:0016591) |  | 110 | 0.61 | 1.09 | 0.000 | 0.361 | 1.000 | 18523 | tags=61%, list=31%, signal=89% |
| 5059 | CENTRIOLE\_ASSEMBLY(GO:0098534) |  | 33 | 0.64 | 1.09 | 0.141 | 0.361 | 1.000 | 19564 | tags=82%, list=33%, signal=122% |
| 5060 | POSITIVE\_REGULATION\_OF\_CELLULAR\_RESPONSE\_TO\_TRANSFORMING\_GROWTH\_FACTOR\_BETA\_STIMULUS(GO:1903846) |  | 26 | 0.68 | 1.09 | 0.141 | 0.361 | 1.000 | 14410 | tags=42%, list=24%, signal=56% |
| 5061 | POSITIVE\_REGULATION\_OF\_TRANSFORMING\_GROWTH\_FACTOR\_BETA\_RECEPTOR\_SIGNALING\_PATHWAY(GO:0030511) |  | 26 | 0.68 | 1.09 | 0.141 | 0.361 | 1.000 | 14410 | tags=42%, list=24%, signal=56% |
| 5062 | AMMONIUM\_TRANSMEMBRANE\_TRANSPORT(GO:0072488) |  | 16 | 0.72 | 1.09 | 0.307 | 0.361 | 1.000 | 10803 | tags=31%, list=18%, signal=38% |
| 5063 | NEGATIVE\_REGULATION\_OF\_CAMP\_MEDIATED\_SIGNALING(GO:0043951) |  | 17 | 0.66 | 1.09 | 0.141 | 0.361 | 1.000 | 10309 | tags=29%, list=17%, signal=36% |
| 5064 | MANGANESE\_ION\_BINDING(GO:0030145) |  | 50 | 0.62 | 1.09 | 0.141 | 0.361 | 1.000 | 18396 | tags=66%, list=31%, signal=96% |
| 5065 | HEMATOPOIETIC\_STEM\_CELL\_PROLIFERATION(GO:0071425) |  | 22 | 0.64 | 1.09 | 0.141 | 0.361 | 1.000 | 17773 | tags=68%, list=30%, signal=97% |
| 5066 | NUCLEOBASE\_METABOLIC\_PROCESS(GO:0009112) |  | 39 | 0.64 | 1.09 | 0.141 | 0.361 | 1.000 | 17343 | tags=69%, list=29%, signal=98% |
| 5067 | REGULATION\_OF\_VIRAL\_ENTRY\_INTO\_HOST\_CELL(GO:0046596) |  | 30 | 0.69 | 1.09 | 0.141 | 0.361 | 1.000 | 6706 | tags=7%, list=11%, signal=8% |
| 5068 | CALCINEURIN\_MEDIATED\_SIGNALING(GO:0097720) |  | 29 | 0.66 | 1.09 | 0.141 | 0.361 | 1.000 | 17187 | tags=59%, list=29%, signal=83% |
| 5069 | POSITIVE\_REGULATION\_OF\_MEMBRANE\_PROTEIN\_ECTODOMAIN\_PROTEOLYSIS(GO:0051044) |  | 16 | 0.64 | 1.09 | 0.141 | 0.361 | 1.000 | 14284 | tags=38%, list=24%, signal=49% |
| 5070 | PIGMENTATION(GO:0043473) |  | 98 | 0.67 | 1.09 | 0.141 | 0.361 | 1.000 | 14583 | tags=42%, list=25%, signal=55% |
| 5071 | SYNAPTIC\_VESICLE\_EXOCYTOSIS(GO:0016079) |  | 80 | 0.65 | 1.09 | 0.141 | 0.361 | 1.000 | 12234 | tags=35%, list=21%, signal=44% |
| 5072 | MACROPHAGE\_MIGRATION(GO:1905517) |  | 38 | 0.67 | 1.09 | 0.141 | 0.361 | 1.000 | 9535 | tags=24%, list=16%, signal=28% |
| 5073 | NEGATIVE\_REGULATION\_OF\_MRNA\_METABOLIC\_PROCESS(GO:1903312) |  | 75 | 0.68 | 1.09 | 0.141 | 0.361 | 1.000 | 16628 | tags=55%, list=28%, signal=76% |
| 5074 | RIBONUCLEOPROTEIN\_COMPLEX\_LOCALIZATION(GO:0071166) |  | 129 | 0.62 | 1.09 | 0.000 | 0.361 | 1.000 | 20302 | tags=81%, list=34%, signal=122% |
| 5075 | SIGNAL\_TRANSDUCTION\_IN\_RESPONSE\_TO\_DNA\_DAMAGE(GO:0042770) |  | 134 | 0.66 | 1.09 | 0.141 | 0.361 | 1.000 | 16028 | tags=44%, list=27%, signal=60% |
| 5076 | POLY(A)+\_MRNA\_EXPORT\_FROM\_NUCLEUS(GO:0016973) |  | 17 | 0.65 | 1.09 | 0.000 | 0.361 | 1.000 | 19336 | tags=71%, list=33%, signal=105% |
| 5077 | POSTREPLICATION\_REPAIR(GO:0006301) |  | 53 | 0.61 | 1.09 | 0.141 | 0.361 | 1.000 | 20078 | tags=74%, list=34%, signal=111% |
| 5078 | NUCLEOBASE\_BIOSYNTHETIC\_PROCESS(GO:0046112) |  | 18 | 0.64 | 1.09 | 0.000 | 0.361 | 1.000 | 19853 | tags=83%, list=34%, signal=125% |
| 5079 | REGULATION\_OF\_CELLULAR\_EXTRAVASATION(GO:0002691) |  | 23 | 0.67 | 1.09 | 0.141 | 0.361 | 1.000 | 8784 | tags=22%, list=15%, signal=26% |
| 5080 | 3'\_5'\_EXONUCLEASE\_ACTIVITY(GO:0008408) |  | 54 | 0.62 | 1.09 | 0.000 | 0.361 | 1.000 | 21004 | tags=83%, list=36%, signal=129% |
| 5081 | REGULATION\_OF\_MONONUCLEAR\_CELL\_PROLIFERATION(GO:0032944) |  | 201 | 0.64 | 1.09 | 0.141 | 0.361 | 1.000 | 16684 | tags=37%, list=28%, signal=51% |
| 5082 | REGULATION\_OF\_LYMPHOCYTE\_PROLIFERATION(GO:0050670) |  | 200 | 0.64 | 1.09 | 0.141 | 0.361 | 1.000 | 16684 | tags=37%, list=28%, signal=51% |
| 5083 | INTERLEUKIN\_2\_PRODUCTION(GO:0032623) |  | 58 | 0.65 | 1.09 | 0.141 | 0.361 | 1.000 | 11034 | tags=31%, list=19%, signal=38% |
| 5084 | CARBOHYDRATE:PROTON\_SYMPORTER\_ACTIVITY(GO:0005351) |  | 16 | 0.62 | 1.09 | 0.141 | 0.361 | 1.000 | 14034 | tags=56%, list=24%, signal=74% |
| 5085 | REGULATION\_OF\_PROTEIN\_SUMOYLATION(GO:0033233) |  | 24 | 0.65 | 1.09 | 0.141 | 0.361 | 1.000 | 12842 | tags=33%, list=22%, signal=43% |
| 5086 | FEMALE\_GAMETE\_GENERATION(GO:0007292) |  | 132 | 0.64 | 1.09 | 0.141 | 0.361 | 1.000 | 14381 | tags=38%, list=24%, signal=50% |
| 5087 | CARBOHYDRATE:CATION\_SYMPORTER\_ACTIVITY(GO:0005402) |  | 24 | 0.62 | 1.09 | 0.141 | 0.361 | 1.000 | 16786 | tags=67%, list=28%, signal=93% |
| 5088 | PHOSPHOLIPID\_TRANSLOCATION(GO:0045332) |  | 23 | 0.68 | 1.09 | 0.141 | 0.361 | 1.000 | 13125 | tags=26%, list=22%, signal=34% |
| 5089 | REGULATION\_OF\_DNA\_DAMAGE\_CHECKPOINT(GO:2000001) |  | 17 | 0.66 | 1.09 | 0.000 | 0.361 | 1.000 | 19800 | tags=88%, list=33%, signal=133% |
| 5090 | PROTEIN\_DISULFIDE\_ISOMERASE\_ACTIVITY(GO:0003756) |  | 20 | 0.64 | 1.09 | 0.000 | 0.361 | 1.000 | 16625 | tags=60%, list=28%, signal=83% |
| 5091 | INTRAMOLECULAR\_OXIDOREDUCTASE\_ACTIVITY\_\_TRANSPOSING\_S\_S\_BONDS(GO:0016864) |  | 20 | 0.64 | 1.09 | 0.000 | 0.361 | 1.000 | 16625 | tags=60%, list=28%, signal=83% |
| 5092 | C\_TERMINAL\_PROTEIN\_LIPIDATION(GO:0006501) |  | 77 | 0.67 | 1.09 | 0.141 | 0.361 | 1.000 | 11467 | tags=29%, list=19%, signal=35% |
| 5093 | CELLULAR\_GLUCURONIDATION(GO:0052695) |  | 20 | 0.63 | 1.09 | 0.000 | 0.361 | 1.000 | 10265 | tags=15%, list=17%, signal=18% |
| 5094 | MRNA\_CONTAINING\_RIBONUCLEOPROTEIN\_COMPLEX\_EXPORT\_FROM\_NUCLEUS(GO:0071427) |  | 109 | 0.62 | 1.09 | 0.000 | 0.361 | 1.000 | 20302 | tags=83%, list=34%, signal=127% |
| 5095 | MRNA\_EXPORT\_FROM\_NUCLEUS(GO:0006406) |  | 109 | 0.62 | 1.09 | 0.000 | 0.361 | 1.000 | 20302 | tags=83%, list=34%, signal=127% |
| 5096 | REGULATION\_OF\_ICOSANOID\_SECRETION(GO:0032303) |  | 17 | 0.69 | 1.09 | 0.141 | 0.361 | 1.000 | 11372 | tags=24%, list=19%, signal=29% |
| 5097 | POSITIVE\_REGULATION\_OF\_NATURAL\_KILLER\_CELL\_MEDIATED\_CYTOTOXICITY(GO:0045954) |  | 20 | 0.65 | 1.09 | 0.141 | 0.361 | 1.000 | 15869 | tags=55%, list=27%, signal=75% |
| 5098 | MATURATION\_OF\_5.8S\_RRNA(GO:0000460) |  | 32 | 0.62 | 1.09 | 0.000 | 0.361 | 1.000 | 17908 | tags=69%, list=30%, signal=99% |
| 5099 | MALE\_GENITALIA\_DEVELOPMENT(GO:0030539) |  | 22 | 0.70 | 1.09 | 0.141 | 0.361 | 1.000 | 8613 | tags=36%, list=15%, signal=43% |
| 5100 | LYMPHOCYTE\_ACTIVATION\_INVOLVED\_IN\_IMMUNE\_RESPONSE(GO:0002285) |  | 161 | 0.64 | 1.09 | 0.141 | 0.361 | 1.000 | 18255 | tags=43%, list=31%, signal=62% |
| 5101 | DENDRITIC\_SPINE\_MAINTENANCE(GO:0097062) |  | 18 | 0.64 | 1.09 | 0.141 | 0.361 | 1.000 | 16236 | tags=39%, list=27%, signal=54% |
| 5102 | U2\_SNRNP(GO:0005686) |  | 19 | 0.64 | 1.09 | 0.000 | 0.361 | 1.000 | 21544 | tags=95%, list=36%, signal=149% |
| 5103 | DNA\_REPLICATION\_CHECKPOINT(GO:0000076) |  | 16 | 0.63 | 1.09 | 0.000 | 0.361 | 1.000 | 19870 | tags=88%, list=34%, signal=132% |
| 5104 | SINGLE\_STRANDED\_DNA\_BINDING(GO:0003697) |  | 103 | 0.66 | 1.09 | 0.141 | 0.361 | 1.000 | 17367 | tags=60%, list=29%, signal=85% |
| 5105 | GLYCINE\_METABOLIC\_PROCESS(GO:0006544) |  | 18 | 0.63 | 1.09 | 0.000 | 0.361 | 1.000 | 16959 | tags=72%, list=29%, signal=101% |
| 5106 | SPECIFIC\_GRANULE\_MEMBRANE(GO:0035579) |  | 90 | 0.62 | 1.09 | 0.141 | 0.361 | 1.000 | 16756 | tags=37%, list=28%, signal=51% |
| 5107 | CELLULAR\_RESPONSE\_TO\_DNA\_DAMAGE\_STIMULUS(GO:0006974) |  | 822 | 0.65 | 1.09 | 0.141 | 0.361 | 1.000 | 16060 | tags=44%, list=27%, signal=60% |
| 5108 | RNA\_POLYMERASE\_COMPLEX(GO:0030880) |  | 140 | 0.61 | 1.09 | 0.000 | 0.361 | 1.000 | 19664 | tags=69%, list=33%, signal=104% |
| 5109 | NEGATIVE\_REGULATION\_OF\_EMBRYONIC\_DEVELOPMENT(GO:0045992) |  | 26 | 0.70 | 1.09 | 0.141 | 0.361 | 1.000 | 10580 | tags=31%, list=18%, signal=37% |
| 5110 | REGULATION\_OF\_FEEDING\_BEHAVIOR(GO:0060259) |  | 24 | 0.68 | 1.09 | 0.141 | 0.361 | 1.000 | 14441 | tags=38%, list=24%, signal=50% |
| 5111 | NEGATIVE\_REGULATION\_OF\_INTERLEUKIN\_10\_PRODUCTION(GO:0032693) |  | 16 | 0.64 | 1.09 | 0.141 | 0.361 | 1.000 | 16562 | tags=38%, list=28%, signal=52% |
| 5112 | EXONUCLEASE\_ACTIVITY\_\_ACTIVE\_WITH\_EITHER\_RIBO\_\_OR\_DEOXYRIBONUCLEIC\_ACIDS\_AND\_PRODUCING\_5'\_PHOSPHOMONOESTERS(GO:0016796) |  | 51 | 0.63 | 1.09 | 0.000 | 0.361 | 1.000 | 19308 | tags=80%, list=33%, signal=119% |
| 5113 | RNA\_POLYMERASE\_II\_CARBOXY\_TERMINAL\_DOMAIN\_KINASE\_ACTIVITY(GO:0008353) |  | 16 | 0.64 | 1.09 | 0.000 | 0.361 | 1.000 | 18883 | tags=75%, list=32%, signal=110% |
| 5114 | POSITIVE\_REGULATION\_OF\_TRANSCRIPTION\_INITIATION\_FROM\_RNA\_POLYMERASE\_II\_PROMOTER(GO:0060261) |  | 22 | 0.69 | 1.09 | 0.141 | 0.361 | 1.000 | 16016 | tags=59%, list=27%, signal=81% |
| 5115 | CALCIUM\_DEPENDENT\_CYSTEINE\_TYPE\_ENDOPEPTIDASE\_ACTIVITY(GO:0004198) |  | 21 | 0.64 | 1.09 | 0.141 | 0.361 | 1.000 | 13706 | tags=29%, list=23%, signal=37% |
| 5116 | ALTERNATIVE\_MRNA\_SPLICING\_\_VIA\_SPLICEOSOME(GO:0000380) |  | 54 | 0.68 | 1.09 | 0.141 | 0.361 | 1.000 | 16628 | tags=70%, list=28%, signal=98% |
| 5117 | DENDRITIC\_CELL\_DIFFERENTIATION(GO:0097028) |  | 40 | 0.67 | 1.09 | 0.141 | 0.361 | 1.000 | 17938 | tags=58%, list=30%, signal=82% |
| 5118 | REGULATION\_OF\_SYNAPTIC\_VESICLE\_TRANSPORT(GO:1902803) |  | 29 | 0.64 | 1.09 | 0.000 | 0.361 | 1.000 | 17682 | tags=66%, list=30%, signal=93% |
| 5119 | C\_TERMINAL\_PROTEIN\_AMINO\_ACID\_MODIFICATION(GO:0018410) |  | 84 | 0.66 | 1.09 | 0.141 | 0.361 | 1.000 | 11467 | tags=30%, list=19%, signal=37% |
| 5120 | SPECIFIC\_GRANULE(GO:0042581) |  | 159 | 0.62 | 1.09 | 0.141 | 0.361 | 1.000 | 16756 | tags=33%, list=28%, signal=46% |
| 5121 | CERAMIDE\_METABOLIC\_PROCESS(GO:0006672) |  | 86 | 0.64 | 1.09 | 0.141 | 0.361 | 1.000 | 8876 | tags=17%, list=15%, signal=20% |
| 5122 | CYTOPLASMIC\_PATTERN\_RECOGNITION\_RECEPTOR\_SIGNALING\_PATHWAY(GO:0002753) |  | 60 | 0.62 | 1.08 | 0.141 | 0.361 | 1.000 | 17194 | tags=50%, list=29%, signal=70% |
| 5123 | UBIQUITIN\_LIGASE\_COMPLEX(GO:0000151) |  | 299 | 0.61 | 1.08 | 0.141 | 0.361 | 1.000 | 19207 | tags=61%, list=32%, signal=90% |
| 5124 | MITOTIC\_SPINDLE(GO:0072686) |  | 91 | 0.61 | 1.08 | 0.000 | 0.361 | 1.000 | 17977 | tags=64%, list=30%, signal=91% |
| 5125 | REGULATION\_OF\_TELOMERE\_MAINTENANCE\_VIA\_TELOMERE\_LENGTHENING(GO:1904356) |  | 60 | 0.62 | 1.08 | 0.000 | 0.361 | 1.000 | 20903 | tags=85%, list=35%, signal=131% |
| 5126 | REGULATION\_OF\_DEFENSE\_RESPONSE\_TO\_VIRUS\_BY\_VIRUS(GO:0050690) |  | 29 | 0.67 | 1.08 | 0.141 | 0.361 | 1.000 | 8700 | tags=21%, list=15%, signal=24% |
| 5127 | REGULATION\_OF\_T\_CELL\_MEDIATED\_CYTOTOXICITY(GO:0001914) |  | 20 | 0.63 | 1.08 | 0.141 | 0.360 | 1.000 | 16562 | tags=45%, list=28%, signal=62% |
| 5128 | UBIQUITIN\_DEPENDENT\_PROTEIN\_CATABOLIC\_PROCESS(GO:0006511) |  | 585 | 0.61 | 1.08 | 0.141 | 0.360 | 1.000 | 19019 | tags=54%, list=32%, signal=78% |
| 5129 | FEMALE\_GENITALIA\_DEVELOPMENT(GO:0030540) |  | 18 | 0.70 | 1.08 | 0.141 | 0.360 | 1.000 | 12860 | tags=33%, list=22%, signal=43% |
| 5130 | RESPONSE\_TO\_EXOGENOUS\_DSRNA(GO:0043330) |  | 47 | 0.64 | 1.08 | 0.141 | 0.360 | 1.000 | 12867 | tags=28%, list=22%, signal=35% |
| 5131 | GENE\_SILENCING(GO:0016458) |  | 444 | 0.64 | 1.08 | 0.141 | 0.360 | 1.000 | 14258 | tags=21%, list=24%, signal=27% |
| 5132 | DNA\_DIRECTED\_RNA\_POLYMERASE\_COMPLEX(GO:0000428) |  | 138 | 0.61 | 1.08 | 0.000 | 0.360 | 1.000 | 19664 | tags=69%, list=33%, signal=103% |
| 5133 | ADA2\_GCN5\_ADA3\_TRANSCRIPTION\_ACTIVATOR\_COMPLEX(GO:0005671) |  | 16 | 0.63 | 1.08 | 0.000 | 0.360 | 1.000 | 20225 | tags=88%, list=34%, signal=133% |
| 5134 | NEGATIVE\_REGULATION\_OF\_CALCIUM\_MEDIATED\_SIGNALING(GO:0050849) |  | 20 | 0.64 | 1.08 | 0.000 | 0.360 | 1.000 | 15321 | tags=50%, list=26%, signal=67% |
| 5135 | MANNOSYLATION(GO:0097502) |  | 33 | 0.61 | 1.08 | 0.141 | 0.360 | 1.000 | 15816 | tags=39%, list=27%, signal=54% |
| 5136 | T\_CELL\_SELECTION(GO:0045058) |  | 45 | 0.66 | 1.08 | 0.141 | 0.360 | 1.000 | 9023 | tags=18%, list=15%, signal=21% |
| 5137 | DNA\_SYNTHESIS\_INVOLVED\_IN\_DNA\_REPAIR(GO:0000731) |  | 76 | 0.61 | 1.08 | 0.141 | 0.360 | 1.000 | 20722 | tags=72%, list=35%, signal=111% |
| 5138 | REGULATION\_OF\_AXON\_EXTENSION(GO:0030516) |  | 76 | 0.66 | 1.08 | 0.141 | 0.360 | 1.000 | 11339 | tags=25%, list=19%, signal=31% |
| 5139 | PSEUDOURIDINE\_SYNTHESIS(GO:0001522) |  | 18 | 0.62 | 1.08 | 0.000 | 0.360 | 1.000 | 20662 | tags=89%, list=35%, signal=137% |
| 5140 | RIBONUCLEASE\_ACTIVITY(GO:0004540) |  | 105 | 0.61 | 1.08 | 0.000 | 0.360 | 1.000 | 20904 | tags=73%, list=35%, signal=113% |
| 5141 | INTRACELLULAR\_ESTROGEN\_RECEPTOR\_SIGNALING\_PATHWAY(GO:0030520) |  | 59 | 0.68 | 1.08 | 0.141 | 0.360 | 1.000 | 14716 | tags=42%, list=25%, signal=56% |
| 5142 | NEGATIVE\_REGULATION\_OF\_MITOTIC\_CELL\_CYCLE(GO:0045930) |  | 317 | 0.65 | 1.08 | 0.141 | 0.360 | 1.000 | 16045 | tags=42%, list=27%, signal=58% |
| 5143 | SPERMATID\_NUCLEUS\_DIFFERENTIATION(GO:0007289) |  | 22 | 0.65 | 1.08 | 0.000 | 0.360 | 1.000 | 18268 | tags=50%, list=31%, signal=72% |
| 5144 | NEGATIVE\_REGULATION\_OF\_MITOTIC\_CELL\_CYCLE\_PHASE\_TRANSITION(GO:1901991) |  | 237 | 0.65 | 1.08 | 0.141 | 0.360 | 1.000 | 16045 | tags=41%, list=27%, signal=57% |
| 5145 | UBIQUITIN\_DEPENDENT\_ERAD\_PATHWAY(GO:0030433) |  | 73 | 0.62 | 1.08 | 0.141 | 0.360 | 1.000 | 19239 | tags=58%, list=33%, signal=85% |
| 5146 | PH\_REDUCTION(GO:0045851) |  | 50 | 0.64 | 1.08 | 0.141 | 0.360 | 1.000 | 8241 | tags=12%, list=14%, signal=14% |
| 5147 | INOSITOL\_PHOSPHATE\_MEDIATED\_SIGNALING(GO:0048016) |  | 38 | 0.66 | 1.08 | 0.141 | 0.360 | 1.000 | 17187 | tags=53%, list=29%, signal=74% |
| 5148 | POSITIVE\_REGULATION\_OF\_CHROMOSOME\_SEGREGATION(GO:0051984) |  | 24 | 0.62 | 1.08 | 0.000 | 0.360 | 1.000 | 19621 | tags=79%, list=33%, signal=118% |
| 5149 | RESPONSE\_TO\_NITRIC\_OXIDE(GO:0071731) |  | 19 | 0.70 | 1.08 | 0.141 | 0.360 | 1.000 | 15560 | tags=53%, list=26%, signal=71% |
| 5150 | REGULATION\_OF\_FIBROBLAST\_GROWTH\_FACTOR\_RECEPTOR\_SIGNALING\_PATHWAY(GO:0040036) |  | 34 | 0.69 | 1.08 | 0.141 | 0.360 | 1.000 | 13062 | tags=32%, list=22%, signal=42% |
| 5151 | DIGESTIVE\_SYSTEM\_PROCESS(GO:0022600) |  | 93 | 0.65 | 1.08 | 0.141 | 0.360 | 1.000 | 12242 | tags=28%, list=21%, signal=35% |
| 5152 | NEGATIVE\_REGULATION\_OF\_FAT\_CELL\_DIFFERENTIATION(GO:0045599) |  | 47 | 0.65 | 1.08 | 0.141 | 0.360 | 1.000 | 15112 | tags=40%, list=26%, signal=54% |
| 5153 | NUCLEAR\_DNA\_DIRECTED\_RNA\_POLYMERASE\_COMPLEX(GO:0055029) |  | 137 | 0.61 | 1.08 | 0.000 | 0.360 | 1.000 | 19664 | tags=69%, list=33%, signal=103% |
| 5154 | MODIFICATION\_DEPENDENT\_MACROMOLECULE\_CATABOLIC\_PROCESS(GO:0043632) |  | 600 | 0.61 | 1.08 | 0.141 | 0.360 | 1.000 | 19019 | tags=54%, list=32%, signal=79% |
| 5155 | ACETYLTRANSFERASE\_ACTIVITY(GO:0016407) |  | 104 | 0.61 | 1.08 | 0.000 | 0.360 | 1.000 | 20745 | tags=76%, list=35%, signal=117% |
| 5156 | TRANSLESION\_SYNTHESIS(GO:0019985) |  | 42 | 0.61 | 1.08 | 0.141 | 0.360 | 1.000 | 19834 | tags=71%, list=34%, signal=107% |
| 5157 | G\_PROTEIN\_COUPLED\_RECEPTOR\_INTERNALIZATION(GO:0002031) |  | 16 | 0.67 | 1.08 | 0.000 | 0.360 | 1.000 | 12916 | tags=50%, list=22%, signal=64% |
| 5158 | CONDENSED\_NUCLEAR\_CHROMOSOME(GO:0000794) |  | 96 | 0.64 | 1.08 | 0.141 | 0.360 | 1.000 | 17140 | tags=55%, list=29%, signal=78% |
| 5159 | PRERIBOSOME(GO:0030684) |  | 77 | 0.60 | 1.08 | 0.000 | 0.360 | 1.000 | 19487 | tags=71%, list=33%, signal=106% |
| 5160 | MODIFICATION\_DEPENDENT\_PROTEIN\_CATABOLIC\_PROCESS(GO:0019941) |  | 591 | 0.61 | 1.08 | 0.141 | 0.360 | 1.000 | 19019 | tags=53%, list=32%, signal=78% |
| 5161 | REGULATION\_OF\_PROTEIN\_POLYUBIQUITINATION(GO:1902914) |  | 17 | 0.65 | 1.08 | 0.141 | 0.360 | 1.000 | 16993 | tags=65%, list=29%, signal=91% |
| 5162 | RNA\_CATABOLIC\_PROCESS(GO:0006401) |  | 391 | 0.65 | 1.08 | 0.141 | 0.360 | 1.000 | 17054 | tags=47%, list=29%, signal=66% |
| 5163 | TOLL\_LIKE\_RECEPTOR\_3\_SIGNALING\_PATHWAY(GO:0034138) |  | 19 | 0.65 | 1.08 | 0.141 | 0.360 | 1.000 | 13678 | tags=26%, list=23%, signal=34% |
| 5164 | EXIT\_FROM\_MITOSIS(GO:0010458) |  | 27 | 0.62 | 1.08 | 0.000 | 0.360 | 1.000 | 20702 | tags=81%, list=35%, signal=125% |
| 5165 | NUCLEAR\_TRANSCRIBED\_MRNA\_CATABOLIC\_PROCESS\_\_EXONUCLEOLYTIC(GO:0000291) |  | 35 | 0.60 | 1.08 | 0.000 | 0.360 | 1.000 | 21067 | tags=80%, list=36%, signal=124% |
| 5166 | PYRIMIDINE\_NUCLEOBASE\_METABOLIC\_PROCESS(GO:0006206) |  | 19 | 0.63 | 1.08 | 0.141 | 0.360 | 1.000 | 17343 | tags=68%, list=29%, signal=97% |
| 5167 | RETINOL\_DEHYDROGENASE\_ACTIVITY(GO:0004745) |  | 18 | 0.66 | 1.08 | 0.141 | 0.360 | 1.000 | 17257 | tags=44%, list=29%, signal=63% |
| 5168 | TOLERANCE\_INDUCTION(GO:0002507) |  | 25 | 0.61 | 1.08 | 0.000 | 0.360 | 1.000 | 17552 | tags=44%, list=30%, signal=63% |
| 5169 | NUCLEOSIDE\_DIPHOSPHATE\_KINASE\_ACTIVITY(GO:0004550) |  | 18 | 0.62 | 1.08 | 0.141 | 0.360 | 1.000 | 17121 | tags=61%, list=29%, signal=86% |
| 5170 | LYMPHOCYTE\_MEDIATED\_IMMUNITY(GO:0002449) |  | 317 | 0.62 | 1.08 | 0.141 | 0.360 | 1.000 | 15869 | tags=25%, list=27%, signal=34% |
| 5171 | PROTEIN\_IMPORT\_INTO\_MITOCHONDRIAL\_MATRIX(GO:0030150) |  | 17 | 0.62 | 1.08 | 0.000 | 0.360 | 1.000 | 19156 | tags=76%, list=32%, signal=113% |
| 5172 | PLACENTA\_BLOOD\_VESSEL\_DEVELOPMENT(GO:0060674) |  | 33 | 0.66 | 1.08 | 0.141 | 0.360 | 1.000 | 13970 | tags=30%, list=24%, signal=40% |
| 5173 | FLUID\_TRANSPORT(GO:0042044) |  | 27 | 0.68 | 1.08 | 0.141 | 0.360 | 1.000 | 10857 | tags=33%, list=18%, signal=41% |
| 5174 | DNA\_INTEGRITY\_CHECKPOINT(GO:0031570) |  | 164 | 0.66 | 1.08 | 0.141 | 0.360 | 1.000 | 16045 | tags=52%, list=27%, signal=72% |
| 5175 | T\_CELL\_LINEAGE\_COMMITMENT(GO:0002360) |  | 25 | 0.65 | 1.08 | 0.141 | 0.360 | 1.000 | 10412 | tags=28%, list=18%, signal=34% |
| 5176 | MITOCHONDRION\_MORPHOGENESIS(GO:0070584) |  | 21 | 0.61 | 1.08 | 0.141 | 0.360 | 1.000 | 20910 | tags=76%, list=35%, signal=118% |
| 5177 | G0\_TO\_G1\_TRANSITION(GO:0045023) |  | 47 | 0.63 | 1.08 | 0.000 | 0.360 | 1.000 | 20745 | tags=79%, list=35%, signal=121% |
| 5178 | DNA\_DAMAGE\_CHECKPOINT(GO:0000077) |  | 155 | 0.66 | 1.08 | 0.141 | 0.360 | 1.000 | 16045 | tags=52%, list=27%, signal=71% |
| 5179 | MITOCHONDRIAL\_SMALL\_RIBOSOMAL\_SUBUNIT(GO:0005763) |  | 28 | 0.62 | 1.08 | 0.000 | 0.360 | 1.000 | 21925 | tags=89%, list=37%, signal=142% |
| 5180 | ORGANELLAR\_SMALL\_RIBOSOMAL\_SUBUNIT(GO:0000314) |  | 28 | 0.62 | 1.08 | 0.000 | 0.360 | 1.000 | 21925 | tags=89%, list=37%, signal=142% |
| 5181 | CELLULAR\_COMPONENT\_DISASSEMBLY(GO:0022411) |  | 558 | 0.63 | 1.08 | 0.141 | 0.360 | 1.000 | 18098 | tags=52%, list=31%, signal=74% |
| 5182 | TRANSCRIPTION\_COUPLED\_NUCLEOTIDE\_EXCISION\_REPAIR(GO:0006283) |  | 73 | 0.60 | 1.08 | 0.000 | 0.360 | 1.000 | 20582 | tags=81%, list=35%, signal=124% |
| 5183 | TELOMERE\_MAINTENANCE\_VIA\_TELOMERE\_LENGTHENING(GO:0010833) |  | 79 | 0.61 | 1.08 | 0.000 | 0.360 | 1.000 | 20903 | tags=81%, list=35%, signal=125% |
| 5184 | CELLULAR\_RESPONSE\_TO\_SUPEROXIDE(GO:0071451) |  | 27 | 0.65 | 1.08 | 0.141 | 0.360 | 1.000 | 15716 | tags=37%, list=27%, signal=50% |
| 5185 | CELLULAR\_RESPONSE\_TO\_OXYGEN\_RADICAL(GO:0071450) |  | 27 | 0.65 | 1.08 | 0.141 | 0.360 | 1.000 | 15716 | tags=37%, list=27%, signal=50% |
| 5186 | MRNA\_CATABOLIC\_PROCESS(GO:0006402) |  | 358 | 0.65 | 1.08 | 0.141 | 0.360 | 1.000 | 17054 | tags=46%, list=29%, signal=64% |
| 5187 | RESPONSE\_TO\_UV\_B(GO:0010224) |  | 17 | 0.63 | 1.08 | 0.307 | 0.360 | 1.000 | 6738 | tags=18%, list=11%, signal=20% |
| 5188 | RESPONSE\_TO\_COLD(GO:0009409) |  | 49 | 0.64 | 1.08 | 0.141 | 0.360 | 1.000 | 17432 | tags=51%, list=29%, signal=72% |
| 5189 | ENERGY\_DERIVATION\_BY\_OXIDATION\_OF\_ORGANIC\_COMPOUNDS(GO:0015980) |  | 284 | 0.62 | 1.08 | 0.141 | 0.360 | 1.000 | 20261 | tags=62%, list=34%, signal=94% |
| 5190 | INTERFERON\_GAMMA\_BIOSYNTHETIC\_PROCESS(GO:0042095) |  | 16 | 0.66 | 1.08 | 0.141 | 0.360 | 1.000 | 12443 | tags=25%, list=21%, signal=32% |
| 5191 | N\_ACETYLTRANSFERASE\_ACTIVITY(GO:0008080) |  | 86 | 0.61 | 1.08 | 0.141 | 0.360 | 1.000 | 20745 | tags=78%, list=35%, signal=120% |
| 5192 | INDOLALKYLAMINE\_METABOLIC\_PROCESS(GO:0006586) |  | 17 | 0.68 | 1.08 | 0.141 | 0.360 | 1.000 | 18727 | tags=59%, list=32%, signal=86% |
| 5193 | UBIQUITIN\_PROTEIN\_LIGASE\_BINDING(GO:0031625) |  | 300 | 0.66 | 1.08 | 0.141 | 0.360 | 1.000 | 11102 | tags=22%, list=19%, signal=27% |
| 5194 | ACTIVATION\_OF\_JUN\_KINASE\_ACTIVITY(GO:0007257) |  | 35 | 0.67 | 1.08 | 0.141 | 0.360 | 1.000 | 7538 | tags=11%, list=13%, signal=13% |
| 5195 | RNA\_DEPENDENT\_DNA\_BIOSYNTHETIC\_PROCESS(GO:0006278) |  | 69 | 0.62 | 1.08 | 0.000 | 0.360 | 1.000 | 20903 | tags=86%, list=35%, signal=132% |
| 5196 | ALPHA\_BETA\_T\_CELL\_LINEAGE\_COMMITMENT(GO:0002363) |  | 18 | 0.64 | 1.08 | 0.307 | 0.360 | 1.000 | 9023 | tags=22%, list=15%, signal=26% |
| 5197 | N\_ACYLTRANSFERASE\_ACTIVITY(GO:0016410) |  | 104 | 0.61 | 1.08 | 0.141 | 0.360 | 1.000 | 20745 | tags=74%, list=35%, signal=114% |
| 5198 | PHOSPHOLIPID\_TRANSPORT(GO:0015914) |  | 64 | 0.66 | 1.08 | 0.141 | 0.360 | 1.000 | 16755 | tags=42%, list=28%, signal=59% |
| 5199 | MANNOSYLTRANSFERASE\_ACTIVITY(GO:0000030) |  | 24 | 0.61 | 1.08 | 0.141 | 0.360 | 1.000 | 13469 | tags=21%, list=23%, signal=27% |
| 5200 | NUCLEAR\_PORE(GO:0005643) |  | 76 | 0.62 | 1.08 | 0.000 | 0.360 | 1.000 | 20253 | tags=82%, list=34%, signal=124% |
| 5201 | SPLICEOSOMAL\_COMPLEX\_ASSEMBLY(GO:0000245) |  | 58 | 0.62 | 1.08 | 0.000 | 0.360 | 1.000 | 21524 | tags=91%, list=36%, signal=144% |
| 5202 | PROTEIN\_IMPORT\_INTO\_NUCLEUS\_\_TRANSLOCATION(GO:0000060) |  | 46 | 0.64 | 1.08 | 0.141 | 0.360 | 1.000 | 17938 | tags=52%, list=30%, signal=75% |
| 5203 | NUCLEOPLASM\_PART(GO:0044451) |  | 1103 | 0.65 | 1.08 | 0.141 | 0.360 | 1.000 | 15212 | tags=42%, list=26%, signal=55% |
| 5204 | DEFENSE\_RESPONSE\_TO\_GRAM\_NEGATIVE\_BACTERIUM(GO:0050829) |  | 82 | 0.63 | 1.08 | 0.141 | 0.360 | 1.000 | 14996 | tags=17%, list=25%, signal=23% |
| 5205 | TRNA\_MODIFICATION(GO:0006400) |  | 74 | 0.60 | 1.08 | 0.000 | 0.360 | 1.000 | 19689 | tags=76%, list=33%, signal=113% |
| 5206 | NF\_KAPPAB\_BINDING(GO:0051059) |  | 28 | 0.63 | 1.08 | 0.141 | 0.360 | 1.000 | 12137 | tags=29%, list=21%, signal=36% |
| 5207 | POSITIVE\_REGULATION\_OF\_IMMUNE\_EFFECTOR\_PROCESS(GO:0002699) |  | 171 | 0.63 | 1.08 | 0.141 | 0.360 | 1.000 | 14247 | tags=32%, list=24%, signal=41% |
| 5208 | CELLULAR\_POLYSACCHARIDE\_BIOSYNTHETIC\_PROCESS(GO:0033692) |  | 64 | 0.65 | 1.08 | 0.141 | 0.360 | 1.000 | 9277 | tags=19%, list=16%, signal=22% |
| 5209 | NAD\_BIOSYNTHETIC\_PROCESS(GO:0009435) |  | 54 | 0.65 | 1.08 | 0.141 | 0.360 | 1.000 | 6424 | tags=6%, list=11%, signal=6% |
| 5210 | GANGLIOSIDE\_METABOLIC\_PROCESS(GO:0001573) |  | 18 | 0.67 | 1.08 | 0.141 | 0.360 | 1.000 | 13970 | tags=33%, list=24%, signal=44% |
| 5211 | ENTRY\_INTO\_OTHER\_ORGANISM\_INVOLVED\_IN\_SYMBIOTIC\_INTERACTION(GO:0051828) |  | 137 | 0.64 | 1.08 | 0.141 | 0.360 | 1.000 | 10550 | tags=16%, list=18%, signal=20% |
| 5212 | ENTRY\_INTO\_HOST\_CELL(GO:0030260) |  | 137 | 0.64 | 1.08 | 0.141 | 0.360 | 1.000 | 10550 | tags=16%, list=18%, signal=20% |
| 5213 | ENTRY\_INTO\_CELL\_OF\_OTHER\_ORGANISM\_INVOLVED\_IN\_SYMBIOTIC\_INTERACTION(GO:0051806) |  | 137 | 0.64 | 1.08 | 0.141 | 0.360 | 1.000 | 10550 | tags=16%, list=18%, signal=20% |
| 5214 | ENTRY\_INTO\_HOST(GO:0044409) |  | 137 | 0.64 | 1.08 | 0.141 | 0.359 | 1.000 | 10550 | tags=16%, list=18%, signal=20% |
| 5215 | RESPONSE\_TO\_CADMIUM\_ION(GO:0046686) |  | 58 | 0.61 | 1.08 | 0.141 | 0.359 | 1.000 | 19731 | tags=60%, list=33%, signal=90% |
| 5216 | REGULATION\_OF\_INTERFERON\_GAMMA\_BIOSYNTHETIC\_PROCESS(GO:0045072) |  | 15 | 0.67 | 1.08 | 0.141 | 0.359 | 1.000 | 12443 | tags=27%, list=21%, signal=34% |
| 5217 | SPINDLE\_MICROTUBULE(GO:0005876) |  | 58 | 0.61 | 1.08 | 0.000 | 0.359 | 1.000 | 20032 | tags=78%, list=34%, signal=117% |
| 5218 | POSITIVE\_REGULATION\_OF\_CELL\_ADHESION\_MEDIATED\_BY\_INTEGRIN(GO:0033630) |  | 17 | 0.67 | 1.08 | 0.141 | 0.359 | 1.000 | 13355 | tags=29%, list=23%, signal=38% |
| 5219 | PHOSPHATASE\_REGULATOR\_ACTIVITY(GO:0019208) |  | 92 | 0.62 | 1.08 | 0.141 | 0.359 | 1.000 | 16246 | tags=40%, list=27%, signal=55% |
| 5220 | NITROGEN\_CYCLE\_METABOLIC\_PROCESS(GO:0071941) |  | 15 | 0.65 | 1.08 | 0.307 | 0.359 | 1.000 | 13132 | tags=40%, list=22%, signal=51% |
| 5221 | REGULATION\_OF\_DNA\_BIOSYNTHETIC\_PROCESS(GO:2000278) |  | 102 | 0.67 | 1.08 | 0.141 | 0.359 | 1.000 | 14468 | tags=49%, list=24%, signal=65% |
| 5222 | MRNA\_DESTABILIZATION(GO:0061157) |  | 22 | 0.64 | 1.08 | 0.141 | 0.359 | 1.000 | 16375 | tags=64%, list=28%, signal=88% |
| 5223 | H4\_HISTONE\_ACETYLTRANSFERASE\_ACTIVITY(GO:0010485) |  | 17 | 0.62 | 1.08 | 0.141 | 0.359 | 1.000 | 20665 | tags=88%, list=35%, signal=136% |
| 5224 | REGULATION\_OF\_ACTIVATED\_T\_CELL\_PROLIFERATION(GO:0046006) |  | 38 | 0.61 | 1.08 | 0.141 | 0.360 | 1.000 | 16562 | tags=42%, list=28%, signal=58% |
| 5225 | SMALL\_NUCLEOLAR\_RIBONUCLEOPROTEIN\_COMPLEX(GO:0005732) |  | 23 | 0.62 | 1.08 | 0.000 | 0.360 | 1.000 | 21511 | tags=96%, list=36%, signal=150% |
| 5226 | RNA\_MODIFICATION(GO:0009451) |  | 136 | 0.60 | 1.08 | 0.000 | 0.360 | 1.000 | 20901 | tags=80%, list=35%, signal=124% |
| 5227 | MICROTUBULE\_ANCHORING(GO:0034453) |  | 24 | 0.61 | 1.08 | 0.000 | 0.360 | 1.000 | 21311 | tags=83%, list=36%, signal=130% |
| 5228 | REGULATION\_OF\_CENTRIOLE\_REPLICATION(GO:0046599) |  | 17 | 0.67 | 1.08 | 0.141 | 0.360 | 1.000 | 19564 | tags=88%, list=33%, signal=132% |
| 5229 | MITOTIC\_CHROMOSOME\_CONDENSATION(GO:0007076) |  | 15 | 0.65 | 1.08 | 0.000 | 0.360 | 1.000 | 20190 | tags=93%, list=34%, signal=142% |
| 5230 | PEROXISOMAL\_MEMBRANE\_TRANSPORT(GO:0015919) |  | 16 | 0.61 | 1.08 | 0.141 | 0.360 | 1.000 | 19319 | tags=69%, list=33%, signal=102% |
| 5231 | DNA\_DIRECTED\_RNA\_POLYMERASE\_III\_COMPLEX(GO:0005666) |  | 18 | 0.61 | 1.08 | 0.141 | 0.360 | 1.000 | 22286 | tags=94%, list=38%, signal=152% |
| 5232 | COENZYME\_BINDING(GO:0050662) |  | 276 | 0.64 | 1.08 | 0.141 | 0.360 | 1.000 | 11786 | tags=24%, list=20%, signal=30% |
| 5233 | HYDROLASE\_ACTIVITY\_\_ACTING\_ON\_CARBON\_NITROGEN\_(BUT\_NOT\_PEPTIDE)\_BONDS\_\_IN\_LINEAR\_AMIDES(GO:0016811) |  | 90 | 0.61 | 1.08 | 0.141 | 0.359 | 1.000 | 19846 | tags=59%, list=34%, signal=88% |
| 5234 | POSITIVE\_REGULATION\_OF\_CARDIOCYTE\_DIFFERENTIATION(GO:1905209) |  | 37 | 0.64 | 1.08 | 0.141 | 0.359 | 1.000 | 17343 | tags=38%, list=29%, signal=54% |
| 5235 | INTRACELLULAR\_PH\_REDUCTION(GO:0051452) |  | 49 | 0.64 | 1.08 | 0.141 | 0.359 | 1.000 | 8241 | tags=12%, list=14%, signal=14% |
| 5236 | NEGATIVE\_REGULATION\_OF\_CELL\_MATRIX\_ADHESION(GO:0001953) |  | 37 | 0.67 | 1.08 | 0.141 | 0.359 | 1.000 | 14735 | tags=38%, list=25%, signal=50% |
| 5237 | REGULATION\_OF\_SISTER\_CHROMATID\_COHESION(GO:0007063) |  | 21 | 0.64 | 1.08 | 0.000 | 0.359 | 1.000 | 20576 | tags=95%, list=35%, signal=146% |
| 5238 | MYD88\_DEPENDENT\_TOLL\_LIKE\_RECEPTOR\_SIGNALING\_PATHWAY(GO:0002755) |  | 37 | 0.62 | 1.08 | 0.141 | 0.359 | 1.000 | 19834 | tags=51%, list=34%, signal=77% |
| 5239 | RESPONSE\_TO\_LEPTIN(GO:0044321) |  | 21 | 0.68 | 1.08 | 0.141 | 0.359 | 1.000 | 13935 | tags=38%, list=24%, signal=50% |
| 5240 | LEUKOCYTE\_MEDIATED\_CYTOTOXICITY(GO:0001909) |  | 85 | 0.62 | 1.08 | 0.141 | 0.359 | 1.000 | 15869 | tags=39%, list=27%, signal=53% |
| 5241 | CELL\_CYCLE\_CHECKPOINT(GO:0000075) |  | 221 | 0.66 | 1.08 | 0.141 | 0.359 | 1.000 | 16045 | tags=51%, list=27%, signal=70% |
| 5242 | ARACHIDONIC\_ACID\_MONOOXYGENASE\_ACTIVITY(GO:0008391) |  | 17 | 0.62 | 1.08 | 0.141 | 0.359 | 1.000 | 11264 | tags=29%, list=19%, signal=36% |
| 5243 | ARACHIDONIC\_ACID\_EPOXYGENASE\_ACTIVITY(GO:0008392) |  | 17 | 0.62 | 1.08 | 0.141 | 0.359 | 1.000 | 11264 | tags=29%, list=19%, signal=36% |
| 5244 | RNA\_PROCESSING(GO:0006396) |  | 956 | 0.62 | 1.08 | 0.141 | 0.359 | 1.000 | 19278 | tags=67%, list=33%, signal=97% |
| 5245 | RNA\_PHOSPHODIESTER\_BOND\_HYDROLYSIS(GO:0090501) |  | 141 | 0.61 | 1.08 | 0.000 | 0.359 | 1.000 | 21004 | tags=74%, list=36%, signal=115% |
| 5246 | H4\_HISTONE\_ACETYLTRANSFERASE\_COMPLEX(GO:1902562) |  | 37 | 0.61 | 1.08 | 0.000 | 0.359 | 1.000 | 20745 | tags=86%, list=35%, signal=133% |
| 5247 | RNA\_CAP\_BINDING(GO:0000339) |  | 17 | 0.63 | 1.08 | 0.000 | 0.359 | 1.000 | 20904 | tags=94%, list=35%, signal=146% |
| 5248 | DICARBOXYLIC\_ACID\_METABOLIC\_PROCESS(GO:0043648) |  | 102 | 0.64 | 1.08 | 0.141 | 0.359 | 1.000 | 15869 | tags=41%, list=27%, signal=56% |
| 5249 | MYELOID\_DENDRITIC\_CELL\_ACTIVATION(GO:0001773) |  | 27 | 0.67 | 1.08 | 0.141 | 0.359 | 1.000 | 17938 | tags=56%, list=30%, signal=80% |
| 5250 | RETROGRADE\_TRANSPORT\_\_ENDOSOME\_TO\_GOLGI(GO:0042147) |  | 80 | 0.60 | 1.08 | 0.141 | 0.359 | 1.000 | 19192 | tags=61%, list=32%, signal=91% |
| 5251 | REPRESSING\_TRANSCRIPTION\_FACTOR\_BINDING(GO:0070491) |  | 60 | 0.62 | 1.08 | 0.000 | 0.359 | 1.000 | 18262 | tags=72%, list=31%, signal=104% |
| 5252 | RNA\_SURVEILLANCE(GO:0071025) |  | 16 | 0.61 | 1.08 | 0.000 | 0.359 | 1.000 | 20709 | tags=75%, list=35%, signal=115% |
| 5253 | REGULATION\_OF\_MEIOTIC\_CELL\_CYCLE(GO:0051445) |  | 44 | 0.65 | 1.08 | 0.141 | 0.359 | 1.000 | 11130 | tags=41%, list=19%, signal=50% |
| 5254 | POSITIVE\_REGULATION\_OF\_STRIATED\_MUSCLE\_CELL\_APOPTOTIC\_PROCESS(GO:0010663) |  | 16 | 0.67 | 1.08 | 0.168 | 0.359 | 1.000 | 19603 | tags=63%, list=33%, signal=93% |
| 5255 | POSITIVE\_REGULATION\_OF\_CARDIAC\_MUSCLE\_CELL\_APOPTOTIC\_PROCESS(GO:0010666) |  | 16 | 0.67 | 1.08 | 0.168 | 0.359 | 1.000 | 19603 | tags=63%, list=33%, signal=93% |
| 5256 | ONE\_CARBON\_METABOLIC\_PROCESS(GO:0006730) |  | 35 | 0.62 | 1.08 | 0.141 | 0.359 | 1.000 | 19306 | tags=57%, list=33%, signal=85% |
| 5257 | NEGATIVE\_REGULATION\_OF\_G0\_TO\_G1\_TRANSITION(GO:0070317) |  | 40 | 0.63 | 1.08 | 0.000 | 0.359 | 1.000 | 19453 | tags=73%, list=33%, signal=108% |
| 5258 | ACETYL\_COA\_BIOSYNTHETIC\_PROCESS\_FROM\_PYRUVATE(GO:0006086) |  | 15 | 0.65 | 1.08 | 0.000 | 0.359 | 1.000 | 17637 | tags=80%, list=30%, signal=114% |
| 5259 | PROTEIN\_O\_LINKED\_GLYCOSYLATION(GO:0006493) |  | 107 | 0.64 | 1.08 | 0.141 | 0.359 | 1.000 | 15853 | tags=42%, list=27%, signal=57% |
| 5260 | FUCOSE\_METABOLIC\_PROCESS(GO:0006004) |  | 16 | 0.69 | 1.08 | 0.141 | 0.359 | 1.000 | 9252 | tags=38%, list=16%, signal=44% |
| 5261 | POSTSYNAPTIC\_NEUROTRANSMITTER\_RECEPTOR\_ACTIVITY(GO:0098960) |  | 22 | 0.66 | 1.08 | 0.141 | 0.359 | 1.000 | 13742 | tags=50%, list=23%, signal=65% |
| 5262 | BONE\_CELL\_DEVELOPMENT(GO:0098751) |  | 32 | 0.67 | 1.08 | 0.141 | 0.359 | 1.000 | 10768 | tags=28%, list=18%, signal=34% |
| 5263 | REGULATION\_OF\_TELOMERE\_CAPPING(GO:1904353) |  | 26 | 0.62 | 1.08 | 0.000 | 0.359 | 1.000 | 20903 | tags=85%, list=35%, signal=131% |
| 5264 | TRANSCYTOSIS(GO:0045056) |  | 17 | 0.66 | 1.08 | 0.141 | 0.359 | 1.000 | 12188 | tags=29%, list=21%, signal=37% |
| 5265 | TELOMERE\_MAINTENANCE(GO:0000723) |  | 158 | 0.61 | 1.08 | 0.141 | 0.359 | 1.000 | 20903 | tags=77%, list=35%, signal=119% |
| 5266 | PSEUDOPODIUM\_ORGANIZATION(GO:0031268) |  | 17 | 0.67 | 1.08 | 0.141 | 0.359 | 1.000 | 15036 | tags=53%, list=25%, signal=71% |
| 5267 | U2\_TYPE\_PRESPLICEOSOME(GO:0071004) |  | 17 | 0.62 | 1.08 | 0.000 | 0.359 | 1.000 | 21067 | tags=88%, list=36%, signal=137% |
| 5268 | EPOXYGENASE\_P450\_PATHWAY(GO:0019373) |  | 21 | 0.63 | 1.08 | 0.141 | 0.359 | 1.000 | 11580 | tags=29%, list=20%, signal=36% |
| 5269 | NEGATIVE\_REGULATION\_OF\_TELOMERE\_MAINTENANCE(GO:0032205) |  | 36 | 0.61 | 1.08 | 0.000 | 0.359 | 1.000 | 20841 | tags=83%, list=35%, signal=129% |
| 5270 | PROTEIN\_TRANSMEMBRANE\_TRANSPORTER\_ACTIVITY(GO:0008320) |  | 17 | 0.60 | 1.08 | 0.000 | 0.359 | 1.000 | 22167 | tags=76%, list=37%, signal=122% |
| 5271 | NEGATIVE\_REGULATION\_OF\_STRIATED\_MUSCLE\_CELL\_DIFFERENTIATION(GO:0051154) |  | 37 | 0.65 | 1.08 | 0.141 | 0.359 | 1.000 | 11569 | tags=30%, list=20%, signal=37% |
| 5272 | REGULATION\_OF\_CELL\_SUBSTRATE\_JUNCTION\_ASSEMBLY(GO:0090109) |  | 54 | 0.66 | 1.08 | 0.141 | 0.359 | 1.000 | 14843 | tags=33%, list=25%, signal=44% |
| 5273 | REGULATION\_OF\_FOCAL\_ADHESION\_ASSEMBLY(GO:0051893) |  | 54 | 0.66 | 1.08 | 0.141 | 0.359 | 1.000 | 14843 | tags=33%, list=25%, signal=44% |
| 5274 | HUMORAL\_IMMUNE\_RESPONSE(GO:0006959) |  | 337 | 0.63 | 1.08 | 0.141 | 0.359 | 1.000 | 12189 | tags=15%, list=21%, signal=19% |
| 5275 | CARBOHYDRATE\_DERIVATIVE\_TRANSMEMBRANE\_TRANSPORTER\_ACTIVITY(GO:1901505) |  | 35 | 0.63 | 1.08 | 0.141 | 0.359 | 1.000 | 13528 | tags=29%, list=23%, signal=37% |
| 5276 | DIGESTION(GO:0007586) |  | 129 | 0.65 | 1.08 | 0.141 | 0.359 | 1.000 | 12242 | tags=26%, list=21%, signal=33% |
| 5277 | PIGMENT\_BIOSYNTHETIC\_PROCESS(GO:0046148) |  | 54 | 0.65 | 1.08 | 0.141 | 0.359 | 1.000 | 14962 | tags=50%, list=25%, signal=67% |
| 5278 | TRNA\_BINDING(GO:0000049) |  | 55 | 0.60 | 1.08 | 0.000 | 0.359 | 1.000 | 21344 | tags=85%, list=36%, signal=134% |
| 5279 | NEGATIVE\_REGULATION\_OF\_CELL\_CYCLE\_PHASE\_TRANSITION(GO:1901988) |  | 257 | 0.65 | 1.08 | 0.141 | 0.359 | 1.000 | 16045 | tags=41%, list=27%, signal=56% |
| 5280 | CLEAVAGE\_FURROW(GO:0032154) |  | 54 | 0.66 | 1.08 | 0.141 | 0.359 | 1.000 | 16994 | tags=48%, list=29%, signal=68% |
| 5281 | NEGATIVE\_REGULATION\_OF\_INTERLEUKIN\_8\_PRODUCTION(GO:0032717) |  | 17 | 0.66 | 1.08 | 0.141 | 0.359 | 1.000 | 6413 | tags=6%, list=11%, signal=7% |
| 5282 | CYTOPLASMIC\_MRNA\_PROCESSING\_BODY\_ASSEMBLY(GO:0033962) |  | 20 | 0.63 | 1.07 | 0.141 | 0.359 | 1.000 | 20042 | tags=90%, list=34%, signal=136% |
| 5283 | AMIDE\_BIOSYNTHETIC\_PROCESS(GO:0043604) |  | 1005 | 0.63 | 1.07 | 0.141 | 0.359 | 1.000 | 18259 | tags=44%, list=31%, signal=63% |
| 5284 | DEOXYRIBONUCLEOTIDE\_METABOLIC\_PROCESS(GO:0009262) |  | 34 | 0.60 | 1.07 | 0.141 | 0.359 | 1.000 | 23049 | tags=94%, list=39%, signal=154% |
| 5285 | RNA\_METHYLTRANSFERASE\_ACTIVITY(GO:0008173) |  | 60 | 0.60 | 1.07 | 0.000 | 0.359 | 1.000 | 19689 | tags=75%, list=33%, signal=112% |
| 5286 | PHOSPHATIDYLINOSITOL\_3\_4\_BISPHOSPHATE\_BINDING(GO:0043325) |  | 24 | 0.63 | 1.07 | 0.141 | 0.359 | 1.000 | 14826 | tags=46%, list=25%, signal=61% |
| 5287 | TELOMERE\_MAINTENANCE\_VIA\_TELOMERE\_TRIMMING(GO:0090737) |  | 15 | 0.63 | 1.07 | 0.000 | 0.359 | 1.000 | 17367 | tags=73%, list=29%, signal=104% |
| 5288 | T\_CIRCLE\_FORMATION(GO:0090656) |  | 15 | 0.63 | 1.07 | 0.000 | 0.359 | 1.000 | 17367 | tags=73%, list=29%, signal=104% |
| 5289 | FORMATION\_OF\_EXTRACHROMOSOMAL\_CIRCULAR\_DNA(GO:0001325) |  | 15 | 0.63 | 1.07 | 0.000 | 0.359 | 1.000 | 17367 | tags=73%, list=29%, signal=104% |
| 5290 | REGULATION\_OF\_REGULATORY\_T\_CELL\_DIFFERENTIATION(GO:0045589) |  | 29 | 0.68 | 1.07 | 0.141 | 0.359 | 1.000 | 6178 | tags=3%, list=10%, signal=4% |
| 5291 | CHOLESTEROL\_STORAGE(GO:0010878) |  | 15 | 0.66 | 1.07 | 0.141 | 0.359 | 1.000 | 11522 | tags=27%, list=19%, signal=33% |
| 5292 | CATALYTIC\_ACTIVITY\_\_ACTING\_ON\_A\_RRNA(GO:0140102) |  | 21 | 0.62 | 1.07 | 0.000 | 0.359 | 1.000 | 19157 | tags=76%, list=32%, signal=113% |
| 5293 | RRNA\_METHYLTRANSFERASE\_ACTIVITY(GO:0008649) |  | 21 | 0.62 | 1.07 | 0.000 | 0.359 | 1.000 | 19157 | tags=76%, list=32%, signal=113% |
| 5294 | POSITIVE\_REGULATION\_OF\_MYELOID\_CELL\_DIFFERENTIATION(GO:0045639) |  | 91 | 0.62 | 1.07 | 0.141 | 0.359 | 1.000 | 16246 | tags=45%, list=27%, signal=62% |
| 5295 | POSITIVE\_REGULATION\_OF\_MRNA\_METABOLIC\_PROCESS(GO:1903313) |  | 65 | 0.61 | 1.07 | 0.141 | 0.359 | 1.000 | 18871 | tags=63%, list=32%, signal=93% |
| 5296 | TRANSLATIONAL\_INITIATION(GO:0006413) |  | 198 | 0.61 | 1.07 | 0.141 | 0.359 | 1.000 | 19174 | tags=58%, list=32%, signal=85% |
| 5297 | ENDORIBONUCLEASE\_ACTIVITY\_\_PRODUCING\_5'\_PHOSPHOMONOESTERS(GO:0016891) |  | 33 | 0.64 | 1.07 | 0.000 | 0.359 | 1.000 | 20821 | tags=88%, list=35%, signal=136% |
| 5298 | CHROMOSOME\_SEPARATION(GO:0051304) |  | 78 | 0.65 | 1.07 | 0.141 | 0.359 | 1.000 | 15966 | tags=60%, list=27%, signal=82% |
| 5299 | RIBONUCLEOPROTEIN\_GRANULE(GO:0035770) |  | 199 | 0.64 | 1.07 | 0.141 | 0.359 | 1.000 | 18344 | tags=60%, list=31%, signal=87% |
| 5300 | ACID\_AMINO\_ACID\_LIGASE\_ACTIVITY(GO:0016881) |  | 24 | 0.61 | 1.07 | 0.141 | 0.359 | 1.000 | 21071 | tags=79%, list=36%, signal=123% |
| 5301 | UBIQUITIN\_LIKE\_PROTEIN\_LIGASE\_BINDING(GO:0044389) |  | 314 | 0.65 | 1.07 | 0.141 | 0.359 | 1.000 | 11102 | tags=21%, list=19%, signal=25% |
| 5302 | NEGATIVE\_REGULATION\_OF\_REPRODUCTIVE\_PROCESS(GO:2000242) |  | 56 | 0.65 | 1.07 | 0.141 | 0.359 | 1.000 | 11130 | tags=36%, list=19%, signal=44% |
| 5303 | CHEMOSENSORY\_BEHAVIOR(GO:0007635) |  | 16 | 0.67 | 1.07 | 0.141 | 0.359 | 1.000 | 9447 | tags=31%, list=16%, signal=37% |
| 5304 | REGULATORY\_T\_CELL\_DIFFERENTIATION(GO:0045066) |  | 31 | 0.67 | 1.07 | 0.141 | 0.359 | 1.000 | 6178 | tags=6%, list=10%, signal=7% |
| 5305 | CYTOPLASMIC\_RIBONUCLEOPROTEIN\_GRANULE(GO:0036464) |  | 188 | 0.64 | 1.07 | 0.141 | 0.359 | 1.000 | 18344 | tags=60%, list=31%, signal=86% |
| 5306 | REGULATION\_OF\_PLASMA\_MEMBRANE\_BOUNDED\_CELL\_PROJECTION\_ASSEMBLY(GO:0120032) |  | 157 | 0.65 | 1.07 | 0.141 | 0.359 | 1.000 | 15397 | tags=39%, list=26%, signal=53% |
| 5307 | ESC\_E(Z)\_COMPLEX(GO:0035098) |  | 18 | 0.65 | 1.07 | 0.000 | 0.359 | 1.000 | 16954 | tags=61%, list=29%, signal=86% |
| 5308 | RNA\_SPLICING\_\_VIA\_TRANSESTERIFICATION\_REACTIONS(GO:0000375) |  | 330 | 0.61 | 1.07 | 0.141 | 0.359 | 1.000 | 18535 | tags=68%, list=31%, signal=99% |
| 5309 | PEPTIDE\_BIOSYNTHETIC\_PROCESS(GO:0043043) |  | 926 | 0.63 | 1.07 | 0.141 | 0.359 | 1.000 | 18259 | tags=44%, list=31%, signal=62% |
| 5310 | RESPONSE\_TO\_IMMOBILIZATION\_STRESS(GO:0035902) |  | 25 | 0.69 | 1.07 | 0.141 | 0.359 | 1.000 | 10387 | tags=24%, list=18%, signal=29% |
| 5311 | ISOPRENOID\_BINDING(GO:0019840) |  | 36 | 0.67 | 1.07 | 0.141 | 0.359 | 1.000 | 11950 | tags=31%, list=20%, signal=38% |
| 5312 | GLUTAMATE\_RECEPTOR\_BINDING(GO:0035254) |  | 41 | 0.62 | 1.07 | 0.141 | 0.359 | 1.000 | 15321 | tags=44%, list=26%, signal=59% |
| 5313 | 90S\_PRERIBOSOME(GO:0030686) |  | 31 | 0.60 | 1.07 | 0.000 | 0.359 | 1.000 | 19119 | tags=71%, list=32%, signal=105% |
| 5314 | HISTONE\_DEUBIQUITINATION(GO:0016578) |  | 24 | 0.64 | 1.07 | 0.000 | 0.359 | 1.000 | 20091 | tags=88%, list=34%, signal=132% |
| 5315 | REGULATION\_OF\_SPINDLE\_ASSEMBLY(GO:0090169) |  | 25 | 0.61 | 1.07 | 0.000 | 0.359 | 1.000 | 15444 | tags=48%, list=26%, signal=65% |
| 5316 | MAGNESIUM\_ION\_TRANSMEMBRANE\_TRANSPORT(GO:1903830) |  | 17 | 0.65 | 1.07 | 0.000 | 0.359 | 1.000 | 18680 | tags=76%, list=32%, signal=112% |
| 5317 | MAGNESIUM\_ION\_TRANSMEMBRANE\_TRANSPORTER\_ACTIVITY(GO:0015095) |  | 17 | 0.65 | 1.07 | 0.000 | 0.359 | 1.000 | 18680 | tags=76%, list=32%, signal=112% |
| 5318 | REGULATION\_OF\_TELOMERASE\_RNA\_LOCALIZATION\_TO\_CAJAL\_BODY(GO:1904872) |  | 18 | 0.63 | 1.07 | 0.000 | 0.359 | 1.000 | 20650 | tags=94%, list=35%, signal=145% |
| 5319 | DETECTION\_OF\_CHEMICAL\_STIMULUS\_INVOLVED\_IN\_SENSORY\_PERCEPTION\_OF\_BITTER\_TASTE(GO:0001580) |  | 36 | 0.65 | 1.07 | 0.141 | 0.359 | 1.000 | 9774 | tags=11%, list=17%, signal=13% |
| 5320 | PIRNA\_METABOLIC\_PROCESS(GO:0034587) |  | 18 | 0.66 | 1.07 | 0.141 | 0.359 | 1.000 | 17636 | tags=61%, list=30%, signal=87% |
| 5321 | POSITIVE\_REGULATION\_OF\_DNA\_BIOSYNTHETIC\_PROCESS(GO:2000573) |  | 65 | 0.67 | 1.07 | 0.141 | 0.359 | 1.000 | 16887 | tags=69%, list=29%, signal=97% |
| 5322 | MICROTUBULE\_PLUS\_END(GO:0035371) |  | 19 | 0.65 | 1.07 | 0.000 | 0.359 | 1.000 | 15497 | tags=47%, list=26%, signal=64% |
| 5323 | DETECTION\_OF\_VISIBLE\_LIGHT(GO:0009584) |  | 58 | 0.62 | 1.07 | 0.141 | 0.359 | 1.000 | 15194 | tags=38%, list=26%, signal=51% |
| 5324 | TRNA\_SPECIFIC\_RIBONUCLEASE\_ACTIVITY(GO:0004549) |  | 17 | 0.63 | 1.07 | 0.000 | 0.359 | 1.000 | 20821 | tags=76%, list=35%, signal=118% |
| 5325 | BRANCHED\_CHAIN\_AMINO\_ACID\_METABOLIC\_PROCESS(GO:0009081) |  | 24 | 0.66 | 1.07 | 0.141 | 0.359 | 1.000 | 17081 | tags=58%, list=29%, signal=82% |
| 5326 | PRONUCLEUS(GO:0045120) |  | 18 | 0.63 | 1.07 | 0.141 | 0.359 | 1.000 | 19782 | tags=67%, list=33%, signal=100% |
| 5327 | RNA\_SPLICING\_\_VIA\_TRANSESTERIFICATION\_REACTIONS\_WITH\_BULGED\_ADENOSINE\_AS\_NUCLEOPHILE(GO:0000377) |  | 327 | 0.61 | 1.07 | 0.141 | 0.359 | 1.000 | 18535 | tags=68%, list=31%, signal=99% |
| 5328 | MRNA\_SPLICING\_\_VIA\_SPLICEOSOME(GO:0000398) |  | 327 | 0.61 | 1.07 | 0.141 | 0.359 | 1.000 | 18535 | tags=68%, list=31%, signal=99% |
| 5329 | TELOMERASE\_RNA\_LOCALIZATION\_TO\_CAJAL\_BODY(GO:0090671) |  | 19 | 0.63 | 1.07 | 0.000 | 0.359 | 1.000 | 20650 | tags=95%, list=35%, signal=146% |
| 5330 | RNA\_LOCALIZATION\_TO\_CAJAL\_BODY(GO:0090670) |  | 19 | 0.63 | 1.07 | 0.000 | 0.359 | 1.000 | 20650 | tags=95%, list=35%, signal=146% |
| 5331 | TELOMERASE\_RNA\_LOCALIZATION(GO:0090672) |  | 19 | 0.63 | 1.07 | 0.000 | 0.359 | 1.000 | 20650 | tags=95%, list=35%, signal=146% |
| 5332 | RNA\_LOCALIZATION\_TO\_NUCLEUS(GO:0090685) |  | 19 | 0.63 | 1.07 | 0.000 | 0.359 | 1.000 | 20650 | tags=95%, list=35%, signal=146% |
| 5333 | EXORIBONUCLEASE\_COMPLEX(GO:1905354) |  | 23 | 0.62 | 1.07 | 0.000 | 0.359 | 1.000 | 19561 | tags=83%, list=33%, signal=123% |
| 5334 | EXOSOME\_(RNASE\_COMPLEX)(GO:0000178) |  | 23 | 0.62 | 1.07 | 0.000 | 0.359 | 1.000 | 19561 | tags=83%, list=33%, signal=123% |
| 5335 | TERTIARY\_GRANULE\_MEMBRANE(GO:0070821) |  | 73 | 0.61 | 1.07 | 0.141 | 0.359 | 1.000 | 8769 | tags=11%, list=15%, signal=13% |
| 5336 | POSITIVE\_REGULATION\_OF\_MITOTIC\_NUCLEAR\_DIVISION(GO:0045840) |  | 50 | 0.64 | 1.07 | 0.141 | 0.359 | 1.000 | 17160 | tags=56%, list=29%, signal=79% |
| 5337 | U2\_TYPE\_SPLICEOSOMAL\_COMPLEX(GO:0005684) |  | 60 | 0.61 | 1.07 | 0.000 | 0.359 | 1.000 | 21067 | tags=82%, list=36%, signal=127% |
| 5338 | POSITIVE\_REGULATION\_OF\_MICROTUBULE\_POLYMERIZATION\_OR\_DEPOLYMERIZATION(GO:0031112) |  | 30 | 0.60 | 1.07 | 0.141 | 0.359 | 1.000 | 20000 | tags=63%, list=34%, signal=96% |
| 5339 | NEGATIVE\_REGULATION\_OF\_BIOMINERAL\_TISSUE\_DEVELOPMENT(GO:0070168) |  | 18 | 0.66 | 1.07 | 0.310 | 0.358 | 1.000 | 18904 | tags=67%, list=32%, signal=98% |
| 5340 | NEUTRAL\_LIPID\_BIOSYNTHETIC\_PROCESS(GO:0046460) |  | 36 | 0.64 | 1.07 | 0.141 | 0.359 | 1.000 | 12925 | tags=39%, list=22%, signal=50% |
| 5341 | ACYLGLYCEROL\_BIOSYNTHETIC\_PROCESS(GO:0046463) |  | 36 | 0.64 | 1.07 | 0.141 | 0.358 | 1.000 | 12925 | tags=39%, list=22%, signal=50% |
| 5342 | ATPASE\_REGULATOR\_ACTIVITY(GO:0060590) |  | 36 | 0.62 | 1.07 | 0.168 | 0.358 | 1.000 | 16634 | tags=58%, list=28%, signal=81% |
| 5343 | U2\_TYPE\_CATALYTIC\_STEP\_2\_SPLICEOSOME(GO:0071007) |  | 29 | 0.62 | 1.07 | 0.000 | 0.358 | 1.000 | 20916 | tags=90%, list=35%, signal=139% |
| 5344 | REGULATION\_OF\_CELL\_PROJECTION\_ASSEMBLY(GO:0060491) |  | 159 | 0.65 | 1.07 | 0.141 | 0.358 | 1.000 | 15397 | tags=39%, list=26%, signal=53% |
| 5345 | MITOCHONDRIAL\_DNA\_METABOLIC\_PROCESS(GO:0032042) |  | 18 | 0.60 | 1.07 | 0.000 | 0.358 | 1.000 | 23468 | tags=83%, list=40%, signal=138% |
| 5346 | POSITIVE\_REGULATION\_OF\_SMOOTH\_MUSCLE\_CELL\_MIGRATION(GO:0014911) |  | 42 | 0.65 | 1.07 | 0.141 | 0.358 | 1.000 | 15941 | tags=43%, list=27%, signal=59% |
| 5347 | REGULATION\_OF\_FILOPODIUM\_ASSEMBLY(GO:0051489) |  | 43 | 0.66 | 1.07 | 0.141 | 0.358 | 1.000 | 15385 | tags=44%, list=26%, signal=60% |
| 5348 | OOGENESIS(GO:0048477) |  | 83 | 0.64 | 1.07 | 0.141 | 0.358 | 1.000 | 15922 | tags=48%, list=27%, signal=66% |
| 5349 | ENDONUCLEOLYTIC\_CLEAVAGE\_INVOLVED\_IN\_RRNA\_PROCESSING(GO:0000478) |  | 15 | 0.63 | 1.07 | 0.000 | 0.358 | 1.000 | 17908 | tags=73%, list=30%, signal=105% |
| 5350 | INNER\_EAR\_AUDITORY\_RECEPTOR\_CELL\_DIFFERENTIATION(GO:0042491) |  | 33 | 0.63 | 1.07 | 0.141 | 0.358 | 1.000 | 18101 | tags=52%, list=31%, signal=74% |
| 5351 | RNA\_5'\_END\_PROCESSING(GO:0000966) |  | 21 | 0.64 | 1.07 | 0.000 | 0.358 | 1.000 | 20821 | tags=90%, list=35%, signal=140% |
| 5352 | RNA\_METHYLATION(GO:0001510) |  | 65 | 0.60 | 1.07 | 0.000 | 0.358 | 1.000 | 20377 | tags=77%, list=34%, signal=117% |
| 5353 | TRNA\_METHYLATION(GO:0030488) |  | 31 | 0.61 | 1.07 | 0.000 | 0.358 | 1.000 | 19689 | tags=74%, list=33%, signal=111% |
| 5354 | COP9\_SIGNALOSOME(GO:0008180) |  | 35 | 0.63 | 1.07 | 0.141 | 0.358 | 1.000 | 20672 | tags=66%, list=35%, signal=101% |
| 5355 | RESPONSE\_TO\_FUNGUS(GO:0009620) |  | 53 | 0.66 | 1.07 | 0.141 | 0.358 | 1.000 | 10014 | tags=13%, list=17%, signal=16% |
| 5356 | SMN\_SM\_PROTEIN\_COMPLEX(GO:0034719) |  | 17 | 0.66 | 1.07 | 0.000 | 0.358 | 1.000 | 20168 | tags=100%, list=34%, signal=152% |
| 5357 | TRANSFERASE\_ACTIVITY\_\_TRANSFERRING\_PENTOSYL\_GROUPS(GO:0016763) |  | 59 | 0.64 | 1.07 | 0.141 | 0.358 | 1.000 | 16374 | tags=47%, list=28%, signal=66% |
| 5358 | POSITIVE\_REGULATION\_OF\_MRNA\_CATABOLIC\_PROCESS(GO:0061014) |  | 39 | 0.63 | 1.07 | 0.141 | 0.358 | 1.000 | 18170 | tags=64%, list=31%, signal=92% |
| 5359 | NEGATIVE\_REGULATION\_OF\_CYCLIN\_DEPENDENT\_PROTEIN\_KINASE\_ACTIVITY(GO:1904030) |  | 32 | 0.63 | 1.07 | 0.141 | 0.358 | 1.000 | 13695 | tags=34%, list=23%, signal=45% |
| 5360 | POSITIVE\_REGULATION\_OF\_EMBRYONIC\_DEVELOPMENT(GO:0040019) |  | 38 | 0.65 | 1.07 | 0.141 | 0.358 | 1.000 | 18222 | tags=50%, list=31%, signal=72% |
| 5361 | POSITIVE\_REGULATION\_OF\_EXTRINSIC\_APOPTOTIC\_SIGNALING\_PATHWAY(GO:2001238) |  | 50 | 0.66 | 1.07 | 0.141 | 0.358 | 1.000 | 15681 | tags=38%, list=27%, signal=52% |
| 5362 | RIG\_I\_SIGNALING\_PATHWAY(GO:0039529) |  | 19 | 0.62 | 1.07 | 0.141 | 0.358 | 1.000 | 14841 | tags=37%, list=25%, signal=49% |
| 5363 | REGULATION\_OF\_GRANULOCYTE\_DIFFERENTIATION(GO:0030852) |  | 16 | 0.67 | 1.07 | 0.141 | 0.358 | 1.000 | 6381 | tags=13%, list=11%, signal=14% |
| 5364 | TRNA\_METHYLTRANSFERASE\_ACTIVITY(GO:0008175) |  | 28 | 0.62 | 1.07 | 0.000 | 0.358 | 1.000 | 20377 | tags=82%, list=34%, signal=125% |
| 5365 | SYNAPTONEMAL\_COMPLEX\_ASSEMBLY(GO:0007130) |  | 21 | 0.62 | 1.07 | 0.141 | 0.358 | 1.000 | 18181 | tags=52%, list=31%, signal=76% |
| 5366 | LIPOPROTEIN\_TRANSPORT(GO:0042953) |  | 20 | 0.64 | 1.07 | 0.000 | 0.358 | 1.000 | 12268 | tags=35%, list=21%, signal=44% |
| 5367 | LIPOPROTEIN\_LOCALIZATION(GO:0044872) |  | 20 | 0.64 | 1.07 | 0.000 | 0.358 | 1.000 | 12268 | tags=35%, list=21%, signal=44% |
| 5368 | ENDOPLASMIC\_RETICULUM\_UNFOLDED\_PROTEIN\_RESPONSE(GO:0030968) |  | 129 | 0.62 | 1.07 | 0.141 | 0.358 | 1.000 | 7261 | tags=5%, list=12%, signal=6% |
| 5369 | IMMUNOGLOBULIN\_MEDIATED\_IMMUNE\_RESPONSE(GO:0016064) |  | 203 | 0.62 | 1.07 | 0.141 | 0.358 | 1.000 | 15749 | tags=19%, list=27%, signal=25% |
| 5370 | TERMINATION\_OF\_RNA\_POLYMERASE\_II\_TRANSCRIPTION(GO:0006369) |  | 68 | 0.61 | 1.07 | 0.000 | 0.358 | 1.000 | 21791 | tags=91%, list=37%, signal=144% |
| 5371 | GLIAL\_CELL\_PROLIFERATION(GO:0014009) |  | 38 | 0.64 | 1.07 | 0.141 | 0.359 | 1.000 | 18003 | tags=45%, list=30%, signal=64% |
| 5372 | REGULATION\_OF\_TOLERANCE\_INDUCTION(GO:0002643) |  | 17 | 0.64 | 1.07 | 0.168 | 0.359 | 1.000 | 19789 | tags=53%, list=33%, signal=80% |
| 5373 | CALCIUM\_ION\_REGULATED\_EXOCYTOSIS(GO:0017156) |  | 105 | 0.63 | 1.07 | 0.141 | 0.359 | 1.000 | 14932 | tags=38%, list=25%, signal=51% |
| 5374 | POSITIVE\_REGULATION\_OF\_STRIATED\_MUSCLE\_CONTRACTION(GO:0045989) |  | 18 | 0.65 | 1.07 | 0.476 | 0.359 | 1.000 | 16226 | tags=44%, list=27%, signal=61% |
| 5375 | SPHINGOLIPID\_METABOLIC\_PROCESS(GO:0006665) |  | 164 | 0.63 | 1.07 | 0.141 | 0.359 | 1.000 | 8876 | tags=16%, list=15%, signal=19% |
| 5376 | WNT\_SIGNALING\_PATHWAY\_\_CALCIUM\_MODULATING\_PATHWAY(GO:0007223) |  | 38 | 0.63 | 1.07 | 0.141 | 0.359 | 1.000 | 18198 | tags=61%, list=31%, signal=87% |
| 5377 | TBP\_CLASS\_PROTEIN\_BINDING(GO:0017025) |  | 25 | 0.65 | 1.07 | 0.141 | 0.359 | 1.000 | 17312 | tags=64%, list=29%, signal=90% |
| 5378 | NEGATIVE\_REGULATION\_OF\_MRNA\_PROCESSING(GO:0050686) |  | 30 | 0.62 | 1.07 | 0.000 | 0.359 | 1.000 | 19278 | tags=77%, list=33%, signal=114% |
| 5379 | ATPASE\_ACTIVITY\_\_COUPLED\_TO\_TRANSMEMBRANE\_MOVEMENT\_OF\_SUBSTANCES(GO:0042626) |  | 116 | 0.63 | 1.07 | 0.141 | 0.359 | 1.000 | 15839 | tags=43%, list=27%, signal=59% |
| 5380 | PRENYLTRANSFERASE\_ACTIVITY(GO:0004659) |  | 17 | 0.61 | 1.07 | 0.000 | 0.359 | 1.000 | 22597 | tags=94%, list=38%, signal=152% |
| 5381 | REGULATION\_OF\_MRNA\_CATABOLIC\_PROCESS(GO:0061013) |  | 181 | 0.64 | 1.07 | 0.141 | 0.359 | 1.000 | 16931 | tags=47%, list=29%, signal=66% |
| 5382 | TRANSCRIPTIONAL\_REPRESSOR\_COMPLEX(GO:0017053) |  | 83 | 0.61 | 1.07 | 0.141 | 0.359 | 1.000 | 21576 | tags=80%, list=36%, signal=125% |
| 5383 | CALCIUM\_RELEASE\_CHANNEL\_ACTIVITY(GO:0015278) |  | 18 | 0.64 | 1.07 | 0.141 | 0.359 | 1.000 | 15094 | tags=50%, list=26%, signal=67% |
| 5384 | CELL\_DIVISION\_SITE\_PART(GO:0032155) |  | 63 | 0.65 | 1.07 | 0.141 | 0.359 | 1.000 | 16994 | tags=49%, list=29%, signal=69% |
| 5385 | CELL\_DIVISION\_SITE(GO:0032153) |  | 63 | 0.65 | 1.07 | 0.141 | 0.359 | 1.000 | 16994 | tags=49%, list=29%, signal=69% |
| 5386 | U1\_SNRNP(GO:0005685) |  | 20 | 0.64 | 1.07 | 0.000 | 0.359 | 1.000 | 20168 | tags=80%, list=34%, signal=121% |
| 5387 | RESPONSE\_TO\_OXYGEN\_RADICAL(GO:0000305) |  | 33 | 0.63 | 1.07 | 0.141 | 0.359 | 1.000 | 18635 | tags=52%, list=32%, signal=75% |
| 5388 | PIGMENT\_METABOLIC\_PROCESS(GO:0042440) |  | 69 | 0.64 | 1.07 | 0.141 | 0.359 | 1.000 | 14962 | tags=42%, list=25%, signal=56% |
| 5389 | PROTEIN\_BINDING\_INVOLVED\_IN\_PROTEIN\_FOLDING(GO:0044183) |  | 27 | 0.63 | 1.07 | 0.141 | 0.359 | 1.000 | 21410 | tags=85%, list=36%, signal=133% |
| 5390 | BRUSH\_BORDER(GO:0005903) |  | 103 | 0.63 | 1.07 | 0.141 | 0.359 | 1.000 | 12090 | tags=25%, list=20%, signal=32% |
| 5391 | STEROID\_CATABOLIC\_PROCESS(GO:0006706) |  | 24 | 0.63 | 1.07 | 0.141 | 0.359 | 1.000 | 11197 | tags=29%, list=19%, signal=36% |
| 5392 | NUCLEAR\_PORE\_ORGANIZATION(GO:0006999) |  | 15 | 0.67 | 1.07 | 0.340 | 0.359 | 1.000 | 17845 | tags=93%, list=30%, signal=134% |
| 5393 | 'DE\_NOVO'\_PROTEIN\_FOLDING(GO:0006458) |  | 39 | 0.61 | 1.07 | 0.000 | 0.359 | 1.000 | 22131 | tags=90%, list=37%, signal=143% |
| 5394 | RESPONSE\_TO\_DEXAMETHASONE(GO:0071548) |  | 41 | 0.64 | 1.07 | 0.141 | 0.359 | 1.000 | 13486 | tags=32%, list=23%, signal=41% |
| 5395 | REGULATION\_OF\_MRNA\_METABOLIC\_PROCESS(GO:1903311) |  | 282 | 0.64 | 1.07 | 0.141 | 0.359 | 1.000 | 16724 | tags=50%, list=28%, signal=69% |
| 5396 | CATALYTIC\_ACTIVITY\_\_ACTING\_ON\_A\_TRNA(GO:0140101) |  | 129 | 0.59 | 1.07 | 0.141 | 0.359 | 1.000 | 22024 | tags=82%, list=37%, signal=131% |
| 5397 | POSITIVE\_REGULATION\_OF\_GLUCOSE\_IMPORT\_IN\_RESPONSE\_TO\_INSULIN\_STIMULUS(GO:2001275) |  | 18 | 0.65 | 1.07 | 0.141 | 0.359 | 1.000 | 18963 | tags=56%, list=32%, signal=82% |
| 5398 | RIBOSOME\_BINDING(GO:0043022) |  | 56 | 0.59 | 1.07 | 0.141 | 0.359 | 1.000 | 22772 | tags=88%, list=39%, signal=142% |
| 5399 | PROTEIN\_TYROSINE\_SERINE\_THREONINE\_PHOSPHATASE\_ACTIVITY(GO:0008138) |  | 51 | 0.62 | 1.07 | 0.141 | 0.359 | 1.000 | 19041 | tags=55%, list=32%, signal=81% |
| 5400 | PROTEIN\_K63\_LINKED\_UBIQUITINATION(GO:0070534) |  | 45 | 0.64 | 1.07 | 0.141 | 0.359 | 1.000 | 17768 | tags=56%, list=30%, signal=79% |
| 5401 | REGULATION\_OF\_RNA\_SPLICING(GO:0043484) |  | 115 | 0.66 | 1.07 | 0.141 | 0.359 | 1.000 | 17789 | tags=65%, list=30%, signal=93% |
| 5402 | UBIQUITIN\_LIKE\_PROTEIN\_BINDING(GO:0032182) |  | 82 | 0.60 | 1.07 | 0.141 | 0.359 | 1.000 | 18224 | tags=55%, list=31%, signal=79% |
| 5403 | RAC\_PROTEIN\_SIGNAL\_TRANSDUCTION(GO:0016601) |  | 33 | 0.62 | 1.07 | 0.000 | 0.359 | 1.000 | 17777 | tags=58%, list=30%, signal=82% |
| 5404 | B\_CELL\_MEDIATED\_IMMUNITY(GO:0019724) |  | 204 | 0.62 | 1.07 | 0.141 | 0.359 | 1.000 | 15749 | tags=19%, list=27%, signal=25% |
| 5405 | PSEUDOPODIUM\_ASSEMBLY(GO:0031269) |  | 16 | 0.67 | 1.07 | 0.141 | 0.359 | 1.000 | 15036 | tags=56%, list=25%, signal=75% |
| 5406 | REGULATION\_OF\_STEROID\_BIOSYNTHETIC\_PROCESS(GO:0050810) |  | 82 | 0.61 | 1.07 | 0.141 | 0.359 | 1.000 | 14948 | tags=37%, list=25%, signal=49% |
| 5407 | POSITIVE\_REGULATION\_OF\_NATURAL\_KILLER\_CELL\_MEDIATED\_IMMUNITY(GO:0002717) |  | 22 | 0.62 | 1.07 | 0.141 | 0.359 | 1.000 | 15869 | tags=50%, list=27%, signal=68% |
| 5408 | NUCLEOBASE\_CONTAINING\_COMPOUND\_TRANSMEMBRANE\_TRANSPORTER\_ACTIVITY(GO:0015932) |  | 39 | 0.62 | 1.07 | 0.141 | 0.359 | 1.000 | 15294 | tags=36%, list=26%, signal=48% |
| 5409 | TRNA\_METABOLIC\_PROCESS(GO:0006399) |  | 194 | 0.60 | 1.07 | 0.000 | 0.359 | 1.000 | 22267 | tags=84%, list=38%, signal=134% |
| 5410 | REGULATION\_OF\_ADHERENS\_JUNCTION\_ORGANIZATION(GO:1903391) |  | 57 | 0.65 | 1.07 | 0.141 | 0.359 | 1.000 | 14843 | tags=33%, list=25%, signal=44% |
| 5411 | GRANULOCYTE\_DIFFERENTIATION(GO:0030851) |  | 32 | 0.65 | 1.07 | 0.141 | 0.359 | 1.000 | 8946 | tags=19%, list=15%, signal=22% |
| 5412 | SEX\_CHROMOSOME(GO:0000803) |  | 29 | 0.64 | 1.07 | 0.168 | 0.359 | 1.000 | 20893 | tags=86%, list=35%, signal=133% |
| 5413 | OUTER\_DYNEIN\_ARM\_ASSEMBLY(GO:0036158) |  | 17 | 0.65 | 1.07 | 0.141 | 0.359 | 1.000 | 15188 | tags=47%, list=26%, signal=63% |
| 5414 | DNA\_DOUBLE\_STRAND\_BREAK\_PROCESSING(GO:0000729) |  | 24 | 0.64 | 1.07 | 0.340 | 0.359 | 1.000 | 14064 | tags=58%, list=24%, signal=77% |
| 5415 | RIBONUCLEOPROTEIN\_COMPLEX(GO:1990904) |  | 783 | 0.63 | 1.07 | 0.141 | 0.359 | 1.000 | 17983 | tags=57%, list=30%, signal=81% |
| 5416 | REGULATION\_OF\_CARDIAC\_MUSCLE\_ADAPTATION(GO:0010612) |  | 18 | 0.63 | 1.07 | 0.141 | 0.359 | 1.000 | 10309 | tags=17%, list=17%, signal=20% |
| 5417 | REGULATION\_OF\_CARDIAC\_MUSCLE\_HYPERTROPHY\_IN\_RESPONSE\_TO\_STRESS(GO:1903242) |  | 18 | 0.63 | 1.07 | 0.141 | 0.359 | 1.000 | 10309 | tags=17%, list=17%, signal=20% |
| 5418 | REGULATION\_OF\_STEM\_CELL\_DIFFERENTIATION(GO:2000736) |  | 143 | 0.63 | 1.07 | 0.141 | 0.359 | 1.000 | 15400 | tags=31%, list=26%, signal=42% |
| 5419 | THYROID\_HORMONE\_METABOLIC\_PROCESS(GO:0042403) |  | 21 | 0.65 | 1.07 | 0.141 | 0.359 | 1.000 | 12505 | tags=33%, list=21%, signal=42% |
| 5420 | NEGATIVE\_REGULATION\_OF\_CELL\_CYCLE\_ARREST(GO:0071157) |  | 23 | 0.61 | 1.07 | 0.141 | 0.359 | 1.000 | 16188 | tags=39%, list=27%, signal=54% |
| 5421 | CATALYTIC\_STEP\_2\_SPLICEOSOME(GO:0071013) |  | 96 | 0.60 | 1.07 | 0.141 | 0.359 | 1.000 | 21067 | tags=80%, list=36%, signal=124% |
| 5422 | RRNA\_MODIFICATION(GO:0000154) |  | 37 | 0.60 | 1.07 | 0.000 | 0.359 | 1.000 | 20860 | tags=81%, list=35%, signal=125% |
| 5423 | NEGATIVE\_REGULATION\_OF\_CELL\_CYCLE\_PROCESS(GO:0010948) |  | 351 | 0.64 | 1.07 | 0.141 | 0.359 | 1.000 | 16969 | tags=47%, list=29%, signal=66% |
| 5424 | NAD+\_ADP\_RIBOSYLTRANSFERASE\_ACTIVITY(GO:0003950) |  | 25 | 0.65 | 1.07 | 0.141 | 0.359 | 1.000 | 16139 | tags=48%, list=27%, signal=66% |
| 5425 | PTERIDINE\_CONTAINING\_COMPOUND\_METABOLIC\_PROCESS(GO:0042558) |  | 34 | 0.61 | 1.07 | 0.141 | 0.359 | 1.000 | 17149 | tags=53%, list=29%, signal=75% |
| 5426 | SPINDLE\_MIDZONE(GO:0051233) |  | 31 | 0.61 | 1.07 | 0.000 | 0.359 | 1.000 | 21647 | tags=77%, list=37%, signal=122% |
| 5427 | NUCLEOTIDE\_EXCISION\_REPAIR(GO:0006289) |  | 117 | 0.59 | 1.07 | 0.141 | 0.359 | 1.000 | 22441 | tags=86%, list=38%, signal=139% |
| 5428 | PERICENTRIC\_HETEROCHROMATIN(GO:0005721) |  | 20 | 0.60 | 1.07 | 0.141 | 0.359 | 1.000 | 22650 | tags=90%, list=38%, signal=146% |
| 5429 | NEGATIVE\_REGULATION\_OF\_MRNA\_SPLICING\_\_VIA\_SPLICEOSOME(GO:0048025) |  | 20 | 0.63 | 1.07 | 0.000 | 0.359 | 1.000 | 19278 | tags=85%, list=33%, signal=126% |
| 5430 | RRNA\_CATABOLIC\_PROCESS(GO:0016075) |  | 18 | 0.62 | 1.07 | 0.000 | 0.359 | 1.000 | 21004 | tags=83%, list=36%, signal=129% |
| 5431 | PRESPLICEOSOME(GO:0071010) |  | 21 | 0.61 | 1.07 | 0.141 | 0.359 | 1.000 | 19278 | tags=81%, list=33%, signal=120% |
| 5432 | ORGANELLE\_LOCALIZATION\_BY\_MEMBRANE\_TETHERING(GO:0140056) |  | 161 | 0.60 | 1.07 | 0.141 | 0.359 | 1.000 | 19348 | tags=68%, list=33%, signal=100% |
| 5433 | PHOSPHATASE\_INHIBITOR\_ACTIVITY(GO:0019212) |  | 41 | 0.60 | 1.06 | 0.141 | 0.359 | 1.000 | 22011 | tags=68%, list=37%, signal=109% |
| 5434 | REGULATION\_OF\_PROTEIN\_UBIQUITINATION(GO:0031396) |  | 184 | 0.63 | 1.06 | 0.141 | 0.359 | 1.000 | 17110 | tags=50%, list=29%, signal=70% |
| 5435 | POSITIVE\_REGULATION\_OF\_SIGNAL\_TRANSDUCTION\_BY\_P53\_CLASS\_MEDIATOR(GO:1901798) |  | 20 | 0.62 | 1.06 | 0.141 | 0.359 | 1.000 | 15865 | tags=50%, list=27%, signal=68% |
| 5436 | 2'\_DEOXYRIBONUCLEOTIDE\_METABOLIC\_PROCESS(GO:0009394) |  | 29 | 0.60 | 1.06 | 0.141 | 0.359 | 1.000 | 23049 | tags=93%, list=39%, signal=152% |
| 5437 | DNA\_DUPLEX\_UNWINDING(GO:0032508) |  | 74 | 0.59 | 1.06 | 0.000 | 0.358 | 1.000 | 22305 | tags=86%, list=38%, signal=139% |
| 5438 | DEOXYRIBOSE\_PHOSPHATE\_METABOLIC\_PROCESS(GO:0019692) |  | 31 | 0.60 | 1.06 | 0.141 | 0.358 | 1.000 | 23049 | tags=94%, list=39%, signal=153% |
| 5439 | ENDOSOMAL\_TRANSPORT(GO:0016197) |  | 191 | 0.64 | 1.06 | 0.141 | 0.358 | 1.000 | 8252 | tags=7%, list=14%, signal=8% |
| 5440 | TYPE\_I\_INTERFERON\_SIGNALING\_PATHWAY(GO:0060337) |  | 85 | 0.63 | 1.06 | 0.141 | 0.358 | 1.000 | 8063 | tags=13%, list=14%, signal=15% |
| 5441 | CELLULAR\_RESPONSE\_TO\_TYPE\_I\_INTERFERON(GO:0071357) |  | 85 | 0.63 | 1.06 | 0.141 | 0.358 | 1.000 | 8063 | tags=13%, list=14%, signal=15% |
| 5442 | REGULATION\_OF\_GASTRULATION(GO:0010470) |  | 43 | 0.64 | 1.06 | 0.141 | 0.359 | 1.000 | 19002 | tags=51%, list=32%, signal=75% |
| 5443 | LIVER\_DEVELOPMENT(GO:0001889) |  | 138 | 0.61 | 1.06 | 0.141 | 0.359 | 1.000 | 17154 | tags=51%, list=29%, signal=72% |
| 5444 | CHROMOSOME\_LOCALIZATION\_TO\_NUCLEAR\_ENVELOPE\_INVOLVED\_IN\_HOMOLOGOUS\_CHROMOSOME\_SEGREGATION(GO:0090220) |  | 15 | 0.68 | 1.06 | 0.141 | 0.359 | 1.000 | 14781 | tags=60%, list=25%, signal=80% |
| 5445 | TELOMERE\_LOCALIZATION(GO:0034397) |  | 16 | 0.69 | 1.06 | 0.141 | 0.359 | 1.000 | 14781 | tags=63%, list=25%, signal=83% |
| 5446 | P\_P\_BOND\_HYDROLYSIS\_DRIVEN\_TRANSMEMBRANE\_TRANSPORTER\_ACTIVITY(GO:0015405) |  | 124 | 0.63 | 1.06 | 0.141 | 0.359 | 1.000 | 15839 | tags=44%, list=27%, signal=60% |
| 5447 | PRIMARY\_ACTIVE\_TRANSMEMBRANE\_TRANSPORTER\_ACTIVITY(GO:0015399) |  | 124 | 0.63 | 1.06 | 0.141 | 0.358 | 1.000 | 15839 | tags=44%, list=27%, signal=60% |
| 5448 | NEGATIVE\_REGULATION\_OF\_LIPID\_LOCALIZATION(GO:1905953) |  | 49 | 0.63 | 1.06 | 0.141 | 0.358 | 1.000 | 8474 | tags=16%, list=14%, signal=19% |
| 5449 | NUCLEOTIDE\_EXCISION\_REPAIR\_\_PREINCISION\_COMPLEX\_ASSEMBLY(GO:0006294) |  | 29 | 0.61 | 1.06 | 0.000 | 0.359 | 1.000 | 18883 | tags=69%, list=32%, signal=101% |
| 5450 | MITOCHONDRIAL\_GENE\_EXPRESSION(GO:0140053) |  | 148 | 0.60 | 1.06 | 0.141 | 0.359 | 1.000 | 21335 | tags=87%, list=36%, signal=136% |
| 5451 | REGULATION\_OF\_METALLOENDOPEPTIDASE\_ACTIVITY(GO:1904683) |  | 18 | 0.66 | 1.06 | 0.141 | 0.359 | 1.000 | 10418 | tags=11%, list=18%, signal=13% |
| 5452 | PROTEIN\_LOCALIZATION\_TO\_PEROXISOME(GO:0072662) |  | 17 | 0.60 | 1.06 | 0.000 | 0.359 | 1.000 | 21459 | tags=82%, list=36%, signal=129% |
| 5453 | PROTEIN\_TARGETING\_TO\_PEROXISOME(GO:0006625) |  | 17 | 0.60 | 1.06 | 0.000 | 0.359 | 1.000 | 21459 | tags=82%, list=36%, signal=129% |
| 5454 | ESTABLISHMENT\_OF\_PROTEIN\_LOCALIZATION\_TO\_PEROXISOME(GO:0072663) |  | 17 | 0.60 | 1.06 | 0.000 | 0.358 | 1.000 | 21459 | tags=82%, list=36%, signal=129% |
| 5455 | RESPONSE\_TO\_SUPEROXIDE(GO:0000303) |  | 30 | 0.63 | 1.06 | 0.141 | 0.358 | 1.000 | 18635 | tags=50%, list=32%, signal=73% |
| 5456 | REGULATION\_OF\_RIG\_I\_SIGNALING\_PATHWAY(GO:0039535) |  | 17 | 0.63 | 1.06 | 0.141 | 0.358 | 1.000 | 14841 | tags=35%, list=25%, signal=47% |
| 5457 | CALCIUM\_ACTIVATED\_POTASSIUM\_CHANNEL\_ACTIVITY(GO:0015269) |  | 20 | 0.67 | 1.06 | 0.141 | 0.358 | 1.000 | 11380 | tags=30%, list=19%, signal=37% |
| 5458 | CYTOPLASMIC\_DYNEIN\_COMPLEX(GO:0005868) |  | 22 | 0.63 | 1.06 | 0.141 | 0.358 | 1.000 | 19263 | tags=73%, list=33%, signal=108% |
| 5459 | RNA\_POLYMERASE\_II\_REPRESSING\_TRANSCRIPTION\_FACTOR\_BINDING(GO:0001103) |  | 29 | 0.62 | 1.06 | 0.000 | 0.358 | 1.000 | 21618 | tags=93%, list=37%, signal=147% |
| 5460 | ENDONUCLEASE\_COMPLEX(GO:1905348) |  | 28 | 0.62 | 1.06 | 0.000 | 0.358 | 1.000 | 20821 | tags=79%, list=35%, signal=121% |
| 5461 | ENDORIBONUCLEASE\_COMPLEX(GO:1902555) |  | 28 | 0.62 | 1.06 | 0.000 | 0.358 | 1.000 | 20821 | tags=79%, list=35%, signal=121% |
| 5462 | CELLULAR\_RESPONSE\_TO\_THYROID\_HORMONE\_STIMULUS(GO:0097067) |  | 16 | 0.63 | 1.06 | 0.141 | 0.358 | 1.000 | 11742 | tags=38%, list=20%, signal=47% |
| 5463 | MIRNA\_BINDING(GO:0035198) |  | 24 | 0.61 | 1.06 | 0.141 | 0.358 | 1.000 | 22367 | tags=88%, list=38%, signal=141% |
| 5464 | DNA\_BIOSYNTHETIC\_PROCESS(GO:0071897) |  | 205 | 0.63 | 1.06 | 0.141 | 0.358 | 1.000 | 17340 | tags=59%, list=29%, signal=83% |
| 5465 | DEOXYRIBONUCLEOSIDE\_TRIPHOSPHATE\_METABOLIC\_PROCESS(GO:0009200) |  | 16 | 0.61 | 1.06 | 0.141 | 0.358 | 1.000 | 22842 | tags=94%, list=39%, signal=153% |
| 5466 | MEIOTIC\_CHROMOSOME\_SEPARATION(GO:0051307) |  | 19 | 0.66 | 1.06 | 0.141 | 0.358 | 1.000 | 16693 | tags=68%, list=28%, signal=95% |
| 5467 | CHROMOSOME\_\_TELOMERIC\_REGION(GO:0000781) |  | 157 | 0.60 | 1.06 | 0.141 | 0.358 | 1.000 | 20818 | tags=71%, list=35%, signal=110% |
| 5468 | UBIQUITIN\_BINDING(GO:0043130) |  | 65 | 0.61 | 1.06 | 0.141 | 0.358 | 1.000 | 18224 | tags=51%, list=31%, signal=73% |
| 5469 | TRNA\_SPLICING\_\_VIA\_ENDONUCLEOLYTIC\_CLEAVAGE\_AND\_LIGATION(GO:0006388) |  | 16 | 0.61 | 1.06 | 0.141 | 0.358 | 1.000 | 18644 | tags=63%, list=32%, signal=91% |
| 5470 | REGULATION\_OF\_FERTILIZATION(GO:0080154) |  | 28 | 0.64 | 1.06 | 0.141 | 0.358 | 1.000 | 14932 | tags=39%, list=25%, signal=53% |
| 5471 | CYTOKINE\_RECEPTOR\_BINDING(GO:0005126) |  | 275 | 0.63 | 1.06 | 0.141 | 0.358 | 1.000 | 15120 | tags=30%, list=26%, signal=40% |
| 5472 | SNRNA\_BINDING(GO:0017069) |  | 40 | 0.60 | 1.06 | 0.141 | 0.358 | 1.000 | 23135 | tags=88%, list=39%, signal=144% |
| 5473 | ATPASE\_ACTIVITY\_\_COUPLED\_TO\_MOVEMENT\_OF\_SUBSTANCES(GO:0043492) |  | 129 | 0.63 | 1.06 | 0.141 | 0.358 | 1.000 | 16279 | tags=44%, list=28%, signal=61% |
| 5474 | TRANSFERASE\_ACTIVITY\_\_TRANSFERRING\_ACYL\_GROUPS\_OTHER\_THAN\_AMINO\_ACYL\_GROUPS(GO:0016747) |  | 222 | 0.62 | 1.06 | 0.141 | 0.358 | 1.000 | 20745 | tags=67%, list=35%, signal=102% |
| 5475 | POSITIVE\_REGULATION\_OF\_MYELOID\_LEUKOCYTE\_DIFFERENTIATION(GO:0002763) |  | 52 | 0.64 | 1.06 | 0.141 | 0.358 | 1.000 | 16246 | tags=48%, list=27%, signal=66% |
| 5476 | REGULATION\_OF\_PROTEIN\_LOCALIZATION\_TO\_MEMBRANE(GO:1905475) |  | 167 | 0.64 | 1.06 | 0.141 | 0.358 | 1.000 | 15321 | tags=34%, list=26%, signal=45% |
| 5477 | T\_CELL\_MEDIATED\_CYTOTOXICITY(GO:0001913) |  | 26 | 0.61 | 1.06 | 0.141 | 0.358 | 1.000 | 18589 | tags=46%, list=31%, signal=67% |
| 5478 | HISTONE\_METHYLATION(GO:0016571) |  | 131 | 0.61 | 1.06 | 0.141 | 0.358 | 1.000 | 19377 | tags=60%, list=33%, signal=88% |
| 5479 | TRANSFERASE\_ACTIVITY\_\_TRANSFERRING\_ACYL\_GROUPS(GO:0016746) |  | 257 | 0.62 | 1.06 | 0.141 | 0.358 | 1.000 | 20745 | tags=63%, list=35%, signal=97% |
| 5480 | ACTIN\_CYTOSKELETON\_REORGANIZATION(GO:0031532) |  | 92 | 0.65 | 1.06 | 0.141 | 0.358 | 1.000 | 9119 | tags=23%, list=15%, signal=27% |
| 5481 | PEROXISOMAL\_TRANSPORT(GO:0043574) |  | 18 | 0.60 | 1.06 | 0.141 | 0.358 | 1.000 | 21459 | tags=78%, list=36%, signal=122% |
| 5482 | HISTONE\_H2A\_ACETYLATION(GO:0043968) |  | 15 | 0.63 | 1.06 | 0.141 | 0.358 | 1.000 | 20745 | tags=93%, list=35%, signal=144% |
| 5483 | MUSCLE\_ALPHA\_ACTININ\_BINDING(GO:0051371) |  | 18 | 0.64 | 1.06 | 0.141 | 0.358 | 1.000 | 18577 | tags=67%, list=31%, signal=97% |
| 5484 | TELOMERASE\_RNA\_BINDING(GO:0070034) |  | 22 | 0.62 | 1.06 | 0.000 | 0.358 | 1.000 | 22365 | tags=100%, list=38%, signal=161% |
| 5485 | POSITIVE\_REGULATION\_OF\_CELL\_MORPHOGENESIS\_INVOLVED\_IN\_DIFFERENTIATION(GO:0010770) |  | 137 | 0.64 | 1.06 | 0.141 | 0.358 | 1.000 | 13510 | tags=30%, list=23%, signal=39% |
| 5486 | POSITIVE\_REGULATION\_OF\_MICROTUBULE\_POLYMERIZATION(GO:0031116) |  | 26 | 0.60 | 1.06 | 0.141 | 0.358 | 1.000 | 20000 | tags=65%, list=34%, signal=99% |
| 5487 | MRNA\_METABOLIC\_PROCESS(GO:0016071) |  | 841 | 0.63 | 1.06 | 0.141 | 0.358 | 1.000 | 18555 | tags=59%, list=31%, signal=84% |
| 5488 | REGULATION\_OF\_RHO\_PROTEIN\_SIGNAL\_TRANSDUCTION(GO:0035023) |  | 138 | 0.63 | 1.06 | 0.141 | 0.358 | 1.000 | 14031 | tags=33%, list=24%, signal=43% |
| 5489 | EMBRYONIC\_PLACENTA\_DEVELOPMENT(GO:0001892) |  | 89 | 0.64 | 1.06 | 0.141 | 0.358 | 1.000 | 13649 | tags=31%, list=23%, signal=41% |
| 5490 | REGULATION\_OF\_CELL\_MATURATION(GO:1903429) |  | 21 | 0.61 | 1.06 | 0.141 | 0.358 | 1.000 | 20420 | tags=71%, list=35%, signal=109% |
| 5491 | CELLULAR\_MODIFIED\_AMINO\_ACID\_CATABOLIC\_PROCESS(GO:0042219) |  | 25 | 0.66 | 1.06 | 0.141 | 0.358 | 1.000 | 15430 | tags=32%, list=26%, signal=43% |
| 5492 | CIRCADIAN\_BEHAVIOR(GO:0048512) |  | 43 | 0.67 | 1.06 | 0.141 | 0.358 | 1.000 | 7172 | tags=21%, list=12%, signal=24% |
| 5493 | SEX\_DETERMINATION(GO:0007530) |  | 23 | 0.67 | 1.06 | 0.141 | 0.358 | 1.000 | 8490 | tags=35%, list=14%, signal=41% |
| 5494 | DISULFIDE\_OXIDOREDUCTASE\_ACTIVITY(GO:0015036) |  | 43 | 0.62 | 1.06 | 0.141 | 0.358 | 1.000 | 18144 | tags=58%, list=31%, signal=84% |
| 5495 | REGULATION\_OF\_CELLULAR\_RESPONSE\_TO\_HEAT(GO:1900034) |  | 77 | 0.60 | 1.06 | 0.000 | 0.358 | 1.000 | 18646 | tags=66%, list=32%, signal=97% |
| 5496 | TRANSLATION(GO:0006412) |  | 901 | 0.61 | 1.06 | 0.141 | 0.358 | 1.000 | 19357 | tags=49%, list=33%, signal=71% |
| 5497 | HEPATICOBILIARY\_SYSTEM\_DEVELOPMENT(GO:0061008) |  | 141 | 0.61 | 1.06 | 0.141 | 0.358 | 1.000 | 17154 | tags=51%, list=29%, signal=72% |
| 5498 | PCG\_PROTEIN\_COMPLEX(GO:0031519) |  | 46 | 0.60 | 1.06 | 0.000 | 0.358 | 1.000 | 20225 | tags=70%, list=34%, signal=106% |
| 5499 | ESTABLISHMENT\_OF\_PROTEIN\_LOCALIZATION\_TO\_MEMBRANE(GO:0090150) |  | 278 | 0.63 | 1.06 | 0.141 | 0.358 | 1.000 | 16763 | tags=41%, list=28%, signal=57% |
| 5500 | REGULATION\_OF\_CARDIOCYTE\_DIFFERENTIATION(GO:1905207) |  | 57 | 0.64 | 1.06 | 0.141 | 0.358 | 1.000 | 17343 | tags=42%, list=29%, signal=60% |
| 5501 | APOLIPOPROTEIN\_BINDING(GO:0034185) |  | 17 | 0.62 | 1.06 | 0.307 | 0.358 | 1.000 | 15911 | tags=47%, list=27%, signal=64% |
| 5502 | TELOMERE\_ORGANIZATION(GO:0032200) |  | 171 | 0.60 | 1.06 | 0.141 | 0.358 | 1.000 | 20903 | tags=73%, list=35%, signal=112% |
| 5503 | CELLULAR\_RESPONSE\_TO\_TOPOLOGICALLY\_INCORRECT\_PROTEIN(GO:0035967) |  | 153 | 0.61 | 1.06 | 0.141 | 0.358 | 1.000 | 19291 | tags=50%, list=33%, signal=74% |
| 5504 | INTRINSIC\_APOPTOTIC\_SIGNALING\_PATHWAY\_BY\_P53\_CLASS\_MEDIATOR(GO:0072332) |  | 81 | 0.62 | 1.06 | 0.141 | 0.358 | 1.000 | 17051 | tags=42%, list=29%, signal=59% |
| 5505 | PEPTIDE\_LYSINE\_N\_ACETYLTRANSFERASE\_ACTIVITY(GO:0061733) |  | 60 | 0.60 | 1.06 | 0.141 | 0.358 | 1.000 | 20745 | tags=73%, list=35%, signal=113% |
| 5506 | RNA\_POLYMERASE\_II\_TRANSCRIPTIONAL\_PREINITIATION\_COMPLEX\_ASSEMBLY(GO:0051123) |  | 32 | 0.66 | 1.06 | 0.141 | 0.358 | 1.000 | 17312 | tags=72%, list=29%, signal=102% |
| 5507 | DEAMINASE\_ACTIVITY(GO:0019239) |  | 33 | 0.60 | 1.06 | 0.141 | 0.358 | 1.000 | 14400 | tags=39%, list=24%, signal=52% |
| 5508 | GLOBAL\_GENOME\_NUCLEOTIDE\_EXCISION\_REPAIR(GO:0070911) |  | 32 | 0.60 | 1.06 | 0.000 | 0.358 | 1.000 | 22198 | tags=84%, list=38%, signal=135% |
| 5509 | PHOSPHORYLATION\_OF\_RNA\_POLYMERASE\_II\_C\_TERMINAL\_DOMAIN(GO:0070816) |  | 16 | 0.62 | 1.06 | 0.000 | 0.358 | 1.000 | 21346 | tags=88%, list=36%, signal=137% |
| 5510 | POSITIVE\_REGULATION\_OF\_TOLL\_LIKE\_RECEPTOR\_SIGNALING\_PATHWAY(GO:0034123) |  | 20 | 0.62 | 1.06 | 0.141 | 0.358 | 1.000 | 15981 | tags=35%, list=27%, signal=48% |
| 5511 | EXON\_EXON\_JUNCTION\_COMPLEX(GO:0035145) |  | 20 | 0.64 | 1.06 | 0.000 | 0.358 | 1.000 | 18239 | tags=80%, list=31%, signal=116% |
| 5512 | TRNA\_PROCESSING(GO:0008033) |  | 125 | 0.59 | 1.06 | 0.141 | 0.358 | 1.000 | 21006 | tags=76%, list=36%, signal=118% |
| 5513 | REGULATION\_OF\_NATURAL\_KILLER\_CELL\_MEDIATED\_CYTOTOXICITY(GO:0042269) |  | 33 | 0.62 | 1.06 | 0.141 | 0.358 | 1.000 | 15869 | tags=48%, list=27%, signal=66% |
| 5514 | REGULATION\_OF\_ACROSOME\_REACTION(GO:0060046) |  | 20 | 0.66 | 1.06 | 0.307 | 0.358 | 1.000 | 19043 | tags=55%, list=32%, signal=81% |
| 5515 | CARDIAC\_MUSCLE\_CELL\_APOPTOTIC\_PROCESS(GO:0010659) |  | 54 | 0.59 | 1.06 | 0.141 | 0.358 | 1.000 | 15300 | tags=28%, list=26%, signal=37% |
| 5516 | CELLULAR\_RESPONSE\_TO\_UNFOLDED\_PROTEIN(GO:0034620) |  | 136 | 0.61 | 1.06 | 0.141 | 0.358 | 1.000 | 19291 | tags=49%, list=33%, signal=72% |
| 5517 | REGULATION\_OF\_RHODOPSIN\_MEDIATED\_SIGNALING\_PATHWAY(GO:0022400) |  | 26 | 0.61 | 1.06 | 0.000 | 0.358 | 1.000 | 21797 | tags=73%, list=37%, signal=116% |
| 5518 | PML\_BODY(GO:0016605) |  | 97 | 0.60 | 1.06 | 0.141 | 0.358 | 1.000 | 20722 | tags=66%, list=35%, signal=101% |
| 5519 | MITOCHONDRIAL\_MEMBRANE\_ORGANIZATION(GO:0007006) |  | 134 | 0.64 | 1.06 | 0.141 | 0.358 | 1.000 | 13041 | tags=20%, list=22%, signal=26% |
| 5520 | ENERGY\_RESERVE\_METABOLIC\_PROCESS(GO:0006112) |  | 93 | 0.63 | 1.06 | 0.141 | 0.358 | 1.000 | 17672 | tags=46%, list=30%, signal=66% |
| 5521 | LAMININ\_BINDING(GO:0043236) |  | 28 | 0.65 | 1.06 | 0.141 | 0.358 | 1.000 | 11467 | tags=18%, list=19%, signal=22% |
| 5522 | POSITIVE\_REGULATION\_OF\_MRNA\_PROCESSING(GO:0050685) |  | 26 | 0.61 | 1.06 | 0.141 | 0.358 | 1.000 | 18871 | tags=65%, list=32%, signal=96% |
| 5523 | CELLULAR\_ALDEHYDE\_METABOLIC\_PROCESS(GO:0006081) |  | 77 | 0.61 | 1.06 | 0.141 | 0.358 | 1.000 | 18741 | tags=47%, list=32%, signal=68% |
| 5524 | NEGATIVE\_REGULATION\_OF\_TRANSFORMING\_GROWTH\_FACTOR\_BETA\_RECEPTOR\_SIGNALING\_PATHWAY(GO:0030512) |  | 74 | 0.62 | 1.06 | 0.141 | 0.358 | 1.000 | 14781 | tags=30%, list=25%, signal=40% |
| 5525 | GTP\_METABOLIC\_PROCESS(GO:0046039) |  | 33 | 0.60 | 1.06 | 0.000 | 0.358 | 1.000 | 17118 | tags=61%, list=29%, signal=85% |
| 5526 | DNA\_POLYMERASE\_COMPLEX(GO:0042575) |  | 18 | 0.61 | 1.06 | 0.141 | 0.358 | 1.000 | 22910 | tags=89%, list=39%, signal=145% |
| 5527 | POSITIVE\_REGULATION\_OF\_IMMUNE\_SYSTEM\_PROCESS(GO:0002684) |  | 1085 | 0.62 | 1.06 | 0.141 | 0.358 | 1.000 | 16724 | tags=32%, list=28%, signal=44% |
| 5528 | REGULATION\_OF\_CARDIAC\_MUSCLE\_CELL\_DIFFERENTIATION(GO:2000725) |  | 42 | 0.64 | 1.06 | 0.141 | 0.358 | 1.000 | 17343 | tags=45%, list=29%, signal=64% |
| 5529 | PROTEIN\_PHOSPHATASE\_INHIBITOR\_ACTIVITY(GO:0004864) |  | 39 | 0.60 | 1.06 | 0.141 | 0.358 | 1.000 | 22011 | tags=67%, list=37%, signal=106% |
| 5530 | CLATHRIN\_COATED\_ENDOCYTIC\_VESICLE\_MEMBRANE(GO:0030669) |  | 39 | 0.62 | 1.06 | 0.141 | 0.358 | 1.000 | 12547 | tags=26%, list=21%, signal=33% |
| 5531 | LUNG\_EPITHELIAL\_CELL\_DIFFERENTIATION(GO:0060487) |  | 22 | 0.68 | 1.06 | 0.141 | 0.358 | 1.000 | 12189 | tags=27%, list=21%, signal=34% |
| 5532 | SULFURIC\_ESTER\_HYDROLASE\_ACTIVITY(GO:0008484) |  | 19 | 0.66 | 1.06 | 0.141 | 0.358 | 1.000 | 13839 | tags=37%, list=23%, signal=48% |
| 5533 | INTRAMOLECULAR\_TRANSFERASE\_ACTIVITY(GO:0016866) |  | 27 | 0.59 | 1.06 | 0.141 | 0.358 | 1.000 | 19634 | tags=67%, list=33%, signal=100% |
| 5534 | SPLICEOSOMAL\_SNRNP\_ASSEMBLY(GO:0000387) |  | 38 | 0.62 | 1.06 | 0.000 | 0.358 | 1.000 | 20582 | tags=87%, list=35%, signal=133% |
| 5535 | POSITIVE\_REGULATION\_OF\_TRANSLATION(GO:0045727) |  | 123 | 0.63 | 1.06 | 0.141 | 0.358 | 1.000 | 16571 | tags=42%, list=28%, signal=59% |
| 5536 | RIBONUCLEOPROTEIN\_COMPLEX\_BINDING(GO:0043021) |  | 120 | 0.60 | 1.06 | 0.141 | 0.358 | 1.000 | 22772 | tags=88%, list=39%, signal=142% |
| 5537 | METHYL\_CPG\_BINDING(GO:0008327) |  | 21 | 0.60 | 1.06 | 0.141 | 0.358 | 1.000 | 22226 | tags=67%, list=38%, signal=107% |
| 5538 | RHYTHMIC\_BEHAVIOR(GO:0007622) |  | 44 | 0.67 | 1.06 | 0.141 | 0.358 | 1.000 | 7172 | tags=20%, list=12%, signal=23% |
| 5539 | TRANSCRIPTION\_INITIATION\_FROM\_RNA\_POLYMERASE\_I\_PROMOTER(GO:0006361) |  | 35 | 0.59 | 1.06 | 0.141 | 0.358 | 1.000 | 20344 | tags=69%, list=34%, signal=104% |
| 5540 | POSITIVE\_REGULATION\_OF\_CELL\_GROWTH(GO:0030307) |  | 158 | 0.63 | 1.06 | 0.141 | 0.358 | 1.000 | 18019 | tags=51%, list=30%, signal=74% |
| 5541 | MITOCHONDRIAL\_RNA\_PROCESSING(GO:0000963) |  | 16 | 0.60 | 1.06 | 0.141 | 0.358 | 1.000 | 21335 | tags=88%, list=36%, signal=137% |
| 5542 | MITOCHONDRIAL\_TRANSLATION(GO:0032543) |  | 122 | 0.60 | 1.06 | 0.141 | 0.358 | 1.000 | 21163 | tags=86%, list=36%, signal=134% |
| 5543 | DNA\_DIRECTED\_5'\_3'\_RNA\_POLYMERASE\_ACTIVITY(GO:0003899) |  | 47 | 0.59 | 1.06 | 0.141 | 0.358 | 1.000 | 22286 | tags=83%, list=38%, signal=133% |
| 5544 | RNA\_HELICASE\_ACTIVITY(GO:0003724) |  | 68 | 0.59 | 1.06 | 0.141 | 0.358 | 1.000 | 23904 | tags=88%, list=40%, signal=148% |
| 5545 | MYOFILAMENT(GO:0036379) |  | 24 | 0.64 | 1.06 | 0.141 | 0.358 | 1.000 | 13148 | tags=46%, list=22%, signal=59% |
| 5546 | INDOLE\_CONTAINING\_COMPOUND\_METABOLIC\_PROCESS(GO:0042430) |  | 24 | 0.65 | 1.06 | 0.141 | 0.358 | 1.000 | 15175 | tags=38%, list=26%, signal=50% |
| 5547 | PEPTIDE\_N\_ACETYLTRANSFERASE\_ACTIVITY(GO:0034212) |  | 68 | 0.60 | 1.06 | 0.141 | 0.358 | 1.000 | 20745 | tags=75%, list=35%, signal=115% |
| 5548 | NUCLEAR\_CYCLIN\_DEPENDENT\_PROTEIN\_KINASE\_HOLOENZYME\_COMPLEX(GO:0019908) |  | 16 | 0.65 | 1.06 | 0.168 | 0.358 | 1.000 | 16370 | tags=69%, list=28%, signal=95% |
| 5549 | TRANSCRIPTION\_BY\_RNA\_POLYMERASE\_I(GO:0006360) |  | 62 | 0.59 | 1.06 | 0.141 | 0.358 | 1.000 | 20344 | tags=73%, list=34%, signal=111% |
| 5550 | REGULATION\_OF\_GLUCOSE\_METABOLIC\_PROCESS(GO:0010906) |  | 95 | 0.63 | 1.06 | 0.141 | 0.358 | 1.000 | 17502 | tags=43%, list=30%, signal=61% |
| 5551 | SPINDLE(GO:0005819) |  | 327 | 0.63 | 1.06 | 0.141 | 0.358 | 1.000 | 18392 | tags=64%, list=31%, signal=92% |
| 5552 | PRODUCTION\_OF\_MOLECULAR\_MEDIATOR\_INVOLVED\_IN\_INFLAMMATORY\_RESPONSE(GO:0002532) |  | 58 | 0.62 | 1.06 | 0.141 | 0.358 | 1.000 | 14390 | tags=34%, list=24%, signal=46% |
| 5553 | RRNA\_METHYLATION(GO:0031167) |  | 24 | 0.60 | 1.06 | 0.000 | 0.358 | 1.000 | 19157 | tags=71%, list=32%, signal=105% |
| 5554 | POSITIVE\_REGULATION\_OF\_ADAPTIVE\_IMMUNE\_RESPONSE(GO:0002821) |  | 80 | 0.62 | 1.06 | 0.141 | 0.358 | 1.000 | 15082 | tags=41%, list=26%, signal=55% |
| 5555 | AMINO\_ACID\_ACTIVATION(GO:0043038) |  | 57 | 0.61 | 1.06 | 0.000 | 0.358 | 1.000 | 22212 | tags=88%, list=38%, signal=140% |
| 5556 | ACUTE\_INFLAMMATORY\_RESPONSE\_TO\_ANTIGENIC\_STIMULUS(GO:0002438) |  | 22 | 0.67 | 1.06 | 0.141 | 0.358 | 1.000 | 18692 | tags=36%, list=32%, signal=53% |
| 5557 | RIBONUCLEOPROTEIN\_COMPLEX\_BIOGENESIS(GO:0022613) |  | 485 | 0.59 | 1.06 | 0.141 | 0.358 | 1.000 | 23272 | tags=88%, list=39%, signal=144% |
| 5558 | CYTOPLASMIC\_PATTERN\_RECOGNITION\_RECEPTOR\_SIGNALING\_PATHWAY\_IN\_RESPONSE\_TO\_VIRUS(GO:0039528) |  | 25 | 0.63 | 1.06 | 0.141 | 0.358 | 1.000 | 15128 | tags=36%, list=26%, signal=48% |
| 5559 | PEPTIDE\_TRANSMEMBRANE\_TRANSPORTER\_ACTIVITY(GO:1904680) |  | 28 | 0.59 | 1.06 | 0.141 | 0.358 | 1.000 | 16449 | tags=43%, list=28%, signal=59% |
| 5560 | POSITIVE\_REGULATION\_OF\_BLOOD\_PRESSURE(GO:0045777) |  | 40 | 0.62 | 1.06 | 0.141 | 0.358 | 1.000 | 17432 | tags=40%, list=29%, signal=57% |
| 5561 | NEGATIVE\_REGULATION\_OF\_TELOMERE\_MAINTENANCE\_VIA\_TELOMERE\_LENGTHENING(GO:1904357) |  | 27 | 0.61 | 1.06 | 0.000 | 0.358 | 1.000 | 20841 | tags=81%, list=35%, signal=126% |
| 5562 | TORC1\_SIGNALING(GO:0038202) |  | 36 | 0.62 | 1.06 | 0.141 | 0.358 | 1.000 | 12070 | tags=28%, list=20%, signal=35% |
| 5563 | RIBOSOMAL\_SMALL\_SUBUNIT\_BIOGENESIS(GO:0042274) |  | 72 | 0.59 | 1.06 | 0.141 | 0.358 | 1.000 | 23262 | tags=88%, list=39%, signal=144% |
| 5564 | HISTONE\_ACETYLTRANSFERASE\_ACTIVITY(GO:0004402) |  | 58 | 0.60 | 1.06 | 0.141 | 0.358 | 1.000 | 20745 | tags=74%, list=35%, signal=114% |
| 5565 | MITOCHONDRIAL\_RESPIRATORY\_CHAIN\_COMPLEX\_I\_ASSEMBLY(GO:0032981) |  | 64 | 0.58 | 1.06 | 0.141 | 0.358 | 1.000 | 24412 | tags=97%, list=41%, signal=165% |
| 5566 | NADH\_DEHYDROGENASE\_COMPLEX\_ASSEMBLY(GO:0010257) |  | 64 | 0.58 | 1.06 | 0.141 | 0.358 | 1.000 | 24412 | tags=97%, list=41%, signal=165% |
| 5567 | CELLULAR\_PROTEIN\_COMPLEX\_DISASSEMBLY(GO:0043624) |  | 202 | 0.62 | 1.06 | 0.141 | 0.358 | 1.000 | 17950 | tags=57%, list=30%, signal=82% |
| 5568 | ERAD\_PATHWAY(GO:0036503) |  | 93 | 0.60 | 1.06 | 0.141 | 0.358 | 1.000 | 19239 | tags=55%, list=33%, signal=81% |
| 5569 | INOSITOL\_PHOSPHATE\_PHOSPHATASE\_ACTIVITY(GO:0052745) |  | 19 | 0.61 | 1.06 | 0.141 | 0.358 | 1.000 | 19903 | tags=63%, list=34%, signal=95% |
| 5570 | TRNA\_5'\_END\_PROCESSING(GO:0099116) |  | 16 | 0.64 | 1.06 | 0.000 | 0.358 | 1.000 | 20821 | tags=88%, list=35%, signal=135% |
| 5571 | STRAND\_DISPLACEMENT(GO:0000732) |  | 26 | 0.62 | 1.06 | 0.340 | 0.358 | 1.000 | 12766 | tags=50%, list=22%, signal=64% |
| 5572 | GLUCAN\_METABOLIC\_PROCESS(GO:0044042) |  | 80 | 0.63 | 1.06 | 0.141 | 0.358 | 1.000 | 17672 | tags=50%, list=30%, signal=71% |
| 5573 | CELLULAR\_GLUCAN\_METABOLIC\_PROCESS(GO:0006073) |  | 80 | 0.63 | 1.06 | 0.141 | 0.358 | 1.000 | 17672 | tags=50%, list=30%, signal=71% |
| 5574 | POSITIVE\_REGULATION\_OF\_INSULIN\_SECRETION\_INVOLVED\_IN\_CELLULAR\_RESPONSE\_TO\_GLUCOSE\_STIMULUS(GO:0035774) |  | 30 | 0.61 | 1.06 | 0.141 | 0.358 | 1.000 | 19716 | tags=67%, list=33%, signal=100% |
| 5575 | NUCLEUS\_ORGANIZATION(GO:0006997) |  | 148 | 0.59 | 1.06 | 0.141 | 0.358 | 1.000 | 21524 | tags=74%, list=36%, signal=117% |
| 5576 | DNA\_REPLICATION\_FACTOR\_A\_COMPLEX(GO:0005662) |  | 15 | 0.62 | 1.06 | 0.000 | 0.358 | 1.000 | 17064 | tags=73%, list=29%, signal=103% |
| 5577 | NEGATIVE\_REGULATION\_OF\_TELOMERE\_MAINTENANCE\_VIA\_TELOMERASE(GO:0032211) |  | 20 | 0.60 | 1.06 | 0.000 | 0.358 | 1.000 | 20841 | tags=80%, list=35%, signal=123% |
| 5578 | NEGATIVE\_REGULATION\_OF\_CALCIUM\_ION\_TRANSMEMBRANE\_TRANSPORTER\_ACTIVITY(GO:1901020) |  | 38 | 0.60 | 1.06 | 0.141 | 0.358 | 1.000 | 15565 | tags=34%, list=26%, signal=46% |
| 5579 | TRNA\_AMINOACYLATION(GO:0043039) |  | 56 | 0.61 | 1.06 | 0.000 | 0.358 | 1.000 | 22212 | tags=88%, list=38%, signal=140% |
| 5580 | SWI\_SNF\_COMPLEX(GO:0016514) |  | 15 | 0.62 | 1.06 | 0.141 | 0.358 | 1.000 | 22085 | tags=80%, list=37%, signal=128% |
| 5581 | MYELOID\_DENDRITIC\_CELL\_DIFFERENTIATION(GO:0043011) |  | 18 | 0.67 | 1.06 | 0.141 | 0.358 | 1.000 | 17938 | tags=61%, list=30%, signal=88% |
| 5582 | REGULATION\_OF\_INSULIN\_SECRETION\_INVOLVED\_IN\_CELLULAR\_RESPONSE\_TO\_GLUCOSE\_STIMULUS(GO:0061178) |  | 55 | 0.60 | 1.06 | 0.141 | 0.358 | 1.000 | 18767 | tags=60%, list=32%, signal=88% |
| 5583 | REGULATION\_OF\_CARDIAC\_MUSCLE\_CELL\_APOPTOTIC\_PROCESS(GO:0010665) |  | 50 | 0.59 | 1.06 | 0.141 | 0.358 | 1.000 | 17567 | tags=30%, list=30%, signal=43% |
| 5584 | REGULATION\_OF\_EXTRACELLULAR\_MATRIX\_ASSEMBLY(GO:1901201) |  | 15 | 0.61 | 1.06 | 0.141 | 0.358 | 1.000 | 18904 | tags=60%, list=32%, signal=88% |
| 5585 | DENDRITIC\_CELL\_CHEMOTAXIS(GO:0002407) |  | 23 | 0.65 | 1.06 | 0.141 | 0.358 | 1.000 | 13828 | tags=26%, list=23%, signal=34% |
| 5586 | SARCOPLASMIC\_RETICULUM\_MEMBRANE(GO:0033017) |  | 40 | 0.61 | 1.06 | 0.141 | 0.358 | 1.000 | 17630 | tags=58%, list=30%, signal=82% |
| 5587 | METALLOAMINOPEPTIDASE\_ACTIVITY(GO:0070006) |  | 20 | 0.62 | 1.06 | 0.141 | 0.358 | 1.000 | 17866 | tags=55%, list=30%, signal=79% |
| 5588 | NEGATIVE\_REGULATION\_OF\_LYMPHOCYTE\_MEDIATED\_IMMUNITY(GO:0002707) |  | 33 | 0.63 | 1.06 | 0.141 | 0.358 | 1.000 | 15529 | tags=39%, list=26%, signal=53% |
| 5589 | REGULATION\_OF\_TRANSCRIPTION\_INITIATION\_FROM\_RNA\_POLYMERASE\_II\_PROMOTER(GO:0060260) |  | 29 | 0.65 | 1.06 | 0.141 | 0.358 | 1.000 | 17119 | tags=59%, list=29%, signal=82% |
| 5590 | REGULATION\_OF\_SHORT\_TERM\_NEURONAL\_SYNAPTIC\_PLASTICITY(GO:0048172) |  | 15 | 0.62 | 1.06 | 0.141 | 0.358 | 1.000 | 18574 | tags=67%, list=31%, signal=97% |
| 5591 | PROTEIN\_ALKYLATION(GO:0008213) |  | 178 | 0.60 | 1.06 | 0.141 | 0.358 | 1.000 | 19904 | tags=62%, list=34%, signal=93% |
| 5592 | PROTEIN\_METHYLATION(GO:0006479) |  | 178 | 0.60 | 1.06 | 0.141 | 0.358 | 1.000 | 19904 | tags=62%, list=34%, signal=93% |
| 5593 | GLYCOGEN\_METABOLIC\_PROCESS(GO:0005977) |  | 79 | 0.63 | 1.06 | 0.141 | 0.358 | 1.000 | 17672 | tags=49%, list=30%, signal=70% |
| 5594 | LIGASE\_ACTIVITY\_\_FORMING\_CARBON\_NITROGEN\_BONDS(GO:0016879) |  | 57 | 0.60 | 1.06 | 0.000 | 0.358 | 1.000 | 21071 | tags=74%, list=36%, signal=114% |
| 5595 | TELOMERIC\_DNA\_BINDING(GO:0042162) |  | 36 | 0.61 | 1.06 | 0.000 | 0.358 | 1.000 | 20697 | tags=86%, list=35%, signal=132% |
| 5596 | POSITIVE\_REGULATION\_OF\_DEFENSE\_RESPONSE\_TO\_VIRUS\_BY\_HOST(GO:0002230) |  | 24 | 0.65 | 1.06 | 0.141 | 0.358 | 1.000 | 13278 | tags=29%, list=22%, signal=38% |
| 5597 | REGULATION\_OF\_MEMBRANE\_PROTEIN\_ECTODOMAIN\_PROTEOLYSIS(GO:0051043) |  | 24 | 0.60 | 1.06 | 0.307 | 0.358 | 1.000 | 14886 | tags=42%, list=25%, signal=56% |
| 5598 | TRNA\_AMINOACYLATION\_FOR\_PROTEIN\_TRANSLATION(GO:0006418) |  | 53 | 0.61 | 1.06 | 0.141 | 0.358 | 1.000 | 22212 | tags=87%, list=38%, signal=139% |
| 5599 | ATPASE\_ACTIVATOR\_ACTIVITY(GO:0001671) |  | 21 | 0.63 | 1.06 | 0.168 | 0.358 | 1.000 | 15823 | tags=57%, list=27%, signal=78% |
| 5600 | SPLICEOSOMAL\_COMPLEX(GO:0005681) |  | 186 | 0.59 | 1.05 | 0.141 | 0.358 | 1.000 | 22162 | tags=83%, list=37%, signal=132% |
| 5601 | REGULATION\_OF\_CHROMATIN\_SILENCING(GO:0031935) |  | 21 | 0.59 | 1.05 | 0.141 | 0.358 | 1.000 | 23965 | tags=95%, list=41%, signal=160% |
| 5602 | MATURATION\_OF\_SSU\_RRNA(GO:0030490) |  | 54 | 0.59 | 1.05 | 0.141 | 0.358 | 1.000 | 23262 | tags=91%, list=39%, signal=149% |
| 5603 | MITOCHONDRIAL\_FUSION(GO:0008053) |  | 23 | 0.60 | 1.05 | 0.141 | 0.358 | 1.000 | 21145 | tags=74%, list=36%, signal=115% |
| 5604 | RNA\_SECONDARY\_STRUCTURE\_UNWINDING(GO:0010501) |  | 46 | 0.59 | 1.05 | 0.141 | 0.358 | 1.000 | 23904 | tags=91%, list=40%, signal=153% |
| 5605 | NEGATIVE\_REGULATION\_OF\_UBIQUITIN\_DEPENDENT\_PROTEIN\_CATABOLIC\_PROCESS(GO:2000059) |  | 42 | 0.62 | 1.05 | 0.141 | 0.358 | 1.000 | 20906 | tags=76%, list=35%, signal=118% |
| 5606 | PROTEIN\_TRANSMEMBRANE\_IMPORT\_INTO\_INTRACELLULAR\_ORGANELLE(GO:0044743) |  | 38 | 0.58 | 1.05 | 0.141 | 0.358 | 1.000 | 19319 | tags=68%, list=33%, signal=102% |
| 5607 | NUCLEOTIDE\_EXCISION\_REPAIR\_\_DNA\_DUPLEX\_UNWINDING(GO:0000717) |  | 22 | 0.60 | 1.05 | 0.141 | 0.358 | 1.000 | 22198 | tags=82%, list=38%, signal=131% |
| 5608 | POSITIVE\_REGULATION\_OF\_PROTEIN\_UBIQUITINATION(GO:0031398) |  | 106 | 0.63 | 1.05 | 0.141 | 0.358 | 1.000 | 17110 | tags=53%, list=29%, signal=74% |
| 5609 | CHD\_TYPE\_COMPLEX(GO:0090545) |  | 16 | 0.62 | 1.05 | 0.000 | 0.358 | 1.000 | 21257 | tags=88%, list=36%, signal=137% |
| 5610 | NURD\_COMPLEX(GO:0016581) |  | 16 | 0.62 | 1.05 | 0.000 | 0.358 | 1.000 | 21257 | tags=88%, list=36%, signal=137% |
| 5611 | RESPONSE\_TO\_SALT(GO:1902074) |  | 19 | 0.63 | 1.05 | 0.141 | 0.358 | 1.000 | 18737 | tags=58%, list=32%, signal=85% |
| 5612 | DNA\_HELICASE\_ACTIVITY(GO:0003678) |  | 49 | 0.59 | 1.05 | 0.141 | 0.358 | 1.000 | 22593 | tags=90%, list=38%, signal=145% |
| 5613 | ORGANELLAR\_RIBOSOME(GO:0000313) |  | 84 | 0.59 | 1.05 | 0.141 | 0.358 | 1.000 | 23709 | tags=94%, list=40%, signal=157% |
| 5614 | MITOCHONDRIAL\_RIBOSOME(GO:0005761) |  | 84 | 0.59 | 1.05 | 0.141 | 0.358 | 1.000 | 23709 | tags=94%, list=40%, signal=157% |
| 5615 | REGULATION\_OF\_MESODERM\_DEVELOPMENT(GO:2000380) |  | 20 | 0.65 | 1.05 | 0.141 | 0.358 | 1.000 | 19172 | tags=40%, list=32%, signal=59% |
| 5616 | O\_METHYLTRANSFERASE\_ACTIVITY(GO:0008171) |  | 23 | 0.59 | 1.05 | 0.141 | 0.358 | 1.000 | 20268 | tags=65%, list=34%, signal=99% |
| 5617 | INTRINSIC\_APOPTOTIC\_SIGNALING\_PATHWAY\_IN\_RESPONSE\_TO\_DNA\_DAMAGE\_BY\_P53\_CLASS\_MEDIATOR(GO:0042771) |  | 43 | 0.63 | 1.05 | 0.141 | 0.358 | 1.000 | 18343 | tags=49%, list=31%, signal=71% |
| 5618 | AMIDE\_TRANSMEMBRANE\_TRANSPORTER\_ACTIVITY(GO:0042887) |  | 40 | 0.58 | 1.05 | 0.141 | 0.358 | 1.000 | 16449 | tags=40%, list=28%, signal=55% |
| 5619 | MISMATCH\_REPAIR(GO:0006298) |  | 32 | 0.65 | 1.05 | 0.141 | 0.358 | 1.000 | 17788 | tags=72%, list=30%, signal=103% |
| 5620 | BRANCHING\_INVOLVED\_IN\_MAMMARY\_GLAND\_DUCT\_MORPHOGENESIS(GO:0060444) |  | 23 | 0.64 | 1.05 | 0.141 | 0.358 | 1.000 | 13967 | tags=26%, list=24%, signal=34% |
| 5621 | RRNA\_CONTAINING\_RIBONUCLEOPROTEIN\_COMPLEX\_EXPORT\_FROM\_NUCLEUS(GO:0071428) |  | 16 | 0.61 | 1.05 | 0.000 | 0.358 | 1.000 | 19487 | tags=69%, list=33%, signal=103% |
| 5622 | CALCINEURIN\_NFAT\_SIGNALING\_CASCADE(GO:0033173) |  | 26 | 0.65 | 1.05 | 0.141 | 0.358 | 1.000 | 17187 | tags=62%, list=29%, signal=87% |
| 5623 | LIPID\_PARTICLE\_ORGANIZATION(GO:0034389) |  | 19 | 0.63 | 1.05 | 0.141 | 0.358 | 1.000 | 18237 | tags=47%, list=31%, signal=68% |
| 5624 | ATP\_DEPENDENT\_RNA\_HELICASE\_ACTIVITY(GO:0004004) |  | 67 | 0.59 | 1.05 | 0.141 | 0.358 | 1.000 | 23904 | tags=88%, list=40%, signal=148% |
| 5625 | RIBOSOME\_BIOGENESIS(GO:0042254) |  | 333 | 0.58 | 1.05 | 0.141 | 0.358 | 1.000 | 23272 | tags=86%, list=39%, signal=141% |
| 5626 | NEGATIVE\_REGULATION\_OF\_CELL\_SUBSTRATE\_ADHESION(GO:0010812) |  | 60 | 0.65 | 1.05 | 0.141 | 0.358 | 1.000 | 14949 | tags=37%, list=25%, signal=49% |
| 5627 | NCRNA\_5'\_END\_PROCESSING(GO:0034471) |  | 19 | 0.64 | 1.05 | 0.000 | 0.358 | 1.000 | 20821 | tags=89%, list=35%, signal=138% |
| 5628 | RNA\_DEPENDENT\_ATPASE\_ACTIVITY(GO:0008186) |  | 68 | 0.59 | 1.05 | 0.141 | 0.358 | 1.000 | 23904 | tags=88%, list=40%, signal=148% |
| 5629 | DETECTION\_OF\_LIGHT\_STIMULUS(GO:0009583) |  | 72 | 0.60 | 1.05 | 0.141 | 0.358 | 1.000 | 14399 | tags=33%, list=24%, signal=44% |
| 5630 | RNA\_CAPPING(GO:0036260) |  | 34 | 0.59 | 1.05 | 0.141 | 0.358 | 1.000 | 21089 | tags=76%, list=36%, signal=119% |
| 5631 | 7\_METHYLGUANOSINE\_RNA\_CAPPING(GO:0009452) |  | 34 | 0.59 | 1.05 | 0.141 | 0.358 | 1.000 | 21089 | tags=76%, list=36%, signal=119% |
| 5632 | SM\_LIKE\_PROTEIN\_FAMILY\_COMPLEX(GO:0120114) |  | 81 | 0.60 | 1.05 | 0.141 | 0.358 | 1.000 | 22418 | tags=84%, list=38%, signal=135% |
| 5633 | NEGATIVE\_REGULATION\_OF\_INTERLEUKIN\_6\_PRODUCTION(GO:0032715) |  | 38 | 0.62 | 1.05 | 0.141 | 0.358 | 1.000 | 14963 | tags=29%, list=25%, signal=39% |
| 5634 | NUCLEAR\_ENVELOPE\_ORGANIZATION(GO:0006998) |  | 85 | 0.60 | 1.05 | 0.141 | 0.358 | 1.000 | 21515 | tags=84%, list=36%, signal=131% |
| 5635 | POSITIVE\_REGULATION\_OF\_NF\_KAPPAB\_TRANSCRIPTION\_FACTOR\_ACTIVITY(GO:0051092) |  | 140 | 0.61 | 1.05 | 0.141 | 0.358 | 1.000 | 18226 | tags=46%, list=31%, signal=66% |
| 5636 | NUCLEOTIDE\_TRANSMEMBRANE\_TRANSPORTER\_ACTIVITY(GO:0015215) |  | 16 | 0.60 | 1.05 | 0.141 | 0.358 | 1.000 | 21199 | tags=75%, list=36%, signal=117% |
| 5637 | NEGATIVE\_REGULATION\_OF\_CYCLIN\_DEPENDENT\_PROTEIN\_SERINE\_THREONINE\_KINASE\_ACTIVITY(GO:0045736) |  | 30 | 0.63 | 1.05 | 0.141 | 0.358 | 1.000 | 13695 | tags=33%, list=23%, signal=43% |
| 5638 | MATURATION\_OF\_LSU\_RRNA(GO:0000470) |  | 24 | 0.61 | 1.05 | 0.141 | 0.358 | 1.000 | 21082 | tags=83%, list=36%, signal=129% |
| 5639 | RNA\_POLYMERASE\_ACTIVITY(GO:0097747) |  | 49 | 0.58 | 1.05 | 0.141 | 0.358 | 1.000 | 22286 | tags=82%, list=38%, signal=131% |
| 5640 | 5'\_3'\_RNA\_POLYMERASE\_ACTIVITY(GO:0034062) |  | 49 | 0.58 | 1.05 | 0.141 | 0.358 | 1.000 | 22286 | tags=82%, list=38%, signal=131% |
| 5641 | HOMOLOGOUS\_RECOMBINATION(GO:0035825) |  | 50 | 0.63 | 1.05 | 0.141 | 0.358 | 1.000 | 15966 | tags=54%, list=27%, signal=74% |
| 5642 | RECIPROCAL\_MEIOTIC\_RECOMBINATION(GO:0007131) |  | 50 | 0.63 | 1.05 | 0.141 | 0.358 | 1.000 | 15966 | tags=54%, list=27%, signal=74% |
| 5643 | VITAMIN\_BINDING(GO:0019842) |  | 138 | 0.63 | 1.05 | 0.141 | 0.358 | 1.000 | 17465 | tags=49%, list=30%, signal=69% |
| 5644 | OXIDOREDUCTASE\_ACTIVITY\_\_ACTING\_ON\_A\_SULFUR\_GROUP\_OF\_DONORS\_\_NAD(P)\_AS\_ACCEPTOR(GO:0016668) |  | 16 | 0.66 | 1.05 | 0.141 | 0.358 | 1.000 | 7054 | tags=13%, list=12%, signal=14% |
| 5645 | CHAPERONE\_MEDIATED\_PROTEIN\_FOLDING(GO:0061077) |  | 65 | 0.60 | 1.05 | 0.141 | 0.358 | 1.000 | 22131 | tags=77%, list=37%, signal=123% |
| 5646 | CULLIN\_FAMILY\_PROTEIN\_BINDING(GO:0097602) |  | 18 | 0.59 | 1.05 | 0.141 | 0.358 | 1.000 | 24144 | tags=100%, list=41%, signal=169% |
| 5647 | RESPONSE\_TO\_TEMPERATURE\_STIMULUS(GO:0009266) |  | 220 | 0.62 | 1.05 | 0.141 | 0.358 | 1.000 | 17646 | tags=54%, list=30%, signal=76% |
| 5648 | LATERAL\_ELEMENT(GO:0000800) |  | 18 | 0.65 | 1.05 | 0.481 | 0.358 | 1.000 | 12034 | tags=39%, list=20%, signal=49% |
| 5649 | REGULATION\_OF\_CENTROSOME\_DUPLICATION(GO:0010824) |  | 39 | 0.60 | 1.05 | 0.141 | 0.358 | 1.000 | 21117 | tags=77%, list=36%, signal=120% |
| 5650 | RIBONUCLEOPROTEIN\_COMPLEX\_ASSEMBLY(GO:0022618) |  | 223 | 0.59 | 1.05 | 0.141 | 0.358 | 1.000 | 23262 | tags=90%, list=39%, signal=147% |
| 5651 | SIGNAL\_TRANSDUCER\_\_DOWNSTREAM\_OF\_RECEPTOR\_\_WITH\_SERINE\_THREONINE\_KINASE\_ACTIVITY(GO:0004702) |  | 69 | 0.60 | 1.05 | 0.141 | 0.358 | 1.000 | 18373 | tags=54%, list=31%, signal=78% |
| 5652 | METHYLATION\_DEPENDENT\_CHROMATIN\_SILENCING(GO:0006346) |  | 20 | 0.60 | 1.05 | 0.000 | 0.358 | 1.000 | 22226 | tags=65%, list=38%, signal=104% |
| 5653 | BLOC\_COMPLEX(GO:0031082) |  | 22 | 0.60 | 1.05 | 0.141 | 0.358 | 1.000 | 21510 | tags=77%, list=36%, signal=121% |
| 5654 | STRIATED\_MUSCLE\_CELL\_APOPTOTIC\_PROCESS(GO:0010658) |  | 57 | 0.59 | 1.05 | 0.141 | 0.358 | 1.000 | 15300 | tags=28%, list=26%, signal=38% |
| 5655 | MODULATION\_BY\_VIRUS\_OF\_HOST\_PROCESS(GO:0019054) |  | 26 | 0.60 | 1.05 | 0.141 | 0.358 | 1.000 | 22440 | tags=77%, list=38%, signal=124% |
| 5656 | DNA\_POLYMERASE\_BINDING(GO:0070182) |  | 16 | 0.59 | 1.05 | 0.141 | 0.358 | 1.000 | 22616 | tags=88%, list=38%, signal=142% |
| 5657 | CENTROMERE\_COMPLEX\_ASSEMBLY(GO:0034508) |  | 54 | 0.62 | 1.05 | 0.141 | 0.358 | 1.000 | 17348 | tags=59%, list=29%, signal=84% |
| 5658 | NUCLEAR\_PERIPHERY(GO:0034399) |  | 130 | 0.62 | 1.05 | 0.141 | 0.358 | 1.000 | 18036 | tags=62%, list=30%, signal=89% |
| 5659 | BLOC\_1\_COMPLEX(GO:0031083) |  | 17 | 0.61 | 1.05 | 0.141 | 0.358 | 1.000 | 21510 | tags=76%, list=36%, signal=120% |
| 5660 | ERYTHROCYTE\_HOMEOSTASIS(GO:0034101) |  | 116 | 0.62 | 1.05 | 0.141 | 0.358 | 1.000 | 12443 | tags=32%, list=21%, signal=40% |
| 5661 | RRNA\_PROCESSING(GO:0006364) |  | 261 | 0.58 | 1.05 | 0.141 | 0.358 | 1.000 | 23272 | tags=87%, list=39%, signal=143% |
| 5662 | REGULATION\_OF\_ADAPTIVE\_IMMUNE\_RESPONSE\_BASED\_ON\_SOMATIC\_RECOMBINATION\_OF\_IMMUNE\_RECEPTORS\_BUILT\_FROM\_IMMUNOGLOBULIN\_SUPERFAMILY\_DOMAINS(GO:0002822) |  | 117 | 0.62 | 1.05 | 0.141 | 0.358 | 1.000 | 19789 | tags=50%, list=33%, signal=74% |
| 5663 | TRNA\_WOBBLE\_BASE\_MODIFICATION(GO:0002097) |  | 17 | 0.63 | 1.05 | 0.141 | 0.358 | 1.000 | 19255 | tags=82%, list=33%, signal=122% |
| 5664 | POSITIVE\_REGULATION\_OF\_TELOMERE\_CAPPING(GO:1904355) |  | 17 | 0.60 | 1.05 | 0.141 | 0.358 | 1.000 | 20903 | tags=76%, list=35%, signal=118% |
| 5665 | IMMUNOGLOBULIN\_PRODUCTION\_INVOLVED\_IN\_IMMUNOGLOBULIN\_MEDIATED\_IMMUNE\_RESPONSE(GO:0002381) |  | 48 | 0.63 | 1.05 | 0.141 | 0.358 | 1.000 | 19540 | tags=67%, list=33%, signal=99% |
| 5666 | POSITIVE\_REGULATION\_OF\_CYTOKINE\_MEDIATED\_SIGNALING\_PATHWAY(GO:0001961) |  | 45 | 0.63 | 1.05 | 0.141 | 0.358 | 1.000 | 16892 | tags=47%, list=29%, signal=65% |
| 5667 | REGULATION\_OF\_STRIATED\_MUSCLE\_CELL\_APOPTOTIC\_PROCESS(GO:0010662) |  | 53 | 0.59 | 1.05 | 0.141 | 0.358 | 1.000 | 17567 | tags=30%, list=30%, signal=43% |
| 5668 | MRNA\_PROCESSING(GO:0006397) |  | 504 | 0.61 | 1.05 | 0.141 | 0.358 | 1.000 | 20318 | tags=72%, list=34%, signal=109% |
| 5669 | SPERM\_CAPACITATION(GO:0048240) |  | 27 | 0.60 | 1.05 | 0.166 | 0.358 | 1.000 | 23421 | tags=70%, list=40%, signal=116% |
| 5670 | CCR4\_NOT\_COMPLEX(GO:0030014) |  | 16 | 0.63 | 1.05 | 0.141 | 0.358 | 1.000 | 16956 | tags=69%, list=29%, signal=96% |
| 5671 | REGULATION\_OF\_MITOCHONDRIAL\_DEPOLARIZATION(GO:0051900) |  | 19 | 0.63 | 1.05 | 0.141 | 0.358 | 1.000 | 10861 | tags=32%, list=18%, signal=39% |
| 5672 | POSITIVE\_REGULATION\_OF\_PROTEIN\_OLIGOMERIZATION(GO:0032461) |  | 25 | 0.68 | 1.05 | 0.307 | 0.358 | 1.000 | 9419 | tags=20%, list=16%, signal=24% |
| 5673 | REGULATION\_OF\_HISTONE\_METHYLATION(GO:0031060) |  | 60 | 0.61 | 1.05 | 0.141 | 0.358 | 1.000 | 19377 | tags=62%, list=33%, signal=92% |
| 5674 | REGULATION\_OF\_NATURAL\_KILLER\_CELL\_MEDIATED\_IMMUNITY(GO:0002715) |  | 36 | 0.61 | 1.05 | 0.141 | 0.358 | 1.000 | 15869 | tags=47%, list=27%, signal=65% |
| 5675 | RESPONSE\_TO\_TOPOLOGICALLY\_INCORRECT\_PROTEIN(GO:0035966) |  | 191 | 0.61 | 1.05 | 0.141 | 0.358 | 1.000 | 19291 | tags=51%, list=33%, signal=76% |
| 5676 | POSITIVE\_REGULATION\_OF\_ADAPTIVE\_IMMUNE\_RESPONSE\_BASED\_ON\_SOMATIC\_RECOMBINATION\_OF\_IMMUNE\_RECEPTORS\_BUILT\_FROM\_IMMUNOGLOBULIN\_SUPERFAMILY\_DOMAINS(GO:0002824) |  | 76 | 0.62 | 1.05 | 0.141 | 0.358 | 1.000 | 15082 | tags=39%, list=26%, signal=53% |
| 5677 | SECRETORY\_GRANULE\_ORGANIZATION(GO:0033363) |  | 31 | 0.64 | 1.05 | 0.141 | 0.358 | 1.000 | 18268 | tags=58%, list=31%, signal=84% |
| 5678 | NCRNA\_3'\_END\_PROCESSING(GO:0043628) |  | 27 | 0.58 | 1.05 | 0.141 | 0.358 | 1.000 | 23883 | tags=85%, list=40%, signal=143% |
| 5679 | REGULATION\_OF\_NUCLEOBASE\_CONTAINING\_COMPOUND\_TRANSPORT(GO:0032239) |  | 16 | 0.61 | 1.05 | 0.141 | 0.358 | 1.000 | 19571 | tags=81%, list=33%, signal=121% |
| 5680 | NUCLEAR\_TRANSCRIPTIONAL\_REPRESSOR\_COMPLEX(GO:0090568) |  | 29 | 0.61 | 1.05 | 0.000 | 0.357 | 1.000 | 21257 | tags=79%, list=36%, signal=124% |
| 5681 | PROTEIN\_SERINE\_THREONINE\_KINASE\_INHIBITOR\_ACTIVITY(GO:0030291) |  | 33 | 0.64 | 1.05 | 0.141 | 0.357 | 1.000 | 10693 | tags=21%, list=18%, signal=26% |
| 5682 | NUCLEOTIDE\_EXCISION\_REPAIR\_\_DNA\_DAMAGE\_RECOGNITION(GO:0000715) |  | 23 | 0.59 | 1.05 | 0.141 | 0.357 | 1.000 | 23234 | tags=91%, list=39%, signal=150% |
| 5683 | VITAMIN\_D\_RECEPTOR\_BINDING(GO:0042809) |  | 15 | 0.60 | 1.05 | 0.141 | 0.357 | 1.000 | 23415 | tags=93%, list=40%, signal=154% |
| 5684 | TETRAPYRROLE\_METABOLIC\_PROCESS(GO:0033013) |  | 60 | 0.60 | 1.05 | 0.141 | 0.357 | 1.000 | 18441 | tags=55%, list=31%, signal=80% |
| 5685 | POSITIVE\_REGULATION\_OF\_CHEMOTAXIS(GO:0050921) |  | 129 | 0.64 | 1.05 | 0.141 | 0.357 | 1.000 | 13967 | tags=28%, list=24%, signal=36% |
| 5686 | IRON\_ION\_TRANSPORT(GO:0006826) |  | 60 | 0.61 | 1.05 | 0.141 | 0.357 | 1.000 | 15375 | tags=30%, list=26%, signal=40% |
| 5687 | POSITIVE\_REGULATION\_OF\_B\_CELL\_ACTIVATION(GO:0050871) |  | 123 | 0.62 | 1.05 | 0.141 | 0.357 | 1.000 | 16856 | tags=28%, list=29%, signal=39% |
| 5688 | NUCLEAR\_CHROMOSOME\_\_TELOMERIC\_REGION(GO:0000784) |  | 123 | 0.60 | 1.05 | 0.141 | 0.357 | 1.000 | 20722 | tags=70%, list=35%, signal=107% |
| 5689 | MITOCHONDRIAL\_TRANSLATIONAL\_TERMINATION(GO:0070126) |  | 86 | 0.59 | 1.05 | 0.141 | 0.357 | 1.000 | 23709 | tags=94%, list=40%, signal=157% |
| 5690 | ACETYLCHOLINE\_GATED\_CHANNEL\_COMPLEX(GO:0005892) |  | 17 | 0.59 | 1.05 | 0.141 | 0.357 | 1.000 | 21485 | tags=71%, list=36%, signal=111% |
| 5691 | ENDORIBONUCLEASE\_ACTIVITY(GO:0004521) |  | 61 | 0.59 | 1.05 | 0.141 | 0.357 | 1.000 | 20828 | tags=67%, list=35%, signal=104% |
| 5692 | UBIQUITIN\_CONJUGATING\_ENZYME\_ACTIVITY(GO:0061631) |  | 27 | 0.61 | 1.05 | 0.141 | 0.357 | 1.000 | 20078 | tags=85%, list=34%, signal=129% |
| 5693 | TRANSLATIONAL\_ELONGATION(GO:0006414) |  | 132 | 0.58 | 1.05 | 0.141 | 0.357 | 1.000 | 23709 | tags=88%, list=40%, signal=146% |
| 5694 | POSITIVE\_REGULATION\_OF\_TRANSLATIONAL\_INITIATION(GO:0045948) |  | 30 | 0.60 | 1.05 | 0.141 | 0.357 | 1.000 | 18192 | tags=53%, list=31%, signal=77% |
| 5695 | NEGATIVE\_REGULATION\_OF\_DEFENSE\_RESPONSE\_TO\_VIRUS(GO:0050687) |  | 24 | 0.60 | 1.05 | 0.141 | 0.357 | 1.000 | 10185 | tags=21%, list=17%, signal=25% |
| 5696 | PURINE\_NTP\_DEPENDENT\_HELICASE\_ACTIVITY(GO:0070035) |  | 97 | 0.58 | 1.05 | 0.141 | 0.357 | 1.000 | 23692 | tags=87%, list=40%, signal=144% |
| 5697 | ATP\_DEPENDENT\_HELICASE\_ACTIVITY(GO:0008026) |  | 97 | 0.58 | 1.05 | 0.141 | 0.357 | 1.000 | 23692 | tags=87%, list=40%, signal=144% |
| 5698 | ESTABLISHMENT\_OF\_VESICLE\_LOCALIZATION(GO:0051650) |  | 233 | 0.64 | 1.05 | 0.141 | 0.357 | 1.000 | 14770 | tags=34%, list=25%, signal=46% |
| 5699 | STORE\_OPERATED\_CALCIUM\_ENTRY(GO:0002115) |  | 15 | 0.62 | 1.05 | 0.141 | 0.357 | 1.000 | 18040 | tags=53%, list=31%, signal=77% |
| 5700 | REGULATION\_OF\_CALCINEURIN\_NFAT\_SIGNALING\_CASCADE(GO:0070884) |  | 17 | 0.65 | 1.05 | 0.141 | 0.357 | 1.000 | 17187 | tags=65%, list=29%, signal=91% |
| 5701 | REGULATION\_OF\_CALCINEURIN\_MEDIATED\_SIGNALING(GO:0106056) |  | 17 | 0.65 | 1.05 | 0.141 | 0.357 | 1.000 | 17187 | tags=65%, list=29%, signal=91% |
| 5702 | DIPEPTIDASE\_ACTIVITY(GO:0016805) |  | 15 | 0.62 | 1.05 | 0.310 | 0.357 | 1.000 | 19324 | tags=67%, list=33%, signal=99% |
| 5703 | RESPONSE\_TO\_UNFOLDED\_PROTEIN(GO:0006986) |  | 172 | 0.61 | 1.05 | 0.141 | 0.357 | 1.000 | 19291 | tags=51%, list=33%, signal=75% |
| 5704 | CELLULAR\_RESPONSE\_TO\_CADMIUM\_ION(GO:0071276) |  | 31 | 0.60 | 1.05 | 0.141 | 0.357 | 1.000 | 17102 | tags=42%, list=29%, signal=59% |
| 5705 | PROTEIN\_CONTAINING\_COMPLEX\_DISASSEMBLY(GO:0032984) |  | 307 | 0.61 | 1.05 | 0.141 | 0.357 | 1.000 | 20010 | tags=64%, list=34%, signal=96% |
| 5706 | PYRIMIDINE\_NUCLEOTIDE\_METABOLIC\_PROCESS(GO:0006220) |  | 46 | 0.59 | 1.05 | 0.141 | 0.357 | 1.000 | 23753 | tags=89%, list=40%, signal=149% |
| 5707 | RNA\_POLYMERASE\_III\_ACTIVITY(GO:0001056) |  | 15 | 0.60 | 1.05 | 0.141 | 0.357 | 1.000 | 22286 | tags=93%, list=38%, signal=150% |
| 5708 | OXIDOREDUCTASE\_ACTIVITY\_\_ACTING\_ON\_NAD(P)H(GO:0016651) |  | 111 | 0.60 | 1.05 | 0.141 | 0.357 | 1.000 | 20227 | tags=57%, list=34%, signal=86% |
| 5709 | RETROGRADE\_VESICLE\_MEDIATED\_TRANSPORT\_\_GOLGI\_TO\_ER(GO:0006890) |  | 83 | 0.59 | 1.05 | 0.141 | 0.357 | 1.000 | 18157 | tags=52%, list=31%, signal=75% |
| 5710 | NUCLEAR\_MATRIX(GO:0016363) |  | 105 | 0.62 | 1.05 | 0.141 | 0.357 | 1.000 | 15649 | tags=51%, list=26%, signal=70% |
| 5711 | RRNA\_METABOLIC\_PROCESS(GO:0016072) |  | 296 | 0.58 | 1.05 | 0.141 | 0.357 | 1.000 | 23364 | tags=87%, list=40%, signal=143% |
| 5712 | REGULATORY\_RNA\_BINDING(GO:0061980) |  | 31 | 0.60 | 1.05 | 0.141 | 0.357 | 1.000 | 22555 | tags=84%, list=38%, signal=136% |
| 5713 | PROTEIN\_SERINE\_THREONINE\_PHOSPHATASE\_ACTIVITY(GO:0004722) |  | 70 | 0.61 | 1.05 | 0.141 | 0.357 | 1.000 | 20733 | tags=70%, list=35%, signal=108% |
| 5714 | MITOCHONDRIAL\_TRANSLATIONAL\_ELONGATION(GO:0070125) |  | 84 | 0.59 | 1.05 | 0.141 | 0.357 | 1.000 | 23709 | tags=94%, list=40%, signal=157% |
| 5715 | LIVER\_MORPHOGENESIS(GO:0072576) |  | 18 | 0.67 | 1.05 | 0.141 | 0.357 | 1.000 | 15437 | tags=44%, list=26%, signal=60% |
| 5716 | 7\_METHYLGUANOSINE\_MRNA\_CAPPING(GO:0006370) |  | 33 | 0.59 | 1.05 | 0.141 | 0.357 | 1.000 | 21089 | tags=76%, list=36%, signal=118% |
| 5717 | INNER\_MITOCHONDRIAL\_MEMBRANE\_ORGANIZATION(GO:0007007) |  | 42 | 0.59 | 1.05 | 0.141 | 0.357 | 1.000 | 22419 | tags=88%, list=38%, signal=142% |
| 5718 | NEGATIVE\_REGULATION\_OF\_MUSCLE\_CELL\_DIFFERENTIATION(GO:0051148) |  | 56 | 0.64 | 1.05 | 0.141 | 0.357 | 1.000 | 19572 | tags=59%, list=33%, signal=88% |
| 5719 | MONONUCLEAR\_CELL\_DIFFERENTIATION(GO:1903131) |  | 33 | 0.63 | 1.05 | 0.141 | 0.357 | 1.000 | 12040 | tags=30%, list=20%, signal=38% |
| 5720 | MONOCYTE\_DIFFERENTIATION(GO:0030224) |  | 33 | 0.63 | 1.05 | 0.141 | 0.357 | 1.000 | 12040 | tags=30%, list=20%, signal=38% |
| 5721 | LIPOPROTEIN\_METABOLIC\_PROCESS(GO:0042157) |  | 196 | 0.60 | 1.05 | 0.141 | 0.357 | 1.000 | 21014 | tags=54%, list=36%, signal=83% |
| 5722 | POSITIVE\_REGULATION\_OF\_MULTI\_ORGANISM\_PROCESS(GO:0043902) |  | 174 | 0.62 | 1.05 | 0.141 | 0.357 | 1.000 | 15831 | tags=33%, list=27%, signal=45% |
| 5723 | HISTONE\_H3\_ACETYLATION(GO:0043966) |  | 59 | 0.61 | 1.05 | 0.141 | 0.357 | 1.000 | 15634 | tags=53%, list=26%, signal=71% |
| 5724 | ERROR\_FREE\_TRANSLESION\_SYNTHESIS(GO:0070987) |  | 19 | 0.61 | 1.05 | 0.000 | 0.357 | 1.000 | 17484 | tags=68%, list=30%, signal=97% |
| 5725 | PROTEIN\_TARGETING(GO:0006605) |  | 390 | 0.63 | 1.05 | 0.141 | 0.357 | 1.000 | 16724 | tags=43%, list=28%, signal=59% |
| 5726 | REGULATION\_OF\_PROTEIN\_MODIFICATION\_BY\_SMALL\_PROTEIN\_CONJUGATION\_OR\_REMOVAL(GO:1903320) |  | 211 | 0.62 | 1.05 | 0.141 | 0.357 | 1.000 | 17110 | tags=48%, list=29%, signal=67% |
| 5727 | CYCLIN\_DEPENDENT\_PROTEIN\_KINASE\_ACTIVITY(GO:0097472) |  | 30 | 0.61 | 1.05 | 0.141 | 0.357 | 1.000 | 16370 | tags=50%, list=28%, signal=69% |
| 5728 | RESPONSE\_TO\_INTERFERON\_ALPHA(GO:0035455) |  | 20 | 0.64 | 1.05 | 0.141 | 0.357 | 1.000 | 18946 | tags=50%, list=32%, signal=74% |
| 5729 | NEGATIVE\_REGULATION\_OF\_ORGANELLE\_ASSEMBLY(GO:1902116) |  | 36 | 0.60 | 1.05 | 0.141 | 0.357 | 1.000 | 20748 | tags=75%, list=35%, signal=115% |
| 5730 | MITOCHONDRIAL\_LARGE\_RIBOSOMAL\_SUBUNIT(GO:0005762) |  | 51 | 0.59 | 1.05 | 0.141 | 0.357 | 1.000 | 23709 | tags=98%, list=40%, signal=164% |
| 5731 | ORGANELLAR\_LARGE\_RIBOSOMAL\_SUBUNIT(GO:0000315) |  | 51 | 0.59 | 1.05 | 0.141 | 0.357 | 1.000 | 23709 | tags=98%, list=40%, signal=164% |
| 5732 | REGULATION\_OF\_CYTOPLASMIC\_TRANSLATION(GO:2000765) |  | 23 | 0.61 | 1.05 | 0.141 | 0.357 | 1.000 | 20011 | tags=74%, list=34%, signal=112% |
| 5733 | NEGATIVE\_REGULATION\_OF\_ENDOTHELIAL\_CELL\_MIGRATION(GO:0010596) |  | 92 | 0.63 | 1.05 | 0.141 | 0.357 | 1.000 | 14175 | tags=28%, list=24%, signal=37% |
| 5734 | REGULATION\_OF\_SMAD\_PROTEIN\_SIGNAL\_TRANSDUCTION(GO:0060390) |  | 16 | 0.62 | 1.05 | 0.141 | 0.357 | 1.000 | 12990 | tags=44%, list=22%, signal=56% |
| 5735 | POSITIVE\_REGULATION\_OF\_AMINO\_ACID\_TRANSPORT(GO:0051957) |  | 17 | 0.63 | 1.05 | 0.141 | 0.357 | 1.000 | 19928 | tags=71%, list=34%, signal=106% |
| 5736 | G\_PROTEIN\_COUPLED\_ACETYLCHOLINE\_RECEPTOR\_SIGNALING\_PATHWAY(GO:0007213) |  | 16 | 0.63 | 1.05 | 0.307 | 0.357 | 1.000 | 6523 | tags=6%, list=11%, signal=7% |
| 5737 | REGULATION\_OF\_INTRINSIC\_APOPTOTIC\_SIGNALING\_PATHWAY\_IN\_RESPONSE\_TO\_DNA\_DAMAGE\_BY\_P53\_CLASS\_MEDIATOR(GO:1902165) |  | 15 | 0.64 | 1.05 | 0.141 | 0.357 | 1.000 | 17869 | tags=53%, list=30%, signal=76% |
| 5738 | NUA4\_HISTONE\_ACETYLTRANSFERASE\_COMPLEX(GO:0035267) |  | 18 | 0.60 | 1.05 | 0.141 | 0.357 | 1.000 | 20745 | tags=83%, list=35%, signal=128% |
| 5739 | H4\_H2A\_HISTONE\_ACETYLTRANSFERASE\_COMPLEX(GO:0043189) |  | 18 | 0.60 | 1.05 | 0.141 | 0.357 | 1.000 | 20745 | tags=83%, list=35%, signal=128% |
| 5740 | NEGATIVE\_REGULATION\_OF\_CELLULAR\_RESPONSE\_TO\_TRANSFORMING\_GROWTH\_FACTOR\_BETA\_STIMULUS(GO:1903845) |  | 76 | 0.61 | 1.05 | 0.141 | 0.357 | 1.000 | 14781 | tags=29%, list=25%, signal=39% |
| 5741 | REGULATION\_OF\_AXON\_REGENERATION(GO:0048679) |  | 25 | 0.62 | 1.05 | 0.141 | 0.357 | 1.000 | 7975 | tags=28%, list=13%, signal=32% |
| 5742 | HISTONE\_H3\_K9\_TRIMETHYLATION(GO:0036124) |  | 15 | 0.61 | 1.05 | 0.141 | 0.357 | 1.000 | 22863 | tags=87%, list=39%, signal=141% |
| 5743 | SMALL\_NUCLEAR\_RIBONUCLEOPROTEIN\_COMPLEX(GO:0030532) |  | 70 | 0.59 | 1.05 | 0.141 | 0.357 | 1.000 | 22418 | tags=81%, list=38%, signal=131% |
| 5744 | PYRIMIDINE\_CONTAINING\_COMPOUND\_BIOSYNTHETIC\_PROCESS(GO:0072528) |  | 42 | 0.59 | 1.05 | 0.141 | 0.357 | 1.000 | 19853 | tags=71%, list=34%, signal=107% |
| 5745 | VITAMIN\_D\_METABOLIC\_PROCESS(GO:0042359) |  | 23 | 0.65 | 1.05 | 0.141 | 0.357 | 1.000 | 13642 | tags=39%, list=23%, signal=51% |
| 5746 | NEGATIVE\_REGULATION\_OF\_RNA\_SPLICING(GO:0033119) |  | 25 | 0.62 | 1.05 | 0.000 | 0.357 | 1.000 | 21605 | tags=92%, list=37%, signal=145% |
| 5747 | HISTONE\_H4\_K16\_ACETYLATION(GO:0043984) |  | 18 | 0.62 | 1.05 | 0.000 | 0.357 | 1.000 | 20665 | tags=89%, list=35%, signal=137% |
| 5748 | MITOTIC\_SISTER\_CHROMATID\_COHESION(GO:0007064) |  | 24 | 0.62 | 1.05 | 0.000 | 0.357 | 1.000 | 20691 | tags=92%, list=35%, signal=141% |
| 5749 | POSITIVE\_REGULATION\_OF\_RESPONSE\_TO\_CYTOKINE\_STIMULUS(GO:0060760) |  | 49 | 0.63 | 1.05 | 0.141 | 0.357 | 1.000 | 16892 | tags=45%, list=29%, signal=63% |
| 5750 | GTP\_RHO\_BINDING(GO:0017049) |  | 19 | 0.61 | 1.05 | 0.141 | 0.357 | 1.000 | 15036 | tags=47%, list=25%, signal=63% |
| 5751 | FILOPODIUM\_ASSEMBLY(GO:0046847) |  | 57 | 0.63 | 1.05 | 0.141 | 0.357 | 1.000 | 16153 | tags=44%, list=27%, signal=60% |
| 5752 | POSITIVE\_REGULATION\_OF\_CARDIAC\_MUSCLE\_CELL\_DIFFERENTIATION(GO:2000727) |  | 26 | 0.64 | 1.05 | 0.141 | 0.357 | 1.000 | 17343 | tags=42%, list=29%, signal=60% |
| 5753 | POSITIVE\_REGULATION\_OF\_CELL\_DIVISION(GO:0051781) |  | 83 | 0.63 | 1.05 | 0.141 | 0.357 | 1.000 | 17160 | tags=45%, list=29%, signal=63% |
| 5754 | CULLIN\_RING\_UBIQUITIN\_LIGASE\_COMPLEX(GO:0031461) |  | 175 | 0.58 | 1.05 | 0.141 | 0.357 | 1.000 | 19678 | tags=59%, list=33%, signal=89% |
| 5755 | DNA\_REPLICATION\_INDEPENDENT\_NUCLEOSOME\_ORGANIZATION(GO:0034724) |  | 54 | 0.61 | 1.05 | 0.141 | 0.357 | 1.000 | 20331 | tags=70%, list=34%, signal=107% |
| 5756 | RNA\_SPLICING(GO:0008380) |  | 431 | 0.61 | 1.05 | 0.141 | 0.357 | 1.000 | 20318 | tags=73%, list=34%, signal=110% |
| 5757 | REGULATION\_OF\_INCLUSION\_BODY\_ASSEMBLY(GO:0090083) |  | 17 | 0.61 | 1.05 | 0.141 | 0.357 | 1.000 | 18684 | tags=53%, list=32%, signal=77% |
| 5758 | ACETYLTRANSFERASE\_COMPLEX(GO:1902493) |  | 92 | 0.59 | 1.05 | 0.141 | 0.357 | 1.000 | 21028 | tags=79%, list=36%, signal=123% |
| 5759 | PROTEIN\_ACETYLTRANSFERASE\_COMPLEX(GO:0031248) |  | 92 | 0.59 | 1.05 | 0.141 | 0.357 | 1.000 | 21028 | tags=79%, list=36%, signal=123% |
| 5760 | P\_BODY(GO:0000932) |  | 82 | 0.59 | 1.05 | 0.141 | 0.357 | 1.000 | 20904 | tags=74%, list=35%, signal=115% |
| 5761 | AMYLOID\_BETA\_METABOLIC\_PROCESS(GO:0050435) |  | 34 | 0.60 | 1.05 | 0.141 | 0.357 | 1.000 | 21613 | tags=74%, list=37%, signal=116% |
| 5762 | METALLOENDOPEPTIDASE\_INHIBITOR\_ACTIVITY(GO:0008191) |  | 16 | 0.66 | 1.05 | 0.141 | 0.357 | 1.000 | 13202 | tags=50%, list=22%, signal=64% |
| 5763 | CELLULAR\_POLYSACCHARIDE\_METABOLIC\_PROCESS(GO:0044264) |  | 95 | 0.62 | 1.05 | 0.141 | 0.357 | 1.000 | 17672 | tags=49%, list=30%, signal=70% |
| 5764 | DNA\_REPLICATION\_INDEPENDENT\_NUCLEOSOME\_ASSEMBLY(GO:0006336) |  | 53 | 0.61 | 1.05 | 0.141 | 0.357 | 1.000 | 20331 | tags=70%, list=34%, signal=106% |
| 5765 | INCLUSION\_BODY\_ASSEMBLY(GO:0070841) |  | 23 | 0.60 | 1.05 | 0.141 | 0.357 | 1.000 | 19177 | tags=48%, list=32%, signal=71% |
| 5766 | PROTON\_TRANSPORTING\_ATP\_SYNTHASE\_COMPLEX(GO:0045259) |  | 23 | 0.60 | 1.05 | 0.141 | 0.357 | 1.000 | 22093 | tags=87%, list=37%, signal=139% |
| 5767 | RNA\_PHOSPHODIESTER\_BOND\_HYDROLYSIS\_\_ENDONUCLEOLYTIC(GO:0090502) |  | 72 | 0.59 | 1.05 | 0.141 | 0.357 | 1.000 | 20828 | tags=69%, list=35%, signal=107% |
| 5768 | POSITIVE\_REGULATION\_OF\_PROTEIN\_MODIFICATION\_BY\_SMALL\_PROTEIN\_CONJUGATION\_OR\_REMOVAL(GO:1903322) |  | 124 | 0.62 | 1.05 | 0.141 | 0.357 | 1.000 | 15458 | tags=42%, list=26%, signal=57% |
| 5769 | HEMATOPOIETIC\_STEM\_CELL\_DIFFERENTIATION(GO:0060218) |  | 114 | 0.60 | 1.05 | 0.141 | 0.357 | 1.000 | 17088 | tags=34%, list=29%, signal=48% |
| 5770 | REGULATION\_OF\_MONOCYTE\_DIFFERENTIATION(GO:0045655) |  | 18 | 0.66 | 1.05 | 0.310 | 0.357 | 1.000 | 15132 | tags=39%, list=26%, signal=52% |
| 5771 | POLYSACCHARIDE\_METABOLIC\_PROCESS(GO:0005976) |  | 108 | 0.62 | 1.05 | 0.141 | 0.357 | 1.000 | 17672 | tags=47%, list=30%, signal=67% |
| 5772 | NEUTRAL\_AMINO\_ACID\_TRANSMEMBRANE\_TRANSPORTER\_ACTIVITY(GO:0015175) |  | 33 | 0.62 | 1.05 | 0.141 | 0.357 | 1.000 | 11579 | tags=24%, list=20%, signal=30% |
| 5773 | PEPTIDE\_CATABOLIC\_PROCESS(GO:0043171) |  | 31 | 0.59 | 1.05 | 0.141 | 0.357 | 1.000 | 19324 | tags=48%, list=33%, signal=72% |
| 5774 | 2\_OXOGLUTARATE\_METABOLIC\_PROCESS(GO:0006103) |  | 19 | 0.62 | 1.05 | 0.141 | 0.357 | 1.000 | 20708 | tags=79%, list=35%, signal=121% |
| 5775 | REGULATION\_OF\_MITOCHONDRIAL\_TRANSLATION(GO:0070129) |  | 18 | 0.61 | 1.05 | 0.141 | 0.357 | 1.000 | 23074 | tags=100%, list=39%, signal=164% |
| 5776 | INSULIN\_SECRETION\_INVOLVED\_IN\_CELLULAR\_RESPONSE\_TO\_GLUCOSE\_STIMULUS(GO:0035773) |  | 62 | 0.59 | 1.05 | 0.141 | 0.357 | 1.000 | 18767 | tags=58%, list=32%, signal=85% |
| 5777 | DAMAGED\_DNA\_BINDING(GO:0003684) |  | 66 | 0.60 | 1.05 | 0.141 | 0.357 | 1.000 | 19305 | tags=73%, list=33%, signal=108% |
| 5778 | POSITIVE\_REGULATION\_OF\_PROTEIN\_BINDING(GO:0032092) |  | 88 | 0.60 | 1.05 | 0.141 | 0.357 | 1.000 | 18685 | tags=49%, list=32%, signal=71% |
| 5779 | SOMATIC\_RECOMBINATION\_OF\_IMMUNOGLOBULIN\_GENE\_SEGMENTS(GO:0016447) |  | 46 | 0.63 | 1.05 | 0.141 | 0.357 | 1.000 | 19540 | tags=67%, list=33%, signal=101% |
| 5780 | AROMATIC\_AMINO\_ACID\_FAMILY\_METABOLIC\_PROCESS(GO:0009072) |  | 29 | 0.64 | 1.05 | 0.141 | 0.357 | 1.000 | 19569 | tags=62%, list=33%, signal=93% |
| 5781 | LIGASE\_ACTIVITY\_\_FORMING\_CARBON\_OXYGEN\_BONDS(GO:0016875) |  | 49 | 0.60 | 1.04 | 0.141 | 0.357 | 1.000 | 22212 | tags=86%, list=38%, signal=137% |
| 5782 | AMINOACYL\_TRNA\_LIGASE\_ACTIVITY(GO:0004812) |  | 49 | 0.60 | 1.04 | 0.141 | 0.357 | 1.000 | 22212 | tags=86%, list=38%, signal=137% |
| 5783 | NEGATIVE\_REGULATION\_OF\_T\_CELL\_MEDIATED\_IMMUNITY(GO:0002710) |  | 15 | 0.65 | 1.04 | 0.141 | 0.357 | 1.000 | 15529 | tags=40%, list=26%, signal=54% |
| 5784 | STRIATED\_MUSCLE\_ADAPTATION(GO:0014888) |  | 54 | 0.60 | 1.04 | 0.141 | 0.357 | 1.000 | 10309 | tags=19%, list=17%, signal=22% |
| 5785 | POSITIVE\_REGULATION\_OF\_IMMUNOGLOBULIN\_MEDIATED\_IMMUNE\_RESPONSE(GO:0002891) |  | 32 | 0.63 | 1.04 | 0.141 | 0.357 | 1.000 | 19540 | tags=59%, list=33%, signal=89% |
| 5786 | POSITIVE\_REGULATION\_OF\_B\_CELL\_MEDIATED\_IMMUNITY(GO:0002714) |  | 32 | 0.63 | 1.04 | 0.141 | 0.357 | 1.000 | 19540 | tags=59%, list=33%, signal=89% |
| 5787 | MEGAKARYOCYTE\_DEVELOPMENT(GO:0035855) |  | 15 | 0.64 | 1.04 | 0.141 | 0.357 | 1.000 | 19272 | tags=67%, list=33%, signal=99% |
| 5788 | REGULATION\_OF\_ADAPTIVE\_IMMUNE\_RESPONSE(GO:0002819) |  | 128 | 0.61 | 1.04 | 0.141 | 0.357 | 1.000 | 19789 | tags=48%, list=33%, signal=71% |
| 5789 | MITOCHONDRIAL\_PROTON\_TRANSPORTING\_ATP\_SYNTHASE\_COMPLEX(GO:0005753) |  | 22 | 0.60 | 1.04 | 0.141 | 0.357 | 1.000 | 22093 | tags=86%, list=37%, signal=138% |
| 5790 | DEOXYRIBOSE\_PHOSPHATE\_CATABOLIC\_PROCESS(GO:0046386) |  | 20 | 0.60 | 1.04 | 0.141 | 0.357 | 1.000 | 23049 | tags=95%, list=39%, signal=156% |
| 5791 | REGULATION\_OF\_PHOSPHOLIPID\_BIOSYNTHETIC\_PROCESS(GO:0071071) |  | 16 | 0.64 | 1.04 | 0.141 | 0.357 | 1.000 | 12916 | tags=38%, list=22%, signal=48% |
| 5792 | REGULATION\_OF\_RAS\_PROTEIN\_SIGNAL\_TRANSDUCTION(GO:0046578) |  | 222 | 0.62 | 1.04 | 0.141 | 0.357 | 1.000 | 16107 | tags=39%, list=27%, signal=53% |
| 5793 | LYSINE\_N\_METHYLTRANSFERASE\_ACTIVITY(GO:0016278) |  | 62 | 0.60 | 1.04 | 0.141 | 0.357 | 1.000 | 21531 | tags=79%, list=36%, signal=124% |
| 5794 | RESPONSE\_TO\_LEAD\_ION(GO:0010288) |  | 23 | 0.64 | 1.04 | 0.141 | 0.357 | 1.000 | 9534 | tags=30%, list=16%, signal=36% |
| 5795 | RESOLUTION\_OF\_MEIOTIC\_RECOMBINATION\_INTERMEDIATES(GO:0000712) |  | 15 | 0.64 | 1.04 | 0.141 | 0.357 | 1.000 | 15966 | tags=60%, list=27%, signal=82% |
| 5796 | POSITIVE\_REGULATION\_OF\_ATPASE\_ACTIVITY(GO:0032781) |  | 51 | 0.60 | 1.04 | 0.141 | 0.357 | 1.000 | 21500 | tags=73%, list=36%, signal=114% |
| 5797 | CHEMOKINE\_MEDIATED\_SIGNALING\_PATHWAY(GO:0070098) |  | 81 | 0.64 | 1.04 | 0.141 | 0.357 | 1.000 | 15381 | tags=27%, list=26%, signal=37% |
| 5798 | MATURATION\_OF\_SSU\_RRNA\_FROM\_TRICISTRONIC\_RRNA\_TRANSCRIPT\_(SSU\_RRNA\_\_5.8S\_RRNA\_\_LSU\_RRNA)(GO:0000462) |  | 38 | 0.58 | 1.04 | 0.141 | 0.357 | 1.000 | 24083 | tags=95%, list=41%, signal=160% |
| 5799 | POSITIVE\_REGULATION\_OF\_T\_CELL\_MIGRATION(GO:2000406) |  | 24 | 0.63 | 1.04 | 0.141 | 0.357 | 1.000 | 10429 | tags=21%, list=18%, signal=25% |
| 5800 | SYNAPTONEMAL\_COMPLEX\_ORGANIZATION(GO:0070193) |  | 24 | 0.59 | 1.04 | 0.141 | 0.357 | 1.000 | 20078 | tags=58%, list=34%, signal=88% |
| 5801 | DENDRITIC\_SHAFT(GO:0043198) |  | 36 | 0.63 | 1.04 | 0.141 | 0.357 | 1.000 | 15752 | tags=39%, list=27%, signal=53% |
| 5802 | SAGA\_TYPE\_COMPLEX(GO:0070461) |  | 27 | 0.60 | 1.04 | 0.141 | 0.357 | 1.000 | 19654 | tags=70%, list=33%, signal=105% |
| 5803 | INTRACELLULAR\_TRANSPORT\_OF\_VIRUS(GO:0075733) |  | 57 | 0.61 | 1.04 | 0.141 | 0.357 | 1.000 | 21893 | tags=81%, list=37%, signal=128% |
| 5804 | MITOTIC\_METAPHASE\_PLATE\_CONGRESSION(GO:0007080) |  | 43 | 0.58 | 1.04 | 0.141 | 0.357 | 1.000 | 21353 | tags=77%, list=36%, signal=120% |
| 5805 | TRANSLATIONAL\_TERMINATION(GO:0006415) |  | 100 | 0.59 | 1.04 | 0.141 | 0.357 | 1.000 | 23709 | tags=93%, list=40%, signal=155% |
| 5806 | INTRACELLULAR\_PROTEIN\_TRANSMEMBRANE\_TRANSPORT(GO:0065002) |  | 49 | 0.57 | 1.04 | 0.141 | 0.357 | 1.000 | 19319 | tags=65%, list=33%, signal=97% |
| 5807 | OXIDOREDUCTASE\_ACTIVITY\_\_ACTING\_ON\_A\_SULFUR\_GROUP\_OF\_DONORS(GO:0016667) |  | 61 | 0.61 | 1.04 | 0.141 | 0.357 | 1.000 | 18144 | tags=51%, list=31%, signal=73% |
| 5808 | SPLICEOSOMAL\_TRI\_SNRNP\_COMPLEX(GO:0097526) |  | 29 | 0.58 | 1.04 | 0.141 | 0.357 | 1.000 | 23254 | tags=79%, list=39%, signal=131% |
| 5809 | MUSCLE\_ADAPTATION(GO:0043500) |  | 107 | 0.59 | 1.04 | 0.141 | 0.357 | 1.000 | 18262 | tags=38%, list=31%, signal=55% |
| 5810 | ATPASE\_COUPLED\_ION\_TRANSMEMBRANE\_TRANSPORTER\_ACTIVITY(GO:0042625) |  | 66 | 0.61 | 1.04 | 0.141 | 0.357 | 1.000 | 18171 | tags=55%, list=31%, signal=79% |
| 5811 | ACTIVE\_ION\_TRANSMEMBRANE\_TRANSPORTER\_ACTIVITY(GO:0022853) |  | 66 | 0.61 | 1.04 | 0.141 | 0.357 | 1.000 | 18171 | tags=55%, list=31%, signal=79% |
| 5812 | POSITIVE\_REGULATION\_OF\_CELLULAR\_AMIDE\_METABOLIC\_PROCESS(GO:0034250) |  | 143 | 0.61 | 1.04 | 0.141 | 0.357 | 1.000 | 18355 | tags=45%, list=31%, signal=66% |
| 5813 | LIVER\_REGENERATION(GO:0097421) |  | 35 | 0.63 | 1.04 | 0.141 | 0.357 | 1.000 | 17154 | tags=57%, list=29%, signal=80% |
| 5814 | BLASTOCYST\_FORMATION(GO:0001825) |  | 31 | 0.60 | 1.04 | 0.141 | 0.357 | 1.000 | 22904 | tags=77%, list=39%, signal=126% |
| 5815 | QUINONE\_BINDING(GO:0048038) |  | 17 | 0.64 | 1.04 | 0.141 | 0.357 | 1.000 | 14288 | tags=41%, list=24%, signal=54% |
| 5816 | SNORNA\_BINDING(GO:0030515) |  | 31 | 0.59 | 1.04 | 0.141 | 0.357 | 1.000 | 20650 | tags=74%, list=35%, signal=114% |
| 5817 | P53\_BINDING(GO:0002039) |  | 69 | 0.59 | 1.04 | 0.141 | 0.357 | 1.000 | 22085 | tags=71%, list=37%, signal=113% |
| 5818 | GLUCAN\_CATABOLIC\_PROCESS(GO:0009251) |  | 26 | 0.59 | 1.04 | 0.141 | 0.357 | 1.000 | 19534 | tags=54%, list=33%, signal=80% |
| 5819 | CELLULAR\_POLYSACCHARIDE\_CATABOLIC\_PROCESS(GO:0044247) |  | 26 | 0.59 | 1.04 | 0.141 | 0.357 | 1.000 | 19534 | tags=54%, list=33%, signal=80% |
| 5820 | NUCLEOID(GO:0009295) |  | 43 | 0.59 | 1.04 | 0.141 | 0.357 | 1.000 | 23756 | tags=95%, list=40%, signal=159% |
| 5821 | MITOCHONDRIAL\_NUCLEOID(GO:0042645) |  | 43 | 0.59 | 1.04 | 0.141 | 0.357 | 1.000 | 23756 | tags=95%, list=40%, signal=159% |
| 5822 | ENDODEOXYRIBONUCLEASE\_ACTIVITY(GO:0004520) |  | 50 | 0.58 | 1.04 | 0.141 | 0.357 | 1.000 | 21853 | tags=74%, list=37%, signal=117% |
| 5823 | REGULATION\_OF\_B\_CELL\_MEDIATED\_IMMUNITY(GO:0002712) |  | 46 | 0.62 | 1.04 | 0.141 | 0.357 | 1.000 | 19789 | tags=52%, list=33%, signal=78% |
| 5824 | REGULATION\_OF\_IMMUNOGLOBULIN\_MEDIATED\_IMMUNE\_RESPONSE(GO:0002889) |  | 46 | 0.62 | 1.04 | 0.141 | 0.357 | 1.000 | 19789 | tags=52%, list=33%, signal=78% |
| 5825 | LIPOPROTEIN\_BIOSYNTHETIC\_PROCESS(GO:0042158) |  | 162 | 0.59 | 1.04 | 0.141 | 0.357 | 1.000 | 20749 | tags=54%, list=35%, signal=83% |
| 5826 | 'DE\_NOVO'\_POSTTRANSLATIONAL\_PROTEIN\_FOLDING(GO:0051084) |  | 22 | 0.60 | 1.04 | 0.141 | 0.357 | 1.000 | 22131 | tags=86%, list=37%, signal=138% |
| 5827 | PROTEIN\_QUALITY\_CONTROL\_FOR\_MISFOLDED\_OR\_INCOMPLETELY\_SYNTHESIZED\_PROTEINS(GO:0006515) |  | 26 | 0.58 | 1.04 | 0.141 | 0.357 | 1.000 | 25069 | tags=100%, list=42%, signal=174% |
| 5828 | POSITIVE\_REGULATION\_OF\_TELOMERASE\_RNA\_LOCALIZATION\_TO\_CAJAL\_BODY(GO:1904874) |  | 15 | 0.63 | 1.04 | 0.000 | 0.357 | 1.000 | 20650 | tags=93%, list=35%, signal=143% |
| 5829 | PROTEIN\_LYSINE\_N\_METHYLTRANSFERASE\_ACTIVITY(GO:0016279) |  | 61 | 0.60 | 1.04 | 0.141 | 0.357 | 1.000 | 21531 | tags=79%, list=36%, signal=124% |
| 5830 | ESTABLISHMENT\_OF\_PROTEIN\_LOCALIZATION\_TO\_CHROMOSOME(GO:0070199) |  | 23 | 0.59 | 1.04 | 0.141 | 0.357 | 1.000 | 23132 | tags=91%, list=39%, signal=150% |
| 5831 | MULTI\_ORGANISM\_LOCALIZATION(GO:1902579) |  | 72 | 0.60 | 1.04 | 0.141 | 0.357 | 1.000 | 19834 | tags=71%, list=34%, signal=106% |
| 5832 | MULTI\_ORGANISM\_TRANSPORT(GO:0044766) |  | 72 | 0.60 | 1.04 | 0.141 | 0.357 | 1.000 | 19834 | tags=71%, list=34%, signal=106% |
| 5833 | CELLULAR\_RESPONSE\_TO\_REACTIVE\_OXYGEN\_SPECIES(GO:0034614) |  | 160 | 0.60 | 1.04 | 0.141 | 0.357 | 1.000 | 19731 | tags=52%, list=33%, signal=78% |
| 5834 | NEURON\_DEATH\_IN\_RESPONSE\_TO\_OXIDATIVE\_STRESS(GO:0036475) |  | 25 | 0.62 | 1.04 | 0.141 | 0.357 | 1.000 | 18880 | tags=60%, list=32%, signal=88% |
| 5835 | MLL1\_COMPLEX(GO:0071339) |  | 31 | 0.59 | 1.04 | 0.141 | 0.357 | 1.000 | 24277 | tags=97%, list=41%, signal=164% |
| 5836 | MLL1\_2\_COMPLEX(GO:0044665) |  | 31 | 0.59 | 1.04 | 0.141 | 0.357 | 1.000 | 24277 | tags=97%, list=41%, signal=164% |
| 5837 | NUCLEAR\_MIGRATION(GO:0007097) |  | 19 | 0.61 | 1.04 | 0.141 | 0.357 | 1.000 | 22456 | tags=84%, list=38%, signal=136% |
| 5838 | PROTEIN\_LIPIDATION(GO:0006497) |  | 157 | 0.59 | 1.04 | 0.141 | 0.357 | 1.000 | 20749 | tags=54%, list=35%, signal=83% |
| 5839 | POSITIVE\_REGULATION\_OF\_LYMPHOCYTE\_APOPTOTIC\_PROCESS(GO:0070230) |  | 19 | 0.63 | 1.04 | 0.141 | 0.357 | 1.000 | 13000 | tags=16%, list=22%, signal=20% |
| 5840 | PROTEIN\_MODIFICATION\_BY\_SMALL\_PROTEIN\_REMOVAL(GO:0070646) |  | 316 | 0.61 | 1.04 | 0.141 | 0.357 | 1.000 | 19867 | tags=60%, list=34%, signal=90% |
| 5841 | B\_CELL\_ACTIVATION\_INVOLVED\_IN\_IMMUNE\_RESPONSE(GO:0002312) |  | 66 | 0.62 | 1.04 | 0.141 | 0.357 | 1.000 | 19138 | tags=52%, list=32%, signal=76% |
| 5842 | MITOCHONDRIAL\_INTERMEMBRANE\_SPACE(GO:0005758) |  | 78 | 0.58 | 1.04 | 0.141 | 0.357 | 1.000 | 24414 | tags=92%, list=41%, signal=157% |
| 5843 | NEGATIVE\_REGULATION\_OF\_ENDOCYTOSIS(GO:0045806) |  | 49 | 0.62 | 1.04 | 0.141 | 0.357 | 1.000 | 9520 | tags=12%, list=16%, signal=15% |
| 5844 | DEOXYRIBONUCLEOTIDE\_CATABOLIC\_PROCESS(GO:0009264) |  | 19 | 0.60 | 1.04 | 0.141 | 0.357 | 1.000 | 23049 | tags=95%, list=39%, signal=155% |
| 5845 | RNA\_POLYMERASE\_II\_TRANSCRIPTION\_COREPRESSOR\_ACTIVITY(GO:0001106) |  | 27 | 0.63 | 1.04 | 0.141 | 0.357 | 1.000 | 19184 | tags=63%, list=32%, signal=93% |
| 5846 | MEDIATOR\_COMPLEX(GO:0016592) |  | 38 | 0.59 | 1.04 | 0.141 | 0.357 | 1.000 | 18276 | tags=66%, list=31%, signal=95% |
| 5847 | DEOXYRIBONUCLEASE\_ACTIVITY(GO:0004536) |  | 69 | 0.58 | 1.04 | 0.141 | 0.357 | 1.000 | 21853 | tags=74%, list=37%, signal=117% |
| 5848 | HISTONE\_ACETYLTRANSFERASE\_COMPLEX(GO:0000123) |  | 82 | 0.59 | 1.04 | 0.141 | 0.357 | 1.000 | 21028 | tags=78%, list=36%, signal=121% |
| 5849 | RIBOSOMAL\_LARGE\_SUBUNIT\_ASSEMBLY(GO:0000027) |  | 30 | 0.59 | 1.04 | 0.141 | 0.357 | 1.000 | 22470 | tags=83%, list=38%, signal=134% |
| 5850 | IMMUNOGLOBULIN\_PRODUCTION(GO:0002377) |  | 153 | 0.60 | 1.04 | 0.141 | 0.357 | 1.000 | 19222 | tags=35%, list=33%, signal=51% |
| 5851 | REGULATION\_OF\_TYPE\_I\_INTERFERON\_MEDIATED\_SIGNALING\_PATHWAY(GO:0060338) |  | 46 | 0.59 | 1.04 | 0.141 | 0.357 | 1.000 | 21073 | tags=59%, list=36%, signal=91% |
| 5852 | RESPONSE\_TO\_NICOTINE(GO:0035094) |  | 49 | 0.62 | 1.04 | 0.141 | 0.357 | 1.000 | 12401 | tags=35%, list=21%, signal=44% |
| 5853 | RIBOSOMAL\_LARGE\_SUBUNIT\_BIOGENESIS(GO:0042273) |  | 70 | 0.58 | 1.04 | 0.141 | 0.357 | 1.000 | 23262 | tags=86%, list=39%, signal=141% |
| 5854 | SOMATIC\_DIVERSIFICATION\_OF\_IMMUNOGLOBULINS\_INVOLVED\_IN\_IMMUNE\_RESPONSE(GO:0002208) |  | 42 | 0.63 | 1.04 | 0.141 | 0.357 | 1.000 | 20770 | tags=74%, list=35%, signal=114% |
| 5855 | SOMATIC\_RECOMBINATION\_OF\_IMMUNOGLOBULIN\_GENES\_INVOLVED\_IN\_IMMUNE\_RESPONSE(GO:0002204) |  | 42 | 0.63 | 1.04 | 0.141 | 0.357 | 1.000 | 20770 | tags=74%, list=35%, signal=114% |
| 5856 | ISOTYPE\_SWITCHING(GO:0045190) |  | 42 | 0.63 | 1.04 | 0.141 | 0.357 | 1.000 | 20770 | tags=74%, list=35%, signal=114% |
| 5857 | TROPHECTODERMAL\_CELL\_DIFFERENTIATION(GO:0001829) |  | 16 | 0.61 | 1.04 | 0.141 | 0.357 | 1.000 | 22159 | tags=81%, list=37%, signal=130% |
| 5858 | LUNG\_CELL\_DIFFERENTIATION(GO:0060479) |  | 23 | 0.67 | 1.04 | 0.141 | 0.357 | 1.000 | 12189 | tags=26%, list=21%, signal=33% |
| 5859 | GLUCOSE\_IMPORT\_IN\_RESPONSE\_TO\_INSULIN\_STIMULUS(GO:0044381) |  | 25 | 0.62 | 1.04 | 0.141 | 0.357 | 1.000 | 20597 | tags=52%, list=35%, signal=80% |
| 5860 | DETECTION\_OF\_CHEMICAL\_STIMULUS\_INVOLVED\_IN\_SENSORY\_PERCEPTION(GO:0050907) |  | 432 | 0.53 | 1.04 | 0.309 | 0.357 | 1.000 | 13514 | tags=7%, list=23%, signal=10% |
| 5861 | ANTIGEN\_PROCESSING\_AND\_PRESENTATION\_OF\_PEPTIDE\_ANTIGEN\_VIA\_MHC\_CLASS\_II(GO:0002495) |  | 98 | 0.61 | 1.04 | 0.141 | 0.357 | 1.000 | 19775 | tags=58%, list=33%, signal=87% |
| 5862 | CELLULAR\_CARBOHYDRATE\_CATABOLIC\_PROCESS(GO:0044275) |  | 45 | 0.58 | 1.04 | 0.141 | 0.357 | 1.000 | 19592 | tags=56%, list=33%, signal=83% |
| 5863 | NUCLEAR\_ENVELOPE\_REASSEMBLY(GO:0031468) |  | 18 | 0.59 | 1.04 | 0.141 | 0.357 | 1.000 | 23801 | tags=94%, list=40%, signal=158% |
| 5864 | RESPONSE\_TO\_TYPE\_I\_INTERFERON(GO:0034340) |  | 90 | 0.62 | 1.04 | 0.141 | 0.357 | 1.000 | 8063 | tags=12%, list=14%, signal=14% |
| 5865 | CYCLIN\_DEPENDENT\_PROTEIN\_SERINE\_THREONINE\_KINASE\_ACTIVITY(GO:0004693) |  | 29 | 0.60 | 1.04 | 0.141 | 0.357 | 1.000 | 16370 | tags=48%, list=28%, signal=67% |
| 5866 | TRANSPORT\_OF\_VIRUS(GO:0046794) |  | 60 | 0.60 | 1.04 | 0.141 | 0.357 | 1.000 | 18624 | tags=67%, list=31%, signal=97% |
| 5867 | PROTON\_TRANSPORTING\_ATP\_SYNTHASE\_ACTIVITY\_\_ROTATIONAL\_MECHANISM(GO:0046933) |  | 17 | 0.59 | 1.04 | 0.141 | 0.357 | 1.000 | 21617 | tags=76%, list=37%, signal=120% |
| 5868 | NEGATIVE\_REGULATION\_OF\_CARDIOCYTE\_DIFFERENTIATION(GO:1905208) |  | 22 | 0.67 | 1.04 | 0.141 | 0.357 | 1.000 | 16108 | tags=50%, list=27%, signal=69% |
| 5869 | POSITIVE\_REGULATION\_OF\_APOPTOTIC\_SIGNALING\_PATHWAY(GO:2001235) |  | 178 | 0.63 | 1.04 | 0.141 | 0.357 | 1.000 | 15681 | tags=34%, list=27%, signal=46% |
| 5870 | RHODOPSIN\_MEDIATED\_SIGNALING\_PATHWAY(GO:0016056) |  | 30 | 0.59 | 1.04 | 0.141 | 0.357 | 1.000 | 17568 | tags=60%, list=30%, signal=85% |
| 5871 | MODULATION\_BY\_HOST\_OF\_VIRAL\_GENOME\_REPLICATION(GO:0044827) |  | 15 | 0.62 | 1.04 | 0.168 | 0.357 | 1.000 | 22174 | tags=93%, list=37%, signal=149% |
| 5872 | TOXIN\_METABOLIC\_PROCESS(GO:0009404) |  | 23 | 0.66 | 1.04 | 0.141 | 0.357 | 1.000 | 14745 | tags=39%, list=25%, signal=52% |
| 5873 | RESPONSE\_TO\_ZINC\_ION(GO:0010043) |  | 51 | 0.61 | 1.04 | 0.141 | 0.357 | 1.000 | 15885 | tags=45%, list=27%, signal=62% |
| 5874 | SERINE\_PHOSPHORYLATION\_OF\_STAT\_PROTEIN(GO:0042501) |  | 27 | 0.60 | 1.04 | 0.141 | 0.357 | 1.000 | 20653 | tags=44%, list=35%, signal=68% |
| 5875 | PYRIMIDINE\_NUCLEOTIDE\_BIOSYNTHETIC\_PROCESS(GO:0006221) |  | 32 | 0.59 | 1.04 | 0.310 | 0.357 | 1.000 | 20715 | tags=75%, list=35%, signal=115% |
| 5876 | 5'\_3'\_EXONUCLEASE\_ACTIVITY(GO:0008409) |  | 15 | 0.62 | 1.04 | 0.340 | 0.357 | 1.000 | 19308 | tags=73%, list=33%, signal=109% |
| 5877 | METHYLTRANSFERASE\_COMPLEX(GO:0034708) |  | 93 | 0.59 | 1.04 | 0.141 | 0.357 | 1.000 | 21089 | tags=77%, list=36%, signal=120% |
| 5878 | SMALL\_RIBOSOMAL\_SUBUNIT(GO:0015935) |  | 73 | 0.58 | 1.04 | 0.141 | 0.357 | 1.000 | 24601 | tags=89%, list=42%, signal=152% |
| 5879 | CLATHRIN\_COATED\_PIT(GO:0005905) |  | 67 | 0.63 | 1.04 | 0.141 | 0.357 | 1.000 | 15436 | tags=40%, list=26%, signal=54% |
| 5880 | CRISTAE\_FORMATION(GO:0042407) |  | 32 | 0.60 | 1.04 | 0.141 | 0.357 | 1.000 | 22419 | tags=91%, list=38%, signal=146% |
| 5881 | INTRA\_S\_DNA\_DAMAGE\_CHECKPOINT(GO:0031573) |  | 16 | 0.62 | 1.04 | 0.166 | 0.357 | 1.000 | 22004 | tags=88%, list=37%, signal=139% |
| 5882 | NADH\_DEHYDROGENASE\_(UBIQUINONE)\_ACTIVITY(GO:0008137) |  | 47 | 0.58 | 1.04 | 0.141 | 0.357 | 1.000 | 24412 | tags=91%, list=41%, signal=156% |
| 5883 | NADH\_DEHYDROGENASE\_(QUINONE)\_ACTIVITY(GO:0050136) |  | 47 | 0.58 | 1.04 | 0.141 | 0.357 | 1.000 | 24412 | tags=91%, list=41%, signal=156% |
| 5884 | DENDRITIC\_CELL\_MIGRATION(GO:0036336) |  | 26 | 0.63 | 1.04 | 0.141 | 0.357 | 1.000 | 13828 | tags=23%, list=23%, signal=30% |
| 5885 | MAINTENANCE\_OF\_PROTEIN\_LOCALIZATION\_IN\_ORGANELLE(GO:0072595) |  | 40 | 0.60 | 1.04 | 0.141 | 0.357 | 1.000 | 22333 | tags=80%, list=38%, signal=128% |
| 5886 | NADH\_DEHYDROGENASE\_ACTIVITY(GO:0003954) |  | 49 | 0.58 | 1.04 | 0.141 | 0.357 | 1.000 | 24636 | tags=92%, list=42%, signal=157% |
| 5887 | NAD\_DEPENDENT\_HISTONE\_DEACETYLASE\_ACTIVITY(GO:0017136) |  | 16 | 0.62 | 1.04 | 0.307 | 0.357 | 1.000 | 19177 | tags=63%, list=32%, signal=92% |
| 5888 | NEGATIVE\_REGULATION\_OF\_CALCIUM\_ION\_TRANSMEMBRANE\_TRANSPORT(GO:1903170) |  | 42 | 0.60 | 1.04 | 0.141 | 0.357 | 1.000 | 15565 | tags=33%, list=26%, signal=45% |
| 5889 | CELLULAR\_RESPONSE\_TO\_HEAT(GO:0034605) |  | 113 | 0.62 | 1.04 | 0.141 | 0.357 | 1.000 | 19220 | tags=65%, list=33%, signal=97% |
| 5890 | RIBOSOME\_ASSEMBLY(GO:0042255) |  | 62 | 0.58 | 1.04 | 0.141 | 0.357 | 1.000 | 23262 | tags=84%, list=39%, signal=138% |
| 5891 | POSITIVE\_REGULATION\_OF\_PROTEIN\_AUTOPHOSPHORYLATION(GO:0031954) |  | 27 | 0.62 | 1.04 | 0.141 | 0.357 | 1.000 | 9498 | tags=19%, list=16%, signal=22% |
| 5892 | CHROMATIN\_REMODELING\_AT\_CENTROMERE(GO:0031055) |  | 45 | 0.62 | 1.04 | 0.141 | 0.357 | 1.000 | 17348 | tags=58%, list=29%, signal=82% |
| 5893 | REGULATION\_OF\_ESTABLISHMENT\_OR\_MAINTENANCE\_OF\_CELL\_POLARITY(GO:0032878) |  | 24 | 0.62 | 1.04 | 0.141 | 0.357 | 1.000 | 20968 | tags=71%, list=35%, signal=110% |
| 5894 | PROTEIN\_DEUBIQUITINATION(GO:0016579) |  | 300 | 0.61 | 1.04 | 0.141 | 0.357 | 1.000 | 19867 | tags=58%, list=34%, signal=87% |
| 5895 | HISTONE\_METHYLTRANSFERASE\_COMPLEX(GO:0035097) |  | 71 | 0.59 | 1.04 | 0.141 | 0.357 | 1.000 | 21077 | tags=76%, list=36%, signal=118% |
| 5896 | PHOSPHATIDYLINOSITOL\_3\_KINASE\_COMPLEX(GO:0005942) |  | 21 | 0.60 | 1.04 | 0.141 | 0.357 | 1.000 | 22821 | tags=71%, list=39%, signal=116% |
| 5897 | PHOSPHOTRANSFERASE\_ACTIVITY\_\_FOR\_OTHER\_SUBSTITUTED\_PHOSPHATE\_GROUPS(GO:0016780) |  | 18 | 0.59 | 1.04 | 0.141 | 0.357 | 1.000 | 16304 | tags=56%, list=28%, signal=77% |
| 5898 | POSITIVE\_REGULATION\_OF\_POSITIVE\_CHEMOTAXIS(GO:0050927) |  | 25 | 0.64 | 1.04 | 0.141 | 0.357 | 1.000 | 6456 | tags=20%, list=11%, signal=22% |
| 5899 | REGULATION\_OF\_OXIDATIVE\_STRESS\_INDUCED\_NEURON\_DEATH(GO:1903203) |  | 24 | 0.63 | 1.04 | 0.141 | 0.357 | 1.000 | 18880 | tags=63%, list=32%, signal=92% |
| 5900 | CYTOSOLIC\_PART(GO:0044445) |  | 245 | 0.58 | 1.04 | 0.141 | 0.357 | 1.000 | 22772 | tags=70%, list=39%, signal=114% |
| 5901 | LIGAND\_GATED\_CALCIUM\_CHANNEL\_ACTIVITY(GO:0099604) |  | 24 | 0.63 | 1.04 | 0.141 | 0.357 | 1.000 | 19199 | tags=58%, list=32%, signal=86% |
| 5902 | POSITIVE\_REGULATION\_OF\_HISTONE\_H3\_K4\_METHYLATION(GO:0051571) |  | 17 | 0.61 | 1.04 | 0.000 | 0.357 | 1.000 | 19138 | tags=65%, list=32%, signal=96% |
| 5903 | POSITIVE\_REGULATION\_OF\_DNA\_RECOMBINATION(GO:0045911) |  | 27 | 0.64 | 1.04 | 0.141 | 0.357 | 1.000 | 12135 | tags=33%, list=21%, signal=42% |
| 5904 | BLASTOCYST\_DEVELOPMENT(GO:0001824) |  | 67 | 0.62 | 1.04 | 0.141 | 0.357 | 1.000 | 16141 | tags=55%, list=27%, signal=76% |
| 5905 | ORGANELLE\_ENVELOPE\_LUMEN(GO:0031970) |  | 86 | 0.58 | 1.04 | 0.141 | 0.357 | 1.000 | 24414 | tags=91%, list=41%, signal=154% |
| 5906 | RESPONSE\_TO\_MINERALOCORTICOID(GO:0051385) |  | 29 | 0.65 | 1.04 | 0.141 | 0.357 | 1.000 | 16848 | tags=52%, list=28%, signal=72% |
| 5907 | RESPONSE\_TO\_HEAT(GO:0009408) |  | 156 | 0.61 | 1.04 | 0.141 | 0.357 | 1.000 | 19294 | tags=63%, list=33%, signal=93% |
| 5908 | MULTI\_ORGANISM\_CELLULAR\_PROCESS(GO:0044764) |  | 46 | 0.62 | 1.04 | 0.141 | 0.357 | 1.000 | 17194 | tags=35%, list=29%, signal=49% |
| 5909 | POSITIVE\_REGULATION\_OF\_ACTIVATED\_T\_CELL\_PROLIFERATION(GO:0042104) |  | 25 | 0.62 | 1.04 | 0.307 | 0.357 | 1.000 | 21696 | tags=56%, list=37%, signal=88% |
| 5910 | MACROMOLECULE\_DEACYLATION(GO:0098732) |  | 110 | 0.58 | 1.04 | 0.141 | 0.357 | 1.000 | 22117 | tags=74%, list=37%, signal=117% |
| 5911 | SOMATIC\_DIVERSIFICATION\_OF\_IMMUNOGLOBULINS(GO:0016445) |  | 51 | 0.63 | 1.04 | 0.141 | 0.357 | 1.000 | 19540 | tags=69%, list=33%, signal=102% |
| 5912 | CELL\_KILLING(GO:0001906) |  | 138 | 0.59 | 1.04 | 0.141 | 0.357 | 1.000 | 16764 | tags=31%, list=28%, signal=43% |
| 5913 | REGULATION\_OF\_ERYTHROCYTE\_DIFFERENTIATION(GO:0045646) |  | 45 | 0.62 | 1.04 | 0.141 | 0.357 | 1.000 | 12195 | tags=36%, list=21%, signal=45% |
| 5914 | PROTEIN\_PHOSPHATASE\_TYPE\_2A\_COMPLEX(GO:0000159) |  | 21 | 0.59 | 1.04 | 0.141 | 0.357 | 1.000 | 20127 | tags=67%, list=34%, signal=101% |
| 5915 | PREASSEMBLY\_OF\_GPI\_ANCHOR\_IN\_ER\_MEMBRANE(GO:0016254) |  | 17 | 0.58 | 1.04 | 0.141 | 0.357 | 1.000 | 21843 | tags=71%, list=37%, signal=112% |
| 5916 | SPLICEOSOMAL\_SNRNP\_COMPLEX(GO:0097525) |  | 63 | 0.58 | 1.04 | 0.141 | 0.357 | 1.000 | 23254 | tags=83%, list=39%, signal=136% |
| 5917 | ANTIBIOTIC\_BIOSYNTHETIC\_PROCESS(GO:0017000) |  | 16 | 0.60 | 1.04 | 0.141 | 0.357 | 1.000 | 22859 | tags=75%, list=39%, signal=122% |
| 5918 | GLYCOGEN\_CATABOLIC\_PROCESS(GO:0005980) |  | 25 | 0.59 | 1.04 | 0.141 | 0.357 | 1.000 | 19534 | tags=52%, list=33%, signal=78% |
| 5919 | REGULATION\_OF\_ASTROCYTE\_DIFFERENTIATION(GO:0048710) |  | 27 | 0.64 | 1.04 | 0.310 | 0.357 | 1.000 | 14425 | tags=41%, list=24%, signal=54% |
| 5920 | DEACETYLASE\_ACTIVITY(GO:0019213) |  | 57 | 0.58 | 1.04 | 0.141 | 0.357 | 1.000 | 21405 | tags=72%, list=36%, signal=113% |
| 5921 | CHROMOSOME\_LOCALIZATION(GO:0050000) |  | 78 | 0.62 | 1.04 | 0.141 | 0.357 | 1.000 | 20205 | tags=74%, list=34%, signal=113% |
| 5922 | REGULATION\_OF\_CYTOKINESIS(GO:0032465) |  | 81 | 0.58 | 1.04 | 0.141 | 0.357 | 1.000 | 20837 | tags=63%, list=35%, signal=97% |
| 5923 | NEGATIVE\_REGULATION\_OF\_DENDRITE\_DEVELOPMENT(GO:2000171) |  | 28 | 0.62 | 1.04 | 0.141 | 0.357 | 1.000 | 13101 | tags=39%, list=22%, signal=50% |
| 5924 | ESTABLISHMENT\_OF\_CHROMOSOME\_LOCALIZATION(GO:0051303) |  | 77 | 0.62 | 1.04 | 0.141 | 0.357 | 1.000 | 20205 | tags=74%, list=34%, signal=112% |
| 5925 | RESPONSE\_TO\_ENDOPLASMIC\_RETICULUM\_STRESS(GO:0034976) |  | 273 | 0.60 | 1.04 | 0.141 | 0.357 | 1.000 | 19291 | tags=54%, list=33%, signal=80% |
| 5926 | TELOMERASE\_HOLOENZYME\_COMPLEX(GO:0005697) |  | 21 | 0.62 | 1.04 | 0.000 | 0.357 | 1.000 | 22365 | tags=100%, list=38%, signal=161% |
| 5927 | FIBROBLAST\_GROWTH\_FACTOR\_RECEPTOR\_SIGNALING\_PATHWAY(GO:0008543) |  | 115 | 0.62 | 1.04 | 0.141 | 0.357 | 1.000 | 16724 | tags=43%, list=28%, signal=59% |
| 5928 | PROTEIN\_K11\_LINKED\_UBIQUITINATION(GO:0070979) |  | 28 | 0.61 | 1.04 | 0.141 | 0.357 | 1.000 | 19018 | tags=79%, list=32%, signal=116% |
| 5929 | NUCLEOBASE\_CONTAINING\_SMALL\_MOLECULE\_INTERCONVERSION(GO:0015949) |  | 30 | 0.59 | 1.04 | 0.141 | 0.357 | 1.000 | 19656 | tags=73%, list=33%, signal=110% |
| 5930 | CARBON\_CARBON\_LYASE\_ACTIVITY(GO:0016830) |  | 55 | 0.58 | 1.04 | 0.141 | 0.357 | 1.000 | 19176 | tags=60%, list=32%, signal=89% |
| 5931 | REGULATION\_OF\_PROTEIN\_OLIGOMERIZATION(GO:0032459) |  | 43 | 0.65 | 1.04 | 0.141 | 0.357 | 1.000 | 12188 | tags=21%, list=21%, signal=26% |
| 5932 | REGULATION\_OF\_EPIDERMAL\_GROWTH\_FACTOR\_RECEPTOR\_SIGNALING\_PATHWAY(GO:0042058) |  | 86 | 0.60 | 1.04 | 0.141 | 0.357 | 1.000 | 17190 | tags=36%, list=29%, signal=51% |
| 5933 | DNA\_TEMPLATED\_TRANSCRIPTION\_\_ELONGATION(GO:0006354) |  | 131 | 0.57 | 1.04 | 0.141 | 0.357 | 1.000 | 22407 | tags=79%, list=38%, signal=126% |
| 5934 | ANTIGEN\_PROCESSING\_AND\_PRESENTATION\_OF\_EXOGENOUS\_PEPTIDE\_ANTIGEN\_VIA\_MHC\_CLASS\_II(GO:0019886) |  | 95 | 0.61 | 1.04 | 0.141 | 0.357 | 1.000 | 19775 | tags=59%, list=33%, signal=88% |
| 5935 | HISTONE\_H4\_K5\_ACETYLATION(GO:0043981) |  | 15 | 0.62 | 1.04 | 0.141 | 0.357 | 1.000 | 20665 | tags=87%, list=35%, signal=133% |
| 5936 | HISTONE\_H4\_K8\_ACETYLATION(GO:0043982) |  | 15 | 0.62 | 1.04 | 0.141 | 0.357 | 1.000 | 20665 | tags=87%, list=35%, signal=133% |
| 5937 | NEGATIVE\_REGULATION\_OF\_CARDIAC\_MUSCLE\_CELL\_DIFFERENTIATION(GO:2000726) |  | 18 | 0.66 | 1.04 | 0.141 | 0.357 | 1.000 | 16108 | tags=44%, list=27%, signal=61% |
| 5938 | UBIQUITIN\_LIKE\_PROTEIN\_CONJUGATING\_ENZYME\_ACTIVITY(GO:0061650) |  | 28 | 0.61 | 1.04 | 0.141 | 0.357 | 1.000 | 21796 | tags=89%, list=37%, signal=141% |
| 5939 | ANDROGEN\_METABOLIC\_PROCESS(GO:0008209) |  | 28 | 0.62 | 1.04 | 0.141 | 0.357 | 1.000 | 14935 | tags=39%, list=25%, signal=53% |
| 5940 | RIBOSOMAL\_SUBUNIT(GO:0044391) |  | 189 | 0.58 | 1.04 | 0.141 | 0.357 | 1.000 | 24309 | tags=91%, list=41%, signal=154% |
| 5941 | POTASSIUM\_CHANNEL\_REGULATOR\_ACTIVITY(GO:0015459) |  | 47 | 0.60 | 1.04 | 0.141 | 0.357 | 1.000 | 17599 | tags=47%, list=30%, signal=67% |
| 5942 | PROTEIN\_HORMONE\_RECEPTOR\_ACTIVITY(GO:0016500) |  | 16 | 0.64 | 1.04 | 0.141 | 0.357 | 1.000 | 16236 | tags=38%, list=27%, signal=52% |
| 5943 | ENDONUCLEASE\_ACTIVITY\_\_ACTIVE\_WITH\_EITHER\_RIBO\_\_OR\_DEOXYRIBONUCLEIC\_ACIDS\_AND\_PRODUCING\_5'\_PHOSPHOMONOESTERS(GO:0016893) |  | 42 | 0.60 | 1.04 | 0.000 | 0.357 | 1.000 | 20821 | tags=76%, list=35%, signal=118% |
| 5944 | MITOCHONDRIAL\_RESPIRATORY\_CHAIN\_COMPLEX\_ASSEMBLY(GO:0033108) |  | 99 | 0.58 | 1.04 | 0.141 | 0.357 | 1.000 | 24640 | tags=95%, list=42%, signal=162% |
| 5945 | REGULATION\_OF\_B\_CELL\_RECEPTOR\_SIGNALING\_PATHWAY(GO:0050855) |  | 21 | 0.63 | 1.04 | 0.141 | 0.357 | 1.000 | 16243 | tags=43%, list=27%, signal=59% |
| 5946 | SNRNA\_PROCESSING(GO:0016180) |  | 23 | 0.58 | 1.04 | 0.141 | 0.357 | 1.000 | 24009 | tags=96%, list=41%, signal=161% |
| 5947 | PROTEIN\_TRANSMEMBRANE\_TRANSPORT(GO:0071806) |  | 56 | 0.57 | 1.04 | 0.141 | 0.357 | 1.000 | 24252 | tags=84%, list=41%, signal=142% |
| 5948 | MRNA\_3'\_END\_PROCESSING(GO:0031124) |  | 90 | 0.58 | 1.04 | 0.141 | 0.357 | 1.000 | 21791 | tags=78%, list=37%, signal=123% |
| 5949 | SELECTIVE\_AUTOPHAGY(GO:0061912) |  | 43 | 0.59 | 1.04 | 0.141 | 0.357 | 1.000 | 22203 | tags=74%, list=38%, signal=119% |
| 5950 | GLUTAMINE\_METABOLIC\_PROCESS(GO:0006541) |  | 24 | 0.60 | 1.04 | 0.141 | 0.357 | 1.000 | 19853 | tags=75%, list=34%, signal=113% |
| 5951 | POSITIVE\_REGULATION\_OF\_PHAGOCYTOSIS(GO:0050766) |  | 47 | 0.60 | 1.04 | 0.141 | 0.357 | 1.000 | 9368 | tags=17%, list=16%, signal=20% |
| 5952 | PORE\_COMPLEX\_ASSEMBLY(GO:0046931) |  | 16 | 0.62 | 1.04 | 0.340 | 0.357 | 1.000 | 20194 | tags=81%, list=34%, signal=123% |
| 5953 | PYRIMIDINE\_DEOXYRIBONUCLEOTIDE\_METABOLIC\_PROCESS(GO:0009219) |  | 18 | 0.61 | 1.04 | 0.310 | 0.357 | 1.000 | 23049 | tags=94%, list=39%, signal=155% |
| 5954 | CAJAL\_BODY(GO:0015030) |  | 55 | 0.59 | 1.04 | 0.141 | 0.357 | 1.000 | 20666 | tags=75%, list=35%, signal=114% |
| 5955 | HISTONE\_H3\_DEACETYLATION(GO:0070932) |  | 20 | 0.60 | 1.04 | 0.307 | 0.357 | 1.000 | 19177 | tags=60%, list=32%, signal=89% |
| 5956 | PHOSPHATIDYLINOSITOL\_4\_PHOSPHATE\_BINDING(GO:0070273) |  | 21 | 0.60 | 1.04 | 0.141 | 0.357 | 1.000 | 20790 | tags=76%, list=35%, signal=117% |
| 5957 | GPI\_ANCHOR\_METABOLIC\_PROCESS(GO:0006505) |  | 35 | 0.57 | 1.04 | 0.141 | 0.357 | 1.000 | 23473 | tags=74%, list=40%, signal=123% |
| 5958 | PYRIMIDINE\_RIBONUCLEOTIDE\_METABOLIC\_PROCESS(GO:0009218) |  | 25 | 0.58 | 1.04 | 0.141 | 0.357 | 1.000 | 23753 | tags=84%, list=40%, signal=140% |
| 5959 | ORGANELLE\_DISASSEMBLY(GO:1903008) |  | 104 | 0.58 | 1.04 | 0.141 | 0.357 | 1.000 | 23032 | tags=83%, list=39%, signal=135% |
| 5960 | IONOTROPIC\_GLUTAMATE\_RECEPTOR\_BINDING(GO:0035255) |  | 27 | 0.60 | 1.04 | 0.141 | 0.357 | 1.000 | 20217 | tags=59%, list=34%, signal=90% |
| 5961 | VASCULAR\_ASSOCIATED\_SMOOTH\_MUSCLE\_CELL\_MIGRATION(GO:1904738) |  | 34 | 0.66 | 1.04 | 0.141 | 0.357 | 1.000 | 13419 | tags=35%, list=23%, signal=46% |
| 5962 | REGULATION\_OF\_VASCULAR\_ASSOCIATED\_SMOOTH\_MUSCLE\_CELL\_MIGRATION(GO:1904752) |  | 34 | 0.66 | 1.04 | 0.141 | 0.357 | 1.000 | 13419 | tags=35%, list=23%, signal=46% |
| 5963 | PATTERN\_RECOGNITION\_RECEPTOR\_SIGNALING\_PATHWAY(GO:0002221) |  | 182 | 0.61 | 1.04 | 0.141 | 0.357 | 1.000 | 17194 | tags=38%, list=29%, signal=54% |
| 5964 | POSITIVE\_REGULATION\_OF\_INNATE\_IMMUNE\_RESPONSE(GO:0045089) |  | 347 | 0.61 | 1.04 | 0.141 | 0.357 | 1.000 | 16724 | tags=38%, list=28%, signal=52% |
| 5965 | TRANS\_GOLGI\_NETWORK(GO:0005802) |  | 199 | 0.60 | 1.04 | 0.141 | 0.357 | 1.000 | 18561 | tags=45%, list=31%, signal=65% |
| 5966 | REGULATION\_OF\_ERBB\_SIGNALING\_PATHWAY(GO:1901184) |  | 99 | 0.60 | 1.04 | 0.141 | 0.357 | 1.000 | 17190 | tags=37%, list=29%, signal=53% |
| 5967 | POSITIVE\_REGULATION\_OF\_CAMP\_BIOSYNTHETIC\_PROCESS(GO:0030819) |  | 21 | 0.59 | 1.04 | 0.141 | 0.357 | 1.000 | 21587 | tags=62%, list=37%, signal=97% |
| 5968 | CHAPERONE\_MEDIATED\_PROTEIN\_COMPLEX\_ASSEMBLY(GO:0051131) |  | 19 | 0.60 | 1.04 | 0.141 | 0.357 | 1.000 | 20585 | tags=63%, list=35%, signal=97% |
| 5969 | FC\_GAMMA\_RECEPTOR\_SIGNALING\_PATHWAY(GO:0038094) |  | 132 | 0.57 | 1.04 | 0.141 | 0.357 | 1.000 | 18981 | tags=27%, list=32%, signal=39% |
| 5970 | SNRNA\_TRANSCRIPTION(GO:0009301) |  | 70 | 0.58 | 1.04 | 0.141 | 0.357 | 1.000 | 24644 | tags=94%, list=42%, signal=161% |
| 5971 | SNRNA\_TRANSCRIPTION\_BY\_RNA\_POLYMERASE\_II(GO:0042795) |  | 70 | 0.58 | 1.04 | 0.141 | 0.357 | 1.000 | 24644 | tags=94%, list=42%, signal=161% |
| 5972 | REGULATION\_OF\_FATTY\_ACID\_TRANSPORT(GO:2000191) |  | 23 | 0.66 | 1.04 | 0.141 | 0.357 | 1.000 | 11372 | tags=22%, list=19%, signal=27% |
| 5973 | ERYTHROCYTE\_DIFFERENTIATION(GO:0030218) |  | 107 | 0.61 | 1.04 | 0.141 | 0.357 | 1.000 | 12443 | tags=32%, list=21%, signal=40% |
| 5974 | HISTONE\_LYSINE\_N\_METHYLTRANSFERASE\_ACTIVITY(GO:0018024) |  | 47 | 0.60 | 1.04 | 0.141 | 0.357 | 1.000 | 23844 | tags=94%, list=40%, signal=157% |
| 5975 | NCRNA\_TRANSCRIPTION(GO:0098781) |  | 102 | 0.57 | 1.04 | 0.141 | 0.357 | 1.000 | 24746 | tags=92%, list=42%, signal=158% |
| 5976 | MITOCHONDRIAL\_DEPOLARIZATION(GO:0051882) |  | 22 | 0.62 | 1.04 | 0.141 | 0.357 | 1.000 | 10861 | tags=27%, list=18%, signal=33% |
| 5977 | ACETYLCHOLINE\_GATED\_CATION\_SELECTIVE\_CHANNEL\_ACTIVITY(GO:0022848) |  | 17 | 0.59 | 1.04 | 0.141 | 0.357 | 1.000 | 21485 | tags=71%, list=36%, signal=111% |
| 5978 | REGULATION\_OF\_EPIDERMAL\_CELL\_DIFFERENTIATION(GO:0045604) |  | 56 | 0.63 | 1.04 | 0.141 | 0.357 | 1.000 | 11480 | tags=23%, list=19%, signal=29% |
| 5979 | RRNA\_BINDING(GO:0019843) |  | 63 | 0.58 | 1.03 | 0.141 | 0.357 | 1.000 | 22455 | tags=81%, list=38%, signal=130% |
| 5980 | CLATHRIN\_COATED\_VESICLE\_MEMBRANE(GO:0030665) |  | 101 | 0.60 | 1.03 | 0.141 | 0.357 | 1.000 | 14383 | tags=29%, list=24%, signal=38% |
| 5981 | SNORNA\_METABOLIC\_PROCESS(GO:0016074) |  | 15 | 0.60 | 1.03 | 0.476 | 0.357 | 1.000 | 20650 | tags=73%, list=35%, signal=113% |
| 5982 | LARGE\_RIBOSOMAL\_SUBUNIT(GO:0015934) |  | 117 | 0.58 | 1.03 | 0.141 | 0.357 | 1.000 | 24309 | tags=94%, list=41%, signal=159% |
| 5983 | ACETYL\_COA\_METABOLIC\_PROCESS(GO:0006084) |  | 34 | 0.62 | 1.03 | 0.141 | 0.357 | 1.000 | 17637 | tags=53%, list=30%, signal=75% |
| 5984 | POSITIVE\_REGULATION\_OF\_UBIQUITIN\_PROTEIN\_TRANSFERASE\_ACTIVITY(GO:0051443) |  | 34 | 0.57 | 1.03 | 0.141 | 0.357 | 1.000 | 24447 | tags=91%, list=41%, signal=155% |
| 5985 | MAINTENANCE\_OF\_PROTEIN\_LOCATION\_IN\_NUCLEUS(GO:0051457) |  | 24 | 0.61 | 1.03 | 0.141 | 0.357 | 1.000 | 22333 | tags=88%, list=38%, signal=141% |
| 5986 | HISTONE\_DEACETYLASE\_ACTIVITY(GO:0004407) |  | 43 | 0.59 | 1.03 | 0.141 | 0.357 | 1.000 | 21405 | tags=77%, list=36%, signal=120% |
| 5987 | NUCLEAR\_EUCHROMATIN(GO:0005719) |  | 25 | 0.62 | 1.03 | 0.141 | 0.357 | 1.000 | 15684 | tags=52%, list=27%, signal=71% |
| 5988 | PROTEIN\_K48\_LINKED\_UBIQUITINATION(GO:0070936) |  | 52 | 0.61 | 1.03 | 0.309 | 0.357 | 1.000 | 19018 | tags=67%, list=32%, signal=99% |
| 5989 | PROTEIN\_DEACETYLASE\_ACTIVITY(GO:0033558) |  | 44 | 0.59 | 1.03 | 0.141 | 0.357 | 1.000 | 21405 | tags=77%, list=36%, signal=121% |
| 5990 | COATED\_VESICLE(GO:0030135) |  | 258 | 0.60 | 1.03 | 0.141 | 0.357 | 1.000 | 18127 | tags=44%, list=31%, signal=63% |
| 5991 | REGULATION\_OF\_RESPONSE\_TO\_ENDOPLASMIC\_RETICULUM\_STRESS(GO:1905897) |  | 78 | 0.61 | 1.03 | 0.141 | 0.357 | 1.000 | 18692 | tags=47%, list=32%, signal=69% |
| 5992 | POLYAMINE\_METABOLIC\_PROCESS(GO:0006595) |  | 17 | 0.60 | 1.03 | 0.141 | 0.357 | 1.000 | 13664 | tags=35%, list=23%, signal=46% |
| 5993 | SPINDLE\_ASSEMBLY(GO:0051225) |  | 96 | 0.61 | 1.03 | 0.141 | 0.357 | 1.000 | 19813 | tags=68%, list=34%, signal=102% |
| 5994 | CELLULAR\_RESPONSE\_TO\_LEPTIN\_STIMULUS(GO:0044320) |  | 17 | 0.67 | 1.03 | 0.141 | 0.357 | 1.000 | 13935 | tags=41%, list=24%, signal=54% |
| 5995 | HYDROGEN\_PEROXIDE\_MEDIATED\_PROGRAMMED\_CELL\_DEATH(GO:0010421) |  | 15 | 0.59 | 1.03 | 0.141 | 0.357 | 1.000 | 24141 | tags=80%, list=41%, signal=135% |
| 5996 | PROGRAMMED\_CELL\_DEATH\_IN\_RESPONSE\_TO\_REACTIVE\_OXYGEN\_SPECIES(GO:0097468) |  | 15 | 0.59 | 1.03 | 0.141 | 0.357 | 1.000 | 24141 | tags=80%, list=41%, signal=135% |
| 5997 | GOLGI\_ORGANIZATION(GO:0007030) |  | 109 | 0.57 | 1.03 | 0.141 | 0.357 | 1.000 | 21966 | tags=75%, list=37%, signal=119% |
| 5998 | EPITHELIAL\_CELL\_PROLIFERATION\_INVOLVED\_IN\_LIVER\_MORPHOGENESIS(GO:0072575) |  | 17 | 0.67 | 1.03 | 0.141 | 0.357 | 1.000 | 15437 | tags=41%, list=26%, signal=56% |
| 5999 | HEPATOCYTE\_PROLIFERATION(GO:0072574) |  | 17 | 0.67 | 1.03 | 0.141 | 0.357 | 1.000 | 15437 | tags=41%, list=26%, signal=56% |
| 6000 | THYROID\_HORMONE\_RECEPTOR\_BINDING(GO:0046966) |  | 28 | 0.60 | 1.03 | 0.141 | 0.357 | 1.000 | 20651 | tags=71%, list=35%, signal=110% |
| 6001 | POSITIVE\_REGULATION\_OF\_CELL\_CYCLE\_G2\_M\_PHASE\_TRANSITION(GO:1902751) |  | 26 | 0.60 | 1.03 | 0.141 | 0.357 | 1.000 | 21405 | tags=77%, list=36%, signal=121% |
| 6002 | CLEAVAGE\_INVOLVED\_IN\_RRNA\_PROCESSING(GO:0000469) |  | 24 | 0.59 | 1.03 | 0.141 | 0.357 | 1.000 | 23262 | tags=92%, list=39%, signal=151% |
| 6003 | PEPTIDYL\_LYSINE\_METHYLATION(GO:0018022) |  | 119 | 0.59 | 1.03 | 0.141 | 0.357 | 1.000 | 21531 | tags=71%, list=36%, signal=112% |
| 6004 | REGULATION\_OF\_CELLULAR\_RESPONSE\_TO\_INSULIN\_STIMULUS(GO:1900076) |  | 70 | 0.58 | 1.03 | 0.141 | 0.357 | 1.000 | 17846 | tags=41%, list=30%, signal=59% |
| 6005 | MITOCHONDRIAL\_ATP\_SYNTHESIS\_COUPLED\_PROTON\_TRANSPORT(GO:0042776) |  | 21 | 0.59 | 1.03 | 0.141 | 0.357 | 1.000 | 22093 | tags=86%, list=37%, signal=137% |
| 6006 | PROTEIN\_METHYLTRANSFERASE\_ACTIVITY(GO:0008276) |  | 89 | 0.58 | 1.03 | 0.141 | 0.357 | 1.000 | 23268 | tags=84%, list=39%, signal=139% |
| 6007 | KINASE\_ACTIVATOR\_ACTIVITY(GO:0019209) |  | 82 | 0.59 | 1.03 | 0.141 | 0.357 | 1.000 | 18981 | tags=60%, list=32%, signal=88% |
| 6008 | TRIF\_DEPENDENT\_TOLL\_LIKE\_RECEPTOR\_SIGNALING\_PATHWAY(GO:0035666) |  | 29 | 0.59 | 1.03 | 0.141 | 0.357 | 1.000 | 19834 | tags=55%, list=34%, signal=83% |
| 6009 | TRANSCRIPTION\_ELONGATION\_FROM\_RNA\_POLYMERASE\_II\_PROMOTER(GO:0006368) |  | 107 | 0.57 | 1.03 | 0.141 | 0.357 | 1.000 | 24644 | tags=90%, list=42%, signal=154% |
| 6010 | CEREBELLAR\_CORTEX\_FORMATION(GO:0021697) |  | 23 | 0.58 | 1.03 | 0.141 | 0.357 | 1.000 | 20668 | tags=57%, list=35%, signal=87% |
| 6011 | ENDODEOXYRIBONUCLEASE\_ACTIVITY\_\_PRODUCING\_3'\_PHOSPHOMONOESTERS(GO:0016889) |  | 22 | 0.58 | 1.03 | 0.141 | 0.357 | 1.000 | 21853 | tags=77%, list=37%, signal=123% |
| 6012 | REGULATION\_OF\_GLYCOPROTEIN\_METABOLIC\_PROCESS(GO:1903018) |  | 43 | 0.60 | 1.03 | 0.141 | 0.357 | 1.000 | 19690 | tags=51%, list=33%, signal=77% |
| 6013 | PROTEIN\_TARGETING\_TO\_LYSOSOME(GO:0006622) |  | 20 | 0.58 | 1.03 | 0.141 | 0.357 | 1.000 | 24672 | tags=95%, list=42%, signal=163% |
| 6014 | REGULATION\_OF\_NEURON\_PROJECTION\_REGENERATION(GO:0070570) |  | 29 | 0.61 | 1.03 | 0.141 | 0.357 | 1.000 | 21439 | tags=72%, list=36%, signal=114% |
| 6015 | CATION\_TRANSPORTING\_ATPASE\_ACTIVITY(GO:0019829) |  | 65 | 0.60 | 1.03 | 0.141 | 0.357 | 1.000 | 18171 | tags=54%, list=31%, signal=78% |
| 6016 | POLYSACCHARIDE\_CATABOLIC\_PROCESS(GO:0000272) |  | 28 | 0.58 | 1.03 | 0.141 | 0.357 | 1.000 | 22338 | tags=68%, list=38%, signal=109% |
| 6017 | MITOCHONDRIAL\_FISSION(GO:0000266) |  | 37 | 0.61 | 1.03 | 0.141 | 0.357 | 1.000 | 19159 | tags=68%, list=32%, signal=100% |
| 6018 | POSITIVE\_REGULATION\_OF\_ISOTYPE\_SWITCHING(GO:0045830) |  | 22 | 0.64 | 1.03 | 0.141 | 0.357 | 1.000 | 19540 | tags=68%, list=33%, signal=102% |
| 6019 | GPI\_ANCHOR\_BIOSYNTHETIC\_PROCESS(GO:0006506) |  | 34 | 0.57 | 1.03 | 0.141 | 0.357 | 1.000 | 23473 | tags=74%, list=40%, signal=122% |
| 6020 | EXODEOXYRIBONUCLEASE\_ACTIVITY(GO:0004529) |  | 19 | 0.61 | 1.03 | 0.141 | 0.357 | 1.000 | 21812 | tags=84%, list=37%, signal=133% |
| 6021 | EXODEOXYRIBONUCLEASE\_ACTIVITY\_\_PRODUCING\_5'\_PHOSPHOMONOESTERS(GO:0016895) |  | 19 | 0.61 | 1.03 | 0.141 | 0.357 | 1.000 | 21812 | tags=84%, list=37%, signal=133% |
| 6022 | MYD88\_INDEPENDENT\_TOLL\_LIKE\_RECEPTOR\_SIGNALING\_PATHWAY(GO:0002756) |  | 32 | 0.59 | 1.03 | 0.141 | 0.357 | 1.000 | 21697 | tags=66%, list=37%, signal=104% |
| 6023 | PHOTOTRANSDUCTION(GO:0007602) |  | 58 | 0.57 | 1.03 | 0.141 | 0.357 | 1.000 | 22324 | tags=57%, list=38%, signal=91% |
| 6024 | REGULATION\_OF\_ISOTYPE\_SWITCHING(GO:0045191) |  | 30 | 0.63 | 1.03 | 0.141 | 0.357 | 1.000 | 19540 | tags=63%, list=33%, signal=95% |
| 6025 | FATTY\_ACID\_DERIVATIVE\_BINDING(GO:1901567) |  | 23 | 0.62 | 1.03 | 0.141 | 0.357 | 1.000 | 11715 | tags=26%, list=20%, signal=33% |
| 6026 | U4\_U6\_X\_U5\_TRI\_SNRNP\_COMPLEX(GO:0046540) |  | 24 | 0.58 | 1.03 | 0.141 | 0.357 | 1.000 | 24161 | tags=79%, list=41%, signal=134% |
| 6027 | DOLICHOL\_LINKED\_OLIGOSACCHARIDE\_BIOSYNTHETIC\_PROCESS(GO:0006488) |  | 20 | 0.60 | 1.03 | 0.141 | 0.357 | 1.000 | 14759 | tags=45%, list=25%, signal=60% |
| 6028 | TRANSLATION\_INITIATION\_FACTOR\_ACTIVITY(GO:0003743) |  | 56 | 0.59 | 1.03 | 0.141 | 0.357 | 1.000 | 21651 | tags=84%, list=37%, signal=132% |
| 6029 | SIN3\_TYPE\_COMPLEX(GO:0070822) |  | 16 | 0.60 | 1.03 | 0.000 | 0.357 | 1.000 | 23743 | tags=100%, list=40%, signal=167% |
| 6030 | PRECATALYTIC\_SPLICEOSOME(GO:0071011) |  | 28 | 0.59 | 1.03 | 0.141 | 0.357 | 1.000 | 23780 | tags=89%, list=40%, signal=149% |
| 6031 | PROTEIN\_DEACYLATION(GO:0035601) |  | 108 | 0.58 | 1.03 | 0.141 | 0.358 | 1.000 | 22117 | tags=73%, list=37%, signal=117% |
| 6032 | POSITIVE\_REGULATION\_OF\_CELLULAR\_RESPONSE\_TO\_INSULIN\_STIMULUS(GO:1900078) |  | 29 | 0.58 | 1.03 | 0.141 | 0.358 | 1.000 | 18963 | tags=48%, list=32%, signal=71% |
| 6033 | POSITIVE\_REGULATION\_OF\_MYOBLAST\_DIFFERENTIATION(GO:0045663) |  | 20 | 0.66 | 1.03 | 0.141 | 0.358 | 1.000 | 9268 | tags=35%, list=16%, signal=41% |
| 6034 | S\_ADENOSYLMETHIONINE\_DEPENDENT\_METHYLTRANSFERASE\_ACTIVITY(GO:0008757) |  | 155 | 0.59 | 1.03 | 0.141 | 0.358 | 1.000 | 21803 | tags=77%, list=37%, signal=122% |
| 6035 | CEREBELLAR\_PURKINJE\_CELL\_LAYER\_DEVELOPMENT(GO:0021680) |  | 27 | 0.58 | 1.03 | 0.141 | 0.358 | 1.000 | 18601 | tags=56%, list=31%, signal=81% |
| 6036 | AMINO\_ACID\_BETAINE\_METABOLIC\_PROCESS(GO:0006577) |  | 17 | 0.61 | 1.03 | 0.310 | 0.358 | 1.000 | 21038 | tags=71%, list=36%, signal=110% |
| 6037 | EMBRYONIC\_HEMOPOIESIS(GO:0035162) |  | 21 | 0.61 | 1.03 | 0.141 | 0.358 | 1.000 | 19756 | tags=62%, list=33%, signal=93% |
| 6038 | HYDRO\_LYASE\_ACTIVITY(GO:0016836) |  | 53 | 0.58 | 1.03 | 0.141 | 0.358 | 1.000 | 20756 | tags=58%, list=35%, signal=90% |
| 6039 | SNRNA\_METABOLIC\_PROCESS(GO:0016073) |  | 87 | 0.57 | 1.03 | 0.141 | 0.358 | 1.000 | 24644 | tags=93%, list=42%, signal=159% |
| 6040 | POSITIVE\_REGULATION\_OF\_KERATINOCYTE\_DIFFERENTIATION(GO:0045618) |  | 15 | 0.65 | 1.03 | 0.141 | 0.358 | 1.000 | 10768 | tags=33%, list=18%, signal=41% |
| 6041 | REGULATION\_OF\_ESTABLISHMENT\_OF\_CELL\_POLARITY(GO:2000114) |  | 21 | 0.61 | 1.03 | 0.141 | 0.358 | 1.000 | 20968 | tags=71%, list=35%, signal=111% |
| 6042 | ANDROGEN\_RECEPTOR\_SIGNALING\_PATHWAY(GO:0030521) |  | 68 | 0.62 | 1.03 | 0.141 | 0.358 | 1.000 | 18416 | tags=59%, list=31%, signal=85% |
| 6043 | PHOTOTRANSDUCTION\_\_VISIBLE\_LIGHT(GO:0007603) |  | 36 | 0.58 | 1.03 | 0.141 | 0.358 | 1.000 | 23921 | tags=75%, list=40%, signal=126% |
| 6044 | GLYCEROL\_METABOLIC\_PROCESS(GO:0006071) |  | 18 | 0.59 | 1.03 | 0.141 | 0.358 | 1.000 | 24352 | tags=83%, list=41%, signal=142% |
| 6045 | POSITIVE\_REGULATION\_OF\_G2\_M\_TRANSITION\_OF\_MITOTIC\_CELL\_CYCLE(GO:0010971) |  | 22 | 0.60 | 1.03 | 0.141 | 0.358 | 1.000 | 21405 | tags=77%, list=36%, signal=121% |
| 6046 | POSITIVE\_REGULATION\_OF\_PROTEIN\_LOCALIZATION\_TO\_PLASMA\_MEMBRANE(GO:1903078) |  | 54 | 0.60 | 1.03 | 0.141 | 0.358 | 1.000 | 19932 | tags=48%, list=34%, signal=73% |
| 6047 | POSITIVE\_REGULATION\_OF\_INTERFERON\_BETA\_PRODUCTION(GO:0032728) |  | 31 | 0.62 | 1.03 | 0.141 | 0.358 | 1.000 | 15297 | tags=39%, list=26%, signal=52% |
| 6048 | FATTY\_ACID\_BETA\_OXIDATION\_USING\_ACYL\_COA\_DEHYDROGENASE(GO:0033539) |  | 16 | 0.62 | 1.03 | 0.141 | 0.358 | 1.000 | 12487 | tags=31%, list=21%, signal=40% |
| 6049 | CENTROSOME\_LOCALIZATION(GO:0051642) |  | 26 | 0.59 | 1.03 | 0.141 | 0.358 | 1.000 | 23520 | tags=96%, list=40%, signal=160% |
| 6050 | INO80\_TYPE\_COMPLEX(GO:0097346) |  | 24 | 0.59 | 1.03 | 0.141 | 0.358 | 1.000 | 23132 | tags=96%, list=39%, signal=157% |
| 6051 | DNA\_LIGATION(GO:0006266) |  | 24 | 0.63 | 1.03 | 0.141 | 0.358 | 1.000 | 20007 | tags=79%, list=34%, signal=120% |
| 6052 | NATURAL\_KILLER\_CELL\_MEDIATED\_IMMUNITY(GO:0002228) |  | 59 | 0.59 | 1.03 | 0.141 | 0.358 | 1.000 | 15869 | tags=41%, list=27%, signal=56% |
| 6053 | REGULATION\_OF\_TRANSCRIPTION\_BY\_RNA\_POLYMERASE\_I(GO:0006356) |  | 33 | 0.57 | 1.03 | 0.141 | 0.358 | 1.000 | 25025 | tags=94%, list=42%, signal=163% |
| 6054 | REGULATION\_OF\_INSULIN\_RECEPTOR\_SIGNALING\_PATHWAY(GO:0046626) |  | 50 | 0.58 | 1.03 | 0.141 | 0.358 | 1.000 | 17387 | tags=40%, list=29%, signal=57% |
| 6055 | VIRAL\_PROCESS(GO:0016032) |  | 778 | 0.61 | 1.03 | 0.141 | 0.358 | 1.000 | 18259 | tags=44%, list=31%, signal=63% |
| 6056 | SEQUESTERING\_OF\_TRIGLYCERIDE(GO:0030730) |  | 15 | 0.60 | 1.03 | 0.141 | 0.358 | 1.000 | 22269 | tags=53%, list=38%, signal=86% |
| 6057 | RHO\_GUANYL\_NUCLEOTIDE\_EXCHANGE\_FACTOR\_ACTIVITY(GO:0005089) |  | 86 | 0.61 | 1.03 | 0.141 | 0.359 | 1.000 | 17265 | tags=45%, list=29%, signal=64% |
| 6058 | REGULATION\_OF\_LYMPHOCYTE\_MEDIATED\_IMMUNITY(GO:0002706) |  | 119 | 0.59 | 1.03 | 0.141 | 0.359 | 1.000 | 16856 | tags=39%, list=29%, signal=55% |
| 6059 | CYTOPLASMIC\_TRANSLATION(GO:0002181) |  | 88 | 0.58 | 1.03 | 0.141 | 0.359 | 1.000 | 22494 | tags=83%, list=38%, signal=134% |
| 6060 | POSITIVE\_REGULATION\_OF\_TELOMERASE\_ACTIVITY(GO:0051973) |  | 36 | 0.64 | 1.03 | 0.141 | 0.359 | 1.000 | 16850 | tags=69%, list=28%, signal=97% |
| 6061 | REGULATION\_OF\_VESICLE\_FUSION(GO:0031338) |  | 70 | 0.59 | 1.03 | 0.307 | 0.359 | 1.000 | 23062 | tags=67%, list=39%, signal=110% |
| 6062 | HISTONE\_METHYLTRANSFERASE\_ACTIVITY(GO:0042054) |  | 59 | 0.59 | 1.03 | 0.141 | 0.359 | 1.000 | 23844 | tags=88%, list=40%, signal=148% |
| 6063 | TERTIARY\_GRANULE\_LUMEN(GO:1904724) |  | 55 | 0.61 | 1.03 | 0.141 | 0.359 | 1.000 | 13076 | tags=20%, list=22%, signal=26% |
| 6064 | RAB\_GTPASE\_BINDING(GO:0017137) |  | 177 | 0.61 | 1.03 | 0.141 | 0.359 | 1.000 | 11933 | tags=22%, list=20%, signal=28% |
| 6065 | ANAPHASE\_PROMOTING\_COMPLEX(GO:0005680) |  | 22 | 0.58 | 1.03 | 0.141 | 0.359 | 1.000 | 24082 | tags=95%, list=41%, signal=161% |
| 6066 | TRANSFORMING\_GROWTH\_FACTOR\_BETA\_BINDING(GO:0050431) |  | 19 | 0.63 | 1.03 | 0.141 | 0.359 | 1.000 | 8744 | tags=16%, list=15%, signal=19% |
| 6067 | ZINC\_ION\_HOMEOSTASIS(GO:0055069) |  | 28 | 0.61 | 1.03 | 0.141 | 0.359 | 1.000 | 17023 | tags=54%, list=29%, signal=75% |
| 6068 | TOXIN\_TRANSPORT(GO:1901998) |  | 38 | 0.59 | 1.03 | 0.141 | 0.359 | 1.000 | 21868 | tags=71%, list=37%, signal=113% |
| 6069 | N\_TERMINAL\_PROTEIN\_AMINO\_ACID\_MODIFICATION(GO:0031365) |  | 28 | 0.58 | 1.03 | 0.141 | 0.359 | 1.000 | 23797 | tags=93%, list=40%, signal=155% |
| 6070 | POLYOL\_CATABOLIC\_PROCESS(GO:0046174) |  | 19 | 0.59 | 1.03 | 0.141 | 0.359 | 1.000 | 19903 | tags=68%, list=34%, signal=103% |
| 6071 | VESICLE\_TRANSPORT\_ALONG\_MICROTUBULE(GO:0047496) |  | 37 | 0.59 | 1.03 | 0.141 | 0.359 | 1.000 | 21510 | tags=81%, list=36%, signal=127% |
| 6072 | MICROTUBULE\_ORGANIZING\_CENTER\_LOCALIZATION(GO:0061842) |  | 27 | 0.59 | 1.03 | 0.141 | 0.359 | 1.000 | 23520 | tags=96%, list=40%, signal=160% |
| 6073 | CELLULAR\_RESPONSE\_TO\_ARSENIC\_CONTAINING\_SUBSTANCE(GO:0071243) |  | 17 | 0.60 | 1.03 | 0.141 | 0.359 | 1.000 | 20696 | tags=76%, list=35%, signal=118% |
| 6074 | REPRODUCTIVE\_BEHAVIOR(GO:0019098) |  | 30 | 0.62 | 1.03 | 0.141 | 0.359 | 1.000 | 10415 | tags=27%, list=18%, signal=32% |
| 6075 | ECTODERMAL\_PLACODE\_DEVELOPMENT(GO:0071696) |  | 15 | 0.62 | 1.03 | 0.141 | 0.359 | 1.000 | 11411 | tags=40%, list=19%, signal=50% |
| 6076 | REGULATION\_OF\_AMYLOID\_PRECURSOR\_PROTEIN\_CATABOLIC\_PROCESS(GO:1902991) |  | 23 | 0.60 | 1.03 | 0.141 | 0.359 | 1.000 | 21613 | tags=74%, list=37%, signal=116% |
| 6077 | PHOTORECEPTOR\_DISC\_MEMBRANE(GO:0097381) |  | 22 | 0.61 | 1.03 | 0.310 | 0.359 | 1.000 | 17145 | tags=45%, list=29%, signal=64% |
| 6078 | PEPTIDYL\_LYSINE\_TRIMETHYLATION(GO:0018023) |  | 40 | 0.58 | 1.03 | 0.141 | 0.359 | 1.000 | 22908 | tags=80%, list=39%, signal=130% |
| 6079 | PEROXISOME\_ORGANIZATION(GO:0007031) |  | 31 | 0.57 | 1.03 | 0.307 | 0.359 | 1.000 | 17954 | tags=52%, list=30%, signal=74% |
| 6080 | OLIGOSACCHARIDE\_LIPID\_INTERMEDIATE\_BIOSYNTHETIC\_PROCESS(GO:0006490) |  | 21 | 0.59 | 1.03 | 0.141 | 0.359 | 1.000 | 15355 | tags=48%, list=26%, signal=64% |
| 6081 | PROTEIN\_DEACETYLATION(GO:0006476) |  | 98 | 0.58 | 1.03 | 0.141 | 0.359 | 1.000 | 22117 | tags=74%, list=37%, signal=119% |
| 6082 | UBIQUINONE\_METABOLIC\_PROCESS(GO:0006743) |  | 17 | 0.60 | 1.03 | 0.141 | 0.359 | 1.000 | 23667 | tags=100%, list=40%, signal=167% |
| 6083 | CELLULAR\_RESPONSE\_TO\_MISFOLDED\_PROTEIN(GO:0071218) |  | 22 | 0.58 | 1.03 | 0.141 | 0.359 | 1.000 | 25069 | tags=100%, list=42%, signal=174% |
| 6084 | CLATHRIN\_COATED\_VESICLE(GO:0030136) |  | 173 | 0.60 | 1.03 | 0.141 | 0.359 | 1.000 | 18127 | tags=40%, list=31%, signal=57% |
| 6085 | SIGNAL\_PEPTIDE\_PROCESSING(GO:0006465) |  | 19 | 0.63 | 1.03 | 0.141 | 0.359 | 1.000 | 16827 | tags=63%, list=28%, signal=88% |
| 6086 | NLS\_BEARING\_PROTEIN\_IMPORT\_INTO\_NUCLEUS(GO:0006607) |  | 22 | 0.61 | 1.03 | 0.340 | 0.359 | 1.000 | 18005 | tags=64%, list=30%, signal=91% |
| 6087 | RIBONUCLEOPROTEIN\_COMPLEX\_SUBUNIT\_ORGANIZATION(GO:0071826) |  | 236 | 0.59 | 1.03 | 0.141 | 0.359 | 1.000 | 22480 | tags=86%, list=38%, signal=138% |
| 6088 | ACROSOME\_REACTION(GO:0007340) |  | 37 | 0.63 | 1.03 | 0.141 | 0.359 | 1.000 | 19043 | tags=57%, list=32%, signal=84% |
| 6089 | REGULATION\_OF\_CARDIAC\_MUSCLE\_HYPERTROPHY(GO:0010611) |  | 65 | 0.59 | 1.03 | 0.141 | 0.359 | 1.000 | 18212 | tags=38%, list=31%, signal=56% |
| 6090 | I\_KAPPAB\_PHOSPHORYLATION(GO:0007252) |  | 15 | 0.61 | 1.03 | 0.141 | 0.359 | 1.000 | 16329 | tags=47%, list=28%, signal=64% |
| 6091 | PROTEIN\_KINASE\_INHIBITOR\_ACTIVITY(GO:0004860) |  | 95 | 0.60 | 1.03 | 0.141 | 0.359 | 1.000 | 10693 | tags=22%, list=18%, signal=27% |
| 6092 | ANTIGEN\_PROCESSING\_AND\_PRESENTATION\_OF\_PEPTIDE\_OR\_POLYSACCHARIDE\_ANTIGEN\_VIA\_MHC\_CLASS\_II(GO:0002504) |  | 101 | 0.61 | 1.03 | 0.141 | 0.359 | 1.000 | 19775 | tags=56%, list=33%, signal=85% |
| 6093 | ESTABLISHMENT\_OF\_PROTEIN\_LOCALIZATION\_TO\_VACUOLE(GO:0072666) |  | 35 | 0.57 | 1.03 | 0.141 | 0.359 | 1.000 | 24672 | tags=89%, list=42%, signal=152% |
| 6094 | G\_PROTEIN\_COUPLED\_CHEMOATTRACTANT\_RECEPTOR\_ACTIVITY(GO:0001637) |  | 26 | 0.64 | 1.03 | 0.141 | 0.359 | 1.000 | 16900 | tags=38%, list=29%, signal=54% |
| 6095 | CHEMOKINE\_RECEPTOR\_ACTIVITY(GO:0004950) |  | 26 | 0.64 | 1.03 | 0.141 | 0.359 | 1.000 | 16900 | tags=38%, list=29%, signal=54% |
| 6096 | PHAGOCYTIC\_CUP(GO:0001891) |  | 22 | 0.59 | 1.03 | 0.310 | 0.359 | 1.000 | 13150 | tags=32%, list=22%, signal=41% |
| 6097 | RESPONSE\_TO\_ISOQUINOLINE\_ALKALOID(GO:0014072) |  | 33 | 0.62 | 1.03 | 0.141 | 0.359 | 1.000 | 6471 | tags=15%, list=11%, signal=17% |
| 6098 | RESPONSE\_TO\_MORPHINE(GO:0043278) |  | 33 | 0.62 | 1.03 | 0.141 | 0.359 | 1.000 | 6471 | tags=15%, list=11%, signal=17% |
| 6099 | NEGATIVE\_REGULATION\_OF\_PLASMA\_MEMBRANE\_BOUNDED\_CELL\_PROJECTION\_ASSEMBLY(GO:0120033) |  | 27 | 0.63 | 1.03 | 0.141 | 0.359 | 1.000 | 16486 | tags=44%, list=28%, signal=62% |
| 6100 | DNA\_HELICASE\_COMPLEX(GO:0033202) |  | 16 | 0.59 | 1.03 | 0.141 | 0.359 | 1.000 | 24171 | tags=100%, list=41%, signal=169% |
| 6101 | INO80\_COMPLEX(GO:0031011) |  | 16 | 0.59 | 1.03 | 0.141 | 0.359 | 1.000 | 24171 | tags=100%, list=41%, signal=169% |
| 6102 | MODULATION\_BY\_VIRUS\_OF\_HOST\_MORPHOLOGY\_OR\_PHYSIOLOGY(GO:0019048) |  | 39 | 0.57 | 1.03 | 0.141 | 0.359 | 1.000 | 21805 | tags=64%, list=37%, signal=101% |
| 6103 | MOLECULAR\_CARRIER\_ACTIVITY(GO:0140104) |  | 42 | 0.62 | 1.03 | 0.141 | 0.359 | 1.000 | 18009 | tags=57%, list=30%, signal=82% |
| 6104 | LYSOSOMAL\_TRANSPORT(GO:0007041) |  | 92 | 0.57 | 1.03 | 0.141 | 0.359 | 1.000 | 24906 | tags=85%, list=42%, signal=146% |
| 6105 | NEGATIVE\_REGULATION\_OF\_PHAGOCYTOSIS(GO:0050765) |  | 16 | 0.59 | 1.03 | 0.309 | 0.360 | 1.000 | 23446 | tags=56%, list=40%, signal=93% |
| 6106 | ENDOPLASMIC\_RETICULUM\_QUALITY\_CONTROL\_COMPARTMENT(GO:0044322) |  | 28 | 0.57 | 1.03 | 0.141 | 0.360 | 1.000 | 20675 | tags=75%, list=35%, signal=115% |
| 6107 | KINASE\_INHIBITOR\_ACTIVITY(GO:0019210) |  | 99 | 0.60 | 1.03 | 0.141 | 0.360 | 1.000 | 19110 | tags=46%, list=32%, signal=69% |
| 6108 | MODIFICATION\_DEPENDENT\_PROTEIN\_BINDING(GO:0140030) |  | 131 | 0.58 | 1.03 | 0.141 | 0.360 | 1.000 | 22759 | tags=76%, list=38%, signal=124% |
| 6109 | CYTOSOLIC\_SMALL\_RIBOSOMAL\_SUBUNIT(GO:0022627) |  | 46 | 0.57 | 1.03 | 0.141 | 0.360 | 1.000 | 24601 | tags=87%, list=42%, signal=149% |
| 6110 | HISTONE\_H2A\_MONOUBIQUITINATION(GO:0035518) |  | 17 | 0.59 | 1.03 | 0.141 | 0.360 | 1.000 | 22028 | tags=76%, list=37%, signal=122% |
| 6111 | POSITIVE\_REGULATION\_OF\_LAMELLIPODIUM\_ORGANIZATION(GO:1902745) |  | 26 | 0.59 | 1.03 | 0.307 | 0.360 | 1.000 | 14664 | tags=31%, list=25%, signal=41% |
| 6112 | REGULATION\_OF\_ANOIKIS(GO:2000209) |  | 23 | 0.60 | 1.03 | 0.141 | 0.360 | 1.000 | 18981 | tags=48%, list=32%, signal=70% |
| 6113 | NEGATIVE\_REGULATION\_OF\_INTERLEUKIN\_1\_SECRETION(GO:0050711) |  | 15 | 0.60 | 1.03 | 0.310 | 0.360 | 1.000 | 20085 | tags=33%, list=34%, signal=50% |
| 6114 | MITOTIC\_RECOMBINATION(GO:0006312) |  | 25 | 0.64 | 1.03 | 0.141 | 0.360 | 1.000 | 19910 | tags=76%, list=34%, signal=115% |
| 6115 | ENDOPLASMIC\_RETICULUM\_TUBULAR\_NETWORK\_ORGANIZATION(GO:0071786) |  | 16 | 0.62 | 1.03 | 0.340 | 0.360 | 1.000 | 21966 | tags=94%, list=37%, signal=149% |
| 6116 | CHAPERONE\_COFACTOR\_DEPENDENT\_PROTEIN\_REFOLDING(GO:0051085) |  | 16 | 0.58 | 1.03 | 0.141 | 0.360 | 1.000 | 22131 | tags=81%, list=37%, signal=130% |
| 6117 | NEGATIVE\_REGULATION\_OF\_MACROAUTOPHAGY(GO:0016242) |  | 32 | 0.59 | 1.03 | 0.141 | 0.360 | 1.000 | 22939 | tags=84%, list=39%, signal=138% |
| 6118 | POSITIVE\_REGULATION\_OF\_LYMPHOCYTE\_MEDIATED\_IMMUNITY(GO:0002708) |  | 76 | 0.60 | 1.03 | 0.141 | 0.360 | 1.000 | 19618 | tags=51%, list=33%, signal=77% |
| 6119 | CIS\_GOLGI\_NETWORK(GO:0005801) |  | 43 | 0.58 | 1.02 | 0.141 | 0.360 | 1.000 | 22051 | tags=74%, list=37%, signal=119% |
| 6120 | RAB\_GUANYL\_NUCLEOTIDE\_EXCHANGE\_FACTOR\_ACTIVITY(GO:0017112) |  | 50 | 0.60 | 1.02 | 0.141 | 0.360 | 1.000 | 18721 | tags=54%, list=32%, signal=79% |
| 6121 | FLAVIN\_ADENINE\_DINUCLEOTIDE\_BINDING(GO:0050660) |  | 75 | 0.59 | 1.02 | 0.141 | 0.360 | 1.000 | 21845 | tags=64%, list=37%, signal=101% |
| 6122 | CELLULAR\_PROTEIN\_CONTAINING\_COMPLEX\_LOCALIZATION(GO:0034629) |  | 25 | 0.59 | 1.02 | 0.141 | 0.360 | 1.000 | 17888 | tags=52%, list=30%, signal=75% |
| 6123 | VESICLE\_DOCKING(GO:0048278) |  | 61 | 0.58 | 1.02 | 0.141 | 0.360 | 1.000 | 21162 | tags=62%, list=36%, signal=97% |
| 6124 | MITOCHONDRION\_DISASSEMBLY(GO:0061726) |  | 83 | 0.58 | 1.02 | 0.141 | 0.360 | 1.000 | 23032 | tags=80%, list=39%, signal=130% |
| 6125 | AUTOPHAGY\_OF\_MITOCHONDRION(GO:0000422) |  | 83 | 0.58 | 1.02 | 0.141 | 0.360 | 1.000 | 23032 | tags=80%, list=39%, signal=130% |
| 6126 | REGULATION\_OF\_AMYLOID\_BETA\_FORMATION(GO:1902003) |  | 18 | 0.60 | 1.02 | 0.141 | 0.360 | 1.000 | 21613 | tags=72%, list=37%, signal=114% |
| 6127 | STRUCTURAL\_CONSTITUENT\_OF\_RIBOSOME(GO:0003735) |  | 176 | 0.57 | 1.02 | 0.141 | 0.360 | 1.000 | 24309 | tags=86%, list=41%, signal=146% |
| 6128 | ENDOPLASMIC\_RETICULUM\_TUBULAR\_NETWORK(GO:0071782) |  | 20 | 0.60 | 1.02 | 0.141 | 0.360 | 1.000 | 21966 | tags=85%, list=37%, signal=135% |
| 6129 | REGULATION\_OF\_INTRINSIC\_APOPTOTIC\_SIGNALING\_PATHWAY\_BY\_P53\_CLASS\_MEDIATOR(GO:1902253) |  | 26 | 0.61 | 1.02 | 0.141 | 0.360 | 1.000 | 17869 | tags=46%, list=30%, signal=66% |
| 6130 | SYNAPTIC\_VESICLE\_CYTOSKELETAL\_TRANSPORT(GO:0099514) |  | 17 | 0.60 | 1.02 | 0.141 | 0.360 | 1.000 | 22122 | tags=88%, list=37%, signal=141% |
| 6131 | SYNAPTIC\_VESICLE\_TRANSPORT\_ALONG\_MICROTUBULE(GO:0099517) |  | 17 | 0.60 | 1.02 | 0.141 | 0.360 | 1.000 | 22122 | tags=88%, list=37%, signal=141% |
| 6132 | ANTEROGRADE\_SYNAPTIC\_VESICLE\_TRANSPORT(GO:0048490) |  | 17 | 0.60 | 1.02 | 0.141 | 0.360 | 1.000 | 22122 | tags=88%, list=37%, signal=141% |
| 6133 | POSITIVE\_REGULATION\_OF\_TRANSCRIPTION\_BY\_RNA\_POLYMERASE\_I(GO:0045943) |  | 22 | 0.59 | 1.02 | 0.141 | 0.360 | 1.000 | 18340 | tags=68%, list=31%, signal=99% |
| 6134 | CLATHRIN\_DEPENDENT\_ENDOCYTOSIS(GO:0072583) |  | 37 | 0.57 | 1.02 | 0.141 | 0.360 | 1.000 | 21521 | tags=68%, list=36%, signal=106% |
| 6135 | VENTRICULAR\_TRABECULA\_MYOCARDIUM\_MORPHOGENESIS(GO:0003222) |  | 16 | 0.59 | 1.02 | 0.141 | 0.361 | 1.000 | 18134 | tags=56%, list=31%, signal=81% |
| 6136 | AMYLOID\_BETA\_FORMATION(GO:0034205) |  | 23 | 0.59 | 1.02 | 0.141 | 0.361 | 1.000 | 21613 | tags=74%, list=37%, signal=116% |
| 6137 | POSITIVE\_REGULATION\_OF\_CALCIUM\_ION\_DEPENDENT\_EXOCYTOSIS(GO:0045956) |  | 21 | 0.60 | 1.02 | 0.307 | 0.361 | 1.000 | 22164 | tags=67%, list=37%, signal=107% |
| 6138 | ATP\_DEPENDENT\_DNA\_HELICASE\_ACTIVITY(GO:0004003) |  | 34 | 0.59 | 1.02 | 0.141 | 0.361 | 1.000 | 24099 | tags=94%, list=41%, signal=159% |
| 6139 | MRNA\_CLEAVAGE\_AND\_POLYADENYLATION\_SPECIFICITY\_FACTOR\_COMPLEX(GO:0005847) |  | 17 | 0.60 | 1.02 | 0.307 | 0.361 | 1.000 | 22758 | tags=88%, list=38%, signal=143% |
| 6140 | NUCLEAR\_UBIQUITIN\_LIGASE\_COMPLEX(GO:0000152) |  | 40 | 0.57 | 1.02 | 0.310 | 0.361 | 1.000 | 23518 | tags=85%, list=40%, signal=141% |
| 6141 | POLYSOME(GO:0005844) |  | 73 | 0.58 | 1.02 | 0.141 | 0.361 | 1.000 | 22494 | tags=79%, list=38%, signal=128% |
| 6142 | PHOSPHATIDYLINOSITOL\_BIOSYNTHETIC\_PROCESS(GO:0006661) |  | 146 | 0.57 | 1.02 | 0.141 | 0.361 | 1.000 | 23473 | tags=71%, list=40%, signal=117% |
| 6143 | HISTONE\_LYSINE\_METHYLATION(GO:0034968) |  | 106 | 0.58 | 1.02 | 0.141 | 0.361 | 1.000 | 23268 | tags=79%, list=39%, signal=130% |
| 6144 | PYRIMIDINE\_NUCLEOSIDE\_BIOSYNTHETIC\_PROCESS(GO:0046134) |  | 29 | 0.57 | 1.02 | 0.310 | 0.361 | 1.000 | 23753 | tags=83%, list=40%, signal=138% |
| 6145 | EMBRYONIC\_PLACENTA\_MORPHOGENESIS(GO:0060669) |  | 24 | 0.65 | 1.02 | 0.307 | 0.361 | 1.000 | 17448 | tags=42%, list=30%, signal=59% |
| 6146 | POSITIVE\_REGULATION\_OF\_IMMUNE\_RESPONSE(GO:0050778) |  | 800 | 0.59 | 1.02 | 0.141 | 0.361 | 1.000 | 18785 | tags=36%, list=32%, signal=52% |
| 6147 | REGULATION\_OF\_TRANSLATION\_IN\_RESPONSE\_TO\_STRESS(GO:0043555) |  | 19 | 0.59 | 1.02 | 0.141 | 0.361 | 1.000 | 22367 | tags=79%, list=38%, signal=127% |
| 6148 | TRANSCRIPTION\_ELONGATION\_FACTOR\_COMPLEX(GO:0008023) |  | 55 | 0.58 | 1.02 | 0.141 | 0.361 | 1.000 | 23679 | tags=85%, list=40%, signal=142% |
| 6149 | HISTONE\_DEACETYLATION(GO:0016575) |  | 85 | 0.58 | 1.02 | 0.141 | 0.361 | 1.000 | 22117 | tags=74%, list=37%, signal=118% |
| 6150 | ANTIGEN\_PROCESSING\_AND\_PRESENTATION\_OF\_EXOGENOUS\_ANTIGEN(GO:0019884) |  | 177 | 0.59 | 1.02 | 0.141 | 0.361 | 1.000 | 19867 | tags=53%, list=34%, signal=79% |
| 6151 | REGULATION\_OF\_CLATHRIN\_DEPENDENT\_ENDOCYTOSIS(GO:2000369) |  | 15 | 0.59 | 1.02 | 0.141 | 0.361 | 1.000 | 17832 | tags=53%, list=30%, signal=76% |
| 6152 | PHOSPHATIDYLINOSITOL\_3\_PHOSPHATASE\_ACTIVITY(GO:0004438) |  | 15 | 0.60 | 1.02 | 0.481 | 0.361 | 1.000 | 23205 | tags=93%, list=39%, signal=154% |
| 6153 | PHOSPHATIDYLINOSITOL\_MONOPHOSPHATE\_PHOSPHATASE\_ACTIVITY(GO:0052744) |  | 15 | 0.60 | 1.02 | 0.481 | 0.361 | 1.000 | 23205 | tags=93%, list=39%, signal=154% |
| 6154 | POSITIVE\_REGULATION\_OF\_AUTOPHAGY\_OF\_MITOCHONDRION(GO:1903599) |  | 19 | 0.60 | 1.02 | 0.307 | 0.361 | 1.000 | 22360 | tags=89%, list=38%, signal=144% |
| 6155 | PHOSPHATIDYLINOSITOL\_DEPHOSPHORYLATION(GO:0046856) |  | 25 | 0.57 | 1.02 | 0.141 | 0.361 | 1.000 | 25291 | tags=96%, list=43%, signal=168% |
| 6156 | PROTEIN\_SERINE\_THREONINE\_KINASE\_ACTIVATOR\_ACTIVITY(GO:0043539) |  | 31 | 0.60 | 1.02 | 0.481 | 0.361 | 1.000 | 23921 | tags=94%, list=40%, signal=157% |
| 6157 | PEPTIDYL\_GLUTAMIC\_ACID\_MODIFICATION(GO:0018200) |  | 34 | 0.62 | 1.02 | 0.141 | 0.361 | 1.000 | 21071 | tags=76%, list=36%, signal=119% |
| 6158 | POSITIVE\_REGULATION\_OF\_NITRIC\_OXIDE\_SYNTHASE\_BIOSYNTHETIC\_PROCESS(GO:0051770) |  | 15 | 0.64 | 1.02 | 0.141 | 0.361 | 1.000 | 13320 | tags=27%, list=23%, signal=34% |
| 6159 | REGULATION\_OF\_CELL\_GROWTH\_INVOLVED\_IN\_CARDIAC\_MUSCLE\_CELL\_DEVELOPMENT(GO:0061050) |  | 18 | 0.63 | 1.02 | 0.141 | 0.361 | 1.000 | 21613 | tags=56%, list=37%, signal=88% |
| 6160 | DIVALENT\_INORGANIC\_CATION\_TRANSMEMBRANE\_TRANSPORTER\_ACTIVITY(GO:0072509) |  | 28 | 0.58 | 1.02 | 0.141 | 0.361 | 1.000 | 18903 | tags=68%, list=32%, signal=100% |
| 6161 | ANTIGEN\_BINDING(GO:0003823) |  | 191 | 0.56 | 1.02 | 0.307 | 0.361 | 1.000 | 21805 | tags=13%, list=37%, signal=21% |
| 6162 | PROTEIN\_LOCALIZATION\_TO\_VACUOLE(GO:0072665) |  | 45 | 0.57 | 1.02 | 0.141 | 0.361 | 1.000 | 24672 | tags=87%, list=42%, signal=149% |
| 6163 | CYTOPLASMIC\_SEQUESTERING\_OF\_TRANSCRIPTION\_FACTOR(GO:0042994) |  | 18 | 0.60 | 1.02 | 0.141 | 0.362 | 1.000 | 17014 | tags=56%, list=29%, signal=78% |
| 6164 | INNATE\_IMMUNE\_RESPONSE\_ACTIVATING\_SIGNAL\_TRANSDUCTION(GO:0002758) |  | 281 | 0.59 | 1.02 | 0.141 | 0.362 | 1.000 | 17194 | tags=36%, list=29%, signal=50% |
| 6165 | SPERM\_MIDPIECE(GO:0097225) |  | 20 | 0.61 | 1.02 | 0.307 | 0.362 | 1.000 | 15522 | tags=45%, list=26%, signal=61% |
| 6166 | 4\_IRON\_\_4\_SULFUR\_CLUSTER\_BINDING(GO:0051539) |  | 42 | 0.57 | 1.02 | 0.141 | 0.362 | 1.000 | 20703 | tags=74%, list=35%, signal=113% |
| 6167 | THIOLESTER\_HYDROLASE\_ACTIVITY(GO:0016790) |  | 37 | 0.59 | 1.02 | 0.141 | 0.362 | 1.000 | 18504 | tags=49%, list=31%, signal=71% |
| 6168 | MRNA\_CLEAVAGE\_FACTOR\_COMPLEX(GO:0005849) |  | 19 | 0.60 | 1.02 | 0.141 | 0.362 | 1.000 | 22758 | tags=89%, list=38%, signal=145% |
| 6169 | OXIDOREDUCTASE\_ACTIVITY\_\_ACTING\_ON\_THE\_CH\_CH\_GROUP\_OF\_DONORS(GO:0016627) |  | 60 | 0.59 | 1.02 | 0.141 | 0.362 | 1.000 | 20971 | tags=57%, list=35%, signal=88% |
| 6170 | REGULATION\_OF\_MUSCLE\_HYPERTROPHY(GO:0014743) |  | 67 | 0.59 | 1.02 | 0.141 | 0.362 | 1.000 | 18212 | tags=39%, list=31%, signal=56% |
| 6171 | HISTONE\_H3\_K9\_METHYLATION(GO:0051567) |  | 30 | 0.58 | 1.02 | 0.141 | 0.362 | 1.000 | 23268 | tags=77%, list=39%, signal=126% |
| 6172 | CORTICOSTEROID\_RECEPTOR\_SIGNALING\_PATHWAY(GO:0031958) |  | 18 | 0.58 | 1.02 | 0.141 | 0.362 | 1.000 | 23558 | tags=89%, list=40%, signal=148% |
| 6173 | UBIQUINONE\_BIOSYNTHETIC\_PROCESS(GO:0006744) |  | 16 | 0.60 | 1.02 | 0.307 | 0.362 | 1.000 | 23667 | tags=100%, list=40%, signal=167% |
| 6174 | QUINONE\_BIOSYNTHETIC\_PROCESS(GO:1901663) |  | 16 | 0.60 | 1.02 | 0.307 | 0.362 | 1.000 | 23667 | tags=100%, list=40%, signal=167% |
| 6175 | FUCOSYLATION(GO:0036065) |  | 27 | 0.65 | 1.02 | 0.310 | 0.362 | 1.000 | 12970 | tags=33%, list=22%, signal=43% |
| 6176 | CENP\_A\_CONTAINING\_NUCLEOSOME\_ASSEMBLY(GO:0034080) |  | 42 | 0.60 | 1.02 | 0.141 | 0.362 | 1.000 | 17348 | tags=55%, list=29%, signal=77% |
| 6177 | CENP\_A\_CONTAINING\_CHROMATIN\_ORGANIZATION(GO:0061641) |  | 42 | 0.60 | 1.02 | 0.141 | 0.362 | 1.000 | 17348 | tags=55%, list=29%, signal=77% |
| 6178 | REGULATION\_OF\_PROTEIN\_IMPORT\_INTO\_NUCLEUS\_\_TRANSLOCATION(GO:0033158) |  | 19 | 0.59 | 1.02 | 0.141 | 0.362 | 1.000 | 23558 | tags=79%, list=40%, signal=131% |
| 6179 | ER\_TO\_GOLGI\_TRANSPORT\_VESICLE\_MEMBRANE(GO:0012507) |  | 56 | 0.57 | 1.02 | 0.141 | 0.362 | 1.000 | 21621 | tags=59%, list=37%, signal=93% |
| 6180 | NUCLEAR\_TRANSCRIBED\_MRNA\_CATABOLIC\_PROCESS\_\_NONSENSE\_MEDIATED\_DECAY(GO:0000184) |  | 125 | 0.57 | 1.02 | 0.141 | 0.362 | 1.000 | 24427 | tags=90%, list=41%, signal=154% |
| 6181 | NUCLEOSOME\_ASSEMBLY(GO:0006334) |  | 144 | 0.61 | 1.02 | 0.141 | 0.362 | 1.000 | 20669 | tags=49%, list=35%, signal=75% |
| 6182 | SIGNAL\_TRANSDUCER\_ACTIVITY\_\_DOWNSTREAM\_OF\_RECEPTOR(GO:0005057) |  | 124 | 0.59 | 1.02 | 0.141 | 0.362 | 1.000 | 18373 | tags=48%, list=31%, signal=69% |
| 6183 | AUTOPHAGOSOME\_ASSEMBLY(GO:0000045) |  | 89 | 0.58 | 1.02 | 0.141 | 0.362 | 1.000 | 20702 | tags=62%, list=35%, signal=95% |
| 6184 | PROTEIN\_ADP\_RIBOSYLATION(GO:0006471) |  | 31 | 0.62 | 1.02 | 0.141 | 0.362 | 1.000 | 11470 | tags=26%, list=19%, signal=32% |
| 6185 | G\_PROTEIN\_COUPLED\_GLUTAMATE\_RECEPTOR\_SIGNALING\_PATHWAY(GO:0007216) |  | 15 | 0.64 | 1.02 | 0.310 | 0.362 | 1.000 | 16259 | tags=40%, list=27%, signal=55% |
| 6186 | TRIGLYCERIDE\_BIOSYNTHETIC\_PROCESS(GO:0019432) |  | 34 | 0.60 | 1.02 | 0.307 | 0.362 | 1.000 | 18054 | tags=47%, list=31%, signal=68% |
| 6187 | MODULATION\_BY\_SYMBIONT\_OF\_HOST\_CELLULAR\_PROCESS(GO:0044068) |  | 30 | 0.59 | 1.02 | 0.141 | 0.362 | 1.000 | 22440 | tags=73%, list=38%, signal=118% |
| 6188 | GLUCOCORTICOID\_RECEPTOR\_SIGNALING\_PATHWAY(GO:0042921) |  | 16 | 0.58 | 1.02 | 0.141 | 0.362 | 1.000 | 23488 | tags=94%, list=40%, signal=155% |
| 6189 | MRNA\_CIS\_SPLICING\_\_VIA\_SPLICEOSOME(GO:0045292) |  | 18 | 0.59 | 1.02 | 0.141 | 0.362 | 1.000 | 24161 | tags=94%, list=41%, signal=160% |
| 6190 | METAL\_CLUSTER\_BINDING(GO:0051540) |  | 66 | 0.57 | 1.02 | 0.141 | 0.362 | 1.000 | 24636 | tags=85%, list=42%, signal=145% |
| 6191 | IRON\_SULFUR\_CLUSTER\_BINDING(GO:0051536) |  | 66 | 0.57 | 1.02 | 0.141 | 0.362 | 1.000 | 24636 | tags=85%, list=42%, signal=145% |
| 6192 | HISTONE\_METHYLTRANSFERASE\_ACTIVITY\_(H3\_K4\_SPECIFIC)(GO:0042800) |  | 17 | 0.60 | 1.02 | 0.141 | 0.362 | 1.000 | 23844 | tags=94%, list=40%, signal=158% |
| 6193 | PROTEIN\_CYSTEINE\_S\_ACYLTRANSFERASE\_ACTIVITY(GO:0019707) |  | 27 | 0.58 | 1.02 | 0.141 | 0.362 | 1.000 | 18819 | tags=63%, list=32%, signal=92% |
| 6194 | PROTEIN\_CYSTEINE\_S\_PALMITOYLTRANSFERASE\_ACTIVITY(GO:0019706) |  | 27 | 0.58 | 1.02 | 0.141 | 0.362 | 1.000 | 18819 | tags=63%, list=32%, signal=92% |
| 6195 | BRANCH\_ELONGATION\_OF\_AN\_EPITHELIUM(GO:0060602) |  | 20 | 0.67 | 1.02 | 0.141 | 0.362 | 1.000 | 15678 | tags=55%, list=27%, signal=75% |
| 6196 | ESTABLISHMENT\_OF\_PROTEIN\_LOCALIZATION\_TO\_ENDOPLASMIC\_RETICULUM(GO:0072599) |  | 105 | 0.57 | 1.02 | 0.141 | 0.362 | 1.000 | 24601 | tags=93%, list=42%, signal=160% |
| 6197 | TRANSCRIPTION\_COACTIVATOR\_BINDING(GO:0001223) |  | 20 | 0.64 | 1.02 | 0.141 | 0.362 | 1.000 | 16715 | tags=55%, list=28%, signal=77% |
| 6198 | PROTEIN\_LOCALIZATION\_TO\_LYSOSOME(GO:0061462) |  | 31 | 0.57 | 1.02 | 0.141 | 0.362 | 1.000 | 24672 | tags=87%, list=42%, signal=149% |
| 6199 | INTRINSIC\_COMPONENT\_OF\_MITOCHONDRIAL\_INNER\_MEMBRANE(GO:0031304) |  | 41 | 0.58 | 1.02 | 0.141 | 0.363 | 1.000 | 24483 | tags=90%, list=41%, signal=154% |
| 6200 | INTEGRAL\_COMPONENT\_OF\_MITOCHONDRIAL\_INNER\_MEMBRANE(GO:0031305) |  | 41 | 0.58 | 1.02 | 0.141 | 0.363 | 1.000 | 24483 | tags=90%, list=41%, signal=154% |
| 6201 | REGULATION\_OF\_INTERFERON\_BETA\_PRODUCTION(GO:0032648) |  | 47 | 0.59 | 1.02 | 0.141 | 0.362 | 1.000 | 16603 | tags=40%, list=28%, signal=56% |
| 6202 | ENDOSOME\_TO\_LYSOSOME\_TRANSPORT(GO:0008333) |  | 50 | 0.57 | 1.02 | 0.141 | 0.363 | 1.000 | 23913 | tags=76%, list=40%, signal=127% |
| 6203 | PHOSPHATIDYLINOSITOL\_3\_5\_BISPHOSPHATE\_PHOSPHATASE\_ACTIVITY(GO:0106018) |  | 15 | 0.60 | 1.02 | 0.481 | 0.363 | 1.000 | 23205 | tags=93%, list=39%, signal=154% |
| 6204 | POSITIVE\_REGULATION\_OF\_NEUTROPHIL\_CHEMOTAXIS(GO:0090023) |  | 30 | 0.62 | 1.02 | 0.307 | 0.363 | 1.000 | 15981 | tags=40%, list=27%, signal=55% |
| 6205 | EXOCYTIC\_PROCESS(GO:0140029) |  | 70 | 0.58 | 1.02 | 0.141 | 0.363 | 1.000 | 19037 | tags=59%, list=32%, signal=86% |
| 6206 | ANTIGEN\_PROCESSING\_AND\_PRESENTATION\_OF\_EXOGENOUS\_PEPTIDE\_ANTIGEN(GO:0002478) |  | 170 | 0.58 | 1.02 | 0.141 | 0.363 | 1.000 | 19867 | tags=53%, list=34%, signal=79% |
| 6207 | REGULATION\_OF\_GLUCOSE\_IMPORT\_IN\_RESPONSE\_TO\_INSULIN\_STIMULUS(GO:2001273) |  | 23 | 0.62 | 1.02 | 0.141 | 0.363 | 1.000 | 20597 | tags=48%, list=35%, signal=73% |
| 6208 | NEGATIVE\_REGULATION\_OF\_CELL\_KILLING(GO:0031342) |  | 17 | 0.59 | 1.02 | 0.141 | 0.363 | 1.000 | 15529 | tags=41%, list=26%, signal=56% |
| 6209 | ENDONUCLEASE\_ACTIVITY\_\_ACTIVE\_WITH\_EITHER\_RIBO\_\_OR\_DEOXYRIBONUCLEIC\_ACIDS\_AND\_PRODUCING\_3'\_PHOSPHOMONOESTERS(GO:0016894) |  | 34 | 0.57 | 1.02 | 0.141 | 0.363 | 1.000 | 23049 | tags=68%, list=39%, signal=111% |
| 6210 | POSITIVE\_REGULATION\_OF\_MUSCLE\_HYPERTROPHY(GO:0014742) |  | 39 | 0.60 | 1.02 | 0.141 | 0.363 | 1.000 | 17738 | tags=44%, list=30%, signal=62% |
| 6211 | POSITIVE\_REGULATION\_OF\_CARDIAC\_MUSCLE\_HYPERTROPHY(GO:0010613) |  | 39 | 0.60 | 1.02 | 0.141 | 0.363 | 1.000 | 17738 | tags=44%, list=30%, signal=62% |
| 6212 | ZINC\_ION\_TRANSPORT(GO:0006829) |  | 27 | 0.58 | 1.02 | 0.141 | 0.363 | 1.000 | 18903 | tags=63%, list=32%, signal=93% |
| 6213 | RESPONSE\_TO\_LIPOPROTEIN\_PARTICLE(GO:0055094) |  | 25 | 0.60 | 1.02 | 0.141 | 0.363 | 1.000 | 6413 | tags=8%, list=11%, signal=9% |
| 6214 | TERTIARY\_GRANULE(GO:0070820) |  | 164 | 0.57 | 1.02 | 0.141 | 0.363 | 1.000 | 19239 | tags=34%, list=33%, signal=50% |
| 6215 | CYTOSOLIC\_RIBOSOME(GO:0022626) |  | 115 | 0.57 | 1.02 | 0.141 | 0.363 | 1.000 | 24601 | tags=89%, list=42%, signal=152% |
| 6216 | ACTIVATION\_OF\_INNATE\_IMMUNE\_RESPONSE(GO:0002218) |  | 299 | 0.59 | 1.02 | 0.141 | 0.363 | 1.000 | 17319 | tags=36%, list=29%, signal=51% |
| 6217 | POSITIVE\_REGULATION\_OF\_LEUKOCYTE\_MEDIATED\_IMMUNITY(GO:0002705) |  | 96 | 0.59 | 1.02 | 0.141 | 0.364 | 1.000 | 18589 | tags=47%, list=31%, signal=68% |
| 6218 | POSITIVE\_REGULATION\_OF\_TRANSCRIPTION\_ELONGATION\_FROM\_RNA\_POLYMERASE\_II\_PROMOTER(GO:0032968) |  | 15 | 0.59 | 1.02 | 0.141 | 0.364 | 1.000 | 23679 | tags=93%, list=40%, signal=156% |
| 6219 | SOLUTE:PROTON\_SYMPORTER\_ACTIVITY(GO:0015295) |  | 30 | 0.56 | 1.02 | 0.141 | 0.364 | 1.000 | 25226 | tags=83%, list=43%, signal=145% |
| 6220 | COTRANSLATIONAL\_PROTEIN\_TARGETING\_TO\_MEMBRANE(GO:0006613) |  | 98 | 0.57 | 1.02 | 0.141 | 0.364 | 1.000 | 24805 | tags=94%, list=42%, signal=161% |
| 6221 | METHYLATED\_HISTONE\_BINDING(GO:0035064) |  | 63 | 0.58 | 1.02 | 0.141 | 0.364 | 1.000 | 22759 | tags=75%, list=38%, signal=121% |
| 6222 | METHYLATION\_DEPENDENT\_PROTEIN\_BINDING(GO:0140034) |  | 63 | 0.58 | 1.02 | 0.141 | 0.364 | 1.000 | 22759 | tags=75%, list=38%, signal=121% |
| 6223 | POSITIVE\_REGULATION\_OF\_CALCIUM\_MEDIATED\_SIGNALING(GO:0050850) |  | 36 | 0.62 | 1.02 | 0.141 | 0.364 | 1.000 | 19067 | tags=50%, list=32%, signal=74% |
| 6224 | SCF\_UBIQUITIN\_LIGASE\_COMPLEX(GO:0019005) |  | 49 | 0.56 | 1.02 | 0.476 | 0.364 | 1.000 | 22746 | tags=71%, list=38%, signal=116% |
| 6225 | NEGATIVE\_REGULATION\_OF\_GLUCOSE\_IMPORT(GO:0046325) |  | 15 | 0.59 | 1.02 | 0.141 | 0.364 | 1.000 | 24352 | tags=73%, list=41%, signal=125% |
| 6226 | VACUOLAR\_TRANSPORT(GO:0007034) |  | 120 | 0.56 | 1.02 | 0.141 | 0.364 | 1.000 | 24906 | tags=81%, list=42%, signal=139% |
| 6227 | ANTIGEN\_PROCESSING\_AND\_PRESENTATION\_OF\_PEPTIDE\_ANTIGEN(GO:0048002) |  | 185 | 0.58 | 1.02 | 0.141 | 0.364 | 1.000 | 19867 | tags=51%, list=34%, signal=77% |
| 6228 | AMINE\_CATABOLIC\_PROCESS(GO:0009310) |  | 25 | 0.62 | 1.02 | 0.141 | 0.364 | 1.000 | 16748 | tags=40%, list=28%, signal=56% |
| 6229 | AMYLOID\_PRECURSOR\_PROTEIN\_CATABOLIC\_PROCESS(GO:0042987) |  | 32 | 0.59 | 1.02 | 0.141 | 0.364 | 1.000 | 21613 | tags=72%, list=37%, signal=113% |
| 6230 | U12\_TYPE\_SPLICEOSOMAL\_COMPLEX(GO:0005689) |  | 26 | 0.60 | 1.02 | 0.476 | 0.364 | 1.000 | 21830 | tags=88%, list=37%, signal=140% |
| 6231 | OOCYTE\_DIFFERENTIATION(GO:0009994) |  | 49 | 0.60 | 1.02 | 0.141 | 0.364 | 1.000 | 20968 | tags=61%, list=35%, signal=95% |
| 6232 | HISTONE\_H3\_K9\_MODIFICATION(GO:0061647) |  | 41 | 0.59 | 1.02 | 0.141 | 0.364 | 1.000 | 23268 | tags=80%, list=39%, signal=133% |
| 6233 | ACTIVATION\_OF\_IMMUNE\_RESPONSE(GO:0002253) |  | 655 | 0.59 | 1.02 | 0.141 | 0.364 | 1.000 | 18659 | tags=33%, list=32%, signal=48% |
| 6234 | RRNA\_TRANSCRIPTION(GO:0009303) |  | 30 | 0.57 | 1.02 | 0.141 | 0.364 | 1.000 | 24746 | tags=93%, list=42%, signal=160% |
| 6235 | PROTEIN\_TARGETING\_TO\_ER(GO:0045047) |  | 101 | 0.57 | 1.02 | 0.141 | 0.364 | 1.000 | 24805 | tags=94%, list=42%, signal=162% |
| 6236 | REGULATION\_OF\_TRANSMISSION\_OF\_NERVE\_IMPULSE(GO:0051969) |  | 16 | 0.58 | 1.02 | 0.307 | 0.365 | 1.000 | 23360 | tags=69%, list=40%, signal=114% |
| 6237 | FC\_GAMMA\_RECEPTOR\_SIGNALING\_PATHWAY\_INVOLVED\_IN\_PHAGOCYTOSIS(GO:0038096) |  | 128 | 0.56 | 1.02 | 0.141 | 0.365 | 1.000 | 18981 | tags=26%, list=32%, signal=38% |
| 6238 | IMMUNE\_RESPONSE\_REGULATING\_CELL\_SURFACE\_RECEPTOR\_SIGNALING\_PATHWAY\_INVOLVED\_IN\_PHAGOCYTOSIS(GO:0002433) |  | 128 | 0.56 | 1.02 | 0.141 | 0.365 | 1.000 | 18981 | tags=26%, list=32%, signal=38% |
| 6239 | PROTEIN\_KINASE\_ACTIVATOR\_ACTIVITY(GO:0030295) |  | 75 | 0.58 | 1.02 | 0.141 | 0.365 | 1.000 | 18981 | tags=57%, list=32%, signal=84% |
| 6240 | NEGATIVE\_REGULATION\_OF\_VOLTAGE\_GATED\_CALCIUM\_CHANNEL\_ACTIVITY(GO:1901386) |  | 17 | 0.61 | 1.02 | 0.307 | 0.365 | 1.000 | 19248 | tags=41%, list=33%, signal=61% |
| 6241 | POSITIVE\_REGULATION\_OF\_NITRIC\_OXIDE\_SYNTHASE\_ACTIVITY(GO:0051000) |  | 19 | 0.57 | 1.02 | 0.141 | 0.365 | 1.000 | 16715 | tags=42%, list=28%, signal=59% |
| 6242 | CYTOSOLIC\_LARGE\_RIBOSOMAL\_SUBUNIT(GO:0022625) |  | 62 | 0.57 | 1.02 | 0.141 | 0.365 | 1.000 | 24309 | tags=90%, list=41%, signal=153% |
| 6243 | ADENYLATE\_CYCLASE\_ACTIVATING\_ADRENERGIC\_RECEPTOR\_SIGNALING\_PATHWAY(GO:0071880) |  | 21 | 0.59 | 1.02 | 0.310 | 0.365 | 1.000 | 17432 | tags=38%, list=29%, signal=54% |
| 6244 | SRP\_DEPENDENT\_COTRANSLATIONAL\_PROTEIN\_TARGETING\_TO\_MEMBRANE(GO:0006614) |  | 92 | 0.57 | 1.01 | 0.141 | 0.365 | 1.000 | 24805 | tags=95%, list=42%, signal=163% |
| 6245 | AUTOPHAGOSOME\_ORGANIZATION(GO:1905037) |  | 92 | 0.57 | 1.01 | 0.141 | 0.365 | 1.000 | 20702 | tags=60%, list=35%, signal=92% |
| 6246 | PYRIMIDINE\_RIBONUCLEOTIDE\_BIOSYNTHETIC\_PROCESS(GO:0009220) |  | 23 | 0.57 | 1.01 | 0.310 | 0.365 | 1.000 | 23753 | tags=83%, list=40%, signal=138% |
| 6247 | WW\_DOMAIN\_BINDING(GO:0050699) |  | 31 | 0.61 | 1.01 | 0.141 | 0.365 | 1.000 | 17215 | tags=42%, list=29%, signal=59% |
| 6248 | NEGATIVE\_REGULATION\_OF\_HEART\_CONTRACTION(GO:0045822) |  | 26 | 0.60 | 1.01 | 0.141 | 0.365 | 1.000 | 12699 | tags=31%, list=21%, signal=39% |
| 6249 | GERMINAL\_CENTER\_FORMATION(GO:0002467) |  | 15 | 0.62 | 1.01 | 0.307 | 0.365 | 1.000 | 19789 | tags=53%, list=33%, signal=80% |
| 6250 | CELLULAR\_RESPONSE\_TO\_HYDROGEN\_PEROXIDE(GO:0070301) |  | 92 | 0.59 | 1.01 | 0.141 | 0.366 | 1.000 | 21146 | tags=57%, list=36%, signal=88% |
| 6251 | REGULATION\_OF\_INNATE\_IMMUNE\_RESPONSE(GO:0045088) |  | 421 | 0.59 | 1.01 | 0.141 | 0.366 | 1.000 | 17194 | tags=38%, list=29%, signal=53% |
| 6252 | NATURAL\_KILLER\_CELL\_MEDIATED\_CYTOTOXICITY(GO:0042267) |  | 55 | 0.58 | 1.01 | 0.141 | 0.366 | 1.000 | 15869 | tags=38%, list=27%, signal=52% |
| 6253 | POSITIVE\_REGULATION\_OF\_IMMUNOGLOBULIN\_PRODUCTION(GO:0002639) |  | 37 | 0.61 | 1.01 | 0.141 | 0.366 | 1.000 | 19540 | tags=57%, list=33%, signal=85% |
| 6254 | REGULATION\_OF\_T\_CELL\_MIGRATION(GO:2000404) |  | 32 | 0.62 | 1.01 | 0.141 | 0.366 | 1.000 | 10429 | tags=22%, list=18%, signal=27% |
| 6255 | ANTIGEN\_RECEPTOR\_MEDIATED\_SIGNALING\_PATHWAY(GO:0050851) |  | 266 | 0.58 | 1.01 | 0.141 | 0.366 | 1.000 | 18785 | tags=35%, list=32%, signal=51% |
| 6256 | REGULATION\_OF\_TRANSLATIONAL\_FIDELITY(GO:0006450) |  | 20 | 0.60 | 1.01 | 0.310 | 0.366 | 1.000 | 18454 | tags=70%, list=31%, signal=102% |
| 6257 | PTERIDINE\_CONTAINING\_COMPOUND\_BIOSYNTHETIC\_PROCESS(GO:0042559) |  | 15 | 0.59 | 1.01 | 0.310 | 0.366 | 1.000 | 24300 | tags=93%, list=41%, signal=158% |
| 6258 | NEGATIVE\_REGULATION\_OF\_GLUCOSE\_TRANSMEMBRANE\_TRANSPORT(GO:0010829) |  | 19 | 0.58 | 1.01 | 0.141 | 0.366 | 1.000 | 24352 | tags=68%, list=41%, signal=116% |
| 6259 | GLUCOCORTICOID\_BIOSYNTHETIC\_PROCESS(GO:0006704) |  | 17 | 0.63 | 1.01 | 0.307 | 0.366 | 1.000 | 10811 | tags=35%, list=18%, signal=43% |
| 6260 | SYNAPTIC\_VESICLE\_FUSION\_TO\_PRESYNAPTIC\_ACTIVE\_ZONE\_MEMBRANE(GO:0031629) |  | 16 | 0.59 | 1.01 | 0.307 | 0.366 | 1.000 | 18759 | tags=56%, list=32%, signal=82% |
| 6261 | VESICLE\_FUSION\_TO\_PLASMA\_MEMBRANE(GO:0099500) |  | 16 | 0.59 | 1.01 | 0.307 | 0.366 | 1.000 | 18759 | tags=56%, list=32%, signal=82% |
| 6262 | ROUGH\_ENDOPLASMIC\_RETICULUM\_MEMBRANE(GO:0030867) |  | 30 | 0.57 | 1.01 | 0.141 | 0.366 | 1.000 | 24601 | tags=83%, list=42%, signal=143% |
| 6263 | RESPIRATORY\_CHAIN\_COMPLEX\_IV\_ASSEMBLY(GO:0008535) |  | 26 | 0.58 | 1.01 | 0.141 | 0.366 | 1.000 | 18359 | tags=69%, list=31%, signal=100% |
| 6264 | AROMATIC\_AMINO\_ACID\_FAMILY\_CATABOLIC\_PROCESS(GO:0009074) |  | 20 | 0.63 | 1.01 | 0.141 | 0.366 | 1.000 | 19569 | tags=65%, list=33%, signal=97% |
| 6265 | PROTEIN\_PHOSPHATASE\_2A\_BINDING(GO:0051721) |  | 28 | 0.57 | 1.01 | 0.141 | 0.366 | 1.000 | 22809 | tags=68%, list=39%, signal=110% |
| 6266 | DNA\_DAMAGE\_RESPONSE\_\_SIGNAL\_TRANSDUCTION\_RESULTING\_IN\_TRANSCRIPTION(GO:0042772) |  | 20 | 0.62 | 1.01 | 0.141 | 0.366 | 1.000 | 20177 | tags=60%, list=34%, signal=91% |
| 6267 | CELLULAR\_GLUCOSE\_HOMEOSTASIS(GO:0001678) |  | 135 | 0.58 | 1.01 | 0.141 | 0.366 | 1.000 | 19378 | tags=50%, list=33%, signal=75% |
| 6268 | ALPHA\_MANNOSIDASE\_ACTIVITY(GO:0004559) |  | 16 | 0.58 | 1.01 | 0.141 | 0.366 | 1.000 | 21303 | tags=69%, list=36%, signal=107% |
| 6269 | TETRAPYRROLE\_BIOSYNTHETIC\_PROCESS(GO:0033014) |  | 29 | 0.58 | 1.01 | 0.141 | 0.366 | 1.000 | 21949 | tags=72%, list=37%, signal=115% |
| 6270 | POSITIVE\_REGULATION\_OF\_NEUTROPHIL\_MIGRATION(GO:1902624) |  | 33 | 0.61 | 1.01 | 0.307 | 0.366 | 1.000 | 16638 | tags=39%, list=28%, signal=55% |
| 6271 | CELLULAR\_RESPONSE\_TO\_GLUCOSE\_STIMULUS(GO:0071333) |  | 122 | 0.58 | 1.01 | 0.141 | 0.367 | 1.000 | 22910 | tags=67%, list=39%, signal=109% |
| 6272 | PRERIBOSOME\_\_LARGE\_SUBUNIT\_PRECURSOR(GO:0030687) |  | 25 | 0.56 | 1.01 | 0.307 | 0.366 | 1.000 | 24676 | tags=84%, list=42%, signal=144% |
| 6273 | 3'\_UTR\_MEDIATED\_MRNA\_STABILIZATION(GO:0070935) |  | 16 | 0.66 | 1.01 | 0.141 | 0.367 | 1.000 | 19321 | tags=56%, list=33%, signal=84% |
| 6274 | CELLULAR\_RESPONSE\_TO\_ACETYLCHOLINE(GO:1905145) |  | 23 | 0.59 | 1.01 | 0.310 | 0.367 | 1.000 | 9607 | tags=17%, list=16%, signal=21% |
| 6275 | RESPONSE\_TO\_ACETYLCHOLINE(GO:1905144) |  | 23 | 0.59 | 1.01 | 0.310 | 0.367 | 1.000 | 9607 | tags=17%, list=16%, signal=21% |
| 6276 | SIGNAL\_TRANSDUCTION\_INVOLVED\_IN\_CELLULAR\_RESPONSE\_TO\_AMMONIUM\_ION(GO:1903831) |  | 23 | 0.59 | 1.01 | 0.310 | 0.367 | 1.000 | 9607 | tags=17%, list=16%, signal=21% |
| 6277 | ACETYLCHOLINE\_RECEPTOR\_SIGNALING\_PATHWAY(GO:0095500) |  | 23 | 0.59 | 1.01 | 0.310 | 0.367 | 1.000 | 9607 | tags=17%, list=16%, signal=21% |
| 6278 | CELLULAR\_RESPONSE\_TO\_HEXOSE\_STIMULUS(GO:0071331) |  | 125 | 0.58 | 1.01 | 0.141 | 0.367 | 1.000 | 22910 | tags=67%, list=39%, signal=109% |
| 6279 | REGULATION\_OF\_METALLOPEPTIDASE\_ACTIVITY(GO:1905048) |  | 25 | 0.59 | 1.01 | 0.141 | 0.367 | 1.000 | 22859 | tags=44%, list=39%, signal=72% |
| 6280 | RNA\_CAP\_BINDING\_COMPLEX(GO:0034518) |  | 15 | 0.60 | 1.01 | 0.309 | 0.367 | 1.000 | 21089 | tags=80%, list=36%, signal=124% |
| 6281 | PLATELET\_MORPHOGENESIS(GO:0036344) |  | 21 | 0.62 | 1.01 | 0.310 | 0.367 | 1.000 | 15485 | tags=38%, list=26%, signal=52% |
| 6282 | S\_ACYLTRANSFERASE\_ACTIVITY(GO:0016417) |  | 31 | 0.58 | 1.01 | 0.307 | 0.367 | 1.000 | 20368 | tags=68%, list=34%, signal=103% |
| 6283 | CYTOPLASMIC\_SEQUESTERING\_OF\_PROTEIN(GO:0051220) |  | 24 | 0.58 | 1.01 | 0.141 | 0.367 | 1.000 | 17014 | tags=54%, list=29%, signal=76% |
| 6284 | REGULATION\_OF\_NUCLEASE\_ACTIVITY(GO:0032069) |  | 27 | 0.56 | 1.01 | 0.307 | 0.367 | 1.000 | 18513 | tags=52%, list=31%, signal=75% |
| 6285 | EUKARYOTIC\_TRANSLATION\_INITIATION\_FACTOR\_3\_COMPLEX(GO:0005852) |  | 17 | 0.59 | 1.01 | 0.141 | 0.367 | 1.000 | 20030 | tags=82%, list=34%, signal=124% |
| 6286 | HSP90\_PROTEIN\_BINDING(GO:0051879) |  | 35 | 0.56 | 1.01 | 0.141 | 0.367 | 1.000 | 22918 | tags=74%, list=39%, signal=121% |
| 6287 | THREONINE\_TYPE\_ENDOPEPTIDASE\_ACTIVITY(GO:0004298) |  | 26 | 0.57 | 1.01 | 0.307 | 0.367 | 1.000 | 24328 | tags=73%, list=41%, signal=124% |
| 6288 | THREONINE\_TYPE\_PEPTIDASE\_ACTIVITY(GO:0070003) |  | 26 | 0.57 | 1.01 | 0.307 | 0.367 | 1.000 | 24328 | tags=73%, list=41%, signal=124% |
| 6289 | NEGATIVE\_REGULATION\_OF\_BEHAVIOR(GO:0048521) |  | 15 | 0.60 | 1.01 | 0.475 | 0.367 | 1.000 | 14441 | tags=33%, list=24%, signal=44% |
| 6290 | TELOMERE\_CAPPING(GO:0016233) |  | 55 | 0.58 | 1.01 | 0.141 | 0.367 | 1.000 | 20903 | tags=62%, list=35%, signal=96% |
| 6291 | IMMUNE\_RESPONSE\_REGULATING\_SIGNALING\_PATHWAY(GO:0002764) |  | 611 | 0.58 | 1.01 | 0.141 | 0.367 | 1.000 | 18785 | tags=34%, list=32%, signal=49% |
| 6292 | RNA\_POLYADENYLATION(GO:0043631) |  | 44 | 0.56 | 1.01 | 0.307 | 0.367 | 1.000 | 24632 | tags=84%, list=42%, signal=144% |
| 6293 | REGULATION\_OF\_MITOCHONDRIAL\_FISSION(GO:0090140) |  | 23 | 0.61 | 1.01 | 0.476 | 0.367 | 1.000 | 19159 | tags=70%, list=32%, signal=103% |
| 6294 | REGULATION\_OF\_INSULIN\_LIKE\_GROWTH\_FACTOR\_RECEPTOR\_SIGNALING\_PATHWAY(GO:0043567) |  | 25 | 0.61 | 1.01 | 0.141 | 0.367 | 1.000 | 15922 | tags=28%, list=27%, signal=38% |
| 6295 | WOUND\_HEALING\_\_SPREADING\_OF\_EPIDERMAL\_CELLS(GO:0035313) |  | 17 | 0.60 | 1.01 | 0.335 | 0.367 | 1.000 | 19603 | tags=76%, list=33%, signal=114% |
| 6296 | T\_CELL\_RECEPTOR\_SIGNALING\_PATHWAY(GO:0050852) |  | 175 | 0.58 | 1.01 | 0.141 | 0.367 | 1.000 | 18785 | tags=42%, list=32%, signal=62% |
| 6297 | PYRIMIDINE\_NUCLEOSIDE\_TRIPHOSPHATE\_METABOLIC\_PROCESS(GO:0009147) |  | 23 | 0.58 | 1.01 | 0.310 | 0.367 | 1.000 | 20753 | tags=70%, list=35%, signal=107% |
| 6298 | PYRIMIDINE\_RIBONUCLEOSIDE\_BIOSYNTHETIC\_PROCESS(GO:0046132) |  | 22 | 0.57 | 1.01 | 0.310 | 0.367 | 1.000 | 23753 | tags=82%, list=40%, signal=137% |
| 6299 | PROTEIN\_LOCALIZATION\_TO\_CHROMOSOME(GO:0034502) |  | 78 | 0.61 | 1.01 | 0.141 | 0.368 | 1.000 | 22014 | tags=91%, list=37%, signal=145% |
| 6300 | GROWTH\_HORMONE\_SECRETION(GO:0030252) |  | 16 | 0.59 | 1.01 | 0.141 | 0.367 | 1.000 | 22379 | tags=75%, list=38%, signal=121% |
| 6301 | NEGATIVE\_REGULATION\_OF\_PROTEIN\_LOCALIZATION\_TO\_CELL\_PERIPHERY(GO:1904376) |  | 24 | 0.58 | 1.01 | 0.310 | 0.368 | 1.000 | 18481 | tags=54%, list=31%, signal=79% |
| 6302 | CELLULAR\_RESPONSE\_TO\_MONOSACCHARIDE\_STIMULUS(GO:0071326) |  | 126 | 0.58 | 1.01 | 0.141 | 0.368 | 1.000 | 22910 | tags=67%, list=39%, signal=109% |
| 6303 | REGULATION\_OF\_IMMUNOGLOBULIN\_PRODUCTION(GO:0002637) |  | 55 | 0.60 | 1.01 | 0.141 | 0.368 | 1.000 | 19540 | tags=51%, list=33%, signal=76% |
| 6304 | HISTONE\_H2A\_UBIQUITINATION(GO:0033522) |  | 25 | 0.58 | 1.01 | 0.481 | 0.368 | 1.000 | 23456 | tags=80%, list=40%, signal=133% |
| 6305 | LEUKOTRIENE\_BIOSYNTHETIC\_PROCESS(GO:0019370) |  | 18 | 0.60 | 1.01 | 0.476 | 0.368 | 1.000 | 16927 | tags=39%, list=29%, signal=54% |
| 6306 | REGULATION\_OF\_MITOPHAGY(GO:1901524) |  | 15 | 0.59 | 1.01 | 0.141 | 0.368 | 1.000 | 22939 | tags=93%, list=39%, signal=152% |
| 6307 | CYTOPLASMIC\_TRANSLATIONAL\_INITIATION(GO:0002183) |  | 26 | 0.59 | 1.01 | 0.141 | 0.368 | 1.000 | 24279 | tags=100%, list=41%, signal=170% |
| 6308 | MRNA\_POLYADENYLATION(GO:0006378) |  | 42 | 0.56 | 1.01 | 0.307 | 0.368 | 1.000 | 24632 | tags=83%, list=42%, signal=143% |
| 6309 | OXIDOREDUCTASE\_ACTIVITY\_\_ACTING\_ON\_NAD(P)H\_\_QUINONE\_OR\_SIMILAR\_COMPOUND\_AS\_ACCEPTOR(GO:0016655) |  | 60 | 0.57 | 1.01 | 0.141 | 0.368 | 1.000 | 24459 | tags=82%, list=41%, signal=139% |
| 6310 | RESPONSE\_TO\_INSECTICIDE(GO:0017085) |  | 16 | 0.63 | 1.01 | 0.141 | 0.368 | 1.000 | 17102 | tags=56%, list=29%, signal=79% |
| 6311 | EPIDERMAL\_GROWTH\_FACTOR\_RECEPTOR\_SIGNALING\_PATHWAY(GO:0007173) |  | 131 | 0.58 | 1.01 | 0.141 | 0.368 | 1.000 | 17190 | tags=35%, list=29%, signal=49% |
| 6312 | INTERACTION\_WITH\_HOST(GO:0051701) |  | 214 | 0.59 | 1.01 | 0.141 | 0.368 | 1.000 | 17188 | tags=33%, list=29%, signal=47% |
| 6313 | FC\_RECEPTOR\_MEDIATED\_STIMULATORY\_SIGNALING\_PATHWAY(GO:0002431) |  | 133 | 0.56 | 1.01 | 0.141 | 0.368 | 1.000 | 23696 | tags=41%, list=40%, signal=69% |
| 6314 | NEGATIVE\_REGULATION\_OF\_PROTEIN\_LOCALIZATION\_TO\_MEMBRANE(GO:1905476) |  | 30 | 0.58 | 1.01 | 0.310 | 0.368 | 1.000 | 19903 | tags=60%, list=34%, signal=90% |
| 6315 | INTRINSIC\_COMPONENT\_OF\_ENDOPLASMIC\_RETICULUM\_MEMBRANE(GO:0031227) |  | 147 | 0.56 | 1.01 | 0.141 | 0.368 | 1.000 | 24073 | tags=70%, list=41%, signal=118% |
| 6316 | MRNA\_CLEAVAGE(GO:0006379) |  | 28 | 0.58 | 1.01 | 0.307 | 0.368 | 1.000 | 22758 | tags=79%, list=38%, signal=128% |
| 6317 | POSITIVE\_REGULATION\_OF\_LEUKOCYTE\_CHEMOTAXIS(GO:0002690) |  | 86 | 0.61 | 1.01 | 0.141 | 0.368 | 1.000 | 15120 | tags=29%, list=26%, signal=39% |
| 6318 | IMMUNE\_RESPONSE\_REGULATING\_CELL\_SURFACE\_RECEPTOR\_SIGNALING\_PATHWAY(GO:0002768) |  | 461 | 0.57 | 1.01 | 0.141 | 0.368 | 1.000 | 18785 | tags=31%, list=32%, signal=45% |
| 6319 | STIMULATORY\_C\_TYPE\_LECTIN\_RECEPTOR\_SIGNALING\_PATHWAY(GO:0002223) |  | 120 | 0.57 | 1.01 | 0.141 | 0.369 | 1.000 | 20050 | tags=46%, list=34%, signal=69% |
| 6320 | AZUROPHIL\_GRANULE\_LUMEN(GO:0035578) |  | 90 | 0.58 | 1.01 | 0.141 | 0.369 | 1.000 | 18821 | tags=37%, list=32%, signal=54% |
| 6321 | PROTEIN\_TARGETING\_TO\_VACUOLE(GO:0006623) |  | 28 | 0.58 | 1.01 | 0.309 | 0.369 | 1.000 | 24672 | tags=93%, list=42%, signal=159% |
| 6322 | LRR\_DOMAIN\_BINDING(GO:0030275) |  | 17 | 0.56 | 1.01 | 0.141 | 0.369 | 1.000 | 24644 | tags=76%, list=42%, signal=131% |
| 6323 | N\_TERMINAL\_PROTEIN\_AMINO\_ACID\_ACETYLATION(GO:0006474) |  | 16 | 0.58 | 1.01 | 0.141 | 0.369 | 1.000 | 23797 | tags=94%, list=40%, signal=157% |
| 6324 | INNATE\_IMMUNE\_RESPONSE\_ACTIVATING\_CELL\_SURFACE\_RECEPTOR\_SIGNALING\_PATHWAY(GO:0002220) |  | 123 | 0.57 | 1.01 | 0.141 | 0.369 | 1.000 | 20050 | tags=45%, list=34%, signal=68% |
| 6325 | PEROXISOMAL\_MATRIX(GO:0005782) |  | 43 | 0.59 | 1.01 | 0.141 | 0.369 | 1.000 | 15758 | tags=47%, list=27%, signal=63% |
| 6326 | MICROBODY\_LUMEN(GO:0031907) |  | 43 | 0.59 | 1.01 | 0.141 | 0.369 | 1.000 | 15758 | tags=47%, list=27%, signal=63% |
| 6327 | EUCHROMATIN(GO:0000791) |  | 33 | 0.60 | 1.01 | 0.310 | 0.369 | 1.000 | 15684 | tags=48%, list=27%, signal=66% |
| 6328 | IMMUNE\_RESPONSE\_ACTIVATING\_SIGNAL\_TRANSDUCTION(GO:0002757) |  | 581 | 0.58 | 1.01 | 0.141 | 0.369 | 1.000 | 18785 | tags=33%, list=32%, signal=48% |
| 6329 | PROTEIN\_KINASE\_C\_BINDING(GO:0005080) |  | 49 | 0.58 | 1.01 | 0.310 | 0.369 | 1.000 | 16610 | tags=41%, list=28%, signal=57% |
| 6330 | CYTOCHROME\_COMPLEX\_ASSEMBLY(GO:0017004) |  | 36 | 0.57 | 1.01 | 0.141 | 0.369 | 1.000 | 23720 | tags=86%, list=40%, signal=144% |
| 6331 | PROTEIN\_O\_LINKED\_FUCOSYLATION(GO:0036066) |  | 15 | 0.67 | 1.01 | 0.476 | 0.369 | 1.000 | 12970 | tags=33%, list=22%, signal=43% |
| 6332 | INTEGRAL\_COMPONENT\_OF\_ENDOPLASMIC\_RETICULUM\_MEMBRANE(GO:0030176) |  | 142 | 0.56 | 1.01 | 0.141 | 0.369 | 1.000 | 24073 | tags=70%, list=41%, signal=118% |
| 6333 | CALCIUM\_ACTIVATED\_CATION\_CHANNEL\_ACTIVITY(GO:0005227) |  | 32 | 0.61 | 1.01 | 0.141 | 0.369 | 1.000 | 15347 | tags=41%, list=26%, signal=55% |
| 6334 | ZINC\_ION\_TRANSMEMBRANE\_TRANSPORTER\_ACTIVITY(GO:0005385) |  | 23 | 0.57 | 1.01 | 0.141 | 0.369 | 1.000 | 24034 | tags=87%, list=41%, signal=146% |
| 6335 | POSITIVE\_REGULATION\_OF\_G\_PROTEIN\_COUPLED\_RECEPTOR\_PROTEIN\_SIGNALING\_PATHWAY(GO:0045745) |  | 30 | 0.59 | 1.01 | 0.310 | 0.369 | 1.000 | 21992 | tags=67%, list=37%, signal=106% |
| 6336 | DNA\_DIRECTED\_RNA\_POLYMERASE\_II\_\_CORE\_COMPLEX(GO:0005665) |  | 19 | 0.57 | 1.01 | 0.476 | 0.369 | 1.000 | 25421 | tags=100%, list=43%, signal=175% |
| 6337 | DISRUPTION\_OF\_CELLS\_OF\_OTHER\_ORGANISM\_INVOLVED\_IN\_SYMBIOTIC\_INTERACTION(GO:0051818) |  | 18 | 0.60 | 1.01 | 0.141 | 0.369 | 1.000 | 23673 | tags=56%, list=40%, signal=93% |
| 6338 | NAD\_DEPENDENT\_PROTEIN\_DEACETYLASE\_ACTIVITY(GO:0034979) |  | 17 | 0.59 | 1.01 | 0.476 | 0.369 | 1.000 | 19177 | tags=59%, list=32%, signal=87% |
| 6339 | PHOTOPERIODISM(GO:0009648) |  | 22 | 0.57 | 1.01 | 0.141 | 0.369 | 1.000 | 22465 | tags=77%, list=38%, signal=125% |
| 6340 | RESPONSE\_TO\_MISFOLDED\_PROTEIN(GO:0051788) |  | 24 | 0.56 | 1.01 | 0.310 | 0.369 | 1.000 | 25069 | tags=92%, list=42%, signal=159% |
| 6341 | TRICARBOXYLIC\_ACID\_CYCLE(GO:0006099) |  | 32 | 0.57 | 1.01 | 0.476 | 0.369 | 1.000 | 21329 | tags=72%, list=36%, signal=112% |
| 6342 | PROTEIN\_DEGLYCOSYLATION(GO:0006517) |  | 36 | 0.56 | 1.01 | 0.141 | 0.369 | 1.000 | 21436 | tags=72%, list=36%, signal=113% |
| 6343 | PROTEIN\_TARGETING\_TO\_MEMBRANE(GO:0006612) |  | 164 | 0.56 | 1.01 | 0.141 | 0.370 | 1.000 | 24861 | tags=84%, list=42%, signal=145% |
| 6344 | PYRIMIDINE\_RIBONUCLEOSIDE\_TRIPHOSPHATE\_METABOLIC\_PROCESS(GO:0009208) |  | 17 | 0.56 | 1.01 | 0.310 | 0.370 | 1.000 | 23753 | tags=76%, list=40%, signal=128% |
| 6345 | AEROBIC\_RESPIRATION(GO:0009060) |  | 85 | 0.56 | 1.01 | 0.141 | 0.370 | 1.000 | 25098 | tags=86%, list=42%, signal=149% |
| 6346 | POSITIVE\_REGULATION\_OF\_PRODUCTION\_OF\_MOLECULAR\_MEDIATOR\_OF\_IMMUNE\_RESPONSE(GO:0002702) |  | 74 | 0.59 | 1.01 | 0.141 | 0.370 | 1.000 | 19540 | tags=49%, list=33%, signal=73% |
| 6347 | INTERFERON\_BETA\_PRODUCTION(GO:0032608) |  | 49 | 0.59 | 1.01 | 0.141 | 0.370 | 1.000 | 16603 | tags=39%, list=28%, signal=54% |
| 6348 | METHYLATION(GO:0032259) |  | 356 | 0.60 | 1.01 | 0.141 | 0.370 | 1.000 | 19904 | tags=61%, list=34%, signal=92% |
| 6349 | MITOCHONDRIAL\_ELECTRON\_TRANSPORT\_\_UBIQUINOL\_TO\_CYTOCHROME\_C(GO:0006122) |  | 15 | 0.57 | 1.01 | 0.141 | 0.370 | 1.000 | 24483 | tags=93%, list=41%, signal=159% |
| 6350 | REGULATION\_OF\_DEFENSE\_RESPONSE\_TO\_VIRUS(GO:0050688) |  | 68 | 0.59 | 1.01 | 0.141 | 0.370 | 1.000 | 15297 | tags=34%, list=26%, signal=46% |
| 6351 | HISTONE\_EXCHANGE(GO:0043486) |  | 58 | 0.59 | 1.01 | 0.141 | 0.370 | 1.000 | 22346 | tags=69%, list=38%, signal=111% |
| 6352 | KILLING\_OF\_CELLS\_IN\_OTHER\_ORGANISM\_INVOLVED\_IN\_SYMBIOTIC\_INTERACTION(GO:0051883) |  | 16 | 0.60 | 1.01 | 0.141 | 0.370 | 1.000 | 23673 | tags=63%, list=40%, signal=104% |
| 6353 | ORGANOPHOSPHATE\_ESTER\_TRANSMEMBRANE\_TRANSPORTER\_ACTIVITY(GO:0015605) |  | 22 | 0.61 | 1.01 | 0.141 | 0.370 | 1.000 | 19384 | tags=59%, list=33%, signal=88% |
| 6354 | ACTIVATION\_OF\_GTPASE\_ACTIVITY(GO:0090630) |  | 90 | 0.58 | 1.01 | 0.141 | 0.370 | 1.000 | 23051 | tags=67%, list=39%, signal=109% |
| 6355 | CARBOXY\_LYASE\_ACTIVITY(GO:0016831) |  | 38 | 0.57 | 1.01 | 0.307 | 0.371 | 1.000 | 19845 | tags=58%, list=34%, signal=87% |
| 6356 | MODIFICATION\_BY\_SYMBIONT\_OF\_HOST\_MORPHOLOGY\_OR\_PHYSIOLOGY(GO:0044003) |  | 48 | 0.57 | 1.01 | 0.141 | 0.371 | 1.000 | 21805 | tags=58%, list=37%, signal=92% |
| 6357 | POSITIVE\_REGULATION\_OF\_VASCULAR\_ASSOCIATED\_SMOOTH\_MUSCLE\_CELL\_MIGRATION(GO:1904754) |  | 20 | 0.63 | 1.01 | 0.141 | 0.371 | 1.000 | 16715 | tags=50%, list=28%, signal=70% |
| 6358 | PROTEIN\_LOCALIZATION\_TO\_GOLGI\_APPARATUS(GO:0034067) |  | 20 | 0.56 | 1.01 | 0.141 | 0.371 | 1.000 | 26064 | tags=95%, list=44%, signal=170% |
| 6359 | DNA\_DAMAGE\_RESPONSE\_\_SIGNAL\_TRANSDUCTION\_BY\_P53\_CLASS\_MEDIATOR\_RESULTING\_IN\_TRANSCRIPTION\_OF\_P21\_CLASS\_MEDIATOR(GO:0006978) |  | 19 | 0.62 | 1.01 | 0.141 | 0.371 | 1.000 | 20177 | tags=58%, list=34%, signal=88% |
| 6360 | VOLTAGE\_GATED\_ANION\_CHANNEL\_ACTIVITY(GO:0008308) |  | 18 | 0.57 | 1.01 | 0.482 | 0.371 | 1.000 | 23017 | tags=78%, list=39%, signal=127% |
| 6361 | KETONE\_BIOSYNTHETIC\_PROCESS(GO:0042181) |  | 41 | 0.59 | 1.01 | 0.307 | 0.371 | 1.000 | 23738 | tags=76%, list=40%, signal=126% |
| 6362 | DNA\_TEMPLATED\_TRANSCRIPTIONAL\_PREINITIATION\_COMPLEX\_ASSEMBLY(GO:0070897) |  | 43 | 0.61 | 1.01 | 0.141 | 0.371 | 1.000 | 18494 | tags=60%, list=31%, signal=88% |
| 6363 | U5\_SNRNP(GO:0005682) |  | 22 | 0.58 | 1.01 | 0.141 | 0.371 | 1.000 | 23078 | tags=73%, list=39%, signal=119% |
| 6364 | RESPONSE\_TO\_ARSENIC\_CONTAINING\_SUBSTANCE(GO:0046685) |  | 31 | 0.58 | 1.01 | 0.141 | 0.371 | 1.000 | 21080 | tags=68%, list=36%, signal=105% |
| 6365 | PROTEASOME\_CORE\_COMPLEX(GO:0005839) |  | 26 | 0.56 | 1.01 | 0.307 | 0.371 | 1.000 | 24328 | tags=73%, list=41%, signal=124% |
| 6366 | N\_METHYLTRANSFERASE\_ACTIVITY(GO:0008170) |  | 94 | 0.59 | 1.01 | 0.141 | 0.371 | 1.000 | 21531 | tags=77%, list=36%, signal=120% |
| 6367 | NUCLEOTIDE\_SUGAR\_METABOLIC\_PROCESS(GO:0009225) |  | 35 | 0.57 | 1.01 | 0.141 | 0.371 | 1.000 | 22842 | tags=66%, list=39%, signal=107% |
| 6368 | CELL\_AGGREGATION(GO:0098743) |  | 21 | 0.63 | 1.01 | 0.141 | 0.371 | 1.000 | 20024 | tags=57%, list=34%, signal=86% |
| 6369 | REGULATION\_OF\_MRNA\_3'\_END\_PROCESSING(GO:0031440) |  | 22 | 0.57 | 1.01 | 0.141 | 0.371 | 1.000 | 18871 | tags=64%, list=32%, signal=93% |
| 6370 | ERBB\_SIGNALING\_PATHWAY(GO:0038127) |  | 153 | 0.58 | 1.01 | 0.141 | 0.371 | 1.000 | 17190 | tags=36%, list=29%, signal=51% |
| 6371 | OXYGEN\_CARRIER\_ACTIVITY(GO:0005344) |  | 15 | 0.66 | 1.01 | 0.307 | 0.371 | 1.000 | 15820 | tags=33%, list=27%, signal=45% |
| 6372 | TRANSITION\_METAL\_ION\_TRANSPORT(GO:0000041) |  | 110 | 0.57 | 1.01 | 0.141 | 0.371 | 1.000 | 19199 | tags=47%, list=32%, signal=70% |
| 6373 | RESPONSE\_TO\_ANTINEOPLASTIC\_AGENT(GO:0097327) |  | 90 | 0.60 | 1.01 | 0.141 | 0.372 | 1.000 | 22051 | tags=61%, list=37%, signal=97% |
| 6374 | NADP\_METABOLIC\_PROCESS(GO:0006739) |  | 37 | 0.57 | 1.01 | 0.476 | 0.372 | 1.000 | 22714 | tags=70%, list=38%, signal=114% |
| 6375 | POSITIVE\_REGULATION\_OF\_LAMELLIPODIUM\_ASSEMBLY(GO:0010592) |  | 18 | 0.58 | 1.01 | 0.307 | 0.372 | 1.000 | 24081 | tags=83%, list=41%, signal=141% |
| 6376 | IMMUNE\_RESPONSE\_ACTIVATING\_CELL\_SURFACE\_RECEPTOR\_SIGNALING\_PATHWAY(GO:0002429) |  | 431 | 0.57 | 1.01 | 0.141 | 0.372 | 1.000 | 18785 | tags=31%, list=32%, signal=45% |
| 6377 | CELLULAR\_BIOGENIC\_AMINE\_CATABOLIC\_PROCESS(GO:0042402) |  | 23 | 0.61 | 1.01 | 0.141 | 0.372 | 1.000 | 16748 | tags=39%, list=28%, signal=55% |
| 6378 | TOLL\_LIKE\_RECEPTOR\_SIGNALING\_PATHWAY(GO:0002224) |  | 136 | 0.59 | 1.01 | 0.141 | 0.372 | 1.000 | 17144 | tags=36%, list=29%, signal=51% |
| 6379 | PHOSPHATIDYLINOSITOL\_PHOSPHATE\_PHOSPHATASE\_ACTIVITY(GO:0052866) |  | 30 | 0.56 | 1.01 | 0.141 | 0.372 | 1.000 | 25291 | tags=90%, list=43%, signal=157% |
| 6380 | DIOL\_BIOSYNTHETIC\_PROCESS(GO:0034312) |  | 15 | 0.57 | 1.01 | 0.141 | 0.372 | 1.000 | 24712 | tags=87%, list=42%, signal=149% |
| 6381 | ORGANELLE\_INHERITANCE(GO:0048308) |  | 15 | 0.57 | 1.01 | 0.141 | 0.372 | 1.000 | 25464 | tags=100%, list=43%, signal=176% |
| 6382 | MANNOSIDASE\_ACTIVITY(GO:0015923) |  | 17 | 0.57 | 1.01 | 0.141 | 0.372 | 1.000 | 22003 | tags=71%, list=37%, signal=112% |
| 6383 | NEGATIVE\_REGULATION\_OF\_ACTIN\_FILAMENT\_DEPOLYMERIZATION(GO:0030835) |  | 39 | 0.57 | 1.01 | 0.141 | 0.372 | 1.000 | 20010 | tags=59%, list=34%, signal=89% |
| 6384 | LIPID\_STORAGE(GO:0019915) |  | 64 | 0.57 | 1.01 | 0.141 | 0.372 | 1.000 | 21621 | tags=47%, list=37%, signal=74% |
| 6385 | ATP\_HYDROLYSIS\_COUPLED\_CATION\_TRANSMEMBRANE\_TRANSPORT(GO:0099132) |  | 57 | 0.58 | 1.00 | 0.141 | 0.372 | 1.000 | 21617 | tags=67%, list=37%, signal=105% |
| 6386 | POSITIVE\_REGULATION\_OF\_AMINE\_TRANSPORT(GO:0051954) |  | 33 | 0.59 | 1.00 | 0.307 | 0.372 | 1.000 | 21674 | tags=61%, list=37%, signal=96% |
| 6387 | NEGATIVE\_REGULATION\_OF\_NEUROLOGICAL\_SYSTEM\_PROCESS(GO:0031645) |  | 15 | 0.59 | 1.00 | 0.309 | 0.372 | 1.000 | 22138 | tags=60%, list=37%, signal=96% |
| 6388 | STEROID\_DEHYDROGENASE\_ACTIVITY\_\_ACTING\_ON\_THE\_CH\_OH\_GROUP\_OF\_DONORS\_\_NAD\_OR\_NADP\_AS\_ACCEPTOR(GO:0033764) |  | 27 | 0.62 | 1.00 | 0.141 | 0.372 | 1.000 | 10115 | tags=30%, list=17%, signal=36% |
| 6389 | CUL4\_RING\_E3\_UBIQUITIN\_LIGASE\_COMPLEX(GO:0080008) |  | 32 | 0.58 | 1.00 | 0.309 | 0.372 | 1.000 | 21198 | tags=75%, list=36%, signal=117% |
| 6390 | PHOSPHATE\_ION\_TRANSPORT(GO:0006817) |  | 21 | 0.58 | 1.00 | 0.310 | 0.373 | 1.000 | 18929 | tags=62%, list=32%, signal=91% |
| 6391 | ENTRAINMENT\_OF\_CIRCADIAN\_CLOCK(GO:0009649) |  | 24 | 0.57 | 1.00 | 0.141 | 0.373 | 1.000 | 22367 | tags=75%, list=38%, signal=121% |
| 6392 | TRANSCRIPTIONALLY\_ACTIVE\_CHROMATIN(GO:0035327) |  | 24 | 0.61 | 1.00 | 0.141 | 0.373 | 1.000 | 19092 | tags=58%, list=32%, signal=86% |
| 6393 | POSITIVE\_REGULATION\_OF\_CAMP\_MEDIATED\_SIGNALING(GO:0043950) |  | 22 | 0.60 | 1.00 | 0.482 | 0.373 | 1.000 | 18573 | tags=59%, list=31%, signal=86% |
| 6394 | NEGATIVE\_REGULATION\_OF\_NEURON\_PROJECTION\_REGENERATION(GO:0070571) |  | 15 | 0.60 | 1.00 | 0.307 | 0.373 | 1.000 | 21439 | tags=73%, list=36%, signal=115% |
| 6395 | BETA\_CATENIN\_TCF\_COMPLEX\_ASSEMBLY(GO:1904837) |  | 44 | 0.60 | 1.00 | 0.141 | 0.373 | 1.000 | 22149 | tags=64%, list=37%, signal=102% |
| 6396 | ZINC\_ION\_TRANSMEMBRANE\_TRANSPORT(GO:0071577) |  | 24 | 0.57 | 1.00 | 0.476 | 0.373 | 1.000 | 24034 | tags=83%, list=41%, signal=140% |
| 6397 | COENZYME\_A\_METABOLIC\_PROCESS(GO:0015936) |  | 15 | 0.58 | 1.00 | 0.141 | 0.373 | 1.000 | 25039 | tags=100%, list=42%, signal=173% |
| 6398 | GLUCOSE\_METABOLIC\_PROCESS(GO:0006006) |  | 191 | 0.59 | 1.00 | 0.141 | 0.373 | 1.000 | 20759 | tags=52%, list=35%, signal=80% |
| 6399 | POSITIVE\_REGULATION\_OF\_RESPONSE\_TO\_DRUG(GO:2001025) |  | 27 | 0.58 | 1.00 | 0.475 | 0.373 | 1.000 | 22606 | tags=63%, list=38%, signal=102% |
| 6400 | CALCIUM\_MEDIATED\_SIGNALING\_USING\_INTRACELLULAR\_CALCIUM\_SOURCE(GO:0035584) |  | 20 | 0.62 | 1.00 | 0.476 | 0.373 | 1.000 | 7544 | tags=20%, list=13%, signal=23% |
| 6401 | AUTOPHAGY\_OF\_NUCLEUS(GO:0044804) |  | 19 | 0.57 | 1.00 | 0.476 | 0.374 | 1.000 | 21821 | tags=68%, list=37%, signal=108% |
| 6402 | NIK\_NF\_KAPPAB\_SIGNALING(GO:0038061) |  | 129 | 0.56 | 1.00 | 0.141 | 0.374 | 1.000 | 24328 | tags=71%, list=41%, signal=120% |
| 6403 | NEGATIVE\_REGULATION\_OF\_INTRINSIC\_APOPTOTIC\_SIGNALING\_PATHWAY\_BY\_P53\_CLASS\_MEDIATOR(GO:1902254) |  | 20 | 0.60 | 1.00 | 0.141 | 0.374 | 1.000 | 17869 | tags=40%, list=30%, signal=57% |
| 6404 | PROTON\_TRANSMEMBRANE\_TRANSPORTER\_ACTIVITY(GO:0015078) |  | 130 | 0.56 | 1.00 | 0.141 | 0.374 | 1.000 | 24260 | tags=74%, list=41%, signal=125% |
| 6405 | POSITIVE\_REGULATION\_OF\_VIRAL\_GENOME\_REPLICATION(GO:0045070) |  | 35 | 0.58 | 1.00 | 0.481 | 0.374 | 1.000 | 22174 | tags=74%, list=37%, signal=119% |
| 6406 | POSITIVE\_REGULATION\_OF\_ERYTHROCYTE\_DIFFERENTIATION(GO:0045648) |  | 31 | 0.58 | 1.00 | 0.310 | 0.374 | 1.000 | 23261 | tags=77%, list=39%, signal=128% |
| 6407 | REGULATION\_OF\_PHAGOCYTOSIS(GO:0050764) |  | 70 | 0.58 | 1.00 | 0.141 | 0.374 | 1.000 | 24115 | tags=57%, list=41%, signal=96% |
| 6408 | DETECTION\_OF\_CHEMICAL\_STIMULUS\_INVOLVED\_IN\_SENSORY\_PERCEPTION\_OF\_TASTE(GO:0050912) |  | 40 | 0.59 | 1.00 | 0.309 | 0.374 | 1.000 | 9774 | tags=13%, list=17%, signal=15% |
| 6409 | POSITIVE\_REGULATION\_OF\_LYMPHOCYTE\_MIGRATION(GO:2000403) |  | 30 | 0.59 | 1.00 | 0.307 | 0.374 | 1.000 | 10429 | tags=17%, list=18%, signal=20% |
| 6410 | BARBED\_END\_ACTIN\_FILAMENT\_CAPPING(GO:0051016) |  | 18 | 0.56 | 1.00 | 0.141 | 0.374 | 1.000 | 25303 | tags=83%, list=43%, signal=146% |
| 6411 | RETROMER\_COMPLEX(GO:0030904) |  | 22 | 0.58 | 1.00 | 0.309 | 0.374 | 1.000 | 22176 | tags=77%, list=38%, signal=124% |
| 6412 | NEGATIVE\_REGULATION\_OF\_ACTIN\_FILAMENT\_POLYMERIZATION(GO:0030837) |  | 52 | 0.57 | 1.00 | 0.141 | 0.374 | 1.000 | 20178 | tags=54%, list=34%, signal=82% |
| 6413 | NEUROLOGICAL\_SYSTEM\_PROCESS\_INVOLVED\_IN\_REGULATION\_OF\_SYSTEMIC\_ARTERIAL\_BLOOD\_PRESSURE(GO:0001976) |  | 15 | 0.60 | 1.00 | 0.476 | 0.374 | 1.000 | 21904 | tags=67%, list=37%, signal=106% |
| 6414 | PHOSPHOLIPID\_DEPHOSPHORYLATION(GO:0046839) |  | 37 | 0.56 | 1.00 | 0.141 | 0.374 | 1.000 | 25291 | tags=86%, list=43%, signal=151% |
| 6415 | GOLGI\_TRANSPORT\_COMPLEX(GO:0017119) |  | 17 | 0.56 | 1.00 | 0.310 | 0.374 | 1.000 | 25886 | tags=94%, list=44%, signal=167% |
| 6416 | RECEPTOR\_SIGNALING\_COMPLEX\_SCAFFOLD\_ACTIVITY(GO:0030159) |  | 23 | 0.60 | 1.00 | 0.141 | 0.374 | 1.000 | 21673 | tags=70%, list=37%, signal=110% |
| 6417 | RNA\_SPLICING\_\_VIA\_ENDONUCLEOLYTIC\_CLEAVAGE\_AND\_LIGATION(GO:0000394) |  | 19 | 0.57 | 1.00 | 0.476 | 0.374 | 1.000 | 18644 | tags=53%, list=32%, signal=77% |
| 6418 | TRANSCRIPTION\_COFACTOR\_BINDING(GO:0001221) |  | 37 | 0.61 | 1.00 | 0.141 | 0.374 | 1.000 | 16956 | tags=46%, list=29%, signal=64% |
| 6419 | NEGATIVE\_REGULATION\_OF\_ENDOPLASMIC\_RETICULUM\_STRESS\_INDUCED\_INTRINSIC\_APOPTOTIC\_SIGNALING\_PATHWAY(GO:1902236) |  | 19 | 0.58 | 1.00 | 0.310 | 0.375 | 1.000 | 18692 | tags=53%, list=32%, signal=77% |
| 6420 | REGULATION\_OF\_LAMELLIPODIUM\_ASSEMBLY(GO:0010591) |  | 33 | 0.58 | 1.00 | 0.476 | 0.375 | 1.000 | 21573 | tags=58%, list=36%, signal=91% |
| 6421 | PROTEIN\_KINASE\_REGULATOR\_ACTIVITY(GO:0019887) |  | 192 | 0.58 | 1.00 | 0.141 | 0.375 | 1.000 | 23403 | tags=71%, list=40%, signal=118% |
| 6422 | SIGNAL\_TRANSDUCTION\_INVOLVED\_IN\_DNA\_INTEGRITY\_CHECKPOINT(GO:0072401) |  | 78 | 0.61 | 1.00 | 0.141 | 0.375 | 1.000 | 21594 | tags=77%, list=37%, signal=121% |
| 6423 | SIGNAL\_TRANSDUCTION\_INVOLVED\_IN\_DNA\_DAMAGE\_CHECKPOINT(GO:0072422) |  | 78 | 0.61 | 1.00 | 0.141 | 0.375 | 1.000 | 21594 | tags=77%, list=37%, signal=121% |
| 6424 | ALDEHYDE\_BIOSYNTHETIC\_PROCESS(GO:0046184) |  | 15 | 0.61 | 1.00 | 0.476 | 0.375 | 1.000 | 21530 | tags=60%, list=36%, signal=94% |
| 6425 | SPINDLE\_POLE(GO:0000922) |  | 152 | 0.60 | 1.00 | 0.141 | 0.375 | 1.000 | 21930 | tags=82%, list=37%, signal=129% |
| 6426 | POSITIVE\_REGULATION\_OF\_ENDOTHELIAL\_CELL\_APOPTOTIC\_PROCESS(GO:2000353) |  | 19 | 0.62 | 1.00 | 0.141 | 0.375 | 1.000 | 19286 | tags=32%, list=33%, signal=47% |
| 6427 | EXTRINSIC\_COMPONENT\_OF\_ORGANELLE\_MEMBRANE(GO:0031312) |  | 33 | 0.57 | 1.00 | 0.310 | 0.375 | 1.000 | 24408 | tags=82%, list=41%, signal=139% |
| 6428 | VESICLE\_COAT(GO:0030120) |  | 52 | 0.56 | 1.00 | 0.307 | 0.375 | 1.000 | 26061 | tags=96%, list=44%, signal=172% |
| 6429 | ANAPHASE\_PROMOTING\_COMPLEX\_DEPENDENT\_CATABOLIC\_PROCESS(GO:0031145) |  | 81 | 0.56 | 1.00 | 0.310 | 0.375 | 1.000 | 24347 | tags=83%, list=41%, signal=140% |
| 6430 | PHOSPHATIDYLINOSITOL\_BISPHOSPHATE\_PHOSPHATASE\_ACTIVITY(GO:0034593) |  | 27 | 0.57 | 1.00 | 0.141 | 0.375 | 1.000 | 21712 | tags=74%, list=37%, signal=117% |
| 6431 | GERM\_PLASM(GO:0060293) |  | 16 | 0.62 | 1.00 | 0.141 | 0.375 | 1.000 | 18259 | tags=56%, list=31%, signal=81% |
| 6432 | POLE\_PLASM(GO:0045495) |  | 16 | 0.62 | 1.00 | 0.141 | 0.375 | 1.000 | 18259 | tags=56%, list=31%, signal=81% |
| 6433 | P\_GRANULE(GO:0043186) |  | 16 | 0.62 | 1.00 | 0.141 | 0.375 | 1.000 | 18259 | tags=56%, list=31%, signal=81% |
| 6434 | REGULATION\_OF\_CHOLESTEROL\_BIOSYNTHETIC\_PROCESS(GO:0045540) |  | 42 | 0.57 | 1.00 | 0.476 | 0.375 | 1.000 | 23289 | tags=76%, list=39%, signal=126% |
| 6435 | LONG\_TERM\_SYNAPTIC\_DEPRESSION(GO:0060292) |  | 28 | 0.61 | 1.00 | 0.141 | 0.375 | 1.000 | 22480 | tags=71%, list=38%, signal=115% |
| 6436 | EXTRACELLULAR\_EXOSOME\_BIOGENESIS(GO:0097734) |  | 20 | 0.57 | 1.00 | 0.307 | 0.375 | 1.000 | 24218 | tags=80%, list=41%, signal=135% |
| 6437 | COATED\_VESICLE\_MEMBRANE(GO:0030662) |  | 163 | 0.57 | 1.00 | 0.141 | 0.375 | 1.000 | 22176 | tags=59%, list=38%, signal=94% |
| 6438 | HEME\_BIOSYNTHETIC\_PROCESS(GO:0006783) |  | 22 | 0.57 | 1.00 | 0.476 | 0.375 | 1.000 | 21949 | tags=73%, list=37%, signal=116% |
| 6439 | CELLULAR\_IRON\_ION\_HOMEOSTASIS(GO:0006879) |  | 64 | 0.57 | 1.00 | 0.141 | 0.375 | 1.000 | 23624 | tags=69%, list=40%, signal=114% |
| 6440 | OOCYTE\_MATURATION(GO:0001556) |  | 25 | 0.57 | 1.00 | 0.141 | 0.375 | 1.000 | 23969 | tags=76%, list=41%, signal=128% |
| 6441 | CELLULAR\_RESPONSE\_TO\_INTERFERON\_BETA(GO:0035458) |  | 15 | 0.58 | 1.00 | 0.141 | 0.375 | 1.000 | 23606 | tags=67%, list=40%, signal=111% |
| 6442 | VITAMIN\_B6\_BINDING(GO:0070279) |  | 56 | 0.60 | 1.00 | 0.141 | 0.375 | 1.000 | 21025 | tags=70%, list=36%, signal=108% |
| 6443 | HISTONE\_DEACETYLASE\_COMPLEX(GO:0000118) |  | 60 | 0.56 | 1.00 | 0.141 | 0.376 | 1.000 | 19631 | tags=62%, list=33%, signal=92% |
| 6444 | MACROMOLECULE\_TRANSMEMBRANE\_TRANSPORTER\_ACTIVITY(GO:0022884) |  | 20 | 0.59 | 1.00 | 0.141 | 0.376 | 1.000 | 22167 | tags=70%, list=37%, signal=112% |
| 6445 | SIGNAL\_TRANSDUCTION\_INVOLVED\_IN\_CELL\_CYCLE\_CHECKPOINT(GO:0072395) |  | 79 | 0.61 | 1.00 | 0.141 | 0.376 | 1.000 | 21594 | tags=76%, list=37%, signal=119% |
| 6446 | PYRIMIDINE\_NUCLEOSIDE\_TRIPHOSPHATE\_BIOSYNTHETIC\_PROCESS(GO:0009148) |  | 20 | 0.57 | 1.00 | 0.310 | 0.376 | 1.000 | 20715 | tags=65%, list=35%, signal=100% |
| 6447 | DICARBOXYLIC\_ACID\_CATABOLIC\_PROCESS(GO:0043649) |  | 18 | 0.62 | 1.00 | 0.307 | 0.376 | 1.000 | 16130 | tags=39%, list=27%, signal=53% |
| 6448 | CLATHRIN\_COATED\_ENDOCYTIC\_VESICLE(GO:0045334) |  | 58 | 0.57 | 1.00 | 0.141 | 0.376 | 1.000 | 12547 | tags=26%, list=21%, signal=33% |
| 6449 | NUCLEOSOME\_ORGANIZATION(GO:0034728) |  | 174 | 0.59 | 1.00 | 0.141 | 0.376 | 1.000 | 20669 | tags=48%, list=35%, signal=73% |
| 6450 | GONADOTROPIN\_SECRETION(GO:0032274) |  | 16 | 0.61 | 1.00 | 0.310 | 0.376 | 1.000 | 16468 | tags=38%, list=28%, signal=52% |
| 6451 | CELLULAR\_ZINC\_ION\_HOMEOSTASIS(GO:0006882) |  | 26 | 0.59 | 1.00 | 0.141 | 0.376 | 1.000 | 21009 | tags=65%, list=36%, signal=101% |
| 6452 | DIOL\_METABOLIC\_PROCESS(GO:0034311) |  | 18 | 0.57 | 1.00 | 0.141 | 0.376 | 1.000 | 24712 | tags=78%, list=42%, signal=134% |
| 6453 | PYRIDOXAL\_PHOSPHATE\_BINDING(GO:0030170) |  | 55 | 0.60 | 1.00 | 0.141 | 0.376 | 1.000 | 21025 | tags=69%, list=36%, signal=107% |
| 6454 | PROTEIN\_PALMITOYLATION(GO:0018345) |  | 28 | 0.57 | 1.00 | 0.141 | 0.376 | 1.000 | 20368 | tags=64%, list=34%, signal=98% |
| 6455 | REGULATION\_OF\_GLUCONEOGENESIS(GO:0006111) |  | 34 | 0.59 | 1.00 | 0.141 | 0.376 | 1.000 | 19589 | tags=50%, list=33%, signal=75% |
| 6456 | NEGATIVE\_REGULATION\_OF\_SIGNALING\_RECEPTOR\_ACTIVITY(GO:2000272) |  | 30 | 0.57 | 1.00 | 0.141 | 0.376 | 1.000 | 23828 | tags=70%, list=40%, signal=117% |
| 6457 | UBIQUITIN\_DEPENDENT\_PROTEIN\_CATABOLIC\_PROCESS\_VIA\_THE\_MULTIVESICULAR\_BODY\_SORTING\_PATHWAY(GO:0043162) |  | 19 | 0.57 | 1.00 | 0.307 | 0.376 | 1.000 | 24304 | tags=95%, list=41%, signal=161% |
| 6458 | SYNAPTONEMAL\_COMPLEX(GO:0000795) |  | 42 | 0.60 | 1.00 | 0.141 | 0.377 | 1.000 | 22865 | tags=64%, list=39%, signal=105% |
| 6459 | SYNAPTONEMAL\_STRUCTURE(GO:0099086) |  | 42 | 0.60 | 1.00 | 0.141 | 0.376 | 1.000 | 22865 | tags=64%, list=39%, signal=105% |
| 6460 | CITRATE\_METABOLIC\_PROCESS(GO:0006101) |  | 36 | 0.58 | 1.00 | 0.476 | 0.377 | 1.000 | 21329 | tags=72%, list=36%, signal=113% |
| 6461 | NEGATIVE\_REGULATION\_OF\_INTRINSIC\_APOPTOTIC\_SIGNALING\_PATHWAY\_IN\_RESPONSE\_TO\_DNA\_DAMAGE(GO:1902230) |  | 29 | 0.60 | 1.00 | 0.141 | 0.377 | 1.000 | 19800 | tags=45%, list=33%, signal=67% |
| 6462 | TRANSLATION\_FACTOR\_ACTIVITY\_\_RNA\_BINDING(GO:0008135) |  | 100 | 0.59 | 1.00 | 0.141 | 0.377 | 1.000 | 21819 | tags=78%, list=37%, signal=123% |
| 6463 | VESICLE\_DOCKING\_INVOLVED\_IN\_EXOCYTOSIS(GO:0006904) |  | 40 | 0.58 | 1.00 | 0.141 | 0.377 | 1.000 | 21162 | tags=63%, list=36%, signal=97% |
| 6464 | EXTRACELLULAR\_VESICLE\_BIOGENESIS(GO:0140112) |  | 22 | 0.57 | 1.00 | 0.307 | 0.377 | 1.000 | 24218 | tags=77%, list=41%, signal=131% |
| 6465 | DETECTION\_OF\_CHEMICAL\_STIMULUS\_INVOLVED\_IN\_SENSORY\_PERCEPTION\_OF\_SMELL(GO:0050911) |  | 389 | 0.50 | 1.00 | 0.481 | 0.378 | 1.000 | 13514 | tags=6%, list=23%, signal=8% |
| 6466 | NEGATIVE\_REGULATION\_OF\_TYPE\_I\_INTERFERON\_PRODUCTION(GO:0032480) |  | 48 | 0.58 | 1.00 | 0.141 | 0.378 | 1.000 | 23234 | tags=67%, list=39%, signal=110% |
| 6467 | ACYL\_COA\_DEHYDROGENASE\_ACTIVITY(GO:0003995) |  | 16 | 0.60 | 1.00 | 0.476 | 0.378 | 1.000 | 11203 | tags=25%, list=19%, signal=31% |
| 6468 | NEGATIVE\_REGULATION\_OF\_PROTEASOMAL\_UBIQUITIN\_DEPENDENT\_PROTEIN\_CATABOLIC\_PROCESS(GO:0032435) |  | 30 | 0.59 | 1.00 | 0.141 | 0.378 | 1.000 | 21505 | tags=77%, list=36%, signal=120% |
| 6469 | HISTONE\_H3\_K27\_METHYLATION(GO:0070734) |  | 19 | 0.61 | 1.00 | 0.307 | 0.378 | 1.000 | 19110 | tags=58%, list=32%, signal=86% |
| 6470 | REGULATION\_OF\_PROTEIN\_AUTOPHOSPHORYLATION(GO:0031952) |  | 42 | 0.58 | 1.00 | 0.141 | 0.378 | 1.000 | 11589 | tags=21%, list=20%, signal=27% |
| 6471 | HISTONE\_H3\_K4\_METHYLATION(GO:0051568) |  | 52 | 0.58 | 1.00 | 0.141 | 0.378 | 1.000 | 23231 | tags=77%, list=39%, signal=127% |
| 6472 | OLFACTORY\_RECEPTOR\_ACTIVITY(GO:0004984) |  | 396 | 0.49 | 1.00 | 0.481 | 0.378 | 1.000 | 13514 | tags=6%, list=23%, signal=8% |
| 6473 | ESTABLISHMENT\_OF\_SKIN\_BARRIER(GO:0061436) |  | 19 | 0.62 | 1.00 | 0.310 | 0.378 | 1.000 | 20077 | tags=37%, list=34%, signal=56% |
| 6474 | REGULATION\_OF\_KERATINOCYTE\_DIFFERENTIATION(GO:0045616) |  | 37 | 0.62 | 1.00 | 0.141 | 0.378 | 1.000 | 17118 | tags=35%, list=29%, signal=49% |
| 6475 | MATURATION\_OF\_5.8S\_RRNA\_FROM\_TRICISTRONIC\_RRNA\_TRANSCRIPT\_(SSU\_RRNA\_\_5.8S\_RRNA\_\_LSU\_RRNA)(GO:0000466) |  | 20 | 0.58 | 1.00 | 0.476 | 0.378 | 1.000 | 17908 | tags=65%, list=30%, signal=93% |
| 6476 | INTRINSIC\_APOPTOTIC\_SIGNALING\_PATHWAY\_IN\_RESPONSE\_TO\_ENDOPLASMIC\_RETICULUM\_STRESS(GO:0070059) |  | 59 | 0.56 | 1.00 | 0.476 | 0.378 | 1.000 | 22540 | tags=66%, list=38%, signal=107% |
| 6477 | OXIDOREDUCTASE\_ACTIVITY\_\_ACTING\_ON\_THE\_CH\_CH\_GROUP\_OF\_DONORS\_\_NAD\_OR\_NADP\_AS\_ACCEPTOR(GO:0016628) |  | 26 | 0.60 | 1.00 | 0.141 | 0.378 | 1.000 | 20971 | tags=50%, list=35%, signal=77% |
| 6478 | IRON\_SULFUR\_CLUSTER\_ASSEMBLY(GO:0016226) |  | 21 | 0.56 | 1.00 | 0.307 | 0.378 | 1.000 | 26235 | tags=95%, list=44%, signal=171% |
| 6479 | METALLO\_SULFUR\_CLUSTER\_ASSEMBLY(GO:0031163) |  | 21 | 0.56 | 1.00 | 0.307 | 0.378 | 1.000 | 26235 | tags=95%, list=44%, signal=171% |
| 6480 | NEGATIVE\_REGULATION\_OF\_OXIDATIVE\_STRESS\_INDUCED\_NEURON\_DEATH(GO:1903204) |  | 18 | 0.60 | 1.00 | 0.309 | 0.379 | 1.000 | 19859 | tags=72%, list=34%, signal=109% |
| 6481 | POSITIVE\_REGULATION\_OF\_DNA\_TEMPLATED\_TRANSCRIPTION\_\_ELONGATION(GO:0032786) |  | 23 | 0.56 | 1.00 | 0.141 | 0.379 | 1.000 | 24632 | tags=91%, list=42%, signal=156% |
| 6482 | CELL\_VOLUME\_HOMEOSTASIS(GO:0006884) |  | 18 | 0.60 | 1.00 | 0.141 | 0.379 | 1.000 | 6374 | tags=6%, list=11%, signal=6% |
| 6483 | POSITIVE\_REGULATION\_OF\_VIRAL\_PROCESS(GO:0048524) |  | 102 | 0.58 | 1.00 | 0.141 | 0.379 | 1.000 | 22407 | tags=72%, list=38%, signal=115% |
| 6484 | REGULATION\_OF\_GLYCOPROTEIN\_BIOSYNTHETIC\_PROCESS(GO:0010559) |  | 36 | 0.58 | 1.00 | 0.141 | 0.379 | 1.000 | 22307 | tags=67%, list=38%, signal=107% |
| 6485 | TRICARBOXYLIC\_ACID\_METABOLIC\_PROCESS(GO:0072350) |  | 40 | 0.57 | 1.00 | 0.310 | 0.379 | 1.000 | 21329 | tags=70%, list=36%, signal=109% |
| 6486 | SAGA\_COMPLEX(GO:0000124) |  | 15 | 0.57 | 1.00 | 0.307 | 0.380 | 1.000 | 25063 | tags=80%, list=42%, signal=139% |
| 6487 | MITOPHAGY(GO:0000423) |  | 20 | 0.58 | 1.00 | 0.141 | 0.380 | 1.000 | 23032 | tags=90%, list=39%, signal=147% |
| 6488 | ENDOPLASMIC\_RETICULUM\_GOLGI\_INTERMEDIATE\_COMPARTMENT(GO:0005793) |  | 112 | 0.55 | 1.00 | 0.141 | 0.380 | 1.000 | 24013 | tags=71%, list=41%, signal=120% |
| 6489 | MACROMOLECULE\_METHYLATION(GO:0043414) |  | 290 | 0.59 | 1.00 | 0.141 | 0.380 | 1.000 | 19804 | tags=61%, list=33%, signal=91% |
| 6490 | PLATELET\_FORMATION(GO:0030220) |  | 20 | 0.61 | 1.00 | 0.310 | 0.380 | 1.000 | 15485 | tags=35%, list=26%, signal=47% |
| 6491 | UBIQUITIN\_SPECIFIC\_PROTEASE\_BINDING(GO:1990381) |  | 16 | 0.58 | 1.00 | 0.141 | 0.380 | 1.000 | 24656 | tags=100%, list=42%, signal=171% |
| 6492 | FLEMMING\_BODY(GO:0090543) |  | 27 | 0.56 | 1.00 | 0.313 | 0.380 | 1.000 | 25747 | tags=96%, list=44%, signal=170% |
| 6493 | POSITIVE\_REGULATION\_OF\_DOUBLE\_STRAND\_BREAK\_REPAIR(GO:2000781) |  | 15 | 0.62 | 1.00 | 0.141 | 0.380 | 1.000 | 19736 | tags=67%, list=33%, signal=100% |
| 6494 | PORPHYRIN\_CONTAINING\_COMPOUND\_BIOSYNTHETIC\_PROCESS(GO:0006779) |  | 26 | 0.57 | 1.00 | 0.476 | 0.380 | 1.000 | 21949 | tags=69%, list=37%, signal=110% |
| 6495 | OOCYTE\_DEVELOPMENT(GO:0048599) |  | 44 | 0.59 | 1.00 | 0.141 | 0.381 | 1.000 | 20968 | tags=61%, list=35%, signal=95% |
| 6496 | ENDOPLASMIC\_RETICULUM\_MANNOSE\_TRIMMING(GO:1904380) |  | 20 | 0.56 | 1.00 | 0.307 | 0.381 | 1.000 | 26213 | tags=100%, list=44%, signal=180% |
| 6497 | ACTIN\_FILAMENT\_CAPPING(GO:0051693) |  | 36 | 0.56 | 1.00 | 0.141 | 0.381 | 1.000 | 24458 | tags=75%, list=41%, signal=128% |
| 6498 | REGULATION\_OF\_CHOLESTEROL\_METABOLIC\_PROCESS(GO:0090181) |  | 53 | 0.56 | 1.00 | 0.476 | 0.381 | 1.000 | 23397 | tags=72%, list=40%, signal=119% |
| 6499 | 2\_IRON\_\_2\_SULFUR\_CLUSTER\_BINDING(GO:0051537) |  | 24 | 0.59 | 1.00 | 0.141 | 0.381 | 1.000 | 23219 | tags=79%, list=39%, signal=130% |
| 6500 | SKELETAL\_MUSCLE\_ADAPTATION(GO:0043501) |  | 25 | 0.59 | 1.00 | 0.141 | 0.381 | 1.000 | 23500 | tags=60%, list=40%, signal=100% |
| 6501 | NON\_MEMBRANE\_SPANNING\_PROTEIN\_TYROSINE\_KINASE\_ACTIVITY(GO:0004715) |  | 51 | 0.57 | 1.00 | 0.141 | 0.381 | 1.000 | 24370 | tags=75%, list=41%, signal=127% |
| 6502 | REGULATION\_OF\_DNA\_TEMPLATED\_TRANSCRIPTION\_\_ELONGATION(GO:0032784) |  | 43 | 0.56 | 1.00 | 0.141 | 0.381 | 1.000 | 24632 | tags=86%, list=42%, signal=147% |
| 6503 | CTP\_BIOSYNTHETIC\_PROCESS(GO:0006241) |  | 15 | 0.56 | 1.00 | 0.310 | 0.381 | 1.000 | 25880 | tags=87%, list=44%, signal=154% |
| 6504 | CTP\_METABOLIC\_PROCESS(GO:0046036) |  | 15 | 0.56 | 1.00 | 0.310 | 0.381 | 1.000 | 25880 | tags=87%, list=44%, signal=154% |
| 6505 | REGULATION\_OF\_PROTEIN\_LOCALIZATION\_TO\_PLASMA\_MEMBRANE(GO:1903076) |  | 88 | 0.59 | 1.00 | 0.141 | 0.381 | 1.000 | 20027 | tags=51%, list=34%, signal=77% |
| 6506 | MAINTENANCE\_OF\_CELL\_POLARITY(GO:0030011) |  | 18 | 0.57 | 0.99 | 0.307 | 0.381 | 1.000 | 19638 | tags=61%, list=33%, signal=91% |
| 6507 | ENDOSOME\_TRANSPORT\_VIA\_MULTIVESICULAR\_BODY\_SORTING\_PATHWAY(GO:0032509) |  | 20 | 0.56 | 0.99 | 0.476 | 0.381 | 1.000 | 26119 | tags=95%, list=44%, signal=170% |
| 6508 | TETHERING\_COMPLEX(GO:0099023) |  | 76 | 0.55 | 0.99 | 0.476 | 0.382 | 1.000 | 22957 | tags=67%, list=39%, signal=110% |
| 6509 | NEGATIVE\_REGULATION\_OF\_G1\_S\_TRANSITION\_OF\_MITOTIC\_CELL\_CYCLE(GO:2000134) |  | 115 | 0.59 | 0.99 | 0.141 | 0.382 | 1.000 | 20085 | tags=53%, list=34%, signal=80% |
| 6510 | REGULATION\_OF\_MONOCYTE\_CHEMOTAXIS(GO:0090025) |  | 22 | 0.60 | 0.99 | 0.141 | 0.382 | 1.000 | 13828 | tags=23%, list=23%, signal=30% |
| 6511 | POLYSOMAL\_RIBOSOME(GO:0042788) |  | 29 | 0.56 | 0.99 | 0.476 | 0.382 | 1.000 | 25293 | tags=93%, list=43%, signal=163% |
| 6512 | ESCRT\_COMPLEX(GO:0036452) |  | 29 | 0.55 | 0.99 | 0.141 | 0.382 | 1.000 | 26168 | tags=97%, list=44%, signal=173% |
| 6513 | NECROPTOTIC\_PROCESS(GO:0070266) |  | 36 | 0.56 | 0.99 | 0.141 | 0.382 | 1.000 | 23284 | tags=64%, list=39%, signal=105% |
| 6514 | RESPONSE\_TO\_CORTICOSTERONE(GO:0051412) |  | 18 | 0.63 | 0.99 | 0.310 | 0.382 | 1.000 | 19286 | tags=50%, list=33%, signal=74% |
| 6515 | NUCLEAR\_EXPORT(GO:0051168) |  | 198 | 0.60 | 0.99 | 0.141 | 0.382 | 1.000 | 22062 | tags=82%, list=37%, signal=131% |
| 6516 | PROTEIN\_DEMANNOSYLATION(GO:0036507) |  | 24 | 0.56 | 0.99 | 0.141 | 0.382 | 1.000 | 20880 | tags=75%, list=35%, signal=116% |
| 6517 | PROTEIN\_ALPHA\_1\_2\_DEMANNOSYLATION(GO:0036508) |  | 24 | 0.56 | 0.99 | 0.141 | 0.382 | 1.000 | 20880 | tags=75%, list=35%, signal=116% |
| 6518 | FC\_RECEPTOR\_SIGNALING\_PATHWAY(GO:0038093) |  | 243 | 0.55 | 0.99 | 0.141 | 0.382 | 1.000 | 25353 | tags=59%, list=43%, signal=103% |
| 6519 | REGULATION\_OF\_AUTOPHAGY\_OF\_MITOCHONDRION(GO:1903146) |  | 47 | 0.56 | 0.99 | 0.476 | 0.382 | 1.000 | 24344 | tags=87%, list=41%, signal=148% |
| 6520 | REGULATION\_OF\_TELOMERASE\_ACTIVITY(GO:0051972) |  | 46 | 0.61 | 0.99 | 0.141 | 0.383 | 1.000 | 20903 | tags=78%, list=35%, signal=121% |
| 6521 | TRANSFERASE\_ACTIVITY\_\_TRANSFERRING\_ONE\_CARBON\_GROUPS(GO:0016741) |  | 228 | 0.59 | 0.99 | 0.141 | 0.383 | 1.000 | 21136 | tags=72%, list=36%, signal=112% |
| 6522 | NEGATIVE\_REGULATION\_OF\_BLOOD\_VESSEL\_ENDOTHELIAL\_CELL\_MIGRATION(GO:0043537) |  | 75 | 0.60 | 0.99 | 0.476 | 0.383 | 1.000 | 20361 | tags=35%, list=34%, signal=53% |
| 6523 | STEROID\_DEHYDROGENASE\_ACTIVITY(GO:0016229) |  | 29 | 0.60 | 0.99 | 0.141 | 0.383 | 1.000 | 19946 | tags=48%, list=34%, signal=73% |
| 6524 | INTEGRAL\_COMPONENT\_OF\_NUCLEAR\_INNER\_MEMBRANE(GO:0005639) |  | 17 | 0.59 | 0.99 | 0.307 | 0.383 | 1.000 | 23437 | tags=82%, list=40%, signal=136% |
| 6525 | INTRINSIC\_COMPONENT\_OF\_NUCLEAR\_INNER\_MEMBRANE(GO:0031229) |  | 17 | 0.59 | 0.99 | 0.307 | 0.383 | 1.000 | 23437 | tags=82%, list=40%, signal=136% |
| 6526 | CARTILAGE\_CONDENSATION(GO:0001502) |  | 20 | 0.61 | 0.99 | 0.141 | 0.383 | 1.000 | 20024 | tags=55%, list=34%, signal=83% |
| 6527 | MRNA\_CLEAVAGE\_INVOLVED\_IN\_MRNA\_PROCESSING(GO:0098787) |  | 15 | 0.58 | 0.99 | 0.307 | 0.383 | 1.000 | 24992 | tags=87%, list=42%, signal=150% |
| 6528 | POSITIVE\_REGULATION\_OF\_TRANSCRIPTION\_OF\_NOTCH\_RECEPTOR\_TARGET(GO:0007221) |  | 18 | 0.56 | 0.99 | 0.476 | 0.383 | 1.000 | 24170 | tags=72%, list=41%, signal=122% |
| 6529 | DNA\_REPAIR\_COMPLEX(GO:1990391) |  | 35 | 0.62 | 0.99 | 0.141 | 0.383 | 1.000 | 18733 | tags=71%, list=32%, signal=104% |
| 6530 | REGULATION\_OF\_PROTEIN\_EXPORT\_FROM\_NUCLEUS(GO:0046825) |  | 40 | 0.61 | 0.99 | 0.141 | 0.384 | 1.000 | 22194 | tags=85%, list=38%, signal=136% |
| 6531 | MITOCHONDRIAL\_RESPIRATORY\_CHAIN\_COMPLEX\_IV\_ASSEMBLY(GO:0033617) |  | 20 | 0.57 | 0.99 | 0.475 | 0.384 | 1.000 | 18359 | tags=65%, list=31%, signal=94% |
| 6532 | TRANSCRIPTIONAL\_REPRESSOR\_ACTIVITY\_\_RNA\_POLYMERASE\_II\_TRANSCRIPTION\_FACTOR\_BINDING(GO:0001191) |  | 41 | 0.61 | 0.99 | 0.141 | 0.384 | 1.000 | 19184 | tags=56%, list=32%, signal=83% |
| 6533 | NEGATIVE\_REGULATION\_OF\_CELL\_CYCLE\_G1\_S\_PHASE\_TRANSITION(GO:1902807) |  | 122 | 0.59 | 0.99 | 0.141 | 0.384 | 1.000 | 19834 | tags=50%, list=34%, signal=75% |
| 6534 | CELLULAR\_TRANSITION\_METAL\_ION\_HOMEOSTASIS(GO:0046916) |  | 103 | 0.56 | 0.99 | 0.141 | 0.384 | 1.000 | 23746 | tags=69%, list=40%, signal=115% |
| 6535 | PROTEIN\_EXPORT\_FROM\_NUCLEUS(GO:0006611) |  | 179 | 0.60 | 0.99 | 0.141 | 0.384 | 1.000 | 22062 | tags=83%, list=37%, signal=131% |
| 6536 | ACROSOMAL\_MEMBRANE(GO:0002080) |  | 23 | 0.59 | 0.99 | 0.476 | 0.384 | 1.000 | 15440 | tags=26%, list=26%, signal=35% |
| 6537 | SIGNAL\_TRANSDUCTION\_INVOLVED\_IN\_MITOTIC\_DNA\_INTEGRITY\_CHECKPOINT(GO:1902403) |  | 65 | 0.60 | 0.99 | 0.141 | 0.385 | 1.000 | 21594 | tags=72%, list=37%, signal=114% |
| 6538 | SIGNAL\_TRANSDUCTION\_INVOLVED\_IN\_MITOTIC\_DNA\_DAMAGE\_CHECKPOINT(GO:1902402) |  | 65 | 0.60 | 0.99 | 0.141 | 0.385 | 1.000 | 21594 | tags=72%, list=37%, signal=114% |
| 6539 | SIGNAL\_TRANSDUCTION\_INVOLVED\_IN\_MITOTIC\_CELL\_CYCLE\_CHECKPOINT(GO:0072413) |  | 65 | 0.60 | 0.99 | 0.141 | 0.385 | 1.000 | 21594 | tags=72%, list=37%, signal=114% |
| 6540 | EXOSOMAL\_SECRETION(GO:1990182) |  | 19 | 0.57 | 0.99 | 0.476 | 0.385 | 1.000 | 24218 | tags=79%, list=41%, signal=134% |
| 6541 | MEMBRANE\_PROTEIN\_ECTODOMAIN\_PROTEOLYSIS(GO:0006509) |  | 43 | 0.55 | 0.99 | 0.307 | 0.385 | 1.000 | 20361 | tags=56%, list=34%, signal=85% |
| 6542 | CIRCADIAN\_REGULATION\_OF\_GENE\_EXPRESSION(GO:0032922) |  | 55 | 0.56 | 0.99 | 0.307 | 0.385 | 1.000 | 20690 | tags=56%, list=35%, signal=87% |
| 6543 | SIGNAL\_TRANSDUCTION\_INVOLVED\_IN\_MITOTIC\_G1\_DNA\_DAMAGE\_CHECKPOINT(GO:0072431) |  | 63 | 0.60 | 0.99 | 0.141 | 0.385 | 1.000 | 21594 | tags=73%, list=37%, signal=115% |
| 6544 | INTRACELLULAR\_SIGNAL\_TRANSDUCTION\_INVOLVED\_IN\_G1\_DNA\_DAMAGE\_CHECKPOINT(GO:1902400) |  | 63 | 0.60 | 0.99 | 0.141 | 0.385 | 1.000 | 21594 | tags=73%, list=37%, signal=115% |
| 6545 | REGULATION\_OF\_HISTONE\_H3\_K9\_METHYLATION(GO:0051570) |  | 22 | 0.56 | 0.99 | 0.313 | 0.385 | 1.000 | 24413 | tags=73%, list=41%, signal=124% |
| 6546 | PYRIMIDINE\_RIBONUCLEOSIDE\_TRIPHOSPHATE\_BIOSYNTHETIC\_PROCESS(GO:0009209) |  | 16 | 0.56 | 0.99 | 0.476 | 0.385 | 1.000 | 23753 | tags=75%, list=40%, signal=125% |
| 6547 | NEGATIVE\_REGULATION\_OF\_MYELOID\_CELL\_APOPTOTIC\_PROCESS(GO:0033033) |  | 16 | 0.58 | 0.99 | 0.307 | 0.385 | 1.000 | 23563 | tags=56%, list=40%, signal=93% |
| 6548 | METAPHASE\_PLATE\_CONGRESSION(GO:0051310) |  | 57 | 0.59 | 0.99 | 0.141 | 0.385 | 1.000 | 21353 | tags=77%, list=36%, signal=121% |
| 6549 | PEPTIDYL\_ARGININE\_METHYLATION(GO:0018216) |  | 16 | 0.57 | 0.99 | 0.476 | 0.385 | 1.000 | 21457 | tags=63%, list=36%, signal=98% |
| 6550 | RETINOL\_BINDING(GO:0019841) |  | 15 | 0.64 | 0.99 | 0.476 | 0.386 | 1.000 | 16098 | tags=47%, list=27%, signal=64% |
| 6551 | B\_CELL\_RECEPTOR\_SIGNALING\_PATHWAY(GO:0050853) |  | 101 | 0.55 | 0.99 | 0.141 | 0.386 | 1.000 | 17501 | tags=20%, list=30%, signal=28% |
| 6552 | DNA\_DAMAGE\_RESPONSE\_\_SIGNAL\_TRANSDUCTION\_BY\_P53\_CLASS\_MEDIATOR\_RESULTING\_IN\_CELL\_CYCLE\_ARREST(GO:0006977) |  | 62 | 0.60 | 0.99 | 0.141 | 0.386 | 1.000 | 21594 | tags=73%, list=37%, signal=114% |
| 6553 | NEGATIVE\_REGULATION\_OF\_CELL\_JUNCTION\_ASSEMBLY(GO:1901889) |  | 20 | 0.61 | 0.99 | 0.141 | 0.386 | 1.000 | 21149 | tags=70%, list=36%, signal=109% |
| 6554 | 1\_PHOSPHATIDYLINOSITOL\_BINDING(GO:0005545) |  | 15 | 0.62 | 0.99 | 0.309 | 0.386 | 1.000 | 14915 | tags=47%, list=25%, signal=62% |
| 6555 | PEPTIDYL\_SERINE\_DEPHOSPHORYLATION(GO:0070262) |  | 16 | 0.57 | 0.99 | 0.476 | 0.386 | 1.000 | 23801 | tags=81%, list=40%, signal=136% |
| 6556 | COPI\_COATED\_VESICLE\_MEMBRANE(GO:0030663) |  | 16 | 0.55 | 0.99 | 0.141 | 0.386 | 1.000 | 26013 | tags=94%, list=44%, signal=167% |
| 6557 | SEQUESTERING\_OF\_METAL\_ION(GO:0051238) |  | 18 | 0.61 | 0.99 | 0.141 | 0.386 | 1.000 | 22774 | tags=72%, list=39%, signal=117% |
| 6558 | MITOTIC\_G1\_DNA\_DAMAGE\_CHECKPOINT(GO:0031571) |  | 71 | 0.60 | 0.99 | 0.141 | 0.387 | 1.000 | 20085 | tags=63%, list=34%, signal=96% |
| 6559 | MITOTIC\_G1\_S\_TRANSITION\_CHECKPOINT(GO:0044819) |  | 71 | 0.60 | 0.99 | 0.141 | 0.386 | 1.000 | 20085 | tags=63%, list=34%, signal=96% |
| 6560 | REGULATION\_OF\_PROTEIN\_DEACETYLATION(GO:0090311) |  | 39 | 0.58 | 0.99 | 0.476 | 0.386 | 1.000 | 22480 | tags=69%, list=38%, signal=112% |
| 6561 | REGULATION\_OF\_ENDOPLASMIC\_RETICULUM\_STRESS\_INDUCED\_INTRINSIC\_APOPTOTIC\_SIGNALING\_PATHWAY(GO:1902235) |  | 32 | 0.57 | 0.99 | 0.310 | 0.386 | 1.000 | 22540 | tags=66%, list=38%, signal=106% |
| 6562 | PLASMA\_MEMBRANE\_FUSION(GO:0045026) |  | 39 | 0.58 | 0.99 | 0.307 | 0.387 | 1.000 | 19755 | tags=44%, list=33%, signal=65% |
| 6563 | LYMPHOID\_PROGENITOR\_CELL\_DIFFERENTIATION(GO:0002320) |  | 19 | 0.57 | 0.99 | 0.141 | 0.387 | 1.000 | 21256 | tags=53%, list=36%, signal=82% |
| 6564 | POSITIVE\_REGULATION\_OF\_RELEASE\_OF\_CYTOCHROME\_C\_FROM\_MITOCHONDRIA(GO:0090200) |  | 29 | 0.59 | 0.99 | 0.476 | 0.387 | 1.000 | 19716 | tags=52%, list=33%, signal=78% |
| 6565 | POSITIVE\_REGULATION\_OF\_PROTEIN\_LOCALIZATION\_TO\_MEMBRANE(GO:1905477) |  | 120 | 0.59 | 0.99 | 0.141 | 0.387 | 1.000 | 22440 | tags=63%, list=38%, signal=102% |
| 6566 | PROTON\_TRANSPORTING\_TWO\_SECTOR\_ATPASE\_COMPLEX\_ASSEMBLY(GO:0070071) |  | 15 | 0.62 | 0.99 | 0.141 | 0.387 | 1.000 | 21646 | tags=87%, list=37%, signal=137% |
| 6567 | CLATHRIN\_VESICLE\_COAT(GO:0030125) |  | 26 | 0.55 | 0.99 | 0.307 | 0.387 | 1.000 | 26061 | tags=96%, list=44%, signal=172% |
| 6568 | NEGATIVE\_REGULATION\_OF\_STRIATED\_MUSCLE\_CELL\_APOPTOTIC\_PROCESS(GO:0010664) |  | 38 | 0.57 | 0.99 | 0.476 | 0.387 | 1.000 | 15300 | tags=21%, list=26%, signal=28% |
| 6569 | PROTEASE\_BINDING(GO:0002020) |  | 133 | 0.56 | 0.99 | 0.141 | 0.387 | 1.000 | 21868 | tags=47%, list=37%, signal=74% |
| 6570 | APICOLATERAL\_PLASMA\_MEMBRANE(GO:0016327) |  | 19 | 0.63 | 0.99 | 0.476 | 0.387 | 1.000 | 6670 | tags=5%, list=11%, signal=6% |
| 6571 | SNAP\_RECEPTOR\_ACTIVITY(GO:0005484) |  | 41 | 0.58 | 0.99 | 0.141 | 0.387 | 1.000 | 24007 | tags=80%, list=41%, signal=135% |
| 6572 | SYNAPSE\_MATURATION(GO:0060074) |  | 19 | 0.58 | 0.99 | 0.476 | 0.388 | 1.000 | 16153 | tags=53%, list=27%, signal=72% |
| 6573 | POSITIVE\_REGULATION\_OF\_MYELOID\_LEUKOCYTE\_MEDIATED\_IMMUNITY(GO:0002888) |  | 22 | 0.60 | 0.99 | 0.476 | 0.388 | 1.000 | 18255 | tags=45%, list=31%, signal=66% |
| 6574 | MELANOSOME\_LOCALIZATION(GO:0032400) |  | 26 | 0.60 | 0.99 | 0.141 | 0.388 | 1.000 | 21618 | tags=62%, list=37%, signal=97% |
| 6575 | AMYLOID\_BETA\_CLEARANCE(GO:0097242) |  | 19 | 0.61 | 0.99 | 0.476 | 0.388 | 1.000 | 21166 | tags=58%, list=36%, signal=90% |
| 6576 | CELLULAR\_RESPONSE\_TO\_AMMONIUM\_ION(GO:0071242) |  | 63 | 0.57 | 0.99 | 0.310 | 0.389 | 1.000 | 22992 | tags=51%, list=39%, signal=83% |
| 6577 | ENDOPLASMIC\_RETICULUM\_GOLGI\_INTERMEDIATE\_COMPARTMENT\_MEMBRANE(GO:0033116) |  | 67 | 0.55 | 0.99 | 0.141 | 0.389 | 1.000 | 24013 | tags=67%, list=41%, signal=113% |
| 6578 | REGULATION\_OF\_POSITIVE\_CHEMOTAXIS(GO:0050926) |  | 26 | 0.60 | 0.99 | 0.310 | 0.389 | 1.000 | 10000 | tags=23%, list=17%, signal=28% |
| 6579 | REGULATION\_OF\_TRANSCRIPTION\_ELONGATION\_FROM\_RNA\_POLYMERASE\_II\_PROMOTER(GO:0034243) |  | 27 | 0.57 | 0.99 | 0.141 | 0.389 | 1.000 | 24346 | tags=85%, list=41%, signal=145% |
| 6580 | EXOCYST(GO:0000145) |  | 20 | 0.57 | 0.99 | 0.307 | 0.389 | 1.000 | 15040 | tags=45%, list=25%, signal=60% |
| 6581 | G1\_DNA\_DAMAGE\_CHECKPOINT(GO:0044783) |  | 72 | 0.60 | 0.99 | 0.141 | 0.389 | 1.000 | 21594 | tags=71%, list=37%, signal=111% |
| 6582 | NEGATIVE\_REGULATION\_OF\_PROTEIN\_LOCALIZATION\_TO\_PLASMA\_MEMBRANE(GO:1903077) |  | 22 | 0.57 | 0.99 | 0.310 | 0.389 | 1.000 | 21805 | tags=68%, list=37%, signal=108% |
| 6583 | REGULATION\_OF\_HORMONE\_BIOSYNTHETIC\_PROCESS(GO:0046885) |  | 26 | 0.59 | 0.99 | 0.476 | 0.389 | 1.000 | 15381 | tags=31%, list=26%, signal=42% |
| 6584 | MRNA\_TRANSCRIPTION\_BY\_RNA\_POLYMERASE\_II(GO:0042789) |  | 19 | 0.57 | 0.99 | 0.476 | 0.390 | 1.000 | 23355 | tags=79%, list=39%, signal=130% |
| 6585 | IMMUNOLOGICAL\_SYNAPSE(GO:0001772) |  | 34 | 0.57 | 0.99 | 0.141 | 0.390 | 1.000 | 24253 | tags=62%, list=41%, signal=105% |
| 6586 | K63\_LINKED\_POLYUBIQUITIN\_MODIFICATION\_DEPENDENT\_PROTEIN\_BINDING(GO:0070530) |  | 21 | 0.56 | 0.99 | 0.476 | 0.390 | 1.000 | 20681 | tags=62%, list=35%, signal=95% |
| 6587 | TYPE\_I\_INTERFERON\_PRODUCTION(GO:0032606) |  | 125 | 0.57 | 0.99 | 0.141 | 0.390 | 1.000 | 24111 | tags=70%, list=41%, signal=119% |
| 6588 | REGULATION\_OF\_TYPE\_I\_INTERFERON\_PRODUCTION(GO:0032479) |  | 124 | 0.57 | 0.99 | 0.141 | 0.390 | 1.000 | 24111 | tags=70%, list=41%, signal=118% |
| 6589 | FC\_EPSILON\_RECEPTOR\_SIGNALING\_PATHWAY(GO:0038095) |  | 177 | 0.55 | 0.99 | 0.141 | 0.391 | 1.000 | 26052 | tags=59%, list=44%, signal=106% |
| 6590 | POSITIVE\_REGULATION\_OF\_GRANULOCYTE\_CHEMOTAXIS(GO:0071624) |  | 32 | 0.61 | 0.99 | 0.307 | 0.391 | 1.000 | 11759 | tags=31%, list=20%, signal=39% |
| 6591 | REGULATION\_OF\_DNA\_METHYLATION(GO:0044030) |  | 26 | 0.61 | 0.99 | 0.141 | 0.392 | 1.000 | 22734 | tags=58%, list=38%, signal=94% |
| 6592 | CYTOKINETIC\_PROCESS(GO:0032506) |  | 19 | 0.56 | 0.99 | 0.476 | 0.392 | 1.000 | 21315 | tags=74%, list=36%, signal=115% |
| 6593 | SH2\_DOMAIN\_BINDING(GO:0042169) |  | 33 | 0.58 | 0.99 | 0.307 | 0.392 | 1.000 | 18388 | tags=42%, list=31%, signal=62% |
| 6594 | REGULATION\_OF\_ACTIN\_NUCLEATION(GO:0051125) |  | 26 | 0.57 | 0.99 | 0.141 | 0.392 | 1.000 | 20257 | tags=38%, list=34%, signal=58% |
| 6595 | NUCLEAR\_MEMBRANE\_PART(GO:0044453) |  | 18 | 0.59 | 0.99 | 0.307 | 0.392 | 1.000 | 23437 | tags=78%, list=40%, signal=129% |
| 6596 | ENDOSOME\_ORGANIZATION(GO:0007032) |  | 75 | 0.54 | 0.99 | 0.141 | 0.392 | 1.000 | 26854 | tags=89%, list=45%, signal=163% |
| 6597 | NEGATIVE\_REGULATION\_OF\_CARDIAC\_MUSCLE\_CELL\_APOPTOTIC\_PROCESS(GO:0010667) |  | 34 | 0.57 | 0.99 | 0.476 | 0.392 | 1.000 | 23743 | tags=38%, list=40%, signal=64% |
| 6598 | POLYUBIQUITIN\_MODIFICATION\_DEPENDENT\_PROTEIN\_BINDING(GO:0031593) |  | 45 | 0.56 | 0.99 | 0.476 | 0.392 | 1.000 | 25644 | tags=91%, list=43%, signal=161% |
| 6599 | METHYLTRANSFERASE\_ACTIVITY(GO:0008168) |  | 217 | 0.59 | 0.99 | 0.141 | 0.392 | 1.000 | 21531 | tags=73%, list=36%, signal=114% |
| 6600 | HEXOSAMINIDASE\_ACTIVITY(GO:0015929) |  | 15 | 0.57 | 0.99 | 0.476 | 0.392 | 1.000 | 24482 | tags=80%, list=41%, signal=136% |
| 6601 | NEGATIVE\_REGULATION\_OF\_CELLULAR\_CARBOHYDRATE\_METABOLIC\_PROCESS(GO:0010677) |  | 36 | 0.59 | 0.99 | 0.141 | 0.392 | 1.000 | 23468 | tags=67%, list=40%, signal=110% |
| 6602 | COFACTOR\_TRANSPORT(GO:0051181) |  | 43 | 0.56 | 0.99 | 0.476 | 0.392 | 1.000 | 23490 | tags=63%, list=40%, signal=104% |
| 6603 | APOPTOTIC\_CELL\_CLEARANCE(GO:0043277) |  | 36 | 0.60 | 0.99 | 0.310 | 0.392 | 1.000 | 21292 | tags=56%, list=36%, signal=87% |
| 6604 | REGULATION\_OF\_HISTONE\_DEACETYLATION(GO:0031063) |  | 27 | 0.58 | 0.99 | 0.476 | 0.392 | 1.000 | 22117 | tags=67%, list=37%, signal=106% |
| 6605 | NEGATIVE\_REGULATION\_OF\_MUSCLE\_CELL\_APOPTOTIC\_PROCESS(GO:0010656) |  | 54 | 0.57 | 0.99 | 0.476 | 0.392 | 1.000 | 23743 | tags=37%, list=40%, signal=62% |
| 6606 | PORE\_COMPLEX(GO:0046930) |  | 18 | 0.57 | 0.99 | 0.476 | 0.392 | 1.000 | 25496 | tags=72%, list=43%, signal=127% |
| 6607 | REGULATION\_OF\_ACTIVIN\_RECEPTOR\_SIGNALING\_PATHWAY(GO:0032925) |  | 23 | 0.57 | 0.98 | 0.141 | 0.393 | 1.000 | 25125 | tags=87%, list=42%, signal=151% |
| 6608 | POSITIVE\_REGULATION\_OF\_ANTIGEN\_RECEPTOR\_MEDIATED\_SIGNALING\_PATHWAY(GO:0050857) |  | 20 | 0.62 | 0.98 | 0.476 | 0.393 | 1.000 | 13707 | tags=40%, list=23%, signal=52% |
| 6609 | POSITIVE\_REGULATION\_OF\_NEUROLOGICAL\_SYSTEM\_PROCESS(GO:0031646) |  | 24 | 0.58 | 0.98 | 0.476 | 0.393 | 1.000 | 24752 | tags=75%, list=42%, signal=129% |
| 6610 | PHOSPHATIDYLETHANOLAMINE\_BIOSYNTHETIC\_PROCESS(GO:0006646) |  | 15 | 0.58 | 0.98 | 0.307 | 0.393 | 1.000 | 16174 | tags=47%, list=27%, signal=64% |
| 6611 | POSITIVE\_REGULATION\_OF\_TYPE\_I\_INTERFERON\_PRODUCTION(GO:0032481) |  | 74 | 0.57 | 0.98 | 0.141 | 0.393 | 1.000 | 24269 | tags=73%, list=41%, signal=124% |
| 6612 | TRANSITION\_METAL\_ION\_TRANSMEMBRANE\_TRANSPORTER\_ACTIVITY(GO:0046915) |  | 41 | 0.56 | 0.98 | 0.476 | 0.393 | 1.000 | 24178 | tags=78%, list=41%, signal=132% |
| 6613 | HEME\_METABOLIC\_PROCESS(GO:0042168) |  | 31 | 0.57 | 0.98 | 0.307 | 0.393 | 1.000 | 21949 | tags=61%, list=37%, signal=97% |
| 6614 | POSITIVE\_REGULATION\_OF\_POSTTRANSCRIPTIONAL\_GENE\_SILENCING(GO:0060148) |  | 21 | 0.60 | 0.98 | 0.476 | 0.393 | 1.000 | 23586 | tags=90%, list=40%, signal=150% |
| 6615 | MELANOSOME\_TRANSPORT(GO:0032402) |  | 23 | 0.59 | 0.98 | 0.141 | 0.393 | 1.000 | 21618 | tags=61%, list=37%, signal=96% |
| 6616 | PROTEIN\_LOCALIZATION\_TO\_CHROMATIN(GO:0071168) |  | 23 | 0.59 | 0.98 | 0.141 | 0.393 | 1.000 | 23132 | tags=91%, list=39%, signal=150% |
| 6617 | NEGATIVE\_REGULATION\_OF\_SMALL\_GTPASE\_MEDIATED\_SIGNAL\_TRANSDUCTION(GO:0051058) |  | 53 | 0.58 | 0.98 | 0.141 | 0.393 | 1.000 | 17757 | tags=51%, list=30%, signal=73% |
| 6618 | PROTEIN\_DNA\_COMPLEX\_DISASSEMBLY(GO:0032986) |  | 20 | 0.57 | 0.98 | 0.476 | 0.394 | 1.000 | 22085 | tags=65%, list=37%, signal=104% |
| 6619 | POSITIVE\_REGULATION\_OF\_PROTEIN\_EXPORT\_FROM\_NUCLEUS(GO:0046827) |  | 21 | 0.60 | 0.98 | 0.141 | 0.394 | 1.000 | 22194 | tags=81%, list=38%, signal=130% |
| 6620 | CLATHRIN\_COAT\_OF\_COATED\_PIT(GO:0030132) |  | 19 | 0.56 | 0.98 | 0.476 | 0.394 | 1.000 | 26061 | tags=100%, list=44%, signal=179% |
| 6621 | ESTABLISHMENT\_OF\_MELANOSOME\_LOCALIZATION(GO:0032401) |  | 24 | 0.59 | 0.98 | 0.141 | 0.395 | 1.000 | 21618 | tags=58%, list=37%, signal=92% |
| 6622 | PEPTIDYL\_LYSINE\_DIMETHYLATION(GO:0018027) |  | 17 | 0.57 | 0.98 | 0.476 | 0.395 | 1.000 | 23844 | tags=71%, list=40%, signal=118% |
| 6623 | ICOSANOID\_RECEPTOR\_ACTIVITY(GO:0004953) |  | 15 | 0.61 | 0.98 | 0.141 | 0.395 | 1.000 | 19010 | tags=53%, list=32%, signal=79% |
| 6624 | POSITIVE\_REGULATION\_OF\_GLYCOPROTEIN\_BIOSYNTHETIC\_PROCESS(GO:0010560) |  | 19 | 0.59 | 0.98 | 0.141 | 0.396 | 1.000 | 19351 | tags=63%, list=33%, signal=94% |
| 6625 | REGULATION\_OF\_HISTONE\_H3\_K4\_METHYLATION(GO:0051569) |  | 27 | 0.60 | 0.98 | 0.310 | 0.396 | 1.000 | 20927 | tags=67%, list=35%, signal=103% |
| 6626 | VESICLE\_BUDDING\_FROM\_MEMBRANE(GO:0006900) |  | 92 | 0.55 | 0.98 | 0.476 | 0.396 | 1.000 | 26061 | tags=85%, list=44%, signal=151% |
| 6627 | INTRINSIC\_APOPTOTIC\_SIGNALING\_PATHWAY\_IN\_RESPONSE\_TO\_DNA\_DAMAGE(GO:0008630) |  | 103 | 0.58 | 0.98 | 0.141 | 0.396 | 1.000 | 22904 | tags=60%, list=39%, signal=98% |
| 6628 | RESPONSE\_TO\_INTERFERON\_BETA(GO:0035456) |  | 23 | 0.58 | 0.98 | 0.141 | 0.396 | 1.000 | 18684 | tags=39%, list=32%, signal=57% |
| 6629 | CELL\_DEATH\_IN\_RESPONSE\_TO\_HYDROGEN\_PEROXIDE(GO:0036474) |  | 35 | 0.58 | 0.98 | 0.141 | 0.397 | 1.000 | 24655 | tags=66%, list=42%, signal=113% |
| 6630 | RESPONSE\_TO\_MITOCHONDRIAL\_DEPOLARISATION(GO:0098780) |  | 19 | 0.58 | 0.98 | 0.476 | 0.397 | 1.000 | 25020 | tags=95%, list=42%, signal=164% |
| 6631 | TRANSMITTER\_GATED\_ION\_CHANNEL\_ACTIVITY\_INVOLVED\_IN\_REGULATION\_OF\_POSTSYNAPTIC\_MEMBRANE\_POTENTIAL(GO:1904315) |  | 20 | 0.58 | 0.98 | 0.141 | 0.397 | 1.000 | 21485 | tags=60%, list=36%, signal=94% |
| 6632 | NEUROTRANSMITTER\_RECEPTOR\_ACTIVITY\_INVOLVED\_IN\_REGULATION\_OF\_POSTSYNAPTIC\_MEMBRANE\_POTENTIAL(GO:0099529) |  | 20 | 0.58 | 0.98 | 0.141 | 0.397 | 1.000 | 21485 | tags=60%, list=36%, signal=94% |
| 6633 | SNARE\_COMPLEX(GO:0031201) |  | 52 | 0.56 | 0.98 | 0.141 | 0.398 | 1.000 | 24163 | tags=79%, list=41%, signal=133% |
| 6634 | REGULATION\_OF\_GOLGI\_ORGANIZATION(GO:1903358) |  | 15 | 0.55 | 0.98 | 0.141 | 0.398 | 1.000 | 25118 | tags=93%, list=42%, signal=162% |
| 6635 | PROGRAMMED\_NECROTIC\_CELL\_DEATH(GO:0097300) |  | 41 | 0.55 | 0.98 | 0.307 | 0.398 | 1.000 | 24562 | tags=68%, list=42%, signal=117% |
| 6636 | NEGATIVE\_REGULATION\_OF\_ORGANIC\_ACID\_TRANSPORT(GO:0032891) |  | 17 | 0.61 | 0.98 | 0.141 | 0.398 | 1.000 | 16608 | tags=29%, list=28%, signal=41% |
| 6637 | POSITIVE\_REGULATION\_OF\_GENE\_SILENCING\_BY\_MIRNA(GO:2000637) |  | 20 | 0.60 | 0.98 | 0.476 | 0.398 | 1.000 | 23586 | tags=90%, list=40%, signal=150% |
| 6638 | REGULATION\_OF\_RESPONSE\_TO\_INTERFERON\_GAMMA(GO:0060330) |  | 24 | 0.59 | 0.98 | 0.141 | 0.399 | 1.000 | 23558 | tags=75%, list=40%, signal=125% |
| 6639 | REGULATION\_OF\_INTERFERON\_GAMMA\_MEDIATED\_SIGNALING\_PATHWAY(GO:0060334) |  | 24 | 0.59 | 0.98 | 0.141 | 0.399 | 1.000 | 23558 | tags=75%, list=40%, signal=125% |
| 6640 | NEGATIVE\_REGULATION\_OF\_CYCLIC\_NUCLEOTIDE\_METABOLIC\_PROCESS(GO:0030800) |  | 35 | 0.56 | 0.98 | 0.310 | 0.399 | 1.000 | 23770 | tags=46%, list=40%, signal=76% |
| 6641 | INTERMEDIATE\_FILAMENT\_ORGANIZATION(GO:0045109) |  | 21 | 0.59 | 0.98 | 0.141 | 0.399 | 1.000 | 21122 | tags=43%, list=36%, signal=67% |
| 6642 | TRANS\_GOLGI\_NETWORK\_TRANSPORT\_VESICLE\_MEMBRANE(GO:0012510) |  | 15 | 0.56 | 0.98 | 0.307 | 0.400 | 1.000 | 25770 | tags=93%, list=44%, signal=165% |
| 6643 | CARGO\_LOADING\_INTO\_VESICLE(GO:0035459) |  | 17 | 0.58 | 0.98 | 0.481 | 0.400 | 1.000 | 23562 | tags=94%, list=40%, signal=156% |
| 6644 | MEMBRANE\_INVAGINATION(GO:0010324) |  | 106 | 0.56 | 0.98 | 0.141 | 0.400 | 1.000 | 24266 | tags=37%, list=41%, signal=62% |
| 6645 | INTRA\_GOLGI\_VESICLE\_MEDIATED\_TRANSPORT(GO:0006891) |  | 34 | 0.56 | 0.98 | 0.481 | 0.400 | 1.000 | 25294 | tags=88%, list=43%, signal=154% |
| 6646 | MULTIVESICULAR\_BODY\_SORTING\_PATHWAY(GO:0071985) |  | 28 | 0.56 | 0.98 | 0.141 | 0.401 | 1.000 | 26119 | tags=93%, list=44%, signal=166% |
| 6647 | NEGATIVE\_REGULATION\_OF\_ADHERENS\_JUNCTION\_ORGANIZATION(GO:1903392) |  | 16 | 0.61 | 0.98 | 0.141 | 0.401 | 1.000 | 21149 | tags=69%, list=36%, signal=107% |
| 6648 | ACYL\_COA\_HYDROLASE\_ACTIVITY(GO:0047617) |  | 17 | 0.61 | 0.98 | 0.307 | 0.402 | 1.000 | 21409 | tags=65%, list=36%, signal=101% |
| 6649 | POSITIVE\_REGULATION\_OF\_SIGNALING\_RECEPTOR\_ACTIVITY(GO:2000273) |  | 32 | 0.57 | 0.98 | 0.307 | 0.402 | 1.000 | 21827 | tags=56%, list=37%, signal=89% |
| 6650 | COA\_HYDROLASE\_ACTIVITY(GO:0016289) |  | 19 | 0.60 | 0.98 | 0.141 | 0.402 | 1.000 | 18374 | tags=53%, list=31%, signal=76% |
| 6651 | EXPLORATION\_BEHAVIOR(GO:0035640) |  | 27 | 0.58 | 0.98 | 0.307 | 0.403 | 1.000 | 21697 | tags=48%, list=37%, signal=76% |
| 6652 | MULTIVESICULAR\_BODY\_ASSEMBLY(GO:0036258) |  | 30 | 0.55 | 0.98 | 0.141 | 0.403 | 1.000 | 26168 | tags=97%, list=44%, signal=173% |
| 6653 | POSITIVE\_REGULATION\_OF\_MITOCHONDRIAL\_FISSION(GO:0090141) |  | 15 | 0.66 | 0.98 | 0.476 | 0.403 | 1.000 | 19159 | tags=80%, list=32%, signal=118% |
| 6654 | PROTON\_TRANSPORTING\_TWO\_SECTOR\_ATPASE\_COMPLEX\_\_CATALYTIC\_DOMAIN(GO:0033178) |  | 16 | 0.56 | 0.98 | 0.476 | 0.403 | 1.000 | 23252 | tags=81%, list=39%, signal=134% |
| 6655 | CELLULAR\_RESPONSE\_TO\_LIPOPROTEIN\_PARTICLE\_STIMULUS(GO:0071402) |  | 27 | 0.57 | 0.98 | 0.307 | 0.403 | 1.000 | 24090 | tags=56%, list=41%, signal=94% |
| 6656 | PRIMARY\_LYSOSOME(GO:0005766) |  | 152 | 0.55 | 0.98 | 0.141 | 0.403 | 1.000 | 22774 | tags=52%, list=39%, signal=84% |
| 6657 | AZUROPHIL\_GRANULE(GO:0042582) |  | 152 | 0.55 | 0.98 | 0.141 | 0.403 | 1.000 | 22774 | tags=52%, list=39%, signal=84% |
| 6658 | AEROBIC\_ELECTRON\_TRANSPORT\_CHAIN(GO:0019646) |  | 22 | 0.56 | 0.98 | 0.313 | 0.403 | 1.000 | 24260 | tags=77%, list=41%, signal=131% |
| 6659 | MITOCHONDRIAL\_ELECTRON\_TRANSPORT\_\_CYTOCHROME\_C\_TO\_OXYGEN(GO:0006123) |  | 22 | 0.56 | 0.98 | 0.313 | 0.403 | 1.000 | 24260 | tags=77%, list=41%, signal=131% |
| 6660 | MULTIVESICULAR\_BODY\_ORGANIZATION(GO:0036257) |  | 31 | 0.55 | 0.98 | 0.141 | 0.404 | 1.000 | 26168 | tags=97%, list=44%, signal=173% |
| 6661 | RIBOSOMAL\_SMALL\_SUBUNIT\_ASSEMBLY(GO:0000028) |  | 19 | 0.57 | 0.98 | 0.476 | 0.404 | 1.000 | 24427 | tags=89%, list=41%, signal=152% |
| 6662 | POSITIVE\_REGULATION\_OF\_LEUKOCYTE\_MIGRATION(GO:0002687) |  | 121 | 0.58 | 0.98 | 0.141 | 0.404 | 1.000 | 21928 | tags=46%, list=37%, signal=73% |
| 6663 | POSITIVE\_REGULATION\_OF\_EXOCYTOSIS(GO:0045921) |  | 79 | 0.56 | 0.98 | 0.141 | 0.404 | 1.000 | 24308 | tags=68%, list=41%, signal=116% |
| 6664 | LYMPHOCYTE\_CHEMOTAXIS(GO:0048247) |  | 59 | 0.60 | 0.98 | 0.482 | 0.405 | 1.000 | 9535 | tags=19%, list=16%, signal=22% |
| 6665 | MRNA\_TRANSCRIPTION(GO:0009299) |  | 25 | 0.56 | 0.98 | 0.476 | 0.405 | 1.000 | 23969 | tags=80%, list=41%, signal=134% |
| 6666 | NEGATIVE\_REGULATION\_OF\_CAMP\_METABOLIC\_PROCESS(GO:0030815) |  | 33 | 0.56 | 0.98 | 0.307 | 0.405 | 1.000 | 23749 | tags=42%, list=40%, signal=71% |
| 6667 | PYRIMIDINE\_NUCLEOSIDE\_MONOPHOSPHATE\_METABOLIC\_PROCESS(GO:0009129) |  | 15 | 0.58 | 0.98 | 0.476 | 0.405 | 1.000 | 23718 | tags=93%, list=40%, signal=156% |
| 6668 | GOLGI\_TO\_ENDOSOME\_TRANSPORT(GO:0006895) |  | 20 | 0.56 | 0.98 | 0.481 | 0.405 | 1.000 | 24869 | tags=90%, list=42%, signal=155% |
| 6669 | DNA\_METHYLATION\_INVOLVED\_IN\_GAMETE\_GENERATION(GO:0043046) |  | 20 | 0.59 | 0.98 | 0.141 | 0.406 | 1.000 | 22150 | tags=70%, list=37%, signal=112% |
| 6670 | REGULATION\_OF\_EXOSOMAL\_SECRETION(GO:1903541) |  | 16 | 0.57 | 0.98 | 0.476 | 0.406 | 1.000 | 24218 | tags=75%, list=41%, signal=127% |
| 6671 | VIRION\_ASSEMBLY(GO:0019068) |  | 37 | 0.54 | 0.98 | 0.476 | 0.406 | 1.000 | 26811 | tags=97%, list=45%, signal=178% |
| 6672 | POSITIVE\_REGULATION\_OF\_NOTCH\_SIGNALING\_PATHWAY(GO:0045747) |  | 47 | 0.57 | 0.98 | 0.141 | 0.406 | 1.000 | 20264 | tags=45%, list=34%, signal=68% |
| 6673 | GLYCOLIPID\_BINDING(GO:0051861) |  | 19 | 0.60 | 0.98 | 0.476 | 0.406 | 1.000 | 19318 | tags=53%, list=33%, signal=78% |
| 6674 | PHOSPHATIDYLINOSITOL\_PHOSPHATE\_KINASE\_ACTIVITY(GO:0016307) |  | 17 | 0.55 | 0.98 | 0.313 | 0.407 | 1.000 | 25512 | tags=76%, list=43%, signal=134% |
| 6675 | POSITIVE\_REGULATION\_OF\_VIRAL\_RELEASE\_FROM\_HOST\_CELL(GO:1902188) |  | 15 | 0.55 | 0.98 | 0.313 | 0.407 | 1.000 | 25358 | tags=93%, list=43%, signal=163% |
| 6676 | REGULATION\_OF\_ANTIGEN\_PROCESSING\_AND\_PRESENTATION(GO:0002577) |  | 17 | 0.64 | 0.98 | 0.476 | 0.407 | 1.000 | 12564 | tags=18%, list=21%, signal=22% |
| 6677 | SMOOTH\_ENDOPLASMIC\_RETICULUM(GO:0005790) |  | 33 | 0.58 | 0.98 | 0.141 | 0.408 | 1.000 | 22540 | tags=76%, list=38%, signal=122% |
| 6678 | GOLGI\_VESICLE\_BUDDING(GO:0048194) |  | 77 | 0.55 | 0.98 | 0.476 | 0.408 | 1.000 | 26218 | tags=86%, list=44%, signal=154% |
| 6679 | DNA\_METHYLATION(GO:0006306) |  | 69 | 0.58 | 0.98 | 0.141 | 0.409 | 1.000 | 23268 | tags=65%, list=39%, signal=107% |
| 6680 | DNA\_ALKYLATION(GO:0006305) |  | 69 | 0.58 | 0.98 | 0.141 | 0.408 | 1.000 | 23268 | tags=65%, list=39%, signal=107% |
| 6681 | NEGATIVE\_REGULATION\_OF\_CARBOHYDRATE\_METABOLIC\_PROCESS(GO:0045912) |  | 39 | 0.57 | 0.98 | 0.307 | 0.409 | 1.000 | 23468 | tags=64%, list=40%, signal=106% |
| 6682 | ANTIGEN\_PROCESSING\_AND\_PRESENTATION\_OF\_PEPTIDE\_ANTIGEN\_VIA\_MHC\_CLASS\_I(GO:0002474) |  | 95 | 0.55 | 0.97 | 0.141 | 0.409 | 1.000 | 24328 | tags=69%, list=41%, signal=118% |
| 6683 | URONIC\_ACID\_METABOLIC\_PROCESS(GO:0006063) |  | 25 | 0.56 | 0.97 | 0.307 | 0.409 | 1.000 | 23172 | tags=28%, list=39%, signal=46% |
| 6684 | GLUCURONATE\_METABOLIC\_PROCESS(GO:0019585) |  | 25 | 0.56 | 0.97 | 0.307 | 0.409 | 1.000 | 23172 | tags=28%, list=39%, signal=46% |
| 6685 | NAD+\_BINDING(GO:0070403) |  | 15 | 0.56 | 0.97 | 0.476 | 0.409 | 1.000 | 24149 | tags=73%, list=41%, signal=124% |
| 6686 | POSITIVE\_REGULATION\_OF\_PROTEIN\_INSERTION\_INTO\_MITOCHONDRIAL\_MEMBRANE\_INVOLVED\_IN\_APOPTOTIC\_SIGNALING\_PATHWAY(GO:1900740) |  | 31 | 0.60 | 0.97 | 0.141 | 0.411 | 1.000 | 21682 | tags=71%, list=37%, signal=112% |
| 6687 | REGULATION\_OF\_PROTEIN\_INSERTION\_INTO\_MITOCHONDRIAL\_MEMBRANE\_INVOLVED\_IN\_APOPTOTIC\_SIGNALING\_PATHWAY(GO:1900739) |  | 31 | 0.60 | 0.97 | 0.141 | 0.410 | 1.000 | 21682 | tags=71%, list=37%, signal=112% |
| 6688 | NEGATIVE\_REGULATION\_OF\_GENE\_SILENCING(GO:0060969) |  | 23 | 0.59 | 0.97 | 0.141 | 0.411 | 1.000 | 21951 | tags=70%, list=37%, signal=111% |
| 6689 | DEMETHYLASE\_ACTIVITY(GO:0032451) |  | 38 | 0.60 | 0.97 | 0.141 | 0.411 | 1.000 | 19369 | tags=61%, list=33%, signal=90% |
| 6690 | CHROMATIN\_DISASSEMBLY(GO:0031498) |  | 20 | 0.56 | 0.97 | 0.476 | 0.411 | 1.000 | 22085 | tags=60%, list=37%, signal=96% |
| 6691 | HEPATOCYTE\_GROWTH\_FACTOR\_RECEPTOR\_SIGNALING\_PATHWAY(GO:0048012) |  | 15 | 0.58 | 0.97 | 0.310 | 0.411 | 1.000 | 17052 | tags=40%, list=29%, signal=56% |
| 6692 | VESICLE\_COATING(GO:0006901) |  | 66 | 0.55 | 0.97 | 0.476 | 0.411 | 1.000 | 25898 | tags=82%, list=44%, signal=145% |
| 6693 | PHAGOCYTOSIS\_\_ENGULFMENT(GO:0006911) |  | 90 | 0.56 | 0.97 | 0.141 | 0.411 | 1.000 | 24241 | tags=28%, list=41%, signal=47% |
| 6694 | REGULATION\_OF\_DEFENSE\_RESPONSE\_TO\_VIRUS\_BY\_HOST(GO:0050691) |  | 37 | 0.57 | 0.97 | 0.141 | 0.411 | 1.000 | 23155 | tags=54%, list=39%, signal=89% |
| 6695 | HISTONE\_DEMETHYLASE\_ACTIVITY(GO:0032452) |  | 28 | 0.61 | 0.97 | 0.141 | 0.411 | 1.000 | 19369 | tags=61%, list=33%, signal=90% |
| 6696 | POSITIVE\_REGULATION\_OF\_HISTONE\_DEACETYLATION(GO:0031065) |  | 17 | 0.58 | 0.97 | 0.476 | 0.411 | 1.000 | 22117 | tags=65%, list=37%, signal=103% |
| 6697 | POSITIVE\_REGULATION\_OF\_T\_HELPER\_1\_TYPE\_IMMUNE\_RESPONSE(GO:0002827) |  | 15 | 0.59 | 0.97 | 0.307 | 0.411 | 1.000 | 20770 | tags=53%, list=35%, signal=82% |
| 6698 | VESICLE\_TARGETING\_\_ROUGH\_ER\_TO\_CIS\_GOLGI(GO:0048207) |  | 64 | 0.54 | 0.97 | 0.476 | 0.411 | 1.000 | 25898 | tags=81%, list=44%, signal=144% |
| 6699 | COPII\_VESICLE\_COATING(GO:0048208) |  | 64 | 0.54 | 0.97 | 0.476 | 0.411 | 1.000 | 25898 | tags=81%, list=44%, signal=144% |
| 6700 | VESICLE\_TARGETING\_\_TO\_\_FROM\_OR\_WITHIN\_GOLGI(GO:0048199) |  | 70 | 0.54 | 0.97 | 0.476 | 0.411 | 1.000 | 25898 | tags=81%, list=44%, signal=145% |
| 6701 | ENERGY\_COUPLED\_PROTON\_TRANSPORT\_\_DOWN\_ELECTROCHEMICAL\_GRADIENT(GO:0015985) |  | 28 | 0.57 | 0.97 | 0.141 | 0.411 | 1.000 | 22093 | tags=75%, list=37%, signal=120% |
| 6702 | ATP\_SYNTHESIS\_COUPLED\_PROTON\_TRANSPORT(GO:0015986) |  | 28 | 0.57 | 0.97 | 0.141 | 0.411 | 1.000 | 22093 | tags=75%, list=37%, signal=120% |
| 6703 | REGULATION\_OF\_MONONUCLEAR\_CELL\_MIGRATION(GO:0071675) |  | 38 | 0.58 | 0.97 | 0.141 | 0.412 | 1.000 | 22699 | tags=47%, list=38%, signal=77% |
| 6704 | PROTON\_TRANSPORTING\_TWO\_SECTOR\_ATPASE\_COMPLEX\_\_PROTON\_TRANSPORTING\_DOMAIN(GO:0033177) |  | 23 | 0.57 | 0.97 | 0.141 | 0.412 | 1.000 | 22093 | tags=78%, list=37%, signal=125% |
| 6705 | NEGATIVE\_REGULATION\_OF\_SMOOTH\_MUSCLE\_CELL\_DIFFERENTIATION(GO:0051151) |  | 19 | 0.61 | 0.97 | 0.141 | 0.412 | 1.000 | 19377 | tags=58%, list=33%, signal=86% |
| 6706 | INSULIN\_LIKE\_GROWTH\_FACTOR\_RECEPTOR\_SIGNALING\_PATHWAY(GO:0048009) |  | 38 | 0.58 | 0.97 | 0.141 | 0.412 | 1.000 | 22355 | tags=47%, list=38%, signal=76% |
| 6707 | HISTONE\_H3\_K4\_TRIMETHYLATION(GO:0080182) |  | 16 | 0.56 | 0.97 | 0.481 | 0.412 | 1.000 | 18261 | tags=56%, list=31%, signal=81% |
| 6708 | REGULATION\_OF\_HORMONE\_METABOLIC\_PROCESS(GO:0032350) |  | 35 | 0.60 | 0.97 | 0.476 | 0.412 | 1.000 | 10811 | tags=26%, list=18%, signal=31% |
| 6709 | NEGATIVE\_REGULATION\_OF\_RAS\_PROTEIN\_SIGNAL\_TRANSDUCTION(GO:0046580) |  | 47 | 0.58 | 0.97 | 0.141 | 0.413 | 1.000 | 19203 | tags=53%, list=32%, signal=79% |
| 6710 | POSITIVE\_REGULATION\_OF\_MAST\_CELL\_ACTIVATION(GO:0033005) |  | 18 | 0.58 | 0.97 | 0.307 | 0.413 | 1.000 | 18255 | tags=50%, list=31%, signal=72% |
| 6711 | TRANSLATION\_ELONGATION\_FACTOR\_ACTIVITY(GO:0003746) |  | 25 | 0.56 | 0.97 | 0.476 | 0.413 | 1.000 | 21819 | tags=64%, list=37%, signal=101% |
| 6712 | COPII\_COATED\_VESICLE\_BUDDING(GO:0090114) |  | 70 | 0.55 | 0.97 | 0.476 | 0.413 | 1.000 | 26218 | tags=84%, list=44%, signal=151% |
| 6713 | REGULATION\_OF\_MITOCHONDRIAL\_OUTER\_MEMBRANE\_PERMEABILIZATION\_INVOLVED\_IN\_APOPTOTIC\_SIGNALING\_PATHWAY(GO:1901028) |  | 48 | 0.58 | 0.97 | 0.141 | 0.413 | 1.000 | 22440 | tags=67%, list=38%, signal=107% |
| 6714 | REGULATION\_OF\_HYDROGEN\_PEROXIDE\_INDUCED\_CELL\_DEATH(GO:1903205) |  | 33 | 0.58 | 0.97 | 0.141 | 0.414 | 1.000 | 24655 | tags=64%, list=42%, signal=109% |
| 6715 | MATURE\_B\_CELL\_DIFFERENTIATION\_INVOLVED\_IN\_IMMUNE\_RESPONSE(GO:0002313) |  | 18 | 0.59 | 0.97 | 0.476 | 0.414 | 1.000 | 18653 | tags=33%, list=32%, signal=49% |
| 6716 | NEGATIVE\_REGULATION\_OF\_MEIOTIC\_CELL\_CYCLE(GO:0051447) |  | 17 | 0.59 | 0.97 | 0.141 | 0.414 | 1.000 | 19143 | tags=71%, list=32%, signal=104% |
| 6717 | ANTIGEN\_PROCESSING\_AND\_PRESENTATION\_OF\_EXOGENOUS\_PEPTIDE\_ANTIGEN\_VIA\_MHC\_CLASS\_I(GO:0042590) |  | 77 | 0.55 | 0.97 | 0.141 | 0.414 | 1.000 | 26127 | tags=79%, list=44%, signal=142% |
| 6718 | VERY\_LONG\_CHAIN\_FATTY\_ACID\_METABOLIC\_PROCESS(GO:0000038) |  | 31 | 0.56 | 0.97 | 0.141 | 0.415 | 1.000 | 21245 | tags=55%, list=36%, signal=86% |
| 6719 | POSITIVE\_REGULATION\_OF\_MITOCHONDRIAL\_OUTER\_MEMBRANE\_PERMEABILIZATION\_INVOLVED\_IN\_APOPTOTIC\_SIGNALING\_PATHWAY(GO:1901030) |  | 39 | 0.58 | 0.97 | 0.141 | 0.415 | 1.000 | 22440 | tags=69%, list=38%, signal=111% |
| 6720 | PROTON\_TRANSPORTING\_TWO\_SECTOR\_ATPASE\_COMPLEX(GO:0016469) |  | 52 | 0.56 | 0.97 | 0.141 | 0.415 | 1.000 | 23371 | tags=77%, list=40%, signal=127% |
| 6721 | RESPONSE\_TO\_ELECTRICAL\_STIMULUS(GO:0051602) |  | 43 | 0.56 | 0.97 | 0.141 | 0.415 | 1.000 | 23675 | tags=60%, list=40%, signal=101% |
| 6722 | MEMBRANE\_LIPID\_CATABOLIC\_PROCESS(GO:0046466) |  | 26 | 0.56 | 0.97 | 0.476 | 0.415 | 1.000 | 24712 | tags=69%, list=42%, signal=119% |
| 6723 | NUCLEOSOME\_DISASSEMBLY(GO:0006337) |  | 18 | 0.57 | 0.97 | 0.476 | 0.415 | 1.000 | 22085 | tags=61%, list=37%, signal=98% |
| 6724 | PLASMA\_MEMBRANE\_INVAGINATION(GO:0099024) |  | 98 | 0.56 | 0.97 | 0.141 | 0.416 | 1.000 | 24266 | tags=33%, list=41%, signal=55% |
| 6725 | EPIDERMAL\_GROWTH\_FACTOR\_RECEPTOR\_BINDING(GO:0005154) |  | 33 | 0.57 | 0.97 | 0.141 | 0.416 | 1.000 | 17190 | tags=42%, list=29%, signal=60% |
| 6726 | PIGMENT\_GRANULE\_LOCALIZATION(GO:0051875) |  | 27 | 0.58 | 0.97 | 0.141 | 0.417 | 1.000 | 21618 | tags=59%, list=37%, signal=93% |
| 6727 | TOLL\_LIKE\_RECEPTOR\_9\_SIGNALING\_PATHWAY(GO:0034162) |  | 23 | 0.57 | 0.97 | 0.141 | 0.417 | 1.000 | 18495 | tags=39%, list=31%, signal=57% |
| 6728 | INTRINSIC\_COMPONENT\_OF\_EXTERNAL\_SIDE\_OF\_PLASMA\_MEMBRANE(GO:0031233) |  | 20 | 0.59 | 0.97 | 0.476 | 0.417 | 1.000 | 20974 | tags=60%, list=35%, signal=93% |
| 6729 | REGULATION\_OF\_CELLULAR\_AMINO\_ACID\_METABOLIC\_PROCESS(GO:0006521) |  | 63 | 0.55 | 0.97 | 0.141 | 0.417 | 1.000 | 24328 | tags=70%, list=41%, signal=119% |
| 6730 | ANOIKIS(GO:0043276) |  | 32 | 0.56 | 0.97 | 0.476 | 0.417 | 1.000 | 21013 | tags=50%, list=36%, signal=78% |
| 6731 | ATP\_HYDROLYSIS\_COUPLED\_ION\_TRANSMEMBRANE\_TRANSPORT(GO:0099131) |  | 28 | 0.57 | 0.97 | 0.141 | 0.418 | 1.000 | 24101 | tags=82%, list=41%, signal=139% |
| 6732 | ANTIMICROBIAL\_HUMORAL\_RESPONSE(GO:0019730) |  | 108 | 0.57 | 0.97 | 0.307 | 0.418 | 1.000 | 20441 | tags=21%, list=35%, signal=32% |
| 6733 | SCF\_DEPENDENT\_PROTEASOMAL\_UBIQUITIN\_DEPENDENT\_PROTEIN\_CATABOLIC\_PROCESS(GO:0031146) |  | 73 | 0.54 | 0.97 | 0.476 | 0.418 | 1.000 | 26052 | tags=82%, list=44%, signal=147% |
| 6734 | NEGATIVE\_REGULATION\_OF\_CIRCADIAN\_RHYTHM(GO:0042754) |  | 15 | 0.56 | 0.97 | 0.644 | 0.418 | 1.000 | 25996 | tags=80%, list=44%, signal=143% |
| 6735 | PHOSPHATIDYLINOSITOL\_3\_5\_BISPHOSPHATE\_BINDING(GO:0080025) |  | 24 | 0.56 | 0.97 | 0.476 | 0.419 | 1.000 | 22125 | tags=71%, list=37%, signal=113% |
| 6736 | ATP\_HYDROLYSIS\_COUPLED\_TRANSMEMBRANE\_TRANSPORT(GO:0090662) |  | 30 | 0.57 | 0.97 | 0.141 | 0.420 | 1.000 | 24101 | tags=83%, list=41%, signal=141% |
| 6737 | ANTIGEN\_PROCESSING\_AND\_PRESENTATION\_OF\_EXOGENOUS\_PEPTIDE\_ANTIGEN\_VIA\_MHC\_CLASS\_I\_\_TAP\_DEPENDENT(GO:0002479) |  | 74 | 0.55 | 0.97 | 0.141 | 0.420 | 1.000 | 26127 | tags=78%, list=44%, signal=140% |
| 6738 | MEMBRANE\_COAT(GO:0030117) |  | 99 | 0.55 | 0.97 | 0.141 | 0.422 | 1.000 | 26293 | tags=93%, list=44%, signal=167% |
| 6739 | COATED\_MEMBRANE(GO:0048475) |  | 99 | 0.55 | 0.97 | 0.141 | 0.422 | 1.000 | 26293 | tags=93%, list=44%, signal=167% |
| 6740 | PORPHYRIN\_CONTAINING\_COMPOUND\_METABOLIC\_PROCESS(GO:0006778) |  | 38 | 0.56 | 0.97 | 0.476 | 0.423 | 1.000 | 24425 | tags=68%, list=41%, signal=116% |
| 6741 | RETINOIC\_ACID\_BINDING(GO:0001972) |  | 18 | 0.62 | 0.97 | 0.476 | 0.423 | 1.000 | 11950 | tags=22%, list=20%, signal=28% |
| 6742 | PROTEASOME\_ASSEMBLY(GO:0043248) |  | 16 | 0.56 | 0.97 | 0.141 | 0.423 | 1.000 | 25816 | tags=88%, list=44%, signal=155% |
| 6743 | NEGATIVE\_REGULATION\_OF\_HISTONE\_METHYLATION(GO:0031061) |  | 18 | 0.58 | 0.97 | 0.476 | 0.423 | 1.000 | 23231 | tags=78%, list=39%, signal=128% |
| 6744 | NEGATIVE\_REGULATION\_OF\_CYTOKINE\_BIOSYNTHETIC\_PROCESS(GO:0042036) |  | 28 | 0.59 | 0.97 | 0.476 | 0.423 | 1.000 | 19789 | tags=32%, list=33%, signal=48% |
| 6745 | REGULATION\_OF\_GENE\_SILENCING(GO:0060968) |  | 130 | 0.56 | 0.97 | 0.307 | 0.423 | 1.000 | 22953 | tags=68%, list=39%, signal=110% |
| 6746 | POSITIVE\_REGULATION\_OF\_GENE\_EXPRESSION\_\_EPIGENETIC(GO:0045815) |  | 79 | 0.54 | 0.97 | 0.141 | 0.423 | 1.000 | 24111 | tags=62%, list=41%, signal=105% |
| 6747 | POSITIVE\_REGULATION\_OF\_VASCULAR\_SMOOTH\_MUSCLE\_CELL\_PROLIFERATION(GO:1904707) |  | 34 | 0.58 | 0.97 | 0.141 | 0.423 | 1.000 | 18825 | tags=56%, list=32%, signal=82% |
| 6748 | OXYGEN\_BINDING(GO:0019825) |  | 48 | 0.57 | 0.97 | 0.310 | 0.423 | 1.000 | 18790 | tags=33%, list=32%, signal=49% |
| 6749 | REGULATION\_OF\_RETINOIC\_ACID\_RECEPTOR\_SIGNALING\_PATHWAY(GO:0048385) |  | 16 | 0.59 | 0.97 | 0.307 | 0.424 | 1.000 | 24182 | tags=81%, list=41%, signal=137% |
| 6750 | ENDOPEPTIDASE\_COMPLEX(GO:1905369) |  | 72 | 0.55 | 0.97 | 0.476 | 0.424 | 1.000 | 26688 | tags=89%, list=45%, signal=162% |
| 6751 | PROTEASOME\_COMPLEX(GO:0000502) |  | 71 | 0.55 | 0.97 | 0.476 | 0.424 | 1.000 | 26688 | tags=89%, list=45%, signal=162% |
| 6752 | HISTONE\_DEMETHYLATION(GO:0016577) |  | 30 | 0.59 | 0.97 | 0.141 | 0.424 | 1.000 | 20078 | tags=60%, list=34%, signal=91% |
| 6753 | REGULATION\_OF\_MACROPHAGE\_DIFFERENTIATION(GO:0045649) |  | 21 | 0.55 | 0.97 | 0.476 | 0.425 | 1.000 | 23092 | tags=67%, list=39%, signal=109% |
| 6754 | REGULATION\_OF\_INTRINSIC\_APOPTOTIC\_SIGNALING\_PATHWAY\_IN\_RESPONSE\_TO\_DNA\_DAMAGE(GO:1902229) |  | 37 | 0.58 | 0.97 | 0.307 | 0.425 | 1.000 | 21812 | tags=54%, list=37%, signal=86% |
| 6755 | PEPTIDYL\_TYROSINE\_AUTOPHOSPHORYLATION(GO:0038083) |  | 53 | 0.56 | 0.97 | 0.476 | 0.425 | 1.000 | 10643 | tags=19%, list=18%, signal=23% |
| 6756 | ENDOPLASMIC\_RETICULUM\_ORGANIZATION(GO:0007029) |  | 46 | 0.58 | 0.97 | 0.141 | 0.425 | 1.000 | 22138 | tags=74%, list=37%, signal=118% |
| 6757 | MEMBRANE\_PROTEIN\_PROTEOLYSIS(GO:0033619) |  | 65 | 0.54 | 0.97 | 0.476 | 0.426 | 1.000 | 25682 | tags=75%, list=43%, signal=133% |
| 6758 | OXIDOREDUCTASE\_ACTIVITY\_\_ACTING\_ON\_A\_HEME\_GROUP\_OF\_DONORS\_\_OXYGEN\_AS\_ACCEPTOR(GO:0016676) |  | 30 | 0.54 | 0.97 | 0.476 | 0.426 | 1.000 | 26337 | tags=83%, list=45%, signal=150% |
| 6759 | HEME\_COPPER\_TERMINAL\_OXIDASE\_ACTIVITY(GO:0015002) |  | 30 | 0.54 | 0.97 | 0.476 | 0.426 | 1.000 | 26337 | tags=83%, list=45%, signal=150% |
| 6760 | CYTOCHROME\_C\_OXIDASE\_ACTIVITY(GO:0004129) |  | 30 | 0.54 | 0.97 | 0.476 | 0.426 | 1.000 | 26337 | tags=83%, list=45%, signal=150% |
| 6761 | FUSION\_OF\_SPERM\_TO\_EGG\_PLASMA\_MEMBRANE\_INVOLVED\_IN\_SINGLE\_FERTILIZATION(GO:0007342) |  | 15 | 0.65 | 0.97 | 0.476 | 0.426 | 1.000 | 8306 | tags=13%, list=14%, signal=16% |
| 6762 | DEFENSE\_RESPONSE\_TO\_GRAM\_POSITIVE\_BACTERIUM(GO:0050830) |  | 92 | 0.55 | 0.97 | 0.141 | 0.426 | 1.000 | 24343 | tags=30%, list=41%, signal=52% |
| 6763 | INTERMEDIATE\_FILAMENT\_BASED\_PROCESS(GO:0045103) |  | 42 | 0.57 | 0.97 | 0.141 | 0.426 | 1.000 | 23181 | tags=52%, list=39%, signal=86% |
| 6764 | ACTIVATION\_OF\_MAPKKK\_ACTIVITY(GO:0000185) |  | 17 | 0.60 | 0.97 | 0.476 | 0.426 | 1.000 | 18373 | tags=47%, list=31%, signal=68% |
| 6765 | INACTIVATION\_OF\_MAPK\_ACTIVITY(GO:0000188) |  | 26 | 0.57 | 0.97 | 0.476 | 0.427 | 1.000 | 24108 | tags=65%, list=41%, signal=110% |
| 6766 | EPITHELIAL\_CELL\_MORPHOGENESIS(GO:0003382) |  | 30 | 0.58 | 0.97 | 0.307 | 0.428 | 1.000 | 21358 | tags=43%, list=36%, signal=68% |
| 6767 | VIRAL\_BUDDING(GO:0046755) |  | 24 | 0.55 | 0.97 | 0.475 | 0.428 | 1.000 | 26811 | tags=100%, list=45%, signal=183% |
| 6768 | POSITIVE\_REGULATION\_OF\_LONG\_TERM\_SYNAPTIC\_POTENTIATION(GO:1900273) |  | 17 | 0.55 | 0.97 | 0.475 | 0.428 | 1.000 | 26487 | tags=88%, list=45%, signal=160% |
| 6769 | NEGATIVE\_REGULATION\_OF\_LIPID\_STORAGE(GO:0010888) |  | 17 | 0.56 | 0.97 | 0.313 | 0.429 | 1.000 | 22269 | tags=53%, list=38%, signal=85% |
| 6770 | TRANSFERASE\_ACTIVITY\_\_TRANSFERRING\_AMINO\_ACYL\_GROUPS(GO:0016755) |  | 16 | 0.58 | 0.96 | 0.476 | 0.431 | 1.000 | 15281 | tags=31%, list=26%, signal=42% |
| 6771 | REGULATION\_OF\_AUTOPHAGOSOME\_ASSEMBLY(GO:2000785) |  | 38 | 0.56 | 0.96 | 0.141 | 0.431 | 1.000 | 20702 | tags=66%, list=35%, signal=101% |
| 6772 | VIRAL\_BUDDING\_VIA\_HOST\_ESCRT\_COMPLEX(GO:0039702) |  | 21 | 0.55 | 0.96 | 0.307 | 0.431 | 1.000 | 26811 | tags=100%, list=45%, signal=183% |
| 6773 | LOW\_DENSITY\_LIPOPROTEIN\_RECEPTOR\_PARTICLE\_METABOLIC\_PROCESS(GO:0032799) |  | 22 | 0.54 | 0.96 | 0.476 | 0.431 | 1.000 | 21005 | tags=55%, list=36%, signal=85% |
| 6774 | POSITIVE\_REGULATION\_OF\_MONOOXYGENASE\_ACTIVITY(GO:0032770) |  | 26 | 0.55 | 0.96 | 0.141 | 0.432 | 1.000 | 21674 | tags=46%, list=37%, signal=73% |
| 6775 | POSITIVE\_REGULATION\_OF\_INTRACELLULAR\_STEROID\_HORMONE\_RECEPTOR\_SIGNALING\_PATHWAY(GO:0033145) |  | 18 | 0.62 | 0.96 | 0.141 | 0.432 | 1.000 | 20916 | tags=72%, list=35%, signal=112% |
| 6776 | PEPTIDASE\_COMPLEX(GO:1905368) |  | 96 | 0.55 | 0.96 | 0.476 | 0.432 | 1.000 | 26688 | tags=88%, list=45%, signal=159% |
| 6777 | PIGMENT\_GRANULE\_TRANSPORT(GO:0051904) |  | 24 | 0.58 | 0.96 | 0.141 | 0.432 | 1.000 | 21618 | tags=58%, list=37%, signal=92% |
| 6778 | ANCHORED\_COMPONENT\_OF\_EXTERNAL\_SIDE\_OF\_PLASMA\_MEMBRANE(GO:0031362) |  | 15 | 0.56 | 0.96 | 0.476 | 0.432 | 1.000 | 23255 | tags=60%, list=39%, signal=99% |
| 6779 | COPI\_COATED\_VESICLE(GO:0030137) |  | 22 | 0.55 | 0.96 | 0.476 | 0.432 | 1.000 | 26013 | tags=86%, list=44%, signal=154% |
| 6780 | NEGATIVE\_REGULATION\_OF\_CAMP\_BIOSYNTHETIC\_PROCESS(GO:0030818) |  | 29 | 0.55 | 0.96 | 0.307 | 0.432 | 1.000 | 23749 | tags=41%, list=40%, signal=69% |
| 6781 | REGULATION\_OF\_ARP2\_3\_COMPLEX\_MEDIATED\_ACTIN\_NUCLEATION(GO:0034315) |  | 16 | 0.53 | 0.96 | 0.476 | 0.432 | 1.000 | 25121 | tags=75%, list=42%, signal=130% |
| 6782 | EPHRIN\_RECEPTOR\_BINDING(GO:0046875) |  | 29 | 0.56 | 0.96 | 0.476 | 0.433 | 1.000 | 24841 | tags=79%, list=42%, signal=137% |
| 6783 | PEPTIDYL\_ARGININE\_MODIFICATION(GO:0018195) |  | 26 | 0.59 | 0.96 | 0.476 | 0.433 | 1.000 | 21457 | tags=54%, list=36%, signal=84% |
| 6784 | DNA\_PACKAGING\_COMPLEX(GO:0044815) |  | 116 | 0.56 | 0.96 | 0.141 | 0.433 | 1.000 | 20540 | tags=30%, list=35%, signal=46% |
| 6785 | CYTOCHROME\_COMPLEX(GO:0070069) |  | 35 | 0.54 | 0.96 | 0.476 | 0.433 | 1.000 | 26608 | tags=83%, list=45%, signal=151% |
| 6786 | REGULATION\_OF\_MITOCHONDRIAL\_MEMBRANE\_PERMEABILITY\_INVOLVED\_IN\_APOPTOTIC\_PROCESS(GO:1902108) |  | 68 | 0.57 | 0.96 | 0.141 | 0.433 | 1.000 | 22440 | tags=62%, list=38%, signal=99% |
| 6787 | ESTABLISHMENT\_OF\_PIGMENT\_GRANULE\_LOCALIZATION(GO:0051905) |  | 25 | 0.58 | 0.96 | 0.141 | 0.433 | 1.000 | 21618 | tags=56%, list=37%, signal=88% |
| 6788 | POSITIVE\_REGULATION\_OF\_MITOCHONDRIAL\_MEMBRANE\_PERMEABILITY\_INVOLVED\_IN\_APOPTOTIC\_PROCESS(GO:1902110) |  | 62 | 0.57 | 0.96 | 0.141 | 0.434 | 1.000 | 22440 | tags=63%, list=38%, signal=101% |
| 6789 | PROTEIN\_INSERTION\_INTO\_MEMBRANE(GO:0051205) |  | 51 | 0.58 | 0.96 | 0.141 | 0.434 | 1.000 | 22745 | tags=69%, list=38%, signal=111% |
| 6790 | COPPER\_ION\_HOMEOSTASIS(GO:0055070) |  | 18 | 0.55 | 0.96 | 0.476 | 0.434 | 1.000 | 24116 | tags=78%, list=41%, signal=131% |
| 6791 | REGULATION\_OF\_PRI\_MIRNA\_TRANSCRIPTION\_BY\_RNA\_POLYMERASE\_II(GO:1902893) |  | 33 | 0.58 | 0.96 | 0.476 | 0.434 | 1.000 | 24079 | tags=76%, list=41%, signal=128% |
| 6792 | NEGATIVE\_REGULATION\_OF\_FOCAL\_ADHESION\_ASSEMBLY(GO:0051895) |  | 15 | 0.60 | 0.96 | 0.141 | 0.434 | 1.000 | 21149 | tags=67%, list=36%, signal=104% |
| 6793 | POSITIVE\_REGULATION\_OF\_PROTEIN\_DEACETYLATION(GO:0090312) |  | 21 | 0.57 | 0.96 | 0.476 | 0.434 | 1.000 | 22117 | tags=62%, list=37%, signal=99% |
| 6794 | SPHINGOLIPID\_CATABOLIC\_PROCESS(GO:0030149) |  | 23 | 0.56 | 0.96 | 0.476 | 0.435 | 1.000 | 24712 | tags=70%, list=42%, signal=119% |
| 6795 | REGULATION\_OF\_EPIDERMAL\_GROWTH\_FACTOR\_ACTIVATED\_RECEPTOR\_ACTIVITY(GO:0007176) |  | 29 | 0.54 | 0.96 | 0.476 | 0.435 | 1.000 | 25905 | tags=79%, list=44%, signal=141% |
| 6796 | TRANSCRIPTION\_BY\_RNA\_POLYMERASE\_III(GO:0006383) |  | 54 | 0.57 | 0.96 | 0.141 | 0.435 | 1.000 | 23679 | tags=85%, list=40%, signal=142% |
| 6797 | POSITIVE\_REGULATION\_OF\_EPITHELIAL\_CELL\_APOPTOTIC\_PROCESS(GO:1904037) |  | 37 | 0.57 | 0.96 | 0.141 | 0.435 | 1.000 | 23558 | tags=41%, list=40%, signal=67% |
| 6798 | REGULATION\_OF\_TRANSCRIPTION\_FROM\_RNA\_POLYMERASE\_II\_PROMOTER\_IN\_RESPONSE\_TO\_HYPOXIA(GO:0061418) |  | 80 | 0.54 | 0.96 | 0.476 | 0.435 | 1.000 | 26149 | tags=81%, list=44%, signal=145% |
| 6799 | PROTEIN\_TYROSINE\_KINASE\_ACTIVATOR\_ACTIVITY(GO:0030296) |  | 19 | 0.57 | 0.96 | 0.310 | 0.436 | 1.000 | 18635 | tags=47%, list=32%, signal=69% |
| 6800 | STEROL\_HOMEOSTASIS(GO:0055092) |  | 70 | 0.54 | 0.96 | 0.307 | 0.436 | 1.000 | 21292 | tags=40%, list=36%, signal=62% |
| 6801 | CHOLESTEROL\_HOMEOSTASIS(GO:0042632) |  | 70 | 0.54 | 0.96 | 0.307 | 0.436 | 1.000 | 21292 | tags=40%, list=36%, signal=62% |
| 6802 | HISTONE\_LYSINE\_DEMETHYLATION(GO:0070076) |  | 28 | 0.59 | 0.96 | 0.141 | 0.436 | 1.000 | 20078 | tags=61%, list=34%, signal=92% |
| 6803 | ENTRAINMENT\_OF\_CIRCADIAN\_CLOCK\_BY\_PHOTOPERIOD(GO:0043153) |  | 18 | 0.55 | 0.96 | 0.476 | 0.436 | 1.000 | 23631 | tags=78%, list=40%, signal=130% |
| 6804 | CYTOPLASMIC\_SIDE\_OF\_ENDOPLASMIC\_RETICULUM\_MEMBRANE(GO:0098554) |  | 15 | 0.55 | 0.96 | 0.476 | 0.436 | 1.000 | 26666 | tags=93%, list=45%, signal=170% |
| 6805 | PALMITOYLTRANSFERASE\_ACTIVITY(GO:0016409) |  | 38 | 0.57 | 0.96 | 0.141 | 0.436 | 1.000 | 20368 | tags=63%, list=34%, signal=96% |
| 6806 | CELLULAR\_MODIFIED\_AMINO\_ACID\_BIOSYNTHETIC\_PROCESS(GO:0042398) |  | 39 | 0.57 | 0.96 | 0.310 | 0.437 | 1.000 | 24300 | tags=79%, list=41%, signal=135% |
| 6807 | OXIDOREDUCTASE\_ACTIVITY\_\_ACTING\_ON\_A\_HEME\_GROUP\_OF\_DONORS(GO:0016675) |  | 31 | 0.54 | 0.96 | 0.476 | 0.437 | 1.000 | 27259 | tags=87%, list=46%, signal=161% |
| 6808 | MHC\_CLASS\_I\_PROTEIN\_BINDING(GO:0042288) |  | 16 | 0.55 | 0.96 | 0.310 | 0.437 | 1.000 | 26127 | tags=75%, list=44%, signal=134% |
| 6809 | MITOGEN\_ACTIVATED\_PROTEIN\_KINASE\_KINASE\_KINASE\_BINDING(GO:0031435) |  | 16 | 0.57 | 0.96 | 0.476 | 0.437 | 1.000 | 17552 | tags=44%, list=30%, signal=62% |
| 6810 | MEGAKARYOCYTE\_DIFFERENTIATION(GO:0030219) |  | 92 | 0.56 | 0.96 | 0.476 | 0.438 | 1.000 | 21405 | tags=52%, list=36%, signal=82% |
| 6811 | RESPIRATORY\_BURST(GO:0045730) |  | 30 | 0.56 | 0.96 | 0.476 | 0.438 | 1.000 | 25883 | tags=70%, list=44%, signal=124% |
| 6812 | PROTEIN\_DEALKYLATION(GO:0008214) |  | 32 | 0.59 | 0.96 | 0.141 | 0.438 | 1.000 | 21422 | tags=66%, list=36%, signal=103% |
| 6813 | PROTEIN\_DEMETHYLATION(GO:0006482) |  | 32 | 0.59 | 0.96 | 0.141 | 0.438 | 1.000 | 21422 | tags=66%, list=36%, signal=103% |
| 6814 | RECEPTOR\_CATABOLIC\_PROCESS(GO:0032801) |  | 33 | 0.54 | 0.96 | 0.476 | 0.438 | 1.000 | 26415 | tags=85%, list=45%, signal=153% |
| 6815 | NEGATIVE\_REGULATION\_OF\_PROTEIN\_TYROSINE\_KINASE\_ACTIVITY(GO:0061099) |  | 27 | 0.55 | 0.96 | 0.476 | 0.438 | 1.000 | 24310 | tags=70%, list=41%, signal=119% |
| 6816 | PROTEIN\_HETEROTETRAMERIZATION(GO:0051290) |  | 48 | 0.56 | 0.96 | 0.141 | 0.438 | 1.000 | 18017 | tags=35%, list=30%, signal=51% |
| 6817 | ERBB2\_SIGNALING\_PATHWAY(GO:0038128) |  | 39 | 0.55 | 0.96 | 0.141 | 0.439 | 1.000 | 26040 | tags=79%, list=44%, signal=142% |
| 6818 | NEUROMUSCULAR\_SYNAPTIC\_TRANSMISSION(GO:0007274) |  | 30 | 0.56 | 0.96 | 0.307 | 0.439 | 1.000 | 19932 | tags=50%, list=34%, signal=75% |
| 6819 | NEUROTRANSMITTER\_CATABOLIC\_PROCESS(GO:0042135) |  | 24 | 0.58 | 0.96 | 0.310 | 0.440 | 1.000 | 21301 | tags=58%, list=36%, signal=91% |
| 6820 | NEGATIVE\_REGULATION\_OF\_INSULIN\_RECEPTOR\_SIGNALING\_PATHWAY(GO:0046627) |  | 31 | 0.55 | 0.96 | 0.310 | 0.440 | 1.000 | 24541 | tags=68%, list=41%, signal=116% |
| 6821 | CELL\_SEPARATION\_AFTER\_CYTOKINESIS(GO:0000920) |  | 17 | 0.55 | 0.96 | 0.647 | 0.441 | 1.000 | 26811 | tags=100%, list=45%, signal=183% |
| 6822 | REGULATION\_OF\_POSTTRANSCRIPTIONAL\_GENE\_SILENCING(GO:0060147) |  | 108 | 0.56 | 0.96 | 0.476 | 0.441 | 1.000 | 20933 | tags=58%, list=35%, signal=90% |
| 6823 | REGULATION\_OF\_GENE\_SILENCING\_BY\_RNA(GO:0060966) |  | 108 | 0.56 | 0.96 | 0.476 | 0.441 | 1.000 | 20933 | tags=58%, list=35%, signal=90% |
| 6824 | EXIT\_FROM\_HOST\_CELL(GO:0035891) |  | 35 | 0.54 | 0.96 | 0.650 | 0.442 | 1.000 | 26728 | tags=86%, list=45%, signal=156% |
| 6825 | VIRAL\_RELEASE\_FROM\_HOST\_CELL(GO:0019076) |  | 35 | 0.54 | 0.96 | 0.650 | 0.442 | 1.000 | 26728 | tags=86%, list=45%, signal=156% |
| 6826 | EXIT\_FROM\_HOST(GO:0035890) |  | 35 | 0.54 | 0.96 | 0.650 | 0.441 | 1.000 | 26728 | tags=86%, list=45%, signal=156% |
| 6827 | MOVEMENT\_IN\_ENVIRONMENT\_OF\_OTHER\_ORGANISM\_INVOLVED\_IN\_SYMBIOTIC\_INTERACTION(GO:0052192) |  | 35 | 0.54 | 0.96 | 0.650 | 0.441 | 1.000 | 26728 | tags=86%, list=45%, signal=156% |
| 6828 | MOVEMENT\_IN\_HOST\_ENVIRONMENT(GO:0052126) |  | 35 | 0.54 | 0.96 | 0.650 | 0.441 | 1.000 | 26728 | tags=86%, list=45%, signal=156% |
| 6829 | PROTEIN\_INSERTION\_INTO\_MITOCHONDRIAL\_MEMBRANE(GO:0051204) |  | 36 | 0.59 | 0.96 | 0.141 | 0.442 | 1.000 | 21682 | tags=69%, list=37%, signal=110% |
| 6830 | PLASMA\_MEMBRANE\_TUBULATION(GO:0097320) |  | 17 | 0.56 | 0.96 | 0.476 | 0.443 | 1.000 | 25682 | tags=88%, list=43%, signal=156% |
| 6831 | REGULATION\_OF\_DNA\_TEMPLATED\_TRANSCRIPTION\_IN\_RESPONSE\_TO\_STRESS(GO:0043620) |  | 119 | 0.54 | 0.96 | 0.476 | 0.443 | 1.000 | 25582 | tags=75%, list=43%, signal=132% |
| 6832 | ENERGY\_COUPLED\_PROTON\_TRANSMEMBRANE\_TRANSPORT\_\_AGAINST\_ELECTROCHEMICAL\_GRADIENT(GO:0015988) |  | 28 | 0.56 | 0.96 | 0.307 | 0.444 | 1.000 | 24101 | tags=82%, list=41%, signal=139% |
| 6833 | NEGATIVE\_REGULATION\_OF\_CYCLIC\_NUCLEOTIDE\_BIOSYNTHETIC\_PROCESS(GO:0030803) |  | 31 | 0.55 | 0.96 | 0.310 | 0.444 | 1.000 | 23770 | tags=45%, list=40%, signal=75% |
| 6834 | OXYGEN\_TRANSPORT(GO:0015671) |  | 17 | 0.61 | 0.96 | 0.476 | 0.444 | 1.000 | 15820 | tags=29%, list=27%, signal=40% |
| 6835 | INTERMEDIATE\_FILAMENT\_CYTOSKELETON\_ORGANIZATION(GO:0045104) |  | 41 | 0.56 | 0.96 | 0.310 | 0.444 | 1.000 | 23181 | tags=51%, list=39%, signal=84% |
| 6836 | REGULATION\_OF\_TRANSCRIPTION\_FROM\_RNA\_POLYMERASE\_II\_PROMOTER\_IN\_RESPONSE\_TO\_STRESS(GO:0043618) |  | 113 | 0.54 | 0.96 | 0.476 | 0.444 | 1.000 | 26149 | tags=79%, list=44%, signal=141% |
| 6837 | PRI\_MIRNA\_TRANSCRIPTION\_BY\_RNA\_POLYMERASE\_II(GO:0061614) |  | 39 | 0.57 | 0.96 | 0.476 | 0.444 | 1.000 | 24079 | tags=74%, list=41%, signal=125% |
| 6838 | MITOCHONDRIAL\_OUTER\_MEMBRANE\_PERMEABILIZATION(GO:0097345) |  | 56 | 0.57 | 0.96 | 0.141 | 0.445 | 1.000 | 22440 | tags=66%, list=38%, signal=106% |
| 6839 | NEGATIVE\_REGULATION\_OF\_CELLULAR\_RESPONSE\_TO\_INSULIN\_STIMULUS(GO:1900077) |  | 34 | 0.55 | 0.96 | 0.310 | 0.445 | 1.000 | 24541 | tags=65%, list=41%, signal=111% |
| 6840 | NEGATIVE\_REGULATION\_OF\_PURINE\_NUCLEOTIDE\_BIOSYNTHETIC\_PROCESS(GO:1900372) |  | 46 | 0.57 | 0.96 | 0.476 | 0.445 | 1.000 | 23770 | tags=54%, list=40%, signal=91% |
| 6841 | NEGATIVE\_REGULATION\_OF\_NUCLEOTIDE\_BIOSYNTHETIC\_PROCESS(GO:0030809) |  | 46 | 0.57 | 0.96 | 0.476 | 0.445 | 1.000 | 23770 | tags=54%, list=40%, signal=91% |
| 6842 | PROTEIN\_INSERTION\_INTO\_MITOCHONDRIAL\_MEMBRANE\_INVOLVED\_IN\_APOPTOTIC\_SIGNALING\_PATHWAY(GO:0001844) |  | 33 | 0.58 | 0.96 | 0.141 | 0.445 | 1.000 | 21682 | tags=67%, list=37%, signal=105% |
| 6843 | PHOSPHATIDYLINOSITOL\_3\_PHOSPHATE\_BINDING(GO:0032266) |  | 35 | 0.54 | 0.96 | 0.476 | 0.447 | 1.000 | 23763 | tags=66%, list=40%, signal=110% |
| 6844 | PHOSPHATIDYLETHANOLAMINE\_METABOLIC\_PROCESS(GO:0046337) |  | 18 | 0.55 | 0.96 | 0.307 | 0.448 | 1.000 | 26546 | tags=89%, list=45%, signal=161% |
| 6845 | CALCIUM\_DEPENDENT\_PROTEIN\_BINDING(GO:0048306) |  | 60 | 0.57 | 0.96 | 0.310 | 0.448 | 1.000 | 22046 | tags=53%, list=37%, signal=85% |
| 6846 | GERM\_CELL\_NUCLEUS(GO:0043073) |  | 19 | 0.62 | 0.96 | 0.309 | 0.448 | 1.000 | 16861 | tags=47%, list=29%, signal=66% |
| 6847 | REGULATION\_OF\_GENE\_SILENCING\_BY\_MIRNA(GO:0060964) |  | 105 | 0.56 | 0.96 | 0.476 | 0.448 | 1.000 | 20933 | tags=57%, list=35%, signal=88% |
| 6848 | NEGATIVE\_REGULATION\_OF\_GENE\_EXPRESSION\_\_EPIGENETIC(GO:0045814) |  | 133 | 0.55 | 0.96 | 0.141 | 0.449 | 1.000 | 22226 | tags=52%, list=38%, signal=83% |
| 6849 | NEGATIVE\_REGULATION\_OF\_INNATE\_IMMUNE\_RESPONSE(GO:0045824) |  | 44 | 0.57 | 0.96 | 0.307 | 0.449 | 1.000 | 20082 | tags=43%, list=34%, signal=65% |
| 6850 | POSITIVE\_REGULATION\_OF\_VIRAL\_LIFE\_CYCLE(GO:1903902) |  | 57 | 0.57 | 0.96 | 0.141 | 0.449 | 1.000 | 22174 | tags=70%, list=37%, signal=112% |
| 6851 | TASTE\_RECEPTOR\_ACTIVITY(GO:0008527) |  | 26 | 0.56 | 0.96 | 0.481 | 0.449 | 1.000 | 24203 | tags=35%, list=41%, signal=59% |
| 6852 | REGULATION\_OF\_VIRAL\_RELEASE\_FROM\_HOST\_CELL(GO:1902186) |  | 31 | 0.54 | 0.96 | 0.482 | 0.449 | 1.000 | 26728 | tags=84%, list=45%, signal=153% |
| 6853 | ATP\_HYDROLYSIS\_COUPLED\_PROTON\_TRANSPORT(GO:0015991) |  | 27 | 0.56 | 0.96 | 0.307 | 0.450 | 1.000 | 24101 | tags=81%, list=41%, signal=137% |
| 6854 | PROTEASOME\_REGULATORY\_PARTICLE(GO:0005838) |  | 22 | 0.56 | 0.96 | 0.476 | 0.451 | 1.000 | 25832 | tags=91%, list=44%, signal=161% |
| 6855 | MITOCHONDRIAL\_OUTER\_MEMBRANE\_PERMEABILIZATION\_INVOLVED\_IN\_PROGRAMMED\_CELL\_DEATH(GO:1902686) |  | 64 | 0.57 | 0.95 | 0.141 | 0.451 | 1.000 | 22440 | tags=63%, list=38%, signal=101% |
| 6856 | THYROID\_HORMONE\_GENERATION(GO:0006590) |  | 17 | 0.58 | 0.95 | 0.476 | 0.451 | 1.000 | 12505 | tags=29%, list=21%, signal=37% |
| 6857 | POSITIVE\_REGULATION\_OF\_EXOSOMAL\_SECRETION(GO:1903543) |  | 15 | 0.56 | 0.95 | 0.476 | 0.453 | 1.000 | 24218 | tags=73%, list=41%, signal=124% |
| 6858 | POSITIVE\_REGULATION\_OF\_AXON\_EXTENSION(GO:0045773) |  | 40 | 0.57 | 0.95 | 0.141 | 0.453 | 1.000 | 23016 | tags=68%, list=39%, signal=110% |
| 6859 | NEGATIVE\_REGULATION\_OF\_RESPONSE\_TO\_REACTIVE\_OXYGEN\_SPECIES(GO:1901032) |  | 26 | 0.58 | 0.95 | 0.476 | 0.453 | 1.000 | 24655 | tags=58%, list=42%, signal=99% |
| 6860 | NEGATIVE\_REGULATION\_OF\_CELLULAR\_RESPONSE\_TO\_DRUG(GO:2001039) |  | 26 | 0.58 | 0.95 | 0.476 | 0.453 | 1.000 | 24655 | tags=58%, list=42%, signal=99% |
| 6861 | NEGATIVE\_REGULATION\_OF\_HYDROGEN\_PEROXIDE\_INDUCED\_CELL\_DEATH(GO:1903206) |  | 26 | 0.58 | 0.95 | 0.476 | 0.453 | 1.000 | 24655 | tags=58%, list=42%, signal=99% |
| 6862 | NEGATIVE\_REGULATION\_OF\_NECROTIC\_CELL\_DEATH(GO:0060547) |  | 15 | 0.57 | 0.95 | 0.476 | 0.454 | 1.000 | 21199 | tags=53%, list=36%, signal=83% |
| 6863 | IRE1\_MEDIATED\_UNFOLDED\_PROTEIN\_RESPONSE(GO:0036498) |  | 70 | 0.55 | 0.95 | 0.476 | 0.454 | 1.000 | 25388 | tags=83%, list=43%, signal=145% |
| 6864 | REGULATION\_OF\_NECROTIC\_CELL\_DEATH(GO:0010939) |  | 30 | 0.55 | 0.95 | 0.476 | 0.454 | 1.000 | 24124 | tags=63%, list=41%, signal=107% |
| 6865 | POSITIVE\_REGULATION\_OF\_MITOCHONDRIAL\_MEMBRANE\_PERMEABILITY(GO:0035794) |  | 66 | 0.57 | 0.95 | 0.141 | 0.454 | 1.000 | 22440 | tags=62%, list=38%, signal=100% |
| 6866 | POSITIVE\_REGULATION\_OF\_MEMBRANE\_PERMEABILITY(GO:1905710) |  | 68 | 0.57 | 0.95 | 0.141 | 0.454 | 1.000 | 22440 | tags=60%, list=38%, signal=97% |
| 6867 | VACUOLAR\_ACIDIFICATION(GO:0007035) |  | 22 | 0.56 | 0.95 | 0.141 | 0.455 | 1.000 | 22005 | tags=68%, list=37%, signal=109% |
| 6868 | NEGATIVE\_REGULATION\_OF\_ADENYLATE\_CYCLASE\_ACTIVITY(GO:0007194) |  | 28 | 0.54 | 0.95 | 0.307 | 0.456 | 1.000 | 23749 | tags=39%, list=40%, signal=66% |
| 6869 | PROTEIN\_O\_LINKED\_MANNOSYLATION(GO:0035269) |  | 15 | 0.56 | 0.95 | 0.476 | 0.457 | 1.000 | 25488 | tags=93%, list=43%, signal=164% |
| 6870 | CELLULAR\_RESPONSE\_TO\_GROWTH\_HORMONE\_STIMULUS(GO:0071378) |  | 24 | 0.56 | 0.95 | 0.481 | 0.459 | 1.000 | 25118 | tags=67%, list=42%, signal=116% |
| 6871 | CXCR\_CHEMOKINE\_RECEPTOR\_BINDING(GO:0045236) |  | 16 | 0.62 | 0.95 | 0.647 | 0.460 | 1.000 | 15120 | tags=38%, list=26%, signal=50% |
| 6872 | PENTOSE\_PHOSPHATE\_SHUNT(GO:0006098) |  | 19 | 0.55 | 0.95 | 0.476 | 0.461 | 1.000 | 25678 | tags=84%, list=43%, signal=149% |
| 6873 | POSITIVE\_REGULATION\_OF\_LEUKOCYTE\_DEGRANULATION(GO:0043302) |  | 21 | 0.57 | 0.95 | 0.476 | 0.462 | 1.000 | 24906 | tags=76%, list=42%, signal=132% |
| 6874 | REGULATION\_OF\_VACUOLE\_ORGANIZATION(GO:0044088) |  | 42 | 0.55 | 0.95 | 0.141 | 0.462 | 1.000 | 22896 | tags=71%, list=39%, signal=116% |
| 6875 | NEGATIVE\_REGULATION\_OF\_CYCLASE\_ACTIVITY(GO:0031280) |  | 30 | 0.55 | 0.95 | 0.310 | 0.463 | 1.000 | 23770 | tags=43%, list=40%, signal=72% |
| 6876 | POSITIVE\_REGULATION\_OF\_TRIGLYCERIDE\_METABOLIC\_PROCESS(GO:0090208) |  | 19 | 0.57 | 0.95 | 0.476 | 0.463 | 1.000 | 22269 | tags=53%, list=38%, signal=84% |
| 6877 | LEUKOCYTE\_TETHERING\_OR\_ROLLING(GO:0050901) |  | 24 | 0.55 | 0.95 | 0.309 | 0.464 | 1.000 | 24352 | tags=54%, list=41%, signal=92% |
| 6878 | POSITIVE\_REGULATION\_OF\_TRANSCRIPTION\_FROM\_RNA\_POLYMERASE\_II\_PROMOTER\_IN\_RESPONSE\_TO\_STRESS(GO:0036003) |  | 24 | 0.54 | 0.95 | 0.476 | 0.465 | 1.000 | 26997 | tags=88%, list=46%, signal=161% |
| 6879 | PHAGOPHORE\_ASSEMBLY\_SITE(GO:0000407) |  | 32 | 0.53 | 0.95 | 0.476 | 0.465 | 1.000 | 26570 | tags=84%, list=45%, signal=153% |
| 6880 | NUCLEOTIDE\_SUGAR\_BIOSYNTHETIC\_PROCESS(GO:0009226) |  | 21 | 0.53 | 0.95 | 0.476 | 0.465 | 1.000 | 24833 | tags=71%, list=42%, signal=123% |
| 6881 | ADENYLYLTRANSFERASE\_ACTIVITY(GO:0070566) |  | 24 | 0.57 | 0.95 | 0.313 | 0.465 | 1.000 | 25013 | tags=79%, list=42%, signal=137% |
| 6882 | PHAGOPHORE\_ASSEMBLY\_SITE\_MEMBRANE(GO:0034045) |  | 16 | 0.54 | 0.95 | 0.476 | 0.466 | 1.000 | 26570 | tags=88%, list=45%, signal=159% |
| 6883 | CHROMATIN\_SILENCING(GO:0006342) |  | 117 | 0.55 | 0.95 | 0.141 | 0.466 | 1.000 | 22226 | tags=50%, list=38%, signal=79% |
| 6884 | LIPOPROTEIN\_PARTICLE\_RECEPTOR\_BINDING(GO:0070325) |  | 26 | 0.54 | 0.95 | 0.476 | 0.468 | 1.000 | 21988 | tags=62%, list=37%, signal=98% |
| 6885 | REGULATION\_OF\_B\_CELL\_APOPTOTIC\_PROCESS(GO:0002902) |  | 18 | 0.57 | 0.95 | 0.310 | 0.469 | 1.000 | 25019 | tags=72%, list=42%, signal=125% |
| 6886 | AP\_TYPE\_MEMBRANE\_COAT\_ADAPTOR\_COMPLEX(GO:0030119) |  | 49 | 0.55 | 0.95 | 0.307 | 0.469 | 1.000 | 26293 | tags=92%, list=44%, signal=165% |
| 6887 | PROTEIN\_PHOSPHORYLATED\_AMINO\_ACID\_BINDING(GO:0045309) |  | 44 | 0.54 | 0.95 | 0.476 | 0.469 | 1.000 | 26758 | tags=80%, list=45%, signal=145% |
| 6888 | MHC\_PROTEIN\_COMPLEX\_BINDING(GO:0023023) |  | 20 | 0.54 | 0.95 | 0.475 | 0.469 | 1.000 | 25096 | tags=70%, list=42%, signal=122% |
| 6889 | POSITIVE\_REGULATION\_OF\_GLYCOPROTEIN\_METABOLIC\_PROCESS(GO:1903020) |  | 23 | 0.56 | 0.95 | 0.141 | 0.469 | 1.000 | 25464 | tags=78%, list=43%, signal=137% |
| 6890 | GLUCONEOGENESIS(GO:0006094) |  | 79 | 0.56 | 0.95 | 0.476 | 0.470 | 1.000 | 23751 | tags=66%, list=40%, signal=110% |
| 6891 | VACUOLAR\_PROTON\_TRANSPORTING\_V\_TYPE\_ATPASE\_COMPLEX(GO:0016471) |  | 18 | 0.57 | 0.95 | 0.307 | 0.470 | 1.000 | 23371 | tags=67%, list=40%, signal=110% |
| 6892 | ACTIVATION\_OF\_PROTEIN\_KINASE\_A\_ACTIVITY(GO:0034199) |  | 17 | 0.55 | 0.95 | 0.476 | 0.471 | 1.000 | 25845 | tags=71%, list=44%, signal=125% |
| 6893 | VESICLE\_TARGETING(GO:0006903) |  | 85 | 0.55 | 0.95 | 0.476 | 0.471 | 1.000 | 21115 | tags=62%, list=36%, signal=97% |
| 6894 | CELLULAR\_RESPONSE\_TO\_GAMMA\_RADIATION(GO:0071480) |  | 28 | 0.58 | 0.95 | 0.141 | 0.471 | 1.000 | 15941 | tags=46%, list=27%, signal=64% |
| 6895 | ENDOLYSOSOME(GO:0036019) |  | 20 | 0.55 | 0.95 | 0.476 | 0.472 | 1.000 | 26061 | tags=80%, list=44%, signal=143% |
| 6896 | PROTEASOME\_ACCESSORY\_COMPLEX(GO:0022624) |  | 25 | 0.56 | 0.95 | 0.476 | 0.472 | 1.000 | 26052 | tags=92%, list=44%, signal=164% |
| 6897 | NEGATIVE\_REGULATION\_OF\_VIRAL\_RELEASE\_FROM\_HOST\_CELL(GO:1902187) |  | 15 | 0.57 | 0.95 | 0.476 | 0.472 | 1.000 | 21411 | tags=60%, list=36%, signal=94% |
| 6898 | CELLULAR\_RESPONSE\_TO\_LOW\_DENSITY\_LIPOPROTEIN\_PARTICLE\_STIMULUS(GO:0071404) |  | 20 | 0.55 | 0.95 | 0.307 | 0.472 | 1.000 | 24090 | tags=60%, list=41%, signal=101% |
| 6899 | LEUKOTRIENE\_METABOLIC\_PROCESS(GO:0006691) |  | 29 | 0.56 | 0.95 | 0.476 | 0.472 | 1.000 | 16927 | tags=34%, list=29%, signal=48% |
| 6900 | REGULATION\_OF\_TRANSCRIPTION\_BY\_RNA\_POLYMERASE\_III(GO:0006359) |  | 27 | 0.57 | 0.95 | 0.310 | 0.472 | 1.000 | 17723 | tags=56%, list=30%, signal=79% |
| 6901 | REGULATION\_OF\_PROTEIN\_TYROSINE\_KINASE\_ACTIVITY(GO:0061097) |  | 76 | 0.54 | 0.95 | 0.476 | 0.473 | 1.000 | 24541 | tags=68%, list=41%, signal=117% |
| 6902 | POSITIVE\_REGULATION\_OF\_PROTEIN\_TARGETING\_TO\_MEMBRANE(GO:0090314) |  | 27 | 0.57 | 0.95 | 0.476 | 0.473 | 1.000 | 22174 | tags=59%, list=37%, signal=95% |
| 6903 | REGULATION\_OF\_RNA\_POLYMERASE\_II\_TRANSCRIPTIONAL\_PREINITIATION\_COMPLEX\_ASSEMBLY(GO:0045898) |  | 15 | 0.58 | 0.95 | 0.141 | 0.473 | 1.000 | 22372 | tags=80%, list=38%, signal=129% |
| 6904 | NEGATIVE\_REGULATION\_OF\_LYASE\_ACTIVITY(GO:0051350) |  | 32 | 0.54 | 0.95 | 0.476 | 0.474 | 1.000 | 23770 | tags=44%, list=40%, signal=73% |
| 6905 | REGULATION\_OF\_HETEROTYPIC\_CELL\_CELL\_ADHESION(GO:0034114) |  | 21 | 0.59 | 0.95 | 0.307 | 0.474 | 1.000 | 19002 | tags=33%, list=32%, signal=49% |
| 6906 | CLATHRIN\_COAT(GO:0030118) |  | 50 | 0.55 | 0.95 | 0.307 | 0.474 | 1.000 | 26227 | tags=94%, list=44%, signal=169% |
| 6907 | HIGH\_DENSITY\_LIPOPROTEIN\_PARTICLE\_ASSEMBLY(GO:0034380) |  | 15 | 0.58 | 0.95 | 0.476 | 0.475 | 1.000 | 20361 | tags=60%, list=34%, signal=91% |
| 6908 | REGULATION\_OF\_HEMATOPOIETIC\_PROGENITOR\_CELL\_DIFFERENTIATION(GO:1901532) |  | 114 | 0.53 | 0.95 | 0.476 | 0.475 | 1.000 | 26113 | tags=69%, list=44%, signal=124% |
| 6909 | POSITIVE\_REGULATION\_OF\_PRI\_MIRNA\_TRANSCRIPTION\_BY\_RNA\_POLYMERASE\_II(GO:1902895) |  | 24 | 0.57 | 0.95 | 0.476 | 0.475 | 1.000 | 23773 | tags=71%, list=40%, signal=118% |
| 6910 | GLUTATHIONE\_BIOSYNTHETIC\_PROCESS(GO:0006750) |  | 15 | 0.57 | 0.95 | 0.476 | 0.476 | 1.000 | 21056 | tags=53%, list=36%, signal=83% |
| 6911 | CELLULAR\_PIGMENTATION(GO:0033059) |  | 52 | 0.57 | 0.95 | 0.141 | 0.476 | 1.000 | 21618 | tags=65%, list=37%, signal=103% |
| 6912 | ATPASE\_ACTIVITY\_\_COUPLED\_TO\_TRANSMEMBRANE\_MOVEMENT\_OF\_IONS\_\_ROTATIONAL\_MECHANISM(GO:0044769) |  | 34 | 0.55 | 0.95 | 0.476 | 0.476 | 1.000 | 23227 | tags=74%, list=39%, signal=121% |
| 6913 | TRAPP\_COMPLEX(GO:0030008) |  | 16 | 0.54 | 0.94 | 0.476 | 0.479 | 1.000 | 25083 | tags=69%, list=42%, signal=119% |
| 6914 | VESICLE\_CYTOSKELETAL\_TRAFFICKING(GO:0099518) |  | 45 | 0.58 | 0.94 | 0.141 | 0.480 | 1.000 | 21510 | tags=73%, list=36%, signal=115% |
| 6915 | GROWTH\_HORMONE\_RECEPTOR\_SIGNALING\_PATHWAY(GO:0060396) |  | 22 | 0.55 | 0.94 | 0.309 | 0.480 | 1.000 | 25118 | tags=64%, list=42%, signal=111% |
| 6916 | ESTROUS\_CYCLE(GO:0044849) |  | 16 | 0.55 | 0.94 | 0.476 | 0.481 | 1.000 | 23287 | tags=63%, list=39%, signal=103% |
| 6917 | DEATH\_RECEPTOR\_BINDING(GO:0005123) |  | 16 | 0.55 | 0.94 | 0.476 | 0.481 | 1.000 | 22606 | tags=63%, list=38%, signal=101% |
| 6918 | REGULATION\_OF\_P38MAPK\_CASCADE(GO:1900744) |  | 40 | 0.56 | 0.94 | 0.310 | 0.481 | 1.000 | 25645 | tags=73%, list=43%, signal=128% |
| 6919 | GLYCOPROTEIN\_CATABOLIC\_PROCESS(GO:0006516) |  | 20 | 0.53 | 0.94 | 0.648 | 0.481 | 1.000 | 26864 | tags=80%, list=45%, signal=147% |
| 6920 | REGULATION\_OF\_PROTEIN\_TARGETING\_TO\_MEMBRANE(GO:0090313) |  | 33 | 0.57 | 0.94 | 0.476 | 0.481 | 1.000 | 22812 | tags=61%, list=39%, signal=99% |
| 6921 | INSULIN\_LIKE\_GROWTH\_FACTOR\_BINDING(GO:0005520) |  | 28 | 0.59 | 0.94 | 0.476 | 0.482 | 1.000 | 19090 | tags=46%, list=32%, signal=69% |
| 6922 | MITOGEN\_ACTIVATED\_PROTEIN\_KINASE\_BINDING(GO:0051019) |  | 27 | 0.55 | 0.94 | 0.476 | 0.482 | 1.000 | 25114 | tags=85%, list=42%, signal=148% |
| 6923 | NEGATIVE\_REGULATION\_OF\_PEPTIDYL\_THREONINE\_PHOSPHORYLATION(GO:0010801) |  | 15 | 0.55 | 0.94 | 0.476 | 0.482 | 1.000 | 25100 | tags=80%, list=42%, signal=139% |
| 6924 | REGULATION\_OF\_WATER\_LOSS\_VIA\_SKIN(GO:0033561) |  | 21 | 0.59 | 0.94 | 0.476 | 0.482 | 1.000 | 22360 | tags=43%, list=38%, signal=69% |
| 6925 | REGULATION\_OF\_ANTIGEN\_RECEPTOR\_MEDIATED\_SIGNALING\_PATHWAY(GO:0050854) |  | 54 | 0.54 | 0.94 | 0.141 | 0.483 | 1.000 | 21983 | tags=50%, list=37%, signal=80% |
| 6926 | FATTY\_ACID\_BETA\_OXIDATION\_USING\_ACYL\_COA\_OXIDASE(GO:0033540) |  | 15 | 0.54 | 0.94 | 0.476 | 0.483 | 1.000 | 27198 | tags=87%, list=46%, signal=160% |
| 6927 | POSITIVE\_REGULATION\_OF\_ACTIN\_CYTOSKELETON\_REORGANIZATION(GO:2000251) |  | 18 | 0.54 | 0.94 | 0.476 | 0.484 | 1.000 | 26338 | tags=72%, list=45%, signal=130% |
| 6928 | MODIFIED\_AMINO\_ACID\_TRANSPORT(GO:0072337) |  | 17 | 0.56 | 0.94 | 0.476 | 0.485 | 1.000 | 24662 | tags=76%, list=42%, signal=131% |
| 6929 | MYOSIN\_V\_BINDING(GO:0031489) |  | 17 | 0.55 | 0.94 | 0.141 | 0.485 | 1.000 | 21645 | tags=71%, list=37%, signal=111% |
| 6930 | EPITHELIAL\_STRUCTURE\_MAINTENANCE(GO:0010669) |  | 24 | 0.59 | 0.94 | 0.476 | 0.486 | 1.000 | 14575 | tags=29%, list=25%, signal=39% |
| 6931 | REGULATION\_OF\_MEMBRANE\_PERMEABILITY(GO:0090559) |  | 87 | 0.55 | 0.94 | 0.476 | 0.486 | 1.000 | 22859 | tags=61%, list=39%, signal=99% |
| 6932 | INTERFERON\_ALPHA\_PRODUCTION(GO:0032607) |  | 29 | 0.56 | 0.94 | 0.309 | 0.487 | 1.000 | 23155 | tags=59%, list=39%, signal=96% |
| 6933 | POSITIVE\_REGULATION\_OF\_ENDOTHELIAL\_CELL\_DIFFERENTIATION(GO:0045603) |  | 21 | 0.57 | 0.94 | 0.476 | 0.487 | 1.000 | 24565 | tags=57%, list=42%, signal=98% |
| 6934 | REGULATION\_OF\_MITOCHONDRIAL\_MEMBRANE\_PERMEABILITY(GO:0046902) |  | 79 | 0.56 | 0.94 | 0.476 | 0.488 | 1.000 | 22859 | tags=62%, list=39%, signal=101% |
| 6935 | POSITIVE\_REGULATION\_OF\_P38MAPK\_CASCADE(GO:1900745) |  | 26 | 0.57 | 0.94 | 0.310 | 0.488 | 1.000 | 17567 | tags=42%, list=30%, signal=60% |
| 6936 | REGULATION\_OF\_CYSTEINE\_TYPE\_ENDOPEPTIDASE\_ACTIVITY\_INVOLVED\_IN\_APOPTOTIC\_SIGNALING\_PATHWAY(GO:2001267) |  | 23 | 0.55 | 0.94 | 0.476 | 0.488 | 1.000 | 24018 | tags=65%, list=41%, signal=110% |
| 6937 | REGULATION\_OF\_HEMATOPOIETIC\_STEM\_CELL\_DIFFERENTIATION(GO:1902036) |  | 104 | 0.53 | 0.94 | 0.476 | 0.491 | 1.000 | 26113 | tags=69%, list=44%, signal=124% |
| 6938 | POSITIVE\_REGULATION\_OF\_REGULATED\_SECRETORY\_PATHWAY(GO:1903307) |  | 46 | 0.55 | 0.94 | 0.476 | 0.491 | 1.000 | 24308 | tags=63%, list=41%, signal=107% |
| 6939 | APICAL\_DENDRITE(GO:0097440) |  | 15 | 0.58 | 0.94 | 0.476 | 0.491 | 1.000 | 21613 | tags=60%, list=37%, signal=95% |
| 6940 | MEMBRANE\_PROTEIN\_INTRACELLULAR\_DOMAIN\_PROTEOLYSIS(GO:0031293) |  | 24 | 0.53 | 0.94 | 0.476 | 0.492 | 1.000 | 27295 | tags=88%, list=46%, signal=162% |
| 6941 | REGULATION\_OF\_SYNCYTIUM\_FORMATION\_BY\_PLASMA\_MEMBRANE\_FUSION(GO:0060142) |  | 26 | 0.55 | 0.94 | 0.476 | 0.492 | 1.000 | 25572 | tags=62%, list=43%, signal=108% |
| 6942 | ENDOPLASMIC\_RETICULUM\_CALCIUM\_ION\_HOMEOSTASIS(GO:0032469) |  | 23 | 0.54 | 0.94 | 0.476 | 0.492 | 1.000 | 27354 | tags=100%, list=46%, signal=186% |
| 6943 | PROTON\_TRANSPORTING\_V\_TYPE\_ATPASE\_COMPLEX(GO:0033176) |  | 27 | 0.55 | 0.94 | 0.307 | 0.493 | 1.000 | 24001 | tags=70%, list=41%, signal=118% |
| 6944 | NADH\_METABOLIC\_PROCESS(GO:0006734) |  | 39 | 0.54 | 0.94 | 0.476 | 0.493 | 1.000 | 26205 | tags=90%, list=44%, signal=161% |
| 6945 | LEUKOCYTE\_ADHESION\_TO\_VASCULAR\_ENDOTHELIAL\_CELL(GO:0061756) |  | 36 | 0.54 | 0.94 | 0.309 | 0.494 | 1.000 | 24352 | tags=44%, list=41%, signal=76% |
| 6946 | NEGATIVE\_REGULATION\_OF\_ATP\_BIOSYNTHETIC\_PROCESS(GO:2001170) |  | 15 | 0.59 | 0.94 | 0.476 | 0.494 | 1.000 | 23468 | tags=73%, list=40%, signal=122% |
| 6947 | LIPOPOLYSACCHARIDE\_BINDING(GO:0001530) |  | 28 | 0.55 | 0.94 | 0.476 | 0.494 | 1.000 | 25572 | tags=54%, list=43%, signal=94% |
| 6948 | TRANSFERRIN\_TRANSPORT(GO:0033572) |  | 36 | 0.54 | 0.94 | 0.476 | 0.495 | 1.000 | 26692 | tags=83%, list=45%, signal=152% |
| 6949 | RESPONSE\_TO\_GROWTH\_HORMONE(GO:0060416) |  | 37 | 0.55 | 0.94 | 0.650 | 0.495 | 1.000 | 25386 | tags=65%, list=43%, signal=114% |
| 6950 | REGULATION\_OF\_LIPID\_STORAGE(GO:0010883) |  | 42 | 0.53 | 0.94 | 0.476 | 0.495 | 1.000 | 23316 | tags=45%, list=39%, signal=75% |
| 6951 | MALE\_GERM\_CELL\_NUCLEUS(GO:0001673) |  | 15 | 0.60 | 0.94 | 0.481 | 0.496 | 1.000 | 16861 | tags=47%, list=29%, signal=65% |
| 6952 | PROTEIN\_CONTAINING\_COMPLEX\_SCAFFOLD\_ACTIVITY(GO:0032947) |  | 71 | 0.57 | 0.94 | 0.141 | 0.497 | 1.000 | 24746 | tags=80%, list=42%, signal=138% |
| 6953 | CLATHRIN\_ADAPTOR\_COMPLEX(GO:0030131) |  | 31 | 0.55 | 0.94 | 0.476 | 0.498 | 1.000 | 26227 | tags=94%, list=44%, signal=168% |
| 6954 | POST\_EMBRYONIC\_ANIMAL\_ORGAN\_DEVELOPMENT(GO:0048569) |  | 17 | 0.55 | 0.94 | 0.476 | 0.499 | 1.000 | 24344 | tags=71%, list=41%, signal=120% |
| 6955 | THYMIC\_T\_CELL\_SELECTION(GO:0045061) |  | 19 | 0.58 | 0.94 | 0.476 | 0.499 | 1.000 | 17052 | tags=21%, list=29%, signal=30% |
| 6956 | REGULATION\_OF\_NECROPTOTIC\_PROCESS(GO:0060544) |  | 20 | 0.54 | 0.94 | 0.476 | 0.499 | 1.000 | 24124 | tags=70%, list=41%, signal=118% |
| 6957 | REGULATION\_OF\_TOLL\_LIKE\_RECEPTOR\_4\_SIGNALING\_PATHWAY(GO:0034143) |  | 18 | 0.54 | 0.94 | 0.476 | 0.499 | 1.000 | 26267 | tags=72%, list=44%, signal=130% |
| 6958 | PIGMENT\_GRANULE\_ORGANIZATION(GO:0048753) |  | 27 | 0.57 | 0.94 | 0.141 | 0.499 | 1.000 | 24245 | tags=81%, list=41%, signal=138% |
| 6959 | LOW\_DENSITY\_LIPOPROTEIN\_PARTICLE\_RECEPTOR\_BINDING(GO:0050750) |  | 21 | 0.54 | 0.94 | 0.476 | 0.499 | 1.000 | 26061 | tags=76%, list=44%, signal=136% |
| 6960 | AZUROPHIL\_GRANULE\_MEMBRANE(GO:0035577) |  | 56 | 0.53 | 0.94 | 0.307 | 0.500 | 1.000 | 26361 | tags=73%, list=45%, signal=132% |
| 6961 | REGULATION\_OF\_TRIGLYCERIDE\_METABOLIC\_PROCESS(GO:0090207) |  | 34 | 0.55 | 0.94 | 0.476 | 0.500 | 1.000 | 22269 | tags=44%, list=38%, signal=71% |
| 6962 | REGULATION\_OF\_MEGAKARYOCYTE\_DIFFERENTIATION(GO:0045652) |  | 76 | 0.54 | 0.94 | 0.476 | 0.502 | 1.000 | 20771 | tags=47%, list=35%, signal=73% |
| 6963 | POSITIVE\_REGULATION\_OF\_NUCLEASE\_ACTIVITY(GO:0032075) |  | 16 | 0.53 | 0.94 | 0.476 | 0.503 | 1.000 | 26020 | tags=81%, list=44%, signal=145% |
| 6964 | MELANOSOME\_ORGANIZATION(GO:0032438) |  | 26 | 0.57 | 0.94 | 0.141 | 0.504 | 1.000 | 24245 | tags=81%, list=41%, signal=137% |
| 6965 | MELANOSOME(GO:0042470) |  | 105 | 0.55 | 0.93 | 0.141 | 0.505 | 1.000 | 25370 | tags=81%, list=43%, signal=142% |
| 6966 | PIGMENT\_GRANULE(GO:0048770) |  | 105 | 0.55 | 0.93 | 0.141 | 0.505 | 1.000 | 25370 | tags=81%, list=43%, signal=142% |
| 6967 | PROGESTERONE\_METABOLIC\_PROCESS(GO:0042448) |  | 16 | 0.58 | 0.93 | 0.476 | 0.505 | 1.000 | 23890 | tags=69%, list=40%, signal=115% |
| 6968 | MITOGEN\_ACTIVATED\_PROTEIN\_KINASE\_KINASE\_BINDING(GO:0031434) |  | 16 | 0.55 | 0.93 | 0.476 | 0.506 | 1.000 | 26225 | tags=94%, list=44%, signal=168% |
| 6969 | CELL\_DIFFERENTIATION\_INVOLVED\_IN\_EMBRYONIC\_PLACENTA\_DEVELOPMENT(GO:0060706) |  | 26 | 0.56 | 0.93 | 0.476 | 0.506 | 1.000 | 18947 | tags=42%, list=32%, signal=62% |
| 6970 | PEPTIDE\_ANTIGEN\_BINDING(GO:0042605) |  | 22 | 0.58 | 0.93 | 0.476 | 0.506 | 1.000 | 8063 | tags=23%, list=14%, signal=26% |
| 6971 | INTESTINAL\_ABSORPTION(GO:0050892) |  | 35 | 0.53 | 0.93 | 0.476 | 0.507 | 1.000 | 27347 | tags=71%, list=46%, signal=133% |
| 6972 | NOTCH\_RECEPTOR\_PROCESSING(GO:0007220) |  | 18 | 0.54 | 0.93 | 0.476 | 0.508 | 1.000 | 18596 | tags=44%, list=31%, signal=65% |
| 6973 | NATURAL\_KILLER\_CELL\_ACTIVATION\_INVOLVED\_IN\_IMMUNE\_RESPONSE(GO:0002323) |  | 30 | 0.55 | 0.93 | 0.476 | 0.510 | 1.000 | 18052 | tags=33%, list=31%, signal=48% |
| 6974 | RAC\_GUANYL\_NUCLEOTIDE\_EXCHANGE\_FACTOR\_ACTIVITY(GO:0030676) |  | 15 | 0.56 | 0.93 | 0.476 | 0.510 | 1.000 | 20862 | tags=53%, list=35%, signal=82% |
| 6975 | PROTON\_EXPORTING\_ATPASE\_ACTIVITY(GO:0036442) |  | 23 | 0.55 | 0.93 | 0.307 | 0.510 | 1.000 | 24001 | tags=74%, list=41%, signal=124% |
| 6976 | PEPTIDYL\_THREONINE\_DEPHOSPHORYLATION(GO:0035970) |  | 16 | 0.56 | 0.93 | 0.816 | 0.511 | 1.000 | 23764 | tags=81%, list=40%, signal=136% |
| 6977 | JAK\_STAT\_CASCADE\_INVOLVED\_IN\_GROWTH\_HORMONE\_SIGNALING\_PATHWAY(GO:0060397) |  | 15 | 0.57 | 0.93 | 0.644 | 0.511 | 1.000 | 25118 | tags=60%, list=42%, signal=104% |
| 6978 | PROTON\_TRANSPORTING\_ATPASE\_ACTIVITY\_\_ROTATIONAL\_MECHANISM(GO:0046961) |  | 21 | 0.55 | 0.93 | 0.476 | 0.513 | 1.000 | 24001 | tags=76%, list=41%, signal=128% |
| 6979 | P38MAPK\_CASCADE(GO:0038066) |  | 46 | 0.55 | 0.93 | 0.310 | 0.514 | 1.000 | 25645 | tags=70%, list=43%, signal=123% |
| 6980 | REGULATION\_OF\_SKELETAL\_MUSCLE\_ADAPTATION(GO:0014733) |  | 15 | 0.56 | 0.93 | 0.476 | 0.516 | 1.000 | 23500 | tags=53%, list=40%, signal=88% |
| 6981 | C\_ACYLTRANSFERASE\_ACTIVITY(GO:0016408) |  | 20 | 0.59 | 0.93 | 0.481 | 0.517 | 1.000 | 21873 | tags=50%, list=37%, signal=79% |
| 6982 | ECTODERM\_DEVELOPMENT(GO:0007398) |  | 22 | 0.56 | 0.93 | 0.476 | 0.518 | 1.000 | 20133 | tags=36%, list=34%, signal=55% |
| 6983 | NEGATIVE\_REGULATION\_OF\_COFACTOR\_METABOLIC\_PROCESS(GO:0051195) |  | 18 | 0.59 | 0.93 | 0.476 | 0.518 | 1.000 | 23468 | tags=61%, list=40%, signal=101% |
| 6984 | POSITIVE\_REGULATION\_OF\_SUBSTRATE\_ADHESION\_DEPENDENT\_CELL\_SPREADING(GO:1900026) |  | 34 | 0.54 | 0.93 | 0.476 | 0.519 | 1.000 | 25800 | tags=74%, list=44%, signal=130% |
| 6985 | POSITIVE\_REGULATION\_OF\_RESPONSE\_TO\_ENDOPLASMIC\_RETICULUM\_STRESS(GO:1905898) |  | 37 | 0.55 | 0.93 | 0.476 | 0.519 | 1.000 | 22919 | tags=65%, list=39%, signal=106% |
| 6986 | MORPHOGENESIS\_OF\_AN\_ENDOTHELIUM(GO:0003159) |  | 16 | 0.54 | 0.93 | 0.476 | 0.521 | 1.000 | 27295 | tags=88%, list=46%, signal=162% |
| 6987 | ENDOTHELIAL\_TUBE\_MORPHOGENESIS(GO:0061154) |  | 16 | 0.54 | 0.93 | 0.476 | 0.521 | 1.000 | 27295 | tags=88%, list=46%, signal=162% |
| 6988 | CARBOHYDRATE\_PHOSPHORYLATION(GO:0046835) |  | 24 | 0.55 | 0.93 | 0.476 | 0.521 | 1.000 | 26672 | tags=96%, list=45%, signal=174% |
| 6989 | CELLULAR\_COPPER\_ION\_HOMEOSTASIS(GO:0006878) |  | 15 | 0.54 | 0.93 | 0.644 | 0.521 | 1.000 | 26713 | tags=87%, list=45%, signal=158% |
| 6990 | MHC\_PROTEIN\_COMPLEX(GO:0042611) |  | 24 | 0.53 | 0.93 | 0.644 | 0.521 | 1.000 | 18589 | tags=42%, list=31%, signal=61% |
| 6991 | INTERLEUKIN\_1\_RECEPTOR\_BINDING(GO:0005149) |  | 19 | 0.56 | 0.93 | 0.475 | 0.522 | 1.000 | 23136 | tags=53%, list=39%, signal=86% |
| 6992 | REGULATION\_OF\_INTERFERON\_ALPHA\_PRODUCTION(GO:0032647) |  | 28 | 0.56 | 0.93 | 0.481 | 0.522 | 1.000 | 23155 | tags=57%, list=39%, signal=94% |
| 6993 | GLYCERALDEHYDE\_3\_PHOSPHATE\_METABOLIC\_PROCESS(GO:0019682) |  | 24 | 0.55 | 0.93 | 0.476 | 0.524 | 1.000 | 25953 | tags=88%, list=44%, signal=156% |
| 6994 | BETA\_CATENIN\_DESTRUCTION\_COMPLEX\_DISASSEMBLY(GO:1904886) |  | 22 | 0.53 | 0.93 | 0.476 | 0.526 | 1.000 | 24590 | tags=73%, list=42%, signal=124% |
| 6995 | ENTRY\_OF\_BACTERIUM\_INTO\_HOST\_CELL(GO:0035635) |  | 19 | 0.55 | 0.93 | 0.476 | 0.528 | 1.000 | 24841 | tags=74%, list=42%, signal=127% |
| 6996 | POSITIVE\_REGULATION\_OF\_SYNCYTIUM\_FORMATION\_BY\_PLASMA\_MEMBRANE\_FUSION(GO:0060143) |  | 24 | 0.55 | 0.93 | 0.476 | 0.530 | 1.000 | 25572 | tags=63%, list=43%, signal=110% |
| 6997 | REGULATION\_OF\_ENDOTHELIAL\_CELL\_CHEMOTAXIS(GO:2001026) |  | 22 | 0.57 | 0.93 | 0.476 | 0.530 | 1.000 | 15720 | tags=27%, list=27%, signal=37% |
| 6998 | B\_CELL\_APOPTOTIC\_PROCESS(GO:0001783) |  | 23 | 0.55 | 0.93 | 0.648 | 0.531 | 1.000 | 25019 | tags=70%, list=42%, signal=121% |
| 6999 | ER\_NUCLEUS\_SIGNALING\_PATHWAY(GO:0006984) |  | 44 | 0.57 | 0.93 | 0.481 | 0.531 | 1.000 | 22919 | tags=73%, list=39%, signal=119% |
| 7000 | NEGATIVE\_REGULATION\_OF\_PROTEIN\_KINASE\_B\_SIGNALING(GO:0051898) |  | 57 | 0.52 | 0.93 | 0.476 | 0.533 | 1.000 | 23270 | tags=46%, list=39%, signal=75% |
| 7001 | GLUCOSE\_CATABOLIC\_PROCESS(GO:0006007) |  | 32 | 0.53 | 0.93 | 0.476 | 0.535 | 1.000 | 27189 | tags=91%, list=46%, signal=168% |
| 7002 | MHC\_CLASS\_II\_PROTEIN\_COMPLEX\_BINDING(GO:0023026) |  | 16 | 0.53 | 0.92 | 0.816 | 0.536 | 1.000 | 27478 | tags=81%, list=46%, signal=152% |
| 7003 | POSITIVE\_REGULATION\_OF\_CYSTEINE\_TYPE\_ENDOPEPTIDASE\_ACTIVITY\_INVOLVED\_IN\_APOPTOTIC\_SIGNALING\_PATHWAY(GO:2001269) |  | 18 | 0.54 | 0.92 | 0.476 | 0.536 | 1.000 | 24018 | tags=61%, list=41%, signal=103% |
| 7004 | NUCLEOSOME(GO:0000786) |  | 109 | 0.53 | 0.92 | 0.476 | 0.538 | 1.000 | 20540 | tags=26%, list=35%, signal=39% |
| 7005 | POSITIVE\_REGULATION\_OF\_PROTEIN\_PROCESSING(GO:0010954) |  | 23 | 0.56 | 0.92 | 0.476 | 0.539 | 1.000 | 17506 | tags=43%, list=30%, signal=62% |
| 7006 | NUCLEAR\_NUCLEOSOME(GO:0000788) |  | 40 | 0.53 | 0.92 | 0.475 | 0.539 | 1.000 | 19761 | tags=23%, list=33%, signal=34% |
| 7007 | INTRINSIC\_COMPONENT\_OF\_PEROXISOMAL\_MEMBRANE(GO:0031231) |  | 17 | 0.54 | 0.92 | 0.816 | 0.540 | 1.000 | 26587 | tags=94%, list=45%, signal=171% |
| 7008 | INTEGRAL\_COMPONENT\_OF\_PEROXISOMAL\_MEMBRANE(GO:0005779) |  | 17 | 0.54 | 0.92 | 0.816 | 0.540 | 1.000 | 26587 | tags=94%, list=45%, signal=171% |
| 7009 | PHOSPHOTYROSINE\_RESIDUE\_BINDING(GO:0001784) |  | 35 | 0.53 | 0.92 | 0.476 | 0.540 | 1.000 | 26758 | tags=74%, list=45%, signal=136% |
| 7010 | MIRNA\_MEDIATED\_INHIBITION\_OF\_TRANSLATION(GO:0035278) |  | 67 | 0.54 | 0.92 | 0.648 | 0.543 | 1.000 | 24391 | tags=30%, list=41%, signal=51% |
| 7011 | NEGATIVE\_REGULATION\_OF\_TRANSLATION\_\_NCRNA\_MEDIATED(GO:0040033) |  | 67 | 0.54 | 0.92 | 0.648 | 0.542 | 1.000 | 24391 | tags=30%, list=41%, signal=51% |
| 7012 | REGULATION\_OF\_TRANSLATION\_\_NCRNA\_MEDIATED(GO:0045974) |  | 67 | 0.54 | 0.92 | 0.648 | 0.542 | 1.000 | 24391 | tags=30%, list=41%, signal=51% |
| 7013 | INTEGRAL\_COMPONENT\_OF\_LUMENAL\_SIDE\_OF\_ENDOPLASMIC\_RETICULUM\_MEMBRANE(GO:0071556) |  | 24 | 0.54 | 0.92 | 0.648 | 0.543 | 1.000 | 26829 | tags=67%, list=45%, signal=122% |
| 7014 | LUMENAL\_SIDE\_OF\_ENDOPLASMIC\_RETICULUM\_MEMBRANE(GO:0098553) |  | 24 | 0.54 | 0.92 | 0.648 | 0.543 | 1.000 | 26829 | tags=67%, list=45%, signal=122% |
| 7015 | RESPIRATORY\_CHAIN\_COMPLEX\_IV(GO:0045277) |  | 21 | 0.54 | 0.92 | 0.816 | 0.544 | 1.000 | 27259 | tags=81%, list=46%, signal=150% |
| 7016 | PARTURITION(GO:0007567) |  | 17 | 0.54 | 0.92 | 0.307 | 0.545 | 1.000 | 22962 | tags=47%, list=39%, signal=77% |
| 7017 | POSITIVE\_REGULATION\_OF\_ERBB\_SIGNALING\_PATHWAY(GO:1901186) |  | 39 | 0.53 | 0.92 | 0.476 | 0.551 | 1.000 | 26548 | tags=77%, list=45%, signal=139% |
| 7018 | REGULATION\_OF\_T\_CELL\_RECEPTOR\_SIGNALING\_PATHWAY(GO:0050856) |  | 37 | 0.53 | 0.92 | 0.307 | 0.551 | 1.000 | 24797 | tags=57%, list=42%, signal=98% |
| 7019 | RUFFLE\_ORGANIZATION(GO:0031529) |  | 47 | 0.53 | 0.92 | 0.476 | 0.555 | 1.000 | 27266 | tags=81%, list=46%, signal=150% |
| 7020 | HYALURONAN\_CATABOLIC\_PROCESS(GO:0030214) |  | 16 | 0.54 | 0.92 | 0.476 | 0.555 | 1.000 | 24451 | tags=69%, list=41%, signal=117% |
| 7021 | POSITIVE\_REGULATION\_OF\_MONONUCLEAR\_CELL\_MIGRATION(GO:0071677) |  | 21 | 0.55 | 0.92 | 0.481 | 0.557 | 1.000 | 24932 | tags=38%, list=42%, signal=66% |
| 7022 | POSITIVE\_REGULATION\_OF\_PROTEIN\_TYROSINE\_KINASE\_ACTIVITY(GO:0061098) |  | 45 | 0.55 | 0.92 | 0.476 | 0.557 | 1.000 | 24541 | tags=67%, list=41%, signal=114% |
| 7023 | DEFENSE\_RESPONSE\_TO\_FUNGUS(GO:0050832) |  | 40 | 0.56 | 0.92 | 0.476 | 0.559 | 1.000 | 24257 | tags=30%, list=41%, signal=51% |
| 7024 | NEGATIVE\_REGULATION\_OF\_CELL\_MIGRATION\_INVOLVED\_IN\_SPROUTING\_ANGIOGENESIS(GO:0090051) |  | 42 | 0.55 | 0.92 | 0.476 | 0.559 | 1.000 | 19002 | tags=31%, list=32%, signal=46% |
| 7025 | NEGATIVE\_REGULATION\_OF\_INTERLEUKIN\_2\_PRODUCTION(GO:0032703) |  | 19 | 0.56 | 0.92 | 0.476 | 0.561 | 1.000 | 24979 | tags=74%, list=42%, signal=128% |
| 7026 | POSITIVE\_REGULATION\_OF\_PROTEIN\_MATURATION(GO:1903319) |  | 25 | 0.56 | 0.91 | 0.476 | 0.570 | 1.000 | 21992 | tags=52%, list=37%, signal=83% |
| 7027 | PROTEIN\_HETEROOLIGOMERIZATION(GO:0051291) |  | 125 | 0.53 | 0.91 | 0.475 | 0.571 | 1.000 | 23923 | tags=55%, list=40%, signal=93% |
| 7028 | FAD\_BINDING(GO:0071949) |  | 22 | 0.52 | 0.91 | 0.476 | 0.572 | 1.000 | 26436 | tags=73%, list=45%, signal=131% |
| 7029 | PINOCYTOSIS(GO:0006907) |  | 19 | 0.55 | 0.91 | 0.476 | 0.580 | 1.000 | 24266 | tags=63%, list=41%, signal=107% |
| 7030 | RUFFLE\_ASSEMBLY(GO:0097178) |  | 36 | 0.54 | 0.91 | 0.476 | 0.580 | 1.000 | 27266 | tags=83%, list=46%, signal=155% |
| 7031 | REGULATION\_OF\_MYOBLAST\_FUSION(GO:1901739) |  | 20 | 0.55 | 0.91 | 0.648 | 0.581 | 1.000 | 26447 | tags=55%, list=45%, signal=99% |
| 7032 | HEXOSE\_CATABOLIC\_PROCESS(GO:0019320) |  | 52 | 0.55 | 0.91 | 0.476 | 0.586 | 1.000 | 26251 | tags=87%, list=44%, signal=155% |
| 7033 | ARF\_GUANYL\_NUCLEOTIDE\_EXCHANGE\_FACTOR\_ACTIVITY(GO:0005086) |  | 21 | 0.54 | 0.91 | 0.476 | 0.587 | 1.000 | 26559 | tags=76%, list=45%, signal=138% |
| 7034 | REGULATION\_OF\_RUFFLE\_ASSEMBLY(GO:1900027) |  | 26 | 0.54 | 0.91 | 0.476 | 0.587 | 1.000 | 27266 | tags=81%, list=46%, signal=150% |
| 7035 | GLUCOSE\_6\_PHOSPHATE\_METABOLIC\_PROCESS(GO:0051156) |  | 28 | 0.54 | 0.91 | 0.476 | 0.588 | 1.000 | 26808 | tags=82%, list=45%, signal=150% |
| 7036 | POSITIVE\_REGULATION\_OF\_MYOBLAST\_FUSION(GO:1901741) |  | 18 | 0.55 | 0.91 | 0.476 | 0.589 | 1.000 | 26447 | tags=56%, list=45%, signal=100% |
| 7037 | REGULATION\_OF\_ENDOPLASMIC\_RETICULUM\_UNFOLDED\_PROTEIN\_RESPONSE(GO:1900101) |  | 28 | 0.52 | 0.91 | 0.476 | 0.591 | 1.000 | 27813 | tags=86%, list=47%, signal=162% |
| 7038 | POSITIVE\_REGULATION\_OF\_INTERFERON\_ALPHA\_PRODUCTION(GO:0032727) |  | 22 | 0.55 | 0.91 | 0.481 | 0.592 | 1.000 | 24269 | tags=64%, list=41%, signal=108% |
| 7039 | POSITIVE\_REGULATION\_OF\_EPIDERMAL\_GROWTH\_FACTOR\_RECEPTOR\_SIGNALING\_PATHWAY(GO:0045742) |  | 37 | 0.53 | 0.91 | 0.476 | 0.592 | 1.000 | 26548 | tags=76%, list=45%, signal=137% |
| 7040 | NEGATIVE\_REGULATION\_OF\_MEGAKARYOCYTE\_DIFFERENTIATION(GO:0045653) |  | 18 | 0.51 | 0.91 | 0.816 | 0.592 | 1.000 | 28357 | tags=78%, list=48%, signal=149% |
| 7041 | MITOCHONDRIAL\_RESPIRATORY\_CHAIN\_COMPLEX\_IV(GO:0005751) |  | 18 | 0.54 | 0.91 | 0.816 | 0.594 | 1.000 | 27259 | tags=83%, list=46%, signal=155% |
| 7042 | REGULATION\_OF\_IRE1\_MEDIATED\_UNFOLDED\_PROTEIN\_RESPONSE(GO:1903894) |  | 15 | 0.54 | 0.91 | 0.816 | 0.595 | 1.000 | 27433 | tags=87%, list=46%, signal=162% |
| 7043 | PERK\_MEDIATED\_UNFOLDED\_PROTEIN\_RESPONSE(GO:0036499) |  | 21 | 0.57 | 0.91 | 0.481 | 0.596 | 1.000 | 22919 | tags=67%, list=39%, signal=109% |
| 7044 | RESPONSE\_TO\_HEPATOCYTE\_GROWTH\_FACTOR(GO:0035728) |  | 16 | 0.54 | 0.91 | 0.648 | 0.596 | 1.000 | 23773 | tags=50%, list=40%, signal=84% |
| 7045 | 14\_3\_3\_PROTEIN\_BINDING(GO:0071889) |  | 26 | 0.53 | 0.91 | 0.476 | 0.599 | 1.000 | 25645 | tags=73%, list=43%, signal=129% |
| 7046 | NEGATIVE\_REGULATION\_OF\_T\_CELL\_RECEPTOR\_SIGNALING\_PATHWAY(GO:0050860) |  | 19 | 0.53 | 0.91 | 0.481 | 0.602 | 1.000 | 27552 | tags=74%, list=47%, signal=138% |
| 7047 | POSITIVE\_REGULATION\_OF\_MONOCYTE\_CHEMOTAXIS(GO:0090026) |  | 17 | 0.56 | 0.91 | 0.481 | 0.603 | 1.000 | 24932 | tags=41%, list=42%, signal=71% |
| 7048 | MONOSACCHARIDE\_CATABOLIC\_PROCESS(GO:0046365) |  | 62 | 0.54 | 0.90 | 0.476 | 0.606 | 1.000 | 26251 | tags=84%, list=44%, signal=151% |
| 7049 | LIPID\_DIGESTION(GO:0044241) |  | 15 | 0.55 | 0.90 | 0.476 | 0.608 | 1.000 | 24352 | tags=53%, list=41%, signal=91% |
| 7050 | NEGATIVE\_REGULATION\_OF\_ANTIGEN\_RECEPTOR\_MEDIATED\_SIGNALING\_PATHWAY(GO:0050858) |  | 23 | 0.53 | 0.90 | 0.481 | 0.608 | 1.000 | 27552 | tags=70%, list=47%, signal=130% |
| 7051 | MAINTENANCE\_OF\_GASTROINTESTINAL\_EPITHELIUM(GO:0030277) |  | 17 | 0.58 | 0.90 | 0.644 | 0.618 | 1.000 | 14575 | tags=35%, list=25%, signal=47% |
| 7052 | N\_GLYCAN\_PROCESSING(GO:0006491) |  | 19 | 0.53 | 0.90 | 0.481 | 0.619 | 1.000 | 27037 | tags=89%, list=46%, signal=165% |
| 7053 | REGULATION\_OF\_ACTIN\_CYTOSKELETON\_REORGANIZATION(GO:2000249) |  | 35 | 0.54 | 0.90 | 0.476 | 0.619 | 1.000 | 24066 | tags=60%, list=41%, signal=101% |
| 7054 | NEGATIVE\_REGULATION\_OF\_COLLAGEN\_METABOLIC\_PROCESS(GO:0010713) |  | 15 | 0.52 | 0.90 | 0.647 | 0.620 | 1.000 | 27490 | tags=60%, list=46%, signal=112% |
| 7055 | ENDODERMAL\_CELL\_FATE\_COMMITMENT(GO:0001711) |  | 16 | 0.54 | 0.90 | 0.816 | 0.624 | 1.000 | 26767 | tags=75%, list=45%, signal=137% |
| 7056 | CARBOHYDRATE\_KINASE\_ACTIVITY(GO:0019200) |  | 21 | 0.53 | 0.90 | 0.476 | 0.625 | 1.000 | 27021 | tags=90%, list=46%, signal=167% |
| 7057 | NEGATIVE\_REGULATION\_OF\_SPROUTING\_ANGIOGENESIS(GO:1903671) |  | 75 | 0.53 | 0.90 | 0.476 | 0.630 | 1.000 | 20939 | tags=25%, list=35%, signal=39% |
| 7058 | REGULATION\_OF\_PRODUCTION\_OF\_SMALL\_RNA\_INVOLVED\_IN\_GENE\_SILENCING\_BY\_RNA(GO:0070920) |  | 16 | 0.55 | 0.90 | 0.476 | 0.632 | 1.000 | 23586 | tags=63%, list=40%, signal=104% |
| 7059 | REGULATION\_OF\_ARF\_PROTEIN\_SIGNAL\_TRANSDUCTION(GO:0032012) |  | 17 | 0.54 | 0.90 | 0.476 | 0.637 | 1.000 | 26559 | tags=76%, list=45%, signal=139% |
| 7060 | ARF\_PROTEIN\_SIGNAL\_TRANSDUCTION(GO:0032011) |  | 17 | 0.54 | 0.90 | 0.476 | 0.636 | 1.000 | 26559 | tags=76%, list=45%, signal=139% |
| 7061 | TOLL\_LIKE\_RECEPTOR\_4\_SIGNALING\_PATHWAY(GO:0034142) |  | 32 | 0.54 | 0.90 | 0.476 | 0.638 | 1.000 | 26267 | tags=66%, list=44%, signal=118% |
| 7062 | DNA\_REPLICATION\_DEPENDENT\_NUCLEOSOME\_ORGANIZATION(GO:0034723) |  | 32 | 0.52 | 0.90 | 0.816 | 0.638 | 1.000 | 27100 | tags=59%, list=46%, signal=110% |
| 7063 | DNA\_REPLICATION\_DEPENDENT\_NUCLEOSOME\_ASSEMBLY(GO:0006335) |  | 32 | 0.52 | 0.90 | 0.816 | 0.638 | 1.000 | 27100 | tags=59%, list=46%, signal=110% |
| 7064 | GLUCOSE\_CATABOLIC\_PROCESS\_TO\_PYRUVATE(GO:0061718) |  | 26 | 0.54 | 0.90 | 0.476 | 0.638 | 1.000 | 26205 | tags=85%, list=44%, signal=152% |
| 7065 | CANONICAL\_GLYCOLYSIS(GO:0061621) |  | 26 | 0.54 | 0.90 | 0.476 | 0.638 | 1.000 | 26205 | tags=85%, list=44%, signal=152% |
| 7066 | NADH\_REGENERATION(GO:0006735) |  | 26 | 0.54 | 0.90 | 0.476 | 0.638 | 1.000 | 26205 | tags=85%, list=44%, signal=152% |
| 7067 | GLYCOLYTIC\_PROCESS\_THROUGH\_FRUCTOSE\_6\_PHOSPHATE(GO:0061615) |  | 27 | 0.53 | 0.89 | 0.476 | 0.651 | 1.000 | 26243 | tags=85%, list=44%, signal=153% |
| 7068 | GLYCOLYTIC\_PROCESS\_THROUGH\_GLUCOSE\_6\_PHOSPHATE(GO:0061620) |  | 27 | 0.53 | 0.89 | 0.476 | 0.651 | 1.000 | 26243 | tags=85%, list=44%, signal=153% |
| 7069 | CCR\_CHEMOKINE\_RECEPTOR\_BINDING(GO:0048020) |  | 41 | 0.54 | 0.89 | 0.648 | 0.651 | 1.000 | 23470 | tags=24%, list=40%, signal=40% |
| 7070 | INSULIN\_LIKE\_GROWTH\_FACTOR\_RECEPTOR\_BINDING(GO:0005159) |  | 16 | 0.52 | 0.89 | 0.476 | 0.655 | 1.000 | 25471 | tags=63%, list=43%, signal=110% |
| 7071 | INTEGRIN\_ACTIVATION(GO:0033622) |  | 19 | 0.52 | 0.89 | 0.476 | 0.658 | 1.000 | 27297 | tags=74%, list=46%, signal=137% |
| 7072 | MACROPHAGE\_CHEMOTAXIS(GO:0048246) |  | 32 | 0.52 | 0.89 | 0.476 | 0.664 | 1.000 | 27957 | tags=69%, list=47%, signal=130% |
| 7073 | REGULATION\_OF\_PRODUCTION\_OF\_MIRNAS\_INVOLVED\_IN\_GENE\_SILENCING\_BY\_MIRNA(GO:1903798) |  | 15 | 0.54 | 0.89 | 0.476 | 0.670 | 1.000 | 23586 | tags=60%, list=40%, signal=100% |
| 7074 | GLUCOCORTICOID\_RECEPTOR\_BINDING(GO:0035259) |  | 15 | 0.53 | 0.89 | 0.816 | 0.672 | 1.000 | 27668 | tags=100%, list=47%, signal=188% |
| 7075 | RESPONSE\_TO\_LAMINAR\_FLUID\_SHEAR\_STRESS(GO:0034616) |  | 15 | 0.53 | 0.89 | 0.816 | 0.674 | 1.000 | 26919 | tags=87%, list=46%, signal=159% |
| 7076 | CHROMATIN\_SILENCING\_AT\_RDNA(GO:0000183) |  | 39 | 0.51 | 0.89 | 0.816 | 0.678 | 1.000 | 28647 | tags=72%, list=48%, signal=139% |
| 7077 | ODORANT\_BINDING(GO:0005549) |  | 85 | 0.46 | 0.89 | 0.650 | 0.682 | 1.000 | 24394 | tags=9%, list=41%, signal=16% |
| 7078 | REGULATION\_OF\_FLAGELLATED\_SPERM\_MOTILITY(GO:1901317) |  | 16 | 0.54 | 0.89 | 0.650 | 0.683 | 1.000 | 26161 | tags=56%, list=44%, signal=101% |
| 7079 | REVERSE\_CHOLESTEROL\_TRANSPORT(GO:0043691) |  | 20 | 0.56 | 0.89 | 0.476 | 0.687 | 1.000 | 20626 | tags=35%, list=35%, signal=54% |
| 7080 | REGULATION\_OF\_MACROPHAGE\_MIGRATION(GO:1905521) |  | 24 | 0.52 | 0.88 | 0.476 | 0.708 | 1.000 | 27957 | tags=71%, list=47%, signal=134% |
| 7081 | REGULATION\_OF\_MACROPHAGE\_CHEMOTAXIS(GO:0010758) |  | 20 | 0.52 | 0.88 | 0.644 | 0.709 | 1.000 | 27957 | tags=75%, list=47%, signal=142% |
| 7082 | POSITIVE\_REGULATION\_OF\_ENDOPLASMIC\_RETICULUM\_UNFOLDED\_PROTEIN\_RESPONSE(GO:1900103) |  | 15 | 0.52 | 0.88 | 0.816 | 0.715 | 1.000 | 27813 | tags=73%, list=47%, signal=138% |
| 7083 | REGULATION\_OF\_TRIGLYCERIDE\_BIOSYNTHETIC\_PROCESS(GO:0010866) |  | 18 | 0.53 | 0.88 | 0.476 | 0.720 | 1.000 | 21292 | tags=44%, list=36%, signal=69% |
| 7084 | REGULATION\_OF\_NUCLEAR\_TRANSCRIBED\_MRNA\_CATABOLIC\_PROCESS\_\_DEADENYLATION\_DEPENDENT\_DECAY(GO:1900151) |  | 16 | 0.52 | 0.87 | 0.816 | 0.740 | 1.000 | 27509 | tags=88%, list=47%, signal=164% |
| 7085 | POSITIVE\_REGULATION\_OF\_NUCLEAR\_TRANSCRIBED\_MRNA\_CATABOLIC\_PROCESS\_\_DEADENYLATION\_DEPENDENT\_DECAY(GO:1900153) |  | 16 | 0.52 | 0.87 | 0.816 | 0.740 | 1.000 | 27509 | tags=88%, list=47%, signal=164% |
| 7086 | RESPONSE\_TO\_INTERLEUKIN\_7(GO:0098760) |  | 31 | 0.51 | 0.86 | 0.476 | 0.791 | 1.000 | 28931 | tags=74%, list=49%, signal=145% |
| 7087 | CELLULAR\_RESPONSE\_TO\_INTERLEUKIN\_7(GO:0098761) |  | 31 | 0.51 | 0.86 | 0.476 | 0.791 | 1.000 | 28931 | tags=74%, list=49%, signal=145% |
| 7088 | INTERLEUKIN\_7\_MEDIATED\_SIGNALING\_PATHWAY(GO:0038111) |  | 30 | 0.51 | 0.86 | 0.476 | 0.798 | 1.000 | 28931 | tags=73%, list=49%, signal=144% |
| 7089 | CHEMOKINE\_RECEPTOR\_BINDING(GO:0042379) |  | 64 | 0.52 | 0.86 | 0.816 | 0.802 | 1.000 | 25496 | tags=28%, list=43%, signal=49% |
| 7090 | MEMBRANE\_RAFT\_ORGANIZATION(GO:0031579) |  | 23 | 0.52 | 0.85 | 0.816 | 0.808 | 1.000 | 28203 | tags=74%, list=48%, signal=141% |
| 7091 | PEPTIDE\_CROSS\_LINKING(GO:0018149) |  | 59 | 0.50 | 0.85 | 0.476 | 0.809 | 1.000 | 26784 | tags=25%, list=45%, signal=46% |
| 7092 | KERATIN\_FILAMENT(GO:0045095) |  | 98 | 0.55 | 0.85 | 0.476 | 0.814 | 1.000 | 20748 | tags=7%, list=35%, signal=11% |
| 7093 | RECEPTOR\_INHIBITOR\_ACTIVITY(GO:0030547) |  | 19 | 0.52 | 0.85 | 0.476 | 0.825 | 1.000 | 27374 | tags=53%, list=46%, signal=98% |
| 7094 | LIPOPROTEIN\_PARTICLE\_RECEPTOR\_ACTIVITY(GO:0030228) |  | 16 | 0.52 | 0.84 | 0.644 | 0.841 | 1.000 | 28344 | tags=88%, list=48%, signal=168% |
| 7095 | POSITIVE\_REGULATION\_OF\_LIPID\_STORAGE(GO:0010884) |  | 22 | 0.51 | 0.84 | 0.476 | 0.841 | 1.000 | 28166 | tags=55%, list=48%, signal=104% |
| 7096 | NARROW\_PORE\_CHANNEL\_ACTIVITY(GO:0022842) |  | 19 | 0.52 | 0.84 | 0.476 | 0.850 | 1.000 | 26732 | tags=42%, list=45%, signal=77% |
| 7097 | LEAK\_CHANNEL\_ACTIVITY(GO:0022840) |  | 19 | 0.52 | 0.84 | 0.476 | 0.850 | 1.000 | 26732 | tags=42%, list=45%, signal=77% |
| 7098 | ENDOTHELIAL\_CELL\_CHEMOTAXIS(GO:0035767) |  | 30 | 0.51 | 0.84 | 0.476 | 0.853 | 1.000 | 28347 | tags=60%, list=48%, signal=115% |
| 7099 | REGULATION\_OF\_PROTEIN\_HOMOOLIGOMERIZATION(GO:0032462) |  | 21 | 0.53 | 0.82 | 0.648 | 0.904 | 1.000 | 23479 | tags=57%, list=40%, signal=95% |
| 7100 | FLAVONOID\_METABOLIC\_PROCESS(GO:0009812) |  | 16 | 0.51 | 0.82 | 0.816 | 0.906 | 1.000 | 28048 | tags=38%, list=47%, signal=71% |
| 7101 | POTASSIUM\_ION\_LEAK\_CHANNEL\_ACTIVITY(GO:0022841) |  | 16 | 0.52 | 0.82 | 0.476 | 0.915 | 1.000 | 26732 | tags=38%, list=45%, signal=68% |
| 7102 | STABILIZATION\_OF\_MEMBRANE\_POTENTIAL(GO:0030322) |  | 16 | 0.52 | 0.81 | 0.476 | 0.927 | 1.000 | 14193 | tags=13%, list=24%, signal=16% |
| 7103 | MHC\_CLASS\_II\_PROTEIN\_COMPLEX(GO:0042613) |  | 16 | 0.49 | 0.78 | 0.816 | 0.985 | 1.000 | 28291 | tags=50%, list=48%, signal=96% |
| 7104 | CHEMOKINE\_ACTIVITY(GO:0008009) |  | 50 | 0.48 | 0.76 | 0.816 | 0.996 | 1.000 | 27827 | tags=28%, list=47%, signal=53% |
| 7105 | IMMUNOGLOBULIN\_COMPLEX(GO:0019814) |  | 57 | 0.40 | 0.76 | 0.816 | 0.997 | 1.000 | 22967 | tags=7%, list=39%, signal=11% |
| 7106 | POSITIVE\_REGULATION\_OF\_HUMORAL\_IMMUNE\_RESPONSE(GO:0002922) |  | 15 | 0.49 | 0.74 | 0.816 | 0.999 | 1.000 | 23238 | tags=20%, list=39%, signal=33% |
| 7107 | MRNA\_BINDING\_INVOLVED\_IN\_POSTTRANSCRIPTIONAL\_GENE\_SILENCING(GO:1903231) |  | 174 | 0.41 | 0.72 | 0.816 | 1.000 | 1.000 | 28306 | tags=16%, list=48%, signal=30% |
| 7108 | REGULATION\_OF\_LEUKOCYTE\_ADHESION\_TO\_VASCULAR\_ENDOTHELIAL\_CELL(GO:1904994) |  | 21 | 0.43 | 0.72 | 0.816 | 1.000 | 1.000 | 20764 | tags=29%, list=35%, signal=44% |
| 7109 | NEGATIVE\_REGULATION\_OF\_BLOOD\_VESSEL\_ENDOTHELIAL\_CELL\_PROLIFERATION\_INVOLVED\_IN\_SPROUTING\_ANGIOGENESIS(GO:1903588) |  | 25 | 0.42 | 0.63 | 0.816 | 1.000 | 1.000 | 28381 | tags=28%, list=48%, signal=54% |
Table: Gene sets enriched in phenotype
 **C328SVIM (2 samples)
 **[plain text format]****

  
