## Supplementary material for "A single cysteine residue in vimentin regulates long non-coding RNA *XIST* to suppress epithelial-mesenchymal transition and stemness in breast cancer": INSULIN_RECEPTOR_BINDING(GO_0005158).html

Details for gene set INSULIN\_RECEPTOR\_BINDING(GO:0005158)[GSEA]


|  || Dataset | fpkm.sample |
| Phenotype | sample.cls |
| Upregulated in class | C328SVIM |
| GeneSet | INSULIN\_RECEPTOR\_BINDING(GO:0005158) |
| Enrichment Score (ES) | 0.89978206 |
| Normalized Enrichment Score (NES) | 1.439941 |
| Nominal p-value | 0.0 |
| FDR q-value | 1.0 |
| FWER p-Value | 0.325 |
Table: GSEA Results Summary

  

Fig 1: Enrichment plot: INSULIN\_RECEPTOR\_BINDING(GO:0005158)
  
Profile of the Running ES Score & Positions of GeneSet Members on the Rank Ordered List

  

| PROBE | DESCRIPTION   (from dataset) | GENE SYMBOL | GENE\_TITLE | RANK IN GENE LIST | RANK METRIC SCORE | RUNNING ES | CORE ENRICHMENT || 1 | ENSG00000133124 | IRS4 |  |  | 5824 | 14620.068 | 0.8998 | Yes |
| 2 | ENSG00000095637 | SORBS1 |  |  | 8594 | 4.504 | 0.8532 | No |
| 3 | ENSG00000106070 | GRB10 |  |  | 8801 | 4.164 | 0.8500 | No |
| 4 | ENSG00000146247 | PHIP |  |  | 11013 | 2.451 | 0.8128 | No |
| 5 | ENSG00000197594 | ENPP1 |  |  | 11589 | 2.241 | 0.8032 | No |
| 6 | ENSG00000205302 | SNX2 |  |  | 13079 | 1.767 | 0.7782 | No |
| 7 | ENSG00000168216 | LMBRD1 |  |  | 15338 | 1.332 | 0.7401 | No |
| 8 | ENSG00000179295 | PTPN11 |  |  | 15435 | 1.312 | 0.7385 | No |
| 9 | ENSG00000017427 | IGF1 |  |  | 15922 | 1.241 | 0.7304 | No |
| 10 | ENSG00000167244 | IGF2 |  |  | 16399 | 1.178 | 0.7224 | No |
| 11 | ENSG00000114520 | SNX4 |  |  | 16632 | 1.150 | 0.7186 | No |
| 12 | ENSG00000028528 | SNX1 |  |  | 21418 | 0.649 | 0.6377 | No |
| 13 | ENSG00000185950 | IRS2 |  |  | 22463 | 0.560 | 0.6201 | No |
| 14 | ENSG00000140992 | PDPK1 |  |  | 23076 | 0.514 | 0.6097 | No |
| 15 | ENSG00000197122 | SRC |  |  | 23479 | 0.482 | 0.6030 | No |
| 16 | ENSG00000196396 | PTPN1 |  |  | 24541 | 0.398 | 0.5850 | No |
| 17 | ENSG00000160691 | SHC1 |  |  | 25353 | 0.339 | 0.5713 | No |
| 18 | ENSG00000140443 | IGF1R |  |  | 26394 | 0.255 | 0.5538 | No |
| 19 | ENSG00000145675 | PIK3R1 |  |  | 27813 | 0.136 | 0.5298 | No |
| 20 | ENSG00000248099 | INSL3 |  |  | 28031 | 0.117 | 0.5261 | No |
| 21 | ENSG00000169047 | IRS1 |  |  | 28746 | 0.054 | 0.5141 | No |
| 22 | ENSG00000254647 | INS |  |  | 32571 | 0.000 | 0.4494 | No |
Table: GSEA details
[plain text format]

  

Fig 2: INSULIN\_RECEPTOR\_BINDING(GO:0005158)
  
Blue-Pink O' Gram in the Space of the Analyzed GeneSet

  

Fig 3: INSULIN\_RECEPTOR\_BINDING(GO:0005158): Random ES distribution
  
Gene set null distribution of ES for
 **INSULIN\_RECEPTOR\_BINDING(GO:0005158)**

  
