## Supplementary material for "A single cysteine residue in vimentin regulates long non-coding RNA *XIST* to suppress epithelial-mesenchymal transition and stemness in breast cancer": MYELIN_SHEATH(GO_0043209).html

Details for gene set MYELIN\_SHEATH(GO:0043209)[GSEA]


|  || Dataset | fpkm.sample |
| Phenotype | sample.cls |
| Upregulated in class | C328SVIM |
| GeneSet | MYELIN\_SHEATH(GO:0043209) |
| Enrichment Score (ES) | 0.8808323 |
| Normalized Enrichment Score (NES) | 1.4296062 |
| Nominal p-value | 0.0 |
| FDR q-value | 1.0 |
| FWER p-Value | 0.325 |
Table: GSEA Results Summary

  

Fig 1: Enrichment plot: MYELIN\_SHEATH(GO:0043209)
  
Profile of the Running ES Score & Positions of GeneSet Members on the Rank Ordered List

  

| PROBE | DESCRIPTION   (from dataset) | GENE SYMBOL | GENE\_TITLE | RANK IN GENE LIST | RANK METRIC SCORE | RUNNING ES | CORE ENRICHMENT || 1 | ENSG00000168314 | MOBP |  |  | 160 | Infinity | -0.0027 | Yes |
| 2 | ENSG00000148798 | INA |  |  | 1410 | Infinity | -0.0239 | Yes |
| 3 | ENSG00000213619 | NDUFS3 |  |  | 2068 | Infinity | -0.0350 | Yes |
| 4 | ENSG00000018236 | CNTN1 |  |  | 2188 | Infinity | -0.0370 | Yes |
| 5 | ENSG00000154096 | THY1 |  |  | 3788 | Infinity | -0.0642 | Yes |
| 6 | ENSG00000119865 | CNRIP1 |  |  | 4194 | Infinity | -0.0710 | Yes |
| 7 | ENSG00000147588 | PMP2 |  |  | 4667 | Infinity | -0.0790 | Yes |
| 8 | ENSG00000277586 | NEFL |  |  | 5211 | Infinity | -0.0882 | Yes |
| 9 | ENSG00000136541 | ERMN |  |  | 5581 | Infinity | -0.0945 | Yes |
| 10 | ENSG00000163631 | ALB |  |  | 5747 | Infinity | -0.0973 | Yes |
| 11 | ENSG00000144834 | TAGLN3 |  |  | 5804 | Infinity | -0.0982 | Yes |
| 12 | ENSG00000147065 | MSN |  |  | 5823 | 15136.710 | 0.7867 | Yes |
| 13 | ENSG00000085662 | AKR1B1 |  |  | 5839 | 1419.529 | 0.8695 | Yes |
| 14 | ENSG00000166086 | JAM3 |  |  | 6011 | 112.234 | 0.8732 | Yes |
| 15 | ENSG00000154277 | UCHL1 |  |  | 6047 | 89.810 | 0.8778 | Yes |
| 16 | ENSG00000111716 | LDHB |  |  | 6164 | 51.630 | 0.8789 | Yes |
| 17 | ENSG00000114450 | GNB4 |  |  | 6245 | 39.860 | 0.8799 | Yes |
| 18 | ENSG00000134198 | TSPAN2 |  |  | 6303 | 33.030 | 0.8808 | Yes |
| 19 | ENSG00000087258 | GNAO1 |  |  | 6471 | 22.359 | 0.8793 | No |
| 20 | ENSG00000185015 | CA13 |  |  | 6660 | 16.073 | 0.8771 | No |
| 21 | ENSG00000166165 | CKB |  |  | 6841 | 13.140 | 0.8748 | No |
| 22 | ENSG00000158887 | MPZ |  |  | 7168 | 9.731 | 0.8698 | No |
| 23 | ENSG00000123560 | PLP1 |  |  | 7818 | 6.236 | 0.8592 | No |
| 24 | ENSG00000104833 | TUBB4A |  |  | 7859 | 6.098 | 0.8589 | No |
| 25 | ENSG00000163531 | NFASC |  |  | 7932 | 5.834 | 0.8580 | No |
| 26 | ENSG00000104267 | CA2 |  |  | 8241 | 5.134 | 0.8531 | No |
| 27 | ENSG00000131095 | GFAP |  |  | 8554 | 4.577 | 0.8480 | No |
| 28 | ENSG00000106976 | DNM1 |  |  | 8763 | 4.227 | 0.8448 | No |
| 29 | ENSG00000120438 | TCP1 |  |  | 10138 | 2.914 | 0.8216 | No |
| 30 | ENSG00000108797 | CNTNAP1 |  |  | 10339 | 2.782 | 0.8184 | No |
| 31 | ENSG00000073969 | NSF |  |  | 10559 | 2.655 | 0.8148 | No |
| 32 | ENSG00000115840 | SLC25A12 |  |  | 11310 | 2.340 | 0.8023 | No |
| 33 | ENSG00000137710 | RDX |  |  | 11742 | 2.181 | 0.7951 | No |
| 34 | ENSG00000080824 | HSP90AA1 |  |  | 11758 | 2.171 | 0.7949 | No |
| 35 | ENSG00000173786 | CNP |  |  | 11879 | 2.124 | 0.7930 | No |
| 36 | ENSG00000125814 | NAPB |  |  | 11885 | 2.122 | 0.7931 | No |
| 37 | ENSG00000150768 | DLAT |  |  | 12039 | 2.064 | 0.7906 | No |
| 38 | ENSG00000136143 | SUCLA2 |  |  | 12438 | 1.944 | 0.7840 | No |
| 39 | ENSG00000075618 | FSCN1 |  |  | 12447 | 1.938 | 0.7839 | No |
| 40 | ENSG00000092964 | DPYSL2 |  |  | 12828 | 1.832 | 0.7776 | No |
| 41 | ENSG00000108387 | SEPT4 |  |  | 12938 | 1.799 | 0.7759 | No |
| 42 | ENSG00000184144 | CNTN2 |  |  | 13179 | 1.741 | 0.7719 | No |
| 43 | ENSG00000171368 | TPPP |  |  | 13336 | 1.701 | 0.7693 | No |
| 44 | ENSG00000014641 | MDH1 |  |  | 13640 | 1.640 | 0.7643 | No |
| 45 | ENSG00000165672 | PRDX3 |  |  | 13697 | 1.628 | 0.7635 | No |
| 46 | ENSG00000116473 | RAP1A |  |  | 14142 | 1.556 | 0.7560 | No |
| 47 | ENSG00000116459 | ATP5PB |  |  | 14263 | 1.529 | 0.7541 | No |
| 48 | ENSG00000125166 | GOT2 |  |  | 14398 | 1.498 | 0.7519 | No |
| 49 | ENSG00000163558 | PRKCI |  |  | 14571 | 1.475 | 0.7491 | No |
| 50 | ENSG00000127022 | CANX |  |  | 14580 | 1.473 | 0.7490 | No |
| 51 | ENSG00000072415 | MPP5 |  |  | 14654 | 1.458 | 0.7479 | No |
| 52 | ENSG00000114573 | ATP6V1A |  |  | 14815 | 1.424 | 0.7452 | No |
| 53 | ENSG00000176402 | GJC3 |  |  | 14885 | 1.412 | 0.7441 | No |
| 54 | ENSG00000142168 | SOD1 |  |  | 14948 | 1.400 | 0.7432 | No |
| 55 | ENSG00000165637 | VDAC2 |  |  | 15063 | 1.378 | 0.7413 | No |
| 56 | ENSG00000147416 | ATP6V1B2 |  |  | 15196 | 1.354 | 0.7392 | No |
| 57 | ENSG00000006451 | RALA |  |  | 15266 | 1.342 | 0.7381 | No |
| 58 | ENSG00000198835 | GJC2 |  |  | 15500 | 1.304 | 0.7342 | No |
| 59 | ENSG00000157870 | FAM213B |  |  | 15577 | 1.292 | 0.7330 | No |
| 60 | ENSG00000132305 | IMMT |  |  | 15636 | 1.285 | 0.7321 | No |
| 61 | ENSG00000167004 | PDIA3 |  |  | 15681 | 1.278 | 0.7314 | No |
| 62 | ENSG00000107186 | MPDZ |  |  | 15696 | 1.275 | 0.7312 | No |
| 63 | ENSG00000117450 | PRDX1 |  |  | 15716 | 1.273 | 0.7310 | No |
| 64 | ENSG00000170248 | PDCD6IP |  |  | 15718 | 1.273 | 0.7310 | No |
| 65 | ENSG00000100285 | NEFH |  |  | 15770 | 1.264 | 0.7303 | No |
| 66 | ENSG00000163468 | CCT3 |  |  | 15907 | 1.241 | 0.7280 | No |
| 67 | ENSG00000126803 | HSPA2 |  |  | 16070 | 1.223 | 0.7253 | No |
| 68 | ENSG00000092621 | PHGDH |  |  | 16110 | 1.215 | 0.7248 | No |
| 69 | ENSG00000135821 | GLUL |  |  | 16130 | 1.213 | 0.7245 | No |
| 70 | ENSG00000143947 | RPS27A |  |  | 16724 | 1.137 | 0.7145 | No |
| 71 | ENSG00000174238 | PITPNA |  |  | 16755 | 1.132 | 0.7141 | No |
| 72 | ENSG00000166226 | CCT2 |  |  | 16850 | 1.121 | 0.7125 | No |
| 73 | ENSG00000091140 | DLD |  |  | 16859 | 1.119 | 0.7125 | No |
| 74 | ENSG00000165629 | ATP5F1C |  |  | 17038 | 1.098 | 0.7095 | No |
| 75 | ENSG00000138107 | ACTR1A |  |  | 17072 | 1.093 | 0.7090 | No |
| 76 | ENSG00000168439 | STIP1 |  |  | 17116 | 1.084 | 0.7084 | No |
| 77 | ENSG00000171314 | PGAM1 |  |  | 17361 | 1.054 | 0.7043 | No |
| 78 | ENSG00000171791 | BCL2 |  |  | 17501 | 1.037 | 0.7020 | No |
| 79 | ENSG00000136854 | STXBP1 |  |  | 17522 | 1.035 | 0.7017 | No |
| 80 | ENSG00000105220 | GPI |  |  | 17565 | 1.029 | 0.7011 | No |
| 81 | ENSG00000131828 | PDHA1 |  |  | 17637 | 1.020 | 0.6999 | No |
| 82 | ENSG00000167863 | ATP5PD |  |  | 17644 | 1.020 | 0.6999 | No |
| 83 | ENSG00000109971 | HSPA8 |  |  | 17646 | 1.019 | 0.6999 | No |
| 84 | ENSG00000182621 | PLCB1 |  |  | 18178 | 0.952 | 0.6910 | No |
| 85 | ENSG00000167552 | TUBA1A |  |  | 18238 | 0.946 | 0.6900 | No |
| 86 | ENSG00000057608 | GDI2 |  |  | 18466 | 0.920 | 0.6862 | No |
| 87 | ENSG00000044574 | HSPA5 |  |  | 18513 | 0.915 | 0.6855 | No |
| 88 | ENSG00000113013 | HSPA9 |  |  | 18525 | 0.914 | 0.6854 | No |
| 89 | ENSG00000013016 | EHD3 |  |  | 18676 | 0.894 | 0.6829 | No |
| 90 | ENSG00000152234 | ATP5F1A |  |  | 18951 | 0.867 | 0.6783 | No |
| 91 | ENSG00000184113 | CLDN5 |  |  | 19080 | 0.851 | 0.6762 | No |
| 92 | ENSG00000075711 | DLG1 |  |  | 19219 | 0.839 | 0.6739 | No |
| 93 | ENSG00000140740 | UQCRC2 |  |  | 19223 | 0.838 | 0.6739 | No |
| 94 | ENSG00000005022 | SLC25A5 |  |  | 19367 | 0.823 | 0.6715 | No |
| 95 | ENSG00000171862 | PTEN |  |  | 19603 | 0.813 | 0.6675 | No |
| 96 | ENSG00000150753 | CCT5 |  |  | 19773 | 0.796 | 0.6647 | No |
| 97 | ENSG00000166411 | IDH3A |  |  | 19992 | 0.778 | 0.6611 | No |
| 98 | ENSG00000176658 | MYO1D |  |  | 19993 | 0.778 | 0.6611 | No |
| 99 | ENSG00000178741 | COX5A |  |  | 20004 | 0.777 | 0.6610 | No |
| 100 | ENSG00000119689 | DLST |  |  | 20058 | 0.771 | 0.6601 | No |
| 101 | ENSG00000075415 | SLC25A3 |  |  | 20216 | 0.753 | 0.6575 | No |
| 102 | ENSG00000023228 | NDUFS1 |  |  | 20686 | 0.723 | 0.6496 | No |
| 103 | ENSG00000100412 | ACO2 |  |  | 20703 | 0.721 | 0.6494 | No |
| 104 | ENSG00000239672 | NME1 |  |  | 20715 | 0.719 | 0.6492 | No |
| 105 | ENSG00000134265 | NAPG |  |  | 20951 | 0.693 | 0.6453 | No |
| 106 | ENSG00000110955 | ATP5F1B |  |  | 20952 | 0.693 | 0.6453 | No |
| 107 | ENSG00000151729 | SLC25A4 |  |  | 21199 | 0.670 | 0.6412 | No |
| 108 | ENSG00000156508 | EEF1A1 |  |  | 21336 | 0.658 | 0.6389 | No |
| 109 | ENSG00000165527 | ARF6 |  |  | 21611 | 0.631 | 0.6343 | No |
| 110 | ENSG00000170027 | YWHAG |  |  | 21682 | 0.624 | 0.6332 | No |
| 111 | ENSG00000167085 | PHB |  |  | 21992 | 0.599 | 0.6280 | No |
| 112 | ENSG00000104419 | NDRG1 |  |  | 22035 | 0.595 | 0.6273 | No |
| 113 | ENSG00000010256 | UQCRC1 |  |  | 22092 | 0.590 | 0.6264 | No |
| 114 | ENSG00000130414 | NDUFA10 |  |  | 22462 | 0.560 | 0.6202 | No |
| 115 | ENSG00000203879 | GDI1 |  |  | 22812 | 0.536 | 0.6143 | No |
| 116 | ENSG00000008056 | SYN1 |  |  | 22865 | 0.532 | 0.6134 | No |
| 117 | ENSG00000163399 | ATP1A1 |  |  | 22943 | 0.524 | 0.6121 | No |
| 118 | ENSG00000184009 | ACTG1 |  |  | 23116 | 0.511 | 0.6093 | No |
| 119 | ENSG00000123416 | TUBA1B |  |  | 23130 | 0.509 | 0.6091 | No |
| 120 | ENSG00000178952 | TUFM |  |  | 23469 | 0.483 | 0.6034 | No |
| 121 | ENSG00000092820 | EZR |  |  | 23520 | 0.479 | 0.6025 | No |
| 122 | ENSG00000067606 | PRKCZ |  |  | 23540 | 0.477 | 0.6022 | No |
| 123 | ENSG00000078369 | GNB1 |  |  | 23704 | 0.464 | 0.5995 | No |
| 124 | ENSG00000146701 | MDH2 |  |  | 23751 | 0.460 | 0.5988 | No |
| 125 | ENSG00000102934 | PLLP |  |  | 23766 | 0.459 | 0.5985 | No |
| 126 | ENSG00000145545 | SRD5A1 |  |  | 23890 | 0.449 | 0.5965 | No |
| 127 | ENSG00000165280 | VCP |  |  | 23913 | 0.447 | 0.5961 | No |
| 128 | ENSG00000103496 | STX4 |  |  | 24007 | 0.439 | 0.5946 | No |
| 129 | ENSG00000150093 | ITGB1 |  |  | 24090 | 0.433 | 0.5932 | No |
| 130 | ENSG00000018625 | ATP1A2 |  |  | 24101 | 0.431 | 0.5931 | No |
| 131 | ENSG00000105402 | NAPA |  |  | 24163 | 0.426 | 0.5921 | No |
| 132 | ENSG00000178127 | NDUFV2 |  |  | 24244 | 0.420 | 0.5907 | No |
| 133 | ENSG00000111674 | ENO2 |  |  | 24342 | 0.411 | 0.5891 | No |
| 134 | ENSG00000073578 | SDHA |  |  | 24405 | 0.410 | 0.5881 | No |
| 135 | ENSG00000180900 | SCRIB |  |  | 24481 | 0.404 | 0.5868 | No |
| 136 | ENSG00000163931 | TKT |  |  | 24652 | 0.389 | 0.5840 | No |
| 137 | ENSG00000131558 | EXOC4 |  |  | 24688 | 0.386 | 0.5834 | No |
| 138 | ENSG00000141736 | ERBB2 |  |  | 24834 | 0.375 | 0.5810 | No |
| 139 | ENSG00000067225 | PKM |  |  | 24975 | 0.369 | 0.5786 | No |
| 140 | ENSG00000188229 | TUBB4B |  |  | 25406 | 0.334 | 0.5713 | No |
| 141 | ENSG00000164300 | SERINC5 |  |  | 25454 | 0.331 | 0.5706 | No |
| 142 | ENSG00000187486 | KCNJ11 |  |  | 25913 | 0.292 | 0.5628 | No |
| 143 | ENSG00000150991 | UBC |  |  | 26040 | 0.281 | 0.5607 | No |
| 144 | ENSG00000109846 | CRYAB |  |  | 26321 | 0.261 | 0.5560 | No |
| 145 | ENSG00000110047 | EHD1 |  |  | 26447 | 0.248 | 0.5539 | No |
| 146 | ENSG00000109099 | PMP22 |  |  | 26542 | 0.243 | 0.5523 | No |
| 147 | ENSG00000130707 | ASS1 |  |  | 26919 | 0.210 | 0.5459 | No |
| 148 | ENSG00000071127 | WDR1 |  |  | 27073 | 0.199 | 0.5433 | No |
| 149 | ENSG00000105357 | MYH14 |  |  | 27263 | 0.183 | 0.5401 | No |
| 150 | ENSG00000172354 | GNB2 |  |  | 27293 | 0.181 | 0.5397 | No |
| 151 | ENSG00000069966 | GNB5 |  |  | 27311 | 0.179 | 0.5394 | No |
| 152 | ENSG00000143153 | ATP1B1 |  |  | 27478 | 0.163 | 0.5366 | No |
| 153 | ENSG00000182718 | ANXA2 |  |  | 27588 | 0.156 | 0.5347 | No |
| 154 | ENSG00000096433 | ITPR3 |  |  | 27876 | 0.130 | 0.5299 | No |
| 155 | ENSG00000148180 | GSN |  |  | 28203 | 0.102 | 0.5244 | No |
| 156 | ENSG00000132639 | SNAP25 |  |  | 28932 | 0.039 | 0.5120 | No |
| 157 | ENSG00000197971 | MBP |  |  | 28944 | 0.038 | 0.5118 | No |
| 158 | ENSG00000152939 | MARVELD2 |  |  | 28973 | 0.036 | 0.5114 | No |
| 159 | ENSG00000124507 | PACSIN1 |  |  | 29172 | 0.023 | 0.5080 | No |
| 160 | ENSG00000101210 | EEF1A2 |  |  | 29343 | 0.012 | 0.5051 | No |
| 161 | ENSG00000105695 | MAG |  |  | 31301 | 0.000 | 0.4719 | No |
| 162 | ENSG00000184454 | NCMAP |  |  | 32589 | 0.000 | 0.4501 | No |
| 163 | ENSG00000240972 | MIF |  |  | 39917 | NaN | 0.3259 | No |
| 164 | ENSG00000068903 | SIRT2 |  |  | 53252 | NaN | 0.0998 | No |
Table: GSEA details
[plain text format]

  

Fig 2: MYELIN\_SHEATH(GO:0043209)
  
Blue-Pink O' Gram in the Space of the Analyzed GeneSet

  

Fig 3: MYELIN\_SHEATH(GO:0043209): Random ES distribution
  
Gene set null distribution of ES for
 **MYELIN\_SHEATH(GO:0043209)**

  
