## Supplementary material for "A single cysteine residue in vimentin regulates long non-coding RNA *XIST* to suppress epithelial-mesenchymal transition and stemness in breast cancer": NEGATIVE_REGULATION_OF_INTERLEUKIN_1_PRODUCTION(GO_0032692).html

Details for gene set NEGATIVE\_REGULATION\_OF\_INTERLEUKIN\_1\_PRODUCTION(GO:0032692)[GSEA]


|  || Dataset | fpkm.sample |
| Phenotype | sample.cls |
| Upregulated in class | C328SVIM |
| GeneSet | NEGATIVE\_REGULATION\_OF\_INTERLEUKIN\_1\_PRODUCTION(GO:0032692) |
| Enrichment Score (ES) | 0.89952904 |
| Normalized Enrichment Score (NES) | 1.4478352 |
| Nominal p-value | 0.0 |
| FDR q-value | 1.0 |
| FWER p-Value | 0.325 |
Table: GSEA Results Summary

  

Fig 1: Enrichment plot: NEGATIVE\_REGULATION\_OF\_INTERLEUKIN\_1\_PRODUCTION(GO:0032692)
  
Profile of the Running ES Score & Positions of GeneSet Members on the Rank Ordered List

  

| PROBE | DESCRIPTION   (from dataset) | GENE SYMBOL | GENE\_TITLE | RANK IN GENE LIST | RANK METRIC SCORE | RUNNING ES | CORE ENRICHMENT || 1 | ENSG00000175344 | CHRNA7 |  |  | 1612 | Infinity | -0.0273 | Yes |
| 2 | ENSG00000084207 | GSTP1 |  |  | 5825 | 8120.758 | 0.8995 | Yes |
| 3 | ENSG00000105483 | CARD8 |  |  | 9648 | 3.241 | 0.8353 | No |
| 4 | ENSG00000118137 | APOA1 |  |  | 10458 | 2.708 | 0.8219 | No |
| 5 | ENSG00000141480 | ARRB2 |  |  | 12549 | 1.909 | 0.7868 | No |
| 6 | ENSG00000103313 | MEFV |  |  | 13873 | 1.596 | 0.7646 | No |
| 7 | ENSG00000183087 | GAS6 |  |  | 16662 | 1.148 | 0.7176 | No |
| 8 | ENSG00000115590 | IL1R2 |  |  | 17771 | 1.003 | 0.6990 | No |
| 9 | ENSG00000157017 | GHRL |  |  | 18462 | 0.921 | 0.6874 | No |
| 10 | ENSG00000140464 | PML |  |  | 20085 | 0.767 | 0.6600 | No |
| 11 | ENSG00000116285 | ERRFI1 |  |  | 21719 | 0.621 | 0.6325 | No |
| 12 | ENSG00000102575 | ACP5 |  |  | 22934 | 0.525 | 0.6120 | No |
| 13 | ENSG00000118503 | TNFAIP3 |  |  | 24022 | 0.438 | 0.5937 | No |
| 14 | ENSG00000079385 | CEACAM1 |  |  | 24797 | 0.378 | 0.5806 | No |
| 15 | ENSG00000224051 | CPTP |  |  | 25094 | 0.358 | 0.5757 | No |
| 16 | ENSG00000163874 | ZC3H12A |  |  | 28166 | 0.104 | 0.5237 | No |
| 17 | ENSG00000142405 | NLRP12 |  |  | 30990 | 0.000 | 0.4760 | No |
| 18 | ENSG00000162711 | NLRP3 |  |  | 31381 | 0.000 | 0.4694 | No |
| 19 | ENSG00000012504 | NR1H4 |  |  | 32894 | 0.000 | 0.4438 | No |
| 20 | ENSG00000255221 | CARD17 |  |  | 36092 | NaN | 0.3897 | No |
| 21 | ENSG00000253548 | PYDC2 |  |  | 36411 | NaN | 0.3843 | No |
| 22 | ENSG00000167634 | NLRP7 |  |  | 47307 | NaN | 0.2000 | No |
| 23 | ENSG00000136634 | IL10 |  |  | 48582 | NaN | 0.1785 | No |
| 24 | ENSG00000121853 | GHSR |  |  | 49655 | NaN | 0.1603 | No |
| 25 | ENSG00000204397 | CARD16 |  |  | 51193 | NaN | 0.1343 | No |
| 26 | ENSG00000283904 | MIR155 |  |  | 56972 | NaN | 0.0366 | No |
| 27 | ENSG00000255501 | CARD18 |  |  | 58939 | NaN | 0.0033 | No |
Table: GSEA details
[plain text format]

  

Fig 2: NEGATIVE\_REGULATION\_OF\_INTERLEUKIN\_1\_PRODUCTION(GO:0032692)
  
Blue-Pink O' Gram in the Space of the Analyzed GeneSet

  

Fig 3: NEGATIVE\_REGULATION\_OF\_INTERLEUKIN\_1\_PRODUCTION(GO:0032692): Random ES distribution
  
Gene set null distribution of ES for
 **NEGATIVE\_REGULATION\_OF\_INTERLEUKIN\_1\_PRODUCTION(GO:0032692)**

  
