## Supplementary material for "A single cysteine residue in vimentin regulates long non-coding RNA *XIST* to suppress epithelial-mesenchymal transition and stemness in breast cancer": PROTEIN_LOCALIZATION_TO_ENDOSOME(GO_0036010).html

Details for gene set PROTEIN\_LOCALIZATION\_TO\_ENDOSOME(GO:0036010)[GSEA]


|  || Dataset | fpkm.sample |
| Phenotype | sample.cls |
| Upregulated in class | C328SVIM |
| GeneSet | PROTEIN\_LOCALIZATION\_TO\_ENDOSOME(GO:0036010) |
| Enrichment Score (ES) | 0.9005573 |
| Normalized Enrichment Score (NES) | 1.4400989 |
| Nominal p-value | 0.0 |
| FDR q-value | 1.0 |
| FWER p-Value | 0.325 |
Table: GSEA Results Summary

  

Fig 1: Enrichment plot: PROTEIN\_LOCALIZATION\_TO\_ENDOSOME(GO:0036010)
  
Profile of the Running ES Score & Positions of GeneSet Members on the Rank Ordered List

  

| PROBE | DESCRIPTION   (from dataset) | GENE SYMBOL | GENE\_TITLE | RANK IN GENE LIST | RANK METRIC SCORE | RUNNING ES | CORE ENRICHMENT || 1 | ENSG00000147065 | MSN |  |  | 5823 | 15136.710 | 0.9006 | Yes |
| 2 | ENSG00000137710 | RDX |  |  | 11742 | 2.181 | 0.8006 | No |
| 3 | ENSG00000114331 | ACAP2 |  |  | 13307 | 1.708 | 0.7743 | No |
| 4 | ENSG00000134318 | ROCK2 |  |  | 13969 | 1.577 | 0.7632 | No |
| 5 | ENSG00000107362 | ABHD17B |  |  | 14619 | 1.467 | 0.7523 | No |
| 6 | ENSG00000100139 | MICALL1 |  |  | 15685 | 1.278 | 0.7344 | No |
| 7 | ENSG00000112697 | TMEM30A |  |  | 18611 | 0.902 | 0.6849 | No |
| 8 | ENSG00000129968 | ABHD17A |  |  | 20065 | 0.770 | 0.6604 | No |
| 9 | ENSG00000137642 | SORL1 |  |  | 20346 | 0.742 | 0.6557 | No |
| 10 | ENSG00000100266 | PACSIN2 |  |  | 21445 | 0.646 | 0.6372 | No |
| 11 | ENSG00000165527 | ARF6 |  |  | 21611 | 0.631 | 0.6345 | No |
| 12 | ENSG00000111737 | RAB35 |  |  | 21807 | 0.618 | 0.6312 | No |
| 13 | ENSG00000078902 | TOLLIP |  |  | 23136 | 0.509 | 0.6088 | No |
| 14 | ENSG00000092820 | EZR |  |  | 23520 | 0.479 | 0.6023 | No |
| 15 | ENSG00000180900 | SCRIB |  |  | 24481 | 0.404 | 0.5861 | No |
| 16 | ENSG00000136379 | ABHD17C |  |  | 26196 | 0.274 | 0.5571 | No |
| 17 | ENSG00000099250 | NRP1 |  |  | 28347 | 0.089 | 0.5208 | No |
| 18 | ENSG00000163840 | DTX3L |  |  | 28908 | 0.040 | 0.5113 | No |
Table: GSEA details
[plain text format]

  

Fig 2: PROTEIN\_LOCALIZATION\_TO\_ENDOSOME(GO:0036010)
  
Blue-Pink O' Gram in the Space of the Analyzed GeneSet

  

Fig 3: PROTEIN\_LOCALIZATION\_TO\_ENDOSOME(GO:0036010): Random ES distribution
  
Gene set null distribution of ES for
 **PROTEIN\_LOCALIZATION\_TO\_ENDOSOME(GO:0036010)**

  
