## Supplementary material for "A single cysteine residue in vimentin regulates long non-coding RNA *XIST* to suppress epithelial-mesenchymal transition and stemness in breast cancer": REGULATION_OF_CELLULAR_PROTEIN_CATABOLIC_PROCESS(GO_1903362).html

Details for gene set REGULATION\_OF\_CELLULAR\_PROTEIN\_CATABOLIC\_PROCESS(GO:1903362)[GSEA]


|  || Dataset | fpkm.sample |
| Phenotype | sample.cls |
| Upregulated in class | C328SVIM |
| GeneSet | REGULATION\_OF\_CELLULAR\_PROTEIN\_CATABOLIC\_PROCESS(GO:1903362) |
| Enrichment Score (ES) | 0.8797938 |
| Normalized Enrichment Score (NES) | 1.4309075 |
| Nominal p-value | 0.0 |
| FDR q-value | 1.0 |
| FWER p-Value | 0.325 |
Table: GSEA Results Summary

  

Fig 1: Enrichment plot: REGULATION\_OF\_CELLULAR\_PROTEIN\_CATABOLIC\_PROCESS(GO:1903362)
  
Profile of the Running ES Score & Positions of GeneSet Members on the Rank Ordered List

  

| PROBE | DESCRIPTION   (from dataset) | GENE SYMBOL | GENE\_TITLE | RANK IN GENE LIST | RANK METRIC SCORE | RUNNING ES | CORE ENRICHMENT || 1 | ENSG00000169174 | PCSK9 |  |  | 439 | Infinity | -0.0075 | Yes |
| 2 | ENSG00000204463 | BAG6 |  |  | 572 | Infinity | -0.0097 | Yes |
| 3 | ENSG00000137033 | IL33 |  |  | 705 | Infinity | -0.0119 | Yes |
| 4 | ENSG00000182533 | CAV3 |  |  | 769 | Infinity | -0.0130 | Yes |
| 5 | ENSG00000157119 | KLHL40 |  |  | 1829 | Infinity | -0.0310 | Yes |
| 6 | ENSG00000165973 | NELL1 |  |  | 4570 | Infinity | -0.0775 | Yes |
| 7 | ENSG00000165617 | DACT1 |  |  | 5294 | Infinity | -0.0898 | Yes |
| 8 | ENSG00000147065 | MSN |  |  | 5823 | 15136.710 | 0.7523 | Yes |
| 9 | ENSG00000233276 | GPX1 |  |  | 5834 | 1919.423 | 0.8600 | Yes |
| 10 | ENSG00000146373 | RNF217 |  |  | 5937 | 212.664 | 0.8702 | Yes |
| 11 | ENSG00000180667 | YOD1 |  |  | 5999 | 120.202 | 0.8760 | Yes |
| 12 | ENSG00000175556 | LONRF3 |  |  | 6133 | 59.845 | 0.8771 | Yes |
| 13 | ENSG00000147889 | CDKN2A |  |  | 6154 | 54.378 | 0.8798 | Yes |
| 14 | ENSG00000133135 | RNF128 |  |  | 6586 | 18.128 | 0.8735 | No |
| 15 | ENSG00000185345 | PRKN |  |  | 7293 | 8.647 | 0.8620 | No |
| 16 | ENSG00000174842 | GLMN |  |  | 7355 | 8.258 | 0.8614 | No |
| 17 | ENSG00000188906 | LRRK2 |  |  | 7426 | 7.825 | 0.8607 | No |
| 18 | ENSG00000134758 | RNF138 |  |  | 7630 | 6.849 | 0.8576 | No |
| 19 | ENSG00000101695 | RNF125 |  |  | 7663 | 6.747 | 0.8575 | No |
| 20 | ENSG00000159197 | KCNE2 |  |  | 7697 | 6.622 | 0.8573 | No |
| 21 | ENSG00000138686 | BBS7 |  |  | 7751 | 6.428 | 0.8567 | No |
| 22 | ENSG00000151692 | RNF144A |  |  | 7795 | 6.309 | 0.8564 | No |
| 23 | ENSG00000189283 | FHIT |  |  | 8483 | 4.679 | 0.8450 | No |
| 24 | ENSG00000174672 | BRSK2 |  |  | 8849 | 4.102 | 0.8390 | No |
| 25 | ENSG00000102081 | FMR1 |  |  | 9105 | 3.818 | 0.8349 | No |
| 26 | ENSG00000071575 | TRIB2 |  |  | 9295 | 3.615 | 0.8319 | No |
| 27 | ENSG00000169884 | WNT10B |  |  | 9633 | 3.254 | 0.8263 | No |
| 28 | ENSG00000101843 | PSMD10 |  |  | 10195 | 2.884 | 0.8170 | No |
| 29 | ENSG00000163743 | RCHY1 |  |  | 10423 | 2.728 | 0.8133 | No |
| 30 | ENSG00000145506 | NKD2 |  |  | 10424 | 2.727 | 0.8134 | No |
| 31 | ENSG00000122406 | RPL5 |  |  | 10469 | 2.702 | 0.8128 | No |
| 32 | ENSG00000109861 | CTSC |  |  | 10763 | 2.560 | 0.8080 | No |
| 33 | ENSG00000123384 | LRP1 |  |  | 11225 | 2.369 | 0.8003 | No |
| 34 | ENSG00000137575 | SDCBP |  |  | 11241 | 2.363 | 0.8002 | No |
| 35 | ENSG00000139324 | TMTC3 |  |  | 11257 | 2.356 | 0.8001 | No |
| 36 | ENSG00000177479 | ARIH2 |  |  | 11335 | 2.330 | 0.7989 | No |
| 37 | ENSG00000173473 | SMARCC1 |  |  | 11520 | 2.268 | 0.7959 | No |
| 38 | ENSG00000256061 | DNAAF4 |  |  | 11685 | 2.208 | 0.7932 | No |
| 39 | ENSG00000102921 | N4BP1 |  |  | 11728 | 2.184 | 0.7926 | No |
| 40 | ENSG00000137710 | RDX |  |  | 11742 | 2.181 | 0.7925 | No |
| 41 | ENSG00000080824 | HSP90AA1 |  |  | 11758 | 2.171 | 0.7924 | No |
| 42 | ENSG00000072849 | DERL2 |  |  | 11976 | 2.086 | 0.7888 | No |
| 43 | ENSG00000154359 | LONRF1 |  |  | 12014 | 2.070 | 0.7883 | No |
| 44 | ENSG00000069329 | VPS35 |  |  | 12188 | 2.019 | 0.7855 | No |
| 45 | ENSG00000079387 | SENP1 |  |  | 12195 | 2.017 | 0.7855 | No |
| 46 | ENSG00000013374 | NUB1 |  |  | 12299 | 1.983 | 0.7839 | No |
| 47 | ENSG00000166167 | BTRC |  |  | 12313 | 1.980 | 0.7838 | No |
| 48 | ENSG00000131023 | LATS1 |  |  | 12316 | 1.979 | 0.7838 | No |
| 49 | ENSG00000139174 | PRICKLE1 |  |  | 12408 | 1.953 | 0.7824 | No |
| 50 | ENSG00000163602 | RYBP |  |  | 12489 | 1.926 | 0.7812 | No |
| 51 | ENSG00000213923 | CSNK1E |  |  | 12586 | 1.899 | 0.7796 | No |
| 52 | ENSG00000153071 | DAB2 |  |  | 12594 | 1.898 | 0.7796 | No |
| 53 | ENSG00000109670 | FBXW7 |  |  | 12908 | 1.808 | 0.7744 | No |
| 54 | ENSG00000130988 | RGN |  |  | 12925 | 1.803 | 0.7742 | No |
| 55 | ENSG00000116514 | RNF19B |  |  | 12997 | 1.787 | 0.7731 | No |
| 56 | ENSG00000136159 | NUDT15 |  |  | 13055 | 1.772 | 0.7723 | No |
| 57 | ENSG00000188612 | SUMO2 |  |  | 13057 | 1.772 | 0.7724 | No |
| 58 | ENSG00000169139 | UBE2V2 |  |  | 13287 | 1.716 | 0.7686 | No |
| 59 | ENSG00000131467 | PSME3 |  |  | 13315 | 1.706 | 0.7682 | No |
| 60 | ENSG00000147162 | OGT |  |  | 13373 | 1.692 | 0.7673 | No |
| 61 | ENSG00000171150 | SOCS5 |  |  | 13675 | 1.634 | 0.7623 | No |
| 62 | ENSG00000014123 | UFL1 |  |  | 13698 | 1.628 | 0.7620 | No |
| 63 | ENSG00000101557 | USP14 |  |  | 13762 | 1.618 | 0.7610 | No |
| 64 | ENSG00000184675 | AMER1 |  |  | 13956 | 1.579 | 0.7579 | No |
| 65 | ENSG00000133028 | SCO1 |  |  | 13995 | 1.572 | 0.7573 | No |
| 66 | ENSG00000171863 | RPS7 |  |  | 14019 | 1.565 | 0.7570 | No |
| 67 | ENSG00000125691 | RPL23 |  |  | 14185 | 1.549 | 0.7543 | No |
| 68 | ENSG00000140543 | DET1 |  |  | 14372 | 1.503 | 0.7512 | No |
| 69 | ENSG00000051108 | HERPUD1 |  |  | 14471 | 1.482 | 0.7496 | No |
| 70 | ENSG00000116750 | UCHL5 |  |  | 14604 | 1.469 | 0.7475 | No |
| 71 | ENSG00000066427 | ATXN3 |  |  | 14612 | 1.468 | 0.7474 | No |
| 72 | ENSG00000180008 | SOCS4 |  |  | 14684 | 1.452 | 0.7463 | No |
| 73 | ENSG00000104969 | SGTA |  |  | 14703 | 1.447 | 0.7461 | No |
| 74 | ENSG00000116030 | SUMO1 |  |  | 14718 | 1.444 | 0.7459 | No |
| 75 | ENSG00000100519 | PSMC6 |  |  | 14846 | 1.420 | 0.7439 | No |
| 76 | ENSG00000100387 | RBX1 |  |  | 14891 | 1.411 | 0.7432 | No |
| 77 | ENSG00000033800 | PIAS1 |  |  | 14983 | 1.392 | 0.7417 | No |
| 78 | ENSG00000125818 | PSMF1 |  |  | 15080 | 1.376 | 0.7402 | No |
| 79 | ENSG00000078140 | UBE2K |  |  | 15098 | 1.371 | 0.7400 | No |
| 80 | ENSG00000105875 | WDR91 |  |  | 15203 | 1.353 | 0.7383 | No |
| 81 | ENSG00000100764 | PSMC1 |  |  | 15400 | 1.318 | 0.7350 | No |
| 82 | ENSG00000168078 | PBK |  |  | 15844 | 1.252 | 0.7276 | No |
| 83 | ENSG00000198833 | UBE2J1 |  |  | 15864 | 1.249 | 0.7273 | No |
| 84 | ENSG00000172046 | USP19 |  |  | 15903 | 1.241 | 0.7267 | No |
| 85 | ENSG00000135679 | MDM2 |  |  | 15941 | 1.240 | 0.7262 | No |
| 86 | ENSG00000147133 | TAF1 |  |  | 16016 | 1.231 | 0.7250 | No |
| 87 | ENSG00000107185 | RGP1 |  |  | 16266 | 1.191 | 0.7208 | No |
| 88 | ENSG00000198171 | DDRGK1 |  |  | 16329 | 1.186 | 0.7198 | No |
| 89 | ENSG00000107882 | SUFU |  |  | 16354 | 1.184 | 0.7195 | No |
| 90 | ENSG00000115233 | PSMD14 |  |  | 16355 | 1.184 | 0.7196 | No |
| 91 | ENSG00000144747 | TMF1 |  |  | 16468 | 1.170 | 0.7177 | No |
| 92 | ENSG00000134882 | UBAC2 |  |  | 16549 | 1.160 | 0.7164 | No |
| 93 | ENSG00000114062 | UBE3A |  |  | 16599 | 1.154 | 0.7157 | No |
| 94 | ENSG00000103549 | RNF40 |  |  | 16604 | 1.154 | 0.7157 | No |
| 95 | ENSG00000166170 | BAG5 |  |  | 16634 | 1.150 | 0.7152 | No |
| 96 | ENSG00000134109 | EDEM1 |  |  | 16796 | 1.128 | 0.7126 | No |
| 97 | ENSG00000155313 | USP25 |  |  | 16833 | 1.123 | 0.7120 | No |
| 98 | ENSG00000166233 | ARIH1 |  |  | 17003 | 1.103 | 0.7092 | No |
| 99 | ENSG00000064393 | HIPK2 |  |  | 17051 | 1.096 | 0.7085 | No |
| 100 | ENSG00000113712 | CSNK1A1 |  |  | 17171 | 1.078 | 0.7065 | No |
| 101 | ENSG00000118564 | FBXL5 |  |  | 17179 | 1.077 | 0.7065 | No |
| 102 | ENSG00000188021 | UBQLN2 |  |  | 17223 | 1.069 | 0.7058 | No |
| 103 | ENSG00000013561 | RNF14 |  |  | 17293 | 1.062 | 0.7047 | No |
| 104 | ENSG00000107036 | RIC1 |  |  | 17328 | 1.057 | 0.7042 | No |
| 105 | ENSG00000171100 | MTM1 |  |  | 17613 | 1.024 | 0.6994 | No |
| 106 | ENSG00000068912 | ERLEC1 |  |  | 17767 | 1.004 | 0.6969 | No |
| 107 | ENSG00000126583 | PRKCG |  |  | 17788 | 1.001 | 0.6966 | No |
| 108 | ENSG00000080815 | PSEN1 |  |  | 17938 | 0.983 | 0.6941 | No |
| 109 | ENSG00000116288 | PARK7 |  |  | 18009 | 0.973 | 0.6930 | No |
| 110 | ENSG00000078081 | LAMP3 |  |  | 18038 | 0.969 | 0.6926 | No |
| 111 | ENSG00000274897 | PANO1 |  |  | 18043 | 0.968 | 0.6925 | No |
| 112 | ENSG00000167196 | FBXO22 |  |  | 18164 | 0.953 | 0.6906 | No |
| 113 | ENSG00000058056 | USP13 |  |  | 18224 | 0.948 | 0.6896 | No |
| 114 | ENSG00000184787 | UBE2G2 |  |  | 18542 | 0.911 | 0.6843 | No |
| 115 | ENSG00000143207 | COP1 |  |  | 18787 | 0.883 | 0.6802 | No |
| 116 | ENSG00000137288 | UQCC2 |  |  | 18858 | 0.876 | 0.6790 | No |
| 117 | ENSG00000158828 | PINK1 |  |  | 18880 | 0.874 | 0.6787 | No |
| 118 | ENSG00000001084 | GCLC |  |  | 18902 | 0.871 | 0.6784 | No |
| 119 | ENSG00000088298 | EDEM2 |  |  | 19007 | 0.861 | 0.6767 | No |
| 120 | ENSG00000072062 | PRKACA |  |  | 19008 | 0.861 | 0.6768 | No |
| 121 | ENSG00000138085 | ATRAID |  |  | 19013 | 0.860 | 0.6767 | No |
| 122 | ENSG00000166851 | PLK1 |  |  | 19193 | 0.841 | 0.6738 | No |
| 123 | ENSG00000081479 | LRP2 |  |  | 19231 | 0.837 | 0.6732 | No |
| 124 | ENSG00000198168 | SVIP |  |  | 19239 | 0.837 | 0.6731 | No |
| 125 | ENSG00000182087 | TMEM259 |  |  | 19594 | 0.814 | 0.6671 | No |
| 126 | ENSG00000171862 | PTEN |  |  | 19603 | 0.813 | 0.6670 | No |
| 127 | ENSG00000130311 | DDA1 |  |  | 19655 | 0.808 | 0.6662 | No |
| 128 | ENSG00000187555 | USP7 |  |  | 19733 | 0.799 | 0.6650 | No |
| 129 | ENSG00000034713 | GABARAPL2 |  |  | 19759 | 0.797 | 0.6646 | No |
| 130 | ENSG00000087586 | AURKA |  |  | 19782 | 0.794 | 0.6643 | No |
| 131 | ENSG00000158941 | CCAR2 |  |  | 19800 | 0.792 | 0.6640 | No |
| 132 | ENSG00000170315 | UBB |  |  | 19834 | 0.789 | 0.6635 | No |
| 133 | ENSG00000142676 | RPL11 |  |  | 19931 | 0.781 | 0.6619 | No |
| 134 | ENSG00000161057 | PSMC2 |  |  | 20050 | 0.772 | 0.6599 | No |
| 135 | ENSG00000140464 | PML |  |  | 20085 | 0.767 | 0.6594 | No |
| 136 | ENSG00000130203 | APOE |  |  | 20361 | 0.741 | 0.6548 | No |
| 137 | ENSG00000096384 | HSP90AB1 |  |  | 20585 | 0.733 | 0.6510 | No |
| 138 | ENSG00000245848 | CEBPA |  |  | 20734 | 0.717 | 0.6486 | No |
| 139 | ENSG00000162191 | UBXN1 |  |  | 20744 | 0.716 | 0.6485 | No |
| 140 | ENSG00000115539 | PDCL3 |  |  | 20906 | 0.698 | 0.6458 | No |
| 141 | ENSG00000120899 | PTK2B |  |  | 20968 | 0.692 | 0.6448 | No |
| 142 | ENSG00000135018 | UBQLN1 |  |  | 21423 | 0.648 | 0.6371 | No |
| 143 | ENSG00000116106 | EPHA4 |  |  | 21439 | 0.647 | 0.6369 | No |
| 144 | ENSG00000095787 | WAC |  |  | 21505 | 0.639 | 0.6358 | No |
| 145 | ENSG00000160803 | UBQLN4 |  |  | 21588 | 0.633 | 0.6345 | No |
| 146 | ENSG00000105723 | GSK3A |  |  | 21613 | 0.631 | 0.6341 | No |
| 147 | ENSG00000179262 | RAD23A |  |  | 21700 | 0.623 | 0.6327 | No |
| 148 | ENSG00000082701 | GSK3B |  |  | 21709 | 0.622 | 0.6326 | No |
| 149 | ENSG00000110651 | CD81 |  |  | 21918 | 0.606 | 0.6291 | No |
| 150 | ENSG00000141551 | CSNK1D |  |  | 21933 | 0.604 | 0.6289 | No |
| 151 | ENSG00000119318 | RAD23B |  |  | 22198 | 0.582 | 0.6244 | No |
| 152 | ENSG00000133265 | HSPBP1 |  |  | 22242 | 0.579 | 0.6237 | No |
| 153 | ENSG00000104341 | LAPTM4B |  |  | 22280 | 0.575 | 0.6231 | No |
| 154 | ENSG00000177628 | GBA |  |  | 22360 | 0.570 | 0.6218 | No |
| 155 | ENSG00000165916 | PSMC3 |  |  | 22372 | 0.569 | 0.6217 | No |
| 156 | ENSG00000169398 | PTK2 |  |  | 22456 | 0.560 | 0.6203 | No |
| 157 | ENSG00000173846 | PLK3 |  |  | 22715 | 0.545 | 0.6159 | No |
| 158 | ENSG00000204628 | RACK1 |  |  | 22728 | 0.545 | 0.6158 | No |
| 159 | ENSG00000185825 | BCAP31 |  |  | 22743 | 0.543 | 0.6155 | No |
| 160 | ENSG00000146535 | GNA12 |  |  | 22854 | 0.532 | 0.6137 | No |
| 161 | ENSG00000116044 | NFE2L2 |  |  | 22919 | 0.527 | 0.6126 | No |
| 162 | ENSG00000038532 | CLEC16A |  |  | 22939 | 0.524 | 0.6124 | No |
| 163 | ENSG00000173334 | TRIB1 |  |  | 23092 | 0.512 | 0.6098 | No |
| 164 | ENSG00000001629 | ANKIB1 |  |  | 23259 | 0.499 | 0.6070 | No |
| 165 | ENSG00000146872 | TLK2 |  |  | 23496 | 0.481 | 0.6030 | No |
| 166 | ENSG00000092820 | EZR |  |  | 23520 | 0.479 | 0.6027 | No |
| 167 | ENSG00000159363 | ATP13A2 |  |  | 23569 | 0.474 | 0.6019 | No |
| 168 | ENSG00000177383 | MAGEF1 |  |  | 23587 | 0.473 | 0.6016 | No |
| 169 | ENSG00000158717 | RNF166 |  |  | 23656 | 0.469 | 0.6005 | No |
| 170 | ENSG00000108465 | CDK5RAP3 |  |  | 23660 | 0.469 | 0.6005 | No |
| 171 | ENSG00000165280 | VCP |  |  | 23913 | 0.447 | 0.5962 | No |
| 172 | ENSG00000101997 | CCDC22 |  |  | 24015 | 0.438 | 0.5945 | No |
| 173 | ENSG00000118503 | TNFAIP3 |  |  | 24022 | 0.438 | 0.5944 | No |
| 174 | ENSG00000107404 | DVL1 |  |  | 24030 | 0.437 | 0.5944 | No |
| 175 | ENSG00000173163 | COMMD1 |  |  | 24116 | 0.431 | 0.5929 | No |
| 176 | ENSG00000198742 | SMURF1 |  |  | 24140 | 0.428 | 0.5926 | No |
| 177 | ENSG00000008710 | PKD1 |  |  | 24191 | 0.424 | 0.5917 | No |
| 178 | ENSG00000103126 | AXIN1 |  |  | 24346 | 0.411 | 0.5892 | No |
| 179 | ENSG00000103266 | STUB1 |  |  | 24447 | 0.406 | 0.5875 | No |
| 180 | ENSG00000034677 | RNF19A |  |  | 24502 | 0.402 | 0.5866 | No |
| 181 | ENSG00000170881 | RNF139 |  |  | 24637 | 0.391 | 0.5843 | No |
| 182 | ENSG00000105373 | NOP53 |  |  | 24639 | 0.390 | 0.5843 | No |
| 183 | ENSG00000079999 | KEAP1 |  |  | 24664 | 0.388 | 0.5840 | No |
| 184 | ENSG00000111667 | USP5 |  |  | 24694 | 0.386 | 0.5835 | No |
| 185 | ENSG00000119283 | TRIM67 |  |  | 25011 | 0.365 | 0.5781 | No |
| 186 | ENSG00000100911 | PSME2 |  |  | 25048 | 0.361 | 0.5775 | No |
| 187 | ENSG00000101665 | SMAD7 |  |  | 25100 | 0.358 | 0.5767 | No |
| 188 | ENSG00000078061 | ARAF |  |  | 25251 | 0.347 | 0.5742 | No |
| 189 | ENSG00000079482 | OPHN1 |  |  | 25299 | 0.343 | 0.5734 | No |
| 190 | ENSG00000013275 | PSMC4 |  |  | 25323 | 0.341 | 0.5730 | No |
| 191 | ENSG00000170500 | LONRF2 |  |  | 25412 | 0.334 | 0.5715 | No |
| 192 | ENSG00000124226 | RNF114 |  |  | 25447 | 0.331 | 0.5710 | No |
| 193 | ENSG00000136848 | DAB2IP |  |  | 25645 | 0.315 | 0.5677 | No |
| 194 | ENSG00000183255 | PTTG1IP |  |  | 25760 | 0.307 | 0.5657 | No |
| 195 | ENSG00000087191 | PSMC5 |  |  | 25832 | 0.300 | 0.5646 | No |
| 196 | ENSG00000092010 | PSME1 |  |  | 26052 | 0.281 | 0.5609 | No |
| 197 | ENSG00000135506 | OS9 |  |  | 26213 | 0.272 | 0.5582 | No |
| 198 | ENSG00000140564 | FURIN |  |  | 26415 | 0.252 | 0.5548 | No |
| 199 | ENSG00000148218 | ALAD |  |  | 26439 | 0.249 | 0.5544 | No |
| 200 | ENSG00000178381 | ZFAND2A |  |  | 26537 | 0.244 | 0.5527 | No |
| 201 | ENSG00000072609 | CHFR |  |  | 26691 | 0.230 | 0.5502 | No |
| 202 | ENSG00000123159 | GIPC1 |  |  | 26798 | 0.222 | 0.5484 | No |
| 203 | ENSG00000007944 | MYLIP |  |  | 26896 | 0.212 | 0.5467 | No |
| 204 | ENSG00000105974 | CAV1 |  |  | 27266 | 0.183 | 0.5405 | No |
| 205 | ENSG00000138798 | EGF |  |  | 27284 | 0.181 | 0.5402 | No |
| 206 | ENSG00000130164 | LDLR |  |  | 27347 | 0.175 | 0.5392 | No |
| 207 | ENSG00000182718 | ANXA2 |  |  | 27588 | 0.156 | 0.5351 | No |
| 208 | ENSG00000101255 | TRIB3 |  |  | 27621 | 0.153 | 0.5346 | No |
| 209 | ENSG00000142208 | AKT1 |  |  | 27630 | 0.152 | 0.5344 | No |
| 210 | ENSG00000120885 | CLU |  |  | 27731 | 0.143 | 0.5327 | No |
| 211 | ENSG00000006025 | OSBPL7 |  |  | 28336 | 0.091 | 0.5225 | No |
| 212 | ENSG00000164197 | RNF180 |  |  | 28439 | 0.082 | 0.5208 | No |
| 213 | ENSG00000100219 | XBP1 |  |  | 28464 | 0.079 | 0.5204 | No |
| 214 | ENSG00000125084 | WNT1 |  |  | 28524 | 0.074 | 0.5194 | No |
| 215 | ENSG00000007384 | RHBDF1 |  |  | 28650 | 0.063 | 0.5173 | No |
| 216 | ENSG00000137393 | RNF144B |  |  | 28959 | 0.037 | 0.5120 | No |
| 217 | ENSG00000169242 | EFNA1 |  |  | 29153 | 0.024 | 0.5088 | No |
| 218 | ENSG00000145632 | PLK2 |  |  | 29161 | 0.023 | 0.5086 | No |
| 219 | ENSG00000010704 | HFE |  |  | 29208 | 0.020 | 0.5079 | No |
| 220 | ENSG00000164690 | SHH |  |  | 29438 | 0.008 | 0.5040 | No |
| 221 | ENSG00000129993 | CBFA2T3 |  |  | 29443 | 0.008 | 0.5039 | No |
| 222 | ENSG00000204389 | HSPA1A |  |  | 32444 | 0.000 | 0.4530 | No |
| 223 | ENSG00000273841 | TAF9 |  |  | 34265 | NaN | 0.4221 | No |
| 224 | ENSG00000160695 | VPS11 |  |  | 35385 | NaN | 0.4031 | No |
| 225 | ENSG00000099958 | DERL3 |  |  | 36499 | NaN | 0.3842 | No |
| 226 | ENSG00000163564 | PYHIN1 |  |  | 38478 | NaN | 0.3506 | No |
| 227 | ENSG00000130770 | ATP5IF1 |  |  | 52680 | NaN | 0.1095 | No |
| 228 | ENSG00000068903 | SIRT2 |  |  | 53252 | NaN | 0.0998 | No |
| 229 | ENSG00000204599 | TRIM39 |  |  | 53424 | NaN | 0.0969 | No |
| 230 | ENSG00000249751 | ECSCR |  |  | 54918 | NaN | 0.0716 | No |
Table: GSEA details
[plain text format]

  

Fig 2: REGULATION\_OF\_CELLULAR\_PROTEIN\_CATABOLIC\_PROCESS(GO:1903362)
  
Blue-Pink O' Gram in the Space of the Analyzed GeneSet

  

Fig 3: REGULATION\_OF\_CELLULAR\_PROTEIN\_CATABOLIC\_PROCESS(GO:1903362): Random ES distribution
  
Gene set null distribution of ES for
 **REGULATION\_OF\_CELLULAR\_PROTEIN\_CATABOLIC\_PROCESS(GO:1903362)**

  
