## Supplementary material for "A single cysteine residue in vimentin regulates long non-coding RNA *XIST* to suppress epithelial-mesenchymal transition and stemness in breast cancer": REGULATION_OF_TUMOR_NECROSIS_FACTOR_MEDIATED_SIGNALING_PATHWAY(GO_0010803).html

Details for gene set REGULATION\_OF\_TUMOR\_NECROSIS\_FACTOR\_MEDIATED\_SIGNALING\_PATHWAY(GO:0010803)[GSEA]


|  || Dataset | fpkm.sample |
| Phenotype | sample.cls |
| Upregulated in class | C328SVIM |
| GeneSet | REGULATION\_OF\_TUMOR\_NECROSIS\_FACTOR\_MEDIATED\_SIGNALING\_PATHWAY(GO:0010803) |
| Enrichment Score (ES) | 0.8915991 |
| Normalized Enrichment Score (NES) | 1.4411669 |
| Nominal p-value | 0.0 |
| FDR q-value | 1.0 |
| FWER p-Value | 0.325 |
Table: GSEA Results Summary

  

Fig 1: Enrichment plot: REGULATION\_OF\_TUMOR\_NECROSIS\_FACTOR\_MEDIATED\_SIGNALING\_PATHWAY(GO:0010803)
  
Profile of the Running ES Score & Positions of GeneSet Members on the Rank Ordered List

  

| PROBE | DESCRIPTION   (from dataset) | GENE SYMBOL | GENE\_TITLE | RANK IN GENE LIST | RANK METRIC SCORE | RUNNING ES | CORE ENRICHMENT || 1 | ENSG00000084207 | GSTP1 |  |  | 5825 | 8120.758 | 0.8618 | Yes |
| 2 | ENSG00000151468 | CCDC3 |  |  | 5922 | 265.492 | 0.8916 | Yes |
| 3 | ENSG00000204388 | HSPA1B |  |  | 6626 | 16.900 | 0.8817 | No |
| 4 | ENSG00000185345 | PRKN |  |  | 7293 | 8.647 | 0.8714 | No |
| 5 | ENSG00000083799 | CYLD |  |  | 8931 | 4.011 | 0.8442 | No |
| 6 | ENSG00000105483 | CARD8 |  |  | 9648 | 3.241 | 0.8325 | No |
| 7 | ENSG00000110330 | BIRC2 |  |  | 10411 | 2.736 | 0.8199 | No |
| 8 | ENSG00000118137 | APOA1 |  |  | 10458 | 2.708 | 0.8194 | No |
| 9 | ENSG00000056558 | TRAF1 |  |  | 10751 | 2.565 | 0.8148 | No |
| 10 | ENSG00000213341 | CHUK |  |  | 11516 | 2.270 | 0.8021 | No |
| 11 | ENSG00000106052 | TAX1BP1 |  |  | 13023 | 1.779 | 0.7769 | No |
| 12 | ENSG00000164251 | F2RL1 |  |  | 13678 | 1.633 | 0.7660 | No |
| 13 | ENSG00000183763 | TRAIP |  |  | 14406 | 1.496 | 0.7538 | No |
| 14 | ENSG00000154124 | OTULIN |  |  | 16031 | 1.229 | 0.7265 | No |
| 15 | ENSG00000104365 | IKBKB |  |  | 16147 | 1.210 | 0.7247 | No |
| 16 | ENSG00000092871 | RFFL |  |  | 16377 | 1.181 | 0.7210 | No |
| 17 | ENSG00000183087 | GAS6 |  |  | 16662 | 1.148 | 0.7163 | No |
| 18 | ENSG00000143947 | RPS27A |  |  | 16724 | 1.137 | 0.7154 | No |
| 19 | ENSG00000151694 | ADAM17 |  |  | 16892 | 1.115 | 0.7127 | No |
| 20 | ENSG00000175354 | PTPN2 |  |  | 17052 | 1.096 | 0.7101 | No |
| 21 | ENSG00000105270 | CLIP3 |  |  | 19318 | 0.828 | 0.6719 | No |
| 22 | ENSG00000119401 | TRIM32 |  |  | 19565 | 0.818 | 0.6678 | No |
| 23 | ENSG00000197372 | ZNF675 |  |  | 19710 | 0.802 | 0.6655 | No |
| 24 | ENSG00000170315 | UBB |  |  | 19834 | 0.789 | 0.6635 | No |
| 25 | ENSG00000105229 | PIAS4 |  |  | 19856 | 0.787 | 0.6632 | No |
| 26 | ENSG00000005206 | SPPL2B |  |  | 20034 | 0.774 | 0.6603 | No |
| 27 | ENSG00000214078 | CPNE1 |  |  | 20339 | 0.743 | 0.6553 | No |
| 28 | ENSG00000064012 | CASP8 |  |  | 21525 | 0.638 | 0.6353 | No |
| 29 | ENSG00000137275 | RIPK1 |  |  | 22606 | 0.555 | 0.6171 | No |
| 30 | ENSG00000204628 | RACK1 |  |  | 22728 | 0.545 | 0.6151 | No |
| 31 | ENSG00000165025 | SYK |  |  | 22874 | 0.531 | 0.6127 | No |
| 32 | ENSG00000221983 | UBA52 |  |  | 23234 | 0.501 | 0.6067 | No |
| 33 | ENSG00000127191 | TRAF2 |  |  | 23284 | 0.496 | 0.6059 | No |
| 34 | ENSG00000118503 | TNFAIP3 |  |  | 24022 | 0.438 | 0.5935 | No |
| 35 | ENSG00000125826 | RBCK1 |  |  | 24124 | 0.430 | 0.5918 | No |
| 36 | ENSG00000176170 | SPHK1 |  |  | 24712 | 0.385 | 0.5819 | No |
| 37 | ENSG00000174516 | PELI3 |  |  | 24838 | 0.375 | 0.5799 | No |
| 38 | ENSG00000179526 | SHARPIN |  |  | 25588 | 0.319 | 0.5672 | No |
| 39 | ENSG00000269335 | IKBKG |  |  | 25644 | 0.315 | 0.5663 | No |
| 40 | ENSG00000150991 | UBC |  |  | 26040 | 0.281 | 0.5597 | No |
| 41 | ENSG00000110514 | MADD |  |  | 26261 | 0.267 | 0.5560 | No |
| 42 | ENSG00000138600 | SPPL2A |  |  | 26609 | 0.237 | 0.5501 | No |
| 43 | ENSG00000102871 | TRADD |  |  | 26880 | 0.213 | 0.5456 | No |
| 44 | ENSG00000067182 | TNFRSF1A |  |  | 27262 | 0.183 | 0.5392 | No |
| 45 | ENSG00000092098 | RNF31 |  |  | 27479 | 0.163 | 0.5355 | No |
| 46 | ENSG00000163349 | HIPK1 |  |  | 27651 | 0.149 | 0.5326 | No |
| 47 | ENSG00000140939 | NOL3 |  |  | 28154 | 0.106 | 0.5242 | No |
| 48 | ENSG00000023445 | BIRC3 |  |  | 28193 | 0.102 | 0.5235 | No |
| 49 | ENSG00000124635 | HIST1H2BJ |  |  | 28626 | 0.066 | 0.5162 | No |
| 50 | ENSG00000169900 | PYDC1 |  |  | 28792 | 0.051 | 0.5134 | No |
| 51 | ENSG00000103490 | PYCARD |  |  | 28865 | 0.044 | 0.5122 | No |
| 52 | ENSG00000204389 | HSPA1A |  |  | 32444 | 0.000 | 0.4517 | No |
| 53 | ENSG00000196954 | CASP4 |  |  | 32683 | 0.000 | 0.4476 | No |
| 54 | ENSG00000012504 | NR1H4 |  |  | 32894 | 0.000 | 0.4441 | No |
| 55 | ENSG00000253548 | PYDC2 |  |  | 36411 | NaN | 0.3846 | No |
| 56 | ENSG00000215174 | NLRP2B |  |  | 50273 | NaN | 0.1499 | No |
| 57 | ENSG00000204397 | CARD16 |  |  | 51193 | NaN | 0.1344 | No |
| 58 | ENSG00000181092 | ADIPOQ |  |  | 53064 | NaN | 0.1027 | No |
| 59 | ENSG00000137752 | CASP1 |  |  | 54443 | NaN | 0.0794 | No |
| 60 | ENSG00000232810 | TNF |  |  | 57062 | NaN | 0.0351 | No |
| 61 | ENSG00000207947 | MIR152 |  |  | 58732 | NaN | 0.0068 | No |
Table: GSEA details
[plain text format]

  

Fig 2: REGULATION\_OF\_TUMOR\_NECROSIS\_FACTOR\_MEDIATED\_SIGNALING\_PATHWAY(GO:0010803)
  
Blue-Pink O' Gram in the Space of the Analyzed GeneSet

  

Fig 3: REGULATION\_OF\_TUMOR\_NECROSIS\_FACTOR\_MEDIATED\_SIGNALING\_PATHWAY(GO:0010803): Random ES distribution
  
Gene set null distribution of ES for
 **REGULATION\_OF\_TUMOR\_NECROSIS\_FACTOR\_MEDIATED\_SIGNALING\_PATHWAY(GO:0010803)**

  
