## Supplementary material for "A single cysteine residue in vimentin regulates long non-coding RNA *XIST* to suppress epithelial-mesenchymal transition and stemness in breast cancer": VESICLE_MEDIATED_TRANSPORT_BETWEEN_ENDOSOMAL_COMPARTMENTS(GO_0098927).html

Details for gene set VESICLE\_MEDIATED\_TRANSPORT\_BETWEEN\_ENDOSOMAL\_COMPARTMENTS(GO:0098927)[GSEA]


|  || Dataset | fpkm.sample |
| Phenotype | sample.cls |
| Upregulated in class | C328SVIM |
| GeneSet | VESICLE\_MEDIATED\_TRANSPORT\_BETWEEN\_ENDOSOMAL\_COMPARTMENTS(GO:0098927) |
| Enrichment Score (ES) | 0.8988307 |
| Normalized Enrichment Score (NES) | 1.4558434 |
| Nominal p-value | 0.0 |
| FDR q-value | 1.0 |
| FWER p-Value | 0.325 |
Table: GSEA Results Summary

  

Fig 1: Enrichment plot: VESICLE\_MEDIATED\_TRANSPORT\_BETWEEN\_ENDOSOMAL\_COMPARTMENTS(GO:0098927)
  
Profile of the Running ES Score & Positions of GeneSet Members on the Rank Ordered List

  

| PROBE | DESCRIPTION   (from dataset) | GENE SYMBOL | GENE\_TITLE | RANK IN GENE LIST | RANK METRIC SCORE | RUNNING ES | CORE ENRICHMENT || 1 | ENSG00000147065 | MSN |  |  | 5823 | 15136.710 | 0.8988 | Yes |
| 2 | ENSG00000168172 | HOOK3 |  |  | 9332 | 3.574 | 0.8397 | No |
| 3 | ENSG00000087053 | MTMR2 |  |  | 9520 | 3.368 | 0.8368 | No |
| 4 | ENSG00000078142 | PIK3C3 |  |  | 9875 | 3.078 | 0.8310 | No |
| 5 | ENSG00000134709 | HOOK1 |  |  | 9987 | 3.013 | 0.8293 | No |
| 6 | ENSG00000137710 | RDX |  |  | 11742 | 2.181 | 0.7998 | No |
| 7 | ENSG00000153071 | DAB2 |  |  | 12594 | 1.898 | 0.7855 | No |
| 8 | ENSG00000170310 | STX8 |  |  | 14439 | 1.489 | 0.7544 | No |
| 9 | ENSG00000102189 | EEA1 |  |  | 14683 | 1.453 | 0.7504 | No |
| 10 | ENSG00000105875 | WDR91 |  |  | 15203 | 1.353 | 0.7417 | No |
| 11 | ENSG00000138246 | DNAJC13 |  |  | 16466 | 1.170 | 0.7204 | No |
| 12 | ENSG00000100030 | MAPK1 |  |  | 16571 | 1.158 | 0.7187 | No |
| 13 | ENSG00000051009 | FAM160A2 |  |  | 17031 | 1.099 | 0.7110 | No |
| 14 | ENSG00000126581 | BECN1 |  |  | 17082 | 1.090 | 0.7102 | No |
| 15 | ENSG00000112335 | SNX3 |  |  | 17188 | 1.076 | 0.7085 | No |
| 16 | ENSG00000105186 | ANKRD27 |  |  | 17576 | 1.027 | 0.7021 | No |
| 17 | ENSG00000089177 | KIF16B |  |  | 17800 | 1.000 | 0.6983 | No |
| 18 | ENSG00000159210 | SNF8 |  |  | 19701 | 0.803 | 0.6662 | No |
| 19 | ENSG00000176658 | MYO1D |  |  | 19993 | 0.778 | 0.6614 | No |
| 20 | ENSG00000137642 | SORL1 |  |  | 20346 | 0.742 | 0.6555 | No |
| 21 | ENSG00000115561 | CHMP3 |  |  | 20348 | 0.742 | 0.6555 | No |
| 22 | ENSG00000080371 | RAB21 |  |  | 20357 | 0.741 | 0.6554 | No |
| 23 | ENSG00000167716 | WDR81 |  |  | 20595 | 0.732 | 0.6514 | No |
| 24 | ENSG00000169032 | MAP2K1 |  |  | 20668 | 0.724 | 0.6503 | No |
| 25 | ENSG00000147164 | SNX12 |  |  | 21048 | 0.685 | 0.6439 | No |
| 26 | ENSG00000164715 | LMTK2 |  |  | 21981 | 0.599 | 0.6282 | No |
| 27 | ENSG00000126934 | MAP2K2 |  |  | 22444 | 0.562 | 0.6204 | No |
| 28 | ENSG00000144566 | RAB5A |  |  | 22605 | 0.555 | 0.6177 | No |
| 29 | ENSG00000104497 | SNX16 |  |  | 22925 | 0.526 | 0.6124 | No |
| 30 | ENSG00000197122 | SRC |  |  | 23479 | 0.482 | 0.6030 | No |
| 31 | ENSG00000092820 | EZR |  |  | 23520 | 0.479 | 0.6024 | No |
| 32 | ENSG00000075785 | RAB7A |  |  | 23825 | 0.454 | 0.5973 | No |
| 33 | ENSG00000076201 | PTPN23 |  |  | 24138 | 0.428 | 0.5920 | No |
| 34 | ENSG00000102882 | MAPK3 |  |  | 25118 | 0.357 | 0.5755 | No |
| 35 | ENSG00000166971 | AKTIP |  |  | 26721 | 0.228 | 0.5484 | No |
| 36 | ENSG00000213853 | EMP2 |  |  | 27502 | 0.161 | 0.5352 | No |
| 37 | ENSG00000095066 | HOOK2 |  |  | 28337 | 0.091 | 0.5211 | No |
| 38 | ENSG00000102879 | CORO1A |  |  | 28572 | 0.070 | 0.5171 | No |
| 39 | ENSG00000160695 | VPS11 |  |  | 35385 | NaN | 0.4019 | No |
| 40 | ENSG00000167705 | RILP |  |  | 52245 | NaN | 0.1166 | No |
Table: GSEA details
[plain text format]

  

Fig 2: VESICLE\_MEDIATED\_TRANSPORT\_BETWEEN\_ENDOSOMAL\_COMPARTMENTS(GO:0098927)
  
Blue-Pink O' Gram in the Space of the Analyzed GeneSet

  

Fig 3: VESICLE\_MEDIATED\_TRANSPORT\_BETWEEN\_ENDOSOMAL\_COMPARTMENTS(GO:0098927): Random ES distribution
  
Gene set null distribution of ES for
 **VESICLE\_MEDIATED\_TRANSPORT\_BETWEEN\_ENDOSOMAL\_COMPARTMENTS(GO:0098927)**

  
