## Supplementary material for "A single cysteine residue in vimentin regulates long non-coding RNA *XIST* to suppress epithelial-mesenchymal transition and stemness in breast cancer": heat_map_corr_plot.html

Heat map and correlation plot for fpkm  

Fig 1: heat\_map      
 Heat Map of the top 50 features for each phenotype in fpkm

  
  

Fig 2: Ranked Gene List Correlation Profile      
 Ranked list correlations for fpkm

  
  
    
