## Supplementary material for "A single cysteine residue in vimentin regulates long non-coding RNA *XIST* to suppress epithelial-mesenchymal transition and stemness in breast cancer": index.html

Index for xtools.gsea.Gsea C328SVIMvsWT\_VIM.Gsea.1625876782425


### GSEA Report for Dataset fpkm

#### Enrichment in phenotype: **C328SVIM (2 samples)**

- 7109 / 7113 gene sets are upregulated in phenotype
   **C328SVIM**
- 0 gene sets are significant at FDR < 25%
- 2116 gene sets are significantly enriched at nominal pvalue < 1%
- 2116 gene sets are significantly enriched at nominal pvalue < 5%
- Snapshot
  of enrichment results
- Detailed
  enrichment results in html
  format
- Detailed
  enrichment results in excel
  format (tab delimited text)
- Guide to
  interpret results

#### Enrichment in phenotype: **WT\_VIM (2 samples)**

- 4 / 7113 gene sets are upregulated in phenotype
   **WT\_VIM**
- 0 gene sets are significantly enriched at FDR < 25%
- 3 gene sets are significantly enriched at nominal pvalue < 1%
- 3 gene sets are significantly enriched at nominal pvalue < 5%
- Snapshot
  of enrichment results
- Detailed
  enrichment results in html
  format
- Detailed
  enrichment results in excel
  format (tab delimited text)
- Guide to
  interpret results

#### Dataset details

- The dataset has 59136 features (genes)
- No probe set => gene symbol collapsing was requested, so all 59136 features were used

#### Gene set details

- Gene set size filters (min=15, max=5000) resulted in filtering out 15264 / 22377 gene sets
- The remaining 7113 gene sets were used in the analysis
- List of
  gene sets used and their sizes
  (restricted to features in the specified dataset)

#### Gene markers for the **C328SVIM** *versus* **WT\_VIM** comparison

- The dataset has 59136 features (genes)
- # of markers for phenotype
   **C328SVIM** 
  : 55815 (94.4% ) with correlation area NaN%
- # of markers for phenotype
   **WT\_VIM** 
  : 3321 (5.6% ) with correlation area NaN%
- Detailed
  rank ordered gene list
  for all features in the dataset
- Heat map and gene list correlation
  profile for all features in the dataset
- Buttefly plot
  of significant genes

#### Global statistics and plots

- Plot of
  p-values
   *vs.* 
  NES
- Global ES
  histogram

#### Other

- Parameters
  used for this analysis

#### Comments

- Timestamp used as random seed: 1625877565679
- Warning: Phenotype permutation was performed but the number of samples in class A is < 7, phenotype: sample.cls
- Warning: Phenotype permutation was performed but the number of samples in class B is < 7, phenotype: sample.cls
- With small datasets, there might not be enough random permutations of sample labels to generate a sufficient null distribution. In such cases, gene\_set randomization might be a better choice.

---

Report: C328SVIMvsWT\_VIM.Gsea.1625876782425.rpt   by user: AI\_ref

xtools.gsea.Gsea [Fri, Jul 9, '21 8 PM 26]

Website:
www.gsea-msigdb.org/gsea
Questions & Suggestions:
Contact page
