## Supplementary material for "A single cysteine residue in vimentin regulates long non-coding RNA *XIST* to suppress epithelial-mesenchymal transition and stemness in breast cancer": gsea_report_for_WT_VIM_1625876782425.html

Report for WT\_VIM 1625876782425 [GSEA]


| GS   follow link to MSigDB | GS DETAILS | SIZE | ES | NES | NOM p-val | FDR q-val | FWER p-val | RANK AT MAX | LEADING EDGE || 1 | POSITIVE\_REGULATION\_OF\_PEPTIDYL\_SERINE\_PHOSPHORYLATION\_OF\_STAT\_PROTEIN(GO:0033141) | Details ... | 21 | -0.29 | -1.00 | 0.000 | 1.000 | 0.141 | 32200 | tags=67%, list=54%, signal=146% |
| 2 | IMMUNOGLOBULIN\_RECEPTOR\_BINDING(GO:0034987) | Details ... | 58 | -0.34 | -1.00 | 0.000 | 1.000 | 0.141 | 38696 | tags=98%, list=65%, signal=284% |
| 3 | IMMUNOGLOBULIN\_COMPLEX\_\_CIRCULATING(GO:0042571) | Details ... | 54 | -0.34 | -1.00 | 0.000 | 1.000 | 0.141 | 38696 | tags=98%, list=65%, signal=284% |
| 4 | TYPE\_I\_INTERFERON\_RECEPTOR\_BINDING(GO:0005132) | Details ... | 17 | -0.29 | -0.70 | 0.434 | 1.000 | 0.325 | 19668 | tags=59%, list=33%, signal=88% |
Table: Gene sets enriched in phenotype
 **WT\_VIM (2 samples)
 **[plain text format]****

  
