## Supplementary material for "A single cysteine residue in vimentin regulates long non-coding RNA *XIST* to suppress epithelial-mesenchymal transition and stemness in breast cancer": IMMUNOGLOBULIN_COMPLEX__CIRCULATING(GO_0042571).html

Details for gene set IMMUNOGLOBULIN\_COMPLEX\_\_CIRCULATING(GO:0042571)[GSEA]


|  || Dataset | fpkm.sample |
| Phenotype | sample.cls |
| Upregulated in class | WT\_VIM |
| GeneSet | IMMUNOGLOBULIN\_COMPLEX\_\_CIRCULATING(GO:0042571) |
| Enrichment Score (ES) | -0.33781597 |
| Normalized Enrichment Score (NES) | -1.0 |
| Nominal p-value | 0.0 |
| FDR q-value | 1.0 |
| FWER p-Value | 0.141 |
Table: GSEA Results Summary

  

Fig 1: Enrichment plot: IMMUNOGLOBULIN\_COMPLEX\_\_CIRCULATING(GO:0042571)
  
Profile of the Running ES Score & Positions of GeneSet Members on the Rank Ordered List

  

| PROBE | DESCRIPTION   (from dataset) | GENE SYMBOL | GENE\_TITLE | RANK IN GENE LIST | RANK METRIC SCORE | RUNNING ES | CORE ENRICHMENT || 1 | ENSG00000128322 | IGLL1 |  |  | 4656 | Infinity | -0.0707 | Yes |
| 2 | ENSG00000132465 | JCHAIN |  |  | 20441 | 0.738 | 0.2631 | Yes |
| 3 | ENSG00000224650 | IGHV3-74 |  |  | 30820 | 0.000 | 0.0875 | No |
| 4 | ENSG00000211893 | IGHG2 |  |  | 32101 | 0.000 | 0.0658 | No |
| 5 | ENSG00000211892 | IGHG4 |  |  | 32604 | 0.000 | 0.0573 | No |
| 6 | ENSG00000211896 | IGHG1 |  |  | 32605 | 0.000 | 0.0573 | No |
| 7 | ENSG00000211947 | IGHV3-21 |  |  | 33401 | NaN | 0.0520 | No |
| 8 | ENSG00000211945 | IGHV1-18 |  |  | 33403 | NaN | 0.0601 | No |
| 9 | ENSG00000211935 | IGHV1-3 |  |  | 34044 | NaN | 0.0574 | No |
| 10 | ENSG00000232216 | IGHV3-43 |  |  | 37879 | NaN | 0.0007 | No |
| 11 | ENSG00000211890 | IGHA2 |  |  | 37893 | NaN | 0.0086 | No |
| 12 | ENSG00000211950 | IGHV1-24 |  |  | 39151 | NaN | -0.0045 | No |
| 13 | ENSG00000211951 | IGHV2-26 |  |  | 40699 | NaN | -0.0226 | No |
| 14 | ENSG00000211952 | IGHV4-28 |  |  | 40700 | NaN | -0.0144 | No |
| 15 | ENSG00000211957 | IGHV3-35 |  |  | 40703 | NaN | -0.0063 | No |
| 16 | ENSG00000211958 | IGHV3-38 |  |  | 40704 | NaN | 0.0018 | No |
| 17 | ENSG00000211979 | IGHV7-81 |  |  | 41796 | NaN | -0.0085 | No |
| 18 | ENSG00000211972 | IGHV3-66 |  |  | 41800 | NaN | -0.0004 | No |
| 19 | ENSG00000231475 | IGHV4-31 |  |  | 42071 | NaN | 0.0032 | No |
| 20 | ENSG00000270472 | IGHV3OR16-9 |  |  | 42516 | NaN | 0.0038 | No |
| 21 | ENSG00000188403 | IGHV1OR15-9 |  |  | 42837 | NaN | 0.0065 | No |
| 22 | ENSG00000223648 | IGHV3-64 |  |  | 43450 | NaN | 0.0043 | No |
| 23 | ENSG00000239951 | IGKV3-20 |  |  | 43867 | NaN | 0.0054 | No |
| 24 | ENSG00000271178 | IGHV3OR16-13 |  |  | 44717 | NaN | -0.0008 | No |
| 25 | ENSG00000259680 | AC136428.1 |  |  | 44852 | NaN | 0.0051 | No |
| 26 | ENSG00000270505 | IGHV1OR15-1 |  |  | 45965 | NaN | -0.0056 | No |
| 27 | ENSG00000281179 | AC135068.8 |  |  | 46591 | NaN | -0.0080 | No |
| 28 | ENSG00000211933 | IGHV6-1 |  |  | 47067 | NaN | -0.0079 | No |
| 29 | ENSG00000211970 | IGHV4-61 |  |  | 47532 | NaN | -0.0076 | No |
| 30 | ENSG00000211976 | IGHV3-73 |  |  | 47533 | NaN | 0.0005 | No |
| 31 | ENSG00000211974 | IGHV2-70D |  |  | 47534 | NaN | 0.0087 | No |
| 32 | ENSG00000225698 | IGHV3-72 |  |  | 48227 | NaN | 0.0051 | No |
| 33 | ENSG00000270467 | IGHV3OR16-12 |  |  | 48449 | NaN | 0.0095 | No |
| 34 | ENSG00000280411 | IGHV1-69D |  |  | 48735 | NaN | 0.0128 | No |
| 35 | ENSG00000278263 | AC135068.2 |  |  | 49083 | NaN | 0.0151 | No |
| 36 | ENSG00000254709 | IGLL5 |  |  | 49907 | NaN | 0.0093 | No |
| 37 | ENSG00000211943 | IGHV3-15 |  |  | 50481 | NaN | 0.0077 | No |
| 38 | ENSG00000211946 | IGHV3-20 |  |  | 50485 | NaN | 0.0158 | No |
| 39 | ENSG00000211944 | IGHV3-16 |  |  | 50486 | NaN | 0.0240 | No |
| 40 | ENSG00000211949 | IGHV3-23 |  |  | 50487 | NaN | 0.0321 | No |
| 41 | ENSG00000271130 | IGHV3OR16-8 |  |  | 53881 | NaN | -0.0172 | No |
| 42 | ENSG00000259490 | IGHV3OR15-7 |  |  | 54306 | NaN | -0.0162 | No |
| 43 | ENSG00000211891 | IGHE |  |  | 55654 | NaN | -0.0309 | No |
| 44 | ENSG00000211895 | IGHA1 |  |  | 55655 | NaN | -0.0227 | No |
| 45 | ENSG00000211899 | IGHM |  |  | 55656 | NaN | -0.0146 | No |
| 46 | ENSG00000211898 | IGHD |  |  | 55657 | NaN | -0.0064 | No |
| 47 | ENSG00000259303 | IGHV2OR16-5 |  |  | 55985 | NaN | -0.0038 | No |
| 48 | ENSG00000211968 | IGHV1-58 |  |  | 57278 | NaN | -0.0175 | No |
| 49 | ENSG00000211965 | IGHV3-49 |  |  | 57279 | NaN | -0.0094 | No |
| 50 | ENSG00000211966 | IGHV5-51 |  |  | 57282 | NaN | -0.0013 | No |
| 51 | ENSG00000211961 | IGHV1-45 |  |  | 57283 | NaN | 0.0069 | No |
| 52 | ENSG00000276775 | IGHV4-4 |  |  | 57880 | NaN | 0.0049 | No |
| 53 | ENSG00000281990 | IGHV1-69-2 |  |  | 58519 | NaN | 0.0023 | No |
| 54 | ENSG00000233732 | IGHV3OR16-10 |  |  | 59093 | NaN | 0.0007 | No |
Table: GSEA details
[plain text format]

  

Fig 2: IMMUNOGLOBULIN\_COMPLEX\_\_CIRCULATING(GO:0042571)
  
Blue-Pink O' Gram in the Space of the Analyzed GeneSet

  

Fig 3: IMMUNOGLOBULIN\_COMPLEX\_\_CIRCULATING(GO:0042571): Random ES distribution
  
Gene set null distribution of ES for
 **IMMUNOGLOBULIN\_COMPLEX\_\_CIRCULATING(GO:0042571)**

  
