## Supplementary material for "A single cysteine residue in vimentin regulates long non-coding RNA *XIST* to suppress epithelial-mesenchymal transition and stemness in breast cancer": IMMUNOGLOBULIN_RECEPTOR_BINDING(GO_0034987).html

Details for gene set IMMUNOGLOBULIN\_RECEPTOR\_BINDING(GO:0034987)[GSEA]


|  || Dataset | fpkm.sample |
| Phenotype | sample.cls |
| Upregulated in class | WT\_VIM |
| GeneSet | IMMUNOGLOBULIN\_RECEPTOR\_BINDING(GO:0034987) |
| Enrichment Score (ES) | -0.34028834 |
| Normalized Enrichment Score (NES) | -1.0 |
| Nominal p-value | 0.0 |
| FDR q-value | 1.0 |
| FWER p-Value | 0.141 |
Table: GSEA Results Summary

  

Fig 1: Enrichment plot: IMMUNOGLOBULIN\_RECEPTOR\_BINDING(GO:0034987)
  
Profile of the Running ES Score & Positions of GeneSet Members on the Rank Ordered List

  

| PROBE | DESCRIPTION   (from dataset) | GENE SYMBOL | GENE\_TITLE | RANK IN GENE LIST | RANK METRIC SCORE | RUNNING ES | CORE ENRICHMENT || 1 | ENSG00000128322 | IGLL1 |  |  | 4656 | Infinity | -0.0731 | Yes |
| 2 | ENSG00000132465 | JCHAIN |  |  | 20441 | 0.738 | 0.0800 | Yes |
| 3 | ENSG00000182511 | FES |  |  | 26338 | 0.260 | 0.1283 | Yes |
| 4 | ENSG00000196924 | FLNA |  |  | 27641 | 0.150 | 0.1918 | Yes |
| 5 | ENSG00000000938 | FGR |  |  | 28133 | 0.108 | 0.2448 | Yes |
| 6 | ENSG00000224650 | IGHV3-74 |  |  | 30820 | 0.000 | 0.1993 | No |
| 7 | ENSG00000211893 | IGHG2 |  |  | 32101 | 0.000 | 0.1777 | No |
| 8 | ENSG00000211892 | IGHG4 |  |  | 32604 | 0.000 | 0.1692 | No |
| 9 | ENSG00000211896 | IGHG1 |  |  | 32605 | 0.000 | 0.1692 | No |
| 10 | ENSG00000211947 | IGHV3-21 |  |  | 33401 | NaN | 0.1614 | No |
| 11 | ENSG00000211945 | IGHV1-18 |  |  | 33403 | NaN | 0.1671 | No |
| 12 | ENSG00000166527 | CLEC4D |  |  | 33979 | NaN | 0.1631 | No |
| 13 | ENSG00000211935 | IGHV1-3 |  |  | 34044 | NaN | 0.1677 | No |
| 14 | ENSG00000232216 | IGHV3-43 |  |  | 37879 | NaN | 0.1085 | No |
| 15 | ENSG00000211890 | IGHA2 |  |  | 37893 | NaN | 0.1139 | No |
| 16 | ENSG00000211950 | IGHV1-24 |  |  | 39151 | NaN | 0.0984 | No |
| 17 | ENSG00000211951 | IGHV2-26 |  |  | 40699 | NaN | 0.0779 | No |
| 18 | ENSG00000211952 | IGHV4-28 |  |  | 40700 | NaN | 0.0836 | No |
| 19 | ENSG00000211957 | IGHV3-35 |  |  | 40703 | NaN | 0.0892 | No |
| 20 | ENSG00000211958 | IGHV3-38 |  |  | 40704 | NaN | 0.0949 | No |
| 21 | ENSG00000211979 | IGHV7-81 |  |  | 41796 | NaN | 0.0822 | No |
| 22 | ENSG00000211972 | IGHV3-66 |  |  | 41800 | NaN | 0.0878 | No |
| 23 | ENSG00000231475 | IGHV4-31 |  |  | 42071 | NaN | 0.0889 | No |
| 24 | ENSG00000270472 | IGHV3OR16-9 |  |  | 42516 | NaN | 0.0871 | No |
| 25 | ENSG00000188403 | IGHV1OR15-9 |  |  | 42837 | NaN | 0.0874 | No |
| 26 | ENSG00000223648 | IGHV3-64 |  |  | 43450 | NaN | 0.0827 | No |
| 27 | ENSG00000271178 | IGHV3OR16-13 |  |  | 44717 | NaN | 0.0670 | No |
| 28 | ENSG00000259680 | AC136428.1 |  |  | 44852 | NaN | 0.0704 | No |
| 29 | ENSG00000270505 | IGHV1OR15-1 |  |  | 45965 | NaN | 0.0573 | No |
| 30 | ENSG00000281179 | AC135068.8 |  |  | 46591 | NaN | 0.0524 | No |
| 31 | ENSG00000211933 | IGHV6-1 |  |  | 47067 | NaN | 0.0501 | No |
| 32 | ENSG00000211970 | IGHV4-61 |  |  | 47532 | NaN | 0.0479 | No |
| 33 | ENSG00000211976 | IGHV3-73 |  |  | 47533 | NaN | 0.0536 | No |
| 34 | ENSG00000211974 | IGHV2-70D |  |  | 47534 | NaN | 0.0593 | No |
| 35 | ENSG00000225698 | IGHV3-72 |  |  | 48227 | NaN | 0.0533 | No |
| 36 | ENSG00000270467 | IGHV3OR16-12 |  |  | 48449 | NaN | 0.0552 | No |
| 37 | ENSG00000280411 | IGHV1-69D |  |  | 48735 | NaN | 0.0561 | No |
| 38 | ENSG00000278263 | AC135068.2 |  |  | 49083 | NaN | 0.0559 | No |
| 39 | ENSG00000254709 | IGLL5 |  |  | 49907 | NaN | 0.0477 | No |
| 40 | ENSG00000211943 | IGHV3-15 |  |  | 50481 | NaN | 0.0437 | No |
| 41 | ENSG00000211946 | IGHV3-20 |  |  | 50485 | NaN | 0.0493 | No |
| 42 | ENSG00000211944 | IGHV3-16 |  |  | 50486 | NaN | 0.0550 | No |
| 43 | ENSG00000211949 | IGHV3-23 |  |  | 50487 | NaN | 0.0607 | No |
| 44 | ENSG00000271130 | IGHV3OR16-8 |  |  | 53881 | NaN | 0.0090 | No |
| 45 | ENSG00000259490 | IGHV3OR15-7 |  |  | 54306 | NaN | 0.0075 | No |
| 46 | ENSG00000248385 | TARM1 |  |  | 55525 | NaN | -0.0074 | No |
| 47 | ENSG00000211891 | IGHE |  |  | 55654 | NaN | -0.0039 | No |
| 48 | ENSG00000211895 | IGHA1 |  |  | 55655 | NaN | 0.0018 | No |
| 49 | ENSG00000211899 | IGHM |  |  | 55656 | NaN | 0.0075 | No |
| 50 | ENSG00000211898 | IGHD |  |  | 55657 | NaN | 0.0132 | No |
| 51 | ENSG00000259303 | IGHV2OR16-5 |  |  | 55985 | NaN | 0.0133 | No |
| 52 | ENSG00000211968 | IGHV1-58 |  |  | 57278 | NaN | -0.0028 | No |
| 53 | ENSG00000211965 | IGHV3-49 |  |  | 57279 | NaN | 0.0029 | No |
| 54 | ENSG00000211966 | IGHV5-51 |  |  | 57282 | NaN | 0.0085 | No |
| 55 | ENSG00000211961 | IGHV1-45 |  |  | 57283 | NaN | 0.0142 | No |
| 56 | ENSG00000276775 | IGHV4-4 |  |  | 57880 | NaN | 0.0098 | No |
| 57 | ENSG00000281990 | IGHV1-69-2 |  |  | 58519 | NaN | 0.0047 | No |
| 58 | ENSG00000233732 | IGHV3OR16-10 |  |  | 59093 | NaN | 0.0007 | No |
Table: GSEA details
[plain text format]

  

Fig 2: IMMUNOGLOBULIN\_RECEPTOR\_BINDING(GO:0034987)
  
Blue-Pink O' Gram in the Space of the Analyzed GeneSet

  

Fig 3: IMMUNOGLOBULIN\_RECEPTOR\_BINDING(GO:0034987): Random ES distribution
  
Gene set null distribution of ES for
 **IMMUNOGLOBULIN\_RECEPTOR\_BINDING(GO:0034987)**

  
