## Supplementary material for "A single cysteine residue in vimentin regulates long non-coding RNA *XIST* to suppress epithelial-mesenchymal transition and stemness in breast cancer": POSITIVE_REGULATION_OF_PEPTIDYL_SERINE_PHOSPHORYLATION_OF_STAT_PROTEIN(GO_0033141).html

Details for gene set POSITIVE\_REGULATION\_OF\_PEPTIDYL\_SERINE\_PHOSPHORYLATION\_OF\_STAT\_PROTEIN(GO:0033141)[GSEA]


|  || Dataset | fpkm.sample |
| Phenotype | sample.cls |
| Upregulated in class | WT\_VIM |
| GeneSet | POSITIVE\_REGULATION\_OF\_PEPTIDYL\_SERINE\_PHOSPHORYLATION\_OF\_STAT\_PROTEIN(GO:0033141) |
| Enrichment Score (ES) | -0.2927491 |
| Normalized Enrichment Score (NES) | -1.0 |
| Nominal p-value | 0.0 |
| FDR q-value | 1.0 |
| FWER p-Value | 0.141 |
Table: GSEA Results Summary

  

Fig 1: Enrichment plot: POSITIVE\_REGULATION\_OF\_PEPTIDYL\_SERINE\_PHOSPHORYLATION\_OF\_STAT\_PROTEIN(GO:0033141)
  
Profile of the Running ES Score & Positions of GeneSet Members on the Rank Ordered List

  

| PROBE | DESCRIPTION   (from dataset) | GENE SYMBOL | GENE\_TITLE | RANK IN GENE LIST | RANK METRIC SCORE | RUNNING ES | CORE ENRICHMENT || 1 | ENSG00000177047 | IFNW1 |  |  | 33 | Infinity | 0.0227 | Yes |
| 2 | ENSG00000147885 | IFNA16 |  |  | 116 | Infinity | 0.0446 | Yes |
| 3 | ENSG00000137080 | IFNA21 |  |  | 1139 | Infinity | 0.0505 | Yes |
| 4 | ENSG00000168546 | GFRA2 |  |  | 2645 | Infinity | 0.0483 | Yes |
| 5 | ENSG00000147873 | IFNA5 |  |  | 4715 | Infinity | 0.0366 | Yes |
| 6 | ENSG00000228083 | IFNA14 |  |  | 5042 | Infinity | 0.0543 | Yes |
| 7 | ENSG00000184995 | IFNE |  |  | 5384 | Infinity | 0.0718 | Yes |
| 8 | ENSG00000128342 | LIF |  |  | 26937 | 0.208 | 0.1912 | Yes |
| 9 | ENSG00000165731 | RET |  |  | 28880 | 0.042 | 0.2558 | Yes |
| 10 | ENSG00000171855 | IFNB1 |  |  | 32947 | 0.000 | 0.1870 | No |
| 11 | ENSG00000120235 | IFNA6 |  |  | 39469 | NaN | 0.0999 | No |
| 12 | ENSG00000234829 | IFNA17 |  |  | 41226 | NaN | 0.0935 | No |
| 13 | ENSG00000147896 | IFNK |  |  | 43124 | NaN | 0.0846 | No |
| 14 | ENSG00000236637 | IFNA4 |  |  | 47027 | NaN | 0.0419 | No |
| 15 | ENSG00000233816 | IFNA13 |  |  | 49000 | NaN | 0.0318 | No |
| 16 | ENSG00000188379 | IFNA2 |  |  | 49060 | NaN | 0.0541 | No |
| 17 | ENSG00000120242 | IFNA8 |  |  | 52035 | NaN | 0.0270 | No |
| 18 | ENSG00000214042 | IFNA7 |  |  | 54878 | NaN | 0.0022 | No |
| 19 | ENSG00000186803 | IFNA10 |  |  | 55594 | NaN | 0.0134 | No |
| 20 | ENSG00000111537 | IFNG |  |  | 56672 | NaN | 0.0184 | No |
| 21 | ENSG00000197919 | IFNA1 |  |  | 58784 | NaN | 0.0059 | No |
Table: GSEA details
[plain text format]

  

Fig 2: POSITIVE\_REGULATION\_OF\_PEPTIDYL\_SERINE\_PHOSPHORYLATION\_OF\_STAT\_PROTEIN(GO:0033141)
  
Blue-Pink O' Gram in the Space of the Analyzed GeneSet

  

Fig 3: POSITIVE\_REGULATION\_OF\_PEPTIDYL\_SERINE\_PHOSPHORYLATION\_OF\_STAT\_PROTEIN(GO:0033141): Random ES distribution
  
Gene set null distribution of ES for
 **POSITIVE\_REGULATION\_OF\_PEPTIDYL\_SERINE\_PHOSPHORYLATION\_OF\_STAT\_PROTEIN(GO:0033141)**

  
