## Supplementary material for "A single cysteine residue in vimentin regulates long non-coding RNA *XIST* to suppress epithelial-mesenchymal transition and stemness in breast cancer": TYPE_I_INTERFERON_RECEPTOR_BINDING(GO_0005132).html

Details for gene set TYPE\_I\_INTERFERON\_RECEPTOR\_BINDING(GO:0005132)[GSEA]


|  || Dataset | fpkm.sample |
| Phenotype | sample.cls |
| Upregulated in class | WT\_VIM |
| GeneSet | TYPE\_I\_INTERFERON\_RECEPTOR\_BINDING(GO:0005132) |
| Enrichment Score (ES) | -0.29250112 |
| Normalized Enrichment Score (NES) | -0.7035457 |
| Nominal p-value | 0.43384615 |
| FDR q-value | 1.0 |
| FWER p-Value | 0.325 |
Table: GSEA Results Summary

  

Fig 1: Enrichment plot: TYPE\_I\_INTERFERON\_RECEPTOR\_BINDING(GO:0005132)
  
Profile of the Running ES Score & Positions of GeneSet Members on the Rank Ordered List

  

| PROBE | DESCRIPTION   (from dataset) | GENE SYMBOL | GENE\_TITLE | RANK IN GENE LIST | RANK METRIC SCORE | RUNNING ES | CORE ENRICHMENT || 1 | ENSG00000177047 | IFNW1 |  |  | 33 | Infinity | 0.0619 | Yes |
| 2 | ENSG00000147885 | IFNA16 |  |  | 116 | Infinity | 0.1231 | Yes |
| 3 | ENSG00000137080 | IFNA21 |  |  | 1139 | Infinity | 0.1683 | Yes |
| 4 | ENSG00000147873 | IFNA5 |  |  | 4715 | Infinity | 0.1703 | Yes |
| 5 | ENSG00000228083 | IFNA14 |  |  | 5042 | Infinity | 0.2273 | Yes |
| 6 | ENSG00000184995 | IFNE |  |  | 5384 | Infinity | 0.2840 | Yes |
| 7 | ENSG00000171855 | IFNB1 |  |  | 32947 | 0.000 | -0.1822 | No |
| 8 | ENSG00000120235 | IFNA6 |  |  | 39469 | NaN | -0.2300 | No |
| 9 | ENSG00000234829 | IFNA17 |  |  | 41226 | NaN | -0.1972 | No |
| 10 | ENSG00000147896 | IFNK |  |  | 43124 | NaN | -0.1668 | No |
| 11 | ENSG00000236637 | IFNA4 |  |  | 47027 | NaN | -0.1703 | No |
| 12 | ENSG00000233816 | IFNA13 |  |  | 49000 | NaN | -0.1412 | No |
| 13 | ENSG00000188379 | IFNA2 |  |  | 49060 | NaN | -0.0796 | No |
| 14 | ENSG00000120242 | IFNA8 |  |  | 52035 | NaN | -0.0675 | No |
| 15 | ENSG00000214042 | IFNA7 |  |  | 54878 | NaN | -0.0530 | No |
| 16 | ENSG00000186803 | IFNA10 |  |  | 55594 | NaN | -0.0026 | No |
| 17 | ENSG00000197919 | IFNA1 |  |  | 58784 | NaN | 0.0059 | No |
Table: GSEA details
[plain text format]

  

Fig 2: TYPE\_I\_INTERFERON\_RECEPTOR\_BINDING(GO:0005132)
  
Blue-Pink O' Gram in the Space of the Analyzed GeneSet

  

Fig 3: TYPE\_I\_INTERFERON\_RECEPTOR\_BINDING(GO:0005132): Random ES distribution
  
Gene set null distribution of ES for
 **TYPE\_I\_INTERFERON\_RECEPTOR\_BINDING(GO:0005132)**

  
