## Supplementary material for "A single cysteine residue in vimentin regulates long non-coding RNA *XIST* to suppress epithelial-mesenchymal transition and stemness in breast cancer": supplementary material C328S_Usman et al.docx

**Running Title: A cysteine residue in vimentin dictates the cancer fate**

**Keywords: Intermediate filaments, cancer stem cells, breast cancer signature, MCF-7 cells, Metastasis**

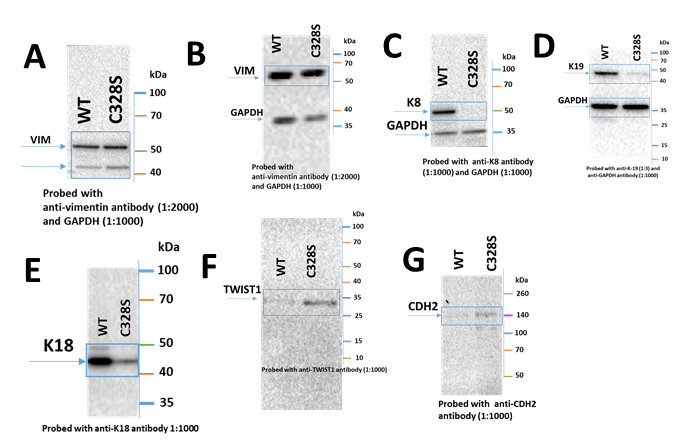

**Figure S1**: (**A**) (**B**) VIM, (**C**) K8, (**D**) K19, (**E**) K18, (**F**) TWIST1, (**G**) CDH2/N-Cadherin expression in MCF-7 expressing WT-VIM and C328S-VIM by western blotting. Ten µg protein was loaded for all samples. GAPDH was used as the loading control. Blue rectangles show the area cropped to regroup bands.

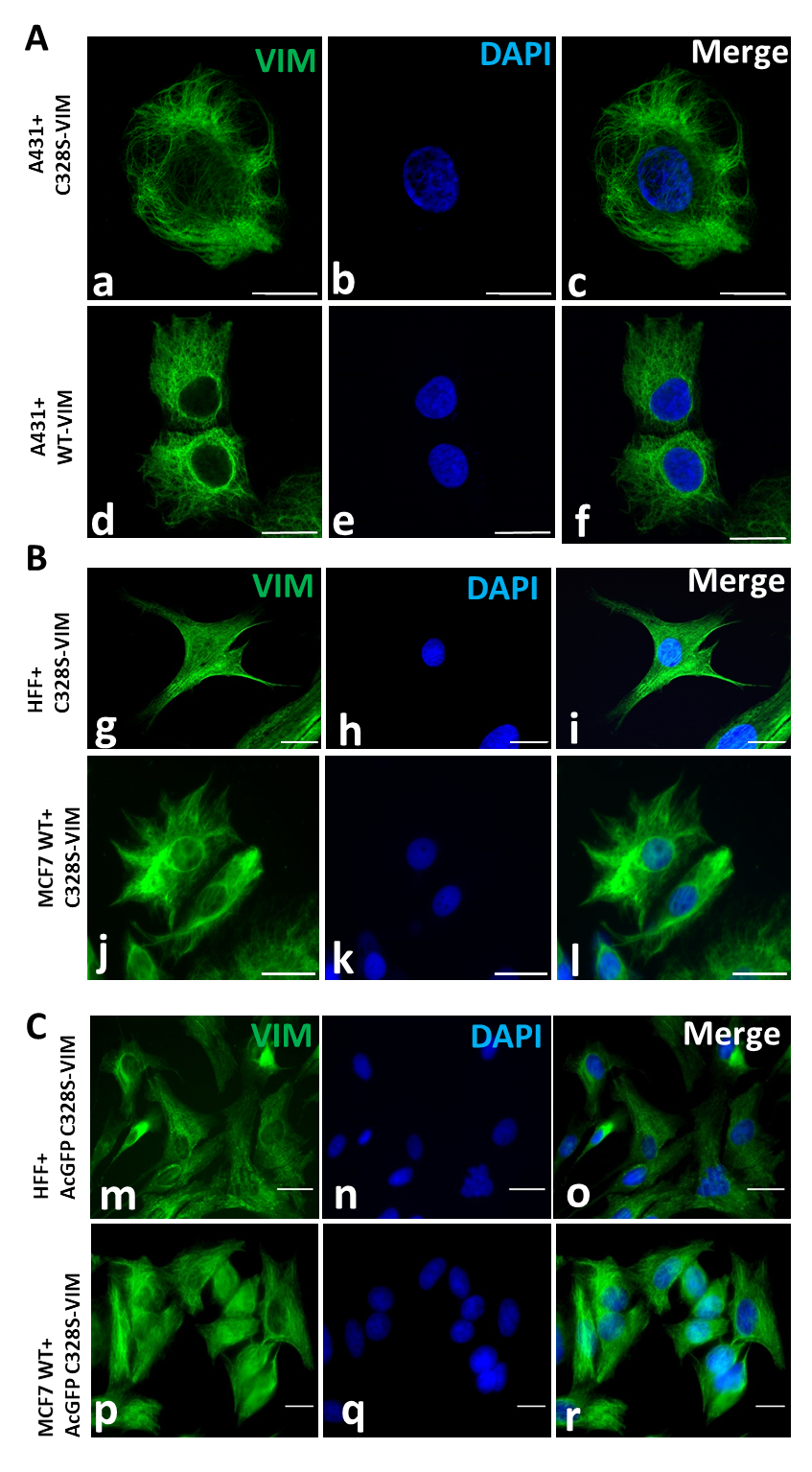

Figure S2: A: Filament formation by C328S-VIM (a-c) and WT-VIM (d-f) in A431. Cells were immunostained with V9 followed by AF-488 labelled anti-mouse secondary antibody. Note that vimentin filament distribution is different from MCF-7C328S-VIM as the filaments are fully extended in the area between nucleus and peripheral cell boundaries. B: Integration of C328S-VIM into the endogenous vimentin network of HFF-1 (g-i) and MCF-7 WT (j-l). Cells were immunostained with V9 followed by AF-488 labelled anti-mouse secondary antibody. C: Integration of AcGFP-C328S-VIM mutant in HFF-1 (m-o) and MCF-7WT cells (p-r). AcGFP-C328S-VIM constructs integrated into pre-existing filament network.

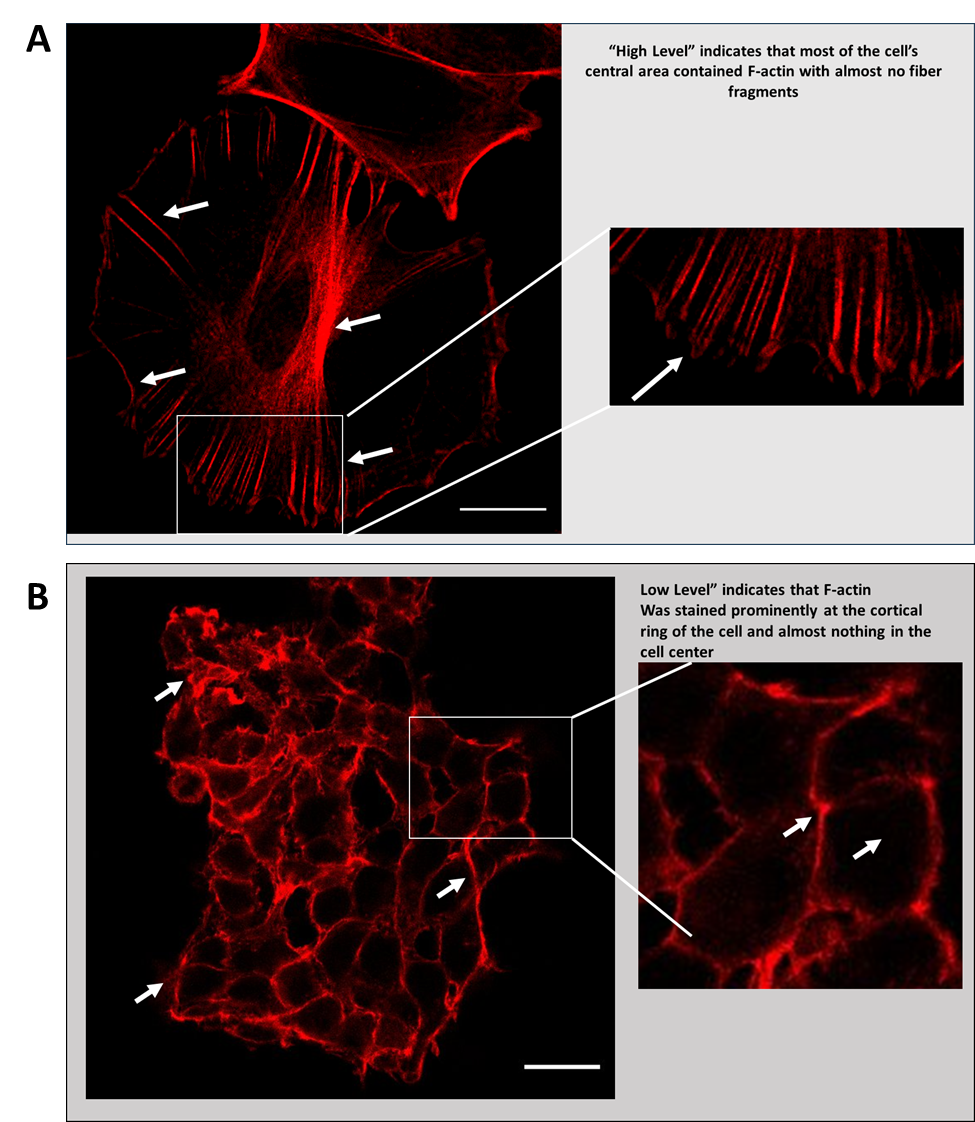

**Figure S3:** **A**: Representative high resolution image showing high level actin stress fibres in the WT-VIM cells. **B**: Representative image showing low level actin staining with aggregates/fragments confined to cortical margins and no staining in cell centres (scale bar = 20 µm).

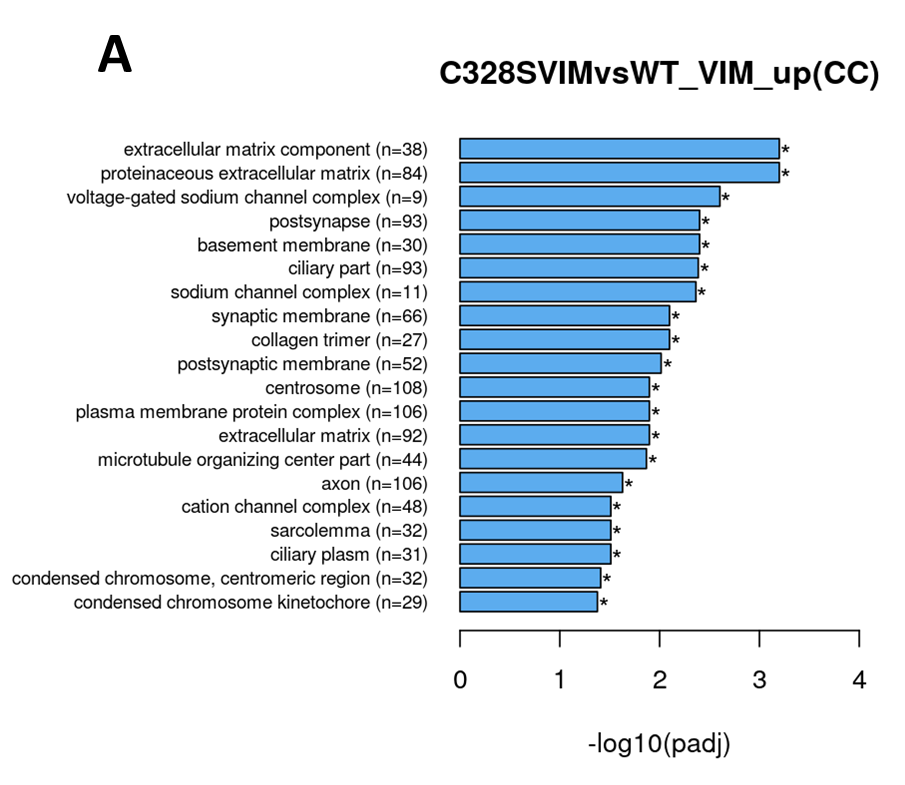

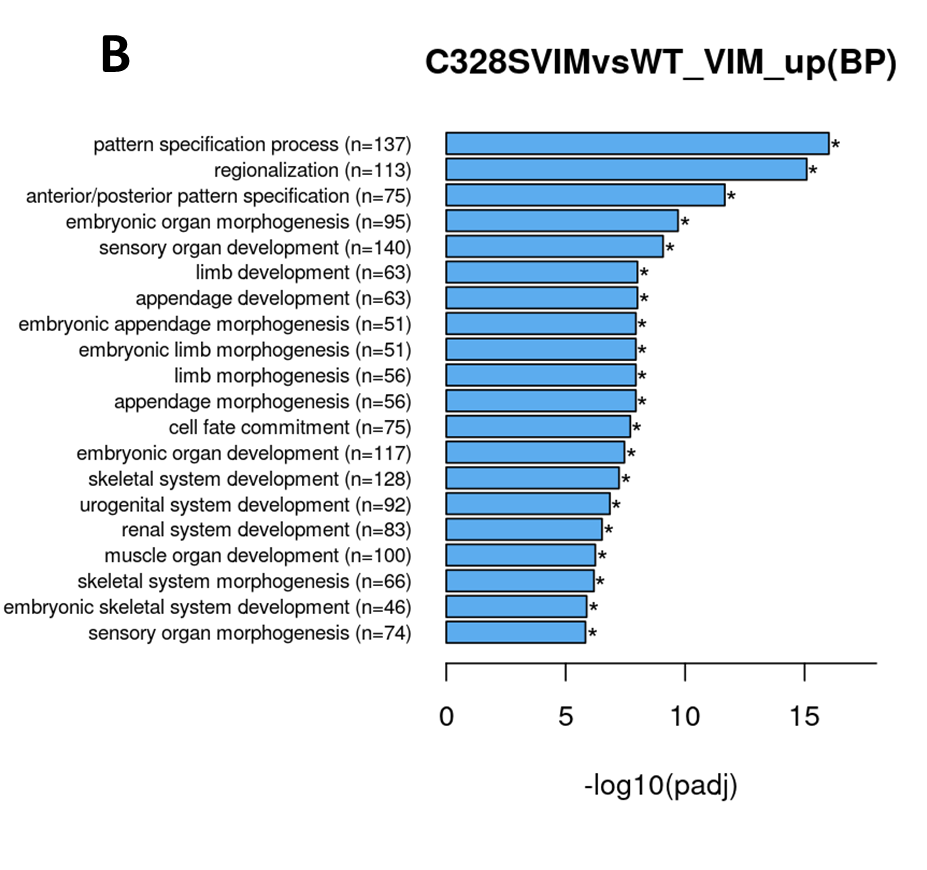

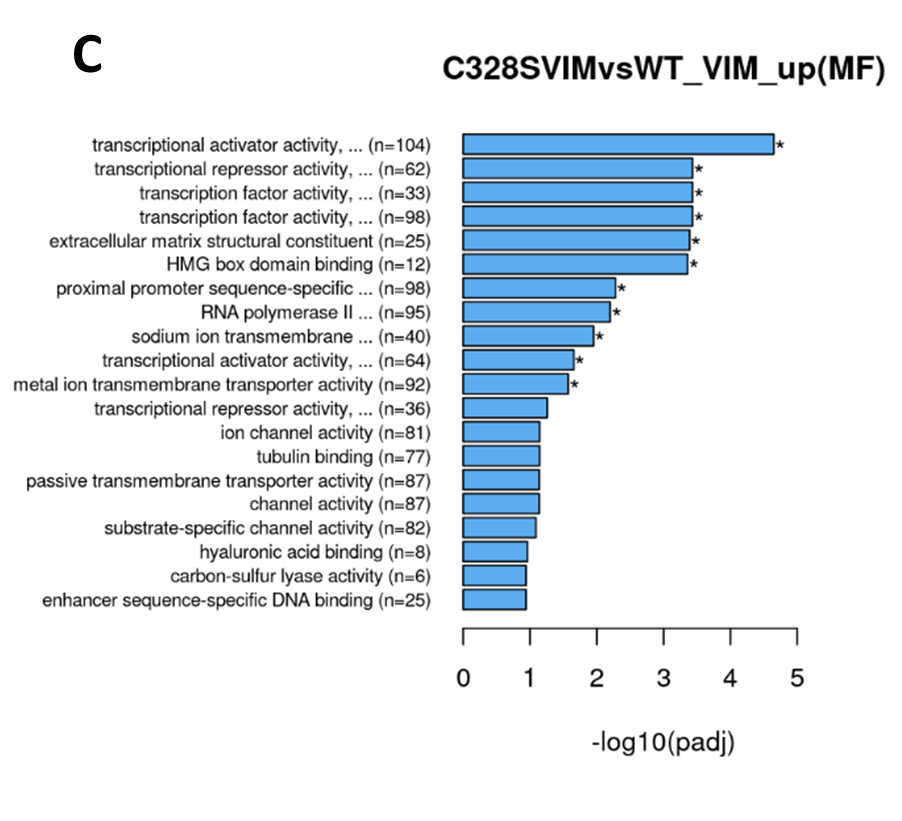

**Figure S4**: **GO pathway enrichment analyses of upregulated DEGs** in WT vs C328S cells by RNA-seq. DEGs were grouped into three major functional types: Molecular Function (MF), Biological Process (BP) and Cellular Component (CC). **A:** GO analysis of CC category showing functions that were upregulated by DEGs in WT vs C328S cells, these include extracellular matrix component, proteinaceous extracellular component, voltage-gated sodium channel complex, post synapse, basement membrane etc. **B**: GO enrichment analysis of BP category showing biological processes that were upregulated by DEGs, these include pattern specification process, regionalisation, ant/post pattern specification etc. **C**: GO analysis of MF category showing functions that were upregulated by DEGs, these include transcriptional activator activity, RNA polymerase II transcription regulatory region sequence-specific DNA binding, transcriptional repressor activity, RNA polymerase II transcription regulatory region sequence-specific DNA binding, extracellular matrix structural constituent, HMG box domain binding, proximal promoter sequence-specific DNA binding etc.

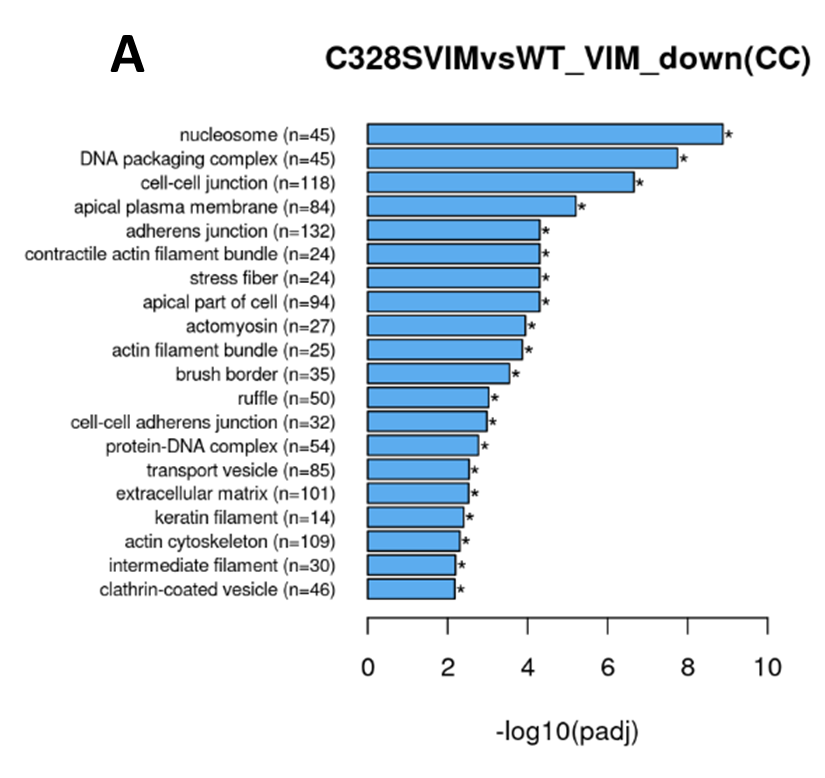

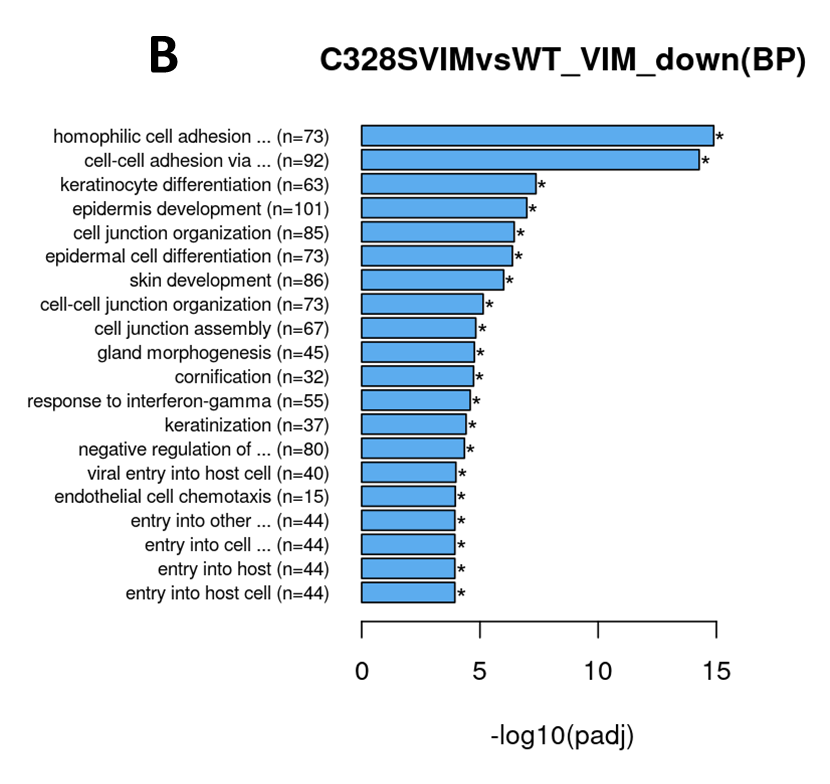

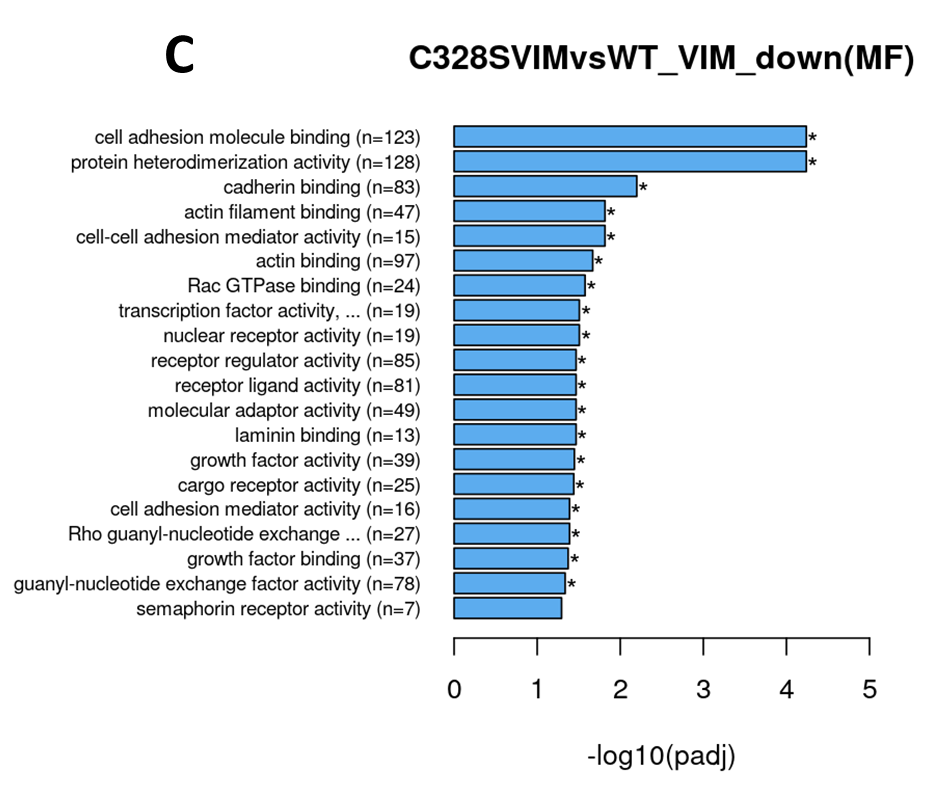

**Figure S5**: **GO pathway enrichment analyses of downregulated DEGs** in WT vs C328S cells by RNA-Seq. DEGs were grouped into three major functional types: Molecular Function (MF), Biological Process (BP) and Cellular Component (CC). **A:** GO analysis of CC category showing functions that were downregulated by DEGs in WT vs C328S cells, these include nucleosome, DNA packaging complex, cell-cell junction, apical plasma membrane, adherens junction, contractile actin filament bundle, stress fibre etc. **B**: GO enrichment analysis of BP category showing biological processes that were downregulated by DEGs, these include homophilic cell adhesion via plasma membrane adhesion molecules, cell-cell adhesion via plasma-membrane adhesion molecules, keratinocyte differentiation, cell junction organization, epidermal cell differentiation, cell junction assembly etc. **C**: GO analysis of MF category showing functions that were downregulated by DEGs, these include cell adhesion molecule binding, protein heterodimerization activity, cadherin binding, cell-cell adhesion mediator activity, actin filament binding, Rac GTPase binding, laminin binding etc.

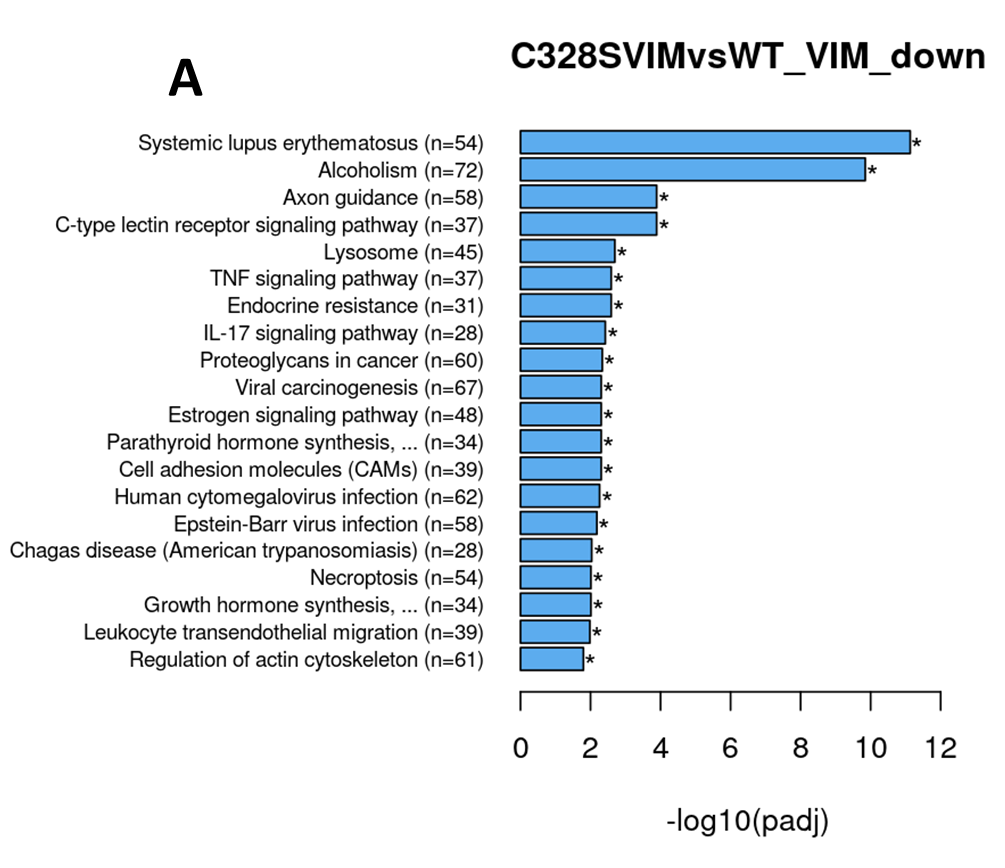

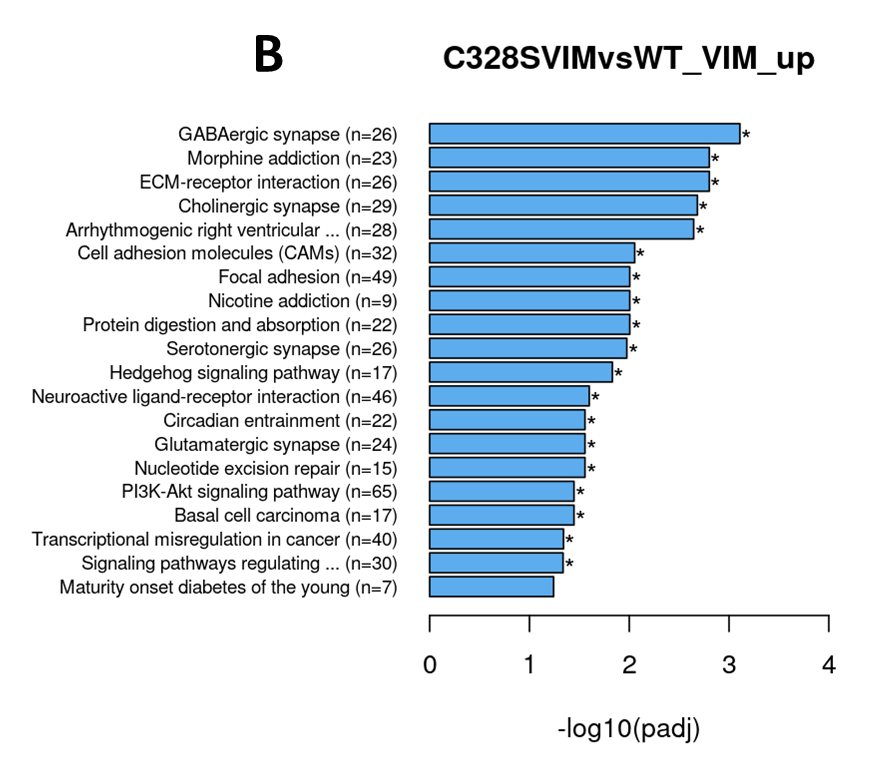

**Figure S6**: **KEGG pathway analysis**. Bar graph **A.** showing downregulated cellular functions by KEGG analysis in WT vs C328S involved in cell adhesion molecules CAMs, regulation of actin skeleton, SLE, alcoholism, axon guidance, TNF signalling, IL17 signalling etc. Bar graph **B** showing upregulated cellular functions by KEGG analysis in WT vs C328S involved ECM receptor interaction, focal adhesion, Pl3K-Akt signalling pathway etc.

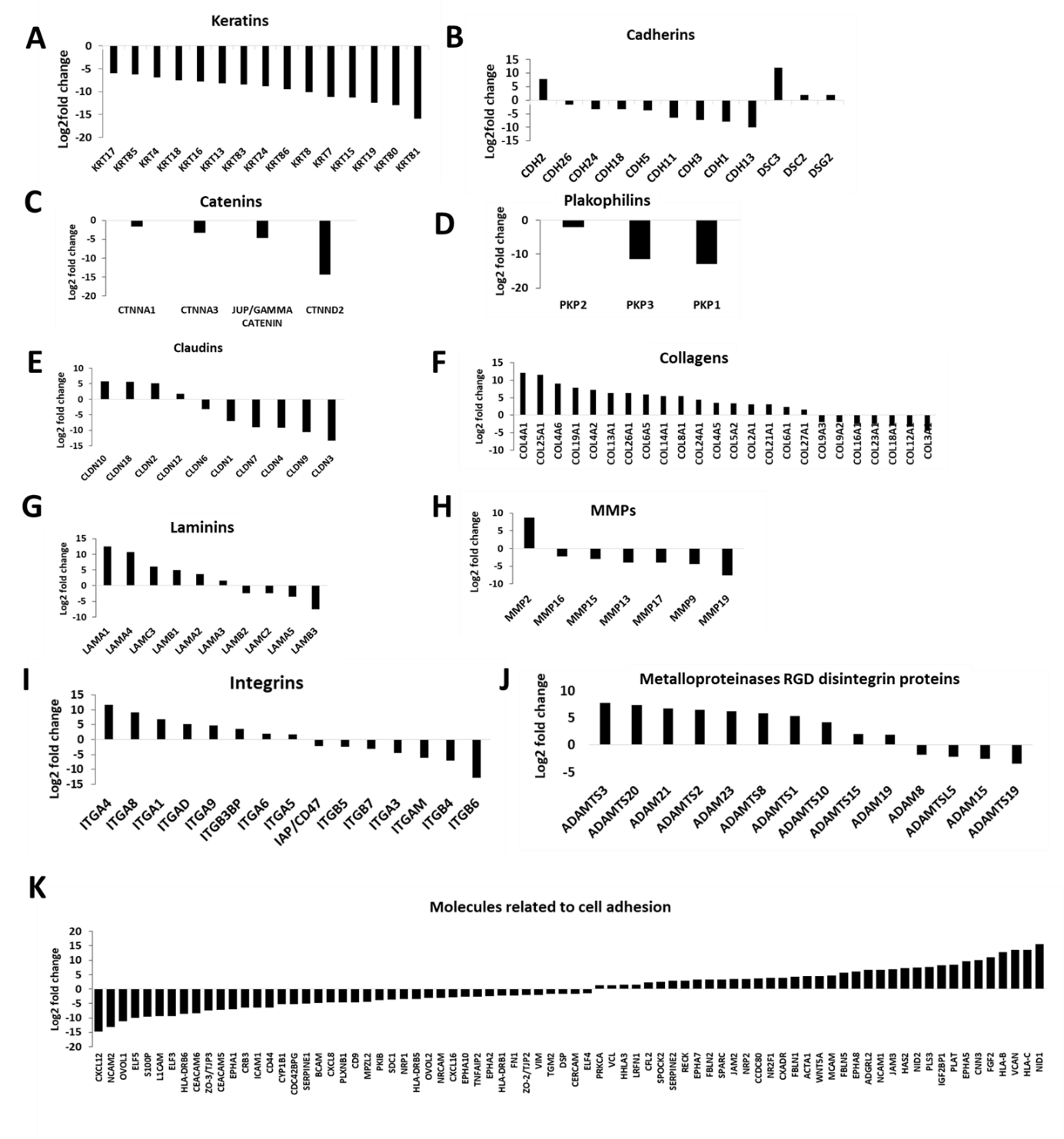

**Figure S7: DEGs deduced from the RNA-Seq data.** Relative log2-fold changes of DEGs in WT vs C328S cells related to cell junctions, cell-ECM junctions including cytokeratins (**A**), cadherins (**B**), catenins (**C**), plakophilins (**D**), claudins (**E**), collagens (**F**), laminins (**G**), MMPs (**H**), integrins (**I**), metalloproteinases RGD disintegrin proteins (**J**) and several genes related to cell adhesion (**K**) by RNA-Seq analysis.

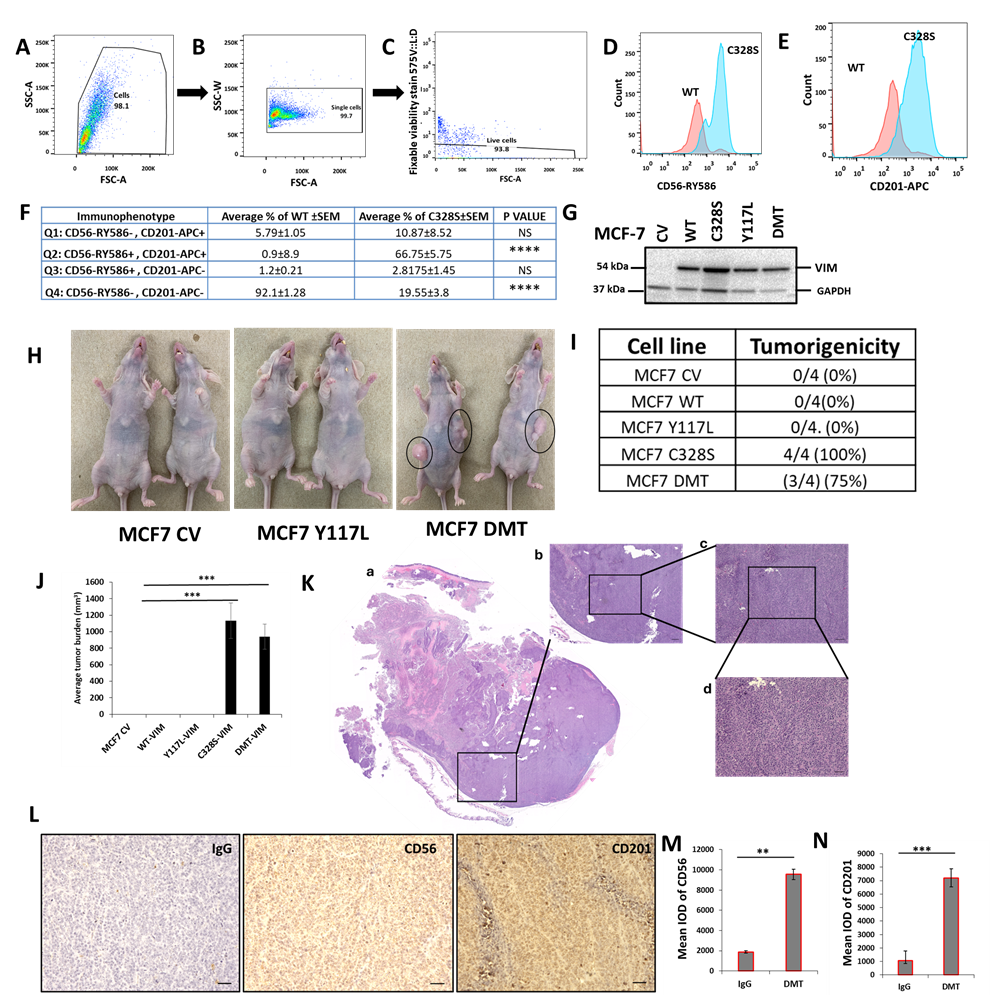

**Figure S8:** Representative gating strategies for flow cytometry analysis of breast cancer stem cell markers CD56/NCAM1 and CD201/PROCR in WT and C328S cells. Cells were stained for surface antigens with RY586–conjugated antibody specific for CD56/NCAM1, APC-conjugated antibody specific for CD201/ PROCR and Fixable Viability Stain FVS575V for live/dead cells. Gates were applied for cells (SSC-A: FSC-A) (**A**), single cells (SSC-W: FSC-A) (**B**), live cells (FVS575V ::L/d:FSC-A) (**C**) as shown. (**D**): Representative histogram showing the MFI of CD56-RY586 in WT and C328S cells by flow cytometry. WT cells are represented by red colour and C328S by blue. (**E**): Representative histogram showing the MFI of CD201-APC in WT and C328S cells by flow cytometry. WT cells are represented by red colour and C328S by blue. (**F**): Immunophenosubtypes in WT and C328S cells in tabulated form. Note that 92.1% WT cells are double negative CD56-RY586-, CD201-APC- (Q4) (p<0.0001) as compared to C328S (19.6%). Similarly 67% of C328S cells are double positive (Q2) CD56-RY586+, CD201-APC+ (p<0.0001) as compared to WT (0.9%). Statistical analyses: n = 3, Error bars= ± SEM, ****=p<0.0001. (**G**): MCF-7 cells were transduced with control vector (CV), wildtype vimentin (WT-VIM), mutant vimentin C328S-VIM, Y117L-VIM and double mutant DMT (C328S-VIM+Y117L-VIM) constructs and the stable expression was confirmed by western blot in these cell lines. (**H**): Transplantation of CV, Y117L-VIM and DMT-VIM expressing MCF-7 cells in nude mice without estrogen supplementation (**I**): Table showing tumorigenicity potential of CV, WT-VIM, C328-VIM, Y117L-VIM and DMT-VIM expressing MCF-7 cells without estrogen supplementation (**J**): Average tumour burden after two weeks in nude mice injected with CV, WT-VIM, C328-VIM, Y117L-VIM and DMT-VIM expressing MCF-7 cells. (**K**): H & E stained tumour section from mouse injected with DMT expressing MCF-7 cells, scale bar= 50 µm. (**L**): Representative image from immunohistochemical staining of CD56 and CD201 in tumour sections from mice injected with DMT expressing MCF-7 cells as compared to IgG control, scale bar= 50 µm. Quantification of CD56 (**M**) and CD201 (**N**) IHC staining in tumour sections from mice injected with DMT expressing MCF-7 cells as compared to IgG control using *ImageJ*. The IOD=optical intensity of positive cells × area of positive cells.

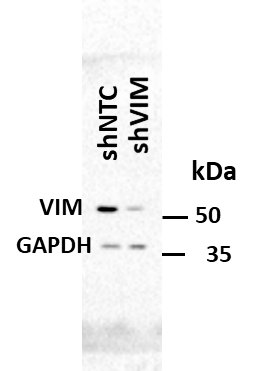

**Figure S9: Vimentin expression in MCF-7C328S-shVIM and shNTC cell lines.** Cell lines were transduced with VIM sh-RNA and NTC retrovirus. Five µg of protein from each transduced cell line was loaded to confirm the transduction efficiency, V9 antibody was used for probing in 1:2000 dilution. GAPDH was used as the loading control.

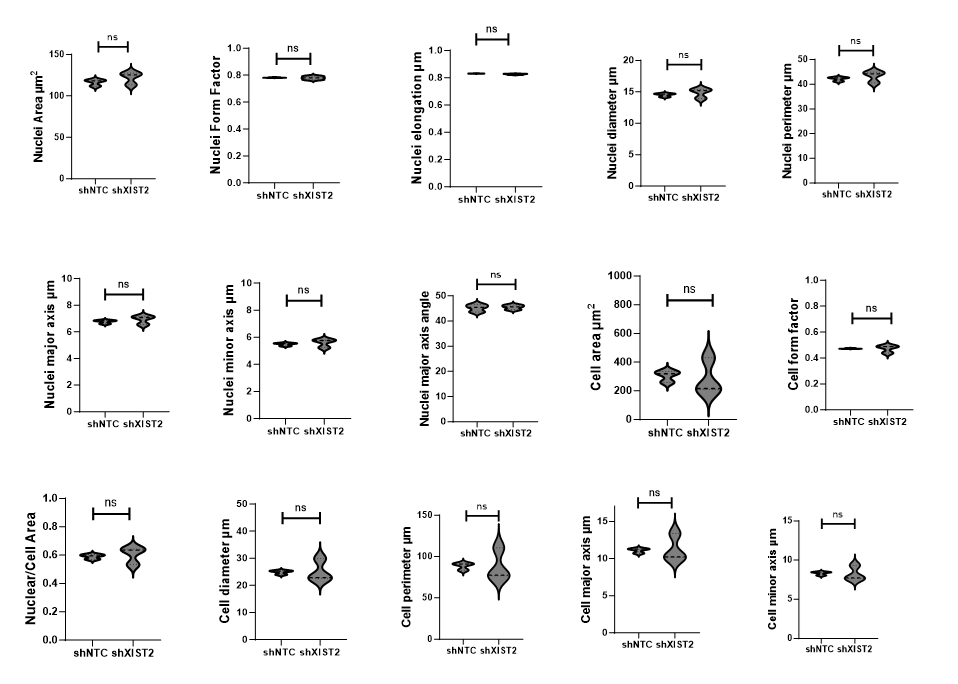

Figure S10: Morphology of C328SVIM_sh*XIST*2 and C328SVIM_shNTC cells. Morphological analysis carried out by using INCarta USA 2200 software showing insignificant differences in nuclei area, nuclei form factor, nuclei elongation, nuclei diameter, nuclei perimeter, nuclei major axis, nuclei minor axis, nuclei major axis angle degree, cell area, cell form factor, nuclei/cell area, cell diameter, cell perimeter, cell major axis and cell minor axis between the two cell lines. Statistical analyses: n = 3, Error bars= ± SEM, Student’s t-test was used to calculate p values using Microsoft Excel (ns = not significant).

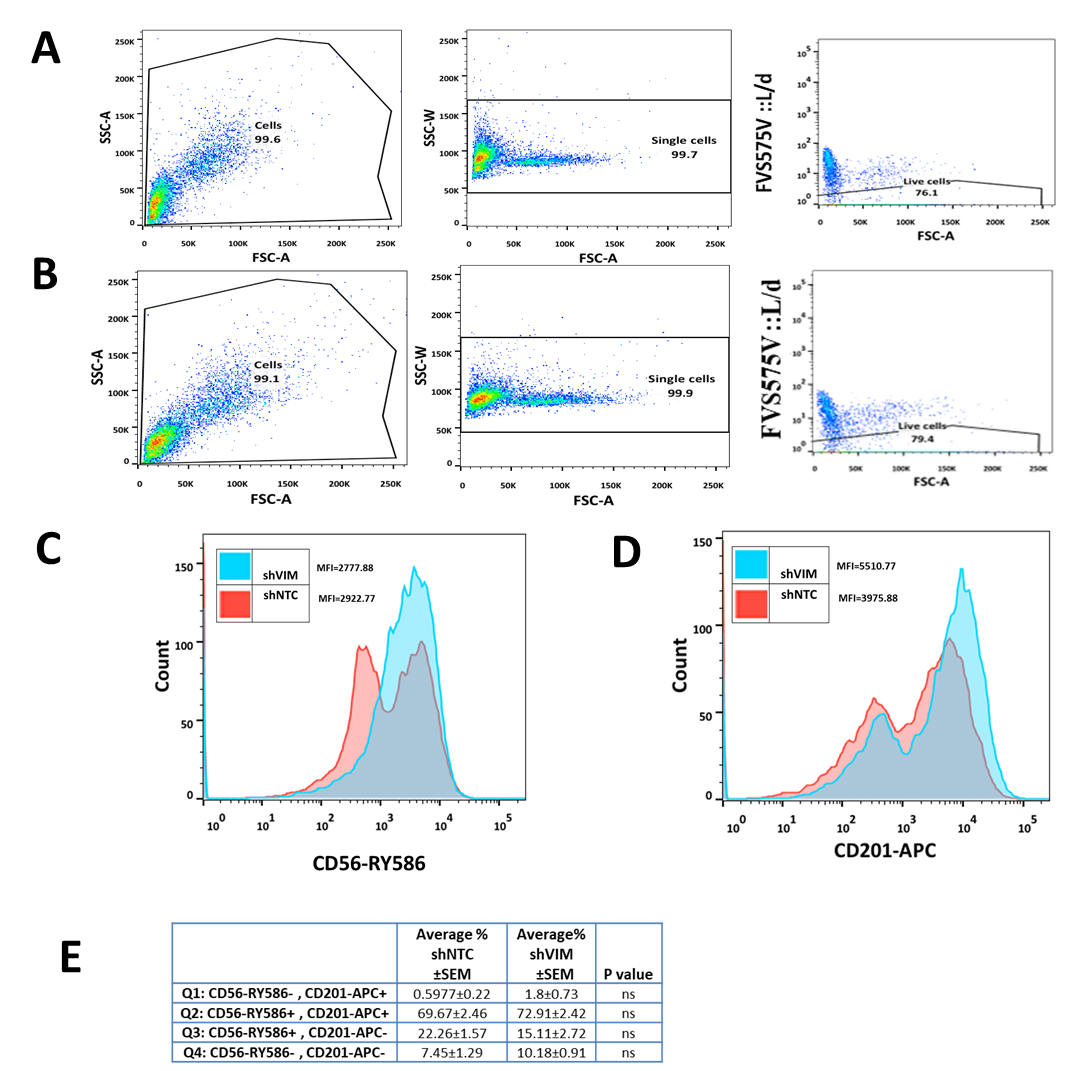

**Figure S11: Flow cytometry analyses of C328S_shVIM and shNTC cells**. (**A**): Representative gating strategies for flow cytometry analysis of breast cancer stem cell markers CD56/NCAM1 and CD201/PROCR in C328SVIM_shNTC. Cells were stained for surface antigens with RY586–conjugated antibody specific for CD56/NCAM1, APC-conjugated antibody specific for CD201/PROCR and Fixable Viability Stain FVS575V for alive/dead cells. Gates were applied for cells (SSC-A:FSC-A), single cells (SSC-W: FSC-A), live cells (FVS575V ::L/d:FSC-A) as shown. (**B**): Representative gating strategies for flow cytometry analysis of breast cancer stem cell markers CD56/NCAM1 and CD201/PROCR in C328SVIM_shVIM. Cells were stained for surface antigens with RY586–conjugated antibody specific for CD56/NCAM1, APC-conjugated antibody specific for CD201/PROCR and viability Stain FVS575V for live/dead cells. Gates were applied for cells (SSC-A:FSC-A), single cells (SSC-W: FSC-A) and alive cells (FVS575V ::L/d:FSC-A) as shown. (**C**): Representative histogram showing the MFI of CD56-RY586 in C328S_shVIM and shNTC cells by flow cytometry. shNTC cells are represented by red colour and shVIM by blue. (**D**): Representative histogram showing the MFI of CD201-APC in C328S_shVIM and shNTC cells by flow cytometry. shNTC cells are represented by red colour and shVIM by blue. (**E**): Flow cytometry overlay dot plot of CD56-RY586 versus CD201-APC (presented in **Figure 4**) after gating on single and live cells for immunophenotype in tabulated form. Statistical analyses: n = 3, Error bars= ± SEM, ns=not significant.

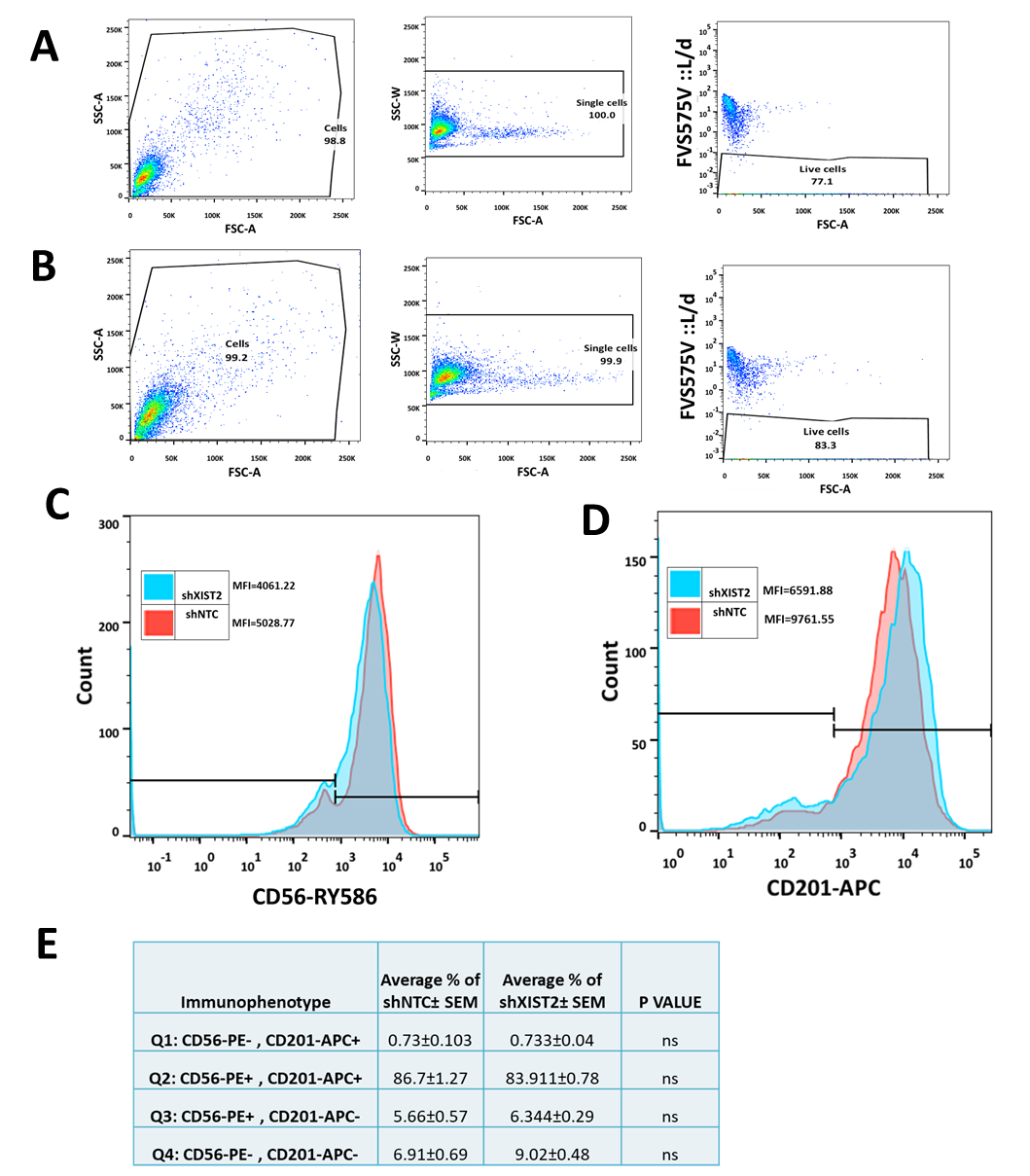

**Figure S12: Flow cytometry analyses of C328S_sh*XIST*2 and shNTC cells**. (**A**): Representative gating strategies for flow cytometry analysis of breast cancer stem cell markers CD56/NCAM1 and CD201/PROCR in C328SVIM_shNTC. Cells were stained for surface antigens with RY586–conjugated antibody specific for CD56/NCAM1, APC-conjugated antibody specific for CD201/PROCR and FVS FVS575V for live/dead cells. Gates were applied for cells (SSC-A:FSC-A), single cells (SSC-W: FSC-A), live cells (FVS575V ::L/d:FSC-A) as shown. (**B**): Representative gating strategies for flow cytometry analysis of breast cancer stem cell markers CD56/NCAM1 and CD201/PROCR in C328SVIM_sh*XIST*2. Cells were stained for surface antigens with RY586–conjugated antibody specific for CD56/NCAM1, APC-conjugated antibody specific for CD201/PROCR and FVS FVS575V for live/dead cells. Gates were applied for cells (SSC-A:FSC-A), single cells (SSC-W: FSC-A) and live cells (FVS575V ::L/d:FSC-A) as shown. (**C**): Representative histogram showing the MFI of CD56-RY586 in C328S_sh*XIST*2 and shNTC cells by flow cytometry. shNTC cells are represented by red colour and sh*XIST*2 by blue. (**D**): Representative histogram showing the MFI of CD201-APC in C328S_sh*XIST*2 and shNTC cells by flow cytometry. shNTC cells are represented by red colour and sh*XIST*2 by blue. (**E**): Flow cytometry overlay dot plot of CD56-RY586 versus CD201-APC (presented in **Figure 4**) after gating on single and live cells for immunophenotype in tabulated form. Statistical analyses: n = 3, Error bars= ± SEM, ns = not significant.

**Table S1: List of primers used for qPCR**

| Gene | Product Size (bp) | Primer Sequence  5`-3` | NM # |
| --- | --- | --- | --- |
| *VIM* | 123 | F>AGGTGGACCAGCTAACCAAC | NM_003380 |
|  |  | R>TTTCGGCTTCCTCTCTCTGA |  |
| *YAP1* | 83 | F>CCCAGATGAACGTCACAGC | NM_006106 |
|  |  | R>GATTCTCTGGTTCATGGCTGA |  |
| *POLR2A* | 73 | F>GCAAATTCACCAAGAGAGAC | NM_000937 |
|  |  | R>CACGTCGACAGGAACATCAG |  |
| *KRT18* | 117 | F>TGATGACACCAATATCACACGA | NM_000224 |
|  |  | R>ATCTGGGCTTGTAGGCCTTT |  |
| *KRT19* | 126 | F>GCCACTACTACACGACCATCC | NM_002276 |
|  |  | R>CAAACTTGGTTCGGAAGTCAT |  |
| *KRT80* | 257 | F> AGGATGCCAAGACCAAGCTG | NM_001081492 |
|  |  | R>CAGCTACAGGGAACATAAGGGG |  |
| *SNAI*1 | 92 | F>TACAGCGAGCTGCAGGACT | NM_005985 |
|  |  | R>ATCTCCGGAGGTGGGATG |  |
| *SNAI2* | 984 | F>TGGTTGCTTCAAGGACACAT | NM_003068.4 |
|  |  | R>GCAAATGCTCTGTTGCAGTG |  |
| *TWIST1* | 124 | F>AGCTACGCCTTCTCGGTCT | NM_000474 |
|  |  | R>CCTTCTCTGGAAACAATGACATC |  |
| *ZEB1* | 74 | F>TGCAGTTTTCAAAGTTAGGAACAA | NM_001128128 |
|  |  | R>TGTTGCTCTCTGAGTCATTAAGGT |  |
| *ZEB2* | 86 | F>TTGCTCCAAGATGTGTGAGG | NM_001171653 |
|  |  | R>TGTGGGGCTCCAGATATACAC |  |
| *KRT8* | 282 | F>AGCTTCTCCGCTCCTTCTAGG | NM_002273 |
|  |  | R>CAGGCTCTGGTTGACCGTAA |  |
| *XIST* | 183 | F>AGATCTTCCTCAGAAGAATAGG | NR_001564.2 |
|  |  | R>TTTATCTTCCTATCTGGGACC |  |

**Table-S2: List of primary and secondary antibodies used in this research work**

| Antibody | Dilution | Host | Catalogue # | Supplier |
| --- | --- | --- | --- | --- |
| Anti-vimentin V9 | IF=1:700 | Mouse | ab8069 | Abcam, UK |
|  | WB=1:2000 |  |  |  |
| Anti-cytokeratin K8 | IF=1:100 | Rabbit | ab53280 | Abcam, UK |
|  | WB=1:500 |  |  |  |
| Anti-cytokeratin K18 | IF=1:500 | Rabbit | ab24561 | Abcam, UK |
|  | WB=1:1000 |  |  |  |
| Anti-CDH2 | WB= 1:500 | Mouse | CA1029 | Merck Millipore, UK |
| Anti-Twist1 | WB= 1:500 | Rabbit | PA5-86070 | Thermo Fisher Scientific, UK |
| Anti-GAPDH | WB= 1:2000 | Rabbit | Ab9485 | Abcam, UK |
| Phalloidin, Alexa Flour568 conjugated | IF= 1:400 | N/A | A12380 | Life Technologies, UK |
| Anti-mouse Alexa Fluor® 488 IgG H+L | IF=1:1000 | Goat | A-11001 | Life Technologies, UK |
| Anti-mouse Alexa Fluor® 594 IgG H+L | IF=1:1000 | Goat | A-11005 | Molecular Probes, UK |
| Anti-rabbit Alexa Fluor®488 IgG H+L | IF=1:1000 | Goat | A-11008 | Life Technologies, UK |
| Anti-rabbit Alexa Fluor®594 IgG H+L | IF=1:1000 | Goat | A-11012 | Life Technologies, UK |
| Mouse IgG peroxidase conjugated | WB= 1:1000 | Goat | NA931V | GE Healthcare, UK |
| Rabbit IgG peroxidase conjugated | WB=1:1000 | Donkey | NA934V | GE Healthcare, UK |
| BD Pharmingen™ APC Rat Anti-Human CD201 | 1:20 | Rat | 563622 | BD Pharmingen™ |
| BD Horizon™ RY586 Mouse Anti-Human CD56 | 1:20 | Mouse | 568150 | BD Pharmingen™ |
| BD Horizon™ Fixable Viability Stain 575V | 1:1000 |  | 565694 | BD Pharmingen™ |
| Rabbit anti-NCAM-1/CD56 antibody (clone E7X9M) | 1:400 | Rabbit | 99746 | Cell Signaling Technology, Danvers, MA |
| Goat anti-CD201 | 10μg/ml | Goat | AF2245 | R&D Systems, Minneapolis, MN |

**Table S3: List of primers used for making *XIST* shRNA constructs and site directed mutagenesis at C328 and Y117.**

| *XIST*_shRNA1 | F:5'GATCTGGAATATTTGCAATTATAAtacctgacccataTTATAATTGCAAATATTCCTTTTTC3'  R:5'TCGAGAAAAAGGAATATTTGCAATTATAAtatgggtcaggtaTTATAATTGCAAATATTCCA3' |
| --- | --- |
| *XIST*_shRNA2 | F:5'GATCTGGATATATTGCTTAATTTAtacctgacccataTAAATTAAGCAATATATCCTTTTTC3'  R:5'TCGAGAAAAAGGATATATTGCTTAATTTAtatgggtcaggtaTAAATTAAGCAATATATCCA3' |
| *XIST*_shRNA3 | F:5'GATCTGAATATTTGCAATTATATAtacctgacccataTATATAATTGCAAATATTCTTTTTC3'  R:5'TCGAGAAAAAGAATATTTGCAATTATATAtatgggtcaggtaTATATAATTGCAAATATTCA3' |
| *XIST*_shRNA4 | F:5'GATCTGCTTTAATTACATTTAATAtacctgacccataTATTAAATGTAATTAAAGCTTTTTC3'  R:5'TCGAGAAAAAGCTTTAATTACATTTAATAtatgggtcaggtaTATTAAATGTAATTAAAGCA3' |
| C328S-VIM | F:5’CAGGTGCAGTCCCTCACCTCTGAAGTGGATGCCCTTAAA3’  R:5’TTTAAGGGCATCCACTTCAGAGGTGAGGGACTGCACCT3’ |
| Y117L-VIM | F:5’AATGACCGCTTCGCCAACCTCATCGACAAGGTGCGCTT3’  R:5’AAGCGCACCTTGTCGATGAGGTTGGCGAAGCGGTCATT3’ |
| The lower case sequence in shRNA constructs forms the loop containing 13 nucleotides before the dicer acts to produce the mature shRNA. | |

**Table S4: List of upregulated genes (cut off padj=0.00009).**

| Gene_name | Gene ID | log2Fold Change | padj |
| --- | --- | --- | --- |
| *XIST* | ENSG00000229807 | 11.79645 | 1.96E-93 |
| *IGF2BP1* | ENSG00000159217 | 8.245026 | 1.30E-50 |
| *FSTL1* | ENSG00000163430 | 8.139257 | 2.61E-40 |
| *CNN3* | ENSG00000117519 | 10.10871 | 1.46E-39 |
| *AKAP12* | ENSG00000131016 | 7.116653 | 3.28E-35 |
| *HMGA2* | ENSG00000149948 | 8.620275 | 3.35E-34 |
| *ADGRL2* | ENSG00000117114 | 6.591145 | 2.79E-32 |
| *Septin 6* | ENSG00000125354 | 7.320673 | 3.65E-32 |
| *CDH2* | ENSG00000170558 | 7.722077 | 9.49E-31 |
| *JAM3* | ENSG00000166086 | 6.779312 | 1.04E-29 |
| *COL4A2* | ENSG00000134871 | 7.216354 | 2.60E-29 |
| *CDKN2A* | ENSG00000147889 | 5.749695 | 2.15E-27 |
| *FLNC* | ENSG00000128591 | 8.10231 | 2.38E-27 |
| *RAB34* | ENSG00000109113 | 8.462863 | 2.76E-27 |
| *MAPRE2* | ENSG00000166974 | 5.772006 | 1.22E-26 |
| *HOXB9* | ENSG00000170689 | 5.695711 | 5.44E-26 |
| *SMOC1* | ENSG00000198732 | 6.677228 | 7.77E-26 |
| *NCAM1* | ENSG00000149294 | 6.737192 | 7.79E-26 |
| *FBN2* | ENSG00000138829 | 5.50309 | 6.04E-24 |
| *ALDH2* | ENSG00000111275 | 8.003519 | 1.01E-23 |
| *LAMC3* | ENSG00000050555 | 5.986133 | 1.31E-21 |
| *CCND2* | ENSG00000118971 | 9.643044 | 7.78E-21 |
| *ANK2* | ENSG00000145362 | 8.760965 | 9.82E-21 |
| *ZEB1* | ENSG00000148516 | 7.687737 | 1.04E-20 |
| *ADAM23* | ENSG00000114948 | 6.225849 | 1.04E-20 |
| *HOXA10* | ENSG00000253293 | 6.876391 | 1.58E-20 |
| *HOXA5* | ENSG00000106004 | 5.081385 | 1.64E-20 |
| *MARK1* | ENSG00000116141 | 5.651885 | 1.66E-20 |
| *ADAMTS3* | ENSG00000156140 | 7.747103 | 4.02E-20 |
| *TCF4* | ENSG00000196628 | 9.505749 | 4.43E-20 |
| *HOXB13* | ENSG00000159184 | 5.58403 | 5.70E-20 |
| *FOXP2* | ENSG00000128573 | 8.044315 | 6.79E-20 |
| *GDF7* | ENSG00000143869 | 6.120914 | 8.16E-20 |
| *LINC02381* | ENSG00000250742 | 6.953446 | 1.78E-19 |
| *PURPL* | ENSG00000250337 | 6.328863 | 2.34E-19 |
| *HOXA3* | ENSG00000105997 | 7.224113 | 3.14E-19 |
| *HOXA11* | ENSG00000005073 | 7.199001 | 4.01E-19 |
| *KIF5A* | ENSG00000155980 | 6.3359 | 4.35E-19 |
| *ITGA1* | ENSG00000213949 | 6.752507 | 9.82E-19 |
| *PLAT* | ENSG00000104368 | 8.445353 | 2.55E-18 |
| *COL4A6* | ENSG00000197565 | 9.021868 | 3.63E-18 |
| *BARX1* | ENSG00000131668 | 10.43802 | 4.06E-18 |
| *NECTIN3* | ENSG00000177707 | 4.969662 | 4.82E-18 |
| *TCF19* | ENSG00000137310 | 5.805573 | 8.18E-18 |
| *LAMB1* | ENSG00000091136 | 4.93954 | 1.32E-17 |
| *CAPN2* | ENSG00000162909 | 4.383822 | 4.43E-17 |
| *ARHGAP22* | ENSG00000128805 | 6.615097 | 5.34E-17 |
| *IGF2BP2* | ENSG00000073792 | 4.670273 | 5.44E-17 |
| *CIB2* | ENSG00000136425 | 5.567477 | 8.61E-17 |
| *ALX4* | ENSG00000052850 | 5.715407 | 8.88E-17 |
| *SNAI2* | ENSG00000019549 | 4.392217 | 1.02E-16 |
| *FMN2* | ENSG00000155816 | 6.063439 | 5.60E-16 |
| *HOTAIRM1* | ENSG00000233429 | 5.231698 | 6.98E-16 |
| *IRF8* | ENSG00000140968 | 6.063818 | 8.63E-16 |
| *COL14A1* | ENSG00000187955 | 5.501122 | 9.83E-16 |
| *WNT5A* | ENSG00000114251 | 4.462759 | 1.74E-15 |
| *HOXA6* | ENSG00000106006 | 6.027171 | 3.08E-15 |
| *IGF2BP3* | ENSG00000136231 | 12.82133 | 3.22E-15 |
| *HLA-B* | ENSG00000234745 | 12.77643 | 4.15E-15 |
| *ADGRL3* | ENSG00000150471 | 5.66715 | 6.18E-15 |
| *INA* | ENSG00000148798 | 12.64546 | 8.20E-15 |
| *C17orf51* | ENSG00000212719 | 12.56534 | 1.26E-14 |
| *PCOLCE2* | ENSG00000163710 | 4.396516 | 1.29E-14 |
| *TFCP2* | ENSG00000135457 | 12.55276 | 1.33E-14 |
| *LAMA1* | ENSG00000101680 | 12.53328 | 1.48E-14 |
| *MIR222HG* | ENSG00000270069 | 5.99406 | 1.56E-14 |
| *COL4A1* | ENSG00000187498 | 12.09111 | 1.71E-14 |
| *MXRA7* | ENSG00000182534 | 4.39984 | 2.28E-14 |
| *DSC3* | ENSG00000134762 | 11.93573 | 3.91E-14 |
| *RAB39A* | ENSG00000179331 | 6.873654 | 4.14E-14 |
| *BCL6B* | ENSG00000161940 | 8.01962 | 5.02E-14 |
| *TP73-AS1* | ENSG00000227372 | 12.29649 | 5.05E-14 |
| *FYN* | ENSG00000010810 | 3.989116 | 6.24E-14 |
| *FLT1* | ENSG00000102755 | 8.037025 | 9.81E-14 |
| *RARB* | ENSG00000077092 | 6.371803 | 1.06E-13 |
| *HOXA9* | ENSG00000078399 | 11.71772 | 1.22E-13 |
| *FOXF2* | ENSG00000137273 | 12.05026 | 1.80E-13 |
| *GSPT2* | ENSG00000189369 | 11.84337 | 5.18E-13 |
| *CASP10* | ENSG00000003400 | 5.502836 | 5.56E-13 |
| *ITGA4* | ENSG00000115232 | 11.69636 | 1.08E-12 |
| *ZEB2* | ENSG00000169554 | 11.64134 | 1.43E-12 |
| *PCDH7* | ENSG00000169851 | 4.022888 | 1.45E-12 |
| *CAMK4* | ENSG00000152495 | 11.60755 | 1.67E-12 |
| *HOXA1* | ENSG00000105991 | 8.702472 | 1.93E-12 |
| *MAP1B* | ENSG00000131711 | 3.759399 | 2.30E-12 |
| *HOXA13* | ENSG00000106031 | 4.546559 | 2.34E-12 |
| *HOXB3* | ENSG00000120093 | 4.563872 | 2.86E-12 |
| *COL25A1* | ENSG00000188517 | 11.48404 | 3.13E-12 |
| *ARPIN* | ENSG00000242498 | 3.758205 | 4.05E-12 |
| *NEXN* | ENSG00000162614 | 6.865973 | 5.42E-12 |
| *TWIST1* | ENSG00000122691 | 8.467432 | 8.16E-12 |
| *ARHGEF6* | ENSG00000129675 | 4.102098 | 8.22E-12 |
| *FOXG1* | ENSG00000176165 | 10.85672 | 9.16E-12 |
| *CDK14* | ENSG00000058091 | 4.146313 | 1.11E-11 |
| *BCL11A* | ENSG00000119866 | 11.21047 | 1.17E-11 |
| *CDH12* | ENSG00000154162 | 11.0138 | 2.99E-11 |
| *COL26A1* | ENSG00000160963 | 6.314866 | 3.26E-11 |
| *FGF2* | ENSG00000138685 | 10.98633 | 3.39E-11 |
| *CNTN1* | ENSG00000018236 | 10.96843 | 3.71E-11 |
| *ADGRB3* | ENSG00000135298 | 7.294488 | 5.22E-11 |
| *AFAP1L1* | ENSG00000157510 | 5.212391 | 6.18E-11 |
| *LAMA4* | ENSG00000112769 | 10.73255 | 1.11E-10 |
| *HOXB4* | ENSG00000182742 | 5.511675 | 1.22E-10 |
| *TWIST2* | ENSG00000233608 | 10.7037 | 1.29E-10 |
| *FNDC1* | ENSG00000164694 | 10.65256 | 1.66E-10 |
| *DOCK3* | ENSG00000088538 | 3.89674 | 1.95E-10 |
| *COL13A1* | ENSG00000197467 | 6.383551 | 2.78E-10 |
| *SOX8* | ENSG00000005513 | 5.255238 | 3.34E-10 |
| *PCDHGA11* | ENSG00000253873 | 4.901264 | 3.81E-10 |
| *ACTN2* | ENSG00000077522 | 7.732494 | 1.08E-09 |
| *SOX6* | ENSG00000110693 | 4.518772 | 1.32E-09 |
| *MAP2K6* | ENSG00000108984 | 3.709073 | 1.63E-09 |
| *PRKCQ* | ENSG00000065675 | 10.14948 | 1.66E-09 |
| *TNFRSF10D* | ENSG00000173530 | 9.71865 | 1.78E-09 |
| *CDH23* | ENSG00000107736 | 4.402438 | 1.95E-09 |
| *COL4A5* | ENSG00000188153 | 3.48696 | 2.16E-09 |
| *CDHR1* | ENSG00000148600 | 9.651466 | 2.21E-09 |
| *PCDH9* | ENSG00000184226 | 3.418713 | 2.38E-09 |
| *PCDH18* | ENSG00000189184 | 9.91912 | 4.70E-09 |
| *COL5A2* | ENSG00000204262 | 3.35551 | 1.45E-08 |
| *ITGA8* | ENSG00000077943 | 9.168304 | 1.30E-07 |
| *LBR* | ENSG00000143815 | 3.593871 | 1.43E-07 |
| *CD40* | ENSG00000101017 | 8.980084 | 2.84E-07 |
| *CDC7* | ENSG00000097046 | 3.559382 | 5.89E-07 |
| *SMAD9* | ENSG00000120693 | 2.784389 | 7.88E-07 |
| *ARHGAP31* | ENSG00000031081 | 3.096402 | 8.21E-07 |
| *MMP2* | ENSG00000087245 | 8.727277 | 8.90E-07 |
| *SKP2* | ENSG00000145604 | 2.771911 | 9.88E-07 |
| *EGFLAM* | ENSG00000164318 | 8.663549 | 1.10E-06 |
| *ADAMTS10* | ENSG00000142303 | 4.201208 | 1.23E-06 |
| *CD109* | ENSG00000156535 | 3.27812 | 1.82E-06 |
| *RASGRF2* | ENSG00000113319 | 3.61194 | 1.87E-06 |
| *COL2A1* | ENSG00000139219 | 3.080835 | 2.28E-06 |
| *MPP6* | ENSG00000105926 | 2.993963 | 2.28E-06 |
| *PINCR* | ENSG00000224294 | 8.499739 | 2.31E-06 |
| *CD70* | ENSG00000125726 | 4.550187 | 2.44E-06 |
| *KRT222* | ENSG00000213424 | 8.451071 | 2.58E-06 |
| *NRXN2* | ENSG00000110076 | 4.226864 | 2.60E-06 |
| *DNM3* | ENSG00000197959 | 3.346947 | 2.99E-06 |
| *CD19* | ENSG00000177455 | 8.405213 | 3.10E-06 |
| *ITGA9* | ENSG00000144668 | 4.800421 | 3.15E-06 |
| *CNTNAP3B* | ENSG00000154529 | 8.391128 | 3.29E-06 |
| *TCL6* | ENSG00000187621 | 8.383689 | 3.38E-06 |
| *CD83* | ENSG00000112149 | 2.894394 | 3.64E-06 |
| *LZTS1* | ENSG00000061337 | 8.35354 | 3.82E-06 |
| *CDCA7* | ENSG00000144354 | 2.785288 | 3.82E-06 |
| *COL24A1* | ENSG00000171502 | 4.430698 | 3.84E-06 |
| *JCAD* | ENSG00000165757 | 2.713746 | 3.88E-06 |
| *KIFAP3* | ENSG00000075945 | 2.698616 | 4.56E-06 |
| *KIF17* | ENSG00000117245 | 3.216778 | 5.82E-06 |
| *LAMA2* | ENSG00000196569 | 3.716597 | 6.18E-06 |
| *FGF5* | ENSG00000138675 | 8.162665 | 8.32E-06 |
| *AKT3* | ENSG00000117020 | 7.905812 | 8.49E-06 |
| *KIF15* | ENSG00000163808 | 3.136117 | 9.94E-06 |
| *ITGB3BP* | ENSG00000142856 | 3.614646 | 1.01E-05 |
| *PAK1* | ENSG00000149269 | 2.421808 | 1.44E-05 |
| *HOXA7* | ENSG00000122592 | 3.302447 | 1.51E-05 |
| *CDON* | ENSG00000064309 | 2.538878 | 1.73E-05 |
| *AMPH* | ENSG00000078053 | 2.580854 | 1.92E-05 |
| *CEP85L* | ENSG00000111860 | 2.728596 | 2.00E-05 |
| *TNFRSF13C* | ENSG00000159958 | 3.154884 | 2.02E-05 |
| *ANGPT1* | ENSG00000154188 | 7.365613 | 2.20E-05 |
| *DNAH6* | ENSG00000115423 | 6.033898 | 2.53E-05 |
| *FAT3* | ENSG00000165323 | 7.329281 | 2.62E-05 |
| *CDC25A* | ENSG00000164045 | 2.269125 | 2.84E-05 |
| *CCSAP* | ENSG00000154429 | 2.767498 | 3.22E-05 |
| *SOX21* | ENSG00000125285 | 5.969039 | 3.49E-05 |
| *HOXA2* | ENSG00000105996 | 7.227463 | 3.50E-05 |
| *HSF2* | ENSG00000025156 | 2.667201 | 3.85E-05 |
| *HOXA4* | ENSG00000197576 | 3.091659 | 4.62E-05 |
| *FGF17* | ENSG00000158815 | 5.801019 | 4.90E-05 |
| *RAB9B* | ENSG00000123570 | 2.95757 | 5.23E-05 |
| *PRKD1* | ENSG00000184304 | 2.453299 | 5.58E-05 |
| *TLR6* | ENSG00000174130 | 4.521907 | 5.69E-05 |
| *MIR221* | ENSG00000207870 | 7.665559 | 5.76E-05 |
| *PLXNA2* | ENSG00000076356 | 3.033146 | 5.80E-05 |
| *ALDH8A1* | ENSG00000118514 | 4.900099 | 6.10E-05 |
| *MN1* | ENSG00000169184 | 2.6547 | 6.91E-05 |
| *TFPI2* | ENSG00000105825 | 3.185966 | 6.97E-05 |
| *ADGRA2* | ENSG00000020181 | 2.740551 | 7.32E-05 |
| *CEP112* | ENSG00000154240 | 2.829766 | 7.36E-05 |
| *FAT4* | ENSG00000196159 | 2.282895 | 7.50E-05 |
| *TTBK2* | ENSG00000128881 | 2.501309 | 7.53E-05 |
| *PAK3* | ENSG00000077264 | 5.712818 | 7.54E-05 |
| *RASSF2* | ENSG00000101265 | 2.731273 | 8.29E-05 |

**Table S5: List of downregulated genes (cut off padj=0.00009).**

| **Gene Name** | **Gene ID** | **Log2fold Change** | **padj** |
| --- | --- | --- | --- |
| *KRT19* | ENSG00000171345 | -12.5017 | 1.04E-92 |
| *KRT80* | ENSG00000167767 | -13.0018 | 3.65E-75 |
| *KRT8* | ENSG00000170421 | -10.1552 | 5.61E-74 |
| *DSCAM-AS1* | ENSG00000235123 | -11.5496 | 1.76E-62 |
| *S100A16* | ENSG00000188643 | -10.7403 | 8.54E-58 |
| *CD24* | ENSG00000272398 | -8.60434 | 9.68E-58 |
| *CDH1* | ENSG00000039068 | -8.03511 | 1.38E-56 |
| *ESRP2* | ENSG00000103067 | -9.60887 | 2.27E-49 |
| *ESRP1* | ENSG00000104413 | -9.95392 | 2.27E-49 |
| *TACSTD2* | ENSG00000184292 | -11.8701 | 3.22E-48 |
| *ERBB3* | ENSG00000065361 | -8.03517 | 4.43E-48 |
| *L1CAM* | ENSG00000198910 | -9.39655 | 3.12E-47 |
| *WISP2* | ENSG00000064205 | -11.8286 | 8.13E-47 |
| *EPPK1* | ENSG00000261150 | -9.79614 | 1.69E-46 |
| *CLDN3* | ENSG00000165215 | -13.4255 | 1.18E-44 |
| *PKP3* | ENSG00000184363 | -11.4093 | 1.18E-44 |
| *FOXA1* | ENSG00000129514 | -7.20191 | 1.27E-44 |
| *CLDN7* | ENSG00000181885 | -9.11937 | 1.02E-42 |
| *CLDN4* | ENSG00000189143 | -9.19594 | 1.62E-42 |
| *WNT7B* | ENSG00000188064 | -9.18573 | 5.58E-42 |
| *RAB25* | ENSG00000132698 | -9.4677 | 7.90E-42 |
| *PPL* | ENSG00000118898 | -8.41424 | 1.51E-41 |
| *KRT18* | ENSG00000111057 | -7.56932 | 6.86E-41 |
| *KRT81* | ENSG00000205426 | -15.9626 | 4.93E-39 |
| *ADGRG1* | ENSG00000205336 | -11.7431 | 6.99E-39 |
| *CDH3* | ENSG00000062038 | -7.39085 | 1.16E-37 |
| *EPCAM* | ENSG00000119888 | -6.7076 | 7.64E-36 |
| *S100A14* | ENSG00000189334 | -11.3462 | 1.93E-33 |
| *PDGFB* | ENSG00000100311 | -8.66131 | 1.65E-32 |
| *KRT86* | ENSG00000170442 | -9.43315 | 1.21E-31 |
| *ITGB4* | ENSG00000132470 | -7.09122 | 8.61E-27 |
| *NUPR1* | ENSG00000176046 | -6.76598 | 1.42E-26 |
| *BCL3* | ENSG00000069399 | -6.41802 | 5.49E-26 |
| *NECTIN4* | ENSG00000143217 | -6.8834 | 1.33E-25 |
| *PTPN6* | ENSG00000111679 | -6.68543 | 9.79E-25 |
| *KRT8P3* | ENSG00000254285 | -10.1481 | 1.30E-24 |
| *CLDN9* | ENSG00000213937 | -10.5792 | 1.55E-24 |
| *APOD* | ENSG00000189058 | -6.66367 | 3.88E-24 |
| *TNFRSF12A* | ENSG00000006327 | -5.95843 | 2.38E-23 |
| *TGFBI* | ENSG00000120708 | -9.49209 | 3.90E-23 |
| *SHH* | ENSG00000164690 | -6.98255 | 6.98E-23 |
| *PCDHGB5* | ENSG00000276547 | -7.79629 | 7.94E-23 |
| *PARP12* | ENSG00000059378 | -7.62925 | 8.78E-23 |
| *CLDN1* | ENSG00000163347 | -7.09195 | 2.18E-22 |
| *HMCN1* | ENSG00000143341 | -5.09272 | 5.33E-22 |
| *CD44* | ENSG00000026508 | -6.34163 | 1.50E-21 |
| *BCAS3* | ENSG00000141376 | -5.08945 | 5.08E-21 |
| *JUNB* | ENSG00000171223 | -4.9864 | 4.86E-20 |
| *IKBKE* | ENSG00000263528 | -6.87871 | 9.30E-20 |
| *LAMB3* | ENSG00000196878 | -7.44425 | 2.08E-19 |
| *YBX2* | ENSG00000006047 | -5.31419 | 2.94E-19 |
| *KIF12* | ENSG00000136883 | -5.68644 | 4.60E-19 |
| *TGFA* | ENSG00000163235 | -5.3384 | 5.31E-19 |
| *PARD6B* | ENSG00000124171 | -5.10334 | 9.58E-19 |
| *ICAM1* | ENSG00000090339 | -6.44375 | 9.71E-19 |
| *CTNND2* | ENSG00000169862 | -14.3521 | 1.25E-18 |
| *COL3A1* | ENSG00000168542 | -4.46134 | 2.29E-18 |
| *PCDH1* | ENSG00000156453 | -5.71508 | 2.59E-18 |
| *PCDHB2* | ENSG00000112852 | -6.21841 | 3.75E-18 |
| *TFAP2C* | ENSG00000087510 | -4.48661 | 3.81E-18 |
| *S100A9* | ENSG00000163220 | -6.7512 | 4.51E-18 |
| *BOK* | ENSG00000176720 | -5.39943 | 4.90E-18 |
| *CD9* | ENSG00000010278 | -4.54939 | 6.14E-18 |
| *CDC42BPG* | ENSG00000171219 | -5.20928 | 9.57E-18 |
| *FIBCD1* | ENSG00000130720 | -5.56018 | 2.00E-17 |
| *S100A11* | ENSG00000163191 | -4.40166 | 3.41E-17 |
| *HSPB8* | ENSG00000152137 | -4.34555 | 5.70E-17 |
| *AIFM2* | ENSG00000042286 | -5.09404 | 1.77E-16 |
| *PARP9* | ENSG00000138496 | -4.99279 | 2.04E-16 |
| *RHOD* | ENSG00000173156 | -13.3707 | 2.27E-16 |
| *SOX3* | ENSG00000134595 | -6.13532 | 2.59E-16 |
| *RAB26* | ENSG00000167964 | -5.14424 | 2.89E-16 |
| *NCAM2* | ENSG00000154654 | -13.2264 | 5.78E-16 |
| *PARP14* | ENSG00000173193 | -9.96466 | 9.67E-16 |
| *BIK* | ENSG00000100290 | -4.95456 | 1.91E-15 |
| *CDKN1A* | ENSG00000124762 | -4.22775 | 3.01E-15 |
| *RAB17* | ENSG00000124839 | -12.7669 | 3.61E-15 |
| *ITGB6* | ENSG00000115221 | -12.736 | 6.79E-15 |
| *PKP1* | ENSG00000081277 | -12.8297 | 7.34E-15 |
| *DSCAM* | ENSG00000171587 | -12.5465 | 7.91E-15 |
| *PCDHB3* | ENSG00000113205 | -9.39072 | 2.70E-14 |
| *S100A6* | ENSG00000197956 | -12.3847 | 3.88E-14 |
| *JUP* | ENSG00000173801 | -4.7403 | 4.40E-14 |
| *BCAM* | ENSG00000187244 | -4.80314 | 6.64E-14 |
| *PCDHA6* | ENSG00000081842 | -12.2947 | 1.27E-13 |
| *KDF1* | ENSG00000175707 | -12.3892 | 1.45E-13 |
| *PLXNB1* | ENSG00000164050 | -4.63775 | 1.88E-13 |
| *AMIGO2* | ENSG00000139211 | -4.19602 | 3.07E-13 |
| *ITGA3* | ENSG00000005884 | -4.4519 | 8.26E-13 |
| *FMN1* | ENSG00000248905 | -6.74342 | 1.27E-12 |
| *MAPK11* | ENSG00000185386 | -4.64408 | 1.34E-12 |
| *PARD6A* | ENSG00000102981 | -5.05535 | 1.39E-12 |
| *RAB27B* | ENSG00000041353 | -5.19445 | 2.15E-12 |
| *KRTCAP3* | ENSG00000157992 | -4.809 | 2.66E-12 |
| *SDC4* | ENSG00000124145 | -3.55913 | 6.44E-12 |
| *DRAM1* | ENSG00000136048 | -3.91881 | 8.11E-12 |
| *PCDHA11* | ENSG00000249158 | -11.3521 | 8.42E-12 |
| *PCDHGA2* | ENSG00000081853 | -11.2075 | 1.70E-11 |
| *KRT7* | ENSG00000135480 | -11.1068 | 1.85E-11 |
| *KRT15* | ENSG00000171346 | -11.2416 | 1.95E-11 |
| *FGF13* | ENSG00000129682 | -5.26285 | 2.46E-11 |
| *TGFB2* | ENSG00000092969 | -4.75549 | 3.11E-11 |
| *BMF* | ENSG00000104081 | -3.66168 | 3.35E-11 |
| *NFKBIA* | ENSG00000100906 | -3.88707 | 4.27E-11 |
| *PCDHA12* | ENSG00000251664 | -8.89581 | 5.35E-11 |
| *SDC1* | ENSG00000115884 | -3.58472 | 9.29E-11 |
| *PCDHB13* | ENSG00000187372 | -10.6909 | 1.89E-10 |
| *MAP10* | ENSG00000212916 | -10.6119 | 2.79E-10 |
| *KRT83* | ENSG00000170523 | -8.418 | 3.58E-10 |
| *MAP2* | ENSG00000078018 | -4.13019 | 3.65E-10 |
| *PCDHB14* | ENSG00000120327 | -6.13142 | 4.48E-10 |
| *ADAMTS19* | ENSG00000145808 | -3.3851 | 5.37E-10 |
| *CCND1* | ENSG00000110092 | -3.27663 | 7.67E-10 |
| *PARP10* | ENSG00000178685 | -4.28497 | 8.90E-10 |
| *NPR3* | ENSG00000113389 | -4.42421 | 1.04E-09 |
| *PCDHB16* | ENSG00000272674 | -10.2591 | 1.70E-09 |
| *MMP17* | ENSG00000198598 | -4.00925 | 3.25E-09 |
| *PCDHA7* | ENSG00000204963 | -10.3054 | 3.99E-09 |
| *CDH13* | ENSG00000140945 | -10.0494 | 4.07E-09 |
| *WNT4* | ENSG00000162552 | -5.39344 | 6.65E-09 |
| *SRMS* | ENSG00000125508 | -6.82755 | 8.40E-09 |
| *RAB20* | ENSG00000139832 | -3.55444 | 8.65E-09 |
| *PCDHGA1* | ENSG00000204956 | -5.43062 | 8.80E-09 |
| *MACC1* | ENSG00000183742 | -9.69882 | 1.00E-08 |
| *SOX11* | ENSG00000176887 | -9.53339 | 1.01E-08 |
| *CD22* | ENSG00000012124 | -6.04625 | 1.29E-08 |
| *BCL6* | ENSG00000113916 | -3.39678 | 1.40E-08 |
| *NRCAM* | ENSG00000091129 | -3.1041 | 4.12E-08 |
| *COL12A1* | ENSG00000111799 | -3.33952 | 4.12E-08 |
| *TP63* | ENSG00000073282 | -5.98185 | 4.95E-08 |
| *PCDHGB1* | ENSG00000254221 | -5.03612 | 5.86E-08 |
| *ADGRF4* | ENSG00000153294 | -5.60247 | 6.66E-08 |
| *TNFRSF18* | ENSG00000186891 | -9.33523 | 6.80E-08 |
| *RAB37* | ENSG00000172794 | -4.28798 | 7.70E-08 |
| *KIF1A* | ENSG00000130294 | -3.36972 | 9.76E-08 |
| *ARHGEF5* | ENSG00000050327 | -9.3588 | 9.81E-08 |
| *LMNA* | ENSG00000160789 | -3.45521 | 9.81E-08 |
| *CDH24* | ENSG00000139880 | -3.24053 | 1.76E-07 |
| *KRT17* | ENSG00000128422 | -6.0269 | 1.82E-07 |
| *AJAP1* | ENSG00000196581 | -9.22878 | 1.85E-07 |
| *PCDHA13* | ENSG00000239389 | -9.39029 | 2.69E-07 |
| *LAMA5* | ENSG00000130702 | -3.45141 | 2.76E-07 |
| *NRBP1* | ENSG00000115216 | -2.8634 | 3.01E-07 |
| *EMP2* | ENSG00000213853 | -2.6466 | 5.67E-07 |
| *ADGRG6* | ENSG00000112414 | -3.10278 | 5.95E-07 |
| *PCDHGB2* | ENSG00000253910 | -4.95149 | 6.53E-07 |
| *COL18A1* | ENSG00000182871 | -2.97089 | 7.64E-07 |
| *ESPN* | ENSG00000187017 | -10.4701 | 8.41E-07 |
| *MMP15* | ENSG00000102996 | -2.90133 | 1.17E-06 |
| *VASP* | ENSG00000125753 | -2.76056 | 1.20E-06 |
| *MUC1* | ENSG00000185499 | -3.0739 | 1.25E-06 |
| *KRT24* | ENSG00000167916 | -8.8525 | 1.40E-06 |
| *MIR4737* | ENSG00000264049 | -4.26976 | 1.67E-06 |
| *PCDHAC1* | ENSG00000248383 | -8.7053 | 1.92E-06 |
| *CEACAM6* | ENSG00000086548 | -8.32546 | 2.00E-06 |
| *PCDHB9* | ENSG00000177839 | -8.63476 | 2.60E-06 |
| *TFPI* | ENSG00000003436 | -3.14726 | 2.68E-06 |
| *TUBB3* | ENSG00000258947 | -4.21155 | 2.96E-06 |
| *TNFSF10* | ENSG00000121858 | -4.99596 | 3.23E-06 |
| *S100P* | ENSG00000163993 | -9.64223 | 3.28E-06 |
| *JDP2* | ENSG00000140044 | -2.69422 | 3.58E-06 |
| *MAMDC4* | ENSG00000177943 | -2.97366 | 3.67E-06 |
| *PCDHB8* | ENSG00000120322 | -8.09131 | 3.92E-06 |
| *NKILA* | ENSG00000278709 | -2.9481 | 4.27E-06 |
| *HSPA6* | ENSG00000173110 | -4.22098 | 4.52E-06 |
| *MAP3K8* | ENSG00000107968 | -2.97908 | 6.03E-06 |
| *MUC3A* | ENSG00000169894 | -4.26317 | 6.56E-06 |
| *JUND* | ENSG00000130522 | -2.45081 | 8.10E-06 |
| *CAMK2B* | ENSG00000058404 | -3.09354 | 9.21E-06 |
| *PCDHA10* | ENSG00000250120 | -6.38713 | 9.52E-06 |
| *CAV1* | ENSG00000105974 | -2.46421 | 9.64E-06 |
| *ABLIM2* | ENSG00000163995 | -3.21067 | 9.64E-06 |
| *PRKCD* | ENSG00000163932 | -2.65035 | 1.29E-05 |
| *NOTCH3* | ENSG00000074181 | -2.93517 | 1.34E-05 |
| *AKT1* | ENSG00000142208 | -2.73676 | 1.74E-05 |
| *KRT13* | ENSG00000171401 | -8.21214 | 1.74E-05 |
| *PCDHB4* | ENSG00000081818 | -7.64107 | 1.94E-05 |
| *NPDC1* | ENSG00000107281 | -2.44297 | 1.94E-05 |
| *TNFAIP2* | ENSG00000185215 | -2.58357 | 1.98E-05 |
| *NECTIN2* | ENSG00000130202 | -2.58125 | 2.00E-05 |
| *ITGAM* | ENSG00000169896 | -6.14166 | 2.39E-05 |
| *ARPC1B* | ENSG00000130429 | -2.79233 | 2.86E-05 |
| *AREG* | ENSG00000109321 | -3.63775 | 2.99E-05 |
| *FILIP1L* | ENSG00000168386 | -3.23019 | 3.01E-05 |
| *MMP19* | ENSG00000123342 | -7.54883 | 3.03E-05 |
| *MMP9* | ENSG00000100985 | -4.37314 | 3.09E-05 |
| *PTPRJ* | ENSG00000149177 | -2.31114 | 3.13E-05 |
| *FLNA* | ENSG00000196924 | -2.75191 | 3.26E-05 |
| *CAPN1* | ENSG00000014216 | -2.69277 | 3.35E-05 |
| *HSPD1P11* | ENSG00000251348 | -4.13806 | 3.55E-05 |
| *MIR149* | ENSG00000207611 | -4.66217 | 3.89E-05 |
| *KISS1* | ENSG00000170498 | -7.93301 | 4.80E-05 |
| *MAPKAPK2* | ENSG00000162889 | -2.35947 | 4.95E-05 |
| *PXN* | ENSG00000089159 | -2.27782 | 4.97E-05 |
| *VDR* | ENSG00000111424 | -2.39643 | 5.42E-05 |
| *SMAGP* | ENSG00000170545 | -2.34217 | 5.44E-05 |
| *CD63* | ENSG00000135404 | -2.34269 | 5.45E-05 |
| *RAB15* | ENSG00000139998 | -2.26252 | 7.38E-05 |
| *CARD9* | ENSG00000187796 | -2.66765 | 7.40E-05 |
| *PCDHGA5* | ENSG00000253485 | -7.29152 | 7.63E-05 |
| *BBC3* | ENSG00000105327 | -2.52634 | 7.68E-05 |
| *TNS2* | ENSG00000111077 | -2.59524 | 7.97E-05 |
| *CEACAM5* | ENSG00000105388 | -7.2521 | 8.32E-05 |
| *KRT16* | ENSG00000186832 | -7.79991 | 9.45E-05 |

**Table S6 : List of upregulated lnRNAs (cut off padj=0.00009).**

| **Gene Name** | **Gene ID** | **Log2fold Change** | **padj** |
| --- | --- | --- | --- |
| *XIST* | ENSG00000229807 | 11.79645137 | 1.96E-93 |
| *LINC02381* | ENSG00000250742 | 6.953445703 | 1.78E-19 |
| *PURPL* | ENSG00000250337 | 6.328863193 | 2.34E-19 |
| *MIR222HG* | ENSG00000270069 | 5.994059507 | 1.56E-14 |
| *AC027031.2* | ENSG00000254615 | 5.128079486 | 9.98E-13 |
| *AL035446.1* | ENSG00000234147 | 7.509471414 | 8.27E-12 |
| *AL133325.3* | ENSG00000278041 | 10.25216513 | 1.60E-10 |
| *FAM239B* | ENSG00000205663 | 9.696877626 | 1.26E-08 |
| *LINC02241* | ENSG00000251629 | 9.620849091 | 1.76E-08 |
| *AC103702.2* | ENSG00000272763 | 9.584780353 | 2.07E-08 |
| *TSIX* | ENSG00000270641 | 8.852012634 | 6.88E-08 |
| *AC015522.1* | ENSG00000254202 | 9.251935664 | 8.91E-08 |
| *LINC00491* | ENSG00000250682 | 9.186133789 | 1.18E-07 |
| *RAMP2-AS1* | ENSG00000197291 | 6.877939113 | 1.77E-07 |
| *LINC01033* | ENSG00000249069 | 8.819571635 | 5.80E-07 |
| *AL356489.2* | ENSG00000260947 | 8.750532639 | 7.92E-07 |
| *AL513318.2* | ENSG00000269994 | 8.756358592 | 8.03E-07 |
| *SOX21-AS1* | ENSG00000227640 | 6.630286843 | 1.45E-06 |
| *PINCR* | ENSG00000224294 | 8.499738862 | 2.31E-06 |
| *LINC00470* | ENSG00000132204 | 6.394021607 | 3.86E-06 |
| *AL445250.1* | ENSG00000225096 | 7.810485755 | 4.19E-06 |
| *LINC01234* | ENSG00000249550 | 7.756791857 | 5.14E-06 |
| *AC233976.1* | ENSG00000229151 | 8.283465103 | 5.28E-06 |
| *SH3RF3-AS1* | ENSG00000259863 | 8.175078669 | 8.17E-06 |
| *FOXCUT* | ENSG00000280916 | 6.149454375 | 9.33E-06 |
| *DSCR8* | ENSG00000198054 | 8.096367971 | 1.08E-05 |
| *AL158055.1* | ENSG00000226530 | 7.990362422 | 1.63E-05 |
| *AC025575.2* | ENSG00000258053 | 7.99019662 | 1.73E-05 |
| *FENDRR* | ENSG00000268388 | 7.880557066 | 2.52E-05 |
| *ERVMER61-1* | ENSG00000230426 | 7.327968096 | 2.59E-05 |
| *ZNF582-AS1* | ENSG00000267454 | 7.854726374 | 2.92E-05 |
| *LINC02315* | ENSG00000251363 | 7.830396631 | 3.02E-05 |
| *Z68871.1* | ENSG00000239407 | 2.943656954 | 3.07E-05 |
| *AC009275.1* | ENSG00000273297 | 4.707067687 | 3.53E-05 |
| *LINC01976* | ENSG00000261514 | 7.788306456 | 3.58E-05 |
| *LINC00896* | ENSG00000236499 | 5.2395894 | 3.58E-05 |
| *AL365361.1* | ENSG00000259834 | 7.769445937 | 3.82E-05 |
| *LINC00648* | ENSG00000259129 | 4.74170971 | 4.18E-05 |
| *AC027020.2* | ENSG00000270127 | 3.530275396 | 4.44E-05 |
| *AC021504.1* | ENSG00000267313 | 7.675041377 | 5.50E-05 |
| *AL022324.3* | ENSG00000272942 | 7.630929262 | 6.47E-05 |
| *LINC01297* | ENSG00000274827 | 7.653421816 | 6.76E-05 |
| *LINC01876* | ENSG00000226383 | 4.271039981 | 7.16E-05 |
| *AL592295.4* | ENSG00000283696 | 5.060160122 | 8.60E-05 |

**Table S7: List of downregulated lnRNAs (cut off padj=0.00009).**

| **Gene name** | **Gene ID** | **log2Fold change** | **padj** |
| --- | --- | --- | --- |
| *GATA3-AS1* | ENSG00000197308 | -8.40024 | 1.71E-29 |
| *MIR9-3HG* | ENSG00000255571 | -7.74173 | 2.47E-17 |
| *AP001816.1* | ENSG00000254531 | -5.80986 | 4.42E-17 |
| *C17orf82* | ENSG00000187013 | -6.65662 | 1.33E-16 |
| *AC144831.1* | ENSG00000261888 | -9.6419 | 9.19E-15 |
| *LINC01293* | ENSG00000230836 | -7.1369 | 1.26E-14 |
| *LINC00992* | ENSG00000248663 | -4.72382 | 3.88E-14 |
| *AC004233.3* | ENSG00000272079 | -5.15436 | 6.35E-12 |
| *AC074135.1* | ENSG00000267886 | -11.218 | 1.73E-11 |
| *AC020916.1* | ENSG00000267519 | -4.66826 | 5.24E-11 |
| *AC141928.1* | ENSG00000250986 | -4.7547 | 1.02E-10 |
| *LINC01468* | ENSG00000231131 | -10.7134 | 2.45E-10 |
| *AL355001.2* | ENSG00000275964 | -3.68048 | 2.62E-10 |
| *AL135818.2* | ENSG00000260810 | -6.20619 | 1.13E-09 |
| *AL035661.1* | ENSG00000274173 | -10.5793 | 1.21E-09 |
| *AC044784.1* | ENSG00000223808 | -10.2426 | 1.94E-09 |
| *FAM225A* | ENSG00000231528 | -10.1514 | 3.87E-09 |
| *AC144450.1* | ENSG00000203635 | -10.0804 | 4.29E-09 |
| *LINC00052* | ENSG00000259527 | -9.73082 | 6.00E-09 |
| *PSMG3-AS1* | ENSG00000230487 | -3.6883 | 6.33E-09 |
| *AC015712.1* | ENSG00000232386 | -6.69834 | 7.38E-09 |
| *AL590004.4* | ENSG00000260604 | -9.9135 | 8.83E-09 |
| *AP000439.2* | ENSG00000255774 | -9.92916 | 9.10E-09 |
| *LINC01503* | ENSG00000233901 | -4.62263 | 1.93E-08 |
| *AC096733.2* | ENSG00000273472 | -3.53195 | 2.25E-08 |
| *AC022034.2* | ENSG00000237807 | -9.66219 | 2.45E-08 |
| *C9orf163* | ENSG00000196366 | -4.3994 | 3.78E-08 |
| *AL365181.2* | ENSG00000272068 | -7.28242 | 4.12E-08 |
| *PVT1* | ENSG00000249859 | -3.06311 | 5.25E-08 |
| *AL390719.2* | ENSG00000272141 | -9.58839 | 5.33E-08 |
| *C9orf106* | ENSG00000179082 | -9.43835 | 6.93E-08 |
| *AC015922.3* | ENSG00000265519 | -9.3775 | 9.11E-08 |
| *FAM111A-DT* | ENSG00000245571 | -9.37063 | 9.83E-08 |
| *LINC00886* | ENSG00000240875 | -4.26358 | 1.17E-07 |
| *MIR200CHG* | ENSG00000257084 | -9.35895 | 1.27E-07 |
| *AP003559.1* | ENSG00000256443 | -5.32785 | 1.80E-07 |
| *AL512625.2* | ENSG00000229422 | -3.34184 | 1.81E-07 |
| *AC012307.1* | ENSG00000228873 | -9.38298 | 2.10E-07 |
| *AC008014.1* | ENSG00000257261 | -4.13124 | 2.36E-07 |
| *AC009237.14* | ENSG00000272913 | -9.18672 | 2.56E-07 |
| *AC008556.1* | ENSG00000277013 | -6.96466 | 2.76E-07 |
| *SNHG19* | ENSG00000260260 | -2.94687 | 4.38E-07 |
| *LINC02568* | ENSG00000259459 | -9.0107 | 5.33E-07 |
| *AC007342.4* | ENSG00000261804 | -8.64953 | 5.72E-07 |
| *LINC00847* | ENSG00000245060 | -3.02844 | 1.03E-06 |
| *AC010735.2* | ENSG00000272622 | -6.74235 | 1.26E-06 |
| *LINC01213* | ENSG00000244541 | -8.76016 | 1.51E-06 |
| *AC061992.1* | ENSG00000266970 | -4.75367 | 1.94E-06 |
| *DIO3OS* | ENSG00000258498 | -5.96001 | 2.22E-06 |
| *AC006206.2* | ENSG00000256417 | -8.65963 | 2.38E-06 |
| *LINC02021* | ENSG00000249846 | -3.41324 | 2.86E-06 |
| *AC090114.2* | ENSG00000273270 | -2.77024 | 3.05E-06 |
| *AC092279.1* | ENSG00000268362 | -3.31659 | 3.28E-06 |
| *AC096888.1* | ENSG00000244564 | -8.56423 | 3.50E-06 |
| *AL021807.1* | ENSG00000272468 | -4.55678 | 3.82E-06 |
| *FAM225B* | ENSG00000225684 | -8.62311 | 4.18E-06 |
| *EPB41L4A-AS2* | ENSG00000278921 | -3.56267 | 4.90E-06 |
| *SNHG18* | ENSG00000250786 | -7.91721 | 7.49E-06 |
| *LINC01750* | ENSG00000231437 | -6.34424 | 8.06E-06 |
| *AC083967.1* | ENSG00000254337 | -7.87222 | 8.32E-06 |
| *LINC02015* | ENSG00000231574 | -7.86906 | 8.34E-06 |
| *LINC01016* | ENSG00000249346 | -7.82734 | 9.95E-06 |
| *AC100860.1* | ENSG00000253266 | -8.34184 | 1.10E-05 |
| *LINC00885* | ENSG00000224652 | -7.93839 | 1.11E-05 |
| *AL645608.9* | ENSG00000273443 | -8.35171 | 1.13E-05 |
| *HAGLROS* | ENSG00000226363 | -4.28912 | 1.18E-05 |
| *UCA1* | ENSG00000214049 | -8.26012 | 1.23E-05 |
| *AC147651.1* | ENSG00000223855 | -4.18837 | 1.30E-05 |
| *C15orf59-AS1* | ENSG00000260469 | -3.70555 | 1.30E-05 |
| *LINC00628* | ENSG00000280924 | -8.21329 | 1.60E-05 |
| *AL357558.1* | ENSG00000228503 | -8.18389 | 1.89E-05 |
| *AL357558.2* | ENSG00000234967 | -8.13677 | 2.22E-05 |
| *AC005332.6* | ENSG00000277476 | -2.89766 | 2.22E-05 |
| *AC091271.1* | ENSG00000273702 | -2.82113 | 2.35E-05 |
| *AL117329.1* | ENSG00000224271 | -8.08866 | 2.67E-05 |
| *AP000439.5* | ENSG00000285094 | -8.09905 | 2.97E-05 |
| *LINC00898* | ENSG00000205634 | -8.1746 | 3.47E-05 |
| *LINC01671* | ENSG00000225431 | -7.96645 | 4.27E-05 |
| *AC011416.3* | ENSG00000283897 | -7.94267 | 4.61E-05 |
| *LINC00994* | ENSG00000189196 | -7.95062 | 4.62E-05 |
| *LINC00865* | ENSG00000232229 | -7.4451 | 4.71E-05 |
| *AP002884.1* | ENSG00000250303 | -3.14553 | 4.71E-05 |
| *AC005618.1* | ENSG00000272070 | -4.40236 | 4.81E-05 |
| *LINC01376* | ENSG00000236204 | -4.25144 | 5.51E-05 |
| *AL133342.1* | ENSG00000278231 | -7.89219 | 5.58E-05 |
| *ERVE-1* | ENSG00000267259 | -7.36674 | 5.74E-05 |
| *AL356740.1* | ENSG00000267868 | -5.99969 | 6.19E-05 |
| *LINC01918* | ENSG00000226508 | -4.15419 | 6.49E-05 |
| *BX539320.1* | ENSG00000278869 | -7.88744 | 7.00E-05 |
| *LINC01665* | ENSG00000235343 | -7.82432 | 7.68E-05 |
| *AC005993.1* | ENSG00000266869 | -7.2285 | 8.47E-05 |
| *LINC01637* | ENSG00000237476 | -5.0912 | 8.73E-05 |
| *LINC01977* | ENSG00000262772 | -3.07166 | 9.91E-05 |
