## Supplementary figures and images for "A single cysteine residue in vimentin regulates long non-coding RNA *XIST* to suppress epithelial-mesenchymal transition and stemness in breast cancer"

### CELLULAR_RESPONSE_TO_INTERLEUKIN_12(GO_0071349)_22.png

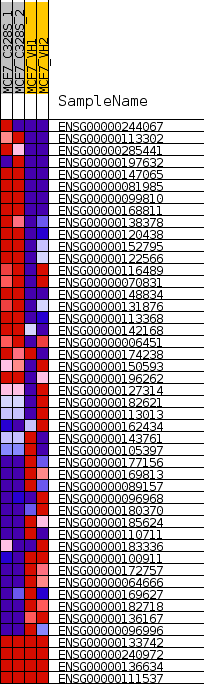

### enplot_CELLULAR_RESPONSE_TO_INTERLEUKIN_12(GO_0071349)_21.png

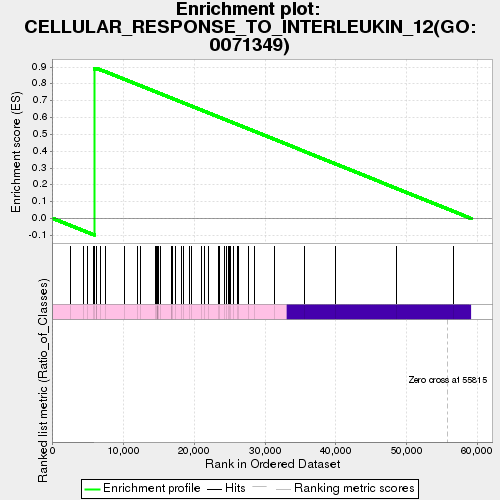

### enplot_INTERLEUKIN_12_MEDIATED_SIGNALING_PATHWAY(GO_0035722)_24.png

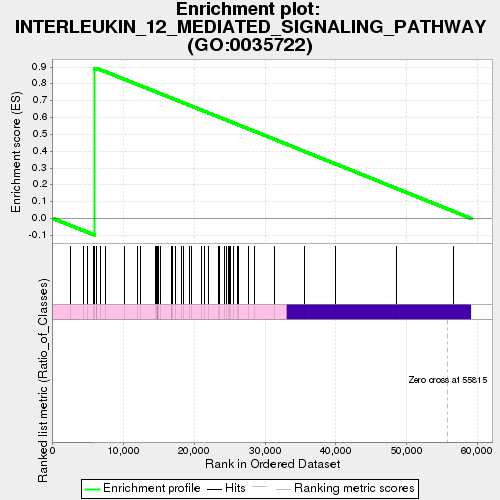

### gset_rnd_es_dist_23.png

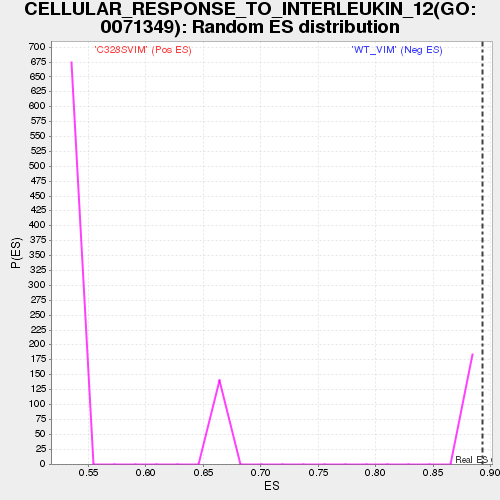

### gset_rnd_es_dist_26.png

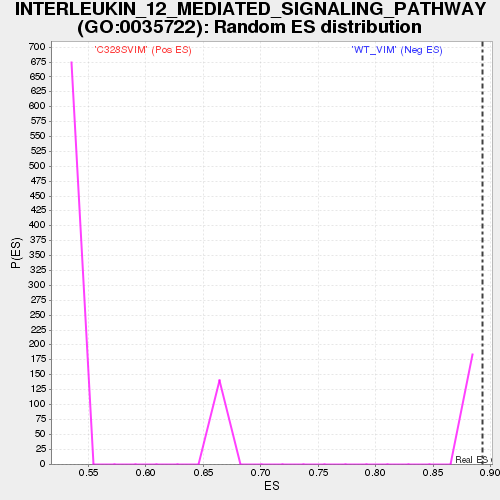

### gset_rnd_es_dist_41.png

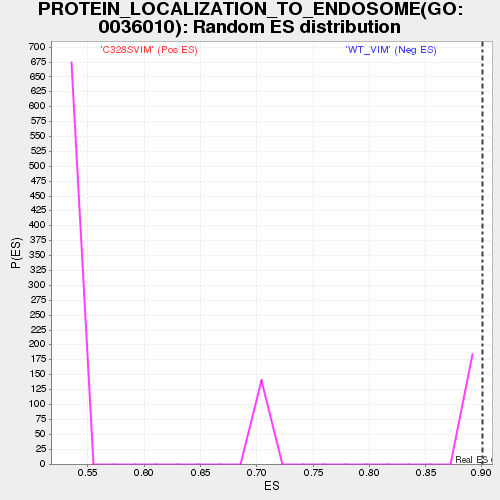

### PROTEIN_LOCALIZATION_TO_ENDOSOME(GO_0036010)_40.png

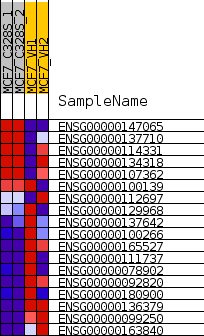
