## Supplementary figures and images for "A single cysteine residue in vimentin regulates long non-coding RNA *XIST* to suppress epithelial-mesenchymal transition and stemness in breast cancer"

### ATRIAL_SEPTUM_DEVELOPMENT(GO_0003283)_52.png

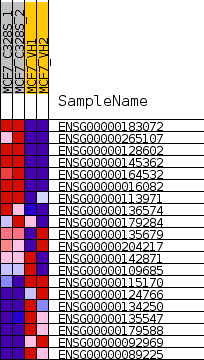

### butterfly_plot.png

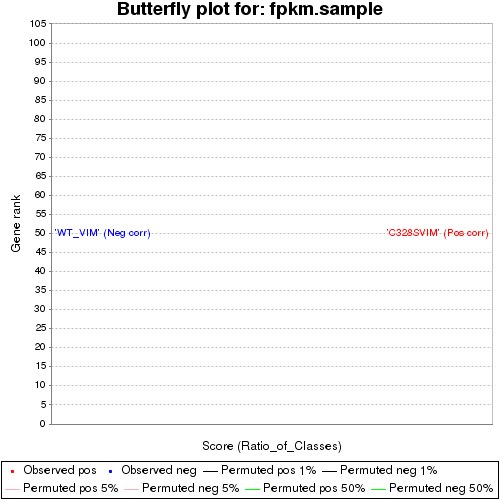

### C328SVIMvsWT_VIM_all.bar.pdf

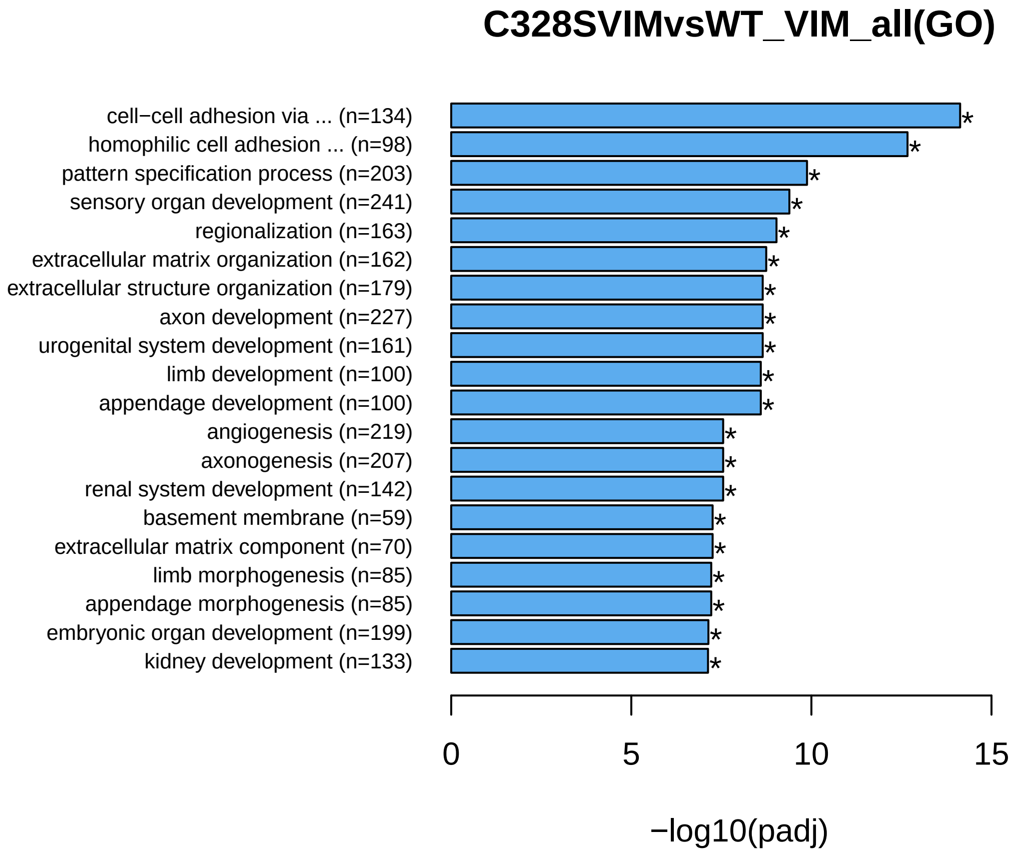

### C328SVIMvsWT_VIM_all.bar.png

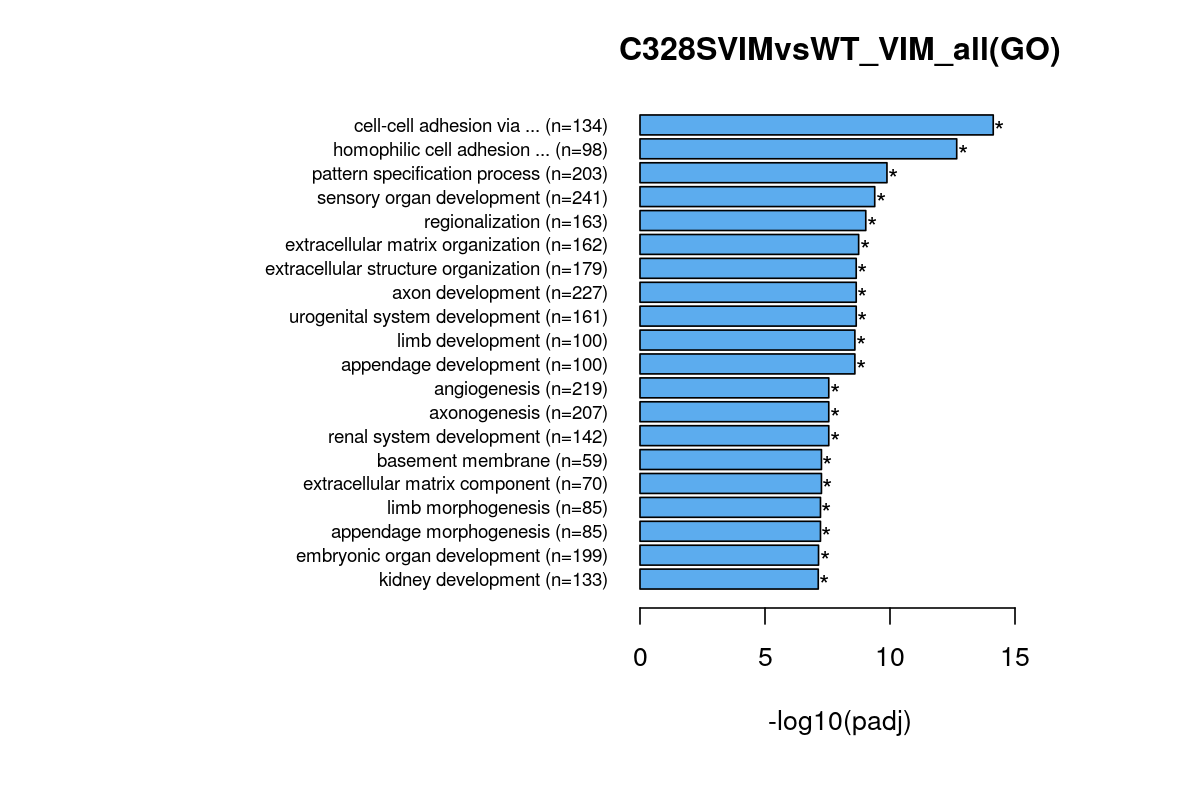

### C328SVIMvsWT_VIM_all.BPbar.pdf

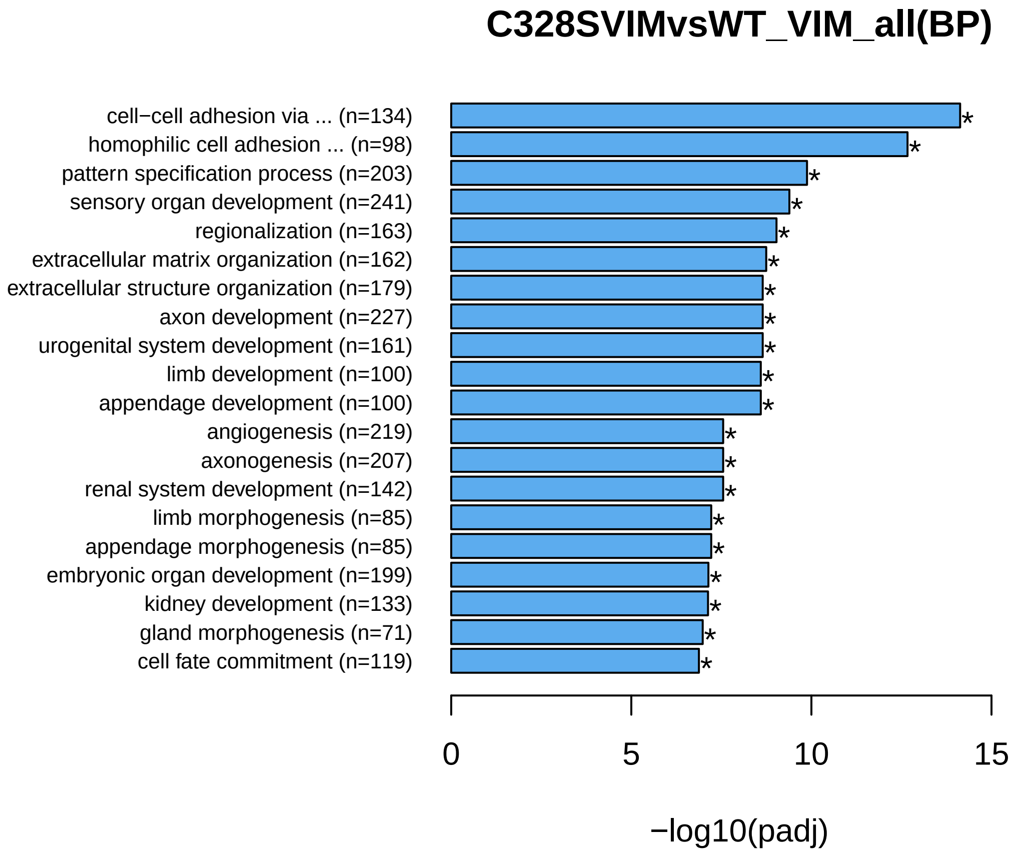

### C328SVIMvsWT_VIM_all.BPbar.png

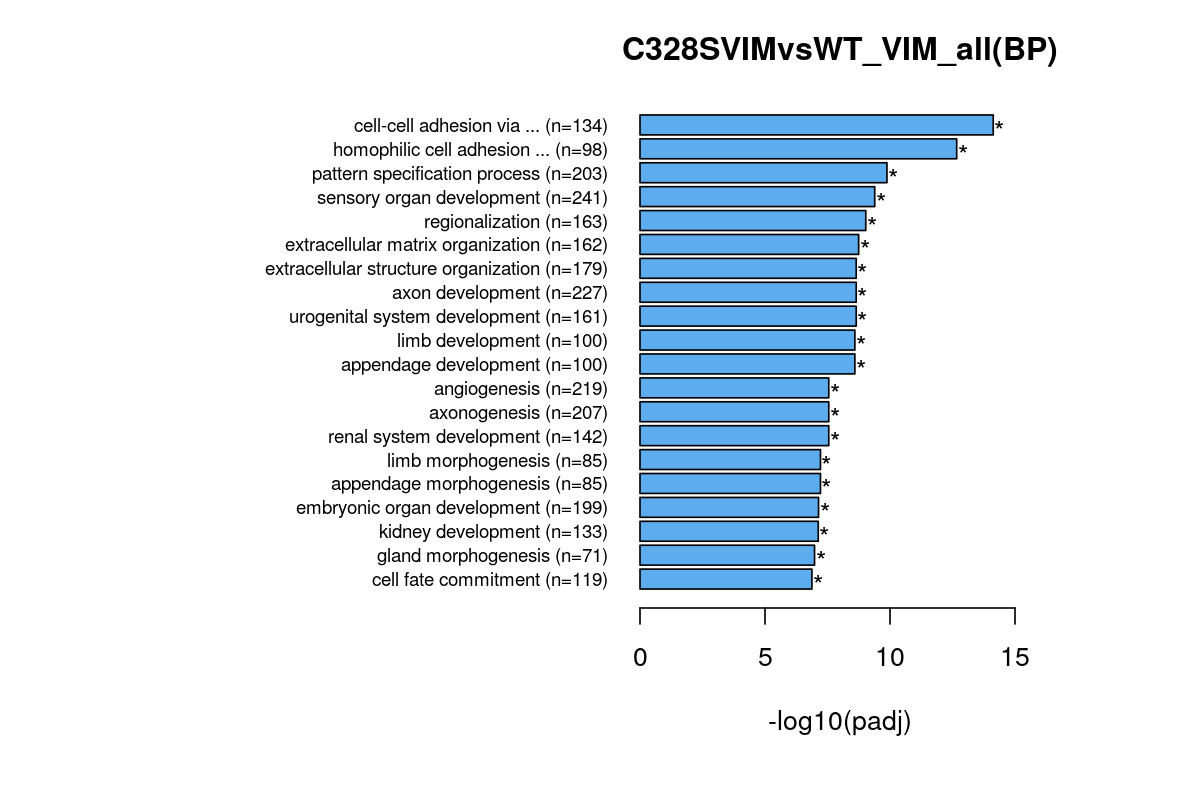

### C328SVIMvsWT_VIM_all.CCbar.pdf

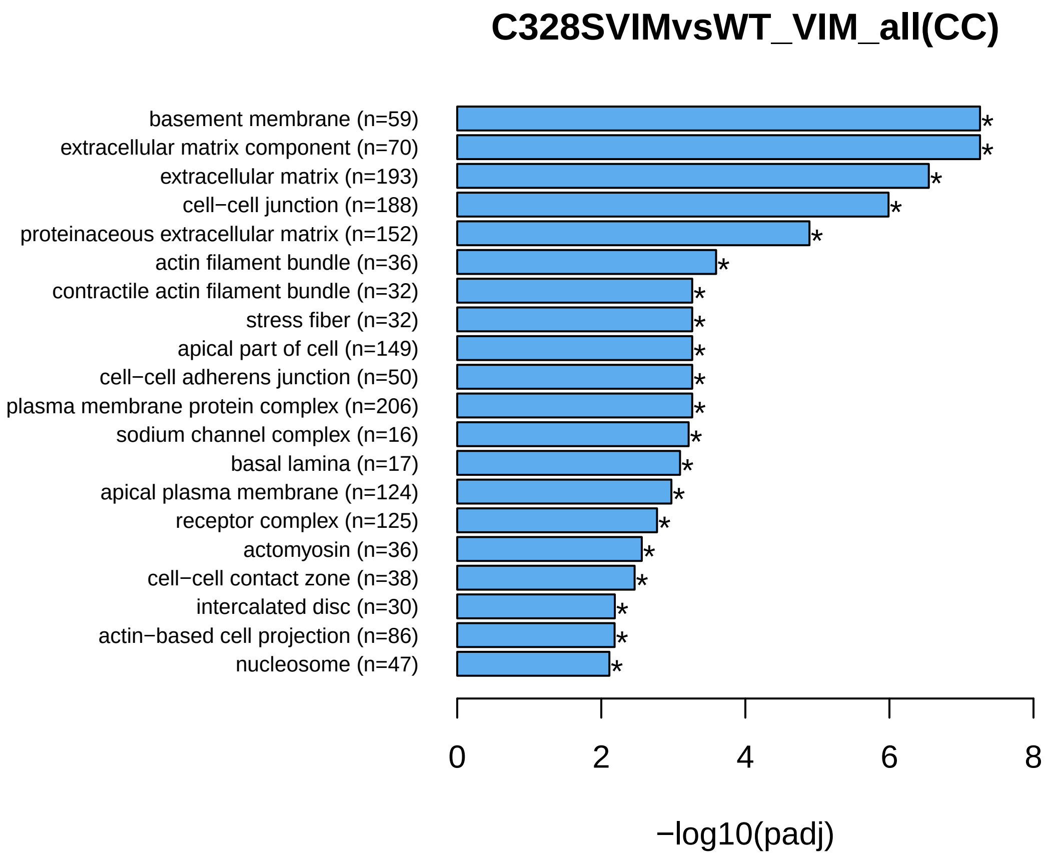

### C328SVIMvsWT_VIM_all.CCbar.png

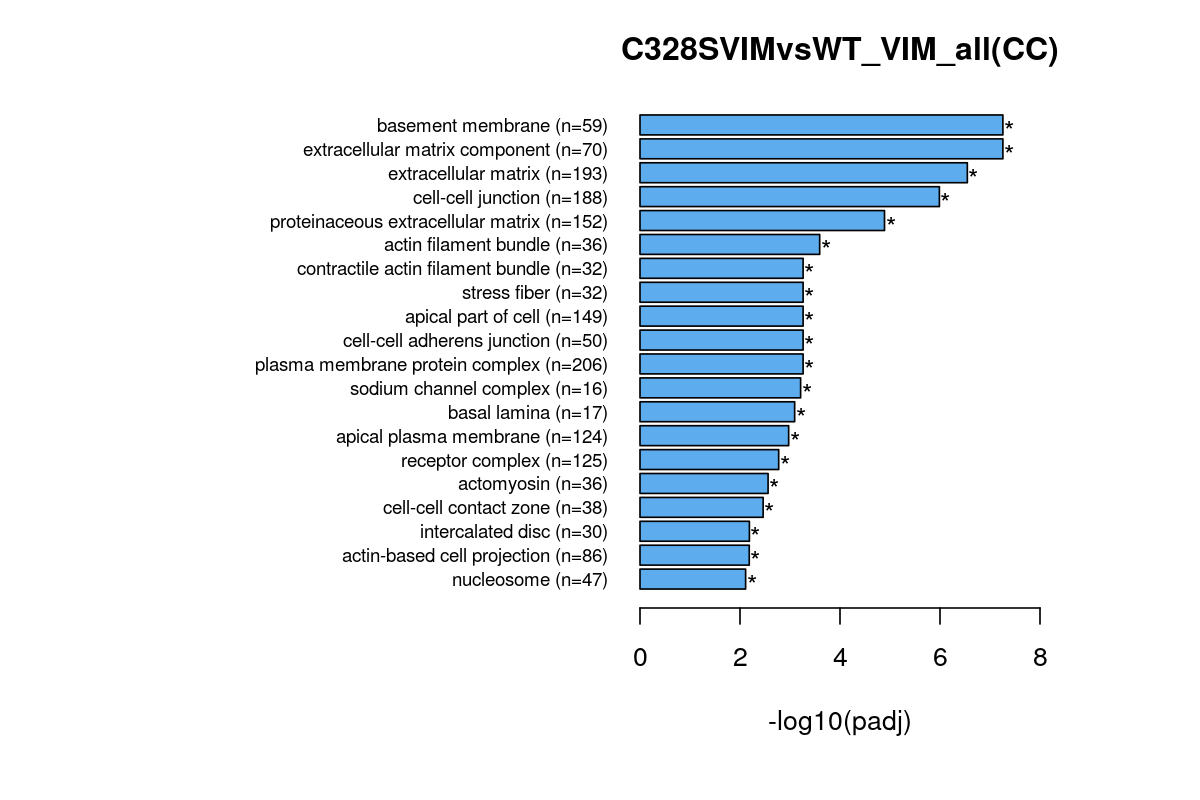

### C328SVIMvsWT_VIM_all.dot.pdf

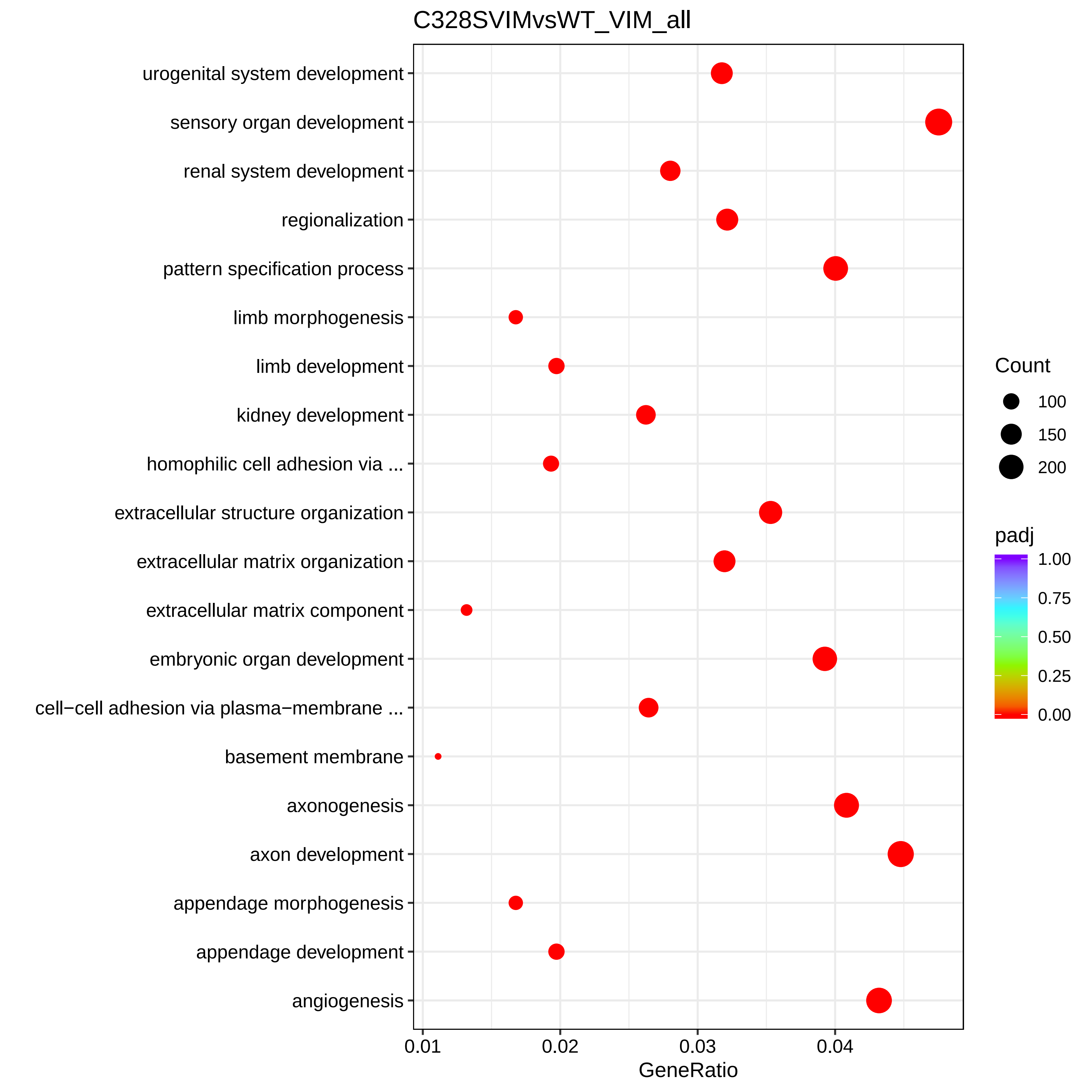

### C328SVIMvsWT_VIM_all.dot.png

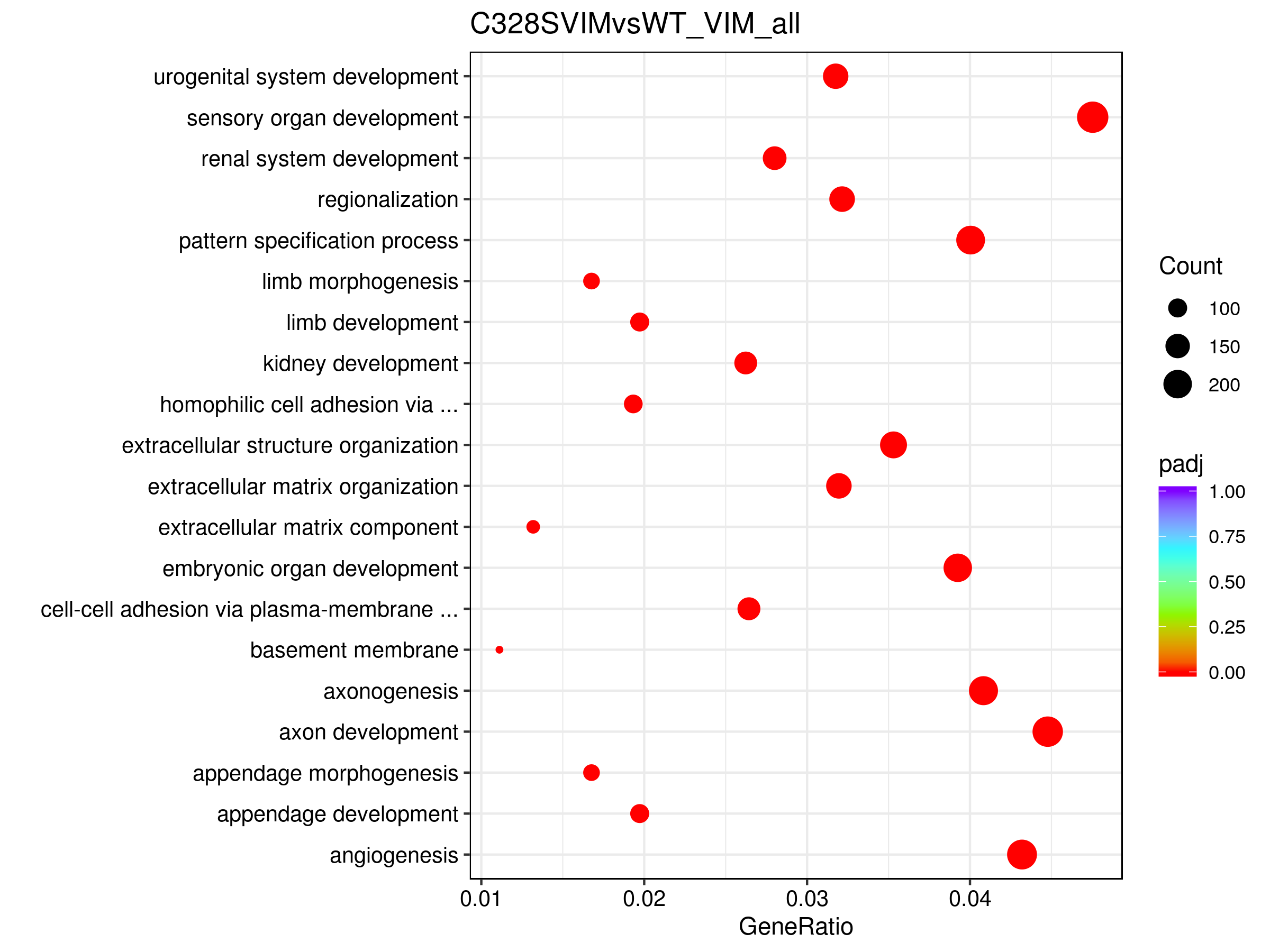

### C328SVIMvsWT_VIM_all.GObp_DAG.pdf

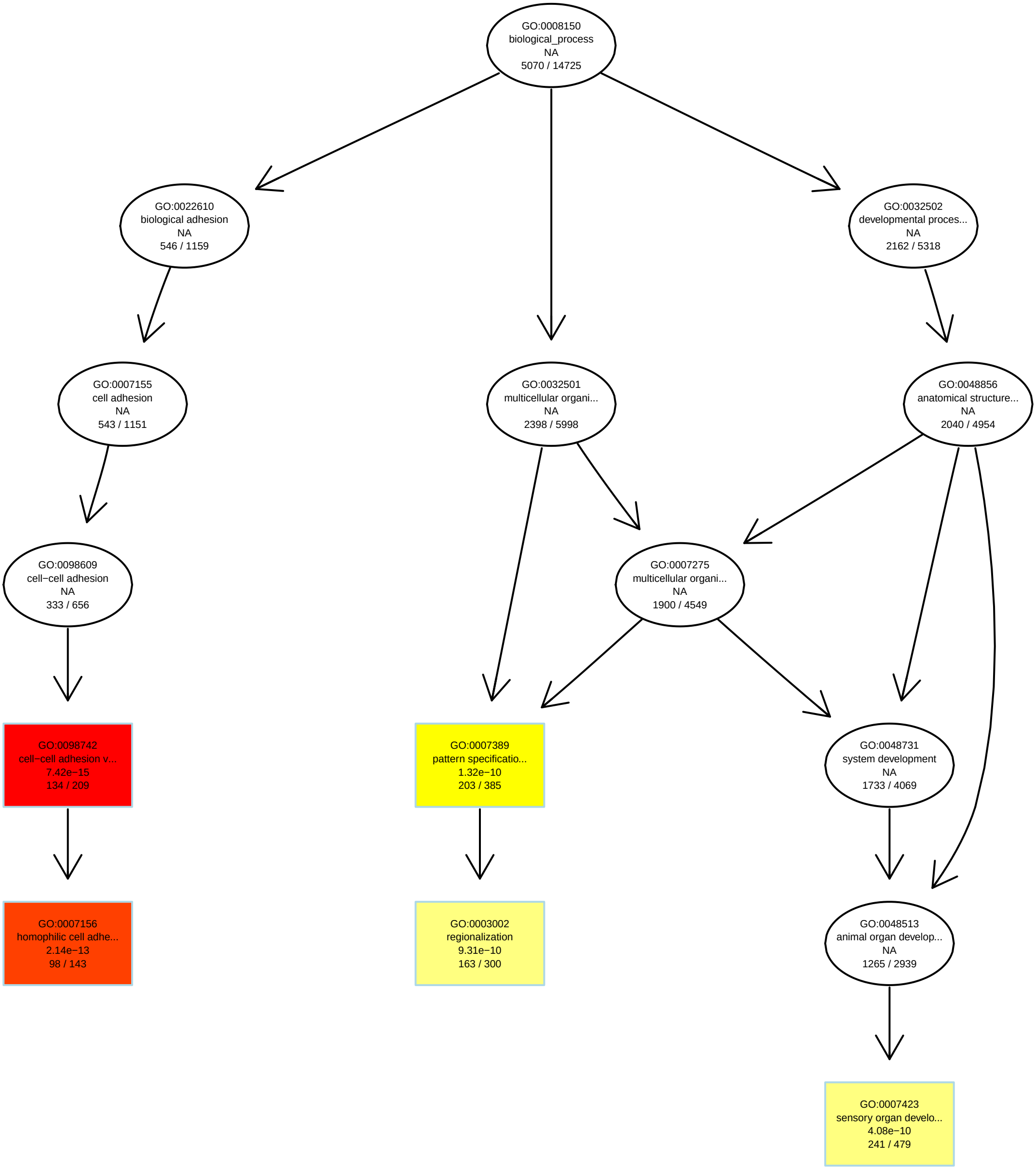

### C328SVIMvsWT_VIM_all.GObp_DAG.png

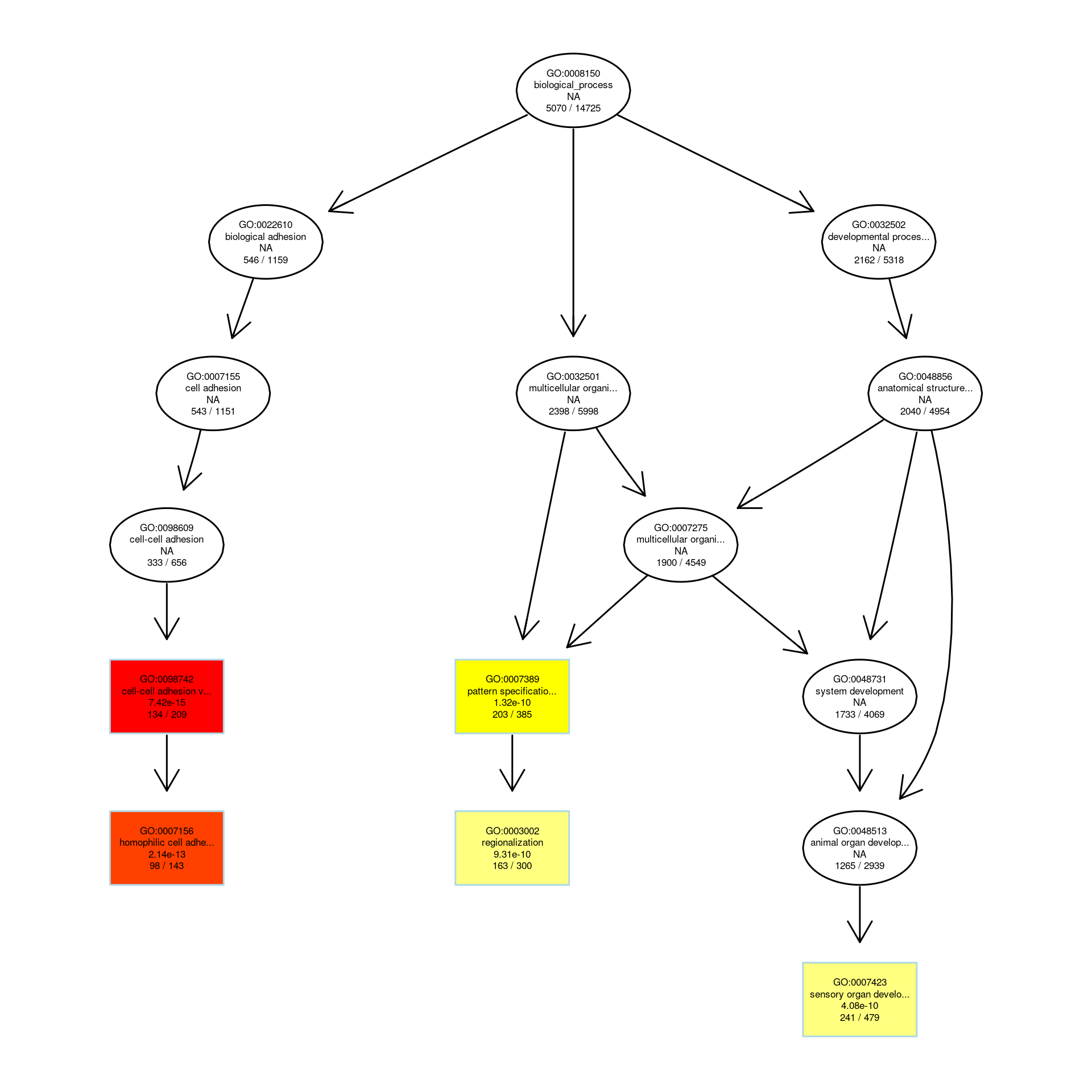

### C328SVIMvsWT_VIM_all.GOmf_DAG.pdf

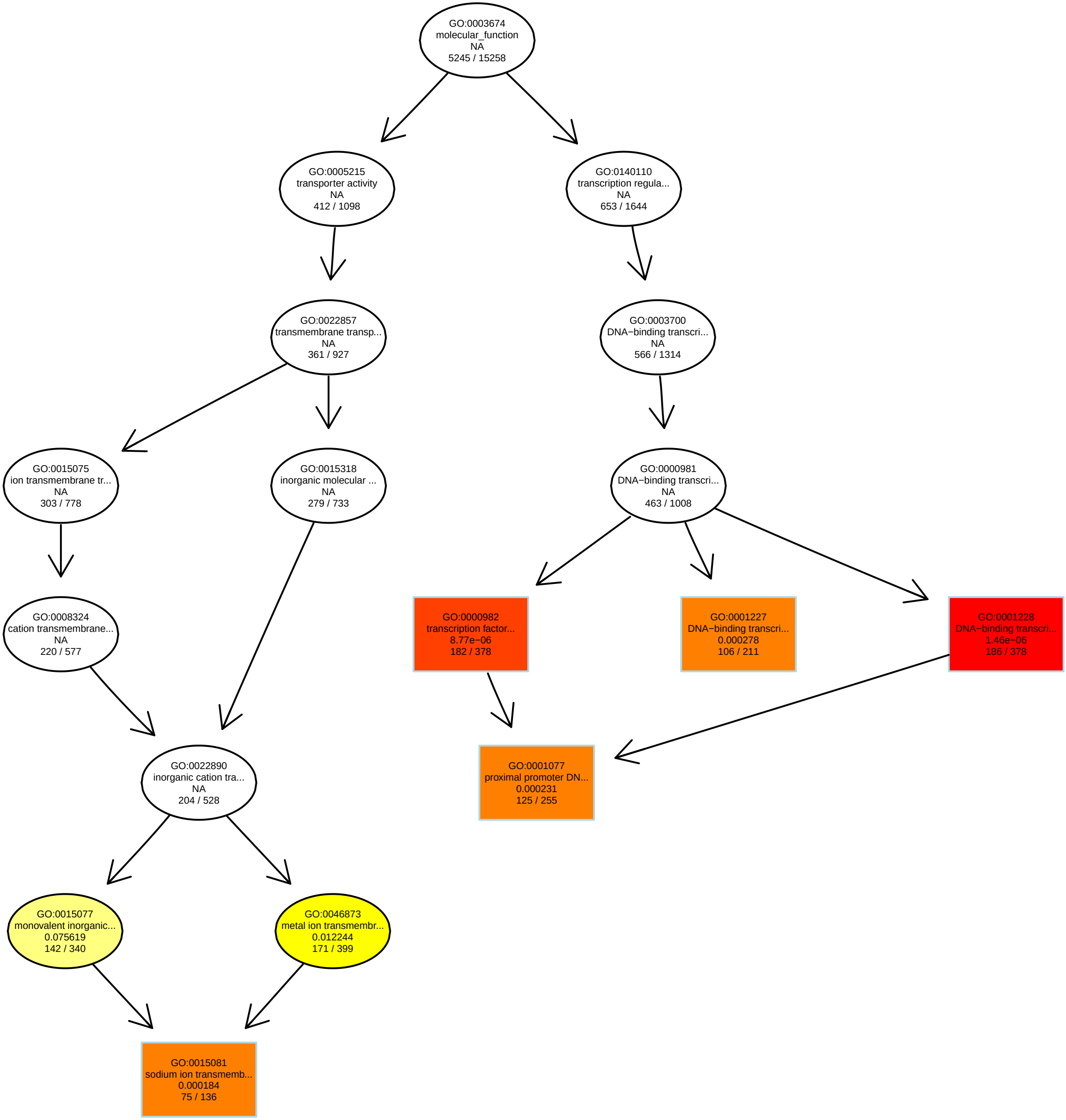

### C328SVIMvsWT_VIM_all.GOmf_DAG.png

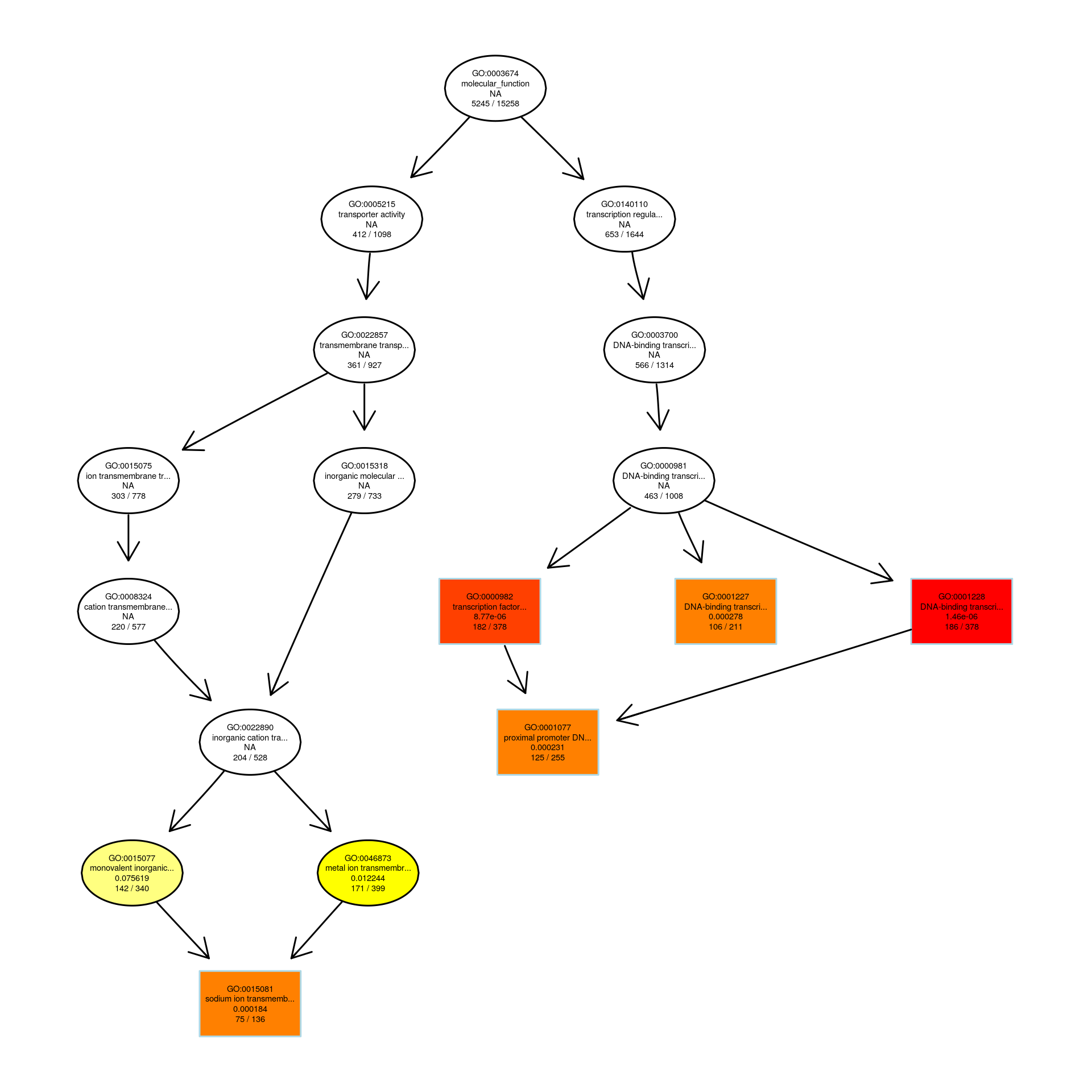

### C328SVIMvsWT_VIM_all.MFbar.pdf

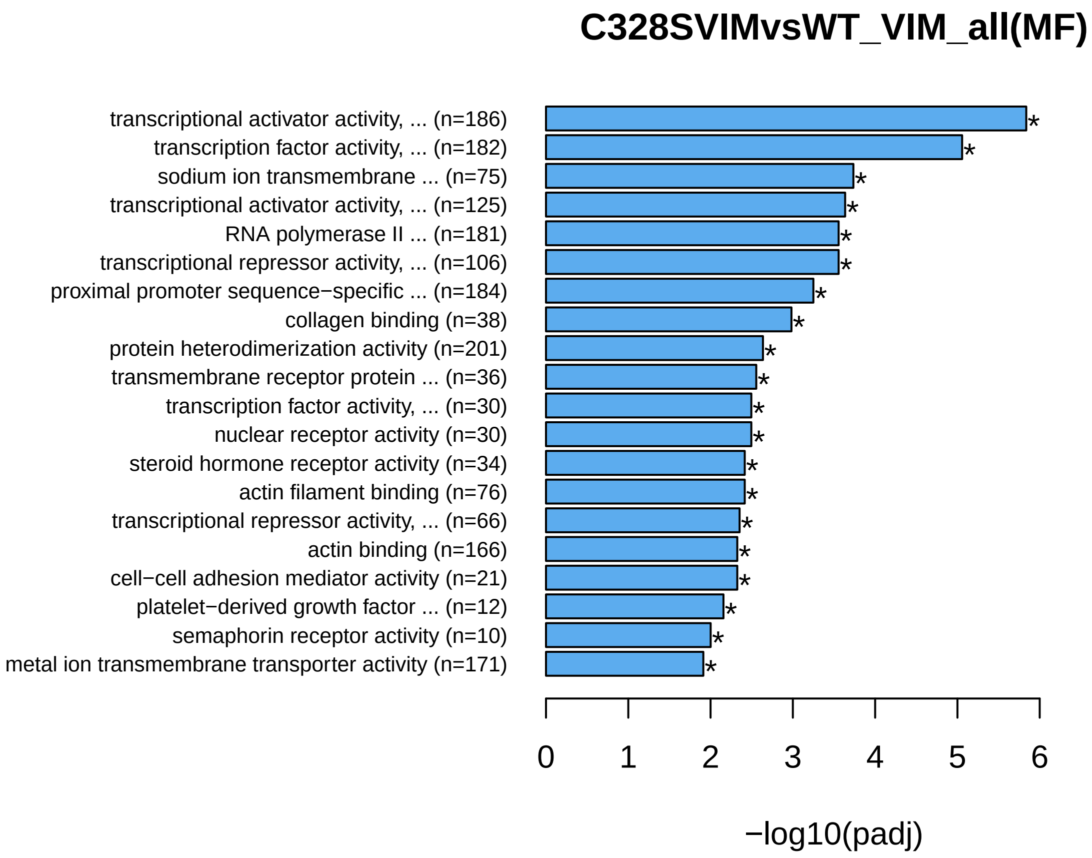

### C328SVIMvsWT_VIM_all.MFbar.png

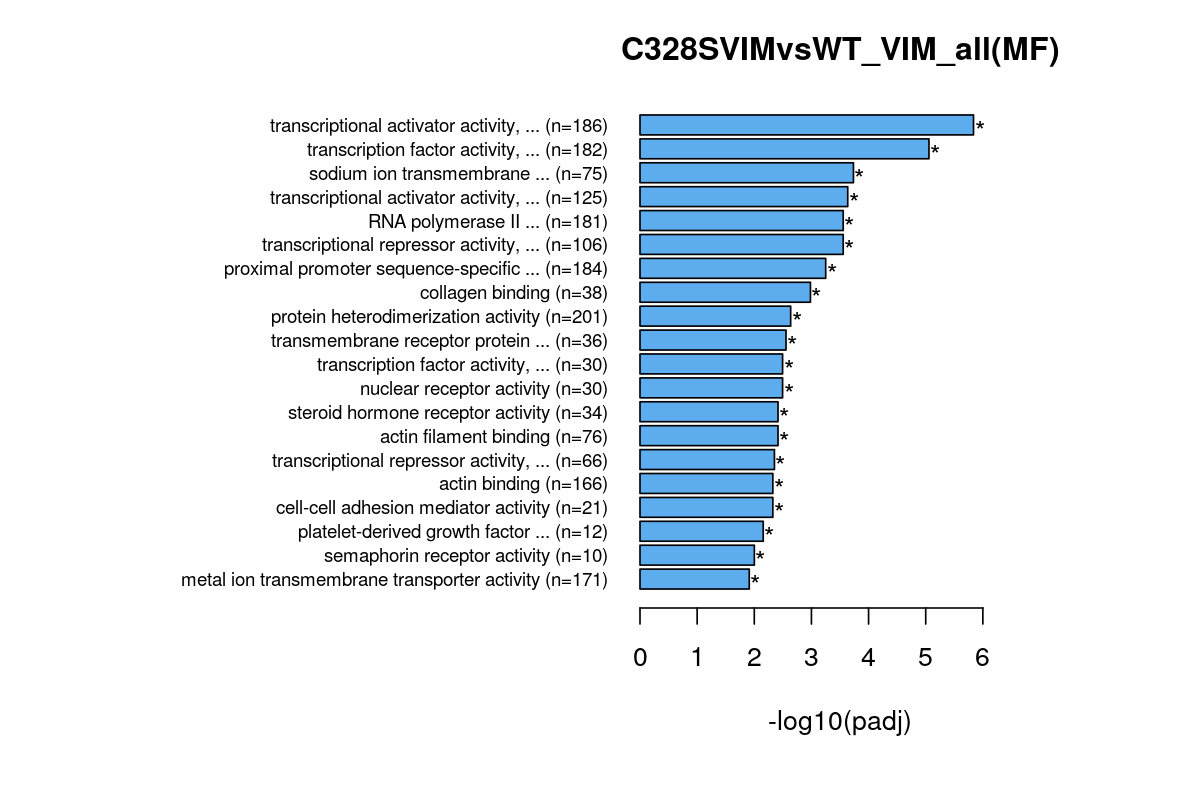

### C328SVIMvsWT_VIM_down.bar.pdf

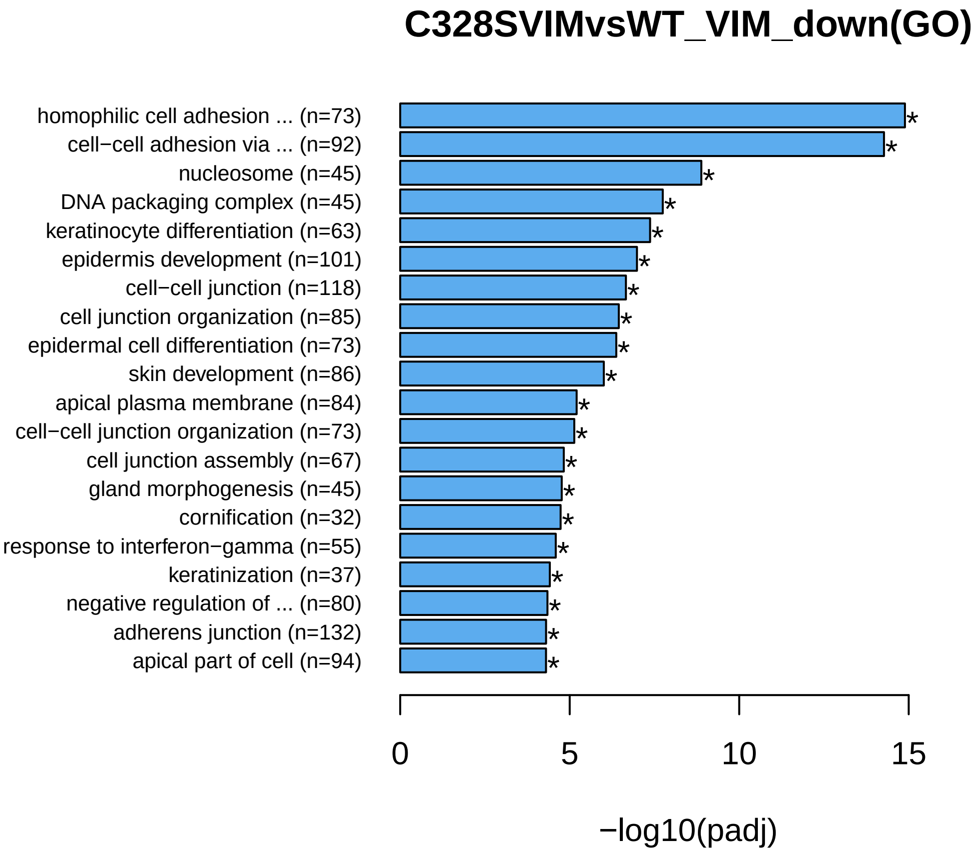

### C328SVIMvsWT_VIM_down.bar.png

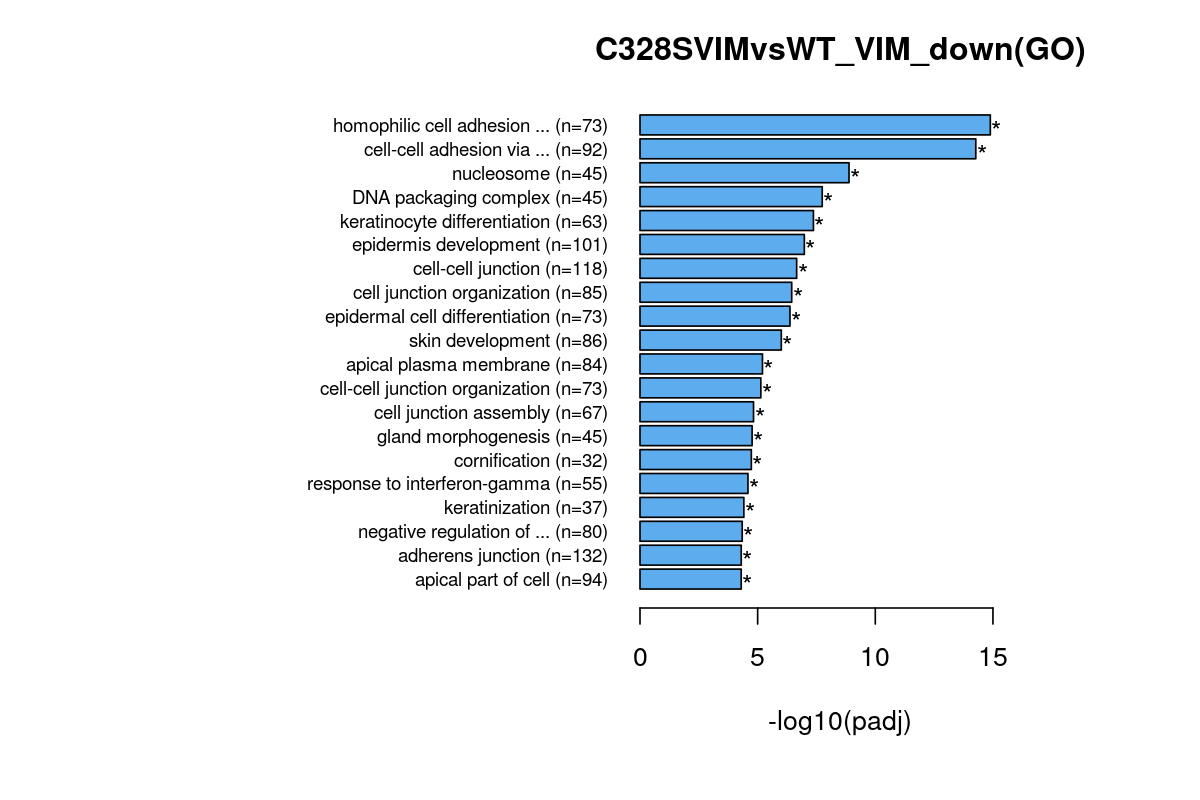

### C328SVIMvsWT_VIM_down.BPbar.pdf

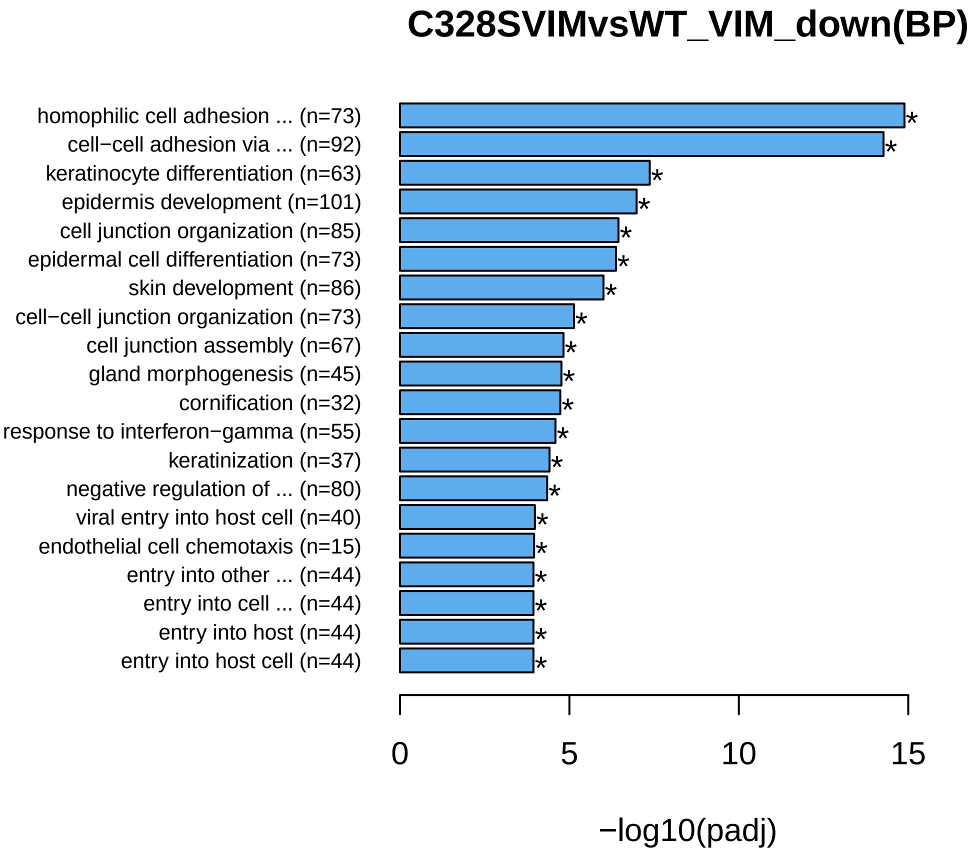

### C328SVIMvsWT_VIM_down.BPbar.png

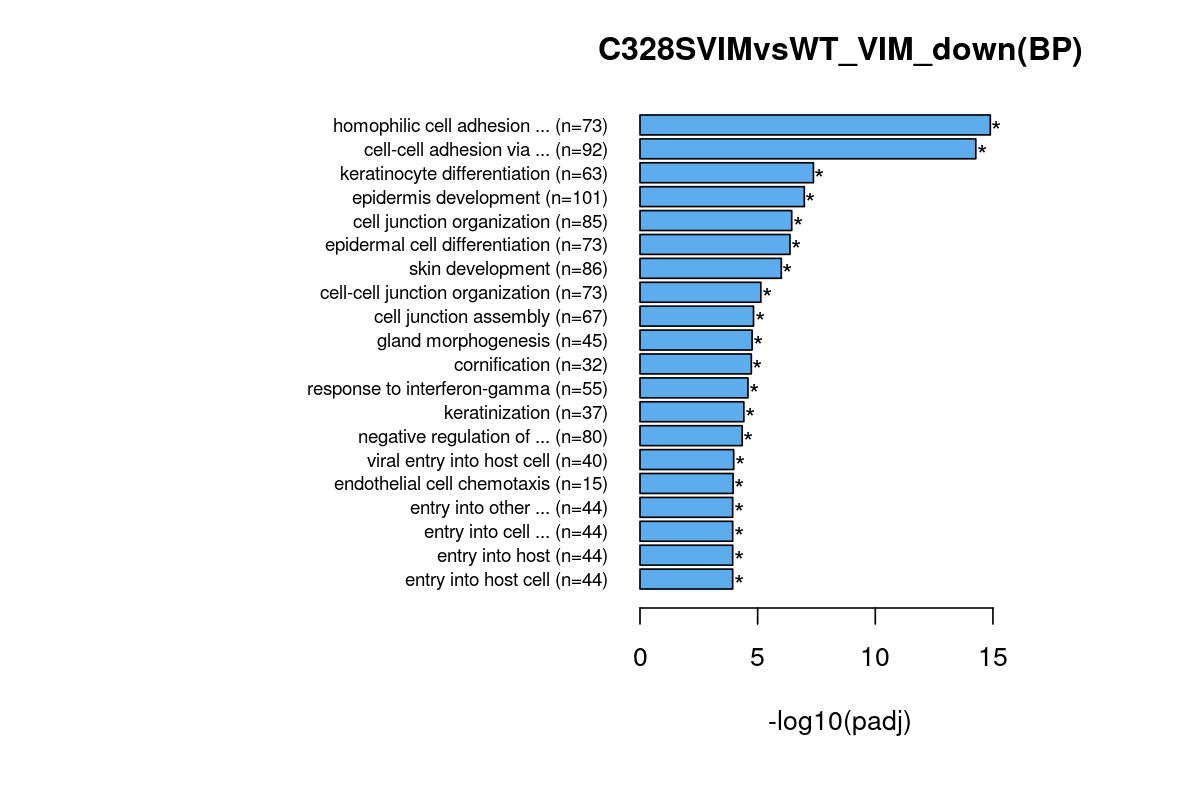

### C328SVIMvsWT_VIM_down.CCbar.pdf

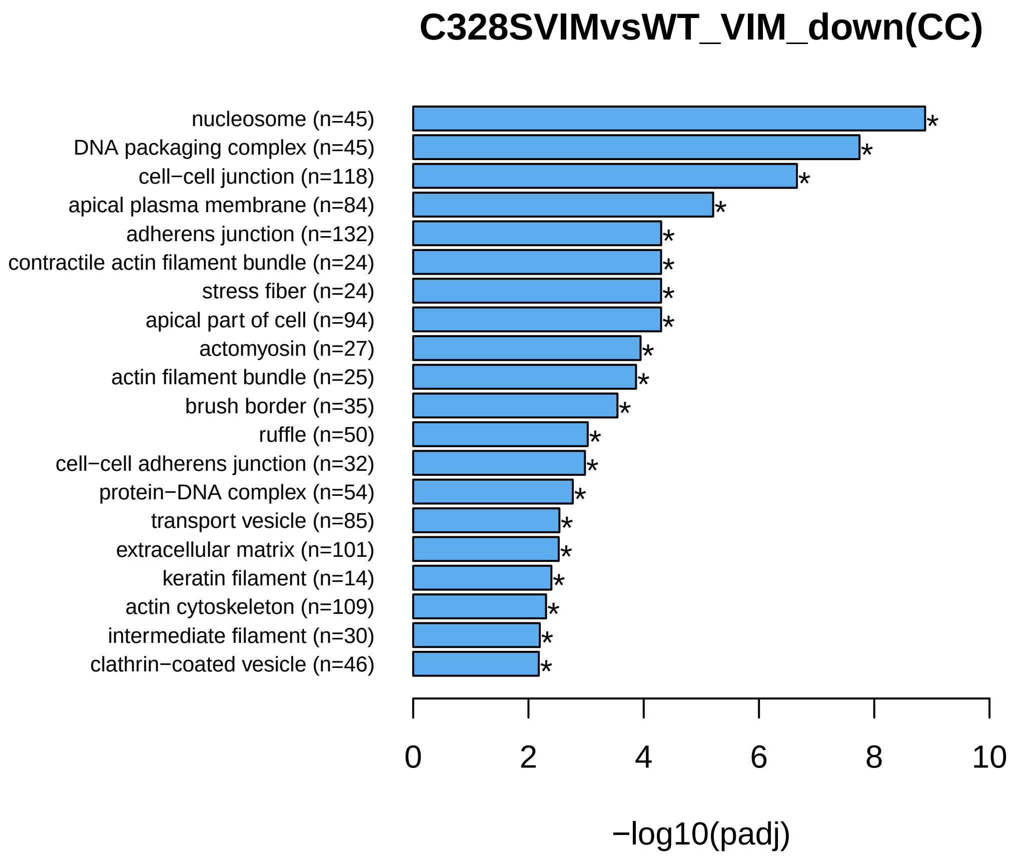

### C328SVIMvsWT_VIM_down.CCbar.png

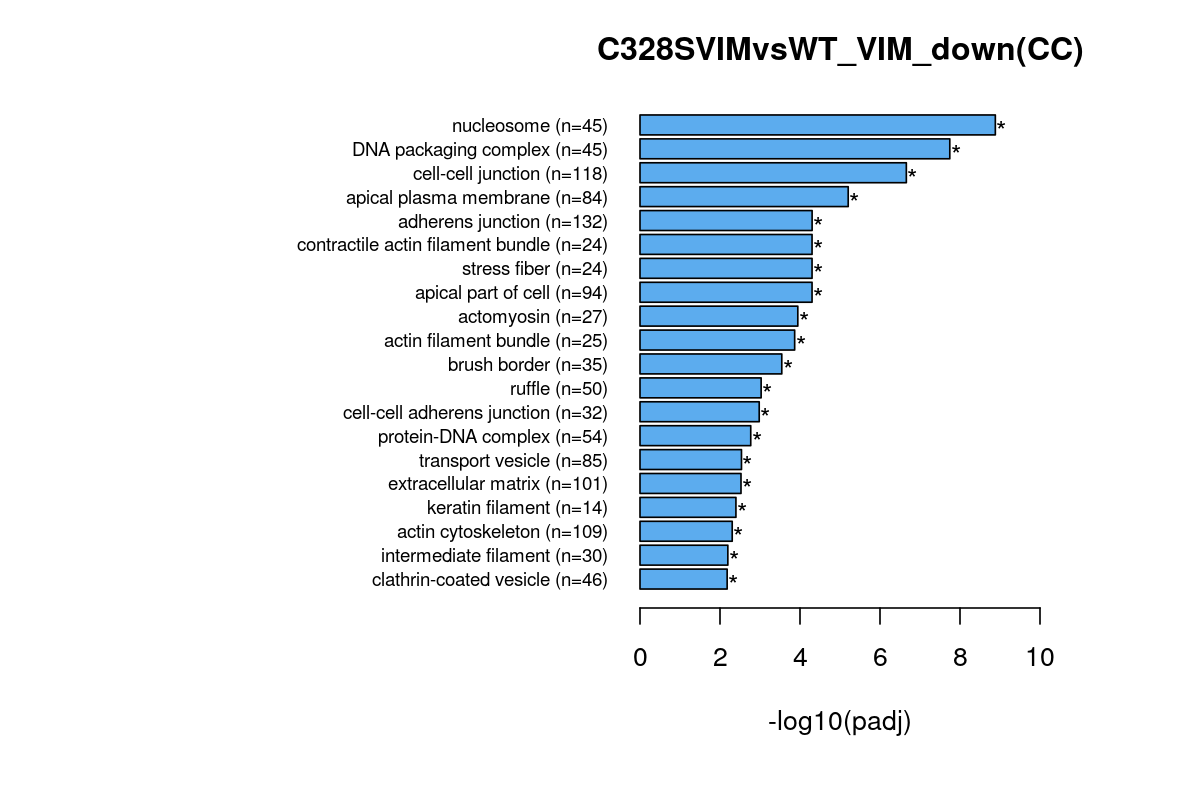

### C328SVIMvsWT_VIM_down.dot.pdf

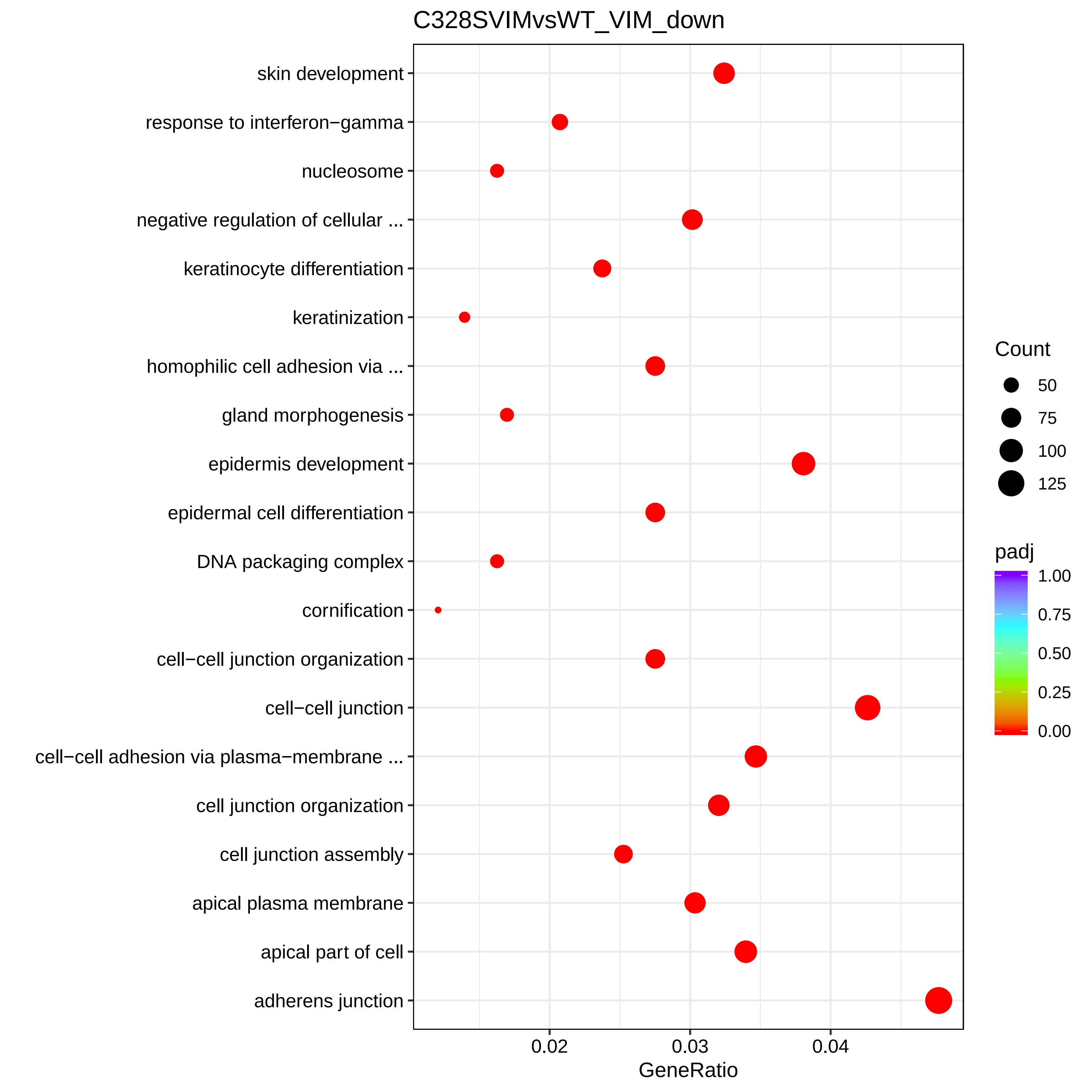

### C328SVIMvsWT_VIM_down.dot.png

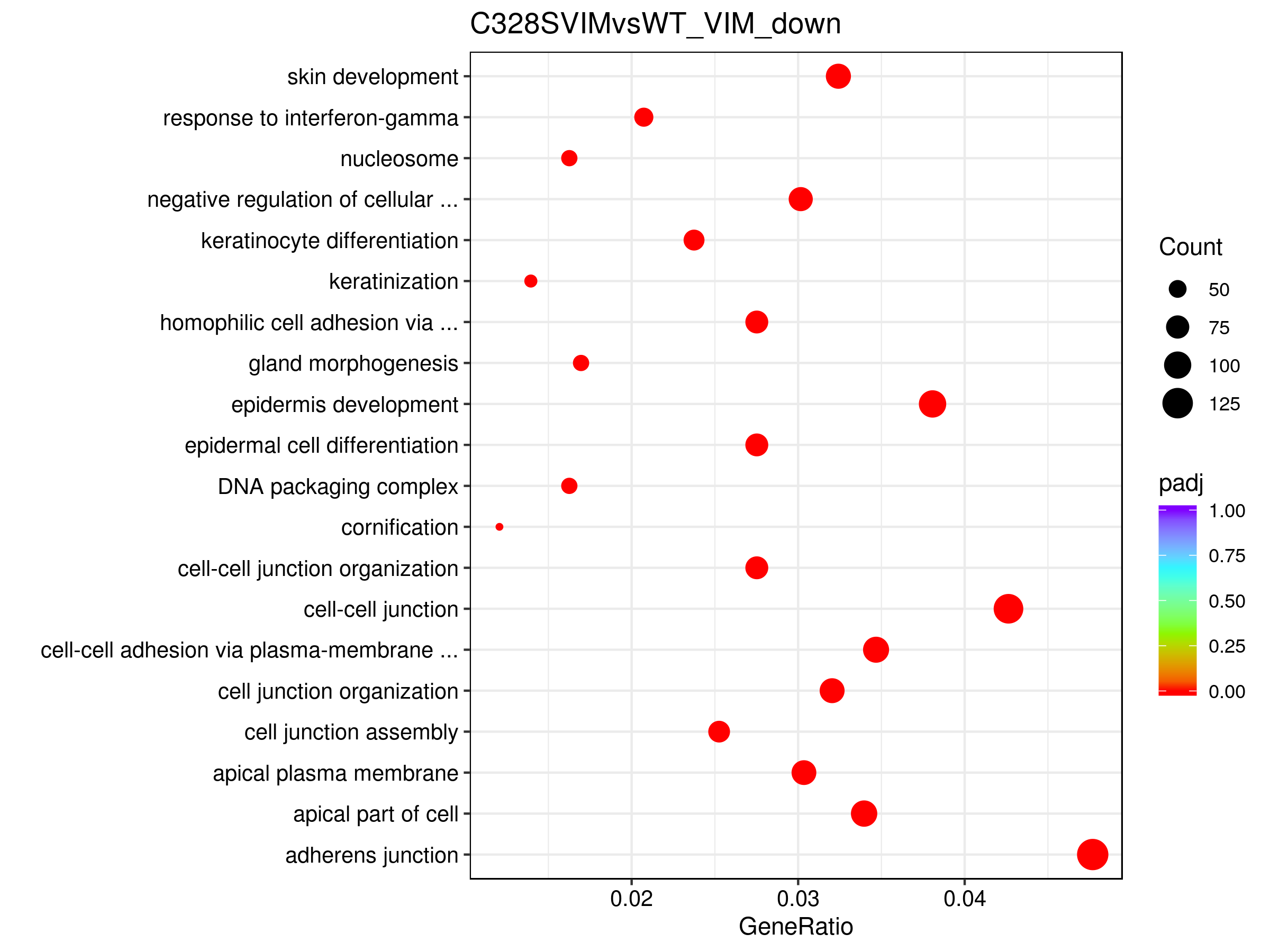

### C328SVIMvsWT_VIM_down.GObp_DAG.pdf

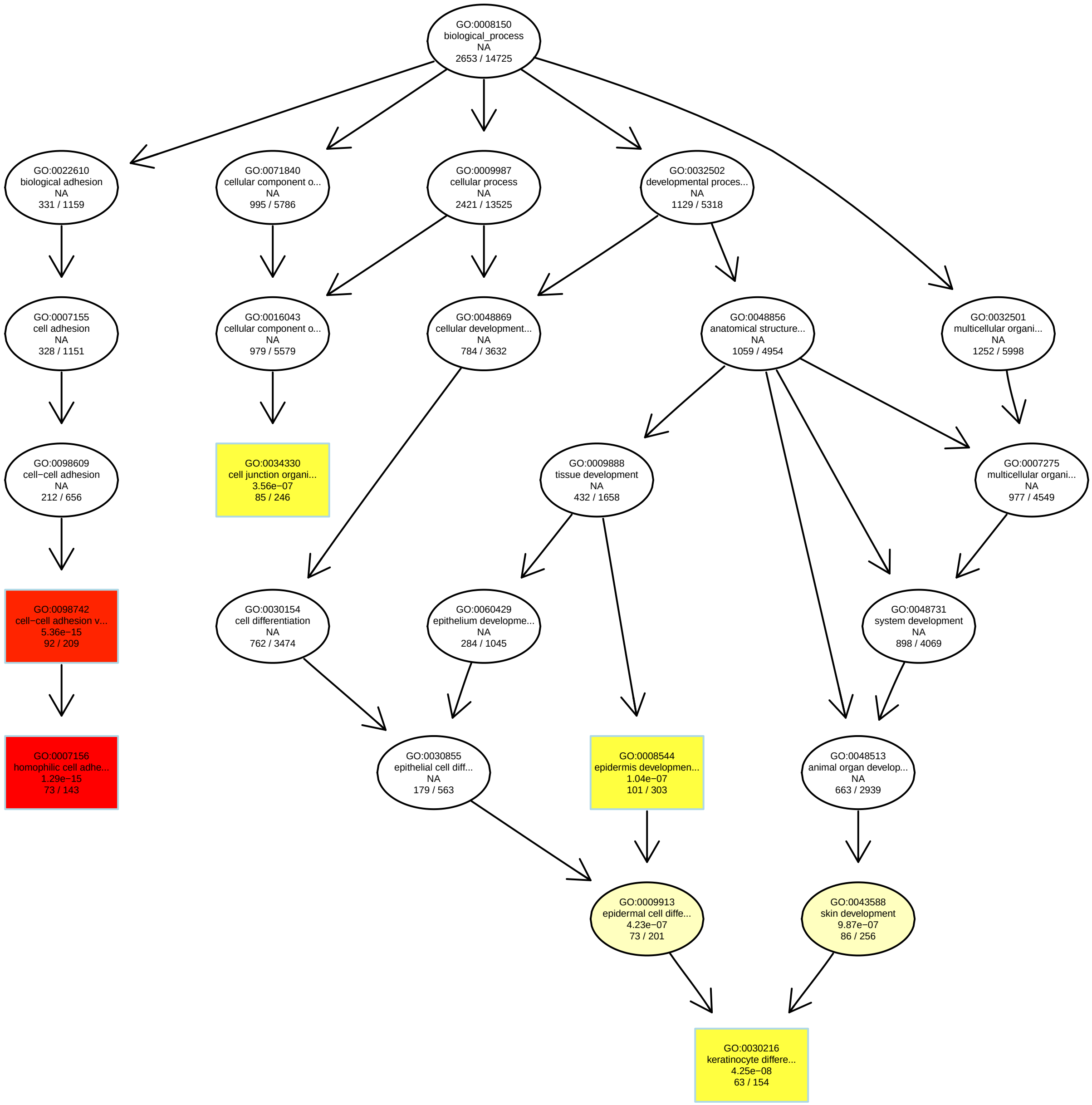

### C328SVIMvsWT_VIM_down.GObp_DAG.png

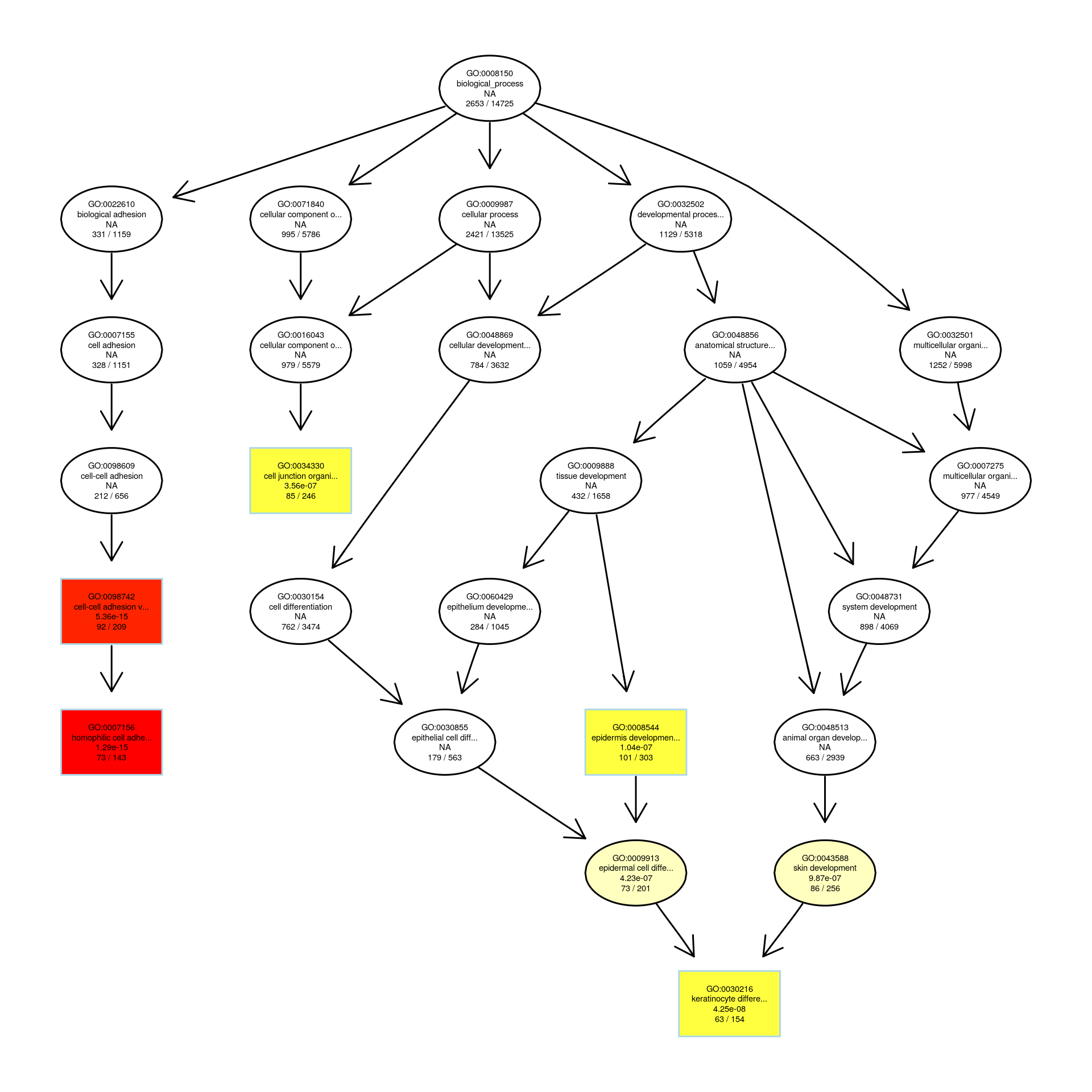

### C328SVIMvsWT_VIM_down.GOcc_DAG.pdf

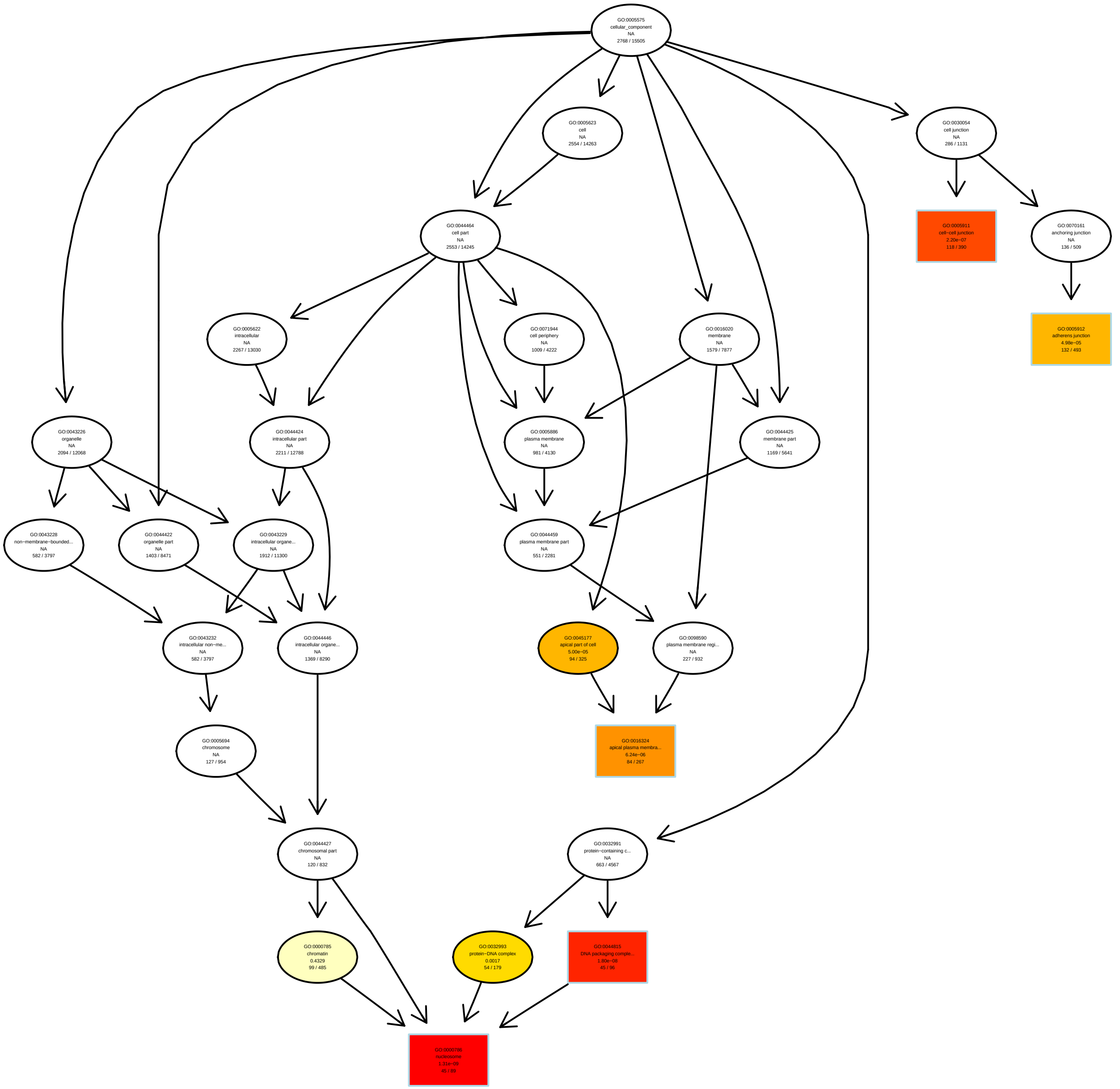

### C328SVIMvsWT_VIM_down.GOcc_DAG.png

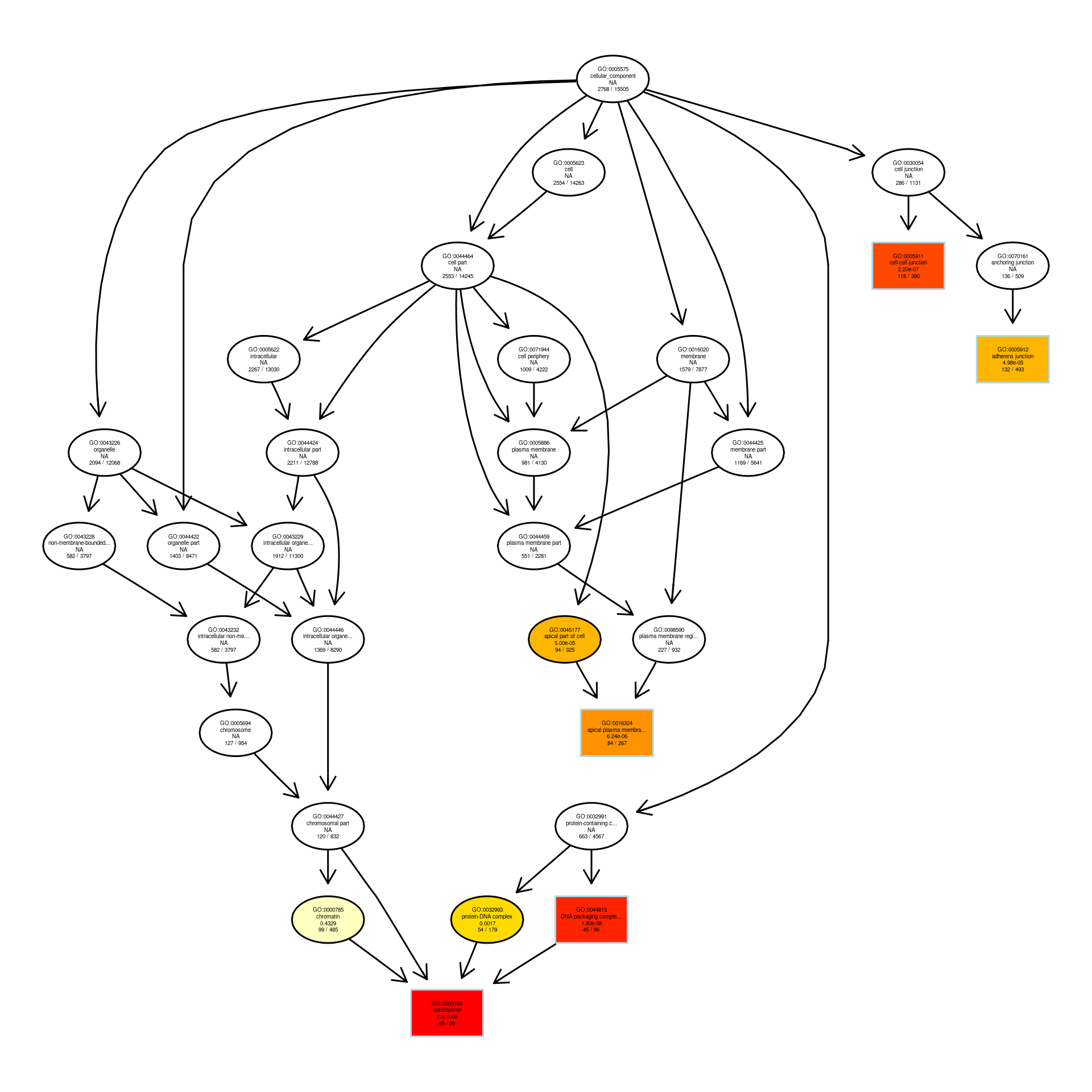

### C328SVIMvsWT_VIM_down.GOmf_DAG.pdf

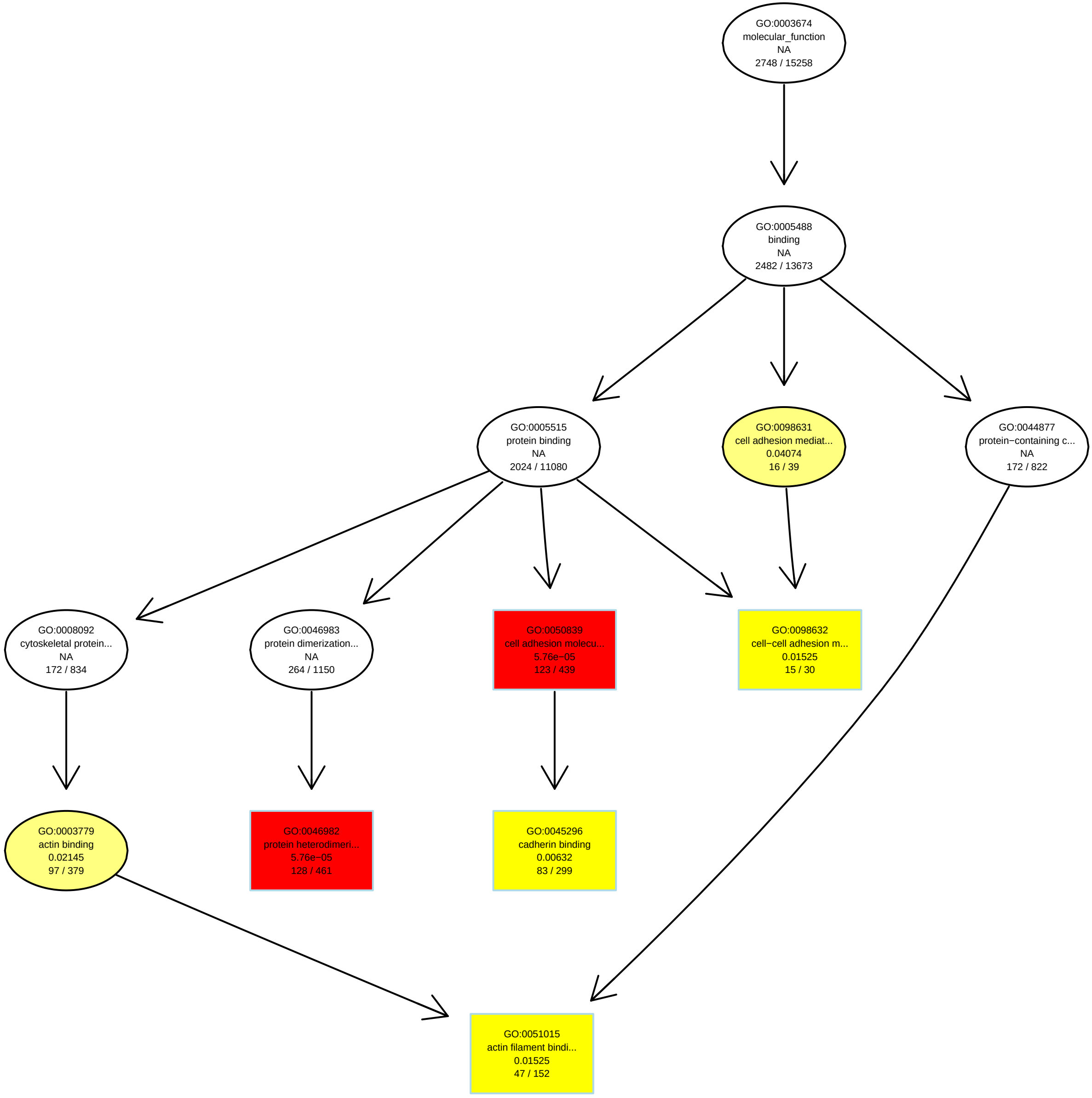

### C328SVIMvsWT_VIM_down.GOmf_DAG.png

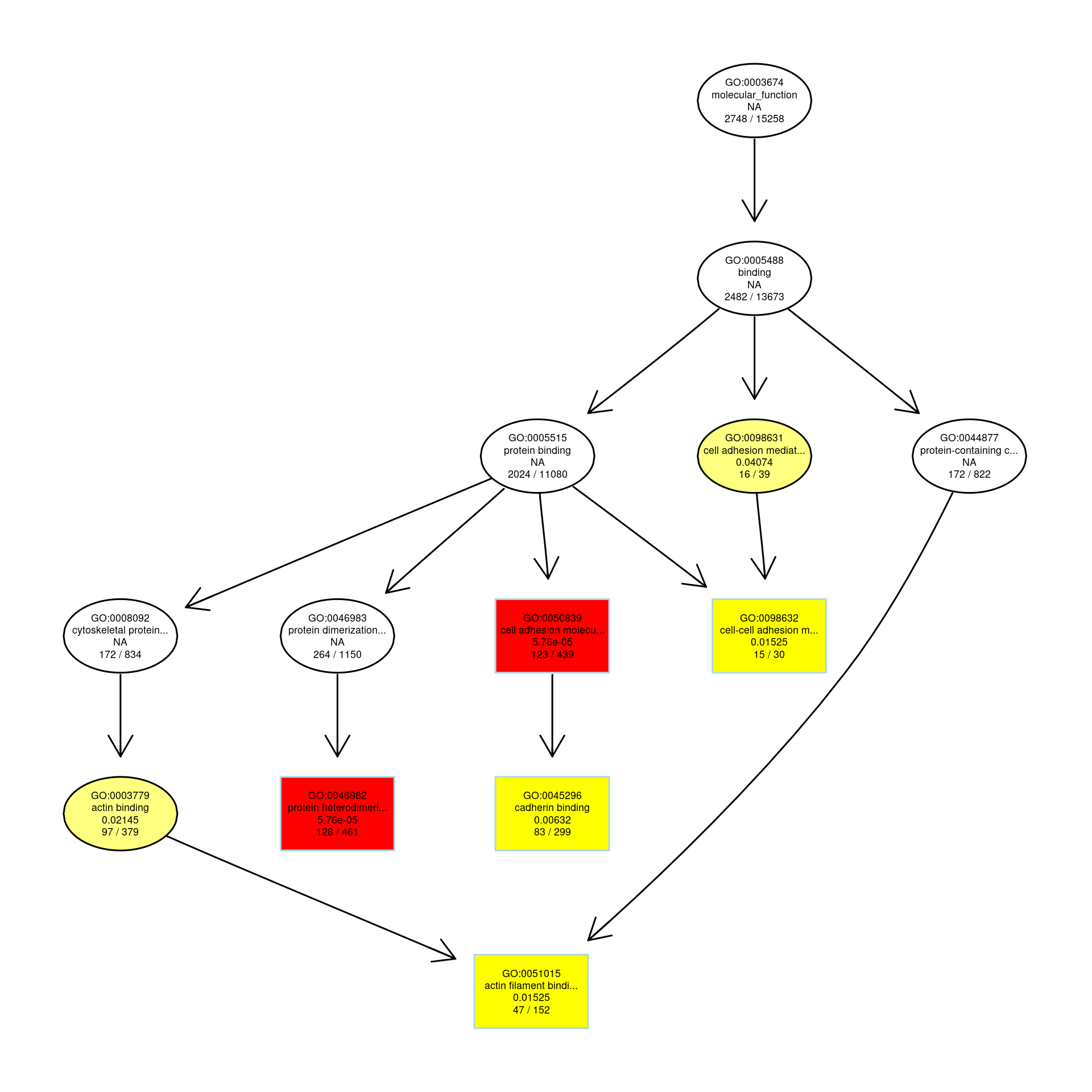

### C328SVIMvsWT_VIM_down.MFbar.pdf

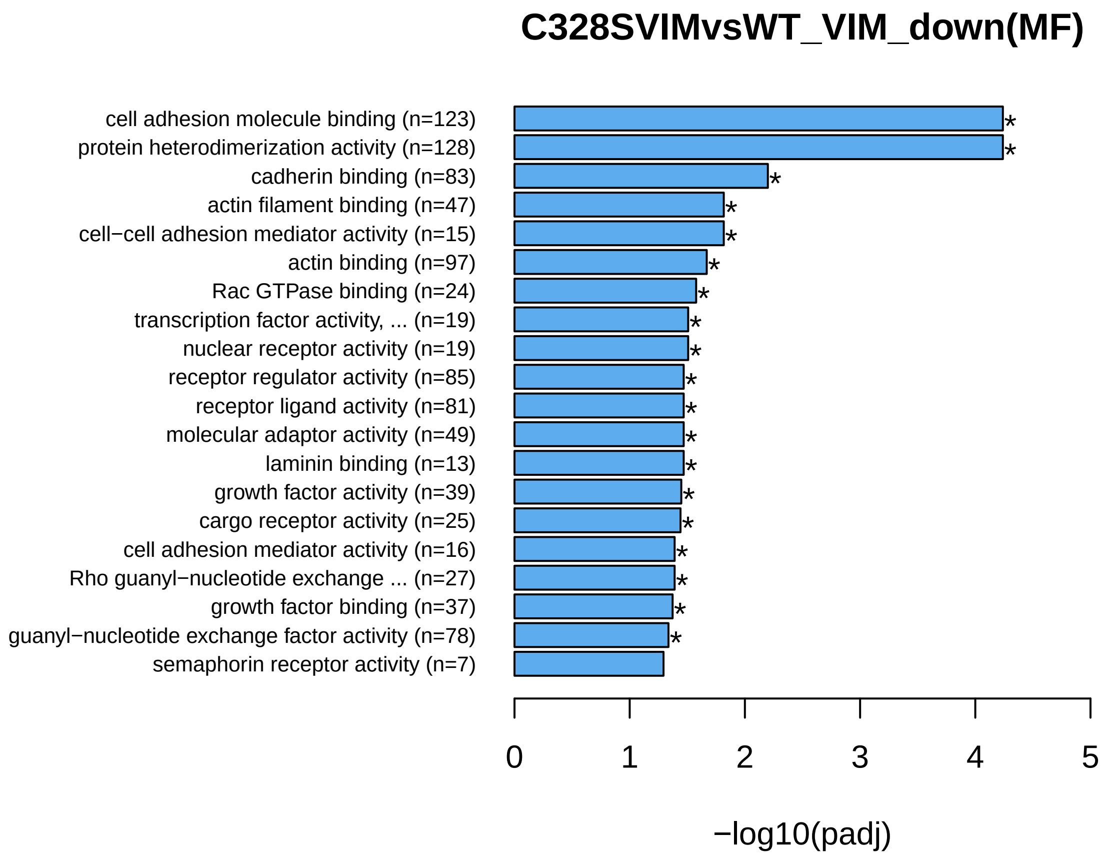

### C328SVIMvsWT_VIM_down.MFbar.png

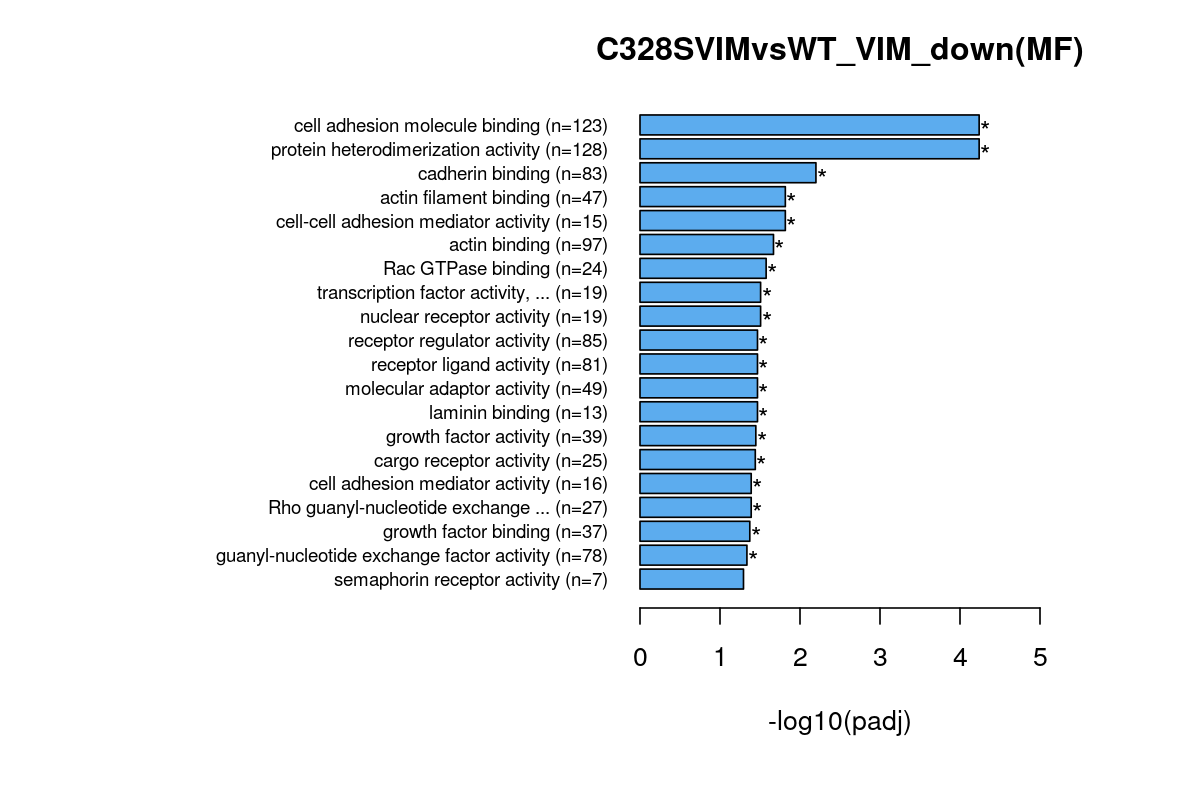

### C328SVIMvsWT_VIM_up.bar.pdf

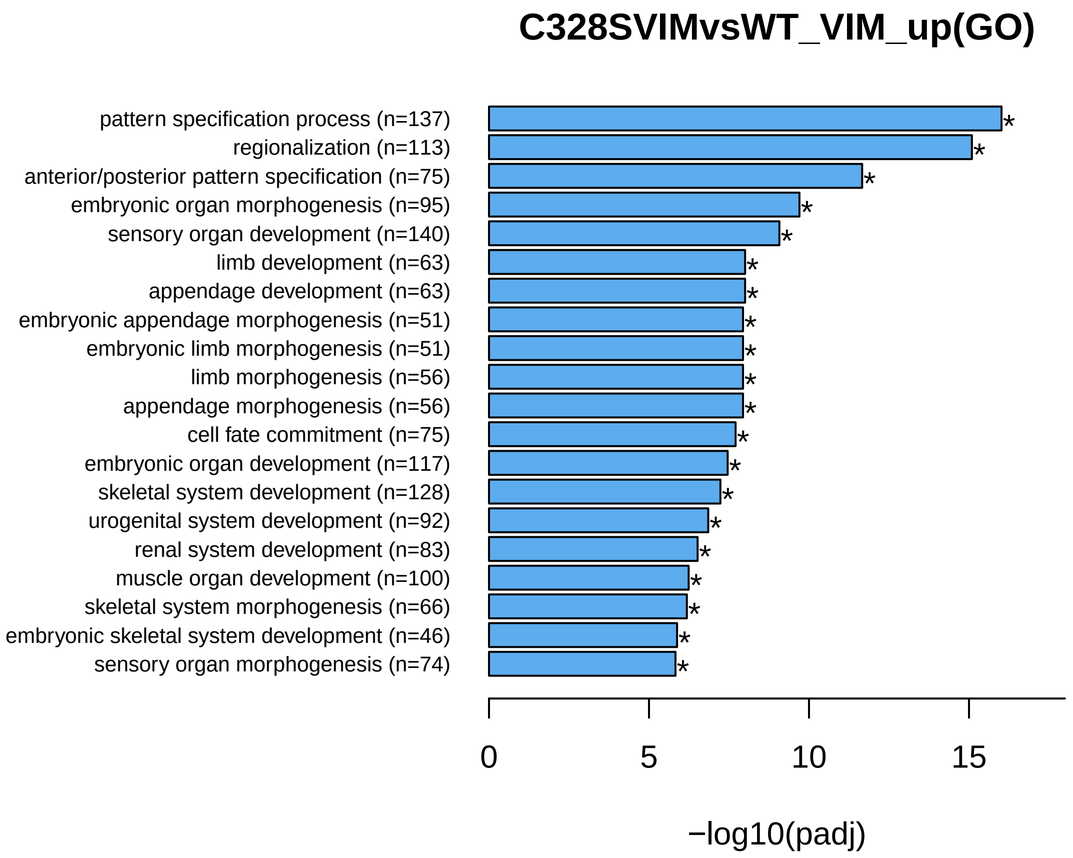

### C328SVIMvsWT_VIM_up.bar.png

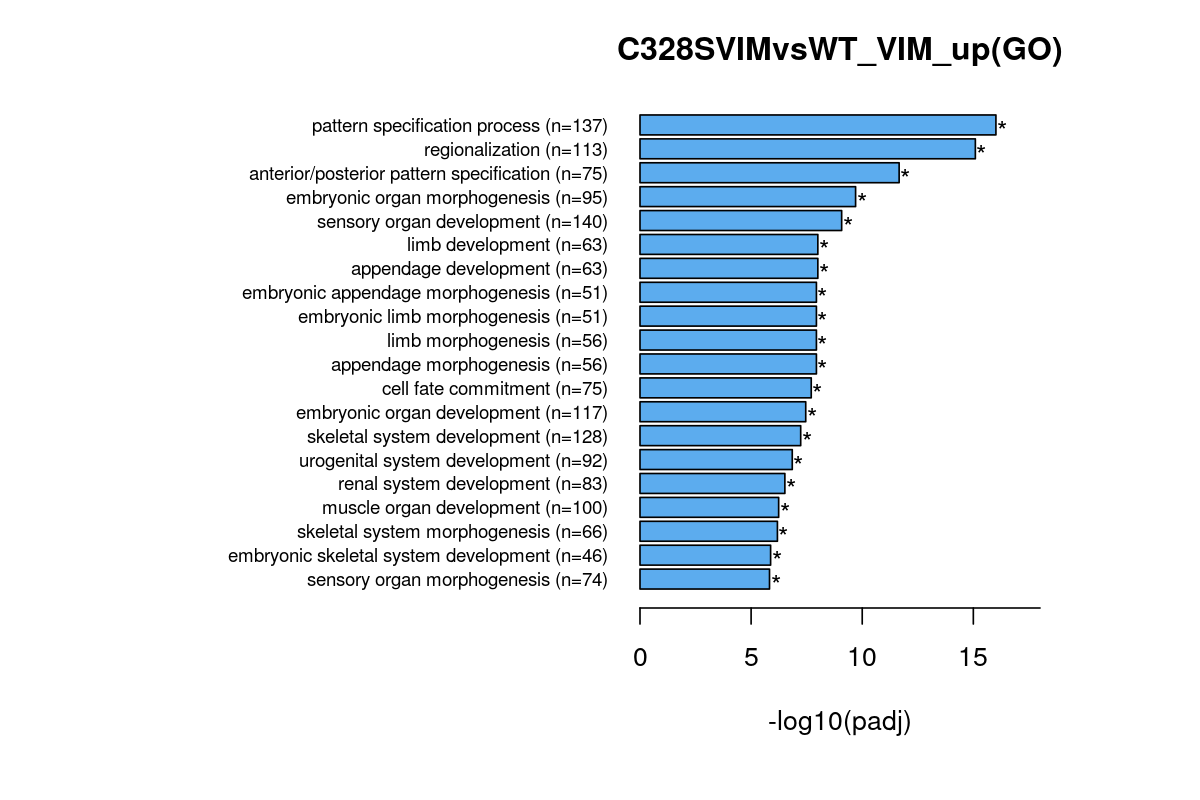

### C328SVIMvsWT_VIM_up.BPbar.pdf

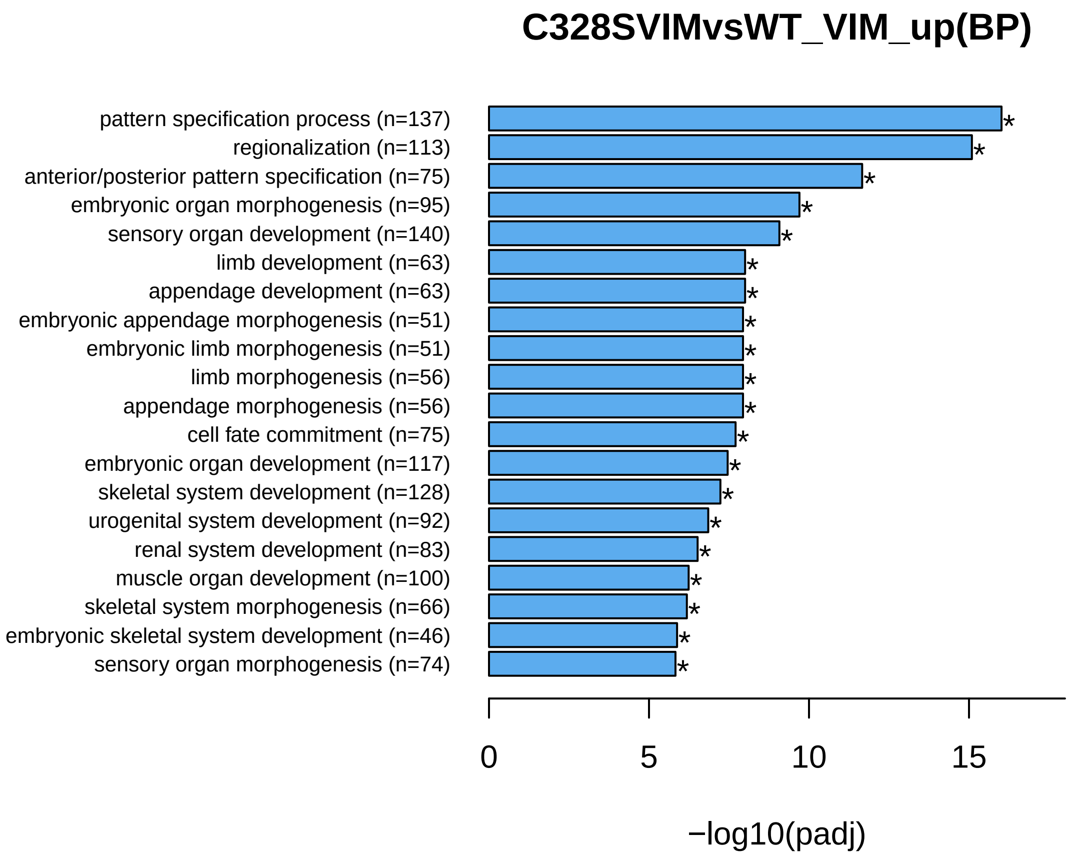

### C328SVIMvsWT_VIM_up.BPbar.png

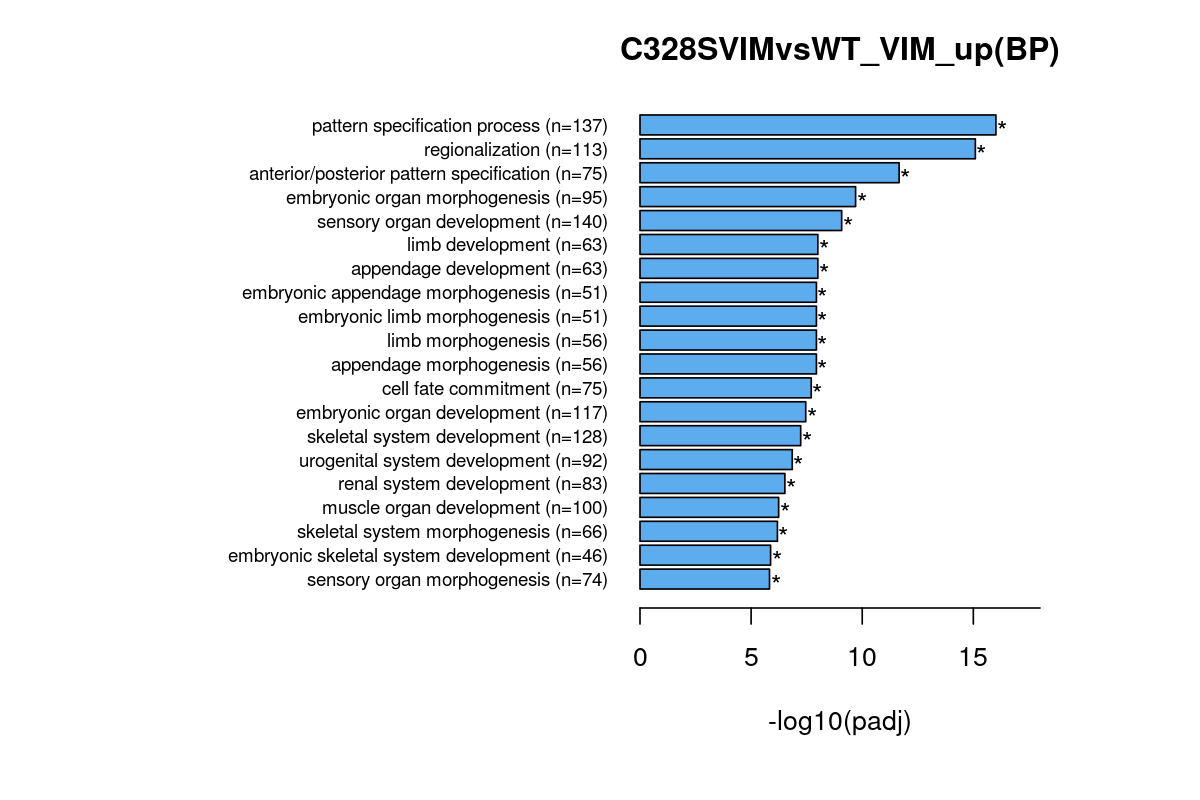

### C328SVIMvsWT_VIM_up.CCbar.pdf

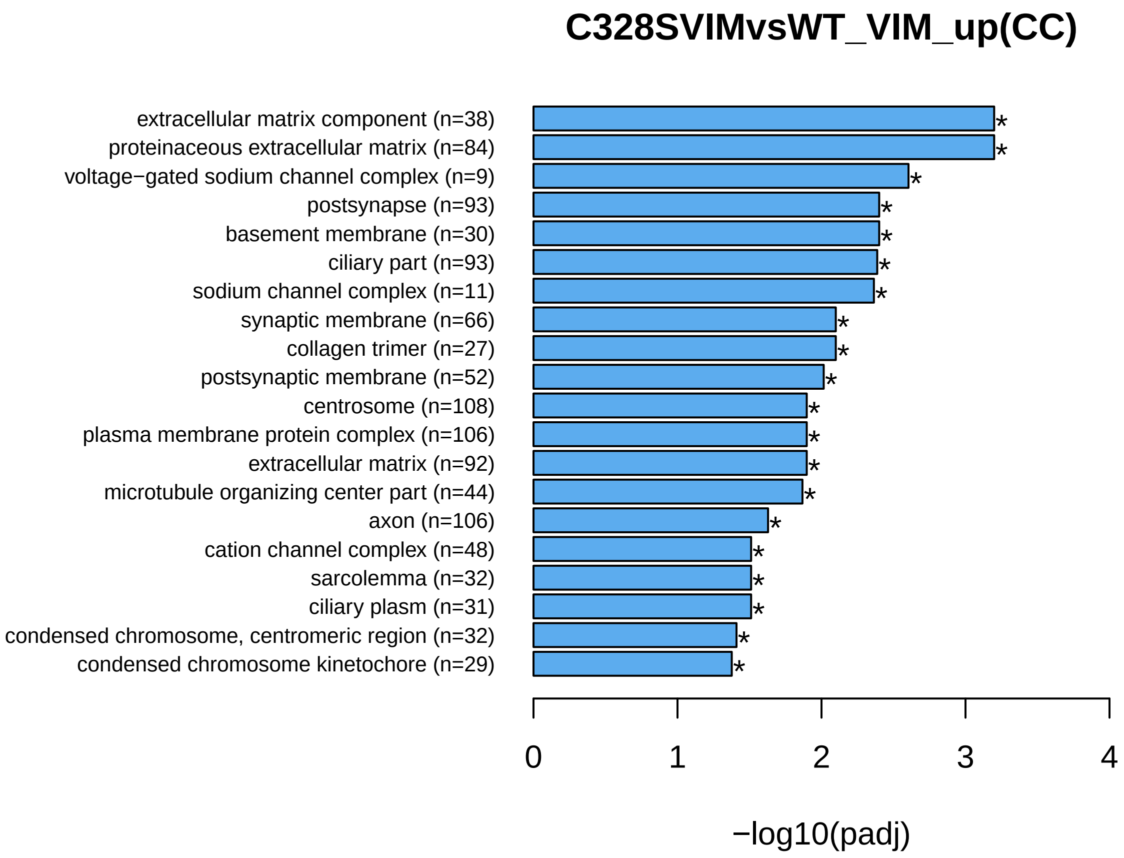

### C328SVIMvsWT_VIM_up.CCbar.png

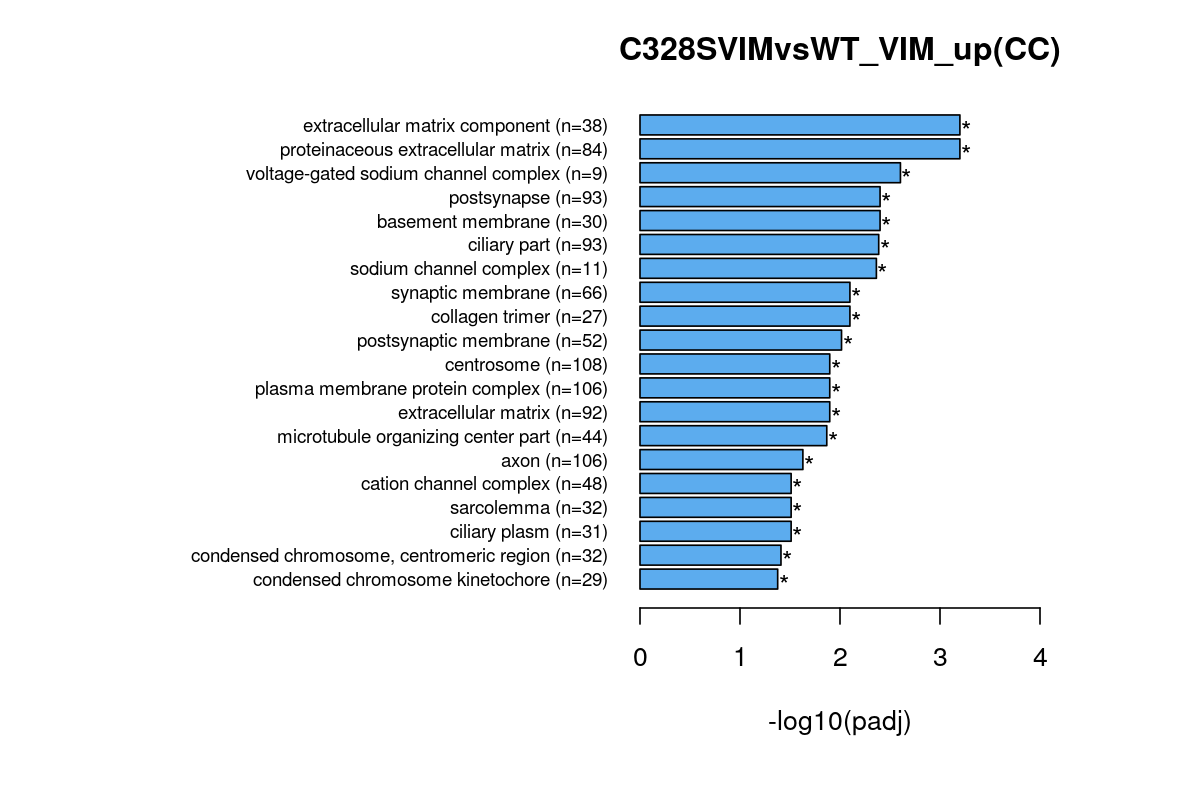

### C328SVIMvsWT_VIM_up.dot.pdf

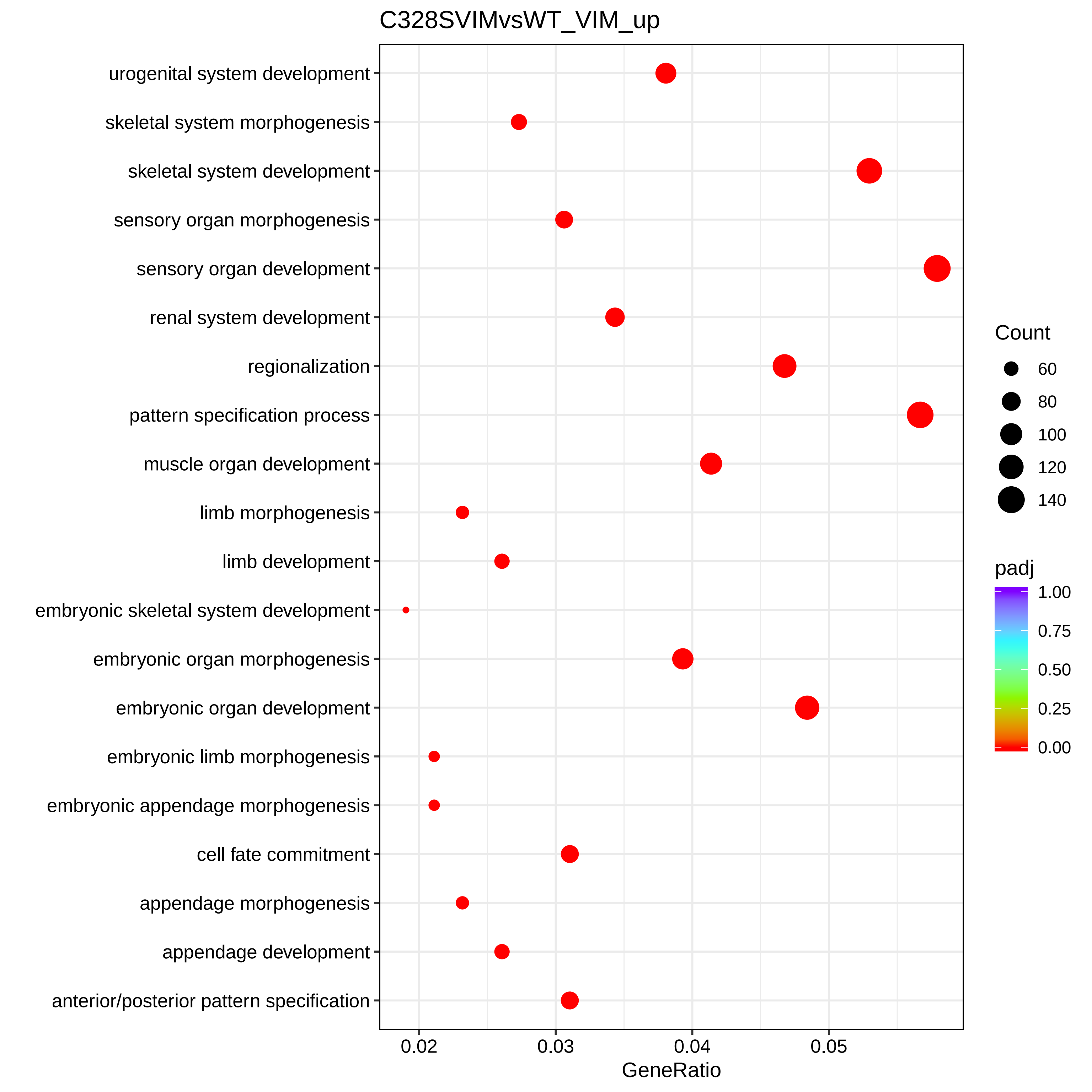

### C328SVIMvsWT_VIM_up.dot.png

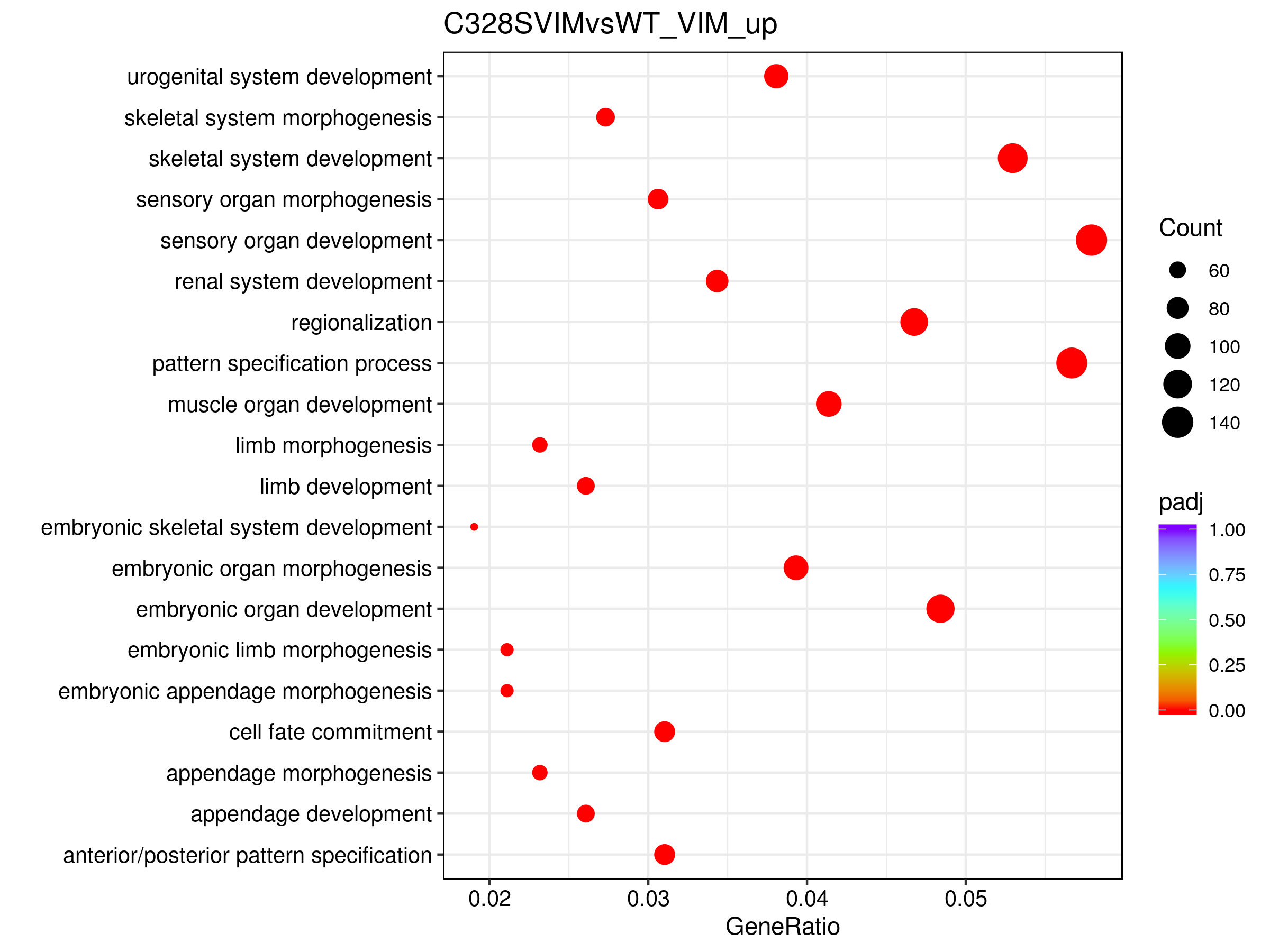

### C328SVIMvsWT_VIM_up.GObp_DAG.pdf

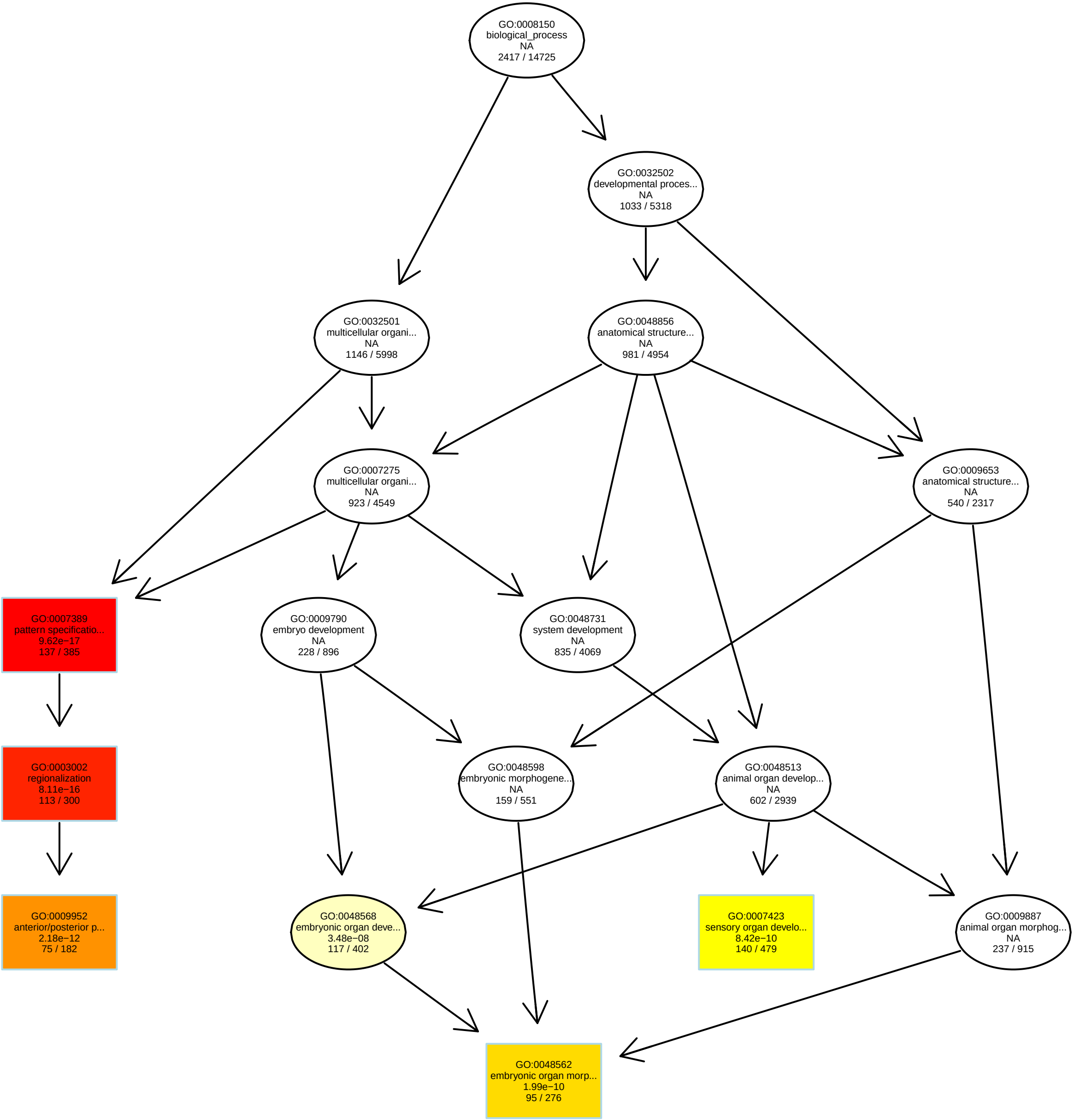

### C328SVIMvsWT_VIM_up.GObp_DAG.png

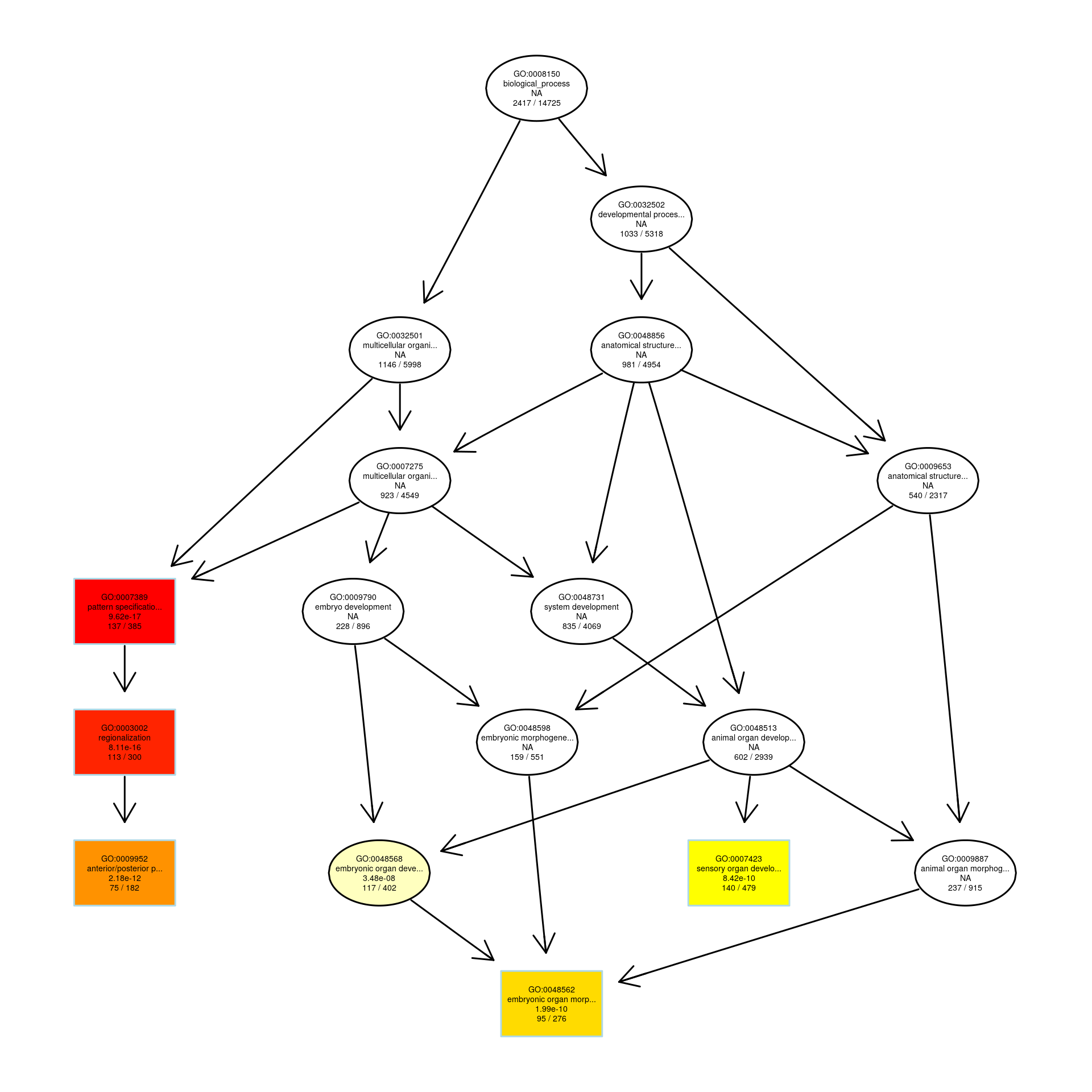

### C328SVIMvsWT_VIM_up.GOmf_DAG.pdf

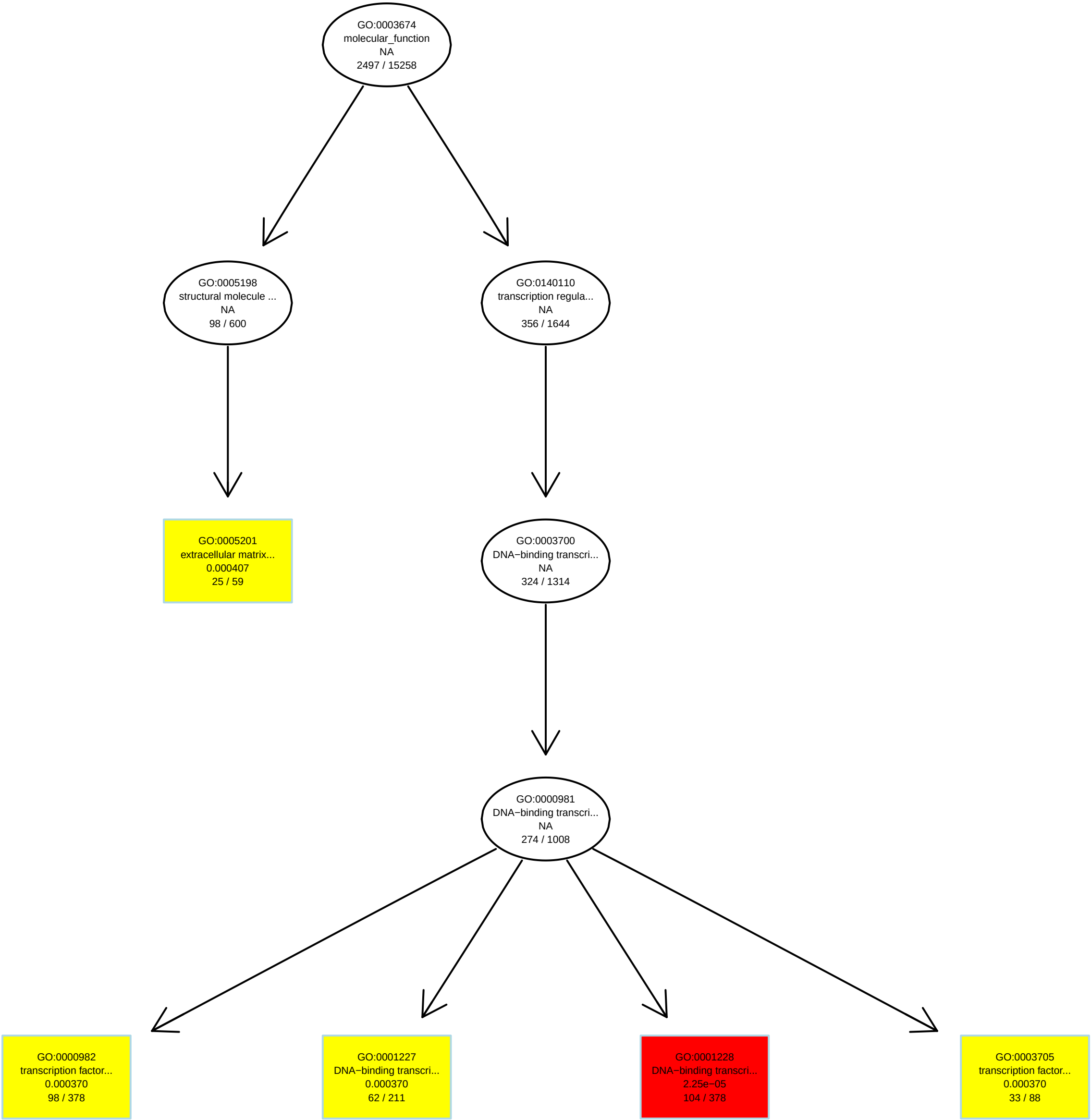

### C328SVIMvsWT_VIM_up.GOmf_DAG.png

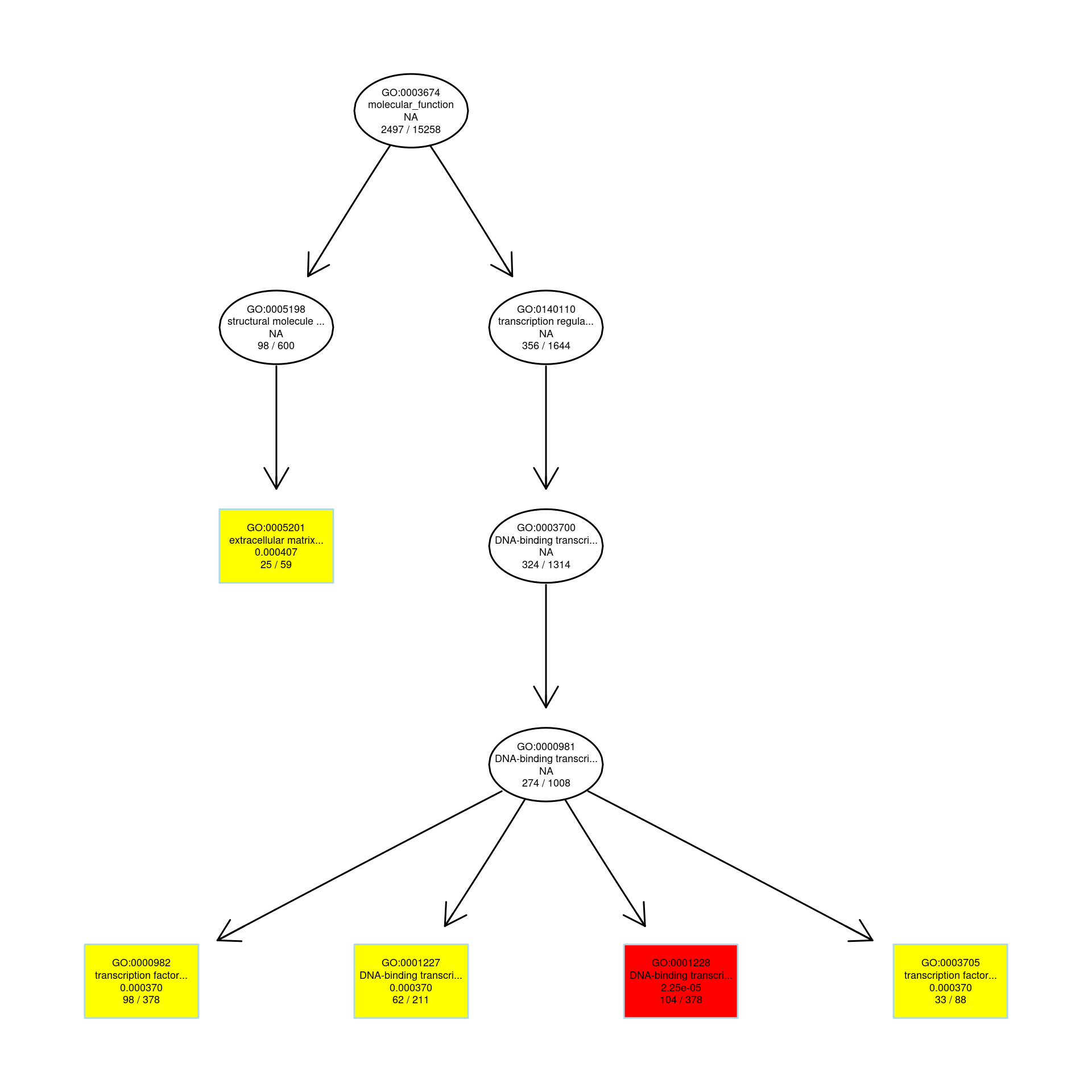

### C328SVIMvsWT_VIM_up.MFbar.pdf

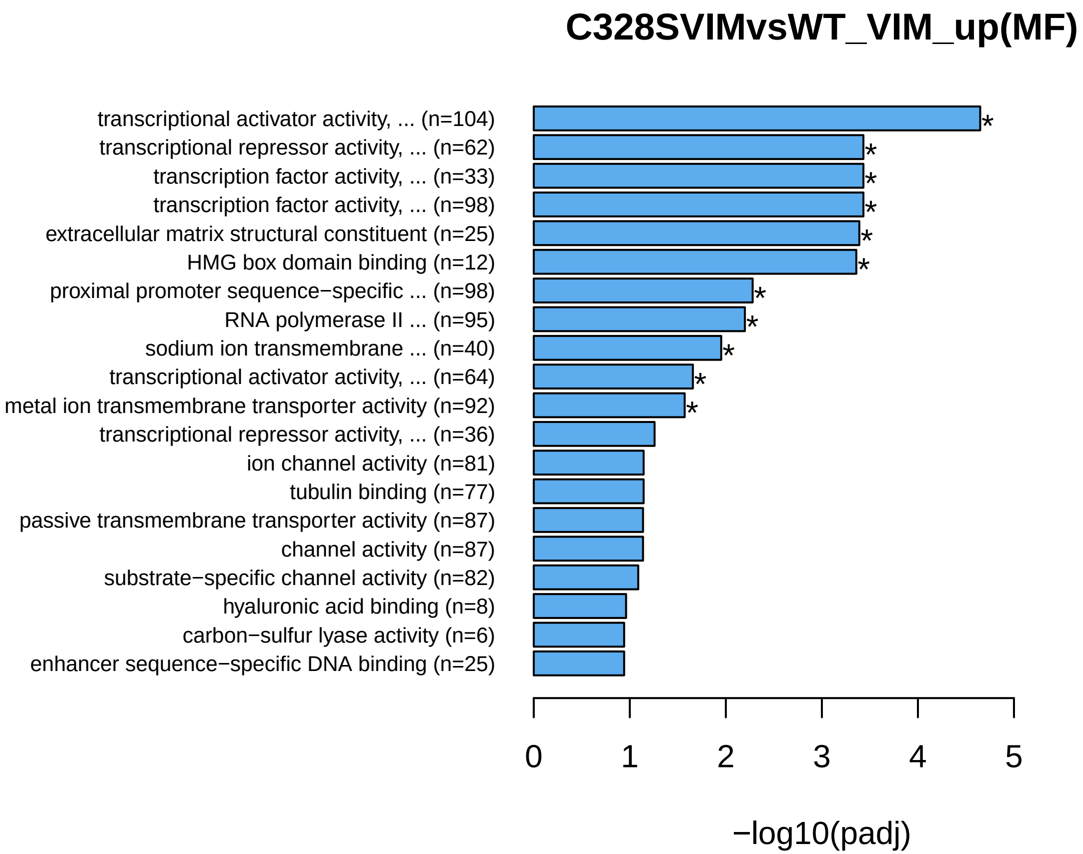

### C328SVIMvsWT_VIM_up.MFbar.png

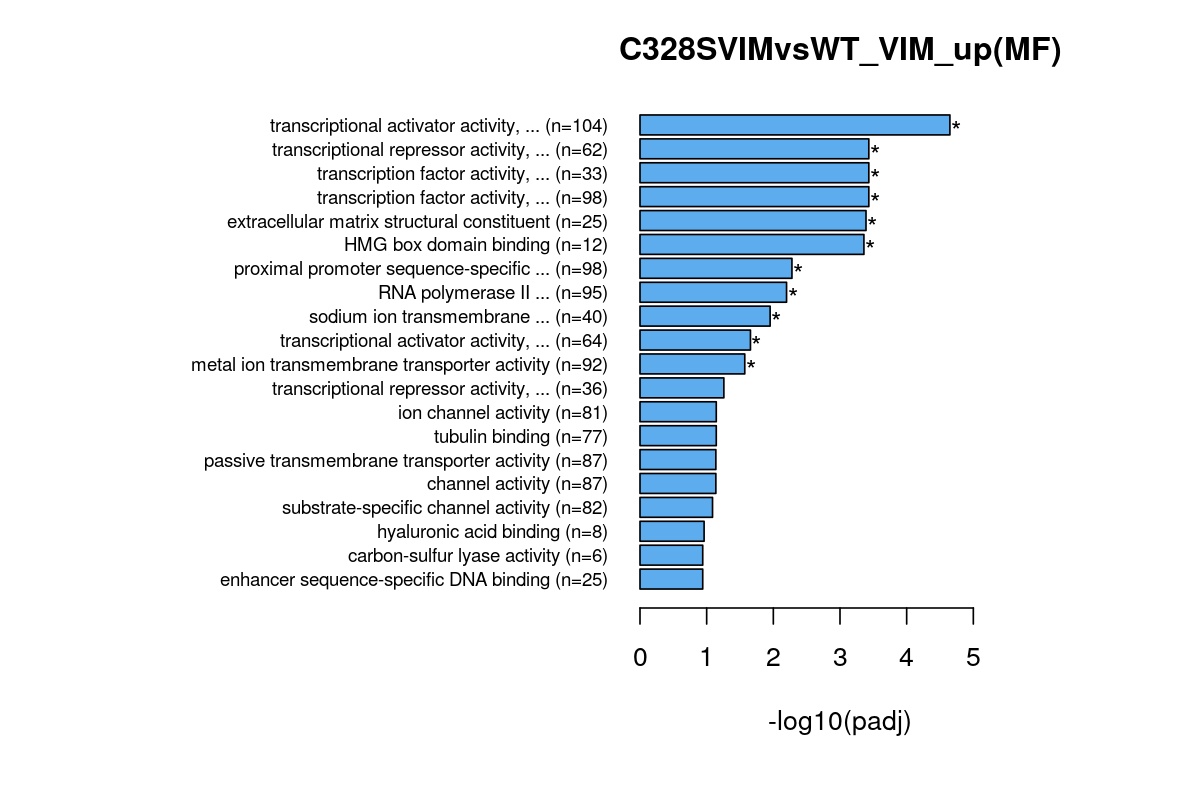

### CEREBELLAR_CORTEX_MORPHOGENESIS(GO_0021696)_49.png

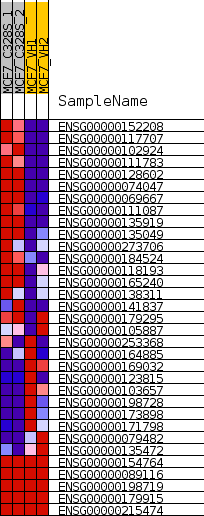

### CYTOSOLIC_TRANSPORT(GO_0016482)_4.png

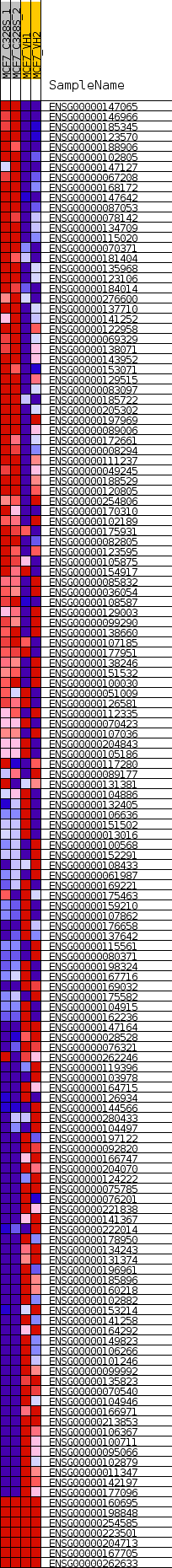

### EARLY_ENDOSOME_TO_LATE_ENDOSOME_TRANSPORT(GO_0045022)_10.png

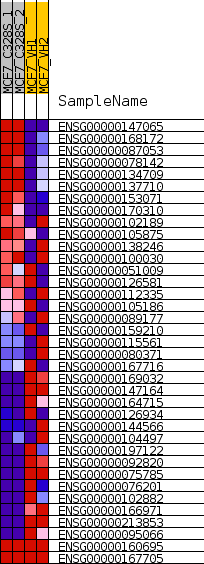

### enplot_ATRIAL_SEPTUM_DEVELOPMENT(GO_0003283)_51.png

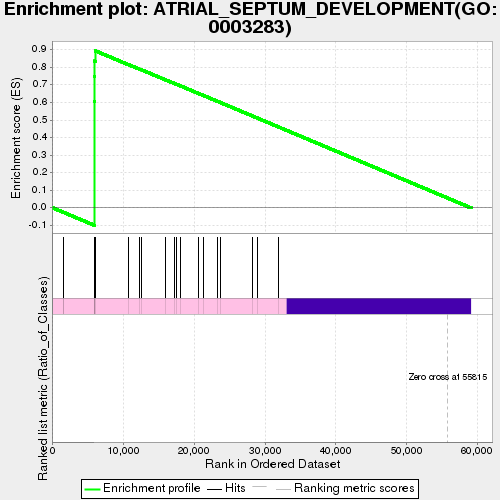

### enplot_CEREBELLAR_CORTEX_MORPHOGENESIS(GO_0021696)_48.png

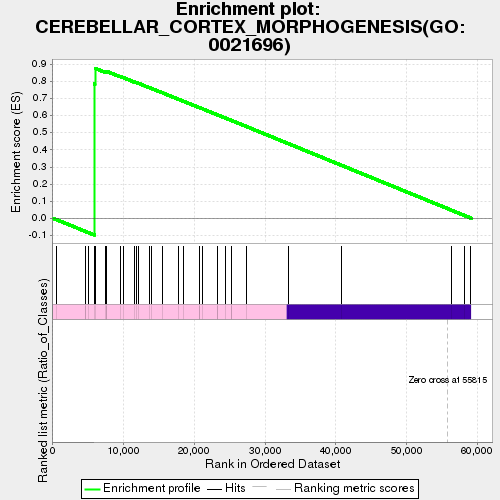

### enplot_CYTOSOLIC_TRANSPORT(GO_0016482)_3.png

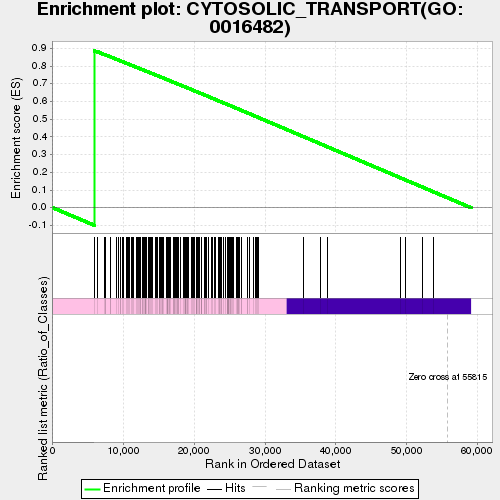

### enplot_EARLY_ENDOSOME_TO_LATE_ENDOSOME_TRANSPORT(GO_0045022)_9.png

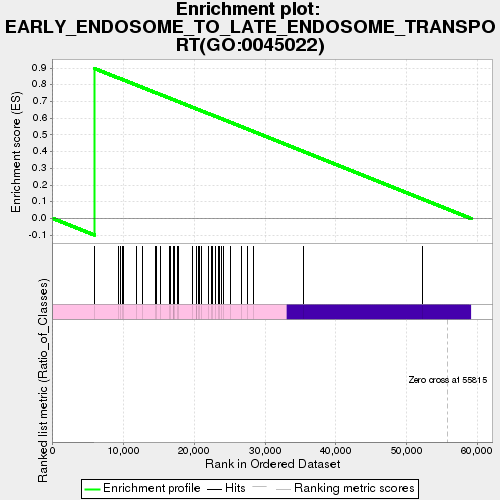

### enplot_ESTABLISHMENT_OR_MAINTENANCE_OF_APICAL_BASAL_CELL_POLARITY(GO_0035088)_18.png

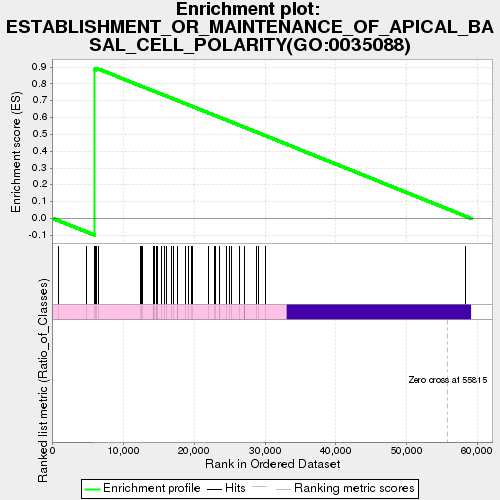

### enplot_ESTABLISHMENT_OR_MAINTENANCE_OF_BIPOLAR_CELL_POLARITY(GO_0061245)_15.png

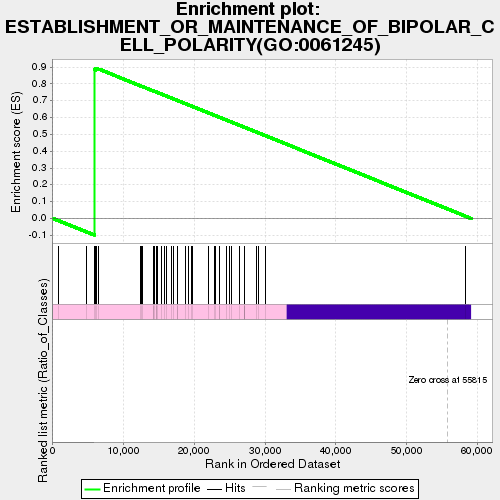

### enplot_ESTABLISHMENT_OR_MAINTENANCE_OF_EPITHELIAL_CELL_APICAL_BASAL_POLARITY(GO_0045197)_36.png

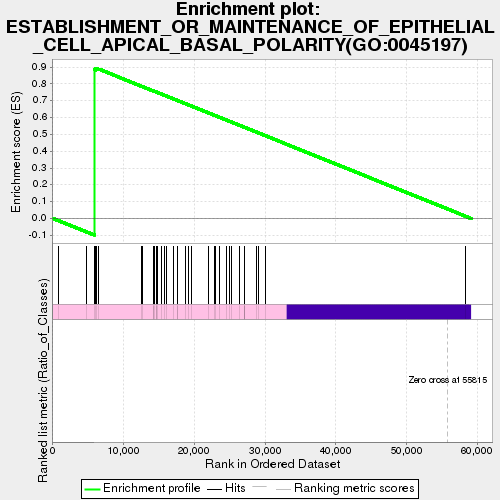

### enplot_IMMUNOGLOBULIN_COMPLEX__CIRCULATING(GO_0042571)_69.png

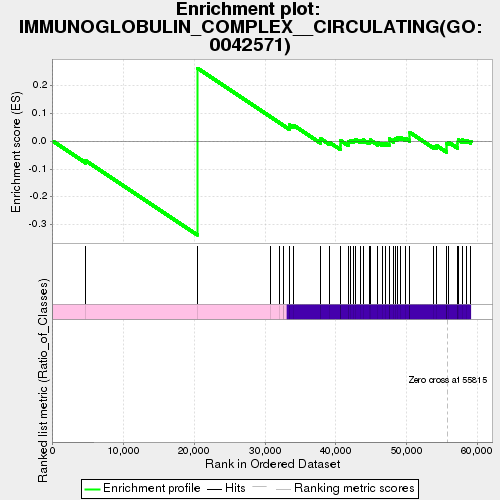

### enplot_IMMUNOGLOBULIN_RECEPTOR_BINDING(GO_0034987)_66.png

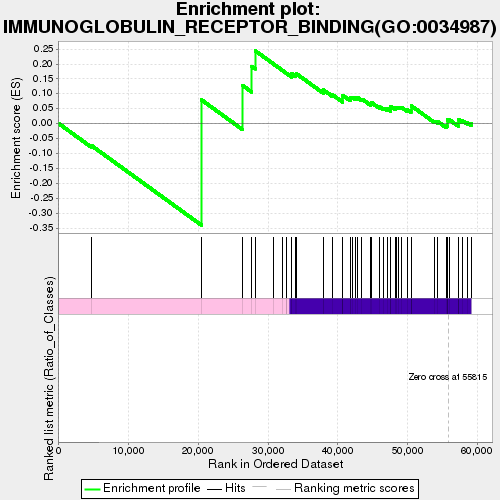

### enplot_MYELIN_SHEATH(GO_0043209)_57.png

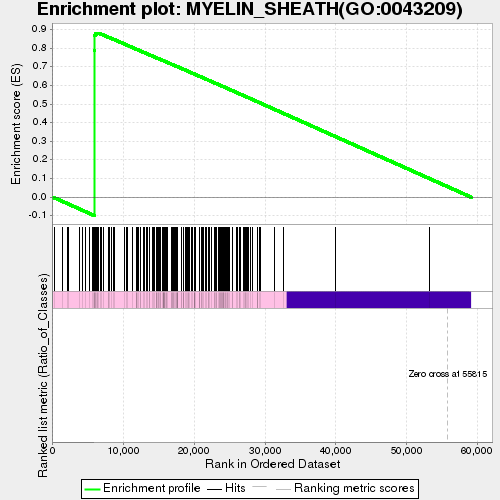

### enplot_NEGATIVE_REGULATION_OF_INTERLEUKIN_1_PRODUCTION(GO_0032692)_12.png

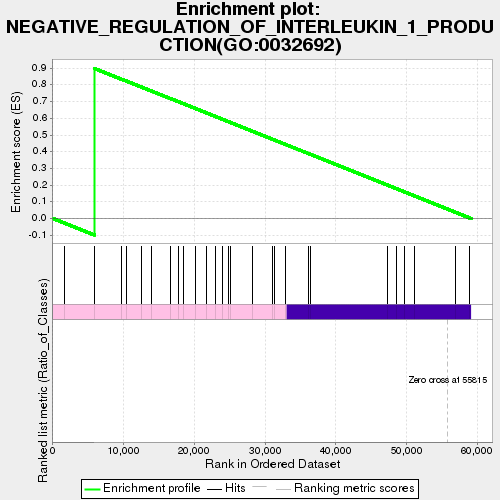

### enplot_POSITIVE_REGULATION_OF_PEPTIDYL_SERINE_PHOSPHORYLATION_OF_STAT_PROTEIN(GO_0033141)_63.png

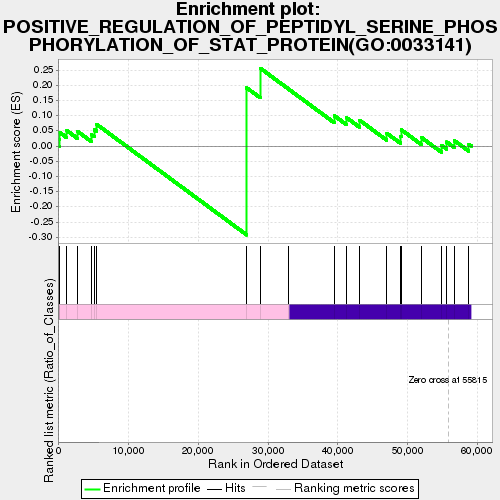

### enplot_PROTEIN_LOCALIZATION_TO_ENDOSOME(GO_0036010)_39.png

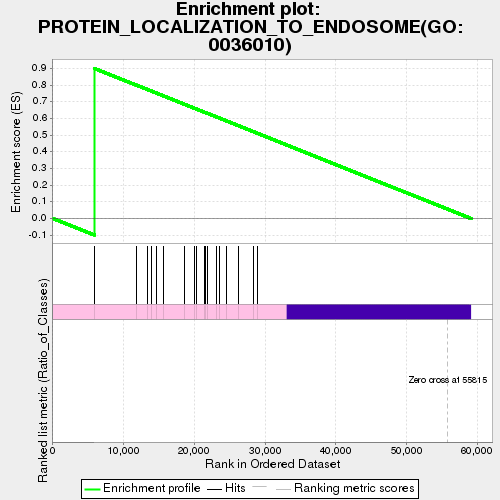

### enplot_REGULATION_OF_CELLULAR_PROTEIN_CATABOLIC_PROCESS(GO_1903362)_54.png

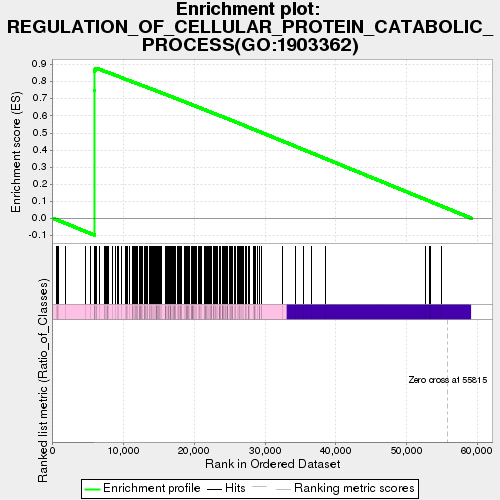

### enplot_REGULATION_OF_TUMOR_NECROSIS_FACTOR_MEDIATED_SIGNALING_PATHWAY(GO_0010803)_33.png

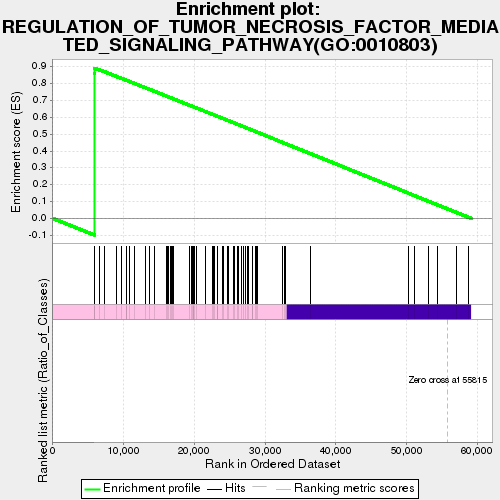

### enplot_RESPONSE_TO_INTERLEUKIN_12(GO_0070671)_27.png

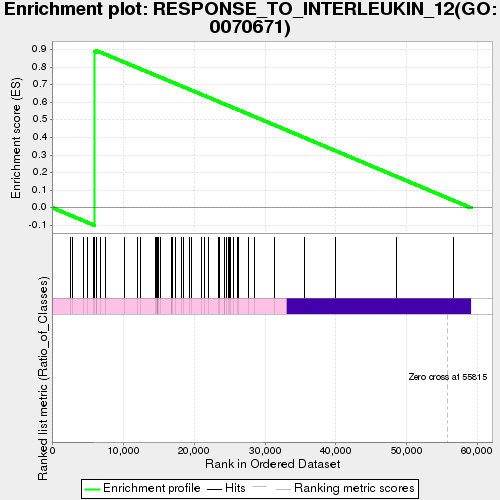

### enplot_TYPE_I_INTERFERON_RECEPTOR_BINDING(GO_0005132)_72.png

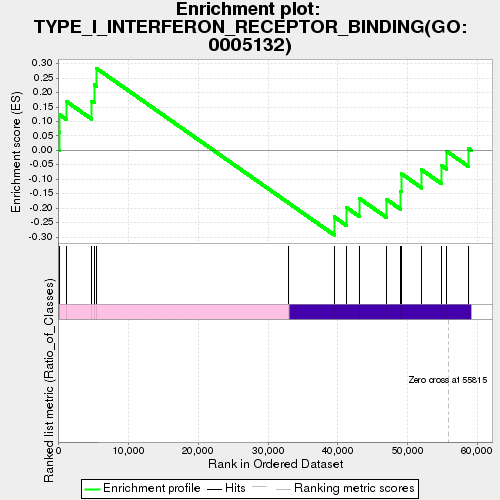

### enplot_VESICLE_MEDIATED_TRANSPORT_BETWEEN_ENDOSOMAL_COMPARTMENTS(GO_0098927)_6.png

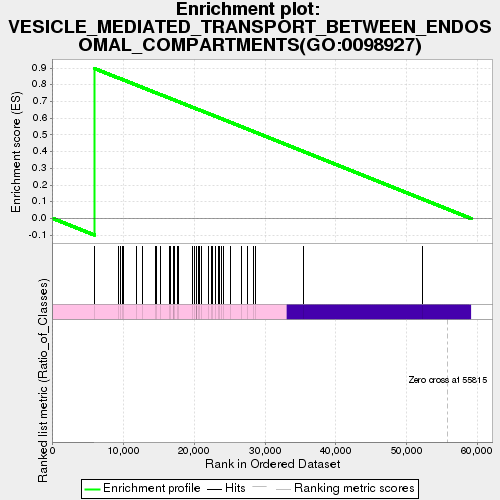

### ESTABLISHMENT_OR_MAINTENANCE_OF_APICAL_BASAL_CELL_POLARITY(GO_0035088)_19.png

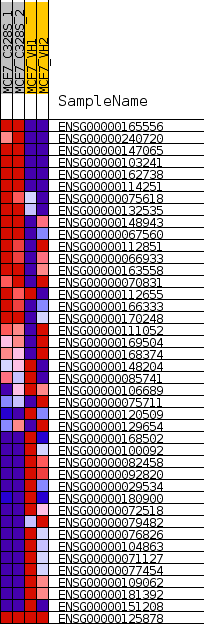

### ESTABLISHMENT_OR_MAINTENANCE_OF_EPITHELIAL_CELL_APICAL_BASAL_POLARITY(GO_0045197)_37.png

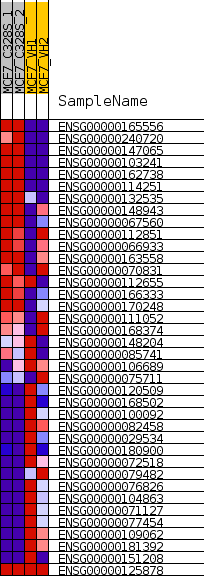

### global_es_histogram.png

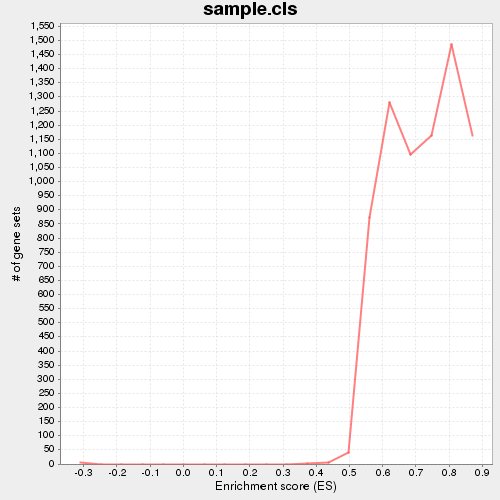

### gset_rnd_es_dist_5.png

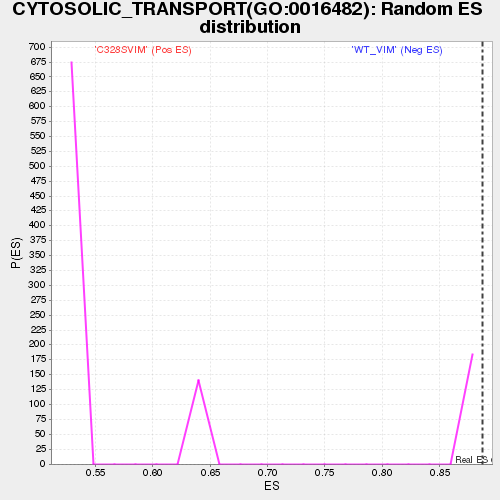

### gset_rnd_es_dist_8.png

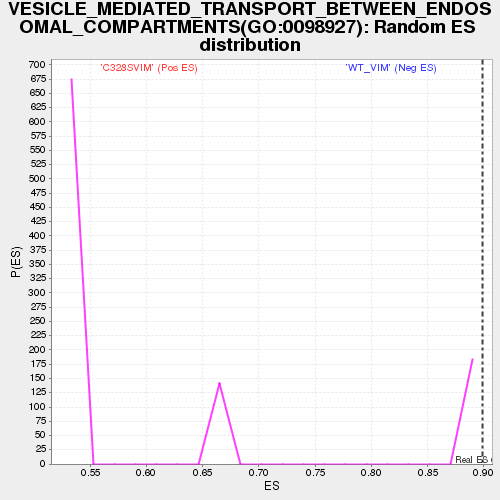

### gset_rnd_es_dist_11.png

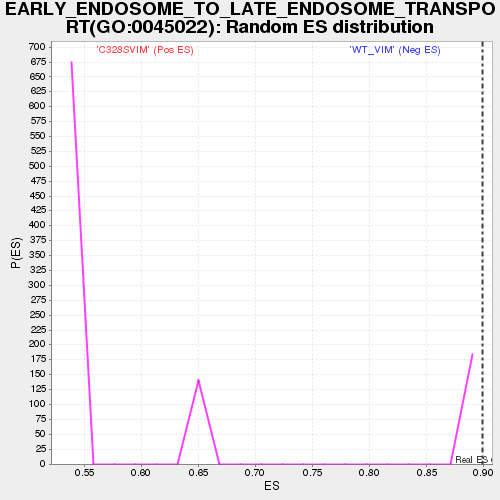

### gset_rnd_es_dist_14.png

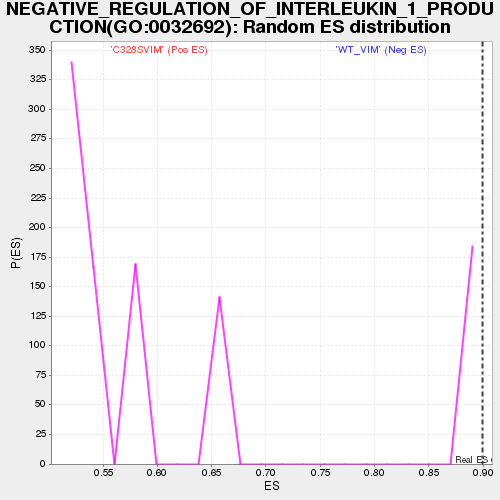

### gset_rnd_es_dist_17.png

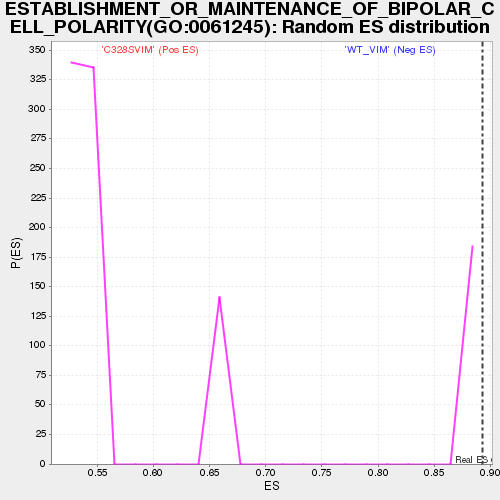

### gset_rnd_es_dist_20.png

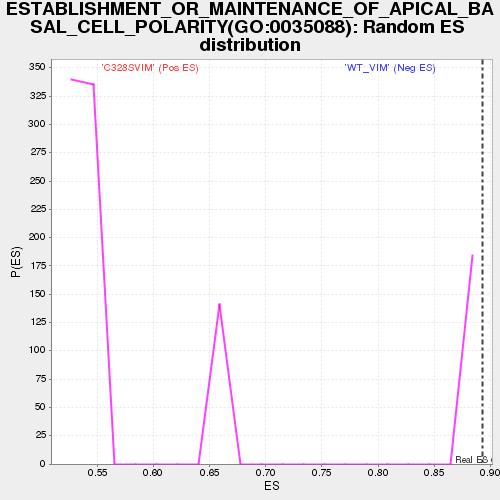

### gset_rnd_es_dist_29.png

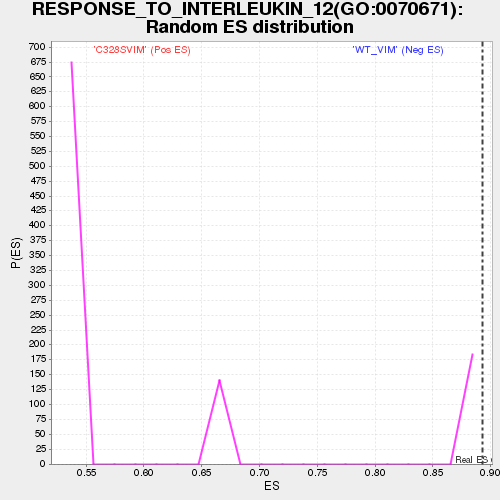

### gset_rnd_es_dist_35.png

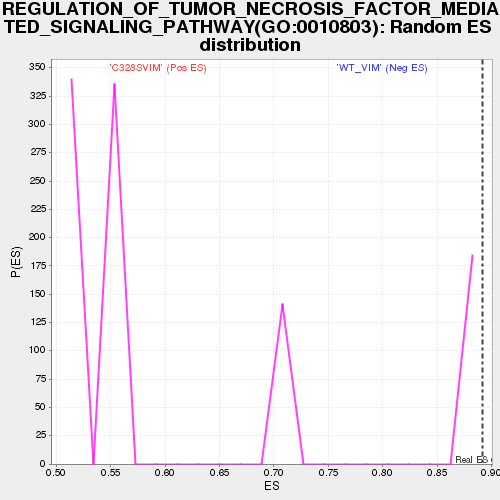

### gset_rnd_es_dist_38.png

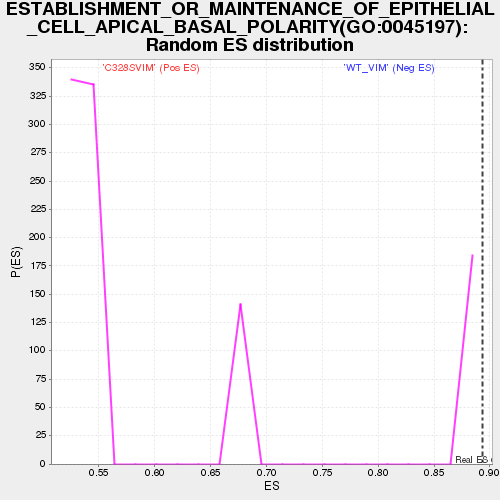

### gset_rnd_es_dist_50.png

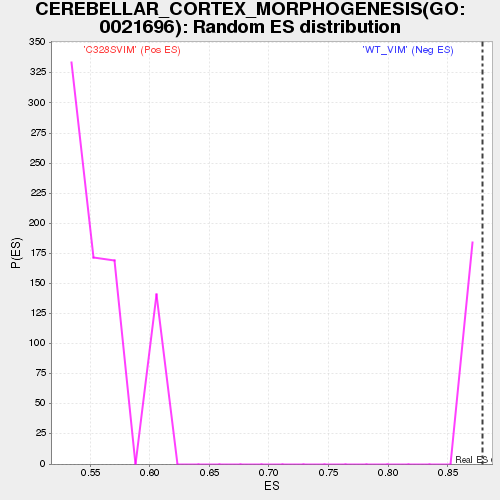

### gset_rnd_es_dist_53.png

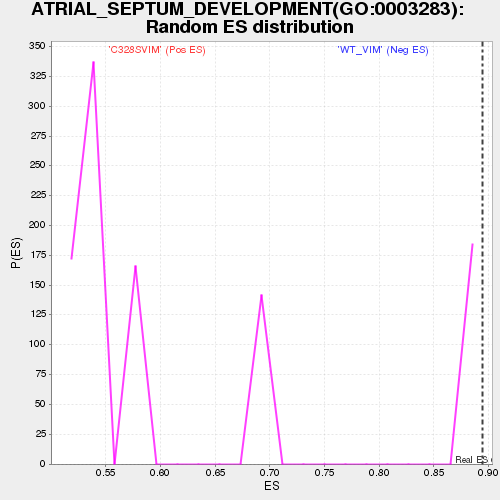

### gset_rnd_es_dist_56.png

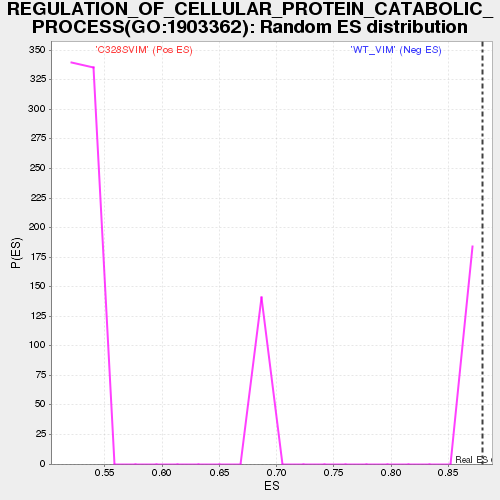

### gset_rnd_es_dist_59.png

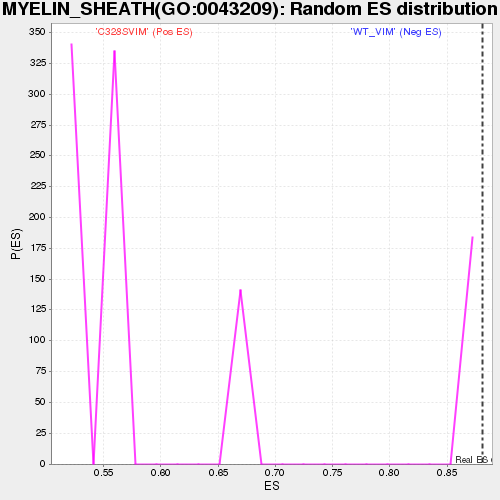

### gset_rnd_es_dist_65.png

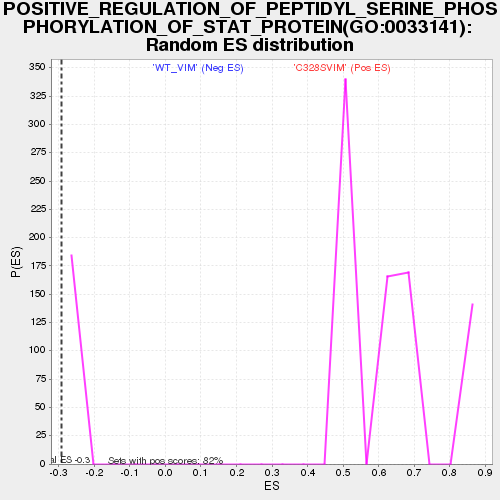

### gset_rnd_es_dist_68.png

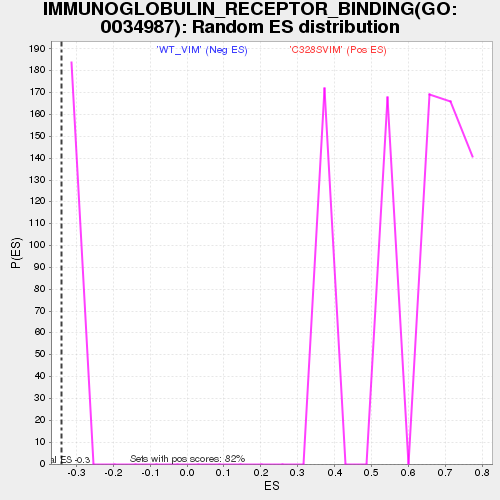

### gset_rnd_es_dist_71.png

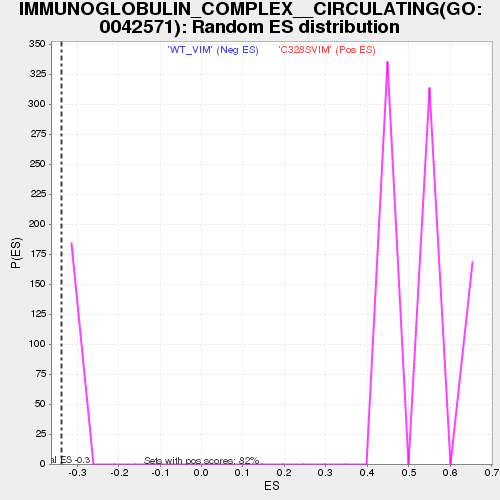

### gset_rnd_es_dist_74.png

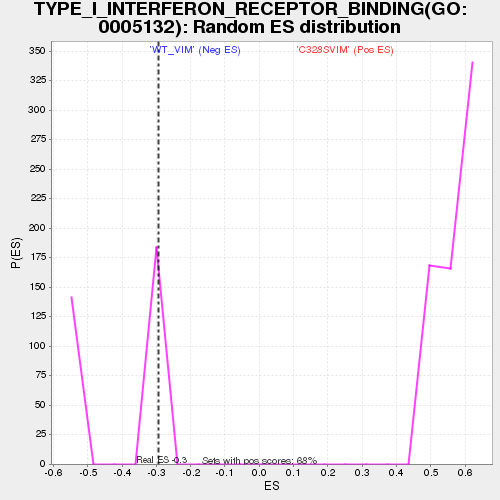

### heat_map_1.png

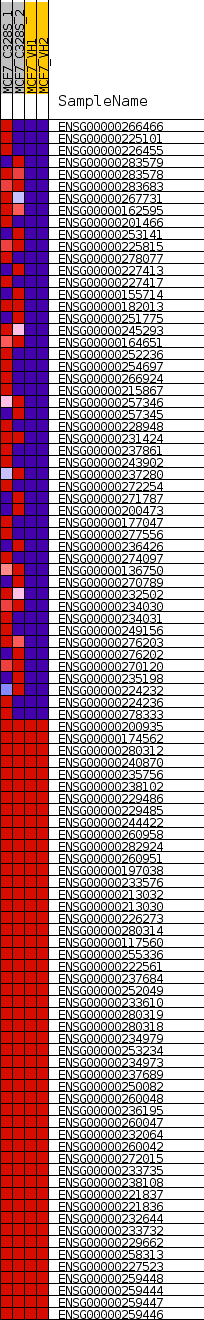

### IMMUNOGLOBULIN_COMPLEX__CIRCULATING(GO_0042571)_70.png

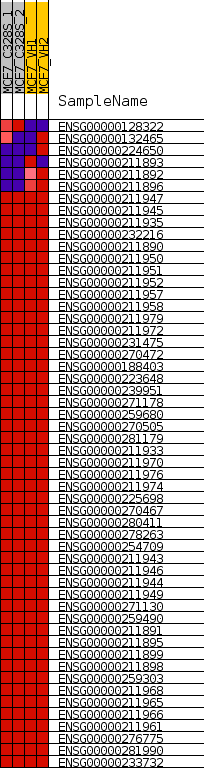

### IMMUNOGLOBULIN_RECEPTOR_BINDING(GO_0034987)_67.png

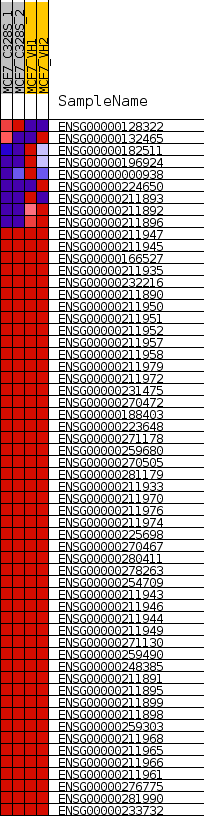

### MYELIN_SHEATH(GO_0043209)_58.png

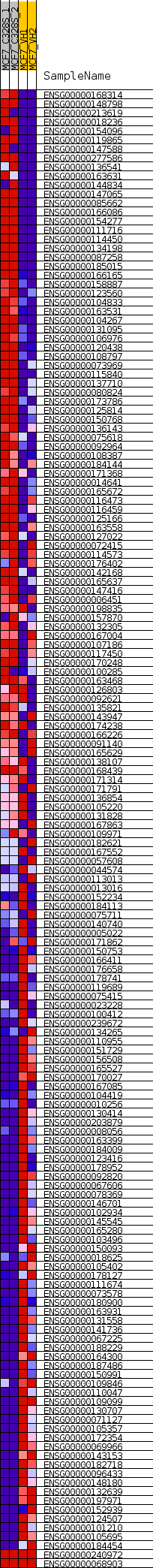

### NEGATIVE_REGULATION_OF_INTERLEUKIN_1_PRODUCTION(GO_0032692)_13.png

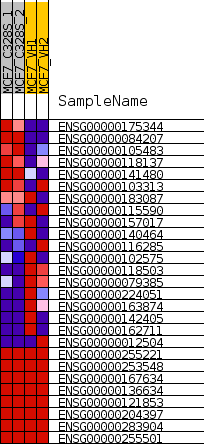

### POSITIVE_REGULATION_OF_PEPTIDYL_SERINE_PHOSPHORYLATION_OF_STAT_PROTEIN(GO_0033141)_64.png

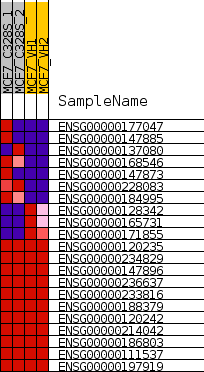

### pvalues_vs_nes_plot.png

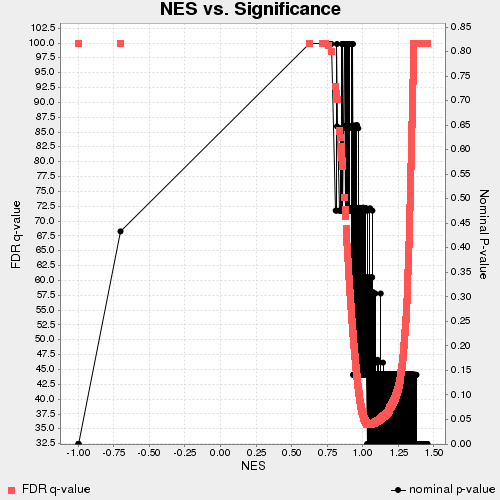

### ranked_list_corr_2.png

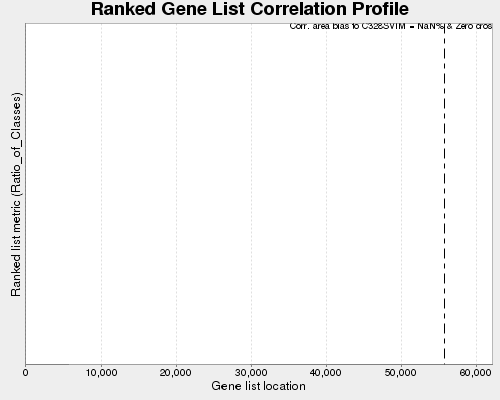

### REGULATION_OF_CELLULAR_PROTEIN_CATABOLIC_PROCESS(GO_1903362)_55.png

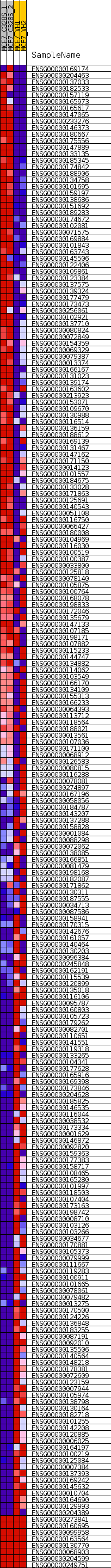

### REGULATION_OF_TUMOR_NECROSIS_FACTOR_MEDIATED_SIGNALING_PATHWAY(GO_0010803)_34.png

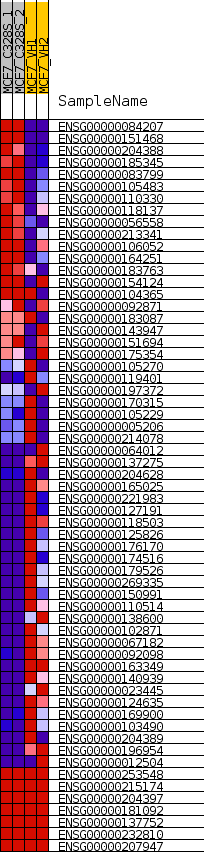

### RESPONSE_TO_INTERLEUKIN_12(GO_0070671)_28.png

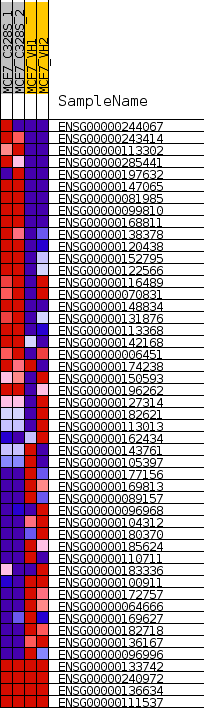

### TYPE_I_INTERFERON_RECEPTOR_BINDING(GO_0005132)_73.png

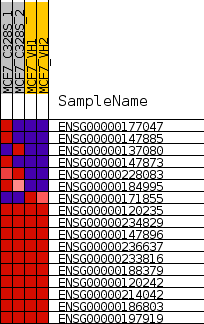

### VESICLE_MEDIATED_TRANSPORT_BETWEEN_ENDOSOMAL_COMPARTMENTS(GO_0098927)_7.png

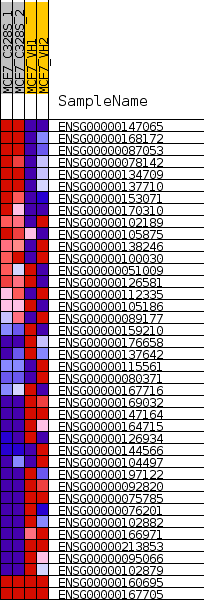
